# ENHANCED GRAVITROPISM 2 coordinates molecular adaptations to gravistimulation in the elongation zone of barley roots

**DOI:** 10.1101/2022.10.11.511704

**Authors:** Li Guo, Alina Klaus, Marcel Baer, Gwendolyn K. Kirschner, Silvio Salvi, Frank Hochholdinger

## Abstract

- Root gravitropism includes gravity perception in the root cap, signal transduction between root cap and elongation zone, and curvature response in the elongation zone. The barley (*Hordeum vulgare*) mutant *enhanced gravitropism 2* (*egt2*) displays a hypergravitropic root phenotype.
- We compared the transcriptomic reprogramming of the root cap, the meristem and the elongation zone of wild type and *egt2* seminal roots upon gravistimulation in a time-course experiment and identified direct interaction partners of EGT2 by yeast-two-hybrid screening and bimolecular fluorescence complementation (BiFC) validation.
- We demonstrated that the elongation zone is subjected to most transcriptomic changes after gravistimulation. Here, 35% of graviregulated genes are also transcriptionally controlled by *EGT2*, suggesting a central role of this gene in controlling the molecular networks associated with gravitropic bending. Gene co-expression analyses suggested a role of *EGT2* in cell wall and reactive oxygen species (ROS)-related processes, in which direct interaction partners of EGT2 regulated by *EGT2* and gravity might be involved.
- Taken together, this study demonstrated the central role of *EGT2* and its interaction partners in the networks controlling root zone-specific transcriptomic reprogramming of barley roots upon gravistimulation. These findings can contribute to the development of novel root idiotypes leading to improved crop performance.

## Introduction

The gravitropic response plays an essential role in controlling plant growth direction and architecture under changing environmental conditions (Nakamura *et al*., 2019). Root gravitropism enables plants to adjust their root growth direction, allowing them to penetrate deeply into the soil, thereby anchoring plants in the soil and optimizing the uptake of water and nutrients (Zhang *et al*., 2019). In flowering plants, root gravitropic response is a three-step process: gravity perception, signal transduction and curvature response (Su *et al*., 2017; Singh *et al*., 2017; Nakamura *et al*., 2019; Jiao *et al*., 2021).

The most prominent explanation of how roots sense gravity is the starch-statolith hypothesis: upon root reorientation, the gravity signal is sensed by starch-filled amyloplasts sedimented to the new bottom-site of the columella cells in the root cap (Su *et al*., 2017; Singh *et al*., 2017). This results in the relocalization of auxin towards the bottom side of columella cells by downward auxin transport and a lateral auxin gradient across the root cap. Auxin is then transported to the lateral root cap and epidermal cells in a shootward direction towards the elongation zone, resulting in asymmetrical auxin distribution across the distal elongation zone with higher auxin levels on the lower flank and lower auxin content on the upper flank. In the epidermal cells on the lower flank of the elongation zone, elevated auxin levels increase the apoplastic pH and inhibit cell elongation. In contrast, on the upper side of the elongation zone, decreased auxin levels lead to a decrease in the pH of the apoplast, which increases cell elongation. The asymmetrical elongation of cells on the two sides of the gravistimulated root eventually leads to the gravitropic root bending in the direction of gravity (Su *et al*., 2020). Nevertheless, cells within the root distal elongation zone have also been shown to play a role in gravity perception with an alternative mechanism (Wolverton *et al*., 2002; Mancuso *et al*., 2006; Su *et al*., 2020).

In a previous study, we identified the barley mutant *egt2* (*enhanced root gravitropism 2*) which displays a narrower root system than wild-type plants, with root angles at least 50% smaller than wild type. Phytohormone treatment showed that *egt2* responds to auxin similarly as the wild type, suggesting that the auxin response is unaffected in *egt2*. Moreover, we did not observe any significant differences between the structure and size of the meristem, the root cap, amyloplast content and the amyloplast sedimentation velocities between *egt2* and wild type. This suggests that *EGT2* is likely involved in root gravitropism after gravity perception. The *EGT2* gene encodes a STERILE ALPHA MOTIF (SAM) domain–containing protein (Kirschner *et al*., 2021). SAM domains are present in thousands of proteins in prokaryotes and diverse eukaryotes (Kim & Bowie, 2003; Qiao & Bowie, 2005; Denay *et al*., 2017). The SAM domain was first considered as an evolutionarily conserved protein binding domain (Schultz *et al*., 1997). Subsequently, a large diversity of SAM functions has been observed, including the binding of proteins, lipids and RNAs (Denay *et al*., 2017; Ray *et al*., 2020). The best characterized plant SAM domain-containing protein is LEAFY which functions as a master regulator of flower development in angiosperms (Eriksson *et al*., 2006; Siriwardana & Lamb, 2012; Yamaguchi, 2021). The SAM domain at the N-terminus of LEAFY mediates oligomerization of LEAFY, which allows the LEAFY dimers to bind to DNA regions without LEAFY-binding sites or closed chromatin regions (Sayou *et al*., 2016). AtSAM5, a homolog of EGT2 in Arabidopsis, has been shown to interact with the Ca^2+^-dependent protein kinase CPK13 (Jones *et al*., 2014), which is involved in the response to light by controlling the stomatal aperture (Ronzier *et al*., 2014). Moreover, narrower lateral root angles were observed in mutants of *AtSAM5*, suggesting a role of *AtSAM5* in regulating lateral root angle in Arabidopsis (Johnson *et al*., 2022). Another homolog of EGT2 is WEEP in peach trees (Hollender *et al*., 2018). A 5’ deletion of the *WEEP* gene caused downward and wandering growth of the mutant shoots that did not bend upward after 90° reorientation. This suggests a role of WEEP in shoot gravitropism, which shows similarities to the function of EGT2 in roots (Hollender *et al*., 2018). In barley, 17 SAM domain-containing proteins were annotated and none of them has been functionally characterized thus far except for *EGT2* (Kirschner *et al*., 2021).

In the present study, the transcriptomes of the root cap, the meristem and the elongation zone of gravistimulated wild type and mutant *egt2* roots were studied by RNA sequencing (RNA-seq) in a time-course experiment. These experiments revealed that EGT2 is a central regulator of the gravitropism-regulated gene expression networks in barley seminal roots. Functional enrichment analyses of differentially expressed genes (DEGs) and weighted gene co-expression network analyses suggested the involvement of plant cell wall organization and reactive oxygen species (ROS) in *EGT2*-regulated root gravitropism. By a combination of yeast-two-hybrid experiments and bimolecular fluorescence complementation (BiFC) we demonstrated that a subset of the genes transcriptionally regulated by gravistimulation and by *EGT2* encoded proteins that directly interact with EGT2. The results of this study highlight the transcriptomic reprogramming of the gravitropic response of barley roots and provide new insights into the molecular function of *EGT2*.

## Materials and Methods

### Plant materials and growth condition

Wild-type barley plants (*Hordeum vulgare* L. cv. Morex) and the *enhanced gravitropism 2* mutant (*egt2*, Kirschner *et al*., 2021) were used for this study. Seeds germination and plant growth were performed as previously described (Kirschner *et al*., 2021).

### Measurement of meristem length

To measure the length of the meristem, 3-day-old wild type and *egt2* plants were rotated by 90° for 6 or 10 hours. Root tip segments of 5 mm length of the most vertical roots before rotation were taken before and after rotation. Root tips were then embedded in 10% agarose and 40 µm sections were prepared with a vibratome. Images of root sections taken with a Zeiss PALM MicroBeam microscope were analyzed by ImageJ. The zone between the first cell in the first cortical layer that doubled in size and the first cell that increased noticeably in length was considered as the transition zone. The length from root cap junction to the middle of transition zone were measured as the length of the meristem (Verbelen *et al*., 2006; Baluška *et al*., 2010). Eleven to nineteen roots were analyzed per genotype by time point combination (Supporting Information Fig. S6).

### Sample collection and RNA isolation

For RNA sequencing, 3-day-old plants were rotated by 90° for 3, 6 and 12 h. The root cap, the meristem (1 mm; Supporting Information Fig. S6) and 1 mm of the distal elongation zone (Fig. 1a) of the most vertically grown seminal roots before rotation were separated by a razor blade under a binocular before (time point 0 h) and after rotation. For RNA-seq, three roots were collected per seedling, root zones from eight plants were pooled per biological replicate. Three biological replicates were used for each genotype by time point combination. RNA extraction was performed with the RNeasy Plant Mini Kit (Qiagen) according to the manufacturer’s protocol. RNA quality was assessed with an Agilent 2100 Bioanalyzer using the Agilent RNA 6000 Nano kit. The RNA integrity number was between 8.3 to 10 for all samples.

**Fig. 1:**
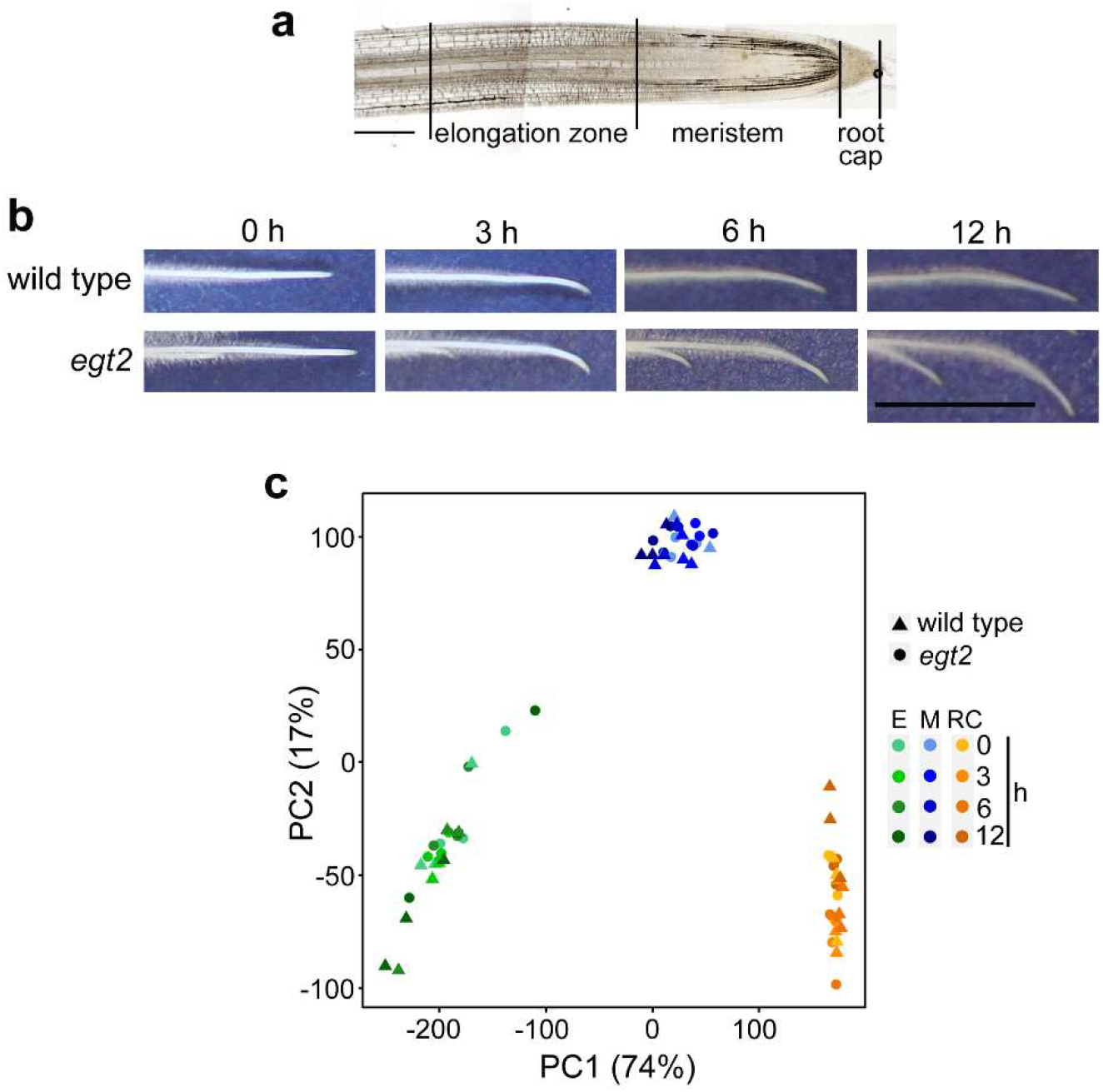
Seminal roots upon rotation and RNA sequencing sample relationship of root zones. **a** Representative section for meristem length measurement, illustrating the elongation zone, the meristem, and the root cap harvested for RNA-seq. Scale bar = 300 µm. **b** Phenotype of wild type and *egt2* mutant roots before (0 h) and after 3 h, 6 h and 12 h of rotation. Scale bar = 1 cm. **c** PCA plot for the transcriptomes of the 72 RNA-seq samples of the two genotypes, three root zones and four time points.

### RNA-seq and data analyzing

Messenger RNA (mRNA) purified from total RNA using poly-T oligo-attached magnetic beads was used to prepare cDNA libraries for Illumina sequencing. Fragmented mRNA and random hexamer primers were used for reverse transcription. The cDNA library was prepared by ligating the cDNA product to adapter, followed by amplification and purification (Novogene, China). Cluster preparation and single read sequencing were performed according to the manufacturer’s instructions (Illumina). Sequencing was performed in a NovaSeq 6000, S4 FlowCell using the paired-end 150 bp sequencing strategy. Raw RNA-seq reads were processed with CLC Genomics Workbench (Version 20.0.1) as previously described with some minor modifications (Osthoff *et al*., 2019). Reads with a length <40 bp were discarded and the remaining reads were mapped to the barley reference genome (Hv_Morex.pgsb.Jul2020) with a similarity of 0.9 and a length fraction of 0.8. Reads mapped to more than one position were regarded as duplicate mapped reads and removed from further analyses (Mascher *et al*., 2021).

Data generated by CLC Genomics Workbench was analyzed using R. To remove lowly expressed genes, the cpm() function from the edgeR library was used to normalize the different sequencing depths for each sample (Robinson *et al*., 2010). A CPM threshold of 1.2 was used and only genes that have at least 3 TRUES in each row of threshold were kept for further steps. A principal component analysis (PCA) was computed by R the functions prcomp() and visualized by the autoplot() function in the ggplot2 package. The contrasts.fit function of the R package limma was used to calculate log_2_ fold change (log_2_FC) values between different root zones and time point combinations in the wild type time course experiment and in comparisons between the *egt2* mutant and wild type (Ritchie *et al*., 2015). The false discovery rate (FDR) was adjusted to <5% to correct *p*-values of contrasts for multiple testing (Benjamini & Hochberg, 1995). Venn diagrams were produced by the VennDiagram package (Chen & Boutros, 2011). A hierarchical clustering analysis was conducted by the cluster package with dynamic tree cutting (‘Package “cluster”‘, 2022). The upset() function in the UpSetR library was used for comparison between multiple gene sets (Conway *et al*., 2017). Gene Ontology (GO) analyses were performed by the R function topGO (Alexa & Rahnenführer, 2015). Redundant GO terms were filtered by REVIGO with a similarity ≤0.5 (Supek *et al*., 2011).

### Gene weighted correlation network analysis (WGCNA)

The R package WGCNA was used for gene co-expression analyses (Langfelder and Horvath, 2008). After filtering out lowly-expressed genes with <50 total reads in the 24 samples of each root zone, the read data generated by CLC Genomics Workbench was used as input. Samples were clustered by using the hclust function. The outlier samples were determined by setting the cutheight to 50 for the root cap, 40 for the meristem and 75 for the elongation zone. After removing outliers, the co-expression modules of each root zone dataset were generated by the one-step network construction function blockwiseModules with a soft threshold of 4 for the root cap and elongation zone and 6 for the meristem, and setting TOMtype to ‘signed’. To avoid small clusters, modules with <30 genes were merged with their closest larger module using the cutreeDynamic function. The eigengene calculation of each module was performed by using the moduleEigengenes function followed by the calculation of the module dissimilarity of eigengenes. Modules whose eigengenes were correlated >0.8 were merged via the mergeCutHeight function with a cutHeight of 0.20. A unique color was then assigned to each merged module via the plotDendroAndColors function. The edge weight determined by the topology overlap measure (TOM) reflected the connectivity between two genes. The weight values across all edges of each gene in each module were calculated and exported for gene network visualization, which was achieved by Cytoscape (version 3.9.1). To determine the hub genes of each module, we sorted the weight values of each edge from the highest to smallest, and nodes with the top 1%, 0.1% or 0.01% of weight were considered as hub genes based on the size of the module. Gene networks were analyzed with the ‘Analyze Network’ tool in Cytoscape and the ‘degree’ values were considered to reduce the numbers of hub genes.

We calculated relationships between module eigengenes and traits using Pearson’s correlation coefficients in R. Significant *p*-values of each module and trait correlation were calculated using the corPvalueStudent function. Modules with *p*-values ⩽0.05 were considered as significantly correlated modules.

### Yeast-two-hybrid library construction

Total RNA extracted from the whole root system of ten 5-day-old seedlings was used to generate the cDNA library, which was conducted with the “Make Your Own “Mate & Plate™” Library System” according to the manufacturer’s instructions with minor modifications (Clontech). Oligo-dT primers provided in the kit were used for first-strand cDNA synthesis with 2 µg total RNA as input. Amplification of cDNA was performed by using long distance PCR with 25 cycles to generate 6-7 µg of ds cDNA. After purifying with CHROMA SPIN + TE-400 columns, ds cDNA library was transformed into yeast (*Saccharomyces cerevisiae*) strain Y187.

Generation of competent yeast cells and library transformation were conducted by using the LiAc/SS-DNA/PEG procedure (Gietz *et al*., 1995). Transformed cell plating and library cell harvesting were conducted according to the manufacturer’s instructions.

### Toxicity and autoactivation testing

To test the toxicity and autoactivation of EGT2, the full-length coding sequence of EGT2 was amplified using the primers specified in Supporting Information Table S9 and then inserted into the vector pGBKT7 using *Eco*RI and *Bam*HI restriction endonucleases. Detection of EGT2 toxicity on yeast growth was performed according to Matchmaker® Gold Yeast Two-Hybrid System User Manual (Takara). Yeast cells expressing pGBKT7-EGT2 displayed a similar growth rate to yeast cells transformed with an empty pGBKT7 vector, indicating that EGT2 is not toxic (Supporting Information Fig. S4a). To test the autoactivation of EGT2, pGBKT7-EGT2 plasmid was co-transformed into Y2HGold yeast cells with pGADT7. pGBKT7-53 and pGBKT7-lam plasmids were transformed together with pGADT7-T as positive and negative controls, respectively. The transformed cells were grown on SD/-Trp/-Leu medium at 30 °C for four days. Three clones of each combination were diluted in ddH_2_O, 1 µl of these diluted cells was dotted on SD/-Trp/-Leu/-His/-Ade medium and grown at 30 °C for four days. Yeast cells transformed with pGBKT7-EGT2 plasmid cannot survive on SD-Trp/-Leu/-Ade/-His medium, suggesting that EGT2 does not display autoactivation (Supporting Information Fig. S4b). Therefore, pGBKT7-EGT2 was used as bait for yeast-two-hybrid screening.

### Yeast-two-hybrid library screening

Yeast-two-hybrid library screening was performed according to the Matchmaker Gold Yeast Two-Hybrid System User Manual (Clontech).

Yeast colony PCR was performed with pGADT7 primers to eliminate duplicated positive colonies containing the same prey plasmid. Among the PCR products from all positive colonies, only DNA bands with different sizes were recovered and sequenced to identify the genes inserted. Subsequently, prey plasmids from library were rescued from yeast clones with “Easy Yeast Plasmid Isolation Kit” (Clontech) and sequenced. By doing so, all prey plasmids contained in one positive colony were identified and further tested. These prey plasmids were further used in one-on-one yeast-two-hybrid assays to confirm the results of yeast-two-hybrid screening.

### Subcellular localization

The vectors constructed using the 2in1 cloning system were used for the subcellular localization analyses (Grefen & Blatt, 2012). The full-length coding sequence of *EGT2* (*HORVU*.*MOREX*.*r3*.*5HG0447830*.*1*), *OMT* (*HORVU*.*MOREX*.*r3*.*3HG0330120*.*1*), *PBP* (*HORVU*.*MOREX*.*r3*.*7HG0709860*.*1*), *GXM* (*HORVU*.*MOREX*.*r3*.*7HG0749870*.*1*), *HMT* (*HORVU*.*MOREX*.*r3*.*2HG0126380*.*1*) and *HORVU*.*MOREX*.*r3*.*1HG0000050*.*1* were amplified using Phusion™ High-Fidelity DNA Polymerase (Thermo Fisher Scientific) with primers specified in Supporting Information Table S9.

The insertion of the *EGT2* amplicons into pDONR221 P1-P4 entry vector carrying the attP1 and attP4 recombination sites were accomplished by BP reactions using the BP ClonaseTM II enzyme mix (Thermo Fisher Scientific). The same enzyme was used for insertion of interaction candidates into pDONR221 P2R-P3 carrying the attP2 and attP3 recombination sites. Subsequently, entry clones were integrated with the corresponding destination vectors pFRETvr-2in1-NN or pFRETvr-2in1-NC. The reaction was catalyzed by LR Clonase™ II Mix (Thermo Fisher Scientific). Finally, we generated expression constructs containing the *EGT2* coding sequence either N- or C-terminally fused to tagRFP (EGT2-tagRFP and tagRFP-EGT2) and the coding sequence of interaction partners N-terminally fused to mVenus. All constructs were verified and transformed into Agrobacterium (*A. tumefaciens*) strain AGI1.

Positive Agrobacterium clones were cultured in lysogeny broth medium with respective antibiotics until OD 600 reach 0.8. Then, cells were precipitated and dissolved in 10 mM MgCl_2_, 10 mM MES (2-morpholinoethanesulfonic acid), and 100 µM AS (Acetosyringone), and infiltrated into tobacco (*Nicotiana benthamiana*) leaves using a syringe. Transgene expression was analyzed using a Zeiss LSM 780 confocal microscopy system three days after infiltration. The fluorescence of tagRFP was excited at 543 nm using a helium neon laser with a laser power of 30%, and detected through the meta-channel at 579 to 633 nm (ChS1). mVenus and chloroplast autofluorescence were excited at 488 nm by the argon laser with a 10% laser power, emission of mVenus was detected at 517 to 553 nm. The emission of chloroplast autofluorescence was detected at 686 to 711 nm via the meta-channel (ChS2).

### Bimolecular fluorescence complementation (BiFC)

The same entry vectors as used for the subcellular localization experiments were integrated with the corresponding destination vectors pBiFCt-2in1-NN or pBiFCt-2in1-NC for BiFC. Both destination vectors contained *tagRFP*, which was driven by the CaMV 35S promoter and used as a positive control for successful infiltration of tobacco cells. Finally, we generate expression constructs containing the *EGT2* coding sequence either N- or C-terminally fused to the C-terminal part of YFP (EGT2-cYFP and cYFP-EGT2) and the coding sequence of the interaction partners N-terminally fused to the N-terminal part of YFP. Constructs containing only the *EGT2* coding sequence either N- or C-terminally fused to the C-terminal part of YFP were used to detect the autofluorescence of EGT2. Constructs containing the coding sequence of interaction partners N-terminally fused to the N-terminal part of YFP and coding sequence of *EGT2* (with stop codon) N-terminally fused to the C-terminal part of YFP were used to detect the autofluorescence of interaction partners. All constructs were verified and transformed into Agrobacterium strain AGI1. Tobacco infiltration and transgene expression analysis was conducted as described above. Excitation and detection of RFP, YFP (similar to mVenus) and chloroplast autofluorescence were performed as described above.

## Results

### Transcriptomic dynamics of three root zones of wild type and *egt2* seminal roots upon gravistimulation

We investigated the transcriptomic dynamics in the root cap, the meristem and the elongation zone (Fig. 1a) in roots of wild type seedlings and the hypergravitropic mutant *egt2* at 0, 3, 6 and 12 h after rotation by 90° (Fig. 1b) by RNA-seq (see methods). We explored the transcriptomic relationship between genotypes, root zones and duration of gravistimulation by a principal component analysis (PCA; Fig. 1c). In the PCA plot, two components (PC1 and PC2) accounted for 91% of the total variance. The first component PC1 explained 74% of the overall variance and distinctly separated the samples of each root zone, indicating larger transcriptomic differences between root zones than between genotypes and time points (Fig. 1c).

### Determination of genes responding to gravistimulation in seminal root zones

To identify genes responding to gravistimulation in barley roots, three pairwise comparisons were computed between control (0 h) and gravistimulated (3 h, 6 h and 12 h) root samples for each of the three zones of wild type roots. We determined differentially expressed genes (DEGs) by using a false discovery rate (FDR) <5% (Fig. 2a). In all three zones, longer gravitropic stimulation resulted in more DEGs (Fig. 2a and Supporting Information Table S1). The lowest number of identified genes at each time point was observed in the meristem (Fig. 2a). Only five genes were differentially regulated in the meristem (M) 12 h after rotation (Fig. 2a). Of 462 distinct DEGs in the root cap (RC), none was differentially expressed at all three time points after gravistimulation (Fig. 2b). Similarly, of 3173 distinct DEGs in the elongation zone (E), only the 7 genes, that were already induced 3 h after treatment were consistently regulated at all three time points (Fig. 2c). This suggests for both root zones, that most gravistimulated genes are specifically stimulated after a certain period of gravitropic bending (Fig. 2b and 2c). By investigating the expression dynamics of genes that or whose homologs have been reported to play a role in regulating root gravitropism, we found that *HORVU*.*MOREX*.*r3*.*2HG0125600*.*1*, a homolog of *AtLZY2* and *AtLZY3* (Taniguchi *et al*., 2017), and *HORVU*.*MOREX*.*r3*.*5HG0509060*.*1*, a homolog of *ZmCIPK15* (Schneider *et al*., 2022) were significantly down-regulated in the root cap after 12 h of gravistimulation (Supporting Information Table S1).

**Fig. 2:**
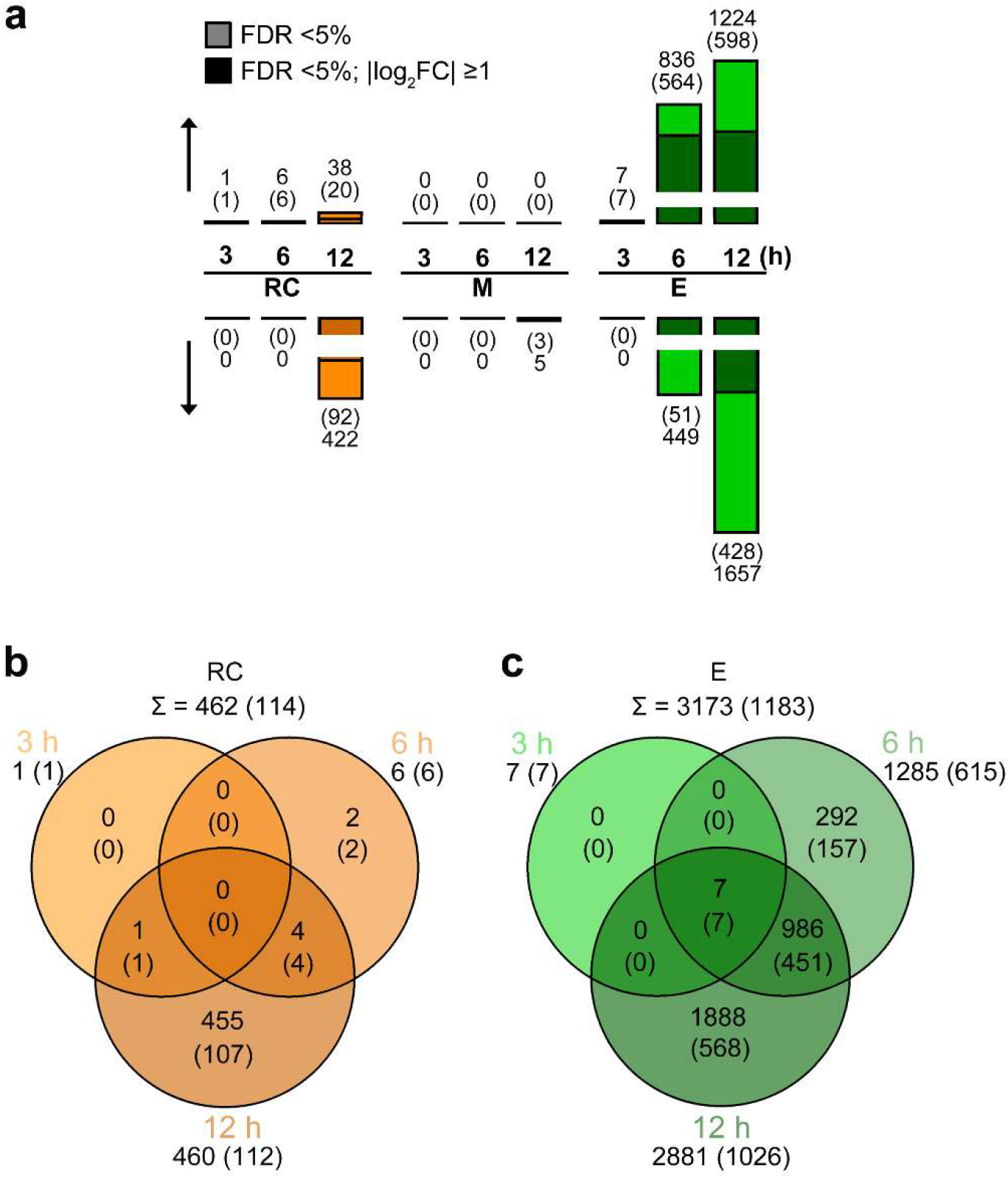
Differential gene expression in gravistimulated wild type roots. **a** Numbers of up- (↑) and down-regulated (↑) genes determined by three pairwise comparisons between the root zones of wild type vertically grown roots and the corresponding zone of wild type gravistimulated roots. RC: root cap. M: meristem, E: elongation zone (light color & numbers without brackets: false discovery rate (FDR) <5%; dark color & numbers in brackets: FDR <5%, log_2_ fold change (|log_2_FC|) ≥1). **b-c**. Venn diagram of differentially regulated genes by gravistimulation in wild type root cap (b) and elongation zone (c) (numbers without brackets: FDR <5%; numbers in brackets: FDR <5%, |log_2_FC| ≥1).

### Expression dynamics of DEGs regulated by gravistimulation in seminal roots

To group genes with similar expression patterns over 12 h of rotation, we performed hierarchical clustering of all DEGs in the root cap and the elongation zone of wild type roots. Hierarchical clustering classified 462 DEGs in the root cap into three clusters (Supporting Information Fig. S1a-b and Supporting Information Table S2) and 3173 DEGs in the elongation zone into nine clusters (Supporting Information Fig. S1c-d and Supporting Information Table S3) according to different patterns of up- and down-regulation in the course of gravistimulation.

Gene Ontology (GO) term analyses assigned a variety of biological functions to the genes enriched in each cluster (Supporting Information Table S4). A number of enriched biological process terms related to cell wall were shared between different clusters in the elongation zone. For instance, the term ‘plant-type cell wall organization’ (GO:0009664); ‘xyloglucan metabolic process’ (GO:0010411); ‘fucose metabolic process’ (GO:0006004); and ‘cellulose biosynthetic process’ (GO:0030244) were shared between several clusters (Supporting Information Table S4b). In addition, the terms ‘response to oxidative stress’ (GO:0006979), ‘hydrogen peroxide catabolic process’ (GO:0042744) and ‘protein phosphorylation’ (GO:0006468) were significantly enriched in multiple clusters (Supporting Information Table S4b).

### Determination of DEGs between gravistimulated roots of *egt2* and wild type

Differential genes expression between *egt2* and wild type was determined for each root zone by time point combination by computing twelve pairwise contrasts (Fig. 3a and Supporting Information Table S5). The root cap displayed 126 (Fig. 3b), the meristem 62 (Fig. 3c) and the elongation zone 2982 unique DEGs (Fig. 3d). In comparison with the wild type, *EGT2* was down-regulated in all three root zones of the *egt2* mutant at all four time points (Supporting Information Table S5).

**Fig. 3:**
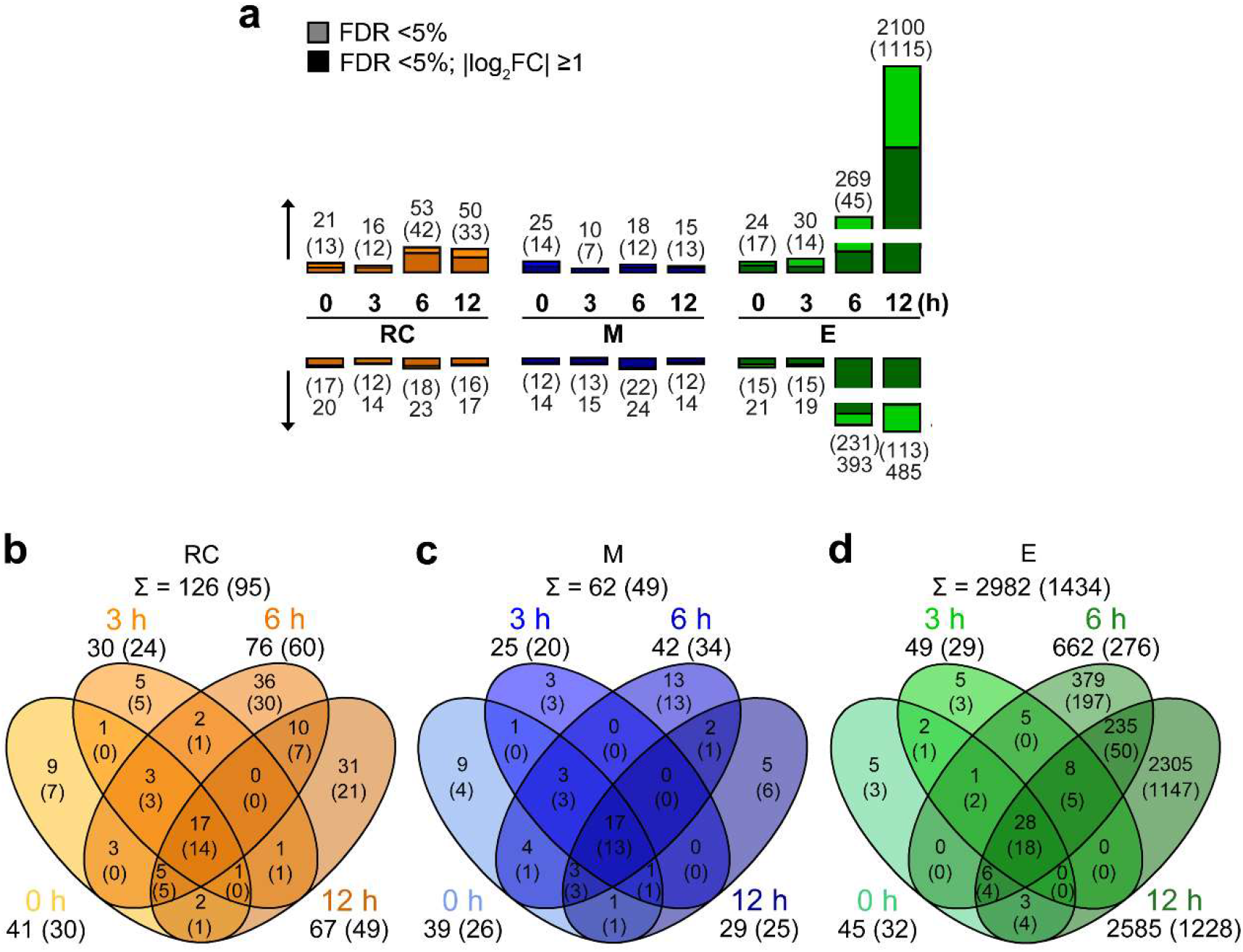
Differential gene expression between *egt2* mutant and wild type. **a** Numbers of up- (↑) and down-regulated (↓) genes identified by pairwise comparisons between the *egt2* mutant and wild type for each of the three root zones: root cap (RC), meristem (M) and elongation zone (E) at each of the four time points: 0 h, 3 h, 6 h and 12 h (light color & numbers without brackets: FDR <5%; dark color & numbers in brackets: FDR <5%, |log_2_FC| ⩾1). **b-d** Overlap of DEGs between the four time points in the root cap (b), the meristem (c) and the elongation zone (d) of *egt2* vs wild type (numbers without brackets: FDR <5%; numbers in brackets: FDR <5%, |log_2_FC| ≥1).

GO analyses revealed that 31 biological process terms were assigned to DEGs in the root cap (Supporting Information Table S6a). Among them, ‘cytoplasmic microtubule organization’ (GO:0031122), ‘trichome branching’ (GO:0010091) and ‘plant-type cell wall biogenesis’ (GO:0009832) were conserved among up-regulated genes at all four time points (Supporting Information Table S6a). In the meristem, 17 GO terms were enriched (Supporting Information Table S6b). As in the root cap, the GO terms ‘cytoplasmic microtubule organization’ (GO:0031122), ‘trichome branching’ (GO:0010091) and ‘plant-type cell wall biogenesis’ (GO:0009832), were conserved among up-regulated genes at all four time points in the meristem. In addition, ‘cell growth’ (GO:0016049) was conserved among up-regulated genes while ‘one-carbon metabolic process’ (GO:0006730) was conserved among down-regulated genes at all four time points (Supporting Information Table S6b). Finally, we identified 113 significantly enriched GO terms in the DEGs of the elongation zone (Supporting Information Table S6c). Among those, five GO terms were conserved among up-regulated genes at 0, 3, and 6 h, while six GO terms were enriched among down-regulated genes at 0 and 3 h, and among up-regulated genes at 12 h (Supporting Information Table S6c).

### Identification of genes regulated by gravity and by *EGT2*

To screen for genes, which are regulated by gravistimulation and *EGT2*, we compared DEGs responding to gravitropic stimulus with DEGs determined in *egt2* vs wild type comparisons. We observed 13 intersecting genes in the root cap (Fig. 4a) and 1038 genes in the elongation zone (Fig. 4b). These genes are listed in Supporting Information Table S7. Among these genes, twelve of 13 in the root cap and 1027 of 1038 in the elongation zone showed opposite regulatory directions in wild type time-course experiments and in *egt2* vs wild type comparisons. No *EGT2*-regulated gene responding to gravistimulation was identified in the meristem. Of the gravity-regulated genes, 3% in the root cap and 33% in the elongation zone were also regulated by *EGT2*.

**Fig. 4:**
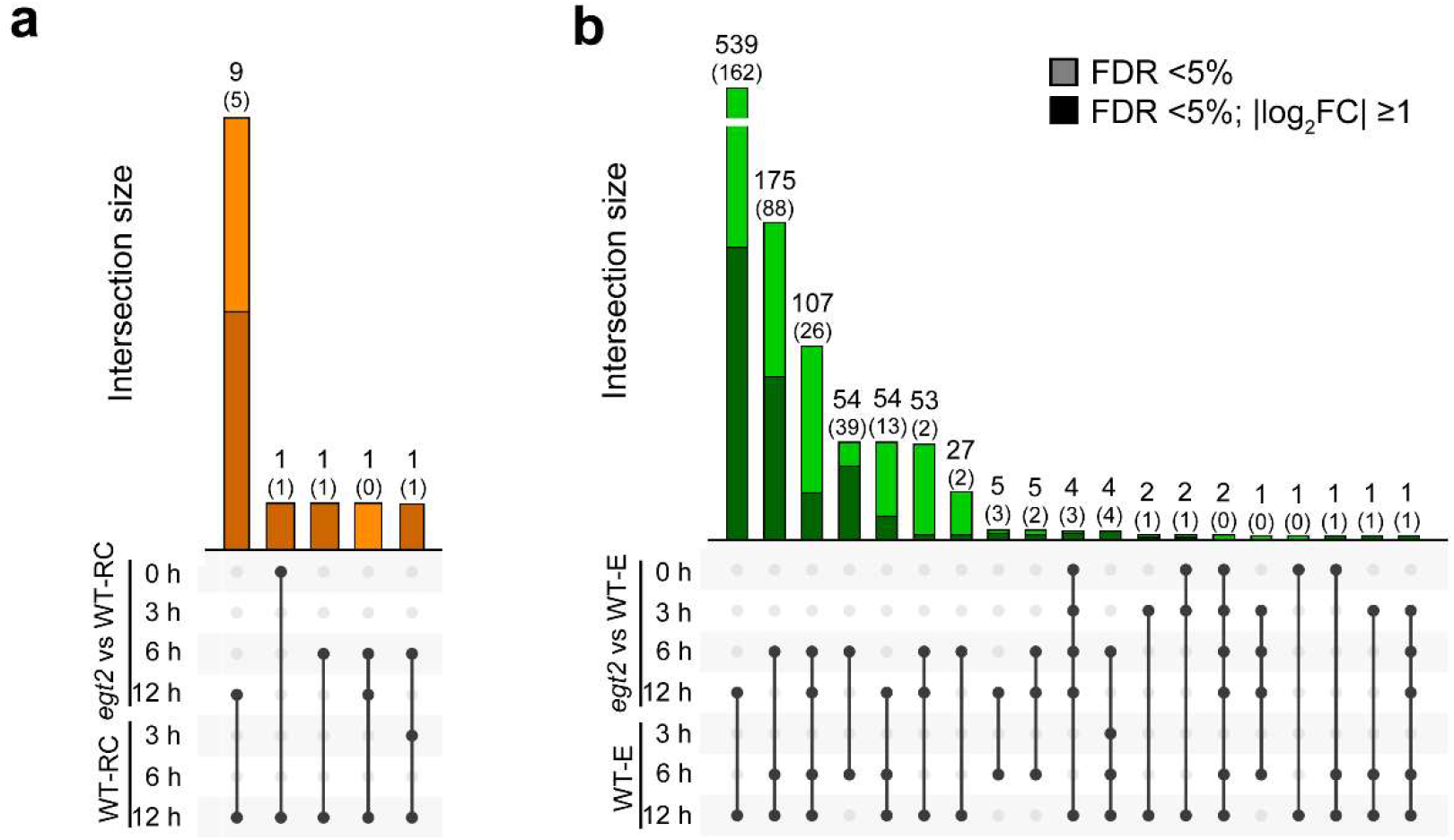
Intersections between differentially expressed genes (DEGs) in the wild type time-course experiment and in *egt2* vs wild type comparisons. **a-b** Overlap of DEGs in the root cap (a) and elongation zone (b) of gravistimulated roots of the wild type time-course and *egt2* vs wild type comparisons (numbers without brackets: FDR <5%; numbers in brackets: FDR <5%, |log_2_FC| ≥1).

We conducted GO analyses for these intersected genes to reveal the biological processes in which *EGT2* might be involved to regulate root gravitropism. Three protein transmembrane transport-related and two mitochondrion-related GO terms were significantly enriched in intersected DEGs in the root cap (Supporting Information Fig. S2a). Of the 26 GO terms significantly enriched among the overlapping genes in the elongation zone, seven terms were associated with plant cell wall organization, four terms were related to protein processing, two terms were ROS-related and two terms were associated with carbohydrate metabolic processes (Supporting Information Fig. S2b).

### Identification of co-expression networks and their association with the traits genotype and time of gravistimulation

To reveal additional genes that are significantly associated with *EGT2* and gravistimulation in the root cap, meristem and elongation zone of barley seminal roots, we performed a weighted gene co-expression network analysis (WGCNA). We constructed a co-expression network for the RNA sequencing data separately for each root zone to eliminate the effect of the large expression differences between the distinct root zones on the following analyses. This analysis identified 13 co-expression modules in the root cap, 21 in the meristem, and 9 in the elongation zone (Supporting Information Fig. S3a-c). These modules were then correlated with the two genotypes (wild type and *egt2*) and the time of gravistimulation (0, 3, 6 and 12 h). This resulted in 9 modules that are strongly correlated (*) with at least one trait in the root cap (Fig. 5a), 14 in the meristem (Fig. 5b) and 6 in the elongation zone (Fig. 5c) with the module size ranging from 38 to 11791 genes.

**Fig. 5:**
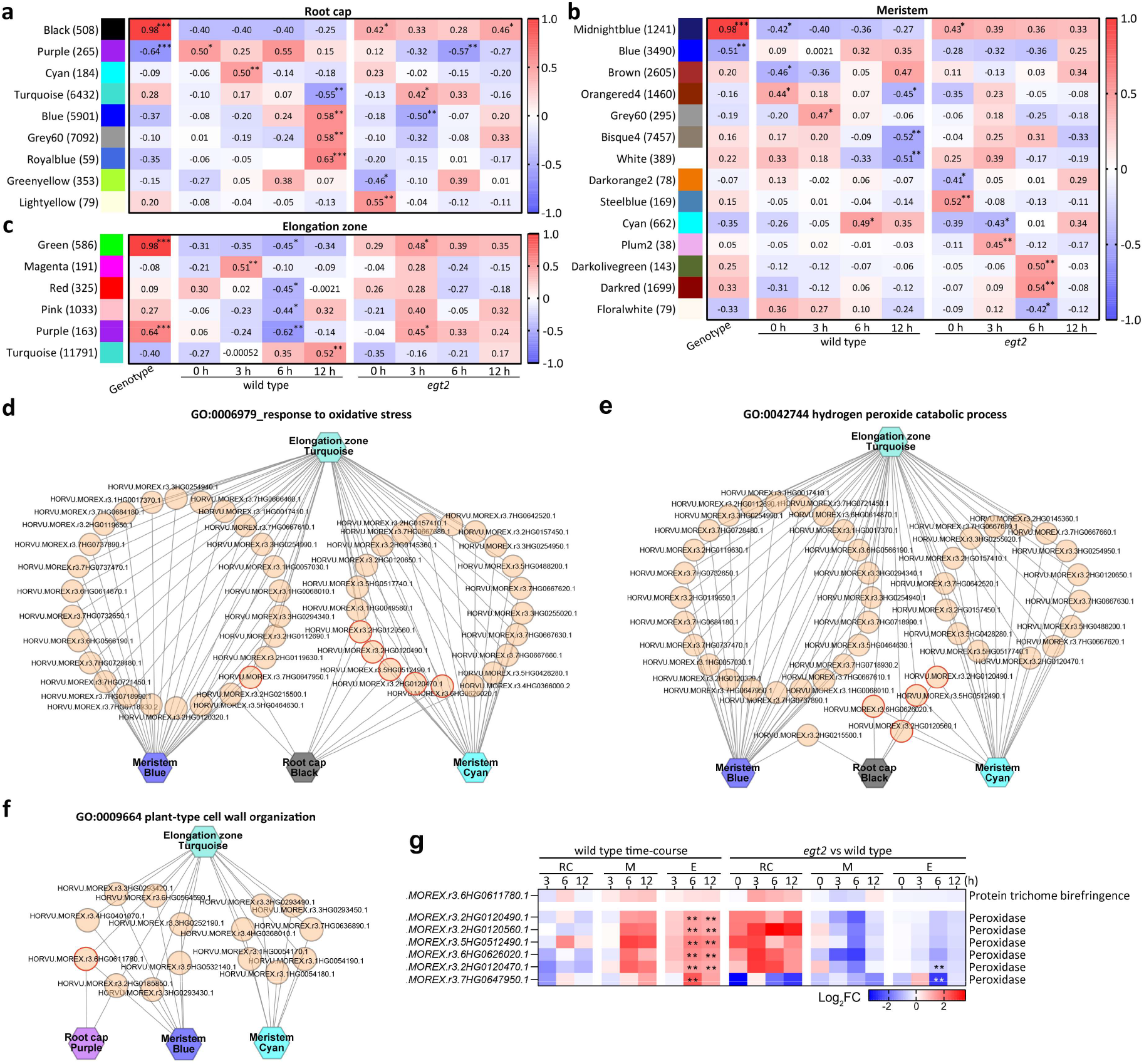
Module-trait associations and genes assigned to plant cell wall and reactive oxygen species (ROS)-related processes in each selected module. **a-c** Module-trait relationships in the root cap (a), the meristem (b) and the elongation zone (c). The genotypes (WT and *egt2*) and time points (0, 3, 6 and 12 h) after gravistimulation are used as traits, each column corresponds to a different trait. Each row corresponds to the characteristic genes of the module. The relationship between the modules and traits is indicated in cell by Pearson correlation coefficients. Asterisks indicate significant values. *, *p* <0.05; **, *p* <0.01; ***, *p* <0.001. Cell color ranges from red (highly positive correlation) to blue (highly negative correlation). The number of genes contained in each module is indicated in brackets next to the module names. **d** Genes associated with ‘response to oxidative stress’. **e** Genes associated with ‘hydrogen peroxide catabolic process’. **f** Genes associated with ‘plant-type cell wall organization’. **d-f** Hexagonal nodes represent modules, circle nodes in orange represent individual genes. Nodes encircled in dark orange represent genes determined in all three root zones. **g** The expression patterns of genes encircled in dark orange in d-f in the wild type gravistimulation time-course experiments (wild type time-course) and in the pairwise comparisons of *egt2* with wild type (*egt2* vs wild type). **, FDR <5%.

We determined two modules that were significantly correlated with genotype, and modules that displayed contrasting correlation values between wild type and *egt2* at some time points in all three root zones (Fig. 5a-c). We performed GO analyses to identify overrepresented biological processes among genes in the three most strongly correlated modules in each root zone. The five most significantly enriched biological process terms in each module are shown in Supporting Information Fig. S3d. Noticeably, ‘plant-type cell wall organization’ (GO: 0009664), ‘response to oxidative stress’ (GO: 0006979) and ‘hydrogen peroxide catabolic processes’ (GO:0042744) were consistently identified in modules that significantly correlated with genotype in all three root zones (Supporting Information Fig. S3d). We compared genes associated with plant cell wall and ROS-related processes between the three root zones (Fig. 5d-f). Results showed that *HORVU*.*MOREX*.*r3*.*6HG0611780* and six genes encoding peroxidases were included in significantly correlated modules in all three root zones (Fig. 5d-f). All six genes encoding peroxidases were regulated by gravistimulation in the elongation zone, two of which were regulated by *EGT2* (Fig. 5g).

The module hub genes are defined as genes that high connected with other genes in the module, and have been shown to be closely correlated with biological processes. We determined hub genes in the three most strongly correlated modules in each root zone and surveyed their expression levels in wild type gravistimulation time-course experiments and in *egt2* vs wild type comparisons (Supporting Information Fig. S3e-g). Among them, *EGT2* was considered as a hub gene in the midnightblue module in the meristem (Supporting Information Fig. S3f). All seven hub genes identified in this module displayed an *EGT2*-regulated while gravity-independent expression pattern (Supporting Information Fig. S3f). Hub genes identified in selected modules in the elongation zone were regulated by either gravity or *EGT2* in the elongation zone, seven of them were regulated by gravity and *EGT2* (Supporting Information Fig. S3g).

### Identification of the interaction partners of EGT2

To determine whether some of these DEGs are encoding for interaction partners of EGT2, we performed a yeast-two-hybrid screening. Two yeast-two-hybrid screens yielded 79 putative interaction partners of EGT2 that were confirmed by Sanger sequencing (Supporting Information Fig. S4c and Supporting Information Table S8). To validate this result in yeast cells, we tested the interaction between EGT2 and 23 putative interaction candidates by a one-on-one yeast-two-hybrid assay. The 23 interaction candidates included 21 candidates that were captured more than twice and two candidates encoded by DEGs in *egt2* vs wild type comparisons (Supporting Information Fig. S4c). All 23 selected interactions were positive in yeast cells (Supporting Information Fig. S4d), which confirmed the validity of the yeast-two-hybrid results.

Of all genes encoding putative interaction candidates, three genes were differentially expressed in the root cap and 24 in the elongation zone of gravity-stimulated wild type roots. Moreover, 16 genes were differentially expressed in the elongation zone of the *egt2* mutant compared with wild type (Supporting Information Fig. S4e and Supporting Information Table S8). Among those, genes encoding eight interaction partners were regulated by gravistimulation and *EGT2* (Supporting Information Table S8). Comparisons between the expected (= number of interaction candidates : number of annotated gene models * number of DEGs) and the observed number of interaction partners in each dataset showed that more interaction partners were observed in the elongation zone of wild type roots than expected after 6 h and 12 h of rotation (Supporting Information Fig. S4e).

Five of eight interaction partners encoded by gravity and *EGT2*-regulated genes were independently surveyed via BiFC (Supporting Information Table S8). We first investigated the subcellular localization of EGT2 and the five interaction partners. Results revealed that EGT2 and the four interaction candidates, HORVU.MOREX.r3.3HG0330120.1 (OMT), HORVU.MOREX.r3.7HG0709860.1 (PBP), HORVU.MOREX.r3.7HG0749870.1 (GXM) and HORVU.MOREX.r3.2HG0126380.1 (HMT) localized to the cytoplasm and the nucleus (Fig. 6a: OMT, GXM and HMT). However, we were not able to detected the subcellular localization signal of HORVU.MOREX.r3.1HG0000050.1. For the BiFC assay, the four interaction candidates that also co-localized with EGT2 were fused with the N-terminal fragment of YFP, while EGT2 was fused with the C-terminal fragment of YFP. Yellow fluorescence signals emitted by the interactions were detected for the proteins OMT, GXM and HMT, but not for the protein PBP (Fig. 6b: OMT, GXM and HMT). We did not detect a yellow fluorescence signal in tobacco epidermal cells infiltrated with constructs containing only the coding sequence of EGT2 fused to the C terminal part of YFP. Nor did we detect YFP signal in tobacco epidermal cells infiltrated with constructs containing coding sequence of interaction partners fused to the N-terminal part of YFP and full-length EGT2 N-terminally fused to the C-terminal part of YFP (Supporting Information Fig. S5).

**Fig. 6:**
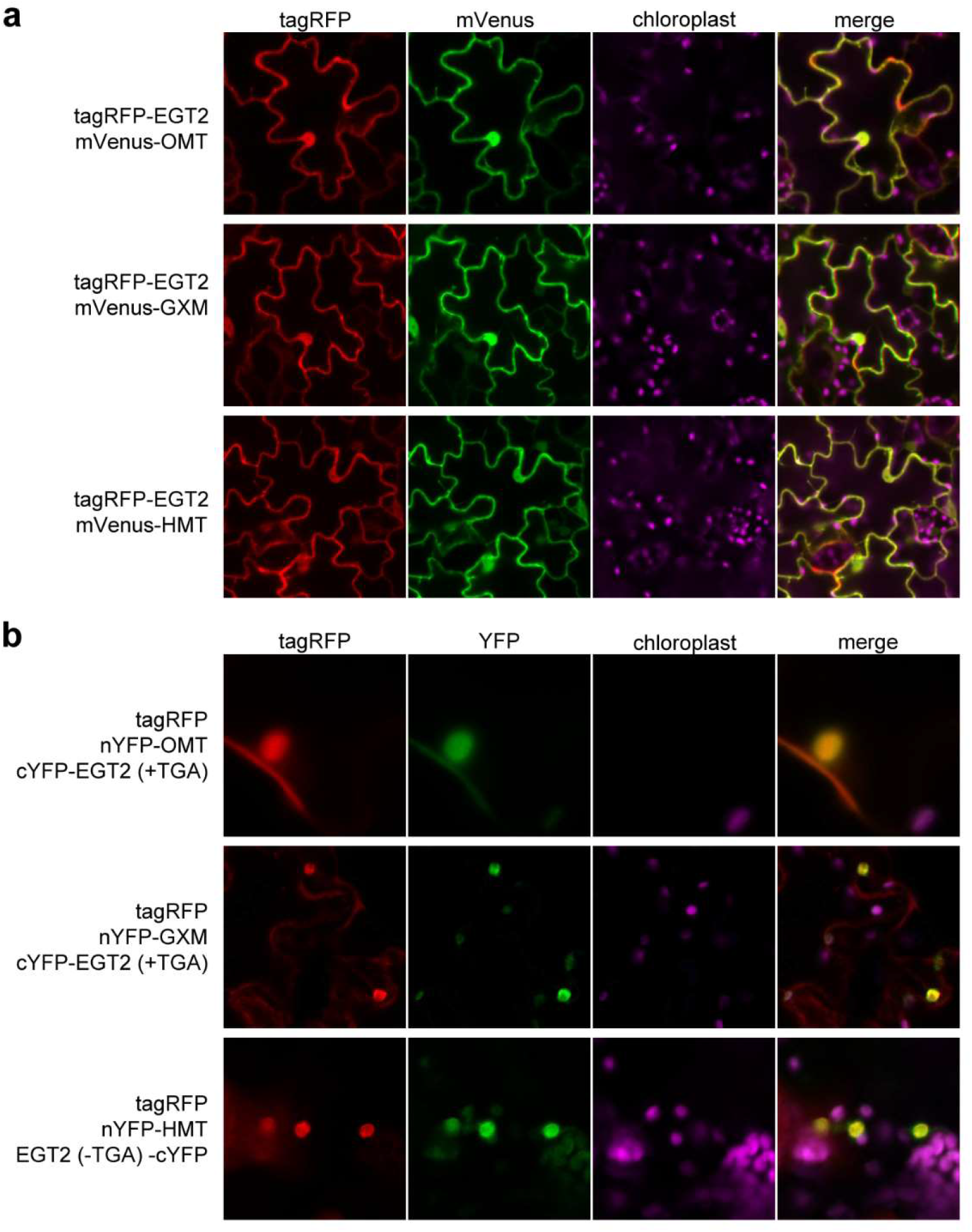
Confirmation of interactions between EGT2 and other proteins. **a** Subcellular localization assay of EGT2 and interaction partners. tagRFP: red fluorescence; mVenus: yellow fluorescence. **b** BiFC assay of EGT2 and interaction partners. tagRFP: red fluorescence; mVenus: yellow fluorescence; nYFP: N-terminal part of YFP; cYFP: C-terminal part of YFP. EGT2 (+TGA): full-length coding sequence of EGT2; EGT2 (-TGA): coding sequence of EGT2 removed stop codon. **a-b** OMT: HORVU.MOREX.r3.3HG0330120.1; GXM: HORVU.MOREX.r3.7HG0749870.1; HMT: HORVU.MOREX.r3.2HG0126380.1; chloroplasts: auto-fluorescence of chloroplasts.

## Discussion

Gravitropism modulates root system architecture by controlling the distribution of roots in different soil layers and thus their ability to absorb water and nutrients. Understanding the molecular mechanisms underlying root gravitropism will help to develop root ideotypes that can improve crop performance under unfavorable environmental conditions.

The root gravitropic response includes the perception of gravity signals mainly in the root cap, signal transduction through the meristem and signal execution translating into a bending response in the elongation zone (Su *et al*., 2017; Singh *et al*., 2017; Nakamura *et al*., 2019). We observed that the angle adjustment of wild-type barley roots was adjusted by 10° after 3 h of rotation and reached 30° (>60% of the total adjusted angle) after 12 h of rotation (Fig. 1b; Kirschner *et al*., 2021). In this study, we performed RNA-seq of the root cap, the meristem and the elongation zone of wild-type barley roots before gravistimulation and 3, 6 and 12 h after reorientation of the roots by 90°. This experiment revealed transcriptomic variation in every step of gravitropic response at different time points. First, only a few DEGs were observed after 3 h of rotation in all three zones (Fig. 2a). This indicates that gravity perception and gravitropic bending caused only little transcriptome adjustment at early stages of the experiment (Fig, 2b and 2c). Secondly, the elongation zone displayed by far the most DEGs (Fig, 2a and 2c), suggesting that gravitropic bending is subject to a more extensive transcriptomic regulation in the elongation zone than gravity perception in the root cap and signal transduction in the meristem. GO term analyses revealed the enrichment of a number of biological processes among the DEGs in the gravistimulated root cap and elongation zone. GO terms related to ROS biosynthesis, cytoplasmic microtubule organization, plant transmembrane transport, carbohydrate metabolic and calcium-mediated signaling were over-represented in down-regulated genes in root cap (Supporting Information Table S4a). These enriched GO terms are in line with the notion that ROS (Krieger *et al*., 2016), starch-related processes (Li *et al*., 2020a), dynamic actin-filament network (Hou *et al*., 2003) and calcium (Bizet *et al*., 2018) are involved in gravity sensing, which occurs primarily in root cap (Su *et al*., 2020). In addition, the homolog of *AtLZY2* and *AtLZY3* and the homolog of *ZmCIPK15* were significantly down-regulated in the root cap after 12 h of gravistimulation (Supporting Information Table S1). This is consistent with the role of *AtLZY2* and *AtLZY3* in root columella cells during gravitropic response and the function of *ZmCIPK15* in regulating maize root angles (Taniguchi *et al*., 2017; Schneider *et al*., 2022). In the elongation zone, seven cell wall, ROS and protein phosphorylation-related terms were assigned to more than three gene clusters with differential gene expression patterns (Supporting Information Table S4b). This is consistent with the notion that cell wall-related processes are closely related to asymmetric cell elongation in the distal elongation zone of gravistimulated roots (Su *et al*., 2017; Lešková *et al*., 2020). It has also been demonstrated that gravistimulation induces an asymmetric distribution of ROS in the distal elongation zone of the root (higher in the lower flanks), which promotes root gravitropic bending (Krieger *et al*., 2016). Moreover, ROS production is also involved in regulating cell wall loosening and cell expansion (Liszkay *et al*., 2004; Xiong *et al*., 2015; Mabuchi *et al*., 2018). Protein phosphorylation functions in mediating auxin biosynthesis, transport and signaling, therefore playing a critical role in many auxin-related processes, including root gravitropism (Tan *et al*., 2021).

In a previous study, we identified the barley mutant *egt2*, which shows enhanced root gravitropism (Kirschner *et al*., 2021). RNA-seq of vertically grown roots showed that cell wall-related processes were affected in the mutant, however, nothing is known about the dynamics of the gravity response in regard to the role of EGT2 (Kirschner *et al*., 2021). Here, we determined DEGs between different root zones of the gravistimulated roots of *egt2* and wild type in a time-course experiment (Fig. 3). In the root cap, only 3% of gravity-responsive genes were regulated by *EGT2* (Fig. 4a), implying that *EGT2* might not play an important role in gravity perception in the root cap. This is in line with our previous observation that there was no significant difference in amyloplast content and their sedimentation rates between wild type and *egt2* (Kirschner *et al*., 2021). Similarly, there was no overlap between gravity-responsive genes and genes regulated by *EGT2* in the meristem suggesting not prominent role of *EGT2* in gravity signal transduction on the transcriptomic level. In our previous study, we suggested that *EGT2* mediates gravity signal transduction at the protein level, further regulating the targets in the elongation zone to control the root gravitropism (Kirschner *et al*., 2021). Consistently, in the elongation zone of *egt2*, gravistimulation induced the differential expression of thousands of *EGT2*-regulated genes (Fig. 3a and 3d). Intriguingly, >1000 *EGT2*-regulated genes in the elongation zone were also regulated by gravistimulation in the wild type roots (Fig. 4b). This implies that around one-third of gravity-responsive genes in the elongation zone of wild type roots were regulated by *EGT2*, suggesting a critical role of *EGT2* in regulating root gravitropic bending. Plant cell wall organization and ROS-related GO terms that were assigned to these intersected genes in the elongation zone (Supporting Information Fig. S2b) support this hypothesis.

Co-expression network analyses revealed that specific modules that have a strong correlation with genotype in all three root zones. These modules were in most cases contrastingly correlated with the gravitropic response of wild type and *egt2* (Fig.5a-c and Supporting Information Fig.S3a-c). Genes in these modules are potential components involved in regulating gravitropic response and anti-gravitropic offset (AGO), which counteracts gravitropism to regulate the root growth direction. (Supporting Information Fig. S3e-g; Kawamoto *et al*., 2020; Fusi *et al*., 2022). Moreover, modules correlated only with wild type or *egt2* at some time point after gravistimulation might be involved in *EGT2*-independent or *EGT2*-dependent gravitropism, respectively. This result complements differential expression analyses, in particular for the root cap and the meristem where fewer gravity or *EGT2*-regulated genes were discovered (Fig. 2 and 3). In the elongation zone, many hub genes of significantly correlated modules were regulated by gravity and/or *EGT2* (Supporting Information Fig. S3g), which is in line with the differential expression analyses. Plant cell wall and ROS-related GO terms were assigned to modules strongly correlated with genotype in all three root zones (Supporting Information Fig.S3d). This is consistent with GO analyses of gravity and *EGT2*-regulated genes in the elongation zone (Supporting Information Fig.S2b), suggesting that plant cell wall and ROS-related processes might be involved in *EGT2*-controlled root gravitropic response. Moreover, six genes encoding peroxidases were present in significantly correlated modules in all three root zones (Fig. 5g). These results suggest that gravitropism regulation of *EGT2* might be similar to that of *EGT1*, which was proposed to control cell wall stiffness by regulating peroxidase and cell wall loosening-related enzymes, therefore mediating AGO (Fusi *et al*., 2022).

Localization of EGT2 in the cytoplasm and nucleus was consistent with the predicted localization of its homologs in Arabidopsis (Denay *et al*., 2017). As the only annotated domain in EGT2, the SAM domain is known as a protein-protein interaction domain in animals (Kim & Bowie, 2003). Recent studies on plant SAM domain-containing proteins have provided initial insights into the interaction capabilities of the plant SAM domain (Sayou *et al*., 2016; Murcha *et al*., 2016). In the current study, we identified 79 proteins as putative interaction partners of EGT2 (Supporting Information Fig. S4c). The number of observed interaction candidates that responded to gravistimulation on basis of the transcriptome analyses was significantly higher than expected by chance (Supporting Information Fig. S4e), highlighting the role of EGT2 in root gravitropic response.

BiFC confirmed the interaction between EGT2 and three interaction candidates encoded by DEGs (Fig. 6). Two candidates were not confirmed, probably due to changes in protein structure caused by inappropriate terminal fusion to fluorescent proteins. Alternatively, the two proteins might be false-positive results of the yeast-two-hybrid experiment (Xing *et al*., 2016). None of the three independently confirmed proteins has already been functionally characterized in barley. OMT, annotated as ‘O-methyltransferase’, shows a high degree of identity with TaOMT2, which displays a broad substrate preference and high catalytic efficiency towards flavonoids and the aldehyde precursors of S lignin (Zhou *et al*., 2006; Ma & Xu, 2008). In addition to lignin, which is an important component of cell walls, flavonoids have been shown to play a role in root gravitropism by redirecting polar auxin fluxes (Santelia *et al*., 2008). GXM is annotated as ‘Glucuronoxylan 4-O-methyltransferase’ and is classified in the Pfam database as a member of the ‘Polysacc_synt_4 (PF04669)’ protein family. Proteins in this family play an essential role in xylan deposition and lignin assembly in plant cell walls (Brown *et al*., 2011; Jensen *et al*., 2011; Urbanowicz *et al*., 2012). The direct interactions of EGT2 with OMT and GXM suggest a potential role of EGT2 in cell wall organization and also imply a role of EGT2 in flavonoid-related processes. HMT is a heavy metal transport/detoxification superfamily protein, also known as ‘copper transport protein atox1-related’. Its homologs in Arabidopsis have been shown to respond to many types of heavy metal stress, including Cu^2+^ (Li *et al*., 2020b). The inhibitory effect of Cu^2+^ treatment on root lignin metabolism and elongation has already been studied in several species (Printz *et al*., 2016). However, the function of EGT2 in cell wall and flavonoid-related processes remains to be investigated.

## Abbreviation

Abbreviation: Definition
AGO: anti-gravitropic offset
BiFC: bimolecular fluorescence complementation
DEGs: differentially expressed genes
FDR: false discovery rate
GO: Gene Ontology
log_2_FC: log_2_ fold change
PCA: principal component analysis
RNA-seq: RNA sequencing
ROS: reactive oxygen species
SAM: sterile alpha motif
WGCNA: weighted gene co-expression network analysis

## Funding information

This work was funded by the Deutsche Forschungsgemeinschaft (DFG) grant HO2249/21-1 (to F.H.).

## Acknowledgments

We thank Andreas Meyer (University of Bonn) and Peng Yu (University of Bonn) for their discussions and suggestions on this project. Many thank also to Helmut Rehkopf (University of Bonn) and Christa Schulz (University of Bonn) for their experimental support of this study.

## Authors Contributions

L.G. performed the experiments, analyzed data and drafted the article. A.K. performed bioinformatic analyses of RNA-seq data. M.B. was involved in the yeast-two-hybrid experiments. G.K.K. and S.S. participated in data interpretation and revised the article. F.H. conceived and coordinated the study, participated in data interpretation and drafting the article. All authors edited the article and approved the final draft. The authors declare no competing interest.

## Data availability

RNA-seq data have been deposited in the SRA under accession number PRJNA858009 (https://www.ncbi.nlm.nih.gov/bioproject/858009).

## Supporting Information

Additional supporting information may be found in the online version of this article.

**Fig. S1**: Dynamics of the expression profiles of graviregulated genes.

**Fig. S2**: Enriched GO terms for gravity regulated genes that are *EGT2* related.

**Fig. S3**: Co-expression analysis.

**Fig. S4**: Overview of yeast-two-hybrid screening.

**Fig. S5**: Control experiments for bimolecular fluorescence complementation analyses of EGT2 and interaction candidates.

**Fig. S6**: Quantification of meristem length of wild type (WT) and *egt2* before and after 6 h or 10 h of gravistimulation.

**Table S1**: Overview of differentially expressed genes (FDR <5%) in gravistimulated wild type roots in a time-course experiment.

**Table S2**: Hierarchical clustering analysis of differentially expressed genes (FDR <5%) in the root cap of gravistimulated wild type roots.

**Table S3a**: Enriched biological process terms among differentially expressed genes in the root cap (RC) of gravistimulated wild type roots.

**Table S3b**: Enriched biological process terms among differentially expressed genes in the elongation zone (E) of gravistimulated wild type roots.

**Table S4**: Hierarchical clustering analysis of differentially expressed genes (FDR <5%) in elongation zone of gravistimulated wild type roots.

**Table S5**: Overview of genes differentially expressed between wild type and *egt2* (FDR <5%) in a gravistimulation time-course experiment.

**Table S6a**: Enriched biological processes terms among genes differentially expressed between wild type and *egt2* (FDR <5%) root caps after a gravistimulation time-course experiment.

**Table S6b**: Enriched biological processes terms among genes differentially expressed between wild type and *egt2* (FDR <5%) meristems after a gravistimulation time-course experiment.

**Table S6c**: Enriched biological processes terms among genes differentially expressed between wild type and *egt2* (FDR <5%) elongation zones after a gravistimulation time-course experiment.

**Table S7**: Overlapping genes among differentially expressed genes in the wild type time-course experiment and *egt2* vs wild type comparisons.

**Table S8**: Overview of the candidates interacting with EGT2 as identified by yeast-two-hybrid screening.

**Table S9**: List of oligonucleotide primers.

## Notes

### Competing Interest Statement

The authors have declared no competing interest.

