## Supporting Information Fig.S1 for "ENHANCED GRAVITROPISM 2 coordinates molecular adaptations to gravistimulation in the elongation zone of barley roots"

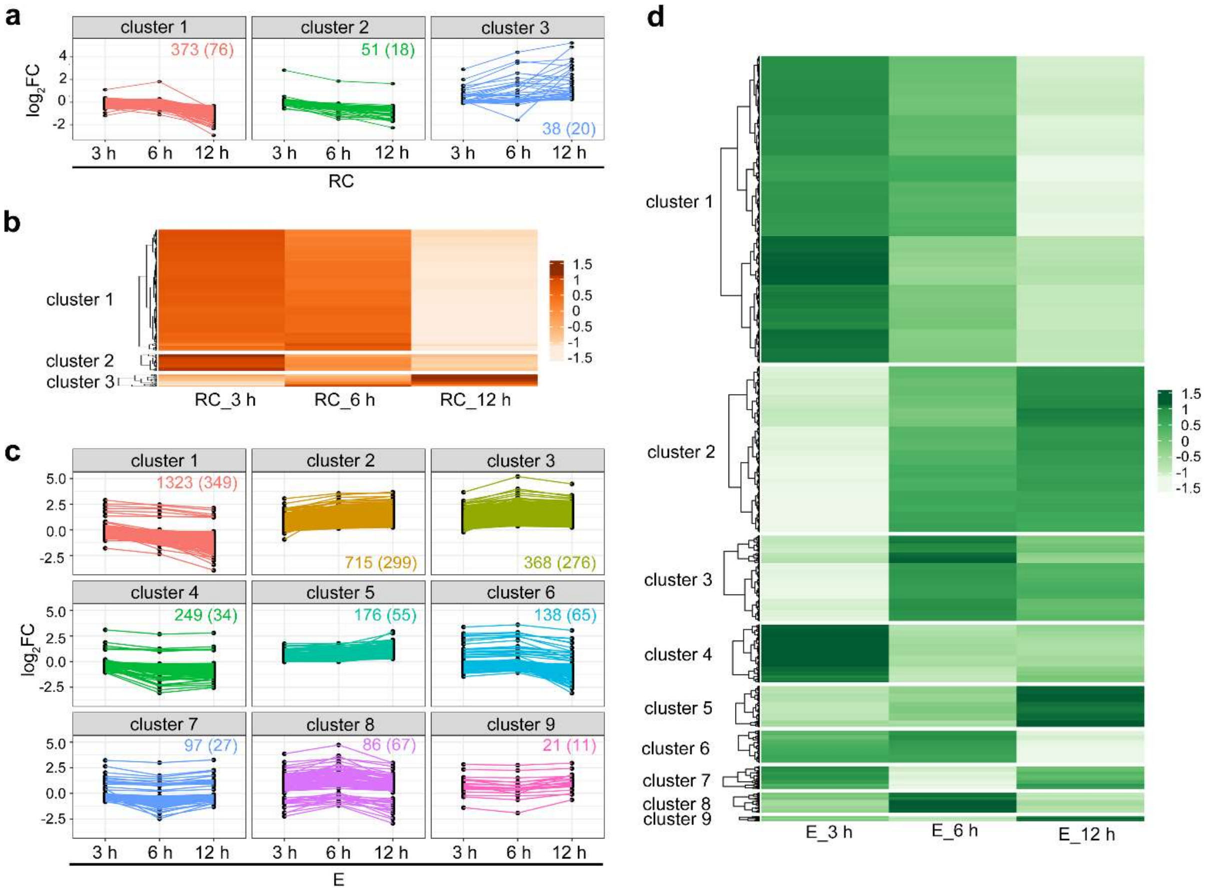


**Fig. S1: Dynamics of the expression profiles of graviregulated genes.**

**a-d** Hierarchical clustering analyses of differentially expressed genes in root cap (a and b) and elongation zone (c and d) of gravistimulated wild type roots.

**a** and **c** Observed expression patterns of differentially expressed genes during 12 h of gravistimulation. Numbers of genes assigned to each cluster are shown (numbers without brackets: FDR <5%; numbers in brackets: FDR <5%, $|log_2FC| \geq 1$).

**b** and **d** Heatmap of differentially expressed genes (FDR <5%) based on average expression values.

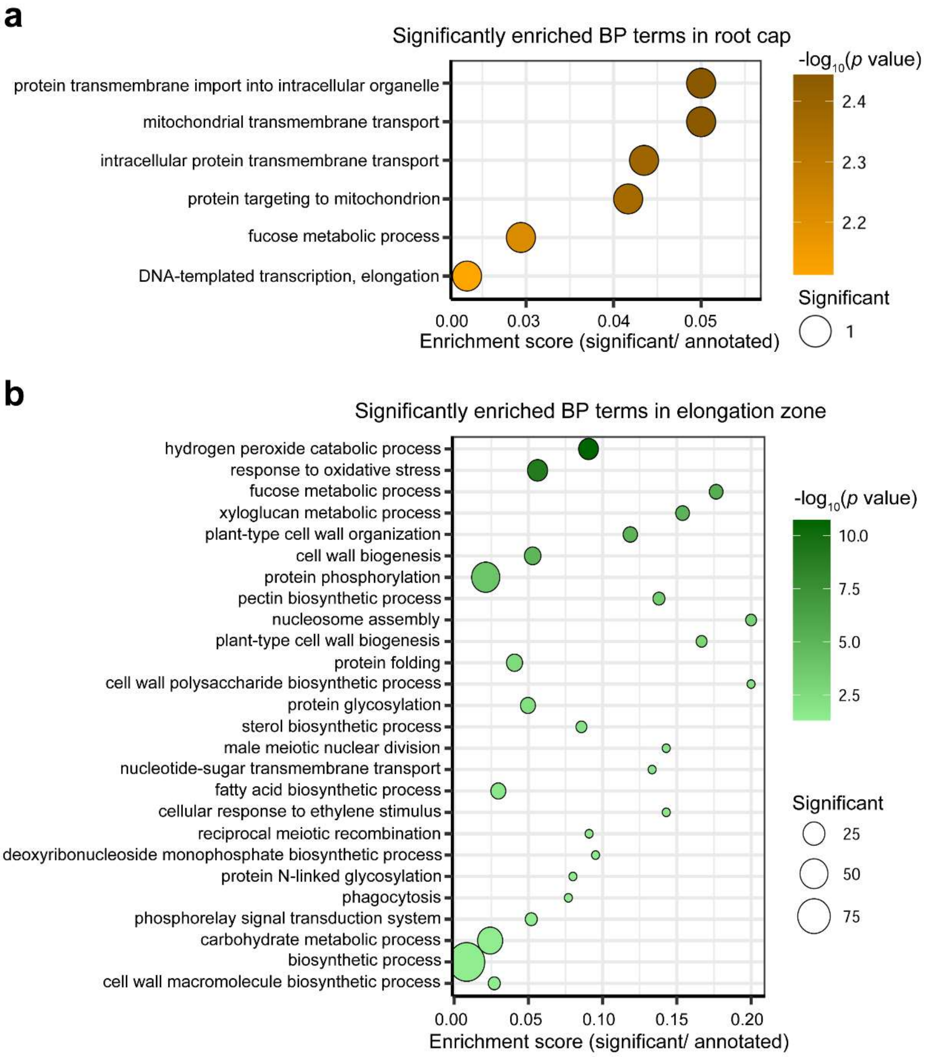


**Fig. S2: Enriched gene ontology (GO) terms for gravity regulated genes that are *EGT2* related.**

**a-b** Significantly enriched GO terms among intersected differentially expressed genes (FDR <5%) in root cap (a) and elongation zone (b).

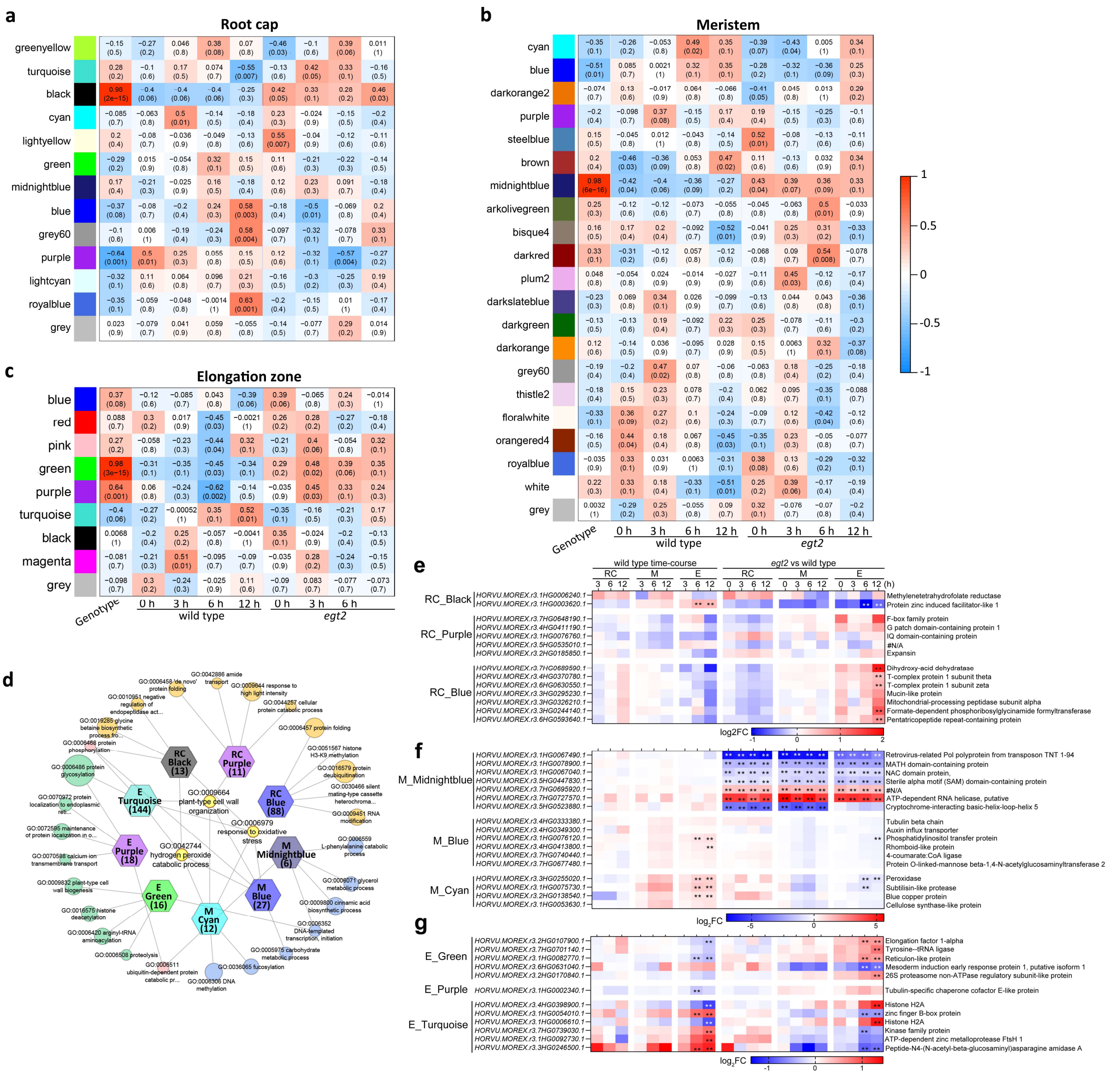


**Fig. S3: Co-expression analyses.**

**a-c** Module-trait relationships in the root cap (a), the meristem (b) and the elongation zone (c). The genotypes (wily type and *egt2*) and time points (0, 3, 6 and 12 h) after gravistimulation are used as traits, each column corresponds to a different trait. Each row corresponds to the characteristic genes of the module. The relationship between the modules and traits is indicated in cell by Pearson correlation coefficients. Numbers in brackets indicate significant levels. Cell color ranges from red (highly positive correlation) to blue (highly negative correction).

**d** Gene ontology analyses of genes in selected significantly correlated modules in the root cap, the meristem and the elongation zone. The five most significantly enriched biological process terms of each module are shown. Hexagonal nodes represent modules, circle nodes represent individual GO terms. Numbers in hexagonal indicate the total number of significantly enriched biological process terms in corresponding module. The size of circle nodes indicates the ratio of the number of genes associated with each GO term to the total number of genes annotated for corresponding term.

**e-g** The expression patterns of hub genes of the black, the purple and the blue modules in the root cap (RC; e), the midnightblue, the blue and cyan modules in the meristem (M; f), and the green, purple and turquoise modules in the elongation zone (E; g) in the WT gravistimulation time-course experiments (wild type time-course) and in the comparisons of *egt2* with wild type (*egt2* vs wild type). **, FDR<5%.

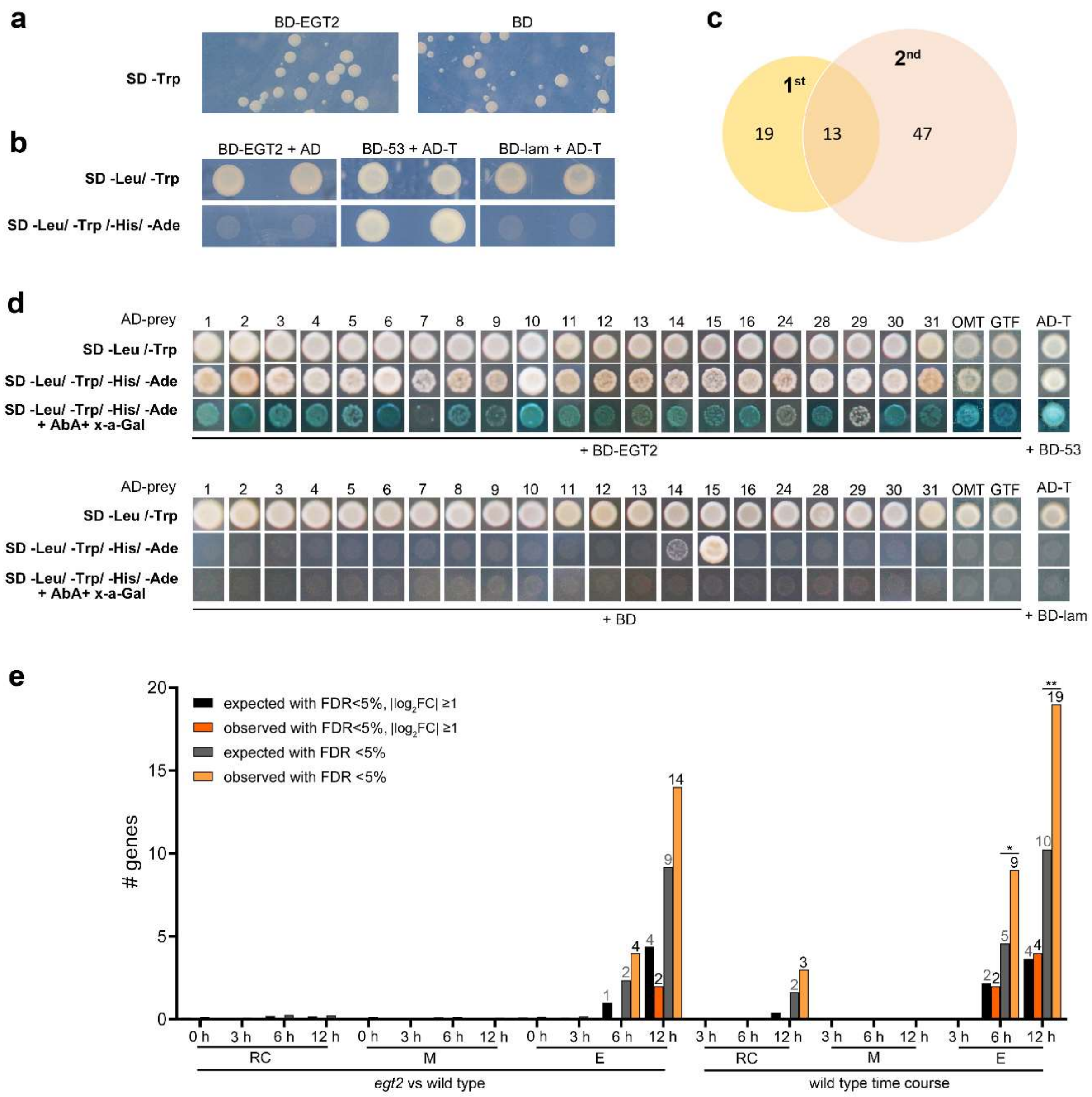


**Fig. S4: Overview of yeast-two-hybrid screening.**

**a**. Toxicity testing of EGT2.

**b**. Autoactivation testing of EGT2.

**c**. Numbers of interaction partners identified by two screens.

**d**. Confirmation of the interaction between EGT2 (BD-EGT2) and the 23 interaction partners (AD-prey) by one-on-one yeast-two-hybrid assays. Co-transformation with each of the AD-preys and pGBKT7 (BD) was used as a negative control. OMT: HORVU.MOREX.r3.3HG0330120.1; GTF: HORVU.MOREX.r3.7HG0736300.1.

**c-d**. Co-transformation of BD-53 and AD-T was shown as positive control, co-transformation of BD-lam and AD-T was shown as negative control,

**e**. Numbers of expected interaction partners and observed interaction candidates in differentially expressed genes. Fisher's exact test, *, $p$ <0.05; **, $p$ <0.01.

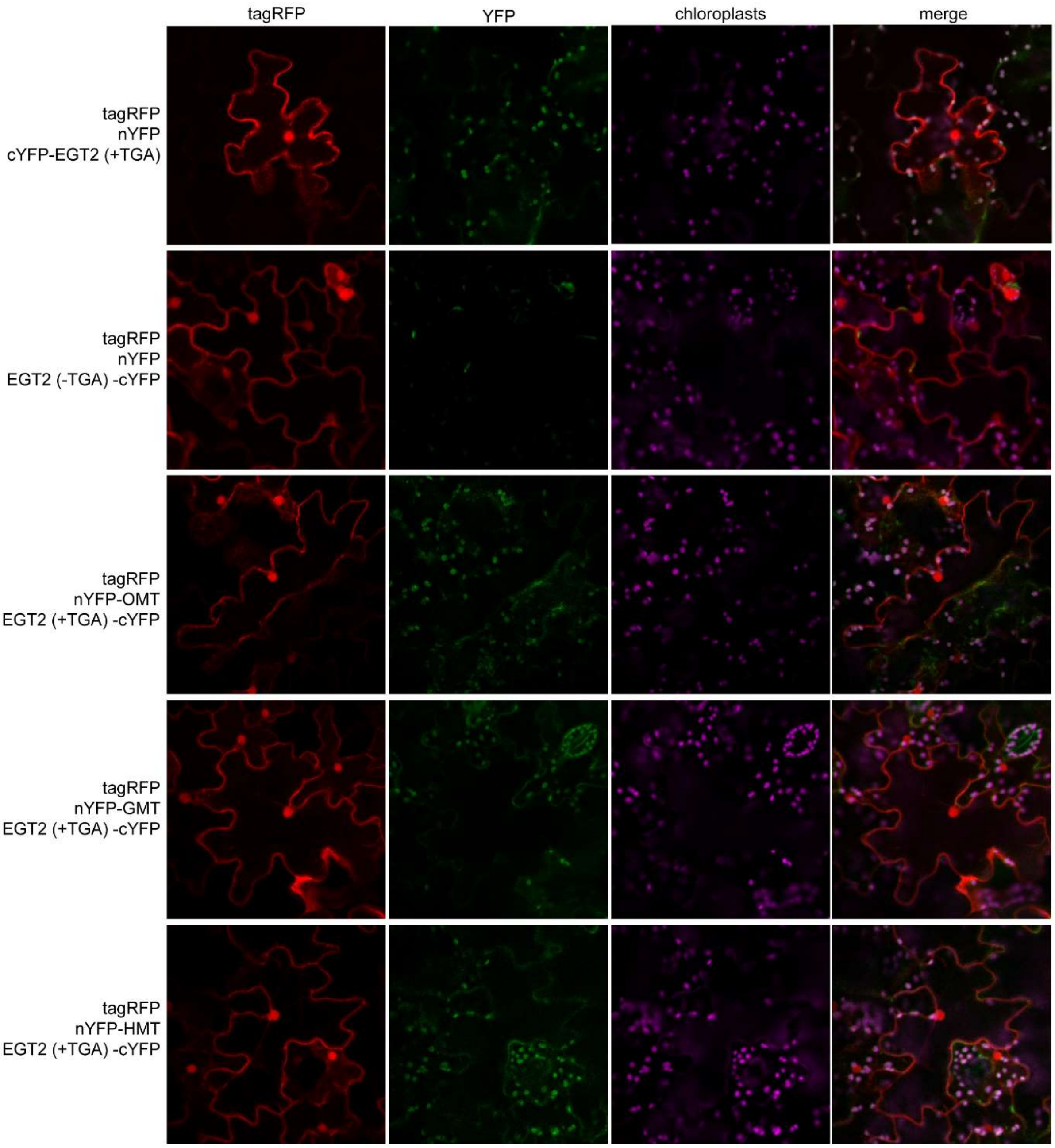


**Fig. S5: Control experiments for bimolecular fluorescence complementation analyses of EGT2 and interaction candidates.**

tagRFP: red fluorescence; YFP: yellow fluorescence; chloroplasts: auto-fluorescence of chloroplasts. nYFP: N-terminal part of YFP; cYFP: C-terminal part of YFP. EGT2 (+TGA): full-length coding sequence of EGT2; EGT2 (-TGA): coding sequence of EGT2 removed stop codon.

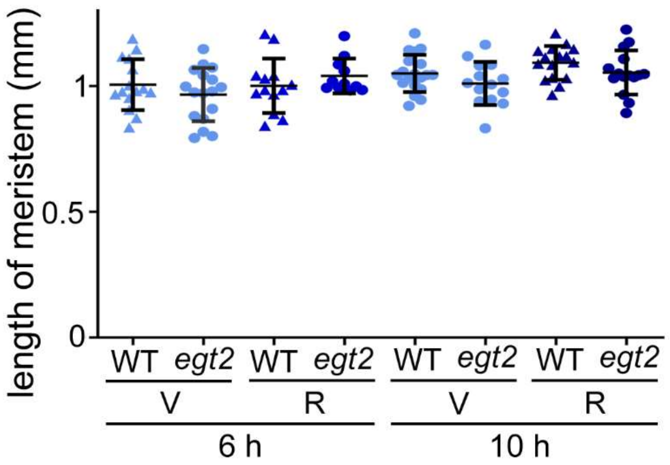


**Fig. S6: Quantification of meristem length of wild type (WT) and *egt2* before and after 6 h or 10 h of gravistimulation.**

(V): vertically grown samples; (R): rotated samples. n = 11-19 per genotype by time point combination.

**Table S1** Overview of differentially expressed genes (FDR <5%) in gravistimulated wild type roots in a time course experiment.

| ID | logFC | AveExpr | t | P.Value | adj.P.Val | B | gene description | root zone | time point |
|---|---|---|---|---|---|---|---|---|---|
| HORVU.MOREX.r3.1HG0003270.1 | 2.823399 | -2.89111 | 7.20642 | 2.10E-09 | 4.30E-05 | -3.57691 | Core-2/I-branching beta-1,6-N-acetylglucosaminyltransferase family protein | root cap | 3 h |
| HORVU.MOREX.r3.4HG0405330.1 | 1.798336 | -1.47892 | 5.69161 | 5.58E-07 | 0.011441 | 5.598813 | Cytochrome P450, putative | root cap | 6 h |
| HORVU.MOREX.r3.4HG0402190.1 | 2.577422 | -1.87054 | 5.34876 | 1.92E-06 | 0.01972 | 4.572588 | Flavin-containing monooxygenase | root cap | 6 h |
| HORVU.MOREX.r3.7HG0656980.1 | 1.490765 | -1.98412 | 4.979942 | 7.12E-06 | 0.029524 | 3.341386 | Transposon protein, putative, CACTA, En/Spm sub-class, expressed | root cap | 6 h |
| HORVU.MOREX.r3.3HG0300110.1 | 1.511206 | -2.56986 | 4.97713 | 7.19E-06 | 0.029524 | 2.824972 | Acidic endochitinase | root cap | 6 h |
| HORVU.MOREX.r3.3HG0276000.2 | 1.810064 | -3.10951 | 5.10213 | 4.63E-06 | 0.029524 | 2.256709 | GDSL esterase/lipase | root cap | 6 h |
| HORVU.MOREX.r3.7HG0739570.1 | 2.549347 | 1.204008 | 4.862519 | 1.07E-05 | 0.03676 | 1.742172 | cotton fiber protein | root cap | 6 h |
| HORVU.MOREX.r3.2HG0125600.1 | -1.28244 | -0.84295 | -6.38892 | 4.33E-08 | 0.000888 | 8.407785 | DNA-directed RNA polymerase subunit alpha | root cap | 12 h |
| HORVU.MOREX.r3.7HG0732860.1 | -0.84978 | 6.992859 | -6.07936 | 1.35E-07 | 0.001387 | 7.343196 | Polyubiquitin | root cap | 12 h |
| HORVU.MOREX.r3.7HG0702990.1 | -0.54851 | 5.286374 | -5.72619 | 4.92E-07 | 0.002233 | 6.156683 | WD-repeat protein, putative | root cap | 12 h |
| HORVU.MOREX.r3.6HG0552230.1 | -0.8487 | 6.698601 | -5.72377 | 4.96E-07 | 0.002233 | 6.12943 | Polyubiquitin | root cap | 12 h |
| HORVU.MOREX.r3.6HG0565880.1 | -0.51588 | 6.39216 | -5.69834 | 5.44E-07 | 0.002233 | 6.059052 | Polyubiquitin | root cap | 12 h |
| HORVU.MOREX.r3.6HG0553150.1 | -0.81679 | 6.876917 | -5.57225 | 8.59E-07 | 0.002396 | 5.612467 | Polyubiquitin | root cap | 12 h |
| HORVU.MOREX.r3.6HG0559390.1 | -1.1202 | 4.844853 | -5.53987 | 9.66E-07 | 0.002396 | 5.511704 | Polyubiquitin protein | root cap | 12 h |
| HORVU.MOREX.r3.7HG0724830.1 | -0.68712 | 6.736593 | -5.54022 | 9.65E-07 | 0.002396 | 5.470374 | Zinc finger A20 and AN1 domain stress-associated protein | root cap | 12 h |
| HORVU.MOREX.r3.1HG0093750.1 | -1.02559 | 3.434371 | -5.51661 | 1.05E-06 | 0.002396 | 5.423285 | ATP-dependent zinc metalloprotease FtsH | root cap | 12 h |
| HORVU.MOREX.r3.7HG0675390.1 | -0.83136 | 3.34579 | -5.47982 | 1.20E-06 | 0.002435 | 5.312858 | Thioredoxin-like family protein | root cap | 12 h |
| HORVU.MOREX.r3.6HG0565890.1 | -0.81717 | 8.056863 | -5.45027 | 1.33E-06 | 0.002435 | 5.156865 | Polyubiquitin | root cap | 12 h |
| HORVU.MOREX.r3.2HG0173440.1 | -0.85101 | 3.881615 | -5.41378 | 1.52E-06 | 0.002435 | 5.08651 | AGAMOUS-like MADS-box transcription factor | root cap | 12 h |
| HORVU.MOREX.r3.2HG0214820.1 | -0.99075 | 3.095886 | -5.41001 | 1.54E-06 | 0.002435 | 5.083794 | Protein kinase, putative | root cap | 12 h |
| HORVU.MOREX.r3.7HG0733110.1 | -0.70107 | 6.267735 | -5.24905 | 2.75E-06 | 0.004024 | 4.523263 | Polyubiquitin | root cap | 12 h |
| HORVU.MOREX.r3.3HG0299100.1 | -0.54346 | 4.96179 | -5.18945 | 3.39E-06 | 0.004274 | 4.346558 | Zinc finger family protein | root cap | 12 h |
| HORVU.MOREX.r3.2HG0203390.1 | -1.08245 | -1.42637 | -5.08974 | 4.84E-06 | 0.004735 | 3.991515 | Auxin-responsive protein | root cap | 12 h |
| HORVU.MOREX.r3.1HG0081860.1 | -0.87883 | -0.61955 | -5.07269 | 5.14E-06 | 0.004735 | 3.961192 | Pollen Ole e 1 allergen/extensin | root cap | 12 h |
| HORVU.MOREX.r3.7HG0709700.1 | -1.08591 | 5.754852 | -5.0935 | 4.77E-06 | 0.004735 | 3.949535 | PENTATRICOPEPTIDE REPEAT 596 | root cap | 12 h |
| HORVU.MOREX.r3.1HG0057800.1 | -1.34225 | 4.113561 | -5.09894 | 4.68E-06 | 0.004735 | 3.916959 | Protein BRANCHLESS TRICHOME | root cap | 12 h |
| HORVU.MOREX.r3.3HG0244080.1 | -0.96407 | 6.147357 | -5.06947 | 5.20E-06 | 0.004735 | 3.888486 | Homeobox leucine-zipper protein | root cap | 12 h |
| HORVU.MOREX.r3.2HG0100060.1 | -1.04214 | 3.172843 | -5.0495 | 5.57E-06 | 0.004735 | 3.874201 | Arginase | root cap | 12 h |
| HORVU.MOREX.r3.1HG0051890.1 | -0.87443 | 2.409167 | -5.03979 | 5.77E-06 | 0.004735 | 3.854096 | Glutathione S-transferase | root cap | 12 h |
| HORVU.MOREX.r3.4HG0352420.1 | -0.64137 | 4.361963 | -5.0399 | 5.77E-06 | 0.004735 | 3.842691 | MACPF domain protein | root cap | 12 h |
| HORVU.MOREX.r3.6HG0614420.1 | -0.70609 | 5.04629 | -5.01674 | 6.26E-06 | 0.004939 | 3.778866 | ethylene-responsive transcription factor | root cap | 12 h |
| HORVU.MOREX.r3.6HG0621190.1 | -1.04126 | 4.746594 | -4.98162 | 7.08E-06 | 0.00519 | 3.662177 | Chaperone protein dnaJ | root cap | 12 h |
| HORVU.MOREX.r3.7HG0671720.1 | -0.87912 | 4.165549 | -4.9496 | 7.92E-06 | 0.005607 | 3.506882 | Amino acid transporter, putative | root cap | 12 h |
| HORVU.MOREX.r3.1HG0015430.1 | -1.59517 | 0.318582 | -5.19004 | 3.39E-06 | 0.004274 | 3.446159 | Trihelix transcription factor GT-3b | root cap | 12 h |
| HORVU.MOREX.r3.6HG0611930.1 | -1.53928 | -2.37556 | -5.1776 | 3.54E-06 | 0.004274 | 3.42081 | RING/U-box superfamily protein, putative | root cap | 12 h |
| HORVU.MOREX.r3.4HG0403930.1 | -1.02401 | -0.63527 | -4.92116 | 8.76E-06 | 0.005796 | 3.413686 | Cysteine proteinase inhibitor | root cap | 12 h |
| HORVU.MOREX.r3.2HG0097600.1 | -0.52797 | 5.036264 | -4.87356 | 1.03E-05 | 0.006632 | 3.31246 | Protein DA1-related 1 | root cap | 12 h |
| HORVU.MOREX.r3.1HG0025520.1 | -1.12521 | 2.003818 | -4.99605 | 6.73E-06 | 0.005115 | 3.302967 | RING/U-box superfamily protein | root cap | 12 h |
| HORVU.MOREX.r3.6HG0568720.1 | -0.81421 | 3.242843 | -4.85594 | 1.10E-05 | 0.006839 | 3.256707 | #N/A | root cap | 12 h |
| HORVU.MOREX.r3.5HG0509060.1 | -0.95964 | 4.702328 | -4.83861 | 1.17E-05 | 0.006945 | 3.196184 | Non-specific serine/threonine protein kinase | root cap | 12 h |
| HORVU.MOREX.r3.3HG0256910.1 | -0.82045 | 2.543019 | -4.82557 | 1.22E-05 | 0.006945 | 3.154161 | Ribonuclease I | root cap | 12 h |

**Table S1** Continued.

| ID | logFC | AveExpr | t | P.Value | adj.P.Val | B | gene description | root zone | time point |
|---|---|---|---|---|---|---|---|---|---|
| HORVU.MOREX.r3.6HG0607560.1 | -0.80876 | 4.13176 | -4.81846 | 1.25E-05 | 0.006945 | 3.127532 | Adenine nucleotide alpha hydrolases-like superfamily protein | root cap | 12 h |
| HORVU.MOREX.r3.6HG0568690.1 | -1.03891 | 4.283634 | -4.81101 | 1.29E-05 | 0.006945 | 3.104319 | Cinnamoyl CoA reductase | root cap | 12 h |
| HORVU.MOREX.r3.3HG0249970.1 | -0.55296 | 3.802328 | -4.80111 | 1.33E-05 | 0.007004 | 3.080605 | Beta-amylase | root cap | 12 h |
| HORVU.MOREX.r3.7HG0707460.1 | -0.91662 | 2.531992 | -4.79321 | 1.37E-05 | 0.007019 | 3.029348 | Protein kinase, putative | root cap | 12 h |
| HORVU.MOREX.r3.1HG0008890.1 | -0.50372 | 5.398609 | -4.78053 | 1.43E-05 | 0.007156 | 2.990593 | Coiled-coil domain-containing protein, putative | root cap | 12 h |
| HORVU.MOREX.r3.2HG0140970.1 | -1.70457 | 2.378717 | -4.92788 | 8.55E-06 | 0.005796 | 2.972836 | Heat-shock protein, putative | root cap | 12 h |
| HORVU.MOREX.r3.1HG0062480.1 | 0.848151 | 5.054697 | 4.747402 | 1.60E-05 | 0.007837 | 2.903511 | 1-deoxy-D-xylulose-5-phosphate synthase | root cap | 12 h |
| HORVU.MOREX.r3.5HG0426180.1 | -0.59936 | 4.726965 | -4.72387 | 1.74E-05 | 0.008115 | 2.835302 | Thioesterase family protein | root cap | 12 h |
| HORVU.MOREX.r3.5HG0495770.1 | -0.48077 | 4.845038 | -4.7046 | 1.86E-05 | 0.008386 | 2.760621 | RING/U-box superfamily protein | root cap | 12 h |
| HORVU.MOREX.r3.2HG0130760.1 | -0.6913 | 4.232936 | -4.68621 | 1.98E-05 | 0.008472 | 2.715463 | Protein yippee-like | root cap | 12 h |
| HORVU.MOREX.r3.1HG0029270.1 | -0.56384 | 4.599412 | -4.64652 | 2.27E-05 | 0.008966 | 2.590698 | Ubiquitin conjugating enzyme E2 | root cap | 12 h |
| HORVU.MOREX.r3.6HG0601740.1 | -1.11478 | -1.96393 | -4.67705 | 2.05E-05 | 0.008566 | 2.554705 | Formate-dependent nitrite reductase complex subunit NrfG | root cap | 12 h |
| HORVU.MOREX.r3.3HG0251900.1 | -0.58184 | 5.429565 | -4.65874 | 2.18E-05 | 0.008941 | 2.553179 | WRKY transcription factor | root cap | 12 h |
| HORVU.MOREX.r3.7HG0659980.1 | -0.75289 | 5.13795 | -4.64851 | 2.26E-05 | 0.008966 | 2.54876 | UAA transporter | root cap | 12 h |
| HORVU.MOREX.r3.2HG0191150.1 | 0.987218 | 5.670797 | 4.624858 | 2.45E-05 | 0.00899 | 2.513627 | Polygalacturonase QRT3 | root cap | 12 h |
| HORVU.MOREX.r3.3HG0281000.1 | -1.13625 | 3.470695 | -4.62746 | 2.43E-05 | 0.00899 | 2.513107 | F-box protein | root cap | 12 h |
| HORVU.MOREX.r3.4HG0332530.1 | -0.56658 | 4.558532 | -4.62005 | 2.49E-05 | 0.00899 | 2.504111 | Aldo-keto reductase/ oxidoreductase | root cap | 12 h |
| HORVU.MOREX.r3.1HG0075220.1 | -1.01928 | 2.657679 | -4.6239 | 2.46E-05 | 0.00899 | 2.497509 | Actin | root cap | 12 h |
| HORVU.MOREX.r3.5HG0487890.1 | -0.81835 | 4.484514 | -4.6133 | 2.55E-05 | 0.00901 | 2.478508 | Protein DETOXIFICATION | root cap | 12 h |
| HORVU.MOREX.r3.1HG0053120.1 | -0.7353 | 4.408808 | -4.619 | 2.50E-05 | 0.00899 | 2.467546 | ADP-ribosylation factor GTPase-activating protein | root cap | 12 h |
| HORVU.MOREX.r3.1HG0029060.1 | 0.331688 | 9.685002 | 4.688856 | 1.96E-05 | 0.008472 | 2.452944 | 70 kDa heat shock protein | root cap | 12 h |
| HORVU.MOREX.r3.2HG0121950.1 | -0.70691 | 3.757635 | -4.59002 | 2.76E-05 | 0.009433 | 2.412075 | RNA binding protein | root cap | 12 h |
| HORVU.MOREX.r3.7HG0663550.1 | -2.12756 | 0.155997 | -4.72681 | 1.72E-05 | 0.008115 | 2.398947 | Glycosyltransferase | root cap | 12 h |
| HORVU.MOREX.r3.1HG0056360.1 | -0.48733 | 6.496535 | -4.60306 | 2.64E-05 | 0.009174 | 2.357061 | Oxysterol-binding protein, expressed | root cap | 12 h |
| HORVU.MOREX.r3.2HG0099980.1 | -0.76137 | 2.832965 | -4.56222 | 3.03E-05 | 0.009444 | 2.325886 | Glyoxylate reductase/hydroxypyruvate reductase | root cap | 12 h |
| HORVU.MOREX.r3.7HG0678530.1 | -0.53528 | 5.729845 | -4.56623 | 2.99E-05 | 0.009444 | 2.300151 | Ubiquitin-associated domain-containing family protein | root cap | 12 h |
| HORVU.MOREX.r3.6HG0557640.2 | -0.85253 | 6.081969 | -4.55992 | 3.06E-05 | 0.009444 | 2.284663 | Polyubiquitin | root cap | 12 h |
| HORVU.MOREX.r3.5HG0427690.1 | -0.85507 | 0.757744 | -4.56751 | 2.98E-05 | 0.009444 | 2.274183 | Exostosin family protein | root cap | 12 h |
| HORVU.MOREX.r3.1HG0058170.1 | -0.49534 | 5.459776 | -4.55165 | 3.14E-05 | 0.009444 | 2.266576 | Autophagy-related protein 3 | root cap | 12 h |
| HORVU.MOREX.r3.3HG0231010.1 | -0.48328 | 5.831789 | -4.55366 | 3.12E-05 | 0.009444 | 2.260216 | NH(3)-dependent NAD(+) synthetase | root cap | 12 h |
| HORVU.MOREX.r3.5HG0501950.1 | -1.18653 | 0.379734 | -4.54541 | 3.21E-05 | 0.009444 | 2.228017 | RNA-binding (RRM/RBD/RNP motifs) family protein | root cap | 12 h |
| HORVU.MOREX.r3.7HG0703000.1 | -0.71136 | 5.112448 | -4.52506 | 3.44E-05 | 0.009522 | 2.194884 | UDP-glucose 4-epimerase, putative | root cap | 12 h |
| HORVU.MOREX.r3.3HG0227890.1 | -1.18155 | 2.984434 | -4.52994 | 3.39E-05 | 0.009522 | 2.183117 | Lactation elevated protein 1 | root cap | 12 h |
| HORVU.MOREX.r3.2HG0201420.1 | -1.09279 | -0.04773 | -4.51186 | 3.60E-05 | 0.009598 | 2.142929 | Protein kinase | root cap | 12 h |
| HORVU.MOREX.r3.2HG0173750.1 | -0.50742 | 8.547687 | -4.56753 | 2.98E-05 | 0.009444 | 2.124164 | Aldehyde dehydrogenase | root cap | 12 h |
| HORVU.MOREX.r3.3HG0329000.1 | -0.54548 | 7.386719 | -4.54458 | 3.22E-05 | 0.009444 | 2.112378 | Haloacid dehalogenase-like hydrolase family protein, putative, expressed | root cap | 12 h |
| HORVU.MOREX.r3.3HG0271550.1 | -0.52196 | 6.477159 | -4.5233 | 3.46E-05 | 0.009522 | 2.090392 | Ubiquitin-like modifier-activating enzyme atg7 | root cap | 12 h |
| HORVU.MOREX.r3.4HG0348850.1 | -0.69046 | 8.009697 | -4.5358 | 3.32E-05 | 0.009522 | 2.084904 | Neutral/alkaline invertase | root cap | 12 h |
| HORVU.MOREX.r3.2HG0186670.1 | -0.67647 | 4.974138 | -4.48168 | 3.99E-05 | 0.010498 | 2.029023 | 3-oxo-5-alpha-steroid 4-dehydrogenase 3 | root cap | 12 h |
| HORVU.MOREX.r3.4HG0397570.1 | -0.95641 | 4.104655 | -4.47622 | 4.07E-05 | 0.010559 | 2.019113 | Glucan endo-1,3-beta-glucosidase 3 | root cap | 12 h |
| HORVU.MOREX.r3.1HG0085780.1 | -1.01066 | 2.653546 | -4.4574 | 4.33E-05 | 0.010585 | 1.999303 | Core-2/I-branching beta-1,6-N-acetylglucosaminyltransferase family protein | root cap | 12 h |

**Table S1** Continued.

| ID | logFC | AveExpr | t | P.Value | adj.P.Val | B | gene description | root zone | time point |
|---|---|---|---|---|---|---|---|---|---|
| HORVU.MOREX.r3.7HG0749800.1 | -0.77039 | 4.620717 | -4.46721 | 4.19E-05 | 0.010585 | 1.989527 | #N/A | root cap | 12 h |
| HORVU.MOREX.r3.2HG0195580.1 | -0.60678 | 7.713102 | -4.52192 | 3.48E-05 | 0.009522 | 1.965649 | UDP-glucose 4-epimerase, putative | root cap | 12 h |
| HORVU.MOREX.r3.7HG0733070.1 | -0.51348 | 6.760905 | -4.47075 | 4.14E-05 | 0.010585 | 1.925919 | Polyubiquitin | root cap | 12 h |
| HORVU.MOREX.r3.3HG0228600.1 | -0.80407 | 3.194099 | -4.43292 | 4.71E-05 | 0.011232 | 1.920616 | transmembrane protein, putative (DUF247) | root cap | 12 h |
| HORVU.MOREX.r3.4HG0384590.1 | -1.4357 | 1.137938 | -4.46206 | 4.27E-05 | 0.010585 | 1.86961 | Chaperone protein DnaJ | root cap | 12 h |
| HORVU.MOREX.r3.5HG0481570.1 | -0.8274 | 3.92217 | -4.4383 | 4.62E-05 | 0.011159 | 1.809097 | Inactive phospholipase C-like protein 1 | root cap | 12 h |
| HORVU.MOREX.r3.2HG0181380.1 | 0.784857 | 2.418376 | 4.420454 | 4.91E-05 | 0.01158 | 1.782252 | Actin-related protein 2/3 complex subunit 2 | root cap | 12 h |
| HORVU.MOREX.r3.5HG0504720.1 | 0.464637 | 6.757338 | 4.410774 | 5.07E-05 | 0.011829 | 1.735987 | Octicosapeptide/Phox/Bem1p domain-containing protein / tetratricopeptide repe | root cap | 12 h |
| HORVU.MOREX.r3.7HG0713100.1 | -0.72286 | 3.920881 | -4.37511 | 5.72E-05 | 0.012723 | 1.72962 | transmembrane protein, putative (DUF594) | root cap | 12 h |
| HORVU.MOREX.r3.2HG0154110.1 | -1.034 | 1.510407 | -4.36463 | 5.93E-05 | 0.012723 | 1.717556 | Succinate dehydrogenase [ubiquinone] flavoprotein subunit, mitochondrial | root cap | 12 h |
| HORVU.MOREX.r3.2HG0175220.1 | -0.43669 | 9.281963 | -4.4605 | 4.29E-05 | 0.010585 | 1.70697 | Amine oxidase | root cap | 12 h |
| HORVU.MOREX.r3.2HG0117540.1 | -0.56241 | 3.822761 | -4.36329 | 5.95E-05 | 0.012723 | 1.703811 | Methyltransferase-like protein | root cap | 12 h |
| HORVU.MOREX.r3.7HG0685620.1 | -0.38926 | 5.51038 | -4.36564 | 5.91E-05 | 0.012723 | 1.695952 | RING/U-box superfamily protein | root cap | 12 h |
| HORVU.MOREX.r3.4HG0342860.1 | -0.83523 | 3.568964 | -4.35774 | 6.06E-05 | 0.012829 | 1.673634 | Kinase family protein | root cap | 12 h |
| HORVU.MOREX.r3.1HG0093880.1 | -0.77277 | 4.163907 | -4.36436 | 5.93E-05 | 0.012723 | 1.672783 | Phosphoglucan phosphatase LSF1 | root cap | 12 h |
| HORVU.MOREX.r3.1HG0002330.1 | -0.66462 | 5.33192 | -4.36457 | 5.93E-05 | 0.012723 | 1.668277 | Ras-like protein | root cap | 12 h |
| HORVU.MOREX.r3.1HG0065120.1 | -0.54994 | 3.907242 | -4.35072 | 6.21E-05 | 0.013001 | 1.657088 | Protein kinase superfamily protein | root cap | 12 h |
| HORVU.MOREX.r3.4HG0391970.1 | -1.21575 | -1.7379 | -4.37105 | 5.80E-05 | 0.012723 | 1.652152 | LOB domain protein | root cap | 12 h |
| HORVU.MOREX.r3.6HG0551620.1 | -0.54924 | 5.566118 | -4.33945 | 6.45E-05 | 0.013232 | 1.599519 | BAX inhibitor 1 | root cap | 12 h |
| HORVU.MOREX.r3.7HG0668190.1 | -0.58408 | 4.882662 | -4.31675 | 6.96E-05 | 0.013841 | 1.53107 | Phytochelatin synthase | root cap | 12 h |
| HORVU.MOREX.r3.5HG0498320.1 | 0.429306 | 5.332211 | 4.314295 | 7.01E-05 | 0.013841 | 1.519631 | Zinc finger, B-box | root cap | 12 h |
| HORVU.MOREX.r3.3HG0271380.1 | -0.54512 | 5.415877 | -4.32755 | 6.71E-05 | 0.013533 | 1.511055 | GDSL esterase/lipase | root cap | 12 h |
| HORVU.MOREX.r3.3HG0291680.1 | -0.97851 | -0.37502 | -4.29504 | 7.48E-05 | 0.01462 | 1.496854 | Tubulin beta-2 chain | root cap | 12 h |
| HORVU.MOREX.r3.6HG0561320.1 | -1.28175 | -0.01134 | -4.28514 | 7.73E-05 | 0.014962 | 1.475521 | RNA-directed RNA polymerase catalytic subunit | root cap | 12 h |
| HORVU.MOREX.r3.1HG0051880.1 | -0.82245 | 4.550064 | -4.34334 | 6.36E-05 | 0.013192 | 1.467889 | Glutathione S-transferase | root cap | 12 h |
| HORVU.MOREX.r3.3HG0305540.1 | -1.31675 | -2.8839 | -4.70151 | 1.88E-05 | 0.008386 | 1.459601 | DUF1685 family protein | root cap | 12 h |
| HORVU.MOREX.r3.6HG0626000.1 | -1.08875 | 1.922137 | -4.28247 | 7.80E-05 | 0.014962 | 1.36707 | Calcineurin B-like protein | root cap | 12 h |
| HORVU.MOREX.r3.4HG0382640.1 | -1.38804 | -2.50777 | -4.51768 | 3.53E-05 | 0.009533 | 1.349406 | Mitochondrial import inner membrane translocase subunit Tim17/Tim22/Tim23 fa | root cap | 12 h |
| HORVU.MOREX.r3.2HG0198000.1 | 0.300141 | 8.426845 | 4.32681 | 6.73E-05 | 0.013533 | 1.346151 | RNA-binding family protein | root cap | 12 h |
| HORVU.MOREX.r3.2HG0191970.1 | -0.87526 | 3.27463 | -4.23744 | 9.06E-05 | 0.016042 | 1.328992 | Protein phosphatase 2c, putative | root cap | 12 h |
| HORVU.MOREX.r3.1HG0069010.1 | -0.8324 | 3.211686 | -4.23768 | 9.06E-05 | 0.016042 | 1.312716 | Glycerol-3-phosphate dehydrogenase [NAD(+)] | root cap | 12 h |
| HORVU.MOREX.r3.2HG0147190.1 | -0.72627 | 3.250226 | -4.23757 | 9.06E-05 | 0.016042 | 1.291305 | Zinc finger A20 and AN1 domain stress-associated protein | root cap | 12 h |
| HORVU.MOREX.r3.1HG0027030.1 | -0.75711 | 1.805042 | -4.25354 | 8.59E-05 | 0.016026 | 1.284667 | Mitochondrial carrier family | root cap | 12 h |
| HORVU.MOREX.r3.5HG0529140.1 | 0.432548 | 10.3593 | 4.366513 | 5.89E-05 | 0.012723 | 1.284433 | Elongation factor | root cap | 12 h |
| HORVU.MOREX.r3.3HG0291590.1 | -0.62888 | 5.468201 | -4.23001 | 9.29E-05 | 0.016292 | 1.27413 | Tunicamycin induced 1 | root cap | 12 h |
| HORVU.MOREX.r3.2HG0189190.1 | -0.56412 | 5.637918 | -4.23726 | 9.07E-05 | 0.016042 | 1.27098 | Hexosyltransferase | root cap | 12 h |
| HORVU.MOREX.r3.7HG0666350.1 | -0.66019 | 2.690709 | -4.22671 | 9.39E-05 | 0.016332 | 1.258411 | FACT complex subunit SPT16 | root cap | 12 h |
| HORVU.MOREX.r3.4HG0336330.1 | -0.82913 | 0.591327 | -4.20935 | 9.95E-05 | 0.016843 | 1.246155 | #N/A | root cap | 12 h |
| HORVU.MOREX.r3.5HG0519200.1 | -0.80827 | 2.210169 | -4.21322 | 9.82E-05 | 0.016795 | 1.228799 | DUF506 family protein | root cap | 12 h |
| HORVU.MOREX.r3.5HG0501370.1 | -0.90534 | -0.59777 | -4.20619 | 0.000101 | 0.016843 | 1.205796 | Cytochrome P450, putative | root cap | 12 h |
| HORVU.MOREX.r3.7HG0658130.1 | -0.78542 | 4.23398 | -4.20489 | 0.000101 | 0.016843 | 1.183352 | Nodulin-like / Major Facilitator Superfamily protein | root cap | 12 h |

**Table S1** Continued.

| ID | logFC | AveExpr | t | P.Value | adj.P.Val | B | gene description | root zone | time point |
|---|---|---|---|---|---|---|---|---|---|
| HORVU.MOREX.r3.2HG0213480.1 | -0.48207 | 8.002679 | -4.24789 | 8.75E-05 | 0.016038 | 1.160298 | Phosphomannomutase | root cap | 12 h |
| HORVU.MOREX.r3.2HG0104110.1 | -0.92422 | 3.073698 | -4.21371 | 9.81E-05 | 0.016795 | 1.144121 | CW7 protein, putative | root cap | 12 h |
| HORVU.MOREX.r3.1HG0073140.1 | -0.86351 | 1.311141 | -4.1712 | 0.000113 | 0.018526 | 1.132761 | Metal tolerance protein | root cap | 12 h |
| HORVU.MOREX.r3.2HG0202110.1 | -0.43979 | 8.934553 | -4.26572 | 8.25E-05 | 0.01553 | 1.072325 | Alpha-1,4-glucan-protein synthase [UDP-forming] | root cap | 12 h |
| HORVU.MOREX.r3.2HG0099750.1 | -0.55729 | 5.148054 | -4.16018 | 0.000117 | 0.018909 | 1.04572 | carbohydrate esterase, putative (DUF303) | root cap | 12 h |
| HORVU.MOREX.r3.2HG0192960.1 | -0.63914 | 3.213874 | -4.16451 | 0.000115 | 0.018789 | 1.027258 | Xyloglucan endotransglucosylase/hydrolase | root cap | 12 h |
| HORVU.MOREX.r3.6HG0604990.1 | -1.34952 | 1.556962 | -4.2482 | 8.74E-05 | 0.016038 | 1.025382 | senescence-associated family protein (DUF581) | root cap | 12 h |
| HORVU.MOREX.r3.7HG0680170.1 | -0.53226 | 4.825321 | -4.15039 | 0.000121 | 0.019244 | 1.014283 | mediator of RNA polymerase II transcription subunit | root cap | 12 h |
| HORVU.MOREX.r3.4HG0411320.1 | -0.49858 | 6.294368 | -4.14747 | 0.000122 | 0.019262 | 0.994595 | Appr-1-p processing enzyme family protein | root cap | 12 h |
| HORVU.MOREX.r3.3HG0253860.1 | -1.40922 | -0.09276 | -4.13021 | 0.000129 | 0.020232 | 0.988279 | Rapid alkalinization factor (RALF) family protein | root cap | 12 h |
| HORVU.MOREX.r3.6HG0598380.1 | -1.04129 | 1.304428 | -4.12426 | 0.000132 | 0.020321 | 0.974873 | Expansin protein | root cap | 12 h |
| HORVU.MOREX.r3.3HG0249710.1 | -0.68048 | 2.92555 | -4.12575 | 0.000131 | 0.020321 | 0.970391 | Heme-binding protein, putative | root cap | 12 h |
| HORVU.MOREX.r3.6HG0630410.1 | -1.20626 | -1.40652 | -4.11023 | 0.000138 | 0.02066 | 0.957122 | LOB domain protein-like | root cap | 12 h |
| HORVU.MOREX.r3.3HG0250330.1 | -0.55949 | 4.018268 | -4.10577 | 0.00014 | 0.02066 | 0.910066 | Phosphate translocator | root cap | 12 h |
| HORVU.MOREX.r3.2HG0189340.1 | -0.91308 | 1.373881 | -4.19685 | 0.000104 | 0.017158 | 0.902253 | #N/A | root cap | 12 h |
| HORVU.MOREX.r3.7HG0668260.1 | -1.47194 | -0.18118 | -4.10861 | 0.000139 | 0.02066 | 0.901284 | Coiled-coil domain-containing protein 21, putative isoform 1 | root cap | 12 h |
| HORVU.MOREX.r3.2HG0151920.1 | -0.85626 | 3.926991 | -4.08544 | 0.00015 | 0.020899 | 0.88491 | Retrovirus-related Pol polyprotein from transposon TNT 1-94 | root cap | 12 h |
| HORVU.MOREX.r3.3HG0308250.1 | -0.53753 | 5.709159 | -4.10929 | 0.000138 | 0.02066 | 0.882982 | Telomere repeat-binding factor like-protein | root cap | 12 h |
| HORVU.MOREX.r3.7HG0735280.1 | -0.43439 | 7.186121 | -4.1501 | 0.000121 | 0.019244 | 0.867029 | Auxin response factor | root cap | 12 h |
| HORVU.MOREX.r3.5HG0453430.1 | -0.84985 | 3.783951 | -4.07719 | 0.000154 | 0.020977 | 0.855892 | Regulator of chromosome condensation (RCC1) family protein | root cap | 12 h |
| HORVU.MOREX.r3.5HG0476220.1 | -0.72929 | 2.950954 | -4.09329 | 0.000146 | 0.020775 | 0.843927 | Major facilitator superfamily transporter | root cap | 12 h |
| HORVU.MOREX.r3.7HG0664810.1 | -1.23154 | 2.590118 | -4.07476 | 0.000155 | 0.020977 | 0.841051 | Heat-shock protein, putative | root cap | 12 h |
| HORVU.MOREX.r3.7HG0706090.1 | -0.51791 | 5.276785 | -4.08517 | 0.00015 | 0.020899 | 0.840887 | methyl esterase 12 | root cap | 12 h |
| HORVU.MOREX.r3.3HG0234880.1 | -0.78762 | 4.615778 | -4.10121 | 0.000142 | 0.020673 | 0.831034 | Guanine-nucleotide-exchange protein | root cap | 12 h |
| HORVU.MOREX.r3.3HG0251700.1 | -0.90786 | 5.703161 | -4.07378 | 0.000155 | 0.020977 | 0.830229 | Glucan endo-1,3-beta-glucosidase | root cap | 12 h |
| HORVU.MOREX.r3.3HG0328340.1 | -0.80723 | 4.602643 | -4.07522 | 0.000155 | 0.020977 | 0.829255 | Two-component response regulator | root cap | 12 h |
| HORVU.MOREX.r3.5HG0458100.1 | -0.80157 | 5.986393 | -4.09586 | 0.000145 | 0.020749 | 0.820963 | Late embryogenesis abundant (LEA) hydroxyproline-rich glycoprotein family | root cap | 12 h |
| HORVU.MOREX.r3.6HG0565830.1 | -0.70583 | 5.523345 | -4.07154 | 0.000157 | 0.020994 | 0.815855 | Polyubiquitin | root cap | 12 h |
| HORVU.MOREX.r3.2HG0122840.1 | -0.57117 | 5.447828 | -4.10855 | 0.000139 | 0.02066 | 0.795163 | 4-alpha-glucanotransferase | root cap | 12 h |
| HORVU.MOREX.r3.7HG0729010.1 | -0.86518 | -0.16187 | -4.07422 | 0.000155 | 0.020977 | 0.774904 | Peroxidase | root cap | 12 h |
| HORVU.MOREX.r3.7HG0752830.1 | -0.42282 | 6.536164 | -4.08555 | 0.00015 | 0.020899 | 0.743067 | Signal peptide peptidase-like protein | root cap | 12 h |
| HORVU.MOREX.r3.3HG0284890.1 | -0.69381 | 4.764663 | -4.10614 | 0.00014 | 0.02066 | 0.733629 | Alpha-amylase | root cap | 12 h |
| HORVU.MOREX.r3.7HG0729510.1 | -0.76062 | 4.709018 | -4.03592 | 0.000176 | 0.022832 | 0.723945 | WEB family protein, chloroplastic | root cap | 12 h |
| HORVU.MOREX.r3.2HG0133060.1 | -0.87061 | 2.031127 | -4.03168 | 0.000178 | 0.022855 | 0.713484 | Vacuole membrane-like protein | root cap | 12 h |
| HORVU.MOREX.r3.5HG0511040.1 | -0.94978 | 2.881944 | -4.05589 | 0.000165 | 0.021808 | 0.693605 | #N/A | root cap | 12 h |
| HORVU.MOREX.r3.2HG0185660.1 | -0.40954 | 4.65136 | -4.03994 | 0.000174 | 0.022679 | 0.689467 | Kinase family protein | root cap | 12 h |
| HORVU.MOREX.r3.3HG0231330.1 | -0.82403 | 4.950299 | -4.02029 | 0.000185 | 0.023143 | 0.685121 | Transferase family protein | root cap | 12 h |
| HORVU.MOREX.r3.3HG0234890.1 | -0.76907 | 3.013939 | -4.00588 | 0.000194 | 0.023321 | 0.636478 | ARF guanine-nucleotide exchange factor GNOM | root cap | 12 h |
| HORVU.MOREX.r3.3HG0293570.1 | -2.92576 | -0.38794 | -4.81622 | 1.26E-05 | 0.006945 | 0.629342 | BTB/POZ domain-containing protein | root cap | 12 h |
| HORVU.MOREX.r3.7HG0635050.1 | -0.76481 | 0.346306 | -4.00553 | 0.000194 | 0.023321 | 0.625437 | NBS-LRR-like resistance protein | root cap | 12 h |
| HORVU.MOREX.r3.5HG0517300.1 | -1.09887 | 3.002936 | -4.00488 | 0.000194 | 0.023321 | 0.624958 | Aldehyde oxidase, putative | root cap | 12 h |

**Table S1** Continued.

| ID | logFC | AveExpr | t | P.Value | adj.P.Val | B | gene description | root zone | time point |
|---|---|---|---|---|---|---|---|---|---|
| HORVU.MOREX.r3.6HG0570660.1 | -1.30055 | 3.599982 | -3.99489 | 0.000201 | 0.023321 | 0.62235 | DUF581 family protein | root cap | 12 h |
| HORVU.MOREX.r3.3HG0252270.1 | -0.96764 | 2.838807 | -4.06773 | 0.000158 | 0.021119 | 0.618632 | Germin-like protein 1 | root cap | 12 h |
| HORVU.MOREX.r3.5HG0480430.1 | -0.75279 | 4.978135 | -4.0126 | 0.00019 | 0.023321 | 0.604252 | Aldehyde dehydrogenase | root cap | 12 h |
| HORVU.MOREX.r3.2HG0119220.1 | -0.69131 | 4.892203 | -3.99638 | 0.0002 | 0.023321 | 0.60106 | Gibberellin receptor GID1a | root cap | 12 h |
| HORVU.MOREX.r3.5HG0446490.1 | -0.80884 | 1.880935 | -3.98706 | 0.000206 | 0.023449 | 0.593292 | Syntaxin | root cap | 12 h |
| HORVU.MOREX.r3.7HG0705240.1 | -0.87314 | 0.811856 | -4.01815 | 0.000186 | 0.023163 | 0.582854 | S-adenosyl-L-methionine-dependent methyltransferases superfamily protein | root cap | 12 h |
| HORVU.MOREX.r3.2HG0177780.1 | -0.49352 | 3.700772 | -3.98039 | 0.000211 | 0.023449 | 0.563349 | Serine incorporator | root cap | 12 h |
| HORVU.MOREX.r3.4HG0403910.1 | -1.17259 | -1.1504 | -3.97882 | 0.000212 | 0.023449 | 0.533893 | Cysteine proteinase inhibitor | root cap | 12 h |
| HORVU.MOREX.r3.5HG0519030.1 | -0.66053 | 3.970546 | -3.9592 | 0.000225 | 0.0236 | 0.513875 | Transmembrane protein 97 | root cap | 12 h |
| HORVU.MOREX.r3.3HG0275260.1 | -0.75245 | 1.027903 | -3.99991 | 0.000198 | 0.023321 | 0.513822 | Mitogen-activated protein kinase | root cap | 12 h |
| HORVU.MOREX.r3.6HG0607470.1 | -0.45716 | 5.597761 | -4.02674 | 0.000181 | 0.022855 | 0.513739 | Inorganic pyrophosphatase, putative | root cap | 12 h |
| HORVU.MOREX.r3.2HG0128490.1 | -0.42898 | 4.434925 | -3.95969 | 0.000225 | 0.0236 | 0.511137 | #N/A | root cap | 12 h |
| HORVU.MOREX.r3.2HG0205250.1 | -0.78193 | 4.533688 | -4.00004 | 0.000198 | 0.023321 | 0.510101 | EamA-like transporter family | root cap | 12 h |
| HORVU.MOREX.r3.5HG0471410.1 | -0.46273 | 5.492756 | -3.98531 | 0.000207 | 0.023449 | 0.509979 | Sodium/hydrogen exchanger | root cap | 12 h |
| HORVU.MOREX.r3.1HG0072100.1 | -0.77492 | 4.063928 | -3.97803 | 0.000212 | 0.023449 | 0.505805 | 2-oxoglutarate-dependent dioxygenase | root cap | 12 h |
| HORVU.MOREX.r3.2HG0183730.1 | -1.20226 | 3.434756 | -3.9564 | 0.000227 | 0.023693 | 0.503296 | Alpha 1,4-glycosyltransferase family protein, putative | root cap | 12 h |
| HORVU.MOREX.r3.7HG0656980.1 | 1.220906 | -1.98412 | 3.997931 | 0.000199 | 0.023321 | 0.496533 | Transposon protein, putative, CACTA, En/Spm sub-class, expressed | root cap | 12 h |
| HORVU.MOREX.r3.5HG0494890.1 | -0.70304 | 5.120042 | -3.9763 | 0.000213 | 0.023449 | 0.485422 | Esterase/lipase/thioesterase family protein | root cap | 12 h |
| HORVU.MOREX.r3.4HG0384300.1 | -1.20119 | -1.67949 | -3.97879 | 0.000212 | 0.023449 | 0.466929 | RING finger protein | root cap | 12 h |
| HORVU.MOREX.r3.2HG0197550.1 | -0.85929 | -0.29036 | -3.96233 | 0.000223 | 0.0236 | 0.459076 | Transmembrane protein, putative | root cap | 12 h |
| HORVU.MOREX.r3.4HG0378900.1 | -0.52369 | 5.843019 | -3.97574 | 0.000214 | 0.023449 | 0.454073 | Kinase, putative | root cap | 12 h |
| HORVU.MOREX.r3.3HG0266950.1 | -0.69055 | 2.505445 | -3.9373 | 0.000242 | 0.024573 | 0.438803 | Calmodulin-binding protein | root cap | 12 h |
| HORVU.MOREX.r3.6HG0608380.1 | -0.47385 | 6.469706 | -3.98334 | 0.000209 | 0.023449 | 0.436209 | TBC domain containing protein | root cap | 12 h |
| HORVU.MOREX.r3.3HG0286640.1 | -0.92172 | 1.754903 | -3.99441 | 0.000201 | 0.023321 | 0.431322 | Chaperone protein dnaJ | root cap | 12 h |
| HORVU.MOREX.r3.1HG0080260.1 | -0.5082 | 3.804751 | -3.9187 | 0.000257 | 0.025457 | 0.40421 | Homogentisate phytyltransferase | root cap | 12 h |
| HORVU.MOREX.r3.2HG0138270.1 | -0.66143 | 3.593157 | -3.91957 | 0.000256 | 0.025457 | 0.403701 | Formin-like protein | root cap | 12 h |
| HORVU.MOREX.r3.4HG0389860.1 | -0.33912 | 6.138184 | -3.96395 | 0.000222 | 0.0236 | 0.403149 | S-ribonuclease binding protein | root cap | 12 h |
| HORVU.MOREX.r3.7HG0646880.1 | -0.48063 | 5.034739 | -3.94552 | 0.000236 | 0.024313 | 0.402696 | Hypersensitive-induced response protein | root cap | 12 h |
| HORVU.MOREX.r3.4HG0385780.1 | -0.53743 | 5.783595 | -3.9437 | 0.000237 | 0.024313 | 0.369497 | Mitogen-activated protein kinase | root cap | 12 h |
| HORVU.MOREX.r3.3HG0227860.1 | -0.38659 | 7.259658 | -3.96709 | 0.00022 | 0.0236 | 0.366759 | Katanin p60 ATPase-containing subunit, putative | root cap | 12 h |
| HORVU.MOREX.r3.3HG0245010.1 | -0.62494 | 4.432877 | -3.91558 | 0.000259 | 0.025485 | 0.352221 | Protein DJ-1 | root cap | 12 h |
| HORVU.MOREX.r3.7HG0733490.1 | -0.68112 | 5.748977 | -3.93253 | 0.000246 | 0.02483 | 0.339054 | Carbonic anhydrase | root cap | 12 h |
| HORVU.MOREX.r3.7HG0639840.1 | -1.98752 | -1.82989 | -4.0958 | 0.000145 | 0.020749 | 0.336419 | 3-ketoacyl-CoA synthase | root cap | 12 h |
| HORVU.MOREX.r3.7HG0743430.1 | -1.27167 | 3.39891 | -3.9427 | 0.000238 | 0.024313 | 0.333197 | Aldehyde oxidase, putative | root cap | 12 h |
| HORVU.MOREX.r3.6HG0577180.1 | -1.00308 | 3.196769 | -3.92969 | 0.000248 | 0.024934 | 0.333024 | ADP,ATP carrier protein | root cap | 12 h |
| HORVU.MOREX.r3.3HG0290130.1 | -0.76123 | 3.667474 | -3.92033 | 0.000255 | 0.025457 | 0.328571 | arginine N-methyltransferase, putative (DUF688) | root cap | 12 h |
| HORVU.MOREX.r3.3HG0220830.1 | -1.40493 | -0.11773 | -4.02603 | 0.000182 | 0.022855 | 0.312429 | Receptor-like kinase | root cap | 12 h |
| HORVU.MOREX.r3.3HG0311480.1 | -0.86407 | 2.639831 | -3.97974 | 0.000211 | 0.023449 | 0.293133 | Pollen Ole e 1 allergen/extensin | root cap | 12 h |
| HORVU.MOREX.r3.3HG0231460.1 | -0.77193 | 7.36546 | -3.96088 | 0.000224 | 0.0236 | 0.289136 | Endosomal targeting BRO1-like domain-containing protein | root cap | 12 h |
| HORVU.MOREX.r3.1HG0057290.1 | -0.74994 | 4.051245 | -3.91475 | 0.00026 | 0.025485 | 0.284943 | SBP (S-ribonuclease-binding protein) family protein | root cap | 12 h |
| HORVU.MOREX.r3.7HG0670910.1 | -0.42936 | 5.156942 | -3.8955 | 0.000277 | 0.02665 | 0.28404 | Aluminum activated malate transporter family protein | root cap | 12 h |

**Table S1** Continued.

| ID | logFC | AveExpr | t | P.Value | adj.P.Val | B | gene description | root zone | time point |
|---|---|---|---|---|---|---|---|---|---|
| HORVU.MOREX.r3.4HG0378750.1 | -0.57172 | 3.963125 | -3.88169 | 0.000289 | 0.027215 | 0.268539 | Transmembrane protein 87A | root cap | 12 h |
| HORVU.MOREX.r3.3HG0236350.1 | -0.8717 | 2.304206 | -3.89332 | 0.000279 | 0.026711 | 0.249493 | F-box family protein | root cap | 12 h |
| HORVU.MOREX.r3.5HG0455820.1 | -0.96891 | 2.941847 | -3.8853 | 0.000286 | 0.027027 | 0.244498 | WD-repeat protein, putative | root cap | 12 h |
| HORVU.MOREX.r3.2HG0211680.1 | -1.01881 | 6.061784 | -3.87344 | 0.000297 | 0.027661 | 0.238673 | Amine oxidase family protein | root cap | 12 h |
| HORVU.MOREX.r3.1HG0080790.1 | -1.09909 | 0.911279 | -3.95955 | 0.000225 | 0.0236 | 0.225685 | Alpha/beta-Hydrolases superfamily protein | root cap | 12 h |
| HORVU.MOREX.r3.1HG0014220.1 | -0.85398 | 1.384051 | -3.85311 | 0.000317 | 0.028759 | 0.218307 | Nuclear pore complex protein Nup98 | root cap | 12 h |
| HORVU.MOREX.r3.3HG0293070.1 | -0.63567 | 5.580025 | -3.89651 | 0.000276 | 0.02665 | 0.217215 | BnaA05g07570D protein | root cap | 12 h |
| HORVU.MOREX.r3.6HG0548800.1 | -1.17493 | -1.57973 | -3.85676 | 0.000313 | 0.028553 | 0.210522 | transcription factor IIIA | root cap | 12 h |
| HORVU.MOREX.r3.4HG0402190.1 | 1.926876 | -1.87054 | 3.871156 | 0.000299 | 0.027661 | 0.208213 | Flavin-containing monooxygenase | root cap | 12 h |
| HORVU.MOREX.r3.7HG0736000.1 | -0.74084 | 9.239643 | -4.01129 | 0.00019 | 0.023321 | 0.208037 | Chitinase | root cap | 12 h |
| HORVU.MOREX.r3.4HG0381820.1 | -0.61682 | 6.627467 | -3.91387 | 0.000261 | 0.025485 | 0.207245 | Fatty acid oxidation complex subunit alpha | root cap | 12 h |
| HORVU.MOREX.r3.3HG0273730.1 | -0.6509 | 6.291958 | -3.88742 | 0.000284 | 0.026969 | 0.184385 | Cytochrome P450 | root cap | 12 h |
| HORVU.MOREX.r3.4HG0413890.1 | -0.42415 | 5.790126 | -3.86633 | 0.000304 | 0.027944 | 0.155217 | Alpha/beta-Hydrolases superfamily protein | root cap | 12 h |
| HORVU.MOREX.r3.1HG0036390.1 | -0.49067 | 4.682709 | -3.84777 | 0.000322 | 0.029124 | 0.146666 | ABC1-like kinase | root cap | 12 h |
| HORVU.MOREX.r3.5HG0481030.1 | -0.80032 | 4.153293 | -3.83551 | 0.000335 | 0.02963 | 0.144373 | Alpha/beta-Hydrolases superfamily protein, putative | root cap | 12 h |
| HORVU.MOREX.r3.2HG0119150.1 | -0.55047 | 4.618655 | -3.83828 | 0.000332 | 0.029626 | 0.129811 | Cytochrome P450, putative | root cap | 12 h |
| HORVU.MOREX.r3.6HG0609220.1 | -0.66757 | 5.489346 | -3.86055 | 0.000309 | 0.028336 | 0.127946 | Diacylglycerol O-acyltransferase 2 | root cap | 12 h |
| HORVU.MOREX.r3.3HG0277400.1 | -0.60714 | 2.488136 | -3.8213 | 0.00035 | 0.030345 | 0.122893 | AMSH-like ubiquitin thioesterase 1 | root cap | 12 h |
| HORVU.MOREX.r3.5HG0462780.1 | -0.53228 | 7.18947 | -3.89016 | 0.000281 | 0.026857 | 0.121624 | Transporter, putative, expressed | root cap | 12 h |
| HORVU.MOREX.r3.5HG0468320.1 | -0.84554 | 2.415447 | -3.82534 | 0.000346 | 0.030163 | 0.107099 | Inosine-5'-monophosphate dehydrogenase | root cap | 12 h |
| HORVU.MOREX.r3.2HG0194790.1 | -0.51495 | 6.395123 | -3.87645 | 0.000294 | 0.027548 | 0.102939 | Amine oxidase family protein | root cap | 12 h |
| HORVU.MOREX.r3.7HG0708320.1 | -0.6994 | 4.603991 | -3.82453 | 0.000347 | 0.030163 | 0.093958 | Stromal cell-derived factor 2-like protein | root cap | 12 h |
| HORVU.MOREX.r3.1HG0059150.1 | -0.75857 | 5.002155 | -3.83296 | 0.000338 | 0.029743 | 0.091702 | Annexin | root cap | 12 h |
| HORVU.MOREX.r3.3HG0290180.1 | -0.61166 | 3.615358 | -3.842 | 0.000328 | 0.029534 | 0.084622 | Non-specific serine/threonine protein kinase | root cap | 12 h |
| HORVU.MOREX.r3.6HG0625490.1 | -0.50042 | 3.526573 | -3.79882 | 0.000376 | 0.031023 | 0.063541 | Cytochrome b5 reductase 4 | root cap | 12 h |
| HORVU.MOREX.r3.2HG0165290.2 | -0.78856 | 2.103536 | -3.80715 | 0.000367 | 0.030889 | 0.055339 | Initiation factor 4F subunit (DUF1350) | root cap | 12 h |
| HORVU.MOREX.r3.6HG0623730.1 | -0.9257 | 1.874032 | -3.79536 | 0.000381 | 0.031023 | 0.054276 | WEB family protein, chloroplastic | root cap | 12 h |
| HORVU.MOREX.r3.5HG0513000.1 | -0.45759 | 4.039212 | -3.80798 | 0.000366 | 0.030889 | 0.051138 | Transcription factor GTE1 | root cap | 12 h |
| HORVU.MOREX.r3.2HG0113590.1 | -0.56087 | 3.800345 | -3.79206 | 0.000385 | 0.031126 | 0.046603 | Zinc finger CCCH domain protein, putative | root cap | 12 h |
| HORVU.MOREX.r3.7HG0704050.1 | -0.74666 | 2.331323 | -3.7966 | 0.000379 | 0.031023 | 0.043818 | Myeloid leukemia factor 1 | root cap | 12 h |
| HORVU.MOREX.r3.6HG0630690.1 | -0.6132 | 3.335143 | -3.78218 | 0.000397 | 0.031576 | 0.020939 | Cyclin-dependent protein kinase inhibitor SMR9 | root cap | 12 h |
| HORVU.MOREX.r3.4HG0397280.1 | -0.54432 | 4.108256 | -3.79582 | 0.00038 | 0.031023 | 0.018921 | Leucine-rich repeat (LRR) family protein | root cap | 12 h |
| HORVU.MOREX.r3.3HG0275450.1 | -0.92711 | 1.752203 | -3.81151 | 0.000362 | 0.030802 | 0.011642 | Elongation factor | root cap | 12 h |
| HORVU.MOREX.r3.1HG0019440.1 | -0.43384 | 5.729684 | -3.83625 | 0.000334 | 0.02963 | 0.008411 | Vegetative cell wall protein gp1 | root cap | 12 h |
| HORVU.MOREX.r3.2HG0096100.1 | -0.86183 | 2.15401 | -3.80558 | 0.000368 | 0.030889 | -0.01152 | 3'-N-debenzoyl-2'-deoxytaxol N-benzoyltransferase | root cap | 12 h |
| HORVU.MOREX.r3.7HG0725940.1 | 1.092577 | -1.94583 | 3.774961 | 0.000406 | 0.031908 | -0.01633 | protein kinase family protein | root cap | 12 h |
| HORVU.MOREX.r3.1HG0054960.1 | -0.73435 | 4.410704 | -3.80468 | 0.000369 | 0.030889 | -0.02526 | Adipocyte plasma membrane-associated protein | root cap | 12 h |
| HORVU.MOREX.r3.7HG0702100.1 | -0.85535 | 0.820309 | -3.8289 | 0.000342 | 0.030001 | -0.02696 | Beta-1,3-N-acetylglucosaminyltransferase lunatic fringe | root cap | 12 h |
| HORVU.MOREX.r3.7HG0713610.1 | -0.48934 | 5.439775 | -3.79867 | 0.000377 | 0.031023 | -0.04321 | Core-2/I-branching beta-1,6-N-acetylglucosaminyltransferase family protein | root cap | 12 h |
| HORVU.MOREX.r3.7HG0720730.1 | -1.03623 | 0.190118 | -3.7542 | 0.000433 | 0.032842 | -0.0559 | Fructose-bisphosphate aldolase | root cap | 12 h |
| HORVU.MOREX.r3.7HG0682960.1 | -0.70444 | 2.362947 | -3.75725 | 0.000429 | 0.032842 | -0.06307 | Rhodanese/cell cycle control phosphatase superfamily protein | root cap | 12 h |

**Table S1** Continued.

| ID | logFC | AveExpr | t | P.Value | adj.P.Val | B | gene description | root zone | time point |
|---|---|---|---|---|---|---|---|---|---|
| HORVU.MOREX.r3.2HG0141900.1 | 0.671938 | 5.940265 | 3.790177 | 0.000387 | 0.031126 | -0.06509 | L-ascorbate oxidase-like protein | root cap | 12 h |
| HORVU.MOREX.r3.7HG0669600.1 | -0.64869 | 3.419699 | -3.77659 | 0.000404 | 0.031908 | -0.07011 | 2-oxoglutarate (2OG) and Fe(II)-dependent oxygenase superfamily protein, putativ | root cap | 12 h |
| HORVU.MOREX.r3.4HG0406900.1 | -0.57107 | 4.724038 | -3.77093 | 0.000411 | 0.03207 | -0.07189 | Ubiquitin carboxyl-terminal hydrolase, putative | root cap | 12 h |
| HORVU.MOREX.r3.3HG0219850.1 | 3.542475 | -1.36231 | 4.032459 | 0.000178 | 0.022855 | -0.07519 | Teneurin-2 | root cap | 12 h |
| HORVU.MOREX.r3.5HG0459810.1 | -0.56357 | 3.493172 | -3.75449 | 0.000433 | 0.032842 | -0.07563 | Protein kinase like protein | root cap | 12 h |
| HORVU.MOREX.r3.6HG0626330.1 | -0.66717 | 3.928068 | -3.81131 | 0.000362 | 0.030802 | -0.08094 | ENTH/ANTH/VHS superfamily protein | root cap | 12 h |
| HORVU.MOREX.r3.7HG0700790.1 | -0.61115 | 3.785936 | -3.74813 | 0.000442 | 0.033181 | -0.08254 | Erythronate-4-phosphate dehydrogenase family protein | root cap | 12 h |
| HORVU.MOREX.r3.4HG0340200.1 | -0.67051 | 4.710006 | -3.77511 | 0.000406 | 0.031908 | -0.08545 | RING finger and CHY zinc finger protein | root cap | 12 h |
| HORVU.MOREX.r3.3HG0235800.1 | -2.32714 | -3.45011 | -4.57065 | 2.95E-05 | 0.009444 | -0.09519 | E3 ubiquitin-protein ligase | root cap | 12 h |
| HORVU.MOREX.r3.1HG0070670.1 | -0.64127 | 4.791315 | -3.75743 | 0.000429 | 0.032842 | -0.09657 | Protein kinase-like protein | root cap | 12 h |
| HORVU.MOREX.r3.4HG0338810.1 | -0.5886 | 4.132368 | -3.77256 | 0.000409 | 0.032027 | -0.10314 | Inositol-tetrakisphosphate 1-kinase | root cap | 12 h |
| HORVU.MOREX.r3.6HG0600190.1 | -0.67023 | 3.392812 | -3.74631 | 0.000444 | 0.033181 | -0.10353 | UPF0565 protein C2orf69 | root cap | 12 h |
| HORVU.MOREX.r3.4HG0393500.1 | -0.83014 | 0.752818 | -3.75422 | 0.000433 | 0.032842 | -0.1104 | #N/A | root cap | 12 h |
| HORVU.MOREX.r3.1HG0019240.1 | -0.9857 | 2.336885 | -3.78663 | 0.000391 | 0.031354 | -0.11077 | N-acetyltransferase, putative | root cap | 12 h |
| HORVU.MOREX.r3.2HG0128270.1 | -0.77611 | 2.338352 | -3.74365 | 0.000448 | 0.033181 | -0.11265 | Zinc finger family protein | root cap | 12 h |
| HORVU.MOREX.r3.2HG0131330.1 | -0.65025 | 7.99791 | -3.83978 | 0.00033 | 0.029613 | -0.12271 | Transporter-related family protein | root cap | 12 h |
| HORVU.MOREX.r3.6HG0591720.1 | -0.48572 | 5.117239 | -3.75186 | 0.000436 | 0.032929 | -0.13432 | Glucose-6-phosphate 1-dehydrogenase | root cap | 12 h |
| HORVU.MOREX.r3.4HG0405830.1 | -0.73613 | 2.230818 | -3.729 | 0.000469 | 0.033952 | -0.1371 | cold regulated protein 27 | root cap | 12 h |
| HORVU.MOREX.r3.7HG0635410.1 | -1.13571 | 0.730461 | -3.79496 | 0.000381 | 0.031023 | -0.15461 | Protein yippee-like | root cap | 12 h |
| HORVU.MOREX.r3.4HG0389580.1 | -0.43409 | 5.135089 | -3.74423 | 0.000447 | 0.033181 | -0.15657 | Mitochondrial carrier family | root cap | 12 h |
| HORVU.MOREX.r3.3HG0242790.1 | -0.49919 | 5.85397 | -3.76035 | 0.000425 | 0.032842 | -0.1629 | Cell number regulator 8 | root cap | 12 h |
| HORVU.MOREX.r3.1HG0077710.1 | -0.75035 | -0.77894 | -3.72225 | 0.000479 | 0.03425 | -0.17275 | Glycosyltransferase | root cap | 12 h |
| HORVU.MOREX.r3.6HG0603150.1 | -0.7566 | 4.113219 | -3.75817 | 0.000428 | 0.032842 | -0.18072 | Argonaute | root cap | 12 h |
| HORVU.MOREX.r3.3HG0295840.1 | -0.58434 | 2.964243 | -3.70959 | 0.000498 | 0.034906 | -0.18264 | Dehydrogenase-like protein | root cap | 12 h |
| HORVU.MOREX.r3.1HG0077230.1 | -0.7893 | 3.396902 | -3.73992 | 0.000453 | 0.033451 | -0.18389 | VQ motif-containing protein | root cap | 12 h |
| HORVU.MOREX.r3.5HG0516730.1 | -0.80689 | 2.88293 | -3.714 | 0.000492 | 0.034545 | -0.18469 | Peroxisomal membrane protein 13 | root cap | 12 h |
| HORVU.MOREX.r3.2HG0125510.1 | -1.40719 | 0.997629 | -3.71554 | 0.000489 | 0.034498 | -0.18493 | SPX domain-containing protein | root cap | 12 h |
| HORVU.MOREX.r3.5HG0455240.1 | -0.78574 | 2.696541 | -3.74491 | 0.000446 | 0.033181 | -0.18736 | O-fucosyltransferase family protein | root cap | 12 h |
| HORVU.MOREX.r3.4HG0390580.1 | -0.73483 | 2.573232 | -3.71765 | 0.000486 | 0.034388 | -0.18741 | #N/A | root cap | 12 h |
| HORVU.MOREX.r3.7HG0725210.3 | -0.69578 | 5.13305 | -3.72919 | 0.000469 | 0.033952 | -0.19062 | Lysophosphatidylcholine acyltransferase 1 | root cap | 12 h |
| HORVU.MOREX.r3.1HG0081120.1 | -1.44593 | 1.005477 | -3.78193 | 0.000397 | 0.031576 | -0.19634 | DNA mismatch repair protein MutL | root cap | 12 h |
| HORVU.MOREX.r3.4HG0392440.1 | -0.6496 | 4.50652 | -3.73128 | 0.000466 | 0.033952 | -0.20368 | Chaperone protein dnaJ | root cap | 12 h |
| HORVU.MOREX.r3.2HG0174560.1 | -0.50891 | 4.366463 | -3.70462 | 0.000506 | 0.03504 | -0.20716 | Ras-related protein | root cap | 12 h |
| HORVU.MOREX.r3.5HG0504340.1 | -0.67217 | 3.865477 | -3.69679 | 0.000519 | 0.035084 | -0.21329 | Mediator of RNA polymerase II transcription subunit 8 | root cap | 12 h |
| HORVU.MOREX.r3.3HG0295550.1 | -0.51865 | 4.950607 | -3.72839 | 0.00047 | 0.033952 | -0.21562 | Stomatal closure-related actin-binding protein 1 | root cap | 12 h |
| HORVU.MOREX.r3.1HG0062030.1 | -1.45568 | -2.69359 | -3.96694 | 0.00022 | 0.0236 | -0.22281 | Dehydration responsive element binding transcription factor | root cap | 12 h |
| HORVU.MOREX.r3.3HG0303670.1 | -0.86762 | 1.411379 | -3.69402 | 0.000523 | 0.035084 | -0.22366 | RNAse E/G-like protein | root cap | 12 h |
| HORVU.MOREX.r3.5HG0445510.1 | -0.47986 | 5.10535 | -3.70146 | 0.000511 | 0.03504 | -0.22612 | Acyl-CoA-binding domain-containing protein 4 | root cap | 12 h |
| HORVU.MOREX.r3.5HG0520850.1 | -0.54163 | 4.065206 | -3.69593 | 0.00052 | 0.035084 | -0.2347 | Kinase, putative | root cap | 12 h |
| HORVU.MOREX.r3.7HG0660830.1 | -0.39304 | 5.341821 | -3.72567 | 0.000474 | 0.034004 | -0.24077 | Ubiquitin-conjugating enzyme, E2 | root cap | 12 h |
| HORVU.MOREX.r3.5HG0464600.1 | -0.6616 | 3.460343 | -3.69436 | 0.000523 | 0.035084 | -0.24739 | Dual specificity protein phosphatase, putative | root cap | 12 h |

**Table S1** Continued.

| ID | logFC | AveExpr | t | P.Value | adj.P.Val | B | gene description | root zone | time point |
|---|---|---|---|---|---|---|---|---|---|
| HORVU.MOREX.r3.5HG0488620.1 | -0.58424 | 3.531829 | -3.73403 | 0.000462 | 0.033952 | -0.25245 | O-fucosyltransferase family protein | root cap | 12 h |
| HORVU.MOREX.r3.2HG0181020.1 | -0.60082 | 7.283571 | -3.81338 | 0.000359 | 0.030802 | -0.25633 | #N/A | root cap | 12 h |
| HORVU.MOREX.r3.2HG0168640.1 | -0.49525 | 4.660135 | -3.7204 | 0.000482 | 0.03433 | -0.26514 | F-box family protein | root cap | 12 h |
| HORVU.MOREX.r3.5HG0497170.1 | -0.56094 | 5.352547 | -3.73035 | 0.000467 | 0.033952 | -0.2746 | Pimeloyl-[acyl-carrier protein] methyl ester esterase | root cap | 12 h |
| HORVU.MOREX.r3.5HG0507370.1 | -0.57749 | 4.402293 | -3.68359 | 0.000541 | 0.035458 | -0.27968 | Protein kinase | root cap | 12 h |
| HORVU.MOREX.r3.7HG0714350.1 | -0.92413 | 3.30207 | -3.67003 | 0.000564 | 0.036145 | -0.28594 | Acyl-[acyl-carrier-protein] hydrolase | root cap | 12 h |
| HORVU.MOREX.r3.7HG0739570.1 | 2.162518 | 1.204008 | 4.026302 | 0.000181 | 0.022855 | -0.30853 | cotton fiber protein | root cap | 12 h |
| HORVU.MOREX.r3.2HG0192920.1 | -0.81411 | 0.782762 | -3.68994 | 0.00053 | 0.035204 | -0.31126 | Xyloglucan endotransglucosylase/hydrolase | root cap | 12 h |
| HORVU.MOREX.r3.5HG0463370.1 | -1.10991 | 2.525221 | -3.6797 | 0.000547 | 0.035604 | -0.3205 | Zinc finger family protein | root cap | 12 h |
| HORVU.MOREX.r3.6HG0570850.1 | -0.60087 | 5.628875 | -3.70163 | 0.000511 | 0.03504 | -0.32198 | Protein kinase | root cap | 12 h |
| HORVU.MOREX.r3.4HG0356780.1 | -0.5458 | 4.400609 | -3.66625 | 0.000571 | 0.036251 | -0.32206 | weak chloroplast movement under blue light protein (DUF827) | root cap | 12 h |
| HORVU.MOREX.r3.4HG0353950.1 | -1.63282 | -1.81142 | -3.80394 | 0.00037 | 0.030889 | -0.3261 | ubiquitin carboxyl-terminal hydrolase | root cap | 12 h |
| HORVU.MOREX.r3.5HG0479000.1 | -1.54475 | 3.962906 | -3.66329 | 0.000576 | 0.036361 | -0.32768 | Alcohol dehydrogenase, putative | root cap | 12 h |
| HORVU.MOREX.r3.5HG0471940.1 | -0.36131 | 6.025728 | -3.70816 | 0.000501 | 0.034928 | -0.33146 | Myb family transcription factor APL | root cap | 12 h |
| HORVU.MOREX.r3.6HG0592790.1 | -0.37766 | 4.902986 | -3.69932 | 0.000515 | 0.03504 | -0.33694 | Nicotinamidase 1 | root cap | 12 h |
| HORVU.MOREX.r3.5HG0458770.1 | 0.716016 | 5.47733 | 3.671716 | 0.000561 | 0.036145 | -0.34979 | B3 domain-containing protein | root cap | 12 h |
| HORVU.MOREX.r3.3HG0256430.1 | -0.65593 | 4.497782 | -3.69869 | 0.000516 | 0.03504 | -0.35008 | Hyp O-arabinosyltransferase-like protein | root cap | 12 h |
| HORVU.MOREX.r3.4HG0406980.1 | -0.91545 | 2.545348 | -3.67033 | 0.000563 | 0.036145 | -0.35059 | Ras-related protein | root cap | 12 h |
| HORVU.MOREX.r3.7HG0713160.1 | -0.61179 | 6.232926 | -3.69874 | 0.000516 | 0.03504 | -0.35862 | Acyl-coenzyme A oxidase | root cap | 12 h |
| HORVU.MOREX.r3.3HG0275500.1 | -1.65646 | -1.87206 | -3.75387 | 0.000434 | 0.032842 | -0.36045 | Flavin-containing monooxygenase | root cap | 12 h |
| HORVU.MOREX.r3.3HG0288830.1 | -0.33014 | 5.399232 | -3.68242 | 0.000543 | 0.035458 | -0.36367 | MACPF domain protein | root cap | 12 h |
| HORVU.MOREX.r3.5HG0496740.1 | -1.66291 | 0.004024 | -3.64074 | 0.000618 | 0.037835 | -0.36974 | Agmatine coumaroyltransferase-2 | root cap | 12 h |
| HORVU.MOREX.r3.5HG0471560.1 | -0.59593 | 3.339465 | -3.68612 | 0.000536 | 0.035389 | -0.3867 | Receptor kinase | root cap | 12 h |
| HORVU.MOREX.r3.5HG0527770.1 | -0.79643 | 1.038275 | -3.72569 | 0.000474 | 0.034004 | -0.39064 | GDP-mannose transporter | root cap | 12 h |
| HORVU.MOREX.r3.6HG0596240.1 | -1.08231 | 0.960124 | -3.69304 | 0.000525 | 0.035084 | -0.39262 | RNA-binding protein | root cap | 12 h |
| HORVU.MOREX.r3.1HG0083840.1 | -0.49752 | 3.99732 | -3.63475 | 0.000629 | 0.037866 | -0.39632 | Serine-rich adhesin for platelets-like protein | root cap | 12 h |
| HORVU.MOREX.r3.7HG0723770.1 | -0.56504 | 6.493213 | -3.68898 | 0.000532 | 0.035204 | -0.39828 | BZIP transcription factor protein | root cap | 12 h |
| HORVU.MOREX.r3.3HG0281780.1 | -0.66413 | 4.822877 | -3.66202 | 0.000578 | 0.036393 | -0.40001 | Protein transport protein Sec61 subunit alpha | root cap | 12 h |
| HORVU.MOREX.r3.2HG0172180.1 | -0.79022 | 4.401326 | -3.64374 | 0.000612 | 0.037598 | -0.41143 | Ring finger protein, putative | root cap | 12 h |
| HORVU.MOREX.r3.4HG0395120.1 | -0.65026 | 2.404038 | -3.62219 | 0.000654 | 0.038466 | -0.42205 | Mitochondrial carrier family | root cap | 12 h |
| HORVU.MOREX.r3.2HG0127370.1 | -0.60917 | 7.932229 | -3.81482 | 0.000358 | 0.030802 | -0.42492 | Alpha-1,4-glucan-protein synthase [UDP-forming] 1 | root cap | 12 h |
| HORVU.MOREX.r3.3HG0257120.1 | -0.7598 | 1.984822 | -3.62405 | 0.00065 | 0.038466 | -0.42492 | Histone-lysine N-methyltransferase | root cap | 12 h |
| HORVU.MOREX.r3.4HG0343380.1 | -0.43066 | 5.245167 | -3.67433 | 0.000556 | 0.036019 | -0.43512 | ATP-dependent (S)-NAD(P)H-hydrate dehydratase | root cap | 12 h |
| HORVU.MOREX.r3.1HG0071320.1 | -1.03784 | 0.645871 | -3.61032 | 0.000679 | 0.039422 | -0.44663 | Transcription elongation factor SPT5 | root cap | 12 h |
| HORVU.MOREX.r3.7HG0748320.1 | -0.75144 | 3.37477 | -3.60771 | 0.000684 | 0.039547 | -0.4568 | Receptor-like kinase | root cap | 12 h |
| HORVU.MOREX.r3.2HG0184620.1 | -0.78739 | 4.117573 | -3.63498 | 0.000629 | 0.037866 | -0.45774 | Hsp70-Hsp90 organizing protein 1 | root cap | 12 h |
| HORVU.MOREX.r3.4HG0347330.1 | -0.55857 | 6.923566 | -3.70722 | 0.000502 | 0.034928 | -0.46016 | Aldehyde dehydrogenase | root cap | 12 h |
| HORVU.MOREX.r3.2HG0180040.2 | -1.10367 | -2.17606 | -3.68259 | 0.000542 | 0.035458 | -0.46236 | Trichome birefringence | root cap | 12 h |
| HORVU.MOREX.r3.3HG0300080.1 | -0.7491 | 5.888104 | -3.66448 | 0.000574 | 0.036339 | -0.46321 | Kinase | root cap | 12 h |
| HORVU.MOREX.r3.2HG0113410.1 | -0.52509 | 4.641504 | -3.6395 | 0.00062 | 0.037866 | -0.46556 | Protein AIG1 | root cap | 12 h |
| HORVU.MOREX.r3.2HG0113860.1 | -1.20673 | 2.66732 | -3.61625 | 0.000666 | 0.039067 | -0.47095 | Uveal autoantigen with coiled-coil domains and ankyrin repeats isoform 2 | root cap | 12 h |

**Table S1** Continued.

| ID | logFC | AveExpr | t | P.Value | adj.P.Val | B | gene description | root zone | time point |
|---|---|---|---|---|---|---|---|---|---|
| HORVU.MOREX.r3.4HG0365970.1 | -0.49877 | 5.950027 | -3.65757 | 0.000586 | 0.036788 | -0.47612 | Phosphatidate phosphatase, Lipin | root cap | 12 h |
| HORVU.MOREX.r3.4HG0417410.1 | -1.34956 | 0.915973 | -3.71816 | 0.000485 | 0.034388 | -0.48195 | Cold regulated protein 27 | root cap | 12 h |
| HORVU.MOREX.r3.7HG0677380.1 | -0.55003 | 5.818397 | -3.70162 | 0.000511 | 0.03504 | -0.48413 | Alpha-glucan water dikinase | root cap | 12 h |
| HORVU.MOREX.r3.6HG0541580.1 | 0.269867 | 8.008196 | 3.688832 | 0.000532 | 0.035204 | -0.49675 | Aspartate aminotransferase | root cap | 12 h |
| HORVU.MOREX.r3.7HG0733020.1 | -0.53077 | 5.546828 | -3.64445 | 0.000611 | 0.037598 | -0.52062 | Polyubiquitin | root cap | 12 h |
| HORVU.MOREX.r3.2HG0135590.1 | -0.63404 | 2.26601 | -3.57977 | 0.000746 | 0.041215 | -0.53028 | Protein THYLAKOID FORMATION1, chloroplastic | root cap | 12 h |
| HORVU.MOREX.r3.5HG0506790.1 | -0.68378 | 5.002887 | -3.6228 | 0.000653 | 0.038466 | -0.53728 | Calcineurin-like metallo-phosphoesterase superfamily | root cap | 12 h |
| HORVU.MOREX.r3.1HG0051740.1 | -0.98734 | 3.433417 | -3.63114 | 0.000636 | 0.037993 | -0.54521 | Glutathione s-transferase, putative | root cap | 12 h |
| HORVU.MOREX.r3.2HG0187980.1 | -0.58131 | 2.469272 | -3.57638 | 0.000754 | 0.041215 | -0.54754 | Transmembrane protein 184C | root cap | 12 h |
| HORVU.MOREX.r3.4HG0407120.1 | -0.57437 | 4.965682 | -3.58914 | 0.000724 | 0.040619 | -0.55774 | methyl-coenzyme M reductase II subunit gamma, putative (DUF3741) | root cap | 12 h |
| HORVU.MOREX.r3.1HG0092900.1 | -1.54857 | -2.96229 | -3.99598 | 0.0002 | 0.023321 | -0.55851 | C3HC4-type RING finger-containing protein | root cap | 12 h |
| HORVU.MOREX.r3.7HG0721080.1 | -0.73373 | 4.617684 | -3.60194 | 0.000696 | 0.039695 | -0.56145 | Zinc finger, B-box | root cap | 12 h |
| HORVU.MOREX.r3.2HG0194600.1 | -0.76373 | 1.177222 | -3.56657 | 0.000777 | 0.041944 | -0.56419 | transmembrane protein, putative (DUF594) | root cap | 12 h |
| HORVU.MOREX.r3.1HG0080420.1 | -0.65181 | 3.160969 | -3.60561 | 0.000689 | 0.039589 | -0.56764 | cotton fiber-like protein (DUF761) | root cap | 12 h |
| HORVU.MOREX.r3.5HG0455770.1 | -0.81391 | 0.761558 | -3.64568 | 0.000608 | 0.037597 | -0.57433 | S-adenosyl-L-methionine-dependent methyltransferases superfamily protein | root cap | 12 h |
| HORVU.MOREX.r3.4HG0391050.1 | -0.67069 | 2.336797 | -3.55928 | 0.000794 | 0.042658 | -0.58446 | Protein FAR1-RELATED SEQUENCE 7 | root cap | 12 h |
| HORVU.MOREX.r3.2HG0119920.1 | -0.67111 | 6.226589 | -3.62221 | 0.000654 | 0.038466 | -0.58496 | protein kinase family protein | root cap | 12 h |
| HORVU.MOREX.r3.2HG0187170.1 | -0.44887 | 6.368636 | -3.63807 | 0.000623 | 0.037866 | -0.59093 | S-adenosyl-L-methionine-dependent methyltransferases superfamily protein | root cap | 12 h |
| HORVU.MOREX.r3.2HG0163200.1 | -0.52 | 5.22535 | -3.60502 | 0.00069 | 0.039589 | -0.59777 | Yellow stripe-like transporter 12 | root cap | 12 h |
| HORVU.MOREX.r3.7HG0723280.1 | -1.88397 | -2.61155 | -3.90003 | 0.000273 | 0.026515 | -0.60034 | PLATZ transcription factor family protein | root cap | 12 h |
| HORVU.MOREX.r3.4HG0381750.1 | -0.60952 | 5.381044 | -3.63085 | 0.000637 | 0.037993 | -0.60124 | Calmodulin | root cap | 12 h |
| HORVU.MOREX.r3.6HG0555250.1 | -0.6987 | 0.376943 | -3.55405 | 0.000807 | 0.042918 | -0.60235 | Esterase/lipase/thioesterase-like protein | root cap | 12 h |
| HORVU.MOREX.r3.7HG0721660.1 | -0.70582 | 4.153249 | -3.61313 | 0.000673 | 0.039221 | -0.60262 | Serine/threonine-protein kinase mTOR | root cap | 12 h |
| HORVU.MOREX.r3.5HG0440700.1 | -0.44424 | 5.639406 | -3.59786 | 0.000705 | 0.039869 | -0.60436 | Diacylglycerol kinase | root cap | 12 h |
| HORVU.MOREX.r3.5HG0512430.1 | -0.81029 | 3.653637 | -3.57565 | 0.000755 | 0.041215 | -0.60756 | Phosphatase 2C family protein | root cap | 12 h |
| HORVU.MOREX.r3.7HG0709580.1 | -1.15886 | -0.69882 | -3.5547 | 0.000805 | 0.042918 | -0.60793 | ADP-ribosylation factor family protein | root cap | 12 h |
| HORVU.MOREX.r3.7HG0676620.1 | -1.44465 | -0.20992 | -3.65387 | 0.000593 | 0.0371 | -0.61199 | Mitogen activated protein kinase kinase kinase-like protein | root cap | 12 h |
| HORVU.MOREX.r3.3HG0256160.1 | -0.5031 | 4.949529 | -3.61459 | 0.00067 | 0.039157 | -0.61408 | Cysteine proteinase inhibitor | root cap | 12 h |
| HORVU.MOREX.r3.1HG0031820.1 | -0.94472 | 1.763108 | -3.54613 | 0.000827 | 0.04339 | -0.61722 | Calmodulin | root cap | 12 h |
| HORVU.MOREX.r3.7HG0725140.1 | 0.628602 | 6.172211 | 3.561705 | 0.000788 | 0.042454 | -0.63299 | Lipid transfer protein | root cap | 12 h |
| HORVU.MOREX.r3.3HG0235720.1 | -1.47724 | 3.057636 | -3.55366 | 0.000808 | 0.042918 | -0.63522 | FMN-dependent NADPH-azoreductase | root cap | 12 h |
| HORVU.MOREX.r3.7HG0656220.1 | -0.58305 | 4.216456 | -3.60107 | 0.000698 | 0.039695 | -0.63526 | Protein DETOXIFICATION | root cap | 12 h |
| HORVU.MOREX.r3.5HG0461790.1 | -1.02717 | -2.4609 | -3.6297 | 0.000639 | 0.038018 | -0.65122 | Ring finger protein, putative | root cap | 12 h |
| HORVU.MOREX.r3.7HG0635330.1 | -1.73805 | 0.867741 | -3.58994 | 0.000723 | 0.040619 | -0.65181 | Protein TolB | root cap | 12 h |
| HORVU.MOREX.r3.2HG0168300.1 | -1.16049 | 0.915123 | -3.63276 | 0.000633 | 0.03799 | -0.65266 | Helicase/SANT-associated, putative isoform 5 | root cap | 12 h |
| HORVU.MOREX.r3.7HG0705130.1 | 1.008475 | 2.637633 | 3.536961 | 0.00085 | 0.043951 | -0.65333 | 3'-N-debenzoyl-2'-deoxytaxol N-benzoyltransferase | root cap | 12 h |
| HORVU.MOREX.r3.1HG0064310.1 | -0.66177 | 4.083368 | -3.55421 | 0.000807 | 0.042918 | -0.65446 | Ribosomal protein L34e superfamily protein | root cap | 12 h |
| HORVU.MOREX.r3.2HG0169000.1 | -1.43216 | 2.146274 | -3.59491 | 0.000712 | 0.040123 | -0.65812 | arginine N-methyltransferase, putative (DUF688) | root cap | 12 h |
| HORVU.MOREX.r3.5HG0481120.1 | -0.98231 | -1.50366 | -3.53107 | 0.000866 | 0.043974 | -0.66068 | RING/U-box superfamily protein | root cap | 12 h |
| HORVU.MOREX.r3.5HG0457960.1 | -0.33819 | 5.424342 | -3.57638 | 0.000754 | 0.041215 | -0.66136 | Tubby-like F-box protein | root cap | 12 h |
| HORVU.MOREX.r3.4HG0400830.1 | -2.27409 | 0.255399 | -4.04009 | 0.000173 | 0.022679 | -0.66164 | ATP-dependent DNA helicase | root cap | 12 h |

**Table S1** Continued.

| ID | logFC | AveExpr | t | P.Value | adj.P.Val | B | gene description | root zone | time point |
|---|---|---|---|---|---|---|---|---|---|
| HORVU.MOREX.r3.UnG0753160.1 | -0.61148 | 7.774397 | -3.65234 | 0.000596 | 0.037163 | -0.66494 | Sucrose-phosphate synthase | root cap | 12 h |
| HORVU.MOREX.r3.5HG0460300.1 | -0.85938 | 4.047693 | -3.52554 | 0.000881 | 0.044002 | -0.67467 | Rho GTPase-activating protein | root cap | 12 h |
| HORVU.MOREX.r3.2HG0110640.1 | -0.31371 | 5.40065 | -3.57589 | 0.000755 | 0.041215 | -0.68016 | transcription factor-like protein | root cap | 12 h |
| HORVU.MOREX.r3.7HG0719840.1 | -0.68762 | 1.628095 | -3.52688 | 0.000877 | 0.044002 | -0.68261 | DUF4050 family protein | root cap | 12 h |
| HORVU.MOREX.r3.4HG0405330.1 | 1.16705 | -1.47892 | 3.532493 | 0.000862 | 0.043974 | -0.68428 | Cytochrome P450, putative | root cap | 12 h |
| HORVU.MOREX.r3.7HG0750850.1 | 3.805186 | -0.5604 | 3.679068 | 0.000548 | 0.035604 | -0.68574 | Cellulose synthase-like protein | root cap | 12 h |
| HORVU.MOREX.r3.5HG0496780.1 | -0.51626 | 4.42219 | -3.54809 | 0.000822 | 0.043324 | -0.6911 | C2 domain-containing protein / GRAM domain-containing protein | root cap | 12 h |
| HORVU.MOREX.r3.7HG0682660.1 | -1.93266 | 1.152818 | -3.94214 | 0.000238 | 0.024313 | -0.69152 | Basic helix-loop-helix transcription factor | root cap | 12 h |
| HORVU.MOREX.r3.2HG0164440.1 | -0.62514 | 4.939875 | -3.59874 | 0.000703 | 0.039869 | -0.69202 | glycosyltransferase family exostosin protein | root cap | 12 h |
| HORVU.MOREX.r3.6HG0634140.1 | -0.82326 | 2.257992 | -3.51938 | 0.000897 | 0.044578 | -0.6938 | Disease resistance protein (NBS-LRR class) family | root cap | 12 h |
| HORVU.MOREX.r3.7HG0677710.1 | -1.33103 | 1.564676 | -3.53791 | 0.000848 | 0.043951 | -0.69393 | BTB/POZ domain-containing protein | root cap | 12 h |
| HORVU.MOREX.r3.1HG0056540.1 | -0.97443 | 2.503604 | -3.51859 | 0.000899 | 0.044578 | -0.6947 | Zinc finger family protein | root cap | 12 h |
| HORVU.MOREX.r3.2HG0169420.1 | -0.84193 | 4.736887 | -3.57953 | 0.000746 | 0.041215 | -0.71092 | Regulator of chromosome condensation (RCC1) family protein | root cap | 12 h |
| HORVU.MOREX.r3.1HG0003270.1 | 1.634235 | -2.89111 | 3.870916 | 0.000299 | 0.027661 | -0.71502 | Core-2/I-branching beta-1,6-N-acetylglucosaminyltransferase family protein | root cap | 12 h |
| HORVU.MOREX.r3.2HG0162620.1 | -1.58942 | -1.43824 | -3.66818 | 0.000567 | 0.036145 | -0.71743 | Auxin response factor | root cap | 12 h |
| HORVU.MOREX.r3.1HG0089180.1 | -0.60941 | 5.019103 | -3.60369 | 0.000693 | 0.039595 | -0.71796 | Cold acclimation protein | root cap | 12 h |
| HORVU.MOREX.r3.3HG0295500.2 | -1.10063 | 1.304608 | -3.54747 | 0.000823 | 0.043324 | -0.72064 | UDP-glycosyltransferase | root cap | 12 h |
| HORVU.MOREX.r3.3HG0252100.1 | -0.77278 | 1.973895 | -3.64859 | 0.000603 | 0.037371 | -0.72096 | Stigma-specific Stig1 family protein, putative | root cap | 12 h |
| HORVU.MOREX.r3.3HG0305840.1 | -0.79566 | 0.764684 | -3.50679 | 0.000932 | 0.045382 | -0.72488 | Calcium sensing receptor, chloroplastic | root cap | 12 h |
| HORVU.MOREX.r3.4HG0394470.1 | -1.1303 | 2.141349 | -3.54143 | 0.000839 | 0.043796 | -0.72507 | Myb/SANT-like DNA-binding domain protein | root cap | 12 h |
| HORVU.MOREX.r3.1HG0082670.1 | -0.74133 | 1.71129 | -3.51438 | 0.000911 | 0.044937 | -0.7404 | 26S protease regulatory subunit 7 | root cap | 12 h |
| HORVU.MOREX.r3.3HG0320710.1 | -0.57633 | 3.737114 | -3.52803 | 0.000874 | 0.044002 | -0.74378 | SNF2 domain-containing protein / helicase domain-containing protein / F-box fam | root cap | 12 h |
| HORVU.MOREX.r3.7HG0644160.1 | -0.92814 | -1.60125 | -3.49636 | 0.000962 | 0.045846 | -0.74775 | GDSL esterase/lipase | root cap | 12 h |
| HORVU.MOREX.r3.3HG0300580.1 | -1.05615 | 5.442061 | -3.50489 | 0.000938 | 0.045382 | -0.74775 | Glucan endo-1,3-beta-glucosidase, putative | root cap | 12 h |
| HORVU.MOREX.r3.2HG0184350.1 | -0.58181 | 3.027133 | -3.49691 | 0.000961 | 0.045846 | -0.74826 | Cytochrome P450 | root cap | 12 h |
| HORVU.MOREX.r3.1HG0095220.1 | -0.78131 | 2.300388 | -3.53186 | 0.000864 | 0.043974 | -0.74959 | Pleckstrin-like (PH) domain protein | root cap | 12 h |
| HORVU.MOREX.r3.3HG0231430.1 | -0.37803 | 4.799815 | -3.5253 | 0.000881 | 0.044002 | -0.7522 | #N/A | root cap | 12 h |
| HORVU.MOREX.r3.5HG0517370.1 | -2.33221 | -3.29058 | -4.10125 | 0.000142 | 0.020673 | -0.76483 | Zinc finger A20 and AN1 domain-containing stress-associated protein | root cap | 12 h |
| HORVU.MOREX.r3.3HG0303190.1 | -0.40578 | 8.19693 | -3.60965 | 0.00068 | 0.039422 | -0.76893 | UDP-glucuronate decarboxylase protein 1 | root cap | 12 h |
| HORVU.MOREX.r3.6HG0560030.1 | -0.5275 | 3.970473 | -3.511 | 0.00092 | 0.045293 | -0.76916 | DUF1644 family protein | root cap | 12 h |
| HORVU.MOREX.r3.7HG0700200.1 | -0.45464 | 4.298926 | -3.52818 | 0.000873 | 0.044002 | -0.77372 | Oxysterol-binding family protein | root cap | 12 h |
| HORVU.MOREX.r3.5HG0494470.1 | -0.46446 | 3.993787 | -3.49723 | 0.00096 | 0.045846 | -0.78799 | Transmembrane emp24 domain-containing protein | root cap | 12 h |
| HORVU.MOREX.r3.7HG0678310.1 | -0.6653 | 3.452075 | -3.55306 | 0.000809 | 0.042918 | -0.79021 | Protein DETOXIFICATION | root cap | 12 h |
| HORVU.MOREX.r3.7HG0708910.1 | -0.58414 | -0.96712 | -3.51695 | 0.000904 | 0.044693 | -0.79655 | NAC domain-containing protein, putative | root cap | 12 h |
| HORVU.MOREX.r3.1HG0014560.1 | -0.8326 | 3.027709 | -3.49471 | 0.000967 | 0.045942 | -0.79656 | Peptide-N4-(N-acetyl-beta-glucosaminyl)asparagine amidase A | root cap | 12 h |
| HORVU.MOREX.r3.6HG0550260.1 | 0.403208 | 5.617383 | 3.536 | 0.000853 | 0.04397 | -0.79944 | aminoalcoholphosphotransferase 1 | root cap | 12 h |
| HORVU.MOREX.r3.7HG0733550.1 | 1.58838 | -0.67914 | 3.473047 | 0.001033 | 0.04717 | -0.81074 | Early nodulin-like protein | root cap | 12 h |
| HORVU.MOREX.r3.2HG0141980.1 | 0.674343 | 5.307117 | 3.480704 | 0.001009 | 0.04706 | -0.81299 | Glucan endo-1,3-beta-glucosidase 3 | root cap | 12 h |
| HORVU.MOREX.r3.3HG0296420.1 | -0.46375 | 7.295023 | -3.57267 | 0.000762 | 0.041484 | -0.8142 | 91A protein | root cap | 12 h |
| HORVU.MOREX.r3.6HG0597880.1 | -0.57049 | 4.866427 | -3.52524 | 0.000881 | 0.044002 | -0.81546 | heat-inducible transcription repressor (DUF639) | root cap | 12 h |
| HORVU.MOREX.r3.2HG0192180.1 | 0.227019 | 9.138579 | 3.636095 | 0.000627 | 0.037866 | -0.8169 | 26S proteasome non-ATPase regulatory subunit 1 | root cap | 12 h |

**Table S1** Continued.

| ID | logFC | AveExpr | t | P.Value | adj.P.Val | B | gene description | root zone | time point |
|---|---|---|---|---|---|---|---|---|---|
| HORVU.MOREX.r3.7HG0671350.1 | -0.41595 | 4.825604 | -3.50651 | 0.000933 | 0.045382 | -0.81861 | Potassium transporter | root cap | 12 h |
| HORVU.MOREX.r3.6HG0617830.1 | -0.90232 | 4.975661 | -3.47789 | 0.001018 | 0.047149 | -0.822 | Cyclin-dependent kinase inhibitor | root cap | 12 h |
| HORVU.MOREX.r3.7HG0648610.1 | -0.29774 | 6.015867 | -3.53799 | 0.000848 | 0.043951 | -0.8275 | Carboxypeptidase | root cap | 12 h |
| HORVU.MOREX.r3.1HG0057610.1 | -0.52206 | 4.023099 | -3.50542 | 0.000936 | 0.045382 | -0.8296 | Peptide methionine sulfoxide reductase MsrA | root cap | 12 h |
| HORVU.MOREX.r3.5HG0482710.1 | 0.623329 | 3.151378 | 3.482151 | 0.001005 | 0.046967 | -0.82962 | Hydroxycinnamoyl-CoA shikimate/quinate hydroxycinnamoyltransferase | root cap | 12 h |
| HORVU.MOREX.r3.2HG0192860.1 | -0.56244 | 6.583932 | -3.5665 | 0.000777 | 0.041944 | -0.85057 | WRKY transcription factor | root cap | 12 h |
| HORVU.MOREX.r3.2HG0124110.1 | -0.73123 | 2.313092 | -3.45546 | 0.001089 | 0.048807 | -0.85464 | GRAM domain protein/ABA-responsive-like protein | root cap | 12 h |
| HORVU.MOREX.r3.3HG0309210.1 | 0.456235 | 5.46384 | 3.505194 | 0.000937 | 0.045382 | -0.86181 | Obg-like ATPase 1 | root cap | 12 h |
| HORVU.MOREX.r3.5HG0510240.1 | 4.866374 | -2.27899 | 4.274457 | 8.01E-05 | 0.015225 | -0.86404 | Dirigent protein | root cap | 12 h |
| HORVU.MOREX.r3.2HG0121380.1 | -0.63645 | 6.42294 | -3.53247 | 0.000862 | 0.043974 | -0.86686 | Cytochrome P450 | root cap | 12 h |
| HORVU.MOREX.r3.1HG0056810.1 | 0.578921 | 5.129894 | 3.480002 | 0.001011 | 0.04706 | -0.86717 | Haloacid dehalogenase-like hydrolase family protein, putative, expressed | root cap | 12 h |
| HORVU.MOREX.r3.5HG0483810.1 | -0.44894 | 4.62245 | -3.479 | 0.001014 | 0.047096 | -0.86849 | Ubiquitin, putative | root cap | 12 h |
| HORVU.MOREX.r3.7HG0740120.1 | -0.86096 | 5.368158 | -3.52063 | 0.000894 | 0.044517 | -0.87049 | Nodulin-like / Major Facilitator Superfamily protein | root cap | 12 h |
| HORVU.MOREX.r3.3HG0306820.1 | -0.59408 | 4.854402 | -3.47278 | 0.001034 | 0.04717 | -0.87212 | ATP-dependent Clp protease ATP-binding subunit ClpX | root cap | 12 h |
| HORVU.MOREX.r3.7HG0648140.1 | -0.58291 | 6.798944 | -3.54237 | 0.000836 | 0.043781 | -0.87367 | Homeobox leucine zipper protein | root cap | 12 h |
| HORVU.MOREX.r3.6HG0622540.1 | -0.58235 | 7.01829 | -3.53697 | 0.00085 | 0.043951 | -0.8768 | Equilibrative nucleoside transporter | root cap | 12 h |
| HORVU.MOREX.r3.5HG0436990.1 | -0.63733 | 6.379696 | -3.50664 | 0.000933 | 0.045382 | -0.90139 | SPX and EXS domain-containing protein 1 | root cap | 12 h |
| HORVU.MOREX.r3.2HG0109400.1 | -1.26891 | 0.929608 | -3.50359 | 0.000941 | 0.045454 | -0.91098 | D-alanine--D-alanine ligase | root cap | 12 h |
| HORVU.MOREX.r3.7HG0728500.1 | -0.92678 | 0.37984 | -3.49259 | 0.000973 | 0.046043 | -0.917 | Protein phosphatase 2C | root cap | 12 h |
| HORVU.MOREX.r3.2HG0100270.2 | -0.59124 | 6.297507 | -3.53236 | 0.000862 | 0.043974 | -0.94183 | Subtilisin-like protease | root cap | 12 h |
| HORVU.MOREX.r3.5HG0517330.1 | -0.50713 | 7.95292 | -3.60464 | 0.000691 | 0.039589 | -0.95431 | Aldehyde oxidase, putative | root cap | 12 h |
| HORVU.MOREX.r3.7HG0720980.1 | -0.74282 | 1.598477 | -3.45992 | 0.001075 | 0.048472 | -0.96245 | Glucan endo-1,3-beta-glucosidase 3 | root cap | 12 h |
| HORVU.MOREX.r3.5HG0439620.1 | -0.48943 | 6.606388 | -3.49247 | 0.000974 | 0.046043 | -0.96398 | Amino acid transporter, putative | root cap | 12 h |
| HORVU.MOREX.r3.2HG0165780.1 | -0.43394 | 6.457413 | -3.50523 | 0.000937 | 0.045382 | -0.9692 | 1,4-alpha-glucan-branching enzyme | root cap | 12 h |
| HORVU.MOREX.r3.3HG0221140.1 | -0.75898 | 2.320691 | -3.48393 | 0.000999 | 0.046928 | -0.97027 | Glutathione S-transferase, putative, expressed | root cap | 12 h |
| HORVU.MOREX.r3.6HG0631080.1 | -0.49979 | 5.818779 | -3.47025 | 0.001042 | 0.047291 | -0.97246 | Hydroxyproline-rich glycoprotein-like | root cap | 12 h |
| HORVU.MOREX.r3.5HG0467230.2 | -0.44861 | 5.749624 | -3.47578 | 0.001024 | 0.04717 | -0.98802 | Alpha/beta-Hydrolases superfamily protein | root cap | 12 h |
| HORVU.MOREX.r3.2HG0188070.1 | -0.8323 | 0.589714 | -3.50039 | 0.000951 | 0.045685 | -0.98881 | L-ascorbate oxidase-like protein | root cap | 12 h |
| HORVU.MOREX.r3.1HG0068060.1 | -0.35504 | 8.078852 | -3.53276 | 0.000861 | 0.043974 | -0.99456 | DCD (Development and cell death) domain protein | root cap | 12 h |
| HORVU.MOREX.r3.1HG0055320.1 | -0.32104 | 5.996621 | -3.45912 | 0.001077 | 0.048482 | -0.99913 | Protein phosphatase 2C | root cap | 12 h |
| HORVU.MOREX.r3.5HG0456950.1 | -0.42893 | 5.604947 | -3.45707 | 0.001084 | 0.048677 | -1.01074 | Proteasome subunit alpha type | root cap | 12 h |
| HORVU.MOREX.r3.3HG0273560.1 | -1.30076 | 0.616289 | -3.47359 | 0.001031 | 0.04717 | -1.0412 | 2-oxoisovalerate dehydrogenase subunit beta | root cap | 12 h |
| HORVU.MOREX.r3.2HG0133930.1 | -2.10388 | -2.62089 | -3.79087 | 0.000386 | 0.031126 | -1.04466 | Cytochrome c6 | root cap | 12 h |
| HORVU.MOREX.r3.7HG0689920.1 | 1.352767 | 2.721231 | 3.496164 | 0.000963 | 0.045846 | -1.05458 | Carboxypeptidase | root cap | 12 h |
| HORVU.MOREX.r3.3HG0232840.1 | -0.46789 | 7.520309 | -3.4733 | 0.001032 | 0.04717 | -1.07205 | Fatty acid oxidation complex subunit alpha | root cap | 12 h |
| HORVU.MOREX.r3.6HG0615840.1 | -1.86416 | -2.2741 | -3.66886 | 0.000566 | 0.036145 | -1.10884 | Metalloendoproteinase 1 | root cap | 12 h |
| HORVU.MOREX.r3.4HG0408300.1 | -0.27223 | 7.884162 | -3.47445 | 0.001029 | 0.04717 | -1.12185 | HR-like lesion-inducing protein-related protein | root cap | 12 h |
| HORVU.MOREX.r3.4HG0406720.1 | -1.74746 | -1.7339 | -3.46505 | 0.001058 | 0.04783 | -1.12225 | nuclear polyadenylated RNA-binding protein | root cap | 12 h |
| HORVU.MOREX.r3.5HG0510700.1 | -0.47072 | 7.762858 | -3.50213 | 0.000946 | 0.04555 | -1.12256 | Beta-glucosidase | root cap | 12 h |
| HORVU.MOREX.r3.4HG0340420.1 | -0.48239 | 5.772276 | -3.47194 | 0.001036 | 0.04717 | -1.13182 | GEM-like protein 1 | root cap | 12 h |
| HORVU.MOREX.r3.5HG0500130.1 | -2.25541 | -2.11769 | -3.48947 | 0.000983 | 0.046251 | -1.13427 | Pathogenesis-related thaumatin-like protein | root cap | 12 h |

**Table S1** Continued.

| ID | logFC | AveExpr | t | P.Value | adj.P.Val | B | gene description | root zone | time point |
|---|---|---|---|---|---|---|---|---|---|
| HORVU.MOREX.r3.3HG0323410.1 | -1.04908 | -2.74988 | -3.4826 | 0.001003 | 0.046967 | -1.13714 | TSA: Wollemia nobilis Ref_Wollemi_Transcript_14908_3298 transcribed RNA sequ | root cap | 12 h |
| HORVU.MOREX.r3.3HG0315300.1 | -0.86865 | 2.211349 | -3.52527 | 0.000881 | 0.044002 | -1.22958 | Late embryogenesis abundant protein | root cap | 12 h |
| HORVU.MOREX.r3.5HG0429390.1 | 3.178481 | -0.95655 | 3.585647 | 0.000732 | 0.040947 | -1.23874 | Germin-like protein | root cap | 12 h |
| HORVU.MOREX.r3.1HG0017350.1 | -0.74546 | 4.123138 | -3.4512 | 0.001103 | 0.049332 | -1.27071 | Peroxidase | root cap | 12 h |
| HORVU.MOREX.r3.5HG0495750.1 | -1.91447 | -2.86879 | -3.63564 | 0.000628 | 0.037866 | -1.30772 | B3 domain-containing protein | root cap | 12 h |
| HORVU.MOREX.r3.2HG0207160.1 | -0.72434 | 2.792582 | -3.47183 | 0.001037 | 0.04717 | -1.32139 | Micronuclear linker histone polyprotein-like protein | root cap | 12 h |
| HORVU.MOREX.r3.1HG0024440.1 | 1.761637 | 2.671204 | 3.566872 | 0.000776 | 0.041944 | -1.36494 | EGG APPARATUS-1 protein | root cap | 12 h |
| HORVU.MOREX.r3.2HG0120640.1 | -1.53762 | 0.112362 | -3.57793 | 0.00075 | 0.041215 | -1.39242 | Protein root UVB sensitive 3 | root cap | 12 h |
| HORVU.MOREX.r3.1HG0072560.1 | 1.884607 | -2.88695 | 3.490882 | 0.000979 | 0.046159 | -1.40075 | Chaperone protein dnaJ | root cap | 12 h |
| HORVU.MOREX.r3.6HG0604200.1 | 2.948316 | 0.762306 | 3.965535 | 0.000221 | 0.0236 | -1.45947 | transcription repressor | root cap | 12 h |
| HORVU.MOREX.r3.2HG0104400.1 | 2.849909 | -1.01887 | 3.577137 | 0.000752 | 0.041215 | -1.48818 | Dirigent protein | root cap | 12 h |
| HORVU.MOREX.r3.6HG0627900.1 | 2.447031 | -3.19226 | 3.650641 | 0.000599 | 0.037246 | -1.49886 | Receptor-like protein kinase | root cap | 12 h |
| HORVU.MOREX.r3.7HG0719310.1 | -2.315 | -2.19972 | -3.4486 | 0.001112 | 0.049612 | -1.59467 | Cytoplasmic tRNA 2-thiolation protein 1 | root cap | 12 h |
| HORVU.MOREX.r3.4HG0399460.1 | -2.11094 | -1.4594 | -3.46788 | 0.001049 | 0.047527 | -1.6223 | transcription repressor | root cap | 12 h |
| HORVU.MOREX.r3.2HG0116730.1 | 3.253115 | 3.133763 | 3.54846 | 0.000821 | 0.043324 | -1.66235 | ABC transporter, putative | root cap | 12 h |
| HORVU.MOREX.r3.4HG0402390.1 | 5.226258 | -1.02408 | 3.576267 | 0.000754 | 0.041215 | -1.80179 | 3-ketoacyl-CoA synthase | root cap | 12 h |
| HORVU.MOREX.r3.3HG0328340.1 | -1.45036 | 4.602643 | -6.7403 | 1.18E-08 | 0.000242 | 9.530264 | Two-component response regulator | meristem | 12 h |
| HORVU.MOREX.r3.5HG0436920.1 | -1.63437 | 1.708407 | -5.87204 | 2.89E-07 | 0.002965 | 5.521741 | RING/U-box superfamily protein | meristem | 12 h |
| HORVU.MOREX.r3.6HG0605060.1 | -0.8199 | 4.908798 | -5.08838 | 4.86E-06 | 0.033236 | 4.011963 | SBP (S-ribonuclease-binding protein) family protein | meristem | 12 h |
| HORVU.MOREX.r3.1HG0089130.1 | -0.46022 | 5.608758 | -4.92789 | 8.55E-06 | 0.039818 | 3.468557 | Serine/Threonine-kinase pakA-like protein | meristem | 12 h |
| HORVU.MOREX.r3.1HG0073350.1 | -1.26259 | 2.878989 | -4.89183 | 9.70E-06 | 0.039818 | 3.351209 | 70 kDa heat shock protein | meristem | 12 h |
| HORVU.MOREX.r3.5HG0517740.1 | 2.426466 | -0.30721 | 6.112302 | 1.20E-07 | 0.002459 | 7.492382 | Peroxidase | elongation zone | 3 h |
| HORVU.MOREX.r3.4HG0395540.1 | 1.75487 | 4.838553 | 5.168475 | 3.66E-06 | 0.025019 | 4.096146 | Glutamine synthetase | elongation zone | 3 h |
| HORVU.MOREX.r3.6HG0623140.1 | 2.893193 | -2.34532 | 5.333784 | 2.03E-06 | 0.020807 | 3.758867 | #N/A | elongation zone | 3 h |
| HORVU.MOREX.r3.5HG0527570.1 | 1.885468 | 4.789839 | 4.919898 | 8.79E-06 | 0.036487 | 3.460428 | UDP-glycosyltransferase | elongation zone | 3 h |
| HORVU.MOREX.r3.5HG0532150.1 | 2.523346 | -2.26204 | 4.864651 | 1.07E-05 | 0.036487 | 3.135388 | Expansin | elongation zone | 3 h |
| HORVU.MOREX.r3.4HG0417920.1 | 2.669337 | -1.69902 | 4.745546 | 1.61E-05 | 0.047323 | 2.848495 | Peroxidase | elongation zone | 3 h |
| HORVU.MOREX.r3.5HG0420460.1 | 3.093241 | -1.03028 | 4.871991 | 1.04E-05 | 0.036487 | 1.921396 | NBS-LRR-like resistance protein | elongation zone | 3 h |
| HORVU.MOREX.r3.5HG0517740.1 | 2.679485 | -0.30721 | 6.828114 | 8.52E-09 | 0.000175 | 10.01505 | Peroxidase | elongation zone | 6 h |
| HORVU.MOREX.r3.7HG0750000.1 | 2.377402 | -2.12041 | 6.515961 | 2.71E-08 | 0.000207 | 8.667624 | Disease resistance protein | elongation zone | 6 h |
| HORVU.MOREX.r3.7HG0663750.1 | 2.397222 | 3.681851 | 6.329002 | 5.40E-08 | 0.000207 | 8.23283 | Galactoside 2-alpha-L-fucosyltransferase | elongation zone | 6 h |
| HORVU.MOREX.r3.1HG0010340.1 | 2.453683 | -2.09413 | 6.363681 | 4.75E-08 | 0.000207 | 8.143603 | Non-specific lipid-transfer protein-like protein | elongation zone | 6 h |
| HORVU.MOREX.r3.4HG0417920.1 | 3.451323 | -1.69902 | 6.328486 | 5.41E-08 | 0.000207 | 8.079794 | Peroxidase | elongation zone | 6 h |
| HORVU.MOREX.r3.7HG0671470.1 | -0.9246 | 4.81657 | -6.28406 | 6.37E-08 | 0.000207 | 8.025028 | Protein trichome birefringence | elongation zone | 6 h |
| HORVU.MOREX.r3.4HG0395540.1 | 2.082668 | 4.838553 | 6.240994 | 7.46E-08 | 0.000207 | 7.797683 | Glutamine synthetase | elongation zone | 6 h |
| HORVU.MOREX.r3.4HG0408270.1 | 2.186194 | -2.27119 | 6.22002 | 8.06E-08 | 0.000207 | 7.64336 | LOB domain-containing protein, putative | elongation zone | 6 h |
| HORVU.MOREX.r3.3HG0318590.1 | 2.82361 | -1.25566 | 6.027641 | 1.63E-07 | 0.000373 | 7.197975 | Endo-1,3-beta-glucanase | elongation zone | 6 h |
| HORVU.MOREX.r3.5HG0518560.1 | 2.741276 | 2.148281 | 5.973574 | 1.99E-07 | 0.000409 | 6.987664 | Leucine-rich repeat protein kinase family protein | elongation zone | 6 h |
| HORVU.MOREX.r3.4HG0415330.1 | 2.298155 | 3.57761 | 5.938845 | 2.26E-07 | 0.000422 | 6.838559 | Cytochrome P450 family cinnamate 4-hydroxylase | elongation zone | 6 h |
| HORVU.MOREX.r3.2HG0217090.1 | 2.254303 | -0.85749 | 5.869515 | 2.92E-07 | 0.000499 | 6.646459 | C2 calcium/lipid-binding and GRAM domain protein | elongation zone | 6 h |
| HORVU.MOREX.r3.5HG0527570.1 | 2.17181 | 4.789839 | 5.755027 | 4.43E-07 | 0.000649 | 6.256887 | UDP-glycosyltransferase | elongation zone | 6 h |

**Table S1** Continued.

| ID | logFC | AveExpr | t | P.Value | adj.P.Val | B | gene description | root zone | time point |
|---|---|---|---|---|---|---|---|---|---|
| HORVU.MOREX.r3.5HG0429900.1 | 1.569007 | 4.093407 | 5.755871 | 4.41E-07 | 0.000649 | 6.254698 | #N/A | elongation zone | 6 h |
| HORVU.MOREX.r3.1HG0054190.1 | 2.243132 | -0.12557 | 5.603621 | 7.67E-07 | 0.000984 | 5.682088 | Expansin-like protein | elongation zone | 6 h |
| HORVU.MOREX.r3.1HG0080180.1 | 2.363438 | -2.35246 | 5.610571 | 7.48E-07 | 0.000984 | 5.545157 | Jacalin lectin family protein | elongation zone | 6 h |
| HORVU.MOREX.r3.7HG0747230.1 | 0.983846 | 2.606945 | 5.493661 | 1.14E-06 | 0.001128 | 5.364691 | Non-specific serine/threonine protein kinase | elongation zone | 6 h |
| HORVU.MOREX.r3.6HG0550600.1 | 1.357649 | 1.720716 | 5.536978 | 9.76E-07 | 0.001113 | 5.309226 | Protein ROOT HAIR DEFECTIVE 3 homolog | elongation zone | 6 h |
| HORVU.MOREX.r3.3HG0329950.1 | 2.017307 | 1.75889 | 5.486008 | 1.17E-06 | 0.001128 | 5.243991 | Pectin acetylesterase | elongation zone | 6 h |
| HORVU.MOREX.r3.5HG0532140.1 | 2.101436 | -1.00113 | 5.448458 | 1.34E-06 | 0.001128 | 5.182938 | Expansin | elongation zone | 6 h |
| HORVU.MOREX.r3.4HG0417010.1 | 1.926109 | -1.71396 | 5.445447 | 1.36E-06 | 0.001128 | 5.130972 | Receptor protein kinase, putative | elongation zone | 6 h |
| HORVU.MOREX.r3.5HG0519120.1 | 1.78141 | -0.8751 | 5.426543 | 1.45E-06 | 0.001128 | 5.124126 | CASP-like protein | elongation zone | 6 h |
| HORVU.MOREX.r3.3HG0222830.1 | -0.48403 | 7.515891 | -5.5027 | 1.10E-06 | 0.001128 | 5.034028 | Heavy metal transport/detoxification superfamily protein | elongation zone | 6 h |
| HORVU.MOREX.r3.1HG0054170.1 | 1.926734 | 0.15708 | 5.422633 | 1.47E-06 | 0.001128 | 5.005012 | Expansin-like protein | elongation zone | 6 h |
| HORVU.MOREX.r3.3HG0234470.1 | -0.56277 | 6.523252 | -5.45501 | 1.31E-06 | 0.001128 | 4.997649 | Kinase interacting (KIP1-like) family protein | elongation zone | 6 h |
| HORVU.MOREX.r3.3HG0297330.1 | -0.63488 | 5.598322 | -5.42073 | 1.48E-06 | 0.001128 | 4.95018 | Receptor-like kinase | elongation zone | 6 h |
| HORVU.MOREX.r3.6HG0626030.1 | 2.162985 | -0.5621 | 5.368023 | 1.79E-06 | 0.001254 | 4.853647 | Peroxidase | elongation zone | 6 h |
| HORVU.MOREX.r3.4HG0413740.1 | 2.335849 | -1.83472 | 5.349929 | 1.91E-06 | 0.001267 | 4.834231 | 12-oxophytodienoate reductase-like protein | elongation zone | 6 h |
| HORVU.MOREX.r3.4HG0413320.1 | 2.408495 | 2.316272 | 5.323825 | 2.10E-06 | 0.001348 | 4.757956 | Subtilisin-like protease | elongation zone | 6 h |
| HORVU.MOREX.r3.3HG0330220.1 | 1.170575 | 6.363412 | 5.363581 | 1.82E-06 | 0.001254 | 4.73745 | GDSL esterase/lipase | elongation zone | 6 h |
| HORVU.MOREX.r3.1HG0070340.1 | 1.597112 | 4.307027 | 5.361894 | 1.83E-06 | 0.001254 | 4.735557 | Protein kinase-like protein | elongation zone | 6 h |
| HORVU.MOREX.r3.3HG0250060.1 | 2.548861 | -2.50642 | 5.542587 | 9.57E-07 | 0.001113 | 4.691105 | CASP-like protein | elongation zone | 6 h |
| HORVU.MOREX.r3.2HG0124850.1 | 1.751525 | -0.21834 | 5.306118 | 2.24E-06 | 0.001351 | 4.672637 | Dirigent protein | elongation zone | 6 h |
| HORVU.MOREX.r3.3HG0294960.1 | 1.785464 | -1.29923 | 5.281463 | 2.45E-06 | 0.001352 | 4.64906 | Eukaryotic translation initiation factor 3 subunit H | elongation zone | 6 h |
| HORVU.MOREX.r3.5HG0535180.1 | 1.675045 | -1.06768 | 5.267504 | 2.57E-06 | 0.001352 | 4.590332 | #N/A | elongation zone | 6 h |
| HORVU.MOREX.r3.3HG0329870.1 | 1.862808 | -1.90322 | 5.259131 | 2.65E-06 | 0.001359 | 4.577099 | Pectin acetylesterase | elongation zone | 6 h |
| HORVU.MOREX.r3.4HG0351750.1 | 2.336208 | -1.8238 | 5.248968 | 2.75E-06 | 0.001363 | 4.541046 | BTB/POZ domain-containing protein | elongation zone | 6 h |
| HORVU.MOREX.r3.2HG0199160.1 | 0.597614 | 6.474544 | 5.274269 | 2.51E-06 | 0.001352 | 4.465054 | Kinesin-like protein | elongation zone | 6 h |
| HORVU.MOREX.r3.2HG0164020.1 | 1.16904 | 1.589544 | 5.204746 | 3.22E-06 | 0.001499 | 4.396228 | E3 ubiquitin protein ligase drip2 | elongation zone | 6 h |
| HORVU.MOREX.r3.2HG0112690.1 | 1.313289 | 1.247173 | 5.274563 | 2.51E-06 | 0.001352 | 4.344444 | Peroxidase | elongation zone | 6 h |
| HORVU.MOREX.r3.5HG0488950.1 | -1.08117 | 2.845933 | -5.1895 | 3.39E-06 | 0.001514 | 4.342797 | Cellulose synthase, putative | elongation zone | 6 h |
| HORVU.MOREX.r3.7HG0642520.1 | 2.284003 | -0.8717 | 5.204781 | 3.21E-06 | 0.001499 | 4.328546 | Peroxidase | elongation zone | 6 h |
| HORVU.MOREX.r3.2HG0195690.1 | 1.669205 | -2.81043 | 5.312093 | 2.19E-06 | 0.001351 | 4.304607 | filamentous hemagglutinin transporter | elongation zone | 6 h |
| HORVU.MOREX.r3.1HG0072760.1 | -0.81927 | 4.906091 | -5.19219 | 3.36E-06 | 0.001514 | 4.19793 | Serpin family protein | elongation zone | 6 h |
| HORVU.MOREX.r3.3HG0293310.1 | 1.921913 | -2.57133 | 5.244511 | 2.79E-06 | 0.001363 | 4.191156 | Protein upstream of flc | elongation zone | 6 h |
| HORVU.MOREX.r3.2HG0112060.1 | -0.78009 | 4.798982 | -5.16625 | 3.69E-06 | 0.001583 | 4.186708 | Transcription factor | elongation zone | 6 h |
| HORVU.MOREX.r3.5HG0477750.1 | 2.461908 | 0.658352 | 5.165091 | 3.70E-06 | 0.001583 | 4.181246 | Homeobox protein, putative | elongation zone | 6 h |
| HORVU.MOREX.r3.3HG0235720.1 | -2.2935 | 3.057636 | -5.085 | 4.92E-06 | 0.001904 | 3.97504 | FMN-dependent NADPH-azoreductase | elongation zone | 6 h |
| HORVU.MOREX.r3.1HG0081950.1 | 3.02783 | -2.05546 | 5.13377 | 4.14E-06 | 0.001733 | 3.96739 | Peroxidase | elongation zone | 6 h |
| HORVU.MOREX.r3.2HG0189670.1 | 1.586645 | 0.283223 | 5.104952 | 4.58E-06 | 0.001826 | 3.951537 | Aquaporin-like protein | elongation zone | 6 h |
| HORVU.MOREX.r3.5HG0537150.1 | 1.4313 | -2.24688 | 5.102156 | 4.63E-06 | 0.001826 | 3.933581 | Protein DETOXIFICATION | elongation zone | 6 h |
| HORVU.MOREX.r3.3HG0245250.1 | 4.001195 | -1.64331 | 5.056812 | 5.43E-06 | 0.002019 | 3.858977 | Cysteine protease, putative | elongation zone | 6 h |
| HORVU.MOREX.r3.3HG0285440.1 | 1.714132 | -1.28505 | 5.028484 | 6.00E-06 | 0.002019 | 3.810065 | #N/A | elongation zone | 6 h |
| HORVU.MOREX.r3.2HG0155750.1 | 1.817294 | -0.29251 | 5.017459 | 6.24E-06 | 0.002019 | 3.775816 | Ribonuclease | elongation zone | 6 h |

**Table S1** Continued.

| ID | logFC | AveExpr | t | P.Value | adj.P.Val | B | gene description | root zone | time point |
|---|---|---|---|---|---|---|---|---|---|
| HORVU.MOREX.r3.2HG0213020.1 | 1.669875 | 3.22145 | 5.018987 | 6.21E-06 | 0.002019 | 3.760502 | Protein F12F1.11-, putative | elongation zone | 6 h |
| HORVU.MOREX.r3.2HG0160560.1 | -1.32161 | 2.799703 | -5.01608 | 6.27E-06 | 0.002019 | 3.753458 | Peptidylprolyl isomerase | elongation zone | 6 h |
| HORVU.MOREX.r3.1HG0094110.1 | 2.621304 | 0.561027 | 4.997221 | 6.70E-06 | 0.002023 | 3.714629 | Sigma factor sigB regulation protein rsbQ | elongation zone | 6 h |
| HORVU.MOREX.r3.6HG0633900.1 | 2.250533 | -0.51261 | 5.013382 | 6.33E-06 | 0.002019 | 3.671062 | Calcium-dependent protein kinase | elongation zone | 6 h |
| HORVU.MOREX.r3.2HG0192100.1 | 1.898231 | 1.501774 | 4.999422 | 6.65E-06 | 0.002023 | 3.660514 | Serine/threonine-protein kinase | elongation zone | 6 h |
| HORVU.MOREX.r3.1HG0006230.1 | -0.62723 | 5.574 | -5.03355 | 5.90E-06 | 0.002019 | 3.654349 | Disease resistance protein RPM1 | elongation zone | 6 h |
| HORVU.MOREX.r3.2HG0112670.1 | 1.793326 | 0.989354 | 5.059305 | 5.39E-06 | 0.002019 | 3.636665 | Peroxidase | elongation zone | 6 h |
| HORVU.MOREX.r3.1HG0066610.1 | 1.140951 | 2.119827 | 5.021741 | 6.15E-06 | 0.002019 | 3.614935 | Dirigent protein | elongation zone | 6 h |
| HORVU.MOREX.r3.2HG0156610.1 | 1.966857 | -0.12875 | 4.978706 | 7.15E-06 | 0.002097 | 3.604955 | ERD (Early-responsive to dehydration stress) family protein | elongation zone | 6 h |
| HORVU.MOREX.r3.3HG0325250.1 | 3.827258 | -1.52184 | 5.115665 | 4.41E-06 | 0.001811 | 3.585871 | Peroxidase | elongation zone | 6 h |
| HORVU.MOREX.r3.5HG0527460.1 | 1.413704 | 1.065374 | 5.026848 | 6.04E-06 | 0.002019 | 3.571092 | Pectin acetylesterase | elongation zone | 6 h |
| HORVU.MOREX.r3.4HG0394330.1 | 1.298373 | 1.645952 | 4.982144 | 7.07E-06 | 0.002097 | 3.571046 | Kinase family protein | elongation zone | 6 h |
| HORVU.MOREX.r3.3HG0312430.1 | 2.027052 | -0.09459 | 4.940226 | 8.19E-06 | 0.002367 | 3.526814 | E3 ubiquitin protein ligase drip2 | elongation zone | 6 h |
| HORVU.MOREX.r3.4HG0382620.1 | 0.645597 | 7.496381 | 5.005042 | 6.52E-06 | 0.002023 | 3.445143 | Argininosuccinate lyase | elongation zone | 6 h |
| HORVU.MOREX.r3.6HG0601830.1 | 1.688595 | -2.05956 | 4.908467 | 9.15E-06 | 0.00245 | 3.421559 | C2 calcium/lipid-binding and GRAM domain protein | elongation zone | 6 h |
| HORVU.MOREX.r3.4HG0350900.1 | -0.44244 | 7.975994 | -5.04518 | 5.66E-06 | 0.002019 | 3.409826 | UDP-glucose 6-dehydrogenase | elongation zone | 6 h |
| HORVU.MOREX.r3.1HG0005700.1 | 2.282402 | -2.12158 | 4.90216 | 9.36E-06 | 0.00245 | 3.379499 | #N/A | elongation zone | 6 h |
| HORVU.MOREX.r3.5HG0470680.1 | 1.809572 | -1.96407 | 4.892452 | 9.68E-06 | 0.00245 | 3.366296 | Heavy metal transport/detoxification protein | elongation zone | 6 h |
| HORVU.MOREX.r3.1HG0047100.1 | 1.16872 | 4.15495 | 4.922399 | 8.72E-06 | 0.00245 | 3.363243 | LanC-like protein | elongation zone | 6 h |
| HORVU.MOREX.r3.3HG0317780.1 | 1.348477 | 0.361248 | 4.88641 | 9.89E-06 | 0.00245 | 3.299334 | 2-oxoglutarate (2OG) and Fe(II)-dependent oxygenase superfamily protein | elongation zone | 6 h |
| HORVU.MOREX.r3.3HG0329040.1 | 2.420593 | -0.45471 | 4.866873 | 1.06E-05 | 0.002454 | 3.290501 | Vacuolar-processing enzyme | elongation zone | 6 h |
| HORVU.MOREX.r3.2HG0179560.1 | 1.265023 | 1.523229 | 4.892207 | 9.69E-06 | 0.00245 | 3.288011 | Nodulin-like / Major Facilitator Superfamily protein | elongation zone | 6 h |
| HORVU.MOREX.r3.4HG0379660.1 | 1.488994 | -1.00939 | 4.869022 | 1.05E-05 | 0.002454 | 3.26633 | Transcription repressor OFP17 | elongation zone | 6 h |
| HORVU.MOREX.r3.3HG0237990.1 | 0.84265 | 4.191674 | 4.903838 | 9.30E-06 | 0.00245 | 3.257984 | #N/A | elongation zone | 6 h |
| HORVU.MOREX.r3.3HG0247440.1 | -1.03879 | 2.109103 | -4.8754 | 1.03E-05 | 0.002454 | 3.252388 | Serine/threonine-protein kinase | elongation zone | 6 h |
| HORVU.MOREX.r3.1HG0073390.1 | 1.680776 | 2.372781 | 4.856169 | 1.10E-05 | 0.002454 | 3.251725 | Respiratory burst oxidase-like protein | elongation zone | 6 h |
| HORVU.MOREX.r3.5HG0472770.1 | 0.935693 | 4.331329 | 4.885725 | 9.91E-06 | 0.00245 | 3.251696 | SIN3-like 2 | elongation zone | 6 h |
| HORVU.MOREX.r3.5HG0485940.1 | 2.28393 | 2.592249 | 4.852771 | 1.11E-05 | 0.002454 | 3.235626 | Hexosyltransferase | elongation zone | 6 h |
| HORVU.MOREX.r3.7HG0747720.1 | 1.022779 | 3.70009 | 4.859879 | 1.08E-05 | 0.002454 | 3.23184 | Cytokinin oxidase/dehydrogenase | elongation zone | 6 h |
| HORVU.MOREX.r3.7HG0667060.1 | 1.645515 | 2.126304 | 4.85024 | 1.12E-05 | 0.002454 | 3.218683 | target of early activation tagged (EAT) 2 | elongation zone | 6 h |
| HORVU.MOREX.r3.5HG0467870.1 | 1.235032 | 2.049337 | 4.846053 | 1.14E-05 | 0.002459 | 3.186763 | Kynurenine formamidase | elongation zone | 6 h |
| HORVU.MOREX.r3.4HG0400920.1 | 1.24527 | 4.093901 | 4.868231 | 1.05E-05 | 0.002454 | 3.144572 | Kinesin-like protein | elongation zone | 6 h |
| HORVU.MOREX.r3.5HG0509120.1 | 1.329413 | 5.607225 | 4.909415 | 9.12E-06 | 0.00245 | 3.139496 | Auxin-responsive protein | elongation zone | 6 h |
| HORVU.MOREX.r3.3HG0300950.1 | -0.85585 | 3.921584 | -4.85595 | 1.10E-05 | 0.002454 | 3.132576 | Amino acid transporter, putative | elongation zone | 6 h |
| HORVU.MOREX.r3.3HG0235350.1 | 2.5297 | 0.004514 | 4.837803 | 1.17E-05 | 0.002504 | 3.131832 | Pollen Ole e 1 allergen and extensin family protein | elongation zone | 6 h |
| HORVU.MOREX.r3.4HG0346830.1 | 1.949416 | -2.75621 | 4.903876 | 9.30E-06 | 0.00245 | 3.098291 | WEB family protein | elongation zone | 6 h |
| HORVU.MOREX.r3.4HG0409230.1 | 2.214958 | -0.03281 | 4.855714 | 1.10E-05 | 0.002454 | 3.09114 | IQ domain-containing protein | elongation zone | 6 h |
| HORVU.MOREX.r3.5HG0517760.1 | 2.078191 | -2.35783 | 4.832731 | 1.19E-05 | 0.002523 | 3.064581 | Zinc finger A20 and AN1 domain-containing stress-associated protein | elongation zone | 6 h |
| HORVU.MOREX.r3.5HG0470700.1 | 1.711283 | -1.11013 | 4.803603 | 1.32E-05 | 0.002593 | 3.054163 | Heavy metal transport/detoxification protein | elongation zone | 6 h |
| HORVU.MOREX.r3.4HG0405400.1 | 1.88462 | 2.976906 | 4.822336 | 1.24E-05 | 0.002589 | 3.048753 | DUF936 family protein | elongation zone | 6 h |
| HORVU.MOREX.r3.7HG0699010.1 | 1.37814 | 4.331849 | 4.802022 | 1.33E-05 | 0.002593 | 3.021367 | LHY | elongation zone | 6 h |

**Table S1** Continued.

| ID | logFC | AveExpr | t | P.Value | adj.P.Val | B | gene description | root zone | time point |
|---|---|---|---|---|---|---|---|---|---|
| HORVU.MOREX.r3.4HG0415490.1 | 2.279941 | -1.78464 | 4.787441 | 1.40E-05 | 0.002644 | 3.01628 | 14 kDa proline-rich protein DC2.15 | elongation zone | 6 h |
| HORVU.MOREX.r3.3HG0298080.1 | 1.206277 | 1.480621 | 4.806876 | 1.30E-05 | 0.002593 | 3.015565 | Amino acid permease | elongation zone | 6 h |
| HORVU.MOREX.r3.4HG0337110.1 | -0.40188 | 7.514078 | -4.90603 | 9.23E-06 | 0.00245 | 3.014211 | Coatomer subunit alpha | elongation zone | 6 h |
| HORVU.MOREX.r3.6HG0622250.1 | 1.439175 | 2.510596 | 4.792629 | 1.37E-05 | 0.002644 | 3.003956 | Alpha/beta-hydrolase superfamily protein | elongation zone | 6 h |
| HORVU.MOREX.r3.5HG0524350.1 | 2.291352 | -1.05304 | 4.781057 | 1.43E-05 | 0.002663 | 2.999931 | Protein MIZU-KUSSEI 1 | elongation zone | 6 h |
| HORVU.MOREX.r3.6HG0600000.1 | 1.619501 | -0.21905 | 4.80384 | 1.32E-05 | 0.002593 | 2.989788 | Blue copper protein | elongation zone | 6 h |
| HORVU.MOREX.r3.7HG0748270.1 | 1.976082 | 0.645561 | 4.769564 | 1.49E-05 | 0.002721 | 2.972423 | Blue copper protein | elongation zone | 6 h |
| HORVU.MOREX.r3.1HG0073550.1 | 2.083101 | -1.6606 | 4.758755 | 1.54E-05 | 0.002728 | 2.939341 | Laccase | elongation zone | 6 h |
| HORVU.MOREX.r3.5HG0429180.1 | 1.588345 | 2.149739 | 4.762823 | 1.52E-05 | 0.002728 | 2.890774 | Glycosyltransferase | elongation zone | 6 h |
| HORVU.MOREX.r3.3HG0254950.1 | 1.458454 | 1.01976 | 4.811486 | 1.28E-05 | 0.002593 | 2.888944 | Peroxidase | elongation zone | 6 h |
| HORVU.MOREX.r3.3HG0321700.1 | 0.826924 | 3.776226 | 4.755356 | 1.56E-05 | 0.002728 | 2.87577 | FMN-dependent NADPH-azoreductase | elongation zone | 6 h |
| HORVU.MOREX.r3.7HG0674720.1 | 2.213087 | -1.46328 | 4.742885 | 1.63E-05 | 0.002754 | 2.866067 | Epoxide hydrolase 2 | elongation zone | 6 h |
| HORVU.MOREX.r3.6HG0629520.1 | 1.814902 | -0.9319 | 4.750709 | 1.59E-05 | 0.002734 | 2.863045 | Ethylene-responsive transcription factor | elongation zone | 6 h |
| HORVU.MOREX.r3.7HG0663920.1 | 2.620663 | -0.12227 | 4.759457 | 1.54E-05 | 0.002728 | 2.850559 | Alpha/beta-Hydrolases superfamily protein, putative | elongation zone | 6 h |
| HORVU.MOREX.r3.7HG0739230.1 | 4.725231 | -3.45098 | 5.275986 | 2.49E-06 | 0.001352 | 2.827608 | Jacalin-like lectin | elongation zone | 6 h |
| HORVU.MOREX.r3.2HG0180010.1 | 1.611356 | 1.125048 | 4.785735 | 1.40E-05 | 0.002644 | 2.826428 | #N/A | elongation zone | 6 h |
| HORVU.MOREX.r3.5HG0532150.1 | 2.45965 | -2.26204 | 4.735246 | 1.67E-05 | 0.002754 | 2.819527 | Expansin | elongation zone | 6 h |
| HORVU.MOREX.r3.1HG0066800.1 | 1.452973 | -2.50287 | 4.733908 | 1.68E-05 | 0.002754 | 2.806418 | Potassium channel | elongation zone | 6 h |
| HORVU.MOREX.r3.1HG0086370.1 | 2.439978 | -3.20911 | 4.899111 | 9.46E-06 | 0.00245 | 2.79446 | Glyoxal oxidase | elongation zone | 6 h |
| HORVU.MOREX.r3.4HG0348800.1 | 1.299582 | 2.044228 | 4.709702 | 1.83E-05 | 0.002842 | 2.784431 | TCP transcription factor | elongation zone | 6 h |
| HORVU.MOREX.r3.5HG0537360.1 | 2.099833 | -1.21127 | 4.707056 | 1.84E-05 | 0.002842 | 2.771019 | transcription factor, putative (Protein of unknown function, DUF547) | elongation zone | 6 h |
| HORVU.MOREX.r3.1HG0054180.1 | 1.743723 | -0.6979 | 4.72537 | 1.73E-05 | 0.002797 | 2.768605 | Expansin-like protein | elongation zone | 6 h |
| HORVU.MOREX.r3.6HG0541940.1 | -0.45334 | 6.764259 | -4.80795 | 1.30E-05 | 0.002593 | 2.751604 | S-adenosyl-L-methionine-dependent methyltransferases superfamily protein | elongation zone | 6 h |
| HORVU.MOREX.r3.1HG0043520.1 | -0.51948 | 4.227432 | -4.7432 | 1.63E-05 | 0.002754 | 2.738959 | F-box protein | elongation zone | 6 h |
| HORVU.MOREX.r3.7HG0730510.1 | 1.366303 | -1.83116 | 4.694704 | 1.92E-05 | 0.002883 | 2.737722 | Germin-like protein 1 | elongation zone | 6 h |
| HORVU.MOREX.r3.7HG0678160.1 | -0.70262 | 6.566933 | -4.76044 | 1.53E-05 | 0.002728 | 2.703576 | Disease resistance protein | elongation zone | 6 h |
| HORVU.MOREX.r3.5HG0488050.1 | 1.895839 | -2.9262 | 4.789925 | 1.38E-05 | 0.002644 | 2.688639 | Blue copper protein | elongation zone | 6 h |
| HORVU.MOREX.r3.3HG0246540.1 | -0.83451 | 4.407087 | -4.70697 | 1.84E-05 | 0.002842 | 2.672307 | Peptide-N4-(N-acetyl-beta-glucosaminyl)asparagine amidase A | elongation zone | 6 h |
| HORVU.MOREX.r3.4HG0339890.1 | 1.693851 | -1.1907 | 4.681314 | 2.02E-05 | 0.002975 | 2.671718 | Pollen Ole e 1 allergen and extensin family protein, putative | elongation zone | 6 h |
| HORVU.MOREX.r3.3HG0330200.1 | 1.317891 | 1.169812 | 4.753844 | 1.57E-05 | 0.002728 | 2.637862 | Aquaporin | elongation zone | 6 h |
| HORVU.MOREX.r3.4HG0405920.1 | 1.58089 | 0.085137 | 4.71022 | 1.82E-05 | 0.002842 | 2.637249 | Hexosyltransferase | elongation zone | 6 h |
| HORVU.MOREX.r3.2HG0113990.1 | 2.278936 | -2.49708 | 4.735486 | 1.67E-05 | 0.002754 | 2.631447 | Calcium-dependent lipid-binding (CaLB domain) family protein | elongation zone | 6 h |
| HORVU.MOREX.r3.4HG0337950.1 | 2.55286 | -0.29631 | 4.70456 | 1.86E-05 | 0.002842 | 2.604523 | 65-kDa microtubule-associated-like protein | elongation zone | 6 h |
| HORVU.MOREX.r3.3HG0275090.1 | 2.292549 | -3.21251 | 4.849632 | 1.12E-05 | 0.002454 | 2.585923 | SWI/SNF-related matrix-associated actin-dependent regulator of chromatin subfa | elongation zone | 6 h |
| HORVU.MOREX.r3.3HG0240310.1 | -1.12244 | 6.614043 | -4.73715 | 1.66E-05 | 0.002754 | 2.575514 | DUF2921 family protein | elongation zone | 6 h |
| HORVU.MOREX.r3.6HG0613360.1 | 1.971445 | -0.60847 | 4.698584 | 1.90E-05 | 0.002865 | 2.565885 | Nucleotidylyl transferase superfamily protein | elongation zone | 6 h |
| HORVU.MOREX.r3.3HG0246670.1 | 1.674864 | -1.91783 | 4.650851 | 2.24E-05 | 0.003069 | 2.542458 | DUF4228 domain protein | elongation zone | 6 h |
| HORVU.MOREX.r3.5HG0527310.1 | 2.503643 | -3.15598 | 4.817665 | 1.26E-05 | 0.002593 | 2.536395 | Protein NRT1/ PTR FAMILY 5.5 | elongation zone | 6 h |
| HORVU.MOREX.r3.4HG0384620.1 | 1.046192 | 3.226323 | 4.654462 | 2.21E-05 | 0.003069 | 2.50729 | Kinase family protein | elongation zone | 6 h |
| HORVU.MOREX.r3.2HG0196800.1 | 2.308663 | -0.20994 | 4.682911 | 2.00E-05 | 0.002975 | 2.496555 | Multidrug resistance protein ABC transporter family | elongation zone | 6 h |
| HORVU.MOREX.r3.6HG0615840.1 | 3.029717 | -2.2741 | 5.010543 | 6.40E-06 | 0.002019 | 2.495812 | Metalloendoproteinase 1 | elongation zone | 6 h |

**Table S1** Continued.

| ID | logFC | AveExpr | t | P.Value | adj.P.Val | B | gene description | root zone | time point |
|---|---|---|---|---|---|---|---|---|---|
| HORVU.MOREX.r3.2HG0097390.1 | -0.67881 | 4.644505 | -4.66435 | 2.14E-05 | 0.003066 | 2.461472 | Inosine-5'-monophosphate dehydrogenase | elongation zone | 6 h |
| HORVU.MOREX.r3.4HG0344370.1 | 1.666446 | -2.72257 | 4.652708 | 2.22E-05 | 0.003069 | 2.449635 | Glucuronoxylan 4-O-methyltransferase | elongation zone | 6 h |
| HORVU.MOREX.r3.2HG0203070.1 | 1.028615 | 3.219858 | 4.654046 | 2.21E-05 | 0.003069 | 2.439331 | Receptor-like kinase | elongation zone | 6 h |
| HORVU.MOREX.r3.2HG0214070.1 | 2.22004 | -2.62397 | 4.674435 | 2.06E-05 | 0.003023 | 2.41469 | AT hook motif DNA-binding family protein, putative | elongation zone | 6 h |
| HORVU.MOREX.r3.6HG0626020.1 | 1.599896 | 1.680187 | 4.703098 | 1.87E-05 | 0.002842 | 2.388228 | Peroxidase | elongation zone | 6 h |
| HORVU.MOREX.r3.1HG0087170.1 | -0.54009 | 5.424188 | -4.6476 | 2.26E-05 | 0.003076 | 2.368309 | 3-deoxy-manno-octulosonate cytidylyltransferase | elongation zone | 6 h |
| HORVU.MOREX.r3.2HG0120830.1 | 2.427585 | -3.15708 | 4.771755 | 1.47E-05 | 0.002721 | 2.358817 | Peroxidase | elongation zone | 6 h |
| HORVU.MOREX.r3.5HG0479500.1 | 1.485594 | 1.850868 | 4.617357 | 2.51E-05 | 0.003262 | 2.354289 | Phosphonates import ATP-binding protein PhnC | elongation zone | 6 h |
| HORVU.MOREX.r3.1HG0055240.1 | 2.238539 | -3.1472 | 4.732093 | 1.69E-05 | 0.002754 | 2.352393 | Acetyl-coenzyme A carboxylase carboxyl transferase subunit beta | elongation zone | 6 h |
| HORVU.MOREX.r3.3HG0234000.1 | 1.95946 | -0.27443 | 4.644075 | 2.29E-05 | 0.003083 | 2.324145 | Xyloglucan endotransglucosylase/hydrolase | elongation zone | 6 h |
| HORVU.MOREX.r3.5HG0496980.1 | -0.62711 | 5.215201 | -4.65024 | 2.24E-05 | 0.003069 | 2.311565 | Protein IQ-DOMAIN 1 | elongation zone | 6 h |
| HORVU.MOREX.r3.5HG0480980.1 | 2.820838 | -1.63821 | 4.564578 | 3.01E-05 | 0.003715 | 2.310783 | Spermidine/putrescine import ATP-binding protein PotA | elongation zone | 6 h |
| HORVU.MOREX.r3.4HG0390770.1 | 1.127831 | 3.149264 | 4.561432 | 3.04E-05 | 0.003715 | 2.289506 | Regulator of Vps4 activity in the MVB pathway protein, putative | elongation zone | 6 h |
| HORVU.MOREX.r3.1HG0056620.1 | 1.936995 | 0.189162 | 4.652306 | 2.23E-05 | 0.003069 | 2.287993 | Expansin protein | elongation zone | 6 h |
| HORVU.MOREX.r3.1HG0068010.1 | 1.638798 | -0.26596 | 4.629764 | 2.41E-05 | 0.003207 | 2.270062 | Peroxidase | elongation zone | 6 h |
| HORVU.MOREX.r3.5HG0458890.1 | 0.613092 | 3.514885 | 4.587915 | 2.78E-05 | 0.003531 | 2.268363 | senescence regulator (Protein of unknown function, DUF584) | elongation zone | 6 h |
| HORVU.MOREX.r3.4HG0403620.1 | 1.680876 | 0.126435 | 4.643159 | 2.30E-05 | 0.003083 | 2.249992 | #N/A | elongation zone | 6 h |
| HORVU.MOREX.r3.4HG0409880.1 | 2.085554 | -2.34755 | 4.623206 | 2.46E-05 | 0.003238 | 2.231724 | Metal tolerance protein | elongation zone | 6 h |
| HORVU.MOREX.r3.1HG0027200.1 | 1.16177 | 4.502879 | 4.567182 | 2.98E-05 | 0.003709 | 2.210766 | Auxin-responsive protein | elongation zone | 6 h |
| HORVU.MOREX.r3.7HG0728820.1 | 1.711457 | 1.672182 | 4.561878 | 3.04E-05 | 0.003715 | 2.203344 | Xyloglucan endotransglucosylase/hydrolase | elongation zone | 6 h |
| HORVU.MOREX.r3.7HG0739080.1 | 2.592859 | -3.17965 | 4.716869 | 1.78E-05 | 0.002836 | 2.142108 | Subtilisin-like protease | elongation zone | 6 h |
| HORVU.MOREX.r3.1HG0075730.1 | 1.142807 | 1.539739 | 4.62608 | 2.44E-05 | 0.003227 | 2.105503 | Subtilisin-like protease | elongation zone | 6 h |
| HORVU.MOREX.r3.5HG0482670.1 | 1.369075 | 0.628979 | 4.584839 | 2.81E-05 | 0.003534 | 2.104686 | Cellulose synthase | elongation zone | 6 h |
| HORVU.MOREX.r3.3HG0307240.1 | 3.047396 | -3.37688 | 4.719442 | 1.77E-05 | 0.002833 | 2.100061 | Amino acid permease | elongation zone | 6 h |
| HORVU.MOREX.r3.3HG0303410.1 | 0.766637 | 6.71103 | 4.658673 | 2.18E-05 | 0.003069 | 2.087438 | Respiratory burst oxidase-like protein | elongation zone | 6 h |
| HORVU.MOREX.r3.2HG0210510.1 | 1.038663 | -0.61919 | 4.526325 | 3.43E-05 | 0.004067 | 2.083709 | Rapid alkalinization factor 1 | elongation zone | 6 h |
| HORVU.MOREX.r3.1HG0028320.1 | -0.51736 | 6.850868 | -4.60113 | 2.66E-05 | 0.003405 | 2.077768 | 70 kDa heat shock protein | elongation zone | 6 h |
| HORVU.MOREX.r3.6HG0600910.1 | 1.525494 | 1.060689 | 4.586951 | 2.79E-05 | 0.003531 | 2.077711 | Aquaporin | elongation zone | 6 h |
| HORVU.MOREX.r3.1HG0050700.1 | 1.470383 | 2.994863 | 4.544497 | 3.22E-05 | 0.00389 | 2.071266 | ATP-dependent zinc metalloprotease FtsH | elongation zone | 6 h |
| HORVU.MOREX.r3.5HG0512230.1 | 1.496443 | 2.04829 | 4.478337 | 4.04E-05 | 0.004424 | 2.062678 | Protein ENHANCED DISEASE RESISTANCE 2-like | elongation zone | 6 h |
| HORVU.MOREX.r3.5HG0470690.1 | 1.690588 | -2.06961 | 4.480688 | 4.00E-05 | 0.004417 | 2.042055 | Heavy metal transport/detoxification protein | elongation zone | 6 h |
| HORVU.MOREX.r3.4HG0345580.1 | 2.430549 | -3.15122 | 4.669612 | 2.10E-05 | 0.003032 | 2.04184 | Dirigent protein | elongation zone | 6 h |
| HORVU.MOREX.r3.4HG0402730.1 | 1.916573 | -1.08162 | 4.494676 | 3.82E-05 | 0.004269 | 2.026897 | Homeobox protein BEL1 like | elongation zone | 6 h |
| HORVU.MOREX.r3.2HG0192410.1 | 1.946545 | 0.742041 | 4.555903 | 3.10E-05 | 0.003763 | 2.017531 | Peroxidase | elongation zone | 6 h |
| HORVU.MOREX.r3.7HG0722260.1 | 1.806626 | -2.17609 | 4.464401 | 4.23E-05 | 0.004481 | 2.015299 | RPM1-interacting protein 4 | elongation zone | 6 h |
| HORVU.MOREX.r3.1HG0066260.1 | -0.44426 | 9.175242 | -4.67254 | 2.08E-05 | 0.003023 | 1.9994 | Tubulin beta chain | elongation zone | 6 h |
| HORVU.MOREX.r3.7HG0742080.1 | 1.368426 | -1.3261 | 4.46957 | 4.16E-05 | 0.004467 | 1.998271 | Receptor-like kinase | elongation zone | 6 h |
| HORVU.MOREX.r3.3HG0230870.1 | 1.663841 | 0.065486 | 4.457413 | 4.33E-05 | 0.004481 | 1.99767 | Protein TRM32 | elongation zone | 6 h |
| HORVU.MOREX.r3.5HG0426110.1 | 2.400586 | -3.22479 | 4.619901 | 2.49E-05 | 0.003254 | 1.979889 | NBS-LRR class disease resistance protein | elongation zone | 6 h |
| HORVU.MOREX.r3.7HG0634520.1 | 1.552893 | -2.23987 | 4.45638 | 4.35E-05 | 0.004481 | 1.978574 | NBS-LRR-like resistance protein | elongation zone | 6 h |
| HORVU.MOREX.r3.1HG0065860.1 | 0.848897 | 4.236426 | 4.524346 | 3.45E-05 | 0.004071 | 1.97413 | MLO-like protein | elongation zone | 6 h |

**Table S1** Continued.

| ID | logFC | AveExpr | t | P.Value | adj.P.Val | B | gene description | root zone | time point |
|---|---|---|---|---|---|---|---|---|---|
| HORVU.MOREX.r3.3HG0264650.1 | 1.800089 | -2.1209 | 4.459394 | 4.30E-05 | 0.004481 | 1.973967 | Retrovirus-related Pol polyprotein from transposon TNT 1-94 | elongation zone | 6 h |
| HORVU.MOREX.r3.3HG0309760.1 | 1.133557 | 3.402606 | 4.455882 | 4.36E-05 | 0.004481 | 1.966706 | ROP-interactive CRIB motif protein | elongation zone | 6 h |
| HORVU.MOREX.r3.1HG0002960.1 | 1.189129 | 0.278873 | 4.514551 | 3.57E-05 | 0.004114 | 1.954537 | NBS-LRR disease resistance protein, putative, expressed | elongation zone | 6 h |
| HORVU.MOREX.r3.1HG0072910.2 | -0.51436 | 4.342105 | -4.47276 | 4.11E-05 | 0.004442 | 1.948413 | Serine/threonine-protein kinase ATM | elongation zone | 6 h |
| HORVU.MOREX.r3.7HG0641030.1 | 1.433632 | -0.36725 | 4.497113 | 3.79E-05 | 0.004269 | 1.948011 | Serpin family protein | elongation zone | 6 h |
| HORVU.MOREX.r3.6HG0622110.1 | 1.621992 | 2.81443 | 4.538426 | 3.29E-05 | 0.003948 | 1.947782 | Pathogenesis-related protein 1 | elongation zone | 6 h |
| HORVU.MOREX.r3.4HG0400740.1 | -0.30488 | 8.244974 | -4.60131 | 2.65E-05 | 0.003405 | 1.933258 | Cell division cycle protein 48-like protein | elongation zone | 6 h |
| HORVU.MOREX.r3.5HG0438460.1 | 1.637099 | 0.165145 | 4.474942 | 4.08E-05 | 0.004433 | 1.933105 | basic helix-loop-helix (bHLH) DNA-binding superfamily protein | elongation zone | 6 h |
| HORVU.MOREX.r3.7HG0636890.1 | 1.968408 | 0.940541 | 4.509718 | 3.63E-05 | 0.004159 | 1.84869 | Expansin-like protein | elongation zone | 6 h |
| HORVU.MOREX.r3.6HG0576010.1 | 0.688546 | 4.663156 | 4.448925 | 4.46E-05 | 0.00453 | 1.844889 | Protein kinase family protein | elongation zone | 6 h |
| HORVU.MOREX.r3.2HG0164850.1 | -0.83779 | 4.574579 | -4.45697 | 4.34E-05 | 0.004481 | 1.837023 | SBP (S-ribonuclease binding protein) family protein | elongation zone | 6 h |
| HORVU.MOREX.r3.7HG0666650.1 | -1.74807 | -0.31405 | -4.40513 | 5.17E-05 | 0.00498 | 1.826821 | Starch synthase, chloroplastic/amyloplastic | elongation zone | 6 h |
| HORVU.MOREX.r3.1HG0057030.1 | -0.85458 | 6.546046 | -4.50545 | 3.68E-05 | 0.004196 | 1.82504 | Peroxidase | elongation zone | 6 h |
| HORVU.MOREX.r3.1HG0084970.1 | 1.228861 | 2.620282 | 4.439007 | 4.61E-05 | 0.004599 | 1.814145 | Phosphatidylserine synthase 2 | elongation zone | 6 h |
| HORVU.MOREX.r3.6HG0550940.1 | -0.3901 | 5.078026 | -4.46072 | 4.28E-05 | 0.004481 | 1.799621 | obscurin-like protein | elongation zone | 6 h |
| HORVU.MOREX.r3.5HG0454660.1 | -0.40023 | 4.270815 | -4.45065 | 4.43E-05 | 0.004526 | 1.782595 | Alpha-(1,4)-fucosyltransferase | elongation zone | 6 h |
| HORVU.MOREX.r3.3HG0257680.1 | -0.50697 | 4.018531 | -4.41962 | 4.92E-05 | 0.004788 | 1.77026 | GDSL esterase/lipase | elongation zone | 6 h |
| HORVU.MOREX.r3.4HG0384390.1 | 1.194255 | 1.574903 | 4.39884 | 5.28E-05 | 0.00498 | 1.764321 | Protein kinase family protein, putative, expressed | elongation zone | 6 h |
| HORVU.MOREX.r3.3HG0238250.1 | -0.98537 | 6.327265 | -4.49393 | 3.83E-05 | 0.004269 | 1.759614 | Gigantea-like protein | elongation zone | 6 h |
| HORVU.MOREX.r3.5HG0480130.1 | 2.755074 | -2.63524 | 4.520606 | 3.50E-05 | 0.00408 | 1.754621 | Kinase family protein | elongation zone | 6 h |
| HORVU.MOREX.r3.4HG0406670.1 | 0.896728 | 6.546221 | 4.529063 | 3.40E-05 | 0.004052 | 1.750733 | Classical arabinogalactan protein 5 | elongation zone | 6 h |
| HORVU.MOREX.r3.3HG0253630.1 | 1.413934 | 3.730748 | 4.404741 | 5.18E-05 | 0.00498 | 1.738963 | Late embryogenesis abundant protein | elongation zone | 6 h |
| HORVU.MOREX.r3.1HG0093050.1 | 1.093019 | 4.062497 | 4.43109 | 4.74E-05 | 0.004673 | 1.737545 | Indole-3-acetic acid-amido synthetase GH3.3 | elongation zone | 6 h |
| HORVU.MOREX.r3.5HG0512210.1 | 3.384525 | -2.87785 | 4.580288 | 2.85E-05 | 0.003568 | 1.728528 | WRKY transcription factor | elongation zone | 6 h |
| HORVU.MOREX.r3.5HG0520990.1 | -0.74157 | 3.822638 | -4.39069 | 5.43E-05 | 0.005017 | 1.728368 | Kinase, putative | elongation zone | 6 h |
| HORVU.MOREX.r3.7HG0751450.1 | 2.087902 | 0.190404 | 4.397294 | 5.31E-05 | 0.00498 | 1.72746 | Glycosyltransferase family 92 protein | elongation zone | 6 h |
| HORVU.MOREX.r3.5HG0466440.1 | -0.49623 | 4.963631 | -4.44012 | 4.59E-05 | 0.004599 | 1.71567 | Glucan endo-1,3-beta-glucosidase, putative | elongation zone | 6 h |
| HORVU.MOREX.r3.6HG0576550.1 | -0.54599 | 4.943295 | -4.43697 | 4.64E-05 | 0.004603 | 1.693978 | F-box protein | elongation zone | 6 h |
| HORVU.MOREX.r3.7HG0724140.1 | 1.052735 | 4.661226 | 4.426327 | 4.81E-05 | 0.004726 | 1.682048 | FAD-binding Berberine family protein | elongation zone | 6 h |
| HORVU.MOREX.r3.5HG0489720.1 | -0.43189 | 6.28355 | -4.4771 | 4.05E-05 | 0.004424 | 1.675934 | WPP domain-interacting protein 1 | elongation zone | 6 h |
| HORVU.MOREX.r3.2HG0178480.1 | 1.158204 | -0.05624 | 4.39474 | 5.35E-05 | 0.004992 | 1.664541 | RING/U-box superfamily protein | elongation zone | 6 h |
| HORVU.MOREX.r3.3HG0307250.1 | 2.116851 | -2.48075 | 4.374902 | 5.72E-05 | 0.005129 | 1.661991 | Amino acid permease | elongation zone | 6 h |
| HORVU.MOREX.r3.2HG0115650.1 | -1.04194 | 6.223826 | -4.48697 | 3.92E-05 | 0.004347 | 1.660044 | #N/A | elongation zone | 6 h |
| HORVU.MOREX.r3.1HG0073330.1 | -0.41018 | 5.588457 | -4.44499 | 4.52E-05 | 0.004568 | 1.648802 | Ubiquitin conjugating enzyme E2 | elongation zone | 6 h |
| HORVU.MOREX.r3.5HG0469300.1 | 1.837129 | 0.250216 | 4.348195 | 6.26E-05 | 0.00546 | 1.644784 | CCCH-type zinc finger protein with ARM repeat domain-containing protein | elongation zone | 6 h |
| HORVU.MOREX.r3.7HG0726170.1 | -0.36883 | 7.833263 | -4.5189 | 3.52E-05 | 0.00408 | 1.637807 | Mitogen-activated protein kinase | elongation zone | 6 h |
| HORVU.MOREX.r3.7HG0637030.1 | 1.199843 | 0.131716 | 4.383858 | 5.55E-05 | 0.005066 | 1.63244 | Cytochrome P450 family protein, expressed | elongation zone | 6 h |
| HORVU.MOREX.r3.5HG0490610.1 | 1.806309 | -0.13337 | 4.337745 | 6.49E-05 | 0.005545 | 1.631143 | Protein kinase | elongation zone | 6 h |
| HORVU.MOREX.r3.5HG0497900.1 | 0.988943 | 3.680571 | 4.393535 | 5.38E-05 | 0.004992 | 1.566644 | Rhomboid-like protein | elongation zone | 6 h |
| HORVU.MOREX.r3.7HG0670900.1 | 1.665796 | 2.29029 | 4.346402 | 6.30E-05 | 0.00546 | 1.563141 | Aspartic proteinase Asp1 | elongation zone | 6 h |
| HORVU.MOREX.r3.5HG0488040.1 | 1.773284 | -1.23014 | 4.344843 | 6.33E-05 | 0.00546 | 1.562854 | Blue copper protein | elongation zone | 6 h |

**Table S1** Continued.

| ID | logFC | AveExpr | t | P.Value | adj.P.Val | B | gene description | root zone | time point |
|---|---|---|---|---|---|---|---|---|---|
| HORVU.MOREX.r3.2HG0175380.1 | 1.628515 | 3.452964 | 4.45505 | 4.37E-05 | 0.004481 | 1.55269 | Sec14p-like phosphatidylinositol transfer family protein | elongation zone | 6 h |
| HORVU.MOREX.r3.7HG0731560.1 | 1.82839 | -0.87401 | 4.381706 | 5.59E-05 | 0.00508 | 1.546359 | Regulator of Vps4 activity in the MVB pathway protein, putative | elongation zone | 6 h |
| HORVU.MOREX.r3.7HG0661830.1 | 2.311953 | -0.86487 | 4.303012 | 7.28E-05 | 0.005869 | 1.53054 | GPI-anchored adhesin-like protein | elongation zone | 6 h |
| HORVU.MOREX.r3.1HG0017280.1 | 1.705673 | -0.33011 | 4.345262 | 6.32E-05 | 0.00546 | 1.516513 | Peroxidase | elongation zone | 6 h |
| HORVU.MOREX.r3.4HG0394460.1 | 1.582218 | 0.734389 | 4.296061 | 7.46E-05 | 0.005869 | 1.500277 | Protein DETOXIFICATION | elongation zone | 6 h |
| HORVU.MOREX.r3.3HG0303210.1 | 1.045578 | 1.990963 | 4.38439 | 5.54E-05 | 0.005066 | 1.496242 | Alpha/beta-Hydrolases superfamily protein | elongation zone | 6 h |
| HORVU.MOREX.r3.3HG0257840.1 | 1.382705 | 0.153713 | 4.378903 | 5.65E-05 | 0.005083 | 1.493426 | S-acyltransferase | elongation zone | 6 h |
| HORVU.MOREX.r3.3HG0267430.2 | -0.70694 | 3.386958 | -4.31704 | 6.95E-05 | 0.005693 | 1.486845 | Receptor-like protein kinase | elongation zone | 6 h |
| HORVU.MOREX.r3.2HG0127260.1 | 1.422613 | -1.06197 | 4.333444 | 6.58E-05 | 0.005579 | 1.485352 | COBRA-like protein | elongation zone | 6 h |
| HORVU.MOREX.r3.6HG0621170.1 | 2.497718 | -0.64178 | 4.295036 | 7.48E-05 | 0.005869 | 1.48204 | GPI-anchored adhesin-like protein | elongation zone | 6 h |
| HORVU.MOREX.r3.6HG0610980.1 | 1.484187 | -1.88326 | 4.287107 | 7.68E-05 | 0.005948 | 1.471212 | Nucleotide/sugar transporter family protein | elongation zone | 6 h |
| HORVU.MOREX.r3.6HG0548710.1 | 2.399406 | -1.47311 | 4.281006 | 7.84E-05 | 0.00598 | 1.461884 | Kinase family protein | elongation zone | 6 h |
| HORVU.MOREX.r3.3HG0310110.1 | 1.932063 | 0.692973 | 4.278796 | 7.90E-05 | 0.00598 | 1.458126 | lysine ketoglutarate reductase trans-splicing-like protein, putative (DUF707) | elongation zone | 6 h |
| HORVU.MOREX.r3.2HG0168700.1 | -1.93174 | 0.035179 | -4.46276 | 4.25E-05 | 0.004481 | 1.454954 | ADP-ribosylation factor GTPase-activating protein AGD4 | elongation zone | 6 h |
| HORVU.MOREX.r3.7HG0715570.1 | -0.48768 | 6.466383 | -4.40172 | 5.23E-05 | 0.00498 | 1.454003 | Transmembrane protein | elongation zone | 6 h |
| HORVU.MOREX.r3.3HG0228600.1 | -0.83809 | 3.194099 | -4.29945 | 7.37E-05 | 0.005869 | 1.451827 | transmembrane protein, putative (DUF247) | elongation zone | 6 h |
| HORVU.MOREX.r3.1HG0083250.1 | -0.79226 | 3.094919 | -4.27951 | 7.88E-05 | 0.00598 | 1.450196 | Protein phosphatase 2c, putative | elongation zone | 6 h |
| HORVU.MOREX.r3.2HG0112190.1 | 1.781051 | -0.62536 | 4.305506 | 7.22E-05 | 0.005859 | 1.44703 | Transcription repressor ofp17 | elongation zone | 6 h |
| HORVU.MOREX.r3.1HG0051110.1 | -0.42943 | 4.84593 | -4.35597 | 6.10E-05 | 0.005389 | 1.434594 | Ankyrin repeat-containing protein, putative | elongation zone | 6 h |
| HORVU.MOREX.r3.6HG0605120.1 | -0.94165 | 2.469228 | -4.28756 | 7.67E-05 | 0.005948 | 1.432225 | RING/U-box superfamily protein | elongation zone | 6 h |
| HORVU.MOREX.r3.4HG0334360.1 | 1.012982 | 4.121321 | 4.37247 | 5.77E-05 | 0.005149 | 1.410669 | Homeobox protein BEL1 like | elongation zone | 6 h |
| HORVU.MOREX.r3.5HG0449620.1 | 0.680084 | 4.972296 | 4.345739 | 6.31E-05 | 0.00546 | 1.409923 | weak chloroplast movement under blue light protein (DUF827) | elongation zone | 6 h |
| HORVU.MOREX.r3.6HG0577040.1 | 2.174545 | -1.97815 | 4.325824 | 6.75E-05 | 0.005655 | 1.409541 | Ycf1 | elongation zone | 6 h |
| HORVU.MOREX.r3.2HG0144760.1 | 1.224079 | -1.38996 | 4.267582 | 8.20E-05 | 0.006162 | 1.407 | tRNA 5-methylaminomethyl-2-thiouridine biosynthesis bifunctional protein MnmC | elongation zone | 6 h |
| HORVU.MOREX.r3.5HG0512490.1 | 1.796482 | 1.425049 | 4.384888 | 5.54E-05 | 0.005066 | 1.398205 | Peroxidase | elongation zone | 6 h |
| HORVU.MOREX.r3.3HG0330980.1 | -0.38188 | 6.458067 | -4.39777 | 5.30E-05 | 0.00498 | 1.38759 | Protein IQ-DOMAIN 1 | elongation zone | 6 h |
| HORVU.MOREX.r3.2HG0157410.1 | 1.282714 | 2.254585 | 4.422845 | 4.87E-05 | 0.004759 | 1.379488 | Peroxidase | elongation zone | 6 h |
| HORVU.MOREX.r3.5HG0512710.1 | 1.953505 | 0.114178 | 4.252321 | 8.63E-05 | 0.006257 | 1.377069 | N-acetyltransferase, putative, expressed | elongation zone | 6 h |
| HORVU.MOREX.r3.3HG0310460.1 | -0.8055 | 0.959034 | -4.31324 | 7.04E-05 | 0.005732 | 1.371256 | Glycosyltransferase | elongation zone | 6 h |
| HORVU.MOREX.r3.5HG0533760.1 | 1.286314 | 3.713664 | 4.402306 | 5.22E-05 | 0.00498 | 1.357668 | Glycosyltransferase | elongation zone | 6 h |
| HORVU.MOREX.r3.7HG0721690.1 | 1.277014 | 0.537157 | 4.317998 | 6.93E-05 | 0.005693 | 1.356246 | Endoglucanase | elongation zone | 6 h |
| HORVU.MOREX.r3.2HG0214240.1 | -0.72002 | 3.183258 | -4.2562 | 8.52E-05 | 0.006257 | 1.350852 | Disease resistance protein (NBS-LRR class) family | elongation zone | 6 h |
| HORVU.MOREX.r3.3HG0306060.1 | -0.40301 | 5.417997 | -4.34218 | 6.39E-05 | 0.005486 | 1.345803 | Thioredoxin-like protein | elongation zone | 6 h |
| HORVU.MOREX.r3.6HG0617970.1 | 1.743656 | -1.86338 | 4.241216 | 8.95E-05 | 0.006432 | 1.345331 | Glycosyltransferase family 92 protein | elongation zone | 6 h |
| HORVU.MOREX.r3.7HG0655210.1 | 1.73036 | -1.68992 | 4.257075 | 8.49E-05 | 0.006257 | 1.330013 | Peroxidase | elongation zone | 6 h |
| HORVU.MOREX.r3.3HG0286100.1 | 2.060245 | 1.587249 | 4.272694 | 8.06E-05 | 0.006081 | 1.328105 | CASP-like protein | elongation zone | 6 h |
| HORVU.MOREX.r3.4HG0412370.1 | 2.248347 | -1.42512 | 4.232964 | 9.20E-05 | 0.006466 | 1.313575 | Homeobox protein knotted-1, putative | elongation zone | 6 h |
| HORVU.MOREX.r3.3HG0331030.1 | -0.42461 | 7.059382 | -4.37941 | 5.64E-05 | 0.005083 | 1.311415 | Chaperone protein dnaJ, putative | elongation zone | 6 h |
| HORVU.MOREX.r3.7HG0647950.1 | 2.328223 | -1.042 | 4.247695 | 8.76E-05 | 0.006329 | 1.309073 | Peroxidase | elongation zone | 6 h |
| HORVU.MOREX.r3.5HG0485780.1 | 3.087395 | -0.7336 | 4.235739 | 9.11E-05 | 0.006466 | 1.294071 | 1-aminocyclopropane-1-carboxylate oxidase | elongation zone | 6 h |
| HORVU.MOREX.r3.3HG0233990.1 | 1.291557 | 1.603331 | 4.396975 | 5.31E-05 | 0.00498 | 1.287834 | Xyloglucan endotransglucosylase/hydrolase | elongation zone | 6 h |

**Table S1** Continued.

| ID | logFC | AveExpr | t | P.Value | adj.P.Val | B | gene description | root zone | time point |
|---|---|---|---|---|---|---|---|---|---|
| HORVU.MOREX.r3.2HG0113720.1 | 1.376604 | 3.679958 | 4.330514 | 6.64E-05 | 0.005611 | 1.277074 | Cytosolic Fe-S cluster assembly factor nbp35 | elongation zone | 6 h |
| HORVU.MOREX.r3.2HG0157710.1 | -0.85214 | 2.534028 | -4.25414 | 8.57E-05 | 0.006257 | 1.276901 | actin cytoskeleton-regulatory complex pan-like protein | elongation zone | 6 h |
| HORVU.MOREX.r3.7HG0736150.1 | 1.217607 | 8.171407 | 4.43866 | 4.62E-05 | 0.004599 | 1.276341 | Phosphatidylinositol transfer protein | elongation zone | 6 h |
| HORVU.MOREX.r3.2HG0208520.1 | 2.349203 | -2.76284 | 4.323872 | 6.79E-05 | 0.005667 | 1.257069 | Heavy metal transport/detoxification superfamily protein | elongation zone | 6 h |
| HORVU.MOREX.r3.6HG0612780.1 | 1.626017 | 1.073388 | 4.214609 | 9.78E-05 | 0.006777 | 1.247681 | Dynein light chain | elongation zone | 6 h |
| HORVU.MOREX.r3.3HG0303240.1 | 1.266074 | 2.978435 | 4.210707 | 9.90E-05 | 0.006797 | 1.228241 | Ankyrin repeat-containing protein | elongation zone | 6 h |
| HORVU.MOREX.r3.4HG0382610.1 | 1.304492 | 0.223451 | 4.298199 | 7.40E-05 | 0.005869 | 1.227494 | cytochrome P450, family 705, subfamily A, polypeptide 21 | elongation zone | 6 h |
| HORVU.MOREX.r3.6HG0585240.1 | 1.030864 | 0.751154 | 4.301068 | 7.33E-05 | 0.005869 | 1.224965 | Kinase superfamily protein | elongation zone | 6 h |
| HORVU.MOREX.r3.7HG0696770.1 | -0.32991 | 5.470938 | -4.29995 | 7.36E-05 | 0.005869 | 1.224712 | Protein kinase | elongation zone | 6 h |
| HORVU.MOREX.r3.7HG0737340.1 | 1.656782 | -0.10666 | 4.318196 | 6.92E-05 | 0.005693 | 1.224188 | GDA1/CD39 nucleoside phosphatase family protein | elongation zone | 6 h |
| HORVU.MOREX.r3.7HG0665040.1 | 3.136968 | -3.45264 | 4.501356 | 3.73E-05 | 0.004232 | 1.220843 | Protein EXORDIUM-like 1 | elongation zone | 6 h |
| HORVU.MOREX.r3.7HG0641750.1 | 2.886043 | -2.73901 | 4.234967 | 9.14E-05 | 0.006466 | 1.212339 | Aspartic proteinase nepenthesin-2 | elongation zone | 6 h |
| HORVU.MOREX.r3.5HG0533430.1 | 1.446154 | 0.093461 | 4.197599 | 0.000103 | 0.007027 | 1.211266 | ROP-interactive CRIB motif protein | elongation zone | 6 h |
| HORVU.MOREX.r3.4HG0393870.1 | 1.509975 | -1.59561 | 4.217434 | 9.69E-05 | 0.006737 | 1.20776 | Peroxidase | elongation zone | 6 h |
| HORVU.MOREX.r3.7HG0667610.1 | 1.533121 | 3.317002 | 4.316481 | 6.96E-05 | 0.005693 | 1.197908 | Peroxidase | elongation zone | 6 h |
| HORVU.MOREX.r3.2HG0161410.1 | -0.33139 | 7.35188 | -4.35509 | 6.12E-05 | 0.005389 | 1.179522 | Acyl-[acyl-carrier-protein] desaturase | elongation zone | 6 h |
| HORVU.MOREX.r3.7HG0640560.1 | -3.05237 | 0.442209 | -4.51862 | 3.52E-05 | 0.00408 | 1.178265 | NBS-LRR disease resistance protein-like protein | elongation zone | 6 h |
| HORVU.MOREX.r3.7HG0721080.1 | -0.88879 | 4.617684 | -4.25844 | 8.45E-05 | 0.006257 | 1.177088 | Zinc finger, B-box | elongation zone | 6 h |
| HORVU.MOREX.r3.1HG0092010.1 | 0.723654 | 6.335202 | 4.318587 | 6.91E-05 | 0.005693 | 1.173644 | #N/A | elongation zone | 6 h |
| HORVU.MOREX.r3.6HG0613020.1 | 0.73279 | 4.924387 | 4.293425 | 7.52E-05 | 0.005869 | 1.17256 | Caleosin | elongation zone | 6 h |
| HORVU.MOREX.r3.3HG0280630.1 | -0.41087 | 5.914468 | -4.29776 | 7.41E-05 | 0.005869 | 1.172527 | Glycosylphosphatidylinositol anchor attachment 1 protein | elongation zone | 6 h |
| HORVU.MOREX.r3.7HG0720100.1 | 0.785019 | 4.342288 | 4.228113 | 9.35E-05 | 0.006547 | 1.17236 | Glucan endo-1,3-beta-glucosidase, putative | elongation zone | 6 h |
| HORVU.MOREX.r3.7HG0722360.1 | 1.657991 | 0.664931 | 4.208263 | 9.98E-05 | 0.006829 | 1.171515 | Receptor-like protein kinase | elongation zone | 6 h |
| HORVU.MOREX.r3.7HG0707700.1 | 1.28134 | -2.42617 | 4.188758 | 0.000106 | 0.007165 | 1.169507 | Late embryogenesis abundant (LEA) hydroxyproline-rich glycoprotein family | elongation zone | 6 h |
| HORVU.MOREX.r3.4HG0415420.1 | -0.71383 | 6.947214 | -4.32574 | 6.75E-05 | 0.005655 | 1.164807 | Tubulin beta chain | elongation zone | 6 h |
| HORVU.MOREX.r3.7HG0667620.1 | 1.390352 | 0.317173 | 4.282499 | 7.80E-05 | 0.005973 | 1.141699 | Peroxidase | elongation zone | 6 h |
| HORVU.MOREX.r3.3HG0218800.1 | 2.036577 | -0.5154 | 4.174834 | 0.000112 | 0.007346 | 1.138679 | hydroxyproline-rich glycoprotein family protein | elongation zone | 6 h |
| HORVU.MOREX.r3.2HG0215060.1 | 2.56843 | -1.59998 | 4.239693 | 9.00E-05 | 0.006432 | 1.135309 | WAT1-related protein | elongation zone | 6 h |
| HORVU.MOREX.r3.4HG0402030.1 | -0.48854 | 5.321938 | -4.25809 | 8.46E-05 | 0.006257 | 1.133581 | Basic helix-loop-helix (BHLH) family transcription factor | elongation zone | 6 h |
| HORVU.MOREX.r3.1HG0078970.1 | 1.606309 | 2.920959 | 4.286019 | 7.71E-05 | 0.005948 | 1.126937 | Protein kinase, putative | elongation zone | 6 h |
| HORVU.MOREX.r3.3HG0235320.1 | 2.090663 | -0.14202 | 4.182294 | 0.000109 | 0.007272 | 1.124894 | Pollen Ole e 1 allergen and extensin family protein | elongation zone | 6 h |
| HORVU.MOREX.r3.1HG0012340.1 | 1.341273 | 2.437594 | 4.258932 | 8.44E-05 | 0.006257 | 1.124798 | Triose-phosphate transporter family protein | elongation zone | 6 h |
| HORVU.MOREX.r3.2HG0212860.1 | 1.881986 | -1.30303 | 4.167517 | 0.000114 | 0.007387 | 1.119368 | #N/A | elongation zone | 6 h |
| HORVU.MOREX.r3.7HG0736960.1 | -0.80623 | 3.87866 | -4.18744 | 0.000107 | 0.007173 | 1.114131 | BTB/POZ domain containing protein, expressed | elongation zone | 6 h |
| HORVU.MOREX.r3.7HG0751300.1 | 1.431848 | -1.39316 | 4.174339 | 0.000112 | 0.007346 | 1.103678 | Phosphatidylinositol N-acetyglucosaminlytransferase subunit P-like protein | elongation zone | 6 h |
| HORVU.MOREX.r3.3HG0221180.1 | 1.198148 | 2.379805 | 4.335977 | 6.52E-05 | 0.005555 | 1.09362 | Glycosyltransferase | elongation zone | 6 h |
| HORVU.MOREX.r3.1HG0094980.1 | -0.59508 | 4.287159 | -4.20004 | 0.000103 | 0.006994 | 1.082257 | Mitochondrial outer membrane porin | elongation zone | 6 h |
| HORVU.MOREX.r3.4HG0378580.1 | -0.5394 | 5.353084 | -4.25469 | 8.56E-05 | 0.006257 | 1.072092 | Transcription factor | elongation zone | 6 h |
| HORVU.MOREX.r3.7HG0724060.1 | 1.472051 | 0.588187 | 4.191324 | 0.000106 | 0.007128 | 1.06404 | U-box domain-containing protein | elongation zone | 6 h |
| HORVU.MOREX.r3.5HG0486660.1 | 1.327783 | -2.06977 | 4.135973 | 0.000127 | 0.008027 | 1.031241 | Calcium-binding protein | elongation zone | 6 h |
| HORVU.MOREX.r3.2HG0205990.1 | 2.277694 | -2.55766 | 4.178281 | 0.00011 | 0.007321 | 1.007939 | Glutaredoxin family protein, putative | elongation zone | 6 h |

**Table S1** Continued.

| ID | logFC | AveExpr | t | P.Value | adj.P.Val | B | gene description | root zone | time point |
|---|---|---|---|---|---|---|---|---|---|
| HORVU.MOREX.r3.6HG0623140.1 | 2.367924 | -2.34532 | 4.233996 | 9.17E-05 | 0.006466 | 1.00097 | #N/A | elongation zone | 6 h |
| HORVU.MOREX.r3.7HG0647300.1 | 2.344324 | -1.17681 | 4.128858 | 0.00013 | 0.008166 | 0.997754 | AT hook motif DNA-binding family protein | elongation zone | 6 h |
| HORVU.MOREX.r3.7HG0729000.1 | 1.037744 | 3.645636 | 4.176245 | 0.000111 | 0.007346 | 0.991496 | RING/U-box superfamily protein | elongation zone | 6 h |
| HORVU.MOREX.r3.5HG0511450.1 | 0.425871 | 5.650127 | 4.239968 | 8.99E-05 | 0.006432 | 0.985226 | ABC transporter B family protein | elongation zone | 6 h |
| HORVU.MOREX.r3.3HG0275060.1 | 1.379367 | 0.275212 | 4.171049 | 0.000113 | 0.007387 | 0.981393 | Isoprenylcysteine alpha-carbonyl methylesterase ICME | elongation zone | 6 h |
| HORVU.MOREX.r3.6HG0567780.1 | -0.73168 | 3.016753 | -4.12133 | 0.000133 | 0.008269 | 0.954213 | disease resistance protein (TIR-NBS-LRR class) | elongation zone | 6 h |
| HORVU.MOREX.r3.3HG0252960.1 | 1.52771 | -1.0236 | 4.106352 | 0.00014 | 0.008505 | 0.944621 | Calmodulin-binding protein | elongation zone | 6 h |
| HORVU.MOREX.r3.2HG0198870.1 | 1.180277 | -0.51668 | 4.163197 | 0.000116 | 0.007453 | 0.943646 | UPF0761 membrane protein Plut_1323 | elongation zone | 6 h |
| HORVU.MOREX.r3.6HG0570230.1 | 1.162509 | 0.733369 | 4.104145 | 0.000141 | 0.008516 | 0.927627 | Aldose 1-epimerase family protein | elongation zone | 6 h |
| HORVU.MOREX.r3.7HG0735700.1 | 1.751337 | 3.307934 | 4.114349 | 0.000136 | 0.00831 | 0.919377 | Patatin | elongation zone | 6 h |
| HORVU.MOREX.r3.1HG0077960.1 | 0.713733 | 7.898538 | 4.283421 | 7.78E-05 | 0.005973 | 0.872625 | #N/A | elongation zone | 6 h |
| HORVU.MOREX.r3.1HG0075260.1 | 3.591717 | 0.393741 | 4.218048 | 9.67E-05 | 0.006737 | 0.872368 | Receptor kinase 2 | elongation zone | 6 h |
| HORVU.MOREX.r3.1HG0000050.1 | -0.48669 | 5.048249 | -4.16686 | 0.000114 | 0.007387 | 0.853426 | RING-finger ubiquitin ligase | elongation zone | 6 h |
| HORVU.MOREX.r3.4HG0412090.1 | 1.838635 | -0.36354 | 4.072261 | 0.000156 | 0.009027 | 0.845801 | C3HC4-type RING finger-containing protein | elongation zone | 6 h |
| HORVU.MOREX.r3.2HG0187450.1 | 1.502526 | 0.035956 | 4.167227 | 0.000114 | 0.007387 | 0.831957 | Rhomboid-like protein | elongation zone | 6 h |
| HORVU.MOREX.r3.1HG0069570.1 | -0.38714 | 8.695958 | -4.29352 | 7.52E-05 | 0.005869 | 0.829353 | Tubulin beta chain | elongation zone | 6 h |
| HORVU.MOREX.r3.5HG0532120.1 | 1.958329 | -1.75561 | 4.080201 | 0.000152 | 0.008905 | 0.828908 | Expansin | elongation zone | 6 h |
| HORVU.MOREX.r3.5HG0511820.1 | 2.863174 | -3.38085 | 4.19273 | 0.000105 | 0.007118 | 0.826277 | Cyclic nucleotide-gated channel | elongation zone | 6 h |
| HORVU.MOREX.r3.3HG0319900.1 | 1.526392 | -0.77823 | 4.087232 | 0.000149 | 0.008845 | 0.82349 | Ankyrin repeat domain-containing protein 1 | elongation zone | 6 h |
| HORVU.MOREX.r3.2HG0164470.1 | 0.885954 | 3.24648 | 4.147607 | 0.000122 | 0.007749 | 0.81663 | Nudix hydrolase | elongation zone | 6 h |
| HORVU.MOREX.r3.5HG0457310.1 | 1.160147 | 0.314313 | 4.151574 | 0.00012 | 0.007696 | 0.802061 | Exocyst complex component, putative | elongation zone | 6 h |
| HORVU.MOREX.r3.7HG0680960.1 | 2.001292 | 0.611498 | 4.052042 | 0.000167 | 0.009235 | 0.786995 | electron transporter, putative (Protein of unknown function, DUF547) | elongation zone | 6 h |
| HORVU.MOREX.r3.5HG0494040.1 | 1.637166 | -3.00086 | 4.080794 | 0.000152 | 0.008905 | 0.766255 | Auxin efflux carrier component | elongation zone | 6 h |
| HORVU.MOREX.r3.4HG0337720.1 | 2.735816 | 1.69822 | 4.357119 | 6.08E-05 | 0.005389 | 0.764459 | PHD finger protein family | elongation zone | 6 h |
| HORVU.MOREX.r3.7HG0730660.1 | 1.69589 | -2.73673 | 4.079872 | 0.000152 | 0.008905 | 0.760753 | #N/A | elongation zone | 6 h |
| HORVU.MOREX.r3.5HG0500360.1 | 3.396968 | 1.290708 | 4.115784 | 0.000135 | 0.008295 | 0.753335 | L-gulonolactone oxidase | elongation zone | 6 h |
| HORVU.MOREX.r3.3HG0309780.1 | 3.054003 | -2.21583 | 4.116906 | 0.000135 | 0.008295 | 0.743116 | RING/U-box superfamily protein | elongation zone | 6 h |
| HORVU.MOREX.r3.7HG0723240.1 | 3.595489 | -0.72296 | 4.211192 | 9.89E-05 | 0.006797 | 0.739836 | TMV response-related protein | elongation zone | 6 h |
| HORVU.MOREX.r3.2HG0100750.1 | 1.493323 | -1.62151 | 4.063312 | 0.000161 | 0.00914 | 0.730312 | Dirigent protein | elongation zone | 6 h |
| HORVU.MOREX.r3.1HG0053060.1 | 1.094966 | 4.011378 | 4.091674 | 0.000147 | 0.008787 | 0.7239 | PLATZ transcription factor | elongation zone | 6 h |
| HORVU.MOREX.r3.6HG0593260.1 | 0.781168 | 5.929813 | 4.120134 | 0.000134 | 0.008277 | 0.722746 | LETM1 and EF-hand domain-containing protein 1, mitochondrial | elongation zone | 6 h |
| HORVU.MOREX.r3.5HG0500120.1 | -1.06176 | 4.270851 | -4.06631 | 0.000159 | 0.009076 | 0.719081 | Thaumatin | elongation zone | 6 h |
| HORVU.MOREX.r3.3HG0293570.1 | -2.42284 | -0.38794 | -4.21086 | 9.90E-05 | 0.006797 | 0.718965 | BTB/POZ domain-containing protein | elongation zone | 6 h |
| HORVU.MOREX.r3.4HG0335450.1 | 2.849427 | -2.95917 | 4.134345 | 0.000127 | 0.008045 | 0.713885 | Nucleobase ascorbate transporter | elongation zone | 6 h |
| HORVU.MOREX.r3.3HG0234970.1 | 0.680893 | 0.091122 | 4.101721 | 0.000142 | 0.008559 | 0.713597 | 6,7-dimethyl-8-ribityllumazine synthase | elongation zone | 6 h |
| HORVU.MOREX.r3.2HG0102670.1 | 2.231459 | -3.01135 | 4.122112 | 0.000133 | 0.008269 | 0.709464 | Dirigent protein | elongation zone | 6 h |
| HORVU.MOREX.r3.5HG0521610.1 | 1.350561 | 0.681734 | 4.158361 | 0.000118 | 0.00755 | 0.706186 | Protein kinase | elongation zone | 6 h |
| HORVU.MOREX.r3.7HG0728480.1 | 1.577404 | 0.878141 | 4.180847 | 0.000109 | 0.007283 | 0.700674 | Peroxidase | elongation zone | 6 h |
| HORVU.MOREX.r3.4HG0333450.1 | 0.677657 | 3.618973 | 4.069688 | 0.000157 | 0.009044 | 0.697865 | Type I inositol-1,4,5-trisphosphate 5-phosphatase CVP2 | elongation zone | 6 h |
| HORVU.MOREX.r3.2HG0118220.1 | 1.200541 | 1.251256 | 4.077872 | 0.000153 | 0.008933 | 0.693078 | En/Spm-like transposon protein | elongation zone | 6 h |
| HORVU.MOREX.r3.2HG0205420.1 | 0.881002 | -0.18096 | 4.077168 | 0.000154 | 0.008933 | 0.686703 | #N/A | elongation zone | 6 h |

**Table S1** Continued.

| ID | logFC | AveExpr | t | P.Value | adj.P.Val | B | gene description | root zone | time point |
|---|---|---|---|---|---|---|---|---|---|
| HORVU.MOREX.r3.1HG0067720.1 | 1.31055 | -2.10602 | 4.01713 | 0.000187 | 0.009858 | 0.680253 | #N/A | elongation zone | 6 h |
| HORVU.MOREX.r3.7HG0735840.1 | -1.04726 | 4.848168 | -4.07252 | 0.000156 | 0.009027 | 0.68008 | serine/arginine repetitive matrix-like protein | elongation zone | 6 h |
| HORVU.MOREX.r3.7HG0641860.1 | 2.105613 | 0.566345 | 4.036788 | 0.000175 | 0.009467 | 0.670204 | Ubiquitin carboxyl-terminal hydrolase 2 | elongation zone | 6 h |
| HORVU.MOREX.r3.2HG0181880.1 | -0.34812 | 6.222963 | -4.16736 | 0.000114 | 0.007387 | 0.666309 | #N/A | elongation zone | 6 h |
| HORVU.MOREX.r3.7HG0662370.1 | 1.725993 | -1.65272 | 4.021453 | 0.000184 | 0.009796 | 0.660686 | Pathogenesis-related protein 1 | elongation zone | 6 h |
| HORVU.MOREX.r3.3HG0298180.1 | 1.714999 | -1.34466 | 4.009429 | 0.000192 | 0.009907 | 0.659972 | #N/A | elongation zone | 6 h |
| HORVU.MOREX.r3.3HG0252170.1 | 1.672983 | -1.70754 | 4.004244 | 0.000195 | 0.009946 | 0.648623 | DUF1685 family protein | elongation zone | 6 h |
| HORVU.MOREX.r3.5HG0509350.1 | 2.091009 | -1.08767 | 4.08212 | 0.000151 | 0.008905 | 0.640197 | Dirigent protein | elongation zone | 6 h |
| HORVU.MOREX.r3.7HG0673330.1 | 1.633347 | -2.01567 | 3.998802 | 0.000198 | 0.010073 | 0.631308 | Embryogenesis transmembrane protein-like | elongation zone | 6 h |
| HORVU.MOREX.r3.4HG0412030.1 | -0.5293 | 4.498999 | -4.06149 | 0.000162 | 0.009144 | 0.630798 | vacuolar sorting-associated protein (DUF946) | elongation zone | 6 h |
| HORVU.MOREX.r3.4HG0342680.1 | 1.265446 | -0.60974 | 4.057766 | 0.000164 | 0.009196 | 0.623076 | Branchpoint-bridging protein | elongation zone | 6 h |
| HORVU.MOREX.r3.2HG0139810.1 | 1.024182 | 0.63654 | 4.119041 | 0.000134 | 0.008281 | 0.621047 | Heavy metal transport/detoxification superfamily protein | elongation zone | 6 h |
| HORVU.MOREX.r3.5HG0483980.1 | 1.49107 | -1.84454 | 3.992059 | 0.000203 | 0.010208 | 0.60216 | Cinnamoyl CoA reductase | elongation zone | 6 h |
| HORVU.MOREX.r3.3HG0281730.1 | -1.52473 | 3.703997 | -4.01298 | 0.000189 | 0.009875 | 0.597608 | AP2/B3 transcription factor family protein | elongation zone | 6 h |
| HORVU.MOREX.r3.3HG0238180.1 | -0.4947 | 4.144587 | -4.0498 | 0.000168 | 0.009235 | 0.596744 | #N/A | elongation zone | 6 h |
| HORVU.MOREX.r3.4HG0380540.1 | 0.71167 | 3.365322 | 4.048467 | 0.000169 | 0.009235 | 0.594237 | Aquaporin-1 | elongation zone | 6 h |
| HORVU.MOREX.r3.6HG0609680.1 | 1.5598 | 2.000697 | 3.987149 | 0.000206 | 0.010307 | 0.591637 | Cobyric acid synthase | elongation zone | 6 h |
| HORVU.MOREX.r3.3HG0234010.1 | 1.626236 | -0.09759 | 4.116028 | 0.000135 | 0.008295 | 0.590752 | Xyloglucan endotransglucosylase/hydrolase | elongation zone | 6 h |
| HORVU.MOREX.r3.3HG0244000.1 | 2.012072 | -2.4581 | 4.091016 | 0.000147 | 0.008787 | 0.581592 | Core-2/I-branching beta-1,6-N-acetylglucosaminyltransferase family protein | elongation zone | 6 h |
| HORVU.MOREX.r3.2HG0193170.1 | 5.202316 | -0.2671 | 4.252191 | 8.63E-05 | 0.006257 | 0.578647 | Pollen Ole e 1 allergen/extensin | elongation zone | 6 h |
| HORVU.MOREX.r3.7HG0648620.1 | 1.334729 | -1.54284 | 3.981217 | 0.00021 | 0.010399 | 0.567671 | 2-oxoglutarate (2OG) and Fe(II)-dependent oxygenase superfamily protein | elongation zone | 6 h |
| HORVU.MOREX.r3.2HG0170230.1 | 0.620513 | 3.492538 | 4.005225 | 0.000194 | 0.009939 | 0.567565 | Smr domain containing protein | elongation zone | 6 h |
| HORVU.MOREX.r3.3HG0255700.1 | 0.936777 | 1.416296 | 3.984093 | 0.000208 | 0.010359 | 0.561064 | PQ-loop repeat family protein / transmembrane family protein | elongation zone | 6 h |
| HORVU.MOREX.r3.5HG0507680.1 | 1.509332 | -1.22844 | 4.005809 | 0.000194 | 0.009939 | 0.557729 | mitogen-activated protein kinase kinase kinase 6 | elongation zone | 6 h |
| HORVU.MOREX.r3.3HG0280060.1 | 1.897938 | -2.86395 | 4.009069 | 0.000192 | 0.009907 | 0.545693 | Heavy metal transport/detoxification superfamily protein | elongation zone | 6 h |
| HORVU.MOREX.r3.5HG0469340.1 | 1.364091 | 0.552227 | 4.003011 | 0.000196 | 0.009961 | 0.542595 | Fasciclin-like arabinogalactan-protein-like | elongation zone | 6 h |
| HORVU.MOREX.r3.6HG0553150.1 | -0.59336 | 6.876917 | -4.12536 | 0.000131 | 0.008235 | 0.534801 | Polyubiquitin | elongation zone | 6 h |
| HORVU.MOREX.r3.3HG0241320.1 | -0.57499 | 3.585418 | -4.01749 | 0.000187 | 0.009858 | 0.532925 | Prolyl 4-hydroxylase alpha subunit, putative | elongation zone | 6 h |
| HORVU.MOREX.r3.2HG0189530.1 | 1.502506 | 0.494093 | 4.084248 | 0.00015 | 0.00888 | 0.52337 | Expansin protein | elongation zone | 6 h |
| HORVU.MOREX.r3.3HG0264640.1 | 1.450044 | -2.14748 | 3.957494 | 0.000227 | 0.010873 | 0.513937 | Retrotransposon protein, putative, unclassified | elongation zone | 6 h |
| HORVU.MOREX.r3.3HG0288710.2 | 1.666455 | -1.63853 | 3.960326 | 0.000225 | 0.010819 | 0.512944 | GDSL esterase/lipase | elongation zone | 6 h |
| HORVU.MOREX.r3.6HG0601670.1 | -1.03885 | 2.45038 | -3.9553 | 0.000228 | 0.010879 | 0.50788 | Phosphatidic acid phosphatase | elongation zone | 6 h |
| HORVU.MOREX.r3.5HG0464440.1 | 1.046278 | 0.420648 | 4.055712 | 0.000165 | 0.009196 | 0.500905 | Kinase, putative | elongation zone | 6 h |
| HORVU.MOREX.r3.2HG0178050.1 | 0.671937 | 4.229626 | 4.049295 | 0.000168 | 0.009235 | 0.498841 | F-box protein | elongation zone | 6 h |
| HORVU.MOREX.r3.7HG0729510.1 | -0.66076 | 4.709018 | -4.05656 | 0.000164 | 0.009196 | 0.493535 | WEB family protein, chloroplastic | elongation zone | 6 h |
| HORVU.MOREX.r3.2HG0157450.1 | 1.0854 | 2.551061 | 4.14807 | 0.000122 | 0.007749 | 0.490267 | Peroxidase | elongation zone | 6 h |
| HORVU.MOREX.r3.3HG0319430.1 | 0.918067 | 2.262365 | 3.953545 | 0.00023 | 0.010879 | 0.482009 | WD40-repeat protein | elongation zone | 6 h |
| HORVU.MOREX.r3.1HG0075320.1 | 1.179116 | 0.461847 | 4.057132 | 0.000164 | 0.009196 | 0.481515 | Myb transcription factor | elongation zone | 6 h |
| HORVU.MOREX.r3.2HG0181680.1 | 1.807609 | -1.90882 | 3.965294 | 0.000221 | 0.010722 | 0.480716 | CASP-like protein | elongation zone | 6 h |
| HORVU.MOREX.r3.1HG0090170.1 | -0.58794 | 5.133692 | -4.0215 | 0.000184 | 0.009796 | 0.473711 | NBS-LRR-like resistance protein | elongation zone | 6 h |
| HORVU.MOREX.r3.4HG0410830.1 | 1.226737 | 1.674186 | 4.045177 | 0.000171 | 0.009285 | 0.473068 | Transcription factor | elongation zone | 6 h |

**Table S1** Continued.

| ID | logFC | AveExpr | t | P.Value | adj.P.Val | B | gene description | root zone | time point |
|---|---|---|---|---|---|---|---|---|---|
| HORVU.MOREX.r3.7HG0727750.1 | 1.271368 | 5.212856 | 4.049506 | 0.000168 | 0.009235 | 0.460302 | Zinc-transporting ATPase | elongation zone | 6 h |
| HORVU.MOREX.r3.5HG0478450.1 | -0.38866 | 5.597721 | -4.05554 | 0.000165 | 0.009196 | 0.439217 | AUGMIN subunit 5 | elongation zone | 6 h |
| HORVU.MOREX.r3.7HG0728870.1 | -0.67588 | 6.913212 | -4.16851 | 0.000114 | 0.007387 | 0.43008 | Xyloglucan endotransglucosylase/hydrolase | elongation zone | 6 h |
| HORVU.MOREX.r3.4HG0397010.1 | 1.588331 | 0.173587 | 3.955603 | 0.000228 | 0.010879 | 0.42269 | Beta-glucosidase | elongation zone | 6 h |
| HORVU.MOREX.r3.3HG0305150.1 | -0.69222 | 4.756714 | -4.0073 | 0.000193 | 0.009922 | 0.42243 | Pectin lyase-like superfamily protein | elongation zone | 6 h |
| HORVU.MOREX.r3.2HG0186480.1 | 2.389649 | -0.52788 | 3.930602 | 0.000247 | 0.011494 | 0.420658 | 1-aminocyclopropane-1-carboxylate synthase | elongation zone | 6 h |
| HORVU.MOREX.r3.3HG0275490.1 | 1.905526 | -2.47526 | 3.933074 | 0.000245 | 0.011462 | 0.420242 | Zinc finger CCCH domain protein | elongation zone | 6 h |
| HORVU.MOREX.r3.6HG0558350.1 | 3.157234 | -0.25422 | 4.069723 | 0.000157 | 0.009044 | 0.410293 | #N/A | elongation zone | 6 h |
| HORVU.MOREX.r3.4HG0335170.1 | 1.363685 | 1.390233 | 4.087215 | 0.000149 | 0.008845 | 0.400681 | Core-2/I-branching beta-1,6-N-acetylglucosaminyltransferase family protein | elongation zone | 6 h |
| HORVU.MOREX.r3.1HG0005730.1 | 2.121992 | -3.38812 | 4.036771 | 0.000175 | 0.009467 | 0.398848 | transcriptional factor B3 family protein | elongation zone | 6 h |
| HORVU.MOREX.r3.2HG0207720.1 | 0.911135 | 0.894337 | 3.981078 | 0.00021 | 0.010399 | 0.398137 | Flavin-containing monooxygenase | elongation zone | 6 h |
| HORVU.MOREX.r3.3HG0304780.1 | -0.51325 | 4.983038 | -4.02358 | 0.000183 | 0.009779 | 0.396785 | Peptidyl serine alpha-galactosyltransferase | elongation zone | 6 h |
| HORVU.MOREX.r3.7HG0688870.1 | 0.781888 | 6.011072 | 4.084672 | 0.00015 | 0.00888 | 0.39521 | Monodehydroascorbate reductase | elongation zone | 6 h |
| HORVU.MOREX.r3.7HG0731490.1 | -0.45292 | 4.892075 | -4.00856 | 0.000192 | 0.009907 | 0.394041 | Transmembrane protein, putative | elongation zone | 6 h |
| HORVU.MOREX.r3.5HG0420460.1 | 2.673472 | -1.03028 | 4.122825 | 0.000132 | 0.008269 | 0.390359 | NBS-LRR-like resistance protein | elongation zone | 6 h |
| HORVU.MOREX.r3.3HG0242950.1 | -0.43332 | 6.032036 | -4.0472 | 0.000169 | 0.009249 | 0.389262 | NF-X1 type zinc finger family protein | elongation zone | 6 h |
| HORVU.MOREX.r3.1HG0023200.1 | 1.268951 | 0.569684 | 3.911926 | 0.000262 | 0.011846 | 0.38519 | LURP-one-like protein | elongation zone | 6 h |
| HORVU.MOREX.r3.4HG0412140.1 | 1.223956 | 2.514643 | 4.01206 | 0.00019 | 0.009875 | 0.381269 | Zinc finger (C3HC4-type RING finger) protein family-like | elongation zone | 6 h |
| HORVU.MOREX.r3.2HG0168070.1 | 1.302242 | 0.66133 | 4.068249 | 0.000158 | 0.009044 | 0.378793 | Peroxidase | elongation zone | 6 h |
| HORVU.MOREX.r3.4HG0400520.1 | 1.361771 | -0.786 | 3.95733 | 0.000227 | 0.010873 | 0.374991 | Lipid transfer protein | elongation zone | 6 h |
| HORVU.MOREX.r3.7HG0684180.1 | 0.646146 | 6.139387 | 4.100743 | 0.000142 | 0.008561 | 0.366996 | Peroxidase | elongation zone | 6 h |
| HORVU.MOREX.r3.4HG0412650.1 | 2.405969 | 1.247202 | 4.068558 | 0.000158 | 0.009044 | 0.365997 | TVP38/TMEM64 family membrane protein | elongation zone | 6 h |
| HORVU.MOREX.r3.4HG0381790.1 | 1.152783 | 3.851367 | 4.049967 | 0.000168 | 0.009235 | 0.363944 | transmembrane protein, putative (DUF247) | elongation zone | 6 h |
| HORVU.MOREX.r3.3HG0326430.1 | -0.3652 | 5.095188 | -4.01458 | 0.000188 | 0.009875 | 0.363147 | Diphosphomevalonate decarboxylase | elongation zone | 6 h |
| HORVU.MOREX.r3.2HG0210610.1 | 0.90861 | 3.089943 | 3.971519 | 0.000217 | 0.010595 | 0.361942 | Acetyl-coenzyme A synthetase | elongation zone | 6 h |
| HORVU.MOREX.r3.3HG0322650.1 | 0.820926 | 3.559006 | 4.015792 | 0.000188 | 0.009875 | 0.36008 | NAD kinase | elongation zone | 6 h |
| HORVU.MOREX.r3.2HG0185600.1 | 1.219804 | 0.732582 | 4.02494 | 0.000182 | 0.009761 | 0.35993 | TRIO and F-actin-binding protein | elongation zone | 6 h |
| HORVU.MOREX.r3.6HG0634130.1 | -0.34476 | 5.21873 | -4.02736 | 0.000181 | 0.009718 | 0.35688 | Alpha-1,3-mannosyl-glycoprotein 2-beta-N-acetylglucosaminyltransferase | elongation zone | 6 h |
| HORVU.MOREX.r3.7HG0649240.1 | 1.654371 | -2.99572 | 3.995882 | 0.0002 | 0.010144 | 0.351149 | protein kinase family protein | elongation zone | 6 h |
| HORVU.MOREX.r3.7HG0727650.1 | 2.785394 | -1.35022 | 4.01979 | 0.000185 | 0.009823 | 0.338712 | BTB/POZ domain-containing protein | elongation zone | 6 h |
| HORVU.MOREX.r3.2HG0098090.1 | -1.22748 | -2.03609 | -3.90116 | 0.000272 | 0.012171 | 0.328123 | Translation initiation factor IF-2 | elongation zone | 6 h |
| HORVU.MOREX.r3.1HG0081620.1 | -0.67083 | 3.155534 | -3.91547 | 0.000259 | 0.01178 | 0.305948 | Glycerophosphodiester phosphodiesterase | elongation zone | 6 h |
| HORVU.MOREX.r3.7HG0739120.1 | 1.056623 | 2.007183 | 3.980317 | 0.000211 | 0.010399 | 0.280998 | Receptor-like protein kinase | elongation zone | 6 h |
| HORVU.MOREX.r3.2HG0214130.1 | 1.478431 | 3.799443 | 3.979911 | 0.000211 | 0.010399 | 0.279119 | CoA ligase | elongation zone | 6 h |
| HORVU.MOREX.r3.2HG0210500.1 | 1.440157 | 3.185342 | 4.104754 | 0.00014 | 0.008516 | 0.277495 | Endoglucanase | elongation zone | 6 h |
| HORVU.MOREX.r3.3HG0230640.1 | 2.149865 | -1.25694 | 3.874376 | 0.000296 | 0.012756 | 0.276546 | Thioesterase family protein | elongation zone | 6 h |
| HORVU.MOREX.r3.2HG0179790.1 | 1.090504 | 3.322266 | 3.890761 | 0.000281 | 0.012448 | 0.275276 | Zinc finger ring-type protein | elongation zone | 6 h |
| HORVU.MOREX.r3.3HG0278170.1 | 1.707695 | 0.346839 | 3.872558 | 0.000298 | 0.01278 | 0.273946 | CCCH-type zinc fingerfamily protein with RNA-binding domain-containing protein | elongation zone | 6 h |
| HORVU.MOREX.r3.7HG0730530.1 | 1.30542 | 0.190357 | 3.91596 | 0.000259 | 0.01178 | 0.264659 | Copper transport protein family | elongation zone | 6 h |
| HORVU.MOREX.r3.2HG0192910.1 | -0.7874 | 6.73787 | -4.05151 | 0.000167 | 0.009235 | 0.262668 | Xyloglucan endotransglucosylase/hydrolase | elongation zone | 6 h |
| HORVU.MOREX.r3.2HG0140170.1 | 0.729722 | 2.981209 | 3.912956 | 0.000262 | 0.011846 | 0.262423 | Ras-related protein Rab-21 | elongation zone | 6 h |

**Table S1** Continued.

| ID | logFC | AveExpr | t | P.Value | adj.P.Val | B | gene description | root zone | time point |
|---|---|---|---|---|---|---|---|---|---|
| HORVU.MOREX.r3.3HG0294950.1 | 1.165905 | -0.01143 | 3.947297 | 0.000234 | 0.011075 | 0.261902 | Deubiquitinase SseL | elongation zone | 6 h |
| HORVU.MOREX.r3.3HG0324180.1 | -0.86838 | 4.528561 | -3.92277 | 0.000253 | 0.011636 | 0.261303 | #N/A | elongation zone | 6 h |
| HORVU.MOREX.r3.4HG0401730.1 | -0.71392 | 2.933703 | -3.88505 | 0.000286 | 0.012542 | 0.259644 | Pentatricopeptide repeat-containing protein | elongation zone | 6 h |
| HORVU.MOREX.r3.4HG0406060.1 | 2.287349 | -1.45049 | 3.874045 | 0.000296 | 0.012756 | 0.250466 | Agmatine coumaroyltransferase-2 | elongation zone | 6 h |
| HORVU.MOREX.r3.6HG0617750.1 | -0.58528 | 4.935682 | -3.99166 | 0.000203 | 0.010208 | 0.243819 | Plant cadmium resistance 2 | elongation zone | 6 h |
| HORVU.MOREX.r3.4HG0407310.1 | 0.93114 | 3.764737 | 3.98741 | 0.000206 | 0.010307 | 0.236033 | Xyloglucan galactosyltransferase KATAMARI1-like protein | elongation zone | 6 h |
| HORVU.MOREX.r3.1HG0059350.1 | -0.58908 | 4.615638 | -3.9447 | 0.000236 | 0.011121 | 0.234833 | S-adenosyl-L-methionine-dependent methyltransferases superfamily protein | elongation zone | 6 h |
| HORVU.MOREX.r3.4HG0395940.1 | 1.431855 | 3.064976 | 3.893856 | 0.000278 | 0.012352 | 0.232016 | Homocysteine S-methyltransferase family protein | elongation zone | 6 h |
| HORVU.MOREX.r3.4HG0349530.1 | -0.39697 | 4.965038 | -3.96357 | 0.000222 | 0.010731 | 0.225711 | Tubby-like F-box protein | elongation zone | 6 h |
| HORVU.MOREX.r3.5HG0420480.1 | 2.172748 | 1.423525 | 3.855048 | 0.000315 | 0.013274 | 0.209779 | BEL1-like homeodomain protein 6 | elongation zone | 6 h |
| HORVU.MOREX.r3.2HG0190260.1 | 1.048626 | 2.916863 | 3.857826 | 0.000312 | 0.013202 | 0.202002 | SHR5-receptor-like kinase | elongation zone | 6 h |
| HORVU.MOREX.r3.7HG0675770.1 | -0.6 | 4.417254 | -3.97472 | 0.000214 | 0.010525 | 0.196545 | basic helix-loop-helix (bHLH) DNA-binding superfamily protein | elongation zone | 6 h |
| HORVU.MOREX.r3.3HG0313900.1 | 1.79453 | 0.291481 | 3.844516 | 0.000326 | 0.013547 | 0.193434 | Methyl esterase 1 | elongation zone | 6 h |
| HORVU.MOREX.r3.6HG0620520.1 | 0.766727 | 3.09452 | 3.977305 | 0.000213 | 0.010462 | 0.19341 | Aminotransferase-related family protein | elongation zone | 6 h |
| HORVU.MOREX.r3.6HG0595190.1 | 1.697851 | -2.07211 | 3.838398 | 0.000332 | 0.013678 | 0.167222 | GDSL esterase/lipase | elongation zone | 6 h |
| HORVU.MOREX.r3.3HG0293320.1 | 1.344097 | 0.363569 | 3.969018 | 0.000218 | 0.010619 | 0.166294 | ALK tyrosine kinase receptor | elongation zone | 6 h |
| HORVU.MOREX.r3.1HG0079020.1 | 2.109964 | 3.471696 | 3.859969 | 0.00031 | 0.013139 | 0.160129 | Protein kinase, putative | elongation zone | 6 h |
| HORVU.MOREX.r3.2HG0096230.1 | 1.3345 | -2.9355 | 3.852368 | 0.000317 | 0.013296 | 0.156505 | Phosphatidylinositol 3- and 4-kinase family protein | elongation zone | 6 h |
| HORVU.MOREX.r3.3HG0228940.1 | 1.160842 | 0.760498 | 3.884121 | 0.000287 | 0.012552 | 0.145085 | Sulfate adenylyltransferase subunit 2 | elongation zone | 6 h |
| HORVU.MOREX.r3.3HG0256860.1 | 2.909361 | -0.92611 | 4.253358 | 8.60E-05 | 0.006257 | 0.143357 | Pentatricopeptide repeat-containing protein | elongation zone | 6 h |
| HORVU.MOREX.r3.3HG0298510.1 | 0.710619 | 4.425452 | 3.930013 | 0.000248 | 0.011494 | 0.142572 | 1-phosphatidylinositol-3-phosphate 5-kinase | elongation zone | 6 h |
| HORVU.MOREX.r3.1HG0044290.1 | 0.806982 | 3.742696 | 3.847944 | 0.000322 | 0.013457 | 0.141382 | electron protein, putative (Protein of unknown function, DUF547) | elongation zone | 6 h |
| HORVU.MOREX.r3.4HG0355210.1 | 2.094068 | -0.13769 | 3.830327 | 0.000341 | 0.013839 | 0.134843 | transcription factor-like protein | elongation zone | 6 h |
| HORVU.MOREX.r3.3HG0255020.1 | 0.949433 | 3.48561 | 4.041865 | 0.000172 | 0.009361 | 0.121671 | Peroxidase | elongation zone | 6 h |
| HORVU.MOREX.r3.3HG0304490.1 | 1.083847 | -1.40069 | 3.832099 | 0.000339 | 0.013839 | 0.117976 | Myosin IC heavy chain | elongation zone | 6 h |
| HORVU.MOREX.r3.3HG0254930.1 | 2.620343 | -0.9917 | 3.816908 | 0.000355 | 0.014243 | 0.117873 | Peroxidase | elongation zone | 6 h |
| HORVU.MOREX.r3.4HG0380330.1 | -0.49912 | 5.401503 | -3.94057 | 0.000239 | 0.01124 | 0.117543 | WRKY transcription factor | elongation zone | 6 h |
| HORVU.MOREX.r3.4HG0349310.1 | -0.411 | 5.375879 | -3.9368 | 0.000242 | 0.011351 | 0.116605 | Sec1 family domain-containing protein 2 | elongation zone | 6 h |
| HORVU.MOREX.r3.4HG0399970.1 | 1.108416 | 4.109756 | 3.954162 | 0.000229 | 0.010879 | 0.107234 | Cytochrome b561 and DOMON domain-containing protein | elongation zone | 6 h |
| HORVU.MOREX.r3.4HG0341050.1 | 1.365415 | 0.610576 | 3.863356 | 0.000307 | 0.013079 | 0.105564 | Protein DETOXIFICATION | elongation zone | 6 h |
| HORVU.MOREX.r3.5HG0436660.1 | -0.47656 | 4.868326 | -3.92783 | 0.000249 | 0.011499 | 0.103439 | #N/A | elongation zone | 6 h |
| HORVU.MOREX.r3.4HG0340030.1 | 1.905496 | -1.31134 | 3.853895 | 0.000316 | 0.013286 | 0.099939 | beta-fructofuranosidase 5 | elongation zone | 6 h |
| HORVU.MOREX.r3.1HG0050220.1 | 1.467589 | -1.46767 | 3.814825 | 0.000358 | 0.01431 | 0.097532 | Polyketide cyclase/dehydrase and lipid transport superfamily protein | elongation zone | 6 h |
| HORVU.MOREX.r3.6HG0606850.1 | -0.45929 | 6.605397 | -3.9712 | 0.000217 | 0.010595 | 0.096013 | Methyltransferase | elongation zone | 6 h |
| HORVU.MOREX.r3.3HG0293760.1 | 1.372775 | 3.466627 | 3.916563 | 0.000259 | 0.01178 | 0.092678 | External alternative NAD(P)H-ubiquinone oxidoreductase B2, mitochondrial | elongation zone | 6 h |
| HORVU.MOREX.r3.3HG0222130.1 | -0.34138 | 8.795276 | -4.06196 | 0.000162 | 0.009144 | 0.092182 | Transmembrane protein 214 | elongation zone | 6 h |
| HORVU.MOREX.r3.2HG0207130.1 | 1.449031 | 1.218085 | 3.920773 | 0.000255 | 0.011684 | 0.088096 | Acetyl-coenzyme A synthetase | elongation zone | 6 h |
| HORVU.MOREX.r3.4HG0376720.1 | 2.748408 | -0.40522 | 4.23288 | 9.20E-05 | 0.006466 | 0.077218 | #N/A | elongation zone | 6 h |
| HORVU.MOREX.r3.3HG0296120.1 | 1.531733 | 0.584762 | 3.862166 | 0.000308 | 0.013101 | 0.075777 | Late embryogenesis abundant (LEA) hydroxyproline-rich glycoprotein family | elongation zone | 6 h |
| HORVU.MOREX.r3.7HG0643260.1 | -0.63975 | 3.282608 | -3.83478 | 0.000336 | 0.013808 | 0.073857 | BTB/POZ domain-containing protein | elongation zone | 6 h |
| HORVU.MOREX.r3.6HG0631710.1 | 1.017095 | 2.878977 | 4.02712 | 0.000181 | 0.009718 | 0.065104 | Aquaporin | elongation zone | 6 h |

**Table S1** Continued.

| ID | logFC | AveExpr | t | P.Value | adj.P.Val | B | gene description | root zone | time point |
|---|---|---|---|---|---|---|---|---|---|
| HORVU.MOREX.r3.4HG0331520.1 | -0.58664 | 3.556196 | -3.89973 | 0.000273 | 0.012201 | 0.063368 | Protein UPSTREAM OF FLC | elongation zone | 6 h |
| HORVU.MOREX.r3.2HG0209680.1 | 1.514912 | 0.534924 | 3.8234 | 0.000348 | 0.014059 | 0.0507 | Isopropylmalate dehydrogenase | elongation zone | 6 h |
| HORVU.MOREX.r3.2HG0164160.2 | 1.454876 | -0.40053 | 3.790717 | 0.000386 | 0.014897 | 0.044823 | Trihelix transcription factor | elongation zone | 6 h |
| HORVU.MOREX.r3.7HG0747940.1 | 0.444076 | 7.327177 | 4.012837 | 0.000189 | 0.009875 | 0.029498 | Polyamine oxidase | elongation zone | 6 h |
| HORVU.MOREX.r3.6HG0573920.1 | 1.414493 | -0.43481 | 3.793134 | 0.000383 | 0.014864 | 0.029255 | Retrotransposon protein, putative, unclassified | elongation zone | 6 h |
| HORVU.MOREX.r3.2HG0105830.1 | 0.792199 | 4.383346 | 3.852965 | 0.000317 | 0.013296 | 0.025007 | #N/A | elongation zone | 6 h |
| HORVU.MOREX.r3.5HG0519660.1 | 0.466698 | 3.342751 | 3.83078 | 0.00034 | 0.013839 | 0.021559 | Basic helix-loop-helix transcription factor | elongation zone | 6 h |
| HORVU.MOREX.r3.6HG0631600.1 | 1.116608 | 1.162908 | 3.92945 | 0.000248 | 0.011494 | 0.016681 | Rop guanine nucleotide exchange factor, putative | elongation zone | 6 h |
| HORVU.MOREX.r3.6HG0547660.1 | 2.243889 | -1.78969 | 3.800782 | 0.000374 | 0.014703 | 0.011529 | Copine-1 | elongation zone | 6 h |
| HORVU.MOREX.r3.2HG0207860.1 | 1.448619 | -2.16402 | 3.778425 | 0.000401 | 0.015004 | 0.010905 | Serine/threonine-protein kinase | elongation zone | 6 h |
| HORVU.MOREX.r3.6HG0624390.1 | 1.263709 | 1.652699 | 3.778864 | 0.000401 | 0.015004 | 0.009373 | Two-component response regulator | elongation zone | 6 h |
| HORVU.MOREX.r3.3HG0285150.1 | 0.858919 | 1.826069 | 3.840902 | 0.000329 | 0.013651 | 0.008922 | Tryptophan aminotransferase | elongation zone | 6 h |
| HORVU.MOREX.r3.5HG0536200.1 | 1.101929 | -0.5546 | 3.886604 | 0.000285 | 0.01251 | 0.004664 | Xyloglucan endotransglucosylase/hydrolase | elongation zone | 6 h |
| HORVU.MOREX.r3.4HG0405860.1 | 2.432806 | 1.28141 | 3.779565 | 0.0004 | 0.015004 | 0.001351 | Protein WEAK CHLOROPLAST MOVEMENT UNDER BLUE LIGHT 1 | elongation zone | 6 h |
| HORVU.MOREX.r3.4HG0402000.1 | -0.71102 | 8.180685 | -3.99194 | 0.000203 | 0.010208 | -0.00288 | Kinase interacting (KIP1-like) family protein, putative | elongation zone | 6 h |
| HORVU.MOREX.r3.2HG0111270.1 | 1.104611 | -0.29425 | 3.788881 | 0.000388 | 0.014941 | -0.00676 | WUSCHEL-related homeobox | elongation zone | 6 h |
| HORVU.MOREX.r3.4HG0338070.1 | 0.975103 | 1.603513 | 3.923496 | 0.000253 | 0.011634 | -0.00773 | Kinase family protein | elongation zone | 6 h |
| HORVU.MOREX.r3.1HG0057990.1 | -0.60781 | 4.42891 | -3.83889 | 0.000331 | 0.013678 | -0.0088 | O-fucosyltransferase family protein | elongation zone | 6 h |
| HORVU.MOREX.r3.5HG0454230.1 | 0.775647 | 2.88731 | 3.82286 | 0.000349 | 0.014059 | -0.01562 | Aminopeptidase | elongation zone | 6 h |
| HORVU.MOREX.r3.1HG0052120.1 | -0.62021 | 3.548831 | -3.79064 | 0.000386 | 0.014897 | -0.01867 | GDP-mannose transporter | elongation zone | 6 h |
| HORVU.MOREX.r3.5HG0461950.1 | 0.850185 | 2.97161 | 3.838985 | 0.000331 | 0.013678 | -0.01903 | 2-isopropylmalate synthase 2 | elongation zone | 6 h |
| HORVU.MOREX.r3.7HG0741080.1 | -0.79685 | 1.851569 | -3.77862 | 0.000401 | 0.015004 | -0.01948 | Disease resistance protein RPM1 | elongation zone | 6 h |
| HORVU.MOREX.r3.4HG0335180.1 | -0.62283 | 5.021928 | -3.85483 | 0.000315 | 0.013274 | -0.02139 | Phosphatidylinositol N-acetyglucosaminlytransferase subunit P-like protein | elongation zone | 6 h |
| HORVU.MOREX.r3.6HG0604200.1 | -1.00506 | 0.762306 | -3.77857 | 0.000401 | 0.015004 | -0.02657 | transcription repressor | elongation zone | 6 h |
| HORVU.MOREX.r3.1HG0051920.1 | 0.957177 | 2.286248 | 3.87961 | 0.000291 | 0.012681 | -0.03212 | Transcription factor | elongation zone | 6 h |
| HORVU.MOREX.r3.3HG0327810.1 | -0.48527 | 7.008521 | -3.97021 | 0.000218 | 0.010603 | -0.03357 | Auxin efflux carrier protein-like | elongation zone | 6 h |
| HORVU.MOREX.r3.5HG0486810.1 | -1.00151 | 1.496685 | -3.76948 | 0.000413 | 0.015322 | -0.04016 | Chaperone DnaJ | elongation zone | 6 h |
| HORVU.MOREX.r3.6HG0555660.2 | -0.4305 | 7.208638 | -3.95416 | 0.000229 | 0.010879 | -0.04056 | Calreticulin/calnexin | elongation zone | 6 h |
| HORVU.MOREX.r3.5HG0513780.1 | 0.602815 | 5.051864 | 3.905561 | 0.000268 | 0.012054 | -0.04911 | Kinesin-like protein | elongation zone | 6 h |
| HORVU.MOREX.r3.2HG0218010.1 | -0.38361 | 8.228324 | -3.98591 | 0.000207 | 0.010324 | -0.04939 | Dolichyl-diphosphooligosaccharide--protein glycosyltransferase subunit 1 | elongation zone | 6 h |
| HORVU.MOREX.r3.4HG0417320.1 | 1.409745 | 1.098725 | 3.911669 | 0.000263 | 0.011846 | -0.05517 | Remorin | elongation zone | 6 h |
| HORVU.MOREX.r3.3HG0305440.1 | 1.566672 | -2.50175 | 3.755657 | 0.000431 | 0.015665 | -0.05772 | Late embryogenesis abundant protein | elongation zone | 6 h |
| HORVU.MOREX.r3.7HG0680410.1 | -0.64709 | 4.544521 | -3.86575 | 0.000304 | 0.013006 | -0.058 | Leucine-rich repeat (LRR) family protein | elongation zone | 6 h |
| HORVU.MOREX.r3.4HG0355460.1 | -0.46368 | 4.5138 | -3.8738 | 0.000297 | 0.012756 | -0.06185 | myosin-4 protein (DUF641) | elongation zone | 6 h |
| HORVU.MOREX.r3.5HG0493080.1 | 0.649714 | 5.851163 | 3.88754 | 0.000284 | 0.01251 | -0.06214 | VWA domain containing protein | elongation zone | 6 h |
| HORVU.MOREX.r3.7HG0675310.1 | 1.456171 | 1.411092 | 3.886514 | 0.000285 | 0.01251 | -0.06712 | N-lysine methyltransferase | elongation zone | 6 h |
| HORVU.MOREX.r3.1HG0079800.1 | -0.50169 | 7.794068 | -3.96403 | 0.000222 | 0.010731 | -0.06984 | Farnesyl diphosphate synthase | elongation zone | 6 h |
| HORVU.MOREX.r3.7HG0707160.1 | 1.688228 | -0.16087 | 3.745022 | 0.000446 | 0.015908 | -0.08672 | NAC domain-containing protein, putative | elongation zone | 6 h |
| HORVU.MOREX.r3.6HG0588520.1 | 0.942761 | 2.56033 | 3.882779 | 0.000288 | 0.01258 | -0.08795 | Rop guanine nucleotide exchange factor | elongation zone | 6 h |
| HORVU.MOREX.r3.6HG0557070.1 | -0.56379 | 6.4953 | -3.89578 | 0.000276 | 0.012309 | -0.08865 | Activator of heat shock protein ATPase | elongation zone | 6 h |
| HORVU.MOREX.r3.4HG0383670.1 | 1.153631 | 2.012771 | 3.861216 | 0.000309 | 0.013114 | -0.09291 | Protein SCAR2 | elongation zone | 6 h |

**Table S1** Continued.

| ID | logFC | AveExpr | t | P.Value | adj.P.Val | B | gene description | root zone | time point |
|---|---|---|---|---|---|---|---|---|---|
| HORVU.MOREX.r3.1HG0062110.1 | 1.355093 | 0.941667 | 3.746878 | 0.000443 | 0.015878 | -0.09391 | Pentatricopeptide repeat-containing protein | elongation zone | 6 h |
| HORVU.MOREX.r3.2HG0185370.1 | 2.612507 | -1.4233 | 3.812331 | 0.000361 | 0.014368 | -0.09839 | Subtilisin-like protease | elongation zone | 6 h |
| HORVU.MOREX.r3.7HG0704450.1 | 0.525161 | 5.126567 | 3.874262 | 0.000296 | 0.012756 | -0.10426 | Hexosyltransferase | elongation zone | 6 h |
| HORVU.MOREX.r3.5HG0481760.1 | -0.29019 | 6.00272 | -3.89561 | 0.000277 | 0.012309 | -0.10428 | Protein AUXIN RESPONSE 4 | elongation zone | 6 h |
| HORVU.MOREX.r3.6HG0559990.1 | 0.815896 | 5.045463 | 3.887002 | 0.000284 | 0.01251 | -0.11135 | Apoptosis-inducing factor-like protein A | elongation zone | 6 h |
| HORVU.MOREX.r3.5HG0484660.1 | 1.489123 | -0.23036 | 3.755081 | 0.000432 | 0.015665 | -0.1296 | Vacuolar-processing enzyme | elongation zone | 6 h |
| HORVU.MOREX.r3.7HG0751340.1 | 1.044816 | 0.25739 | 3.843936 | 0.000326 | 0.013547 | -0.13244 | Core-2/I-branching beta-1,6-N-acetylglucosaminyltransferase family protein | elongation zone | 6 h |
| HORVU.MOREX.r3.7HG0700270.1 | -0.39241 | 7.724735 | -3.94456 | 0.000236 | 0.011121 | -0.13247 | Transaldolase | elongation zone | 6 h |
| HORVU.MOREX.r3.4HG0356860.1 | 0.993279 | 1.759753 | 3.751236 | 0.000437 | 0.01578 | -0.1355 | Ulp1 protease family, C-terminal catalytic domain containing protein, expressed | elongation zone | 6 h |
| HORVU.MOREX.r3.7HG0671360.1 | -0.33715 | 9.214028 | -4.01186 | 0.00019 | 0.009875 | -0.13979 | Aldehyde dehydrogenase | elongation zone | 6 h |
| HORVU.MOREX.r3.5HG0531070.1 | 3.405701 | -3.02117 | 3.929373 | 0.000248 | 0.011494 | -0.14075 | Protein DETOXIFICATION | elongation zone | 6 h |
| HORVU.MOREX.r3.4HG0340910.2 | 2.331663 | -1.88941 | 3.793259 | 0.000383 | 0.014864 | -0.14156 | p-loop containing nucleoside triphosphate hydrolases superfamily protein | elongation zone | 6 h |
| HORVU.MOREX.r3.2HG0202250.1 | 1.350384 | 1.379273 | 3.74269 | 0.000449 | 0.01595 | -0.14444 | Kelch repeat-containing protein | elongation zone | 6 h |
| HORVU.MOREX.r3.2HG0177790.1 | 2.107683 | -2.43363 | 3.764456 | 0.00042 | 0.0154 | -0.14944 | Methyl-CpG-binding domain protein 6 | elongation zone | 6 h |
| HORVU.MOREX.r3.4HG0342660.1 | 1.245412 | 1.013503 | 3.877014 | 0.000293 | 0.012756 | -0.15427 | Calcium-binding EF-hand family protein | elongation zone | 6 h |
| HORVU.MOREX.r3.1HG0014680.1 | 1.397217 | -0.77974 | 3.710919 | 0.000496 | 0.017032 | -0.17483 | Tetraspanin family protein | elongation zone | 6 h |
| HORVU.MOREX.r3.5HG0469120.1 | 1.58902 | -1.16488 | 3.740688 | 0.000452 | 0.016007 | -0.18193 | E3 ubiquitin-protein ligase SINA-like 10 | elongation zone | 6 h |
| HORVU.MOREX.r3.3HG0246300.1 | 0.485266 | 4.385187 | 3.808929 | 0.000364 | 0.014439 | -0.20381 | RNA-binding protein, putative | elongation zone | 6 h |
| HORVU.MOREX.r3.7HG0653380.1 | 1.360238 | 1.304596 | 3.916315 | 0.000259 | 0.01178 | -0.2097 | GDSL esterase/lipase | elongation zone | 6 h |
| HORVU.MOREX.r3.6HG0608960.1 | 0.675326 | 3.939137 | 3.781867 | 0.000397 | 0.015004 | -0.21182 | Galactan beta-1,4-galactosyltransferase GALS1 | elongation zone | 6 h |
| HORVU.MOREX.r3.4HG0386490.1 | 2.000337 | -0.76068 | 3.731226 | 0.000466 | 0.016253 | -0.21763 | Receptor-like kinase, putative | elongation zone | 6 h |
| HORVU.MOREX.r3.4HG0412790.1 | 0.657907 | 3.163327 | 3.779355 | 0.0004 | 0.015004 | -0.22337 | Remorin | elongation zone | 6 h |
| HORVU.MOREX.r3.6HG0567150.1 | 1.074184 | 2.844231 | 3.694699 | 0.000522 | 0.017624 | -0.22725 | Exosome complex component | elongation zone | 6 h |
| HORVU.MOREX.r3.3HG0299020.1 | 1.221284 | -1.58695 | 3.692077 | 0.000526 | 0.01771 | -0.22832 | Protein NRT1/ PTR FAMILY 5.5 | elongation zone | 6 h |
| HORVU.MOREX.r3.6HG0609720.1 | -0.70447 | 3.853974 | -3.74045 | 0.000452 | 0.016007 | -0.22834 | Glucosamine 6-phosphate N-acetyltransferase | elongation zone | 6 h |
| HORVU.MOREX.r3.4HG0416760.1 | -0.58607 | 4.391267 | -3.78399 | 0.000394 | 0.015004 | -0.23116 | Non-structural protein 1 | elongation zone | 6 h |
| HORVU.MOREX.r3.7HG0719830.1 | -1.73302 | 0.487996 | -3.69789 | 0.000517 | 0.017564 | -0.23631 | #N/A | elongation zone | 6 h |
| HORVU.MOREX.r3.6HG0597590.1 | -0.40757 | 6.568014 | -3.86957 | 0.000301 | 0.012875 | -0.2374 | Protein disulfide-isomerase | elongation zone | 6 h |
| HORVU.MOREX.r3.3HG0288710.1 | 1.482542 | -1.93307 | 3.689136 | 0.000531 | 0.01781 | -0.23991 | GDSL esterase/lipase | elongation zone | 6 h |
| HORVU.MOREX.r3.2HG0099900.1 | 0.580535 | 5.512063 | 3.81764 | 0.000355 | 0.014238 | -0.24151 | Glyoxylate reductase/hydroxypyruvate reductase | elongation zone | 6 h |
| HORVU.MOREX.r3.4HG0332930.1 | -1.6159 | 0.588838 | -3.8102 | 0.000363 | 0.014437 | -0.24713 | Short-chain dehydrogenase/reductase family protein | elongation zone | 6 h |
| HORVU.MOREX.r3.7HG0654480.1 | -0.82074 | 4.98012 | -3.79482 | 0.000381 | 0.014854 | -0.25522 | Histone H3 | elongation zone | 6 h |
| HORVU.MOREX.r3.5HG0529500.1 | -0.31885 | 7.05237 | -3.87496 | 0.000295 | 0.012756 | -0.25544 | Protein kinase | elongation zone | 6 h |
| HORVU.MOREX.r3.3HG0276120.1 | 1.93491 | -1.33075 | 3.685006 | 0.000538 | 0.017871 | -0.25849 | Phospholipase A1 | elongation zone | 6 h |
| HORVU.MOREX.r3.5HG0434380.1 | -0.48255 | 4.006199 | -3.75519 | 0.000432 | 0.015665 | -0.25871 | Receptor-like kinase | elongation zone | 6 h |
| HORVU.MOREX.r3.3HG0287690.1 | 0.906034 | 2.646815 | 3.846112 | 0.000324 | 0.013509 | -0.25885 | Profilin | elongation zone | 6 h |
| HORVU.MOREX.r3.7HG0668560.1 | 0.976069 | 0.264899 | 3.684469 | 0.000539 | 0.017872 | -0.26132 | Pyruvate dehydrogenase E1 component subunit alpha | elongation zone | 6 h |
| HORVU.MOREX.r3.3HG0293130.1 | 1.061527 | 3.772343 | 3.782887 | 0.000396 | 0.015004 | -0.26216 | Eukaryotic aspartyl protease family protein | elongation zone | 6 h |
| HORVU.MOREX.r3.6HG0616450.1 | 1.387171 | 0.245809 | 3.737615 | 0.000456 | 0.016067 | -0.26267 | Na(+)/H(+) exchange regulatory cofactor NHE-RF3 | elongation zone | 6 h |
| HORVU.MOREX.r3.5HG0477180.1 | -0.38979 | 5.597571 | -3.82679 | 0.000344 | 0.01394 | -0.26532 | LAG1 longevity assurance-like protein | elongation zone | 6 h |
| HORVU.MOREX.r3.5HG0485220.1 | -0.50725 | 5.159201 | -3.79229 | 0.000384 | 0.014876 | -0.26698 | Gamma-tubulin complex component | elongation zone | 6 h |

**Table S1** Continued.

| ID | logFC | AveExpr | t | P.Value | adj.P.Val | B | gene description | root zone | time point |
|---|---|---|---|---|---|---|---|---|---|
| HORVU.MOREX.r3.7HG0712000.1 | 1.999225 | -3.03639 | 3.732828 | 0.000463 | 0.016199 | -0.27386 | Transposon protein | elongation zone | 6 h |
| HORVU.MOREX.r3.1HG0038800.1 | 0.634516 | 2.879103 | 3.758877 | 0.000427 | 0.015589 | -0.27694 | Protein kinase | elongation zone | 6 h |
| HORVU.MOREX.r3.2HG0204480.1 | -0.63084 | -0.53341 | -3.73371 | 0.000462 | 0.016195 | -0.28093 | Cysteine protease | elongation zone | 6 h |
| HORVU.MOREX.r3.5HG0508880.1 | 1.71581 | -0.73312 | 3.671846 | 0.000561 | 0.018236 | -0.28234 | Leucine-rich repeat protein kinase family protein | elongation zone | 6 h |
| HORVU.MOREX.r3.2HG0211730.1 | -0.50183 | 4.52794 | -3.79723 | 0.000378 | 0.014813 | -0.28439 | Phosphoribosylformylglycinamidine synthase subunit PurQ | elongation zone | 6 h |
| HORVU.MOREX.r3.5HG0420820.1 | 1.512927 | -2.28387 | 3.670049 | 0.000564 | 0.01828 | -0.28606 | TIFY domain/Divergent CCT motif family protein | elongation zone | 6 h |
| HORVU.MOREX.r3.2HG0182990.1 | -0.79277 | 2.816822 | -3.74692 | 0.000443 | 0.015878 | -0.29387 | UDP-glycosyltransferase | elongation zone | 6 h |
| HORVU.MOREX.r3.1HG0026080.1 | 0.669772 | 4.010774 | 3.75311 | 0.000435 | 0.015735 | -0.29681 | RING finger protein | elongation zone | 6 h |
| HORVU.MOREX.r3.7HG0649950.1 | 0.604118 | 3.836695 | 3.775402 | 0.000405 | 0.015121 | -0.29914 | Nudix hydrolase | elongation zone | 6 h |
| HORVU.MOREX.r3.4HG0417660.1 | 2.790655 | -3.39591 | 3.832708 | 0.000338 | 0.013839 | -0.30419 | rRNA N-glycosidase | elongation zone | 6 h |
| HORVU.MOREX.r3.1HG0090710.1 | -0.39678 | 4.396711 | -3.78302 | 0.000396 | 0.015004 | -0.3071 | transmembrane protein | elongation zone | 6 h |
| HORVU.MOREX.r3.7HG0714660.1 | 1.511632 | -2.48817 | 3.67658 | 0.000553 | 0.018074 | -0.31137 | Retrovirus-related Pol polyprotein from transposon TNT 1-94 | elongation zone | 6 h |
| HORVU.MOREX.r3.1HG0069030.1 | 1.620953 | 1.48038 | 3.682516 | 0.000542 | 0.017926 | -0.31488 | Calcium-transporting ATPase | elongation zone | 6 h |
| HORVU.MOREX.r3.3HG0298790.1 | 0.828883 | 4.93163 | 3.688708 | 0.000532 | 0.01781 | -0.31866 | Amidophosphoribosyltransferase | elongation zone | 6 h |
| HORVU.MOREX.r3.1HG0002570.1 | 1.329815 | -1.7761 | 3.679139 | 0.000548 | 0.017997 | -0.32178 | GDSL esterase/lipase | elongation zone | 6 h |
| HORVU.MOREX.r3.4HG0384230.1 | -0.57981 | 3.949266 | -3.73346 | 0.000462 | 0.016195 | -0.32965 | RING/U-box superfamily protein | elongation zone | 6 h |
| HORVU.MOREX.r3.2HG0126960.1 | -0.56668 | 4.780474 | -3.80139 | 0.000373 | 0.014703 | -0.3311 | Longifolia protein | elongation zone | 6 h |
| HORVU.MOREX.r3.7HG0671410.1 | 1.292059 | -1.71062 | 3.652605 | 0.000595 | 0.01894 | -0.3333 | Glycosyltransferase | elongation zone | 6 h |
| HORVU.MOREX.r3.2HG0128490.1 | -0.34746 | 4.434925 | -3.77366 | 0.000408 | 0.015165 | -0.33667 | #N/A | elongation zone | 6 h |
| HORVU.MOREX.r3.1HG0071420.1 | 0.917509 | 3.587411 | 3.69943 | 0.000515 | 0.017538 | -0.33871 | TOM1-like protein 2 | elongation zone | 6 h |
| HORVU.MOREX.r3.1HG0001010.1 | -0.79498 | 4.233711 | -3.72648 | 0.000473 | 0.016469 | -0.33888 | Pm3-like disease resistance protein | elongation zone | 6 h |
| HORVU.MOREX.r3.5HG0488200.1 | 1.124769 | 0.584662 | 3.65932 | 0.000583 | 0.018648 | -0.34028 | Peroxidase | elongation zone | 6 h |
| HORVU.MOREX.r3.5HG0483860.1 | -0.54771 | 3.316157 | -3.68124 | 0.000545 | 0.017937 | -0.34447 | RING/U-box superfamily protein | elongation zone | 6 h |
| HORVU.MOREX.r3.5HG0533230.1 | -0.52821 | 5.643177 | -3.80229 | 0.000372 | 0.01469 | -0.34779 | Pirin-like protein | elongation zone | 6 h |
| HORVU.MOREX.r3.7HG0664720.1 | 1.172021 | 0.421259 | 3.809012 | 0.000364 | 0.014439 | -0.35336 | Blue copper protein | elongation zone | 6 h |
| HORVU.MOREX.r3.1HG0067980.1 | 1.480567 | -1.60301 | 3.686909 | 0.000535 | 0.017814 | -0.36761 | Peroxidase | elongation zone | 6 h |
| HORVU.MOREX.r3.3HG0246550.1 | -0.79275 | 2.643364 | -3.77987 | 0.0004 | 0.015004 | -0.37021 | Peroxidase | elongation zone | 6 h |
| HORVU.MOREX.r3.2HG0184230.1 | 2.484088 | -3.03591 | 3.686687 | 0.000535 | 0.017814 | -0.37139 | #N/A | elongation zone | 6 h |
| HORVU.MOREX.r3.5HG0477220.1 | 1.014635 | 1.858254 | 3.668133 | 0.000567 | 0.018303 | -0.37942 | F-box protein | elongation zone | 6 h |
| HORVU.MOREX.r3.4HG0351290.1 | 1.050914 | 5.034448 | 3.806721 | 0.000367 | 0.014513 | -0.39111 | B3 domain-containing protein | elongation zone | 6 h |
| HORVU.MOREX.r3.2HG0119080.1 | 0.820267 | 0.931759 | 3.67621 | 0.000553 | 0.018074 | -0.39953 | Endonuclease/exonuclease/phosphatase family protein | elongation zone | 6 h |
| HORVU.MOREX.r3.1HG0092860.1 | 1.378859 | -1.39568 | 3.671 | 0.000562 | 0.018255 | -0.40427 | S-type anion channel | elongation zone | 6 h |
| HORVU.MOREX.r3.5HG0461280.1 | 0.518097 | 6.01476 | 3.788538 | 0.000389 | 0.014941 | -0.41204 | Calcium-binding EF-hand protein | elongation zone | 6 h |
| HORVU.MOREX.r3.6HG0613430.1 | 1.926202 | -1.24134 | 3.623302 | 0.000652 | 0.020088 | -0.41209 | RNA-dependent RNA polymerase | elongation zone | 6 h |
| HORVU.MOREX.r3.2HG0129450.1 | 0.962481 | 1.029346 | 3.760547 | 0.000425 | 0.015563 | -0.41329 | basic helix-loop-helix (bHLH) DNA-binding superfamily protein | elongation zone | 6 h |
| HORVU.MOREX.r3.7HG0742720.1 | -0.35549 | 4.763908 | -3.74818 | 0.000442 | 0.015869 | -0.41389 | Magnesium transporter MRS2-like protein | elongation zone | 6 h |
| HORVU.MOREX.r3.1HG0035400.1 | 0.540672 | 4.194434 | 3.759679 | 0.000426 | 0.015578 | -0.41987 | Ras-related protein, expressed | elongation zone | 6 h |
| HORVU.MOREX.r3.5HG0460630.1 | -0.98074 | 1.842017 | -3.63182 | 0.000635 | 0.019743 | -0.42185 | DTW domain containing protein | elongation zone | 6 h |
| HORVU.MOREX.r3.5HG0493330.1 | 1.085278 | 0.977391 | 3.711273 | 0.000496 | 0.017032 | -0.42506 | UDP-glucuronate 4-epimerase 4 | elongation zone | 6 h |
| HORVU.MOREX.r3.1HG0006660.1 | 0.953973 | 5.455877 | 3.744067 | 0.000447 | 0.015908 | -0.43968 | Inorganic pyrophosphatase family protein | elongation zone | 6 h |
| HORVU.MOREX.r3.1HG0015750.1 | 1.767899 | -2.91215 | 3.614636 | 0.00067 | 0.020541 | -0.45028 | transcription factor, putative (Protein of unknown function, DUF547) | elongation zone | 6 h |

**Table S1** Continued.

| ID | logFC | AveExpr | t | P.Value | adj.P.Val | B | gene description | root zone | time point |
|---|---|---|---|---|---|---|---|---|---|
| HORVU.MOREX.r3.3HG0323750.1 | 0.916829 | 2.450618 | 3.625955 | 0.000647 | 0.020044 | -0.4527 | bZIP transcription factor, putative (DUF630 and DUF632) | elongation zone | 6 h |
| HORVU.MOREX.r3.5HG0430050.1 | 0.638041 | 4.495639 | 3.723105 | 0.000478 | 0.01656 | -0.45876 | Stomatal closure-related actin-binding protein 1 | elongation zone | 6 h |
| HORVU.MOREX.r3.1HG0087620.1 | 1.00565 | 3.06372 | 3.795221 | 0.000381 | 0.014854 | -0.4615 | Xyloglucan endotransglucosylase/hydrolase | elongation zone | 6 h |
| HORVU.MOREX.r3.5HG0422080.1 | 0.422951 | 5.370925 | 3.768683 | 0.000414 | 0.015333 | -0.46151 | RING/U-box superfamily protein | elongation zone | 6 h |
| HORVU.MOREX.r3.2HG0203080.1 | 2.09661 | 0.039803 | 3.608613 | 0.000682 | 0.020703 | -0.46359 | NA | elongation zone | 6 h |
| HORVU.MOREX.r3.4HG0382280.1 | 1.208932 | 2.916703 | 3.60375 | 0.000693 | 0.020828 | -0.46421 | Calcium-dependent phosphotriesterase superfamily protein | elongation zone | 6 h |
| HORVU.MOREX.r3.7HG0676570.1 | -0.67091 | 6.054358 | -3.75108 | 0.000438 | 0.01578 | -0.46437 | Deaminase-related family protein | elongation zone | 6 h |
| HORVU.MOREX.r3.1HG0088530.1 | 0.959256 | 0.052643 | 3.611159 | 0.000677 | 0.020693 | -0.46732 | Cytochrome b561 domain-containing protein | elongation zone | 6 h |
| HORVU.MOREX.r3.6HG0553580.1 | 1.274208 | -1.47778 | 3.633182 | 0.000632 | 0.019703 | -0.47278 | Methionine--tRNA ligase | elongation zone | 6 h |
| HORVU.MOREX.r3.7HG0638340.1 | 1.291694 | 2.325425 | 3.606821 | 0.000686 | 0.020703 | -0.47566 | S-adenosyl-L-methionine-dependent methyltransferases superfamily protein | elongation zone | 6 h |
| HORVU.MOREX.r3.5HG0423790.1 | -0.54515 | 1.914718 | -3.66221 | 0.000578 | 0.018556 | -0.48154 | Casein kinase II subunit beta | elongation zone | 6 h |
| HORVU.MOREX.r3.5HG0490800.1 | 1.247526 | 0.125384 | 3.601607 | 0.000697 | 0.020915 | -0.48602 | IRK-interacting protein | elongation zone | 6 h |
| HORVU.MOREX.r3.4HG0369880.1 | -1.07041 | 2.555153 | -3.60982 | 0.00068 | 0.020694 | -0.49772 | Flavin-binding kelch domain F box protein | elongation zone | 6 h |
| HORVU.MOREX.r3.7HG0728000.1 | 0.629734 | 5.007313 | 3.779383 | 0.0004 | 0.015004 | -0.49935 | Glycosyltransferase | elongation zone | 6 h |
| HORVU.MOREX.r3.2HG0174750.1 | 1.205453 | 1.498345 | 3.764811 | 0.000419 | 0.0154 | -0.50093 | Fasciclin-like arabinogalactan protein | elongation zone | 6 h |
| HORVU.MOREX.r3.1HG0075340.1 | 0.974552 | 1.815297 | 3.645735 | 0.000608 | 0.0192 | -0.50159 | Alpha/beta-Hydrolases superfamily protein | elongation zone | 6 h |
| HORVU.MOREX.r3.3HG0219380.1 | 1.046041 | 4.367645 | 3.799171 | 0.000376 | 0.01475 | -0.50261 | Glutamate receptor ionotropic, NMDA 3A | elongation zone | 6 h |
| HORVU.MOREX.r3.5HG0447610.1 | 0.942992 | 3.765571 | 3.611691 | 0.000676 | 0.020693 | -0.50509 | #N/A | elongation zone | 6 h |
| HORVU.MOREX.r3.6HG0557920.1 | 0.923139 | 4.693871 | 3.713366 | 0.000493 | 0.017016 | -0.50783 | Auxin response factor | elongation zone | 6 h |
| HORVU.MOREX.r3.3HG0310390.1 | 1.633224 | 0.671699 | 3.646103 | 0.000607 | 0.0192 | -0.51643 | Transcription and mRNA export factor SUS1 | elongation zone | 6 h |
| HORVU.MOREX.r3.7HG0748070.1 | 1.352132 | -0.49696 | 3.672831 | 0.000559 | 0.018209 | -0.51803 | Heavy metal transport/detoxification superfamily protein, putative | elongation zone | 6 h |
| HORVU.MOREX.r3.3HG0218100.1 | -0.33079 | 6.103028 | -3.74992 | 0.000439 | 0.01581 | -0.51969 | Chaperone protein DnaJ | elongation zone | 6 h |
| HORVU.MOREX.r3.7HG0746770.1 | -0.49877 | 10.27031 | -3.90228 | 0.000271 | 0.012154 | -0.5232 | Chaperone protein htpG family protein | elongation zone | 6 h |
| HORVU.MOREX.r3.1HG0041280.1 | 0.864441 | 4.323201 | 3.784782 | 0.000393 | 0.015004 | -0.52324 | Cellulose synthase | elongation zone | 6 h |
| HORVU.MOREX.r3.2HG0140970.1 | -0.80443 | 2.378717 | -3.6533 | 0.000594 | 0.018929 | -0.52666 | Heat-shock protein, putative | elongation zone | 6 h |
| HORVU.MOREX.r3.4HG0407230.1 | 1.421274 | -0.9449 | 3.583553 | 0.000737 | 0.021545 | -0.52796 | Flowering-promoting factor 1-like protein 1 | elongation zone | 6 h |
| HORVU.MOREX.r3.1HG0015960.1 | 1.247268 | -0.2749 | 3.591304 | 0.00072 | 0.021192 | -0.52838 | Protein kinase family protein | elongation zone | 6 h |
| HORVU.MOREX.r3.7HG0669590.1 | 1.463324 | -0.99729 | 3.624475 | 0.00065 | 0.020075 | -0.53524 | Caleosin | elongation zone | 6 h |
| HORVU.MOREX.r3.3HG0289020.1 | 1.176905 | 2.050529 | 3.703174 | 0.000509 | 0.017421 | -0.53571 | Receptor-like kinase | elongation zone | 6 h |
| HORVU.MOREX.r3.3HG0307850.1 | -0.69387 | 6.891759 | -3.81341 | 0.000359 | 0.014346 | -0.53617 | Basic 7S globulin | elongation zone | 6 h |
| HORVU.MOREX.r3.7HG0642940.1 | 1.901718 | -1.68507 | 3.609264 | 0.000681 | 0.020699 | -0.541 | Tryptophan decarboxylase | elongation zone | 6 h |
| HORVU.MOREX.r3.5HG0485800.1 | -0.4179 | 8.255741 | -3.82919 | 0.000342 | 0.013861 | -0.54375 | Protein disulfide isomerase | elongation zone | 6 h |
| HORVU.MOREX.r3.1HG0089220.1 | -0.84934 | 0.50855 | -3.58229 | 0.00074 | 0.021553 | -0.54644 | Cell surface antigen I/II | elongation zone | 6 h |
| HORVU.MOREX.r3.1HG0055500.1 | 1.205796 | 2.091951 | 3.773311 | 0.000408 | 0.015165 | -0.54674 | Pollen Ole e 1 allergen and extensin family protein | elongation zone | 6 h |
| HORVU.MOREX.r3.7HG0661480.1 | -0.29565 | 7.365152 | -3.79455 | 0.000381 | 0.014854 | -0.55229 | Alpha-amylase 2 | elongation zone | 6 h |
| HORVU.MOREX.r3.6HG0619500.1 | 1.446488 | -0.34034 | 3.593252 | 0.000715 | 0.021158 | -0.55699 | #N/A | elongation zone | 6 h |
| HORVU.MOREX.r3.6HG0545620.1 | -0.37959 | 11.02319 | -3.92861 | 0.000249 | 0.011496 | -0.55711 | Heat shock 70 kDa protein | elongation zone | 6 h |
| HORVU.MOREX.r3.6HG0610500.1 | 0.95895 | 1.298618 | 3.767986 | 0.000415 | 0.015336 | -0.57013 | Eukaryotic aspartyl protease family protein | elongation zone | 6 h |
| HORVU.MOREX.r3.2HG0120490.1 | 1.315125 | 1.876399 | 3.76578 | 0.000418 | 0.015391 | -0.57648 | Peroxidase | elongation zone | 6 h |
| HORVU.MOREX.r3.1HG0039370.1 | -0.40294 | 4.945966 | -3.68247 | 0.000542 | 0.017926 | -0.57682 | Leucine-rich repeat (LRR) family protein | elongation zone | 6 h |
| HORVU.MOREX.r3.5HG0512250.1 | 1.49733 | -0.16991 | 3.689718 | 0.00053 | 0.01781 | -0.57713 | Receptor-like kinase, putative | elongation zone | 6 h |

**Table S1** Continued.

| ID | logFC | AveExpr | t | P.Value | adj.P.Val | B | gene description | root zone | time point |
|---|---|---|---|---|---|---|---|---|---|
| HORVU.MOREX.r3.4HG0391330.1 | -0.4174 | 6.457456 | -3.74568 | 0.000445 | 0.015908 | -0.58242 | Phosphomevalonate kinase | elongation zone | 6 h |
| HORVU.MOREX.r3.4HG0392230.1 | 1.071412 | 0.189187 | 3.696101 | 0.00052 | 0.017591 | -0.58442 | RNA-binding family protein, putative | elongation zone | 6 h |
| HORVU.MOREX.r3.6HG0549990.1 | -0.91081 | 2.338996 | -3.56085 | 0.00079 | 0.022277 | -0.58738 | Pentatricopeptide repeat-containing protein | elongation zone | 6 h |
| HORVU.MOREX.r3.6HG0601260.1 | 1.042577 | 3.050364 | 3.725452 | 0.000474 | 0.016494 | -0.59399 | Sulfotransferase | elongation zone | 6 h |
| HORVU.MOREX.r3.4HG0353390.1 | 0.806003 | 1.790635 | 3.637572 | 0.000624 | 0.019542 | -0.59562 | Type I inositol-1,4,5-trisphosphate 5-phosphatase | elongation zone | 6 h |
| HORVU.MOREX.r3.7HG0750170.1 | 1.280081 | 1.590474 | 3.575925 | 0.000755 | 0.021777 | -0.59792 | #N/A | elongation zone | 6 h |
| HORVU.MOREX.r3.5HG0485790.1 | 0.767141 | 1.583435 | 3.69823 | 0.000516 | 0.017564 | -0.60043 | 1-aminocyclopropane-1-carboxylate oxidase | elongation zone | 6 h |
| HORVU.MOREX.r3.4HG0342480.1 | 0.99918 | 4.010766 | 3.702286 | 0.00051 | 0.017427 | -0.60472 | Subtilisin-like protease | elongation zone | 6 h |
| HORVU.MOREX.r3.4HG0345840.1 | 1.474028 | 1.556535 | 3.551083 | 0.000814 | 0.022588 | -0.60809 | Receptor lectin kinase | elongation zone | 6 h |
| HORVU.MOREX.r3.3HG0291040.1 | 0.761351 | 3.480757 | 3.607446 | 0.000685 | 0.020703 | -0.60849 | Glutathione S-transferase | elongation zone | 6 h |
| HORVU.MOREX.r3.3HG0231000.1 | 0.97974 | 3.745714 | 3.723371 | 0.000477 | 0.01656 | -0.60856 | Gamma-glutamyltranspeptidase | elongation zone | 6 h |
| HORVU.MOREX.r3.7HG0680990.1 | -0.3482 | 5.931399 | -3.73878 | 0.000455 | 0.016064 | -0.61316 | Laminin-like protein epi-1 | elongation zone | 6 h |
| HORVU.MOREX.r3.4HG0364050.1 | 0.755507 | 2.580176 | 3.591534 | 0.000719 | 0.021192 | -0.61774 | Sodium/hydrogen exchanger | elongation zone | 6 h |
| HORVU.MOREX.r3.6HG0620200.1 | 1.402455 | -0.686 | 3.560933 | 0.00079 | 0.022277 | -0.61987 | Kinase family protein | elongation zone | 6 h |
| HORVU.MOREX.r3.1HG0037910.1 | -0.70404 | 4.060046 | -3.59945 | 0.000702 | 0.020963 | -0.62639 | Starch synthase, chloroplastic/amyloplastic | elongation zone | 6 h |
| HORVU.MOREX.r3.4HG0338290.1 | 0.836673 | 1.18244 | 3.564574 | 0.000781 | 0.02217 | -0.62903 | Glucan endo-1,3-beta-glucosidase | elongation zone | 6 h |
| HORVU.MOREX.r3.2HG0124550.1 | 1.388435 | 2.362034 | 3.544183 | 0.000832 | 0.022817 | -0.63085 | Protein WEAK CHLOROPLAST MOVEMENT UNDER BLUE LIGHT 1 | elongation zone | 6 h |
| HORVU.MOREX.r3.2HG0158260.1 | -0.99133 | 2.424618 | -3.54054 | 0.000841 | 0.02292 | -0.63896 | Glycogen synthase | elongation zone | 6 h |
| HORVU.MOREX.r3.5HG0477040.1 | -0.40892 | 5.55814 | -3.68124 | 0.000545 | 0.017937 | -0.64307 | MND1-interacting protein 1 | elongation zone | 6 h |
| HORVU.MOREX.r3.6HG0582480.1 | 0.74213 | 2.53548 | 3.577 | 0.000752 | 0.021767 | -0.66403 | Ethylene-responsive transcription factor-like protein | elongation zone | 6 h |
| HORVU.MOREX.r3.3HG0253860.1 | 2.235977 | -0.09276 | 3.542259 | 0.000837 | 0.02283 | -0.66418 | Rapid alkalinization factor (RALF) family protein | elongation zone | 6 h |
| HORVU.MOREX.r3.3HG0314070.1 | 1.386924 | -1.61281 | 3.534046 | 0.000858 | 0.023177 | -0.66733 | Protein kinase family protein | elongation zone | 6 h |
| HORVU.MOREX.r3.3HG0235030.1 | -0.51603 | 3.06592 | -3.56421 | 0.000782 | 0.02217 | -0.66983 | Dehydration-responsive element binding protein | elongation zone | 6 h |
| HORVU.MOREX.r3.6HG0620720.1 | 0.925278 | 0.656099 | 3.6936 | 0.000524 | 0.017655 | -0.67209 | Ribonuclease | elongation zone | 6 h |
| HORVU.MOREX.r3.2HG0197540.1 | 1.248544 | 3.09996 | 3.570153 | 0.000768 | 0.022074 | -0.67247 | MYB protein | elongation zone | 6 h |
| HORVU.MOREX.r3.2HG0170210.1 | 1.703154 | 2.50277 | 3.530589 | 0.000867 | 0.023177 | -0.67729 | Pentatricopeptide repeat-containing protein | elongation zone | 6 h |
| HORVU.MOREX.r3.1HG0059340.1 | -0.34212 | 6.875668 | -3.74414 | 0.000447 | 0.015908 | -0.67983 | Conserved oligomeric Golgi complex subunit 5 | elongation zone | 6 h |
| HORVU.MOREX.r3.5HG0513980.1 | 1.257644 | -0.36473 | 3.548567 | 0.000821 | 0.022653 | -0.68289 | Auxin-responsive protein | elongation zone | 6 h |
| HORVU.MOREX.r3.3HG0269130.1 | 1.229899 | 4.115188 | 3.78538 | 0.000393 | 0.015004 | -0.68398 | Leucine-rich repeat protein kinase family protein | elongation zone | 6 h |
| HORVU.MOREX.r3.1HG0024040.1 | -0.35973 | 9.734968 | -3.83314 | 0.000338 | 0.013839 | -0.68892 | Disulfide-isomerase | elongation zone | 6 h |
| HORVU.MOREX.r3.2HG0117830.1 | 1.828193 | -2.70524 | 3.521576 | 0.000891 | 0.023455 | -0.69073 | Protein trichome birefringence | elongation zone | 6 h |
| HORVU.MOREX.r3.3HG0245120.1 | 3.008632 | -2.69428 | 3.579122 | 0.000747 | 0.021656 | -0.69076 | Cysteine protease, putative | elongation zone | 6 h |
| HORVU.MOREX.r3.7HG0634430.1 | -0.82024 | 3.249235 | -3.54489 | 0.00083 | 0.022798 | -0.69287 | NBS-LRR-like resistance protein | elongation zone | 6 h |
| HORVU.MOREX.r3.5HG0517190.1 | 0.612034 | 2.990144 | 3.562931 | 0.000785 | 0.022226 | -0.69664 | Calcium-dependent protein kinase | elongation zone | 6 h |
| HORVU.MOREX.r3.4HG0405990.1 | 1.490237 | 0.091283 | 3.530995 | 0.000866 | 0.023177 | -0.69696 | Transmembrane protein, putative | elongation zone | 6 h |
| HORVU.MOREX.r3.3HG0288700.1 | -0.59049 | 3.838111 | -3.59364 | 0.000715 | 0.021158 | -0.69744 | #N/A | elongation zone | 6 h |
| HORVU.MOREX.r3.3HG0242150.1 | 2.816029 | -3.21832 | 3.594624 | 0.000712 | 0.021154 | -0.69914 | Glycine-rich cell wall structural protein | elongation zone | 6 h |
| HORVU.MOREX.r3.3HG0293490.1 | 1.09879 | -0.53962 | 3.630886 | 0.000637 | 0.01977 | -0.70004 | Expansin | elongation zone | 6 h |
| HORVU.MOREX.r3.4HG0383530.1 | 1.160899 | -0.0177 | 3.66083 | 0.00058 | 0.018607 | -0.70623 | Pathogenesis-related protein 1 | elongation zone | 6 h |
| HORVU.MOREX.r3.5HG0465810.1 | 2.584797 | -0.29224 | 3.552432 | 0.000811 | 0.022568 | -0.70879 | Calcium-binding protein | elongation zone | 6 h |
| HORVU.MOREX.r3.3HG0265140.1 | 2.429957 | -1.23714 | 3.568326 | 0.000772 | 0.022075 | -0.71205 | DUF3511 domain protein | elongation zone | 6 h |

**Table S1** Continued.

| ID | logFC | AveExpr | t | P.Value | adj.P.Val | B | gene description | root zone | time point |
|---|---|---|---|---|---|---|---|---|---|
| HORVU.MOREX.r3.3HG0241000.1 | -1.14573 | 6.471225 | -3.66746 | 0.000568 | 0.018312 | -0.71256 | HXXXD-type acyl-transferase-like protein | elongation zone | 6 h |
| HORVU.MOREX.r3.6HG0632270.1 | -0.51155 | 3.993705 | -3.59315 | 0.000716 | 0.021158 | -0.72051 | Leucine-rich repeat family protein | elongation zone | 6 h |
| HORVU.MOREX.r3.7HG0636630.1 | 1.069984 | -0.12855 | 3.596159 | 0.000709 | 0.021085 | -0.72197 | WAT1-related protein | elongation zone | 6 h |
| HORVU.MOREX.r3.4HG0391440.1 | 2.444945 | 2.5321 | 3.566854 | 0.000776 | 0.022144 | -0.7237 | Transcription initiation factor IIF subunit alpha | elongation zone | 6 h |
| HORVU.MOREX.r3.3HG0328430.1 | 1.715402 | -0.79684 | 3.504925 | 0.000938 | 0.024245 | -0.728 | Zinc transporter, putative | elongation zone | 6 h |
| HORVU.MOREX.r3.4HG0354010.1 | 2.103758 | -0.99752 | 3.50354 | 0.000942 | 0.024273 | -0.72936 | LURP-one-like protein | elongation zone | 6 h |
| HORVU.MOREX.r3.7HG0684880.1 | 0.720825 | 3.761267 | 3.610797 | 0.000678 | 0.020693 | -0.72937 | meiosis chromosome segregation family protein | elongation zone | 6 h |
| HORVU.MOREX.r3.5HG0520650.1 | -0.68725 | 5.200623 | -3.6436 | 0.000612 | 0.019279 | -0.73148 | Argonaute family protein | elongation zone | 6 h |
| HORVU.MOREX.r3.3HG0313020.1 | -0.35711 | 5.21659 | -3.65089 | 0.000599 | 0.018997 | -0.73458 | Alcohol dehydrogenase, putative | elongation zone | 6 h |
| HORVU.MOREX.r3.2HG0184620.1 | -1.00724 | 4.117573 | -3.56447 | 0.000782 | 0.02217 | -0.73656 | Hsp70-Hsp90 organizing protein 1 | elongation zone | 6 h |
| HORVU.MOREX.r3.5HG0532560.1 | 1.841864 | -0.15138 | 3.5015 | 0.000947 | 0.024363 | -0.73656 | Ras-related protein, expressed | elongation zone | 6 h |
| HORVU.MOREX.r3.7HG0670690.1 | 0.617126 | 8.088776 | 3.819768 | 0.000352 | 0.01417 | -0.7401 | HIPL1 protein | elongation zone | 6 h |
| HORVU.MOREX.r3.7HG0721320.1 | 1.386436 | -3.12942 | 3.518006 | 0.000901 | 0.023642 | -0.74814 | Auxin-responsive family protein | elongation zone | 6 h |
| HORVU.MOREX.r3.1HG0006400.1 | 0.703199 | 6.387241 | 3.757925 | 0.000428 | 0.015608 | -0.75216 | Beta-1,3-galactosyltransferase-like protein | elongation zone | 6 h |
| HORVU.MOREX.r3.2HG0212570.1 | -0.40516 | 10.53927 | -3.83098 | 0.00034 | 0.013839 | -0.76313 | Calreticulin | elongation zone | 6 h |
| HORVU.MOREX.r3.3HG0288960.1 | 1.047837 | 4.129323 | 3.655136 | 0.000591 | 0.018851 | -0.76325 | Cellulose synthase | elongation zone | 6 h |
| HORVU.MOREX.r3.7HG0725290.1 | -0.60065 | 3.61352 | -3.55994 | 0.000793 | 0.022309 | -0.76686 | YGL010w-like protein | elongation zone | 6 h |
| HORVU.MOREX.r3.2HG0124450.1 | -0.54969 | 4.909528 | -3.64212 | 0.000615 | 0.019328 | -0.76896 | calmodulin-binding protein (DUF1645) | elongation zone | 6 h |
| HORVU.MOREX.r3.6HG0621190.1 | -0.67317 | 4.746594 | -3.63638 | 0.000626 | 0.019584 | -0.76978 | Chaperone protein dnaJ | elongation zone | 6 h |
| HORVU.MOREX.r3.6HG0609830.1 | 0.771268 | 4.638207 | 3.737827 | 0.000456 | 0.016067 | -0.77069 | Pollen-specific arabinogalacta protein BAN102 | elongation zone | 6 h |
| HORVU.MOREX.r3.6HG0606530.1 | 0.657707 | 4.864638 | 3.677174 | 0.000552 | 0.018074 | -0.77447 | Myb family transcription factor APL | elongation zone | 6 h |
| HORVU.MOREX.r3.2HG0096800.1 | 0.75183 | 4.6918 | 3.625038 | 0.000648 | 0.020071 | -0.77531 | Flavin-containing monooxygenase | elongation zone | 6 h |
| HORVU.MOREX.r3.6HG0542640.1 | 1.2642 | 2.086759 | 3.581257 | 0.000742 | 0.021565 | -0.77778 | Plant/F1M20-13 protein | elongation zone | 6 h |
| HORVU.MOREX.r3.2HG0192270.1 | 2.365035 | -2.74995 | 3.523578 | 0.000886 | 0.023423 | -0.78273 | Sodium transporter | elongation zone | 6 h |
| HORVU.MOREX.r3.6HG0616880.1 | -1.49604 | -0.67725 | -3.48233 | 0.001004 | 0.025247 | -0.78438 | Upf0503 protein, chloroplastic | elongation zone | 6 h |
| HORVU.MOREX.r3.1HG0025320.1 | 1.216938 | 0.572545 | 3.487242 | 0.000989 | 0.025053 | -0.78755 | Receptor-like kinase, putative | elongation zone | 6 h |
| HORVU.MOREX.r3.2HG0194620.1 | 2.066231 | -2.14798 | 3.512418 | 0.000916 | 0.023957 | -0.79004 | F-box protein | elongation zone | 6 h |
| HORVU.MOREX.r3.7HG0644220.1 | 0.581232 | 8.36113 | 3.778945 | 0.000401 | 0.015004 | -0.79369 | Arginine decarboxylase | elongation zone | 6 h |
| HORVU.MOREX.r3.1HG0087590.1 | 1.456679 | -1.00814 | 3.493205 | 0.000972 | 0.024706 | -0.79414 | transmembrane protein, putative (DUF679 domain membrane protein 2) | elongation zone | 6 h |
| HORVU.MOREX.r3.5HG0486960.1 | -1.50136 | 2.507872 | -3.47795 | 0.001018 | 0.025374 | -0.79567 | Alpha/beta-Hydrolases superfamily protein, putative | elongation zone | 6 h |
| HORVU.MOREX.r3.1HG0002630.1 | 1.989681 | 4.599685 | 3.477997 | 0.001018 | 0.025374 | -0.79615 | Formin-like protein | elongation zone | 6 h |
| HORVU.MOREX.r3.7HG0707220.1 | 0.614353 | 4.707516 | 3.675721 | 0.000554 | 0.018074 | -0.79734 | #N/A | elongation zone | 6 h |
| HORVU.MOREX.r3.4HG0400530.1 | 1.3693 | 0.815449 | 3.475798 | 0.001024 | 0.025509 | -0.80126 | #N/A | elongation zone | 6 h |
| HORVU.MOREX.r3.6HG0542320.1 | 1.273934 | 2.291237 | 3.474933 | 0.001027 | 0.025525 | -0.80567 | Kinase family protein | elongation zone | 6 h |
| HORVU.MOREX.r3.3HG0295640.1 | 1.319709 | -1.27257 | 3.533983 | 0.000858 | 0.023177 | -0.80582 | Protein MIZU-KUSSEI 1 | elongation zone | 6 h |
| HORVU.MOREX.r3.6HG0625570.1 | 0.571664 | 5.941941 | 3.701985 | 0.00051 | 0.017427 | -0.81157 | Transmembrane protein | elongation zone | 6 h |
| HORVU.MOREX.r3.1HG0055380.1 | -0.47664 | 6.771434 | -3.67949 | 0.000548 | 0.017997 | -0.81415 | Methylsterol monooxygenase 1-2 | elongation zone | 6 h |
| HORVU.MOREX.r3.2HG0216470.1 | -0.58313 | 6.666412 | -3.68778 | 0.000534 | 0.017814 | -0.81435 | Kinase family protein | elongation zone | 6 h |
| HORVU.MOREX.r3.6HG0570520.1 | -0.25122 | 6.220455 | -3.68654 | 0.000536 | 0.017814 | -0.81499 | Core-2/I-branching beta-1,6-N-acetylglucosaminyltransferase family protein | elongation zone | 6 h |
| HORVU.MOREX.r3.3HG0225880.1 | 0.908065 | 4.380821 | 3.545766 | 0.000828 | 0.022768 | -0.8166 | Receptor-like kinase | elongation zone | 6 h |
| HORVU.MOREX.r3.7HG0709270.1 | 0.887614 | 3.012432 | 3.588321 | 0.000726 | 0.021292 | -0.82139 | RING/U-box superfamily protein | elongation zone | 6 h |

**Table S1** Continued.

| ID | logFC | AveExpr | t | P.Value | adj.P.Val | B | gene description | root zone | time point |
|---|---|---|---|---|---|---|---|---|---|
| HORVU.MOREX.r3.6HG0589860.1 | 0.795652 | 4.121931 | 3.584453 | 0.000735 | 0.021516 | -0.82207 | Kinase family protein | elongation zone | 6 h |
| HORVU.MOREX.r3.1HG0063810.1 | -0.41341 | 4.973905 | -3.59673 | 0.000708 | 0.021078 | -0.82523 | Cyclin family protein | elongation zone | 6 h |
| HORVU.MOREX.r3.4HG0382550.1 | 0.659022 | 5.636635 | 3.64992 | 0.0006 | 0.019011 | -0.83025 | Xyloglucan 6-xylosyltransferase 1 | elongation zone | 6 h |
| HORVU.MOREX.r3.1HG0078750.1 | -0.50118 | 3.941561 | -3.53535 | 0.000855 | 0.023177 | -0.83367 | Tesmin/TSO1-like CXC domain containing protein | elongation zone | 6 h |
| HORVU.MOREX.r3.2HG0117030.1 | 0.624057 | 7.823219 | 3.710981 | 0.000496 | 0.017032 | -0.84019 | Glutamate synthase, putative | elongation zone | 6 h |
| HORVU.MOREX.r3.2HG0120650.1 | 1.012572 | 1.67255 | 3.711013 | 0.000496 | 0.017032 | -0.84356 | Peroxidase | elongation zone | 6 h |
| HORVU.MOREX.r3.1HG0049450.1 | -0.45109 | 5.425708 | -3.63297 | 0.000633 | 0.019703 | -0.84449 | #N/A | elongation zone | 6 h |
| HORVU.MOREX.r3.7HG0658270.1 | -0.60436 | 2.885178 | -3.53702 | 0.00085 | 0.023137 | -0.8503 | Hyp O-arabinosyltransferase-like protein | elongation zone | 6 h |
| HORVU.MOREX.r3.3HG0294670.1 | -1.5833 | -0.36802 | -3.52646 | 0.000878 | 0.023338 | -0.85976 | demeter-like protein 3 | elongation zone | 6 h |
| HORVU.MOREX.r3.5HG0528570.1 | 1.082949 | 0.170023 | 3.54314 | 0.000834 | 0.022819 | -0.8704 | Protein PLANT CADMIUM RESISTANCE 2 | elongation zone | 6 h |
| HORVU.MOREX.r3.6HG0542030.1 | 1.090227 | 1.995705 | 3.722414 | 0.000479 | 0.016568 | -0.87062 | Acid beta-fructofuranosidase | elongation zone | 6 h |
| HORVU.MOREX.r3.4HG0343540.1 | -1.86821 | -0.26772 | -3.551 | 0.000815 | 0.022588 | -0.87555 | BSD domain containing protein, expressed | elongation zone | 6 h |
| HORVU.MOREX.r3.7HG0641180.1 | 0.477929 | 5.725264 | 3.633743 | 0.000631 | 0.019703 | -0.87672 | Protein ENHANCED DISEASE RESISTANCE 2-like | elongation zone | 6 h |
| HORVU.MOREX.r3.5HG0537090.1 | -0.3276 | 6.791678 | -3.66542 | 0.000572 | 0.0184 | -0.87699 | WD40 repeat-containing protein | elongation zone | 6 h |
| HORVU.MOREX.r3.3HG0313090.1 | 0.808062 | 2.944421 | 3.521623 | 0.000891 | 0.023455 | -0.8778 | Thiamin pyrophosphokinase 1 | elongation zone | 6 h |
| HORVU.MOREX.r3.1HG0050440.1 | 1.630883 | -1.13792 | 3.445783 | 0.001122 | 0.027022 | -0.88028 | DUF538 family protein (Protein of unknown function, DUF538) | elongation zone | 6 h |
| HORVU.MOREX.r3.5HG0426280.1 | 0.802689 | 2.961285 | 3.450813 | 0.001105 | 0.026745 | -0.88209 | GTP binding protein | elongation zone | 6 h |
| HORVU.MOREX.r3.5HG0500180.1 | -0.41309 | 4.872313 | -3.58302 | 0.000738 | 0.02155 | -0.88392 | Peptidyl-prolyl cis-trans isomerase | elongation zone | 6 h |
| HORVU.MOREX.r3.7HG0638940.1 | 2.403498 | 2.010885 | 3.533601 | 0.000859 | 0.023177 | -0.88518 | DnaJ domain-containing protein | elongation zone | 6 h |
| HORVU.MOREX.r3.2HG0100900.2 | 1.466048 | 0.62711 | 3.445935 | 0.001121 | 0.027022 | -0.89236 | Phosphoglycerate mutase-like protein | elongation zone | 6 h |
| HORVU.MOREX.r3.5HG0527500.1 | -0.51146 | 4.452639 | -3.53065 | 0.000867 | 0.023177 | -0.89468 | Ribosomal L11 methyltransferase | elongation zone | 6 h |
| HORVU.MOREX.r3.6HG0629810.2 | 1.014768 | 1.891863 | 3.442642 | 0.001132 | 0.027022 | -0.89728 | Auxin-responsive protein | elongation zone | 6 h |
| HORVU.MOREX.r3.1HG0083730.1 | 1.06344 | 0.340392 | 3.482628 | 0.001003 | 0.025247 | -0.90119 | Lipid transfer protein | elongation zone | 6 h |
| HORVU.MOREX.r3.1HG0000140.1 | -0.36426 | 5.300482 | -3.59039 | 0.000722 | 0.021192 | -0.90275 | Glycosyltransferase-like KOBITO 1 | elongation zone | 6 h |
| HORVU.MOREX.r3.4HG0356090.1 | -0.62258 | 2.309877 | -3.44397 | 0.001128 | 0.027022 | -0.90891 | DELLA domain GRAS family transcription factor GAI | elongation zone | 6 h |
| HORVU.MOREX.r3.1HG0054510.1 | 1.021317 | 2.568148 | 3.494517 | 0.000968 | 0.024639 | -0.91572 | Protein PLASTID MOVEMENT IMPAIRED 1 | elongation zone | 6 h |
| HORVU.MOREX.r3.2HG0173960.1 | 0.683328 | 2.814915 | 3.479501 | 0.001013 | 0.025374 | -0.9163 | Peroxidase | elongation zone | 6 h |
| HORVU.MOREX.r3.4HG0377070.1 | -0.46487 | 5.867895 | -3.59897 | 0.000703 | 0.020964 | -0.91789 | Kinase family protein | elongation zone | 6 h |
| HORVU.MOREX.r3.6HG0618760.1 | -0.74765 | 4.331592 | -3.52446 | 0.000883 | 0.023391 | -0.91933 | WRKY transcription factor, putative | elongation zone | 6 h |
| HORVU.MOREX.r3.6HG0628410.1 | -0.32229 | 8.572245 | -3.76747 | 0.000416 | 0.015336 | -0.91991 | Germin-like protein 1-1 | elongation zone | 6 h |
| HORVU.MOREX.r3.6HG0626650.1 | -0.26756 | 8.259289 | -3.69582 | 0.00052 | 0.017591 | -0.92463 | 26S protease regulatory subunit-like protein | elongation zone | 6 h |
| HORVU.MOREX.r3.7HG0651110.1 | -0.35594 | 7.468711 | -3.66923 | 0.000565 | 0.018298 | -0.92934 | AT4G29520-like protein | elongation zone | 6 h |
| HORVU.MOREX.r3.2HG0102600.1 | -0.41046 | 5.97563 | -3.60586 | 0.000688 | 0.020734 | -0.92939 | Peptidylprolyl isomerase | elongation zone | 6 h |
| HORVU.MOREX.r3.4HG0354980.1 | 0.687205 | 2.412831 | 3.498746 | 0.000955 | 0.024445 | -0.93222 | #N/A | elongation zone | 6 h |
| HORVU.MOREX.r3.4HG0331440.1 | 1.502997 | -0.62465 | 3.435984 | 0.001155 | 0.027375 | -0.93919 | LOB domain-containing protein, putative | elongation zone | 6 h |
| HORVU.MOREX.r3.2HG0152370.1 | -0.42073 | 4.701662 | -3.55696 | 0.0008 | 0.02239 | -0.93946 | MD-2-related lipid recognition domain-containing protein / ML domain-containing | elongation zone | 6 h |
| HORVU.MOREX.r3.3HG0240350.1 | 0.666715 | 2.152333 | 3.491601 | 0.000976 | 0.024796 | -0.94279 | Two-component response regulator | elongation zone | 6 h |
| HORVU.MOREX.r3.7HG0715170.1 | 1.505029 | -0.05365 | 3.428796 | 0.001181 | 0.027814 | -0.94361 | Glycosyltransferase | elongation zone | 6 h |
| HORVU.MOREX.r3.4HG0348450.1 | -1.24325 | 1.119942 | -3.42634 | 0.001189 | 0.027891 | -0.94508 | O-fucosyltransferase family protein | elongation zone | 6 h |
| HORVU.MOREX.r3.4HG0331730.1 | -0.30864 | 6.082061 | -3.60717 | 0.000685 | 0.020703 | -0.94565 | Zinc finger, C2H2 | elongation zone | 6 h |
| HORVU.MOREX.r3.7HG0644540.1 | 0.755999 | 3.792744 | 3.463956 | 0.001062 | 0.026155 | -0.96051 | 1-aminocyclopropane-1-carboxylate synthase | elongation zone | 6 h |

**Table S1** Continued.

| ID | logFC | AveExpr | t | P.Value | adj.P.Val | B | gene description | root zone | time point |
|---|---|---|---|---|---|---|---|---|---|
| HORVU.MOREX.r3.6HG0595580.1 | 2.387881 | -0.31218 | 3.417633 | 0.001221 | 0.028437 | -0.96489 | Ammonium transporter | elongation zone | 6 h |
| HORVU.MOREX.r3.6HG0633960.1 | 0.977583 | 2.11648 | 3.650654 | 0.000599 | 0.018997 | -0.96976 | Monosaccharide-sensing protein 2 | elongation zone | 6 h |
| HORVU.MOREX.r3.2HG0204340.1 | -2.38704 | -0.82148 | -3.73562 | 0.000459 | 0.01614 | -0.97123 | ERD (early-responsive to dehydration stress) family protein | elongation zone | 6 h |
| HORVU.MOREX.r3.1HG0004560.1 | 0.877794 | 5.305216 | 3.530018 | 0.000869 | 0.023177 | -0.97362 | ERD (Early-responsive to dehydration stress) family protein | elongation zone | 6 h |
| HORVU.MOREX.r3.4HG0398550.1 | 0.95697 | 6.045114 | 3.697042 | 0.000518 | 0.017582 | -0.97875 | Calcium-transporting ATPase | elongation zone | 6 h |
| HORVU.MOREX.r3.7HG0748670.1 | 2.330437 | -1.85496 | 3.406744 | 0.001261 | 0.028987 | -0.97913 | GDSL esterase/lipase | elongation zone | 6 h |
| HORVU.MOREX.r3.2HG0160130.1 | 0.595919 | 3.66281 | 3.603441 | 0.000693 | 0.020828 | -0.98055 | Basic helix-loop-helix transcription factor | elongation zone | 6 h |
| HORVU.MOREX.r3.7HG0699020.1 | 1.492684 | -0.26924 | 3.442819 | 0.001132 | 0.027022 | -0.98574 | Retrotransposon protein, putative, Ty3-gypsy subclass | elongation zone | 6 h |
| HORVU.MOREX.r3.5HG0462200.1 | 1.66813 | -0.85589 | 3.404127 | 0.001271 | 0.02915 | -0.98797 | Metacaspase | elongation zone | 6 h |
| HORVU.MOREX.r3.1HG0061660.1 | 0.885936 | 3.333271 | 3.580949 | 0.000743 | 0.021565 | -0.98837 | RING/FYVE/PHD zinc finger superfamily protein | elongation zone | 6 h |
| HORVU.MOREX.r3.5HG0423820.1 | 0.952504 | 2.29448 | 3.412191 | 0.001241 | 0.028698 | -0.99031 | #N/A | elongation zone | 6 h |
| HORVU.MOREX.r3.5HG0528520.1 | -0.37801 | 4.086447 | -3.52817 | 0.000873 | 0.023277 | -0.99413 | Epoxide hydrolase 2 | elongation zone | 6 h |
| HORVU.MOREX.r3.2HG0193000.1 | -0.71756 | 5.418419 | -3.53252 | 0.000862 | 0.023177 | -0.99747 | HTH-type transcriptional regulator YidZ | elongation zone | 6 h |
| HORVU.MOREX.r3.6HG0606930.1 | -0.37452 | 5.680326 | -3.57274 | 0.000762 | 0.02193 | -0.99998 | 3-ketodihydrosphingosine reductase | elongation zone | 6 h |
| HORVU.MOREX.r3.7HG0669410.1 | 0.508302 | 3.917601 | 3.504741 | 0.000938 | 0.024245 | -1.00593 | Cytochrome P450 | elongation zone | 6 h |
| HORVU.MOREX.r3.1HG0032930.1 | 0.670299 | 5.503727 | 3.619237 | 0.00066 | 0.020312 | -1.00638 | lipase, putative (DUF620) | elongation zone | 6 h |
| HORVU.MOREX.r3.3HG0288370.1 | -0.97215 | 2.377529 | -3.40334 | 0.001274 | 0.029154 | -1.00691 | Chaperone clpb, putative | elongation zone | 6 h |
| HORVU.MOREX.r3.2HG0132590.1 | -0.3315 | 6.039953 | -3.61772 | 0.000663 | 0.020376 | -1.01251 | GDP-fucose protein O-fucosyltransferase 2 | elongation zone | 6 h |
| HORVU.MOREX.r3.3HG0292080.1 | -0.49333 | 4.723449 | -3.53021 | 0.000868 | 0.023177 | -1.01265 | F-box family protein-like | elongation zone | 6 h |
| HORVU.MOREX.r3.6HG0622550.1 | -0.34993 | 5.355981 | -3.56854 | 0.000772 | 0.022075 | -1.01301 | Cyclic nucleotide-gated channel | elongation zone | 6 h |
| HORVU.MOREX.r3.2HG0186750.1 | 2.919088 | 1.34332 | 3.638218 | 0.000623 | 0.019533 | -1.01355 | Growth-regulating factor | elongation zone | 6 h |
| HORVU.MOREX.r3.7HG0747690.1 | 3.555856 | 0.558808 | 3.609986 | 0.000679 | 0.020694 | -1.01965 | Hydroxycinnamoyl-CoA shikimate/quinate hydroxycinnamoyltransferase | elongation zone | 6 h |
| HORVU.MOREX.r3.3HG0280920.1 | -0.734 | 3.219452 | -3.55946 | 0.000794 | 0.022311 | -1.01971 | DUF4228 domain protein | elongation zone | 6 h |
| HORVU.MOREX.r3.6HG0620630.1 | 0.917238 | 3.35364 | 3.643419 | 0.000613 | 0.019279 | -1.02228 | Subtilisin-like protease | elongation zone | 6 h |
| HORVU.MOREX.r3.4HG0391160.1 | 0.961782 | 4.078357 | 3.437601 | 0.00115 | 0.027274 | -1.0225 | Patatin | elongation zone | 6 h |
| HORVU.MOREX.r3.7HG0726770.1 | 0.936261 | 2.29225 | 3.496706 | 0.000961 | 0.024536 | -1.02467 | Glycine-rich protein | elongation zone | 6 h |
| HORVU.MOREX.r3.4HG0341800.1 | 0.651379 | 2.209476 | 3.565182 | 0.00078 | 0.02217 | -1.02687 | Receptor-like kinase | elongation zone | 6 h |
| HORVU.MOREX.r3.2HG0182400.1 | 0.787318 | 5.649464 | 3.606868 | 0.000686 | 0.020703 | -1.03023 | Aquaporin | elongation zone | 6 h |
| HORVU.MOREX.r3.7HG0711890.1 | 1.034972 | 2.553034 | 3.524749 | 0.000883 | 0.023391 | -1.03035 | O-methyltransferase | elongation zone | 6 h |
| HORVU.MOREX.r3.4HG0364210.1 | 2.506742 | -2.34248 | 3.582055 | 0.00074 | 0.021553 | -1.03304 | #N/A | elongation zone | 6 h |
| HORVU.MOREX.r3.6HG0577770.1 | -1.91662 | -0.27649 | -3.52521 | 0.000881 | 0.023391 | -1.03415 | P-loop containing nucleoside triphosphate hydrolases superfamily protein | elongation zone | 6 h |
| HORVU.MOREX.r3.2HG0197230.1 | 1.239964 | -1.20366 | 3.412461 | 0.00124 | 0.028698 | -1.04106 | disease resistance protein (TIR-NBS-LRR class) | elongation zone | 6 h |
| HORVU.MOREX.r3.7HG0655170.1 | 1.356424 | 0.095219 | 3.426857 | 0.001187 | 0.027891 | -1.04256 | Late embryogenesis abundant protein | elongation zone | 6 h |
| HORVU.MOREX.r3.5HG0517640.1 | -0.6841 | 6.451472 | -3.57632 | 0.000754 | 0.021777 | -1.04518 | Glucan endo-1,3-beta-glucosidase, putative | elongation zone | 6 h |
| HORVU.MOREX.r3.2HG0120470.1 | 1.316735 | 3.222129 | 3.668206 | 0.000567 | 0.018303 | -1.05145 | Peroxidase | elongation zone | 6 h |
| HORVU.MOREX.r3.4HG0398090.1 | 0.671999 | 4.062819 | 3.552775 | 0.00081 | 0.022568 | -1.05179 | Syntaxin protein | elongation zone | 6 h |
| HORVU.MOREX.r3.4HG0388310.1 | 0.411842 | 3.870759 | 3.542848 | 0.000835 | 0.022819 | -1.05405 | Protein kinase | elongation zone | 6 h |
| HORVU.MOREX.r3.3HG0244040.1 | 1.22859 | -2.29384 | 3.390814 | 0.001323 | 0.029902 | -1.05708 | ethylene-responsive transcription factor | elongation zone | 6 h |
| HORVU.MOREX.r3.4HG0355390.1 | -0.74552 | 1.970371 | -3.40081 | 0.001284 | 0.02931 | -1.05918 | RING/U-box superfamily protein | elongation zone | 6 h |
| HORVU.MOREX.r3.5HG0465400.1 | 1.191877 | 0.747502 | 3.548547 | 0.000821 | 0.022653 | -1.07019 | Trichome birefringence-like protein | elongation zone | 6 h |
| HORVU.MOREX.r3.7HG0674440.1 | -0.45381 | 5.321799 | -3.54829 | 0.000821 | 0.022653 | -1.07201 | Aluminum activated malate transporter family protein | elongation zone | 6 h |

**Table S1** Continued.

| ID | logFC | AveExpr | t | P.Value | adj.P.Val | B | gene description | root zone | time point |
|---|---|---|---|---|---|---|---|---|---|
| HORVU.MOREX.r3.6HG0572400.1 | -0.41327 | 3.75498 | -3.47036 | 0.001041 | 0.025776 | -1.07302 | F-box family protein | elongation zone | 6 h |
| HORVU.MOREX.r3.4HG0381150.1 | 2.074882 | -0.56149 | 3.549958 | 0.000817 | 0.022629 | -1.08111 | nuclear pore complex protein (DUF3414) | elongation zone | 6 h |
| HORVU.MOREX.r3.4HG0343710.1 | -0.34929 | 4.292721 | -3.50719 | 0.000931 | 0.024187 | -1.08281 | Calcineurin-like metallo-phosphoesterase superfamily protein | elongation zone | 6 h |
| HORVU.MOREX.r3.5HG0510160.1 | -0.45995 | 4.316907 | -3.46305 | 0.001065 | 0.026178 | -1.08575 | Myb/SANT-like DNA-binding domain protein | elongation zone | 6 h |
| HORVU.MOREX.r3.5HG0524060.1 | 1.293089 | 3.909294 | 3.498747 | 0.000955 | 0.024445 | -1.08979 | Remorin | elongation zone | 6 h |
| HORVU.MOREX.r3.1HG0056240.1 | 0.671893 | 4.721848 | 3.552163 | 0.000812 | 0.022568 | -1.09102 | Trehalose 6-phosphate phosphatase | elongation zone | 6 h |
| HORVU.MOREX.r3.3HG0278630.1 | 0.519308 | 5.03935 | 3.532936 | 0.000861 | 0.023177 | -1.09385 | Ras-related protein, expressed | elongation zone | 6 h |
| HORVU.MOREX.r3.1HG0069380.1 | 0.824246 | 2.129643 | 3.486991 | 0.00099 | 0.025053 | -1.09471 | Cysteine proteinase inhibitor | elongation zone | 6 h |
| HORVU.MOREX.r3.1HG0092200.1 | 1.364471 | 1.592343 | 3.361325 | 0.001445 | 0.031276 | -1.09884 | Solanesyl diphosphate synthase family protein | elongation zone | 6 h |
| HORVU.MOREX.r3.2HG0121570.1 | -0.50747 | 4.65733 | -3.48316 | 0.001002 | 0.025247 | -1.10642 | Protein LTV1 | elongation zone | 6 h |
| HORVU.MOREX.r3.2HG0182650.1 | 2.04427 | -1.3514 | 3.370063 | 0.001408 | 0.030961 | -1.10742 | Cytokinin oxidase/dehydrogenase | elongation zone | 6 h |
| HORVU.MOREX.r3.6HG0609520.1 | -0.46892 | 8.966633 | -3.65911 | 0.000583 | 0.018648 | -1.10792 | Fatty acid desaturase | elongation zone | 6 h |
| HORVU.MOREX.r3.2HG0211880.1 | -0.4562 | 4.077958 | -3.45819 | 0.00108 | 0.026331 | -1.10829 | Exostosin family protein | elongation zone | 6 h |
| HORVU.MOREX.r3.4HG0373800.1 | -1.0656 | 1.360295 | -3.35658 | 0.001465 | 0.031424 | -1.10939 | helicase with zinc finger protein | elongation zone | 6 h |
| HORVU.MOREX.r3.1HG0044800.1 | 2.156857 | -1.71771 | 3.359186 | 0.001454 | 0.031376 | -1.11211 | DnaJ heat shock amino-terminal domain protein | elongation zone | 6 h |
| HORVU.MOREX.r3.3HG0309410.1 | -0.97268 | 3.365238 | -3.38368 | 0.001352 | 0.03018 | -1.11324 | Subtilisin-like protease | elongation zone | 6 h |
| HORVU.MOREX.r3.1HG0052250.1 | 1.481307 | 0.582616 | 3.358468 | 0.001457 | 0.031377 | -1.11426 | Membrane-associated kinase regulator-like protein, putative | elongation zone | 6 h |
| HORVU.MOREX.r3.6HG0579050.1 | 1.256213 | 0.455547 | 3.356607 | 0.001465 | 0.031424 | -1.11451 | Pentatricopeptide repeat-containing protein At5g14820, mitochondrial | elongation zone | 6 h |
| HORVU.MOREX.r3.3HG0308350.1 | 0.910081 | 4.926407 | 3.500607 | 0.00095 | 0.024398 | -1.11725 | Cysteine-rich receptor kinase | elongation zone | 6 h |
| HORVU.MOREX.r3.1HG0051740.1 | -1.90775 | 3.433417 | -3.36841 | 0.001415 | 0.030961 | -1.12137 | Glutathione s-transferase, putative | elongation zone | 6 h |
| HORVU.MOREX.r3.7HG0717610.1 | 0.589725 | 4.008622 | 3.510342 | 0.000922 | 0.024017 | -1.12243 | Pollen-specific protein SF21 | elongation zone | 6 h |
| HORVU.MOREX.r3.3HG0280960.1 | 0.713944 | 1.805356 | 3.462876 | 0.001065 | 0.026178 | -1.12475 | Glycosyltransferase | elongation zone | 6 h |
| HORVU.MOREX.r3.4HG0379290.1 | -0.31691 | 5.28798 | -3.51106 | 0.00092 | 0.024016 | -1.12964 | COP1-interacting-like protein, putative | elongation zone | 6 h |
| HORVU.MOREX.r3.7HG0739910.1 | -0.47919 | 4.210714 | -3.4685 | 0.001047 | 0.025891 | -1.1365 | Disease resistance protein (NBS-LRR class) family | elongation zone | 6 h |
| HORVU.MOREX.r3.5HG0508610.1 | 1.210921 | 3.825671 | 3.507445 | 0.00093 | 0.024187 | -1.1369 | Patatin | elongation zone | 6 h |
| HORVU.MOREX.r3.1HG0072580.1 | 0.507449 | 5.490114 | 3.510775 | 0.000921 | 0.024016 | -1.13814 | Aminotransferase-related family protein | elongation zone | 6 h |
| HORVU.MOREX.r3.5HG0528310.1 | 0.571033 | 3.716139 | 3.482021 | 0.001005 | 0.025247 | -1.13839 | SPX domain-containing family protein | elongation zone | 6 h |
| HORVU.MOREX.r3.2HG0117100.1 | 0.914701 | 2.815111 | 3.344698 | 0.001518 | 0.031854 | -1.13904 | NHL repeat-containing protein | elongation zone | 6 h |
| HORVU.MOREX.r3.2HG0198320.1 | 1.713974 | -0.19189 | 3.380128 | 0.001366 | 0.030437 | -1.14285 | Protein translation factor SUI1-like protein | elongation zone | 6 h |
| HORVU.MOREX.r3.6HG0558840.1 | -0.26251 | 6.872948 | -3.55747 | 0.000799 | 0.02239 | -1.14679 | Kinetochore protein nuf2, putative | elongation zone | 6 h |
| HORVU.MOREX.r3.4HG0410910.1 | 2.274625 | -2.20066 | 3.478218 | 0.001017 | 0.025374 | -1.15344 | Ubiquitin carboxyl-terminal hydrolase CYLD | elongation zone | 6 h |
| HORVU.MOREX.r3.5HG0444060.1 | -0.88566 | 1.863328 | -3.35509 | 0.001472 | 0.031497 | -1.15513 | Calmodulin-binding family protein, putative, expressed | elongation zone | 6 h |
| HORVU.MOREX.r3.3HG0276130.1 | 1.05447 | -0.22255 | 3.388953 | 0.001331 | 0.029969 | -1.15932 | Phospholipase A1 | elongation zone | 6 h |
| HORVU.MOREX.r3.6HG0546160.1 | 0.788552 | 2.97611 | 3.513646 | 0.000913 | 0.023928 | -1.16082 | Acylamino-acid-releasing enzyme | elongation zone | 6 h |
| HORVU.MOREX.r3.3HG0290300.1 | 2.995613 | -2.72546 | 3.56112 | 0.00079 | 0.022277 | -1.16252 | Homeobox leucine zipper protein | elongation zone | 6 h |
| HORVU.MOREX.r3.2HG0171780.1 | 1.569453 | -1.26245 | 3.335709 | 0.001559 | 0.032443 | -1.16258 | transcription repressor | elongation zone | 6 h |
| HORVU.MOREX.r3.2HG0140700.1 | -0.31445 | 5.904168 | -3.53193 | 0.000864 | 0.023177 | -1.16261 | #N/A | elongation zone | 6 h |
| HORVU.MOREX.r3.6HG0542050.1 | 1.293229 | 1.403897 | 3.600901 | 0.000699 | 0.02093 | -1.16408 | Acid beta-fructofuranosidase | elongation zone | 6 h |
| HORVU.MOREX.r3.5HG0485960.1 | 1.767777 | -2.16732 | 3.350303 | 0.001493 | 0.031694 | -1.16548 | Transmembrane emp24 domain-containing protein | elongation zone | 6 h |
| HORVU.MOREX.r3.5HG0528280.1 | 0.424964 | 5.95615 | 3.546665 | 0.000825 | 0.022736 | -1.16667 | Organic solute transporter-like | elongation zone | 6 h |
| HORVU.MOREX.r3.2HG0189780.1 | 1.683591 | 0.49991 | 3.368277 | 0.001415 | 0.030961 | -1.17027 | ethylene-responsive transcription factor | elongation zone | 6 h |

**Table S1** Continued.

| ID | logFC | AveExpr | t | P.Value | adj.P.Val | B | gene description | root zone | time point |
|---|---|---|---|---|---|---|---|---|---|
| HORVU.MOREX.r3.1HG0086480.1 | 1.367622 | 0.13169 | 3.358812 | 0.001456 | 0.031377 | -1.17405 | UV-B-induced protein, chloroplastic | elongation zone | 6 h |
| HORVU.MOREX.r3.3HG0242870.1 | -0.29248 | 7.618528 | -3.59984 | 0.000701 | 0.020963 | -1.17846 | Carboxypeptidase | elongation zone | 6 h |
| HORVU.MOREX.r3.6HG0627920.1 | -0.28575 | 7.364979 | -3.59031 | 0.000722 | 0.021192 | -1.18198 | Ankyrin repeat protein-like | elongation zone | 6 h |
| HORVU.MOREX.r3.7HG0667180.1 | -0.79845 | 1.133168 | -3.32841 | 0.001593 | 0.03286 | -1.19533 | F-box protein | elongation zone | 6 h |
| HORVU.MOREX.r3.4HG0403370.1 | -0.40266 | 4.841977 | -3.46513 | 0.001058 | 0.026094 | -1.19667 | O-fucosyltransferase family protein | elongation zone | 6 h |
| HORVU.MOREX.r3.1HG0089130.1 | -0.35304 | 5.608758 | -3.47388 | 0.00103 | 0.025565 | -1.20081 | Serine/Threonine-kinase pakA-like protein | elongation zone | 6 h |
| HORVU.MOREX.r3.4HG0393520.1 | 0.781571 | 3.79819 | 3.422568 | 0.001203 | 0.028114 | -1.20317 | Microtubule associated protein family protein, putative, expressed | elongation zone | 6 h |
| HORVU.MOREX.r3.2HG0172730.1 | -0.43328 | 7.548613 | -3.56918 | 0.00077 | 0.022075 | -1.20709 | Alpha-ketoglutarate-dependent dioxygenase alkB-like protein 2 | elongation zone | 6 h |
| HORVU.MOREX.r3.3HG0273730.1 | -0.55109 | 6.291958 | -3.53371 | 0.000859 | 0.023177 | -1.20766 | Cytochrome P450 | elongation zone | 6 h |
| HORVU.MOREX.r3.2HG0179320.1 | 0.799008 | 3.064606 | 3.430059 | 0.001176 | 0.02774 | -1.21157 | BTB/POZ domain-containing protein | elongation zone | 6 h |
| HORVU.MOREX.r3.5HG0522630.1 | 0.721851 | 1.481024 | 3.521879 | 0.00089 | 0.023455 | -1.21186 | Blue copper protein | elongation zone | 6 h |
| HORVU.MOREX.r3.5HG0472810.1 | 0.883584 | 2.371978 | 3.554182 | 0.000807 | 0.022551 | -1.21234 | Aquaporin | elongation zone | 6 h |
| HORVU.MOREX.r3.4HG0410640.1 | -0.37826 | 8.151713 | -3.59083 | 0.000721 | 0.021192 | -1.21394 | Dolichyl-diphosphooligosaccharide--protein glycosyltransferase subunit 1 | elongation zone | 6 h |
| HORVU.MOREX.r3.2HG0109340.1 | 1.1237 | 0.874826 | 3.369099 | 0.001412 | 0.030961 | -1.21485 | Wound-induced protein 1, putative, expressed | elongation zone | 6 h |
| HORVU.MOREX.r3.7HG0660720.1 | 1.67228 | 1.415323 | 3.352681 | 0.001483 | 0.031653 | -1.22023 | Ring finger protein, putative | elongation zone | 6 h |
| HORVU.MOREX.r3.5HG0438280.1 | -1.80554 | 0.696137 | -3.35361 | 0.001478 | 0.031602 | -1.22094 | Protein translocase subunit SecA | elongation zone | 6 h |
| HORVU.MOREX.r3.1HG0073010.1 | 0.83823 | 1.113004 | 3.520944 | 0.000893 | 0.023461 | -1.22218 | Kinase family protein | elongation zone | 6 h |
| HORVU.MOREX.r3.7HG0710170.1 | -2.23823 | 0.072486 | -3.45827 | 0.00108 | 0.026331 | -1.22528 | ATP-dependent DNA helicase 2 subunit KU70 | elongation zone | 6 h |
| HORVU.MOREX.r3.1HG0083040.1 | -0.74817 | 1.526854 | -3.39189 | 0.001319 | 0.029839 | -1.22565 | Retrovirus-related Pol polyprotein from transposon TNT 1-94 | elongation zone | 6 h |
| HORVU.MOREX.r3.7HG0667280.1 | -0.43161 | 4.293106 | -3.44295 | 0.001131 | 0.027022 | -1.226 | Hexosyltransferase | elongation zone | 6 h |
| HORVU.MOREX.r3.6HG0613230.1 | -0.36781 | 6.009357 | -3.52752 | 0.000875 | 0.023293 | -1.23323 | Allergen, putative | elongation zone | 6 h |
| HORVU.MOREX.r3.6HG0566940.1 | 0.575621 | 4.632192 | 3.456992 | 0.001084 | 0.026395 | -1.23533 | Sulfurtransferase | elongation zone | 6 h |
| HORVU.MOREX.r3.2HG0195600.1 | -0.3589 | 4.30012 | -3.44221 | 0.001134 | 0.027022 | -1.23689 | GPI mannosyltransferase 2 | elongation zone | 6 h |
| HORVU.MOREX.r3.3HG0322740.1 | -0.90268 | 0.784975 | -3.35622 | 0.001467 | 0.031424 | -1.23717 | transmembrane protein, putative (DUF247) | elongation zone | 6 h |
| HORVU.MOREX.r3.7HG0751290.1 | -0.90823 | 0.60944 | -3.30604 | 0.001702 | 0.03438 | -1.23788 | Receptor-kinase, putative | elongation zone | 6 h |
| HORVU.MOREX.r3.7HG0745700.1 | -0.30534 | 5.461365 | -3.49791 | 0.000958 | 0.024477 | -1.23959 | Phox (PX) domain-containing protein | elongation zone | 6 h |
| HORVU.MOREX.r3.3HG0252250.1 | 1.398626 | -1.6584 | 3.372568 | 0.001397 | 0.03093 | -1.23999 | Expansin | elongation zone | 6 h |
| HORVU.MOREX.r3.7HG0656570.1 | 0.910474 | 1.40656 | 3.301451 | 0.001726 | 0.034545 | -1.24512 | Pentatricopeptide repeat-containing protein | elongation zone | 6 h |
| HORVU.MOREX.r3.2HG0120560.1 | 1.319028 | 3.65156 | 3.623809 | 0.000651 | 0.020087 | -1.24529 | Peroxidase | elongation zone | 6 h |
| HORVU.MOREX.r3.5HG0474540.1 | -0.36376 | 7.609556 | -3.56527 | 0.00078 | 0.02217 | -1.24578 | Protein transport protein Sec61 subunit alpha | elongation zone | 6 h |
| HORVU.MOREX.r3.7HG0722380.1 | -0.50336 | 2.715102 | -3.35625 | 0.001467 | 0.031424 | -1.24987 | Nudix hydrolase | elongation zone | 6 h |
| HORVU.MOREX.r3.2HG0137140.1 | -1.13362 | 0.734361 | -3.30229 | 0.001721 | 0.034545 | -1.25066 | Acyl-CoA dehydrogenase family member 10 | elongation zone | 6 h |
| HORVU.MOREX.r3.7HG0679050.1 | 0.565689 | 4.002868 | 3.404806 | 0.001269 | 0.029123 | -1.25557 | Upstream activation factor subunit spp27 | elongation zone | 6 h |
| HORVU.MOREX.r3.6HG0593200.1 | -0.87124 | 4.784544 | -3.4036 | 0.001273 | 0.029154 | -1.25682 | Short-chain dehydrogenase/reductase family protein | elongation zone | 6 h |
| HORVU.MOREX.r3.1HG0030290.1 | 2.990007 | -1.84835 | 3.512989 | 0.000915 | 0.023945 | -1.25899 | PRA1 family protein | elongation zone | 6 h |
| HORVU.MOREX.r3.5HG0488300.1 | 1.483763 | 2.76736 | 3.39983 | 0.001288 | 0.029352 | -1.25958 | ABC transporter G family member | elongation zone | 6 h |
| HORVU.MOREX.r3.6HG0560140.1 | 2.173685 | -1.9167 | 3.294894 | 0.001759 | 0.03479 | -1.2647 | Endoglucanase | elongation zone | 6 h |
| HORVU.MOREX.r3.4HG0353120.1 | 1.422343 | -2.86086 | 3.295926 | 0.001754 | 0.034787 | -1.26878 | Transposon protein, putative, CACTA, En/Spm sub-class | elongation zone | 6 h |
| HORVU.MOREX.r3.3HG0239900.1 | 1.36656 | 3.375419 | 3.356822 | 0.001464 | 0.031424 | -1.27127 | Auxin-responsive protein | elongation zone | 6 h |
| HORVU.MOREX.r3.2HG0193510.2 | -0.36012 | 3.804131 | -3.41196 | 0.001242 | 0.028698 | -1.276 | Gpcr-type g protein 2 | elongation zone | 6 h |
| HORVU.MOREX.r3.6HG0611050.1 | 2.46585 | -0.65229 | 3.569275 | 0.00077 | 0.022075 | -1.27667 | Rubisco methyltransferase family protein | elongation zone | 6 h |

**Table S1** Continued.

| ID | logFC | AveExpr | t | P.Value | adj.P.Val | B | gene description | root zone | time point |
|---|---|---|---|---|---|---|---|---|---|
| HORVU.MOREX.r3.5HG0473970.1 | 2.298764 | -3.04844 | 3.330914 | 0.001582 | 0.032759 | -1.27748 | Glycosyltransferase | elongation zone | 6 h |
| HORVU.MOREX.r3.1HG0025870.1 | 0.56367 | 4.903164 | 3.453644 | 0.001095 | 0.026631 | -1.27978 | 6,7-dimethyl-8-ribityllumazine synthase | elongation zone | 6 h |
| HORVU.MOREX.r3.7HG0663940.1 | 0.835316 | 0.764913 | 3.444763 | 0.001125 | 0.027022 | -1.28354 | Alpha/beta-Hydrolases superfamily protein, putative | elongation zone | 6 h |
| HORVU.MOREX.r3.4HG0396030.1 | 1.938505 | 0.582741 | 3.293492 | 0.001767 | 0.034855 | -1.2851 | Tubulin alpha chain | elongation zone | 6 h |
| HORVU.MOREX.r3.7HG0636030.1 | 0.839192 | -1.10938 | 3.386966 | 0.001338 | 0.030049 | -1.28672 | hydroxysteroid dehydrogenase 3 | elongation zone | 6 h |
| HORVU.MOREX.r3.7HG0728080.1 | 0.821239 | 4.103954 | 3.327381 | 0.001598 | 0.032894 | -1.28779 | Protein CHLOROPLAST IMPORT APPARATUS 2 | elongation zone | 6 h |
| HORVU.MOREX.r3.3HG0254940.1 | 0.860821 | 2.451212 | 3.543125 | 0.000834 | 0.022819 | -1.28993 | Peroxidase | elongation zone | 6 h |
| HORVU.MOREX.r3.1HG0054010.1 | 0.571617 | 5.841657 | 3.535391 | 0.000854 | 0.023177 | -1.29355 | zinc finger B-box protein | elongation zone | 6 h |
| HORVU.MOREX.r3.7HG0701080.1 | 0.984388 | -1.51889 | 3.304935 | 0.001708 | 0.034427 | -1.29449 | MYB transcription factor | elongation zone | 6 h |
| HORVU.MOREX.r3.6HG0597320.1 | -0.28729 | 6.618535 | -3.50628 | 0.000934 | 0.024223 | -1.29535 | ATP-dependent Clp protease proteolytic subunit | elongation zone | 6 h |
| HORVU.MOREX.r3.7HG0661490.1 | -0.30737 | 6.094894 | -3.49597 | 0.000964 | 0.024561 | -1.29574 | Stromal cell-derived factor 2 | elongation zone | 6 h |
| HORVU.MOREX.r3.2HG0171810.1 | -0.48908 | 3.837575 | -3.36737 | 0.001419 | 0.031012 | -1.29725 | DUF1644 family protein | elongation zone | 6 h |
| HORVU.MOREX.r3.1HG0075440.1 | 0.716354 | 1.650312 | 3.414336 | 0.001233 | 0.028622 | -1.30319 | Protein NRT1/ PTR FAMILY 5.5 | elongation zone | 6 h |
| HORVU.MOREX.r3.5HG0485610.1 | 1.185152 | 0.79472 | 3.335437 | 0.00156 | 0.032443 | -1.30544 | Cytochrome P450, putative | elongation zone | 6 h |
| HORVU.MOREX.r3.3HG0258240.1 | 1.351762 | 1.126706 | 3.299747 | 0.001734 | 0.03465 | -1.30547 | Sacsin | elongation zone | 6 h |
| HORVU.MOREX.r3.7HG0711230.1 | 0.58968 | 3.91603 | 3.376543 | 0.001381 | 0.030664 | -1.3062 | Auxin-responsive protein | elongation zone | 6 h |
| HORVU.MOREX.r3.7HG0714530.1 | -1.36198 | 0.583064 | -3.2808 | 0.001834 | 0.03577 | -1.30637 | F-box protein PP2 | elongation zone | 6 h |
| HORVU.MOREX.r3.3HG0307340.1 | 1.299012 | 1.041748 | 3.278696 | 0.001845 | 0.035889 | -1.30674 | Receptor-like kinase | elongation zone | 6 h |
| HORVU.MOREX.r3.2HG0205200.1 | -0.45711 | 4.551525 | -3.41464 | 0.001232 | 0.028622 | -1.30802 | Ubiquitin carboxyl-terminal hydrolase | elongation zone | 6 h |
| HORVU.MOREX.r3.3HG0257130.1 | -0.50669 | 4.342452 | -3.39935 | 0.00129 | 0.029352 | -1.30857 | Presenilin | elongation zone | 6 h |
| HORVU.MOREX.r3.2HG0106260.1 | 2.138318 | -2.23574 | 3.330361 | 0.001584 | 0.032759 | -1.30885 | F-box family protein | elongation zone | 6 h |
| HORVU.MOREX.r3.5HG0484180.1 | 0.408465 | 5.636527 | 3.473067 | 0.001033 | 0.025597 | -1.3108 | DUF868 family protein | elongation zone | 6 h |
| HORVU.MOREX.r3.7HG0738770.1 | 0.926577 | 1.993948 | 3.272677 | 0.001878 | 0.036323 | -1.31682 | Receptor-like kinase | elongation zone | 6 h |
| HORVU.MOREX.r3.2HG0121670.1 | 0.737294 | 2.510007 | 3.460128 | 0.001074 | 0.026284 | -1.31969 | Ribosome-releasing factor 2, mitochondrial | elongation zone | 6 h |
| HORVU.MOREX.r3.1HG0046210.1 | 2.432764 | -0.44242 | 3.311924 | 0.001673 | 0.034187 | -1.32193 | Receptor kinase | elongation zone | 6 h |
| HORVU.MOREX.r3.3HG0236500.1 | 0.687458 | 5.370928 | 3.442402 | 0.001133 | 0.027022 | -1.324 | Pollen-specific protein SF21 | elongation zone | 6 h |
| HORVU.MOREX.r3.2HG0213250.1 | -0.53848 | 6.347479 | -3.50197 | 0.000946 | 0.024358 | -1.3309 | Choline transporter-related family protein | elongation zone | 6 h |
| HORVU.MOREX.r3.2HG0190300.1 | 0.597955 | 3.611092 | 3.425313 | 0.001193 | 0.027914 | -1.33675 | SHR5-receptor-like kinase | elongation zone | 6 h |
| HORVU.MOREX.r3.3HG0307950.1 | -0.303 | 6.122502 | -3.48244 | 0.001004 | 0.025247 | -1.34039 | GPI transamidase component PIG-S | elongation zone | 6 h |
| HORVU.MOREX.r3.7HG0706130.1 | 0.695062 | 4.046121 | 3.451295 | 0.001103 | 0.026745 | -1.34131 | Kinase family protein | elongation zone | 6 h |
| HORVU.MOREX.r3.4HG0403680.1 | -1.26457 | 3.612484 | -3.37319 | 0.001395 | 0.030906 | -1.34211 | #N/A | elongation zone | 6 h |
| HORVU.MOREX.r3.4HG0396760.1 | -0.40032 | 5.597414 | -3.4748 | 0.001027 | 0.025525 | -1.34253 | Protein root UVB sensitive 3 | elongation zone | 6 h |
| HORVU.MOREX.r3.3HG0292490.1 | 0.897386 | 0.56764 | 3.443305 | 0.00113 | 0.027022 | -1.34584 | Core-2/I-branching beta-1,6-N-acetylglucosaminyltransferase family protein, puta | elongation zone | 6 h |
| HORVU.MOREX.r3.1HG0040930.1 | 1.788286 | -2.88506 | 3.306536 | 0.0017 | 0.03438 | -1.34999 | F-box family protein | elongation zone | 6 h |
| HORVU.MOREX.r3.1HG0062630.1 | 0.521188 | 5.379794 | 3.426236 | 0.00119 | 0.027891 | -1.35007 | #N/A | elongation zone | 6 h |
| HORVU.MOREX.r3.5HG0495910.1 | 1.915482 | -0.71833 | 3.2621 | 0.001937 | 0.036879 | -1.35581 | Respiratory burst oxidase homolog | elongation zone | 6 h |
| HORVU.MOREX.r3.4HG0378080.1 | 2.8965 | -1.12859 | 3.505461 | 0.000936 | 0.024245 | -1.35662 | #N/A | elongation zone | 6 h |
| HORVU.MOREX.r3.4HG0383780.1 | 1.843386 | -0.06269 | 3.333248 | 0.001571 | 0.032621 | -1.36249 | Laccase | elongation zone | 6 h |
| HORVU.MOREX.r3.6HG0577220.1 | 1.847585 | -2.2339 | 3.252914 | 0.00199 | 0.037535 | -1.3659 | Glycosyltransferase | elongation zone | 6 h |
| HORVU.MOREX.r3.7HG0662420.1 | 1.086645 | 1.123246 | 3.308235 | 0.001691 | 0.034363 | -1.36782 | Kinase-like protein | elongation zone | 6 h |
| HORVU.MOREX.r3.7HG0676690.1 | 0.802091 | 4.543618 | 3.557152 | 0.000799 | 0.02239 | -1.36808 | Receptor-like kinase | elongation zone | 6 h |

**Table S1** Continued.

| ID | logFC | AveExpr | t | P.Value | adj.P.Val | B | gene description | root zone | time point |
|---|---|---|---|---|---|---|---|---|---|
| HORVU.MOREX.r3.1HG0082630.1 | -0.36116 | 5.023124 | -3.42597 | 0.001191 | 0.027891 | -1.36863 | Glycosyltransferase | elongation zone | 6 h |
| HORVU.MOREX.r3.4HG0409180.1 | -0.32848 | 6.8545 | -3.49902 | 0.000955 | 0.024445 | -1.36869 | Glutathione S-transferase | elongation zone | 6 h |
| HORVU.MOREX.r3.6HG0543540.1 | 1.505351 | -0.02472 | 3.342677 | 0.001527 | 0.032013 | -1.37011 | Expansin protein | elongation zone | 6 h |
| HORVU.MOREX.r3.5HG0458440.1 | -1.06742 | 1.256617 | -3.24885 | 0.002014 | 0.037845 | -1.3773 | phosphoglycerate/bisphosphoglycerate mutase family protein | elongation zone | 6 h |
| HORVU.MOREX.r3.5HG0501200.1 | -0.49089 | 4.695239 | -3.40731 | 0.001259 | 0.028987 | -1.37857 | Thioredoxin superfamily protein | elongation zone | 6 h |
| HORVU.MOREX.r3.3HG0293010.1 | -0.41184 | 4.239633 | -3.40709 | 0.00126 | 0.028987 | -1.37863 | WD-repeat protein, putative | elongation zone | 6 h |
| HORVU.MOREX.r3.2HG0168270.1 | -0.53513 | 3.16772 | -3.30572 | 0.001704 | 0.03438 | -1.37984 | Ubiquitin carboxyl-terminal hydrolase | elongation zone | 6 h |
| HORVU.MOREX.r3.6HG0588350.1 | -0.49933 | 4.125371 | -3.34559 | 0.001514 | 0.031802 | -1.38241 | RING finger family protein | elongation zone | 6 h |
| HORVU.MOREX.r3.2HG0141140.1 | 1.060458 | -2.85799 | 3.243919 | 0.002043 | 0.03815 | -1.38719 | DUF4228 domain protein | elongation zone | 6 h |
| HORVU.MOREX.r3.5HG0495730.1 | 0.985288 | 2.468419 | 3.381677 | 0.00136 | 0.030329 | -1.38732 | Kinase family protein | elongation zone | 6 h |
| HORVU.MOREX.r3.6HG0554050.1 | 0.865174 | -1.56957 | 3.294882 | 0.001759 | 0.03479 | -1.38751 | cyclin-dependent kinase inhibitor | elongation zone | 6 h |
| HORVU.MOREX.r3.4HG0342080.1 | 0.693973 | 4.067325 | 3.430642 | 0.001174 | 0.027723 | -1.38771 | Adenine/guanine permease | elongation zone | 6 h |
| HORVU.MOREX.r3.7HG0749090.1 | 1.083227 | -0.58865 | 3.369963 | 0.001408 | 0.030961 | -1.38909 | Expansin-like protein | elongation zone | 6 h |
| HORVU.MOREX.r3.6HG0582120.1 | 0.474716 | 3.287652 | 3.369458 | 0.00141 | 0.030961 | -1.3893 | magnesium transporter NIPA (DUF803) | elongation zone | 6 h |
| HORVU.MOREX.r3.3HG0249640.1 | 0.598829 | 4.297068 | 3.434269 | 0.001161 | 0.027485 | -1.39086 | E3 ubiquitin-protein ligase | elongation zone | 6 h |
| HORVU.MOREX.r3.6HG0555700.1 | -0.36921 | 8.875975 | -3.5732 | 0.000761 | 0.02193 | -1.39114 | Calreticulin/calnexin | elongation zone | 6 h |
| HORVU.MOREX.r3.6HG0633650.1 | -1.17282 | 0.935723 | -3.24671 | 0.002027 | 0.037944 | -1.39236 | #N/A | elongation zone | 6 h |
| HORVU.MOREX.r3.5HG0486750.1 | 0.734142 | 2.037011 | 3.384947 | 0.001347 | 0.030132 | -1.39395 | Phosphatase 2C family protein | elongation zone | 6 h |
| HORVU.MOREX.r3.7HG0638850.1 | -0.42458 | 7.446158 | -3.53269 | 0.000861 | 0.023177 | -1.3948 | Cellulose synthase-like protein | elongation zone | 6 h |
| HORVU.MOREX.r3.1HG0057780.1 | -0.40647 | 5.901425 | -3.45204 | 0.001101 | 0.026729 | -1.39646 | bZIP transcription factor (DUF630 and DUF632) | elongation zone | 6 h |
| HORVU.MOREX.r3.2HG0178160.1 | -0.56924 | 7.879945 | -3.5041 | 0.00094 | 0.024262 | -1.39871 | HTH-type transcriptional regulator YidZ | elongation zone | 6 h |
| HORVU.MOREX.r3.6HG0607360.1 | -0.55585 | 4.502873 | -3.34854 | 0.001501 | 0.031694 | -1.39955 | WAT1-related protein | elongation zone | 6 h |
| HORVU.MOREX.r3.4HG0417670.1 | 1.218021 | 0.032065 | 3.409555 | 0.001251 | 0.028873 | -1.40445 | Lipid transfer protein | elongation zone | 6 h |
| HORVU.MOREX.r3.3HG0279100.1 | 1.189175 | -1.86348 | 3.253361 | 0.001988 | 0.03752 | -1.40817 | Serine/threonine-protein kinase | elongation zone | 6 h |
| HORVU.MOREX.r3.4HG0337870.1 | 0.822274 | 4.45297 | 3.420684 | 0.00121 | 0.028241 | -1.4105 | Homeobox-leucine zipper family protein / lipid-binding START domain-containing | elongation zone | 6 h |
| HORVU.MOREX.r3.1HG0055320.1 | -0.27754 | 5.996621 | -3.45996 | 0.001075 | 0.026284 | -1.4147 | Protein phosphatase 2C | elongation zone | 6 h |
| HORVU.MOREX.r3.1HG0073170.1 | -0.61217 | 5.178736 | -3.36559 | 0.001427 | 0.031106 | -1.41553 | F-box protein | elongation zone | 6 h |
| HORVU.MOREX.r3.6HG0631660.1 | 0.692153 | 5.110572 | 3.387391 | 0.001337 | 0.030044 | -1.41608 | RING/U-box superfamily protein | elongation zone | 6 h |
| HORVU.MOREX.r3.3HG0330040.1 | -0.65876 | 2.332376 | -3.25861 | 0.001957 | 0.037085 | -1.4162 | Kelch repeat-containing F-box family protein | elongation zone | 6 h |
| HORVU.MOREX.r3.2HG0122880.1 | 1.452158 | 1.048128 | 3.242228 | 0.002053 | 0.038269 | -1.41718 | Phytoene desaturase | elongation zone | 6 h |
| HORVU.MOREX.r3.1HG0020270.1 | -0.37999 | 5.457119 | -3.39682 | 0.0013 | 0.029498 | -1.41768 | SAD1/UNC-84 domain protein, putative | elongation zone | 6 h |
| HORVU.MOREX.r3.5HG0528910.1 | 0.915646 | 1.611089 | 3.365978 | 0.001425 | 0.031106 | -1.41779 | DUF3511 domain protein | elongation zone | 6 h |
| HORVU.MOREX.r3.5HG0468580.1 | 1.417051 | -0.71714 | 3.307261 | 0.001696 | 0.03438 | -1.41789 | Cinnamoyl-CoA reductase | elongation zone | 6 h |
| HORVU.MOREX.r3.3HG0288070.1 | 0.565253 | 0.496177 | 3.395473 | 0.001305 | 0.029585 | -1.41861 | 2-oxoglutarate (2OG) and Fe(II)-dependent oxygenase superfamily protein, putativ | elongation zone | 6 h |
| HORVU.MOREX.r3.4HG0361680.1 | -0.87521 | 4.471436 | -3.33245 | 0.001574 | 0.032666 | -1.42034 | Nuclear distribution protein nudE-like 1 | elongation zone | 6 h |
| HORVU.MOREX.r3.5HG0420830.1 | 1.128876 | -1.8789 | 3.239579 | 0.002069 | 0.038427 | -1.42154 | cycling DOF factor 1 | elongation zone | 6 h |
| HORVU.MOREX.r3.3HG0234030.1 | 0.932613 | 0.965447 | 3.438866 | 0.001145 | 0.027233 | -1.42454 | Xyloglucan galactosyltransferase KATAMARI1-like protein | elongation zone | 6 h |
| HORVU.MOREX.r3.1HG0025430.1 | 1.272739 | 0.604962 | 3.235551 | 0.002094 | 0.038776 | -1.42521 | WEB family protein (DUF827) | elongation zone | 6 h |
| HORVU.MOREX.r3.7HG0648390.1 | 0.931378 | -0.60992 | 3.3019 | 0.001723 | 0.034545 | -1.4256 | Protein casein kinase I-like 4 | elongation zone | 6 h |
| HORVU.MOREX.r3.1HG0003090.7 | -1.19155 | 6.089425 | -3.36909 | 0.001412 | 0.030961 | -1.42645 | Thionin-2.2 | elongation zone | 6 h |
| HORVU.MOREX.r3.5HG0448080.1 | -0.55616 | 5.913862 | -3.48048 | 0.00101 | 0.025334 | -1.42749 | Polyol transporter | elongation zone | 6 h |

**Table S1** Continued.

| ID | logFC | AveExpr | t | P.Value | adj.P.Val | B | gene description | root zone | time point |
|---|---|---|---|---|---|---|---|---|---|
| HORVU.MOREX.r3.7HG0676040.1 | 0.657176 | -0.53383 | 3.301728 | 0.001724 | 0.034545 | -1.43062 | Decaprenyl-phosphate N-acetylglucosaminephosphotransferase | elongation zone | 6 h |
| HORVU.MOREX.r3.7HG0650250.1 | -0.51392 | 3.840838 | -3.32591 | 0.001605 | 0.033004 | -1.43066 | Protein-O-fucosyltransferase 1 | elongation zone | 6 h |
| HORVU.MOREX.r3.6HG0613460.1 | 1.160272 | 2.574467 | 3.317967 | 0.001643 | 0.033722 | -1.43285 | MACPF domain-containing protein | elongation zone | 6 h |
| HORVU.MOREX.r3.5HG0526200.1 | -0.27222 | 5.485729 | -3.4075 | 0.001259 | 0.028987 | -1.43675 | Sec14 cytosolic factor | elongation zone | 6 h |
| HORVU.MOREX.r3.7HG0639930.1 | -0.51382 | 3.436078 | -3.29629 | 0.001752 | 0.034787 | -1.43694 | Protein FAR1-RELATED SEQUENCE 5 | elongation zone | 6 h |
| HORVU.MOREX.r3.2HG0161580.1 | -0.55261 | 2.675596 | -3.26116 | 0.001943 | 0.03691 | -1.43772 | Alpha/beta-Hydrolases superfamily protein | elongation zone | 6 h |
| HORVU.MOREX.r3.5HG0526160.1 | -0.49719 | 4.252818 | -3.3639 | 0.001434 | 0.031117 | -1.44504 | NBS-LRR resistance-like protein | elongation zone | 6 h |
| HORVU.MOREX.r3.7HG0650310.1 | 1.34066 | 2.884324 | 3.219994 | 0.002191 | 0.039786 | -1.44519 | RNA-binding family protein | elongation zone | 6 h |
| HORVU.MOREX.r3.4HG0417710.1 | 0.957589 | 0.768626 | 3.419588 | 0.001214 | 0.028302 | -1.44627 | Protease inhibitor/seed storage/lipid transfer protein family protein | elongation zone | 6 h |
| HORVU.MOREX.r3.3HG0326330.1 | 0.989929 | 0.282326 | 3.393048 | 0.001314 | 0.029768 | -1.44669 | Rho GTPase-activating protein | elongation zone | 6 h |
| HORVU.MOREX.r3.2HG0152430.1 | -0.47782 | 6.139317 | -3.45067 | 0.001105 | 0.026745 | -1.44713 | Ubiquitin carboxyl-terminal hydrolase, putative | elongation zone | 6 h |
| HORVU.MOREX.r3.6HG0618180.1 | 1.443268 | -2.40297 | 3.221915 | 0.002179 | 0.039605 | -1.44815 | Universal stress protein family protein | elongation zone | 6 h |
| HORVU.MOREX.r3.4HG0347840.1 | 0.48229 | 3.393693 | 3.305944 | 0.001703 | 0.03438 | -1.44907 | Receptor-type tyrosine-protein phosphatase | elongation zone | 6 h |
| HORVU.MOREX.r3.2HG0121630.1 | 0.738785 | 3.093911 | 3.24361 | 0.002045 | 0.03815 | -1.45083 | Mitochondrial carrier family protein | elongation zone | 6 h |
| HORVU.MOREX.r3.5HG0511090.1 | 0.552882 | 5.583411 | 3.399218 | 0.00129 | 0.029352 | -1.45167 | Phytochrome | elongation zone | 6 h |
| HORVU.MOREX.r3.2HG0117880.1 | 2.617431 | -0.66123 | 3.231312 | 0.00212 | 0.039083 | -1.46232 | GDSL esterase/lipase | elongation zone | 6 h |
| HORVU.MOREX.r3.3HG0218560.1 | 1.025919 | 0.309747 | 3.287218 | 0.001799 | 0.035369 | -1.46421 | Receptor-like protein kinase | elongation zone | 6 h |
| HORVU.MOREX.r3.4HG0406600.1 | 0.744867 | 0.071867 | 3.384001 | 0.00135 | 0.03018 | -1.46458 | Heavy metal transport/detoxification superfamily protein | elongation zone | 6 h |
| HORVU.MOREX.r3.4HG0400990.1 | -0.52699 | 5.330008 | -3.39018 | 0.001326 | 0.029926 | -1.46468 | 3-hydroxybenzoate 6-hydroxylase 1 | elongation zone | 6 h |
| HORVU.MOREX.r3.7HG0716640.1 | 0.831281 | 2.750789 | 3.478107 | 0.001017 | 0.025374 | -1.465 | Xyloglucan endotransglucosylase/hydrolase | elongation zone | 6 h |
| HORVU.MOREX.r3.5HG0485410.1 | 2.23351 | -0.44467 | 3.440032 | 0.001141 | 0.027169 | -1.46659 | #N/A | elongation zone | 6 h |
| HORVU.MOREX.r3.5HG0441420.1 | 2.301652 | -0.42861 | 3.33893 | 0.001544 | 0.032239 | -1.46695 | Ankyrin repeat-containing protein, putative | elongation zone | 6 h |
| HORVU.MOREX.r3.3HG0309460.1 | -0.28496 | 5.431182 | -3.41381 | 0.001235 | 0.028635 | -1.46731 | Trichome birefringence-like protein | elongation zone | 6 h |
| HORVU.MOREX.r3.7HG0688730.1 | 0.515643 | 4.635127 | 3.339143 | 0.001543 | 0.032239 | -1.47174 | Myotubularin | elongation zone | 6 h |
| HORVU.MOREX.r3.1HG0082770.1 | -0.28673 | 7.702552 | -3.48415 | 0.000999 | 0.025239 | -1.47447 | Reticulon-like protein | elongation zone | 6 h |
| HORVU.MOREX.r3.6HG0552230.1 | -0.52885 | 6.698601 | -3.44735 | 0.001116 | 0.026982 | -1.47521 | Polyubiquitin | elongation zone | 6 h |
| HORVU.MOREX.r3.5HG0524800.1 | -1.37206 | 1.969957 | -3.20739 | 0.002273 | 0.040716 | -1.47557 | RNA polymerase II transcription mediator | elongation zone | 6 h |
| HORVU.MOREX.r3.3HG0264440.1 | -0.65319 | 2.161098 | -3.22657 | 0.002149 | 0.039274 | -1.47801 | Condensin complex subunit 2 | elongation zone | 6 h |
| HORVU.MOREX.r3.2HG0145360.1 | 0.99074 | 3.483304 | 3.521457 | 0.000892 | 0.023455 | -1.47842 | Peroxidase | elongation zone | 6 h |
| HORVU.MOREX.r3.7HG0731320.1 | -0.34152 | 5.022738 | -3.39779 | 0.001296 | 0.029445 | -1.48169 | O-fucosyltransferase family protein | elongation zone | 6 h |
| HORVU.MOREX.r3.7HG0659040.1 | 0.83727 | 3.074402 | 3.25216 | 0.001995 | 0.037583 | -1.48313 | Formin-like protein | elongation zone | 6 h |
| HORVU.MOREX.r3.7HG0683600.1 | 1.85156 | 0.200809 | 3.222443 | 0.002175 | 0.039605 | -1.48589 | F-box protein | elongation zone | 6 h |
| HORVU.MOREX.r3.5HG0470170.1 | -0.52632 | 3.126834 | -3.2699 | 0.001893 | 0.036511 | -1.48661 | F-box family protein | elongation zone | 6 h |
| HORVU.MOREX.r3.1HG0055590.1 | 1.461738 | -0.98036 | 3.207102 | 0.002275 | 0.040716 | -1.48807 | RING/U-box superfamily protein | elongation zone | 6 h |
| HORVU.MOREX.r3.4HG0389250.1 | 1.309859 | 3.018686 | 3.256512 | 0.001969 | 0.037279 | -1.49206 | lipase, putative (DUF620) | elongation zone | 6 h |
| HORVU.MOREX.r3.6HG0607590.1 | 1.035755 | 1.682477 | 3.229268 | 0.002133 | 0.039141 | -1.49349 | U-box domain-containing family protein | elongation zone | 6 h |
| HORVU.MOREX.r3.4HG0335160.1 | 0.801062 | 3.629312 | 3.312649 | 0.001669 | 0.034187 | -1.49638 | Hedgehog-interacting-like protein | elongation zone | 6 h |
| HORVU.MOREX.r3.3HG0302340.1 | -2.74254 | -0.09437 | -3.34737 | 0.001506 | 0.031731 | -1.49728 | Eukaryotic aspartyl protease family protein, expressed | elongation zone | 6 h |
| HORVU.MOREX.r3.7HG0705500.1 | 1.092654 | 0.351075 | 3.19715 | 0.002341 | 0.041422 | -1.50384 | Non-lysosomal glucosylceramidase | elongation zone | 6 h |
| HORVU.MOREX.r3.2HG0188490.1 | -0.80805 | 2.129721 | -3.22186 | 0.002179 | 0.039605 | -1.51267 | No-apical-meristem-associated carboxy-terminal domain protein | elongation zone | 6 h |
| HORVU.MOREX.r3.7HG0737470.1 | 1.027893 | 3.292476 | 3.553377 | 0.000809 | 0.022568 | -1.51293 | Peroxidase | elongation zone | 6 h |

**Table S1** Continued.

| ID | logFC | AveExpr | t | P.Value | adj.P.Val | B | gene description | root zone | time point |
|---|---|---|---|---|---|---|---|---|---|
| HORVU.MOREX.r3.3HG0296990.1 | 0.924845 | -1.49078 | 3.224981 | 0.002159 | 0.039386 | -1.51338 | LURP-one-like protein | elongation zone | 6 h |
| HORVU.MOREX.r3.5HG0498240.1 | 2.677685 | -1.97942 | 3.2285 | 0.002137 | 0.039149 | -1.51359 | Cytochrome P450, putative | elongation zone | 6 h |
| HORVU.MOREX.r3.5HG0439190.1 | -0.50769 | 2.950971 | -3.31166 | 0.001674 | 0.034187 | -1.5165 | Cytochrome P450 | elongation zone | 6 h |
| HORVU.MOREX.r3.6HG0627570.1 | -0.53916 | 4.766957 | -3.34614 | 0.001512 | 0.031783 | -1.51679 | Immunoglobulin A1 protease autotransporter | elongation zone | 6 h |
| HORVU.MOREX.r3.7HG0668820.1 | 0.563417 | 4.620228 | 3.468103 | 0.001049 | 0.025891 | -1.52697 | Endoglucanase | elongation zone | 6 h |
| HORVU.MOREX.r3.7HG0720790.1 | -0.87663 | 3.44449 | -3.21698 | 0.00221 | 0.040038 | -1.53092 | Myb family transcription factor family protein | elongation zone | 6 h |
| HORVU.MOREX.r3.7HG0738680.1 | 0.700665 | 3.55287 | 3.359412 | 0.001453 | 0.031376 | -1.53145 | Vacuolar sorting receptor family protein | elongation zone | 6 h |
| HORVU.MOREX.r3.2HG0132160.1 | 2.326143 | -1.56452 | 3.29579 | 0.001755 | 0.034787 | -1.53285 | Ethylene-responsive transcription factor | elongation zone | 6 h |
| HORVU.MOREX.r3.4HG0408780.1 | 2.085153 | 0.649042 | 3.362806 | 0.001439 | 0.031171 | -1.53523 | Pentatricopeptide repeat-containing protein | elongation zone | 6 h |
| HORVU.MOREX.r3.7HG0674990.1 | -0.46509 | 3.286295 | -3.27395 | 0.001871 | 0.036221 | -1.53723 | Mannan endo-1,4-beta-mannosidase-like protein | elongation zone | 6 h |
| HORVU.MOREX.r3.6HG0569610.1 | -0.72343 | 5.321736 | -3.41496 | 0.001231 | 0.028622 | -1.54425 | #N/A | elongation zone | 6 h |
| HORVU.MOREX.r3.4HG0415590.1 | -1.43555 | -0.04239 | -3.20186 | 0.00231 | 0.041032 | -1.5512 | Bifunctional inhibitor/lipid-transfer protein/seed storage 2S albumin superfamily p | elongation zone | 6 h |
| HORVU.MOREX.r3.7HG0674320.1 | -0.36868 | 4.709886 | -3.3693 | 0.001411 | 0.030961 | -1.55234 | Vesicle transport protein | elongation zone | 6 h |
| HORVU.MOREX.r3.4HG0399530.1 | 1.75006 | 1.156384 | 3.180918 | 0.002454 | 0.042679 | -1.55264 | Zinc finger protein | elongation zone | 6 h |
| HORVU.MOREX.r3.7HG0667260.1 | 0.801391 | 2.036947 | 3.241502 | 0.002058 | 0.038281 | -1.55386 | Trichome birefringence-like protein | elongation zone | 6 h |
| HORVU.MOREX.r3.3HG0219750.1 | -1.73555 | -0.71619 | -3.20495 | 0.002289 | 0.040793 | -1.55811 | Disease resistance protein (NBS-LRR class) family | elongation zone | 6 h |
| HORVU.MOREX.r3.4HG0345610.1 | -0.28595 | 8.214926 | -3.49056 | 0.000979 | 0.024844 | -1.55951 | Protein transport protein Sec24-like family | elongation zone | 6 h |
| HORVU.MOREX.r3.3HG0294390.1 | 0.716963 | -1.06639 | 3.212456 | 0.00224 | 0.040418 | -1.5635 | Pectinesterase | elongation zone | 6 h |
| HORVU.MOREX.r3.1HG0045770.1 | 1.745873 | -2.26851 | 3.180885 | 0.002454 | 0.042679 | -1.57286 | GRAM domain-containing protein / ABA-responsive protein-related | elongation zone | 6 h |
| HORVU.MOREX.r3.1HG0053450.1 | 0.857729 | 3.313219 | 3.444959 | 0.001124 | 0.027022 | -1.57323 | WEB family protein, chloroplastic | elongation zone | 6 h |
| HORVU.MOREX.r3.5HG0459600.1 | 0.671997 | 3.96473 | 3.348872 | 0.001499 | 0.031694 | -1.57442 | Serine/threonine-protein kinase | elongation zone | 6 h |
| HORVU.MOREX.r3.7HG0679760.1 | -0.47293 | 6.056962 | -3.38543 | 0.001345 | 0.030122 | -1.57527 | Basic helix-loop-helix transcription factor | elongation zone | 6 h |
| HORVU.MOREX.r3.4HG0381620.1 | 1.270875 | -2.76226 | 3.165465 | 0.002566 | 0.044285 | -1.57728 | Transcription factor, putative | elongation zone | 6 h |
| HORVU.MOREX.r3.2HG0160700.1 | 2.049175 | -0.50356 | 3.210289 | 0.002254 | 0.040566 | -1.5801 | L-threonine 3-dehydrogenase | elongation zone | 6 h |
| HORVU.MOREX.r3.2HG0151790.1 | 2.587146 | -1.72838 | 3.400937 | 0.001284 | 0.02931 | -1.58237 | Ankyrin repeat-containing protein | elongation zone | 6 h |
| HORVU.MOREX.r3.5HG0495860.1 | 0.777665 | 4.058331 | 3.351854 | 0.001486 | 0.031669 | -1.58429 | glycosyltransferase family exostosin protein | elongation zone | 6 h |
| HORVU.MOREX.r3.3HG0219410.1 | 0.753885 | 3.869903 | 3.330397 | 0.001584 | 0.032759 | -1.58707 | Hydroxyproline-rich glycoprotein family protein | elongation zone | 6 h |
| HORVU.MOREX.r3.3HG0307400.1 | 1.254974 | 1.770343 | 3.206033 | 0.002282 | 0.040785 | -1.59143 | GRAS family transcription factor | elongation zone | 6 h |
| HORVU.MOREX.r3.6HG0617730.1 | -0.33983 | 7.183954 | -3.45838 | 0.00108 | 0.026331 | -1.59159 | Galactoside 2-alpha-L-fucosyltransferase | elongation zone | 6 h |
| HORVU.MOREX.r3.1HG0083550.1 | 2.808128 | 0.710912 | 3.348935 | 0.001499 | 0.031694 | -1.59392 | Pentatricopeptide repeat-containing protein | elongation zone | 6 h |
| HORVU.MOREX.r3.7HG0751060.1 | -0.3863 | 4.948458 | -3.3515 | 0.001488 | 0.031669 | -1.59426 | Disease resistance protein (NBS-LRR class) family | elongation zone | 6 h |
| HORVU.MOREX.r3.6HG0581020.1 | -0.32994 | 5.648771 | -3.37037 | 0.001406 | 0.030961 | -1.5956 | Xyloglucan 6-xylosyltransferase 1 | elongation zone | 6 h |
| HORVU.MOREX.r3.1HG0047570.1 | 0.705153 | 2.998289 | 3.278716 | 0.001845 | 0.035889 | -1.59895 | Di-glucose binding protein with Kinesin motor domain-containing protein | elongation zone | 6 h |
| HORVU.MOREX.r3.7HG0677510.1 | -0.43265 | 5.971824 | -3.37803 | 0.001375 | 0.030595 | -1.60079 | Glycosyltransferase | elongation zone | 6 h |
| HORVU.MOREX.r3.2HG0207790.1 | 1.182283 | -2.59422 | 3.154097 | 0.002651 | 0.045332 | -1.60406 | Serine/threonine-protein kinase | elongation zone | 6 h |
| HORVU.MOREX.r3.5HG0476690.1 | -0.3216 | 5.143097 | -3.33998 | 0.00154 | 0.032205 | -1.60892 | #N/A | elongation zone | 6 h |
| HORVU.MOREX.r3.2HG0096520.1 | 1.579066 | 0.617281 | 3.160341 | 0.002604 | 0.044719 | -1.60904 | Pentatricopeptide repeat-containing protein | elongation zone | 6 h |
| HORVU.MOREX.r3.7HG0652820.1 | -2.25153 | -0.08916 | -3.28557 | 0.001808 | 0.035474 | -1.60999 | Disease resistance protein (NBS-LRR class) family | elongation zone | 6 h |
| HORVU.MOREX.r3.6HG0558430.1 | 2.253591 | -1.96916 | 3.153871 | 0.002653 | 0.045332 | -1.61194 | SAUR-like auxin-responsive protein family, putative | elongation zone | 6 h |
| HORVU.MOREX.r3.1HG0095170.1 | 1.583626 | 5.086192 | 3.298625 | 0.00174 | 0.03465 | -1.61387 | UDP-glucose-4-epimerase | elongation zone | 6 h |
| HORVU.MOREX.r3.7HG0742370.1 | 1.338747 | -1.39541 | 3.152168 | 0.002666 | 0.045479 | -1.6143 | Calmodulin | elongation zone | 6 h |

**Table S1** Continued.

| ID | logFC | AveExpr | t | P.Value | adj.P.Val | B | gene description | root zone | time point |
|---|---|---|---|---|---|---|---|---|---|
| HORVU.MOREX.r3.2HG0159540.1 | -0.82915 | 3.353367 | -3.22968 | 0.00213 | 0.039141 | -1.6157 | Glycosyltransferase | elongation zone | 6 h |
| HORVU.MOREX.r3.7HG0728060.1 | 0.478751 | 5.512337 | 3.348454 | 0.001501 | 0.031694 | -1.6202 | Aminodeoxychorismate synthase | elongation zone | 6 h |
| HORVU.MOREX.r3.3HG0307310.1 | -0.30788 | 5.321617 | -3.34904 | 0.001499 | 0.031694 | -1.62717 | magnesium transporter, putative (DUF803) | elongation zone | 6 h |
| HORVU.MOREX.r3.5HG0524120.1 | -0.36691 | 5.963425 | -3.3774 | 0.001377 | 0.030619 | -1.62904 | #N/A | elongation zone | 6 h |
| HORVU.MOREX.r3.2HG0204940.1 | -0.49608 | 6.058217 | -3.34 | 0.00154 | 0.032205 | -1.6315 | Crossover junction endonuclease EME1 | elongation zone | 6 h |
| HORVU.MOREX.r3.3HG0295330.1 | -0.40556 | 7.613979 | -3.46125 | 0.00107 | 0.026245 | -1.63226 | Sugar phosphate/phosphate translocator | elongation zone | 6 h |
| HORVU.MOREX.r3.3HG0288940.1 | -0.48121 | 6.05706 | -3.36492 | 0.001429 | 0.031106 | -1.63319 | WRKY transcription factor, putative, expressed | elongation zone | 6 h |
| HORVU.MOREX.r3.1HG0056510.1 | -1.46064 | 0.442591 | -3.17049 | 0.002529 | 0.043758 | -1.63686 | Cytochrome P450, putative | elongation zone | 6 h |
| HORVU.MOREX.r3.6HG0606360.1 | 2.08108 | -2.06608 | 3.143467 | 0.002734 | 0.046171 | -1.63714 | Peptide transporter | elongation zone | 6 h |
| HORVU.MOREX.r3.7HG0687630.1 | -0.53741 | 7.969317 | -3.43782 | 0.001149 | 0.027274 | -1.64063 | Acyl carrier protein | elongation zone | 6 h |
| HORVU.MOREX.r3.1HG0017770.1 | 0.751788 | 2.765713 | 3.302164 | 0.001722 | 0.034545 | -1.64412 | Germin-like protein 1 | elongation zone | 6 h |
| HORVU.MOREX.r3.5HG0525430.1 | 0.708665 | 4.268757 | 3.385954 | 0.001343 | 0.030107 | -1.64467 | Trichome birefringence-like protein | elongation zone | 6 h |
| HORVU.MOREX.r3.7HG0708080.1 | 1.686414 | 0.105537 | 3.149528 | 0.002687 | 0.045781 | -1.64475 | Molybdate transporter 1 | elongation zone | 6 h |
| HORVU.MOREX.r3.3HG0313320.1 | 1.501797 | -2.45652 | 3.148285 | 0.002696 | 0.045838 | -1.64695 | Potassium transporter | elongation zone | 6 h |
| HORVU.MOREX.r3.3HG0257310.1 | -0.27332 | 7.693219 | -3.43252 | 0.001167 | 0.027599 | -1.64825 | Glucosidase 2 subunit beta | elongation zone | 6 h |
| HORVU.MOREX.r3.3HG0315130.1 | -1.20057 | 1.022238 | -3.16227 | 0.00259 | 0.044578 | -1.64855 | RING/U-box superfamily protein, putative | elongation zone | 6 h |
| HORVU.MOREX.r3.7HG0735210.1 | 1.506838 | -3.00596 | 3.173506 | 0.002507 | 0.043525 | -1.65015 | Peptide transporter | elongation zone | 6 h |
| HORVU.MOREX.r3.4HG0391730.1 | -0.31715 | 6.58587 | -3.38974 | 0.001327 | 0.029931 | -1.65606 | UDP-glucuronate 4-epimerase 4 | elongation zone | 6 h |
| HORVU.MOREX.r3.5HG0432950.1 | 0.421233 | 5.320519 | 3.327414 | 0.001598 | 0.032894 | -1.66302 | L-galactose dehydrogenase | elongation zone | 6 h |
| HORVU.MOREX.r3.1HG0051960.1 | -1.091 | 1.903866 | -3.13208 | 0.002825 | 0.047264 | -1.66367 | DNA topoisomerase 2 | elongation zone | 6 h |
| HORVU.MOREX.r3.6HG0607160.1 | 0.758549 | 2.134391 | 3.265697 | 0.001917 | 0.036694 | -1.67238 | chitin synthase, putative (DUF1218) | elongation zone | 6 h |
| HORVU.MOREX.r3.1HG0073430.1 | 0.422695 | 3.344559 | 3.232771 | 0.002111 | 0.038952 | -1.67374 | O-fucosyltransferase family protein | elongation zone | 6 h |
| HORVU.MOREX.r3.5HG0522130.1 | -0.48394 | 4.373396 | -3.28454 | 0.001814 | 0.035513 | -1.67841 | #N/A | elongation zone | 6 h |
| HORVU.MOREX.r3.5HG0473920.1 | -0.46191 | 3.263199 | -3.22935 | 0.002132 | 0.039141 | -1.67891 | OBP32pep, putative (DUF220) | elongation zone | 6 h |
| HORVU.MOREX.r3.4HG0344550.1 | 1.692325 | 1.167724 | 3.123978 | 0.002891 | 0.047996 | -1.68072 | UPF0496 protein 4 | elongation zone | 6 h |
| HORVU.MOREX.r3.6HG0571750.1 | 1.142623 | 0.529808 | 3.125012 | 0.002882 | 0.047932 | -1.68478 | #N/A | elongation zone | 6 h |
| HORVU.MOREX.r3.5HG0463550.1 | 0.847008 | 4.670129 | 3.230927 | 0.002122 | 0.039091 | -1.68583 | RNA-binding domain CCCH-type zinc finger protein | elongation zone | 6 h |
| HORVU.MOREX.r3.6HG0597620.1 | -0.3814 | 5.241895 | -3.33561 | 0.00156 | 0.032443 | -1.68609 | #N/A | elongation zone | 6 h |
| HORVU.MOREX.r3.7HG0752940.1 | -0.32484 | 6.359204 | -3.34932 | 0.001497 | 0.031694 | -1.68728 | RING/FYVE/PHD zinc finger protein | elongation zone | 6 h |
| HORVU.MOREX.r3.2HG0210650.1 | 1.725412 | 0.042671 | 3.122555 | 0.002903 | 0.048115 | -1.68841 | BTB/POZ domain-containing protein | elongation zone | 6 h |
| HORVU.MOREX.r3.2HG0121200.1 | 0.743407 | 0.643147 | 3.306121 | 0.001702 | 0.03438 | -1.68962 | Dirigent protein | elongation zone | 6 h |
| HORVU.MOREX.r3.7HG0693590.1 | -0.40945 | 4.705719 | -3.29854 | 0.00174 | 0.03465 | -1.69015 | Exocyst complex component, putative | elongation zone | 6 h |
| HORVU.MOREX.r3.3HG0325670.1 | -0.62185 | 3.462193 | -3.17655 | 0.002485 | 0.04318 | -1.69568 | #N/A | elongation zone | 6 h |
| HORVU.MOREX.r3.6HG0572960.1 | -1.03781 | 1.634383 | -3.11559 | 0.002961 | 0.048729 | -1.69655 | Chlorophyll a-b binding protein, chloroplastic | elongation zone | 6 h |
| HORVU.MOREX.r3.3HG0239980.1 | 1.044211 | 1.292154 | 3.247599 | 0.002021 | 0.037914 | -1.6971 | Alpha/beta-Hydrolases superfamily protein, putative | elongation zone | 6 h |
| HORVU.MOREX.r3.1HG0042120.1 | -0.47008 | 4.166727 | -3.2458 | 0.002032 | 0.03801 | -1.69808 | WD repeat protein, putative | elongation zone | 6 h |
| HORVU.MOREX.r3.6HG0597770.1 | -0.46077 | 4.348024 | -3.24712 | 0.002024 | 0.037933 | -1.69842 | F-box/LRR protein | elongation zone | 6 h |
| HORVU.MOREX.r3.2HG0119800.1 | 1.49005 | -1.32815 | 3.113822 | 0.002976 | 0.04885 | -1.70002 | GDSL esterase/lipase | elongation zone | 6 h |
| HORVU.MOREX.r3.7HG0737890.1 | 1.01253 | 1.23238 | 3.371189 | 0.001403 | 0.030961 | -1.70131 | Peroxidase | elongation zone | 6 h |
| HORVU.MOREX.r3.1HG0073460.1 | 0.458291 | 2.975475 | 3.215736 | 0.002218 | 0.040113 | -1.70142 | Microneme/rhoptry antigen | elongation zone | 6 h |
| HORVU.MOREX.r3.3HG0253450.1 | -0.369 | 5.357439 | -3.30846 | 0.00169 | 0.034363 | -1.70681 | FHA domain containing protein, expressed | elongation zone | 6 h |

**Table S1** Continued.

| ID | logFC | AveExpr | t | P.Value | adj.P.Val | B | gene description | root zone | time point |
|---|---|---|---|---|---|---|---|---|---|
| HORVU.MOREX.r3.3HG0302230.1 | 0.611238 | 4.001482 | 3.306541 | 0.0017 | 0.03438 | -1.70687 | Avr9/Cf-9 rapidly elicited protein 137 | elongation zone | 6 h |
| HORVU.MOREX.r3.1HG0079920.1 | 0.617331 | 3.500385 | 3.299659 | 0.001735 | 0.03465 | -1.71035 | Disease resistance protein (NBS-LRR class) family | elongation zone | 6 h |
| HORVU.MOREX.r3.1HG0069210.1 | -0.70361 | 2.90701 | -3.21437 | 0.002227 | 0.040229 | -1.71171 | LOW protein: E3 ubiquitin ligase-like protein | elongation zone | 6 h |
| HORVU.MOREX.r3.5HG0481580.1 | 0.763379 | 1.426704 | 3.268229 | 0.001903 | 0.036589 | -1.71246 | Hexosyltransferase | elongation zone | 6 h |
| HORVU.MOREX.r3.2HG0109580.1 | -1.39181 | -0.60672 | -3.11359 | 0.002978 | 0.04885 | -1.71271 | Liprin-alpha-3 | elongation zone | 6 h |
| HORVU.MOREX.r3.5HG0532960.1 | -0.40744 | 5.01578 | -3.29445 | 0.001762 | 0.03479 | -1.71704 | ER lumen protein-retaining receptor | elongation zone | 6 h |
| HORVU.MOREX.r3.2HG0213840.1 | 1.037211 | -0.58232 | 3.191846 | 0.002378 | 0.041914 | -1.72036 | transcription repressor | elongation zone | 6 h |
| HORVU.MOREX.r3.4HG0406140.1 | 0.462274 | 5.424725 | 3.317398 | 0.001646 | 0.033745 | -1.7221 | Transmembrane protein 131 | elongation zone | 6 h |
| HORVU.MOREX.r3.7HG0718190.1 | 0.690847 | 6.009061 | 3.426105 | 0.00119 | 0.027891 | -1.72223 | ABC transporter family protein | elongation zone | 6 h |
| HORVU.MOREX.r3.5HG0420630.1 | -0.48088 | 3.720927 | -3.24786 | 0.00202 | 0.037914 | -1.72394 | Ankyrin repeat family protein | elongation zone | 6 h |
| HORVU.MOREX.r3.3HG0256400.1 | -0.49994 | 3.232954 | -3.19112 | 0.002383 | 0.04193 | -1.72736 | rRNA N-glycosidase | elongation zone | 6 h |
| HORVU.MOREX.r3.7HG0714400.1 | 0.87393 | 2.702391 | 3.1813 | 0.002451 | 0.042679 | -1.73113 | Auxin-responsive protein | elongation zone | 6 h |
| HORVU.MOREX.r3.5HG0499000.1 | 2.121898 | -0.43495 | 3.225887 | 0.002154 | 0.039317 | -1.73274 | Purple acid phosphatase | elongation zone | 6 h |
| HORVU.MOREX.r3.7HG0636540.1 | 1.968688 | 1.811078 | 3.12362 | 0.002894 | 0.048007 | -1.73411 | Pectin lyase-like superfamily protein | elongation zone | 6 h |
| HORVU.MOREX.r3.3HG0224440.1 | 0.810164 | 1.43625 | 3.262086 | 0.001937 | 0.036879 | -1.73437 | DUF4228 domain protein | elongation zone | 6 h |
| HORVU.MOREX.r3.6HG0540020.1 | 1.54027 | -2.7011 | 3.11184 | 0.002993 | 0.048862 | -1.73508 | Ubiquitin carboxyl-terminal hydrolase 2 | elongation zone | 6 h |
| HORVU.MOREX.r3.7HG0704060.1 | 0.555967 | 4.974659 | 3.205005 | 0.002289 | 0.040793 | -1.73709 | AT hook motif DNA-binding family protein | elongation zone | 6 h |
| HORVU.MOREX.r3.2HG0193490.1 | 1.41193 | -1.27815 | 3.102071 | 0.003078 | 0.049581 | -1.73712 | Growth-regulating factor | elongation zone | 6 h |
| HORVU.MOREX.r3.4HG0352760.1 | -0.34431 | 5.845969 | -3.32978 | 0.001587 | 0.032759 | -1.74084 | GPI transamidase component PIG-T | elongation zone | 6 h |
| HORVU.MOREX.r3.6HG0573290.1 | 0.315378 | 6.719302 | 3.365131 | 0.001429 | 0.031106 | -1.744 | Zinc finger A20 and AN1 domain stress-associated protein | elongation zone | 6 h |
| HORVU.MOREX.r3.2HG0156810.1 | -1.60509 | 1.309434 | -3.12132 | 0.002913 | 0.048242 | -1.74517 | Sodium/potassium/calcium exchanger 3 | elongation zone | 6 h |
| HORVU.MOREX.r3.2HG0196940.1 | -0.3414 | 4.564537 | -3.27171 | 0.001883 | 0.036357 | -1.74837 | Alpha-1,3-mannosyltransferase, putative | elongation zone | 6 h |
| HORVU.MOREX.r3.6HG0609390.1 | 0.72323 | 5.890658 | 3.239785 | 0.002068 | 0.038427 | -1.75154 | Ribose-phosphate pyrophosphokinase, putative | elongation zone | 6 h |
| HORVU.MOREX.r3.5HG0425880.1 | -0.54849 | 2.819479 | -3.15506 | 0.002644 | 0.045269 | -1.7542 | cDNA clone:J013058P10, full insert sequence | elongation zone | 6 h |
| HORVU.MOREX.r3.2HG0162300.1 | -0.43156 | 3.631707 | -3.28496 | 0.001811 | 0.035503 | -1.75461 | Intracellular protein transporter USO1-like protein | elongation zone | 6 h |
| HORVU.MOREX.r3.4HG0410820.1 | 1.271354 | 3.960298 | 3.097813 | 0.003115 | 0.049823 | -1.76431 | SMAD/FHA domain-containing family protein | elongation zone | 6 h |
| HORVU.MOREX.r3.7HG0637760.1 | -0.42383 | 3.556847 | -3.19914 | 0.002328 | 0.041321 | -1.76811 | #N/A | elongation zone | 6 h |
| HORVU.MOREX.r3.7HG0682460.1 | -0.35679 | 5.61607 | -3.31115 | 0.001677 | 0.034204 | -1.77171 | ADP-ribosylation factor, putative | elongation zone | 6 h |
| HORVU.MOREX.r3.4HG0394830.1 | 0.824201 | 0.997429 | 3.171291 | 0.002523 | 0.043737 | -1.77635 | Lecithin-cholesterol acyltransferase-like 1 | elongation zone | 6 h |
| HORVU.MOREX.r3.2HG0196520.1 | 0.736155 | 4.252659 | 3.26772 | 0.001906 | 0.036589 | -1.77792 | senescence-associated family protein (DUF581) | elongation zone | 6 h |
| HORVU.MOREX.r3.1HG0079750.1 | 1.009409 | -1.15538 | 3.135125 | 0.0028 | 0.046943 | -1.77811 | MYB transcription factor | elongation zone | 6 h |
| HORVU.MOREX.r3.3HG0249430.1 | -2.4815 | -0.56924 | -3.26405 | 0.001926 | 0.036837 | -1.7817 | Protein kinase | elongation zone | 6 h |
| HORVU.MOREX.r3.7HG0658560.1 | 0.739428 | 3.664515 | 3.143569 | 0.002733 | 0.046171 | -1.78172 | Kinase family protein | elongation zone | 6 h |
| HORVU.MOREX.r3.5HG0493380.1 | -0.32008 | 4.391009 | -3.25104 | 0.002001 | 0.03764 | -1.7826 | Cyclin family protein | elongation zone | 6 h |
| HORVU.MOREX.r3.3HG0316640.1 | -0.64098 | 4.162627 | -3.28315 | 0.001821 | 0.035591 | -1.78488 | Pentatricopeptide repeat (PPR) superfamily protein | elongation zone | 6 h |
| HORVU.MOREX.r3.1HG0002340.1 | -0.30519 | 6.091944 | -3.30332 | 0.001716 | 0.034524 | -1.78669 | Tubulin-specific chaperone cofactor E-like protein | elongation zone | 6 h |
| HORVU.MOREX.r3.5HG0497490.1 | 1.347614 | -1.08137 | 3.14929 | 0.002688 | 0.045781 | -1.79059 | SPX domain-like protein | elongation zone | 6 h |
| HORVU.MOREX.r3.3HG0309900.1 | 1.148609 | -1.42396 | 3.11083 | 0.003002 | 0.048898 | -1.79141 | ROP-interactive CRIB motif protein | elongation zone | 6 h |
| HORVU.MOREX.r3.1HG0039670.1 | -0.23814 | 7.408425 | -3.36879 | 0.001413 | 0.030961 | -1.79147 | Calcium ion-binding protein | elongation zone | 6 h |
| HORVU.MOREX.r3.2HG0128240.1 | -0.40735 | 5.300426 | -3.26963 | 0.001895 | 0.036511 | -1.79227 | Aldose 1-epimerase | elongation zone | 6 h |
| HORVU.MOREX.r3.6HG0602230.1 | -0.51499 | 5.050411 | -3.26216 | 0.001937 | 0.036879 | -1.79255 | Protein kinase | elongation zone | 6 h |

**Table S1** Continued.

| ID | logFC | AveExpr | t | P.Value | adj.P.Val | B | gene description | root zone | time point |
|---|---|---|---|---|---|---|---|---|---|
| HORVU.MOREX.r3.3HG0250940.1 | -0.60542 | 3.302703 | -3.18121 | 0.002452 | 0.042679 | -1.79257 | Polynucleotidyl transferase ribonuclease H-like superfamily protein | elongation zone | 6 h |
| HORVU.MOREX.r3.4HG0344830.1 | -0.44922 | 4.535568 | -3.23382 | 0.002104 | 0.038877 | -1.79345 | NBS-LRR disease resistance protein, putative, expressed | elongation zone | 6 h |
| HORVU.MOREX.r3.1HG0090100.1 | 1.254066 | 1.392953 | 3.103352 | 0.003067 | 0.049561 | -1.79688 | DUF1645 family protein | elongation zone | 6 h |
| HORVU.MOREX.r3.3HG0243130.1 | -0.75678 | 2.631713 | -3.15594 | 0.002637 | 0.045214 | -1.80082 | Peroxidase | elongation zone | 6 h |
| HORVU.MOREX.r3.2HG0208960.1 | -0.8505 | 2.446722 | -3.0987 | 0.003108 | 0.049818 | -1.80373 | Exocyst complex component EXO70B1 | elongation zone | 6 h |
| HORVU.MOREX.r3.2HG0165840.1 | -0.55924 | 6.204491 | -3.26593 | 0.001916 | 0.036694 | -1.80573 | Protein kinase | elongation zone | 6 h |
| HORVU.MOREX.r3.2HG0117980.1 | -0.35959 | 6.654309 | -3.32984 | 0.001587 | 0.032759 | -1.80714 | Transmembrane protein 115 | elongation zone | 6 h |
| HORVU.MOREX.r3.5HG0531390.1 | -0.29748 | 6.128355 | -3.32464 | 0.001611 | 0.033096 | -1.80992 | Kinase family protein | elongation zone | 6 h |
| HORVU.MOREX.r3.5HG0466510.1 | -0.96959 | 3.504705 | -3.10606 | 0.003043 | 0.049244 | -1.81025 | Retrovirus-related Pol polyprotein from transposon TNT 1-94 | elongation zone | 6 h |
| HORVU.MOREX.r3.2HG0153420.1 | 0.741351 | 6.452276 | 3.352372 | 0.001484 | 0.031653 | -1.81087 | #N/A | elongation zone | 6 h |
| HORVU.MOREX.r3.2HG0189010.1 | 1.011391 | 3.632309 | 3.197106 | 0.002342 | 0.041422 | -1.81098 | Glycerol-3-phosphate transporter, putative | elongation zone | 6 h |
| HORVU.MOREX.r3.3HG0277100.1 | -0.41939 | 4.350493 | -3.22962 | 0.00213 | 0.039141 | -1.81281 | Calcium-dependent lipid-binding (CaLB domain) family protein | elongation zone | 6 h |
| HORVU.MOREX.r3.1HG0081220.1 | -0.41721 | 5.709055 | -3.29222 | 0.001773 | 0.034953 | -1.81452 | Galactose-6-phosphate isomerase subunit LacB | elongation zone | 6 h |
| HORVU.MOREX.r3.1HG0051860.1 | -0.64618 | 5.340638 | -3.2371 | 0.002084 | 0.038637 | -1.81616 | Glutathione S-transferase | elongation zone | 6 h |
| HORVU.MOREX.r3.2HG0169700.1 | -0.42524 | 4.388016 | -3.23374 | 0.002105 | 0.038877 | -1.817 | transmembrane protein | elongation zone | 6 h |
| HORVU.MOREX.r3.2HG0188300.1 | -0.42572 | 4.693992 | -3.21566 | 0.002219 | 0.040113 | -1.81923 | CsAtPR5 | elongation zone | 6 h |
| HORVU.MOREX.r3.4HG0348380.1 | 0.828904 | 1.737869 | 3.207005 | 0.002275 | 0.040716 | -1.81929 | phox (PX) domain-containing protein | elongation zone | 6 h |
| HORVU.MOREX.r3.4HG0389200.1 | 0.700295 | 0.876157 | 3.143738 | 0.002732 | 0.046171 | -1.81948 | LIM domain-containing protein 1 | elongation zone | 6 h |
| HORVU.MOREX.r3.3HG0291030.1 | 0.614211 | 1.945138 | 3.312102 | 0.001672 | 0.034187 | -1.82418 | Glucan endo-1,3-beta-glucosidase 1 | elongation zone | 6 h |
| HORVU.MOREX.r3.3HG0292680.1 | 0.687349 | 2.459133 | 3.254365 | 0.001982 | 0.037445 | -1.82681 | Replicase polyprotein 1a | elongation zone | 6 h |
| HORVU.MOREX.r3.5HG0488650.1 | 0.700777 | 4.071835 | 3.282384 | 0.001825 | 0.035637 | -1.82711 | Dof zinc finger protein | elongation zone | 6 h |
| HORVU.MOREX.r3.6HG0616240.1 | 0.837459 | -0.52276 | 3.239264 | 0.002071 | 0.038428 | -1.83191 | Endoglucanase | elongation zone | 6 h |
| HORVU.MOREX.r3.3HG0327170.1 | -0.42388 | 7.966567 | -3.35975 | 0.001452 | 0.031376 | -1.83443 | Bifunctional polymyxin resistance arnA protein | elongation zone | 6 h |
| HORVU.MOREX.r3.3HG0314240.1 | -0.41732 | 5.164101 | -3.25989 | 0.00195 | 0.03698 | -1.83528 | Potassium transporter | elongation zone | 6 h |
| HORVU.MOREX.r3.7HG0665990.1 | 0.579124 | 6.156682 | 3.37477 | 0.001388 | 0.030794 | -1.84092 | Shikimate kinase | elongation zone | 6 h |
| HORVU.MOREX.r3.3HG0246370.1 | -0.4324 | 5.820127 | -3.26366 | 0.001928 | 0.036846 | -1.84285 | Mannosyltransferase | elongation zone | 6 h |
| HORVU.MOREX.r3.4HG0379400.1 | 0.812035 | 1.850081 | 3.337366 | 0.001552 | 0.032356 | -1.84646 | Protein kinase-like protein | elongation zone | 6 h |
| HORVU.MOREX.r3.5HG0485760.1 | 1.397883 | -1.3336 | 3.117853 | 0.002942 | 0.048492 | -1.84867 | 65-kDa microtubule-associated-like protein | elongation zone | 6 h |
| HORVU.MOREX.r3.2HG0171550.1 | -0.64018 | 4.111529 | -3.15923 | 0.002613 | 0.044825 | -1.84942 | Major facilitator superfamily transporter | elongation zone | 6 h |
| HORVU.MOREX.r3.7HG0680970.1 | -0.79884 | 3.415292 | -3.10299 | 0.00307 | 0.049561 | -1.8564 | Mitochondrial transcription termination factor family protein | elongation zone | 6 h |
| HORVU.MOREX.r3.5HG0477800.1 | 0.693459 | 5.596819 | 3.29854 | 0.00174 | 0.03465 | -1.85777 | Endo-1,3(4)-beta-glucanase 1 | elongation zone | 6 h |
| HORVU.MOREX.r3.1HG0076120.1 | 0.568455 | 7.792381 | 3.461231 | 0.001071 | 0.026245 | -1.85944 | Phosphatidylinositol transfer protein | elongation zone | 6 h |
| HORVU.MOREX.r3.3HG0230880.1 | -0.2135 | 7.037661 | -3.35025 | 0.001493 | 0.031694 | -1.86093 | Steroid 5-alpha reductase | elongation zone | 6 h |
| HORVU.MOREX.r3.1HG0050920.1 | 0.834189 | 2.478719 | 3.277063 | 0.001854 | 0.03596 | -1.86675 | Trehalase | elongation zone | 6 h |
| HORVU.MOREX.r3.1HG0090200.1 | -0.51011 | 5.563212 | -3.22797 | 0.002141 | 0.039149 | -1.87046 | NBS-LRR-like resistance protein | elongation zone | 6 h |
| HORVU.MOREX.r3.2HG0121830.1 | -0.24655 | 7.00528 | -3.34793 | 0.001504 | 0.031712 | -1.87179 | Calcium ion-binding protein | elongation zone | 6 h |
| HORVU.MOREX.r3.1HG0062450.1 | -0.8201 | 7.022636 | -3.27805 | 0.001849 | 0.035889 | -1.87181 | Pyrimidine-specific ribonucleoside hydrolase | elongation zone | 6 h |
| HORVU.MOREX.r3.5HG0491250.1 | -0.43929 | 3.850964 | -3.14751 | 0.002702 | 0.045882 | -1.87421 | Cinnamoyl-CoA reductase 4 | elongation zone | 6 h |
| HORVU.MOREX.r3.2HG0134300.1 | 2.069302 | 1.307766 | 3.097573 | 0.003118 | 0.049823 | -1.88001 | Skp1, putative | elongation zone | 6 h |
| HORVU.MOREX.r3.1HG0011820.1 | 0.973091 | -1.28428 | 3.127317 | 0.002863 | 0.04772 | -1.88002 | Protein ABIL1 | elongation zone | 6 h |
| HORVU.MOREX.r3.2HG0141980.1 | -0.40365 | 5.307117 | -3.29845 | 0.001741 | 0.03465 | -1.88139 | Glucan endo-1,3-beta-glucosidase 3 | elongation zone | 6 h |

**Table S1** Continued.

| ID | logFC | AveExpr | t | P.Value | adj.P.Val | B | gene description | root zone | time point |
|---|---|---|---|---|---|---|---|---|---|
| HORVU.MOREX.r3.3HG0258280.1 | -0.28742 | 6.470164 | -3.3035 | 0.001715 | 0.034524 | -1.88715 | Coatomer subunit beta | elongation zone | 6 h |
| HORVU.MOREX.r3.3HG0298540.1 | 0.583325 | 7.885451 | 3.443447 | 0.00113 | 0.027022 | -1.89307 | Phosphatidylinositol transfer protein | elongation zone | 6 h |
| HORVU.MOREX.r3.4HG0390140.1 | -0.26214 | 5.370404 | -3.24183 | 0.002056 | 0.038279 | -1.90153 | Vacuolar protein sorting-associated protein 9A | elongation zone | 6 h |
| HORVU.MOREX.r3.2HG0115930.1 | -0.50204 | 9.174331 | -3.38766 | 0.001336 | 0.030044 | -1.90383 | 4-alpha-glucanotransferase | elongation zone | 6 h |
| HORVU.MOREX.r3.2HG0168330.1 | 0.435261 | 5.088967 | 3.271911 | 0.001882 | 0.036357 | -1.90456 | Nicotinate phosphoribosyltransferase | elongation zone | 6 h |
| HORVU.MOREX.r3.5HG0444350.1 | 0.752919 | 2.597523 | 3.110473 | 0.003005 | 0.048898 | -1.90532 | Sn1-specific diacylglycerol lipase alpha | elongation zone | 6 h |
| HORVU.MOREX.r3.2HG0160690.1 | 1.038193 | -0.41896 | 3.13547 | 0.002797 | 0.046935 | -1.90533 | RALF | elongation zone | 6 h |
| HORVU.MOREX.r3.1HG0003430.1 | 0.458613 | 4.894308 | 3.235062 | 0.002097 | 0.038797 | -1.90577 | TBC1 domain family member | elongation zone | 6 h |
| HORVU.MOREX.r3.2HG0206920.1 | 0.937904 | 4.999661 | 3.102677 | 0.003073 | 0.049566 | -1.90945 | DNA polymerase I | elongation zone | 6 h |
| HORVU.MOREX.r3.3HG0287000.1 | 0.778647 | 3.768611 | 3.206916 | 0.002276 | 0.040716 | -1.90996 | Respiratory burst oxidase | elongation zone | 6 h |
| HORVU.MOREX.r3.1HG0072510.1 | 1.043202 | -0.03799 | 3.111979 | 0.002992 | 0.048862 | -1.91435 | Protein-methionine-s-oxide reductase | elongation zone | 6 h |
| HORVU.MOREX.r3.1HG0094770.1 | -0.5512 | 3.670278 | -3.10685 | 0.003036 | 0.049244 | -1.91684 | Transmembrane protein, putative | elongation zone | 6 h |
| HORVU.MOREX.r3.3HG0316130.1 | -0.69532 | 7.033938 | -3.34656 | 0.00151 | 0.031775 | -1.91747 | Basic 7S globulin 2 | elongation zone | 6 h |
| HORVU.MOREX.r3.3HG0305800.1 | 0.724486 | 4.447 | 3.1809 | 0.002454 | 0.042679 | -1.92296 | Ribonuclease | elongation zone | 6 h |
| HORVU.MOREX.r3.1HG0076060.1 | -0.36366 | 6.353402 | -3.29178 | 0.001775 | 0.034965 | -1.92936 | C2 calcium/lipid-binding and GRAM domain protein | elongation zone | 6 h |
| HORVU.MOREX.r3.6HG0575530.1 | 0.490052 | 6.1314 | 3.278125 | 0.001848 | 0.035889 | -1.92997 | Alcohol dehydrogenase, putative | elongation zone | 6 h |
| HORVU.MOREX.r3.3HG0246500.1 | 0.768424 | 3.040522 | 3.289425 | 0.001788 | 0.035174 | -1.93165 | Peptide-N4-(N-acetyl-beta-glucosaminyl)asparagine amidase A | elongation zone | 6 h |
| HORVU.MOREX.r3.2HG0152890.1 | -0.65663 | 3.809628 | -3.12706 | 0.002866 | 0.04772 | -1.94191 | TCP transcription factor | elongation zone | 6 h |
| HORVU.MOREX.r3.3HG0284780.1 | 0.355958 | 5.323435 | 3.228951 | 0.002134 | 0.039142 | -1.94987 | Protein PLASTID MOVEMENT IMPAIRED 1-RELATED 1 | elongation zone | 6 h |
| HORVU.MOREX.r3.5HG0519810.1 | -0.39195 | 7.617925 | -3.31051 | 0.00168 | 0.034234 | -1.95141 | Diphosphomevalonate decarboxylase | elongation zone | 6 h |
| HORVU.MOREX.r3.7HG0679350.1 | -0.50858 | 3.668408 | -3.22802 | 0.00214 | 0.039149 | -1.95307 | Leucine-rich repeat receptor-like protein kinase family | elongation zone | 6 h |
| HORVU.MOREX.r3.5HG0527720.1 | 0.979501 | 0.951915 | 3.096354 | 0.003128 | 0.049957 | -1.95442 | Receptor protein kinase, putative,expressed | elongation zone | 6 h |
| HORVU.MOREX.r3.5HG0493200.1 | 0.427709 | 8.020324 | 3.364467 | 0.001431 | 0.031116 | -1.9545 | Fasciclin-like arabinogalactan-protein-like | elongation zone | 6 h |
| HORVU.MOREX.r3.4HG0384110.1 | -0.30907 | 6.300031 | -3.26797 | 0.001904 | 0.036589 | -1.95779 | RING/U-box superfamily protein | elongation zone | 6 h |
| HORVU.MOREX.r3.3HG0304710.1 | -0.28797 | 8.270238 | -3.36374 | 0.001435 | 0.031117 | -1.95899 | Fructose-bisphosphate aldolase | elongation zone | 6 h |
| HORVU.MOREX.r3.7HG0651010.1 | -0.85539 | 4.932391 | -3.18358 | 0.002435 | 0.0426 | -1.96309 | Xyloglucan endotransglucosylase/hydrolase | elongation zone | 6 h |
| HORVU.MOREX.r3.6HG0568170.1 | 0.852894 | 3.002871 | 3.15493 | 0.002645 | 0.045269 | -1.96711 | Transmembrane protein | elongation zone | 6 h |
| HORVU.MOREX.r3.1HG0042740.1 | -0.29675 | 6.415285 | -3.26637 | 0.001913 | 0.036691 | -1.96834 | Adaptin ear-binding coat-associated protein, putative | elongation zone | 6 h |
| HORVU.MOREX.r3.1HG0086390.1 | 0.796769 | -0.26089 | 3.121064 | 0.002915 | 0.048242 | -1.97279 | #N/A | elongation zone | 6 h |
| HORVU.MOREX.r3.4HG0383400.1 | -0.35575 | 5.833159 | -3.28416 | 0.001816 | 0.035519 | -1.97728 | Protein COBRA, putative | elongation zone | 6 h |
| HORVU.MOREX.r3.2HG0192000.1 | 0.786542 | 0.357064 | 3.204967 | 0.002289 | 0.040793 | -1.97752 | HXXXD-type acyl-transferase family protein, putative | elongation zone | 6 h |
| HORVU.MOREX.r3.7HG0741640.1 | -0.75666 | 4.20973 | -3.12684 | 0.002867 | 0.04772 | -1.97862 | Histone H2A | elongation zone | 6 h |
| HORVU.MOREX.r3.7HG0734380.1 | -0.4554 | 4.659788 | -3.16334 | 0.002582 | 0.044482 | -1.97938 | Mechanosensitive ion channel-like protein | elongation zone | 6 h |
| HORVU.MOREX.r3.1HG0058470.1 | -0.37992 | 5.359187 | -3.20794 | 0.002269 | 0.040716 | -1.98097 | RING/U-box superfamily protein, putative | elongation zone | 6 h |
| HORVU.MOREX.r3.7HG0734670.1 | -0.44022 | 6.626504 | -3.27951 | 0.001841 | 0.035871 | -1.98806 | L-ascorbate oxidase | elongation zone | 6 h |
| HORVU.MOREX.r3.4HG0377020.1 | -0.29423 | 6.726202 | -3.29446 | 0.001762 | 0.03479 | -1.98904 | Hexosyltransferase | elongation zone | 6 h |
| HORVU.MOREX.r3.6HG0616890.1 | -0.91748 | 0.931971 | -3.1074 | 0.003031 | 0.049213 | -1.99113 | UPF0503 protein, chloroplastic | elongation zone | 6 h |
| HORVU.MOREX.r3.2HG0116990.1 | 0.555538 | 4.613527 | 3.212102 | 0.002242 | 0.040424 | -1.99897 | Phosphatidylinositol 4-phosphate 5-kinase 4 | elongation zone | 6 h |
| HORVU.MOREX.r3.6HG0543400.1 | 1.327886 | -0.34964 | 3.140341 | 0.002758 | 0.046473 | -2.00517 | High affinity nitrate transporter | elongation zone | 6 h |
| HORVU.MOREX.r3.1HG0021880.1 | 2.444903 | 0.747345 | 3.14276 | 0.002739 | 0.046227 | -2.00635 | SKP1-like protein 4 | elongation zone | 6 h |
| HORVU.MOREX.r3.3HG0271380.1 | 0.474111 | 5.415877 | 3.185576 | 0.002421 | 0.042416 | -2.0065 | GDSL esterase/lipase | elongation zone | 6 h |

**Table S1** Continued.

| ID | logFC | AveExpr | t | P.Value | adj.P.Val | B | gene description | root zone | time point |
|---|---|---|---|---|---|---|---|---|---|
| HORVU.MOREX.r3.6HG0615130.1 | 0.663016 | 3.141376 | 3.218471 | 0.002201 | 0.039927 | -2.00906 | TCP transcription factor | elongation zone | 6 h |
| HORVU.MOREX.r3.2HG0197110.1 | 0.355576 | 6.332954 | 3.262124 | 0.001937 | 0.036879 | -2.0123 | AT hook motif DNA-binding family protein | elongation zone | 6 h |
| HORVU.MOREX.r3.4HG0331420.1 | 1.3604 | 6.431216 | 3.10655 | 0.003039 | 0.049244 | -2.01882 | Peptidyl-prolyl cis-trans isomerase | elongation zone | 6 h |
| HORVU.MOREX.r3.1HG0042870.1 | -0.24621 | 6.875706 | -3.26763 | 0.001906 | 0.036589 | -2.02184 | Serine/threonine-protein phosphatase 2A regulatory subunit B'' subunit alpha | elongation zone | 6 h |
| HORVU.MOREX.r3.7HG0745360.1 | -0.31756 | 4.449287 | -3.16202 | 0.002592 | 0.044578 | -2.02431 | Actin-related protein 2/3 complex subunit 3 | elongation zone | 6 h |
| HORVU.MOREX.r3.7HG0718830.1 | 0.72123 | 4.155941 | 3.285978 | 0.001806 | 0.035465 | -2.03253 | FAD-binding Berberine family protein, putative | elongation zone | 6 h |
| HORVU.MOREX.r3.7HG0668440.1 | 0.559021 | 5.028289 | 3.196218 | 0.002348 | 0.041493 | -2.03449 | #N/A | elongation zone | 6 h |
| HORVU.MOREX.r3.4HG0403510.1 | -0.4133 | 4.248358 | -3.14488 | 0.002723 | 0.046126 | -2.0376 | Protein kinase | elongation zone | 6 h |
| HORVU.MOREX.r3.4HG0407470.1 | 0.865251 | 2.42529 | 3.223382 | 0.002169 | 0.039535 | -2.04401 | Serpin | elongation zone | 6 h |
| HORVU.MOREX.r3.4HG0352200.1 | 0.624645 | 3.627838 | 3.139401 | 0.002766 | 0.046484 | -2.04431 | Methyltransferase-related protein, putative | elongation zone | 6 h |
| HORVU.MOREX.r3.3HG0219860.1 | -0.44448 | 3.216471 | -3.10118 | 0.003086 | 0.049584 | -2.0446 | Disease resistance protein (NBS-LRR class) family | elongation zone | 6 h |
| HORVU.MOREX.r3.7HG0687540.1 | -0.31187 | 5.801521 | -3.21049 | 0.002252 | 0.040566 | -2.05101 | Proteasome subunit alpha type | elongation zone | 6 h |
| HORVU.MOREX.r3.1HG0003620.1 | 0.592803 | 2.573513 | 3.100608 | 0.003091 | 0.049625 | -2.05571 | Protein ZINC INDUCED FACILITATOR-LIKE 1 | elongation zone | 6 h |
| HORVU.MOREX.r3.7HG0718930.2 | 1.333753 | -0.79034 | 3.145132 | 0.002721 | 0.046126 | -2.05726 | Peroxidase | elongation zone | 6 h |
| HORVU.MOREX.r3.2HG0183780.1 | -0.4371 | 5.376384 | -3.18678 | 0.002413 | 0.042315 | -2.05813 | Sphingolipid delta-4 desaturase | elongation zone | 6 h |
| HORVU.MOREX.r3.1HG0042290.1 | 0.406695 | 4.21134 | 3.17124 | 0.002524 | 0.043737 | -2.06071 | NBS-LRR disease resistance protein | elongation zone | 6 h |
| HORVU.MOREX.r3.2HG0168090.1 | -0.33916 | 5.385495 | -3.18295 | 0.00244 | 0.042629 | -2.06769 | Sphingomyelin synthase-like domain-containing protein | elongation zone | 6 h |
| HORVU.MOREX.r3.6HG0554070.1 | 3.668711 | 1.92611 | 3.097922 | 0.003114 | 0.049823 | -2.06807 | HECT-like ubiquitin-conjugating enzyme (E2)-binding protein | elongation zone | 6 h |
| HORVU.MOREX.r3.3HG0242770.1 | 0.653721 | 1.656653 | 3.119969 | 0.002924 | 0.048315 | -2.0787 | Glycosyltransferases | elongation zone | 6 h |
| HORVU.MOREX.r3.1HG0046180.1 | 0.409217 | 5.918665 | 3.244527 | 0.00204 | 0.038117 | -2.07895 | Ser/Thr protein phosphatase family protein, expressed | elongation zone | 6 h |
| HORVU.MOREX.r3.1HG0069410.1 | 1.113651 | 3.463735 | 3.166476 | 0.002559 | 0.044197 | -2.08305 | Aldehyde dehydrogenase | elongation zone | 6 h |
| HORVU.MOREX.r3.6HG0599080.1 | -0.38683 | 8.233534 | -3.27648 | 0.001857 | 0.035987 | -2.08854 | Hsp70-Hsp90 organizing protein 1 | elongation zone | 6 h |
| HORVU.MOREX.r3.7HG0730870.1 | 0.374378 | 7.493886 | 3.308519 | 0.00169 | 0.034363 | -2.09011 | Glycosyltransferases | elongation zone | 6 h |
| HORVU.MOREX.r3.3HG0255040.1 | -0.3733 | 4.904219 | -3.12011 | 0.002923 | 0.048315 | -2.09233 | Mediator of RNA polymerase II transcription subunit 27 | elongation zone | 6 h |
| HORVU.MOREX.r3.1HG0053680.1 | -0.30219 | 5.78418 | -3.20829 | 0.002267 | 0.040716 | -2.09586 | Choline-phosphate cytidylyltransferase | elongation zone | 6 h |
| HORVU.MOREX.r3.6HG0548360.1 | -0.59509 | 3.627752 | -3.11204 | 0.002991 | 0.048862 | -2.09843 | F-box family protein | elongation zone | 6 h |
| HORVU.MOREX.r3.3HG0221440.1 | 0.806842 | 2.700751 | 3.297347 | 0.001747 | 0.034729 | -2.10235 | Glycosyltransferase | elongation zone | 6 h |
| HORVU.MOREX.r3.1HG0079870.1 | -0.82007 | 5.830169 | -3.19521 | 0.002355 | 0.041579 | -2.10358 | Pectin lyase-like superfamily protein | elongation zone | 6 h |
| HORVU.MOREX.r3.1HG0058140.1 | -0.40767 | 4.033555 | -3.15332 | 0.002657 | 0.045366 | -2.10444 | ethylene-responsive transcription factor | elongation zone | 6 h |
| HORVU.MOREX.r3.6HG0622740.1 | 0.452234 | 4.313476 | 3.182758 | 0.002441 | 0.042629 | -2.11316 | Protein DETOXIFICATION | elongation zone | 6 h |
| HORVU.MOREX.r3.3HG0252610.1 | -0.3373 | 7.233439 | -3.21691 | 0.002211 | 0.040038 | -2.11581 | Protein kinase | elongation zone | 6 h |
| HORVU.MOREX.r3.3HG0270910.1 | -0.25347 | 6.963859 | -3.25102 | 0.002001 | 0.03764 | -2.1161 | Protein YIPF | elongation zone | 6 h |
| HORVU.MOREX.r3.2HG0119220.1 | -0.46391 | 4.892203 | -3.16335 | 0.002582 | 0.044482 | -2.12344 | Gibberellin receptor GID1a | elongation zone | 6 h |
| HORVU.MOREX.r3.6HG0608780.1 | -0.41697 | 7.822442 | -3.26174 | 0.001939 | 0.036882 | -2.1274 | Heat shock 70 kDa protein, putative | elongation zone | 6 h |
| HORVU.MOREX.r3.6HG0632320.1 | -0.36064 | 5.523777 | -3.14874 | 0.002693 | 0.045815 | -2.16011 | Leucine-rich repeat protein | elongation zone | 6 h |
| HORVU.MOREX.r3.5HG0501120.1 | -0.26871 | 6.923122 | -3.20468 | 0.002291 | 0.040793 | -2.16127 | 26S proteasome regulatory subunit | elongation zone | 6 h |
| HORVU.MOREX.r3.7HG0667630.1 | 1.074915 | 5.380802 | 3.186901 | 0.002412 | 0.042315 | -2.16602 | Peroxidase | elongation zone | 6 h |
| HORVU.MOREX.r3.6HG0634200.1 | 0.950669 | 0.006824 | 3.112597 | 0.002987 | 0.048862 | -2.17338 | Syntaxin | elongation zone | 6 h |
| HORVU.MOREX.r3.5HG0430940.1 | 0.928441 | 5.193365 | 3.187535 | 0.002407 | 0.042295 | -2.17376 | Glucan 1,3-beta-glucosidase | elongation zone | 6 h |
| HORVU.MOREX.r3.7HG0711200.1 | -0.38 | 5.29275 | -3.18537 | 0.002423 | 0.042416 | -2.19147 | Hexosyltransferase | elongation zone | 6 h |
| HORVU.MOREX.r3.3HG0310540.1 | -0.49862 | 5.764879 | -3.13204 | 0.002825 | 0.047264 | -2.19569 | Mannosyl-oligosaccharide glucosidase | elongation zone | 6 h |

**Table S1** Continued.

| ID | logFC | AveExpr | t | P.Value | adj.P.Val | B | gene description | root zone | time point |
|---|---|---|---|---|---|---|---|---|---|
| HORVU.MOREX.r3.2HG0105990.1 | -0.41256 | 6.154762 | -3.19789 | 0.002336 | 0.041399 | -2.19646 | Cytochrome P450 | elongation zone | 6 h |
| HORVU.MOREX.r3.2HG0184180.1 | 0.884669 | 4.475955 | 3.19131 | 0.002381 | 0.04193 | -2.20257 | Remorin | elongation zone | 6 h |
| HORVU.MOREX.r3.6HG0577360.1 | -0.66197 | 5.665724 | -3.26011 | 0.001949 | 0.03698 | -2.20989 | transmembrane protein, putative (Protein of unknown function, DUF642) | elongation zone | 6 h |
| HORVU.MOREX.r3.5HG0454860.1 | -0.3381 | 5.797971 | -3.13975 | 0.002763 | 0.046475 | -2.21359 | Trehalose-6-phosphate synthase, putative | elongation zone | 6 h |
| HORVU.MOREX.r3.4HG0342160.1 | -0.36356 | 5.308472 | -3.11332 | 0.002981 | 0.04885 | -2.21486 | Presenilin | elongation zone | 6 h |
| HORVU.MOREX.r3.5HG0519040.1 | -0.36819 | 4.736037 | -3.11184 | 0.002993 | 0.048862 | -2.21578 | Serine/arginine repetitive matrix protein 2, putative isoform 1 | elongation zone | 6 h |
| HORVU.MOREX.r3.6HG0541250.1 | -0.20957 | 7.143195 | -3.18755 | 0.002407 | 0.042295 | -2.24353 | Acetate kinase | elongation zone | 6 h |
| HORVU.MOREX.r3.1HG0026830.1 | 0.505722 | 3.566935 | 3.1115 | 0.002996 | 0.048871 | -2.24463 | Leucine-rich repeat receptor-like protein kinase family | elongation zone | 6 h |
| HORVU.MOREX.r3.7HG0662650.1 | -0.71282 | 6.460162 | -3.13987 | 0.002762 | 0.046475 | -2.24544 | Histone H3 | elongation zone | 6 h |
| HORVU.MOREX.r3.1HG0022860.1 | -0.3017 | 5.022173 | -3.10159 | 0.003082 | 0.049584 | -2.24629 | Xylosyltransferase 1 | elongation zone | 6 h |
| HORVU.MOREX.r3.4HG0347400.1 | -0.45285 | 6.883926 | -3.19863 | 0.002331 | 0.041347 | -2.2529 | U-box domain-containing protein | elongation zone | 6 h |
| HORVU.MOREX.r3.2HG0121410.1 | 0.925683 | 3.844656 | 3.127803 | 0.00286 | 0.04772 | -2.25388 | Cytochrome P450 | elongation zone | 6 h |
| HORVU.MOREX.r3.1HG0072840.1 | -0.34283 | 5.890668 | -3.12399 | 0.002891 | 0.047996 | -2.27082 | Ectonucleoside triphosphate diphosphohydrolase 1 | elongation zone | 6 h |
| HORVU.MOREX.r3.7HG0739180.1 | -0.30909 | 7.099598 | -3.20888 | 0.002263 | 0.040697 | -2.28083 | Core-2/I-branching beta-1,6-N-acetylglucosaminyltransferase family protein | elongation zone | 6 h |
| HORVU.MOREX.r3.3HG0284100.1 | 0.38402 | 6.109021 | 3.16049 | 0.002603 | 0.044719 | -2.29999 | 5'-nucleotidase SurE | elongation zone | 6 h |
| HORVU.MOREX.r3.2HG0190290.1 | 0.601391 | 5.33841 | 3.110596 | 0.003004 | 0.048898 | -2.30441 | SHR5-receptor-like kinase | elongation zone | 6 h |
| HORVU.MOREX.r3.3HG0262210.1 | -0.28094 | 5.498655 | -3.10775 | 0.003028 | 0.049213 | -2.30876 | Phosphatase 2C family protein | elongation zone | 6 h |
| HORVU.MOREX.r3.3HG0288360.1 | -0.2874 | 5.883309 | -3.11697 | 0.00295 | 0.048576 | -2.31613 | Dolichyl-diphosphooligosaccharide--protein glycosyltransferase subunit STT3A | elongation zone | 6 h |
| HORVU.MOREX.r3.5HG0463000.1 | -0.44824 | 4.936645 | -3.11969 | 0.002927 | 0.048315 | -2.32535 | Ulp1 protease family, C-terminal catalytic domain containing protein, expressed | elongation zone | 6 h |
| HORVU.MOREX.r3.5HG0446900.1 | -0.21894 | 6.633774 | -3.14737 | 0.002703 | 0.045882 | -2.32789 | Dipeptidyl peptidase | elongation zone | 6 h |
| HORVU.MOREX.r3.4HG0335700.1 | -0.23936 | 5.55838 | -3.09807 | 0.003113 | 0.049823 | -2.33937 | Alkaline ceramidase 3 | elongation zone | 6 h |
| HORVU.MOREX.r3.7HG0677940.1 | -0.28967 | 7.210054 | -3.16645 | 0.002559 | 0.044197 | -2.34109 | Transmembrane 9 superfamily member | elongation zone | 6 h |
| HORVU.MOREX.r3.2HG0205750.1 | -0.45601 | 6.410155 | -3.11506 | 0.002966 | 0.048764 | -2.34485 | 3-oxoacyl-[acyl-carrier-protein] synthase 3 | elongation zone | 6 h |
| HORVU.MOREX.r3.6HG0614100.1 | 0.831645 | 4.236522 | 3.103064 | 0.003069 | 0.049561 | -2.35241 | TPR repeat-containing thioredoxin TTL4 | elongation zone | 6 h |
| HORVU.MOREX.r3.5HG0527280.1 | -0.25161 | 6.528337 | -3.14466 | 0.002724 | 0.046126 | -2.35717 | Ras-related protein, expressed | elongation zone | 6 h |
| HORVU.MOREX.r3.3HG0287070.1 | -0.83892 | 6.192473 | -3.10631 | 0.003041 | 0.049244 | -2.37004 | Glycosyltransferase | elongation zone | 6 h |
| HORVU.MOREX.r3.5HG0419520.1 | -0.26213 | 7.603138 | -3.20361 | 0.002298 | 0.040859 | -2.37753 | Microtubule-associated protein 70-2 | elongation zone | 6 h |
| HORVU.MOREX.r3.1HG0061710.1 | 0.424179 | 5.632386 | 3.130881 | 0.002834 | 0.047363 | -2.39222 | Trichome birefringence-like protein | elongation zone | 6 h |
| HORVU.MOREX.r3.5HG0529660.1 | -0.27169 | 6.908166 | -3.11468 | 0.002969 | 0.048778 | -2.39388 | Squamosa promoter-binding protein, putative | elongation zone | 6 h |
| HORVU.MOREX.r3.3HG0301750.1 | -0.45649 | 9.943131 | -3.255 | 0.001978 | 0.037409 | -2.39511 | Alpha-1,4 glucan phosphorylase | elongation zone | 6 h |
| HORVU.MOREX.r3.5HG0425680.1 | -0.33898 | 7.233115 | -3.13553 | 0.002797 | 0.046935 | -2.39732 | No exine formation 1 | elongation zone | 6 h |
| HORVU.MOREX.r3.7HG0726790.1 | -0.30144 | 7.796481 | -3.13189 | 0.002826 | 0.047264 | -2.45362 | Dolichyl-diphosphooligosaccharide--protein glycosyltransferase 48 kDa subunit | elongation zone | 6 h |
| HORVU.MOREX.r3.2HG0108780.1 | -0.20548 | 8.923708 | -3.19223 | 0.002375 | 0.041904 | -2.45811 | 26S protease regulatory subunit | elongation zone | 6 h |
| HORVU.MOREX.r3.5HG0430480.1 | -0.27297 | 7.401068 | -3.14184 | 0.002747 | 0.046311 | -2.4776 | Protein kinase | elongation zone | 6 h |
| HORVU.MOREX.r3.4HG0381800.1 | -0.27972 | 7.086406 | -3.10202 | 0.003078 | 0.049581 | -2.48719 | AP-2 complex subunit mu | elongation zone | 6 h |
| HORVU.MOREX.r3.6HG0539460.1 | -0.22919 | 7.146504 | -3.09918 | 0.003103 | 0.049789 | -2.49466 | FG-GAP repeat-containing protein | elongation zone | 6 h |
| HORVU.MOREX.r3.2HG0199550.1 | -0.52199 | 7.232601 | -3.10747 | 0.003031 | 0.049213 | -2.50743 | Glycosyltransferase | elongation zone | 6 h |
| HORVU.MOREX.r3.3HG0221490.1 | -0.34942 | 7.438494 | -3.12734 | 0.002863 | 0.04772 | -2.50905 | Sodium/hydrogen exchanger 1 | elongation zone | 6 h |
| HORVU.MOREX.r3.2HG0138540.1 | 0.579899 | 4.435354 | 3.170758 | 0.002527 | 0.043758 | -2.52317 | Blue copper protein | elongation zone | 6 h |
| HORVU.MOREX.r3.2HG0135210.1 | 0.672772 | 2.447492 | 3.101421 | 0.003084 | 0.049584 | -2.53729 | Late embryogenesis abundant (LEA) hydroxyproline-rich glycoprotein family | elongation zone | 6 h |
| HORVU.MOREX.r3.5HG0470030.1 | -0.40454 | 7.999201 | -3.11882 | 0.002934 | 0.048397 | -2.56747 | 3-ketoacyl-CoA thiolase | elongation zone | 6 h |

**Table S1** Continued.

| ID | logFC | AveExpr | t | P.Value | adj.P.Val | B | gene description | root zone | time point |
|---|---|---|---|---|---|---|---|---|---|
| HORVU.MOREX.r3.4HG0346720.1 | -0.41162 | 10.60861 | -3.20447 | 0.002292 | 0.040793 | -2.5856 | Protein disulfide-isomerase | elongation zone | 6 h |
| HORVU.MOREX.r3.7HG0699010.1 | 2.219936 | 4.331849 | 8.039164 | 9.67E-11 | 1.99E-06 | 14.06542 | LHY | elongation zone | 12 h |
| HORVU.MOREX.r3.3HG0328340.1 | -1.61312 | 4.602643 | -7.59881 | 4.90E-10 | 5.03E-06 | 12.60193 | Two-component response regulator | elongation zone | 12 h |
| HORVU.MOREX.r3.6HG0550600.1 | 1.560832 | 1.720716 | 6.401342 | 4.13E-08 | 0.000274 | 8.492075 | Protein ROOT HAIR DEFECTIVE 3 homolog | elongation zone | 12 h |
| HORVU.MOREX.r3.6HG0609520.1 | -0.83778 | 8.966633 | -6.33217 | 5.33E-08 | 0.000274 | 8.133236 | Fatty acid desaturase | elongation zone | 12 h |
| HORVU.MOREX.r3.2HG0097390.1 | -0.9076 | 4.644505 | -6.05012 | 1.51E-07 | 0.000515 | 7.277088 | Inosine-5'-monophosphate dehydrogenase | elongation zone | 12 h |
| HORVU.MOREX.r3.5HG0517740.1 | 2.365849 | -0.30721 | 5.929291 | 2.34E-07 | 0.000675 | 6.855633 | Peroxidase | elongation zone | 12 h |
| HORVU.MOREX.r3.5HG0537150.1 | 1.700402 | -2.24688 | 6.145168 | 1.06E-07 | 0.000436 | 6.729744 | Protein DETOXIFICATION | elongation zone | 12 h |
| HORVU.MOREX.r3.7HG0636030.1 | 1.403702 | -1.10938 | 5.84935 | 3.14E-07 | 0.000675 | 6.556341 | hydroxysteroid dehydrogenase 3 | elongation zone | 12 h |
| HORVU.MOREX.r3.4HG0415420.1 | -0.98235 | 6.947214 | -5.80147 | 3.74E-07 | 0.000675 | 6.357551 | Tubulin beta chain | elongation zone | 12 h |
| HORVU.MOREX.r3.5HG0477040.1 | -0.65743 | 5.55814 | -5.75481 | 4.43E-07 | 0.000675 | 6.254191 | MND1-interacting protein 1 | elongation zone | 12 h |
| HORVU.MOREX.r3.7HG0715570.1 | -0.64331 | 6.466383 | -5.70699 | 5.27E-07 | 0.000675 | 6.044857 | Transmembrane protein | elongation zone | 12 h |
| HORVU.MOREX.r3.1HG0072760.1 | -0.91262 | 4.906091 | -5.69425 | 5.52E-07 | 0.000675 | 6.040728 | Serpin family protein | elongation zone | 12 h |
| HORVU.MOREX.r3.1HG0065860.1 | 1.059627 | 4.236426 | 5.684903 | 5.71E-07 | 0.000675 | 6.007964 | MLO-like protein | elongation zone | 12 h |
| HORVU.MOREX.r3.4HG0415330.1 | 2.206769 | 3.57761 | 5.655959 | 6.35E-07 | 0.000675 | 5.917139 | Cytochrome P450 family cinnamate 4-hydroxylase | elongation zone | 12 h |
| HORVU.MOREX.r3.3HG0329870.1 | 2.009599 | -1.90322 | 5.688373 | 5.64E-07 | 0.000675 | 5.861495 | Pectin acetylesterase | elongation zone | 12 h |
| HORVU.MOREX.r3.7HG0736960.1 | -1.15281 | 3.87866 | -5.62384 | 7.13E-07 | 0.000675 | 5.692605 | BTB/POZ domain containing protein, expressed | elongation zone | 12 h |
| HORVU.MOREX.r3.6HG0541940.1 | -0.53468 | 6.764259 | -5.60401 | 7.66E-07 | 0.000675 | 5.659529 | S-adenosyl-L-methionine-dependent methyltransferases superfamily protein | elongation zone | 12 h |
| HORVU.MOREX.r3.6HG0576550.1 | -0.70013 | 4.943295 | -5.57261 | 8.58E-07 | 0.000675 | 5.626732 | F-box protein | elongation zone | 12 h |
| HORVU.MOREX.r3.5HG0477180.1 | -0.57894 | 5.597571 | -5.56653 | 8.77E-07 | 0.000675 | 5.589844 | LAG1 longevity assurance-like protein | elongation zone | 12 h |
| HORVU.MOREX.r3.2HG0100750.1 | 2.000371 | -1.62151 | 5.572159 | 8.60E-07 | 0.000675 | 5.583729 | Dirigent protein | elongation zone | 12 h |
| HORVU.MOREX.r3.3HG0318590.1 | 2.636097 | -1.25566 | 5.567612 | 8.74E-07 | 0.000675 | 5.555983 | Endo-1,3-beta-glucanase | elongation zone | 12 h |
| HORVU.MOREX.r3.6HG0545620.1 | -0.55782 | 11.02319 | -5.66803 | 6.07E-07 | 0.000675 | 5.544028 | Heat shock 70 kDa protein | elongation zone | 12 h |
| HORVU.MOREX.r3.7HG0709860.1 | -0.69289 | 6.368461 | -5.55152 | 9.26E-07 | 0.000675 | 5.54022 | Polyadenylate-binding protein | elongation zone | 12 h |
| HORVU.MOREX.r3.3HG0240640.1 | -1.47936 | 5.859448 | -5.5435 | 9.54E-07 | 0.000675 | 5.535402 | Glycosyltransferase | elongation zone | 12 h |
| HORVU.MOREX.r3.7HG0750000.1 | 2.113151 | -2.12041 | 5.672838 | 5.97E-07 | 0.000675 | 5.532886 | Disease resistance protein | elongation zone | 12 h |
| HORVU.MOREX.r3.4HG0372280.1 | -1.64456 | 3.216377 | -5.67916 | 5.83E-07 | 0.000675 | 5.505877 | xylulose kinase-2 | elongation zone | 12 h |
| HORVU.MOREX.r3.2HG0155750.1 | 2.014688 | -0.29251 | 5.595662 | 7.90E-07 | 0.000675 | 5.478731 | Ribonuclease | elongation zone | 12 h |
| HORVU.MOREX.r3.2HG0205200.1 | -0.76884 | 4.551525 | -5.50324 | 1.10E-06 | 0.000754 | 5.397647 | Ubiquitin carboxyl-terminal hydrolase | elongation zone | 12 h |
| HORVU.MOREX.r3.1HG0010340.1 | 2.200324 | -2.09413 | 5.595111 | 7.91E-07 | 0.000675 | 5.270722 | Non-specific lipid-transfer protein-like protein | elongation zone | 12 h |
| HORVU.MOREX.r3.1HG0079800.1 | -0.70852 | 7.794068 | -5.48428 | 1.18E-06 | 0.000777 | 5.187997 | Farnesyl diphosphate synthase | elongation zone | 12 h |
| HORVU.MOREX.r3.4HG0395540.1 | 1.851854 | 4.838553 | 5.462982 | 1.27E-06 | 0.000793 | 5.187795 | Glutamine synthetase | elongation zone | 12 h |
| HORVU.MOREX.r3.2HG0169700.1 | -0.74087 | 4.388016 | -5.40701 | 1.56E-06 | 0.000889 | 5.074029 | transmembrane protein | elongation zone | 12 h |
| HORVU.MOREX.r3.5HG0535180.1 | 1.724447 | -1.06768 | 5.40731 | 1.56E-06 | 0.000889 | 5.051941 | #N/A | elongation zone | 12 h |
| HORVU.MOREX.r3.1HG0073390.1 | 1.866204 | 2.372781 | 5.420606 | 1.49E-06 | 0.000889 | 5.034035 | Respiratory burst oxidase-like protein | elongation zone | 12 h |
| HORVU.MOREX.r3.1HG0078970.1 | 1.991718 | 2.920959 | 5.391788 | 1.65E-06 | 0.000913 | 4.962337 | Protein kinase, putative | elongation zone | 12 h |
| HORVU.MOREX.r3.4HG0417010.1 | 1.947177 | -1.71396 | 5.47725 | 1.21E-06 | 0.000777 | 4.95958 | Receptor protein kinase, putative | elongation zone | 12 h |
| HORVU.MOREX.r3.2HG0160560.1 | -1.46809 | 2.799703 | -5.38108 | 1.71E-06 | 0.000924 | 4.945492 | Peptidylprolyl isomerase | elongation zone | 12 h |
| HORVU.MOREX.r3.5HG0527570.1 | 2.048334 | 4.789839 | 5.37112 | 1.77E-06 | 0.000933 | 4.925615 | UDP-glycosyltransferase | elongation zone | 12 h |
| HORVU.MOREX.r3.1HG0059350.1 | -0.82523 | 4.615638 | -5.33851 | 1.99E-06 | 0.000974 | 4.843988 | S-adenosyl-L-methionine-dependent methyltransferases superfamily protein | elongation zone | 12 h |
| HORVU.MOREX.r3.4HG0378580.1 | -0.68702 | 5.353084 | -5.31861 | 2.14E-06 | 0.001022 | 4.745139 | Transcription factor | elongation zone | 12 h |

**Table S1** Continued.

| ID | logFC | AveExpr | t | P.Value | adj.P.Val | B | gene description | root zone | time point |
|---|---|---|---|---|---|---|---|---|---|
| HORVU.MOREX.r3.3HG0329040.1 | 2.624444 | -0.45471 | 5.299649 | 2.29E-06 | 0.001061 | 4.620698 | Vacuolar-processing enzyme | elongation zone | 12 h |
| HORVU.MOREX.r3.6HG0555700.1 | -0.56356 | 8.875975 | -5.34932 | 1.92E-06 | 0.00097 | 4.609942 | Calreticulin/calnexin | elongation zone | 12 h |
| HORVU.MOREX.r3.3HG0305150.1 | -0.94468 | 4.756714 | -5.26829 | 2.56E-06 | 0.001143 | 4.609055 | Pectin lyase-like superfamily protein | elongation zone | 12 h |
| HORVU.MOREX.r3.1HG0028320.1 | -0.60288 | 6.850868 | -5.29541 | 2.33E-06 | 0.001061 | 4.600616 | 70 kDa heat shock protein | elongation zone | 12 h |
| HORVU.MOREX.r3.7HG0746770.1 | -0.69703 | 10.27031 | -5.3463 | 1.94E-06 | 0.00097 | 4.505818 | Chaperone protein htpG family protein | elongation zone | 12 h |
| HORVU.MOREX.r3.1HG0070340.1 | 1.566317 | 4.307027 | 5.224526 | 3.00E-06 | 0.001254 | 4.403143 | Protein kinase-like protein | elongation zone | 12 h |
| HORVU.MOREX.r3.5HG0482670.1 | 1.549392 | 0.628979 | 5.210589 | 3.15E-06 | 0.001254 | 4.35227 | Cellulose synthase | elongation zone | 12 h |
| HORVU.MOREX.r3.3HG0241000.1 | -1.73226 | 6.471225 | -5.19664 | 3.31E-06 | 0.001254 | 4.34874 | HXXXD-type acyl-transferase-like protein | elongation zone | 12 h |
| HORVU.MOREX.r3.7HG0731490.1 | -0.5994 | 4.892075 | -5.18792 | 3.41E-06 | 0.001254 | 4.334328 | Transmembrane protein, putative | elongation zone | 12 h |
| HORVU.MOREX.r3.7HG0742080.1 | 1.576026 | -1.3261 | 5.184107 | 3.46E-06 | 0.001254 | 4.296466 | Receptor-like kinase | elongation zone | 12 h |
| HORVU.MOREX.r3.4HG0384390.1 | 1.395961 | 1.574903 | 5.179502 | 3.52E-06 | 0.001254 | 4.29185 | Protein kinase family protein, putative, expressed | elongation zone | 12 h |
| HORVU.MOREX.r3.1HG0081220.1 | -0.67397 | 5.709055 | -5.18155 | 3.49E-06 | 0.001254 | 4.270927 | Galactose-6-phosphate isomerase subunit LacB | elongation zone | 12 h |
| HORVU.MOREX.r3.1HG0006230.1 | -0.65106 | 5.574 | -5.17619 | 3.56E-06 | 0.001254 | 4.267941 | Disease resistance protein RPM1 | elongation zone | 12 h |
| HORVU.MOREX.r3.2HG0097360.3 | -1.1548 | 5.485237 | -5.16189 | 3.74E-06 | 0.001254 | 4.22101 | Laccase | elongation zone | 12 h |
| HORVU.MOREX.r3.1HG0042740.1 | -0.479 | 6.415285 | -5.1747 | 3.58E-06 | 0.001254 | 4.202821 | Adaptin ear-binding coat-associated protein, putative | elongation zone | 12 h |
| HORVU.MOREX.r3.2HG0112670.1 | 1.837357 | 0.989354 | 5.170071 | 3.64E-06 | 0.001254 | 4.178453 | Peroxidase | elongation zone | 12 h |
| HORVU.MOREX.r3.3HG0237990.1 | 0.885957 | 4.191674 | 5.140066 | 4.05E-06 | 0.001254 | 4.164326 | #N/A | elongation zone | 12 h |
| HORVU.MOREX.r3.4HG0400920.1 | 1.313937 | 4.093901 | 5.128498 | 4.22E-06 | 0.001254 | 4.12767 | Kinesin-like protein | elongation zone | 12 h |
| HORVU.MOREX.r3.1HG0067720.1 | 1.657385 | -2.10602 | 5.177819 | 3.54E-06 | 0.001254 | 4.104761 | #N/A | elongation zone | 12 h |
| HORVU.MOREX.r3.7HG0730510.1 | 1.492597 | -1.83116 | 5.13854 | 4.07E-06 | 0.001254 | 4.098203 | Germin-like protein 1 | elongation zone | 12 h |
| HORVU.MOREX.r3.3HG0320840.1 | -0.43634 | 7.266524 | -5.15112 | 3.89E-06 | 0.001254 | 4.083994 | Signal peptide peptidase family protein | elongation zone | 12 h |
| HORVU.MOREX.r3.5HG0537360.1 | 2.289942 | -1.21127 | 5.158664 | 3.79E-06 | 0.001254 | 3.999524 | transcription factor, putative (Protein of unknown function, DUF547) | elongation zone | 12 h |
| HORVU.MOREX.r3.2HG0153420.1 | 1.117386 | 6.452276 | 5.133567 | 4.14E-06 | 0.001254 | 3.993957 | #N/A | elongation zone | 12 h |
| HORVU.MOREX.r3.7HG0677230.1 | -1.48294 | 4.893498 | -5.14153 | 4.02E-06 | 0.001254 | 3.967566 | Receptor protein kinase, putative | elongation zone | 12 h |
| HORVU.MOREX.r3.4HG0346720.1 | -0.684 | 10.60861 | -5.1948 | 3.33E-06 | 0.001254 | 3.952302 | Protein disulfide-isomerase | elongation zone | 12 h |
| HORVU.MOREX.r3.2HG0207720.1 | 1.145726 | 0.894337 | 5.052416 | 5.52E-06 | 0.001514 | 3.89533 | Flavin-containing monooxygenase | elongation zone | 12 h |
| HORVU.MOREX.r3.4HG0354980.1 | 0.976228 | 2.412831 | 5.047897 | 5.61E-06 | 0.001514 | 3.872566 | #N/A | elongation zone | 12 h |
| HORVU.MOREX.r3.3HG0291490.1 | -0.96114 | 7.190585 | -5.08932 | 4.84E-06 | 0.00142 | 3.857396 | #N/A | elongation zone | 12 h |
| HORVU.MOREX.r3.4HG0402030.1 | -0.58859 | 5.321938 | -5.04782 | 5.61E-06 | 0.001514 | 3.852842 | Basic helix-loop-helix (BHLH) family transcription factor | elongation zone | 12 h |
| HORVU.MOREX.r3.6HG0550700.1 | -2.29686 | 1.442253 | -5.56153 | 8.93E-07 | 0.000675 | 3.844752 | F-box family protein | elongation zone | 12 h |
| HORVU.MOREX.r3.2HG0156610.1 | 1.995905 | -0.12875 | 5.034818 | 5.87E-06 | 0.001545 | 3.837548 | ERD (Early-responsive to dehydration stress) family protein | elongation zone | 12 h |
| HORVU.MOREX.r3.6HG0601830.1 | 1.749784 | -2.05956 | 5.070699 | 5.17E-06 | 0.001495 | 3.799878 | C2 calcium/lipid-binding and GRAM domain protein | elongation zone | 12 h |
| HORVU.MOREX.r3.5HG0532140.1 | 1.958298 | -1.00113 | 5.020638 | 6.17E-06 | 0.001545 | 3.791737 | Expansin | elongation zone | 12 h |
| HORVU.MOREX.r3.7HG0636370.1 | 1.410918 | 2.619393 | 5.030212 | 5.97E-06 | 0.001545 | 3.771879 | MYB-related transcription factor | elongation zone | 12 h |
| HORVU.MOREX.r3.2HG0152370.1 | -0.60828 | 4.701662 | -5.00819 | 6.45E-06 | 0.001557 | 3.73872 | MD-2-related lipid recognition domain-containing protein / ML domain-containing | elongation zone | 12 h |
| HORVU.MOREX.r3.3HG0257130.1 | -0.77606 | 4.342452 | -4.99665 | 6.72E-06 | 0.001584 | 3.713397 | Presenilin | elongation zone | 12 h |
| HORVU.MOREX.r3.3HG0295330.1 | -0.60561 | 7.613979 | -5.06587 | 5.26E-06 | 0.0015 | 3.69374 | Sugar phosphate/phosphate translocator | elongation zone | 12 h |
| HORVU.MOREX.r3.3HG0270400.1 | -0.88988 | 4.405759 | -4.99016 | 6.87E-06 | 0.001596 | 3.691468 | Six-bladed beta-propeller, TolB-like protein | elongation zone | 12 h |
| HORVU.MOREX.r3.1HG0080180.1 | 2.199377 | -2.35246 | 5.135327 | 4.11E-06 | 0.001254 | 3.68756 | Jacalin lectin family protein | elongation zone | 12 h |
| HORVU.MOREX.r3.5HG0479500.1 | 1.602566 | 1.850868 | 4.985214 | 6.99E-06 | 0.001596 | 3.665587 | Phosphonates import ATP-binding protein PhnC | elongation zone | 12 h |
| HORVU.MOREX.r3.2HG0112690.1 | 1.260737 | 1.247173 | 5.023686 | 6.11E-06 | 0.001545 | 3.658484 | Peroxidase | elongation zone | 12 h |

**Table S1** Continued.

| ID | logFC | AveExpr | t | P.Value | adj.P.Val | B | gene description | root zone | time point |
|---|---|---|---|---|---|---|---|---|---|
| HORVU.MOREX.r3.2HG0124850.1 | 1.659319 | -0.21834 | 4.978488 | 7.16E-06 | 0.001596 | 3.648927 | Dirigent protein | elongation zone | 12 h |
| HORVU.MOREX.r3.1HG0055380.1 | -0.6613 | 6.771434 | -5.00868 | 6.44E-06 | 0.001557 | 3.627812 | Methylsterol monooxygenase 1-2 | elongation zone | 12 h |
| HORVU.MOREX.r3.1HG0057990.1 | -0.8162 | 4.42891 | -4.96602 | 7.48E-06 | 0.001633 | 3.612353 | O-fucosyltransferase family protein | elongation zone | 12 h |
| HORVU.MOREX.r3.7HG0679350.1 | -0.80904 | 3.668408 | -4.97652 | 7.21E-06 | 0.001596 | 3.60021 | Leucine-rich repeat receptor-like protein kinase family | elongation zone | 12 h |
| HORVU.MOREX.r3.3HG0330980.1 | -0.43852 | 6.458067 | -4.99926 | 6.66E-06 | 0.001584 | 3.573263 | Protein IQ-DOMAIN 1 | elongation zone | 12 h |
| HORVU.MOREX.r3.2HG0218010.1 | -0.49035 | 8.228324 | -5.0271 | 6.03E-06 | 0.001545 | 3.57213 | Dolichyl-diphosphooligosaccharide--protein glycosyltransferase subunit 1 | elongation zone | 12 h |
| HORVU.MOREX.r3.3HG0240310.1 | -1.19501 | 6.614043 | -4.98186 | 7.08E-06 | 0.001596 | 3.565678 | DUF2921 family protein | elongation zone | 12 h |
| HORVU.MOREX.r3.4HG0394830.1 | 1.25434 | 0.997429 | 4.954452 | 7.79E-06 | 0.00167 | 3.560137 | Lecithin-cholesterol acyltransferase-like 1 | elongation zone | 12 h |
| HORVU.MOREX.r3.5HG0527460.1 | 1.406926 | 1.065374 | 4.975588 | 7.23E-06 | 0.001596 | 3.554157 | Pectin acetylesterase | elongation zone | 12 h |
| HORVU.MOREX.r3.4HG0408270.1 | 1.826412 | -2.27119 | 5.04187 | 5.73E-06 | 0.001526 | 3.551263 | LOB domain-containing protein, putative | elongation zone | 12 h |
| HORVU.MOREX.r3.1HG0002960.1 | 1.302114 | 0.278873 | 4.947682 | 7.98E-06 | 0.00167 | 3.513685 | NBS-LRR disease resistance protein, putative, expressed | elongation zone | 12 h |
| HORVU.MOREX.r3.2HG0112060.1 | -0.75031 | 4.798982 | -4.93592 | 8.31E-06 | 0.001723 | 3.505475 | Transcription factor | elongation zone | 12 h |
| HORVU.MOREX.r3.4HG0346830.1 | 2.077443 | -2.75621 | 5.236067 | 2.88E-06 | 0.001254 | 3.501348 | WEB family protein | elongation zone | 12 h |
| HORVU.MOREX.r3.1HG0073330.1 | -0.46147 | 5.588457 | -4.94924 | 7.93E-06 | 0.00167 | 3.477857 | Ubiquitin conjugating enzyme E2 | elongation zone | 12 h |
| HORVU.MOREX.r3.4HG0344370.1 | 1.830537 | -2.72257 | 5.132489 | 4.16E-06 | 0.001254 | 3.420227 | Glucuronoxylan 4-O-methyltransferase | elongation zone | 12 h |
| HORVU.MOREX.r3.3HG0303210.1 | 1.171749 | 1.990963 | 4.921456 | 8.75E-06 | 0.001789 | 3.405827 | Alpha/beta-Hydrolases superfamily protein | elongation zone | 12 h |
| HORVU.MOREX.r3.6HG0612180.1 | -0.82757 | 6.264663 | -4.91289 | 9.01E-06 | 0.001809 | 3.378267 | NHL repeat-containing protein-like | elongation zone | 12 h |
| HORVU.MOREX.r3.7HG0726790.1 | -0.48551 | 7.796481 | -4.95152 | 7.87E-06 | 0.00167 | 3.368215 | Dolichyl-diphosphooligosaccharide--protein glycosyltransferase 48 kDa subunit | elongation zone | 12 h |
| HORVU.MOREX.r3.1HG0066610.1 | 1.12183 | 2.119827 | 4.906046 | 9.23E-06 | 0.001821 | 3.363552 | Dirigent protein | elongation zone | 12 h |
| HORVU.MOREX.r3.1HG0059790.1 | -0.98655 | 4.636699 | -4.88498 | 9.94E-06 | 0.001875 | 3.35111 | Hydroxyproline-rich glycoprotein-like | elongation zone | 12 h |
| HORVU.MOREX.r3.3HG0246540.1 | -0.87824 | 4.407087 | -4.87722 | 1.02E-05 | 0.001875 | 3.321623 | Peptide-N4-(N-acetyl-beta-glucosaminyl)asparagine amidase A | elongation zone | 12 h |
| HORVU.MOREX.r3.6HG0631400.1 | 1.211369 | 2.336066 | 4.878503 | 1.02E-05 | 0.001875 | 3.316019 | Disease resistance protein RPM1 | elongation zone | 12 h |
| HORVU.MOREX.r3.4HG0397010.1 | 1.928371 | 0.173587 | 4.871128 | 1.04E-05 | 0.001875 | 3.292291 | Beta-glucosidase | elongation zone | 12 h |
| HORVU.MOREX.r3.4HG0400990.1 | -0.78055 | 5.330008 | -4.87219 | 1.04E-05 | 0.001875 | 3.272578 | 3-hydroxybenzoate 6-hydroxylase 1 | elongation zone | 12 h |
| HORVU.MOREX.r3.3HG0288940.1 | -0.71571 | 6.05706 | -4.88622 | 9.89E-06 | 0.001875 | 3.269132 | WRKY transcription factor, putative, expressed | elongation zone | 12 h |
| HORVU.MOREX.r3.6HG0631760.1 | -0.53342 | 5.489051 | -4.8577 | 1.09E-05 | 0.001875 | 3.210717 | Signal peptide peptidase-like protein | elongation zone | 12 h |
| HORVU.MOREX.r3.4HG0350900.1 | -0.4339 | 7.975994 | -4.91958 | 8.80E-06 | 0.001789 | 3.186687 | UDP-glucose 6-dehydrogenase | elongation zone | 12 h |
| HORVU.MOREX.r3.3HG0246370.1 | -0.65896 | 5.820127 | -4.84876 | 1.13E-05 | 0.001897 | 3.180684 | Mannosyltransferase | elongation zone | 12 h |
| HORVU.MOREX.r3.6HG0546160.1 | 1.077283 | 2.97611 | 4.853043 | 1.11E-05 | 0.001884 | 3.172305 | Acylamino-acid-releasing enzyme | elongation zone | 12 h |
| HORVU.MOREX.r3.6HG0604200.1 | -1.39831 | 0.762306 | -4.87926 | 1.01E-05 | 0.001875 | 3.170489 | transcription repressor | elongation zone | 12 h |
| HORVU.MOREX.r3.3HG0331030.1 | -0.47748 | 7.059382 | -4.8765 | 1.02E-05 | 0.001875 | 3.147658 | Chaperone protein dnaJ, putative | elongation zone | 12 h |
| HORVU.MOREX.r3.2HG0189670.1 | 1.512791 | 0.283223 | 4.823733 | 1.23E-05 | 0.001973 | 3.125547 | Aquaporin-like protein | elongation zone | 12 h |
| HORVU.MOREX.r3.1HG0092010.1 | 0.814228 | 6.335202 | 4.858143 | 1.09E-05 | 0.001875 | 3.123286 | #N/A | elongation zone | 12 h |
| HORVU.MOREX.r3.2HG0168070.1 | 1.543833 | 0.66133 | 4.859148 | 1.09E-05 | 0.001875 | 3.120066 | Peroxidase | elongation zone | 12 h |
| HORVU.MOREX.r3.3HG0293310.1 | 1.870295 | -2.57133 | 5.049878 | 5.57E-06 | 0.001514 | 3.113111 | Protein upstream of flc | elongation zone | 12 h |
| HORVU.MOREX.r3.3HG0299200.1 | -0.66194 | 5.845565 | -4.83275 | 1.19E-05 | 0.001973 | 3.105584 | Ion channel DMI1 | elongation zone | 12 h |
| HORVU.MOREX.r3.1HG0003090.7 | -1.88664 | 6.089425 | -4.80881 | 1.30E-05 | 0.00203 | 3.105354 | Thionin-2.2 | elongation zone | 12 h |
| HORVU.MOREX.r3.1HG0047100.1 | 1.150871 | 4.15495 | 4.81334 | 1.28E-05 | 0.002014 | 3.104576 | LanC-like protein | elongation zone | 12 h |
| HORVU.MOREX.r3.4HG0392230.1 | 1.382213 | 0.189187 | 4.824216 | 1.23E-05 | 0.001973 | 3.102065 | RNA-binding family protein, putative | elongation zone | 12 h |
| HORVU.MOREX.r3.5HG0490800.1 | 1.646216 | 0.125384 | 4.865824 | 1.06E-05 | 0.001875 | 3.09782 | IRK-interacting protein | elongation zone | 12 h |
| HORVU.MOREX.r3.3HG0327810.1 | -0.60339 | 7.008521 | -4.86578 | 1.06E-05 | 0.001875 | 3.076393 | Auxin efflux carrier protein-like | elongation zone | 12 h |

**Table S1** Continued.

| ID | logFC | AveExpr | t | P.Value | adj.P.Val | B | gene description | root zone | time point |
|---|---|---|---|---|---|---|---|---|---|
| HORVU.MOREX.r3.2HG0160130.1 | 0.793328 | 3.66281 | 4.825686 | 1.22E-05 | 0.001973 | 3.037468 | Basic helix-loop-helix transcription factor | elongation zone | 12 h |
| HORVU.MOREX.r3.1HG0024040.1 | -0.46705 | 9.734968 | -4.91081 | 9.08E-06 | 0.001809 | 3.036831 | Disulfide-isomerase | elongation zone | 12 h |
| HORVU.MOREX.r3.5HG0536490.1 | -1.51207 | 4.431144 | -4.7867 | 1.40E-05 | 0.002142 | 3.031499 | Acyl carrier protein | elongation zone | 12 h |
| HORVU.MOREX.r3.7HG0654480.1 | -1.06939 | 4.98012 | -4.7749 | 1.46E-05 | 0.002168 | 2.972611 | Histone H3 | elongation zone | 12 h |
| HORVU.MOREX.r3.7HG0719720.1 | -0.71868 | 5.938268 | -4.79057 | 1.38E-05 | 0.002131 | 2.971259 | Phospholipase D | elongation zone | 12 h |
| HORVU.MOREX.r3.1HG0073550.1 | 2.09899 | -1.6606 | 4.772611 | 1.47E-05 | 0.002169 | 2.958888 | Laccase | elongation zone | 12 h |
| HORVU.MOREX.r3.5HG0517640.1 | -0.94095 | 6.451472 | -4.7923 | 1.37E-05 | 0.002131 | 2.952024 | Glucan endo-1,3-beta-glucosidase, putative | elongation zone | 12 h |
| HORVU.MOREX.r3.4HG0369880.1 | -1.56807 | 2.555153 | -4.81491 | 1.27E-05 | 0.002014 | 2.946173 | Flavin-binding kelch domain F box protein | elongation zone | 12 h |
| HORVU.MOREX.r3.7HG0739910.1 | -0.67872 | 4.210714 | -4.7607 | 1.53E-05 | 0.002229 | 2.944522 | Disease resistance protein (NBS-LRR class) family | elongation zone | 12 h |
| HORVU.MOREX.r3.3HG0246670.1 | 1.756703 | -1.91783 | 4.873725 | 1.03E-05 | 0.001875 | 2.940686 | DUF4228 domain protein | elongation zone | 12 h |
| HORVU.MOREX.r3.4HG0334360.1 | 1.108594 | 4.121321 | 4.784683 | 1.41E-05 | 0.002142 | 2.923695 | Homeobox protein BEL1 like | elongation zone | 12 h |
| HORVU.MOREX.r3.2HG0217090.1 | 1.865928 | -0.85749 | 4.748702 | 1.60E-05 | 0.00227 | 2.914013 | C2 calcium/lipid-binding and GRAM domain protein | elongation zone | 12 h |
| HORVU.MOREX.r3.7HG0735840.1 | -1.26875 | 4.848168 | -4.73673 | 1.66E-05 | 0.002283 | 2.875179 | serine/arginine repetitive matrix-like protein | elongation zone | 12 h |
| HORVU.MOREX.r3.5HG0518560.1 | 2.229337 | 2.148281 | 4.738783 | 1.65E-05 | 0.002283 | 2.869386 | Leucine-rich repeat protein kinase family protein | elongation zone | 12 h |
| HORVU.MOREX.r3.1HG0066260.1 | -0.46587 | 9.175242 | -4.86211 | 1.08E-05 | 0.001875 | 2.865838 | Tubulin beta chain | elongation zone | 12 h |
| HORVU.MOREX.r3.3HG0275060.1 | 1.557367 | 0.275212 | 4.730526 | 1.70E-05 | 0.002284 | 2.850348 | Isoprenylcysteine alpha-carbonyl methylesterase ICME | elongation zone | 12 h |
| HORVU.MOREX.r3.5HG0528520.1 | -0.51911 | 4.086447 | -4.73398 | 1.68E-05 | 0.002283 | 2.849679 | Epoxide hydrolase 2 | elongation zone | 12 h |
| HORVU.MOREX.r3.1HG0072580.1 | 0.682639 | 5.490114 | 4.745939 | 1.61E-05 | 0.00227 | 2.838814 | Aminotransferase-related family protein | elongation zone | 12 h |
| HORVU.MOREX.r3.1HG0073170.1 | -0.89724 | 5.178736 | -4.72631 | 1.73E-05 | 0.002284 | 2.837119 | F-box protein | elongation zone | 12 h |
| HORVU.MOREX.r3.4HG0393870.1 | 1.684954 | -1.59561 | 4.72629 | 1.73E-05 | 0.002284 | 2.837034 | Peroxidase | elongation zone | 12 h |
| HORVU.MOREX.r3.1HG0086090.1 | 1.04016 | 3.988991 | 4.737984 | 1.66E-05 | 0.002283 | 2.82294 | Ubiquitin-conjugating enzyme E2 | elongation zone | 12 h |
| HORVU.MOREX.r3.2HG0172790.1 | -0.81425 | 7.313554 | -4.78148 | 1.43E-05 | 0.00215 | 2.80017 | Lipoxygenase domain-containing 1 | elongation zone | 12 h |
| HORVU.MOREX.r3.2HG0212570.1 | -0.52089 | 10.53927 | -4.85682 | 1.10E-05 | 0.001875 | 2.795552 | Calreticulin | elongation zone | 12 h |
| HORVU.MOREX.r3.7HG0736150.1 | 1.323904 | 8.171407 | 4.827574 | 1.21E-05 | 0.001973 | 2.788451 | Phosphatidylinositol transfer protein | elongation zone | 12 h |
| HORVU.MOREX.r3.6HG0568390.1 | 0.653533 | 4.337129 | 4.734731 | 1.68E-05 | 0.002283 | 2.786735 | Kinase family protein | elongation zone | 12 h |
| HORVU.MOREX.r3.6HG0597590.1 | -0.50688 | 6.568014 | -4.74996 | 1.59E-05 | 0.00227 | 2.765242 | Protein disulfide-isomerase | elongation zone | 12 h |
| HORVU.MOREX.r3.1HG0058390.1 | -1.20169 | 4.869952 | -4.70869 | 1.83E-05 | 0.002397 | 2.748496 | GATA transcription factor | elongation zone | 12 h |
| HORVU.MOREX.r3.5HG0508190.1 | -0.71198 | 8.08234 | -4.77714 | 1.45E-05 | 0.002167 | 2.747192 | 1-deoxy-D-xylulose 5-phosphate synthase 1 | elongation zone | 12 h |
| HORVU.MOREX.r3.7HG0687630.1 | -0.76226 | 7.969317 | -4.77064 | 1.48E-05 | 0.002169 | 2.741723 | Acyl carrier protein | elongation zone | 12 h |
| HORVU.MOREX.r3.3HG0235720.1 | -2.07204 | 3.057636 | -4.68947 | 1.96E-05 | 0.002467 | 2.725641 | FMN-dependent NADPH-azoreductase | elongation zone | 12 h |
| HORVU.MOREX.r3.7HG0706330.1 | -0.52817 | 4.543867 | -4.69332 | 1.93E-05 | 0.002465 | 2.722804 | TOM1-like protein 2 | elongation zone | 12 h |
| HORVU.MOREX.r3.4HG0391330.1 | -0.53457 | 6.457456 | -4.7282 | 1.71E-05 | 0.002284 | 2.71273 | Phosphomevalonate kinase | elongation zone | 12 h |
| HORVU.MOREX.r3.4HG0405400.1 | 1.842353 | 2.976906 | 4.681294 | 2.02E-05 | 0.002487 | 2.686026 | DUF936 family protein | elongation zone | 12 h |
| HORVU.MOREX.r3.4HG0412030.1 | -0.62225 | 4.498999 | -4.67814 | 2.04E-05 | 0.002487 | 2.683197 | vacuolar sorting-associated protein (DUF946) | elongation zone | 12 h |
| HORVU.MOREX.r3.7HG0647520.1 | -0.77042 | 6.632237 | -4.701 | 1.88E-05 | 0.00243 | 2.679955 | GDP-mannose 4,6 dehydratase-like protein | elongation zone | 12 h |
| HORVU.MOREX.r3.3HG0304490.1 | 1.3125 | -1.40069 | 4.691007 | 1.95E-05 | 0.002467 | 2.678233 | Myosin IC heavy chain | elongation zone | 12 h |
| HORVU.MOREX.r3.6HG0621690.1 | -0.74275 | 4.664753 | -4.66821 | 2.11E-05 | 0.002487 | 2.655533 | E3 ubiquitin-protein ligase | elongation zone | 12 h |
| HORVU.MOREX.r3.3HG0222830.1 | -0.41894 | 7.515891 | -4.75409 | 1.57E-05 | 0.002265 | 2.654079 | Heavy metal transport/detoxification superfamily protein | elongation zone | 12 h |
| HORVU.MOREX.r3.4HG0418530.1 | -1.47421 | 4.704049 | -4.6711 | 2.09E-05 | 0.002487 | 2.620025 | ATP-dependent chaperone ClpB | elongation zone | 12 h |
| HORVU.MOREX.r3.1HG0025870.1 | 0.758286 | 4.903164 | 4.674898 | 2.06E-05 | 0.002487 | 2.617095 | 6,7-dimethyl-8-ribityllumazine synthase | elongation zone | 12 h |
| HORVU.MOREX.r3.7HG0667620.1 | 1.518926 | 0.317173 | 4.686416 | 1.98E-05 | 0.002478 | 2.612572 | Peroxidase | elongation zone | 12 h |

**Table S1** Continued.

| ID | logFC | AveExpr | t | P.Value | adj.P.Val | B | gene description | root zone | time point |
|---|---|---|---|---|---|---|---|---|---|
| HORVU.MOREX.r3.4HG0394330.1 | 1.225618 | 1.645952 | 4.655838 | 2.20E-05 | 0.002522 | 2.600121 | Kinase family protein | elongation zone | 12 h |
| HORVU.MOREX.r3.5HG0429900.1 | 1.296121 | 4.093407 | 4.647836 | 2.26E-05 | 0.002546 | 2.59445 | #N/A | elongation zone | 12 h |
| HORVU.MOREX.r3.7HG0728220.1 | -0.52196 | 4.414112 | -4.6502 | 2.24E-05 | 0.002546 | 2.594088 | GATA transcription factor | elongation zone | 12 h |
| HORVU.MOREX.r3.6HG0620520.1 | 0.900079 | 3.09452 | 4.678573 | 2.03E-05 | 0.002487 | 2.590735 | Aminotransferase-related family protein | elongation zone | 12 h |
| HORVU.MOREX.r3.3HG0303410.1 | 0.784391 | 6.71103 | 4.745354 | 1.62E-05 | 0.00227 | 2.589263 | Respiratory burst oxidase-like protein | elongation zone | 12 h |
| HORVU.MOREX.r3.6HG0587140.1 | -0.70674 | 4.797451 | -4.64993 | 2.25E-05 | 0.002546 | 2.587235 | Transmembrane 9 superfamily member | elongation zone | 12 h |
| HORVU.MOREX.r3.2HG0200640.1 | -0.52165 | 7.509809 | -4.69925 | 1.89E-05 | 0.00243 | 2.578985 | Carboxypeptidase | elongation zone | 12 h |
| HORVU.MOREX.r3.3HG0330220.1 | 1.033583 | 6.363412 | 4.67461 | 2.06E-05 | 0.002487 | 2.566267 | GDSL esterase/lipase | elongation zone | 12 h |
| HORVU.MOREX.r3.2HG0199160.1 | 0.533679 | 6.474544 | 4.664283 | 2.14E-05 | 0.002506 | 2.56045 | Kinesin-like protein | elongation zone | 12 h |
| HORVU.MOREX.r3.4HG0379660.1 | 1.430816 | -1.00939 | 4.63139 | 2.39E-05 | 0.002546 | 2.542873 | Transcription repressor OFP17 | elongation zone | 12 h |
| HORVU.MOREX.r3.1HG0044530.1 | -0.47527 | 5.667948 | -4.65952 | 2.17E-05 | 0.002513 | 2.539596 | L-type lectin-domain containing receptor kinase VIII.2 | elongation zone | 12 h |
| HORVU.MOREX.r3.1HG0039670.1 | -0.33646 | 7.408425 | -4.6995 | 1.89E-05 | 0.00243 | 2.531827 | Calcium ion-binding protein | elongation zone | 12 h |
| HORVU.MOREX.r3.3HG0226110.1 | 1.862092 | -3.10127 | 5.008302 | 6.45E-06 | 0.001557 | 2.531262 | Heat shock protein | elongation zone | 12 h |
| HORVU.MOREX.r3.4HG0339890.1 | 1.685825 | -1.1907 | 4.628685 | 2.42E-05 | 0.002555 | 2.530358 | Pollen Ole e 1 allergen and extensin family protein, putative | elongation zone | 12 h |
| HORVU.MOREX.r3.3HG0218560.1 | 1.412843 | 0.309747 | 4.623788 | 2.46E-05 | 0.002585 | 2.514539 | Receptor-like protein kinase | elongation zone | 12 h |
| HORVU.MOREX.r3.1HG0087170.1 | -0.54378 | 5.424188 | -4.6443 | 2.29E-05 | 0.002546 | 2.508447 | 3-deoxy-manno-octulosonate cytidylyltransferase | elongation zone | 12 h |
| HORVU.MOREX.r3.5HG0449620.1 | 0.728408 | 4.972296 | 4.64018 | 2.32E-05 | 0.002546 | 2.506699 | weak chloroplast movement under blue light protein (DUF827) | elongation zone | 12 h |
| HORVU.MOREX.r3.2HG0178050.1 | 0.769183 | 4.229626 | 4.635321 | 2.36E-05 | 0.002546 | 2.500998 | F-box protein | elongation zone | 12 h |
| HORVU.MOREX.r3.1HG0075320.1 | 1.343706 | 0.461847 | 4.639994 | 2.32E-05 | 0.002546 | 2.490724 | Myb transcription factor | elongation zone | 12 h |
| HORVU.MOREX.r3.3HG0294970.2 | 0.852795 | 2.769527 | 4.66849 | 2.11E-05 | 0.002487 | 2.488079 | DNA-3-methyladenine glycosylase | elongation zone | 12 h |
| HORVU.MOREX.r3.6HG0607360.1 | -0.79761 | 4.502873 | -4.61164 | 2.56E-05 | 0.002647 | 2.480611 | WAT1-related protein | elongation zone | 12 h |
| HORVU.MOREX.r3.3HG0238180.1 | -0.57461 | 4.144587 | -4.6127 | 2.55E-05 | 0.002647 | 2.475081 | #N/A | elongation zone | 12 h |
| HORVU.MOREX.r3.7HG0720670.1 | -0.88297 | 2.879624 | -4.63468 | 2.37E-05 | 0.002546 | 2.471714 | Nucleoside diphosphate kinase | elongation zone | 12 h |
| HORVU.MOREX.r3.6HG0588350.1 | -0.71335 | 4.125371 | -4.60191 | 2.65E-05 | 0.002677 | 2.450539 | RING finger family protein | elongation zone | 12 h |
| HORVU.MOREX.r3.1HG0054190.1 | 1.88453 | -0.12557 | 4.610995 | 2.57E-05 | 0.002647 | 2.449717 | Expansin-like protein | elongation zone | 12 h |
| HORVU.MOREX.r3.1HG0085140.1 | -0.54429 | 6.163123 | -4.63673 | 2.35E-05 | 0.002546 | 2.446441 | Serine/threonine protein phosphatase 2A regulatory subunit B | elongation zone | 12 h |
| HORVU.MOREX.r3.4HG0380540.1 | 0.80901 | 3.365322 | 4.6036 | 2.63E-05 | 0.002677 | 2.442916 | Aquaporin-1 | elongation zone | 12 h |
| HORVU.MOREX.r3.4HG0417920.1 | 2.624034 | -1.69902 | 4.64325 | 2.30E-05 | 0.002546 | 2.440617 | Peroxidase | elongation zone | 12 h |
| HORVU.MOREX.r3.1HG0084970.1 | 1.277967 | 2.620282 | 4.602197 | 2.65E-05 | 0.002677 | 2.438186 | Phosphatidylserine synthase 2 | elongation zone | 12 h |
| HORVU.MOREX.r3.7HG0637760.1 | -0.63236 | 3.556847 | -4.5978 | 2.69E-05 | 0.002684 | 2.437452 | #N/A | elongation zone | 12 h |
| HORVU.MOREX.r3.5HG0533760.1 | 1.365941 | 3.713664 | 4.670418 | 2.09E-05 | 0.002487 | 2.434186 | Glycosyltransferase | elongation zone | 12 h |
| HORVU.MOREX.r3.7HG0717610.1 | 0.771857 | 4.008622 | 4.618543 | 2.50E-05 | 0.002619 | 2.433176 | Pollen-specific protein SF21 | elongation zone | 12 h |
| HORVU.MOREX.r3.4HG0413740.1 | 2.066764 | -1.83472 | 4.634178 | 2.37E-05 | 0.002546 | 2.421984 | 12-oxophytodienoate reductase-like protein | elongation zone | 12 h |
| HORVU.MOREX.r3.1HG0057780.1 | -0.55468 | 5.901425 | -4.63124 | 2.39E-05 | 0.002546 | 2.421229 | bZIP transcription factor (DUF630 and DUF632) | elongation zone | 12 h |
| HORVU.MOREX.r3.4HG0405780.1 | 0.639315 | 7.951421 | 4.713832 | 1.80E-05 | 0.00237 | 2.416231 | Receptor-like protein kinase | elongation zone | 12 h |
| HORVU.MOREX.r3.2HG0151920.1 | -0.93756 | 3.926991 | -4.5916 | 2.74E-05 | 0.002684 | 2.415362 | Retrovirus-related Pol polyprotein from transposon TNT 1-94 | elongation zone | 12 h |
| HORVU.MOREX.r3.7HG0638850.1 | -0.57145 | 7.446158 | -4.6779 | 2.04E-05 | 0.002487 | 2.409563 | Cellulose synthase-like protein | elongation zone | 12 h |
| HORVU.MOREX.r3.2HG0179560.1 | 1.197882 | 1.523229 | 4.585916 | 2.80E-05 | 0.00269 | 2.385472 | Nodulin-like / Major Facilitator Superfamily protein | elongation zone | 12 h |
| HORVU.MOREX.r3.1HG0035400.1 | 0.66107 | 4.194434 | 4.603968 | 2.63E-05 | 0.002677 | 2.375066 | Ras-related protein, expressed | elongation zone | 12 h |
| HORVU.MOREX.r3.6HG0608780.1 | -0.6072 | 7.822442 | -4.65956 | 2.17E-05 | 0.002513 | 2.374493 | Heat shock 70 kDa protein, putative | elongation zone | 12 h |
| HORVU.MOREX.r3.5HG0454660.1 | -0.41697 | 4.270815 | -4.59161 | 2.74E-05 | 0.002684 | 2.373581 | Alpha-(1,4)-fucosyltransferase | elongation zone | 12 h |

**Table S1** Continued.

| ID | logFC | AveExpr | t | P.Value | adj.P.Val | B | gene description | root zone | time point |
|---|---|---|---|---|---|---|---|---|---|
| HORVU.MOREX.r3.4HG0340200.1 | -1.00517 | 4.710006 | -4.5789 | 2.87E-05 | 0.002722 | 2.37174 | RING finger and CHY zinc finger protein | elongation zone | 12 h |
| HORVU.MOREX.r3.3HG0300950.1 | -0.81202 | 3.921584 | -4.58506 | 2.81E-05 | 0.00269 | 2.362893 | Amino acid transporter, putative | elongation zone | 12 h |
| HORVU.MOREX.r3.4HG0333450.1 | 0.761677 | 3.618973 | 4.569582 | 2.96E-05 | 0.002794 | 2.34585 | Type I inositol-1,4,5-trisphosphate 5-phosphatase CVP2 | elongation zone | 12 h |
| HORVU.MOREX.r3.1HG0031180.1 | -0.48647 | 4.951 | -4.58893 | 2.77E-05 | 0.002684 | 2.333401 | Hexosyltransferase | elongation zone | 12 h |
| HORVU.MOREX.r3.4HG0341800.1 | 0.835853 | 2.209476 | 4.598434 | 2.68E-05 | 0.002684 | 2.332246 | Receptor-like kinase | elongation zone | 12 h |
| HORVU.MOREX.r3.7HG0712850.1 | -0.51263 | 4.698391 | -4.56861 | 2.97E-05 | 0.002794 | 2.313984 | Fiber protein Fb34 | elongation zone | 12 h |
| HORVU.MOREX.r3.4HG0361680.1 | -1.28306 | 4.471436 | -4.55765 | 3.08E-05 | 0.002848 | 2.311686 | Nuclear distribution protein nudE-like 1 | elongation zone | 12 h |
| HORVU.MOREX.r3.3HG0254950.1 | 1.402237 | 1.01976 | 4.588473 | 2.77E-05 | 0.002684 | 2.311441 | Peroxidase | elongation zone | 12 h |
| HORVU.MOREX.r3.7HG0724060.1 | 1.597101 | 0.588187 | 4.553952 | 3.12E-05 | 0.002858 | 2.294427 | U-box domain-containing protein | elongation zone | 12 h |
| HORVU.MOREX.r3.7HG0677510.1 | -0.59865 | 5.971824 | -4.58905 | 2.77E-05 | 0.002684 | 2.28694 | Glycosyltransferase | elongation zone | 12 h |
| HORVU.MOREX.r3.3HG0287070.1 | -1.29173 | 6.192473 | -4.58916 | 2.77E-05 | 0.002684 | 2.280294 | Glycosyltransferase | elongation zone | 12 h |
| HORVU.MOREX.r3.3HG0312430.1 | 1.900038 | -0.09459 | 4.565209 | 3.00E-05 | 0.0028 | 2.272136 | E3 ubiquitin protein ligase drip2 | elongation zone | 12 h |
| HORVU.MOREX.r3.3HG0291030.1 | 0.844483 | 1.945138 | 4.589331 | 2.76E-05 | 0.002684 | 2.246325 | Glucan endo-1,3-beta-glucosidase 1 | elongation zone | 12 h |
| HORVU.MOREX.r3.1HG0002570.1 | 1.619696 | -1.7761 | 4.546267 | 3.20E-05 | 0.002921 | 2.243113 | GDSL esterase/lipase | elongation zone | 12 h |
| HORVU.MOREX.r3.4HG0349080.1 | 0.728322 | 3.881326 | 4.538024 | 3.29E-05 | 0.002965 | 2.242856 | CBS domain containing protein | elongation zone | 12 h |
| HORVU.MOREX.r3.2HG0134700.1 | -0.77196 | 2.881388 | -4.53404 | 3.34E-05 | 0.002967 | 2.238545 | Ras-related protein Rab-2 | elongation zone | 12 h |
| HORVU.MOREX.r3.3HG0230870.1 | 1.70205 | 0.065486 | 4.540328 | 3.27E-05 | 0.002958 | 2.211962 | Protein TRM32 | elongation zone | 12 h |
| HORVU.MOREX.r3.5HG0470700.1 | 1.62798 | -1.11013 | 4.522245 | 3.48E-05 | 0.003 | 2.202157 | Heavy metal transport/detoxification protein | elongation zone | 12 h |
| HORVU.MOREX.r3.6HG0552860.1 | -1.32499 | 6.65591 | -4.54001 | 3.27E-05 | 0.002958 | 2.199129 | Histone H2B | elongation zone | 12 h |
| HORVU.MOREX.r3.1HG0043520.1 | -0.50014 | 4.227432 | -4.53601 | 3.32E-05 | 0.002967 | 2.195043 | F-box protein | elongation zone | 12 h |
| HORVU.MOREX.r3.4HG0405920.1 | 1.531469 | 0.085137 | 4.527025 | 3.42E-05 | 0.002987 | 2.171278 | Hexosyltransferase | elongation zone | 12 h |
| HORVU.MOREX.r3.5HG0429180.1 | 1.518471 | 2.149739 | 4.5098 | 3.63E-05 | 0.003013 | 2.15744 | Glycosyltransferase | elongation zone | 12 h |
| HORVU.MOREX.r3.2HG0127530.1 | -1.48823 | 2.912589 | -4.52238 | 3.47E-05 | 0.003 | 2.152992 | Glucan endo-1,3-beta-glucosidase 3 | elongation zone | 12 h |
| HORVU.MOREX.r3.7HG0726770.1 | 1.195641 | 2.29225 | 4.512265 | 3.60E-05 | 0.003013 | 2.152855 | Glycine-rich protein | elongation zone | 12 h |
| HORVU.MOREX.r3.5HG0519810.1 | -0.55181 | 7.617925 | -4.58172 | 2.84E-05 | 0.002708 | 2.148553 | Diphosphomevalonate decarboxylase | elongation zone | 12 h |
| HORVU.MOREX.r3.1HG0052120.1 | -0.76288 | 3.548831 | -4.51071 | 3.62E-05 | 0.003013 | 2.132419 | GDP-mannose transporter | elongation zone | 12 h |
| HORVU.MOREX.r3.7HG0711890.1 | 1.311822 | 2.553034 | 4.513798 | 3.58E-05 | 0.003013 | 2.121476 | O-methyltransferase | elongation zone | 12 h |
| HORVU.MOREX.r3.6HG0622250.1 | 1.387109 | 2.510596 | 4.566791 | 2.99E-05 | 0.002798 | 2.119721 | Alpha/beta-hydrolase superfamily protein | elongation zone | 12 h |
| HORVU.MOREX.r3.2HG0175320.1 | -0.85077 | 4.008441 | -4.4887 | 3.90E-05 | 0.003173 | 2.091719 | Protein FAR1-RELATED SEQUENCE 6 | elongation zone | 12 h |
| HORVU.MOREX.r3.6HG0548670.1 | -1.16237 | 5.25208 | -4.49163 | 3.86E-05 | 0.003167 | 2.082928 | Histone H4 | elongation zone | 12 h |
| HORVU.MOREX.r3.6HG0582480.1 | 0.92359 | 2.53548 | 4.484342 | 3.95E-05 | 0.003202 | 2.081204 | Ethylene-responsive transcription factor-like protein | elongation zone | 12 h |
| HORVU.MOREX.r3.1HG0068010.1 | 1.610494 | -0.26596 | 4.520322 | 3.50E-05 | 0.003 | 2.079158 | Peroxidase | elongation zone | 12 h |
| HORVU.MOREX.r3.3HG0294960.1 | 1.551704 | -1.29923 | 4.489588 | 3.88E-05 | 0.003173 | 2.069261 | Eukaryotic translation initiation factor 3 subunit H | elongation zone | 12 h |
| HORVU.MOREX.r3.6HG0627570.1 | -0.74427 | 4.766957 | -4.48357 | 3.96E-05 | 0.003202 | 2.045827 | Immunoglobulin A1 protease autotransporter | elongation zone | 12 h |
| HORVU.MOREX.r3.2HG0213250.1 | -0.70922 | 6.347479 | -4.53005 | 3.39E-05 | 0.002987 | 2.040743 | Choline transporter-related family protein | elongation zone | 12 h |
| HORVU.MOREX.r3.6HG0559990.1 | 0.945935 | 5.045463 | 4.514801 | 3.57E-05 | 0.003013 | 2.039135 | Apoptosis-inducing factor-like protein A | elongation zone | 12 h |
| HORVU.MOREX.r3.3HG0301000.1 | 0.916534 | 4.492782 | 4.47807 | 4.04E-05 | 0.003217 | 2.033298 | ABC transporter B family-like protein | elongation zone | 12 h |
| HORVU.MOREX.r3.3HG0297330.1 | -0.52724 | 5.598322 | -4.50692 | 3.66E-05 | 0.00303 | 2.026252 | Receptor-like kinase | elongation zone | 12 h |
| HORVU.MOREX.r3.7HG0665400.1 | -0.78064 | 4.54259 | -4.46267 | 4.26E-05 | 0.003301 | 2.012797 | Mitochondrial 39S ribosomal protein L53 | elongation zone | 12 h |
| HORVU.MOREX.r3.7HG0724140.1 | 1.071058 | 4.661226 | 4.48254 | 3.98E-05 | 0.003202 | 2.009443 | FAD-binding Berberine family protein | elongation zone | 12 h |
| HORVU.MOREX.r3.5HG0438160.1 | -0.54153 | 5.336636 | -4.4766 | 4.06E-05 | 0.003217 | 2.008393 | #N/A | elongation zone | 12 h |

**Table S1** Continued.

| ID | logFC | AveExpr | t | P.Value | adj.P.Val | B | gene description | root zone | time point |
|---|---|---|---|---|---|---|---|---|---|
| HORVU.MOREX.r3.2HG0136670.1 | -1.9059 | 4.713901 | -4.52731 | 3.42E-05 | 0.002987 | 2.001968 | Cellulose synthase-like protein | elongation zone | 12 h |
| HORVU.MOREX.r3.5HG0494040.1 | 1.853113 | -3.00086 | 4.658566 | 2.18E-05 | 0.002513 | 1.982849 | Auxin efflux carrier component | elongation zone | 12 h |
| HORVU.MOREX.r3.7HG0732290.1 | -1.7091 | 2.965224 | -4.5195 | 3.51E-05 | 0.003 | 1.981645 | Dihydroflavonol-4-reductase | elongation zone | 12 h |
| HORVU.MOREX.r3.5HG0485800.1 | -0.5032 | 8.255741 | -4.55542 | 3.10E-05 | 0.002857 | 1.981036 | Protein disulfide isomerase | elongation zone | 12 h |
| HORVU.MOREX.r3.1HG0055240.1 | 2.288179 | -3.1472 | 4.824912 | 1.23E-05 | 0.001973 | 1.964779 | Acetyl-coenzyme A carboxylase carboxyl transferase subunit beta | elongation zone | 12 h |
| HORVU.MOREX.r3.5HG0519120.1 | 1.491953 | -0.8751 | 4.44632 | 4.50E-05 | 0.003412 | 1.964473 | CASP-like protein | elongation zone | 12 h |
| HORVU.MOREX.r3.5HG0485220.1 | -0.60757 | 5.159201 | -4.46301 | 4.25E-05 | 0.003301 | 1.962926 | Gamma-tubulin complex component | elongation zone | 12 h |
| HORVU.MOREX.r3.3HG0269960.1 | 0.562759 | 4.098981 | 4.472878 | 4.11E-05 | 0.003241 | 1.959505 | Protein kinase family protein | elongation zone | 12 h |
| HORVU.MOREX.r3.2HG0175380.1 | 1.656332 | 3.452964 | 4.514708 | 3.57E-05 | 0.003013 | 1.944856 | Sec14p-like phosphatidylinositol transfer family protein | elongation zone | 12 h |
| HORVU.MOREX.r3.7HG0661480.1 | -0.35586 | 7.365152 | -4.5208 | 3.49E-05 | 0.003 | 1.944699 | Alpha-amylase 2 | elongation zone | 12 h |
| HORVU.MOREX.r3.6HG0574950.1 | 0.652988 | 6.289342 | 4.534867 | 3.33E-05 | 0.002967 | 1.940068 | Receptor-like protein kinase | elongation zone | 12 h |
| HORVU.MOREX.r3.2HG0113990.1 | 2.25069 | -2.49708 | 4.63673 | 2.35E-05 | 0.002546 | 1.929458 | Calcium-dependent lipid-binding (CaLB domain) family protein | elongation zone | 12 h |
| HORVU.MOREX.r3.3HG0295020.1 | -0.56766 | 4.746066 | -4.45227 | 4.41E-05 | 0.003388 | 1.924047 | Polyadenylate-binding protein 1-B-binding protein | elongation zone | 12 h |
| HORVU.MOREX.r3.4HG0351750.1 | 2.021135 | -1.8238 | 4.440438 | 4.59E-05 | 0.003449 | 1.920398 | BTB/POZ domain-containing protein | elongation zone | 12 h |
| HORVU.MOREX.r3.2HG0181680.1 | 2.009347 | -1.90882 | 4.432275 | 4.72E-05 | 0.003494 | 1.915151 | CASP-like protein | elongation zone | 12 h |
| HORVU.MOREX.r3.7HG0674720.1 | 2.117226 | -1.46328 | 4.479718 | 4.02E-05 | 0.003217 | 1.90847 | Epoxide hydrolase 2 | elongation zone | 12 h |
| HORVU.MOREX.r3.3HG0293320.1 | 1.508086 | 0.363569 | 4.468004 | 4.18E-05 | 0.003274 | 1.907066 | ALK tyrosine kinase receptor | elongation zone | 12 h |
| HORVU.MOREX.r3.7HG0750170.1 | 1.566794 | 1.590474 | 4.43599 | 4.66E-05 | 0.003464 | 1.899452 | #N/A | elongation zone | 12 h |
| HORVU.MOREX.r3.1HG0051860.1 | -0.9201 | 5.340638 | -4.43662 | 4.65E-05 | 0.003464 | 1.89934 | Glutathione S-transferase | elongation zone | 12 h |
| HORVU.MOREX.r3.2HG0121830.1 | -0.33619 | 7.00528 | -4.50983 | 3.63E-05 | 0.003013 | 1.89168 | Calcium ion-binding protein | elongation zone | 12 h |
| HORVU.MOREX.r3.2HG0188300.1 | -0.60263 | 4.693992 | -4.42395 | 4.85E-05 | 0.003506 | 1.877333 | CsAtPR5 | elongation zone | 12 h |
| HORVU.MOREX.r3.7HG0704450.1 | 0.603667 | 5.126567 | 4.44854 | 4.46E-05 | 0.003412 | 1.859229 | Hexosyltransferase | elongation zone | 12 h |
| HORVU.MOREX.r3.1HG0082550.1 | 1.500181 | -2.24502 | 4.445407 | 4.51E-05 | 0.003412 | 1.843821 | Glycosyltransferase | elongation zone | 12 h |
| HORVU.MOREX.r3.1HG0079140.1 | 0.692729 | 4.435815 | 4.436066 | 4.66E-05 | 0.003464 | 1.822233 | Kinase family protein | elongation zone | 12 h |
| HORVU.MOREX.r3.5HG0489720.1 | -0.43318 | 6.28355 | -4.46225 | 4.26E-05 | 0.003301 | 1.81881 | WPP domain-interacting protein 1 | elongation zone | 12 h |
| HORVU.MOREX.r3.5HG0429930.1 | -1.05885 | 4.099119 | -4.39356 | 5.38E-05 | 0.003739 | 1.799766 | Protein DEK | elongation zone | 12 h |
| HORVU.MOREX.r3.2HG0104480.1 | -0.79134 | 3.899784 | -4.39202 | 5.40E-05 | 0.003746 | 1.799526 | F-box protein | elongation zone | 12 h |
| HORVU.MOREX.r3.2HG0168090.1 | -0.47994 | 5.385495 | -4.42483 | 4.84E-05 | 0.003506 | 1.799282 | Sphingomyelin synthase-like domain-containing protein | elongation zone | 12 h |
| HORVU.MOREX.r3.7HG0643230.1 | -0.7595 | 5.512465 | -4.42855 | 4.78E-05 | 0.003501 | 1.791268 | Transmembrane protein | elongation zone | 12 h |
| HORVU.MOREX.r3.7HG0660310.1 | -0.64109 | 4.34026 | -4.40531 | 5.17E-05 | 0.003656 | 1.787041 | Protein kinase | elongation zone | 12 h |
| HORVU.MOREX.r3.1HG0072910.2 | -0.5099 | 4.342105 | -4.38971 | 5.45E-05 | 0.003763 | 1.78251 | Serine/threonine-protein kinase ATM | elongation zone | 12 h |
| HORVU.MOREX.r3.6HG0542030.1 | 1.30864 | 1.995705 | 4.497603 | 3.78E-05 | 0.003115 | 1.772732 | Acid beta-fructofuranosidase | elongation zone | 12 h |
| HORVU.MOREX.r3.2HG0204940.1 | -0.66903 | 6.058217 | -4.41049 | 5.08E-05 | 0.00363 | 1.760745 | Crossover junction endonuclease EME1 | elongation zone | 12 h |
| HORVU.MOREX.r3.3HG0307120.1 | -0.63587 | 8.364634 | -4.47696 | 4.06E-05 | 0.003217 | 1.752829 | S-adenosyl-L-methionine-dependent methyltransferases superfamily protein | elongation zone | 12 h |
| HORVU.MOREX.r3.5HG0497900.1 | 0.996187 | 3.680571 | 4.402332 | 5.22E-05 | 0.00368 | 1.748422 | Rhomboid-like protein | elongation zone | 12 h |
| HORVU.MOREX.r3.4HG0417320.1 | 1.588295 | 1.098725 | 4.424929 | 4.84E-05 | 0.003506 | 1.735275 | Remorin | elongation zone | 12 h |
| HORVU.MOREX.r3.1HG0082630.1 | -0.47086 | 5.023124 | -4.39468 | 5.36E-05 | 0.003738 | 1.722179 | Glycosyltransferase | elongation zone | 12 h |
| HORVU.MOREX.r3.7HG0679050.1 | 0.722855 | 4.002868 | 4.371043 | 5.80E-05 | 0.003852 | 1.720978 | Upstream activation factor subunit spp27 | elongation zone | 12 h |
| HORVU.MOREX.r3.4HG0385880.1 | -0.93223 | 3.891411 | -4.36053 | 6.01E-05 | 0.00391 | 1.695249 | Pentatricopeptide repeat-containing protein | elongation zone | 12 h |
| HORVU.MOREX.r3.2HG0192100.1 | 1.683259 | 1.501774 | 4.362834 | 5.96E-05 | 0.003896 | 1.694509 | Serine/threonine-protein kinase | elongation zone | 12 h |
| HORVU.MOREX.r3.3HG0330200.1 | 1.233486 | 1.169812 | 4.40707 | 5.14E-05 | 0.003647 | 1.68287 | Aquaporin | elongation zone | 12 h |

**Table S1** Continued.

| ID | logFC | AveExpr | t | P.Value | adj.P.Val | B | gene description | root zone | time point |
|---|---|---|---|---|---|---|---|---|---|
| HORVU.MOREX.r3.3HG0309410.1 | -1.35125 | 3.365238 | -4.3764 | 5.70E-05 | 0.003832 | 1.672083 | Subtilisin-like protease | elongation zone | 12 h |
| HORVU.MOREX.r3.4HG0406060.1 | 2.606463 | -1.45049 | 4.472125 | 4.12E-05 | 0.003241 | 1.662728 | Agmatine coumaroyltransferase-2 | elongation zone | 12 h |
| HORVU.MOREX.r3.6HG0542040.1 | 1.328814 | 1.468748 | 4.444699 | 4.52E-05 | 0.003412 | 1.659916 | Acid beta-fructofuranosidase | elongation zone | 12 h |
| HORVU.MOREX.r3.3HG0257840.1 | 1.389722 | 0.153713 | 4.379772 | 5.63E-05 | 0.003829 | 1.658474 | S-acyltransferase | elongation zone | 12 h |
| HORVU.MOREX.r3.4HG0379400.1 | 1.067998 | 1.850081 | 4.428695 | 4.77E-05 | 0.003501 | 1.656569 | Protein kinase-like protein | elongation zone | 12 h |
| HORVU.MOREX.r3.3HG0317780.1 | 1.216718 | 0.361248 | 4.345065 | 6.33E-05 | 0.00401 | 1.647161 | 2-oxoglutarate (2OG) and Fe(II)-dependent oxygenase superfamily protein | elongation zone | 12 h |
| HORVU.MOREX.r3.7HG0696770.1 | -0.33854 | 5.470938 | -4.37909 | 5.64E-05 | 0.003829 | 1.646095 | Protein kinase | elongation zone | 12 h |
| HORVU.MOREX.r3.5HG0532960.1 | -0.55076 | 5.01578 | -4.36438 | 5.93E-05 | 0.003888 | 1.637258 | ER lumen protein-retaining receptor | elongation zone | 12 h |
| HORVU.MOREX.r3.5HG0447720.1 | -0.63709 | 7.79832 | -4.41897 | 4.93E-05 | 0.003553 | 1.628473 | Dicarboxylate transporter 2.1, chloroplastic | elongation zone | 12 h |
| HORVU.MOREX.r3.6HG0581010.1 | -0.79523 | 4.49799 | -4.33694 | 6.50E-05 | 0.004044 | 1.624034 | Poly(A) polymerase | elongation zone | 12 h |
| HORVU.MOREX.r3.4HG0411960.1 | -0.9173 | 3.220574 | -4.33341 | 6.58E-05 | 0.004054 | 1.622814 | #N/A | elongation zone | 12 h |
| HORVU.MOREX.r3.1HG0086010.1 | -0.6194 | 5.131587 | -4.3508 | 6.21E-05 | 0.003971 | 1.616267 | TSA: Wollemia nobilis Ref_Wollemi_Transcript_14910_3291 transcribed RNA sequ | elongation zone | 12 h |
| HORVU.MOREX.r3.4HG0390770.1 | 1.08226 | 3.149264 | 4.32974 | 6.66E-05 | 0.004056 | 1.611764 | Regulator of Vps4 activity in the MVB pathway protein, putative | elongation zone | 12 h |
| HORVU.MOREX.r3.2HG0107470.1 | -0.65221 | 4.149979 | -4.33169 | 6.62E-05 | 0.004054 | 1.609288 | Pectate lyase | elongation zone | 12 h |
| HORVU.MOREX.r3.7HG0651110.1 | -0.43342 | 7.468711 | -4.41758 | 4.96E-05 | 0.003557 | 1.607905 | AT4G29520-like protein | elongation zone | 12 h |
| HORVU.MOREX.r3.6HG0600000.1 | 1.481651 | -0.21905 | 4.338054 | 6.48E-05 | 0.004041 | 1.604949 | Blue copper protein | elongation zone | 12 h |
| HORVU.MOREX.r3.3HG0288960.1 | 1.24567 | 4.129323 | 4.368993 | 5.84E-05 | 0.003853 | 1.597375 | Cellulose synthase | elongation zone | 12 h |
| HORVU.MOREX.r3.7HG0654410.1 | -0.88402 | 6.620821 | -4.37492 | 5.72E-05 | 0.003839 | 1.591348 | Histone H3 | elongation zone | 12 h |
| HORVU.MOREX.r3.4HG0349530.1 | -0.44049 | 4.965038 | -4.34796 | 6.27E-05 | 0.003993 | 1.5837 | Tubby-like F-box protein | elongation zone | 12 h |
| HORVU.MOREX.r3.1HG0091870.1 | -0.61113 | 5.262379 | -4.34488 | 6.33E-05 | 0.00401 | 1.576011 | Protein kinase superfamily protein | elongation zone | 12 h |
| HORVU.MOREX.r3.5HG0522060.1 | -0.80368 | 4.084626 | -4.31502 | 7.00E-05 | 0.004083 | 1.567232 | #N/A | elongation zone | 12 h |
| HORVU.MOREX.r3.7HG0678160.1 | -0.64516 | 6.566933 | -4.35952 | 6.03E-05 | 0.00391 | 1.563259 | Disease resistance protein | elongation zone | 12 h |
| HORVU.MOREX.r3.4HG0383670.1 | 1.294062 | 2.012771 | 4.341282 | 6.41E-05 | 0.004025 | 1.550733 | Protein SCAR2 | elongation zone | 12 h |
| HORVU.MOREX.r3.1HG0080760.1 | -1.07021 | 7.93903 | -4.3776 | 5.67E-05 | 0.003829 | 1.541427 | Histone H3 | elongation zone | 12 h |
| HORVU.MOREX.r3.1HG0050700.1 | 1.413469 | 2.994863 | 4.332111 | 6.61E-05 | 0.004054 | 1.540069 | ATP-dependent zinc metalloprotease FtsH | elongation zone | 12 h |
| HORVU.MOREX.r3.1HG0089220.1 | -1.07794 | 0.50855 | -4.341 | 6.41E-05 | 0.004025 | 1.531276 | Cell surface antigen I/II | elongation zone | 12 h |
| HORVU.MOREX.r3.6HG0543780.1 | -0.94232 | 7.397821 | -4.37157 | 5.79E-05 | 0.003852 | 1.525851 | Histone H2B | elongation zone | 12 h |
| HORVU.MOREX.r3.1HG0092810.1 | -1.15487 | 7.116497 | -4.33209 | 6.61E-05 | 0.004054 | 1.515014 | Late embryogenesis abundant protein Lea14 | elongation zone | 12 h |
| HORVU.MOREX.r3.2HG0207790.1 | 1.619224 | -2.59422 | 4.445222 | 4.52E-05 | 0.003412 | 1.508772 | Serine/threonine-protein kinase | elongation zone | 12 h |
| HORVU.MOREX.r3.5HG0489130.1 | -1.38223 | 6.654428 | -4.38072 | 5.61E-05 | 0.003829 | 1.506566 | Receptor protein kinase, putative | elongation zone | 12 h |
| HORVU.MOREX.r3.1HG0006400.1 | 0.823963 | 6.387241 | 4.409322 | 5.10E-05 | 0.003632 | 1.500205 | Beta-1,3-galactosyltransferase-like protein | elongation zone | 12 h |
| HORVU.MOREX.r3.7HG0666340.1 | -0.66114 | 4.728876 | -4.3042 | 7.26E-05 | 0.004172 | 1.497311 | Hexosyltransferase | elongation zone | 12 h |
| HORVU.MOREX.r3.6HG0609560.1 | -1.12616 | 3.881425 | -4.29089 | 7.59E-05 | 0.00423 | 1.493876 | Alpha/beta hydrolase | elongation zone | 12 h |
| HORVU.MOREX.r3.1HG0095210.1 | -0.34876 | 8.682671 | -4.42742 | 4.80E-05 | 0.003502 | 1.493832 | Nucleoside diphosphate kinase | elongation zone | 12 h |
| HORVU.MOREX.r3.7HG0751300.1 | 1.477716 | -1.39316 | 4.291546 | 7.57E-05 | 0.00423 | 1.493222 | Phosphatidylinositol N-acetyglucosaminlytransferase subunit P-like protein | elongation zone | 12 h |
| HORVU.MOREX.r3.1HG0056620.1 | 1.829984 | 0.189162 | 4.350602 | 6.21E-05 | 0.003971 | 1.493145 | Expansin protein | elongation zone | 12 h |
| HORVU.MOREX.r3.1HG0047250.1 | -0.86174 | 2.716377 | -4.32164 | 6.84E-05 | 0.004083 | 1.490614 | Major facilitator superfamily transporter | elongation zone | 12 h |
| HORVU.MOREX.r3.3HG0280630.1 | -0.41773 | 5.914468 | -4.33891 | 6.46E-05 | 0.004041 | 1.484714 | Glycosylphosphatidylinositol anchor attachment 1 protein | elongation zone | 12 h |
| HORVU.MOREX.r3.6HG0557920.1 | 1.069601 | 4.693871 | 4.314849 | 7.00E-05 | 0.004083 | 1.483056 | Auxin response factor | elongation zone | 12 h |
| HORVU.MOREX.r3.4HG0415720.1 | -0.71083 | 4.208548 | -4.30183 | 7.31E-05 | 0.00418 | 1.470103 | Endoglucanase | elongation zone | 12 h |
| HORVU.MOREX.r3.2HG0170560.1 | -1.21823 | 3.704875 | -4.32875 | 6.68E-05 | 0.004058 | 1.458503 | Ras-like protein | elongation zone | 12 h |

**Table S1** Continued.

| ID | logFC | AveExpr | t | P.Value | adj.P.Val | B | gene description | root zone | time point |
|---|---|---|---|---|---|---|---|---|---|
| HORVU.MOREX.r3.5HG0420410.1 | 0.550715 | 5.049102 | 4.321666 | 6.84E-05 | 0.004083 | 1.448437 | Ubiquitin-conjugating enzyme E2 | elongation zone | 12 h |
| HORVU.MOREX.r3.3HG0305270.1 | -0.62745 | 5.993243 | -4.31568 | 6.98E-05 | 0.004083 | 1.445082 | U-box domain-containing family protein | elongation zone | 12 h |
| HORVU.MOREX.r3.4HG0398090.1 | 0.814604 | 4.062819 | 4.318349 | 6.92E-05 | 0.004083 | 1.444254 | Syntaxin protein | elongation zone | 12 h |
| HORVU.MOREX.r3.5HG0478450.1 | -0.41743 | 5.597721 | -4.31729 | 6.94E-05 | 0.004083 | 1.438873 | AUGMIN subunit 5 | elongation zone | 12 h |
| HORVU.MOREX.r3.4HG0341050.1 | 1.507336 | 0.610576 | 4.2742 | 8.02E-05 | 0.004331 | 1.437845 | Protein DETOXIFICATION | elongation zone | 12 h |
| HORVU.MOREX.r3.6HG0542970.1 | -1.09532 | 4.467056 | -4.27183 | 8.08E-05 | 0.004342 | 1.42845 | Receptor-like kinase | elongation zone | 12 h |
| HORVU.MOREX.r3.7HG0662650.1 | -1.00965 | 6.460162 | -4.3167 | 6.96E-05 | 0.004083 | 1.42349 | Histone H3 | elongation zone | 12 h |
| HORVU.MOREX.r3.4HG0402730.1 | 1.837201 | -1.08162 | 4.268691 | 8.17E-05 | 0.004344 | 1.413591 | Homeobox protein BEL1 like | elongation zone | 12 h |
| HORVU.MOREX.r3.1HG0054170.1 | 1.561465 | 0.15708 | 4.295755 | 7.46E-05 | 0.004207 | 1.411281 | Expansin-like protein | elongation zone | 12 h |
| HORVU.MOREX.r3.4HG0338070.1 | 1.073526 | 1.603513 | 4.319566 | 6.89E-05 | 0.004083 | 1.410607 | Kinase family protein | elongation zone | 12 h |
| HORVU.MOREX.r3.5HG0461280.1 | 0.591328 | 6.01476 | 4.318145 | 6.93E-05 | 0.004083 | 1.409356 | Calcium-binding EF-hand protein | elongation zone | 12 h |
| HORVU.MOREX.r3.3HG0247440.1 | -0.91441 | 2.109103 | -4.31397 | 7.02E-05 | 0.004083 | 1.407777 | Serine/threonine-protein kinase | elongation zone | 12 h |
| HORVU.MOREX.r3.6HG0542050.1 | 1.554948 | 1.403897 | 4.367007 | 5.88E-05 | 0.003866 | 1.402642 | Acid beta-fructofuranosidase | elongation zone | 12 h |
| HORVU.MOREX.r3.5HG0511090.1 | 0.696828 | 5.583411 | 4.297216 | 7.43E-05 | 0.004198 | 1.400671 | Phytochrome | elongation zone | 12 h |
| HORVU.MOREX.r3.7HG0738770.1 | 1.211448 | 1.993948 | 4.358917 | 6.04E-05 | 0.00391 | 1.400567 | Receptor-like kinase | elongation zone | 12 h |
| HORVU.MOREX.r3.3HG0285440.1 | 1.491529 | -1.28505 | 4.272794 | 8.06E-05 | 0.00434 | 1.400422 | #N/A | elongation zone | 12 h |
| HORVU.MOREX.r3.3HG0298080.1 | 1.086173 | 1.480621 | 4.268551 | 8.17E-05 | 0.004344 | 1.400229 | Amino acid permease | elongation zone | 12 h |
| HORVU.MOREX.r3.3HG0280060.1 | 2.094624 | -2.86395 | 4.45809 | 4.32E-05 | 0.003335 | 1.388218 | Heavy metal transport/detoxification superfamily protein | elongation zone | 12 h |
| HORVU.MOREX.r3.2HG0110840.1 | -0.62257 | 6.010911 | -4.28947 | 7.62E-05 | 0.004238 | 1.38683 | 2-dehydro-3-deoxyphosphooctonate aldolase | elongation zone | 12 h |
| HORVU.MOREX.r3.7HG0709110.1 | -1.56963 | 3.79338 | -4.25991 | 8.41E-05 | 0.004437 | 1.380343 | Histone H3 | elongation zone | 12 h |
| HORVU.MOREX.r3.3HG0329950.1 | 1.631461 | 1.75889 | 4.291207 | 7.58E-05 | 0.00423 | 1.378215 | Pectin acetylesterase | elongation zone | 12 h |
| HORVU.MOREX.r3.1HG0019410.1 | -0.96466 | 2.787892 | -4.28398 | 7.76E-05 | 0.004293 | 1.372074 | Transducin/WD-like repeat-protein | elongation zone | 12 h |
| HORVU.MOREX.r3.4HG0403070.1 | -0.672 | 6.7424 | -4.30813 | 7.16E-05 | 0.004139 | 1.366264 | Serine/threonine-protein phosphatase | elongation zone | 12 h |
| HORVU.MOREX.r3.2HG0205750.1 | -0.64261 | 6.410155 | -4.30409 | 7.26E-05 | 0.004172 | 1.354776 | 3-oxoacyl-[acyl-carrier-protein] synthase 3 | elongation zone | 12 h |
| HORVU.MOREX.r3.5HG0511450.1 | 0.434001 | 5.650127 | 4.298297 | 7.40E-05 | 0.004195 | 1.352635 | ABC transporter B family protein | elongation zone | 12 h |
| HORVU.MOREX.r3.2HG0172730.1 | -0.53292 | 7.548613 | -4.33352 | 6.58E-05 | 0.004054 | 1.351163 | Alpha-ketoglutarate-dependent dioxygenase alkB-like protein 2 | elongation zone | 12 h |
| HORVU.MOREX.r3.1HG0072540.1 | -1.20754 | 5.751313 | -4.2615 | 8.37E-05 | 0.004425 | 1.349429 | Histone H4 | elongation zone | 12 h |
| HORVU.MOREX.r3.6HG0585240.1 | 1.031982 | 0.751154 | 4.282334 | 7.81E-05 | 0.004294 | 1.337296 | Kinase superfamily protein | elongation zone | 12 h |
| HORVU.MOREX.r3.7HG0674440.1 | -0.55543 | 5.321799 | -4.28011 | 7.86E-05 | 0.004314 | 1.328447 | Aluminum activated malate transporter family protein | elongation zone | 12 h |
| HORVU.MOREX.r3.3HG0221180.1 | 1.205783 | 2.379805 | 4.342396 | 6.38E-05 | 0.004025 | 1.327074 | Glycosyltransferase | elongation zone | 12 h |
| HORVU.MOREX.r3.2HG0209300.1 | -1.18247 | 4.575756 | -4.23881 | 9.02E-05 | 0.00464 | 1.324291 | Sugar transporter, putative | elongation zone | 12 h |
| HORVU.MOREX.r3.1HG0068370.1 | -1.27704 | 4.947166 | -4.23626 | 9.10E-05 | 0.004668 | 1.313667 | #N/A | elongation zone | 12 h |
| HORVU.MOREX.r3.7HG0718190.1 | 0.873213 | 6.009061 | 4.35186 | 6.19E-05 | 0.003971 | 1.310351 | ABC transporter family protein | elongation zone | 12 h |
| HORVU.MOREX.r3.6HG0634200.1 | 1.283575 | 0.006824 | 4.259056 | 8.43E-05 | 0.004438 | 1.309729 | Syntaxin | elongation zone | 12 h |
| HORVU.MOREX.r3.2HG0183740.1 | -1.21852 | 5.211569 | -4.23012 | 9.29E-05 | 0.004702 | 1.309581 | Sugar transporter, putative | elongation zone | 12 h |
| HORVU.MOREX.r3.7HG0677580.1 | -1.49919 | 5.782253 | -4.23991 | 8.99E-05 | 0.004635 | 1.30143 | Protein O-linked-mannose beta-1,4-N-acetylglucosaminyltransferase 2 | elongation zone | 12 h |
| HORVU.MOREX.r3.1HG0079870.1 | -1.13274 | 5.830169 | -4.27854 | 7.90E-05 | 0.004314 | 1.299823 | Pectin lyase-like superfamily protein | elongation zone | 12 h |
| HORVU.MOREX.r3.5HG0497490.1 | 1.76946 | -1.08137 | 4.224964 | 9.45E-05 | 0.004721 | 1.29578 | SPX domain-like protein | elongation zone | 12 h |
| HORVU.MOREX.r3.2HG0210420.1 | -0.57174 | 7.910637 | -4.322 | 6.84E-05 | 0.004083 | 1.29342 | Dolichyl-diphosphooligosaccharide--protein glycosyltransferase subunit STT3 | elongation zone | 12 h |
| HORVU.MOREX.r3.7HG0648620.1 | 1.421385 | -1.54284 | 4.233649 | 9.18E-05 | 0.004697 | 1.288073 | 2-oxoglutarate (2OG) and Fe(II)-dependent oxygenase superfamily protein | elongation zone | 12 h |
| HORVU.MOREX.r3.4HG0417410.1 | -1.68956 | 0.915973 | -4.3783 | 5.66E-05 | 0.003829 | 1.283171 | Cold regulated protein 27 | elongation zone | 12 h |

**Table S1** Continued.

| ID | logFC | AveExpr | t | P.Value | adj.P.Val | B | gene description | root zone | time point |
|---|---|---|---|---|---|---|---|---|---|
| HORVU.MOREX.r3.6HG0610500.1 | 1.092672 | 1.298618 | 4.302095 | 7.31E-05 | 0.00418 | 1.281725 | Eukaryotic aspartyl protease family protein | elongation zone | 12 h |
| HORVU.MOREX.r3.7HG0667610.1 | 1.529334 | 3.317002 | 4.283131 | 7.78E-05 | 0.004294 | 1.279977 | Peroxidase | elongation zone | 12 h |
| HORVU.MOREX.r3.5HG0524350.1 | 2.055719 | -1.05304 | 4.218576 | 9.65E-05 | 0.004752 | 1.277835 | Protein MIZU-KUSSEI 1 | elongation zone | 12 h |
| HORVU.MOREX.r3.1HG0041280.1 | 0.980912 | 4.323201 | 4.299521 | 7.37E-05 | 0.004195 | 1.271428 | Cellulose synthase | elongation zone | 12 h |
| HORVU.MOREX.r3.7HG0641750.1 | 2.979122 | -2.73901 | 4.369425 | 5.83E-05 | 0.003853 | 1.265579 | Aspartic proteinase nepenthesin-2 | elongation zone | 12 h |
| HORVU.MOREX.r3.3HG0327170.1 | -0.55247 | 7.966567 | -4.31388 | 7.02E-05 | 0.004083 | 1.265145 | Bifunctional polymyxin resistance arnA protein | elongation zone | 12 h |
| HORVU.MOREX.r3.5HG0528280.1 | 0.512705 | 5.95615 | 4.276712 | 7.95E-05 | 0.004329 | 1.244628 | Organic solute transporter-like | elongation zone | 12 h |
| HORVU.MOREX.r3.5HG0459270.1 | -0.84855 | 3.649219 | -4.20887 | 9.96E-05 | 0.004816 | 1.241247 | Cation calcium exchanger | elongation zone | 12 h |
| HORVU.MOREX.r3.6HG0572960.1 | -1.57567 | 1.634383 | -4.38224 | 5.58E-05 | 0.003829 | 1.23804 | Chlorophyll a-b binding protein, chloroplastic | elongation zone | 12 h |
| HORVU.MOREX.r3.1HG0018810.1 | -0.96919 | 5.360039 | -4.2115 | 9.88E-05 | 0.004807 | 1.232275 | #N/A | elongation zone | 12 h |
| HORVU.MOREX.r3.2HG0157260.1 | 1.445115 | -0.94189 | 4.207568 | 0.0001 | 0.004816 | 1.230818 | NAD(P)H dehydrogenase (Quinone) | elongation zone | 12 h |
| HORVU.MOREX.r3.7HG0667280.1 | -0.53944 | 4.293106 | -4.21833 | 9.66E-05 | 0.004752 | 1.225249 | Hexosyltransferase | elongation zone | 12 h |
| HORVU.MOREX.r3.3HG0301750.1 | -0.62386 | 9.943131 | -4.37094 | 5.80E-05 | 0.003852 | 1.220849 | Alpha-1,4 glucan phosphorylase | elongation zone | 12 h |
| HORVU.MOREX.r3.4HG0400740.1 | -0.28746 | 8.244974 | -4.31653 | 6.96E-05 | 0.004083 | 1.218679 | Cell division cycle protein 48-like protein | elongation zone | 12 h |
| HORVU.MOREX.r3.7HG0721690.1 | 1.257552 | 0.537157 | 4.224495 | 9.46E-05 | 0.004721 | 1.213527 | Endoglucanase | elongation zone | 12 h |
| HORVU.MOREX.r3.4HG0412790.1 | 0.733626 | 3.163327 | 4.207108 | 0.0001 | 0.004816 | 1.202035 | Remorin | elongation zone | 12 h |
| HORVU.MOREX.r3.1HG0053530.1 | -1.45226 | 2.523226 | -4.38005 | 5.63E-05 | 0.003829 | 1.197995 | Glycerol-3-phosphate acyltransferase, chloroplastic | elongation zone | 12 h |
| HORVU.MOREX.r3.2HG0115930.1 | -0.65234 | 9.174331 | -4.32974 | 6.66E-05 | 0.004056 | 1.193772 | 4-alpha-glucanotransferase | elongation zone | 12 h |
| HORVU.MOREX.r3.7HG0662160.1 | -0.58652 | 5.721315 | -4.22042 | 9.59E-05 | 0.004752 | 1.191973 | Phosphopantothenoylcysteine decarboxylase | elongation zone | 12 h |
| HORVU.MOREX.r3.2HG0117030.1 | 0.722651 | 7.823219 | 4.298367 | 7.40E-05 | 0.004195 | 1.188672 | Glutamate synthase, putative | elongation zone | 12 h |
| HORVU.MOREX.r3.3HG0249400.1 | 1.212766 | 4.937663 | 4.310132 | 7.11E-05 | 0.004123 | 1.184833 | Plasma membrane-associated cation-binding protein 1 | elongation zone | 12 h |
| HORVU.MOREX.r3.5HG0530750.1 | -0.4972 | 5.706474 | -4.22831 | 9.34E-05 | 0.004702 | 1.182274 | Post-GPI attachment to proteins factor 3 | elongation zone | 12 h |
| HORVU.MOREX.r3.5HG0498320.1 | 0.421419 | 5.332211 | 4.228061 | 9.35E-05 | 0.004702 | 1.180629 | Zinc finger, B-box | elongation zone | 12 h |
| HORVU.MOREX.r3.5HG0432360.1 | 0.637144 | 6.047915 | 4.269128 | 8.16E-05 | 0.004344 | 1.170388 | Inter-alpha-trypsin inhibitor heavy chain, putative | elongation zone | 12 h |
| HORVU.MOREX.r3.5HG0420150.1 | -0.441 | 6.344897 | -4.24068 | 8.97E-05 | 0.004634 | 1.158981 | Core-2/I-branching beta-1,6-N-acetylglucosaminyltransferase family protein | elongation zone | 12 h |
| HORVU.MOREX.r3.6HG0609720.1 | -0.80756 | 3.853974 | -4.17823 | 0.00011 | 0.005113 | 1.156809 | Glucosamine 6-phosphate N-acetyltransferase | elongation zone | 12 h |
| HORVU.MOREX.r3.1HG0036370.1 | -0.39076 | 6.141758 | -4.24102 | 8.96E-05 | 0.004634 | 1.150968 | Ras-related protein, expressed | elongation zone | 12 h |
| HORVU.MOREX.r3.5HG0466010.1 | -0.66964 | 3.637084 | -4.17721 | 0.000111 | 0.005113 | 1.150669 | Glutaredoxin | elongation zone | 12 h |
| HORVU.MOREX.r3.3HG0302140.1 | -0.4769 | 5.575661 | -4.21705 | 9.70E-05 | 0.004761 | 1.147805 | S-adenosyl-L-methionine-dependent methyltransferases superfamily protein | elongation zone | 12 h |
| HORVU.MOREX.r3.1HG0082770.1 | -0.3557 | 7.702552 | -4.27588 | 7.98E-05 | 0.004329 | 1.147785 | Reticulon-like protein | elongation zone | 12 h |
| HORVU.MOREX.r3.2HG0161410.1 | -0.32757 | 7.35188 | -4.27923 | 7.89E-05 | 0.004314 | 1.147354 | Acyl-[acyl-carrier-protein] desaturase | elongation zone | 12 h |
| HORVU.MOREX.r3.6HG0625570.1 | 0.659825 | 5.941941 | 4.27068 | 8.11E-05 | 0.004344 | 1.147226 | Transmembrane protein | elongation zone | 12 h |
| HORVU.MOREX.r3.5HG0498150.1 | -1.28562 | 5.56412 | -4.1845 | 0.000108 | 0.005063 | 1.141376 | 2-keto-3-deoxy-L-rhamnonate aldolase | elongation zone | 12 h |
| HORVU.MOREX.r3.2HG0174820.1 | -0.49585 | 6.649571 | -4.22977 | 9.30E-05 | 0.004702 | 1.141142 | F-box family protein | elongation zone | 12 h |
| HORVU.MOREX.r3.1HG0080370.1 | -0.48419 | 5.91275 | -4.22394 | 9.48E-05 | 0.004721 | 1.134648 | F-box family protein | elongation zone | 12 h |
| HORVU.MOREX.r3.1HG0064740.1 | 0.757373 | 3.101051 | 4.184285 | 0.000108 | 0.005063 | 1.125888 | Phospholipase A1 | elongation zone | 12 h |
| HORVU.MOREX.r3.4HG0392160.1 | -1.26162 | 6.711088 | -4.20599 | 0.000101 | 0.004823 | 1.125839 | Glucan endo-1,3-beta-glucosidase, putative | elongation zone | 12 h |
| HORVU.MOREX.r3.3HG0240710.1 | -1.40334 | 3.472687 | -4.17606 | 0.000111 | 0.005113 | 1.120306 | Boron transporter | elongation zone | 12 h |
| HORVU.MOREX.r3.1HG0057900.1 | -0.66246 | 4.053556 | -4.17406 | 0.000112 | 0.005113 | 1.118737 | MYB-related transcription factor | elongation zone | 12 h |
| HORVU.MOREX.r3.7HG0641030.1 | 1.349055 | -0.36725 | 4.188688 | 0.000107 | 0.005013 | 1.11698 | Serpin family protein | elongation zone | 12 h |
| HORVU.MOREX.r3.7HG0727750.1 | 1.323645 | 5.212856 | 4.204674 | 0.000101 | 0.004833 | 1.111823 | Zinc-transporting ATPase | elongation zone | 12 h |

**Table S1** Continued.

| ID | logFC | AveExpr | t | P.Value | adj.P.Val | B | gene description | root zone | time point |
|---|---|---|---|---|---|---|---|---|---|
| HORVU.MOREX.r3.2HG0182400.1 | 0.924203 | 5.649464 | 4.243957 | 8.87E-05 | 0.004634 | 1.10986 | Aquaporin | elongation zone | 12 h |
| HORVU.MOREX.r3.3HG0325250.1 | 3.373214 | -1.52184 | 4.40057 | 5.25E-05 | 0.00369 | 1.094327 | Peroxidase | elongation zone | 12 h |
| HORVU.MOREX.r3.7HG0707700.1 | 1.299693 | -2.42617 | 4.218608 | 9.65E-05 | 0.004752 | 1.088988 | Late embryogenesis abundant (LEA) hydroxyproline-rich glycoprotein family | elongation zone | 12 h |
| HORVU.MOREX.r3.7HG0635320.1 | -0.59826 | 8.584452 | -4.27486 | 8.00E-05 | 0.004331 | 1.083184 | BAG family molecular chaperone regulator 7 | elongation zone | 12 h |
| HORVU.MOREX.r3.5HG0500120.1 | -1.105 | 4.270851 | -4.15476 | 0.000119 | 0.005253 | 1.081867 | Thaumatin | elongation zone | 12 h |
| HORVU.MOREX.r3.3HG0304780.1 | -0.53973 | 4.983038 | -4.18997 | 0.000106 | 0.005013 | 1.081655 | Peptidyl serine alpha-galactosyltransferase | elongation zone | 12 h |
| HORVU.MOREX.r3.7HG0636630.1 | 1.233529 | -0.12855 | 4.161349 | 0.000117 | 0.005234 | 1.076926 | WAT1-related protein | elongation zone | 12 h |
| HORVU.MOREX.r3.5HG0433570.1 | -0.5807 | 4.985176 | -4.15947 | 0.000117 | 0.005234 | 1.076758 | E3 ubiquitin protein ligase DRIP2 | elongation zone | 12 h |
| HORVU.MOREX.r3.5HG0504710.1 | -0.3963 | 6.006762 | -4.21054 | 9.91E-05 | 0.004807 | 1.06976 | Signal peptidase subunit | elongation zone | 12 h |
| HORVU.MOREX.r3.1HG0081950.1 | 2.576402 | -2.05546 | 4.242161 | 8.92E-05 | 0.004634 | 1.064484 | Peroxidase | elongation zone | 12 h |
| HORVU.MOREX.r3.3HG0288710.2 | 1.75276 | -1.63853 | 4.159693 | 0.000117 | 0.005234 | 1.063382 | GDSL esterase/lipase | elongation zone | 12 h |
| HORVU.MOREX.r3.3HG0287000.1 | 1.00542 | 3.768611 | 4.173841 | 0.000112 | 0.005113 | 1.056305 | Respiratory burst oxidase | elongation zone | 12 h |
| HORVU.MOREX.r3.3HG0251830.1 | -1.12154 | 3.41269 | -4.15745 | 0.000118 | 0.005245 | 1.055573 | Plant regulator RWP-RK | elongation zone | 12 h |
| HORVU.MOREX.r3.3HG0294950.1 | 1.231536 | -0.01143 | 4.160938 | 0.000117 | 0.005234 | 1.052113 | Deubiquitinase SseL | elongation zone | 12 h |
| HORVU.MOREX.r3.6HG0626030.1 | 1.722439 | -0.5621 | 4.168357 | 0.000114 | 0.005183 | 1.051979 | Peroxidase | elongation zone | 12 h |
| HORVU.MOREX.r3.2HG0210510.1 | 0.964903 | -0.61919 | 4.15782 | 0.000118 | 0.005245 | 1.051041 | Rapid alkalinization factor 1 | elongation zone | 12 h |
| HORVU.MOREX.r3.7HG0737870.1 | -1.71847 | 1.699649 | -4.25021 | 8.69E-05 | 0.004559 | 1.040195 | E3 ubiquitin-protein ligase RNF12-A | elongation zone | 12 h |
| HORVU.MOREX.r3.3HG0230090.1 | 0.940606 | 1.525122 | 4.228639 | 9.33E-05 | 0.004702 | 1.037877 | Cysteine protease, putative | elongation zone | 12 h |
| HORVU.MOREX.r3.6HG0600500.1 | -0.86376 | 3.118887 | -4.13759 | 0.000126 | 0.0054 | 1.032825 | Ceramide kinase | elongation zone | 12 h |
| HORVU.MOREX.r3.6HG0539460.1 | -0.31637 | 7.146504 | -4.22727 | 9.37E-05 | 0.004703 | 1.026815 | FG-GAP repeat-containing protein | elongation zone | 12 h |
| HORVU.MOREX.r3.3HG0234210.1 | 0.989786 | 3.873833 | 4.159386 | 0.000117 | 0.005234 | 1.022203 | Receptor protein kinase, putative | elongation zone | 12 h |
| HORVU.MOREX.r3.7HG0739120.1 | 1.108937 | 2.007183 | 4.16673 | 0.000115 | 0.005199 | 1.017973 | Receptor-like protein kinase | elongation zone | 12 h |
| HORVU.MOREX.r3.5HG0513780.1 | 0.648523 | 5.051864 | 4.189825 | 0.000106 | 0.005013 | 1.01478 | Kinesin-like protein | elongation zone | 12 h |
| HORVU.MOREX.r3.2HG0204920.1 | -1.38459 | 3.638603 | -4.16375 | 0.000116 | 0.005234 | 1.012207 | Crossover junction endonuclease EME1B | elongation zone | 12 h |
| HORVU.MOREX.r3.2HG0102670.1 | 2.391367 | -3.01135 | 4.431283 | 4.73E-05 | 0.003494 | 1.009149 | Dirigent protein | elongation zone | 12 h |
| HORVU.MOREX.r3.1HG0038800.1 | 0.700493 | 2.879103 | 4.140907 | 0.000125 | 0.005398 | 1.003947 | Protein kinase | elongation zone | 12 h |
| HORVU.MOREX.r3.2HG0120650.1 | 1.154291 | 1.67255 | 4.241326 | 8.95E-05 | 0.004634 | 1.002559 | Peroxidase | elongation zone | 12 h |
| HORVU.MOREX.r3.4HG0357230.1 | -1.22682 | 5.256363 | -4.13104 | 0.000129 | 0.005458 | 1.001035 | Pyridoxal 5'-phosphate synthase subunit PdxS | elongation zone | 12 h |
| HORVU.MOREX.r3.5HG0529500.1 | -0.34939 | 7.05237 | -4.21102 | 9.89E-05 | 0.004807 | 0.994542 | Protein kinase | elongation zone | 12 h |
| HORVU.MOREX.r3.5HG0446210.1 | -1.06958 | 3.497682 | -4.13087 | 0.000129 | 0.005458 | 0.994488 | Agenet domain, putative | elongation zone | 12 h |
| HORVU.MOREX.r3.4HG0403620.1 | 1.536981 | 0.126435 | 4.192795 | 0.000105 | 0.004991 | 0.990315 | #N/A | elongation zone | 12 h |
| HORVU.MOREX.r3.6HG0543800.1 | -0.92812 | 7.420452 | -4.20718 | 0.0001 | 0.004816 | 0.990216 | Histone H2A | elongation zone | 12 h |
| HORVU.MOREX.r3.7HG0722360.1 | 1.636206 | 0.664931 | 4.12172 | 0.000133 | 0.005517 | 0.986935 | Receptor-like protein kinase | elongation zone | 12 h |
| HORVU.MOREX.r3.7HG0700270.1 | -0.42456 | 7.724735 | -4.23203 | 9.23E-05 | 0.004702 | 0.985323 | Transaldolase | elongation zone | 12 h |
| HORVU.MOREX.r3.2HG0165840.1 | -0.72761 | 6.204491 | -4.15667 | 0.000118 | 0.005247 | 0.984828 | Protein kinase | elongation zone | 12 h |
| HORVU.MOREX.r3.7HG0676570.1 | -0.75448 | 6.054358 | -4.16064 | 0.000117 | 0.005234 | 0.966551 | Deaminase-related family protein | elongation zone | 12 h |
| HORVU.MOREX.r3.7HG0739400.1 | -1.21056 | 4.93511 | -4.124 | 0.000132 | 0.005517 | 0.963995 | Histone H2B | elongation zone | 12 h |
| HORVU.MOREX.r3.2HG0198870.1 | 1.177674 | -0.51668 | 4.127474 | 0.00013 | 0.005469 | 0.963587 | UPF0761 membrane protein Plut_1323 | elongation zone | 12 h |
| HORVU.MOREX.r3.6HG0568880.1 | -1.26447 | 5.953717 | -4.13798 | 0.000126 | 0.0054 | 0.961847 | #N/A | elongation zone | 12 h |
| HORVU.MOREX.r3.2HG0110580.1 | -0.55957 | 3.365388 | -4.11625 | 0.000135 | 0.005554 | 0.961828 | Ras-responsive element-binding protein 1 | elongation zone | 12 h |
| HORVU.MOREX.r3.3HG0306420.1 | -1.04302 | 3.782406 | -4.11041 | 0.000138 | 0.005596 | 0.951762 | TPR repeat-containing protein ZIP4 | elongation zone | 12 h |

**Table S1** Continued.

| ID | logFC | AveExpr | t | P.Value | adj.P.Val | B | gene description | root zone | time point |
|---|---|---|---|---|---|---|---|---|---|
| HORVU.MOREX.r3.3HG0293760.1 | 1.453863 | 3.466627 | 4.143802 | 0.000124 | 0.005393 | 0.950534 | External alternative NAD(P)H-ubiquinone oxidoreductase B2, mitochondrial | elongation zone | 12 h |
| HORVU.MOREX.r3.2HG0207130.1 | 1.534229 | 1.218085 | 4.147631 | 0.000122 | 0.005349 | 0.95016 | Acetyl-coenzyme A synthetase | elongation zone | 12 h |
| HORVU.MOREX.r3.7HG0663920.1 | 2.354114 | -0.12227 | 4.181561 | 0.000109 | 0.005098 | 0.947156 | Alpha/beta-Hydrolases superfamily protein, putative | elongation zone | 12 h |
| HORVU.MOREX.r3.2HG0207860.1 | 1.587118 | -2.16402 | 4.154338 | 0.000119 | 0.005253 | 0.943939 | Serine/threonine-protein kinase | elongation zone | 12 h |
| HORVU.MOREX.r3.4HG0337110.1 | -0.34603 | 7.514078 | -4.21378 | 9.80E-05 | 0.004801 | 0.943096 | Coatomer subunit alpha | elongation zone | 12 h |
| HORVU.MOREX.r3.7HG0729150.1 | 1.260149 | -2.07184 | 4.107247 | 0.000139 | 0.005596 | 0.943004 | GDSL esterase/lipase | elongation zone | 12 h |
| HORVU.MOREX.r3.2HG0199560.1 | -1.82807 | 5.029114 | -4.10998 | 0.000138 | 0.005596 | 0.942071 | Organic cation transporter-like protein | elongation zone | 12 h |
| HORVU.MOREX.r3.3HG0275770.1 | 0.728263 | 5.894882 | 4.201195 | 0.000102 | 0.004877 | 0.938511 | Alpha/beta hydrolase-like | elongation zone | 12 h |
| HORVU.MOREX.r3.5HG0469120.1 | 1.737777 | -1.16488 | 4.102936 | 0.000141 | 0.005625 | 0.936066 | E3 ubiquitin-protein ligase SINA-like 10 | elongation zone | 12 h |
| HORVU.MOREX.r3.1HG0075730.1 | 1.047776 | 1.539739 | 4.196987 | 0.000104 | 0.004934 | 0.924583 | Subtilisin-like protease | elongation zone | 12 h |
| HORVU.MOREX.r3.7HG0676350.1 | -0.60217 | 4.38572 | -4.12953 | 0.000129 | 0.005466 | 0.920845 | Maternal effect embryo arrest protein | elongation zone | 12 h |
| HORVU.MOREX.r3.3HG0305500.1 | -1.31455 | 2.378582 | -4.17435 | 0.000112 | 0.005113 | 0.914269 | 50S ribosomal protein-related, putative | elongation zone | 12 h |
| HORVU.MOREX.r3.7HG0648460.1 | 0.620827 | 2.500172 | 4.111429 | 0.000137 | 0.005593 | 0.912195 | Glycosyltransferase | elongation zone | 12 h |
| HORVU.MOREX.r3.7HG0720100.1 | 0.768927 | 4.342288 | 4.105142 | 0.00014 | 0.005599 | 0.912009 | Glucan endo-1,3-beta-glucosidase, putative | elongation zone | 12 h |
| HORVU.MOREX.r3.1HG0051740.1 | -2.71044 | 3.433417 | -4.14521 | 0.000123 | 0.005379 | 0.908551 | Glutathione s-transferase, putative | elongation zone | 12 h |
| HORVU.MOREX.r3.2HG0190230.1 | -1.47377 | 4.385744 | -4.09308 | 0.000146 | 0.005731 | 0.906254 | Histone H4 | elongation zone | 12 h |
| HORVU.MOREX.r3.7HG0660760.1 | -1.15356 | 5.301372 | -4.10691 | 0.000139 | 0.005596 | 0.904214 | Histone H3 | elongation zone | 12 h |
| HORVU.MOREX.r3.1HG0054180.1 | 1.538617 | -0.6979 | 4.100335 | 0.000142 | 0.005654 | 0.903462 | Expansin-like protein | elongation zone | 12 h |
| HORVU.MOREX.r3.4HG0380330.1 | -0.52912 | 5.401503 | -4.138 | 0.000126 | 0.0054 | 0.90305 | WRKY transcription factor | elongation zone | 12 h |
| HORVU.MOREX.r3.4HG0395220.1 | -0.65223 | 6.119297 | -4.1417 | 0.000124 | 0.005398 | 0.902652 | Trehalose-6-phosphate synthase | elongation zone | 12 h |
| HORVU.MOREX.r3.5HG0517190.1 | 0.702954 | 2.990144 | 4.089167 | 0.000148 | 0.005751 | 0.89564 | Calcium-dependent protein kinase | elongation zone | 12 h |
| HORVU.MOREX.r3.4HG0337950.1 | 2.271565 | -0.29631 | 4.119137 | 0.000134 | 0.00553 | 0.89075 | 65-kDa microtubule-associated-like protein | elongation zone | 12 h |
| HORVU.MOREX.r3.5HG0458890.1 | 0.554844 | 3.514885 | 4.105111 | 0.00014 | 0.005599 | 0.880751 | senescence regulator (Protein of unknown function, DUF584) | elongation zone | 12 h |
| HORVU.MOREX.r3.6HG0552800.1 | -1.13398 | 7.010449 | -4.13639 | 0.000127 | 0.005411 | 0.874778 | Histone H2B | elongation zone | 12 h |
| HORVU.MOREX.r3.1HG0089130.1 | -0.4243 | 5.608758 | -4.11581 | 0.000135 | 0.005554 | 0.873458 | Serine/Threonine-kinase pakA-like protein | elongation zone | 12 h |
| HORVU.MOREX.r3.7HG0748270.1 | 1.725878 | 0.645561 | 4.081136 | 0.000152 | 0.005814 | 0.872048 | Blue copper protein | elongation zone | 12 h |
| HORVU.MOREX.r3.3HG0289020.1 | 1.305574 | 2.050529 | 4.11506 | 0.000136 | 0.005554 | 0.866394 | Receptor-like kinase | elongation zone | 12 h |
| HORVU.MOREX.r3.3HG0304370.1 | 0.608195 | 4.367817 | 4.12174 | 0.000133 | 0.005517 | 0.863869 | Glutamyl-tRNA(Gln) amidotransferase subunit A | elongation zone | 12 h |
| HORVU.MOREX.r3.3HG0319900.1 | 1.531919 | -0.77823 | 4.07715 | 0.000154 | 0.005819 | 0.859784 | Ankyrin repeat domain-containing protein 1 | elongation zone | 12 h |
| HORVU.MOREX.r3.5HG0481440.1 | -0.47662 | 4.469677 | -4.10645 | 0.00014 | 0.005596 | 0.859431 | glucan synthase-like 10 | elongation zone | 12 h |
| HORVU.MOREX.r3.3HG0264650.1 | 1.692176 | -2.1209 | 4.127697 | 0.00013 | 0.005469 | 0.857287 | Retrovirus-related Pol polyprotein from transposon TNT 1-94 | elongation zone | 12 h |
| HORVU.MOREX.r3.5HG0510940.1 | -1.46151 | 5.282693 | -4.0755 | 0.000155 | 0.005829 | 0.850169 | HMG-Y-related protein A | elongation zone | 12 h |
| HORVU.MOREX.r3.3HG0244690.1 | -0.83906 | 4.95279 | -4.08165 | 0.000151 | 0.005814 | 0.839301 | Tubulin alpha chain | elongation zone | 12 h |
| HORVU.MOREX.r3.3HG0233990.1 | 1.24044 | 1.603331 | 4.189007 | 0.000106 | 0.005013 | 0.838882 | Xyloglucan endotransglucosylase/hydrolase | elongation zone | 12 h |
| HORVU.MOREX.r3.2HG0097950.1 | -0.87049 | 10.61692 | -4.24093 | 8.96E-05 | 0.004634 | 0.836316 | Apyrase | elongation zone | 12 h |
| HORVU.MOREX.r3.3HG0297180.1 | -1.7609 | 2.307652 | -4.21181 | 9.87E-05 | 0.004807 | 0.833175 | Pentatricopeptide repeat-containing protein | elongation zone | 12 h |
| HORVU.MOREX.r3.5HG0490610.1 | 1.717845 | -0.13337 | 4.073125 | 0.000156 | 0.005863 | 0.833139 | Protein kinase | elongation zone | 12 h |
| HORVU.MOREX.r3.7HG0699020.1 | 1.85112 | -0.26924 | 4.351227 | 6.20E-05 | 0.003971 | 0.831186 | Retrotransposon protein, putative, Ty3-gypsy subclass | elongation zone | 12 h |
| HORVU.MOREX.r3.3HG0222130.1 | -0.3574 | 8.795276 | -4.22182 | 9.55E-05 | 0.004743 | 0.828263 | Transmembrane protein 214 | elongation zone | 12 h |
| HORVU.MOREX.r3.7HG0709270.1 | 1.010671 | 3.012432 | 4.092512 | 0.000146 | 0.005731 | 0.828025 | RING/U-box superfamily protein | elongation zone | 12 h |
| HORVU.MOREX.r3.2HG0120320.1 | 0.824766 | 1.685244 | 4.169557 | 0.000113 | 0.005174 | 0.823334 | Peroxidase | elongation zone | 12 h |

**Table S1** Continued.

| ID | logFC | AveExpr | t | P.Value | adj.P.Val | B | gene description | root zone | time point |
|---|---|---|---|---|---|---|---|---|---|
| HORVU.MOREX.r3.4HG0385780.1 | -0.56087 | 5.783595 | -4.12205 | 0.000133 | 0.005517 | 0.822654 | Mitogen-activated protein kinase | elongation zone | 12 h |
| HORVU.MOREX.r3.1HG0023610.1 | -1.32683 | 5.369444 | -4.0785 | 0.000153 | 0.005814 | 0.821833 | #N/A | elongation zone | 12 h |
| HORVU.MOREX.r3.6HG0551740.1 | -0.70258 | 5.295694 | -4.08498 | 0.00015 | 0.0058 | 0.821437 | Early light-inducible protein | elongation zone | 12 h |
| HORVU.MOREX.r3.1HG0050920.1 | 1.043125 | 2.478719 | 4.123174 | 0.000132 | 0.005517 | 0.803006 | Trehalase | elongation zone | 12 h |
| HORVU.MOREX.r3.5HG0425680.1 | -0.45559 | 7.233115 | -4.15499 | 0.000119 | 0.005253 | 0.801983 | No exine formation 1 | elongation zone | 12 h |
| HORVU.MOREX.r3.1HG0005700.1 | 1.949907 | -2.12158 | 4.078759 | 0.000153 | 0.005814 | 0.798959 | #N/A | elongation zone | 12 h |
| HORVU.MOREX.r3.6HG0558800.1 | -0.64398 | 5.008691 | -4.0695 | 0.000158 | 0.00589 | 0.79346 | DNA-binding protein BIN4 | elongation zone | 12 h |
| HORVU.MOREX.r3.4HG0404750.1 | -2.00695 | 2.885556 | -4.05386 | 0.000166 | 0.006055 | 0.787843 | Hydrolase, alpha/beta fold family protein, expressed | elongation zone | 12 h |
| HORVU.MOREX.r3.3HG0271380.1 | 0.607629 | 5.415877 | 4.092597 | 0.000146 | 0.005731 | 0.785605 | GDSL esterase/lipase | elongation zone | 12 h |
| HORVU.MOREX.r3.3HG0295680.1 | 0.616236 | 4.416665 | 4.113964 | 0.000136 | 0.005558 | 0.785088 | Dihydropyrimidinase | elongation zone | 12 h |
| HORVU.MOREX.r3.7HG0720790.1 | -1.17463 | 3.44449 | -4.06893 | 0.000158 | 0.00589 | 0.781788 | Myb family transcription factor family protein | elongation zone | 12 h |
| HORVU.MOREX.r3.6HG0626020.1 | 1.428647 | 1.680187 | 4.141812 | 0.000124 | 0.005398 | 0.778046 | Peroxidase | elongation zone | 12 h |
| HORVU.MOREX.r3.1HG0008200.1 | -2.45219 | 1.354368 | -4.05613 | 0.000165 | 0.006032 | 0.775839 | Cytochrome P450 | elongation zone | 12 h |
| HORVU.MOREX.r3.2HG0184180.1 | 1.137619 | 4.475955 | 4.139182 | 0.000125 | 0.0054 | 0.77234 | Remorin | elongation zone | 12 h |
| HORVU.MOREX.r3.7HG0670900.1 | 1.570995 | 2.29029 | 4.054332 | 0.000166 | 0.006055 | 0.771381 | Aspartic proteinase Asp1 | elongation zone | 12 h |
| HORVU.MOREX.r3.7HG0642520.1 | 1.830252 | -0.8717 | 4.065528 | 0.00016 | 0.005945 | 0.770686 | Peroxidase | elongation zone | 12 h |
| HORVU.MOREX.r3.1HG0060320.1 | -1.05585 | 7.084889 | -4.10794 | 0.000139 | 0.005596 | 0.769387 | Histone H3 | elongation zone | 12 h |
| HORVU.MOREX.r3.7HG0747230.1 | 0.746133 | 2.606945 | 4.044909 | 0.000171 | 0.006089 | 0.766735 | Non-specific serine/threonine protein kinase | elongation zone | 12 h |
| HORVU.MOREX.r3.3HG0255020.1 | 0.984176 | 3.48561 | 4.17494 | 0.000111 | 0.005113 | 0.759937 | Peroxidase | elongation zone | 12 h |
| HORVU.MOREX.r3.5HG0513740.1 | 1.598995 | -3.01674 | 4.231653 | 9.24E-05 | 0.004702 | 0.758283 | Alpha/beta-Hydrolases superfamily protein, putative | elongation zone | 12 h |
| HORVU.MOREX.r3.1HG0084540.1 | -0.51556 | 4.682873 | -4.06461 | 0.00016 | 0.005948 | 0.757601 | F-box family protein | elongation zone | 12 h |
| HORVU.MOREX.r3.3HG0308420.1 | -0.8279 | 3.844691 | -4.04158 | 0.000173 | 0.00612 | 0.757262 | Integrator complex subunit | elongation zone | 12 h |
| HORVU.MOREX.r3.3HG0246560.1 | -0.76999 | 5.025368 | -4.04832 | 0.000169 | 0.006075 | 0.756947 | Multiprotein-bridging factor, putative | elongation zone | 12 h |
| HORVU.MOREX.r3.7HG0666650.1 | -1.60043 | -0.31405 | -4.0693 | 0.000158 | 0.00589 | 0.752726 | Starch synthase, chloroplastic/amyloplastic | elongation zone | 12 h |
| HORVU.MOREX.r3.6HG0631710.1 | 1.057823 | 2.878977 | 4.175195 | 0.000111 | 0.005113 | 0.750974 | Aquaporin | elongation zone | 12 h |
| HORVU.MOREX.r3.1HG0001010.1 | -0.88425 | 4.233711 | -4.04897 | 0.000168 | 0.006075 | 0.750273 | Pm3-like disease resistance protein | elongation zone | 12 h |
| HORVU.MOREX.r3.6HG0586850.1 | -1.26756 | 0.435847 | -4.10252 | 0.000141 | 0.005625 | 0.746832 | Replication-associated protein | elongation zone | 12 h |
| HORVU.MOREX.r3.2HG0139910.1 | -1.00909 | 2.439496 | -4.04606 | 0.00017 | 0.006081 | 0.744843 | RNA-binding CRS1 / YhbY (CRM) domain protein | elongation zone | 12 h |
| HORVU.MOREX.r3.4HG0377070.1 | -0.53464 | 5.867895 | -4.08874 | 0.000148 | 0.005751 | 0.738742 | Kinase family protein | elongation zone | 12 h |
| HORVU.MOREX.r3.7HG0661060.1 | -0.44743 | 6.051504 | -4.08894 | 0.000148 | 0.005751 | 0.736993 | Ethylene-responsive transcription factor | elongation zone | 12 h |
| HORVU.MOREX.r3.6HG0614840.1 | 0.569248 | 5.33259 | 4.114761 | 0.000136 | 0.005554 | 0.732255 | Shikimate kinase | elongation zone | 12 h |
| HORVU.MOREX.r3.4HG0384620.1 | 0.92359 | 3.226323 | 4.047899 | 0.000169 | 0.006075 | 0.725588 | Kinase family protein | elongation zone | 12 h |
| HORVU.MOREX.r3.3HG0282170.1 | -1.82596 | 6.992997 | -4.09211 | 0.000146 | 0.005731 | 0.712133 | Methyltransferase | elongation zone | 12 h |
| HORVU.MOREX.r3.5HG0463760.1 | -0.74771 | 3.113818 | -4.02506 | 0.000182 | 0.00626 | 0.709355 | Ethanolamine-phosphate cytidylyltransferase | elongation zone | 12 h |
| HORVU.MOREX.r3.3HG0309900.1 | 1.463688 | -1.42396 | 4.034118 | 0.000177 | 0.006165 | 0.706816 | ROP-interactive CRIB motif protein | elongation zone | 12 h |
| HORVU.MOREX.r3.7HG0734670.1 | -0.5608 | 6.626504 | -4.11818 | 0.000134 | 0.005536 | 0.702896 | L-ascorbate oxidase | elongation zone | 12 h |
| HORVU.MOREX.r3.4HG0384230.1 | -0.63771 | 3.949266 | -4.03514 | 0.000176 | 0.006165 | 0.702046 | RING/U-box superfamily protein | elongation zone | 12 h |
| HORVU.MOREX.r3.7HG0661830.1 | 2.193351 | -0.86487 | 4.032858 | 0.000178 | 0.006165 | 0.70134 | GPI-anchored adhesin-like protein | elongation zone | 12 h |
| HORVU.MOREX.r3.2HG0194840.1 | -1.85975 | 3.704881 | -4.02801 | 0.00018 | 0.006231 | 0.701099 | Glucans biosynthesis glucosyltransferase H | elongation zone | 12 h |
| HORVU.MOREX.r3.2HG0157410.1 | 1.212013 | 2.254585 | 4.141316 | 0.000125 | 0.005398 | 0.696567 | Peroxidase | elongation zone | 12 h |
| HORVU.MOREX.r3.7HG0653380.1 | 1.439011 | 1.304596 | 4.138632 | 0.000126 | 0.0054 | 0.693981 | GDSL esterase/lipase | elongation zone | 12 h |

**Table S1** Continued.

| ID | logFC | AveExpr | t | P.Value | adj.P.Val | B | gene description | root zone | time point |
|---|---|---|---|---|---|---|---|---|---|
| HORVU.MOREX.r3.5HG0485860.1 | -0.6617 | 3.795871 | -4.04914 | 0.000168 | 0.006075 | 0.685731 | LysM domain-containing GPI-anchored protein 1 | elongation zone | 12 h |
| HORVU.MOREX.r3.2HG0164160.2 | 1.553915 | -0.40053 | 4.049615 | 0.000168 | 0.006075 | 0.682135 | Trihelix transcription factor | elongation zone | 12 h |
| HORVU.MOREX.r3.3HG0238250.1 | -0.89848 | 6.327265 | -4.09553 | 0.000145 | 0.005722 | 0.680184 | Gigantea-like protein | elongation zone | 12 h |
| HORVU.MOREX.r3.2HG0164650.1 | -0.47352 | 5.166801 | -4.0469 | 0.00017 | 0.006075 | 0.680128 | Sucrose nonfermenting 4-like protein | elongation zone | 12 h |
| HORVU.MOREX.r3.4HG0404740.1 | 0.80758 | 2.889075 | 4.079976 | 0.000152 | 0.005814 | 0.67668 | Hydrolase, alpha/beta fold family protein, expressed | elongation zone | 12 h |
| HORVU.MOREX.r3.5HG0486750.1 | 0.873877 | 2.037011 | 4.040656 | 0.000173 | 0.006125 | 0.671235 | Phosphatase 2C family protein | elongation zone | 12 h |
| HORVU.MOREX.r3.2HG0146580.1 | -0.5505 | 4.815547 | -4.03643 | 0.000176 | 0.006165 | 0.668194 | PHD finger alfin-like protein | elongation zone | 12 h |
| HORVU.MOREX.r3.4HG0382610.1 | 1.243522 | 0.223451 | 4.062262 | 0.000161 | 0.005955 | 0.666417 | cytochrome P450, family 705, subfamily A, polypeptide 21 | elongation zone | 12 h |
| HORVU.MOREX.r3.4HG0348800.1 | 1.130201 | 2.044228 | 4.010239 | 0.000191 | 0.006387 | 0.665813 | TCP transcription factor | elongation zone | 12 h |
| HORVU.MOREX.r3.7HG0650970.1 | -1.17613 | 7.468679 | -4.07605 | 0.000154 | 0.005829 | 0.661654 | Histone H3 | elongation zone | 12 h |
| HORVU.MOREX.r3.3HG0319960.1 | -0.94542 | 2.661966 | -4.03427 | 0.000177 | 0.006165 | 0.658603 | F-box-like protein | elongation zone | 12 h |
| HORVU.MOREX.r3.3HG0255700.1 | 0.949034 | 1.416296 | 4.007633 | 0.000193 | 0.006402 | 0.653542 | PQ-loop repeat family protein / transmembrane family protein | elongation zone | 12 h |
| HORVU.MOREX.r3.1HG0017280.1 | 1.59651 | -0.33011 | 4.022299 | 0.000184 | 0.006278 | 0.650143 | Peroxidase | elongation zone | 12 h |
| HORVU.MOREX.r3.2HG0210610.1 | 0.925465 | 3.089943 | 4.022117 | 0.000184 | 0.006278 | 0.650022 | Acetyl-coenzyme A synthetase | elongation zone | 12 h |
| HORVU.MOREX.r3.7HG0749170.1 | -1.3517 | 5.305591 | -4.02946 | 0.00018 | 0.006217 | 0.649194 | DUF241 domain protein (DUF241) | elongation zone | 12 h |
| HORVU.MOREX.r3.4HG0406600.1 | 0.888365 | 0.071867 | 4.046912 | 0.00017 | 0.006075 | 0.646893 | Heavy metal transport/detoxification superfamily protein | elongation zone | 12 h |
| HORVU.MOREX.r3.6HG0597620.1 | -0.47097 | 5.241895 | -4.0637 | 0.000161 | 0.005948 | 0.646402 | #N/A | elongation zone | 12 h |
| HORVU.MOREX.r3.3HG0218100.1 | -0.36221 | 6.103028 | -4.0702 | 0.000157 | 0.00589 | 0.646235 | Chaperone protein DnaJ | elongation zone | 12 h |
| HORVU.MOREX.r3.5HG0463550.1 | 1.043959 | 4.670129 | 4.01103 | 0.000191 | 0.006387 | 0.64457 | RNA-binding domain CCCH-type zinc finger protein | elongation zone | 12 h |
| HORVU.MOREX.r3.6HG0598750.1 | 1.171516 | 0.858146 | 4.097067 | 0.000144 | 0.005704 | 0.644096 | Remorin | elongation zone | 12 h |
| HORVU.MOREX.r3.6HG0627920.1 | -0.3309 | 7.364979 | -4.1197 | 0.000134 | 0.00553 | 0.642997 | Ankyrin repeat protein-like | elongation zone | 12 h |
| HORVU.MOREX.r3.3HG0272160.2 | -0.706 | 3.90619 | -4.007 | 0.000193 | 0.006402 | 0.640427 | Aldo/keto reductase family oxidoreductase | elongation zone | 12 h |
| HORVU.MOREX.r3.5HG0461830.1 | -1.41496 | 1.937434 | -4.1337 | 0.000128 | 0.005436 | 0.638145 | F-box/kelch-repeat protein | elongation zone | 12 h |
| HORVU.MOREX.r3.7HG0649950.1 | 0.647727 | 3.836695 | 4.033277 | 0.000177 | 0.006165 | 0.636184 | Nudix hydrolase | elongation zone | 12 h |
| HORVU.MOREX.r3.4HG0342160.1 | -0.48024 | 5.308472 | -4.04338 | 0.000172 | 0.006102 | 0.635113 | Presenilin | elongation zone | 12 h |
| HORVU.MOREX.r3.1HG0069570.1 | -0.37717 | 8.695958 | -4.15957 | 0.000117 | 0.005234 | 0.631104 | Tubulin beta chain | elongation zone | 12 h |
| HORVU.MOREX.r3.1HG0052810.1 | -0.66093 | 7.005109 | -4.07861 | 0.000153 | 0.005814 | 0.62667 | Mitochondrial carrier protein | elongation zone | 12 h |
| HORVU.MOREX.r3.6HG0586290.1 | -0.87785 | 2.563314 | -4.00053 | 0.000197 | 0.006444 | 0.62289 | CDT1-like protein a, chloroplastic | elongation zone | 12 h |
| HORVU.MOREX.r3.3HG0218800.1 | 1.971084 | -0.5154 | 3.999058 | 0.000198 | 0.006444 | 0.621042 | hydroxyproline-rich glycoprotein family protein | elongation zone | 12 h |
| HORVU.MOREX.r3.5HG0497360.1 | -0.56078 | 3.787521 | -3.9983 | 0.000199 | 0.006448 | 0.619229 | Bile acid sodium symporter | elongation zone | 12 h |
| HORVU.MOREX.r3.2HG0099010.1 | 1.17563 | -2.64978 | 4.082181 | 0.000151 | 0.005814 | 0.616661 | Glutamyl-tRNA (Gln) amidotransferase subunit A | elongation zone | 12 h |
| HORVU.MOREX.r3.7HG0747940.1 | 0.459113 | 7.327177 | 4.128528 | 0.00013 | 0.005469 | 0.613541 | Polyamine oxidase | elongation zone | 12 h |
| HORVU.MOREX.r3.1HG0026430.1 | -0.41505 | 4.788835 | -4.03732 | 0.000175 | 0.00616 | 0.604281 | Proline-rich receptor-like kinase, putative (DUF1421) | elongation zone | 12 h |
| HORVU.MOREX.r3.4HG0335690.1 | -0.64334 | 6.238374 | -4.05177 | 0.000167 | 0.006064 | 0.602143 | Zinc finger CCCH domain-containing protein | elongation zone | 12 h |
| HORVU.MOREX.r3.1HG0008510.2 | 0.543767 | 3.338317 | 3.999991 | 0.000198 | 0.006444 | 0.597615 | Beta-hexosaminidase | elongation zone | 12 h |
| HORVU.MOREX.r3.3HG0288700.1 | -0.67011 | 3.838111 | -3.99334 | 0.000202 | 0.006513 | 0.595243 | #N/A | elongation zone | 12 h |
| HORVU.MOREX.r3.3HG0242950.1 | -0.4367 | 6.032036 | -4.0512 | 0.000167 | 0.006064 | 0.591728 | NF-X1 type zinc finger family protein | elongation zone | 12 h |
| HORVU.MOREX.r3.3HG0307850.1 | -0.75535 | 6.891759 | -4.10701 | 0.000139 | 0.005596 | 0.590726 | Basic 7S globulin | elongation zone | 12 h |
| HORVU.MOREX.r3.2HG0121200.1 | 0.902639 | 0.643147 | 4.029244 | 0.00018 | 0.006217 | 0.585822 | Dirigent protein | elongation zone | 12 h |
| HORVU.MOREX.r3.5HG0477750.1 | 1.981434 | 0.658352 | 4.013485 | 0.000189 | 0.006367 | 0.576746 | Homeobox protein, putative | elongation zone | 12 h |
| HORVU.MOREX.r3.4HG0401730.1 | -0.7434 | 2.933703 | -3.97939 | 0.000211 | 0.006618 | 0.575755 | Pentatricopeptide repeat-containing protein | elongation zone | 12 h |

**Table S1** Continued.

| ID | logFC | AveExpr | t | P.Value | adj.P.Val | B | gene description | root zone | time point |
|---|---|---|---|---|---|---|---|---|---|
| HORVU.MOREX.r3.4HG0413320.1 | 1.859757 | 2.316272 | 3.991716 | 0.000203 | 0.006534 | 0.574251 | Subtilisin-like protease | elongation zone | 12 h |
| HORVU.MOREX.r3.1HG0018190.2 | -1.18149 | 3.737279 | -3.97592 | 0.000214 | 0.00667 | 0.561255 | Aspartic proteinase | elongation zone | 12 h |
| HORVU.MOREX.r3.7HG0684880.1 | 0.79924 | 3.761267 | 3.997829 | 0.000199 | 0.006448 | 0.560572 | meiosis chromosome segregation family protein | elongation zone | 12 h |
| HORVU.MOREX.r3.1HG0091330.1 | -0.97883 | 4.42056 | -3.97754 | 0.000212 | 0.006645 | 0.560416 | Flavin-containing monooxygenase | elongation zone | 12 h |
| HORVU.MOREX.r3.7HG0669580.1 | 0.7114 | 1.551164 | 4.041416 | 0.000173 | 0.00612 | 0.559434 | Caleosin | elongation zone | 12 h |
| HORVU.MOREX.r3.4HG0412090.1 | 1.817709 | -0.36354 | 3.988402 | 0.000205 | 0.006554 | 0.557106 | C3HC4-type RING finger-containing protein | elongation zone | 12 h |
| HORVU.MOREX.r3.5HG0454720.1 | 0.569329 | 5.752069 | 4.01507 | 0.000188 | 0.006364 | 0.555617 | Filament-like plant protein | elongation zone | 12 h |
| HORVU.MOREX.r3.2HG0142740.1 | 0.365846 | 6.623245 | 4.091156 | 0.000147 | 0.005738 | 0.554294 | Protein phosphatase 2c | elongation zone | 12 h |
| HORVU.MOREX.r3.5HG0514950.1 | -0.5784 | 5.805968 | -4.0236 | 0.000183 | 0.006269 | 0.54917 | Actin-depolymerizing factor | elongation zone | 12 h |
| HORVU.MOREX.r3.7HG0730840.1 | -1.08969 | 4.5286 | -3.98201 | 0.000209 | 0.0066 | 0.548401 | Histone H3 | elongation zone | 12 h |
| HORVU.MOREX.r3.1HG0076120.1 | 0.68528 | 7.792381 | 4.177781 | 0.00011 | 0.005113 | 0.546564 | Phosphatidylinositol transfer protein | elongation zone | 12 h |
| HORVU.MOREX.r3.7HG0650250.1 | -0.62992 | 3.840838 | -3.9741 | 0.000215 | 0.006679 | 0.54407 | Protein-O-fucosyltransferase 1 | elongation zone | 12 h |
| HORVU.MOREX.r3.3HG0245250.1 | 3.239054 | -1.64331 | 3.988561 | 0.000205 | 0.006554 | 0.537673 | Cysteine protease, putative | elongation zone | 12 h |
| HORVU.MOREX.r3.2HG0110130.1 | -1.02012 | 3.822081 | -4.00269 | 0.000196 | 0.00643 | 0.534689 | Transcription factor protein | elongation zone | 12 h |
| HORVU.MOREX.r3.2HG0180010.1 | 1.375117 | 1.125048 | 4.01451 | 0.000188 | 0.006364 | 0.533217 | #N/A | elongation zone | 12 h |
| HORVU.MOREX.r3.3HG0313900.1 | 1.873062 | 0.291481 | 4.005222 | 0.000194 | 0.006419 | 0.526921 | Methyl esterase 1 | elongation zone | 12 h |
| HORVU.MOREX.r3.3HG0296010.1 | 1.968773 | -0.38601 | 3.961341 | 0.000224 | 0.006805 | 0.525691 | WAT1-related protein | elongation zone | 12 h |
| HORVU.MOREX.r3.6HG0543770.1 | -0.9087 | 7.792354 | -4.06286 | 0.000161 | 0.005954 | 0.52526 | Histone H2A | elongation zone | 12 h |
| HORVU.MOREX.r3.6HG0567150.1 | 1.159328 | 2.844231 | 3.979896 | 0.000211 | 0.006618 | 0.523284 | Exosome complex component | elongation zone | 12 h |
| HORVU.MOREX.r3.5HG0464440.1 | 1.039304 | 0.420648 | 4.004921 | 0.000194 | 0.006419 | 0.517211 | Kinase, putative | elongation zone | 12 h |
| HORVU.MOREX.r3.6HG0590430.1 | -0.41192 | 5.249518 | -4.00282 | 0.000196 | 0.00643 | 0.513924 | Multidrug resistance protein MdtC | elongation zone | 12 h |
| HORVU.MOREX.r3.3HG0308350.1 | 1.03734 | 4.926407 | 3.997424 | 0.000199 | 0.006448 | 0.511026 | Cysteine-rich receptor kinase | elongation zone | 12 h |
| HORVU.MOREX.r3.5HG0457310.1 | 1.126551 | 0.314313 | 4.001557 | 0.000197 | 0.006443 | 0.510447 | Exocyst complex component, putative | elongation zone | 12 h |
| HORVU.MOREX.r3.2HG0096600.1 | -0.59642 | 8.083915 | -4.08285 | 0.000151 | 0.005814 | 0.508985 | Pyruvate dehydrogenase E1 component subunit alpha | elongation zone | 12 h |
| HORVU.MOREX.r3.1HG0075650.1 | 1.18635 | 4.442331 | 4.080471 | 0.000152 | 0.005814 | 0.506425 | transmembrane protein, putative (DUF1218) | elongation zone | 12 h |
| HORVU.MOREX.r3.1HG0073010.1 | 0.958928 | 1.113004 | 4.033381 | 0.000177 | 0.006165 | 0.500115 | Kinase family protein | elongation zone | 12 h |
| HORVU.MOREX.r3.7HG0722260.1 | 1.63999 | -2.17609 | 3.973294 | 0.000215 | 0.006686 | 0.499731 | RPM1-interacting protein 4 | elongation zone | 12 h |
| HORVU.MOREX.r3.3HG0314270.1 | -0.78876 | 2.38297 | -3.96934 | 0.000218 | 0.006691 | 0.498499 | F-box-like protein | elongation zone | 12 h |
| HORVU.MOREX.r3.2HG0112030.1 | 0.516722 | 5.79888 | 4.034679 | 0.000177 | 0.006165 | 0.498297 | B3 domain-containing transcription factor VAL3 | elongation zone | 12 h |
| HORVU.MOREX.r3.6HG0615840.1 | 2.786594 | -2.2741 | 4.528701 | 3.40E-05 | 0.002987 | 0.496025 | Metalloendoproteinase 1 | elongation zone | 12 h |
| HORVU.MOREX.r3.4HG0406670.1 | 0.814044 | 6.546221 | 4.069503 | 0.000158 | 0.00589 | 0.495065 | Classical arabinogalactan protein 5 | elongation zone | 12 h |
| HORVU.MOREX.r3.3HG0255220.1 | -0.99854 | 5.459644 | -3.9754 | 0.000214 | 0.006671 | 0.493668 | Geranylgeranyl reductase | elongation zone | 12 h |
| HORVU.MOREX.r3.6HG0545970.1 | -0.48011 | 5.654969 | -4.01071 | 0.000191 | 0.006387 | 0.491545 | Short-chain dehydrogenase/reductase family protein | elongation zone | 12 h |
| HORVU.MOREX.r3.2HG0186680.1 | -0.69402 | 7.556619 | -4.06168 | 0.000162 | 0.005955 | 0.489932 | Histone H1 | elongation zone | 12 h |
| HORVU.MOREX.r3.2HG0165920.1 | 1.323692 | 1.852926 | 3.95265 | 0.00023 | 0.006891 | 0.489524 | Purple acid phosphatase | elongation zone | 12 h |
| HORVU.MOREX.r3.7HG0661450.1 | 0.669138 | 6.337893 | 4.07787 | 0.000153 | 0.005816 | 0.487059 | Phosphoinositide phosphatase family protein | elongation zone | 12 h |
| HORVU.MOREX.r3.4HG0409230.1 | 1.858508 | -0.03281 | 3.989212 | 0.000205 | 0.006554 | 0.486208 | IQ domain-containing protein | elongation zone | 12 h |
| HORVU.MOREX.r3.5HG0524010.1 | -1.3108 | 6.200507 | -3.98427 | 0.000208 | 0.006583 | 0.48512 | Glycosyltransferase | elongation zone | 12 h |
| HORVU.MOREX.r3.5HG0495730.1 | 1.153289 | 2.468419 | 3.974701 | 0.000214 | 0.006676 | 0.482125 | Kinase family protein | elongation zone | 12 h |
| HORVU.MOREX.r3.3HG0306060.1 | -0.3737 | 5.417997 | -4.00749 | 0.000193 | 0.006402 | 0.47904 | Thioredoxin-like protein | elongation zone | 12 h |
| HORVU.MOREX.r3.2HG0183780.1 | -0.55786 | 5.376384 | -4.00045 | 0.000197 | 0.006444 | 0.478889 | Sphingolipid delta-4 desaturase | elongation zone | 12 h |

**Table S1** Continued.

| ID | logFC | AveExpr | t | P.Value | adj.P.Val | B | gene description | root zone | time point |
|---|---|---|---|---|---|---|---|---|---|
| HORVU.MOREX.r3.5HG0532150.1 | 2.123181 | -2.26204 | 3.983079 | 0.000209 | 0.006588 | 0.478194 | Expansin | elongation zone | 12 h |
| HORVU.MOREX.r3.6HG0633900.1 | 1.826189 | -0.51261 | 3.970636 | 0.000217 | 0.006691 | 0.477093 | Calcium-dependent protein kinase | elongation zone | 12 h |
| HORVU.MOREX.r3.7HG0679760.1 | -0.56827 | 6.056962 | -4.01322 | 0.000189 | 0.006367 | 0.471735 | Basic helix-loop-helix transcription factor | elongation zone | 12 h |
| HORVU.MOREX.r3.6HG0618630.1 | -0.60317 | 6.66233 | -4.06035 | 0.000162 | 0.00597 | 0.467752 | Glucan endo-1,3-beta-glucosidase 3 | elongation zone | 12 h |
| HORVU.MOREX.r3.1HG0075440.1 | 0.832065 | 1.650312 | 3.970108 | 0.000218 | 0.006691 | 0.465883 | Protein NRT1/ PTR FAMILY 5.5 | elongation zone | 12 h |
| HORVU.MOREX.r3.6HG0609100.1 | -1.11456 | 1.570415 | -3.99474 | 0.000201 | 0.006494 | 0.464721 | 3-hydroxy-3-methylglutaryl coenzyme A reductase | elongation zone | 12 h |
| HORVU.MOREX.r3.4HG0383870.1 | -0.79619 | 7.088832 | -4.01989 | 0.000185 | 0.006303 | 0.463059 | Malonyl CoA-acyl carrier protein transacylase containing protein, expressed | elongation zone | 12 h |
| HORVU.MOREX.r3.4HG0395940.1 | 1.45687 | 3.064976 | 3.941134 | 0.000239 | 0.006937 | 0.460473 | Homocysteine S-methyltransferase family protein | elongation zone | 12 h |
| HORVU.MOREX.r3.6HG0543740.1 | -0.9818 | 7.842285 | -4.0446 | 0.000171 | 0.006089 | 0.460458 | Histone H2A | elongation zone | 12 h |
| HORVU.MOREX.r3.2HG0140170.1 | 0.740702 | 2.981209 | 3.944158 | 0.000237 | 0.006926 | 0.459744 | Ras-related protein Rab-21 | elongation zone | 12 h |
| HORVU.MOREX.r3.6HG0568910.1 | 0.82335 | 3.99892 | 4.025904 | 0.000182 | 0.00626 | 0.455537 | L-gulonolactone oxidase | elongation zone | 12 h |
| HORVU.MOREX.r3.2HG0214090.1 | -1.07901 | 7.152075 | -3.98803 | 0.000205 | 0.006554 | 0.450429 | C2 calcium/lipid-binding and GRAM domain protein | elongation zone | 12 h |
| HORVU.MOREX.r3.1HG0056280.1 | -1.05304 | 3.39368 | -3.93948 | 0.00024 | 0.006944 | 0.449166 | Xyloglucan galactosyltransferase KATAMARI1 | elongation zone | 12 h |
| HORVU.MOREX.r3.6HG0547660.1 | 2.391812 | -1.78969 | 4.063885 | 0.00016 | 0.005948 | 0.440347 | Copine-1 | elongation zone | 12 h |
| HORVU.MOREX.r3.2HG0127260.1 | 1.313234 | -1.06197 | 3.949596 | 0.000233 | 0.006891 | 0.438906 | COBRA-like protein | elongation zone | 12 h |
| HORVU.MOREX.r3.4HG0415490.1 | 1.916614 | -1.78464 | 3.931274 | 0.000247 | 0.007038 | 0.434288 | 14 kDa proline-rich protein DC2.15 | elongation zone | 12 h |
| HORVU.MOREX.r3.2HG0192900.1 | -1.32098 | 6.268762 | -3.95666 | 0.000227 | 0.006853 | 0.432205 | Xyloglucan endotransglucosylase/hydrolase | elongation zone | 12 h |
| HORVU.MOREX.r3.7HG0714910.1 | -0.70352 | 7.21685 | -4.02456 | 0.000182 | 0.00626 | 0.426256 | #N/A | elongation zone | 12 h |
| HORVU.MOREX.r3.5HG0512490.1 | 1.663332 | 1.425049 | 4.01258 | 0.00019 | 0.006369 | 0.426193 | Peroxidase | elongation zone | 12 h |
| HORVU.MOREX.r3.3HG0242150.1 | 3.271829 | -3.21832 | 4.261719 | 8.36E-05 | 0.004425 | 0.422949 | Glycine-rich cell wall structural protein | elongation zone | 12 h |
| HORVU.MOREX.r3.5HG0495840.1 | -0.39103 | 5.657016 | -3.98949 | 0.000204 | 0.006554 | 0.422198 | Protein farnesyltransferase/geranylgeranyltransferase type-1 subunit alpha | elongation zone | 12 h |
| HORVU.MOREX.r3.7HG0714930.1 | -0.39796 | 7.749002 | -4.05239 | 0.000167 | 0.006062 | 0.420028 | UDP-galactose transporter, putative | elongation zone | 12 h |
| HORVU.MOREX.r3.1HG0064220.1 | -0.75906 | 4.418073 | -3.92612 | 0.000251 | 0.007078 | 0.416623 | Isoamylase-like glucan debranching enzyme | elongation zone | 12 h |
| HORVU.MOREX.r3.6HG0575690.1 | -0.85327 | 3.926859 | -3.92597 | 0.000251 | 0.007078 | 0.415734 | Exostosin-like | elongation zone | 12 h |
| HORVU.MOREX.r3.6HG0588520.1 | 0.971566 | 2.56033 | 3.985751 | 0.000207 | 0.006561 | 0.411471 | Rop guanine nucleotide exchange factor | elongation zone | 12 h |
| HORVU.MOREX.r3.6HG0595860.1 | -0.45682 | 5.620865 | -3.9715 | 0.000217 | 0.006691 | 0.409036 | E3 ubiquitin-protein ligase | elongation zone | 12 h |
| HORVU.MOREX.r3.1HG0085420.1 | -0.75789 | 3.718027 | -3.92016 | 0.000256 | 0.007125 | 0.407661 | DNA replication complex GINS protein PSF2 | elongation zone | 12 h |
| HORVU.MOREX.r3.1HG0061660.1 | 0.986173 | 3.333271 | 3.986796 | 0.000206 | 0.006561 | 0.406461 | RING/FYVE/PHD zinc finger superfamily protein | elongation zone | 12 h |
| HORVU.MOREX.r3.2HG0206470.1 | 0.55066 | 6.2777 | 4.039803 | 0.000174 | 0.006131 | 0.401113 | Loricrin-like protein | elongation zone | 12 h |
| HORVU.MOREX.r3.5HG0444060.1 | -1.08418 | 1.863328 | -3.93464 | 0.000244 | 0.007006 | 0.399591 | Calmodulin-binding family protein, putative, expressed | elongation zone | 12 h |
| HORVU.MOREX.r3.5HG0465690.1 | 0.828296 | 3.37211 | 3.945746 | 0.000235 | 0.006919 | 0.398279 | Phytol kinase | elongation zone | 12 h |
| HORVU.MOREX.r3.6HG0626070.1 | 0.569575 | 3.613021 | 3.954274 | 0.000229 | 0.006881 | 0.393097 | UDP-N-acetylglucosamine--N-acetylmuramyl-(Pentapeptide) pyrophosphoryl-unde | elongation zone | 12 h |
| HORVU.MOREX.r3.7HG0739480.1 | -0.88395 | 7.565794 | -4.00421 | 0.000195 | 0.006419 | 0.392692 | Histone H2B | elongation zone | 12 h |
| HORVU.MOREX.r3.3HG0252170.1 | 1.656456 | -1.70754 | 3.930232 | 0.000247 | 0.007043 | 0.390328 | DUF1685 family protein | elongation zone | 12 h |
| HORVU.MOREX.r3.5HG0511860.1 | -0.92938 | 6.516686 | -3.98609 | 0.000207 | 0.006561 | 0.384364 | Blue copper protein | elongation zone | 12 h |
| HORVU.MOREX.r3.5HG0488040.1 | 1.623253 | -1.23014 | 3.921476 | 0.000255 | 0.007125 | 0.383407 | Blue copper protein | elongation zone | 12 h |
| HORVU.MOREX.r3.7HG0674750.1 | -0.4465 | 5.335659 | -3.96543 | 0.000221 | 0.006756 | 0.383282 | Dual specificity protein phosphatase, putative | elongation zone | 12 h |
| HORVU.MOREX.r3.3HG0225610.1 | -0.86364 | 2.476199 | -3.93357 | 0.000245 | 0.007006 | 0.382941 | Mediator of RNA polymerase II transcription subunit 22 | elongation zone | 12 h |
| HORVU.MOREX.r3.4HG0345990.1 | 0.511432 | 6.344145 | 4.00819 | 0.000192 | 0.006402 | 0.380012 | GDP-L-galactose phosphorylase 1 | elongation zone | 12 h |
| HORVU.MOREX.r3.7HG0650300.1 | 0.67397 | 3.514135 | 3.942036 | 0.000238 | 0.006937 | 0.37908 | Protein kinase | elongation zone | 12 h |
| HORVU.MOREX.r3.2HG0182020.2 | -1.34584 | 3.196873 | -3.90942 | 0.000265 | 0.00721 | 0.378139 | Phenylalanine ammonia-lyase | elongation zone | 12 h |

**Table S1** Continued.

| ID | logFC | AveExpr | t | P.Value | adj.P.Val | B | gene description | root zone | time point |
|---|---|---|---|---|---|---|---|---|---|
| HORVU.MOREX.r3.4HG0381990.1 | 1.46516 | 1.592907 | 3.972147 | 0.000216 | 0.006691 | 0.377003 | Pre-mRNA splicing factor, putative | elongation zone | 12 h |
| HORVU.MOREX.r3.5HG0426280.1 | 0.913367 | 2.961285 | 3.930876 | 0.000247 | 0.007038 | 0.375538 | GTP binding protein | elongation zone | 12 h |
| HORVU.MOREX.r3.6HG0606140.1 | 2.005058 | -0.36014 | 3.958801 | 0.000226 | 0.006833 | 0.375461 | Pentatricopeptide repeat-containing protein | elongation zone | 12 h |
| HORVU.MOREX.r3.5HG0531450.1 | 0.914179 | -1.15012 | 3.925691 | 0.000251 | 0.007078 | 0.373971 | Speckle-type POZ protein-like protein | elongation zone | 12 h |
| HORVU.MOREX.r3.1HG0056560.1 | -0.39174 | 5.466997 | -3.96952 | 0.000218 | 0.006691 | 0.373742 | AP-3 complex subunit delta | elongation zone | 12 h |
| HORVU.MOREX.r3.1HG0048030.1 | -0.46033 | 7.472589 | -4.01437 | 0.000189 | 0.006364 | 0.372158 | Membrane steroid-binding protein | elongation zone | 12 h |
| HORVU.MOREX.r3.7HG0721760.1 | -0.3845 | 8.4211 | -4.05685 | 0.000164 | 0.006028 | 0.368885 | Translocon-associated protein subunit alpha | elongation zone | 12 h |
| HORVU.MOREX.r3.6HG0629520.1 | 1.528497 | -0.9319 | 3.915365 | 0.00026 | 0.007125 | 0.365732 | Ethylene-responsive transcription factor | elongation zone | 12 h |
| HORVU.MOREX.r3.1HG0074500.1 | -1.07864 | 7.370525 | -3.98045 | 0.00021 | 0.006618 | 0.360713 | Histone H3 | elongation zone | 12 h |
| HORVU.MOREX.r3.2HG0128890.1 | -0.68362 | 5.994209 | -3.95872 | 0.000226 | 0.006833 | 0.360124 | RAN GTPASE ACTIVATING protein 2 | elongation zone | 12 h |
| HORVU.MOREX.r3.1HG0077880.1 | -1.07543 | 3.552074 | -3.91005 | 0.000264 | 0.007205 | 0.358976 | BAG family molecular chaperone regulator 7 | elongation zone | 12 h |
| HORVU.MOREX.r3.7HG0705700.1 | -0.6989 | 3.946655 | -3.91098 | 0.000263 | 0.007193 | 0.35868 | Calcium ion-binding protein | elongation zone | 12 h |
| HORVU.MOREX.r3.5HG0438750.1 | -0.72831 | 5.420678 | -3.94797 | 0.000234 | 0.006891 | 0.357 | Disease resistance protein (NBS-LRR class) family | elongation zone | 12 h |
| HORVU.MOREX.r3.1HG0054500.1 | -0.97984 | 5.920469 | -3.9493 | 0.000233 | 0.006891 | 0.353256 | #N/A | elongation zone | 12 h |
| HORVU.MOREX.r3.1HG0078750.1 | -0.56448 | 3.941561 | -3.90589 | 0.000268 | 0.007263 | 0.350547 | Tesmin/TSO1-like CXC domain containing protein | elongation zone | 12 h |
| HORVU.MOREX.r3.4HG0354970.1 | -0.68025 | 6.732308 | -3.98592 | 0.000207 | 0.006561 | 0.350426 | Omega-3 fatty acid desaturase | elongation zone | 12 h |
| HORVU.MOREX.r3.3HG0299010.1 | -1.07148 | 2.065005 | -3.91814 | 0.000257 | 0.007125 | 0.346533 | Protein NRT1/ PTR FAMILY 5.5 | elongation zone | 12 h |
| HORVU.MOREX.r3.7HG0650980.1 | -0.9585 | 5.5668 | -3.93719 | 0.000242 | 0.006974 | 0.343837 | Histone H3 | elongation zone | 12 h |
| HORVU.MOREX.r3.4HG0340920.1 | -0.43794 | 4.789685 | -3.9192 | 0.000256 | 0.007125 | 0.341981 | RNA-binding KH domain-containing protein | elongation zone | 12 h |
| HORVU.MOREX.r3.7HG0741080.1 | -0.83663 | 1.851569 | -3.90006 | 0.000273 | 0.007364 | 0.340277 | Disease resistance protein RPM1 | elongation zone | 12 h |
| HORVU.MOREX.r3.3HG0250060.1 | 1.991888 | -2.50642 | 4.147566 | 0.000122 | 0.005349 | 0.338129 | CASP-like protein | elongation zone | 12 h |
| HORVU.MOREX.r3.1HG0074260.1 | -0.40016 | 7.572556 | -4.01556 | 0.000188 | 0.006364 | 0.335415 | Beta-glucosidase | elongation zone | 12 h |
| HORVU.MOREX.r3.4HG0370520.1 | -0.74284 | 3.910658 | -3.89647 | 0.000276 | 0.007381 | 0.335302 | inactive purple acid phosphatase-like protein | elongation zone | 12 h |
| HORVU.MOREX.r3.4HG0378330.1 | 1.261165 | 0.037978 | 3.920412 | 0.000255 | 0.007125 | 0.331573 | transcription repressor | elongation zone | 12 h |
| HORVU.MOREX.r3.6HG0620630.1 | 1.012137 | 3.35364 | 4.02034 | 0.000185 | 0.006303 | 0.331391 | Subtilisin-like protease | elongation zone | 12 h |
| HORVU.MOREX.r3.3HG0244240.1 | 1.222215 | 3.630904 | 3.914963 | 0.00026 | 0.007125 | 0.330616 | Alpha/beta-Hydrolases superfamily protein | elongation zone | 12 h |
| HORVU.MOREX.r3.4HG0398550.1 | 1.049275 | 6.045114 | 4.052829 | 0.000166 | 0.006062 | 0.330555 | Calcium-transporting ATPase | elongation zone | 12 h |
| HORVU.MOREX.r3.1HG0055230.1 | -0.88924 | 7.590256 | -4.00424 | 0.000195 | 0.006419 | 0.330418 | Methyltransferase | elongation zone | 12 h |
| HORVU.MOREX.r3.5HG0513810.1 | 0.629718 | 4.245645 | 3.952317 | 0.00023 | 0.006891 | 0.330305 | Malate dehydrogenase | elongation zone | 12 h |
| HORVU.MOREX.r3.5HG0460160.1 | 0.430942 | 6.54551 | 4.024865 | 0.000182 | 0.00626 | 0.330281 | Bifunctional uridylyltransferase/uridylyl-removing enzyme | elongation zone | 12 h |
| HORVU.MOREX.r3.5HG0533230.1 | -0.55534 | 5.643177 | -3.96252 | 0.000223 | 0.00679 | 0.330258 | Pirin-like protein | elongation zone | 12 h |
| HORVU.MOREX.r3.3HG0230190.1 | -1.17966 | 2.957931 | -3.94024 | 0.00024 | 0.006944 | 0.32855 | #N/A | elongation zone | 12 h |
| HORVU.MOREX.r3.3HG0309460.1 | -0.33386 | 5.431182 | -3.95887 | 0.000226 | 0.006833 | 0.326114 | Trichome birefringence-like protein | elongation zone | 12 h |
| HORVU.MOREX.r3.1HG0016810.1 | -0.81163 | 5.521689 | -3.91477 | 0.00026 | 0.007125 | 0.320822 | Agenet domain-containing protein | elongation zone | 12 h |
| HORVU.MOREX.r3.7HG0730070.1 | -1.33555 | 6.723992 | -3.94509 | 0.000236 | 0.006919 | 0.318158 | Thaumatin-like protein | elongation zone | 12 h |
| HORVU.MOREX.r3.3HG0252250.1 | 1.603359 | -1.6584 | 3.891444 | 0.00028 | 0.007449 | 0.318073 | Expansin | elongation zone | 12 h |
| HORVU.MOREX.r3.4HG0335180.1 | -0.63969 | 5.021928 | -3.91484 | 0.00026 | 0.007125 | 0.31323 | Phosphatidylinositol N-acetyglucosaminlytransferase subunit P-like protein | elongation zone | 12 h |
| HORVU.MOREX.r3.1HG0012340.1 | 1.253337 | 2.437594 | 3.939378 | 0.00024 | 0.006944 | 0.312979 | Triose-phosphate transporter family protein | elongation zone | 12 h |
| HORVU.MOREX.r3.2HG0197560.1 | -0.76776 | 5.104956 | -3.89722 | 0.000275 | 0.00738 | 0.303155 | RNA-binding protein 39 | elongation zone | 12 h |
| HORVU.MOREX.r3.3HG0254930.1 | 2.689498 | -0.9917 | 3.908716 | 0.000265 | 0.007214 | 0.300402 | Peroxidase | elongation zone | 12 h |
| HORVU.MOREX.r3.2HG0196800.1 | 2.024398 | -0.20994 | 3.999539 | 0.000198 | 0.006444 | 0.298898 | Multidrug resistance protein ABC transporter family | elongation zone | 12 h |

**Table S1** Continued.

| ID | logFC | AveExpr | t | P.Value | adj.P.Val | B | gene description | root zone | time point |
|---|---|---|---|---|---|---|---|---|---|
| HORVU.MOREX.r3.5HG0490830.1 | -0.41664 | 5.180095 | -3.92195 | 0.000254 | 0.007125 | 0.296014 | 2-oxoglutarate (2OG) and Fe(II)-dependent oxygenase superfamily protein | elongation zone | 12 h |
| HORVU.MOREX.r3.4HG0340910.2 | 2.527565 | -1.88941 | 4.135578 | 0.000127 | 0.005414 | 0.292021 | p-loop containing nucleoside triphosphate hydrolases superfamily protein | elongation zone | 12 h |
| HORVU.MOREX.r3.1HG0016710.2 | 1.755892 | -3.14485 | 4.085564 | 0.00015 | 0.0058 | 0.290984 | Annexin | elongation zone | 12 h |
| HORVU.MOREX.r3.1HG0067980.1 | 1.560748 | -1.60301 | 3.881814 | 0.000289 | 0.007575 | 0.289972 | Peroxidase | elongation zone | 12 h |
| HORVU.MOREX.r3.7HG0746230.1 | -0.60374 | 6.353103 | -3.95642 | 0.000227 | 0.006853 | 0.28833 | Receptor protein kinase-like protein | elongation zone | 12 h |
| HORVU.MOREX.r3.6HG0612780.1 | 1.533597 | 1.073388 | 3.917698 | 0.000258 | 0.007125 | 0.287468 | Dynein light chain | elongation zone | 12 h |
| HORVU.MOREX.r3.3HG0256580.1 | -0.49073 | 6.795309 | -3.95493 | 0.000229 | 0.006876 | 0.285959 | NEDD8-activating enzyme E1 catalytic subunit | elongation zone | 12 h |
| HORVU.MOREX.r3.7HG0729000.1 | 0.979307 | 3.645636 | 3.898164 | 0.000274 | 0.007375 | 0.285877 | RING/U-box superfamily protein | elongation zone | 12 h |
| HORVU.MOREX.r3.6HG0606940.1 | -3.08142 | 2.180677 | -4.39605 | 5.33E-05 | 0.003734 | 0.281793 | Pentatricopeptide repeat-containing protein | elongation zone | 12 h |
| HORVU.MOREX.r3.5HG0480980.1 | 2.465734 | -1.63821 | 3.899867 | 0.000273 | 0.007364 | 0.279174 | Spermidine/putrescine import ATP-binding protein PotA | elongation zone | 12 h |
| HORVU.MOREX.r3.3HG0288240.1 | -0.69902 | 5.202677 | -3.89632 | 0.000276 | 0.007381 | 0.274004 | ubiquitin fusion degradation 1 | elongation zone | 12 h |
| HORVU.MOREX.r3.2HG0171550.1 | -0.81251 | 4.111529 | -3.87687 | 0.000294 | 0.007636 | 0.271834 | Major facilitator superfamily transporter | elongation zone | 12 h |
| HORVU.MOREX.r3.5HG0520990.1 | -0.65535 | 3.822638 | -3.8797 | 0.000291 | 0.007597 | 0.270986 | Kinase, putative | elongation zone | 12 h |
| HORVU.MOREX.r3.6HG0601260.1 | 1.107052 | 3.050364 | 3.948217 | 0.000234 | 0.006891 | 0.268642 | Sulfotransferase | elongation zone | 12 h |
| HORVU.MOREX.r3.6HG0624630.1 | -0.44344 | 6.985603 | -3.97928 | 0.000211 | 0.006618 | 0.264482 | Aspartate aminotransferase | elongation zone | 12 h |
| HORVU.MOREX.r3.4HG0342850.1 | -0.4054 | 5.711387 | -3.92415 | 0.000252 | 0.007103 | 0.262195 | Serpin family protein | elongation zone | 12 h |
| HORVU.MOREX.r3.6HG0604860.1 | -1.09537 | 1.903058 | -3.93412 | 0.000244 | 0.007006 | 0.258698 | Glycosyltransferase | elongation zone | 12 h |
| HORVU.MOREX.r3.1HG0069290.1 | -0.33331 | 8.209881 | -4.01783 | 0.000186 | 0.006334 | 0.257858 | Nucleotide-sugar transporter family protein | elongation zone | 12 h |
| HORVU.MOREX.r3.6HG0573920.1 | 1.475431 | -0.43481 | 3.944927 | 0.000236 | 0.006919 | 0.252712 | Retrotransposon protein, putative, unclassified | elongation zone | 12 h |
| HORVU.MOREX.r3.6HG0603530.1 | -1.09117 | 2.451293 | -3.87968 | 0.000291 | 0.007597 | 0.252432 | Protein DETOXIFICATION | elongation zone | 12 h |
| HORVU.MOREX.r3.1HG0087620.1 | 1.053131 | 3.06372 | 3.962767 | 0.000223 | 0.00679 | 0.251206 | Xyloglucan endotransglucosylase/hydrolase | elongation zone | 12 h |
| HORVU.MOREX.r3.2HG0115780.1 | -1.18166 | 6.198436 | -3.95077 | 0.000232 | 0.006891 | 0.247731 | FAD-binding Berberine family protein, putative | elongation zone | 12 h |
| HORVU.MOREX.r3.2HG0187440.1 | -0.53745 | 5.452931 | -3.91678 | 0.000258 | 0.007125 | 0.241392 | S-adenosyl-L-methionine-dependent methyltransferases superfamily protein | elongation zone | 12 h |
| HORVU.MOREX.r3.6HG0631600.1 | 1.124704 | 1.162908 | 3.938525 | 0.000241 | 0.006954 | 0.23913 | Rop guanine nucleotide exchange factor, putative | elongation zone | 12 h |
| HORVU.MOREX.r3.2HG0155810.1 | -0.94308 | 3.567204 | -3.86139 | 0.000309 | 0.007759 | 0.238582 | Replication protein A 70 kDa DNA-binding subunit A | elongation zone | 12 h |
| HORVU.MOREX.r3.1HG0023200.1 | 1.271124 | 0.569684 | 3.884635 | 0.000286 | 0.007556 | 0.23679 | LURP-one-like protein | elongation zone | 12 h |
| HORVU.MOREX.r3.3HG0287690.1 | 0.932619 | 2.646815 | 3.943227 | 0.000237 | 0.006937 | 0.234015 | Profilin | elongation zone | 12 h |
| HORVU.MOREX.r3.1HG0001510.1 | -1.06579 | 3.140127 | -3.86222 | 0.000308 | 0.007759 | 0.230565 | NBS-LRR-like resistance protein | elongation zone | 12 h |
| HORVU.MOREX.r3.4HG0384110.1 | -0.37757 | 6.300031 | -3.94887 | 0.000233 | 0.006891 | 0.229437 | RING/U-box superfamily protein | elongation zone | 12 h |
| HORVU.MOREX.r3.2HG0097360.1 | -0.82577 | 5.914899 | -3.90289 | 0.00027 | 0.007323 | 0.22628 | Laccase | elongation zone | 12 h |
| HORVU.MOREX.r3.3HG0286160.1 | -1.74814 | 6.290215 | -3.88114 | 0.00029 | 0.007581 | 0.226209 | Pollen Ole e 1 allergen/extensin | elongation zone | 12 h |
| HORVU.MOREX.r3.3HG0242030.1 | -1.66538 | 2.331268 | -3.94825 | 0.000234 | 0.006891 | 0.222702 | Alpha/beta-Hydrolases superfamily protein | elongation zone | 12 h |
| HORVU.MOREX.r3.6HG0555660.2 | -0.43508 | 7.208638 | -3.96976 | 0.000218 | 0.006691 | 0.221947 | Calreticulin/calnexin | elongation zone | 12 h |
| HORVU.MOREX.r3.2HG0113880.1 | 0.62138 | 4.823614 | 3.908391 | 0.000265 | 0.007214 | 0.220577 | Heavy metal transport/detoxification superfamily protein, putative | elongation zone | 12 h |
| HORVU.MOREX.r3.5HG0426210.3 | 0.861042 | 1.98152 | 3.861529 | 0.000308 | 0.007759 | 0.219907 | Protein IQ-DOMAIN 1 | elongation zone | 12 h |
| HORVU.MOREX.r3.2HG0192400.1 | 0.785664 | 3.506293 | 3.862664 | 0.000307 | 0.007759 | 0.20101 | Receptor protein kinase, putative | elongation zone | 12 h |
| HORVU.MOREX.r3.6HG0622110.1 | 1.423774 | 2.81443 | 3.922937 | 0.000253 | 0.007121 | 0.200877 | Pathogenesis-related protein 1 | elongation zone | 12 h |
| HORVU.MOREX.r3.6HG0593200.1 | -1.018 | 4.784544 | -3.86135 | 0.000309 | 0.007759 | 0.200011 | Short-chain dehydrogenase/reductase family protein | elongation zone | 12 h |
| HORVU.MOREX.r3.2HG0157290.1 | -1.07392 | 3.397681 | -3.85029 | 0.00032 | 0.007921 | 0.198771 | Calcineurin-like metallo-phosphoesterase superfamily | elongation zone | 12 h |
| HORVU.MOREX.r3.1HG0086370.1 | 2.105225 | -3.20911 | 4.108594 | 0.000139 | 0.005596 | 0.198412 | Glyoxal oxidase | elongation zone | 12 h |
| HORVU.MOREX.r3.4HG0342660.1 | 1.266999 | 1.013503 | 3.92803 | 0.000249 | 0.007073 | 0.19738 | Calcium-binding EF-hand family protein | elongation zone | 12 h |

**Table S1** Continued.

| ID | logFC | AveExpr | t | P.Value | adj.P.Val | B | gene description | root zone | time point |
|---|---|---|---|---|---|---|---|---|---|
| HORVU.MOREX.r3.5HG0485610.1 | 1.357976 | 0.79472 | 3.845829 | 0.000324 | 0.007986 | 0.195993 | Cytochrome P450, putative | elongation zone | 12 h |
| HORVU.MOREX.r3.5HG0420460.1 | 2.796521 | -1.03028 | 4.319064 | 6.90E-05 | 0.004083 | 0.195936 | NBS-LRR-like resistance protein | elongation zone | 12 h |
| HORVU.MOREX.r3.3HG0256380.1 | -0.85666 | 6.689113 | -3.91745 | 0.000258 | 0.007125 | 0.193908 | Ubiquitin-conjugating enzyme E2 | elongation zone | 12 h |
| HORVU.MOREX.r3.2HG0139370.1 | -0.79417 | 6.338385 | -3.92966 | 0.000248 | 0.007046 | 0.188684 | Xyloglucan endotransglucosylase/hydrolase | elongation zone | 12 h |
| HORVU.MOREX.r3.2HG0203070.1 | 0.872154 | 3.219858 | 3.885153 | 0.000286 | 0.007556 | 0.186942 | Receptor-like kinase | elongation zone | 12 h |
| HORVU.MOREX.r3.5HG0489840.1 | -1.11502 | 2.789323 | -3.88458 | 0.000286 | 0.007556 | 0.186721 | #N/A | elongation zone | 12 h |
| HORVU.MOREX.r3.3HG0304710.1 | -0.34666 | 8.270238 | -4.0082 | 0.000192 | 0.006402 | 0.184934 | Fructose-bisphosphate aldolase | elongation zone | 12 h |
| HORVU.MOREX.r3.3HG0321700.1 | 0.683543 | 3.776226 | 3.853653 | 0.000316 | 0.007898 | 0.183987 | FMN-dependent NADPH-azoreductase | elongation zone | 12 h |
| HORVU.MOREX.r3.7HG0739080.1 | 2.344023 | -3.17965 | 4.174016 | 0.000112 | 0.005113 | 0.182638 | Subtilisin-like protease | elongation zone | 12 h |
| HORVU.MOREX.r3.2HG0128840.1 | -0.59474 | 4.89057 | -3.89289 | 0.000279 | 0.007434 | 0.179281 | Son of sevenless | elongation zone | 12 h |
| HORVU.MOREX.r3.3HG0248440.1 | 0.696104 | 4.338634 | 3.945978 | 0.000235 | 0.006919 | 0.179012 | Ras-like protein | elongation zone | 12 h |
| HORVU.MOREX.r3.1HG0056240.1 | 0.741885 | 4.721848 | 3.916231 | 0.000259 | 0.007125 | 0.178954 | Trehalose 6-phosphate phosphatase | elongation zone | 12 h |
| HORVU.MOREX.r3.4HG0337040.1 | 0.991278 | 2.764027 | 3.867206 | 0.000303 | 0.007757 | 0.17738 | Accelerated cell death 11 family protein | elongation zone | 12 h |
| HORVU.MOREX.r3.2HG0190680.1 | -0.48109 | 6.930995 | -3.92636 | 0.000251 | 0.007078 | 0.173461 | Aminotransferase | elongation zone | 12 h |
| HORVU.MOREX.r3.2HG0178160.1 | -0.65043 | 7.879945 | -3.95685 | 0.000227 | 0.006853 | 0.169735 | HTH-type transcriptional regulator YidZ | elongation zone | 12 h |
| HORVU.MOREX.r3.6HG0600910.1 | 1.324317 | 1.060689 | 3.919899 | 0.000256 | 0.007125 | 0.165547 | Aquaporin | elongation zone | 12 h |
| HORVU.MOREX.r3.2HG0182010.1 | -0.852 | 6.691165 | -3.93288 | 0.000245 | 0.007012 | 0.164458 | Phenylalanine ammonia-lyase | elongation zone | 12 h |
| HORVU.MOREX.r3.3HG0283080.2 | 1.024447 | 1.384373 | 3.879057 | 0.000292 | 0.007602 | 0.160765 | suppressor | elongation zone | 12 h |
| HORVU.MOREX.r3.6HG0632020.2 | 1.448126 | -2.23983 | 3.869996 | 0.0003 | 0.007754 | 0.160369 | Photosystem II 22 kDa, chloroplastic | elongation zone | 12 h |
| HORVU.MOREX.r3.7HG0685970.1 | 0.769168 | 4.80192 | 3.969522 | 0.000218 | 0.006691 | 0.157807 | Kinase family protein | elongation zone | 12 h |
| HORVU.MOREX.r3.3HG0299280.1 | -0.73257 | 4.635338 | -3.85836 | 0.000311 | 0.007805 | 0.155665 | WD-repeat protein, putative | elongation zone | 12 h |
| HORVU.MOREX.r3.5HG0486660.1 | 1.249554 | -2.06977 | 3.836147 | 0.000334 | 0.008157 | 0.151516 | Calcium-binding protein | elongation zone | 12 h |
| HORVU.MOREX.r3.1HG0053060.1 | 1.041453 | 4.011378 | 3.852947 | 0.000317 | 0.007902 | 0.148877 | PLATZ transcription factor | elongation zone | 12 h |
| HORVU.MOREX.r3.5HG0432640.1 | -0.38942 | 5.94978 | -3.89248 | 0.000279 | 0.007434 | 0.139617 | Protein O-linked-mannose beta-1,4-N-acetylglucosaminyltransferase 2 | elongation zone | 12 h |
| HORVU.MOREX.r3.5HG0469300.1 | 1.644501 | 0.250216 | 3.824648 | 0.000347 | 0.008288 | 0.138985 | CCCH-type zinc finger protein with ARM repeat domain-containing protein | elongation zone | 12 h |
| HORVU.MOREX.r3.6HG0622300.1 | -0.79426 | 7.348551 | -3.92727 | 0.00025 | 0.007078 | 0.134636 | Pathogenesis-related protein 1 | elongation zone | 12 h |
| HORVU.MOREX.r3.7HG0741640.1 | -0.96463 | 4.20973 | -3.83382 | 0.000337 | 0.008177 | 0.133135 | Histone H2A | elongation zone | 12 h |
| HORVU.MOREX.r3.1HG0065520.1 | 0.525843 | 5.125584 | 3.915944 | 0.000259 | 0.007125 | 0.132262 | KH domain-containing protein | elongation zone | 12 h |
| HORVU.MOREX.r3.3HG0293130.1 | 1.088174 | 3.772343 | 3.860562 | 0.000309 | 0.007759 | 0.131482 | Eukaryotic aspartyl protease family protein | elongation zone | 12 h |
| HORVU.MOREX.r3.6HG0558840.1 | -0.29144 | 6.872948 | -3.91756 | 0.000258 | 0.007125 | 0.130073 | Kinetochore protein nuf2, putative | elongation zone | 12 h |
| HORVU.MOREX.r3.6HG0548710.1 | 2.180602 | -1.47311 | 3.823767 | 0.000348 | 0.008288 | 0.128125 | Kinase family protein | elongation zone | 12 h |
| HORVU.MOREX.r3.2HG0193930.1 | -0.69923 | 3.655261 | -3.83366 | 0.000337 | 0.008177 | 0.126367 | ATP synthase epsilon chain | elongation zone | 12 h |
| HORVU.MOREX.r3.2HG0122080.1 | 0.592342 | 3.732882 | 3.899137 | 0.000273 | 0.007364 | 0.123185 | DUF630 family protein, putative (DUF630 and DUF632) | elongation zone | 12 h |
| HORVU.MOREX.r3.5HG0513440.1 | -0.40915 | 5.701505 | -3.89517 | 0.000277 | 0.007399 | 0.122818 | #N/A | elongation zone | 12 h |
| HORVU.MOREX.r3.1HG0026830.1 | 0.629926 | 3.566935 | 3.881972 | 0.000289 | 0.007575 | 0.121332 | Leucine-rich repeat receptor-like protein kinase family | elongation zone | 12 h |
| HORVU.MOREX.r3.4HG0337120.1 | -0.65051 | 5.989451 | -3.88586 | 0.000285 | 0.007554 | 0.116931 | Histone H1, putative | elongation zone | 12 h |
| HORVU.MOREX.r3.5HG0485940.1 | 1.857338 | 2.592249 | 3.824069 | 0.000347 | 0.008288 | 0.116865 | Hexosyltransferase | elongation zone | 12 h |
| HORVU.MOREX.r3.3HG0278170.1 | 1.706706 | 0.346839 | 3.840083 | 0.00033 | 0.008104 | 0.116858 | CCCH-type zinc fingerfamily protein with RNA-binding domain-containing protein | elongation zone | 12 h |
| HORVU.MOREX.r3.5HG0510590.4 | -1.47342 | 4.084011 | -3.8693 | 0.000301 | 0.007754 | 0.11046 | Beta-glucosidase | elongation zone | 12 h |
| HORVU.MOREX.r3.7HG0686000.1 | -0.65705 | 5.976004 | -3.87175 | 0.000298 | 0.007733 | 0.106776 | Biotin synthase | elongation zone | 12 h |
| HORVU.MOREX.r3.4HG0377020.1 | -0.35522 | 6.726202 | -3.93658 | 0.000242 | 0.006978 | 0.106333 | Hexosyltransferase | elongation zone | 12 h |

**Table S1** Continued.

| ID | logFC | AveExpr | t | P.Value | adj.P.Val | B | gene description | root zone | time point |
|---|---|---|---|---|---|---|---|---|---|
| HORVU.MOREX.r3.7HG0656750.1 | -0.78725 | 3.218229 | -3.81435 | 0.000358 | 0.008428 | 0.105458 | Exocyst complex component, putative | elongation zone | 12 h |
| HORVU.MOREX.r3.3HG0231780.1 | -1.06119 | 5.499284 | -3.82742 | 0.000344 | 0.008287 | 0.103727 | Charged multivesicular body protein 5 | elongation zone | 12 h |
| HORVU.MOREX.r3.2HG0140970.1 | -0.85919 | 2.378717 | -3.82426 | 0.000347 | 0.008288 | 0.102609 | Heat-shock protein, putative | elongation zone | 12 h |
| HORVU.MOREX.r3.3HG0244010.1 | -0.66104 | 4.561972 | -3.81872 | 0.000353 | 0.008389 | 0.101781 | Kinase superfamily protein | elongation zone | 12 h |
| HORVU.MOREX.r3.2HG0212860.1 | 1.760564 | -1.30303 | 3.834701 | 0.000336 | 0.008176 | 0.094258 | #N/A | elongation zone | 12 h |
| HORVU.MOREX.r3.3HG0242870.1 | -0.32352 | 7.618528 | -3.94909 | 0.000233 | 0.006891 | 0.09409 | Carboxypeptidase | elongation zone | 12 h |
| HORVU.MOREX.r3.3HG0298390.1 | -1.15585 | 3.722188 | -3.81008 | 0.000363 | 0.008498 | 0.092909 | Adenine nucleotide alpha hydrolases-like superfamily protein | elongation zone | 12 h |
| HORVU.MOREX.r3.1HG0054510.1 | 1.113899 | 2.568148 | 3.81085 | 0.000362 | 0.008498 | 0.085359 | Protein PLASTID MOVEMENT IMPAIRED 1 | elongation zone | 12 h |
| HORVU.MOREX.r3.2HG0118580.1 | -1.06141 | 4.186861 | -3.81487 | 0.000358 | 0.008427 | 0.084972 | Glycosyltransferase | elongation zone | 12 h |
| HORVU.MOREX.r3.2HG0182210.2 | 0.863928 | 0.342142 | 3.869403 | 0.000301 | 0.007754 | 0.084239 | Transporter-related family protein | elongation zone | 12 h |
| HORVU.MOREX.r3.4HG0402600.2 | -1.22584 | 2.451456 | -3.87387 | 0.000296 | 0.0077 | 0.083807 | 5-dehydro-2-deoxygluconokinase | elongation zone | 12 h |
| HORVU.MOREX.r3.2HG0197680.1 | -0.39683 | 7.691665 | -3.91983 | 0.000256 | 0.007125 | 0.083345 | actin cytoskeleton-regulatory complex pan-like protein | elongation zone | 12 h |
| HORVU.MOREX.r3.2HG0124450.1 | -0.58966 | 4.909528 | -3.86569 | 0.000304 | 0.007757 | 0.081338 | calmodulin-binding protein (DUF1645) | elongation zone | 12 h |
| HORVU.MOREX.r3.6HG0597640.1 | 0.766519 | 3.081429 | 3.823937 | 0.000348 | 0.008288 | 0.079915 | Nitrilase | elongation zone | 12 h |
| HORVU.MOREX.r3.5HG0420970.1 | 0.891505 | 2.170731 | 3.817384 | 0.000355 | 0.008389 | 0.074953 | ERD (Early-responsive to dehydration stress) family protein | elongation zone | 12 h |
| HORVU.MOREX.r3.5HG0514110.1 | -0.6477 | 6.24431 | -3.86686 | 0.000303 | 0.007757 | 0.071163 | #N/A | elongation zone | 12 h |
| HORVU.MOREX.r3.1HG0050440.1 | 1.805239 | -1.13792 | 3.838733 | 0.000332 | 0.00812 | 0.071021 | DUF538 family protein (Protein of unknown function, DUF538) | elongation zone | 12 h |
| HORVU.MOREX.r3.2HG0180090.1 | -0.59663 | 3.817498 | -3.80935 | 0.000364 | 0.008498 | 0.070296 | F-box protein | elongation zone | 12 h |
| HORVU.MOREX.r3.2HG0098090.1 | -1.23275 | -2.03609 | -3.86936 | 0.000301 | 0.007754 | 0.070028 | Translation initiation factor IF-2 | elongation zone | 12 h |
| HORVU.MOREX.r3.6HG0577040.1 | 2.017784 | -1.97815 | 3.9414 | 0.000239 | 0.006937 | 0.069154 | Ycf1 | elongation zone | 12 h |
| HORVU.MOREX.r3.6HG0581020.1 | -0.38317 | 5.648771 | -3.87321 | 0.000297 | 0.007707 | 0.063591 | Xyloglucan 6-xylosyltransferase 1 | elongation zone | 12 h |
| HORVU.MOREX.r3.1HG0047130.1 | -0.50123 | 5.781989 | -3.85025 | 0.00032 | 0.007921 | 0.060804 | Pre-mRNA splicing factor-related | elongation zone | 12 h |
| HORVU.MOREX.r3.4HG0353390.1 | 0.849076 | 1.790635 | 3.815567 | 0.000357 | 0.008418 | 0.059068 | Type I inositol-1,4,5-trisphosphate 5-phosphatase | elongation zone | 12 h |
| HORVU.MOREX.r3.6HG0543250.1 | -0.79371 | 6.018336 | -3.96496 | 0.000221 | 0.006756 | 0.05802 | Receptor-like protein kinase | elongation zone | 12 h |
| HORVU.MOREX.r3.4HG0407990.1 | 0.870734 | 3.438278 | 3.867611 | 0.000302 | 0.007757 | 0.056614 | C2 calcium/lipid-binding and GRAM domain protein | elongation zone | 12 h |
| HORVU.MOREX.r3.5HG0467870.1 | 0.993588 | 2.049337 | 3.804563 | 0.00037 | 0.00855 | 0.055267 | Kynurenine formamidase | elongation zone | 12 h |
| HORVU.MOREX.r3.2HG0144760.1 | 1.107751 | -1.38996 | 3.793804 | 0.000382 | 0.008703 | 0.053624 | tRNA 5-methylaminomethyl-2-thiouridine biosynthesis bifunctional protein MnmC | elongation zone | 12 h |
| HORVU.MOREX.r3.3HG0310110.1 | 1.750342 | 0.692973 | 3.800338 | 0.000375 | 0.008587 | 0.049864 | lysine ketoglutarate reductase trans-splicing-like protein, putative (DUF707) | elongation zone | 12 h |
| HORVU.MOREX.r3.6HG0560290.1 | -2.45486 | -0.18724 | -4.29478 | 7.49E-05 | 0.00421 | 0.049857 | Cyclin | elongation zone | 12 h |
| HORVU.MOREX.r3.2HG0159540.1 | -1.01294 | 3.353367 | -3.79365 | 0.000383 | 0.008703 | 0.048468 | Glycosyltransferase | elongation zone | 12 h |
| HORVU.MOREX.r3.6HG0603410.1 | -0.76039 | 6.46923 | -3.85117 | 0.000319 | 0.007921 | 0.046696 | Homeobox-leucine zipper protein | elongation zone | 12 h |
| HORVU.MOREX.r3.4HG0382760.1 | -0.93305 | 4.231724 | -3.80304 | 0.000371 | 0.008553 | 0.045854 | CoA ligase | elongation zone | 12 h |
| HORVU.MOREX.r3.2HG0192410.1 | 1.68137 | 0.742041 | 3.865903 | 0.000304 | 0.007757 | 0.042609 | Peroxidase | elongation zone | 12 h |
| HORVU.MOREX.r3.2HG0158620.1 | -0.42124 | 4.942048 | -3.84544 | 0.000325 | 0.007986 | 0.04228 | Hydroxyproline-rich glycoprotein-like | elongation zone | 12 h |
| HORVU.MOREX.r3.1HG0090170.1 | -0.56275 | 5.133692 | -3.82451 | 0.000347 | 0.008288 | 0.039166 | NBS-LRR-like resistance protein | elongation zone | 12 h |
| HORVU.MOREX.r3.1HG0054010.1 | 0.633748 | 5.841657 | 3.91165 | 0.000263 | 0.007187 | 0.037799 | zinc finger B-box protein | elongation zone | 12 h |
| HORVU.MOREX.r3.2HG0106260.1 | 2.573854 | -2.23574 | 4.093316 | 0.000146 | 0.005731 | 0.037307 | F-box family protein | elongation zone | 12 h |
| HORVU.MOREX.r3.1HG0085700.1 | -0.96337 | 4.979026 | -3.7999 | 0.000375 | 0.008587 | 0.036446 | Sodium Bile acid symporter family | elongation zone | 12 h |
| HORVU.MOREX.r3.1HG0072510.1 | 1.259198 | -0.03799 | 3.790654 | 0.000386 | 0.008739 | 0.034658 | Protein-methionine-s-oxide reductase | elongation zone | 12 h |
| HORVU.MOREX.r3.3HG0283990.1 | -0.52523 | 7.009142 | -3.88298 | 0.000288 | 0.007575 | 0.030692 | Glycylpeptide N-tetradecanoyltransferase | elongation zone | 12 h |
| HORVU.MOREX.r3.1HG0046180.1 | 0.492342 | 5.918665 | 3.900792 | 0.000272 | 0.007363 | 0.029938 | Ser/Thr protein phosphatase family protein, expressed | elongation zone | 12 h |

**Table S1** Continued.

| ID | logFC | AveExpr | t | P.Value | adj.P.Val | B | gene description | root zone | time point |
|---|---|---|---|---|---|---|---|---|---|
| HORVU.MOREX.r3.1HG0024140.1 | -1.02861 | 6.990532 | -3.8978 | 0.000275 | 0.007375 | 0.02989 | Cellulose synthase-like protein | elongation zone | 12 h |
| HORVU.MOREX.r3.5HG0477220.1 | 1.053019 | 1.858254 | 3.785855 | 0.000392 | 0.008794 | 0.029522 | F-box protein | elongation zone | 12 h |
| HORVU.MOREX.r3.7HG0739460.1 | -1.04859 | 6.558798 | -3.8487 | 0.000321 | 0.007942 | 0.025994 | Histone H2B | elongation zone | 12 h |
| HORVU.MOREX.r3.5HG0509390.1 | 1.095672 | 0.147864 | 3.81986 | 0.000352 | 0.008372 | 0.023972 | LEM3 (Ligand-effect modulator 3)-like | elongation zone | 12 h |
| HORVU.MOREX.r3.6HG0616130.1 | 0.588645 | 6.04228 | 3.933843 | 0.000245 | 0.007006 | 0.02372 | Hexosyltransferase | elongation zone | 12 h |
| HORVU.MOREX.r3.7HG0673610.1 | -0.55871 | 3.691119 | -3.79665 | 0.000379 | 0.00865 | 0.022371 | CONSTANS-like zinc finger protein | elongation zone | 12 h |
| HORVU.MOREX.r3.2HG0199520.1 | -1.66919 | 1.070689 | -3.98346 | 0.000208 | 0.006588 | 0.022084 | mechanosensitive channel of small conductance-like 5 | elongation zone | 12 h |
| HORVU.MOREX.r3.6HG0606930.1 | -0.40773 | 5.680326 | -3.85348 | 0.000316 | 0.007898 | 0.021112 | 3-ketodihydrosphingosine reductase | elongation zone | 12 h |
| HORVU.MOREX.r3.1HG0038960.1 | -1.03759 | 2.949857 | -3.7923 | 0.000384 | 0.008721 | 0.020549 | #N/A | elongation zone | 12 h |
| HORVU.MOREX.r3.5HG0501980.1 | -0.44855 | 5.561069 | -3.84225 | 0.000328 | 0.008058 | 0.01989 | Rop guanine nucleotide exchange factor, putative | elongation zone | 12 h |
| HORVU.MOREX.r3.3HG0283740.1 | 0.545755 | 5.066973 | 3.850321 | 0.00032 | 0.007921 | 0.017527 | Anoctamin-like protein | elongation zone | 12 h |
| HORVU.MOREX.r3.1HG0066530.1 | -1.19262 | 4.906097 | -3.78038 | 0.000399 | 0.008889 | 0.014176 | DUF630 family protein, putative (DUF630 and DUF632) | elongation zone | 12 h |
| HORVU.MOREX.r3.5HG0513980.1 | 1.341517 | -0.36473 | 3.778592 | 0.000401 | 0.00891 | 0.011307 | Auxin-responsive protein | elongation zone | 12 h |
| HORVU.MOREX.r3.5HG0531070.1 | 3.684604 | -3.02117 | 4.284451 | 7.75E-05 | 0.004293 | 0.009433 | Protein DETOXIFICATION | elongation zone | 12 h |
| HORVU.MOREX.r3.5HG0461170.1 | -0.68308 | 4.201237 | -3.7825 | 0.000396 | 0.008859 | 0.00682 | Protein FAR1-RELATED SEQUENCE 5 | elongation zone | 12 h |
| HORVU.MOREX.r3.3HG0266960.1 | -0.6095 | 4.636036 | -3.80266 | 0.000372 | 0.008553 | 0.003626 | #N/A | elongation zone | 12 h |
| HORVU.MOREX.r3.2HG0204020.1 | 0.741716 | 1.435954 | 3.833165 | 0.000338 | 0.008177 | 0.001965 | Sulfate transporter | elongation zone | 12 h |
| HORVU.MOREX.r3.2HG0107740.1 | -0.56434 | 9.485909 | -3.95132 | 0.000231 | 0.006891 | -0.00255 | Ascorbate peroxidase | elongation zone | 12 h |
| HORVU.MOREX.r3.5HG0533430.1 | 1.330724 | 0.093461 | 3.792933 | 0.000383 | 0.008713 | -0.0028 | ROP-interactive CRIB motif protein | elongation zone | 12 h |
| HORVU.MOREX.r3.5HG0533630.1 | -1.44445 | 5.193358 | -3.84687 | 0.000323 | 0.007978 | -0.00521 | Expansin | elongation zone | 12 h |
| HORVU.MOREX.r3.5HG0421310.1 | 1.073256 | 0.680461 | 3.818482 | 0.000354 | 0.008389 | -0.00908 | Receptor protein kinase, putative | elongation zone | 12 h |
| HORVU.MOREX.r3.3HG0250630.1 | -0.69859 | 3.543167 | -3.77211 | 0.00041 | 0.008991 | -0.01311 | tRNA(His) guanylyltransferase | elongation zone | 12 h |
| HORVU.MOREX.r3.7HG0688870.1 | 0.748556 | 6.011072 | 3.881764 | 0.000289 | 0.007575 | -0.01714 | Monodehydroascorbate reductase | elongation zone | 12 h |
| HORVU.MOREX.r3.4HG0339750.1 | -0.61166 | 7.537128 | -3.88796 | 0.000283 | 0.007523 | -0.02015 | Cathepsin B-like cysteine protease | elongation zone | 12 h |
| HORVU.MOREX.r3.3HG0289970.1 | -0.60452 | 4.641812 | -3.78611 | 0.000392 | 0.008794 | -0.02199 | Riboflavin kinase/fmn hydrolase | elongation zone | 12 h |
| HORVU.MOREX.r3.2HG0110060.1 | -0.57746 | 6.526409 | -3.85064 | 0.000319 | 0.007921 | -0.02366 | Aldehyde dehydrogenase | elongation zone | 12 h |
| HORVU.MOREX.r3.6HG0555500.1 | -0.61887 | 5.078092 | -3.83348 | 0.000337 | 0.008177 | -0.03804 | Subtilisin-like protease | elongation zone | 12 h |
| HORVU.MOREX.r3.6HG0550940.1 | -0.33463 | 5.078026 | -3.81411 | 0.000359 | 0.008428 | -0.0383 | obscurin-like protein | elongation zone | 12 h |
| HORVU.MOREX.r3.1HG0045770.1 | 2.120943 | -2.26851 | 3.941014 | 0.000239 | 0.006937 | -0.04256 | GRAM domain-containing protein / ABA-responsive protein-related | elongation zone | 12 h |
| HORVU.MOREX.r3.5HG0517760.1 | 1.696398 | -2.35783 | 3.809692 | 0.000364 | 0.008498 | -0.04465 | Zinc finger A20 and AN1 domain-containing stress-associated protein | elongation zone | 12 h |
| HORVU.MOREX.r3.7HG0661890.1 | -0.57398 | 7.916049 | -3.89911 | 0.000273 | 0.007364 | -0.04657 | 3-oxoacyl-[acyl-carrier-protein] synthase | elongation zone | 12 h |
| HORVU.MOREX.r3.1HG0081620.1 | -0.65092 | 3.155534 | -3.76157 | 0.000423 | 0.009125 | -0.04755 | Glycerophosphodiester phosphodiesterase | elongation zone | 12 h |
| HORVU.MOREX.r3.3HG0236000.1 | -0.44653 | 6.001989 | -3.8213 | 0.00035 | 0.008343 | -0.0506 | Ectonucleotide pyrophosphatase/phosphodiesterase, putative | elongation zone | 12 h |
| HORVU.MOREX.r3.3HG0231880.1 | -1.08129 | 2.985187 | -3.77529 | 0.000405 | 0.008965 | -0.05068 | Leucine-rich repeat (LRR) family protein | elongation zone | 12 h |
| HORVU.MOREX.r3.6HG0617970.1 | 1.5786 | -1.86338 | 3.765043 | 0.000419 | 0.009082 | -0.05076 | Glycosyltransferase family 92 protein | elongation zone | 12 h |
| HORVU.MOREX.r3.1HG0052080.1 | -0.45021 | 6.314334 | -3.83259 | 0.000338 | 0.008182 | -0.05111 | C-terminal binding protein | elongation zone | 12 h |
| HORVU.MOREX.r3.6HG0633160.1 | -0.77726 | 3.333953 | -3.75595 | 0.000431 | 0.00924 | -0.05698 | U-box domain-containing protein 62 | elongation zone | 12 h |
| HORVU.MOREX.r3.5HG0485760.1 | 1.663329 | -1.3336 | 3.753877 | 0.000434 | 0.009279 | -0.05826 | 65-kDa microtubule-associated-like protein | elongation zone | 12 h |
| HORVU.MOREX.r3.2HG0128230.1 | -0.94154 | 5.522587 | -3.78823 | 0.000389 | 0.008785 | -0.05836 | Thaumatin | elongation zone | 12 h |
| HORVU.MOREX.r3.1HG0073050.1 | -1.46272 | 3.873682 | -3.75239 | 0.000436 | 0.009279 | -0.06085 | Histone H4 | elongation zone | 12 h |
| HORVU.MOREX.r3.7HG0636660.1 | -0.40995 | 6.587599 | -3.86332 | 0.000307 | 0.007759 | -0.06167 | SKP1-like protein | elongation zone | 12 h |

**Table S1** Continued.

| ID | logFC | AveExpr | t | P.Value | adj.P.Val | B | gene description | root zone | time point |
|---|---|---|---|---|---|---|---|---|---|
| HORVU.MOREX.r3.2HG0148170.1 | -1.31696 | 0.675747 | -3.86147 | 0.000308 | 0.007759 | -0.06802 | transmembrane protein | elongation zone | 12 h |
| HORVU.MOREX.r3.3HG0307240.1 | 2.701127 | -3.37688 | 4.083101 | 0.000151 | 0.005814 | -0.07366 | Amino acid permease | elongation zone | 12 h |
| HORVU.MOREX.r3.3HG0231050.1 | -0.88093 | 7.166338 | -3.8456 | 0.000324 | 0.007986 | -0.07371 | Histone H2B | elongation zone | 12 h |
| HORVU.MOREX.r3.4HG0412370.1 | 2.027924 | -1.42512 | 3.74849 | 0.000441 | 0.009306 | -0.07444 | Homeobox protein knotted-1, putative | elongation zone | 12 h |
| HORVU.MOREX.r3.2HG0122510.1 | 0.706896 | 2.203854 | 3.829761 | 0.000341 | 0.008246 | -0.07524 | Sphingoid base hydroxylase 2 | elongation zone | 12 h |
| HORVU.MOREX.r3.7HG0718540.1 | 0.78858 | 3.335782 | 3.752632 | 0.000435 | 0.009279 | -0.07805 | Heat shock transcription factor | elongation zone | 12 h |
| HORVU.MOREX.r3.2HG0168010.1 | -0.55596 | 4.55186 | -3.77198 | 0.00041 | 0.008991 | -0.08103 | Aldo-keto reductase | elongation zone | 12 h |
| HORVU.MOREX.r3.3HG0276160.2 | 1.557918 | -3.02913 | 3.948098 | 0.000234 | 0.006891 | -0.08266 | GDSL esterase/lipase | elongation zone | 12 h |
| HORVU.MOREX.r3.5HG0420980.1 | 0.791941 | 2.046048 | 3.750927 | 0.000438 | 0.009279 | -0.0855 | ERD (Early-responsive to dehydration stress) family protein | elongation zone | 12 h |
| HORVU.MOREX.r3.1HG0061390.1 | 0.721862 | 1.721091 | 3.866324 | 0.000304 | 0.007757 | -0.08796 | Trichome birefringence-like protein | elongation zone | 12 h |
| HORVU.MOREX.r3.1HG0065370.1 | -0.77917 | 5.483156 | -3.78667 | 0.000391 | 0.008794 | -0.08809 | TPX2 (Targeting protein for Xklp2) family protein | elongation zone | 12 h |
| HORVU.MOREX.r3.7HG0659040.1 | 0.960934 | 3.074402 | 3.741749 | 0.000451 | 0.009442 | -0.09014 | Formin-like protein | elongation zone | 12 h |
| HORVU.MOREX.r3.5HG0466190.1 | -0.70948 | 5.829313 | -3.83487 | 0.000336 | 0.008176 | -0.0939 | Protein transport protein Sec61 subunit beta, putative | elongation zone | 12 h |
| HORVU.MOREX.r3.2HG0157450.1 | 1.025741 | 2.551061 | 3.886335 | 0.000285 | 0.007552 | -0.09658 | Peroxidase | elongation zone | 12 h |
| HORVU.MOREX.r3.2HG0124130.1 | -0.72369 | 4.407551 | -3.76214 | 0.000423 | 0.009118 | -0.0974 | Cyclin family protein | elongation zone | 12 h |
| HORVU.MOREX.r3.1HG0000050.1 | -0.44534 | 5.048249 | -3.79621 | 0.000379 | 0.008652 | -0.10018 | RING-finger ubiquitin ligase | elongation zone | 12 h |
| HORVU.MOREX.r3.6HG0608180.1 | 1.109853 | 0.307405 | 3.73753 | 0.000457 | 0.009532 | -0.10373 | Protein MIZU-KUSSEI 1 | elongation zone | 12 h |
| HORVU.MOREX.r3.1HG0065020.1 | -1.1703 | 5.184531 | -3.74213 | 0.00045 | 0.009442 | -0.10509 | BZIP transcription factor | elongation zone | 12 h |
| HORVU.MOREX.r3.2HG0185370.1 | 2.719499 | -1.4233 | 3.968401 | 0.000219 | 0.006702 | -0.10599 | Subtilisin-like protease | elongation zone | 12 h |
| HORVU.MOREX.r3.2HG0205420.1 | 0.822282 | -0.18096 | 3.762954 | 0.000422 | 0.009111 | -0.10643 | #N/A | elongation zone | 12 h |
| HORVU.MOREX.r3.2HG0097010.1 | 0.397998 | 5.998881 | 3.849915 | 0.00032 | 0.007921 | -0.10649 | Sterol 3-beta-glucosyltransferase | elongation zone | 12 h |
| HORVU.MOREX.r3.7HG0677940.1 | -0.35754 | 7.210054 | -3.86706 | 0.000303 | 0.007757 | -0.10717 | Transmembrane 9 superfamily member | elongation zone | 12 h |
| HORVU.MOREX.r3.2HG0168330.1 | 0.509769 | 5.088967 | 3.827093 | 0.000344 | 0.008287 | -0.10947 | Nicotinate phosphoribosyltransferase | elongation zone | 12 h |
| HORVU.MOREX.r3.4HG0415600.1 | -1.68475 | 5.888147 | -3.75847 | 0.000428 | 0.009191 | -0.11147 | HXXXD-type acyl-transferase family protein | elongation zone | 12 h |
| HORVU.MOREX.r3.5HG0484660.1 | 1.491417 | -0.23036 | 3.732868 | 0.000463 | 0.009604 | -0.11754 | Vacuolar-processing enzyme | elongation zone | 12 h |
| HORVU.MOREX.r3.2HG0181660.1 | -0.83461 | 5.953972 | -3.78834 | 0.000389 | 0.008785 | -0.12216 | 3-hydroxyacyl-[acyl-carrier-protein] dehydratase FabZ | elongation zone | 12 h |
| HORVU.MOREX.r3.1HG0024900.1 | 0.348583 | 5.998911 | 3.839151 | 0.000331 | 0.008118 | -0.12418 | Macrophage erythroblast attacher | elongation zone | 12 h |
| HORVU.MOREX.r3.2HG0214240.1 | -0.63166 | 3.183258 | -3.73123 | 0.000466 | 0.009644 | -0.12751 | Disease resistance protein (NBS-LRR class) family | elongation zone | 12 h |
| HORVU.MOREX.r3.5HG0469340.1 | 1.288979 | 0.552227 | 3.736539 | 0.000458 | 0.009546 | -0.12937 | Fasciclin-like arabinogalactan-protein-like | elongation zone | 12 h |
| HORVU.MOREX.r3.1HG0072110.1 | -0.814 | 5.194691 | -3.7796 | 0.0004 | 0.008901 | -0.131 | #N/A | elongation zone | 12 h |
| HORVU.MOREX.r3.6HG0620650.1 | -0.57696 | 7.259903 | -3.86063 | 0.000309 | 0.007759 | -0.13238 | Subtilisin protease | elongation zone | 12 h |
| HORVU.MOREX.r3.7HG0684180.1 | 0.613883 | 6.139387 | 3.866545 | 0.000303 | 0.007757 | -0.1337 | Peroxidase | elongation zone | 12 h |
| HORVU.MOREX.r3.6HG0549810.1 | 0.522353 | 6.037644 | 3.856865 | 0.000313 | 0.007832 | -0.13693 | Rac-like GTP binding protein | elongation zone | 12 h |
| HORVU.MOREX.r3.1HG0045490.1 | -0.4149 | 5.161254 | -3.77338 | 0.000408 | 0.008984 | -0.13865 | RING/U-box superfamily protein | elongation zone | 12 h |
| HORVU.MOREX.r3.5HG0484180.1 | 0.449292 | 5.636527 | 3.807438 | 0.000366 | 0.00853 | -0.13976 | DUF868 family protein | elongation zone | 12 h |
| HORVU.MOREX.r3.4HG0400040.1 | 0.603727 | 1.083906 | 3.804939 | 0.000369 | 0.00855 | -0.14096 | Cytochrome b561 and DOMON domain-containing protein | elongation zone | 12 h |
| HORVU.MOREX.r3.1HG0051920.1 | 0.939664 | 2.286248 | 3.78247 | 0.000396 | 0.008859 | -0.14117 | Transcription factor | elongation zone | 12 h |
| HORVU.MOREX.r3.4HG0418640.1 | -0.69142 | 4.260686 | -3.72788 | 0.000471 | 0.009706 | -0.14227 | C2 domain-containing protein | elongation zone | 12 h |
| HORVU.MOREX.r3.2HG0182280.1 | -0.72761 | 6.433901 | -3.80187 | 0.000373 | 0.008557 | -0.14387 | Auxin response factor | elongation zone | 12 h |
| HORVU.MOREX.r3.7HG0654510.1 | -1.34959 | 5.364113 | -3.7472 | 0.000443 | 0.009332 | -0.14645 | Histone H3 | elongation zone | 12 h |
| HORVU.MOREX.r3.5HG0537090.1 | -0.34553 | 6.791678 | -3.83786 | 0.000333 | 0.008132 | -0.14818 | WD40 repeat-containing protein | elongation zone | 12 h |

**Table S1** Continued.

| ID | logFC | AveExpr | t | P.Value | adj.P.Val | B | gene description | root zone | time point |
|---|---|---|---|---|---|---|---|---|---|
| HORVU.MOREX.r3.7HG0718830.1 | 0.846116 | 4.155941 | 3.86129 | 0.000309 | 0.007759 | -0.14822 | FAD-binding Berberine family protein, putative | elongation zone | 12 h |
| HORVU.MOREX.r3.5HG0526250.1 | -0.32537 | 12.03818 | -3.99139 | 0.000203 | 0.006534 | -0.14836 | 70 kDa heat shock protein | elongation zone | 12 h |
| HORVU.MOREX.r3.5HG0534340.1 | -1.05251 | 3.235501 | -3.72043 | 0.000482 | 0.009857 | -0.15343 | Transferase, transferring glycosyl groups | elongation zone | 12 h |
| HORVU.MOREX.r3.7HG0720130.1 | -0.59159 | 6.1996 | -3.80356 | 0.000371 | 0.008553 | -0.1553 | Phosphatidylcholine:diacylglycerol cholinephosphotransferase 1 | elongation zone | 12 h |
| HORVU.MOREX.r3.2HG0168730.1 | -0.52815 | 7.047952 | -3.80181 | 0.000373 | 0.008557 | -0.15926 | Kinase family protein | elongation zone | 12 h |
| HORVU.MOREX.r3.6HG0620720.1 | 0.955463 | 0.656099 | 3.799664 | 0.000375 | 0.008587 | -0.16162 | Ribonuclease | elongation zone | 12 h |
| HORVU.MOREX.r3.4HG0351440.1 | 0.460798 | 5.31189 | 3.816949 | 0.000355 | 0.008391 | -0.16217 | Rho GTPase-activating protein | elongation zone | 12 h |
| HORVU.MOREX.r3.4HG0376720.1 | 2.944864 | -0.40522 | 4.559089 | 3.07E-05 | 0.002847 | -0.16243 | #N/A | elongation zone | 12 h |
| HORVU.MOREX.r3.5HG0536200.1 | 1.076437 | -0.5546 | 3.769039 | 0.000413 | 0.009031 | -0.16782 | Xyloglucan endotransglucosylase/hydrolase | elongation zone | 12 h |
| HORVU.MOREX.r3.1HG0067410.1 | -0.41497 | 5.646383 | -3.77791 | 0.000402 | 0.00892 | -0.17055 | 1-acyl-sn-glycerol-3-phosphate acyltransferase | elongation zone | 12 h |
| HORVU.MOREX.r3.3HG0235350.1 | 2.006446 | 0.004514 | 3.733848 | 0.000462 | 0.009584 | -0.17067 | Pollen Ole e 1 allergen and extensin family protein | elongation zone | 12 h |
| HORVU.MOREX.r3.3HG0288420.1 | -0.92497 | 7.285552 | -3.78607 | 0.000392 | 0.008794 | -0.17552 | Mannan endo-1,4-beta-mannosidase-like protein | elongation zone | 12 h |
| HORVU.MOREX.r3.2HG0184620.1 | -1.07628 | 4.117573 | -3.71607 | 0.000488 | 0.009916 | -0.18349 | Hsp70-Hsp90 organizing protein 1 | elongation zone | 12 h |
| HORVU.MOREX.r3.7HG0679060.1 | 0.797325 | 1.156649 | 3.728877 | 0.000469 | 0.009696 | -0.18381 | Nuclear transport factor 2 family protein, putative | elongation zone | 12 h |
| HORVU.MOREX.r3.3HG0221240.1 | 0.781353 | 1.030061 | 3.791279 | 0.000385 | 0.008739 | -0.1908 | Glycosyltransferase | elongation zone | 12 h |
| HORVU.MOREX.r3.3HG0285780.1 | 1.031952 | 1.233022 | 3.803871 | 0.00037 | 0.008553 | -0.19082 | Protein kinase, putative | elongation zone | 12 h |
| HORVU.MOREX.r3.4HG0342080.1 | 0.765648 | 4.067325 | 3.778952 | 0.000401 | 0.00891 | -0.1911 | Adenine/guanine permease | elongation zone | 12 h |
| HORVU.MOREX.r3.6HG0543730.1 | -0.85299 | 7.651801 | -3.81804 | 0.000354 | 0.008389 | -0.19365 | Histone H2B | elongation zone | 12 h |
| HORVU.MOREX.r3.1HG0072530.1 | -1.14492 | 4.730817 | -3.72543 | 0.000474 | 0.009762 | -0.19607 | Histone H4 | elongation zone | 12 h |
| HORVU.MOREX.r3.6HG0616480.1 | -0.60564 | 5.832986 | -3.76911 | 0.000413 | 0.009031 | -0.19632 | #N/A | elongation zone | 12 h |
| HORVU.MOREX.r3.1HG0094110.1 | 2.009173 | 0.561027 | 3.703053 | 0.000509 | 0.010183 | -0.19812 | Sigma factor sigB regulation protein rsbQ | elongation zone | 12 h |
| HORVU.MOREX.r3.5HG0420830.1 | 1.288318 | -1.8789 | 3.717478 | 0.000486 | 0.009899 | -0.20402 | cycling DOF factor 1 | elongation zone | 12 h |
| HORVU.MOREX.r3.2HG0192910.1 | -0.7466 | 6.73787 | -3.82888 | 0.000342 | 0.00826 | -0.20522 | Xyloglucan endotransglucosylase/hydrolase | elongation zone | 12 h |
| HORVU.MOREX.r3.1HG0077950.1 | -0.3894 | 4.840037 | -3.75103 | 0.000438 | 0.009279 | -0.21039 | BSD domain containing protein | elongation zone | 12 h |
| HORVU.MOREX.r3.2HG0168270.1 | -0.61213 | 3.16772 | -3.69799 | 0.000517 | 0.010257 | -0.21314 | Ubiquitin carboxyl-terminal hydrolase | elongation zone | 12 h |
| HORVU.MOREX.r3.7HG0685920.1 | 0.53821 | 4.571683 | 3.787257 | 0.00039 | 0.008794 | -0.21325 | Solute carrier family 35 protein | elongation zone | 12 h |
| HORVU.MOREX.r3.1HG0079020.1 | 2.039497 | 3.471696 | 3.699655 | 0.000514 | 0.010233 | -0.21388 | Protein kinase, putative | elongation zone | 12 h |
| HORVU.MOREX.r3.5HG0510160.1 | -0.50168 | 4.316907 | -3.71785 | 0.000486 | 0.009897 | -0.2143 | Myb/SANT-like DNA-binding domain protein | elongation zone | 12 h |
| HORVU.MOREX.r3.1HG0010470.1 | -0.58354 | 5.658792 | -3.75063 | 0.000438 | 0.009279 | -0.21643 | Arginine/serine-rich splicing factor, putative | elongation zone | 12 h |
| HORVU.MOREX.r3.6HG0560930.1 | 0.640376 | 5.592543 | 3.808424 | 0.000365 | 0.008513 | -0.21881 | ternary complex factor MIP1 leucine-zipper protein (Protein of unknown function, | elongation zone | 12 h |
| HORVU.MOREX.r3.3HG0241630.1 | -0.65151 | 5.456663 | -3.80494 | 0.000369 | 0.00855 | -0.21969 | Embryo-specific 3 | elongation zone | 12 h |
| HORVU.MOREX.r3.2HG0190620.1 | 0.959315 | 0.132535 | 3.700251 | 0.000513 | 0.010224 | -0.21982 | Ceramide kinase | elongation zone | 12 h |
| HORVU.MOREX.r3.4HG0381790.1 | 1.08853 | 3.851367 | 3.790603 | 0.000386 | 0.008739 | -0.22082 | transmembrane protein, putative (DUF247) | elongation zone | 12 h |
| HORVU.MOREX.r3.7HG0728820.1 | 1.427083 | 1.672182 | 3.725005 | 0.000475 | 0.009765 | -0.22165 | Xyloglucan endotransglucosylase/hydrolase | elongation zone | 12 h |
| HORVU.MOREX.r3.3HG0318400.1 | -0.96279 | 6.359366 | -3.74014 | 0.000453 | 0.009464 | -0.22238 | S-formylglutathione hydrolase | elongation zone | 12 h |
| HORVU.MOREX.r3.1HG0018590.1 | 0.841734 | 2.172649 | 3.804832 | 0.000369 | 0.00855 | -0.22336 | MYB transcription factor | elongation zone | 12 h |
| HORVU.MOREX.r3.5HG0422080.1 | 0.426725 | 5.370925 | 3.780828 | 0.000398 | 0.008886 | -0.22584 | RING/U-box superfamily protein | elongation zone | 12 h |
| HORVU.MOREX.r3.5HG0501120.1 | -0.32218 | 6.923122 | -3.80519 | 0.000369 | 0.00855 | -0.22891 | 26S proteasome regulatory subunit | elongation zone | 12 h |
| HORVU.MOREX.r3.5HG0493850.1 | -0.52792 | 7.419096 | -3.81748 | 0.000355 | 0.008389 | -0.22991 | Acyl-[acyl-carrier-protein] hydrolase | elongation zone | 12 h |
| HORVU.MOREX.r3.4HG0396710.1 | -1.15706 | 6.334866 | -3.75229 | 0.000436 | 0.009279 | -0.23058 | Histone H3 | elongation zone | 12 h |
| HORVU.MOREX.r3.7HG0662210.1 | -0.57909 | 8.387754 | -3.82453 | 0.000347 | 0.008288 | -0.23366 | T-complex protein 1 subunit delta | elongation zone | 12 h |

**Table S1** Continued.

| ID | logFC | AveExpr | t | P.Value | adj.P.Val | B | gene description | root zone | time point |
|---|---|---|---|---|---|---|---|---|---|
| HORVU.MOREX.r3.3HG0291040.1 | 0.785987 | 3.480757 | 3.698682 | 0.000516 | 0.010244 | -0.23402 | Glutathione S-transferase | elongation zone | 12 h |
| HORVU.MOREX.r3.3HG0228940.1 | 1.119244 | 0.760498 | 3.708278 | 0.0005 | 0.010068 | -0.23468 | Sulfate adenylyltransferase subunit 2 | elongation zone | 12 h |
| HORVU.MOREX.r3.4HG0350800.1 | -2.538 | 1.29696 | -4.03893 | 0.000174 | 0.006138 | -0.23574 | FBD-associated F-box protein | elongation zone | 12 h |
| HORVU.MOREX.r3.4HG0398900.1 | -1.04009 | 8.042635 | -3.80941 | 0.000364 | 0.008498 | -0.23689 | Histone H2A | elongation zone | 12 h |
| HORVU.MOREX.r3.1HG0066800.1 | 1.176372 | -2.50287 | 3.714313 | 0.000491 | 0.009948 | -0.23997 | Potassium channel | elongation zone | 12 h |
| HORVU.MOREX.r3.5HG0512350.1 | -1.1281 | 7.007306 | -3.75658 | 0.00043 | 0.009231 | -0.24167 | Cytochrome P450 | elongation zone | 12 h |
| HORVU.MOREX.r3.7HG0636890.1 | 1.679045 | 0.940541 | 3.774152 | 0.000407 | 0.008984 | -0.24497 | Expansin-like protein | elongation zone | 12 h |
| HORVU.MOREX.r3.5HG0488950.1 | -0.74427 | 2.845933 | -3.69656 | 0.000519 | 0.010292 | -0.24581 | Cellulose synthase, putative | elongation zone | 12 h |
| HORVU.MOREX.r3.6HG0619500.1 | 1.490131 | -0.34034 | 3.684658 | 0.000539 | 0.01044 | -0.24618 | #N/A | elongation zone | 12 h |
| HORVU.MOREX.r3.1HG0091240.1 | 0.568875 | 4.910104 | 3.787215 | 0.00039 | 0.008794 | -0.24724 | Mitogen-activated protein kinase | elongation zone | 12 h |
| HORVU.MOREX.r3.3HG0275090.1 | 1.921845 | -3.21251 | 3.942268 | 0.000238 | 0.006937 | -0.24893 | SWI/SNF-related matrix-associated actin-dependent regulator of chromatin subfa | elongation zone | 12 h |
| HORVU.MOREX.r3.1HG0090760.1 | -0.60365 | 3.106359 | -3.68826 | 0.000533 | 0.010373 | -0.24914 | Nuclear transcription factor Y subunit B | elongation zone | 12 h |
| HORVU.MOREX.r3.2HG0187640.1 | -0.79541 | 2.595046 | -3.69064 | 0.000529 | 0.010355 | -0.24963 | Gamma-tubulin complex component | elongation zone | 12 h |
| HORVU.MOREX.r3.5HG0454860.1 | -0.41026 | 5.797971 | -3.76474 | 0.000419 | 0.009082 | -0.24986 | Trehalose-6-phosphate synthase, putative | elongation zone | 12 h |
| HORVU.MOREX.r3.2HG0107820.1 | -0.6777 | 4.968047 | -3.74132 | 0.000451 | 0.009442 | -0.25135 | Glycosyltransferases | elongation zone | 12 h |
| HORVU.MOREX.r3.2HG0099900.1 | 0.574116 | 5.512063 | 3.751124 | 0.000438 | 0.009279 | -0.2518 | Glyoxylate reductase/hydroxypyruvate reductase | elongation zone | 12 h |
| HORVU.MOREX.r3.2HG0100900.2 | 1.598476 | 0.62711 | 3.76838 | 0.000414 | 0.009031 | -0.25697 | Phosphoglycerate mutase-like protein | elongation zone | 12 h |
| HORVU.MOREX.r3.4HG0389200.1 | 0.819065 | 0.876157 | 3.683972 | 0.00054 | 0.010443 | -0.25872 | LIM domain-containing protein 1 | elongation zone | 12 h |
| HORVU.MOREX.r3.1HG0055400.1 | -0.50514 | 5.192784 | -3.71993 | 0.000483 | 0.009862 | -0.26034 | Gag polyprotein | elongation zone | 12 h |
| HORVU.MOREX.r3.4HG0401720.1 | -1.12859 | 7.267357 | -3.76644 | 0.000417 | 0.009058 | -0.26139 | Histone H3 | elongation zone | 12 h |
| HORVU.MOREX.r3.7HG0714510.1 | -1.15591 | 5.004338 | -3.68279 | 0.000542 | 0.010462 | -0.2656 | Protein phosphatase 2C | elongation zone | 12 h |
| HORVU.MOREX.r3.3HG0224680.1 | -0.41086 | 5.580543 | -3.75075 | 0.000438 | 0.009279 | -0.27335 | Ferredoxin reductase-like protein | elongation zone | 12 h |
| HORVU.MOREX.r3.1HG0046650.1 | -0.60389 | 6.618439 | -3.80584 | 0.000368 | 0.00855 | -0.2757 | Carboxypeptidase | elongation zone | 12 h |
| HORVU.MOREX.r3.7HG0708320.1 | -0.67687 | 4.603991 | -3.72159 | 0.00048 | 0.009831 | -0.27764 | Stromal cell-derived factor 2-like protein | elongation zone | 12 h |
| HORVU.MOREX.r3.4HG0358310.1 | -0.7158 | 4.422454 | -3.69606 | 0.00052 | 0.010292 | -0.2795 | Scarecrow transcription factor family protein | elongation zone | 12 h |
| HORVU.MOREX.r3.5HG0430940.1 | 1.096859 | 5.193365 | 3.781257 | 0.000398 | 0.008884 | -0.28226 | Glucan 1,3-beta-glucosidase | elongation zone | 12 h |
| HORVU.MOREX.r3.2HG0112190.1 | 1.553885 | -0.62536 | 3.686549 | 0.000536 | 0.010399 | -0.28382 | Transcription repressor ofp17 | elongation zone | 12 h |
| HORVU.MOREX.r3.2HG0189940.1 | -1.32041 | 5.951644 | -3.71266 | 0.000494 | 0.00998 | -0.28399 | #N/A | elongation zone | 12 h |
| HORVU.MOREX.r3.2HG0151910.1 | -1.0244 | 2.112464 | -3.67767 | 0.000551 | 0.010541 | -0.28525 | Retrovirus-related Pol polyprotein from transposon TNT 1-94 | elongation zone | 12 h |
| HORVU.MOREX.r3.2HG0199600.1 | -2.21726 | 2.139352 | -3.77558 | 0.000405 | 0.008965 | -0.28719 | Organic cation transporter protein | elongation zone | 12 h |
| HORVU.MOREX.r3.3HG0285840.1 | 0.989928 | -0.69484 | 3.681722 | 0.000544 | 0.010463 | -0.28777 | Senescence/dehydration-associated protein-like protein | elongation zone | 12 h |
| HORVU.MOREX.r3.2HG0190220.1 | -0.99801 | 6.32386 | -3.74121 | 0.000451 | 0.009442 | -0.288 | #N/A | elongation zone | 12 h |
| HORVU.MOREX.r3.1HG0089750.1 | -1.13685 | 2.726337 | -3.71159 | 0.000495 | 0.009994 | -0.28966 | DNA-3-methyladenine glycosylase 1 | elongation zone | 12 h |
| HORVU.MOREX.r3.3HG0324180.1 | -0.82122 | 4.528561 | -3.68775 | 0.000534 | 0.010379 | -0.29375 | #N/A | elongation zone | 12 h |
| HORVU.MOREX.r3.5HG0510320.1 | 0.722219 | 5.613209 | 3.862994 | 0.000307 | 0.007759 | -0.29404 | Myosin family protein, putative, expressed | elongation zone | 12 h |
| HORVU.MOREX.r3.6HG0618760.1 | -0.79667 | 4.331592 | -3.69076 | 0.000529 | 0.010355 | -0.29407 | WRKY transcription factor, putative | elongation zone | 12 h |
| HORVU.MOREX.r3.1HG0001460.1 | -0.83629 | 5.414195 | -3.73424 | 0.000461 | 0.009584 | -0.29623 | Transmembrane protein, putative (DUF247) | elongation zone | 12 h |
| HORVU.MOREX.r3.7HG0737890.1 | 1.135165 | 1.23238 | 3.78358 | 0.000395 | 0.008847 | -0.2992 | Peroxidase | elongation zone | 12 h |
| HORVU.MOREX.r3.3HG0299820.1 | -1.06121 | 7.690345 | -3.77117 | 0.000411 | 0.009004 | -0.29949 | Histone H3 | elongation zone | 12 h |
| HORVU.MOREX.r3.5HG0436660.1 | -0.45509 | 4.868326 | -3.73039 | 0.000467 | 0.009659 | -0.30247 | #N/A | elongation zone | 12 h |
| HORVU.MOREX.r3.3HG0306840.1 | 0.55811 | 5.072726 | 3.770131 | 0.000412 | 0.009024 | -0.30293 | KH domain-containing protein | elongation zone | 12 h |

**Table S1** Continued.

| ID | logFC | AveExpr | t | P.Value | adj.P.Val | B | gene description | root zone | time point |
|---|---|---|---|---|---|---|---|---|---|
| HORVU.MOREX.r3.1HG0062110.1 | 1.357883 | 0.941667 | 3.722814 | 0.000478 | 0.009812 | -0.3052 | Pentatricopeptide repeat-containing protein | elongation zone | 12 h |
| HORVU.MOREX.r3.7HG0663150.1 | -0.39192 | 4.624346 | -3.71596 | 0.000489 | 0.009916 | -0.30995 | Protein trichome birefringence | elongation zone | 12 h |
| HORVU.MOREX.r3.6HG0543720.1 | -0.90855 | 2.548383 | -3.67052 | 0.000563 | 0.01063 | -0.31027 | Histone H2A | elongation zone | 12 h |
| HORVU.MOREX.r3.4HG0381310.1 | -1.02883 | -0.06914 | -3.67211 | 0.00056 | 0.010623 | -0.31068 | Cell surface antigen I/II | elongation zone | 12 h |
| HORVU.MOREX.r3.4HG0399970.1 | 1.059518 | 4.109756 | 3.748828 | 0.000441 | 0.009306 | -0.31111 | Cytochrome b561 and DOMON domain-containing protein | elongation zone | 12 h |
| HORVU.MOREX.r3.5HG0521610.1 | 1.234662 | 0.681734 | 3.758285 | 0.000428 | 0.009191 | -0.31131 | Protein kinase | elongation zone | 12 h |
| HORVU.MOREX.r3.3HG0320040.1 | -1.47022 | 0.715118 | -3.87817 | 0.000292 | 0.007614 | -0.31505 | Phytochromobilin:ferredoxin oxidoreductase | elongation zone | 12 h |
| HORVU.MOREX.r3.6HG0597320.1 | -0.31118 | 6.618535 | -3.7682 | 0.000415 | 0.009031 | -0.31802 | ATP-dependent Clp protease proteolytic subunit | elongation zone | 12 h |
| HORVU.MOREX.r3.2HG0129450.1 | 0.959651 | 1.029346 | 3.72803 | 0.00047 | 0.009706 | -0.32099 | basic helix-loop-helix (bHLH) DNA-binding superfamily protein | elongation zone | 12 h |
| HORVU.MOREX.r3.2HG0150450.1 | -0.6592 | 7.245664 | -3.77386 | 0.000407 | 0.008984 | -0.32102 | Pyruvate kinase | elongation zone | 12 h |
| HORVU.MOREX.r3.3HG0314070.1 | 1.444048 | -1.61281 | 3.663376 | 0.000576 | 0.010779 | -0.32441 | Protein kinase family protein | elongation zone | 12 h |
| HORVU.MOREX.r3.3HG0221460.1 | -0.59807 | 7.255591 | -3.76626 | 0.000417 | 0.009058 | -0.32665 | Glycosyltransferase | elongation zone | 12 h |
| HORVU.MOREX.r3.2HG0187450.1 | 1.359735 | 0.035956 | 3.723143 | 0.000478 | 0.009812 | -0.32773 | Rhomboid-like protein | elongation zone | 12 h |
| HORVU.MOREX.r3.6HG0610980.1 | 1.293514 | -1.88326 | 3.653501 | 0.000594 | 0.011015 | -0.3306 | Nucleotide/sugar transporter family protein | elongation zone | 12 h |
| HORVU.MOREX.r3.7HG0751340.1 | 1.017261 | 0.25739 | 3.715102 | 0.00049 | 0.009933 | -0.33169 | Core-2/I-branching beta-1,6-N-acetylglucosaminyltransferase family protein | elongation zone | 12 h |
| HORVU.MOREX.r3.3HG0323530.1 | -0.76598 | 3.205547 | -3.65464 | 0.000592 | 0.010996 | -0.33233 | cysteine-rich RLK (RECEPTOR-like protein kinase) 30 | elongation zone | 12 h |
| HORVU.MOREX.r3.2HG0170230.1 | 0.574424 | 3.492538 | 3.658786 | 0.000584 | 0.010905 | -0.33679 | Smr domain containing protein | elongation zone | 12 h |
| HORVU.MOREX.r3.4HG0354010.1 | 2.217604 | -0.99752 | 3.692471 | 0.000526 | 0.010342 | -0.33698 | LURP-one-like protein | elongation zone | 12 h |
| HORVU.MOREX.r3.5HG0527500.1 | -0.53972 | 4.452639 | -3.67417 | 0.000557 | 0.010588 | -0.33813 | Ribosomal L11 methyltransferase | elongation zone | 12 h |
| HORVU.MOREX.r3.2HG0190470.1 | -0.58734 | 4.623548 | -3.68135 | 0.000544 | 0.010463 | -0.3434 | Argonaute family protein | elongation zone | 12 h |
| HORVU.MOREX.r3.6HG0541250.1 | -0.25052 | 7.143195 | -3.77686 | 0.000403 | 0.00894 | -0.34402 | Acetate kinase | elongation zone | 12 h |
| HORVU.MOREX.r3.1HG0076060.1 | -0.42041 | 6.353402 | -3.76748 | 0.000416 | 0.009042 | -0.3454 | C2 calcium/lipid-binding and GRAM domain protein | elongation zone | 12 h |
| HORVU.MOREX.r3.5HG0454230.1 | 0.751017 | 2.88731 | 3.664062 | 0.000575 | 0.010776 | -0.34639 | Aminopeptidase | elongation zone | 12 h |
| HORVU.MOREX.r3.4HG0396220.1 | -0.38499 | 8.114652 | -3.8104 | 0.000363 | 0.008498 | -0.34877 | Alpha-glucosidase like protein | elongation zone | 12 h |
| HORVU.MOREX.r3.5HG0501200.1 | -0.54024 | 4.695239 | -3.70311 | 0.000509 | 0.010183 | -0.34917 | Thioredoxin superfamily protein | elongation zone | 12 h |
| HORVU.MOREX.r3.7HG0707160.1 | 1.662232 | -0.16087 | 3.654331 | 0.000592 | 0.010997 | -0.35173 | NAC domain-containing protein, putative | elongation zone | 12 h |
| HORVU.MOREX.r3.5HG0509120.1 | 1.036665 | 5.607225 | 3.74911 | 0.00044 | 0.009306 | -0.35427 | Auxin-responsive protein | elongation zone | 12 h |
| HORVU.MOREX.r3.1HG0026010.1 | 0.648876 | 4.391694 | 3.688693 | 0.000532 | 0.010373 | -0.35443 | Kinase interacting (KIP1-like) family protein | elongation zone | 12 h |
| HORVU.MOREX.r3.4HG0394870.1 | 1.106204 | -0.77161 | 3.658296 | 0.000585 | 0.010911 | -0.35459 | Formin-like protein | elongation zone | 12 h |
| HORVU.MOREX.r3.3HG0298540.1 | 0.658454 | 7.885451 | 3.881932 | 0.000289 | 0.007575 | -0.35631 | Phosphatidylinositol transfer protein | elongation zone | 12 h |
| HORVU.MOREX.r3.1HG0039370.1 | -0.40796 | 4.945966 | -3.69566 | 0.000521 | 0.010292 | -0.36083 | Leucine-rich repeat (LRR) family protein | elongation zone | 12 h |
| HORVU.MOREX.r3.5HG0476380.1 | -0.66147 | 4.634573 | -3.66663 | 0.00057 | 0.010701 | -0.36397 | Major facilitator superfamily transporter | elongation zone | 12 h |
| HORVU.MOREX.r3.5HG0512210.1 | 2.998356 | -2.87785 | 3.952795 | 0.00023 | 0.006891 | -0.36414 | WRKY transcription factor | elongation zone | 12 h |
| HORVU.MOREX.r3.1HG0005750.1 | -0.44392 | 7.06453 | -3.76884 | 0.000414 | 0.009031 | -0.36662 | Ankyrin repeat family protein | elongation zone | 12 h |
| HORVU.MOREX.r3.2HG0192270.1 | 2.550248 | -2.74995 | 3.825406 | 0.000346 | 0.008288 | -0.36918 | Sodium transporter | elongation zone | 12 h |
| HORVU.MOREX.r3.7HG0749090.1 | 1.181462 | -0.58865 | 3.674674 | 0.000556 | 0.010588 | -0.37218 | Expansin-like protein | elongation zone | 12 h |
| HORVU.MOREX.r3.4HG0416070.1 | -2.14611 | 7.030234 | -3.69419 | 0.000523 | 0.010309 | -0.37402 | GATA transcription factor | elongation zone | 12 h |
| HORVU.MOREX.r3.1HG0079810.1 | -0.54028 | 3.696599 | -3.64346 | 0.000612 | 0.011222 | -0.37467 | WD repeat protein-like | elongation zone | 12 h |
| HORVU.MOREX.r3.2HG0172580.1 | 1.652595 | -0.45214 | 3.674805 | 0.000556 | 0.010588 | -0.38097 | Sugar transporter, putative | elongation zone | 12 h |
| HORVU.MOREX.r3.3HG0247250.1 | -1.32408 | 3.952989 | -3.63603 | 0.000627 | 0.011422 | -0.38407 | V-type proton ATPase subunit E | elongation zone | 12 h |
| HORVU.MOREX.r3.3HG0303240.1 | 1.11421 | 2.978435 | 3.632505 | 0.000634 | 0.01149 | -0.38983 | Ankyrin repeat-containing protein | elongation zone | 12 h |

**Table S1** Continued.

| ID | logFC | AveExpr | t | P.Value | adj.P.Val | B | gene description | root zone | time point |
|---|---|---|---|---|---|---|---|---|---|
| HORVU.MOREX.r3.2HG0160550.1 | -2.71679 | 2.252308 | -3.86636 | 0.000304 | 0.007757 | -0.39115 | At4g33800-like protein | elongation zone | 12 h |
| HORVU.MOREX.r3.7HG0641860.1 | 1.969302 | 0.566345 | 3.710691 | 0.000497 | 0.010012 | -0.39221 | Ubiquitin carboxyl-terminal hydrolase 2 | elongation zone | 12 h |
| HORVU.MOREX.r3.1HG0090710.1 | -0.39015 | 4.396711 | -3.69213 | 0.000526 | 0.010342 | -0.39238 | transmembrane protein | elongation zone | 12 h |
| HORVU.MOREX.r3.5HG0425250.1 | 1.550422 | -1.96474 | 3.633741 | 0.000631 | 0.011477 | -0.39559 | Disease resistance protein RPH8A | elongation zone | 12 h |
| HORVU.MOREX.r3.6HG0597400.2 | 1.318778 | 3.448997 | 3.674268 | 0.000557 | 0.010588 | -0.39827 | Carboxypeptidase | elongation zone | 12 h |
| HORVU.MOREX.r3.7HG0639360.1 | 0.74913 | 7.224542 | 3.751994 | 0.000436 | 0.009279 | -0.40094 | Autophagy-related protein 22-1 | elongation zone | 12 h |
| HORVU.MOREX.r3.6HG0545630.1 | -1.49407 | 3.70973 | -3.62804 | 0.000642 | 0.011595 | -0.40303 | Histone H2A | elongation zone | 12 h |
| HORVU.MOREX.r3.1HG0076800.1 | 0.647108 | 5.255637 | 3.750939 | 0.000438 | 0.009279 | -0.41338 | Kinesin-like protein | elongation zone | 12 h |
| HORVU.MOREX.r3.5HG0465520.1 | -1.86399 | 0.949637 | -3.86064 | 0.000309 | 0.007759 | -0.41439 | Transposon protein, putative, CACTA, En/Spm sub-class, expressed | elongation zone | 12 h |
| HORVU.MOREX.r3.6HG0612070.1 | -0.52599 | 5.864582 | -3.69081 | 0.000529 | 0.010355 | -0.41489 | RNA-binding family protein | elongation zone | 12 h |
| HORVU.MOREX.r3.3HG0281730.1 | -1.36518 | 3.703997 | -3.63133 | 0.000636 | 0.011508 | -0.41507 | AP2/B3 transcription factor family protein | elongation zone | 12 h |
| HORVU.MOREX.r3.7HG0751030.1 | 1.318873 | 2.711562 | 3.621869 | 0.000655 | 0.011707 | -0.42333 | MYB-related protein | elongation zone | 12 h |
| HORVU.MOREX.r3.1HG0079950.1 | 0.311174 | 5.644065 | 3.718651 | 0.000484 | 0.009892 | -0.42777 | Lys-63-specific deubiquitinase BRCC36 | elongation zone | 12 h |
| HORVU.MOREX.r3.6HG0606990.1 | 1.427945 | -0.1834 | 3.799081 | 0.000376 | 0.008593 | -0.42865 | RNA-binding protein Nova-1 | elongation zone | 12 h |
| HORVU.MOREX.r3.5HG0528310.1 | 0.604431 | 3.716139 | 3.669076 | 0.000566 | 0.010658 | -0.4291 | SPX domain-containing family protein | elongation zone | 12 h |
| HORVU.MOREX.r3.3HG0303970.1 | -0.67794 | 5.51606 | -3.6709 | 0.000562 | 0.01063 | -0.42917 | Internal alternative NAD(P)H-ubiquinone oxidoreductase A1, mitochondrial | elongation zone | 12 h |
| HORVU.MOREX.r3.1HG0068710.1 | -1.07378 | 4.871841 | -3.67635 | 0.000553 | 0.010558 | -0.42968 | Ethylene-responsive factor-like transcription factor | elongation zone | 12 h |
| HORVU.MOREX.r3.7HG0685360.1 | 0.604994 | 4.544409 | 3.70219 | 0.00051 | 0.010189 | -0.43073 | Receptor protein kinase, putative | elongation zone | 12 h |
| HORVU.MOREX.r3.1HG0057610.1 | 0.698136 | 4.023099 | 3.633624 | 0.000631 | 0.011477 | -0.43241 | Peptide methionine sulfoxide reductase MsrA | elongation zone | 12 h |
| HORVU.MOREX.r3.3HG0270910.1 | -0.29532 | 6.963859 | -3.75394 | 0.000434 | 0.009279 | -0.45039 | Protein YIPF | elongation zone | 12 h |
| HORVU.MOREX.r3.1HG0006660.1 | 0.94286 | 5.455877 | 3.676244 | 0.000553 | 0.010558 | -0.45247 | Inorganic pyrophosphatase family protein | elongation zone | 12 h |
| HORVU.MOREX.r3.1HG0060340.1 | -1.41274 | 6.073412 | -3.6478 | 0.000604 | 0.011171 | -0.45447 | Histone H2B | elongation zone | 12 h |
| HORVU.MOREX.r3.7HG0667730.1 | 0.459143 | 5.429237 | 3.713196 | 0.000493 | 0.009973 | -0.45624 | Chaperone protein dnaJ | elongation zone | 12 h |
| HORVU.MOREX.r3.7HG0739640.1 | -0.96227 | 8.298465 | -3.73639 | 0.000458 | 0.009546 | -0.45636 | Auxin efflux carrier component | elongation zone | 12 h |
| HORVU.MOREX.r3.2HG0174750.1 | 1.195019 | 1.498345 | 3.709999 | 0.000498 | 0.010024 | -0.45749 | Fasciclin-like arabinogalactan protein | elongation zone | 12 h |
| HORVU.MOREX.r3.7HG0664720.1 | 1.147644 | 0.421259 | 3.70465 | 0.000506 | 0.010163 | -0.45932 | Blue copper protein | elongation zone | 12 h |
| HORVU.MOREX.r3.6HG0616710.1 | -0.98545 | 5.40807 | -3.64205 | 0.000615 | 0.011251 | -0.45937 | Cyclin family protein | elongation zone | 12 h |
| HORVU.MOREX.r3.6HG0554840.1 | 0.877771 | 2.671575 | 3.622247 | 0.000654 | 0.011707 | -0.46238 | Phosphopantetheine adenylyltransferase | elongation zone | 12 h |
| HORVU.MOREX.r3.4HG0400520.1 | 1.263484 | -0.786 | 3.623587 | 0.000651 | 0.011674 | -0.46458 | Lipid transfer protein | elongation zone | 12 h |
| HORVU.MOREX.r3.2HG0169720.1 | -0.49615 | 4.773186 | -3.6329 | 0.000633 | 0.01149 | -0.46727 | Zinc finger CCCH domain-containing protein 28 | elongation zone | 12 h |
| HORVU.MOREX.r3.1HG0014680.1 | 1.378384 | -0.77974 | 3.624964 | 0.000649 | 0.011649 | -0.4716 | Tetraspanin family protein | elongation zone | 12 h |
| HORVU.MOREX.r3.6HG0607960.1 | 0.530257 | 6.253417 | 3.763963 | 0.00042 | 0.009095 | -0.47449 | Protein KINESIN LIGHT CHAIN-RELATED 3 | elongation zone | 12 h |
| HORVU.MOREX.r3.2HG0109340.1 | 1.204394 | 0.874826 | 3.603828 | 0.000692 | 0.012162 | -0.47543 | Wound-induced protein 1, putative, expressed | elongation zone | 12 h |
| HORVU.MOREX.r3.1HG0050220.1 | 1.403533 | -1.46767 | 3.601413 | 0.000698 | 0.012162 | -0.47556 | Polyketide cyclase/dehydrase and lipid transport superfamily protein | elongation zone | 12 h |
| HORVU.MOREX.r3.5HG0470030.1 | -0.4954 | 7.999201 | -3.77223 | 0.000409 | 0.008991 | -0.47611 | 3-ketoacyl-CoA thiolase | elongation zone | 12 h |
| HORVU.MOREX.r3.5HG0465260.1 | -0.77701 | 2.270829 | -3.61344 | 0.000672 | 0.011932 | -0.47707 | Zinc finger HIT domain-containing protein 1 | elongation zone | 12 h |
| HORVU.MOREX.r3.2HG0119360.1 | -2.1031 | -1.10965 | -3.83677 | 0.000334 | 0.008151 | -0.47711 | Gibberellin receptor GID1A | elongation zone | 12 h |
| HORVU.MOREX.r3.3HG0246030.1 | 1.088338 | 0.793866 | 3.609912 | 0.00068 | 0.012034 | -0.47865 | Blue copper protein | elongation zone | 12 h |
| HORVU.MOREX.r3.2HG0139810.1 | 0.927478 | 0.63654 | 3.689556 | 0.000531 | 0.01036 | -0.47904 | Heavy metal transport/detoxification superfamily protein | elongation zone | 12 h |
| HORVU.MOREX.r3.2HG0105050.1 | 1.856826 | 0.102901 | 3.802656 | 0.000372 | 0.008553 | -0.48014 | 2-oxoglutarate (2OG) and Fe(II)-dependent oxygenase superfamily protein | elongation zone | 12 h |
| HORVU.MOREX.r3.5HG0470170.1 | -0.59204 | 3.126834 | -3.60155 | 0.000697 | 0.012162 | -0.48019 | F-box family protein | elongation zone | 12 h |

**Table S1** Continued.

| ID | logFC | AveExpr | t | P.Value | adj.P.Val | B | gene description | root zone | time point |
|---|---|---|---|---|---|---|---|---|---|
| HORVU.MOREX.r3.1HG0057030.1 | -0.69874 | 6.546046 | -3.70199 | 0.00051 | 0.010189 | -0.48257 | Peroxidase | elongation zone | 12 h |
| HORVU.MOREX.r3.4HG0414000.1 | 0.769809 | 1.7924 | 3.626084 | 0.000646 | 0.011635 | -0.4838 | Short-chain dehydrogenase/reductase family protein | elongation zone | 12 h |
| HORVU.MOREX.r3.5HG0510500.1 | 0.463597 | 5.572952 | 3.700993 | 0.000512 | 0.01021 | -0.48473 | RNA binding family protein isoform 1 | elongation zone | 12 h |
| HORVU.MOREX.r3.7HG0670690.1 | 0.621261 | 8.088776 | 3.824432 | 0.000347 | 0.008288 | -0.48534 | HIPL1 protein | elongation zone | 12 h |
| HORVU.MOREX.r3.3HG0295450.1 | -0.82213 | 2.043228 | -3.60571 | 0.000688 | 0.01213 | -0.48624 | Cytochrome P450 family protein | elongation zone | 12 h |
| HORVU.MOREX.r3.5HG0470940.1 | 0.50071 | 5.957094 | 3.71712 | 0.000487 | 0.0099 | -0.48669 | Protein ENHANCED DISEASE RESISTANCE 2-like | elongation zone | 12 h |
| HORVU.MOREX.r3.2HG0118220.1 | 1.083065 | 1.251256 | 3.627374 | 0.000644 | 0.011609 | -0.48824 | En/Spm-like transposon protein | elongation zone | 12 h |
| HORVU.MOREX.r3.4HG0400800.1 | -0.42632 | 4.757552 | -3.64677 | 0.000606 | 0.011192 | -0.48902 | Transcription factor | elongation zone | 12 h |
| HORVU.MOREX.r3.2HG0194450.1 | -0.38186 | 5.508153 | -3.65006 | 0.0006 | 0.011123 | -0.49132 | Secretory carrier-associated membrane protein | elongation zone | 12 h |
| HORVU.MOREX.r3.3HG0285150.1 | 0.817734 | 1.826069 | 3.617495 | 0.000664 | 0.011834 | -0.49522 | Tryptophan aminotransferase | elongation zone | 12 h |
| HORVU.MOREX.r3.4HG0394690.1 | -0.32971 | 5.03075 | -3.65977 | 0.000582 | 0.010887 | -0.49828 | #N/A | elongation zone | 12 h |
| HORVU.MOREX.r3.7HG0749150.1 | -2.05551 | 3.418595 | -3.6497 | 0.000601 | 0.011126 | -0.49894 | DUF241 domain protein (DUF241) | elongation zone | 12 h |
| HORVU.MOREX.r3.3HG0309780.1 | 2.840115 | -2.21583 | 3.759706 | 0.000426 | 0.009169 | -0.49927 | RING/U-box superfamily protein | elongation zone | 12 h |
| HORVU.MOREX.r3.1HG0012170.1 | -0.84196 | 3.51455 | -3.59266 | 0.000717 | 0.012358 | -0.49982 | Disease resistance protein RPM1 | elongation zone | 12 h |
| HORVU.MOREX.r3.2HG0119220.1 | -0.54563 | 4.892203 | -3.67081 | 0.000563 | 0.01063 | -0.50163 | Gibberellin receptor GID1a | elongation zone | 12 h |
| HORVU.MOREX.r3.3HG0287800.1 | -0.68859 | 4.734584 | -3.61619 | 0.000666 | 0.011861 | -0.50487 | 6-phosphogluconate dehydrogenase family protein | elongation zone | 12 h |
| HORVU.MOREX.r3.1HG0074600.1 | -1.22446 | 3.896702 | -3.58829 | 0.000726 | 0.012414 | -0.50703 | Cytochrome P450 | elongation zone | 12 h |
| HORVU.MOREX.r3.3HG0251950.1 | -0.69158 | 3.77745 | -3.60677 | 0.000686 | 0.012117 | -0.50932 | Polynucleotidyl transferase, ribonuclease H-like superfamily protein, putative | elongation zone | 12 h |
| HORVU.MOREX.r3.4HG0364050.1 | 0.761651 | 2.580176 | 3.593111 | 0.000716 | 0.012358 | -0.50985 | Sodium/hydrogen exchanger | elongation zone | 12 h |
| HORVU.MOREX.r3.1HG0006650.1 | -0.87831 | 8.905369 | -3.74455 | 0.000447 | 0.00939 | -0.51025 | Histone H2A | elongation zone | 12 h |
| HORVU.MOREX.r3.4HG0386000.2 | -0.51779 | 3.642466 | -3.60281 | 0.000695 | 0.012162 | -0.51296 | Transthyretin-like S-allantoin synthase protein | elongation zone | 12 h |
| HORVU.MOREX.r3.3HG0318700.1 | 0.508878 | 6.088388 | 3.727488 | 0.000471 | 0.009709 | -0.51805 | Phosphotransferase | elongation zone | 12 h |
| HORVU.MOREX.r3.3HG0285570.1 | -0.72853 | 3.375913 | -3.60943 | 0.000681 | 0.012039 | -0.51832 | Serine acetyltransferase | elongation zone | 12 h |
| HORVU.MOREX.r3.2HG0129200.1 | -0.58836 | 5.391664 | -3.64399 | 0.000611 | 0.011213 | -0.51916 | glutamyl-tRNA (Gln) amidotransferase subunit A (DUF620) | elongation zone | 12 h |
| HORVU.MOREX.r3.7HG0674590.1 | 1.129929 | 1.003958 | 3.591586 | 0.000719 | 0.012373 | -0.52179 | Retrovirus-related Pol polyprotein from transposon TNT 1-94 | elongation zone | 12 h |
| HORVU.MOREX.r3.7HG0707220.1 | 0.621046 | 4.707516 | 3.695759 | 0.00052 | 0.010292 | -0.52764 | #N/A | elongation zone | 12 h |
| HORVU.MOREX.r3.7HG0683940.1 | 0.891844 | 2.232556 | 3.58418 | 0.000736 | 0.012507 | -0.52933 | 65-kDa microtubule-associated-like protein | elongation zone | 12 h |
| HORVU.MOREX.r3.5HG0522090.1 | 0.601137 | 2.891849 | 3.614038 | 0.000671 | 0.011927 | -0.53187 | Exocyst complex component EXO84B | elongation zone | 12 h |
| HORVU.MOREX.r3.1HG0049550.1 | 0.399156 | 3.543555 | 3.594221 | 0.000713 | 0.01234 | -0.53676 | Protein BPS1, chloroplastic | elongation zone | 12 h |
| HORVU.MOREX.r3.6HG0570960.1 | -1.1965 | 6.423531 | -3.63423 | 0.00063 | 0.011476 | -0.5393 | Histone H3 | elongation zone | 12 h |
| HORVU.MOREX.r3.4HG0410640.1 | -0.39703 | 8.151713 | -3.74119 | 0.000451 | 0.009442 | -0.54112 | Dolichyl-diphosphooligosaccharide--protein glycosyltransferase subunit 1 | elongation zone | 12 h |
| HORVU.MOREX.r3.5HG0532560.1 | 1.897846 | -0.15138 | 3.59502 | 0.000711 | 0.012335 | -0.54124 | Ras-related protein, expressed | elongation zone | 12 h |
| HORVU.MOREX.r3.4HG0335170.1 | 1.250295 | 1.390233 | 3.7052 | 0.000505 | 0.010156 | -0.54189 | Core-2/I-branching beta-1,6-N-acetylglucosaminyltransferase family protein | elongation zone | 12 h |
| HORVU.MOREX.r3.7HG0723730.1 | 0.903529 | 2.888777 | 3.644966 | 0.00061 | 0.011213 | -0.54216 | RNA binding protein, putative | elongation zone | 12 h |
| HORVU.MOREX.r3.7HG0656430.1 | -0.37147 | 6.915192 | -3.67976 | 0.000547 | 0.010502 | -0.54297 | Zinc finger CCCH domain protein | elongation zone | 12 h |
| HORVU.MOREX.r3.7HG0731950.1 | 0.785413 | 3.076597 | 3.684006 | 0.00054 | 0.010443 | -0.54671 | #N/A | elongation zone | 12 h |
| HORVU.MOREX.r3.6HG0616300.1 | 0.550224 | 5.182691 | 3.695279 | 0.000521 | 0.010294 | -0.54724 | Histidine kinase | elongation zone | 12 h |
| HORVU.MOREX.r3.5HG0482360.1 | -0.78437 | 2.02018 | -3.63944 | 0.00062 | 0.011322 | -0.54849 | WAT1-related protein | elongation zone | 12 h |
| HORVU.MOREX.r3.7HG0687080.1 | -0.55476 | 4.583869 | -3.60789 | 0.000684 | 0.012086 | -0.54971 | RNA-binding family protein | elongation zone | 12 h |
| HORVU.MOREX.r3.3HG0248280.1 | -0.39368 | 8.50615 | -3.7484 | 0.000441 | 0.009306 | -0.54999 | Reticulon-like protein | elongation zone | 12 h |
| HORVU.MOREX.r3.5HG0524800.1 | -1.62524 | 1.969957 | -3.64575 | 0.000608 | 0.011202 | -0.55103 | RNA polymerase II transcription mediator | elongation zone | 12 h |

**Table S1** Continued.

| ID | logFC | AveExpr | t | P.Value | adj.P.Val | B | gene description | root zone | time point |
|---|---|---|---|---|---|---|---|---|---|
| HORVU.MOREX.r3.6HG0567330.1 | -1.2531 | 5.748869 | -3.60195 | 0.000696 | 0.012162 | -0.55194 | aspartyl/glutamyl-tRNA(Asn/Gln) amidotransferase subunit | elongation zone | 12 h |
| HORVU.MOREX.r3.1HG0023620.1 | -0.9138 | 6.596829 | -3.66366 | 0.000575 | 0.010779 | -0.55195 | Histone H3 | elongation zone | 12 h |
| HORVU.MOREX.r3.1HG0072520.1 | -1.06453 | 4.709601 | -3.60084 | 0.000699 | 0.012173 | -0.55386 | #N/A | elongation zone | 12 h |
| HORVU.MOREX.r3.1HG0064610.1 | -0.97295 | 5.770306 | -3.60178 | 0.000697 | 0.012162 | -0.5576 | membrane-associated kinase regulator | elongation zone | 12 h |
| HORVU.MOREX.r3.3HG0280310.1 | -0.68124 | 2.629974 | -3.56879 | 0.000771 | 0.012872 | -0.55765 | #N/A | elongation zone | 12 h |
| HORVU.MOREX.r3.5HG0480130.1 | 2.366623 | -2.63524 | 3.773247 | 0.000408 | 0.008984 | -0.55856 | Kinase family protein | elongation zone | 12 h |
| HORVU.MOREX.r3.7HG0718200.1 | 0.854267 | 3.590015 | 3.67245 | 0.00056 | 0.010623 | -0.56015 | Kinesin-like protein | elongation zone | 12 h |
| HORVU.MOREX.r3.1HG0036930.1 | -1.08193 | 6.161598 | -3.61623 | 0.000666 | 0.011861 | -0.56334 | Histone H2A | elongation zone | 12 h |
| HORVU.MOREX.r3.1HG0061710.1 | 0.502223 | 5.632386 | 3.702871 | 0.000509 | 0.010183 | -0.5659 | Trichome birefringence-like protein | elongation zone | 12 h |
| HORVU.MOREX.r3.4HG0407560.1 | 0.414982 | 3.734856 | 3.630866 | 0.000637 | 0.011514 | -0.56594 | Ras family protein | elongation zone | 12 h |
| HORVU.MOREX.r3.7HG0650210.1 | -0.41276 | 6.623351 | -3.68352 | 0.000541 | 0.010448 | -0.56774 | Oligosaccharyltransferase complex subunit ostc | elongation zone | 12 h |
| HORVU.MOREX.r3.3HG0246500.1 | 0.862979 | 3.040522 | 3.693171 | 0.000525 | 0.010332 | -0.56803 | Peptide-N4-(N-acetyl-beta-glucosaminyl)asparagine amidase A | elongation zone | 12 h |
| HORVU.MOREX.r3.3HG0319760.1 | -0.45627 | 3.423519 | -3.58045 | 0.000744 | 0.012578 | -0.57121 | Alpha-1,3-mannosyl-glycoprotein 2-beta-N-acetylglucosaminyltransferase | elongation zone | 12 h |
| HORVU.MOREX.r3.7HG0706140.1 | -0.56111 | 4.415559 | -3.58963 | 0.000723 | 0.012401 | -0.57234 | Bromodomain-containing factor 2 | elongation zone | 12 h |
| HORVU.MOREX.r3.5HG0420670.1 | -0.7737 | 3.163154 | -3.56246 | 0.000786 | 0.013014 | -0.57409 | Translation elongation factor EF1A/initiation factor IF2gamma family protein | elongation zone | 12 h |
| HORVU.MOREX.r3.6HG0553150.1 | -0.53218 | 6.876917 | -3.69196 | 0.000527 | 0.010342 | -0.57422 | Polyubiquitin | elongation zone | 12 h |
| HORVU.MOREX.r3.4HG0344990.1 | -1.23095 | 2.816049 | -3.58428 | 0.000735 | 0.012507 | -0.57447 | NBS-LRR disease resistance protein family-1 | elongation zone | 12 h |
| HORVU.MOREX.r3.5HG0462450.1 | 1.304696 | -0.84666 | 3.594265 | 0.000713 | 0.01234 | -0.57629 | S-acyltransferase | elongation zone | 12 h |
| HORVU.MOREX.r3.1HG0053450.1 | 0.925669 | 3.313219 | 3.711733 | 0.000495 | 0.009994 | -0.57879 | WEB family protein, chloroplastic | elongation zone | 12 h |
| HORVU.MOREX.r3.7HG0752370.1 | 0.765482 | 2.088249 | 3.667256 | 0.000569 | 0.010699 | -0.58343 | NBS-LRR-like resistance protein | elongation zone | 12 h |
| HORVU.MOREX.r3.4HG0388310.1 | 0.425172 | 3.870759 | 3.638158 | 0.000623 | 0.011357 | -0.58599 | Protein kinase | elongation zone | 12 h |
| HORVU.MOREX.r3.7HG0728910.1 | -0.83454 | 4.539344 | -3.58285 | 0.000739 | 0.012516 | -0.58664 | Ring finger protein, putative | elongation zone | 12 h |
| HORVU.MOREX.r3.2HG0130180.1 | -0.30301 | 5.922255 | -3.65611 | 0.000589 | 0.010966 | -0.58836 | Glycosyltransferase | elongation zone | 12 h |
| HORVU.MOREX.r3.3HG0234010.1 | 1.464655 | -0.09759 | 3.657394 | 0.000587 | 0.010932 | -0.58922 | Xyloglucan endotransglucosylase/hydrolase | elongation zone | 12 h |
| HORVU.MOREX.r3.2HG0116540.1 | -0.59965 | 3.495583 | -3.58655 | 0.00073 | 0.012447 | -0.58968 | 5-aminolevulinate synthase, mitochondrial | elongation zone | 12 h |
| HORVU.MOREX.r3.3HG0288710.1 | 1.449009 | -1.93307 | 3.563459 | 0.000784 | 0.012995 | -0.59141 | GDSL esterase/lipase | elongation zone | 12 h |
| HORVU.MOREX.r3.3HG0279760.1 | -0.78786 | 3.223061 | -3.57086 | 0.000766 | 0.012828 | -0.59223 | NAC domain-containing protein | elongation zone | 12 h |
| HORVU.MOREX.r3.1HG0093750.1 | -0.93936 | 3.434371 | -3.56017 | 0.000792 | 0.013047 | -0.59257 | ATP-dependent zinc metalloprotease FtsH | elongation zone | 12 h |
| HORVU.MOREX.r3.7HG0663750.1 | 1.429923 | 3.681851 | 3.560953 | 0.00079 | 0.013047 | -0.59285 | Galactoside 2-alpha-L-fucosyltransferase | elongation zone | 12 h |
| HORVU.MOREX.r3.5HG0461950.1 | 0.803167 | 2.97161 | 3.588095 | 0.000727 | 0.012414 | -0.59325 | 2-isopropylmalate synthase 2 | elongation zone | 12 h |
| HORVU.MOREX.r3.6HG0601680.1 | 0.862149 | 2.148004 | 3.588378 | 0.000726 | 0.012414 | -0.59343 | Vesicle-associated 1-1-like protein | elongation zone | 12 h |
| HORVU.MOREX.r3.3HG0316110.1 | -1.05048 | 5.235108 | -3.6431 | 0.000613 | 0.011224 | -0.59368 | Basic 7S globulin 2 | elongation zone | 12 h |
| HORVU.MOREX.r3.5HG0527650.1 | -0.69457 | 4.818573 | -3.59969 | 0.000701 | 0.012206 | -0.59387 | Succinate--CoA ligase [ADP-forming] subunit alpha-1, mitochondrial | elongation zone | 12 h |
| HORVU.MOREX.r3.5HG0490720.1 | -3.94475 | 0.668588 | -4.11974 | 0.000134 | 0.00553 | -0.5982 | Retrovirus-related Pol polyprotein from transposon TNT 1-94 | elongation zone | 12 h |
| HORVU.MOREX.r3.4HG0403680.1 | -1.38283 | 3.612484 | -3.58074 | 0.000743 | 0.012578 | -0.59901 | #N/A | elongation zone | 12 h |
| HORVU.MOREX.r3.2HG0164850.1 | -0.67246 | 4.574579 | -3.60552 | 0.000689 | 0.01213 | -0.60202 | SBP (S-ribonuclease binding protein) family protein | elongation zone | 12 h |
| HORVU.MOREX.r3.1HG0063430.1 | 0.395673 | 7.658435 | 3.745901 | 0.000445 | 0.00936 | -0.60257 | Oxidoreductase, 2OG-Fe(II) oxygenase family protein | elongation zone | 12 h |
| HORVU.MOREX.r3.4HG0382620.1 | 0.481021 | 7.496381 | 3.671908 | 0.000561 | 0.010623 | -0.60364 | Argininosuccinate lyase | elongation zone | 12 h |
| HORVU.MOREX.r3.4HG0386830.1 | -1.04086 | 3.806924 | -3.54789 | 0.000822 | 0.01322 | -0.61257 | UDP-glucuronate decarboxylase protein 1 | elongation zone | 12 h |
| HORVU.MOREX.r3.3HG0231240.1 | 1.05285 | 0.82508 | 3.571363 | 0.000765 | 0.012828 | -0.61354 | Short-chain dehydrogenase/reductase family protein | elongation zone | 12 h |
| HORVU.MOREX.r3.2HG0115480.1 | -0.69267 | 3.022508 | -3.5479 | 0.000822 | 0.01322 | -0.61643 | Tesmin/TSO1-like CXC domain protein | elongation zone | 12 h |

**Table S1** Continued.

| ID | logFC | AveExpr | t | P.Value | adj.P.Val | B | gene description | root zone | time point |
|---|---|---|---|---|---|---|---|---|---|
| HORVU.MOREX.r3.1HG0087150.1 | -0.87603 | 6.174716 | -3.68177 | 0.000544 | 0.010463 | -0.62285 | Purple acid phosphatase | elongation zone | 12 h |
| HORVU.MOREX.r3.3HG0310540.1 | -0.58524 | 5.764879 | -3.62503 | 0.000648 | 0.011649 | -0.62468 | Mannosyl-oligosaccharide glucosidase | elongation zone | 12 h |
| HORVU.MOREX.r3.4HG0410090.1 | 0.598135 | 5.519883 | 3.646626 | 0.000607 | 0.011192 | -0.62652 | Hypersensitive-induced response protein 1 | elongation zone | 12 h |
| HORVU.MOREX.r3.2HG0202250.1 | 1.294398 | 1.379273 | 3.543478 | 0.000834 | 0.013311 | -0.62724 | Kelch repeat-containing protein | elongation zone | 12 h |
| HORVU.MOREX.r3.7HG0718170.1 | 1.022393 | 1.273375 | 3.546403 | 0.000826 | 0.013244 | -0.62731 | Pleiotropic drug resistance ABC transporter | elongation zone | 12 h |
| HORVU.MOREX.r3.7HG0751290.1 | -1.01768 | 0.60944 | -3.62183 | 0.000655 | 0.011707 | -0.62765 | Receptor-kinase, putative | elongation zone | 12 h |
| HORVU.MOREX.r3.4HG0398420.1 | -0.55096 | 3.560721 | -3.55753 | 0.000798 | 0.013094 | -0.62797 | Membrane steroid-binding protein | elongation zone | 12 h |
| HORVU.MOREX.r3.1HG0054070.1 | 1.20247 | 0.426324 | 3.554136 | 0.000807 | 0.01314 | -0.62927 | CASP-like protein | elongation zone | 12 h |
| HORVU.MOREX.r3.3HG0290510.1 | -0.89133 | 3.665191 | -3.54157 | 0.000838 | 0.013368 | -0.6311 | Tesmin/TSO1-like CXC domain-containing protein | elongation zone | 12 h |
| HORVU.MOREX.r3.1HG0026080.1 | 0.644875 | 4.010774 | 3.579919 | 0.000745 | 0.012588 | -0.63456 | RING finger protein | elongation zone | 12 h |
| HORVU.MOREX.r3.7HG0737440.1 | -1.17613 | 5.648436 | -3.56962 | 0.000769 | 0.012856 | -0.63481 | carboxyl-terminal peptidase (DUF239) | elongation zone | 12 h |
| HORVU.MOREX.r3.7HG0702660.1 | 1.161249 | 1.220256 | 3.554263 | 0.000806 | 0.01314 | -0.63575 | CoA ligase | elongation zone | 12 h |
| HORVU.MOREX.r3.5HG0508870.1 | 0.75041 | 4.361487 | 3.673465 | 0.000558 | 0.010601 | -0.63619 | Kinase interacting (KIP1-like) family protein, putative | elongation zone | 12 h |
| HORVU.MOREX.r3.2HG0155450.1 | -0.85433 | 3.209398 | -3.5534 | 0.000809 | 0.01314 | -0.64026 | Major facilitator superfamily protein | elongation zone | 12 h |
| HORVU.MOREX.r3.1HG0050990.1 | -2.28703 | 2.072399 | -3.8939 | 0.000278 | 0.00742 | -0.64109 | Membrane protein insertase YidC | elongation zone | 12 h |
| HORVU.MOREX.r3.5HG0426060.1 | -0.62578 | 3.90766 | -3.54972 | 0.000818 | 0.013192 | -0.64352 | Protein FAR1-RELATED SEQUENCE 3 | elongation zone | 12 h |
| HORVU.MOREX.r3.7HG0739380.1 | -0.94239 | 6.500354 | -3.62628 | 0.000646 | 0.011635 | -0.64457 | Histone H2B | elongation zone | 12 h |
| HORVU.MOREX.r3.7HG0728480.1 | 1.403506 | 0.878141 | 3.667433 | 0.000569 | 0.010699 | -0.6451 | Peroxidase | elongation zone | 12 h |
| HORVU.MOREX.r3.6HG0616790.1 | -0.5905 | 7.277735 | -3.65503 | 0.000591 | 0.010993 | -0.64552 | Transport inhibitor response 1-like protein | elongation zone | 12 h |
| HORVU.MOREX.r3.1HG0072680.1 | 0.905093 | 1.174684 | 3.542965 | 0.000835 | 0.013321 | -0.64731 | Respiratory burst oxidase-like protein | elongation zone | 12 h |
| HORVU.MOREX.r3.2HG0161670.1 | -0.52234 | 5.960687 | -3.60984 | 0.00068 | 0.012034 | -0.6477 | SWI/SNF complex component SNF12-like protein | elongation zone | 12 h |
| HORVU.MOREX.r3.1HG0024280.1 | -0.71937 | 7.969902 | -3.68535 | 0.000538 | 0.010428 | -0.64898 | Dihydrolipoyl dehydrogenase | elongation zone | 12 h |
| HORVU.MOREX.r3.1HG0095440.1 | -0.34329 | 7.430324 | -3.67131 | 0.000562 | 0.01063 | -0.64946 | #N/A | elongation zone | 12 h |
| HORVU.MOREX.r3.3HG0219410.1 | 0.816398 | 3.869903 | 3.5989 | 0.000703 | 0.012226 | -0.65044 | Hydroxyproline-rich glycoprotein family protein | elongation zone | 12 h |
| HORVU.MOREX.r3.3HG0261980.1 | -0.59022 | 3.054779 | -3.53553 | 0.000854 | 0.013502 | -0.65065 | Serine/threonine-protein phosphatase | elongation zone | 12 h |
| HORVU.MOREX.r3.7HG0687050.1 | -0.46362 | 5.185144 | -3.64444 | 0.000611 | 0.011213 | -0.65323 | NC domain-containing protein | elongation zone | 12 h |
| HORVU.MOREX.r3.3HG0313490.1 | -0.90704 | 3.185804 | -3.55614 | 0.000802 | 0.013121 | -0.65434 | Rapid alkalinization factor (RALF) family protein | elongation zone | 12 h |
| HORVU.MOREX.r3.1HG0057620.1 | -0.74779 | 6.017316 | -3.60344 | 0.000693 | 0.012162 | -0.65494 | Kinase like protein | elongation zone | 12 h |
| HORVU.MOREX.r3.3HG0277330.1 | -0.62245 | 4.294396 | -3.55351 | 0.000808 | 0.01314 | -0.66099 | CASP-like protein | elongation zone | 12 h |
| HORVU.MOREX.r3.3HG0253440.1 | -0.832 | 1.953551 | -3.54775 | 0.000823 | 0.01322 | -0.66142 | Protein RETICULATA, chloroplastic | elongation zone | 12 h |
| HORVU.MOREX.r3.6HG0629220.1 | -0.75231 | 5.709587 | -3.58779 | 0.000728 | 0.012414 | -0.66925 | methyl-coenzyme M reductase II subunit gamma, putative (DUF3741) | elongation zone | 12 h |
| HORVU.MOREX.r3.2HG0161020.1 | -0.86615 | 7.262436 | -3.64116 | 0.000617 | 0.011272 | -0.6702 | #N/A | elongation zone | 12 h |
| HORVU.MOREX.r3.7HG0640560.1 | -2.53466 | 0.442209 | -3.91514 | 0.00026 | 0.007125 | -0.6742 | NBS-LRR disease resistance protein-like protein | elongation zone | 12 h |
| HORVU.MOREX.r3.6HG0554790.1 | -0.3005 | 7.933943 | -3.68963 | 0.000531 | 0.01036 | -0.67552 | Proteasome subunit alpha type | elongation zone | 12 h |
| HORVU.MOREX.r3.5HG0504670.1 | -1.30216 | 3.865033 | -3.53927 | 0.000844 | 0.01341 | -0.67563 | Acetyl-coenzyme A synthetase | elongation zone | 12 h |
| HORVU.MOREX.r3.5HG0527670.1 | -1.94866 | 3.348267 | -3.54701 | 0.000825 | 0.013239 | -0.68071 | Cytochrome P450 | elongation zone | 12 h |
| HORVU.MOREX.r3.4HG0394980.1 | -0.83162 | 8.862444 | -3.68138 | 0.000544 | 0.010463 | -0.68237 | Allene oxide synthase | elongation zone | 12 h |
| HORVU.MOREX.r3.2HG0209810.1 | 0.432063 | 6.643137 | 3.66659 | 0.00057 | 0.010701 | -0.68602 | Katanin p80 WD40 repeat-containing subunit B1 homolog | elongation zone | 12 h |
| HORVU.MOREX.r3.5HG0532120.1 | 1.726027 | -1.75561 | 3.522887 | 0.000888 | 0.013841 | -0.68705 | Expansin | elongation zone | 12 h |
| HORVU.MOREX.r3.2HG0099740.1 | -0.39325 | 5.374952 | -3.58363 | 0.000737 | 0.012516 | -0.68991 | Zinc finger family protein | elongation zone | 12 h |
| HORVU.MOREX.r3.2HG0179410.1 | -0.56969 | 2.81034 | -3.53231 | 0.000862 | 0.013594 | -0.69063 | transmembrane protein, putative (DUF1068) | elongation zone | 12 h |

**Table S1** Continued.

| ID | logFC | AveExpr | t | P.Value | adj.P.Val | B | gene description | root zone | time point |
|---|---|---|---|---|---|---|---|---|---|
| HORVU.MOREX.r3.3HG0245400.1 | 0.618669 | 5.359013 | 3.722118 | 0.000479 | 0.009824 | -0.69191 | GDSL esterase/lipase | elongation zone | 12 h |
| HORVU.MOREX.r3.6HG0605130.1 | 1.024442 | -2.05206 | 3.529754 | 0.000869 | 0.013618 | -0.69271 | Invertase inhibitor | elongation zone | 12 h |
| HORVU.MOREX.r3.3HG0293490.1 | 1.087833 | -0.53962 | 3.570972 | 0.000766 | 0.012828 | -0.69674 | Expansin | elongation zone | 12 h |
| HORVU.MOREX.r3.7HG0730820.1 | -0.96964 | 5.001622 | -3.55021 | 0.000817 | 0.013183 | -0.69858 | Histone H2A | elongation zone | 12 h |
| HORVU.MOREX.r3.7HG0748070.1 | 1.319155 | -0.49696 | 3.553542 | 0.000808 | 0.01314 | -0.6992 | Heavy metal transport/detoxification superfamily protein, putative | elongation zone | 12 h |
| HORVU.MOREX.r3.5HG0419590.1 | 0.511222 | 6.133126 | 3.678625 | 0.000549 | 0.010519 | -0.7007 | #N/A | elongation zone | 12 h |
| HORVU.MOREX.r3.3HG0297000.1 | 1.160655 | -0.77719 | 3.515845 | 0.000907 | 0.013962 | -0.7012 | LURP-one-like protein | elongation zone | 12 h |
| HORVU.MOREX.r3.3HG0324070.1 | 1.763822 | 4.131542 | 3.529704 | 0.000869 | 0.013618 | -0.70858 | Beta-1,3-glucanase | elongation zone | 12 h |
| HORVU.MOREX.r3.1HG0062980.1 | 1.0336 | 1.135611 | 3.528358 | 0.000873 | 0.013663 | -0.71081 | L-gulonolactone oxidase | elongation zone | 12 h |
| HORVU.MOREX.r3.5HG0517490.1 | 0.764187 | 1.614895 | 3.550775 | 0.000815 | 0.01317 | -0.71093 | Protein kinase | elongation zone | 12 h |
| HORVU.MOREX.r3.4HG0348380.1 | 0.918556 | 1.737869 | 3.551674 | 0.000813 | 0.013153 | -0.71126 | phox (PX) domain-containing protein | elongation zone | 12 h |
| HORVU.MOREX.r3.5HG0521630.1 | 0.66458 | 4.704079 | 3.590578 | 0.000721 | 0.012386 | -0.71491 | Beta-1,3-galactosyltransferase-like protein | elongation zone | 12 h |
| HORVU.MOREX.r3.5HG0482590.1 | -0.92635 | 4.122873 | -3.51587 | 0.000907 | 0.013962 | -0.71602 | Major facilitator superfamily protein | elongation zone | 12 h |
| HORVU.MOREX.r3.3HG0308820.1 | -0.36137 | 9.031851 | -3.71794 | 0.000486 | 0.009897 | -0.71727 | Dolichyl-diphosphooligosaccharide--protein glycosyltransferase subunit 2 | elongation zone | 12 h |
| HORVU.MOREX.r3.6HG0634130.1 | -0.30969 | 5.21873 | -3.60215 | 0.000696 | 0.012162 | -0.71755 | Alpha-1,3-mannosyl-glycoprotein 2-beta-N-acetylglucosaminyltransferase | elongation zone | 12 h |
| HORVU.MOREX.r3.7HG0695500.1 | -0.44661 | 6.592723 | -3.61823 | 0.000662 | 0.011818 | -0.72822 | Transmembrane 9 superfamily member | elongation zone | 12 h |
| HORVU.MOREX.r3.6HG0593050.1 | 0.910784 | 0.467029 | 3.540487 | 0.000841 | 0.013377 | -0.72958 | C2 domain-containing protein / GRAM domain-containing protein | elongation zone | 12 h |
| HORVU.MOREX.r3.5HG0465400.1 | 1.213326 | 0.747502 | 3.597097 | 0.000707 | 0.012273 | -0.73118 | Trichome birefringence-like protein | elongation zone | 12 h |
| HORVU.MOREX.r3.3HG0252380.1 | 1.021872 | 2.240647 | 3.50746 | 0.00093 | 0.014174 | -0.73463 | DNA-binding storekeeper protein-related transcriptional regulator | elongation zone | 12 h |
| HORVU.MOREX.r3.3HG0276130.1 | 1.096246 | -0.22255 | 3.507177 | 0.000931 | 0.014175 | -0.73559 | Phospholipase A1 | elongation zone | 12 h |
| HORVU.MOREX.r3.2HG0113720.1 | 1.164831 | 3.679958 | 3.603302 | 0.000694 | 0.012162 | -0.73623 | Cytosolic Fe-S cluster assembly factor nbp35 | elongation zone | 12 h |
| HORVU.MOREX.r3.6HG0570430.1 | 0.374683 | 5.175773 | 3.592767 | 0.000716 | 0.012358 | -0.74176 | Ubiquitin-conjugating enzyme E2 | elongation zone | 12 h |
| HORVU.MOREX.r3.3HG0288370.1 | -1.02207 | 2.377529 | -3.50667 | 0.000933 | 0.014176 | -0.74243 | Chaperone clpb, putative | elongation zone | 12 h |
| HORVU.MOREX.r3.3HG0262210.1 | -0.32843 | 5.498655 | -3.59766 | 0.000706 | 0.012262 | -0.74455 | Phosphatase 2C family protein | elongation zone | 12 h |
| HORVU.MOREX.r3.4HG0412140.1 | 1.102548 | 2.514643 | 3.568541 | 0.000772 | 0.012872 | -0.74522 | Zinc finger (C3HC4-type RING finger) protein family-like | elongation zone | 12 h |
| HORVU.MOREX.r3.5HG0524060.1 | 1.320151 | 3.909294 | 3.55735 | 0.000799 | 0.013094 | -0.74553 | Remorin | elongation zone | 12 h |
| HORVU.MOREX.r3.2HG0172400.1 | -1.75025 | 5.128092 | -3.5214 | 0.000892 | 0.013872 | -0.74644 | Glycosyltransferase | elongation zone | 12 h |
| HORVU.MOREX.r3.2HG0171810.1 | -0.5188 | 3.837575 | -3.51605 | 0.000906 | 0.013962 | -0.74715 | DUF1644 family protein | elongation zone | 12 h |
| HORVU.MOREX.r3.6HG0601670.1 | -0.9144 | 2.45038 | -3.49908 | 0.000954 | 0.014386 | -0.74785 | Phosphatidic acid phosphatase | elongation zone | 12 h |
| HORVU.MOREX.r3.1HG0017720.1 | -0.7434 | 7.063635 | -3.62126 | 0.000656 | 0.011718 | -0.74834 | Glutathione S-transferase | elongation zone | 12 h |
| HORVU.MOREX.r3.2HG0152140.1 | -0.66342 | 5.200814 | -3.53024 | 0.000868 | 0.013618 | -0.75053 | D-amino acid dehydrogenase | elongation zone | 12 h |
| HORVU.MOREX.r3.4HG0334600.1 | 0.605566 | 2.850253 | 3.494527 | 0.000968 | 0.014469 | -0.75257 | Magnesium transporter MRS2-like protein | elongation zone | 12 h |
| HORVU.MOREX.r3.2HG0186480.1 | 2.160718 | -0.52788 | 3.494388 | 0.000968 | 0.014469 | -0.75289 | 1-aminocyclopropane-1-carboxylate synthase | elongation zone | 12 h |
| HORVU.MOREX.r3.1HG0049840.1 | -0.71647 | 3.999244 | -3.49784 | 0.000958 | 0.0144 | -0.75411 | protein kinase family protein | elongation zone | 12 h |
| HORVU.MOREX.r3.6HG0574510.1 | -1.53475 | 7.321758 | -3.56425 | 0.000782 | 0.012985 | -0.75538 | #N/A | elongation zone | 12 h |
| HORVU.MOREX.r3.4HG0375760.1 | -0.77233 | 3.175902 | -3.52006 | 0.000895 | 0.013908 | -0.75578 | Oxidoreductase/ transition metal ion binding protein | elongation zone | 12 h |
| HORVU.MOREX.r3.6HG0589860.1 | 0.792761 | 4.121931 | 3.546392 | 0.000826 | 0.013244 | -0.75657 | Kinase family protein | elongation zone | 12 h |
| HORVU.MOREX.r3.7HG0642940.1 | 1.860453 | -1.68507 | 3.497425 | 0.000959 | 0.0144 | -0.75664 | Tryptophan decarboxylase | elongation zone | 12 h |
| HORVU.MOREX.r3.7HG0741650.1 | -0.83815 | 6.190632 | -3.57654 | 0.000753 | 0.012678 | -0.7589 | Histone H2A | elongation zone | 12 h |
| HORVU.MOREX.r3.5HG0493070.1 | -0.43426 | 6.006758 | -3.60419 | 0.000692 | 0.012162 | -0.76005 | Peptidylprolyl isomerase | elongation zone | 12 h |
| HORVU.MOREX.r3.4HG0356860.1 | 0.938395 | 1.759753 | 3.494598 | 0.000968 | 0.014469 | -0.76248 | Ulp1 protease family, C-terminal catalytic domain containing protein, expressed | elongation zone | 12 h |

**Table S1** Continued.

| ID | logFC | AveExpr | t | P.Value | adj.P.Val | B | gene description | root zone | time point |
|---|---|---|---|---|---|---|---|---|---|
| HORVU.MOREX.r3.2HG0145360.1 | 1.040134 | 3.483304 | 3.687049 | 0.000535 | 0.010392 | -0.7629 | Peroxidase | elongation zone | 12 h |
| HORVU.MOREX.r3.4HG0361420.2 | -0.4902 | 5.485836 | -3.55363 | 0.000808 | 0.01314 | -0.76292 | ATP dependent RNA helicase | elongation zone | 12 h |
| HORVU.MOREX.r3.7HG0727650.1 | 2.627481 | -1.35022 | 3.734055 | 0.000462 | 0.009584 | -0.76329 | BTB/POZ domain-containing protein | elongation zone | 12 h |
| HORVU.MOREX.r3.3HG0253360.1 | -1.0775 | 1.395191 | -3.51498 | 0.000909 | 0.013987 | -0.76599 | F-box domain containing protein | elongation zone | 12 h |
| HORVU.MOREX.r3.4HG0335150.1 | -0.30934 | 8.47371 | -3.67639 | 0.000553 | 0.010558 | -0.76616 | Nuclease domain-containing protein 1 | elongation zone | 12 h |
| HORVU.MOREX.r3.7HG0708500.1 | -0.90047 | 2.950817 | -3.48915 | 0.000984 | 0.014606 | -0.76664 | Chloride channel protein | elongation zone | 12 h |
| HORVU.MOREX.r3.5HG0466440.1 | -0.39841 | 4.963631 | -3.5667 | 0.000776 | 0.012908 | -0.77124 | Glucan endo-1,3-beta-glucosidase, putative | elongation zone | 12 h |
| HORVU.MOREX.r3.4HG0403250.1 | -1.08069 | 4.254713 | -3.49054 | 0.00098 | 0.014565 | -0.77392 | 50S ribosomal L32 | elongation zone | 12 h |
| HORVU.MOREX.r3.1HG0032930.1 | 0.67474 | 5.503727 | 3.624031 | 0.000651 | 0.011668 | -0.77472 | lipase, putative (DUF620) | elongation zone | 12 h |
| HORVU.MOREX.r3.7HG0649240.1 | 1.54596 | -2.99572 | 3.681241 | 0.000545 | 0.010463 | -0.77523 | protein kinase family protein | elongation zone | 12 h |
| HORVU.MOREX.r3.7HG0667540.1 | 0.627999 | 6.898857 | 3.702772 | 0.000509 | 0.010183 | -0.77702 | Pollen-specific protein SF3 | elongation zone | 12 h |
| HORVU.MOREX.r3.4HG0400840.1 | 0.906314 | 4.455067 | 3.582888 | 0.000739 | 0.012516 | -0.77756 | 4-coumarate--CoA ligase family protein | elongation zone | 12 h |
| HORVU.MOREX.r3.2HG0132700.1 | -1.79534 | 1.569987 | -3.50223 | 0.000945 | 0.014306 | -0.77808 | Phenylalanine ammonia-lyase | elongation zone | 12 h |
| HORVU.MOREX.r3.2HG0164700.2 | -0.67274 | 5.234489 | -3.55173 | 0.000813 | 0.013153 | -0.77812 | Potassium transporter | elongation zone | 12 h |
| HORVU.MOREX.r3.7HG0674480.1 | -0.2996 | 6.371176 | -3.60579 | 0.000688 | 0.01213 | -0.77841 | RING/U-box superfamily protein | elongation zone | 12 h |
| HORVU.MOREX.r3.7HG0663180.1 | -1.00667 | 1.342669 | -3.49131 | 0.000977 | 0.014552 | -0.78005 | Growth-regulating factor | elongation zone | 12 h |
| HORVU.MOREX.r3.2HG0173340.1 | -3.40117 | 1.078294 | -4.13056 | 0.000129 | 0.005458 | -0.78042 | Pentatricopeptide repeat-containing protein, putative | elongation zone | 12 h |
| HORVU.MOREX.r3.2HG0195110.2 | -1.87546 | 0.674913 | -3.69891 | 0.000515 | 0.010244 | -0.78085 | Remorin family protein | elongation zone | 12 h |
| HORVU.MOREX.r3.5HG0470680.1 | 1.343119 | -1.96407 | 3.484063 | 0.000999 | 0.014726 | -0.78107 | Heavy metal transport/detoxification protein | elongation zone | 12 h |
| HORVU.MOREX.r3.1HG0037910.1 | -0.69128 | 4.060046 | -3.49894 | 0.000955 | 0.014386 | -0.7812 | Starch synthase, chloroplastic/amyloplastic | elongation zone | 12 h |
| HORVU.MOREX.r3.6HG0608390.1 | 0.374186 | 8.330662 | 3.688327 | 0.000533 | 0.010373 | -0.78261 | Nuclear matrix constituent protein 1 | elongation zone | 12 h |
| HORVU.MOREX.r3.1HG0022860.1 | -0.35035 | 5.022173 | -3.56185 | 0.000788 | 0.013028 | -0.78379 | Xylosyltransferase 1 | elongation zone | 12 h |
| HORVU.MOREX.r3.3HG0319430.1 | 0.823866 | 2.262365 | 3.482422 | 0.001004 | 0.014726 | -0.78445 | WD40-repeat protein | elongation zone | 12 h |
| HORVU.MOREX.r3.5HG0516720.1 | -1.82664 | 6.439288 | -3.49136 | 0.000977 | 0.014552 | -0.78538 | Glutathione-S-transferase | elongation zone | 12 h |
| HORVU.MOREX.r3.3HG0316640.1 | -0.70542 | 4.162627 | -3.56778 | 0.000774 | 0.012886 | -0.78543 | Pentatricopeptide repeat (PPR) superfamily protein | elongation zone | 12 h |
| HORVU.MOREX.r3.4HG0349980.1 | 0.615035 | 0.310535 | 3.531143 | 0.000866 | 0.013618 | -0.78577 | DUF241 domain protein, putative (DUF241) | elongation zone | 12 h |
| HORVU.MOREX.r3.5HG0462220.1 | 0.943213 | 1.070356 | 3.545417 | 0.000829 | 0.013267 | -0.78656 | Calmodulin protein | elongation zone | 12 h |
| HORVU.MOREX.r3.4HG0355150.1 | -0.6149 | 5.515568 | -3.56842 | 0.000772 | 0.012872 | -0.79279 | Protein transport protein Sec61 subunit beta, putative | elongation zone | 12 h |
| HORVU.MOREX.r3.6HG0597770.1 | -0.50701 | 4.348024 | -3.51627 | 0.000906 | 0.013962 | -0.79292 | F-box/LRR protein | elongation zone | 12 h |
| HORVU.MOREX.r3.5HG0520650.1 | -0.67566 | 5.200623 | -3.55569 | 0.000803 | 0.013128 | -0.79386 | Argonaute family protein | elongation zone | 12 h |
| HORVU.MOREX.r3.3HG0240450.1 | -0.75347 | 4.298635 | -3.49168 | 0.000976 | 0.014552 | -0.79422 | Protein gamma response 1 | elongation zone | 12 h |
| HORVU.MOREX.r3.6HG0549520.1 | 0.634171 | 3.372434 | 3.516825 | 0.000904 | 0.013962 | -0.80234 | Beta-fructofuranosidase, insoluble protein | elongation zone | 12 h |
| HORVU.MOREX.r3.3HG0277200.1 | -1.12157 | 6.184722 | -3.54787 | 0.000822 | 0.01322 | -0.80306 | Pyruvate kinase | elongation zone | 12 h |
| HORVU.MOREX.r3.2HG0195190.1 | -0.87082 | 4.061362 | -3.50389 | 0.000941 | 0.014244 | -0.80334 | Multidrug resistance protein MdtC | elongation zone | 12 h |
| HORVU.MOREX.r3.3HG0326330.1 | 1.0398 | 0.282326 | 3.553807 | 0.000808 | 0.01314 | -0.80341 | Rho GTPase-activating protein | elongation zone | 12 h |
| HORVU.MOREX.r3.2HG0205760.1 | -0.91344 | 5.328219 | -3.49832 | 0.000957 | 0.0144 | -0.80548 | Glutamyl-tRNA(Gln) amidotransferase subunit A | elongation zone | 12 h |
| HORVU.MOREX.r3.5HG0493080.1 | 0.601523 | 5.851163 | 3.56725 | 0.000775 | 0.012897 | -0.80667 | VWA domain containing protein | elongation zone | 12 h |
| HORVU.MOREX.r3.2HG0178480.1 | 0.946154 | -0.05624 | 3.518066 | 0.000901 | 0.013961 | -0.80742 | RING/U-box superfamily protein | elongation zone | 12 h |
| HORVU.MOREX.r3.4HG0364310.1 | -0.51374 | 4.81568 | -3.51176 | 0.000918 | 0.014073 | -0.80804 | Protein SUPPRESSOR OF GENE SILENCING 3 | elongation zone | 12 h |
| HORVU.MOREX.r3.7HG0728870.1 | -0.59884 | 6.913212 | -3.68965 | 0.00053 | 0.01036 | -0.80931 | Xyloglucan endotransglucosylase/hydrolase | elongation zone | 12 h |
| HORVU.MOREX.r3.6HG0605290.1 | -0.6543 | 4.144726 | -3.47923 | 0.001014 | 0.014816 | -0.80977 | Protein phosphatase 2C | elongation zone | 12 h |

**Table S1** Continued.

| ID | logFC | AveExpr | t | P.Value | adj.P.Val | B | gene description | root zone | time point |
|---|---|---|---|---|---|---|---|---|---|
| HORVU.MOREX.r3.6HG0610700.1 | -0.90684 | 3.726004 | -3.5107 | 0.000921 | 0.014087 | -0.81128 | Blue copper protein | elongation zone | 12 h |
| HORVU.MOREX.r3.7HG0714900.1 | -0.72309 | 4.447401 | -3.49205 | 0.000975 | 0.014552 | -0.81828 | S-acyltransferase | elongation zone | 12 h |
| HORVU.MOREX.r3.3HG0234000.1 | 1.542139 | -0.27443 | 3.563474 | 0.000784 | 0.012995 | -0.81846 | Xyloglucan endotransglucosylase/hydrolase | elongation zone | 12 h |
| HORVU.MOREX.r3.1HG0030090.1 | -1.09987 | 1.881613 | -3.4833 | 0.001001 | 0.014726 | -0.81915 | Leucine-rich repeat protein kinase family protein | elongation zone | 12 h |
| HORVU.MOREX.r3.5HG0503950.1 | -0.48232 | 4.442886 | -3.50611 | 0.000934 | 0.014181 | -0.82028 | Kinase family protein | elongation zone | 12 h |
| HORVU.MOREX.r3.4HG0410830.1 | 1.089344 | 1.674186 | 3.544583 | 0.000831 | 0.013276 | -0.82121 | Transcription factor | elongation zone | 12 h |
| HORVU.MOREX.r3.6HG0539590.1 | 0.983647 | 3.210813 | 3.530163 | 0.000868 | 0.013618 | -0.82461 | Protein disulfide-isomerase | elongation zone | 12 h |
| HORVU.MOREX.r3.3HG0249640.1 | 0.623515 | 4.297068 | 3.560118 | 0.000792 | 0.013047 | -0.83302 | E3 ubiquitin-protein ligase | elongation zone | 12 h |
| HORVU.MOREX.r3.5HG0432950.1 | 0.451544 | 5.320519 | 3.551868 | 0.000812 | 0.013153 | -0.83577 | L-galactose dehydrogenase | elongation zone | 12 h |
| HORVU.MOREX.r3.1HG0086390.1 | 0.89146 | -0.26089 | 3.489637 | 0.000982 | 0.014595 | -0.83637 | #N/A | elongation zone | 12 h |
| HORVU.MOREX.r3.1HG0005840.1 | -1.42188 | 1.736344 | -3.59147 | 0.000719 | 0.012373 | -0.83709 | Embryo defective 2735 | elongation zone | 12 h |
| HORVU.MOREX.r3.5HG0510060.1 | -0.65642 | 7.466645 | -3.59092 | 0.000721 | 0.012383 | -0.83862 | Histone H2B | elongation zone | 12 h |
| HORVU.MOREX.r3.6HG0604620.1 | -1.34418 | 6.017915 | -3.51445 | 0.000911 | 0.013989 | -0.84237 | #N/A | elongation zone | 12 h |
| HORVU.MOREX.r3.5HG0486070.1 | -0.64751 | 6.538472 | -3.55259 | 0.000811 | 0.013149 | -0.84375 | Aminotransferase | elongation zone | 12 h |
| HORVU.MOREX.r3.7HG0715170.1 | 1.530379 | -0.05365 | 3.463691 | 0.001063 | 0.015289 | -0.8438 | Glycosyltransferase | elongation zone | 12 h |
| HORVU.MOREX.r3.5HG0514100.1 | -0.62927 | 4.963873 | -3.49799 | 0.000958 | 0.0144 | -0.84535 | #N/A | elongation zone | 12 h |
| HORVU.MOREX.r3.1HG0060260.1 | -0.97756 | 5.874721 | -3.52813 | 0.000874 | 0.013663 | -0.84628 | Histone H2B | elongation zone | 12 h |
| HORVU.MOREX.r3.2HG0215060.1 | 2.22913 | -1.59998 | 3.574152 | 0.000759 | 0.01274 | -0.84643 | WAT1-related protein | elongation zone | 12 h |
| HORVU.MOREX.r3.6HG0568900.1 | -1.11865 | 6.529769 | -3.53831 | 0.000847 | 0.013429 | -0.84667 | Histone H3 | elongation zone | 12 h |
| HORVU.MOREX.r3.2HG0105980.1 | -0.67326 | 2.766067 | -3.45831 | 0.00108 | 0.015382 | -0.84694 | #N/A | elongation zone | 12 h |
| HORVU.MOREX.r3.6HG0550950.1 | -0.48676 | 7.177294 | -3.58609 | 0.000731 | 0.012454 | -0.84931 | NADP dependent sorbitol 6-phosphate dehydrogenase family protein | elongation zone | 12 h |
| HORVU.MOREX.r3.6HG0573850.1 | -0.32853 | 5.097663 | -3.5447 | 0.00083 | 0.013276 | -0.852 | Protein YIF1B | elongation zone | 12 h |
| HORVU.MOREX.r3.2HG0136010.1 | -0.71292 | 2.505497 | -3.46064 | 0.001072 | 0.015346 | -0.854 | DNA repair family protein | elongation zone | 12 h |
| HORVU.MOREX.r3.5HG0524120.1 | -0.3915 | 5.963425 | -3.57496 | 0.000757 | 0.012729 | -0.85445 | #N/A | elongation zone | 12 h |
| HORVU.MOREX.r3.2HG0189800.1 | -0.68678 | 3.770835 | -3.4838 | 0.001 | 0.014726 | -0.85511 | ethylene-responsive transcription factor | elongation zone | 12 h |
| HORVU.MOREX.r3.3HG0277100.1 | -0.46209 | 4.350493 | -3.51088 | 0.000921 | 0.014087 | -0.85578 | Calcium-dependent lipid-binding (CaLB domain) family protein | elongation zone | 12 h |
| HORVU.MOREX.r3.1HG0053720.1 | -0.90305 | 1.572526 | -3.46748 | 0.00105 | 0.015148 | -0.85699 | Protein kinase family protein | elongation zone | 12 h |
| HORVU.MOREX.r3.6HG0617730.1 | -0.35964 | 7.183954 | -3.63239 | 0.000634 | 0.01149 | -0.85734 | Galactoside 2-alpha-L-fucosyltransferase | elongation zone | 12 h |
| HORVU.MOREX.r3.1HG0091770.1 | -1.21309 | 4.911247 | -3.48479 | 0.000997 | 0.014726 | -0.85938 | Histone H2B | elongation zone | 12 h |
| HORVU.MOREX.r3.6HG0601180.1 | 0.918442 | 3.243543 | 3.645766 | 0.000608 | 0.011202 | -0.8614 | Expansin protein | elongation zone | 12 h |
| HORVU.MOREX.r3.7HG0730660.1 | 1.50246 | -2.73673 | 3.530432 | 0.000867 | 0.013618 | -0.86439 | #N/A | elongation zone | 12 h |
| HORVU.MOREX.r3.1HG0075590.1 | -0.27657 | 8.230864 | -3.6482 | 0.000604 | 0.011167 | -0.86511 | Coiled-coil domain-containing protein 47 | elongation zone | 12 h |
| HORVU.MOREX.r3.4HG0335110.1 | -0.9457 | 5.132212 | -3.48246 | 0.001004 | 0.014726 | -0.8654 | BHLH family protein, putative, expressed | elongation zone | 12 h |
| HORVU.MOREX.r3.5HG0462380.1 | -0.63423 | 3.998698 | -3.53134 | 0.000865 | 0.013618 | -0.86576 | LYR family of Fe/S cluster biogenesis protein | elongation zone | 12 h |
| HORVU.MOREX.r3.3HG0302090.1 | -0.58639 | 3.520126 | -3.4596 | 0.001076 | 0.015372 | -0.86613 | CASP-like protein | elongation zone | 12 h |
| HORVU.MOREX.r3.5HG0504700.1 | -0.91605 | 3.74855 | -3.47709 | 0.00102 | 0.014881 | -0.86652 | Acetyl-coenzyme A synthetase | elongation zone | 12 h |
| HORVU.MOREX.r3.5HG0502130.1 | -0.77611 | 6.54341 | -3.52979 | 0.000869 | 0.013618 | -0.86718 | T-complex protein 1 subunit beta | elongation zone | 12 h |
| HORVU.MOREX.r3.6HG0557140.1 | -0.64475 | 3.245091 | -3.4603 | 0.001074 | 0.015351 | -0.86901 | Myb family transcription factor-like | elongation zone | 12 h |
| HORVU.MOREX.r3.5HG0478050.1 | 0.607932 | 5.803016 | 3.613842 | 0.000671 | 0.011927 | -0.87005 | Nucleobase ascorbate transporter | elongation zone | 12 h |
| HORVU.MOREX.r3.7HG0737340.1 | 1.392017 | -0.10666 | 3.559862 | 0.000793 | 0.013047 | -0.87148 | GDA1/CD39 nucleoside phosphatase family protein | elongation zone | 12 h |
| HORVU.MOREX.r3.3HG0284100.1 | 0.437751 | 6.109021 | 3.593535 | 0.000715 | 0.012356 | -0.87177 | 5'-nucleotidase SurE | elongation zone | 12 h |

**Table S1** Continued.

| ID | logFC | AveExpr | t | P.Value | adj.P.Val | B | gene description | root zone | time point |
|---|---|---|---|---|---|---|---|---|---|
| HORVU.MOREX.r3.7HG0705860.1 | 0.476177 | 7.982366 | 3.678827 | 0.000549 | 0.010519 | -0.87219 | Non-lysosomal glucosylceramidase | elongation zone | 12 h |
| HORVU.MOREX.r3.3HG0296490.1 | -0.631 | 6.526217 | -3.56309 | 0.000785 | 0.013 | -0.87223 | NADH-cytochrome b5 reductase | elongation zone | 12 h |
| HORVU.MOREX.r3.5HG0420480.1 | 1.979522 | 1.423525 | 3.448414 | 0.001113 | 0.015672 | -0.87256 | BEL1-like homeodomain protein 6 | elongation zone | 12 h |
| HORVU.MOREX.r3.4HG0331730.1 | -0.30634 | 6.082061 | -3.55804 | 0.000797 | 0.013094 | -0.87951 | Zinc finger, C2H2 | elongation zone | 12 h |
| HORVU.MOREX.r3.5HG0525980.1 | -0.44145 | 8.509545 | -3.64399 | 0.000611 | 0.011213 | -0.88 | Dihydrolipoamide acetyltransferase component of pyruvate dehydrogenase comp | elongation zone | 12 h |
| HORVU.MOREX.r3.7HG0665990.1 | 0.624684 | 6.156682 | 3.629205 | 0.00064 | 0.011564 | -0.88211 | Shikimate kinase | elongation zone | 12 h |
| HORVU.MOREX.r3.1HG0007360.1 | -0.54863 | 5.263881 | -3.56081 | 0.00079 | 0.013047 | -0.88285 | Type I inositol-1,4,5-trisphosphate 5-phosphatase 1 | elongation zone | 12 h |
| HORVU.MOREX.r3.7HG0736860.1 | -2.17186 | 3.347913 | -3.5357 | 0.000854 | 0.013502 | -0.88359 | Aspartyl/glutamyl-tRNA(Asn/Gln) amidotransferase subunit B, putative isoform 2 | elongation zone | 12 h |
| HORVU.MOREX.r3.2HG0128490.1 | -0.32614 | 4.434925 | -3.52034 | 0.000895 | 0.013907 | -0.88373 | #N/A | elongation zone | 12 h |
| HORVU.MOREX.r3.7HG0674990.1 | -0.50001 | 3.286295 | -3.4631 | 0.001065 | 0.015297 | -0.88381 | Mannan endo-1,4-beta-mannosidase-like protein | elongation zone | 12 h |
| HORVU.MOREX.r3.5HG0496220.1 | -1.19608 | 4.965587 | -3.47561 | 0.001025 | 0.014916 | -0.88425 | #N/A | elongation zone | 12 h |
| HORVU.MOREX.r3.5HG0428840.1 | -0.73186 | 5.358224 | -3.48337 | 0.001001 | 0.014726 | -0.88884 | BTB/POZ domain-containing protein | elongation zone | 12 h |
| HORVU.MOREX.r3.7HG0712650.1 | -0.85576 | 3.870701 | -3.44688 | 0.001118 | 0.015713 | -0.8952 | DnaJ subfamily C member 2 | elongation zone | 12 h |
| HORVU.MOREX.r3.6HG0624240.1 | 0.865736 | 1.910661 | 3.475242 | 0.001026 | 0.014922 | -0.89588 | Basic helix-loop-helix transcription factor | elongation zone | 12 h |
| HORVU.MOREX.r3.5HG0498770.1 | -0.41008 | 6.313633 | -3.55308 | 0.000809 | 0.01314 | -0.89618 | Hexosyltransferase | elongation zone | 12 h |
| HORVU.MOREX.r3.6HG0618100.1 | -1.83706 | 2.509582 | -3.48519 | 0.000996 | 0.014726 | -0.89838 | Protein trichome birefringence | elongation zone | 12 h |
| HORVU.MOREX.r3.3HG0243560.1 | -0.4069 | 5.438018 | -3.51583 | 0.000907 | 0.013962 | -0.89896 | BES1/BZR1 homolog 1 | elongation zone | 12 h |
| HORVU.MOREX.r3.2HG0209440.1 | -0.48026 | 5.469564 | -3.52567 | 0.00088 | 0.013745 | -0.90553 | Sugar transporter, putative | elongation zone | 12 h |
| HORVU.MOREX.r3.3HG0298180.1 | 1.503506 | -1.34466 | 3.435325 | 0.001158 | 0.01605 | -0.90763 | #N/A | elongation zone | 12 h |
| HORVU.MOREX.r3.3HG0255580.1 | 0.489729 | 5.497552 | 3.579479 | 0.000746 | 0.012595 | -0.90778 | Respiratory burst oxidase, putative | elongation zone | 12 h |
| HORVU.MOREX.r3.3HG0322660.1 | -1.31269 | 5.027667 | -3.44748 | 0.001116 | 0.015695 | -0.90833 | Receptor protein kinase, putative | elongation zone | 12 h |
| HORVU.MOREX.r3.5HG0494470.1 | -0.5211 | 3.993787 | -3.45668 | 0.001085 | 0.015413 | -0.90936 | Transmembrane emp24 domain-containing protein | elongation zone | 12 h |
| HORVU.MOREX.r3.7HG0655180.1 | -0.53486 | 3.708393 | -3.45054 | 0.001106 | 0.015604 | -0.90981 | F-box protein | elongation zone | 12 h |
| HORVU.MOREX.r3.1HG0003430.1 | 0.501143 | 4.894308 | 3.52258 | 0.000889 | 0.013844 | -0.91151 | TBC1 domain family member | elongation zone | 12 h |
| HORVU.MOREX.r3.4HG0415590.1 | -1.60417 | -0.04239 | -3.43533 | 0.001158 | 0.01605 | -0.91174 | Bifunctional inhibitor/lipid-transfer protein/seed storage 2S albumin superfamily p | elongation zone | 12 h |
| HORVU.MOREX.r3.7HG0689580.1 | -0.60683 | 4.115082 | -3.44808 | 0.001114 | 0.015677 | -0.91199 | Signal peptide peptidase family protein | elongation zone | 12 h |
| HORVU.MOREX.r3.4HG0379960.1 | -0.42667 | 5.175068 | -3.50568 | 0.000935 | 0.014187 | -0.91247 | Exostosin family protein | elongation zone | 12 h |
| HORVU.MOREX.r3.5HG0495790.1 | -0.63472 | 4.487486 | -3.46516 | 0.001058 | 0.015244 | -0.91308 | D-amino acid dehydrogenase | elongation zone | 12 h |
| HORVU.MOREX.r3.1HG0015960.1 | 1.206401 | -0.2749 | 3.432282 | 0.001168 | 0.016079 | -0.91409 | Protein kinase family protein | elongation zone | 12 h |
| HORVU.MOREX.r3.2HG0136570.1 | -2.14977 | 1.302768 | -3.54033 | 0.000842 | 0.013377 | -0.91516 | Cellulose synthase-like protein | elongation zone | 12 h |
| HORVU.MOREX.r3.4HG0379290.1 | -0.31983 | 5.28798 | -3.51719 | 0.000903 | 0.013962 | -0.91611 | COP1-interacting-like protein, putative | elongation zone | 12 h |
| HORVU.MOREX.r3.6HG0564590.1 | -0.48695 | 6.44927 | -3.53334 | 0.00086 | 0.013572 | -0.91639 | O-fucosyltransferase family protein | elongation zone | 12 h |
| HORVU.MOREX.r3.3HG0307250.1 | 1.74247 | -2.48075 | 3.474127 | 0.00103 | 0.014951 | -0.9164 | Amino acid permease | elongation zone | 12 h |
| HORVU.MOREX.r3.1HG0006610.1 | -0.95894 | 8.69333 | -3.58972 | 0.000723 | 0.012401 | -0.91966 | Histone H2A | elongation zone | 12 h |
| HORVU.MOREX.r3.3HG0252610.1 | -0.37714 | 7.233439 | -3.56514 | 0.00078 | 0.01296 | -0.92076 | Protein kinase | elongation zone | 12 h |
| HORVU.MOREX.r3.7HG0637030.1 | 0.971635 | 0.131716 | 3.475625 | 0.001025 | 0.014916 | -0.92188 | Cytochrome P450 family protein, expressed | elongation zone | 12 h |
| HORVU.MOREX.r3.5HG0523560.1 | -0.87058 | 6.843578 | -3.55316 | 0.000809 | 0.01314 | -0.92358 | Histone H2A | elongation zone | 12 h |
| HORVU.MOREX.r3.3HG0330120.1 | 2.289758 | -3.00967 | 3.479617 | 0.001013 | 0.014809 | -0.92632 | O-methyltransferase | elongation zone | 12 h |
| HORVU.MOREX.r3.3HG0287840.1 | -0.70993 | 3.05117 | -3.43477 | 0.00116 | 0.016058 | -0.92686 | peroxisomal ABC transporter 1 | elongation zone | 12 h |
| HORVU.MOREX.r3.2HG0099780.1 | 1.103458 | -0.01779 | 3.42734 | 0.001186 | 0.016221 | -0.92693 | Maternal effect embryo arrest protein | elongation zone | 12 h |
| HORVU.MOREX.r3.4HG0356690.1 | 1.103072 | 0.745555 | 3.43352 | 0.001164 | 0.016068 | -0.92893 | Protein ZINC INDUCED FACILITATOR-LIKE 1 | elongation zone | 12 h |

**Table S1** Continued.

| ID | logFC | AveExpr | t | P.Value | adj.P.Val | B | gene description | root zone | time point |
|---|---|---|---|---|---|---|---|---|---|
| HORVU.MOREX.r3.3HG0235030.1 | -0.50211 | 3.06592 | -3.43204 | 0.001169 | 0.016079 | -0.93102 | Dehydration-responsive element binding protein | elongation zone | 12 h |
| HORVU.MOREX.r3.5HG0525440.1 | -0.37273 | 6.964938 | -3.58046 | 0.000744 | 0.012578 | -0.93217 | Trichome birefringence-like protein | elongation zone | 12 h |
| HORVU.MOREX.r3.5HG0512230.1 | 1.183108 | 2.04829 | 3.425771 | 0.001191 | 0.016287 | -0.93298 | Protein ENHANCED DISEASE RESISTANCE 2-like | elongation zone | 12 h |
| HORVU.MOREX.r3.4HG0358880.2 | -0.52281 | 5.351205 | -3.51931 | 0.000897 | 0.01393 | -0.93563 | Succinate--CoA ligase [ADP-forming] subunit beta | elongation zone | 12 h |
| HORVU.MOREX.r3.2HG0211880.1 | -0.46183 | 4.077958 | -3.46207 | 0.001068 | 0.015301 | -0.93577 | Exostosin family protein | elongation zone | 12 h |
| HORVU.MOREX.r3.1HG0095000.1 | -1.58078 | 0.995323 | -3.49102 | 0.000978 | 0.014555 | -0.93582 | S-adenosyl-L-methionine-dependent methyltransferases superfamily protein | elongation zone | 12 h |
| HORVU.MOREX.r3.2HG0111570.1 | 0.736155 | 1.289254 | 3.427838 | 0.001184 | 0.016219 | -0.93916 | Polynucleotidyl transferase, ribonuclease H-like superfamily protein | elongation zone | 12 h |
| HORVU.MOREX.r3.3HG0224440.1 | 0.867778 | 1.43625 | 3.48481 | 0.000997 | 0.014726 | -0.9411 | DUF4228 domain protein | elongation zone | 12 h |
| HORVU.MOREX.r3.5HG0477800.1 | 0.747897 | 5.596819 | 3.548386 | 0.000821 | 0.01322 | -0.94315 | Endo-1,3(4)-beta-glucanase 1 | elongation zone | 12 h |
| HORVU.MOREX.r3.5HG0510920.1 | -0.38934 | 5.904242 | -3.50607 | 0.000934 | 0.014181 | -0.94597 | Transcriptional adapter ADA2 | elongation zone | 12 h |
| HORVU.MOREX.r3.2HG0194850.1 | 0.684492 | 6.889643 | 3.594903 | 0.000712 | 0.012335 | -0.94691 | Cysteine/Histidine-rich C1 domain family protein | elongation zone | 12 h |
| HORVU.MOREX.r3.7HG0726800.1 | 1.133392 | 2.279935 | 3.434718 | 0.00116 | 0.016058 | -0.94699 | 3,4-dihydroxy-2-butanone 4-phosphate synthase | elongation zone | 12 h |
| HORVU.MOREX.r3.4HG0331850.1 | -0.60822 | 3.521487 | -3.43982 | 0.001142 | 0.015898 | -0.95072 | Sn1-specific diacylglycerol lipase alpha | elongation zone | 12 h |
| HORVU.MOREX.r3.7HG0751110.1 | -0.61381 | 4.358149 | -3.45778 | 0.001082 | 0.015394 | -0.95118 | Kinesin-like | elongation zone | 12 h |
| HORVU.MOREX.r3.3HG0267150.1 | -0.35217 | 5.178731 | -3.49654 | 0.000962 | 0.014417 | -0.95301 | Calcium binding protein-like | elongation zone | 12 h |
| HORVU.MOREX.r3.5HG0509020.1 | 1.013104 | 0.642934 | 3.419563 | 0.001214 | 0.016441 | -0.9552 | guanine nucleotide-binding protein subunit gamma | elongation zone | 12 h |
| HORVU.MOREX.r3.3HG0314820.1 | 0.703931 | 3.291172 | 3.522295 | 0.000889 | 0.013845 | -0.95819 | 26S proteasome regulatory subunit RPN2 | elongation zone | 12 h |
| HORVU.MOREX.r3.5HG0525430.1 | 0.745961 | 4.268757 | 3.551468 | 0.000813 | 0.013153 | -0.95917 | Trichome birefringence-like protein | elongation zone | 12 h |
| HORVU.MOREX.r3.1HG0091670.1 | -0.56351 | 4.229991 | -3.43262 | 0.001167 | 0.016079 | -0.96454 | Peroxisomal membrane PEX14-like protein | elongation zone | 12 h |
| HORVU.MOREX.r3.3HG0284170.1 | -0.57584 | 4.134399 | -3.51593 | 0.000907 | 0.013962 | -0.96677 | cotton fiber-like protein (DUF761) | elongation zone | 12 h |
| HORVU.MOREX.r3.3HG0275490.1 | 1.707844 | -2.47526 | 3.444522 | 0.001126 | 0.01576 | -0.96898 | Zinc finger CCCH domain protein | elongation zone | 12 h |
| HORVU.MOREX.r3.6HG0624650.1 | -0.35452 | 8.612308 | -3.63135 | 0.000636 | 0.011508 | -0.96929 | Transmembrane 9 superfamily member | elongation zone | 12 h |
| HORVU.MOREX.r3.7HG0668450.1 | -0.49651 | 5.247544 | -3.48298 | 0.001002 | 0.014726 | -0.97067 | Alpha-mannosidase | elongation zone | 12 h |
| HORVU.MOREX.r3.7HG0642550.1 | -0.32762 | 6.014924 | -3.51713 | 0.000903 | 0.013962 | -0.97563 | Peroxisomal membrane protein 11 homolog | elongation zone | 12 h |
| HORVU.MOREX.r3.5HG0527720.1 | 1.082186 | 0.951915 | 3.420705 | 0.00121 | 0.016416 | -0.97796 | Receptor protein kinase, putative,expressed | elongation zone | 12 h |
| HORVU.MOREX.r3.3HG0247260.1 | -1.16939 | 0.519017 | -3.47746 | 0.001019 | 0.014875 | -0.97855 | Leucine-rich repeat protein kinase family protein | elongation zone | 12 h |
| HORVU.MOREX.r3.2HG0195910.1 | 0.703522 | 2.102422 | 3.523828 | 0.000885 | 0.013812 | -0.9787 | Protease inhibitor/seed storage/lipid transfer family protein | elongation zone | 12 h |
| HORVU.MOREX.r3.5HG0439090.1 | -0.72701 | 2.651507 | -3.40873 | 0.001254 | 0.016818 | -0.97877 | ERD (Early-responsive to dehydration stress) family protein | elongation zone | 12 h |
| HORVU.MOREX.r3.7HG0727460.1 | -0.83413 | 8.013727 | -3.53864 | 0.000846 | 0.013426 | -0.981 | 60S acidic ribosomal protein P3 | elongation zone | 12 h |
| HORVU.MOREX.r3.1HG0092730.1 | 0.773468 | 7.867854 | 3.624833 | 0.000649 | 0.011649 | -0.986 | ATP-dependent zinc metalloprotease FtsH 1 | elongation zone | 12 h |
| HORVU.MOREX.r3.2HG0212900.1 | -0.61294 | 6.207725 | -3.48368 | 0.001 | 0.014726 | -0.98864 | Methylthioribose kinase | elongation zone | 12 h |
| HORVU.MOREX.r3.3HG0324480.1 | 1.356772 | -1.11625 | 3.431356 | 0.001172 | 0.016094 | -0.98963 | UDP-glycosyltransferase | elongation zone | 12 h |
| HORVU.MOREX.r3.3HG0245310.1 | -0.74754 | 4.920306 | -3.47834 | 0.001016 | 0.014846 | -0.99074 | GDSL esterase/lipase | elongation zone | 12 h |
| HORVU.MOREX.r3.1HG0084760.1 | 1.315812 | -3.22969 | 3.541218 | 0.000839 | 0.013372 | -0.99216 | Myb transcription factor | elongation zone | 12 h |
| HORVU.MOREX.r3.7HG0742070.1 | -0.65421 | 6.57434 | -3.49741 | 0.000959 | 0.0144 | -0.99389 | UDP-glucose 4-epimerase 1 | elongation zone | 12 h |
| HORVU.MOREX.r3.2HG0104080.1 | 1.006087 | -1.54865 | 3.401109 | 0.001283 | 0.01705 | -0.99409 | Glycosyltransferase | elongation zone | 12 h |
| HORVU.MOREX.r3.2HG0115300.2 | -0.28072 | 7.926726 | -3.57463 | 0.000758 | 0.012732 | -0.99726 | Superoxide dismutase [Cu-Zn] | elongation zone | 12 h |
| HORVU.MOREX.r3.7HG0654680.1 | -0.93333 | 5.771912 | -3.43324 | 0.001165 | 0.016068 | -1.00165 | Starch synthase, chloroplastic/amyloplastic | elongation zone | 12 h |
| HORVU.MOREX.r3.5HG0488050.1 | 1.446708 | -2.9262 | 3.501809 | 0.000947 | 0.014313 | -1.00374 | Blue copper protein | elongation zone | 12 h |
| HORVU.MOREX.r3.7HG0676690.1 | 0.816004 | 4.543618 | 3.601768 | 0.000697 | 0.012162 | -1.00438 | Receptor-like kinase | elongation zone | 12 h |
| HORVU.MOREX.r3.3HG0302910.1 | -0.9604 | 7.267731 | -3.51097 | 0.000921 | 0.014087 | -1.01003 | Histone H2B | elongation zone | 12 h |

**Table S1** Continued.

| ID | logFC | AveExpr | t | P.Value | adj.P.Val | B | gene description | root zone | time point |
|---|---|---|---|---|---|---|---|---|---|
| HORVU.MOREX.r3.3HG0252960.1 | 1.299096 | -1.0236 | 3.399728 | 0.001288 | 0.017063 | -1.0106 | Calmodulin-binding protein | elongation zone | 12 h |
| HORVU.MOREX.r3.3HG0234560.1 | -1.34087 | 2.285408 | -3.44296 | 0.001131 | 0.015774 | -1.01265 | Ethylene receptor 2 | elongation zone | 12 h |
| HORVU.MOREX.r3.3HG0328480.1 | -1.73065 | 1.933392 | -3.51224 | 0.000917 | 0.014073 | -1.01588 | Rotundifolia-like protein | elongation zone | 12 h |
| HORVU.MOREX.r3.4HG0406140.1 | 0.48942 | 5.424725 | 3.497041 | 0.00096 | 0.014406 | -1.0164 | Transmembrane protein 131 | elongation zone | 12 h |
| HORVU.MOREX.r3.4HG0402000.1 | -0.63812 | 8.180685 | -3.57848 | 0.000749 | 0.012613 | -1.01895 | Kinase interacting (KIP1-like) family protein, putative | elongation zone | 12 h |
| HORVU.MOREX.r3.6HG0614500.1 | -0.38563 | 5.054555 | -3.46401 | 0.001062 | 0.015287 | -1.02196 | 2,3-bisphosphoglycerate-dependent phosphoglycerate mutase | elongation zone | 12 h |
| HORVU.MOREX.r3.7HG0640800.1 | 2.180269 | 0.469292 | 3.694259 | 0.000523 | 0.010309 | -1.02229 | Cell division cycle protein 123 like | elongation zone | 12 h |
| HORVU.MOREX.r3.7HG0706860.1 | -0.64394 | 4.054763 | -3.39882 | 0.001292 | 0.017089 | -1.02275 | PHD finger family protein | elongation zone | 12 h |
| HORVU.MOREX.r3.3HG0254940.1 | 0.868714 | 2.451212 | 3.557318 | 0.000799 | 0.013094 | -1.02351 | Peroxidase | elongation zone | 12 h |
| HORVU.MOREX.r3.7HG0676000.1 | -1.24175 | 0.270183 | -3.4446 | 0.001126 | 0.01576 | -1.02532 | Jasmonate O-methyltransferase | elongation zone | 12 h |
| HORVU.MOREX.r3.2HG0115970.1 | -1.05766 | 3.355368 | -3.45353 | 0.001096 | 0.015528 | -1.02592 | NAD(P)H dehydrogenase (Quinone) | elongation zone | 12 h |
| HORVU.MOREX.r3.5HG0475250.1 | -0.69647 | 3.753494 | -3.39696 | 0.001299 | 0.017131 | -1.02729 | Translation initiation factor IF-2 | elongation zone | 12 h |
| HORVU.MOREX.r3.5HG0421460.1 | -0.68188 | 11.22444 | -3.66956 | 0.000565 | 0.010652 | -1.02788 | RNA-binding protein | elongation zone | 12 h |
| HORVU.MOREX.r3.6HG0566940.1 | 0.58032 | 4.632192 | 3.463486 | 0.001063 | 0.015289 | -1.02831 | Sulfurtransferase | elongation zone | 12 h |
| HORVU.MOREX.r3.7HG0652830.1 | -1.91475 | 0.863491 | -3.6596 | 0.000583 | 0.010887 | -1.029 | Disease resistance protein RPP13 | elongation zone | 12 h |
| HORVU.MOREX.r3.5HG0444350.1 | 0.824653 | 2.597523 | 3.398644 | 0.001292 | 0.017089 | -1.02962 | Sn1-specific diacylglycerol lipase alpha | elongation zone | 12 h |
| HORVU.MOREX.r3.7HG0669590.1 | 1.389504 | -0.99729 | 3.400606 | 0.001285 | 0.01705 | -1.03221 | Caleosin | elongation zone | 12 h |
| HORVU.MOREX.r3.7HG0739030.1 | 0.549526 | 5.253412 | 3.508076 | 0.000929 | 0.014168 | -1.03378 | Kinase family protein | elongation zone | 12 h |
| HORVU.MOREX.r3.4HG0352200.1 | 0.687494 | 3.627838 | 3.446324 | 0.00112 | 0.015728 | -1.03446 | Methyltransferase-related protein, putative | elongation zone | 12 h |
| HORVU.MOREX.r3.1HG0069030.1 | 1.513426 | 1.48038 | 3.388332 | 0.001333 | 0.017367 | -1.0349 | Calcium-transporting ATPase | elongation zone | 12 h |
| HORVU.MOREX.r3.1HG0075230.1 | -1.20937 | 6.583831 | -3.46261 | 0.001066 | 0.015301 | -1.04058 | Histone H3 | elongation zone | 12 h |
| HORVU.MOREX.r3.5HG0509970.1 | 1.778014 | -0.93022 | 3.418401 | 0.001218 | 0.016477 | -1.04279 | Heat shock transcription factor | elongation zone | 12 h |
| HORVU.MOREX.r3.7HG0716900.1 | -0.35492 | 8.192568 | -3.5702 | 0.000768 | 0.012843 | -1.04375 | Transmembrane 9 superfamily member | elongation zone | 12 h |
| HORVU.MOREX.r3.2HG0135650.1 | 2.770416 | -3.48765 | 3.605312 | 0.000689 | 0.01213 | -1.04376 | Laccase | elongation zone | 12 h |
| HORVU.MOREX.r3.2HG0135210.1 | 0.771811 | 2.447492 | 3.558142 | 0.000797 | 0.013094 | -1.04606 | Late embryogenesis abundant (LEA) hydroxyproline-rich glycoprotein family | elongation zone | 12 h |
| HORVU.MOREX.r3.3HG0323750.1 | 0.866479 | 2.450618 | 3.380858 | 0.001363 | 0.017603 | -1.04738 | bZIP transcription factor, putative (DUF630 and DUF632) | elongation zone | 12 h |
| HORVU.MOREX.r3.2HG0134990.1 | 0.685889 | 5.499939 | 3.44915 | 0.00111 | 0.015648 | -1.048 | Myosin-related family protein | elongation zone | 12 h |
| HORVU.MOREX.r3.4HG0382550.1 | 0.636612 | 5.636635 | 3.50115 | 0.000948 | 0.014332 | -1.0485 | Xyloglucan 6-xylosyltransferase 1 | elongation zone | 12 h |
| HORVU.MOREX.r3.1HG0087590.1 | 1.424365 | -1.00814 | 3.37951 | 0.001369 | 0.01763 | -1.04851 | transmembrane protein, putative (DUF679 domain membrane protein 2) | elongation zone | 12 h |
| HORVU.MOREX.r3.1HG0079920.1 | 0.651371 | 3.500385 | 3.467743 | 0.00105 | 0.015148 | -1.04895 | Disease resistance protein (NBS-LRR class) family | elongation zone | 12 h |
| HORVU.MOREX.r3.4HG0398050.1 | 1.595613 | -2.71986 | 3.437184 | 0.001151 | 0.015993 | -1.05051 | #N/A | elongation zone | 12 h |
| HORVU.MOREX.r3.4HG0381880.1 | -0.47828 | 5.392899 | -3.46244 | 0.001067 | 0.015301 | -1.05266 | Phytochrome | elongation zone | 12 h |
| HORVU.MOREX.r3.4HG0398910.1 | -1.01886 | 7.14368 | -3.49462 | 0.000967 | 0.014469 | -1.05291 | Histone H2B | elongation zone | 12 h |
| HORVU.MOREX.r3.2HG0205050.1 | 2.025911 | -2.49223 | 3.473297 | 0.001032 | 0.014978 | -1.05356 | Receptor-like protein kinase | elongation zone | 12 h |
| HORVU.MOREX.r3.5HG0522130.1 | -0.51327 | 4.373396 | -3.44312 | 0.001131 | 0.015774 | -1.05477 | #N/A | elongation zone | 12 h |
| HORVU.MOREX.r3.2HG0185850.1 | -0.4627 | 5.208449 | -3.46753 | 0.00105 | 0.015148 | -1.05613 | Expansin | elongation zone | 12 h |
| HORVU.MOREX.r3.4HG0338400.1 | -0.86073 | 8.453677 | -3.53775 | 0.000848 | 0.013441 | -1.05763 | Histone H2B | elongation zone | 12 h |
| HORVU.MOREX.r3.3HG0233150.1 | -0.82771 | 5.953619 | -3.45738 | 0.001083 | 0.015401 | -1.06234 | Histone H2B | elongation zone | 12 h |
| HORVU.MOREX.r3.6HG0608910.1 | 1.218664 | -1.72298 | 3.374192 | 0.001391 | 0.017856 | -1.06241 | Heavy metal transport/detoxification superfamily protein | elongation zone | 12 h |
| HORVU.MOREX.r3.6HG0547970.1 | -0.92694 | 5.20494 | -3.44288 | 0.001132 | 0.015774 | -1.0625 | #N/A | elongation zone | 12 h |
| HORVU.MOREX.r3.4HG0403260.1 | 0.346316 | 4.85718 | 3.458257 | 0.00108 | 0.015382 | -1.0652 | Ribulose-phosphate 3-epimerase | elongation zone | 12 h |

**Table S1** Continued.

| ID | logFC | AveExpr | t | P.Value | adj.P.Val | B | gene description | root zone | time point |
|---|---|---|---|---|---|---|---|---|---|
| HORVU.MOREX.r3.1HG0069380.1 | 0.817388 | 2.129643 | 3.433418 | 0.001164 | 0.016068 | -1.06624 | Cysteine proteinase inhibitor | elongation zone | 12 h |
| HORVU.MOREX.r3.1HG0020980.1 | 0.544452 | 7.252013 | 3.589018 | 0.000725 | 0.012414 | -1.06749 | Endo-1,31,4-beta-D-glucanase | elongation zone | 12 h |
| HORVU.MOREX.r3.6HG0564510.1 | -0.91871 | 4.576465 | -3.381 | 0.001363 | 0.017603 | -1.06769 | Arabinose 5-phosphate isomerase, putative | elongation zone | 12 h |
| HORVU.MOREX.r3.2HG0184230.1 | 2.393572 | -3.03591 | 3.507479 | 0.00093 | 0.014174 | -1.06813 | #N/A | elongation zone | 12 h |
| HORVU.MOREX.r3.3HG0223950.1 | -0.66356 | 4.288991 | -3.38873 | 0.001331 | 0.017357 | -1.06813 | #N/A | elongation zone | 12 h |
| HORVU.MOREX.r3.2HG0132440.2 | -0.45464 | 4.455297 | -3.40713 | 0.00126 | 0.016857 | -1.07039 | #N/A | elongation zone | 12 h |
| HORVU.MOREX.r3.5HG0513020.1 | -0.33397 | 6.516882 | -3.50869 | 0.000927 | 0.014152 | -1.07144 | Transmembrane protein 147 | elongation zone | 12 h |
| HORVU.MOREX.r3.7HG0709720.1 | -0.46542 | 5.925323 | -3.51831 | 0.0009 | 0.013961 | -1.0729 | plant/protein (Protein of unknown function, DUF538) | elongation zone | 12 h |
| HORVU.MOREX.r3.1HG0000140.1 | -0.35343 | 5.300482 | -3.46137 | 0.00107 | 0.015323 | -1.07342 | Glycosyltransferase-like KOBITO 1 | elongation zone | 12 h |
| HORVU.MOREX.r3.4HG0352780.1 | 0.751161 | 1.138745 | 3.413052 | 0.001238 | 0.016656 | -1.07666 | WRKY transcription factor | elongation zone | 12 h |
| HORVU.MOREX.r3.7HG0749070.1 | 1.223174 | -1.82628 | 3.372802 | 0.001396 | 0.017896 | -1.07794 | nodulin MtN21 /EamA-like transporter family protein | elongation zone | 12 h |
| HORVU.MOREX.r3.6HG0554050.1 | 0.891348 | -1.56957 | 3.373736 | 0.001392 | 0.017869 | -1.07815 | cyclin-dependent kinase inhibitor | elongation zone | 12 h |
| HORVU.MOREX.r3.6HG0605940.1 | 0.717678 | 3.179648 | 3.427365 | 0.001186 | 0.016221 | -1.07968 | Xyloglucan endotransglucosylase/hydrolase | elongation zone | 12 h |
| HORVU.MOREX.r3.7HG0730800.1 | -1.12616 | 5.096289 | -3.39957 | 0.001289 | 0.017063 | -1.08195 | Histone H3 | elongation zone | 12 h |
| HORVU.MOREX.r3.2HG0141980.1 | -0.43246 | 5.307117 | -3.50485 | 0.000938 | 0.014213 | -1.08289 | Glucan endo-1,3-beta-glucosidase 3 | elongation zone | 12 h |
| HORVU.MOREX.r3.7HG0726700.1 | -0.86121 | 7.183777 | -3.47659 | 0.001022 | 0.014893 | -1.08493 | Leucine-rich repeat protein kinase family protein | elongation zone | 12 h |
| HORVU.MOREX.r3.7HG0667630.1 | 1.180154 | 5.380802 | 3.500075 | 0.000952 | 0.014357 | -1.08656 | Peroxidase | elongation zone | 12 h |
| HORVU.MOREX.r3.5HG0497940.1 | -1.2614 | 4.128013 | -3.3681 | 0.001416 | 0.018059 | -1.08767 | Homeobox leucine zipper protein | elongation zone | 12 h |
| HORVU.MOREX.r3.5HG0460720.1 | -0.50172 | 6.552351 | -3.47281 | 0.001034 | 0.01499 | -1.08905 | L-galactono-1,4-lactone dehydrogenase | elongation zone | 12 h |
| HORVU.MOREX.r3.2HG0167570.1 | -1.31702 | 5.317382 | -3.39706 | 0.001299 | 0.017131 | -1.08935 | Histone H3 | elongation zone | 12 h |
| HORVU.MOREX.r3.2HG0141140.1 | 1.118787 | -2.85799 | 3.407892 | 0.001257 | 0.016849 | -1.08957 | DUF4228 domain protein | elongation zone | 12 h |
| HORVU.MOREX.r3.3HG0313040.1 | -1.42151 | 6.108078 | -3.47473 | 0.001028 | 0.014935 | -1.08965 | Hydroxycinnamoyl-CoA shikimate/quinate hydroxycinnamoyltransferase | elongation zone | 12 h |
| HORVU.MOREX.r3.7HG0685350.1 | 0.602232 | 3.413791 | 3.434361 | 0.001161 | 0.016061 | -1.09306 | LysM receptor-like kinase protein | elongation zone | 12 h |
| HORVU.MOREX.r3.5HG0500680.1 | -0.68109 | 6.785877 | -3.48185 | 0.001006 | 0.01473 | -1.09416 | Acyl carrier protein | elongation zone | 12 h |
| HORVU.MOREX.r3.3HG0231000.1 | 0.924133 | 3.745714 | 3.481856 | 0.001006 | 0.01473 | -1.09688 | Gamma-glutamyltranspeptidase | elongation zone | 12 h |
| HORVU.MOREX.r3.7HG0738680.1 | 0.722108 | 3.55287 | 3.445839 | 0.001121 | 0.01573 | -1.09755 | Vacuolar sorting receptor family protein | elongation zone | 12 h |
| HORVU.MOREX.r3.3HG0256360.1 | -0.37438 | 5.280798 | -3.45633 | 0.001087 | 0.015419 | -1.09811 | BSD domain-containing protein 1 | elongation zone | 12 h |
| HORVU.MOREX.r3.1HG0077840.1 | -0.46114 | 7.930641 | -3.53727 | 0.00085 | 0.013451 | -1.09969 | Glucan endo-1,3-beta-glucosidase 1 | elongation zone | 12 h |
| HORVU.MOREX.r3.7HG0665520.1 | -0.53998 | 3.332246 | -3.42527 | 0.001193 | 0.016301 | -1.10037 | MBOAT (Membrane bound O-acyl transferase) family protein | elongation zone | 12 h |
| HORVU.MOREX.r3.3HG0325760.1 | -0.49518 | 5.438001 | -3.43883 | 0.001145 | 0.015935 | -1.10344 | Mitochondrial carrier protein-like | elongation zone | 12 h |
| HORVU.MOREX.r3.4HG0406630.1 | 0.719518 | 4.673873 | 3.526775 | 0.000877 | 0.013709 | -1.10434 | Classical arabinogalactan protein 5 | elongation zone | 12 h |
| HORVU.MOREX.r3.7HG0647930.1 | -1.39828 | 1.498899 | -3.44308 | 0.001131 | 0.015774 | -1.10637 | S-adenosyl-L-methionine-dependent methyltransferases superfamily protein | elongation zone | 12 h |
| HORVU.MOREX.r3.6HG0613270.1 | -0.44926 | 9.613904 | -3.58334 | 0.000738 | 0.012516 | -1.1064 | Glutamine synthetase | elongation zone | 12 h |
| HORVU.MOREX.r3.7HG0661490.1 | -0.30853 | 6.094894 | -3.48695 | 0.00099 | 0.014693 | -1.10709 | Stromal cell-derived factor 2 | elongation zone | 12 h |
| HORVU.MOREX.r3.7HG0701640.1 | 0.807464 | 2.542885 | 3.469902 | 0.001043 | 0.01508 | -1.10717 | Vesicle-associated protein 4-2 | elongation zone | 12 h |
| HORVU.MOREX.r3.5HG0519360.1 | 0.968408 | 2.680303 | 3.354868 | 0.001473 | 0.018499 | -1.11127 | Retrovirus-related Pol polyprotein from transposon TNT 1-94 | elongation zone | 12 h |
| HORVU.MOREX.r3.3HG0309820.1 | -0.73799 | 5.758875 | -3.42975 | 0.001177 | 0.016136 | -1.11128 | Small ubiquitin-related modifier | elongation zone | 12 h |
| HORVU.MOREX.r3.2HG0206910.1 | -1.00295 | 3.37302 | -3.35461 | 0.001474 | 0.0185 | -1.1122 | Mitochondrial transcription termination factor family protein | elongation zone | 12 h |
| HORVU.MOREX.r3.6HG0599450.1 | 1.723706 | -2.13758 | 3.386836 | 0.001339 | 0.017425 | -1.11388 | Yellow stripe 1A transporter | elongation zone | 12 h |
| HORVU.MOREX.r3.7HG0705490.1 | -1.52851 | 2.356127 | -3.42283 | 0.001202 | 0.016359 | -1.11538 | HXXXD-type acyl-transferase family protein | elongation zone | 12 h |
| HORVU.MOREX.r3.2HG0100040.1 | -2.09234 | -0.29926 | -3.45829 | 0.00108 | 0.015382 | -1.11801 | HXXXD-type acyl-transferase family protein | elongation zone | 12 h |

**Table S1** Continued.

| ID | logFC | AveExpr | t | P.Value | adj.P.Val | B | gene description | root zone | time point |
|---|---|---|---|---|---|---|---|---|---|
| HORVU.MOREX.r3.7HG0718720.1 | 1.046375 | 2.503323 | 3.366125 | 0.001424 | 0.018114 | -1.11809 | DUF1685 family protein | elongation zone | 12 h |
| HORVU.MOREX.r3.5HG0485780.1 | 2.537986 | -0.7336 | 3.371804 | 0.0014 | 0.017938 | -1.11911 | 1-aminocyclopropane-1-carboxylate oxidase | elongation zone | 12 h |
| HORVU.MOREX.r3.2HG0108490.1 | -1.27031 | 3.17005 | -3.35058 | 0.001492 | 0.018644 | -1.12235 | Xyloglucan endotransglucosylase/hydrolase | elongation zone | 12 h |
| HORVU.MOREX.r3.5HG0476460.1 | -0.75976 | 2.596562 | -3.3543 | 0.001475 | 0.018506 | -1.12369 | VQ motif-containing family protein | elongation zone | 12 h |
| HORVU.MOREX.r3.2HG0173960.1 | 0.667257 | 2.814915 | 3.360621 | 0.001448 | 0.018262 | -1.12407 | Peroxidase | elongation zone | 12 h |
| HORVU.MOREX.r3.2HG0203500.1 | -0.82561 | 6.139708 | -3.44457 | 0.001126 | 0.01576 | -1.12437 | UPF0136 membrane protein | elongation zone | 12 h |
| HORVU.MOREX.r3.7HG0638340.1 | 1.233699 | 2.325425 | 3.395256 | 0.001306 | 0.017152 | -1.12886 | S-adenosyl-L-methionine-dependent methyltransferases superfamily protein | elongation zone | 12 h |
| HORVU.MOREX.r3.3HG0323600.1 | 0.532598 | 4.355593 | 3.450713 | 0.001105 | 0.015604 | -1.13286 | Phosphatidylserine decarboxylase proenzyme 2 | elongation zone | 12 h |
| HORVU.MOREX.r3.3HG0272430.1 | -0.40368 | 5.81708 | -3.45887 | 0.001078 | 0.015382 | -1.13319 | Pectin lyase-like superfamily protein | elongation zone | 12 h |
| HORVU.MOREX.r3.4HG0342640.1 | -1.80842 | 4.773275 | -3.34393 | 0.001522 | 0.018912 | -1.13833 | Cyclin | elongation zone | 12 h |
| HORVU.MOREX.r3.7HG0727290.1 | 0.693282 | 3.998416 | 3.462295 | 0.001067 | 0.015301 | -1.14211 | #N/A | elongation zone | 12 h |
| HORVU.MOREX.r3.7HG0651010.1 | -0.93294 | 4.932391 | -3.41349 | 0.001236 | 0.016645 | -1.14292 | Xyloglucan endotransglucosylase/hydrolase | elongation zone | 12 h |
| HORVU.MOREX.r3.3HG0292080.1 | -0.48101 | 4.723449 | -3.41699 | 0.001223 | 0.016525 | -1.14348 | F-box family protein-like | elongation zone | 12 h |
| HORVU.MOREX.r3.3HG0283250.1 | 0.617689 | 3.003629 | 3.453024 | 0.001097 | 0.015536 | -1.14386 | MYB transcription factor | elongation zone | 12 h |
| HORVU.MOREX.r3.4HG0416760.1 | -0.52908 | 4.391267 | -3.40337 | 0.001274 | 0.016993 | -1.14601 | Non-structural protein 1 | elongation zone | 12 h |
| HORVU.MOREX.r3.4HG0416390.1 | 1.347277 | 0.67132 | 3.420693 | 0.00121 | 0.016416 | -1.14712 | Sugar transporter, putative | elongation zone | 12 h |
| HORVU.MOREX.r3.7HG0672400.1 | 1.070717 | 3.007592 | 3.349056 | 0.001499 | 0.018717 | -1.14892 | DNA repair protein complementing XP-C cells-like protein | elongation zone | 12 h |
| HORVU.MOREX.r3.1HG0030290.1 | 3.260825 | -1.84835 | 3.863468 | 0.000306 | 0.007759 | -1.15015 | PRA1 family protein | elongation zone | 12 h |
| HORVU.MOREX.r3.1HG0076980.1 | -0.34441 | 8.500486 | -3.54531 | 0.000829 | 0.013267 | -1.15123 | Dolichyl-diphosphooligosaccharide--protein glycosyltransferase subunit STT3 | elongation zone | 12 h |
| HORVU.MOREX.r3.1HG0066730.1 | -0.68177 | 4.534097 | -3.41379 | 0.001235 | 0.016641 | -1.15222 | Indole-3-glycerol phosphate synthase | elongation zone | 12 h |
| HORVU.MOREX.r3.6HG0595190.1 | 1.519507 | -2.07211 | 3.359425 | 0.001453 | 0.018316 | -1.15482 | GDSL esterase/lipase | elongation zone | 12 h |
| HORVU.MOREX.r3.5HG0490960.1 | -0.64319 | 3.86387 | -3.36423 | 0.001432 | 0.018164 | -1.15504 | DNA-binding bromodomain-containing protein | elongation zone | 12 h |
| HORVU.MOREX.r3.5HG0438460.1 | 1.2713 | 0.165145 | 3.385245 | 0.001345 | 0.017473 | -1.15654 | basic helix-loop-helix (bHLH) DNA-binding superfamily protein | elongation zone | 12 h |
| HORVU.MOREX.r3.1HG0075260.1 | 3.105787 | 0.393741 | 3.535071 | 0.000855 | 0.01351 | -1.158 | Receptor kinase 2 | elongation zone | 12 h |
| HORVU.MOREX.r3.4HG0375690.1 | -1.15219 | 1.261321 | -3.39574 | 0.001304 | 0.017152 | -1.15963 | DENN domain-containing protein 5B | elongation zone | 12 h |
| HORVU.MOREX.r3.2HG0124820.1 | -0.85368 | 3.652036 | -3.34875 | 0.0015 | 0.018723 | -1.16037 | F-box protein | elongation zone | 12 h |
| HORVU.MOREX.r3.7HG0739490.1 | -0.9539 | 7.817058 | -3.48126 | 0.001008 | 0.014746 | -1.16056 | Histone H2B | elongation zone | 12 h |
| HORVU.MOREX.r3.4HG0345580.1 | 1.914774 | -3.15122 | 3.540894 | 0.00084 | 0.013375 | -1.16065 | Dirigent protein | elongation zone | 12 h |
| HORVU.MOREX.r3.6HG0587790.1 | -1.59516 | -0.11015 | -3.49174 | 0.000976 | 0.014552 | -1.16238 | Transposon Ty3-G Gag-Pol polyprotein | elongation zone | 12 h |
| HORVU.MOREX.r3.4HG0416220.1 | -0.31577 | 6.683316 | -3.4828 | 0.001003 | 0.014726 | -1.16485 | Cell number regulator 6 | elongation zone | 12 h |
| HORVU.MOREX.r3.4HG0416730.1 | -0.80843 | 6.869699 | -3.43656 | 0.001153 | 0.016012 | -1.16539 | Kinesin-like protein | elongation zone | 12 h |
| HORVU.MOREX.r3.4HG0362450.1 | -0.9434 | 2.708144 | -3.33292 | 0.001572 | 0.019344 | -1.17196 | calmodulin-binding protein (DUF1645) | elongation zone | 12 h |
| HORVU.MOREX.r3.5HG0450100.1 | -0.52611 | 5.024744 | -3.3848 | 0.001347 | 0.017485 | -1.17573 | Quinolinate synthase A | elongation zone | 12 h |
| HORVU.MOREX.r3.1HG0024230.1 | -0.5737 | 5.422255 | -3.38679 | 0.001339 | 0.017425 | -1.1785 | Tubulin beta chain | elongation zone | 12 h |
| HORVU.MOREX.r3.5HG0486610.1 | 0.831506 | 3.213671 | 3.485919 | 0.000993 | 0.014718 | -1.17957 | Leucine-rich repeat receptor-like protein kinase family protein | elongation zone | 12 h |
| HORVU.MOREX.r3.7HG0728000.1 | 0.583007 | 5.007313 | 3.469666 | 0.001044 | 0.01508 | -1.1833 | Glycosyltransferase | elongation zone | 12 h |
| HORVU.MOREX.r3.5HG0504800.1 | -0.81346 | 5.65419 | -3.39331 | 0.001313 | 0.017213 | -1.18409 | Histone-lysine N-methyltransferase | elongation zone | 12 h |
| HORVU.MOREX.r3.5HG0466020.1 | -0.76269 | 7.677987 | -3.48304 | 0.001002 | 0.014726 | -1.18833 | Omega-3 fatty acid desaturase | elongation zone | 12 h |
| HORVU.MOREX.r3.1HG0060270.1 | -1.05896 | 5.417861 | -3.38206 | 0.001358 | 0.017584 | -1.19144 | Histone H3 | elongation zone | 12 h |
| HORVU.MOREX.r3.1HG0055730.1 | -0.43234 | 4.544955 | -3.38035 | 0.001365 | 0.017608 | -1.19155 | Translocase of chloroplast | elongation zone | 12 h |
| HORVU.MOREX.r3.5HG0529130.1 | -0.58652 | 9.571785 | -3.53229 | 0.000863 | 0.013594 | -1.19252 | Elongation factor | elongation zone | 12 h |

**Table S1** Continued.

| ID | logFC | AveExpr | t | P.Value | adj.P.Val | B | gene description | root zone | time point |
|---|---|---|---|---|---|---|---|---|---|
| HORVU.MOREX.r3.3HG0230970.1 | -0.81634 | 7.763065 | -3.47254 | 0.001034 | 0.014991 | -1.19277 | Histone H2B | elongation zone | 12 h |
| HORVU.MOREX.r3.2HG0164470.1 | 0.737877 | 3.24648 | 3.402184 | 0.001279 | 0.017029 | -1.19281 | Nudix hydrolase | elongation zone | 12 h |
| HORVU.MOREX.r3.2HG0126440.1 | 1.162297 | 2.011284 | 3.366031 | 0.001425 | 0.018114 | -1.19635 | ATP-dependent RNA helicase-like protein | elongation zone | 12 h |
| HORVU.MOREX.r3.7HG0737470.1 | 1.040212 | 3.292476 | 3.578796 | 0.000748 | 0.012611 | -1.19675 | Peroxidase | elongation zone | 12 h |
| HORVU.MOREX.r3.5HG0507680.1 | 1.287413 | -1.22844 | 3.341747 | 0.001532 | 0.019016 | -1.19914 | mitogen-activated protein kinase kinase kinase 6 | elongation zone | 12 h |
| HORVU.MOREX.r3.6HG0543390.1 | 1.225568 | 0.250443 | 3.400451 | 0.001285 | 0.01705 | -1.20079 | High affinity nitrate transporter | elongation zone | 12 h |
| HORVU.MOREX.r3.3HG0255040.1 | -0.40994 | 4.904219 | -3.38232 | 0.001357 | 0.017582 | -1.20151 | Mediator of RNA polymerase II transcription subunit 27 | elongation zone | 12 h |
| HORVU.MOREX.r3.1HG0004560.1 | 0.848348 | 5.305216 | 3.383604 | 0.001352 | 0.017536 | -1.20216 | ERD (Early-responsive to dehydration stress) family protein | elongation zone | 12 h |
| HORVU.MOREX.r3.3HG0243920.1 | -1.18597 | 3.290866 | -3.31894 | 0.001639 | 0.019852 | -1.20383 | VQ motif-containing protein | elongation zone | 12 h |
| HORVU.MOREX.r3.3HG0304640.1 | -0.58208 | 4.177695 | -3.34439 | 0.00152 | 0.018909 | -1.20406 | Tetratricopeptide repeat-containing family protein | elongation zone | 12 h |
| HORVU.MOREX.r3.7HG0730310.1 | -0.8961 | 7.32045 | -3.44934 | 0.00111 | 0.015648 | -1.20523 | Histone H2A | elongation zone | 12 h |
| HORVU.MOREX.r3.1HG0002800.1 | -1.35957 | 3.206208 | -3.3385 | 0.001546 | 0.01915 | -1.20583 | Powdery mildew resistance protein Pm3 | elongation zone | 12 h |
| HORVU.MOREX.r3.3HG0316130.1 | -0.7369 | 7.033938 | -3.51456 | 0.00091 | 0.013989 | -1.20603 | Basic 7S globulin 2 | elongation zone | 12 h |
| HORVU.MOREX.r3.6HG0573870.1 | -0.64366 | 4.15012 | -3.3358 | 0.001559 | 0.019246 | -1.20625 | Histone H3 | elongation zone | 12 h |
| HORVU.MOREX.r3.2HG0138620.1 | -1.11624 | 3.379812 | -3.32057 | 0.001631 | 0.019852 | -1.20689 | MBOAT (Membrane bound O-acyl transferase) family protein | elongation zone | 12 h |
| HORVU.MOREX.r3.2HG0185600.1 | 1.052814 | 0.732582 | 3.422538 | 0.001203 | 0.016359 | -1.20698 | TRIO and F-actin-binding protein | elongation zone | 12 h |
| HORVU.MOREX.r3.2HG0185840.1 | -0.96076 | 7.644996 | -3.45406 | 0.001094 | 0.015514 | -1.209 | #N/A | elongation zone | 12 h |
| HORVU.MOREX.r3.7HG0674790.1 | -0.74093 | 5.490517 | -3.37668 | 0.00138 | 0.017746 | -1.20936 | Microtubule associated family protein | elongation zone | 12 h |
| HORVU.MOREX.r3.7HG0654460.1 | -1.07171 | 6.42811 | -3.40905 | 0.001253 | 0.016813 | -1.20979 | Histone H3 | elongation zone | 12 h |
| HORVU.MOREX.r3.1HG0008190.1 | -1.16374 | -0.63382 | -3.31941 | 0.001636 | 0.019852 | -1.2125 | Cytochrome P450 | elongation zone | 12 h |
| HORVU.MOREX.r3.3HG0266530.1 | 0.854064 | 1.443645 | 3.315997 | 0.001653 | 0.019898 | -1.21349 | carboxyl-terminal peptidase, putative (DUF239) | elongation zone | 12 h |
| HORVU.MOREX.r3.5HG0479970.1 | -0.39042 | 6.75595 | -3.45197 | 0.001101 | 0.015569 | -1.21368 | Aspartyl aminopeptidase, putative, expressed | elongation zone | 12 h |
| HORVU.MOREX.r3.2HG0165250.1 | -0.40338 | 6.077437 | -3.44323 | 0.00113 | 0.015774 | -1.21642 | OJ000315_02.19 protein | elongation zone | 12 h |
| HORVU.MOREX.r3.3HG0259960.1 | -1.96866 | -0.28204 | -3.55981 | 0.000793 | 0.013047 | -1.21715 | Leucine-rich repeat protein kinase family protein | elongation zone | 12 h |
| HORVU.MOREX.r3.3HG0252210.1 | 1.172954 | 0.169661 | 3.362795 | 0.001439 | 0.018178 | -1.21848 | Transcription factor GLABRA 3 | elongation zone | 12 h |
| HORVU.MOREX.r3.4HG0374640.1 | -0.71951 | 4.330972 | -3.33722 | 0.001552 | 0.0192 | -1.22234 | DNA-binding storekeeper protein-related transcriptional regulator | elongation zone | 12 h |
| HORVU.MOREX.r3.3HG0218330.1 | -1.57496 | 8.028798 | -3.41701 | 0.001223 | 0.016525 | -1.22237 | #N/A | elongation zone | 12 h |
| HORVU.MOREX.r3.4HG0415230.1 | -0.90584 | 2.026714 | -3.30905 | 0.001687 | 0.020145 | -1.22578 | basic helix-loop-helix (bHLH) DNA-binding superfamily protein | elongation zone | 12 h |
| HORVU.MOREX.r3.2HG0215250.1 | -1.13212 | 6.918385 | -3.43058 | 0.001174 | 0.016107 | -1.22638 | Peroxidase | elongation zone | 12 h |
| HORVU.MOREX.r3.4HG0383530.1 | 1.090938 | -0.0177 | 3.407607 | 0.001258 | 0.016853 | -1.22725 | Pathogenesis-related protein 1 | elongation zone | 12 h |
| HORVU.MOREX.r3.3HG0297140.1 | -0.26881 | 7.212598 | -3.50952 | 0.000925 | 0.014127 | -1.2276 | Auxin efflux carrier family protein-like | elongation zone | 12 h |
| HORVU.MOREX.r3.2HG0196960.1 | -2.61213 | 1.756153 | -3.59214 | 0.000718 | 0.012368 | -1.22847 | Y' element ATP-dependent helicase YEL077C | elongation zone | 12 h |
| HORVU.MOREX.r3.7HG0735210.1 | 1.656602 | -3.00596 | 3.500743 | 0.00095 | 0.014339 | -1.2313 | Peptide transporter | elongation zone | 12 h |
| HORVU.MOREX.r3.5HG0463350.1 | -1.63397 | 5.140919 | -3.32274 | 0.00162 | 0.019769 | -1.23208 | Cell division control 45-like protein | elongation zone | 12 h |
| HORVU.MOREX.r3.6HG0582120.1 | 0.477315 | 3.287652 | 3.363101 | 0.001437 | 0.018178 | -1.23218 | magnesium transporter NIPA (DUF803) | elongation zone | 12 h |
| HORVU.MOREX.r3.1HG0049450.1 | -0.42734 | 5.425708 | -3.42486 | 0.001195 | 0.01631 | -1.23262 | #N/A | elongation zone | 12 h |
| HORVU.MOREX.r3.6HG0622080.1 | 0.692966 | 1.073855 | 3.381616 | 0.00136 | 0.017585 | -1.23291 | MYB-related transcription factor | elongation zone | 12 h |
| HORVU.MOREX.r3.2HG0157160.1 | -0.83514 | 4.380598 | -3.31791 | 0.001644 | 0.019852 | -1.23441 | Receptor kinase | elongation zone | 12 h |
| HORVU.MOREX.r3.5HG0500360.1 | 2.916843 | 1.290708 | 3.423819 | 0.001198 | 0.01635 | -1.23467 | L-gulonolactone oxidase | elongation zone | 12 h |
| HORVU.MOREX.r3.2HG0147770.1 | -0.44361 | 5.593653 | -3.40056 | 0.001285 | 0.01705 | -1.23771 | Growth-regulating factor | elongation zone | 12 h |
| HORVU.MOREX.r3.4HG0339430.2 | -1.84443 | 3.805151 | -3.31663 | 0.00165 | 0.019892 | -1.24232 | Growth-regulating factor | elongation zone | 12 h |

**Table S1** Continued.

| ID | logFC | AveExpr | t | P.Value | adj.P.Val | B | gene description | root zone | time point |
|---|---|---|---|---|---|---|---|---|---|
| HORVU.MOREX.r3.5HG0473920.1 | -0.48361 | 3.263199 | -3.33396 | 0.001567 | 0.019305 | -1.24358 | OBP32pep, putative (DUF220) | elongation zone | 12 h |
| HORVU.MOREX.r3.6HG0558330.1 | -1.16252 | 6.152461 | -3.39323 | 0.001314 | 0.017213 | -1.24483 | Calcium-dependent lipid-binding domain protein | elongation zone | 12 h |
| HORVU.MOREX.r3.5HG0493330.1 | 0.994298 | 0.977391 | 3.358806 | 0.001456 | 0.018338 | -1.24564 | UDP-glucuronate 4-epimerase 4 | elongation zone | 12 h |
| HORVU.MOREX.r3.6HG0606360.1 | 2.198272 | -2.06608 | 3.317656 | 0.001645 | 0.019855 | -1.24677 | Peptide transporter | elongation zone | 12 h |
| HORVU.MOREX.r3.5HG0472770.1 | 0.66082 | 4.331329 | 3.361641 | 0.001444 | 0.018229 | -1.24744 | SIN3-like 2 | elongation zone | 12 h |
| HORVU.MOREX.r3.3HG0306210.1 | -0.6006 | 7.280706 | -3.42273 | 0.001202 | 0.016359 | -1.24981 | RAN GTPase activating protein 2 | elongation zone | 12 h |
| HORVU.MOREX.r3.7HG0642770.1 | -0.35383 | 5.619529 | -3.39588 | 0.001303 | 0.017152 | -1.24991 | RNA-binding family protein | elongation zone | 12 h |
| HORVU.MOREX.r3.5HG0530240.1 | -1.14968 | 0.885356 | -3.39153 | 0.00132 | 0.017256 | -1.25161 | Cytochrome P450 family protein | elongation zone | 12 h |
| HORVU.MOREX.r3.6HG0596460.1 | -0.41541 | 6.392262 | -3.42254 | 0.001203 | 0.016359 | -1.25331 | Cryptochrome | elongation zone | 12 h |
| HORVU.MOREX.r3.3HG0252240.1 | -0.90173 | 7.225249 | -3.4184 | 0.001218 | 0.016477 | -1.25776 | Histone H2B | elongation zone | 12 h |
| HORVU.MOREX.r3.5HG0496930.1 | 0.836419 | 1.51856 | 3.297381 | 0.001746 | 0.02056 | -1.25807 | Inositol-tetrakisphosphate 1-kinase 4 | elongation zone | 12 h |
| HORVU.MOREX.r3.1HG0063870.1 | -0.62243 | 6.880931 | -3.42016 | 0.001212 | 0.016422 | -1.25822 | Kinesin-like protein | elongation zone | 12 h |
| HORVU.MOREX.r3.7HG0690090.1 | 0.837973 | 0.298554 | 3.353458 | 0.001479 | 0.01853 | -1.25918 | PLATZ transcription factor family protein | elongation zone | 12 h |
| HORVU.MOREX.r3.4HG0343040.1 | 0.776077 | 2.501277 | 3.407078 | 0.00126 | 0.016857 | -1.26159 | basic helix-loop-helix (bHLH) DNA-binding superfamily protein | elongation zone | 12 h |
| HORVU.MOREX.r3.5HG0528340.1 | -0.97974 | 3.051746 | -3.29437 | 0.001762 | 0.020637 | -1.26216 | F-box family protein | elongation zone | 12 h |
| HORVU.MOREX.r3.6HG0616680.1 | -1.34067 | 0.45557 | -3.39527 | 0.001306 | 0.017152 | -1.26378 | F-box family protein | elongation zone | 12 h |
| HORVU.MOREX.r3.3HG0284760.1 | 0.568664 | 6.914607 | 3.511768 | 0.000918 | 0.014073 | -1.26665 | Myosin | elongation zone | 12 h |
| HORVU.MOREX.r3.4HG0358580.1 | -1.04932 | 3.228773 | -3.29395 | 0.001764 | 0.020639 | -1.26725 | Harbinger transposase-derived nuclease | elongation zone | 12 h |
| HORVU.MOREX.r3.3HG0315000.1 | -0.47241 | 3.461758 | -3.31846 | 0.001641 | 0.019852 | -1.26766 | F-box protein, putative (DUF295) | elongation zone | 12 h |
| HORVU.MOREX.r3.7HG0674320.1 | -0.37467 | 4.709886 | -3.39807 | 0.001295 | 0.017096 | -1.26778 | Vesicle transport protein | elongation zone | 12 h |
| HORVU.MOREX.r3.1HG0015750.1 | 1.657205 | -2.91215 | 3.329527 | 0.001588 | 0.019467 | -1.26831 | transcription factor, putative (Protein of unknown function, DUF547) | elongation zone | 12 h |
| HORVU.MOREX.r3.7HG0682120.1 | -0.38256 | 4.669456 | -3.37961 | 0.001368 | 0.01763 | -1.26842 | Methyltransferase-like protein 13 | elongation zone | 12 h |
| HORVU.MOREX.r3.2HG0104510.1 | -0.35808 | 7.315942 | -3.43316 | 0.001165 | 0.016068 | -1.26856 | FACT complex subunit SPT16 | elongation zone | 12 h |
| HORVU.MOREX.r3.5HG0471960.1 | -1.20112 | 3.222285 | -3.29323 | 0.001768 | 0.020671 | -1.26927 | Chromosome transmission fidelity protein 8 | elongation zone | 12 h |
| HORVU.MOREX.r3.7HG0654580.1 | -1.5678 | 4.602405 | -3.29765 | 0.001745 | 0.02056 | -1.26969 | Histone H3 | elongation zone | 12 h |
| HORVU.MOREX.r3.2HG0173190.1 | -1.43972 | 7.090069 | -3.35186 | 0.001486 | 0.018584 | -1.27026 | ABC transporter ATP-binding protein | elongation zone | 12 h |
| HORVU.MOREX.r3.6HG0622030.1 | -0.5847 | 3.435933 | -3.30412 | 0.001712 | 0.020365 | -1.2734 | tRNA pseudouridine synthase | elongation zone | 12 h |
| HORVU.MOREX.r3.4HG0384350.1 | 0.378712 | 5.925988 | 3.445886 | 0.001121 | 0.01573 | -1.27347 | Rab3 GTPase-activating protein catalytic subunit | elongation zone | 12 h |
| HORVU.MOREX.r3.2HG0186980.1 | 1.099659 | 1.712815 | 3.301087 | 0.001727 | 0.020487 | -1.27423 | Vacuolar protein sorting-associated protein 53 A | elongation zone | 12 h |
| HORVU.MOREX.r3.5HG0466890.1 | -0.57769 | 3.242376 | -3.30024 | 0.001732 | 0.020487 | -1.27596 | Alcohol dehydrogenase 2 | elongation zone | 12 h |
| HORVU.MOREX.r3.2HG0105680.3 | -1.02681 | 4.721415 | -3.31054 | 0.00168 | 0.020135 | -1.27611 | Translation initiation factor IF-2 | elongation zone | 12 h |
| HORVU.MOREX.r3.2HG0205990.1 | 1.897341 | -2.55766 | 3.362788 | 0.001439 | 0.018178 | -1.27657 | Glutaredoxin family protein, putative | elongation zone | 12 h |
| HORVU.MOREX.r3.7HG0654120.1 | -1.19858 | 1.753011 | -3.28907 | 0.00179 | 0.020878 | -1.27662 | Basic helix-loop-helix transcription factor | elongation zone | 12 h |
| HORVU.MOREX.r3.4HG0376950.1 | -2.2875 | -0.73491 | -3.6447 | 0.00061 | 0.011213 | -1.27762 | Transposon protein, putative, CACTA, En/Spm sub-class | elongation zone | 12 h |
| HORVU.MOREX.r3.2HG0097350.1 | -2.09498 | 5.612431 | -3.30093 | 0.001728 | 0.020487 | -1.28079 | Agmatine coumaroyltransferase-2 | elongation zone | 12 h |
| HORVU.MOREX.r3.3HG0305800.1 | 0.766478 | 4.447 | 3.352212 | 0.001485 | 0.018576 | -1.28158 | Ribonuclease | elongation zone | 12 h |
| HORVU.MOREX.r3.5HG0487910.1 | 2.001237 | -2.5953 | 3.405349 | 0.001267 | 0.016918 | -1.28226 | Purine permease-like protein | elongation zone | 12 h |
| HORVU.MOREX.r3.4HG0401130.1 | 0.362452 | 6.245539 | 3.45112 | 0.001104 | 0.015598 | -1.28242 | La-related protein | elongation zone | 12 h |
| HORVU.MOREX.r3.5HG0536900.1 | -1.47371 | 0.743448 | -3.41094 | 0.001246 | 0.016739 | -1.28312 | F-box only 46 | elongation zone | 12 h |
| HORVU.MOREX.r3.3HG0276120.1 | 1.77701 | -1.33075 | 3.315923 | 0.001653 | 0.019898 | -1.28392 | Phospholipase A1 | elongation zone | 12 h |
| HORVU.MOREX.r3.7HG0636750.1 | -0.86208 | 7.072571 | -3.42275 | 0.001202 | 0.016359 | -1.28546 | carbohydrate esterase, putative (DUF303) | elongation zone | 12 h |

**Table S1** Continued.

| ID | logFC | AveExpr | t | P.Value | adj.P.Val | B | gene description | root zone | time point |
|---|---|---|---|---|---|---|---|---|---|
| HORVU.MOREX.r3.3HG0310210.1 | -0.53852 | 7.031206 | -3.42051 | 0.00121 | 0.016416 | -1.28954 | Acyl-[acyl-carrier-protein] desaturase | elongation zone | 12 h |
| HORVU.MOREX.r3.2HG0173680.1 | -0.4292 | 6.209185 | -3.39528 | 0.001306 | 0.017152 | -1.29008 | Anthranilate synthase | elongation zone | 12 h |
| HORVU.MOREX.r3.4HG0339180.1 | -1.95132 | 3.295927 | -3.31914 | 0.001638 | 0.019852 | -1.29376 | F-box-like protein | elongation zone | 12 h |
| HORVU.MOREX.r3.4HG0413410.1 | -0.96731 | 7.171548 | -3.43074 | 0.001174 | 0.016107 | -1.29488 | Hydroxymethylglutaryl-CoA synthase | elongation zone | 12 h |
| HORVU.MOREX.r3.6HG0557070.1 | -0.49462 | 6.4953 | -3.41184 | 0.001242 | 0.016705 | -1.29519 | Activator of heat shock protein ATPase | elongation zone | 12 h |
| HORVU.MOREX.r3.7HG0676890.1 | -1.4801 | 1.263909 | -3.31805 | 0.001643 | 0.019852 | -1.29612 | Transmembrane protein, putative | elongation zone | 12 h |
| HORVU.MOREX.r3.4HG0383530.3 | 0.947781 | 1.457399 | 3.363326 | 0.001436 | 0.018178 | -1.2962 | Pathogenesis-related protein 1 | elongation zone | 12 h |
| HORVU.MOREX.r3.4HG0333130.1 | -1.10703 | 1.113277 | -3.33661 | 0.001555 | 0.019223 | -1.29626 | #N/A | elongation zone | 12 h |
| HORVU.MOREX.r3.6HG0632210.1 | -0.54573 | 4.522362 | -3.34629 | 0.001511 | 0.018826 | -1.29675 | Electron transfer flavoprotein beta-subunit | elongation zone | 12 h |
| HORVU.MOREX.r3.3HG0322650.1 | 0.700248 | 3.559006 | 3.380429 | 0.001365 | 0.017608 | -1.29827 | NAD kinase | elongation zone | 12 h |
| HORVU.MOREX.r3.3HG0291670.1 | 1.294579 | -0.21921 | 3.311651 | 0.001674 | 0.020092 | -1.29841 | Transmembrane protein 56 | elongation zone | 12 h |
| HORVU.MOREX.r3.7HG0745980.1 | -0.40314 | 3.968558 | -3.33287 | 0.001572 | 0.019344 | -1.29978 | Mutator-like transposase | elongation zone | 12 h |
| HORVU.MOREX.r3.6HG0548820.1 | -1.23534 | 0.468673 | -3.31637 | 0.001651 | 0.019896 | -1.30386 | #N/A | elongation zone | 12 h |
| HORVU.MOREX.r3.2HG0168790.1 | -0.72427 | 4.208617 | -3.31995 | 0.001634 | 0.019852 | -1.304 | Haloacid dehalogenase-like hydrolase family protein | elongation zone | 12 h |
| HORVU.MOREX.r3.3HG0313020.1 | -0.33221 | 5.21659 | -3.37852 | 0.001373 | 0.017671 | -1.30965 | Alcohol dehydrogenase, putative | elongation zone | 12 h |
| HORVU.MOREX.r3.3HG0219290.1 | -0.42496 | 5.927719 | -3.45697 | 0.001084 | 0.01541 | -1.31095 | WAT1-related protein | elongation zone | 12 h |
| HORVU.MOREX.r3.4HG0343050.1 | 0.494764 | 6.230075 | 3.434201 | 0.001162 | 0.016061 | -1.31102 | N-acetyl-gamma-glutamyl-phosphate reductase | elongation zone | 12 h |
| HORVU.MOREX.r3.2HG0189530.1 | 1.269362 | 0.494093 | 3.390192 | 0.001326 | 0.017304 | -1.31323 | Expansin protein | elongation zone | 12 h |
| HORVU.MOREX.r3.6HG0629240.1 | 1.74505 | -0.90421 | 3.276776 | 0.001856 | 0.021386 | -1.31506 | O-methyltransferase | elongation zone | 12 h |
| HORVU.MOREX.r3.2HG0130770.1 | -0.5732 | 3.563473 | -3.30888 | 0.001688 | 0.020145 | -1.31578 | maternal effect embryo arrest 60 | elongation zone | 12 h |
| HORVU.MOREX.r3.7HG0645480.1 | -0.50336 | 4.421668 | -3.32057 | 0.001631 | 0.019852 | -1.31609 | RNA 2'-phosphotransferase | elongation zone | 12 h |
| HORVU.MOREX.r3.1HG0091360.1 | -0.47807 | 6.214277 | -3.39527 | 0.001306 | 0.017152 | -1.31911 | Delta(14)-sterol reductase | elongation zone | 12 h |
| HORVU.MOREX.r3.1HG0094940.1 | -0.42519 | 7.388621 | -3.43129 | 0.001172 | 0.016094 | -1.32142 | ATP-dependent zinc metalloprotease FTSH protein | elongation zone | 12 h |
| HORVU.MOREX.r3.5HG0527890.1 | 0.319499 | 6.194702 | 3.42097 | 0.001209 | 0.016416 | -1.32368 | Anthranilate synthase | elongation zone | 12 h |
| HORVU.MOREX.r3.4HG0393710.1 | 1.001294 | 2.016364 | 3.309986 | 0.001683 | 0.020137 | -1.32778 | Peptide transporter | elongation zone | 12 h |
| HORVU.MOREX.r3.7HG0716320.1 | 0.530567 | 4.550005 | 3.362842 | 0.001438 | 0.018178 | -1.32954 | Kinase family protein | elongation zone | 12 h |
| HORVU.MOREX.r3.2HG0106430.1 | -0.4843 | 5.367597 | -3.39325 | 0.001313 | 0.017213 | -1.33114 | Sulfotransferase | elongation zone | 12 h |
| HORVU.MOREX.r3.4HG0383280.1 | 0.706146 | 6.188142 | 3.471119 | 0.001039 | 0.015035 | -1.33118 | Kinesin-related protein | elongation zone | 12 h |
| HORVU.MOREX.r3.4HG0414200.1 | -0.49057 | 3.59178 | -3.28571 | 0.001807 | 0.021026 | -1.33203 | Pyridoxine/pyridoxamine 5'-phosphate oxidase | elongation zone | 12 h |
| HORVU.MOREX.r3.3HG0231630.1 | -1.46627 | 3.593295 | -3.27945 | 0.001841 | 0.02129 | -1.33246 | Zinc-finger domain of monoamine-oxidase A repressor R1 protein | elongation zone | 12 h |
| HORVU.MOREX.r3.4HG0355390.1 | -0.72352 | 1.970371 | -3.26734 | 0.001908 | 0.021791 | -1.33314 | RING/U-box superfamily protein | elongation zone | 12 h |
| HORVU.MOREX.r3.7HG0749940.3 | -0.71176 | 5.387912 | -3.40311 | 0.001275 | 0.016993 | -1.3344 | Delta(24)-sterol reductase | elongation zone | 12 h |
| HORVU.MOREX.r3.1HG0067630.1 | 0.396125 | 5.282271 | 3.432074 | 0.001169 | 0.016079 | -1.33548 | Serine/threonine-protein kinase | elongation zone | 12 h |
| HORVU.MOREX.r3.2HG0192000.1 | 0.829105 | 0.357064 | 3.367043 | 0.00142 | 0.018093 | -1.33609 | HXXXD-type acyl-transferase family protein, putative | elongation zone | 12 h |
| HORVU.MOREX.r3.6HG0611060.1 | -0.40704 | 6.39405 | -3.39836 | 0.001294 | 0.017092 | -1.3366 | Cellulose synthase, putative | elongation zone | 12 h |
| HORVU.MOREX.r3.2HG0193000.1 | -0.68266 | 5.418419 | -3.34393 | 0.001522 | 0.018912 | -1.3371 | HTH-type transcriptional regulator YidZ | elongation zone | 12 h |
| HORVU.MOREX.r3.2HG0170300.1 | -0.5693 | 3.481684 | -3.28713 | 0.0018 | 0.020962 | -1.33963 | Protein FAR1-RELATED SEQUENCE 11 | elongation zone | 12 h |
| HORVU.MOREX.r3.1HG0067550.1 | -0.72865 | 6.102888 | -3.37043 | 0.001406 | 0.01799 | -1.34022 | Adenine nucleotide alpha hydrolases-like superfamily protein, putative | elongation zone | 12 h |
| HORVU.MOREX.r3.2HG0107450.1 | -0.92971 | 3.576515 | -3.26816 | 0.001903 | 0.021791 | -1.34075 | methyl-coenzyme M reductase II subunit gamma, putative (DUF3741) | elongation zone | 12 h |
| HORVU.MOREX.r3.3HG0240670.1 | -1.16159 | 5.35839 | -3.32539 | 0.001608 | 0.019683 | -1.34103 | #N/A | elongation zone | 12 h |
| HORVU.MOREX.r3.4HG0388470.1 | -2.27351 | 0.659213 | -3.57332 | 0.000761 | 0.012762 | -1.34267 | Rotundifolia-like protein | elongation zone | 12 h |

**Table S1** Continued.

| ID | logFC | AveExpr | t | P.Value | adj.P.Val | B | gene description | root zone | time point |
|---|---|---|---|---|---|---|---|---|---|
| HORVU.MOREX.r3.6HG0605150.1 | 0.904917 | 3.747441 | 3.324754 | 0.001611 | 0.019705 | -1.34316 | Purine permease family protein | elongation zone | 12 h |
| HORVU.MOREX.r3.2HG0165180.1 | 0.488914 | 4.87636 | 3.389211 | 0.001329 | 0.017343 | -1.34437 | ATP-dependent Clp protease ATP-binding subunit | elongation zone | 12 h |
| HORVU.MOREX.r3.1HG0080780.1 | -0.99698 | 6.438004 | -3.35361 | 0.001478 | 0.01853 | -1.34512 | Histone H3 | elongation zone | 12 h |
| HORVU.MOREX.r3.2HG0095710.1 | -0.89614 | -0.29298 | -3.26164 | 0.00194 | 0.022073 | -1.34533 | #N/A | elongation zone | 12 h |
| HORVU.MOREX.r3.1HG0064320.1 | -0.55232 | 6.073378 | -3.37113 | 0.001403 | 0.017964 | -1.34625 | Argininosuccinate synthase | elongation zone | 12 h |
| HORVU.MOREX.r3.5HG0442250.1 | 0.92212 | 3.294825 | 3.279356 | 0.001842 | 0.02129 | -1.34701 | Receptor-like protein kinase | elongation zone | 12 h |
| HORVU.MOREX.r3.4HG0418690.1 | -1.18844 | 0.241039 | -3.30192 | 0.001723 | 0.020463 | -1.34747 | Invertase inhibitor | elongation zone | 12 h |
| HORVU.MOREX.r3.5HG0502750.1 | -0.90944 | 6.985367 | -3.38583 | 0.001343 | 0.017454 | -1.35277 | #N/A | elongation zone | 12 h |
| HORVU.MOREX.r3.1HG0090100.1 | 1.319255 | 1.392953 | 3.255854 | 0.001973 | 0.022307 | -1.35728 | DUF1645 family protein | elongation zone | 12 h |
| HORVU.MOREX.r3.3HG0223980.1 | -0.49591 | 3.988579 | -3.29821 | 0.001742 | 0.020545 | -1.35745 | UDP-N-acetylmuramate--L-alanine ligase | elongation zone | 12 h |
| HORVU.MOREX.r3.5HG0501170.1 | 0.642768 | 0.879425 | 3.364262 | 0.001432 | 0.018164 | -1.35855 | Eukaryotic aspartyl protease family protein | elongation zone | 12 h |
| HORVU.MOREX.r3.3HG0303330.1 | -0.81135 | 6.555669 | -3.3658 | 0.001426 | 0.018116 | -1.36036 | #N/A | elongation zone | 12 h |
| HORVU.MOREX.r3.5HG0500560.2 | -1.00447 | 3.167577 | -3.25635 | 0.00197 | 0.022287 | -1.36147 | DUF630 family protein, putative (DUF630 and DUF632) | elongation zone | 12 h |
| HORVU.MOREX.r3.7HG0703580.1 | -0.62693 | 4.485599 | -3.31808 | 0.001643 | 0.019852 | -1.36217 | Glyceraldehyde-3-phosphate dehydrogenase | elongation zone | 12 h |
| HORVU.MOREX.r3.1HG0058640.1 | -1.42593 | 5.97699 | -3.30423 | 0.001711 | 0.020365 | -1.3635 | Histone H2B | elongation zone | 12 h |
| HORVU.MOREX.r3.6HG0542320.1 | 1.212697 | 2.291237 | 3.260863 | 0.001944 | 0.02208 | -1.36527 | Kinase family protein | elongation zone | 12 h |
| HORVU.MOREX.r3.2HG0201250.1 | -0.56517 | 6.181228 | -3.36851 | 0.001414 | 0.018048 | -1.36673 | GTP cyclohydrolase 1 | elongation zone | 12 h |
| HORVU.MOREX.r3.2HG0137670.1 | -1.08072 | 0.728871 | -3.26402 | 0.001926 | 0.021937 | -1.36828 | Eukaryotic translation initiation factor-related family protein | elongation zone | 12 h |
| HORVU.MOREX.r3.6HG0540280.1 | -0.83258 | 5.482685 | -3.29472 | 0.00176 | 0.020627 | -1.3696 | F-box family protein | elongation zone | 12 h |
| HORVU.MOREX.r3.2HG0208520.1 | 1.906727 | -2.76284 | 3.383407 | 0.001353 | 0.017536 | -1.37136 | Heavy metal transport/detoxification superfamily protein | elongation zone | 12 h |
| HORVU.MOREX.r3.7HG0718930.2 | 1.41421 | -0.79034 | 3.330855 | 0.001582 | 0.019414 | -1.37154 | Peroxidase | elongation zone | 12 h |
| HORVU.MOREX.r3.6HG0542640.1 | 1.180014 | 2.086759 | 3.303612 | 0.001715 | 0.020382 | -1.37541 | Plant/F1M20-13 protein | elongation zone | 12 h |
| HORVU.MOREX.r3.3HG0307400.1 | 1.281587 | 1.770343 | 3.25311 | 0.001989 | 0.022401 | -1.37576 | GRAS family transcription factor | elongation zone | 12 h |
| HORVU.MOREX.r3.2HG0170880.1 | -1.30178 | 3.770157 | -3.24796 | 0.002019 | 0.022545 | -1.37674 | Histone H2B | elongation zone | 12 h |
| HORVU.MOREX.r3.3HG0256620.1 | -1.02546 | 6.771761 | -3.33396 | 0.001567 | 0.019305 | -1.37962 | Expansin | elongation zone | 12 h |
| HORVU.MOREX.r3.2HG0124550.1 | 1.293051 | 2.362034 | 3.248702 | 0.002015 | 0.022519 | -1.3799 | Protein WEAK CHLOROPLAST MOVEMENT UNDER BLUE LIGHT 1 | elongation zone | 12 h |
| HORVU.MOREX.r3.7HG0728060.1 | 0.48326 | 5.512337 | 3.360778 | 0.001447 | 0.018262 | -1.3802 | Aminodeoxychorismate synthase | elongation zone | 12 h |
| HORVU.MOREX.r3.6HG0565950.1 | 1.393586 | -1.88296 | 3.248825 | 0.002014 | 0.022519 | -1.38387 | ROP interacting CRIB motif protein | elongation zone | 12 h |
| HORVU.MOREX.r3.1HG0043640.1 | 0.423387 | 5.423575 | 3.406197 | 0.001264 | 0.016891 | -1.38422 | Pleckstrin-like (PH) and lipid-binding START domain protein | elongation zone | 12 h |
| HORVU.MOREX.r3.7HG0657400.1 | -0.67192 | 5.076089 | -3.29559 | 0.001756 | 0.020598 | -1.38424 | Cystathionine beta-lyase, putative | elongation zone | 12 h |
| HORVU.MOREX.r3.6HG0597360.1 | -0.5114 | 5.075212 | -3.30686 | 0.001698 | 0.020239 | -1.38437 | SET domain-containing protein 9 | elongation zone | 12 h |
| HORVU.MOREX.r3.6HG0632320.1 | -0.38804 | 5.523777 | -3.35747 | 0.001462 | 0.018377 | -1.3853 | Leucine-rich repeat protein | elongation zone | 12 h |
| HORVU.MOREX.r3.5HG0472880.1 | -0.50889 | 4.879838 | -3.30997 | 0.001683 | 0.020137 | -1.38766 | GDP-mannose transporter, putative | elongation zone | 12 h |
| HORVU.MOREX.r3.7HG0710180.1 | -1.38509 | 1.647984 | -3.32045 | 0.001631 | 0.019852 | -1.39136 | related to KPC1 | elongation zone | 12 h |
| HORVU.MOREX.r3.6HG0547030.1 | 1.114231 | -2.48918 | 3.247287 | 0.002023 | 0.022545 | -1.393 | Harbinger transposase-derived nuclease | elongation zone | 12 h |
| HORVU.MOREX.r3.2HG0177830.1 | 0.620315 | 4.208721 | 3.301504 | 0.001725 | 0.020476 | -1.39498 | Transporter | elongation zone | 12 h |
| HORVU.MOREX.r3.5HG0472810.1 | 0.853867 | 2.371978 | 3.4104 | 0.001248 | 0.016756 | -1.3967 | Aquaporin | elongation zone | 12 h |
| HORVU.MOREX.r3.5HG0495640.1 | 1.909416 | -2.03513 | 3.298976 | 0.001738 | 0.020523 | -1.39993 | Solute carrier family 35 protein | elongation zone | 12 h |
| HORVU.MOREX.r3.6HG0601690.1 | -0.47857 | 4.701066 | -3.29938 | 0.001736 | 0.02051 | -1.40005 | YbaK/aminoacyl-tRNA synthetase-associated domain-containing protein | elongation zone | 12 h |
| HORVU.MOREX.r3.3HG0280960.1 | 0.686407 | 1.805356 | 3.297401 | 0.001746 | 0.02056 | -1.40291 | Glycosyltransferase | elongation zone | 12 h |
| HORVU.MOREX.r3.1HG0051960.1 | -1.17548 | 1.903866 | -3.30921 | 0.001686 | 0.020145 | -1.40514 | DNA topoisomerase 2 | elongation zone | 12 h |

**Table S1** Continued.

| ID | logFC | AveExpr | t | P.Value | adj.P.Val | B | gene description | root zone | time point |
|---|---|---|---|---|---|---|---|---|---|
| HORVU.MOREX.r3.4HG0353450.1 | -2.11518 | 2.283433 | -3.35811 | 0.001459 | 0.018354 | -1.40716 | Cell division control protein 45-like protein | elongation zone | 12 h |
| HORVU.MOREX.r3.4HG0403370.1 | -0.38828 | 4.841977 | -3.31791 | 0.001644 | 0.019852 | -1.41314 | O-fucosyltransferase family protein | elongation zone | 12 h |
| HORVU.MOREX.r3.6HG0632780.1 | -0.65717 | 1.908878 | -3.23862 | 0.002075 | 0.022947 | -1.41402 | E3 ubiquitin-protein ligase | elongation zone | 12 h |
| HORVU.MOREX.r3.6HG0602210.1 | 0.842927 | 4.713608 | 3.364083 | 0.001433 | 0.018164 | -1.41481 | Aquaporin | elongation zone | 12 h |
| HORVU.MOREX.r3.2HG0105830.1 | 0.685766 | 4.383346 | 3.282626 | 0.001824 | 0.021158 | -1.42003 | #N/A | elongation zone | 12 h |
| HORVU.MOREX.r3.3HG0306440.1 | -1.16629 | -0.68673 | -3.23502 | 0.002097 | 0.023123 | -1.42029 | Serine/threonine-protein kinase | elongation zone | 12 h |
| HORVU.MOREX.r3.2HG0214130.1 | 1.256699 | 3.799443 | 3.324615 | 0.001611 | 0.019705 | -1.42096 | CoA ligase | elongation zone | 12 h |
| HORVU.MOREX.r3.7HG0742450.1 | -0.64112 | 3.881739 | -3.24892 | 0.002014 | 0.022519 | -1.42205 | HVA22-like protein | elongation zone | 12 h |
| HORVU.MOREX.r3.6HG0554520.1 | -0.98749 | 5.131199 | -3.25077 | 0.002003 | 0.022493 | -1.42477 | BRCT domain-containing protein | elongation zone | 12 h |
| HORVU.MOREX.r3.3HG0230640.1 | 1.844244 | -1.25694 | 3.234378 | 0.002101 | 0.023141 | -1.42797 | Thioesterase family protein | elongation zone | 12 h |
| HORVU.MOREX.r3.6HG0631580.1 | -0.57234 | 4.298528 | -3.2714 | 0.001885 | 0.021637 | -1.4283 | Protein SMG8 | elongation zone | 12 h |
| HORVU.MOREX.r3.7HG0681050.1 | 0.429679 | 4.075358 | 3.346504 | 0.00151 | 0.018825 | -1.42944 | Chitinase | elongation zone | 12 h |
| HORVU.MOREX.r3.3HG0236500.1 | 0.670059 | 5.370928 | 3.331683 | 0.001578 | 0.019378 | -1.42948 | Pollen-specific protein SF21 | elongation zone | 12 h |
| HORVU.MOREX.r3.7HG0658270.1 | -0.56183 | 2.885178 | -3.26844 | 0.001902 | 0.021787 | -1.42994 | Hyp O-arabinosyltransferase-like protein | elongation zone | 12 h |
| HORVU.MOREX.r3.5HG0485410.1 | 2.429921 | -0.44467 | 3.762735 | 0.000422 | 0.009111 | -1.43019 | #N/A | elongation zone | 12 h |
| HORVU.MOREX.r3.2HG0215220.1 | -0.74946 | 4.050855 | -3.2523 | 0.001994 | 0.022429 | -1.43219 | Tubby-like F-box protein | elongation zone | 12 h |
| HORVU.MOREX.r3.6HG0543790.1 | -1.06692 | 7.131519 | -3.34657 | 0.00151 | 0.018825 | -1.43324 | Histone H2A | elongation zone | 12 h |
| HORVU.MOREX.r3.3HG0219380.1 | 0.946576 | 4.367645 | 3.400258 | 0.001286 | 0.01705 | -1.43376 | Glutamate receptor ionotropic, NMDA 3A | elongation zone | 12 h |
| HORVU.MOREX.r3.1HG0020270.1 | -0.37399 | 5.457119 | -3.31822 | 0.001642 | 0.019852 | -1.43695 | SAD1/UNC-84 domain protein, putative | elongation zone | 12 h |
| HORVU.MOREX.r3.3HG0270070.1 | -0.87242 | 2.011026 | -3.22561 | 0.002155 | 0.023536 | -1.43849 | Polyadenylation and cleavage factor homolog 1 | elongation zone | 12 h |
| HORVU.MOREX.r3.6HG0626650.1 | -0.25016 | 8.259289 | -3.43821 | 0.001148 | 0.015954 | -1.43866 | 26S protease regulatory subunit-like protein | elongation zone | 12 h |
| HORVU.MOREX.r3.1HG0078910.1 | 0.523941 | 4.676362 | 3.358439 | 0.001457 | 0.018347 | -1.43946 | p-loop containing nucleoside triphosphate hydrolases superfamily protein | elongation zone | 12 h |
| HORVU.MOREX.r3.7HG0634660.1 | -0.57651 | 4.623502 | -3.2961 | 0.001753 | 0.020579 | -1.44227 | Disease resistance protein (NBS-LRR class) family | elongation zone | 12 h |
| HORVU.MOREX.r3.3HG0254850.1 | -0.33702 | 4.808159 | -3.29628 | 0.001752 | 0.020579 | -1.4429 | sucrose synthase 1 | elongation zone | 12 h |
| HORVU.MOREX.r3.2HG0134220.1 | -0.33399 | 5.083549 | -3.3319 | 0.001577 | 0.019378 | -1.44293 | Exostosin family protein-like protein | elongation zone | 12 h |
| HORVU.MOREX.r3.6HG0573290.1 | 0.319681 | 6.719302 | 3.392113 | 0.001318 | 0.017237 | -1.44383 | Zinc finger A20 and AN1 domain stress-associated protein | elongation zone | 12 h |
| HORVU.MOREX.r3.2HG0141490.1 | -0.66815 | 3.164747 | -3.23447 | 0.0021 | 0.023141 | -1.4443 | ATP synthase gamma chain | elongation zone | 12 h |
| HORVU.MOREX.r3.2HG0113700.1 | -1.12158 | 4.9871 | -3.29193 | 0.001775 | 0.020739 | -1.4457 | Rubber elongation factor family protein | elongation zone | 12 h |
| HORVU.MOREX.r3.5HG0516490.2 | -0.5225 | 3.181218 | -3.24298 | 0.002049 | 0.022752 | -1.44606 | #N/A | elongation zone | 12 h |
| HORVU.MOREX.r3.7HG0698910.1 | -0.39278 | 6.327155 | -3.35316 | 0.00148 | 0.018535 | -1.44844 | DnaJ homolog subfamily C member 2 | elongation zone | 12 h |
| HORVU.MOREX.r3.5HG0529120.1 | -0.71326 | 7.801287 | -3.36603 | 0.001425 | 0.018114 | -1.45139 | Elongation factor | elongation zone | 12 h |
| HORVU.MOREX.r3.7HG0723240.1 | 3.038616 | -0.72296 | 3.443958 | 0.001128 | 0.015774 | -1.45358 | TMV response-related protein | elongation zone | 12 h |
| HORVU.MOREX.r3.2HG0131280.1 | 0.892256 | 2.514635 | 3.241325 | 0.002059 | 0.022811 | -1.4554 | Homeobox-leucine zipper family protein | elongation zone | 12 h |
| HORVU.MOREX.r3.2HG0209680.1 | 1.308685 | 0.534924 | 3.232343 | 0.002113 | 0.023229 | -1.45773 | Isopropylmalate dehydrogenase | elongation zone | 12 h |
| HORVU.MOREX.r3.2HG0121530.1 | -1.05297 | 2.335771 | -3.22485 | 0.00216 | 0.023553 | -1.45824 | B3 domain-containing protein | elongation zone | 12 h |
| HORVU.MOREX.r3.5HG0525560.1 | -0.50842 | 4.392032 | -3.2489 | 0.002014 | 0.022519 | -1.46037 | PHD finger alfin-like protein | elongation zone | 12 h |
| HORVU.MOREX.r3.2HG0164020.1 | 0.756723 | 1.589544 | 3.221437 | 0.002182 | 0.0237 | -1.46098 | E3 ubiquitin protein ligase drip2 | elongation zone | 12 h |
| HORVU.MOREX.r3.7HG0671470.1 | -0.47825 | 4.81657 | -3.3407 | 0.001536 | 0.019048 | -1.46526 | Protein trichome birefringence | elongation zone | 12 h |
| HORVU.MOREX.r3.7HG0742470.1 | -0.87696 | 2.928405 | -3.21224 | 0.002241 | 0.024012 | -1.46625 | Zinc finger protein | elongation zone | 12 h |
| HORVU.MOREX.r3.4HG0417670.1 | 1.192284 | 0.032065 | 3.313765 | 0.001664 | 0.019979 | -1.46809 | Lipid transfer protein | elongation zone | 12 h |
| HORVU.MOREX.r3.4HG0364520.1 | -0.38018 | 4.639723 | -3.29997 | 0.001733 | 0.020487 | -1.46935 | Nuclear factor Y subunit C | elongation zone | 12 h |

**Table S1** Continued.

| ID | logFC | AveExpr | t | P.Value | adj.P.Val | B | gene description | root zone | time point |
|---|---|---|---|---|---|---|---|---|---|
| HORVU.MOREX.r3.1HG0016560.1 | -1.42562 | 0.290348 | -3.38162 | 0.00136 | 0.017585 | -1.46951 | Beta-carotene isomerase d27, chloroplastic | elongation zone | 12 h |
| HORVU.MOREX.r3.2HG0208170.1 | 1.562628 | -2.43737 | 3.219017 | 0.002197 | 0.023779 | -1.46999 | Non-specific serine/threonine protein kinase | elongation zone | 12 h |
| HORVU.MOREX.r3.7HG0638370.1 | 0.729296 | 4.319061 | 3.300274 | 0.001732 | 0.020487 | -1.47071 | Membrane protein of ER body-like protein | elongation zone | 12 h |
| HORVU.MOREX.r3.5HG0446330.1 | -0.71471 | 1.961051 | -3.20836 | 0.002266 | 0.02416 | -1.47322 | SAGA-associated factor 11 | elongation zone | 12 h |
| HORVU.MOREX.r3.6HG0597790.1 | 0.738277 | 2.912708 | 3.254169 | 0.001983 | 0.022356 | -1.47396 | GRAM domain-containing protein / ABA-responsive | elongation zone | 12 h |
| HORVU.MOREX.r3.5HG0537000.1 | 1.008214 | -2.10915 | 3.213349 | 0.002234 | 0.023973 | -1.47412 | Oxidoreductase/transition metal ion-binding protein | elongation zone | 12 h |
| HORVU.MOREX.r3.3HG0267000.1 | 0.819897 | 3.929222 | 3.40524 | 0.001267 | 0.016918 | -1.47419 | Regulator of chromosome condensation (RCC1) family protein | elongation zone | 12 h |
| HORVU.MOREX.r3.2HG0167500.2 | -1.67188 | 4.031522 | -3.20935 | 0.00226 | 0.02416 | -1.47501 | Repetitive proline-rich cell wall protein | elongation zone | 12 h |
| HORVU.MOREX.r3.4HG0383780.1 | 1.804277 | -0.06269 | 3.236303 | 0.002089 | 0.023073 | -1.47646 | Laccase | elongation zone | 12 h |
| HORVU.MOREX.r3.2HG0193170.1 | 4.478357 | -0.2671 | 3.556571 | 0.000801 | 0.013114 | -1.47762 | Pollen Ole e 1 allergen/extensin | elongation zone | 12 h |
| HORVU.MOREX.r3.7HG0727980.1 | 0.597201 | 1.541509 | 3.338026 | 0.001549 | 0.019165 | -1.47868 | ENTH/ANTH/VHS superfamily protein | elongation zone | 12 h |
| HORVU.MOREX.r3.7HG0702190.1 | -1.63898 | 3.843343 | -3.24151 | 0.002058 | 0.022811 | -1.47899 | Fertilization independent endosperm 1 protein | elongation zone | 12 h |
| HORVU.MOREX.r3.7HG0668850.1 | -0.61305 | 6.214378 | -3.34168 | 0.001532 | 0.019016 | -1.4793 | UDP-glycosyltransferase | elongation zone | 12 h |
| HORVU.MOREX.r3.4HG0333540.1 | 0.94212 | 3.467572 | 3.23987 | 0.002068 | 0.022883 | -1.48121 | Acetyltransferase, GNAT family protein, expressed | elongation zone | 12 h |
| HORVU.MOREX.r3.6HG0615130.1 | 0.689772 | 3.141376 | 3.334292 | 0.001566 | 0.019305 | -1.48251 | TCP transcription factor | elongation zone | 12 h |
| HORVU.MOREX.r3.3HG0316000.1 | -0.73856 | 6.548487 | -3.31122 | 0.001677 | 0.020106 | -1.48394 | Salicylate O-methyltransferase | elongation zone | 12 h |
| HORVU.MOREX.r3.1HG0072840.1 | -0.36914 | 5.890668 | -3.33543 | 0.001561 | 0.019256 | -1.48413 | Ectonucleoside triphosphate diphosphohydrolase 1 | elongation zone | 12 h |
| HORVU.MOREX.r3.6HG0634260.1 | -0.75186 | 4.589475 | -3.22628 | 0.002151 | 0.023517 | -1.48635 | TSA: Wollemia nobilis Ref_Wollemi_Transcript_29242_1275 transcribed RNA sequ | elongation zone | 12 h |
| HORVU.MOREX.r3.5HG0509460.1 | -0.94485 | 2.420051 | -3.21613 | 0.002216 | 0.023861 | -1.48646 | WRKY transcription factor | elongation zone | 12 h |
| HORVU.MOREX.r3.6HG0617860.1 | -0.29822 | 5.538253 | -3.33004 | 0.001586 | 0.019449 | -1.48693 | Peptidyl-prolyl cis-trans isomerase | elongation zone | 12 h |
| HORVU.MOREX.r3.7HG0634520.1 | 1.163989 | -2.23987 | 3.205429 | 0.002286 | 0.0242 | -1.48743 | NBS-LRR-like resistance protein | elongation zone | 12 h |
| HORVU.MOREX.r3.4HG0410720.1 | -0.86024 | 2.267394 | -3.20987 | 0.002256 | 0.024141 | -1.48962 | Alpha-2C adrenergic receptor | elongation zone | 12 h |
| HORVU.MOREX.r3.7HG0719070.1 | -0.68554 | 5.233224 | -3.24143 | 0.002058 | 0.022811 | -1.49118 | SUMO-activating enzyme subunit 1A | elongation zone | 12 h |
| HORVU.MOREX.r3.5HG0492650.1 | -1.03647 | 7.121607 | -3.31462 | 0.00166 | 0.01994 | -1.49128 | O-fucosyltransferase family protein | elongation zone | 12 h |
| HORVU.MOREX.r3.7HG0721930.1 | -0.55565 | 4.955958 | -3.26019 | 0.001948 | 0.022103 | -1.49469 | B-cell receptor-associated protein 31-like containing protein | elongation zone | 12 h |
| HORVU.MOREX.r3.2HG0191510.1 | 0.841887 | -0.88853 | 3.250404 | 0.002005 | 0.022505 | -1.49482 | transmembrane protein, putative (DUF247) | elongation zone | 12 h |
| HORVU.MOREX.r3.7HG0719830.1 | -1.48299 | 0.487996 | -3.22444 | 0.002163 | 0.023553 | -1.49496 | #N/A | elongation zone | 12 h |
| HORVU.MOREX.r3.7HG0716170.1 | -1.01362 | 2.519767 | -3.21671 | 0.002212 | 0.023852 | -1.49544 | F-box family protein | elongation zone | 12 h |
| HORVU.MOREX.r3.2HG0096800.1 | 0.690551 | 4.6918 | 3.296775 | 0.00175 | 0.020579 | -1.49861 | Flavin-containing monooxygenase | elongation zone | 12 h |
| HORVU.MOREX.r3.5HG0503200.1 | -0.60979 | 3.671203 | -3.22349 | 0.002169 | 0.023571 | -1.50176 | Bromodomain-containing protein | elongation zone | 12 h |
| HORVU.MOREX.r3.6HG0564690.1 | 0.41004 | 5.759195 | 3.373257 | 0.001394 | 0.017883 | -1.50581 | Regulator of Vps4 activity in the MVB pathway protein, putative | elongation zone | 12 h |
| HORVU.MOREX.r3.1HG0080230.1 | -1.09595 | 6.650954 | -3.29995 | 0.001733 | 0.020487 | -1.50605 | #N/A | elongation zone | 12 h |
| HORVU.MOREX.r3.5HG0459680.1 | -0.38438 | 3.895086 | -3.24766 | 0.002021 | 0.022545 | -1.50658 | RING/U-box superfamily protein | elongation zone | 12 h |
| HORVU.MOREX.r3.1HG0007370.1 | -0.48707 | 3.297626 | -3.21996 | 0.002191 | 0.023739 | -1.50865 | B3 domain-containing protein | elongation zone | 12 h |
| HORVU.MOREX.r3.3HG0285270.1 | -2.00538 | 0.025552 | -3.33896 | 0.001544 | 0.019135 | -1.51033 | Guanine nucleotide exchange family protein | elongation zone | 12 h |
| HORVU.MOREX.r3.2HG0111130.1 | -0.77583 | 3.835075 | -3.19939 | 0.002326 | 0.024453 | -1.51168 | Short chain dehydrogenase/reductase | elongation zone | 12 h |
| HORVU.MOREX.r3.5HG0526200.1 | -0.26585 | 5.485729 | -3.3059 | 0.001703 | 0.020282 | -1.51257 | Sec14 cytosolic factor | elongation zone | 12 h |
| HORVU.MOREX.r3.3HG0302860.1 | -1.95639 | 4.061598 | -3.19826 | 0.002334 | 0.024484 | -1.51326 | Pathogenesis-related thaumatin-like protein | elongation zone | 12 h |
| HORVU.MOREX.r3.7HG0734380.1 | -0.47567 | 4.659788 | -3.26683 | 0.001911 | 0.021793 | -1.51346 | Mechanosensitive ion channel-like protein | elongation zone | 12 h |
| HORVU.MOREX.r3.5HG0465270.1 | 0.741706 | 4.111151 | 3.414849 | 0.001231 | 0.01661 | -1.5135 | Laccase | elongation zone | 12 h |
| HORVU.MOREX.r3.4HG0389410.1 | 0.751985 | 2.32327 | 3.194799 | 0.002357 | 0.024642 | -1.5145 | SH3 domain-containing protein 1 | elongation zone | 12 h |

**Table S1** Continued.

| ID | logFC | AveExpr | t | P.Value | adj.P.Val | B | gene description | root zone | time point |
|---|---|---|---|---|---|---|---|---|---|
| HORVU.MOREX.r3.4HG0393520.1 | 0.74874 | 3.79819 | 3.244959 | 0.002037 | 0.022668 | -1.51461 | Microtubule associated protein family protein, putative, expressed | elongation zone | 12 h |
| HORVU.MOREX.r3.6HG0539410.1 | -1.74118 | 1.804569 | -3.27822 | 0.001848 | 0.021343 | -1.51748 | LOW protein: UPF0503-like protein, putative (DUF740) | elongation zone | 12 h |
| HORVU.MOREX.r3.7HG0667150.1 | 0.91636 | -0.51034 | 3.230115 | 0.002127 | 0.023343 | -1.51771 | Rac-like GTP-binding protein | elongation zone | 12 h |
| HORVU.MOREX.r3.6HG0623140.1 | 1.922545 | -2.34532 | 3.319168 | 0.001638 | 0.019852 | -1.51824 | #N/A | elongation zone | 12 h |
| HORVU.MOREX.r3.7HG0660660.1 | -0.48051 | 4.772651 | -3.25383 | 0.001985 | 0.022366 | -1.51857 | 26S protease regulatory subunit | elongation zone | 12 h |
| HORVU.MOREX.r3.6HG0617080.1 | 0.507097 | 4.45228 | 3.323498 | 0.001617 | 0.019747 | -1.51956 | Lipid transfer protein | elongation zone | 12 h |
| HORVU.MOREX.r3.3HG0239940.1 | -0.60537 | 3.204602 | -3.21826 | 0.002202 | 0.023794 | -1.52002 | Transmembrane protein 19 | elongation zone | 12 h |
| HORVU.MOREX.r3.1HG0056570.1 | 0.515125 | 7.070581 | 3.48484 | 0.000997 | 0.014726 | -1.52096 | Expansin protein | elongation zone | 12 h |
| HORVU.MOREX.r3.7HG0749640.1 | 0.613005 | 4.119977 | 3.275536 | 0.001862 | 0.021446 | -1.521 | Transmembrane protein 53 | elongation zone | 12 h |
| HORVU.MOREX.r3.7HG0749080.1 | 1.128564 | -1.66836 | 3.199606 | 0.002325 | 0.02445 | -1.52236 | G-protein coupled receptor 98 | elongation zone | 12 h |
| HORVU.MOREX.r3.4HG0354540.1 | -2.4083 | 4.811312 | -3.1987 | 0.002331 | 0.024477 | -1.52409 | Receptor-like kinase | elongation zone | 12 h |
| HORVU.MOREX.r3.6HG0614350.1 | -0.52437 | 8.627624 | -3.44254 | 0.001133 | 0.015779 | -1.52455 | Lipoxygenase domain-containing 1 | elongation zone | 12 h |
| HORVU.MOREX.r3.3HG0230500.1 | 1.226877 | 1.089665 | 3.206526 | 0.002278 | 0.024194 | -1.52468 | NBS-LRR disease resistance protein | elongation zone | 12 h |
| HORVU.MOREX.r3.4HG0355460.1 | -0.39537 | 4.5138 | -3.29611 | 0.001753 | 0.020579 | -1.52575 | myosin-4 protein (DUF641) | elongation zone | 12 h |
| HORVU.MOREX.r3.7HG0641160.1 | -0.37998 | 5.342895 | -3.28213 | 0.001827 | 0.021176 | -1.52711 | Guanine nucleotide-binding protein subunit alpha-like protein | elongation zone | 12 h |
| HORVU.MOREX.r3.7HG0668440.1 | 0.581003 | 5.028289 | 3.30683 | 0.001698 | 0.020239 | -1.52749 | #N/A | elongation zone | 12 h |
| HORVU.MOREX.r3.1HG0060240.1 | -0.83038 | 2.446107 | -3.18627 | 0.002416 | 0.025092 | -1.53188 | Thiamin pyrophosphokinase 1 | elongation zone | 12 h |
| HORVU.MOREX.r3.1HG0082750.1 | -1.89983 | 0.305019 | -3.3858 | 0.001343 | 0.017454 | -1.53298 | Ring finger protein, putative | elongation zone | 12 h |
| HORVU.MOREX.r3.5HG0461400.1 | -1.30105 | 2.587492 | -3.20813 | 0.002268 | 0.02416 | -1.53541 | UDP-glycosyltransferase | elongation zone | 12 h |
| HORVU.MOREX.r3.7HG0644240.1 | 1.20748 | -0.03364 | 3.198881 | 0.00233 | 0.024477 | -1.53555 | F-box protein PP2 | elongation zone | 12 h |
| HORVU.MOREX.r3.2HG0210500.1 | 1.219187 | 3.185342 | 3.41642 | 0.001225 | 0.016542 | -1.53686 | Endoglucanase | elongation zone | 12 h |
| HORVU.MOREX.r3.7HG0747690.1 | 3.568482 | 0.558808 | 3.612775 | 0.000674 | 0.011946 | -1.53734 | Hydroxycinnamoyl-CoA shikimate/quinate hydroxycinnamoyltransferase | elongation zone | 12 h |
| HORVU.MOREX.r3.7HG0647550.1 | -0.50875 | 4.057694 | -3.22015 | 0.00219 | 0.023739 | -1.5409 | bZIP transcription factor, putative (DUF630 and DUF632) | elongation zone | 12 h |
| HORVU.MOREX.r3.2HG0154760.1 | -0.91302 | 1.184214 | -3.18221 | 0.002445 | 0.025246 | -1.54153 | cysteine-rich RLK (RECEPTOR-like protein kinase) 11 | elongation zone | 12 h |
| HORVU.MOREX.r3.4HG0379260.1 | 0.636392 | 6.080607 | 3.369825 | 0.001409 | 0.018011 | -1.54182 | Calcium binding protein | elongation zone | 12 h |
| HORVU.MOREX.r3.1HG0069870.1 | 0.680526 | 2.552246 | 3.260128 | 0.001949 | 0.022103 | -1.54227 | #N/A | elongation zone | 12 h |
| HORVU.MOREX.r3.4HG0402650.1 | 0.825072 | 1.552903 | 3.242955 | 0.002049 | 0.022752 | -1.54237 | Annexin | elongation zone | 12 h |
| HORVU.MOREX.r3.6HG0558810.1 | -0.96612 | 5.76557 | -3.2678 | 0.001905 | 0.021791 | -1.54244 | Histone H2A | elongation zone | 12 h |
| HORVU.MOREX.r3.3HG0244000.1 | 1.683681 | -2.4581 | 3.326078 | 0.001604 | 0.019655 | -1.54308 | Core-2/I-branching beta-1,6-N-acetylglucosaminyltransferase family protein | elongation zone | 12 h |
| HORVU.MOREX.r3.5HG0450470.1 | -0.55904 | 3.631343 | -3.19713 | 0.002341 | 0.024507 | -1.54499 | Basic-leucine zipper (bZIP) transcription factor family protein | elongation zone | 12 h |
| HORVU.MOREX.r3.1HG0095170.1 | 1.57227 | 5.086192 | 3.255443 | 0.001975 | 0.022314 | -1.54503 | UDP-glucose-4-epimerase | elongation zone | 12 h |
| HORVU.MOREX.r3.4HG0386490.1 | 1.786727 | -0.76068 | 3.257435 | 0.001964 | 0.022241 | -1.54751 | Receptor-like kinase, putative | elongation zone | 12 h |
| HORVU.MOREX.r3.6HG0547020.1 | 1.15425 | -2.68848 | 3.201668 | 0.002311 | 0.024379 | -1.54896 | #N/A | elongation zone | 12 h |
| HORVU.MOREX.r3.2HG0195690.1 | 1.074979 | -2.81043 | 3.233481 | 0.002106 | 0.023189 | -1.54944 | filamentous hemagglutinin transporter | elongation zone | 12 h |
| HORVU.MOREX.r3.3HG0294840.1 | -0.52459 | 3.760264 | -3.21369 | 0.002231 | 0.023972 | -1.54961 | Haloacid dehalogenase (HAD) superfamily protein | elongation zone | 12 h |
| HORVU.MOREX.r3.5HG0525850.1 | 0.685206 | 2.978562 | 3.247369 | 0.002023 | 0.022545 | -1.55241 | Phosphatidylinositol N-acetyglucosaminlytransferase subunit P-like protein | elongation zone | 12 h |
| HORVU.MOREX.r3.6HG0628790.1 | 0.728557 | 2.686977 | 3.191279 | 0.002381 | 0.024831 | -1.55682 | Myb-like transcription factor family protein | elongation zone | 12 h |
| HORVU.MOREX.r3.4HG0385300.1 | -0.49922 | 9.307024 | -3.40323 | 0.001275 | 0.016993 | -1.55972 | Elongation factor 1-alpha | elongation zone | 12 h |
| HORVU.MOREX.r3.2HG0190710.1 | -0.34368 | 4.642345 | -3.2487 | 0.002015 | 0.022519 | -1.55988 | #N/A | elongation zone | 12 h |
| HORVU.MOREX.r3.3HG0295640.1 | 1.213561 | -1.27257 | 3.201458 | 0.002312 | 0.024382 | -1.56006 | Protein MIZU-KUSSEI 1 | elongation zone | 12 h |
| HORVU.MOREX.r3.3HG0303660.1 | 0.54769 | 6.126297 | 3.365016 | 0.001429 | 0.018147 | -1.56071 | Carboxypeptidase | elongation zone | 12 h |

**Table S1** Continued.

| ID | logFC | AveExpr | t | P.Value | adj.P.Val | B | gene description | root zone | time point |
|---|---|---|---|---|---|---|---|---|---|
| HORVU.MOREX.r3.1HG0003090.4 | -1.69798 | 2.96451 | -3.19774 | 0.002337 | 0.024495 | -1.56196 | Thionin-2.2 | elongation zone | 12 h |
| HORVU.MOREX.r3.2HG0214070.1 | 1.618596 | -2.62397 | 3.231547 | 0.002118 | 0.023258 | -1.56367 | AT hook motif DNA-binding family protein, putative | elongation zone | 12 h |
| HORVU.MOREX.r3.3HG0219810.1 | -0.91791 | 3.950558 | -3.17152 | 0.002522 | 0.025832 | -1.564 | Agenet domain containing protein | elongation zone | 12 h |
| HORVU.MOREX.r3.5HG0520260.1 | -2.4668 | 1.284182 | -3.48643 | 0.000992 | 0.014705 | -1.56461 | GTPase obg | elongation zone | 12 h |
| HORVU.MOREX.r3.5HG0491380.1 | -0.53889 | 8.151632 | -3.39243 | 0.001317 | 0.017232 | -1.56462 | Proton pump-interactor 1 | elongation zone | 12 h |
| HORVU.MOREX.r3.7HG0705500.1 | 1.096897 | 0.351075 | 3.182922 | 0.00244 | 0.025209 | -1.56826 | Non-lysosomal glucosylceramidase | elongation zone | 12 h |
| HORVU.MOREX.r3.5HG0472870.1 | 0.639122 | 3.59022 | 3.266934 | 0.00191 | 0.021793 | -1.56906 | Disease resistance protein (NBS-LRR class) family | elongation zone | 12 h |
| HORVU.MOREX.r3.2HG0199590.1 | -1.26472 | -1.68344 | -3.20345 | 0.002299 | 0.024291 | -1.56997 | Organic cation transporter protein | elongation zone | 12 h |
| HORVU.MOREX.r3.7HG0676880.1 | -0.38292 | 5.477475 | -3.27137 | 0.001885 | 0.021637 | -1.57119 | BolA-like | elongation zone | 12 h |
| HORVU.MOREX.r3.7HG0654810.1 | -2.40425 | -1.68567 | -3.73461 | 0.000461 | 0.009584 | -1.57122 | actin cross-linking protein (DUF569) | elongation zone | 12 h |
| HORVU.MOREX.r3.4HG0407470.1 | 0.893497 | 2.42529 | 3.315449 | 0.001656 | 0.019915 | -1.57625 | Serpin | elongation zone | 12 h |
| HORVU.MOREX.r3.5HG0483880.1 | -1.60123 | -0.16233 | -3.37584 | 0.001384 | 0.017779 | -1.57664 | RING/U-box superfamily protein | elongation zone | 12 h |
| HORVU.MOREX.r3.1HG0063020.1 | 1.295173 | 0.508236 | 3.276675 | 0.001856 | 0.021386 | -1.5777 | Resistance to phytophthora 1 protein | elongation zone | 12 h |
| HORVU.MOREX.r3.7HG0655100.1 | 0.526096 | 3.752132 | 3.277469 | 0.001852 | 0.02136 | -1.57794 | Peroxyureidoacrylate/ureidoacrylate amidohydrolase RutB | elongation zone | 12 h |
| HORVU.MOREX.r3.2HG0200980.1 | -0.40613 | 5.67298 | -3.29513 | 0.001758 | 0.020614 | -1.57928 | Peptidyl-prolyl cis-trans isomerase | elongation zone | 12 h |
| HORVU.MOREX.r3.3HG0304420.1 | -1.07403 | 5.851635 | -3.23855 | 0.002076 | 0.022947 | -1.58016 | #N/A | elongation zone | 12 h |
| HORVU.MOREX.r3.6HG0616960.1 | -0.61419 | 4.492257 | -3.20033 | 0.00232 | 0.024424 | -1.58047 | Polyadenylate-binding protein family protein | elongation zone | 12 h |
| HORVU.MOREX.r3.3HG0325670.1 | -0.62984 | 3.462193 | -3.17531 | 0.002494 | 0.025643 | -1.58151 | #N/A | elongation zone | 12 h |
| HORVU.MOREX.r3.5HG0468460.1 | -0.60009 | 6.084231 | -3.27813 | 0.001848 | 0.021343 | -1.5842 | FCA protein | elongation zone | 12 h |
| HORVU.MOREX.r3.2HG0111620.1 | 0.757251 | 2.992838 | 3.414201 | 0.001234 | 0.016631 | -1.58494 | Heavy metal transport/detoxification superfamily protein | elongation zone | 12 h |
| HORVU.MOREX.r3.1HG0070850.1 | 0.89347 | 0.443712 | 3.225875 | 0.002154 | 0.023533 | -1.5857 | Extra-large G-like protein, putative (DUF3133) | elongation zone | 12 h |
| HORVU.MOREX.r3.2HG0177440.1 | -0.84542 | 5.076842 | -3.23326 | 0.002108 | 0.023192 | -1.58661 | Ankyrin repeat | elongation zone | 12 h |
| HORVU.MOREX.r3.6HG0633650.1 | -1.17675 | 0.935723 | -3.21759 | 0.002206 | 0.023823 | -1.58701 | #N/A | elongation zone | 12 h |
| HORVU.MOREX.r3.1HG0079760.1 | 0.796728 | 0.18048 | 3.228958 | 0.002134 | 0.023397 | -1.58789 | MYB transcription factor | elongation zone | 12 h |
| HORVU.MOREX.r3.6HG0557340.1 | -0.47246 | 7.977821 | -3.36917 | 0.001411 | 0.018035 | -1.58879 | Terpene cyclase/mutase family member | elongation zone | 12 h |
| HORVU.MOREX.r3.5HG0446350.1 | -0.36156 | 5.803999 | -3.27372 | 0.001872 | 0.021548 | -1.59024 | RNA polymerase-associated protein RTF1-like protein | elongation zone | 12 h |
| HORVU.MOREX.r3.2HG0119800.1 | 1.533778 | -1.32815 | 3.185614 | 0.002421 | 0.025127 | -1.59056 | GDSL esterase/lipase | elongation zone | 12 h |
| HORVU.MOREX.r3.5HG0486330.1 | -0.3159 | 5.745088 | -3.27303 | 0.001876 | 0.021568 | -1.59104 | UPF0396 protein | elongation zone | 12 h |
| HORVU.MOREX.r3.2HG0208960.1 | -0.88158 | 2.446722 | -3.15948 | 0.002611 | 0.026441 | -1.59113 | Exocyst complex component EXO70B1 | elongation zone | 12 h |
| HORVU.MOREX.r3.3HG0305520.1 | 0.695335 | 1.413662 | 3.164047 | 0.002577 | 0.026239 | -1.59254 | DUF1685 family protein | elongation zone | 12 h |
| HORVU.MOREX.r3.1HG0018730.1 | -0.32082 | 6.708783 | -3.30901 | 0.001687 | 0.020145 | -1.59419 | RNA-binding family protein | elongation zone | 12 h |
| HORVU.MOREX.r3.1HG0040930.1 | 1.803732 | -2.88506 | 3.317077 | 0.001648 | 0.019877 | -1.59493 | F-box family protein | elongation zone | 12 h |
| HORVU.MOREX.r3.2HG0200630.1 | -0.77758 | 9.635165 | -3.39279 | 0.001315 | 0.017224 | -1.59608 | Adenosylhomocysteinase | elongation zone | 12 h |
| HORVU.MOREX.r3.7HG0680960.1 | 1.615506 | 0.611498 | 3.158215 | 0.00262 | 0.026447 | -1.59625 | electron transporter, putative (Protein of unknown function, DUF547) | elongation zone | 12 h |
| HORVU.MOREX.r3.1HG0087190.1 | -0.56194 | 4.047935 | -3.18442 | 0.002429 | 0.025148 | -1.59666 | Structural constituent of cell wall | elongation zone | 12 h |
| HORVU.MOREX.r3.6HG0620240.1 | -0.50722 | 3.057466 | -3.18977 | 0.002392 | 0.024902 | -1.59698 | FRIGIDA interacting protein 1 | elongation zone | 12 h |
| HORVU.MOREX.r3.5HG0435800.1 | -2.09664 | 0.267696 | -3.40101 | 0.001283 | 0.01705 | -1.60142 | Leucine-rich repeat protein kinase family protein | elongation zone | 12 h |
| HORVU.MOREX.r3.5HG0437110.1 | 0.508929 | 5.656876 | 3.340786 | 0.001536 | 0.019048 | -1.60484 | Kinase family protein | elongation zone | 12 h |
| HORVU.MOREX.r3.6HG0550690.1 | -1.02142 | 4.80697 | -3.20765 | 0.002271 | 0.02416 | -1.60535 | #N/A | elongation zone | 12 h |
| HORVU.MOREX.r3.6HG0619260.1 | -0.4589 | 6.188825 | -3.28023 | 0.001837 | 0.021272 | -1.60898 | Transmembrane emp24 domain-containing protein | elongation zone | 12 h |
| HORVU.MOREX.r3.6HG0543320.1 | -0.68552 | 3.154801 | -3.16011 | 0.002606 | 0.026433 | -1.60921 | Mitochondrial transcription termination factor-like | elongation zone | 12 h |

**Table S1** Continued.

| ID | logFC | AveExpr | t | P.Value | adj.P.Val | B | gene description | root zone | time point |
|---|---|---|---|---|---|---|---|---|---|
| HORVU.MOREX.r3.5HG0495860.1 | 0.76335 | 4.058331 | 3.267239 | 0.001908 | 0.021791 | -1.61031 | glycosyltransferase family exostosin protein | elongation zone | 12 h |
| HORVU.MOREX.r3.7HG0723420.1 | -0.37736 | 6.130958 | -3.28395 | 0.001817 | 0.0211 | -1.61057 | Eukaryotic aspartyl protease family protein | elongation zone | 12 h |
| HORVU.MOREX.r3.1HG0082530.1 | 0.468307 | 7.395893 | 3.393551 | 0.001312 | 0.017213 | -1.61063 | Respiratory burst oxidase | elongation zone | 12 h |
| HORVU.MOREX.r3.4HG0358890.1 | -0.57961 | 6.54155 | -3.26739 | 0.001907 | 0.021791 | -1.61183 | #N/A | elongation zone | 12 h |
| HORVU.MOREX.r3.3HG0285010.1 | -2.01788 | 1.558759 | -3.28799 | 0.001795 | 0.020933 | -1.61464 | indeterminate(ID)-domain 4 | elongation zone | 12 h |
| HORVU.MOREX.r3.3HG0308840.1 | 0.62646 | 2.063856 | 3.184294 | 0.00243 | 0.025148 | -1.61732 | Aldehyde dehydrogenase, putative | elongation zone | 12 h |
| HORVU.MOREX.r3.6HG0593390.1 | 1.365032 | 0.157725 | 3.15156 | 0.002671 | 0.026814 | -1.61747 | Bidirectional sugar transporter SWEET | elongation zone | 12 h |
| HORVU.MOREX.r3.6HG0558880.1 | -1.29199 | 5.027918 | -3.17399 | 0.002504 | 0.02572 | -1.62659 | Histone H2A | elongation zone | 12 h |
| HORVU.MOREX.r3.7HG0747720.1 | 0.693604 | 3.70009 | 3.18407 | 0.002432 | 0.025148 | -1.62866 | Cytokinin oxidase/dehydrogenase | elongation zone | 12 h |
| HORVU.MOREX.r3.2HG0180500.1 | 0.363372 | 5.818938 | 3.300639 | 0.00173 | 0.020487 | -1.62868 | Glutaredoxin | elongation zone | 12 h |
| HORVU.MOREX.r3.4HG0392150.1 | -1.50616 | 3.360314 | -3.1432 | 0.002736 | 0.027326 | -1.62991 | #N/A | elongation zone | 12 h |
| HORVU.MOREX.r3.6HG0575410.1 | -2.05578 | 3.790358 | -3.15667 | 0.002632 | 0.026526 | -1.63023 | Glycosyltransferase | elongation zone | 12 h |
| HORVU.MOREX.r3.3HG0223160.1 | 0.612972 | 5.278307 | 3.376808 | 0.00138 | 0.017746 | -1.63584 | aberrant root formation protein | elongation zone | 12 h |
| HORVU.MOREX.r3.4HG0400530.1 | 1.264247 | 0.815449 | 3.15079 | 0.002677 | 0.02686 | -1.63718 | #N/A | elongation zone | 12 h |
| HORVU.MOREX.r3.5HG0508880.1 | 1.504618 | -0.73312 | 3.141222 | 0.002752 | 0.027347 | -1.639 | Leucine-rich repeat protein kinase family protein | elongation zone | 12 h |
| HORVU.MOREX.r3.2HG0208280.1 | -0.49204 | 7.445128 | -3.32272 | 0.00162 | 0.019769 | -1.63922 | Sulfotransferase | elongation zone | 12 h |
| HORVU.MOREX.r3.7HG0661470.1 | 0.71511 | 3.159382 | 3.171676 | 0.00252 | 0.025832 | -1.63929 | 2-oxoglutarate (2OG) and Fe(II)-dependent oxygenase superfamily protein | elongation zone | 12 h |
| HORVU.MOREX.r3.7HG0722690.1 | 0.509382 | 4.955916 | 3.314643 | 0.00166 | 0.01994 | -1.63956 | Amino acid transporter-like protein | elongation zone | 12 h |
| HORVU.MOREX.r3.4HG0342480.1 | 0.89056 | 4.010766 | 3.261533 | 0.001941 | 0.022073 | -1.64133 | Subtilisin-like protease | elongation zone | 12 h |
| HORVU.MOREX.r3.4HG0415170.1 | 1.544236 | -2.40064 | 3.138206 | 0.002775 | 0.027553 | -1.64375 | 3'-N-debenzoyl-2'-deoxytaxol N-benzoyltransferase | elongation zone | 12 h |
| HORVU.MOREX.r3.1HG0091820.1 | -0.60967 | 5.321523 | -3.21694 | 0.00221 | 0.023848 | -1.64465 | Glutamyl-tRNA (Gln) amidotransferase subunit A | elongation zone | 12 h |
| HORVU.MOREX.r3.4HG0362350.1 | -1.21447 | 2.471284 | -3.17966 | 0.002463 | 0.025392 | -1.64922 | Pentatricopeptide repeat-containing protein | elongation zone | 12 h |
| HORVU.MOREX.r3.4HG0343260.1 | 1.417524 | -2.7306 | 3.162116 | 0.002591 | 0.026328 | -1.65015 | NAD(P)H dehydrogenase subunit 48 | elongation zone | 12 h |
| HORVU.MOREX.r3.3HG0245320.1 | -0.50118 | 5.238204 | -3.21859 | 0.0022 | 0.02379 | -1.65177 | GDSL esterase/lipase | elongation zone | 12 h |
| HORVU.MOREX.r3.1HG0092860.1 | 1.211127 | -1.39568 | 3.159287 | 0.002612 | 0.026441 | -1.65479 | S-type anion channel | elongation zone | 12 h |
| HORVU.MOREX.r3.4HG0335160.1 | 0.77858 | 3.629312 | 3.188838 | 0.002398 | 0.024957 | -1.65496 | Hedgehog-interacting-like protein | elongation zone | 12 h |
| HORVU.MOREX.r3.2HG0180880.1 | -0.90096 | 1.957679 | -3.13266 | 0.00282 | 0.027847 | -1.65571 | GH3.3 | elongation zone | 12 h |
| HORVU.MOREX.r3.6HG0605090.1 | -0.3172 | 5.055557 | -3.25154 | 0.001998 | 0.022467 | -1.65637 | Alpha-1,3-glucosyltransferase | elongation zone | 12 h |
| HORVU.MOREX.r3.2HG0120490.1 | 1.171234 | 1.876399 | 3.309907 | 0.001683 | 0.020137 | -1.65717 | Peroxidase | elongation zone | 12 h |
| HORVU.MOREX.r3.7HG0749870.1 | 0.545306 | 4.57 | 3.298238 | 0.001742 | 0.020545 | -1.65731 | Glucuronoxylan 4-O-methyltransferase | elongation zone | 12 h |
| HORVU.MOREX.r3.6HG0552030.1 | -0.73035 | 3.851578 | -3.15622 | 0.002635 | 0.026547 | -1.65774 | Phosphatidate cytidylyltransferase | elongation zone | 12 h |
| HORVU.MOREX.r3.2HG0124990.1 | -0.63401 | 6.152886 | -3.25819 | 0.00196 | 0.022216 | -1.66018 | 3-oxoacyl-[acyl-carrier-protein] synthase | elongation zone | 12 h |
| HORVU.MOREX.r3.5HG0470690.1 | 1.236516 | -2.06961 | 3.133835 | 0.00281 | 0.027806 | -1.6604 | Heavy metal transport/detoxification protein | elongation zone | 12 h |
| HORVU.MOREX.r3.6HG0619760.1 | -0.47856 | 6.476084 | -3.26087 | 0.001944 | 0.02208 | -1.66262 | Serine--tRNA ligase | elongation zone | 12 h |
| HORVU.MOREX.r3.5HG0428270.2 | 0.690779 | 2.536061 | 3.231597 | 0.002118 | 0.023258 | -1.66496 | Bidirectional sugar transporter SWEET | elongation zone | 12 h |
| HORVU.MOREX.r3.2HG0116990.1 | 0.566387 | 4.613527 | 3.25748 | 0.001964 | 0.022241 | -1.66634 | Phosphatidylinositol 4-phosphate 5-kinase 4 | elongation zone | 12 h |
| HORVU.MOREX.r3.7HG0710980.1 | -1.14958 | 1.521079 | -3.19503 | 0.002356 | 0.024638 | -1.667 | Mitochondrial transcription termination factor-like | elongation zone | 12 h |
| HORVU.MOREX.r3.7HG0675310.1 | 1.239978 | 1.411092 | 3.254688 | 0.00198 | 0.022347 | -1.66749 | N-lysine methyltransferase | elongation zone | 12 h |
| HORVU.MOREX.r3.1HG0083980.1 | -0.993 | 4.411839 | -3.13283 | 0.002819 | 0.027846 | -1.66837 | Thymidine kinase | elongation zone | 12 h |
| HORVU.MOREX.r3.2HG0181510.2 | 0.451529 | 4.821103 | 3.272152 | 0.001881 | 0.021612 | -1.66847 | Beta-glucosidase | elongation zone | 12 h |
| HORVU.MOREX.r3.6HG0547010.1 | 0.880449 | -1.51289 | 3.141499 | 0.002749 | 0.027347 | -1.66871 | NBS-LRR-like resistance protein | elongation zone | 12 h |

**Table S1** Continued.

| ID | logFC | AveExpr | t | P.Value | adj.P.Val | B | gene description | root zone | time point |
|---|---|---|---|---|---|---|---|---|---|
| HORVU.MOREX.r3.3HG0307280.1 | 1.134433 | -2.743 | 3.153533 | 0.002656 | 0.026714 | -1.66925 | #N/A | elongation zone | 12 h |
| HORVU.MOREX.r3.5HG0493040.1 | 0.552633 | 5.221386 | 3.31858 | 0.00164 | 0.019852 | -1.66936 | Basic helix-loop-helix transcription factor | elongation zone | 12 h |
| HORVU.MOREX.r3.6HG0621190.1 | -0.60189 | 4.746594 | -3.24419 | 0.002042 | 0.022694 | -1.66938 | Chaperone protein dnaJ | elongation zone | 12 h |
| HORVU.MOREX.r3.5HG0502760.1 | -0.74648 | 7.161462 | -3.27956 | 0.001841 | 0.02129 | -1.67331 | Histone H4 | elongation zone | 12 h |
| HORVU.MOREX.r3.3HG0313090.1 | 0.737376 | 2.944421 | 3.167903 | 0.002548 | 0.026026 | -1.67422 | Thiamin pyrophosphokinase 1 | elongation zone | 12 h |
| HORVU.MOREX.r3.1HG0080690.1 | -2.17717 | 4.834533 | -3.12402 | 0.002891 | 0.028313 | -1.67654 | Endoglucanase | elongation zone | 12 h |
| HORVU.MOREX.r3.4HG0353200.1 | 0.524688 | 2.886286 | 3.23998 | 0.002067 | 0.022883 | -1.67738 | heat-inducible transcription repressor | elongation zone | 12 h |
| HORVU.MOREX.r3.1HG0004300.1 | -1.92616 | 1.490539 | -3.2847 | 0.001813 | 0.021077 | -1.68169 | Disease resistance protein (TIR-NBS-LRR class) family | elongation zone | 12 h |
| HORVU.MOREX.r3.3HG0295440.1 | -0.81047 | 5.026864 | -3.18394 | 0.002433 | 0.025148 | -1.68372 | Cytochrome P450 family protein | elongation zone | 12 h |
| HORVU.MOREX.r3.3HG0280160.1 | 0.422379 | 4.904831 | 3.270734 | 0.001889 | 0.021665 | -1.68414 | Evolutionarily conserved C-terminal region 2 | elongation zone | 12 h |
| HORVU.MOREX.r3.4HG0384780.1 | 3.253521 | -0.01492 | 3.15882 | 0.002616 | 0.026447 | -1.68634 | Trichome birefringence-like protein | elongation zone | 12 h |
| HORVU.MOREX.r3.3HG0315410.1 | 0.359796 | 5.906295 | 3.283693 | 0.001818 | 0.021103 | -1.6889 | Vesicle-associated protein 1-2 | elongation zone | 12 h |
| HORVU.MOREX.r3.1HG0093050.1 | 0.81023 | 4.062497 | 3.206431 | 0.002279 | 0.024194 | -1.6902 | Indole-3-acetic acid-amido synthetase GH3.3 | elongation zone | 12 h |
| HORVU.MOREX.r3.4HG0385080.1 | -0.57534 | 3.22216 | -3.1476 | 0.002701 | 0.02705 | -1.69117 | RING/U-box superfamily protein | elongation zone | 12 h |
| HORVU.MOREX.r3.4HG0378960.1 | -0.99437 | 5.668628 | -3.17066 | 0.002528 | 0.025884 | -1.69188 | Peroxidase | elongation zone | 12 h |
| HORVU.MOREX.r3.7HG0728630.1 | -1.00506 | 1.895272 | -3.12981 | 0.002843 | 0.028035 | -1.69421 | Receptor-like protein kinase | elongation zone | 12 h |
| HORVU.MOREX.r3.6HG0616890.1 | -0.94787 | 0.931971 | -3.15765 | 0.002624 | 0.026466 | -1.69565 | UPF0503 protein, chloroplastic | elongation zone | 12 h |
| HORVU.MOREX.r3.2HG0171850.1 | -0.48872 | 5.054708 | -3.17955 | 0.002464 | 0.025392 | -1.69609 | mediator of RNA polymerase II transcription subunit-like protein | elongation zone | 12 h |
| HORVU.MOREX.r3.7HG0655260.1 | -0.96356 | 1.989817 | -3.11677 | 0.002951 | 0.028675 | -1.7011 | Cortactin-binding protein 2 | elongation zone | 12 h |
| HORVU.MOREX.r3.1HG0003280.1 | -2.2261 | -1.64899 | -3.34482 | 0.001518 | 0.018897 | -1.7013 | #N/A | elongation zone | 12 h |
| HORVU.MOREX.r3.5HG0534420.1 | 0.841071 | 1.580423 | 3.242733 | 0.00205 | 0.022754 | -1.70159 | #N/A | elongation zone | 12 h |
| HORVU.MOREX.r3.7HG0702070.1 | 1.293035 | -1.23872 | 3.126692 | 0.002869 | 0.02816 | -1.70193 | Blue copper protein | elongation zone | 12 h |
| HORVU.MOREX.r3.7HG0636510.1 | -0.54177 | 3.279937 | -3.14678 | 0.002708 | 0.027079 | -1.70196 | PPPDE putative thiol peptidase family protein | elongation zone | 12 h |
| HORVU.MOREX.r3.1HG0068890.1 | -0.55352 | 3.912321 | -3.13405 | 0.002809 | 0.027803 | -1.70219 | Hydroxyproline-rich glycoprotein family protein, putative | elongation zone | 12 h |
| HORVU.MOREX.r3.7HG0663940.1 | 0.787758 | 0.764913 | 3.218496 | 0.0022 | 0.02379 | -1.70261 | Alpha/beta-Hydrolases superfamily protein, putative | elongation zone | 12 h |
| HORVU.MOREX.r3.6HG0609750.1 | -0.51401 | 4.811837 | -3.21632 | 0.002214 | 0.023861 | -1.70301 | PRELI/MSF1-like protein | elongation zone | 12 h |
| HORVU.MOREX.r3.6HG0543750.1 | -0.80048 | 7.794738 | -3.29396 | 0.001764 | 0.020639 | -1.70305 | Histone H2A | elongation zone | 12 h |
| HORVU.MOREX.r3.7HG0650530.2 | 1.666946 | -2.50586 | 3.123733 | 0.002893 | 0.028323 | -1.70352 | 2-oxoglutarate (2OG) and Fe(II)-dependent oxygenase superfamily protein | elongation zone | 12 h |
| HORVU.MOREX.r3.1HG0079280.1 | -0.93957 | 8.411999 | -3.29609 | 0.001753 | 0.020579 | -1.70461 | Histone H2B | elongation zone | 12 h |
| HORVU.MOREX.r3.1HG0032910.1 | -0.57602 | 5.350159 | -3.20829 | 0.002267 | 0.02416 | -1.70545 | Lysine-specific demethylase | elongation zone | 12 h |
| HORVU.MOREX.r3.7HG0728190.1 | -0.42379 | 4.089813 | -3.18467 | 0.002428 | 0.025148 | -1.70662 | Purple acid phosphatase | elongation zone | 12 h |
| HORVU.MOREX.r3.2HG0188200.1 | 0.783338 | 0.241735 | 3.161528 | 0.002595 | 0.026351 | -1.70919 | Calcium-dependent protein kinase | elongation zone | 12 h |
| HORVU.MOREX.r3.2HG0181390.1 | -1.49649 | 5.181001 | -3.13312 | 0.002816 | 0.027837 | -1.70957 | Protein BREAST CANCER SUSCEPTIBILITY 1-like protein | elongation zone | 12 h |
| HORVU.MOREX.r3.1HG0053770.1 | -1.30372 | 0.507378 | -3.20531 | 0.002286 | 0.0242 | -1.71105 | Protein kinase family protein | elongation zone | 12 h |
| HORVU.MOREX.r3.1HG0007930.1 | -0.7667 | 2.116239 | -3.11435 | 0.002972 | 0.028819 | -1.71223 | calpain-type cysteine protease family | elongation zone | 12 h |
| HORVU.MOREX.r3.3HG0237910.1 | -0.99223 | 2.053847 | -3.11722 | 0.002947 | 0.028675 | -1.71339 | P53/DNA damage-regulated protein | elongation zone | 12 h |
| HORVU.MOREX.r3.5HG0462200.1 | 1.559192 | -0.85589 | 3.128152 | 0.002857 | 0.028074 | -1.71353 | Metacaspase | elongation zone | 12 h |
| HORVU.MOREX.r3.7HG0719170.1 | 0.699368 | 5.542866 | 3.303456 | 0.001715 | 0.020382 | -1.71368 | O-fucosyltransferase-like protein | elongation zone | 12 h |
| HORVU.MOREX.r3.1HG0084850.1 | -1.19552 | 4.250962 | -3.11163 | 0.002995 | 0.029003 | -1.71493 | Histone chaperone | elongation zone | 12 h |
| HORVU.MOREX.r3.4HG0385100.1 | -0.33817 | 5.669475 | -3.25538 | 0.001976 | 0.022314 | -1.71626 | Protein YIPF | elongation zone | 12 h |
| HORVU.MOREX.r3.7HG0673330.1 | 1.312537 | -2.01567 | 3.105167 | 0.003051 | 0.029335 | -1.72069 | Embryogenesis transmembrane protein-like | elongation zone | 12 h |

**Table S1** Continued.

| ID | logFC | AveExpr | t | P.Value | adj.P.Val | B | gene description | root zone | time point |
|---|---|---|---|---|---|---|---|---|---|
| HORVU.MOREX.r3.4HG0411270.1 | -0.78687 | 6.581905 | -3.2251 | 0.002159 | 0.023548 | -1.72077 | N-acetyltransferase | elongation zone | 12 h |
| HORVU.MOREX.r3.4HG0342950.1 | -0.95721 | 6.384536 | -3.1878 | 0.002406 | 0.025019 | -1.72288 | 50S ribosomal protein L25 | elongation zone | 12 h |
| HORVU.MOREX.r3.7HG0725440.1 | 0.888939 | 1.519211 | 3.108432 | 0.003022 | 0.029118 | -1.7235 | Pre-mRNA-splicing factor syf2 | elongation zone | 12 h |
| HORVU.MOREX.r3.5HG0472140.1 | 0.692391 | 2.018509 | 3.123358 | 0.002896 | 0.028327 | -1.72505 | Chlororespiratory reduction 3 | elongation zone | 12 h |
| HORVU.MOREX.r3.6HG0595810.1 | -0.92617 | 1.111513 | -3.1245 | 0.002887 | 0.028299 | -1.7256 | Pre-mRNA-processing protein 40A | elongation zone | 12 h |
| HORVU.MOREX.r3.2HG0185570.1 | 0.736473 | 1.316705 | 3.182074 | 0.002446 | 0.025246 | -1.72583 | transcription repressor | elongation zone | 12 h |
| HORVU.MOREX.r3.6HG0614460.1 | -0.71042 | 3.048312 | -3.10254 | 0.003074 | 0.029497 | -1.72829 | DNA-directed RNA polymerase subunit beta | elongation zone | 12 h |
| HORVU.MOREX.r3.4HG0336310.1 | -0.67415 | 6.50287 | -3.22411 | 0.002165 | 0.023553 | -1.72871 | Zinc finger family protein | elongation zone | 12 h |
| HORVU.MOREX.r3.1HG0033450.1 | -1.60612 | 0.194648 | -3.21289 | 0.002237 | 0.023992 | -1.72938 | MBOAT (membrane bound O-acyl transferase) family protein | elongation zone | 12 h |
| HORVU.MOREX.r3.7HG0687190.1 | -0.51904 | 3.62232 | -3.12658 | 0.00287 | 0.02816 | -1.73079 | Glucosamine-6-phosphate deaminase | elongation zone | 12 h |
| HORVU.MOREX.r3.4HG0387120.1 | -1.01903 | 6.929288 | -3.22547 | 0.002156 | 0.023536 | -1.73232 | Histone H2A | elongation zone | 12 h |
| HORVU.MOREX.r3.5HG0430050.1 | 0.55481 | 4.495639 | 3.198111 | 0.002335 | 0.024484 | -1.73295 | Stomatal closure-related actin-binding protein 1 | elongation zone | 12 h |
| HORVU.MOREX.r3.7HG0635700.1 | -0.33382 | 5.807806 | -3.22974 | 0.00213 | 0.023356 | -1.73458 | Transposon protein, putative, CACTA, En/Spm sub-class | elongation zone | 12 h |
| HORVU.MOREX.r3.1HG0079250.1 | -0.82686 | 7.964833 | -3.26979 | 0.001894 | 0.021713 | -1.73567 | Histone H2B | elongation zone | 12 h |
| HORVU.MOREX.r3.1HG0017310.1 | -1.10015 | 6.737539 | -3.18395 | 0.002433 | 0.025148 | -1.74004 | Auxin influx transporter | elongation zone | 12 h |
| HORVU.MOREX.r3.7HG0726350.1 | -0.58695 | 4.52716 | -3.16431 | 0.002575 | 0.026239 | -1.74107 | Serine/threonine-protein kinase | elongation zone | 12 h |
| HORVU.MOREX.r3.1HG0049520.1 | -0.75271 | 3.446094 | -3.11102 | 0.003 | 0.029025 | -1.74357 | Cytochrome P450 family protein | elongation zone | 12 h |
| HORVU.MOREX.r3.2HG0117880.1 | 2.599524 | -0.66123 | 3.184875 | 0.002426 | 0.025148 | -1.74359 | GDSL esterase/lipase | elongation zone | 12 h |
| HORVU.MOREX.r3.7HG0671880.1 | -0.74751 | 3.670438 | -3.10093 | 0.003088 | 0.029609 | -1.74522 | RING/U-box protein | elongation zone | 12 h |
| HORVU.MOREX.r3.5HG0504160.1 | -1.07219 | 6.884631 | -3.21748 | 0.002207 | 0.023823 | -1.74538 | #N/A | elongation zone | 12 h |
| HORVU.MOREX.r3.7HG0732330.1 | -1.52362 | 0.94 | -3.18991 | 0.002391 | 0.024902 | -1.74565 | Phosphosulfolactate synthase-related protein | elongation zone | 12 h |
| HORVU.MOREX.r3.7HG0663820.1 | -0.83397 | 2.698491 | -3.10072 | 0.00309 | 0.029613 | -1.7468 | Protein TIC 22, chloroplastic | elongation zone | 12 h |
| HORVU.MOREX.r3.4HG0407980.1 | -0.7173 | 2.024558 | -3.09458 | 0.003144 | 0.02993 | -1.74686 | Myc-type, basic helix-loop-helix (BHLH) domain-containing protein | elongation zone | 12 h |
| HORVU.MOREX.r3.5HG0423820.1 | 0.878324 | 2.29448 | 3.093612 | 0.003153 | 0.029967 | -1.74717 | #N/A | elongation zone | 12 h |
| HORVU.MOREX.r3.4HG0385240.1 | 0.621024 | 6.413717 | 3.30879 | 0.001689 | 0.020145 | -1.74819 | Alpha/beta-Hydrolases superfamily protein | elongation zone | 12 h |
| HORVU.MOREX.r3.7HG0706310.1 | 0.481027 | 5.228141 | 3.208683 | 0.002264 | 0.02416 | -1.74936 | Alcohol dehydrogenase | elongation zone | 12 h |
| HORVU.MOREX.r3.6HG0546180.1 | -0.70925 | 4.078621 | -3.1105 | 0.003005 | 0.029025 | -1.75013 | Remorin family protein | elongation zone | 12 h |
| HORVU.MOREX.r3.2HG0181550.1 | -1.01065 | 3.831202 | -3.11047 | 0.003005 | 0.029025 | -1.75035 | Imidazole glycerol phosphate synthase subunit HisF | elongation zone | 12 h |
| HORVU.MOREX.r3.3HG0275530.1 | -0.75369 | 3.98589 | -3.10973 | 0.003011 | 0.02903 | -1.75262 | Sodium Bile acid symporter family | elongation zone | 12 h |
| HORVU.MOREX.r3.7HG0744790.1 | -0.58148 | 5.538256 | -3.16416 | 0.002576 | 0.026239 | -1.7527 | XH/XS domain-containing family protein | elongation zone | 12 h |
| HORVU.MOREX.r3.5HG0497990.1 | 1.11839 | 3.554356 | 3.134921 | 0.002802 | 0.027765 | -1.7527 | Heat shock transcription factor | elongation zone | 12 h |
| HORVU.MOREX.r3.6HG0602230.1 | -0.50919 | 5.050411 | -3.20099 | 0.002315 | 0.02439 | -1.7542 | Protein kinase | elongation zone | 12 h |
| HORVU.MOREX.r3.5HG0516310.1 | -0.97831 | 5.306262 | -3.20802 | 0.002269 | 0.02416 | -1.75433 | Patatin | elongation zone | 12 h |
| HORVU.MOREX.r3.6HG0573340.1 | -0.38354 | 4.889095 | -3.19705 | 0.002342 | 0.024507 | -1.75433 | Alpha/beta hydrolase family protein | elongation zone | 12 h |
| HORVU.MOREX.r3.5HG0459660.1 | -0.78159 | 3.348001 | -3.22388 | 0.002166 | 0.023557 | -1.75488 | Stress responsive A/B barrel domain family protein | elongation zone | 12 h |
| HORVU.MOREX.r3.3HG0234240.1 | 0.97395 | 0.193181 | 3.143618 | 0.002733 | 0.027313 | -1.75535 | Alpha/beta-hydrolase superfamily protein | elongation zone | 12 h |
| HORVU.MOREX.r3.6HG0603820.1 | -0.24723 | 8.222526 | -3.35484 | 0.001473 | 0.018499 | -1.75757 | 3,5-epimerase/4-reductase | elongation zone | 12 h |
| HORVU.MOREX.r3.2HG0215310.1 | -1.45184 | 7.0907 | -3.19438 | 0.00236 | 0.024647 | -1.75791 | Peroxidase | elongation zone | 12 h |
| HORVU.MOREX.r3.6HG0599900.1 | 0.629818 | 3.934569 | 3.207628 | 0.002271 | 0.02416 | -1.75813 | Vesicle-associated protein 1-1 | elongation zone | 12 h |
| HORVU.MOREX.r3.5HG0487660.1 | -1.05829 | 6.052415 | -3.19225 | 0.002375 | 0.024774 | -1.7591 | Histone H2A | elongation zone | 12 h |
| HORVU.MOREX.r3.4HG0378460.1 | 0.778869 | 2.042348 | 3.128521 | 0.002854 | 0.028071 | -1.75922 | 1-(5-phosphoribosyl)-5-[(5-phosphoribosylamino)methylideneamino] imidazole-4- | elongation zone | 12 h |

**Table S1** Continued.

| ID | logFC | AveExpr | t | P.Value | adj.P.Val | B | gene description | root zone | time point |
|---|---|---|---|---|---|---|---|---|---|
| HORVU.MOREX.r3.7HG0727070.1 | -0.64752 | 4.936044 | -3.15869 | 0.002617 | 0.026447 | -1.76161 | Histone H2A | elongation zone | 12 h |
| HORVU.MOREX.r3.5HG0420630.1 | -0.47273 | 3.720927 | -3.16376 | 0.002579 | 0.026247 | -1.76166 | Ankyrin repeat family protein | elongation zone | 12 h |
| HORVU.MOREX.r3.7HG0635160.1 | -0.39648 | 6.815127 | -3.23568 | 0.002093 | 0.023091 | -1.76229 | Monocopper oxidase-like protein SKU5 | elongation zone | 12 h |
| HORVU.MOREX.r3.5HG0480540.1 | -0.36845 | 7.034207 | -3.27749 | 0.001852 | 0.02136 | -1.76235 | Transmembrane protein 87A | elongation zone | 12 h |
| HORVU.MOREX.r3.6HG0604310.1 | -1.12904 | 2.696154 | -3.13368 | 0.002812 | 0.027806 | -1.7628 | RING finger protein | elongation zone | 12 h |
| HORVU.MOREX.r3.6HG0596770.1 | 0.737545 | 4.952847 | 3.367488 | 0.001419 | 0.018081 | -1.76339 | Aquaporin | elongation zone | 12 h |
| HORVU.MOREX.r3.7HG0656030.1 | 0.61465 | 3.605835 | 3.21505 | 0.002223 | 0.023917 | -1.7687 | Receptor-like kinase | elongation zone | 12 h |
| HORVU.MOREX.r3.4HG0393630.1 | -0.29714 | 9.32692 | -3.36875 | 0.001413 | 0.018046 | -1.77077 | Transmembrane 9 superfamily member | elongation zone | 12 h |
| HORVU.MOREX.r3.1HG0037020.1 | -0.43364 | 3.762682 | -3.12505 | 0.002882 | 0.028271 | -1.77154 | F-box/kelch-repeat protein | elongation zone | 12 h |
| HORVU.MOREX.r3.1HG0077790.1 | 1.008805 | 3.495979 | 3.200036 | 0.002322 | 0.024432 | -1.77312 | Calcium-transporting ATPase | elongation zone | 12 h |
| HORVU.MOREX.r3.5HG0471320.1 | -1.73437 | 0.737353 | -3.22015 | 0.00219 | 0.023739 | -1.77451 | Amino acid transporter, putative | elongation zone | 12 h |
| HORVU.MOREX.r3.2HG0102600.1 | -0.36884 | 5.97563 | -3.22841 | 0.002138 | 0.023409 | -1.77519 | Peptidylprolyl isomerase | elongation zone | 12 h |
| HORVU.MOREX.r3.6HG0613430.1 | 1.684689 | -1.24134 | 3.086207 | 0.00322 | 0.030422 | -1.77808 | RNA-dependent RNA polymerase | elongation zone | 12 h |
| HORVU.MOREX.r3.5HG0477360.1 | -0.68158 | 2.537497 | -3.08278 | 0.003252 | 0.030649 | -1.77819 | Pentatricopeptide repeat-containing protein | elongation zone | 12 h |
| HORVU.MOREX.r3.2HG0190290.1 | 0.626476 | 5.33841 | 3.227377 | 0.002144 | 0.023455 | -1.77826 | SHR5-receptor-like kinase | elongation zone | 12 h |
| HORVU.MOREX.r3.2HG0157340.1 | 0.496722 | 7.39472 | 3.400577 | 0.001285 | 0.01705 | -1.77924 | Blue copper protein | elongation zone | 12 h |
| HORVU.MOREX.r3.6HG0549510.1 | -0.67131 | 8.771369 | -3.32104 | 0.001628 | 0.019852 | -1.7821 | Peptidyl-prolyl cis-trans isomerase | elongation zone | 12 h |
| HORVU.MOREX.r3.2HG0121480.1 | 0.765263 | 1.064359 | 3.261259 | 0.001942 | 0.022079 | -1.7826 | MYB transcription factor | elongation zone | 12 h |
| HORVU.MOREX.r3.1HG0014410.1 | -0.63575 | 5.218845 | -3.16087 | 0.0026 | 0.026389 | -1.78343 | Receptor-like kinase | elongation zone | 12 h |
| HORVU.MOREX.r3.1HG0072320.1 | -0.82668 | 1.113845 | -3.07916 | 0.003285 | 0.030825 | -1.7839 | S-adenosyl-L-methionine-dependent methyltransferases superfamily protein | elongation zone | 12 h |
| HORVU.MOREX.r3.3HG0245120.1 | 2.751055 | -2.69428 | 3.205406 | 0.002286 | 0.0242 | -1.78464 | Cysteine protease, putative | elongation zone | 12 h |
| HORVU.MOREX.r3.3HG0314240.1 | -0.4127 | 5.164101 | -3.20122 | 0.002314 | 0.024386 | -1.78543 | Potassium transporter | elongation zone | 12 h |
| HORVU.MOREX.r3.4HG0401450.1 | -0.44327 | 7.107053 | -3.26795 | 0.001904 | 0.021791 | -1.78569 | Anthocyanin 5-aromatic acyltransferase | elongation zone | 12 h |
| HORVU.MOREX.r3.2HG0155290.1 | 1.482062 | -0.36543 | 3.134857 | 0.002802 | 0.027765 | -1.7886 | Retrotransposon protein, putative, unclassified | elongation zone | 12 h |
| HORVU.MOREX.r3.6HG0611300.1 | -0.98231 | 2.278681 | -3.09201 | 0.003167 | 0.030076 | -1.78871 | AMP-dependent synthetase and ligase family protein | elongation zone | 12 h |
| HORVU.MOREX.r3.3HG0232780.1 | -0.56892 | 2.836303 | -3.08171 | 0.003262 | 0.030701 | -1.79009 | R3H domain containing protein | elongation zone | 12 h |
| HORVU.MOREX.r3.5HG0525020.1 | -0.80314 | 2.149516 | -3.07988 | 0.003279 | 0.030803 | -1.79105 | Pentatricopeptide repeat-containing protein | elongation zone | 12 h |
| HORVU.MOREX.r3.5HG0499490.1 | -0.28473 | 6.586701 | -3.23663 | 0.002087 | 0.023064 | -1.792 | Transcription initiation factor IIB | elongation zone | 12 h |
| HORVU.MOREX.r3.1HG0000670.1 | -0.84436 | 2.483716 | -3.07379 | 0.003336 | 0.031156 | -1.79438 | translocase subunit seca | elongation zone | 12 h |
| HORVU.MOREX.r3.1HG0018390.1 | -0.98561 | 5.772475 | -3.16959 | 0.002536 | 0.025938 | -1.79629 | #N/A | elongation zone | 12 h |
| HORVU.MOREX.r3.7HG0662220.1 | -1.00768 | 3.864095 | -3.07778 | 0.003298 | 0.030931 | -1.79637 | UDP-glycosyltransferase | elongation zone | 12 h |
| HORVU.MOREX.r3.3HG0240390.2 | -1.25813 | 2.367221 | -3.07555 | 0.003319 | 0.031043 | -1.79889 | Purine nucleoside phosphorylase DeoD-type | elongation zone | 12 h |
| HORVU.MOREX.r3.3HG0220250.1 | -0.48567 | 3.555795 | -3.12045 | 0.00292 | 0.028505 | -1.79923 | Receptor-like kinase | elongation zone | 12 h |
| HORVU.MOREX.r3.5HG0467950.1 | -0.47774 | 6.475334 | -3.22459 | 0.002162 | 0.023553 | -1.80033 | Divalent metal cation transporter MntH 2 | elongation zone | 12 h |
| HORVU.MOREX.r3.7HG0680170.1 | -0.4793 | 4.825321 | -3.15763 | 0.002625 | 0.026466 | -1.80043 | mediator of RNA polymerase II transcription subunit | elongation zone | 12 h |
| HORVU.MOREX.r3.5HG0512250.1 | 1.309951 | -0.16991 | 3.178281 | 0.002473 | 0.025473 | -1.80131 | Receptor-like kinase, putative | elongation zone | 12 h |
| HORVU.MOREX.r3.4HG0382500.1 | -0.39502 | 5.221396 | -3.1819 | 0.002447 | 0.025246 | -1.80152 | Seed maturation protein PM36, putative, expressed | elongation zone | 12 h |
| HORVU.MOREX.r3.4HG0353330.1 | -0.93577 | 5.045545 | -3.11907 | 0.002932 | 0.028562 | -1.80231 | Protein kinase | elongation zone | 12 h |
| HORVU.MOREX.r3.1HG0003130.1 | -0.49332 | 5.529036 | -3.21196 | 0.002243 | 0.02402 | -1.80483 | Regulator of chromosome condensation (RCC1) family with FYVE zinc finger doma | elongation zone | 12 h |
| HORVU.MOREX.r3.4HG0405860.1 | 2.050549 | 1.28141 | 3.081837 | 0.00326 | 0.030701 | -1.80521 | Protein WEAK CHLOROPLAST MOVEMENT UNDER BLUE LIGHT 1 | elongation zone | 12 h |
| HORVU.MOREX.r3.6HG0629810.2 | 0.921281 | 1.891863 | 3.069158 | 0.00338 | 0.031411 | -1.8054 | Auxin-responsive protein | elongation zone | 12 h |

**Table S1** Continued.

| ID | logFC | AveExpr | t | P.Value | adj.P.Val | B | gene description | root zone | time point |
|---|---|---|---|---|---|---|---|---|---|
| HORVU.MOREX.r3.6HG0549830.1 | -0.74227 | 2.038518 | -3.06998 | 0.003372 | 0.031367 | -1.80548 | #N/A | elongation zone | 12 h |
| HORVU.MOREX.r3.5HG0511820.1 | 2.279487 | -3.38085 | 3.205578 | 0.002285 | 0.0242 | -1.80714 | Cyclic nucleotide-gated channel | elongation zone | 12 h |
| HORVU.MOREX.r3.3HG0309760.1 | 0.81508 | 3.402606 | 3.0931 | 0.003158 | 0.029997 | -1.80818 | ROP-interactive CRIB motif protein | elongation zone | 12 h |
| HORVU.MOREX.r3.4HG0354480.1 | -0.30043 | 6.236734 | -3.22759 | 0.002143 | 0.023453 | -1.81059 | Aquaporin-like protein | elongation zone | 12 h |
| HORVU.MOREX.r3.2HG0200710.1 | 1.833556 | 1.897406 | 3.06686 | 0.003402 | 0.031531 | -1.81076 | Cytochrome P450 family protein | elongation zone | 12 h |
| HORVU.MOREX.r3.2HG0120030.1 | -0.72384 | 4.633702 | -3.14283 | 0.002739 | 0.027326 | -1.81082 | Xylulose kinase | elongation zone | 12 h |
| HORVU.MOREX.r3.2HG0155460.1 | -0.65291 | 2.860269 | -3.08629 | 0.003219 | 0.030422 | -1.81236 | #N/A | elongation zone | 12 h |
| HORVU.MOREX.r3.7HG0726210.1 | -0.72097 | 2.875567 | -3.06578 | 0.003412 | 0.031613 | -1.81265 | Two-component response regulator | elongation zone | 12 h |
| HORVU.MOREX.r3.5HG0511930.1 | -0.8946 | 2.612278 | -3.07011 | 0.003371 | 0.031367 | -1.815 | Ring finger protein, putative | elongation zone | 12 h |
| HORVU.MOREX.r3.2HG0204340.1 | -2.2925 | -0.82148 | -3.58767 | 0.000728 | 0.012414 | -1.81523 | ERD (early-responsive to dehydration stress) family protein | elongation zone | 12 h |
| HORVU.MOREX.r3.5HG0487040.1 | -0.49437 | 7.455763 | -3.28418 | 0.001816 | 0.021097 | -1.81563 | Alpha/beta-Hydrolases superfamily protein, putative | elongation zone | 12 h |
| HORVU.MOREX.r3.3HG0220360.1 | -0.82439 | 4.616465 | -3.09826 | 0.003111 | 0.029766 | -1.81724 | Disease resistance protein | elongation zone | 12 h |
| HORVU.MOREX.r3.2HG0180210.1 | -1.20188 | 2.207614 | -3.07169 | 0.003356 | 0.0313 | -1.8176 | methyl-coenzyme M reductase II subunit gamma, putative (DUF3741) | elongation zone | 12 h |
| HORVU.MOREX.r3.6HG0624390.1 | 1.053902 | 1.652699 | 3.067491 | 0.003396 | 0.031493 | -1.8194 | Two-component response regulator | elongation zone | 12 h |
| HORVU.MOREX.r3.6HG0615710.1 | -0.53607 | 6.427382 | -3.18511 | 0.002424 | 0.025148 | -1.82032 | Nucleobase ascorbate transporter | elongation zone | 12 h |
| HORVU.MOREX.r3.2HG0159580.1 | 0.654808 | 4.426208 | 3.207669 | 0.002271 | 0.02416 | -1.82202 | Carboxypeptidase | elongation zone | 12 h |
| HORVU.MOREX.r3.5HG0480600.1 | 1.846016 | -0.20786 | 3.221218 | 0.002183 | 0.023703 | -1.82344 | Jasmonate zim-domain protein | elongation zone | 12 h |
| HORVU.MOREX.r3.6HG0555250.1 | 2.168061 | 0.376943 | 3.391173 | 0.001322 | 0.017264 | -1.82347 | Esterase/lipase/thioesterase-like protein | elongation zone | 12 h |
| HORVU.MOREX.r3.4HG0379530.1 | -0.417 | 3.959492 | -3.13122 | 0.002832 | 0.027935 | -1.82548 | O-fucosyltransferase family protein | elongation zone | 12 h |
| HORVU.MOREX.r3.6HG0613360.1 | 1.369878 | -0.60847 | 3.087913 | 0.003205 | 0.030372 | -1.8275 | Nucleotidylyl transferase superfamily protein | elongation zone | 12 h |
| HORVU.MOREX.r3.2HG0175350.1 | 2.260985 | 1.060585 | 3.266471 | 0.001913 | 0.021804 | -1.8288 | C2 and GRAM domain-containing protein | elongation zone | 12 h |
| HORVU.MOREX.r3.3HG0303560.2 | 1.618222 | -1.42721 | 3.14127 | 0.002751 | 0.027347 | -1.82894 | Interactor of constitutive active ROPs 2, chloroplastic | elongation zone | 12 h |
| HORVU.MOREX.r3.7HG0749660.1 | -0.65669 | 5.553628 | -3.14278 | 0.002739 | 0.027326 | -1.83583 | DNA-binding protein RHL1 | elongation zone | 12 h |
| HORVU.MOREX.r3.1HG0070480.1 | 0.326891 | 6.291081 | 3.244411 | 0.00204 | 0.022692 | -1.83611 | 60 kDa chaperonin | elongation zone | 12 h |
| HORVU.MOREX.r3.2HG0122720.1 | -0.98085 | 3.012374 | -3.05461 | 0.003522 | 0.032179 | -1.83805 | Serine hydroxymethyltransferase | elongation zone | 12 h |
| HORVU.MOREX.r3.5HG0480060.1 | -1.18427 | 0.892295 | -3.06491 | 0.003421 | 0.031662 | -1.83915 | Protein GAMETE EXPRESSED 1 | elongation zone | 12 h |
| HORVU.MOREX.r3.4HG0399470.1 | 2.529684 | -1.76622 | 3.319681 | 0.001635 | 0.019852 | -1.84382 | Clathrin heavy chain | elongation zone | 12 h |
| HORVU.MOREX.r3.7HG0682460.1 | -0.34739 | 5.61607 | -3.20489 | 0.002289 | 0.024215 | -1.84405 | ADP-ribosylation factor, putative | elongation zone | 12 h |
| HORVU.MOREX.r3.4HG0351070.1 | 0.746998 | 2.367322 | 3.146889 | 0.002707 | 0.027079 | -1.84562 | STRUBBELIG-receptor family 5 | elongation zone | 12 h |
| HORVU.MOREX.r3.4HG0338180.1 | -0.40324 | 5.846029 | -3.21603 | 0.002216 | 0.023861 | -1.84751 | Transmembrane and coiled-coil domains protein 1 | elongation zone | 12 h |
| HORVU.MOREX.r3.4HG0333550.2 | 0.70912 | 1.181427 | 3.072298 | 0.00335 | 0.03126 | -1.84811 | senescence-associated family protein, putative (DUF581) | elongation zone | 12 h |
| HORVU.MOREX.r3.7HG0712000.1 | 1.730225 | -3.03639 | 3.14822 | 0.002697 | 0.02702 | -1.84844 | Transposon protein | elongation zone | 12 h |
| HORVU.MOREX.r3.2HG0182800.1 | -1.17278 | 3.942974 | -3.06925 | 0.003379 | 0.031411 | -1.85178 | Histone H3 | elongation zone | 12 h |
| HORVU.MOREX.r3.5HG0476690.1 | -0.30753 | 5.143097 | -3.17372 | 0.002506 | 0.02572 | -1.85236 | #N/A | elongation zone | 12 h |
| HORVU.MOREX.r3.6HG0633960.1 | 0.881693 | 2.11648 | 3.256514 | 0.001969 | 0.022287 | -1.85346 | Monosaccharide-sensing protein 2 | elongation zone | 12 h |
| HORVU.MOREX.r3.1HG0078370.1 | -1.78543 | 4.280641 | -3.05097 | 0.003559 | 0.032357 | -1.85361 | breast cancer associated RING 1 | elongation zone | 12 h |
| HORVU.MOREX.r3.7HG0645350.1 | -0.30248 | 9.301069 | -3.33171 | 0.001578 | 0.019378 | -1.85419 | 1-deoxy-D-xylulose-5-phosphate synthase, Transketolase | elongation zone | 12 h |
| HORVU.MOREX.r3.2HG0194620.1 | 1.879527 | -2.14798 | 3.127957 | 0.002858 | 0.028076 | -1.85449 | F-box protein | elongation zone | 12 h |
| HORVU.MOREX.r3.1HG0081110.1 | 0.650019 | 4.269156 | 3.212564 | 0.002239 | 0.024002 | -1.85467 | Type I inositol-1,4,5-trisphosphate 5-phosphatase 1 | elongation zone | 12 h |
| HORVU.MOREX.r3.3HG0309930.1 | 1.053411 | -0.53911 | 3.067551 | 0.003395 | 0.031493 | -1.8558 | Sucrose-phosphate synthase | elongation zone | 12 h |
| HORVU.MOREX.r3.7HG0719410.1 | -0.89347 | 6.134816 | -3.11897 | 0.002933 | 0.028562 | -1.85673 | Trichome birefringence-like protein | elongation zone | 12 h |

**Table S1** Continued.

| ID | logFC | AveExpr | t | P.Value | adj.P.Val | B | gene description | root zone | time point |
|---|---|---|---|---|---|---|---|---|---|
| HORVU.MOREX.r3.5HG0456510.1 | -0.44608 | 7.286365 | -3.24905 | 0.002013 | 0.022519 | -1.85958 | Dihydrolipoamide acetyltransferase component of pyruvate dehydrogenase comp | elongation zone | 12 h |
| HORVU.MOREX.r3.3HG0291570.1 | 0.404537 | 6.662916 | 3.24719 | 0.002024 | 0.022545 | -1.86159 | Nuclear matrix constituent-like protein 1 | elongation zone | 12 h |
| HORVU.MOREX.r3.2HG0130200.1 | -0.42795 | 4.823465 | -3.14154 | 0.002749 | 0.027347 | -1.8622 | ZCW7 | elongation zone | 12 h |
| HORVU.MOREX.r3.6HG0547550.1 | -0.36026 | 3.546337 | -3.08671 | 0.003216 | 0.030407 | -1.8636 | F-box family protein | elongation zone | 12 h |
| HORVU.MOREX.r3.5HG0512320.1 | -0.74987 | 8.205588 | -3.2896 | 0.001787 | 0.020869 | -1.86364 | Cytochrome P450 | elongation zone | 12 h |
| HORVU.MOREX.r3.1HG0039230.1 | -0.66126 | 5.567693 | -3.12033 | 0.002921 | 0.028505 | -1.86703 | AT-rich interactive domain-containing protein 2 | elongation zone | 12 h |
| HORVU.MOREX.r3.3HG0286100.1 | 1.549848 | 1.587249 | 3.068679 | 0.003384 | 0.03144 | -1.87185 | CASP-like protein | elongation zone | 12 h |
| HORVU.MOREX.r3.3HG0329510.1 | -0.96261 | 1.401423 | -3.05994 | 0.003469 | 0.031904 | -1.87227 | Vacuolar protein sorting protein 25 | elongation zone | 12 h |
| HORVU.MOREX.r3.6HG0559000.1 | -0.43074 | 4.951771 | -3.11632 | 0.002955 | 0.028698 | -1.87248 | Homeobox protein | elongation zone | 12 h |
| HORVU.MOREX.r3.5HG0459600.1 | 0.639745 | 3.96473 | 3.162002 | 0.002592 | 0.026328 | -1.87307 | Serine/threonine-protein kinase | elongation zone | 12 h |
| HORVU.MOREX.r3.7HG0659420.1 | -2.53338 | -0.00292 | -3.16431 | 0.002575 | 0.026239 | -1.87311 | Tetratricopeptide repeat protein | elongation zone | 12 h |
| HORVU.MOREX.r3.7HG0708050.1 | -2.63562 | -1.4308 | -3.45939 | 0.001077 | 0.015372 | -1.87337 | Cytochrome b561 | elongation zone | 12 h |
| HORVU.MOREX.r3.5HG0455080.1 | -0.95704 | 2.618738 | -3.04008 | 0.00367 | 0.03307 | -1.87349 | 5-formyltetrahydrofolate cyclo-ligase | elongation zone | 12 h |
| HORVU.MOREX.r3.3HG0231900.1 | -0.24249 | 8.11606 | -3.2875 | 0.001798 | 0.020951 | -1.87432 | Translocon-associated protein subunit beta | elongation zone | 12 h |
| HORVU.MOREX.r3.6HG0564910.1 | -1.01006 | -0.11724 | -3.04036 | 0.003667 | 0.033059 | -1.87646 | Myosin-related family protein | elongation zone | 12 h |
| HORVU.MOREX.r3.5HG0532630.1 | -1.2751 | 4.235353 | -3.0447 | 0.003622 | 0.032772 | -1.87676 | Sister chromatid cohesion protein DCC1 | elongation zone | 12 h |
| HORVU.MOREX.r3.3HG0292490.1 | 0.834397 | 0.56764 | 3.172027 | 0.002518 | 0.025821 | -1.87684 | Core-2/I-branching beta-1,6-N-acetylglucosaminyltransferase family protein, putat | elongation zone | 12 h |
| HORVU.MOREX.r3.4HG0356280.1 | 1.067174 | 0.330026 | 3.03784 | 0.003693 | 0.033251 | -1.87684 | Serine/threonine-protein kinase | elongation zone | 12 h |
| HORVU.MOREX.r3.4HG0353950.1 | 2.947045 | -1.81142 | 3.506945 | 0.000932 | 0.014175 | -1.87707 | ubiquitin carboxyl-terminal hydrolase | elongation zone | 12 h |
| HORVU.MOREX.r3.2HG0182990.1 | -0.65441 | 2.816822 | -3.11347 | 0.002979 | 0.028878 | -1.87831 | UDP-glycosyltransferase | elongation zone | 12 h |
| HORVU.MOREX.r3.2HG0125050.1 | -0.45196 | 4.076346 | -3.10272 | 0.003072 | 0.029497 | -1.87834 | Protein FAR1-RELATED SEQUENCE 5 | elongation zone | 12 h |
| HORVU.MOREX.r3.3HG0298090.1 | 0.803517 | 2.186755 | 3.074308 | 0.003331 | 0.031125 | -1.88048 | Leucine-rich repeat receptor-like protein kinase | elongation zone | 12 h |
| HORVU.MOREX.r3.2HG0152890.1 | -0.65497 | 3.809628 | -3.08598 | 0.003222 | 0.030428 | -1.88136 | TCP transcription factor | elongation zone | 12 h |
| HORVU.MOREX.r3.3HG0296810.1 | -0.90733 | 1.903174 | -3.05141 | 0.003554 | 0.032357 | -1.88247 | 60S ribosomal protein l35a | elongation zone | 12 h |
| HORVU.MOREX.r3.1HG0057000.1 | -1.22528 | 1.397457 | -3.05856 | 0.003483 | 0.031953 | -1.88341 | Pentatricopeptide repeat-containing protein | elongation zone | 12 h |
| HORVU.MOREX.r3.2HG0149940.1 | -0.47089 | 5.130258 | -3.12939 | 0.002847 | 0.028054 | -1.88478 | RNA-binding family protein | elongation zone | 12 h |
| HORVU.MOREX.r3.2HG0127020.1 | 0.685112 | 4.632959 | 3.193464 | 0.002366 | 0.0247 | -1.88525 | methyl esterase 11 | elongation zone | 12 h |
| HORVU.MOREX.r3.1HG0073030.1 | -0.81021 | 5.962504 | -3.15825 | 0.00262 | 0.026447 | -1.88619 | Histone H4 | elongation zone | 12 h |
| HORVU.MOREX.r3.3HG0234800.1 | -0.27897 | 6.705697 | -3.20528 | 0.002287 | 0.0242 | -1.88743 | Carboxymethylenebutenolidase 1 | elongation zone | 12 h |
| HORVU.MOREX.r3.2HG0189970.1 | -0.71522 | 2.22472 | -3.03292 | 0.003744 | 0.033538 | -1.88946 | NBS-LRR disease resistance protein homologue | elongation zone | 12 h |
| HORVU.MOREX.r3.7HG0656330.1 | -1.49235 | 0.123232 | -3.12871 | 0.002852 | 0.028069 | -1.88976 | actin cross-linking protein, putative (DUF569) | elongation zone | 12 h |
| HORVU.MOREX.r3.2HG0217610.1 | -0.30704 | 5.812248 | -3.20198 | 0.002309 | 0.02437 | -1.8903 | S-adenosyl-L-methionine-dependent methyltransferases superfamily protein | elongation zone | 12 h |
| HORVU.MOREX.r3.4HG0399530.1 | 1.708693 | 1.156384 | 3.080203 | 0.003276 | 0.03079 | -1.89185 | Zinc finger protein | elongation zone | 12 h |
| HORVU.MOREX.r3.3HG0264700.1 | -0.34001 | 4.24631 | -3.11145 | 0.002996 | 0.029004 | -1.89256 | ELL-associated factor, putative | elongation zone | 12 h |
| HORVU.MOREX.r3.4HG0407290.1 | -1.41032 | 5.558259 | -3.07974 | 0.00328 | 0.030803 | -1.89278 | Xyloglucan galactosyltransferase KATAMARI1 | elongation zone | 12 h |
| HORVU.MOREX.r3.5HG0536710.1 | -0.4416 | 6.624729 | -3.22099 | 0.002185 | 0.023706 | -1.89362 | NAD/NADP-dependent betaine aldehyde dehydrogenase | elongation zone | 12 h |
| HORVU.MOREX.r3.2HG0184780.1 | -0.76014 | 4.07506 | -3.05744 | 0.003494 | 0.032008 | -1.89394 | Required to maintain repression 2 | elongation zone | 12 h |
| HORVU.MOREX.r3.4HG0396030.1 | 1.838722 | 0.582741 | 3.077569 | 0.0033 | 0.030936 | -1.89638 | Tubulin alpha chain | elongation zone | 12 h |
| HORVU.MOREX.r3.2HG0170390.1 | -0.31913 | 4.904965 | -3.13926 | 0.002767 | 0.027483 | -1.8984 | Protein phosphatase methylesterase 1 | elongation zone | 12 h |
| HORVU.MOREX.r3.7HG0685550.1 | -0.73095 | 3.473275 | -3.05012 | 0.003567 | 0.032388 | -1.9004 | S-acyltransferase | elongation zone | 12 h |
| HORVU.MOREX.r3.7HG0745120.1 | 0.353556 | 6.029656 | 3.198071 | 0.002335 | 0.024484 | -1.90161 | Sucrose phosphate synthase | elongation zone | 12 h |

**Table S1** Continued.

| ID | logFC | AveExpr | t | P.Value | adj.P.Val | B | gene description | root zone | time point |
|---|---|---|---|---|---|---|---|---|---|
| HORVU.MOREX.r3.4HG0392690.1 | 0.444601 | 3.767902 | 3.134347 | 0.002806 | 0.027793 | -1.90172 | Acyl-CoA-binding domain protein | elongation zone | 12 h |
| HORVU.MOREX.r3.5HG0528570.1 | 0.960661 | 0.170023 | 3.094207 | 0.003148 | 0.02993 | -1.90498 | Protein PLANT CADMIUM RESISTANCE 2 | elongation zone | 12 h |
| HORVU.MOREX.r3.7HG0702050.1 | 1.015866 | -0.45045 | 3.037414 | 0.003697 | 0.033276 | -1.90601 | Blue copper protein | elongation zone | 12 h |
| HORVU.MOREX.r3.3HG0304080.1 | -0.86263 | 4.173396 | -3.05506 | 0.003518 | 0.032152 | -1.90686 | Glutaredoxin-like | elongation zone | 12 h |
| HORVU.MOREX.r3.3HG0275300.1 | -0.96952 | 0.578299 | -3.0535 | 0.003533 | 0.032266 | -1.90777 | Major facilitator superfamily protein, putative | elongation zone | 12 h |
| HORVU.MOREX.r3.7HG0657010.1 | 0.521406 | 4.684137 | 3.206649 | 0.002278 | 0.024194 | -1.90819 | Vesicle-associated membrane protein | elongation zone | 12 h |
| HORVU.MOREX.r3.7HG0721790.1 | -0.78653 | 2.548734 | -3.02466 | 0.003833 | 0.034006 | -1.90903 | Pentatricopeptide repeat-containing protein, putative | elongation zone | 12 h |
| HORVU.MOREX.r3.1HG0067930.1 | 0.559631 | 2.533288 | 3.080227 | 0.003275 | 0.03079 | -1.91217 | Peroxidase | elongation zone | 12 h |
| HORVU.MOREX.r3.3HG0230880.1 | -0.20813 | 7.037661 | -3.24681 | 0.002026 | 0.022557 | -1.91241 | Steroid 5-alpha reductase | elongation zone | 12 h |
| HORVU.MOREX.r3.3HG0316870.1 | 0.366972 | 6.819357 | 3.25282 | 0.001991 | 0.022408 | -1.91268 | RNA binding protein | elongation zone | 12 h |
| HORVU.MOREX.r3.3HG0226060.1 | -1.26342 | 4.441713 | -3.09588 | 0.003133 | 0.029866 | -1.91294 | Calcium binding family protein | elongation zone | 12 h |
| HORVU.MOREX.r3.7HG0700930.1 | -1.43826 | 4.508977 | -3.02159 | 0.003866 | 0.034222 | -1.91517 | RING/U-box superfamily protein | elongation zone | 12 h |
| HORVU.MOREX.r3.3HG0294040.1 | 0.779442 | 3.082547 | 3.213537 | 0.002232 | 0.023972 | -1.9154 | Serine/threonine-protein kinase | elongation zone | 12 h |
| HORVU.MOREX.r3.7HG0732610.1 | -1.24274 | 4.827785 | -3.03096 | 0.003765 | 0.03365 | -1.91553 | DNA polymerase alpha subunit B | elongation zone | 12 h |
| HORVU.MOREX.r3.6HG0553290.1 | -0.60792 | 5.490398 | -3.1102 | 0.003007 | 0.029025 | -1.91929 | Glycosyltransferase | elongation zone | 12 h |
| HORVU.MOREX.r3.6HG0622120.1 | 0.573799 | 4.581543 | 3.251271 | 0.002 | 0.022473 | -1.91954 | Pathogenesis-related protein 1 | elongation zone | 12 h |
| HORVU.MOREX.r3.2HG0123650.1 | -0.39028 | 5.958028 | -3.14166 | 0.002748 | 0.027347 | -1.92009 | Serine/arginine rich splicing factor, putative | elongation zone | 12 h |
| HORVU.MOREX.r3.1HG0057860.1 | 0.657909 | 3.261596 | 3.235977 | 0.002091 | 0.023083 | -1.92297 | BTB/POZ domain-containing protein | elongation zone | 12 h |
| HORVU.MOREX.r3.5HG0476000.1 | -0.3452 | 5.038015 | -3.16323 | 0.002583 | 0.026274 | -1.92532 | Molybdopterin synthase catalytic subunit | elongation zone | 12 h |
| HORVU.MOREX.r3.1HG0074490.1 | -1.22124 | 4.901731 | -3.05192 | 0.003549 | 0.032342 | -1.92568 | #N/A | elongation zone | 12 h |
| HORVU.MOREX.r3.7HG0725450.1 | 1.580461 | -0.1554 | 3.207722 | 0.002271 | 0.02416 | -1.92572 | Mitochondrial carrier protein, putative | elongation zone | 12 h |
| HORVU.MOREX.r3.6HG0587460.1 | 0.953828 | -2.65712 | 3.029513 | 0.003781 | 0.0337 | -1.92644 | D-alanine--D-alanine ligase | elongation zone | 12 h |
| HORVU.MOREX.r3.7HG0744970.1 | -0.29501 | 5.768018 | -3.16901 | 0.00254 | 0.025968 | -1.9269 | Protein OS-9 | elongation zone | 12 h |
| HORVU.MOREX.r3.2HG0154500.1 | -1.03982 | 0.570465 | -3.06325 | 0.003437 | 0.03178 | -1.92699 | Protein PHLOEM PROTEIN 2-LIKE A10 | elongation zone | 12 h |
| HORVU.MOREX.r3.1HG0008400.1 | -0.8133 | 2.111607 | -3.01726 | 0.003913 | 0.034492 | -1.92885 | F-box family protein | elongation zone | 12 h |
| HORVU.MOREX.r3.2HG0171860.1 | -0.61819 | 4.629161 | -3.0593 | 0.003476 | 0.031939 | -1.93009 | mediator of RNA polymerase II transcription subunit-like protein | elongation zone | 12 h |
| HORVU.MOREX.r3.1HG0027200.1 | 0.811802 | 4.502879 | 3.098443 | 0.00311 | 0.029766 | -1.93023 | Auxin-responsive protein | elongation zone | 12 h |
| HORVU.MOREX.r3.5HG0523860.1 | 0.518181 | 4.167581 | 3.137463 | 0.002781 | 0.027598 | -1.93129 | Calcium-dependent protein kinase | elongation zone | 12 h |
| HORVU.MOREX.r3.4HG0405830.1 | -0.78982 | 2.230818 | -3.01406 | 0.003949 | 0.03467 | -1.93154 | cold regulated protein 27 | elongation zone | 12 h |
| HORVU.MOREX.r3.3HG0328990.1 | -0.41816 | 4.328412 | -3.10572 | 0.003046 | 0.02931 | -1.93252 | Rab5-interacting family protein | elongation zone | 12 h |
| HORVU.MOREX.r3.6HG0577170.1 | -1.73295 | 1.196593 | -3.16468 | 0.002572 | 0.026239 | -1.93461 | Argonaute family protein | elongation zone | 12 h |
| HORVU.MOREX.r3.7HG0677860.1 | -2.40034 | 2.903041 | -3.07054 | 0.003367 | 0.031359 | -1.93566 | RNA polymerase II elongation factor | elongation zone | 12 h |
| HORVU.MOREX.r3.5HG0514790.3 | -0.47526 | 10.92396 | -3.33591 | 0.001558 | 0.019246 | -1.9358 | Elongation factor 1-alpha | elongation zone | 12 h |
| HORVU.MOREX.r3.4HG0417160.1 | 0.573336 | 4.344914 | 3.214704 | 0.002225 | 0.023928 | -1.93581 | Flavin-containing monooxygenase | elongation zone | 12 h |
| HORVU.MOREX.r3.4HG0378080.1 | 2.881676 | -1.12859 | 3.471349 | 0.001038 | 0.015035 | -1.9363 | #N/A | elongation zone | 12 h |
| HORVU.MOREX.r3.1HG0029080.1 | -0.64928 | 2.288361 | -3.02701 | 0.003807 | 0.033836 | -1.93714 | 60S ribosomal protein L18a-like protein | elongation zone | 12 h |
| HORVU.MOREX.r3.5HG0479210.1 | -1.01622 | 7.833409 | -3.17528 | 0.002494 | 0.025643 | -1.93764 | HMG-Y-related protein A | elongation zone | 12 h |
| HORVU.MOREX.r3.2HG0150530.1 | 0.566041 | 3.135946 | 3.070867 | 0.003364 | 0.031344 | -1.93828 | E3 ubiquitin-protein ligase family | elongation zone | 12 h |
| HORVU.MOREX.r3.3HG0301250.1 | -1.17061 | 1.880477 | -3.0167 | 0.003919 | 0.034516 | -1.93842 | #N/A | elongation zone | 12 h |
| HORVU.MOREX.r3.4HG0417710.1 | 0.893247 | 0.768626 | 3.159737 | 0.002609 | 0.026441 | -1.9395 | Protease inhibitor/seed storage/lipid transfer protein family protein | elongation zone | 12 h |
| HORVU.MOREX.r3.1HG0073320.1 | -0.91593 | 2.923796 | -3.03513 | 0.003721 | 0.033412 | -1.9404 | Histone H2B | elongation zone | 12 h |

**Table S1** Continued.

| ID | logFC | AveExpr | t | P.Value | adj.P.Val | B | gene description | root zone | time point |
|---|---|---|---|---|---|---|---|---|---|
| HORVU.MOREX.r3.2HG0135510.1 | -0.34359 | 5.626394 | -3.15915 | 0.002613 | 0.026441 | -1.94176 | Vesicle-associated 4-2-like protein | elongation zone | 12 h |
| HORVU.MOREX.r3.1HG0083110.1 | -0.54962 | 2.265521 | -3.02756 | 0.003801 | 0.033827 | -1.945 | UDP-D-apiose/UDP-D-xylose synthase | elongation zone | 12 h |
| HORVU.MOREX.r3.3HG0255900.1 | -0.29148 | 6.088112 | -3.17611 | 0.002488 | 0.025607 | -1.94579 | SUN domain-containing protein 2 | elongation zone | 12 h |
| HORVU.MOREX.r3.6HG0632270.1 | -0.43922 | 3.993705 | -3.07726 | 0.003303 | 0.03095 | -1.94886 | Leucine-rich repeat family protein | elongation zone | 12 h |
| HORVU.MOREX.r3.7HG0714120.1 | -0.33138 | 6.206134 | -3.15205 | 0.002667 | 0.026789 | -1.94913 | Eukaryotic translation initiation factor 3 subunit I | elongation zone | 12 h |
| HORVU.MOREX.r3.1HG0025320.1 | 1.07233 | 0.572545 | 3.006327 | 0.004035 | 0.035207 | -1.94964 | Receptor-like kinase, putative | elongation zone | 12 h |
| HORVU.MOREX.r3.6HG0633420.1 | -0.7963 | 7.127371 | -3.14873 | 0.002693 | 0.027002 | -1.95035 | Pentatricopeptide repeat-containing protein | elongation zone | 12 h |
| HORVU.MOREX.r3.4HG0331630.1 | -0.39182 | 4.435596 | -3.08841 | 0.0032 | 0.030344 | -1.95121 | Short-chain dehydrogenase/reductase family protein | elongation zone | 12 h |
| HORVU.MOREX.r3.7HG0708820.1 | -0.6856 | 5.143803 | -3.05743 | 0.003494 | 0.032008 | -1.95134 | PHD finger family protein | elongation zone | 12 h |
| HORVU.MOREX.r3.6HG0540620.1 | -1.53898 | 4.022998 | -3.00629 | 0.004035 | 0.035207 | -1.95156 | Malignant brain tumor repeat protein 1 | elongation zone | 12 h |
| HORVU.MOREX.r3.5HG0528910.1 | 0.851303 | 1.611089 | 3.094268 | 0.003147 | 0.02993 | -1.952 | DUF3511 domain protein | elongation zone | 12 h |
| HORVU.MOREX.r3.2HG0198320.1 | 1.558602 | -0.19189 | 3.019617 | 0.003887 | 0.03428 | -1.95255 | Protein translation factor SUI1-like protein | elongation zone | 12 h |
| HORVU.MOREX.r3.3HG0298510.1 | 0.572642 | 4.425452 | 3.118993 | 0.002933 | 0.028562 | -1.95301 | 1-phosphatidylinositol-3-phosphate 5-kinase | elongation zone | 12 h |
| HORVU.MOREX.r3.3HG0275100.1 | 1.283142 | -2.73635 | 3.032277 | 0.003751 | 0.033585 | -1.95377 | Integrase-type DNA-binding superfamily protein | elongation zone | 12 h |
| HORVU.MOREX.r3.2HG0095520.1 | -0.5947 | 4.42382 | -3.10815 | 0.003025 | 0.029127 | -1.95454 | Histone deacetylase complex subunit SAP30 | elongation zone | 12 h |
| HORVU.MOREX.r3.5HG0483980.1 | 1.165309 | -1.84454 | 3.007284 | 0.004024 | 0.035139 | -1.95566 | Cinnamoyl CoA reductase | elongation zone | 12 h |
| HORVU.MOREX.r3.1HG0029730.1 | 0.470925 | 7.923005 | 3.280547 | 0.001835 | 0.021263 | -1.95575 | Malic enzyme | elongation zone | 12 h |
| HORVU.MOREX.r3.3HG0249590.1 | -0.69958 | 3.60705 | -3.01198 | 0.003972 | 0.034843 | -1.95701 | Vesicle transport protein-like | elongation zone | 12 h |
| HORVU.MOREX.r3.5HG0510210.1 | -0.37319 | 4.966015 | -3.14137 | 0.00275 | 0.027347 | -1.95737 | Tryptophan RNA-binding attenuator-like protein | elongation zone | 12 h |
| HORVU.MOREX.r3.7HG0647300.1 | 1.785206 | -1.17681 | 3.005199 | 0.004048 | 0.03527 | -1.9576 | AT hook motif DNA-binding family protein | elongation zone | 12 h |
| HORVU.MOREX.r3.2HG0189980.1 | 0.501743 | 3.503027 | 3.104946 | 0.003053 | 0.02934 | -1.95806 | PPPDE thiol peptidase family protein, putative | elongation zone | 12 h |
| HORVU.MOREX.r3.4HG0401710.1 | -1.27885 | 4.660057 | -3.01313 | 0.003959 | 0.034745 | -1.95808 | Histone H2B | elongation zone | 12 h |
| HORVU.MOREX.r3.6HG0631700.1 | 0.441617 | 8.078937 | 3.324292 | 0.001613 | 0.019712 | -1.96123 | Aquaporin | elongation zone | 12 h |
| HORVU.MOREX.r3.2HG0190260.1 | 0.841578 | 2.916863 | 3.011066 | 0.003982 | 0.034902 | -1.96192 | SHR5-receptor-like kinase | elongation zone | 12 h |
| HORVU.MOREX.r3.3HG0229480.1 | -0.38936 | 5.27277 | -3.09645 | 0.003128 | 0.029864 | -1.96504 | Actin/actin-like family protein | elongation zone | 12 h |
| HORVU.MOREX.r3.2HG0099160.1 | 0.984643 | -0.22736 | 3.051875 | 0.003549 | 0.032342 | -1.96606 | 2-oxoglutarate (2OG) and Fe(II)-dependent oxygenase superfamily protein, putativ | elongation zone | 12 h |
| HORVU.MOREX.r3.5HG0526940.1 | -0.36095 | 7.55668 | -3.20302 | 0.002302 | 0.024309 | -1.96613 | Alpha-mannosidase | elongation zone | 12 h |
| HORVU.MOREX.r3.6HG0623920.1 | -0.20249 | 9.176211 | -3.28919 | 0.001789 | 0.020878 | -1.96622 | Leucine aminopeptidase, putative | elongation zone | 12 h |
| HORVU.MOREX.r3.1HG0008140.1 | -1.05263 | 2.579787 | -3.04228 | 0.003647 | 0.032938 | -1.9678 | Glycosyltransferase | elongation zone | 12 h |
| HORVU.MOREX.r3.2HG0130000.1 | -0.35093 | 4.729393 | -3.09766 | 0.003117 | 0.029789 | -1.96871 | Effector of transcription protein | elongation zone | 12 h |
| HORVU.MOREX.r3.7HG0732270.1 | -0.565 | 2.663349 | -3.00074 | 0.004099 | 0.035578 | -1.97072 | F-box protein | elongation zone | 12 h |
| HORVU.MOREX.r3.4HG0367380.1 | -2.69387 | -1.8835 | -3.45291 | 0.001098 | 0.015536 | -1.97145 | zinc finger (ubiquitin-hydrolase) domain-containing protein | elongation zone | 12 h |
| HORVU.MOREX.r3.2HG0180260.1 | -0.69862 | 1.836043 | -2.99652 | 0.004147 | 0.035879 | -1.97155 | Myb/SANT-like DNA-binding domain protein | elongation zone | 12 h |
| HORVU.MOREX.r3.5HG0446900.1 | -0.22366 | 6.633774 | -3.19451 | 0.002359 | 0.024647 | -1.97253 | Dipeptidyl peptidase | elongation zone | 12 h |
| HORVU.MOREX.r3.7HG0751000.1 | -1.25265 | 3.800431 | -3.01423 | 0.003947 | 0.034668 | -1.97319 | UDP-glycosyltransferase | elongation zone | 12 h |
| HORVU.MOREX.r3.3HG0305580.1 | -1.38155 | 7.017223 | -3.07602 | 0.003315 | 0.031043 | -1.98 | Alcohol dehydrogenase class-3 | elongation zone | 12 h |
| HORVU.MOREX.r3.6HG0553310.1 | -0.78567 | 4.173624 | -3.00442 | 0.004057 | 0.035331 | -1.98108 | Pentatricopeptide repeat-containing protein | elongation zone | 12 h |
| HORVU.MOREX.r3.3HG0294670.1 | -1.38307 | -0.36802 | -3.11435 | 0.002972 | 0.028819 | -1.98174 | demeter-like protein 3 | elongation zone | 12 h |
| HORVU.MOREX.r3.5HG0441840.1 | 1.318125 | 1.936883 | 3.02988 | 0.003777 | 0.03368 | -1.98207 | Dof zinc finger protein | elongation zone | 12 h |
| HORVU.MOREX.r3.2HG0197230.1 | 1.110772 | -1.20366 | 3.00241 | 0.00408 | 0.035442 | -1.98435 | disease resistance protein (TIR-NBS-LRR class) | elongation zone | 12 h |
| HORVU.MOREX.r3.3HG0303430.1 | -0.4641 | 7.726619 | -3.21965 | 0.002193 | 0.023748 | -1.98449 | TBC1 domain family member 8 | elongation zone | 12 h |

**Table S1** Continued.

| ID | logFC | AveExpr | t | P.Value | adj.P.Val | B | gene description | root zone | time point |
|---|---|---|---|---|---|---|---|---|---|
| HORVU.MOREX.r3.3HG0288070.1 | 0.522583 | 0.496177 | 3.109651 | 0.003012 | 0.02903 | -1.98525 | 2-oxoglutarate (2OG) and Fe(II)-dependent oxygenase superfamily protein, putati | elongation zone | 12 h |
| HORVU.MOREX.r3.1HG0088990.1 | -0.51988 | 4.115581 | -3.05264 | 0.003542 | 0.032316 | -1.98549 | #N/A | elongation zone | 12 h |
| HORVU.MOREX.r3.7HG0711850.1 | -0.38883 | 4.991548 | -3.08707 | 0.003212 | 0.03039 | -1.986 | RNA-binding KH domain-containing protein | elongation zone | 12 h |
| HORVU.MOREX.r3.1HG0044470.1 | 0.770877 | 2.284476 | 3.035122 | 0.003721 | 0.033412 | -1.98638 | Cytokinin oxidase/dehydrogenase | elongation zone | 12 h |
| HORVU.MOREX.r3.7HG0672960.1 | 1.534157 | -2.45339 | 3.036797 | 0.003704 | 0.033319 | -1.98698 | U-box domain-containing family protein | elongation zone | 12 h |
| HORVU.MOREX.r3.2HG0177200.1 | -0.53736 | 3.273869 | -3.05288 | 0.003539 | 0.032308 | -1.98709 | Cytochrome f | elongation zone | 12 h |
| HORVU.MOREX.r3.7HG0650950.1 | 0.442875 | 4.47289 | 3.120185 | 0.002923 | 0.028505 | -1.99007 | Cysteine synthase | elongation zone | 12 h |
| HORVU.MOREX.r3.1HG0074010.1 | 1.002605 | 1.165265 | 3.009735 | 0.003997 | 0.034974 | -1.99413 | Guanylyl cyclase | elongation zone | 12 h |
| HORVU.MOREX.r3.3HG0221440.1 | 0.799814 | 2.700751 | 3.24899 | 0.002013 | 0.022519 | -1.9951 | Glycosyltransferase | elongation zone | 12 h |
| HORVU.MOREX.r3.2HG0095970.1 | -1.27234 | 3.317154 | -2.98691 | 0.00426 | 0.036533 | -1.99651 | NBS-LRR disease resistance protein-like protein | elongation zone | 12 h |
| HORVU.MOREX.r3.1HG0073500.1 | 1.886354 | -2.89345 | 2.99601 | 0.004153 | 0.035915 | -1.99676 | Laccase | elongation zone | 12 h |
| HORVU.MOREX.r3.6HG0614470.2 | 0.50449 | 0.971508 | 3.136743 | 0.002787 | 0.027642 | -1.99818 | Beta-xylosidase, putative | elongation zone | 12 h |
| HORVU.MOREX.r3.5HG0506290.1 | -0.88126 | 1.60619 | -2.9913 | 0.004208 | 0.036196 | -2.00106 | Splicing factor u2af large subunit, putative | elongation zone | 12 h |
| HORVU.MOREX.r3.1HG0072410.1 | -0.26945 | 6.475901 | -3.17261 | 0.002514 | 0.02579 | -2.00136 | Calcium-dependent protein kinase | elongation zone | 12 h |
| HORVU.MOREX.r3.5HG0471520.1 | 0.549076 | 3.519872 | 3.080304 | 0.003275 | 0.03079 | -2.00458 | Ubiquitin-conjugating enzyme E2 | elongation zone | 12 h |
| HORVU.MOREX.r3.7HG0743900.1 | 0.510369 | 5.143712 | 3.170434 | 0.002529 | 0.025888 | -2.00664 | Kinase family protein | elongation zone | 12 h |
| HORVU.MOREX.r3.6HG0596240.1 | -1.16534 | 0.960124 | -3.01564 | 0.003931 | 0.034575 | -2.00818 | RNA-binding protein | elongation zone | 12 h |
| HORVU.MOREX.r3.1HG0094980.1 | -0.42992 | 4.287159 | -3.05997 | 0.003469 | 0.031904 | -2.00897 | Mitochondrial outer membrane porin | elongation zone | 12 h |
| HORVU.MOREX.r3.5HG0478390.1 | -0.4779 | 5.551367 | -3.08716 | 0.003211 | 0.03039 | -2.00944 | PCI domain-containing protein 2 | elongation zone | 12 h |
| HORVU.MOREX.r3.5HG0495240.1 | 1.058 | -0.87388 | 2.984361 | 0.004291 | 0.036763 | -2.01204 | Kinase family protein | elongation zone | 12 h |
| HORVU.MOREX.r3.6HG0594290.1 | -1.0226 | 2.905971 | -2.98083 | 0.004333 | 0.036957 | -2.01382 | histone acetyltransferase of the CBP family 12 | elongation zone | 12 h |
| HORVU.MOREX.r3.3HG0291340.1 | -0.96811 | 3.400685 | -2.98248 | 0.004313 | 0.036879 | -2.01426 | Deoxycytidylate deaminase-like | elongation zone | 12 h |
| HORVU.MOREX.r3.2HG0193220.1 | -0.64547 | 4.02011 | -3.00363 | 0.004066 | 0.035366 | -2.01432 | Putrescine-binding periplasmic protein | elongation zone | 12 h |
| HORVU.MOREX.r3.2HG0196630.1 | 0.966007 | -0.81111 | 3.019608 | 0.003887 | 0.03428 | -2.01454 | Invertase/pectin methylesterase inhibitor family protein | elongation zone | 12 h |
| HORVU.MOREX.r3.6HG0613460.1 | 1.074621 | 2.574467 | 3.031068 | 0.003764 | 0.03365 | -2.01474 | MACPF domain-containing protein | elongation zone | 12 h |
| HORVU.MOREX.r3.7HG0656250.1 | -0.39682 | 7.976001 | -3.21062 | 0.002251 | 0.0241 | -2.0149 | Proteasome subunit alpha type | elongation zone | 12 h |
| HORVU.MOREX.r3.6HG0601160.1 | 0.683812 | 4.335499 | 3.28678 | 0.001802 | 0.020972 | -2.01504 | Expansin protein | elongation zone | 12 h |
| HORVU.MOREX.r3.2HG0119060.1 | -0.47205 | 4.049026 | -3.04068 | 0.003663 | 0.033043 | -2.01763 | zinc transporter-like protein | elongation zone | 12 h |
| HORVU.MOREX.r3.7HG0714870.1 | -0.61942 | 5.526539 | -3.12134 | 0.002913 | 0.028463 | -2.01958 | Glycosyltransferase | elongation zone | 12 h |
| HORVU.MOREX.r3.4HG0386480.1 | -0.39888 | 8.895935 | -3.24913 | 0.002012 | 0.022519 | -2.01982 | Calcium-transporting ATPase | elongation zone | 12 h |
| HORVU.MOREX.r3.6HG0633510.1 | 0.580872 | 4.690627 | 3.158419 | 0.002619 | 0.026447 | -2.0202 | integral membrane metal-binding family protein (DUF2296) | elongation zone | 12 h |
| HORVU.MOREX.r3.2HG0138540.1 | 0.600067 | 4.435354 | 3.266212 | 0.001914 | 0.021808 | -2.02032 | Blue copper protein | elongation zone | 12 h |
| HORVU.MOREX.r3.2HG0202220.1 | 0.348704 | 4.425224 | 3.057861 | 0.00349 | 0.031998 | -2.02085 | Protein pelota homolog | elongation zone | 12 h |
| HORVU.MOREX.r3.1HG0079750.1 | 0.974438 | -1.15538 | 2.988952 | 0.004236 | 0.036371 | -2.02354 | MYB transcription factor | elongation zone | 12 h |
| HORVU.MOREX.r3.5HG0500730.1 | -0.66802 | 4.668649 | -3.03017 | 0.003774 | 0.03368 | -2.02488 | Mediator of RNA polymerase II transcription subunit 4 | elongation zone | 12 h |
| HORVU.MOREX.r3.5HG0479810.1 | 0.237483 | 7.043626 | 3.20346 | 0.002299 | 0.024291 | -2.02534 | Phospholipid:diacylglycerol acyltransferase | elongation zone | 12 h |
| HORVU.MOREX.r3.3HG0245010.1 | 0.470893 | 4.432877 | 3.098402 | 0.00311 | 0.029766 | -2.0271 | Protein DJ-1 | elongation zone | 12 h |
| HORVU.MOREX.r3.4HG0396660.1 | -1.61252 | 6.770443 | -3.05121 | 0.003556 | 0.032357 | -2.02792 | Replication protein A 70 kDa DNA-binding subunit | elongation zone | 12 h |
| HORVU.MOREX.r3.4HG0404450.1 | -0.28227 | 5.835227 | -3.11273 | 0.002986 | 0.028925 | -2.03102 | Nitrilase/cyanide hydratase and apolipoprotein N-acyltransferase family protein | elongation zone | 12 h |
| HORVU.MOREX.r3.3HG0234970.1 | 0.519161 | 0.091122 | 3.070163 | 0.00337 | 0.031367 | -2.03224 | 6,7-dimethyl-8-ribityllumazine synthase | elongation zone | 12 h |
| HORVU.MOREX.r3.5HG0429230.1 | -0.49938 | 4.516929 | -3.03577 | 0.003714 | 0.033401 | -2.03238 | Protein FAR1-RELATED SEQUENCE 5 | elongation zone | 12 h |

**Table S1** Continued.

| ID | logFC | AveExpr | t | P.Value | adj.P.Val | B | gene description | root zone | time point |
|---|---|---|---|---|---|---|---|---|---|
| HORVU.MOREX.r3.5HG0473520.1 | 0.391991 | 6.096551 | 3.197493 | 0.002339 | 0.0245 | -2.03286 | UDP-glucose 4-epimerase family protein | elongation zone | 12 h |
| HORVU.MOREX.r3.3HG0297940.1 | -1.178 | 3.507795 | -2.97299 | 0.004429 | 0.037408 | -2.03338 | Transcription initiation factor TFIID subunit 9 | elongation zone | 12 h |
| HORVU.MOREX.r3.1HG0084890.1 | 0.610711 | 1.342406 | 3.027806 | 0.003799 | 0.033827 | -2.03734 | Protein DETOXIFICATION | elongation zone | 12 h |
| HORVU.MOREX.r3.7HG0718710.1 | -0.2782 | 6.996212 | -3.18991 | 0.002391 | 0.024902 | -2.03864 | Ras-related protein, expressed | elongation zone | 12 h |
| HORVU.MOREX.r3.5HG0433490.1 | -0.50341 | 5.319229 | -3.07278 | 0.003345 | 0.031231 | -2.04014 | F-box/LRR protein, putative | elongation zone | 12 h |
| HORVU.MOREX.r3.6HG0582230.1 | -0.51329 | 3.581325 | -3.01698 | 0.003916 | 0.034505 | -2.04063 | GDSL esterase/lipase | elongation zone | 12 h |
| HORVU.MOREX.r3.4HG0338210.1 | -0.77598 | 3.760864 | -3.02357 | 0.003844 | 0.034053 | -2.04139 | Histone H2A | elongation zone | 12 h |
| HORVU.MOREX.r3.2HG0120470.1 | 1.17548 | 3.222129 | 3.232344 | 0.002113 | 0.023229 | -2.04479 | Peroxidase | elongation zone | 12 h |
| HORVU.MOREX.r3.3HG0225760.1 | 1.188559 | -0.95266 | 3.034175 | 0.003731 | 0.033458 | -2.04491 | ABC transporter G family member | elongation zone | 12 h |
| HORVU.MOREX.r3.2HG0189950.1 | -0.43769 | 4.476014 | -3.06744 | 0.003396 | 0.031493 | -2.04613 | Receptor-like kinase | elongation zone | 12 h |
| HORVU.MOREX.r3.1HG0025430.1 | 1.184581 | 0.604962 | 2.963721 | 0.004545 | 0.03803 | -2.04629 | WEB family protein (DUF827) | elongation zone | 12 h |
| HORVU.MOREX.r3.2HG0189110.1 | 1.061076 | 0.028785 | 2.967114 | 0.004502 | 0.037785 | -2.04849 | Vesicle-associated protein 1-1 | elongation zone | 12 h |
| HORVU.MOREX.r3.3HG0281860.1 | -0.59301 | 3.606298 | -2.97797 | 0.004368 | 0.037145 | -2.04947 | Peptidyl-tRNA hydrolase | elongation zone | 12 h |
| HORVU.MOREX.r3.3HG0300810.1 | -0.48959 | 6.316915 | -3.16274 | 0.002586 | 0.026299 | -2.05079 | Sialyltransferase-like protein 2 | elongation zone | 12 h |
| HORVU.MOREX.r3.1HG0046770.1 | -0.7079 | 4.616852 | -3.00044 | 0.004102 | 0.035593 | -2.05309 | #N/A | elongation zone | 12 h |
| HORVU.MOREX.r3.6HG0550740.1 | -0.50018 | 3.020677 | -2.97923 | 0.004353 | 0.037092 | -2.05325 | F-box family protein | elongation zone | 12 h |
| HORVU.MOREX.r3.5HG0480890.1 | 1.451091 | -1.46482 | 2.961452 | 0.004574 | 0.038197 | -2.05332 | Dirigent protein | elongation zone | 12 h |
| HORVU.MOREX.r3.5HG0465810.1 | 2.271032 | -0.29224 | 3.039807 | 0.003672 | 0.033081 | -2.0546 | Calcium-binding protein | elongation zone | 12 h |
| HORVU.MOREX.r3.3HG0219650.1 | -0.44238 | 6.043804 | -3.08761 | 0.003207 | 0.030372 | -2.05669 | RNA polymerase II-associated factor 1 | elongation zone | 12 h |
| HORVU.MOREX.r3.6HG0555810.1 | -1.22175 | 0.610303 | -3.01089 | 0.003984 | 0.034905 | -2.05895 | 5'-AMP-activated protein kinase subunit beta-1 | elongation zone | 12 h |
| HORVU.MOREX.r3.5HG0515610.1 | 1.235955 | -3.36667 | 3.023503 | 0.003845 | 0.034053 | -2.05983 | Collagen alpha chain | elongation zone | 12 h |
| HORVU.MOREX.r3.2HG0181800.1 | -0.7995 | 3.895018 | -2.97983 | 0.004345 | 0.037045 | -2.0604 | Protodermal factor 1 | elongation zone | 12 h |
| HORVU.MOREX.r3.5HG0531850.1 | -0.64019 | 4.723485 | -3.00277 | 0.004076 | 0.035421 | -2.06386 | #N/A | elongation zone | 12 h |
| HORVU.MOREX.r3.5HG0514490.1 | -0.50507 | 5.559766 | -3.07135 | 0.003359 | 0.031316 | -2.06504 | Hexosyltransferase | elongation zone | 12 h |
| HORVU.MOREX.r3.3HG0250870.1 | -0.50008 | 4.187521 | -3.03548 | 0.003718 | 0.033412 | -2.06638 | Polynucleotidyl transferase ribonuclease H-like superfamily protein | elongation zone | 12 h |
| HORVU.MOREX.r3.2HG0106750.1 | -1.09118 | 0.325037 | -3.00522 | 0.004048 | 0.03527 | -2.06673 | Alpha-1,4 glucan phosphorylase | elongation zone | 12 h |
| HORVU.MOREX.r3.2HG0216610.1 | -0.33206 | 6.040531 | -3.14862 | 0.002694 | 0.027002 | -2.06684 | Protein NRT1/ PTR FAMILY 5.5 | elongation zone | 12 h |
| HORVU.MOREX.r3.5HG0536610.1 | 0.656104 | 2.755965 | 3.128276 | 0.002856 | 0.028074 | -2.06916 | Gibberellin 20 oxidase | elongation zone | 12 h |
| HORVU.MOREX.r3.3HG0250940.1 | -0.57739 | 3.302703 | -3.00826 | 0.004013 | 0.035058 | -2.07025 | Polynucleotidyl transferase ribonuclease H-like superfamily protein | elongation zone | 12 h |
| HORVU.MOREX.r3.3HG0226150.1 | 1.694694 | 2.218815 | 2.995816 | 0.004156 | 0.035915 | -2.07043 | Protein kinase family protein | elongation zone | 12 h |
| HORVU.MOREX.r3.7HG0640570.1 | -0.42152 | 6.096045 | -3.09619 | 0.00313 | 0.029866 | -2.07112 | Plant/F18O14-17 protein | elongation zone | 12 h |
| HORVU.MOREX.r3.1HG0088900.1 | -0.92274 | 2.109858 | -2.96176 | 0.00457 | 0.038196 | -2.07136 | Orphans transcription factor | elongation zone | 12 h |
| HORVU.MOREX.r3.3HG0278630.1 | 0.459883 | 5.03935 | 3.097956 | 0.003114 | 0.029778 | -2.07242 | Ras-related protein, expressed | elongation zone | 12 h |
| HORVU.MOREX.r3.3HG0237870.1 | -1.65932 | 5.343811 | -2.97548 | 0.004398 | 0.037281 | -2.07297 | Histone-lysine N-methyltransferase | elongation zone | 12 h |
| HORVU.MOREX.r3.1HG0072160.1 | 0.705046 | 2.908685 | 2.974581 | 0.004409 | 0.037331 | -2.07337 | UDP-glucose pyrophosphorylase | elongation zone | 12 h |
| HORVU.MOREX.r3.2HG0162940.1 | -0.33006 | 6.349043 | -3.11722 | 0.002947 | 0.028675 | -2.07343 | Chaperone protein dnaJ | elongation zone | 12 h |
| HORVU.MOREX.r3.5HG0464630.1 | 0.567498 | 3.359325 | 3.141149 | 0.002752 | 0.027347 | -2.07388 | Peroxidase | elongation zone | 12 h |
| HORVU.MOREX.r3.7HG0704760.1 | 1.212351 | -2.61956 | 2.982818 | 0.004309 | 0.036879 | -2.0744 | Myb family transcription factor family protein | elongation zone | 12 h |
| HORVU.MOREX.r3.2HG0210880.1 | -1.15579 | 2.678515 | -2.97387 | 0.004418 | 0.037371 | -2.07455 | #N/A | elongation zone | 12 h |
| HORVU.MOREX.r3.4HG0349170.1 | -1.02342 | 4.645229 | -3.05091 | 0.003559 | 0.032357 | -2.07542 | Long-Chain Acyl-CoA Synthetase | elongation zone | 12 h |
| HORVU.MOREX.r3.3HG0249890.1 | 0.769399 | 2.405613 | 3.074969 | 0.003325 | 0.03108 | -2.0768 | Auxin response factor | elongation zone | 12 h |

**Table S1** Continued.

| ID | logFC | AveExpr | t | P.Value | adj.P.Val | B | gene description | root zone | time point |
|---|---|---|---|---|---|---|---|---|---|
| HORVU.MOREX.r3.6HG0583730.1 | -0.38431 | 4.393564 | -3.02399 | 0.00384 | 0.034035 | -2.07729 | Loricrin-like | elongation zone | 12 h |
| HORVU.MOREX.r3.7HG0653850.1 | -0.6948 | 4.845223 | -3.03346 | 0.003739 | 0.033502 | -2.07815 | Succinate dehydrogenase assembly factor 4, mitochondrial | elongation zone | 12 h |
| HORVU.MOREX.r3.7HG0721800.1 | -0.34807 | 7.386378 | -3.16489 | 0.00257 | 0.026239 | -2.07957 | Malate dehydrogenase | elongation zone | 12 h |
| HORVU.MOREX.r3.4HG0337870.1 | 0.751168 | 4.45297 | 3.094331 | 0.003147 | 0.02993 | -2.08058 | Homeobox-leucine zipper family protein / lipid-binding START domain-containing | elongation zone | 12 h |
| HORVU.MOREX.r3.6HG0573150.1 | 0.389177 | 5.184389 | 3.06493 | 0.003421 | 0.031662 | -2.08374 | Copper-transporting ATPase | elongation zone | 12 h |
| HORVU.MOREX.r3.4HG0388270.1 | -0.41612 | 6.243595 | -3.123 | 0.002899 | 0.028342 | -2.08697 | Ankyrin repeat family protein, putative, expressed | elongation zone | 12 h |
| HORVU.MOREX.r3.3HG0290180.1 | -1.13009 | 3.615358 | -2.94556 | 0.00478 | 0.039249 | -2.08842 | Non-specific serine/threonine protein kinase | elongation zone | 12 h |
| HORVU.MOREX.r3.2HG0176920.1 | -1.23594 | 3.864016 | -2.94683 | 0.004763 | 0.039158 | -2.08871 | ATP-dependent RNA helicase DeaD | elongation zone | 12 h |
| HORVU.MOREX.r3.3HG0273790.2 | 0.668958 | 2.302445 | 2.95523 | 0.004653 | 0.038674 | -2.08881 | CTD small phosphatase-like protein | elongation zone | 12 h |
| HORVU.MOREX.r3.2HG0107300.1 | 1.649842 | -1.31881 | 2.958236 | 0.004615 | 0.038477 | -2.08935 | HXXXD-type acyl-transferase family protein | elongation zone | 12 h |
| HORVU.MOREX.r3.3HG0306170.1 | -0.32995 | 5.38776 | -3.08549 | 0.003227 | 0.030457 | -2.09074 | Selenoprotein | elongation zone | 12 h |
| HORVU.MOREX.r3.7HG0655210.1 | 1.261293 | -1.68992 | 2.985127 | 0.004282 | 0.0367 | -2.0912 | Peroxidase | elongation zone | 12 h |
| HORVU.MOREX.r3.2HG0110730.1 | -0.4073 | 4.019032 | -3.00844 | 0.004011 | 0.035058 | -2.0921 | DNA primase | elongation zone | 12 h |
| HORVU.MOREX.r3.2HG0161580.1 | -0.50391 | 2.675596 | -2.9567 | 0.004634 | 0.038595 | -2.09308 | Alpha/beta-Hydrolases superfamily protein | elongation zone | 12 h |
| HORVU.MOREX.r3.5HG0478610.1 | 0.399512 | 4.6112 | 3.102419 | 0.003075 | 0.029497 | -2.09472 | Malonyl-CoA decarboxylase | elongation zone | 12 h |
| HORVU.MOREX.r3.5HG0502310.1 | -0.71053 | 3.718859 | -2.98275 | 0.00431 | 0.036879 | -2.09548 | NBS-LRR disease resistance protein, putative | elongation zone | 12 h |
| HORVU.MOREX.r3.6HG0577360.1 | -0.65634 | 5.665724 | -3.21381 | 0.002231 | 0.023972 | -2.09549 | transmembrane protein, putative (Protein of unknown function, DUF642) | elongation zone | 12 h |
| HORVU.MOREX.r3.1HG0005730.1 | 1.670036 | -3.38812 | 3.064627 | 0.003424 | 0.031674 | -2.09578 | transcriptional factor B3 family protein | elongation zone | 12 h |
| HORVU.MOREX.r3.5HG0518490.1 | -0.87275 | 0.764022 | -2.94837 | 0.004743 | 0.03909 | -2.09611 | C2 domain-containing protein / GRAM domain-containing protein | elongation zone | 12 h |
| HORVU.MOREX.r3.6HG0558350.1 | 2.504283 | -0.25422 | 3.095776 | 0.003134 | 0.029866 | -2.09926 | #N/A | elongation zone | 12 h |
| HORVU.MOREX.r3.4HG0347760.1 | 0.666669 | 2.347769 | 2.96361 | 0.004546 | 0.03803 | -2.09986 | MYB transcription factor | elongation zone | 12 h |
| HORVU.MOREX.r3.2HG0174650.1 | 0.967072 | 1.398683 | 2.940732 | 0.004844 | 0.039587 | -2.10301 | WPP domain-interacting protein 1 | elongation zone | 12 h |
| HORVU.MOREX.r3.3HG0281000.1 | -0.85124 | 3.470695 | -2.97455 | 0.00441 | 0.037331 | -2.10337 | F-box protein | elongation zone | 12 h |
| HORVU.MOREX.r3.5HG0432040.1 | -0.63153 | 4.077273 | -2.97714 | 0.004378 | 0.037198 | -2.1042 | #N/A | elongation zone | 12 h |
| HORVU.MOREX.r3.2HG0209090.1 | -0.60056 | 3.138841 | -2.94935 | 0.00473 | 0.03901 | -2.10474 | Tetraspanin family protein | elongation zone | 12 h |
| HORVU.MOREX.r3.3HG0250170.1 | -0.58714 | 3.50268 | -3.01135 | 0.003979 | 0.03489 | -2.10484 | Branched-chain amino acid aminotransferase-like | elongation zone | 12 h |
| HORVU.MOREX.r3.2HG0123170.1 | 0.993826 | 0.673299 | 2.964982 | 0.004529 | 0.037933 | -2.10561 | Chaperone protein dnaJ | elongation zone | 12 h |
| HORVU.MOREX.r3.2HG0126380.1 | 0.47792 | 5.874038 | 3.206504 | 0.002279 | 0.024194 | -2.10576 | #N/A | elongation zone | 12 h |
| HORVU.MOREX.r3.5HG0472840.1 | 0.510954 | 2.862571 | 3.056943 | 0.003499 | 0.032032 | -2.10612 | NBS-LRR disease resistance protein | elongation zone | 12 h |
| HORVU.MOREX.r3.6HG0572400.1 | -0.3608 | 3.75498 | -3.01624 | 0.003924 | 0.034532 | -2.10615 | F-box family protein | elongation zone | 12 h |
| HORVU.MOREX.r3.1HG0056810.1 | 0.417771 | 5.129894 | 3.10272 | 0.003072 | 0.029497 | -2.10836 | Haloacid dehalogenase-like hydrolase family protein, putative, expressed | elongation zone | 12 h |
| HORVU.MOREX.r3.6HG0566930.1 | -0.75993 | 4.262539 | -2.96609 | 0.004515 | 0.037877 | -2.10868 | Biotin holocarboxylase synthetase 1-like protein | elongation zone | 12 h |
| HORVU.MOREX.r3.7HG0744890.1 | 1.095906 | 0.128022 | 2.940827 | 0.004843 | 0.039587 | -2.10896 | Methyl-coenzyme M reductase subunit beta | elongation zone | 12 h |
| HORVU.MOREX.r3.4HG0392940.1 | 0.520224 | 5.529744 | 3.123413 | 0.002896 | 0.028327 | -2.109 | Kinase family protein | elongation zone | 12 h |
| HORVU.MOREX.r3.3HG0274100.1 | -1.26865 | 1.731697 | -3.00287 | 0.004074 | 0.035421 | -2.1092 | 50S ribosomal protein L31 | elongation zone | 12 h |
| HORVU.MOREX.r3.3HG0280970.1 | -0.41452 | 5.971206 | -3.09662 | 0.003126 | 0.029864 | -2.10954 | Protein methyltransferase | elongation zone | 12 h |
| HORVU.MOREX.r3.6HG0624580.1 | -1.1248 | 4.006778 | -2.93989 | 0.004856 | 0.039601 | -2.11073 | Translation initiation factor IF-2, putative isoform 2 | elongation zone | 12 h |
| HORVU.MOREX.r3.1HG0021830.1 | -0.33933 | 5.177385 | -3.04236 | 0.003646 | 0.032938 | -2.11081 | SKP1-like protein 4 | elongation zone | 12 h |
| HORVU.MOREX.r3.6HG0620200.1 | 1.189975 | -0.686 | 2.942733 | 0.004817 | 0.039433 | -2.11136 | Kinase family protein | elongation zone | 12 h |
| HORVU.MOREX.r3.1HG0091860.1 | -0.32371 | 7.126684 | -3.18645 | 0.002415 | 0.025092 | -2.11234 | Sugar transporter protein | elongation zone | 12 h |
| HORVU.MOREX.r3.4HG0407410.1 | -0.94514 | -0.65632 | -2.93313 | 0.004947 | 0.040109 | -2.11621 | Glycosyltransferase | elongation zone | 12 h |

**Table S1** Continued.

| ID | logFC | AveExpr | t | P.Value | adj.P.Val | B | gene description | root zone | time point |
|---|---|---|---|---|---|---|---|---|---|
| HORVU.MOREX.r3.2HG0160760.1 | 0.665852 | 3.8459 | 3.045811 | 0.003611 | 0.032683 | -2.11746 | L-threonine 3-dehydrogenase | elongation zone | 12 h |
| HORVU.MOREX.r3.1HG0061220.1 | -1.13579 | 4.835261 | -2.95265 | 0.004687 | 0.038809 | -2.11769 | DNA primase | elongation zone | 12 h |
| HORVU.MOREX.r3.3HG0306220.1 | -0.47163 | 4.083588 | -2.99188 | 0.004202 | 0.036195 | -2.11789 | Transmembrane protein 62 | elongation zone | 12 h |
| HORVU.MOREX.r3.2HG0193080.1 | -0.44137 | 3.176199 | -2.97359 | 0.004422 | 0.037375 | -2.11861 | Protein-S-isoprenylcysteine O-methyltransferase | elongation zone | 12 h |
| HORVU.MOREX.r3.1HG0041250.1 | -0.37019 | 4.039793 | -2.99989 | 0.004109 | 0.035609 | -2.12034 | Peroxisome biogenesis protein 12 | elongation zone | 12 h |
| HORVU.MOREX.r3.1HG0059340.1 | -0.28908 | 6.875668 | -3.15506 | 0.002644 | 0.02661 | -2.12046 | Conserved oligomeric Golgi complex subunit 5 | elongation zone | 12 h |
| HORVU.MOREX.r3.3HG0236690.1 | 0.668023 | 1.038969 | 3.043804 | 0.003631 | 0.03284 | -2.12458 | Myb transcription factor | elongation zone | 12 h |
| HORVU.MOREX.r3.1HG0088870.1 | -0.26953 | 5.735596 | -3.07935 | 0.003283 | 0.030822 | -2.12514 | ABC transporter ATP-binding protein ARB1 | elongation zone | 12 h |
| HORVU.MOREX.r3.2HG0173380.1 | 0.415622 | 4.283255 | 3.083436 | 0.003246 | 0.030621 | -2.12539 | Amino acid transporter, putative | elongation zone | 12 h |
| HORVU.MOREX.r3.1HG0055580.1 | -0.41079 | 5.226651 | -3.0511 | 0.003557 | 0.032357 | -2.12641 | Kinesin-like protein | elongation zone | 12 h |
| HORVU.MOREX.r3.5HG0516470.1 | -1.40758 | 4.720536 | -2.92977 | 0.004993 | 0.040388 | -2.12697 | GDSL esterase/lipase | elongation zone | 12 h |
| HORVU.MOREX.r3.5HG0444860.1 | -1.36506 | 2.653302 | -2.92727 | 0.005028 | 0.040572 | -2.12748 | Kinesin-related protein | elongation zone | 12 h |
| HORVU.MOREX.r3.3HG0298340.1 | -0.73899 | 4.243389 | -2.96005 | 0.004592 | 0.038331 | -2.1291 | Beta-galactosidase | elongation zone | 12 h |
| HORVU.MOREX.r3.6HG0631050.1 | -1.12145 | 6.885939 | -3.06055 | 0.003463 | 0.031884 | -2.12957 | Histone H2A | elongation zone | 12 h |
| HORVU.MOREX.r3.5HG0520670.1 | -1.38496 | -0.02366 | -2.9785 | 0.004362 | 0.037137 | -2.13022 | beta glucosidase 41 | elongation zone | 12 h |
| HORVU.MOREX.r3.4HG0409880.1 | 1.419768 | -2.34755 | 2.968916 | 0.00448 | 0.03765 | -2.13266 | Metal tolerance protein | elongation zone | 12 h |
| HORVU.MOREX.r3.1HG0046210.1 | 2.264067 | -0.44242 | 3.03006 | 0.003775 | 0.03368 | -2.13293 | Receptor kinase | elongation zone | 12 h |
| HORVU.MOREX.r3.5HG0513900.1 | 0.785464 | -0.25185 | 2.928667 | 0.005009 | 0.040463 | -2.13311 | Leucine carboxyl methyltransferase | elongation zone | 12 h |
| HORVU.MOREX.r3.3HG0293570.1 | -1.60574 | -0.38794 | -3.02074 | 0.003875 | 0.03426 | -2.1348 | BTB/POZ domain-containing protein | elongation zone | 12 h |
| HORVU.MOREX.r3.2HG0128240.1 | -0.38362 | 5.300426 | -3.06193 | 0.00345 | 0.031816 | -2.13532 | Aldose 1-epimerase | elongation zone | 12 h |
| HORVU.MOREX.r3.2HG0122500.2 | -1.83143 | -0.50803 | -3.14198 | 0.002745 | 0.027347 | -2.13745 | GTPase obg | elongation zone | 12 h |
| HORVU.MOREX.r3.5HG0426110.1 | 1.661491 | -3.22479 | 3.027572 | 0.003801 | 0.033827 | -2.13805 | NBS-LRR class disease resistance protein | elongation zone | 12 h |
| HORVU.MOREX.r3.6HG0560140.1 | 1.980317 | -1.9167 | 2.940566 | 0.004846 | 0.03959 | -2.13886 | Endoglucanase | elongation zone | 12 h |
| HORVU.MOREX.r3.3HG0307950.1 | -0.2717 | 6.122502 | -3.10986 | 0.00301 | 0.02903 | -2.14091 | GPI transamidase component PIG-S | elongation zone | 12 h |
| HORVU.MOREX.r3.7HG0676230.1 | -0.86943 | 0.616952 | -2.92505 | 0.005059 | 0.040741 | -2.14137 | Zinc finger C-x8-C-x5-C-x3-H type family protein | elongation zone | 12 h |
| HORVU.MOREX.r3.5HG0501500.1 | 1.126821 | 3.165714 | 2.991273 | 0.004209 | 0.036196 | -2.14205 | Glucan endo-1,3-beta-glucosidase 3 | elongation zone | 12 h |
| HORVU.MOREX.r3.7HG0674600.1 | 1.115929 | 1.082795 | 2.949616 | 0.004726 | 0.03901 | -2.1431 | Aconitate hydratase | elongation zone | 12 h |
| HORVU.MOREX.r3.2HG0110920.1 | -1.88635 | 1.02517 | -2.99268 | 0.004192 | 0.036159 | -2.14538 | VQ motif-containing family protein | elongation zone | 12 h |
| HORVU.MOREX.r3.4HG0388780.1 | 1.83723 | 0.089382 | 3.177136 | 0.002481 | 0.025544 | -2.14542 | RNA polymerase sigma factor | elongation zone | 12 h |
| HORVU.MOREX.r3.7HG0656340.1 | -1.97221 | -0.36508 | -3.11022 | 0.003007 | 0.029025 | -2.1456 | Adhesion G protein-coupled receptor L3 | elongation zone | 12 h |
| HORVU.MOREX.r3.6HG0609830.1 | 0.661773 | 4.638207 | 3.168655 | 0.002543 | 0.025982 | -2.14586 | Pollen-specific arabinogalacta protein BAN102 | elongation zone | 12 h |
| HORVU.MOREX.r3.7HG0697980.1 | -0.93761 | 0.647065 | -2.93945 | 0.004861 | 0.039613 | -2.15486 | Fatty acid hydroxylase superfamily | elongation zone | 12 h |
| HORVU.MOREX.r3.4HG0337500.1 | -0.18104 | 9.290144 | -3.22861 | 0.002137 | 0.023408 | -2.15718 | NADH-ubiquinone oxidoreductase subunit | elongation zone | 12 h |
| HORVU.MOREX.r3.1HG0055500.1 | 1.013156 | 2.091951 | 3.120162 | 0.002923 | 0.028505 | -2.15753 | Pollen Ole e 1 allergen and extensin family protein | elongation zone | 12 h |
| HORVU.MOREX.r3.6HG0620040.1 | -1.29021 | 1.386644 | -2.93496 | 0.004922 | 0.040007 | -2.16053 | Protein kinase family protein | elongation zone | 12 h |
| HORVU.MOREX.r3.7HG0714950.1 | -0.59358 | 3.547437 | -2.95277 | 0.004685 | 0.038809 | -2.16152 | Cytokinin riboside 5'-monophosphate phosphoribohydrolase | elongation zone | 12 h |
| HORVU.MOREX.r3.2HG0122880.1 | 1.329168 | 1.048128 | 2.912038 | 0.005244 | 0.041806 | -2.1619 | Phytoene desaturase | elongation zone | 12 h |
| HORVU.MOREX.r3.2HG0159920.1 | -0.56459 | 3.687678 | -2.9461 | 0.004773 | 0.039205 | -2.16382 | receptor-like kinase in flowers 1 | elongation zone | 12 h |
| HORVU.MOREX.r3.4HG0341810.1 | -0.31501 | 7.757023 | -3.14283 | 0.002739 | 0.027326 | -2.16492 | Cycloeucalenol cycloisomerase | elongation zone | 12 h |
| HORVU.MOREX.r3.5HG0426480.1 | -1.1734 | 2.819296 | -2.91225 | 0.005241 | 0.041806 | -2.16493 | 1-deoxy-D-xylulose 5-phosphate synthase 1 | elongation zone | 12 h |
| HORVU.MOREX.r3.1HG0058800.1 | -0.53823 | 3.603853 | -2.94377 | 0.004804 | 0.039373 | -2.16522 | Glycosyltransferase | elongation zone | 12 h |

**Table S1** Continued.

| ID | logFC | AveExpr | t | P.Value | adj.P.Val | B | gene description | root zone | time point |
|---|---|---|---|---|---|---|---|---|---|
| HORVU.MOREX.r3.7HG0667680.1 | 1.009409 | 4.54919 | 3.09828 | 0.003111 | 0.029766 | -2.16721 | Peroxidase | elongation zone | 12 h |
| HORVU.MOREX.r3.3HG0299300.1 | -1.10794 | 0.206257 | -2.9694 | 0.004474 | 0.037622 | -2.16836 | #N/A | elongation zone | 12 h |
| HORVU.MOREX.r3.5HG0434380.1 | -0.3843 | 4.006199 | -2.99286 | 0.00419 | 0.036157 | -2.16892 | Receptor-like kinase | elongation zone | 12 h |
| HORVU.MOREX.r3.6HG0616450.1 | 1.129299 | 0.245809 | 2.968129 | 0.004489 | 0.037709 | -2.17079 | Na(+)/H(+) exchange regulatory cofactor NHE-RF3 | elongation zone | 12 h |
| HORVU.MOREX.r3.7HG0657330.1 | 0.426851 | 6.822879 | 3.22414 | 0.002165 | 0.023553 | -2.17354 | ENTH/ANTH/VHS superfamily protein | elongation zone | 12 h |
| HORVU.MOREX.r3.7HG0656420.1 | -1.06389 | 0.082394 | -2.9437 | 0.004805 | 0.039373 | -2.17518 | SWI/SNF complex subunit SMARCC2 | elongation zone | 12 h |
| HORVU.MOREX.r3.6HG0607000.1 | -1.73278 | 0.642043 | -3.17377 | 0.002505 | 0.02572 | -2.17601 | Pectin acetylesterase | elongation zone | 12 h |
| HORVU.MOREX.r3.3HG0219740.1 | -0.61527 | 6.408651 | -3.05647 | 0.003504 | 0.032039 | -2.17849 | 50S ribosomal protein L2, putative | elongation zone | 12 h |
| HORVU.MOREX.r3.7HG0744260.1 | -2.98097 | 0.783361 | -3.20564 | 0.002284 | 0.0242 | -2.18088 | NHL domain-containing protein, putative | elongation zone | 12 h |
| HORVU.MOREX.r3.5HG0500220.1 | -0.31621 | 6.295637 | -3.09065 | 0.00318 | 0.030179 | -2.18135 | Protein PAT1 1 | elongation zone | 12 h |
| HORVU.MOREX.r3.2HG0158020.1 | -0.41197 | 4.300593 | -2.97683 | 0.004382 | 0.037198 | -2.18254 | BTB-POZ and MATH domain protein | elongation zone | 12 h |
| HORVU.MOREX.r3.1HG0018430.1 | -1.0677 | 3.282493 | -2.90264 | 0.005381 | 0.042386 | -2.18361 | Histone H3 | elongation zone | 12 h |
| HORVU.MOREX.r3.7HG0711270.1 | 0.807358 | 3.15201 | 3.186582 | 0.002414 | 0.025092 | -2.18431 | Xyloglucan endotransglucosylase/hydrolase | elongation zone | 12 h |
| HORVU.MOREX.r3.2HG0190800.1 | 1.708104 | 0.966626 | 3.049107 | 0.003577 | 0.032459 | -2.18469 | F-box protein | elongation zone | 12 h |
| HORVU.MOREX.r3.4HG0395770.1 | -0.74065 | 1.202249 | -2.90392 | 0.005362 | 0.042352 | -2.18617 | Hsp70 nucleotide exchange factor fes1 | elongation zone | 12 h |
| HORVU.MOREX.r3.6HG0627090.1 | 0.662526 | 1.38286 | 2.91679 | 0.005175 | 0.041436 | -2.18619 | Alcohol dehydrogenase 3 | elongation zone | 12 h |
| HORVU.MOREX.r3.7HG0727350.1 | -0.40269 | 10.95622 | -3.24775 | 0.00202 | 0.022545 | -2.18675 | #N/A | elongation zone | 12 h |
| HORVU.MOREX.r3.3HG0316280.1 | -0.85384 | 4.501603 | -2.94017 | 0.004852 | 0.039601 | -2.18967 | Transcription initiation factor TFIID subunit 9 | elongation zone | 12 h |
| HORVU.MOREX.r3.3HG0327100.1 | 1.167641 | -3.0577 | 2.933764 | 0.004939 | 0.040086 | -2.19108 | #N/A | elongation zone | 12 h |
| HORVU.MOREX.r3.5HG0512510.1 | -0.60982 | 7.202032 | -3.11074 | 0.003003 | 0.029025 | -2.19166 | Subtilisin-like protease | elongation zone | 12 h |
| HORVU.MOREX.r3.6HG0546200.1 | -0.55843 | 4.682471 | -2.97816 | 0.004366 | 0.037141 | -2.19229 | DNA-directed RNA polymerase II subunit rpb4 | elongation zone | 12 h |
| HORVU.MOREX.r3.1HG0062630.1 | 0.466177 | 5.379794 | 3.034513 | 0.003728 | 0.033447 | -2.19356 | #N/A | elongation zone | 12 h |
| HORVU.MOREX.r3.6HG0543540.1 | 1.35374 | -0.02472 | 2.958892 | 0.004606 | 0.038423 | -2.19473 | Expansin protein | elongation zone | 12 h |
| HORVU.MOREX.r3.3HG0327630.1 | -1.15984 | 4.246104 | -2.90491 | 0.005347 | 0.042285 | -2.1961 | Cyclin family protein | elongation zone | 12 h |
| HORVU.MOREX.r3.3HG0296990.1 | 0.847045 | -1.49078 | 2.906294 | 0.005327 | 0.042206 | -2.1969 | LURP-one-like protein | elongation zone | 12 h |
| HORVU.MOREX.r3.6HG0605960.1 | 0.687148 | 3.198592 | 2.996644 | 0.004146 | 0.035879 | -2.19846 | Lysine-specific demethylase | elongation zone | 12 h |
| HORVU.MOREX.r3.4HG0341080.1 | -1.81 | 4.721757 | -2.89833 | 0.005445 | 0.042719 | -2.19939 | interferon-activable protein | elongation zone | 12 h |
| HORVU.MOREX.r3.2HG0176680.1 | -0.23745 | 6.632089 | -3.0902 | 0.003184 | 0.030203 | -2.20084 | Serine/threonine protein phosphatase 2A regulatory subunit B | elongation zone | 12 h |
| HORVU.MOREX.r3.6HG0618110.1 | -0.70438 | 3.25054 | -2.97672 | 0.004383 | 0.037198 | -2.20279 | growth-regulating factor 9 | elongation zone | 12 h |
| HORVU.MOREX.r3.1HG0070110.1 | 0.347249 | 5.55369 | 3.087594 | 0.003207 | 0.030372 | -2.20454 | Type I inositol-1,4,5-trisphosphate 5-phosphatase | elongation zone | 12 h |
| HORVU.MOREX.r3.1HG0056990.1 | 0.457771 | 3.797448 | 3.019746 | 0.003886 | 0.03428 | -2.20571 | Camphor resistance CrcB family protein | elongation zone | 12 h |
| HORVU.MOREX.r3.4HG0409600.1 | 0.487457 | 4.470889 | 3.096104 | 0.003131 | 0.029866 | -2.20833 | Protein kinase family protein, putative, expressed | elongation zone | 12 h |
| HORVU.MOREX.r3.3HG0278710.1 | 0.660082 | 6.634206 | 3.254345 | 0.001982 | 0.022356 | -2.21108 | Fasciclin-like arabinogalactan protein | elongation zone | 12 h |
| HORVU.MOREX.r3.1HG0018110.1 | 0.374711 | 8.069235 | 3.20901 | 0.002262 | 0.02416 | -2.2121 | ABC transporter B family protein | elongation zone | 12 h |
| HORVU.MOREX.r3.6HG0565800.1 | 0.770302 | 0.673146 | 2.925295 | 0.005055 | 0.040734 | -2.21338 | Peroxidase | elongation zone | 12 h |
| HORVU.MOREX.r3.1HG0011820.1 | 0.924023 | -1.28428 | 2.931672 | 0.004967 | 0.040223 | -2.21464 | Protein ABIL1 | elongation zone | 12 h |
| HORVU.MOREX.r3.3HG0293700.1 | -0.66041 | 6.606906 | -3.04098 | 0.00366 | 0.03303 | -2.21532 | Protein downstream neighbor of Son | elongation zone | 12 h |
| HORVU.MOREX.r3.3HG0294100.1 | -0.3761 | 4.223869 | -3.03025 | 0.003773 | 0.03368 | -2.21643 | NAD(P)H dehydrogenase (Quinone) | elongation zone | 12 h |
| HORVU.MOREX.r3.2HG0101220.2 | -0.49806 | 2.581199 | -2.92005 | 0.005129 | 0.041178 | -2.21667 | Formin-like protein | elongation zone | 12 h |
| HORVU.MOREX.r3.7HG0688770.1 | -0.68814 | 3.39936 | -2.91467 | 0.005206 | 0.041597 | -2.2176 | #N/A | elongation zone | 12 h |
| HORVU.MOREX.r3.6HG0567780.1 | -0.50974 | 3.016753 | -2.92241 | 0.005096 | 0.041023 | -2.2189 | disease resistance protein (TIR-NBS-LRR class) | elongation zone | 12 h |

**Table S1** Continued.

| ID | logFC | AveExpr | t | P.Value | adj.P.Val | B | gene description | root zone | time point |
|---|---|---|---|---|---|---|---|---|---|
| HORVU.MOREX.r3.2HG0159500.1 | -0.75882 | 5.303207 | -2.96293 | 0.004555 | 0.038087 | -2.21906 | tRNA (Adenine-N(1)-)-methyltransferase catalytic subunit trmt61a | elongation zone | 12 h |
| HORVU.MOREX.r3.5HG0476390.1 | -0.85495 | 1.081391 | -2.88765 | 0.005607 | 0.04358 | -2.21941 | Transporter-related family protein | elongation zone | 12 h |
| HORVU.MOREX.r3.4HG0332710.1 | -2.92507 | -2.51339 | -3.27302 | 0.001876 | 0.021568 | -2.21966 | DNA mismatch repair protein MutL | elongation zone | 12 h |
| HORVU.MOREX.r3.3HG0294110.1 | 0.927177 | -1.24039 | 2.885214 | 0.005644 | 0.043691 | -2.22067 | #N/A | elongation zone | 12 h |
| HORVU.MOREX.r3.4HG0375440.1 | -1.16763 | 2.464559 | -2.90668 | 0.005322 | 0.042178 | -2.22113 | ATP-dependent Clp protease proteolytic subunit | elongation zone | 12 h |
| HORVU.MOREX.r3.3HG0239110.1 | -0.92548 | 4.789763 | -3.05077 | 0.003561 | 0.032357 | -2.22151 | Glycosyltransferase | elongation zone | 12 h |
| HORVU.MOREX.r3.6HG0611290.1 | -0.8512 | 7.158294 | -3.02463 | 0.003833 | 0.034006 | -2.22329 | Acetolactate synthase | elongation zone | 12 h |
| HORVU.MOREX.r3.6HG0550670.1 | -1.81029 | 7.005258 | -2.99238 | 0.004196 | 0.036159 | -2.22545 | Glycine-rich cell wall structural protein 2 | elongation zone | 12 h |
| HORVU.MOREX.r3.7HG0673710.1 | 1.43696 | -0.09914 | 2.89192 | 0.005542 | 0.043254 | -2.2257 | Receptor-like protein kinase | elongation zone | 12 h |
| HORVU.MOREX.r3.2HG0125780.1 | -1.2602 | 0.405475 | -2.95685 | 0.004632 | 0.038594 | -2.22594 | #N/A | elongation zone | 12 h |
| HORVU.MOREX.r3.7HG0749100.1 | 0.92186 | -1.03535 | 2.948325 | 0.004743 | 0.03909 | -2.22818 | arginine/serine-rich splicing factor 35 | elongation zone | 12 h |
| HORVU.MOREX.r3.3HG0319570.1 | -1.15683 | 3.275641 | -2.8859 | 0.005634 | 0.043682 | -2.22826 | Crossover junction endonuclease mus81 | elongation zone | 12 h |
| HORVU.MOREX.r3.3HG0280920.1 | -0.62641 | 3.219452 | -3.04127 | 0.003657 | 0.033017 | -2.2287 | DUF4228 domain protein | elongation zone | 12 h |
| HORVU.MOREX.r3.7HG0730530.1 | 1.009407 | 0.190357 | 2.942306 | 0.004823 | 0.039462 | -2.22973 | Copper transport protein family | elongation zone | 12 h |
| HORVU.MOREX.r3.1HG0037920.1 | -1.92316 | 0.68135 | -2.98252 | 0.004313 | 0.036879 | -2.23033 | Protein restricted tev movement 2 | elongation zone | 12 h |
| HORVU.MOREX.r3.4HG0395120.1 | -0.63162 | 2.404038 | -2.88534 | 0.005643 | 0.043691 | -2.23084 | Mitochondrial carrier family | elongation zone | 12 h |
| HORVU.MOREX.r3.3HG0234030.1 | 0.838639 | 0.965447 | 3.059054 | 0.003478 | 0.031944 | -2.23106 | Xyloglucan galactosyltransferase KATAMARI1-like protein | elongation zone | 12 h |
| HORVU.MOREX.r3.2HG0167620.1 | -1.25165 | 2.679481 | -2.89101 | 0.005556 | 0.043297 | -2.2313 | Plastid-lipid associated protein PAP / fibrillin family protein | elongation zone | 12 h |
| HORVU.MOREX.r3.7HG0686830.1 | -0.82458 | 2.110337 | -2.87968 | 0.005731 | 0.044174 | -2.23279 | TraB family protein | elongation zone | 12 h |
| HORVU.MOREX.r3.6HG0608800.1 | -0.56853 | 4.150298 | -2.91858 | 0.00515 | 0.041296 | -2.23495 | Pentatricopeptide repeat-containing protein | elongation zone | 12 h |
| HORVU.MOREX.r3.4HG0345260.1 | 1.209781 | -1.43865 | 2.878385 | 0.005751 | 0.044248 | -2.23636 | Phosphatidylinositol 3,4,5-trisphosphate 5-phosphatase 1 | elongation zone | 12 h |
| HORVU.MOREX.r3.4HG0333840.1 | -0.80044 | 1.907765 | -2.87906 | 0.00574 | 0.044183 | -2.23653 | Histone-lysine N-methyltransferase, H3 lysine-9 specific SUVH5 | elongation zone | 12 h |
| HORVU.MOREX.r3.2HG0203330.1 | 0.432961 | 5.510492 | 3.0629 | 0.00344 | 0.031786 | -2.23694 | Sec14p-like phosphatidylinositol transfer family protein | elongation zone | 12 h |
| HORVU.MOREX.r3.1HG0003150.1 | -0.62639 | 4.554906 | -2.92156 | 0.005108 | 0.041076 | -2.23752 | Chaperone protein DnaJ | elongation zone | 12 h |
| HORVU.MOREX.r3.4HG0344790.1 | 0.453177 | 4.360827 | 2.999819 | 0.004109 | 0.035609 | -2.23801 | Non-specific serine/threonine protein kinase | elongation zone | 12 h |
| HORVU.MOREX.r3.6HG0594420.1 | -0.39558 | 3.225218 | -2.91799 | 0.005158 | 0.041348 | -2.24071 | Transporter-related family protein | elongation zone | 12 h |
| HORVU.MOREX.r3.5HG0514020.1 | 0.441112 | 3.72423 | 2.968843 | 0.004481 | 0.03765 | -2.2424 | GRAS transcription factor | elongation zone | 12 h |
| HORVU.MOREX.r3.6HG0601560.1 | -1.1885 | 0.277948 | -2.92919 | 0.005001 | 0.040437 | -2.24279 | Pentatricopeptide repeat-containing protein | elongation zone | 12 h |
| HORVU.MOREX.r3.7HG0701080.1 | 0.875563 | -1.51889 | 2.882938 | 0.00568 | 0.043865 | -2.24322 | MYB transcription factor | elongation zone | 12 h |
| HORVU.MOREX.r3.2HG0160790.1 | -0.52785 | 4.130952 | -2.93017 | 0.004988 | 0.040374 | -2.24325 | Alcohol dehydrogenase 3 | elongation zone | 12 h |
| HORVU.MOREX.r3.7HG0679730.1 | -1.40074 | -0.42256 | -2.93893 | 0.004868 | 0.039627 | -2.24333 | AP-1 complex subunit gamma-2 | elongation zone | 12 h |
| HORVU.MOREX.r3.3HG0296940.1 | -0.77807 | 2.177147 | -2.87404 | 0.00582 | 0.04466 | -2.24494 | Calcium uniporter, mitochondrial | elongation zone | 12 h |
| HORVU.MOREX.r3.4HG0395130.1 | -1.68299 | -0.45189 | -3.11682 | 0.002951 | 0.028675 | -2.24535 | Cyclin, putative | elongation zone | 12 h |
| HORVU.MOREX.r3.4HG0345810.1 | -1.26043 | 1.299714 | -2.9212 | 0.005113 | 0.041095 | -2.24656 | #N/A | elongation zone | 12 h |
| HORVU.MOREX.r3.2HG0140380.1 | 1.766269 | -1.74806 | 2.90582 | 0.005334 | 0.042229 | -2.24677 | CTD small phosphatase-like protein 2, putative | elongation zone | 12 h |
| HORVU.MOREX.r3.7HG0669750.1 | 0.408166 | 4.895319 | 3.081726 | 0.003261 | 0.030701 | -2.24758 | Sn1-specific diacylglycerol lipase alpha | elongation zone | 12 h |
| HORVU.MOREX.r3.2HG0158180.1 | 0.93866 | 3.62095 | 2.953343 | 0.004678 | 0.038805 | -2.24971 | Proline transporter | elongation zone | 12 h |
| HORVU.MOREX.r3.2HG0096290.1 | 0.608155 | 1.613406 | 2.952847 | 0.004684 | 0.038809 | -2.25141 | Late embryogenesis abundant (LEA) hydroxyproline-rich glycoprotein family meml | elongation zone | 12 h |
| HORVU.MOREX.r3.3HG0326430.1 | -0.27487 | 5.095188 | -3.02093 | 0.003873 | 0.03426 | -2.25278 | Diphosphomevalonate decarboxylase | elongation zone | 12 h |
| HORVU.MOREX.r3.2HG0189010.1 | 0.945507 | 3.632309 | 2.952995 | 0.004682 | 0.038809 | -2.2528 | Glycerol-3-phosphate transporter, putative | elongation zone | 12 h |
| HORVU.MOREX.r3.1HG0004290.1 | -1.05573 | 6.306577 | -3.00847 | 0.004011 | 0.035058 | -2.25383 | Chymotrypsin inhibitor | elongation zone | 12 h |

**Table S1** Continued.

| ID | logFC | AveExpr | t | P.Value | adj.P.Val | B | gene description | root zone | time point |
|---|---|---|---|---|---|---|---|---|---|
| HORVU.MOREX.r3.4HG0375550.1 | -2.26771 | 0.494963 | -3.12907 | 0.002849 | 0.028054 | -2.25622 | Rapid ALkalinization Factor family protein | elongation zone | 12 h |
| HORVU.MOREX.r3.6HG0632980.1 | -1.5627 | 5.615728 | -2.91436 | 0.00521 | 0.041617 | -2.25884 | Replication protein A 32 kDa subunit | elongation zone | 12 h |
| HORVU.MOREX.r3.6HG0568870.1 | -0.79519 | 4.739053 | -2.94756 | 0.004753 | 0.039139 | -2.2596 | Histone H2A | elongation zone | 12 h |
| HORVU.MOREX.r3.4HG0407310.1 | 0.720376 | 3.764737 | 3.029891 | 0.003777 | 0.03368 | -2.26017 | Xyloglucan galactosyltransferase KATAMARI1-like protein | elongation zone | 12 h |
| HORVU.MOREX.r3.1HG0088530.1 | 0.783643 | 0.052643 | 2.872872 | 0.005838 | 0.044752 | -2.26111 | Cytochrome b561 domain-containing protein | elongation zone | 12 h |
| HORVU.MOREX.r3.4HG0342680.1 | 0.947958 | -0.60974 | 2.957978 | 0.004618 | 0.038489 | -2.26301 | Branchpoint-bridging protein | elongation zone | 12 h |
| HORVU.MOREX.r3.3HG0228010.1 | -0.39623 | 4.500471 | -2.96497 | 0.004529 | 0.037933 | -2.26545 | EKC/KEOPS complex subunit CGI121 | elongation zone | 12 h |
| HORVU.MOREX.r3.6HG0618170.1 | -0.25586 | 6.062292 | -3.03503 | 0.003722 | 0.033412 | -2.26558 | 1-aminocyclopropane-1-carboxylate deaminase/D-cysteine desulfhydrase | elongation zone | 12 h |
| HORVU.MOREX.r3.4HG0409670.1 | 1.171815 | 1.37934 | 2.970553 | 0.004459 | 0.037548 | -2.26702 | Respiratory burst oxidase homolog | elongation zone | 12 h |
| HORVU.MOREX.r3.2HG0163090.1 | -0.83342 | 7.015883 | -3.01481 | 0.00394 | 0.034626 | -2.26805 | Heavy metal-associated protein | elongation zone | 12 h |
| HORVU.MOREX.r3.6HG0586570.1 | -0.86434 | 0.923324 | -2.88752 | 0.005609 | 0.04358 | -2.26882 | Transposon Ty3-G Gag-Pol polyprotein | elongation zone | 12 h |
| HORVU.MOREX.r3.3HG0225640.1 | -1.29815 | 1.458957 | -2.92075 | 0.005119 | 0.041131 | -2.26984 | Peptidyl-tRNA hydrolase family protein | elongation zone | 12 h |
| HORVU.MOREX.r3.6HG0543400.1 | 1.263887 | -0.34964 | 2.961449 | 0.004574 | 0.038197 | -2.27088 | High affinity nitrate transporter | elongation zone | 12 h |
| HORVU.MOREX.r3.5HG0491230.1 | -0.92335 | 2.284228 | -2.86224 | 0.00601 | 0.045661 | -2.27102 | Dihydroflavonol-4-reductase | elongation zone | 12 h |
| HORVU.MOREX.r3.1HG0073160.1 | 0.42732 | 5.05488 | 3.058927 | 0.003479 | 0.031944 | -2.27177 | Histone H2A | elongation zone | 12 h |
| HORVU.MOREX.r3.1HG0010920.1 | 0.564321 | 2.101242 | 2.972919 | 0.00443 | 0.037408 | -2.27243 | Protein aluminum sensitive 3 | elongation zone | 12 h |
| HORVU.MOREX.r3.6HG0619540.1 | -0.35122 | 5.673476 | -2.99152 | 0.004206 | 0.036196 | -2.27257 | THO complex subunit 5 | elongation zone | 12 h |
| HORVU.MOREX.r3.6HG0556240.1 | 0.455526 | 4.26639 | 2.987018 | 0.004259 | 0.036533 | -2.27359 | #N/A | elongation zone | 12 h |
| HORVU.MOREX.r3.3HG0289070.1 | -1.20542 | 2.310764 | -2.90585 | 0.005334 | 0.042229 | -2.27416 | Protein TIC 22-like, chloroplastic | elongation zone | 12 h |
| HORVU.MOREX.r3.2HG0098420.1 | -1.14149 | 5.161437 | -2.9029 | 0.005377 | 0.042386 | -2.27687 | CDT1-like protein a, chloroplastic | elongation zone | 12 h |
| HORVU.MOREX.r3.1HG0043350.1 | -0.49482 | 3.574077 | -2.92851 | 0.005011 | 0.040464 | -2.27758 | Transmembrane protein, putative | elongation zone | 12 h |
| HORVU.MOREX.r3.3HG0310460.1 | -0.54787 | 0.959034 | -2.98394 | 0.004296 | 0.036791 | -2.27929 | Glycosyltransferase | elongation zone | 12 h |
| HORVU.MOREX.r3.7HG0662830.1 | -0.92379 | 4.730198 | -2.88582 | 0.005635 | 0.043682 | -2.27939 | Mechanosensitive ion channel family protein | elongation zone | 12 h |
| HORVU.MOREX.r3.5HG0495490.1 | -0.42215 | 4.866247 | -2.99107 | 0.004211 | 0.036201 | -2.27964 | CCR4-NOT transcription complex subunit 2 | elongation zone | 12 h |
| HORVU.MOREX.r3.5HG0446420.1 | 0.765283 | 4.215118 | 3.155351 | 0.002642 | 0.026601 | -2.27982 | GDSL esterase/lipase | elongation zone | 12 h |
| HORVU.MOREX.r3.7HG0675770.1 | -0.45552 | 4.417254 | -3.02885 | 0.003788 | 0.033749 | -2.27996 | basic helix-loop-helix (bHLH) DNA-binding superfamily protein | elongation zone | 12 h |
| HORVU.MOREX.r3.6HG0607590.1 | 0.937508 | 1.682477 | 2.867464 | 0.005925 | 0.045249 | -2.28 | U-box domain-containing family protein | elongation zone | 12 h |
| HORVU.MOREX.r3.5HG0535780.1 | -0.71126 | 6.219129 | -2.99167 | 0.004204 | 0.036196 | -2.28069 | Transmembrane protein 70, mitochondrial | elongation zone | 12 h |
| HORVU.MOREX.r3.2HG0212990.1 | -0.66075 | 5.148472 | -2.91159 | 0.00525 | 0.041806 | -2.28074 | Acyl-protein thioesterase 1 | elongation zone | 12 h |
| HORVU.MOREX.r3.6HG0552720.1 | -0.41058 | 3.733307 | -2.92037 | 0.005125 | 0.041158 | -2.28244 | Beta-amylase | elongation zone | 12 h |
| HORVU.MOREX.r3.2HG0154740.1 | -0.78356 | 1.125076 | -2.85799 | 0.00608 | 0.045953 | -2.28321 | cysteine-rich RLK (RECEPTOR-like protein kinase) 11 | elongation zone | 12 h |
| HORVU.MOREX.r3.4HG0406380.1 | -0.38527 | 5.163919 | -2.9752 | 0.004402 | 0.037294 | -2.28393 | GEM-like protein 1 | elongation zone | 12 h |
| HORVU.MOREX.r3.6HG0544350.1 | 0.423123 | 5.04997 | 3.019794 | 0.003885 | 0.03428 | -2.2846 | Carboxypeptidase | elongation zone | 12 h |
| HORVU.MOREX.r3.3HG0235320.1 | 1.506909 | -0.14202 | 2.891324 | 0.005551 | 0.043292 | -2.2854 | Pollen Ole e 1 allergen and extensin family protein | elongation zone | 12 h |
| HORVU.MOREX.r3.2HG0139050.1 | -1.00938 | 3.432928 | -2.86438 | 0.005975 | 0.045496 | -2.28621 | Protein kinase | elongation zone | 12 h |
| HORVU.MOREX.r3.3HG0306820.2 | -1.32584 | 0.136032 | -2.88881 | 0.005589 | 0.043492 | -2.28721 | ATP-dependent Clp protease ATP-binding subunit ClpX | elongation zone | 12 h |
| HORVU.MOREX.r3.6HG0621170.1 | 1.754667 | -0.64178 | 2.854987 | 0.00613 | 0.046179 | -2.28721 | GPI-anchored adhesin-like protein | elongation zone | 12 h |
| HORVU.MOREX.r3.5HG0437610.1 | -0.82768 | 5.573525 | -2.94937 | 0.00473 | 0.03901 | -2.28908 | caspase-6 protein | elongation zone | 12 h |
| HORVU.MOREX.r3.5HG0501270.1 | -1.40767 | 6.400081 | -2.93344 | 0.004943 | 0.04009 | -2.29121 | Chromatin assembly factor-1 | elongation zone | 12 h |
| HORVU.MOREX.r3.6HG0569490.1 | -0.44483 | 6.2556 | -3.00429 | 0.004058 | 0.035331 | -2.29166 | Rad23 UV excision repair protein family | elongation zone | 12 h |
| HORVU.MOREX.r3.6HG0548460.1 | -1.10854 | 4.93645 | -2.91491 | 0.005202 | 0.041594 | -2.29204 | Histone H4 | elongation zone | 12 h |

**Table S1** Continued.

| ID | logFC | AveExpr | t | P.Value | adj.P.Val | B | gene description | root zone | time point |
|---|---|---|---|---|---|---|---|---|---|
| HORVU.MOREX.r3.7HG0742370.1 | 1.241051 | -1.39541 | 2.876372 | 0.005783 | 0.044393 | -2.29447 | Calmodulin | elongation zone | 12 h |
| HORVU.MOREX.r3.3HG0242520.1 | -0.54075 | 4.276799 | -2.95569 | 0.004647 | 0.038656 | -2.29639 | Protein O-mannosyl-transferase 2 | elongation zone | 12 h |
| HORVU.MOREX.r3.2HG0126010.1 | -0.35596 | 4.262083 | -2.9474 | 0.004756 | 0.039139 | -2.29656 | RING/U-box superfamily protein | elongation zone | 12 h |
| HORVU.MOREX.r3.1HG0077960.1 | 0.530065 | 7.898538 | 3.129066 | 0.002849 | 0.028054 | -2.29665 | #N/A | elongation zone | 12 h |
| HORVU.MOREX.r3.3HG0299530.1 | -0.5169 | 3.978312 | -2.92151 | 0.005108 | 0.041076 | -2.29666 | F-box protein | elongation zone | 12 h |
| HORVU.MOREX.r3.5HG0510370.1 | -0.30185 | 4.654596 | -2.97163 | 0.004446 | 0.037513 | -2.29705 | PHD finger protein ING | elongation zone | 12 h |
| HORVU.MOREX.r3.2HG0199330.1 | 0.628715 | 1.351775 | 3.031737 | 0.003757 | 0.033621 | -2.2999 | Anthocyanidin reductase | elongation zone | 12 h |
| HORVU.MOREX.r3.6HG0570230.1 | 0.8401 | 0.733369 | 2.84818 | 0.006245 | 0.046631 | -2.30134 | Aldose 1-epimerase family protein | elongation zone | 12 h |
| HORVU.MOREX.r3.5HG0515370.1 | -0.50243 | 3.30293 | -2.91764 | 0.005163 | 0.041371 | -2.30577 | F-box family protein | elongation zone | 12 h |
| HORVU.MOREX.r3.2HG0213430.1 | 0.76089 | 0.482455 | 2.996904 | 0.004143 | 0.035871 | -2.3073 | Calcium-dependent lipid-binding domain-containing protein | elongation zone | 12 h |
| HORVU.MOREX.r3.7HG0729510.1 | -0.48554 | 4.709018 | -3.00109 | 0.004095 | 0.035557 | -2.30734 | WEB family protein, chloroplastic | elongation zone | 12 h |
| HORVU.MOREX.r3.1HG0074650.2 | -1.02941 | 0.819721 | -2.84529 | 0.006294 | 0.046811 | -2.30742 | Solute carrier family 35 protein | elongation zone | 12 h |
| HORVU.MOREX.r3.3HG0240350.1 | 0.56487 | 2.152333 | 2.905165 | 0.005344 | 0.042285 | -2.30877 | Two-component response regulator | elongation zone | 12 h |
| HORVU.MOREX.r3.2HG0132160.1 | 2.189127 | -1.56452 | 3.061343 | 0.003456 | 0.031841 | -2.30891 | Ethylene-responsive transcription factor | elongation zone | 12 h |
| HORVU.MOREX.r3.6HG0634070.1 | -0.42937 | 4.157253 | -2.93927 | 0.004864 | 0.039613 | -2.30896 | Metal tolerance-like protein | elongation zone | 12 h |
| HORVU.MOREX.r3.3HG0303380.1 | 0.67693 | -0.86738 | 2.907599 | 0.005308 | 0.042087 | -2.30956 | Zinc finger protein CONSTANS | elongation zone | 12 h |
| HORVU.MOREX.r3.1HG0054770.1 | -0.57242 | 3.561935 | -2.90334 | 0.005371 | 0.042365 | -2.31011 | #N/A | elongation zone | 12 h |
| HORVU.MOREX.r3.4HG0335650.1 | -1.33168 | 8.653964 | -3.05852 | 0.003483 | 0.031953 | -2.31164 | Antimicrobial peptide MBP-1 | elongation zone | 12 h |
| HORVU.MOREX.r3.5HG0517260.1 | 0.540547 | 3.552392 | 2.886096 | 0.005631 | 0.043682 | -2.3121 | Glutamate carboxypeptidase 2 | elongation zone | 12 h |
| HORVU.MOREX.r3.5HG0504250.1 | -0.73654 | 4.308218 | -2.91379 | 0.005218 | 0.04165 | -2.3131 | #N/A | elongation zone | 12 h |
| HORVU.MOREX.r3.1HG0042440.1 | 1.39212 | -1.0109 | 2.855717 | 0.006118 | 0.046138 | -2.31543 | Long-chain-alcohol oxidase | elongation zone | 12 h |
| HORVU.MOREX.r3.6HG0622980.1 | -0.39238 | 4.462563 | -2.93185 | 0.004965 | 0.040219 | -2.31634 | Protein RFT1-like protein | elongation zone | 12 h |
| HORVU.MOREX.r3.1HG0033360.1 | -0.28475 | 4.66696 | -2.97354 | 0.004422 | 0.037375 | -2.31853 | Cysteine synthase | elongation zone | 12 h |
| HORVU.MOREX.r3.4HG0393260.1 | -0.32201 | 7.041152 | -3.0246 | 0.003833 | 0.034006 | -2.31905 | Oxidoreductase, 2OG-Fe(II) oxygenase family protein | elongation zone | 12 h |
| HORVU.MOREX.r3.1HG0090200.1 | -0.47267 | 5.563212 | -2.9777 | 0.004371 | 0.037158 | -2.31927 | NBS-LRR-like resistance protein | elongation zone | 12 h |
| HORVU.MOREX.r3.2HG0138300.1 | -0.70876 | 7.540969 | -3.04829 | 0.003586 | 0.032498 | -2.3197 | Histone H2A | elongation zone | 12 h |
| HORVU.MOREX.r3.6HG0546140.1 | -0.70236 | 0.639454 | -2.8475 | 0.006256 | 0.046701 | -2.32027 | F-box protein | elongation zone | 12 h |
| HORVU.MOREX.r3.4HG0388440.1 | 0.875687 | 4.312002 | 3.060968 | 0.003459 | 0.03186 | -2.32134 | #N/A | elongation zone | 12 h |
| HORVU.MOREX.r3.5HG0470800.1 | -2.18216 | 1.425735 | -2.97882 | 0.004358 | 0.037119 | -2.32311 | #N/A | elongation zone | 12 h |
| HORVU.MOREX.r3.4HG0408780.1 | 1.977515 | 0.649042 | 3.152131 | 0.002667 | 0.026789 | -2.32533 | Pentatricopeptide repeat-containing protein | elongation zone | 12 h |
| HORVU.MOREX.r3.1HG0054200.1 | -0.33308 | 5.861579 | -3.0037 | 0.004065 | 0.035366 | -2.32606 | E2 ubiquitin-conjugating-like enzyme | elongation zone | 12 h |
| HORVU.MOREX.r3.4HG0381150.1 | 1.851211 | -0.56149 | 3.105635 | 0.003047 | 0.02931 | -2.32777 | nuclear pore complex protein (DUF3414) | elongation zone | 12 h |
| HORVU.MOREX.r3.2HG0165200.1 | 0.628891 | 5.318501 | 3.117027 | 0.002949 | 0.028675 | -2.33121 | #N/A | elongation zone | 12 h |
| HORVU.MOREX.r3.3HG0298750.1 | -0.47868 | 5.760333 | -3.00551 | 0.004044 | 0.03527 | -2.33165 | magnesium transporter NIPA (DUF803) | elongation zone | 12 h |
| HORVU.MOREX.r3.6HG0569990.1 | 0.274275 | 6.145643 | 3.061362 | 0.003455 | 0.031841 | -2.33294 | Calmodulin binding protein | elongation zone | 12 h |
| HORVU.MOREX.r3.2HG0120560.1 | 1.161665 | 3.65156 | 3.147496 | 0.002702 | 0.02705 | -2.33341 | Peroxidase | elongation zone | 12 h |
| HORVU.MOREX.r3.7HG0696220.1 | -0.94967 | 1.255372 | -2.84248 | 0.006342 | 0.047069 | -2.33509 | #N/A | elongation zone | 12 h |
| HORVU.MOREX.r3.6HG0633640.1 | -0.27235 | 5.286476 | -3.00972 | 0.003997 | 0.034974 | -2.33552 | Thiaminase-1 | elongation zone | 12 h |
| HORVU.MOREX.r3.4HG0340650.1 | 0.399662 | 3.253441 | 2.909831 | 0.005276 | 0.041943 | -2.33604 | Peptidyl-prolyl cis-trans isomerase | elongation zone | 12 h |
| HORVU.MOREX.r3.3HG0249810.1 | -0.39998 | 5.280301 | -2.95375 | 0.004673 | 0.038786 | -2.3367 | Basic helix-loop-helix (BHLH) family transcription factor | elongation zone | 12 h |
| HORVU.MOREX.r3.7HG0744250.1 | -0.73466 | 5.1276 | -2.91532 | 0.005196 | 0.041571 | -2.33681 | Indole-3-glycerol phosphate synthase-like | elongation zone | 12 h |

**Table S1** Continued.

| ID | logFC | AveExpr | t | P.Value | adj.P.Val | B | gene description | root zone | time point |
|---|---|---|---|---|---|---|---|---|---|
| HORVU.MOREX.r3.6HG0539200.1 | -0.3625 | 5.866339 | -2.98103 | 0.004331 | 0.036952 | -2.33708 | Translation initiation factor IF-3 | elongation zone | 12 h |
| HORVU.MOREX.r3.6HG0618180.1 | 1.303284 | -2.40297 | 2.852517 | 0.006172 | 0.046337 | -2.33901 | Universal stress protein family protein | elongation zone | 12 h |
| HORVU.MOREX.r3.7HG0751450.1 | 1.429302 | 0.190404 | 2.837129 | 0.006435 | 0.047408 | -2.34023 | Glycosyltransferase family 92 protein | elongation zone | 12 h |
| HORVU.MOREX.r3.2HG0095650.1 | -1.45853 | 1.475239 | -2.87765 | 0.005763 | 0.044304 | -2.34027 | Mitochondrial inner membrane protease subunit 1 | elongation zone | 12 h |
| HORVU.MOREX.r3.4HG0391260.1 | 0.6147 | 2.259948 | 3.034078 | 0.003732 | 0.033458 | -2.34133 | Clathrin heavy chain | elongation zone | 12 h |
| HORVU.MOREX.r3.2HG0205680.1 | -1.94321 | 1.160765 | -3.04655 | 0.003603 | 0.032644 | -2.34155 | Pentatricopeptide repeat-containing protein | elongation zone | 12 h |
| HORVU.MOREX.r3.2HG0156140.1 | -1.14665 | 0.390799 | -2.86506 | 0.005964 | 0.045453 | -2.34255 | D-alanine--D-alanine ligase | elongation zone | 12 h |
| HORVU.MOREX.r3.7HG0663730.1 | -1.95767 | 2.17959 | -2.85614 | 0.006111 | 0.046117 | -2.34258 | Galactoside 2-alpha-L-fucosyltransferase | elongation zone | 12 h |
| HORVU.MOREX.r3.2HG0104260.1 | -0.66387 | 3.968231 | -2.88387 | 0.005665 | 0.043799 | -2.34312 | Carboxypeptidase | elongation zone | 12 h |
| HORVU.MOREX.r3.5HG0496780.1 | 0.436708 | 4.42219 | 2.952154 | 0.004693 | 0.038833 | -2.34365 | C2 domain-containing protein / GRAM domain-containing protein | elongation zone | 12 h |
| HORVU.MOREX.r3.6HG0630650.1 | -1.69052 | 3.863031 | -2.83906 | 0.006401 | 0.047335 | -2.34477 | Protein DETOXIFICATION | elongation zone | 12 h |
| HORVU.MOREX.r3.4HG0403630.1 | 1.562826 | -3.22747 | 2.877447 | 0.005766 | 0.044312 | -2.34488 | 3-ketoacyl-CoA synthase 19 | elongation zone | 12 h |
| HORVU.MOREX.r3.2HG0124660.1 | 0.296124 | 4.762294 | 2.9816 | 0.004324 | 0.036952 | -2.34519 | Dihydropteroate synthase, putative | elongation zone | 12 h |
| HORVU.MOREX.r3.5HG0490810.1 | -0.26348 | 7.111815 | -3.06782 | 0.003393 | 0.031493 | -2.34553 | C2 calcium/lipid-binding and GRAM domain protein | elongation zone | 12 h |
| HORVU.MOREX.r3.4HG0408000.1 | 2.639891 | -1.23574 | 3.124366 | 0.002888 | 0.028299 | -2.34677 | Cytokinin riboside 5'-monophosphate phosphoribohydrolase | elongation zone | 12 h |
| HORVU.MOREX.r3.4HG0378710.1 | -0.77229 | 2.281537 | -2.82765 | 0.006602 | 0.048163 | -2.34773 | F-box/SPRY domain-containing protein 1 | elongation zone | 12 h |
| HORVU.MOREX.r3.6HG0605120.1 | -0.61425 | 2.469228 | -2.88939 | 0.00558 | 0.04344 | -2.34813 | RING/U-box superfamily protein | elongation zone | 12 h |
| HORVU.MOREX.r3.5HG0427370.1 | -0.83484 | 1.304181 | -2.85749 | 0.006088 | 0.045983 | -2.34836 | #N/A | elongation zone | 12 h |
| HORVU.MOREX.r3.4HG0393890.1 | -0.6528 | 3.825718 | -2.8484 | 0.006241 | 0.046621 | -2.34921 | Hydroxyproline-rich glycoprotein | elongation zone | 12 h |
| HORVU.MOREX.r3.1HG0008440.1 | -0.70841 | 7.032439 | -2.99571 | 0.004157 | 0.035915 | -2.35482 | Glutathione S-transferase | elongation zone | 12 h |
| HORVU.MOREX.r3.2HG0101250.1 | 0.803399 | 0.35887 | 2.844063 | 0.006315 | 0.046917 | -2.35518 | Protein kinase superfamily protein | elongation zone | 12 h |
| HORVU.MOREX.r3.7HG0656440.1 | -0.45916 | 5.106986 | -2.91871 | 0.005148 | 0.041296 | -2.3562 | Adenylyltransferase and sulfurtransferase MOCS3 | elongation zone | 12 h |
| HORVU.MOREX.r3.5HG0430460.1 | 0.489467 | 1.634629 | 2.974328 | 0.004413 | 0.037339 | -2.35785 | Proline-rich family protein | elongation zone | 12 h |
| HORVU.MOREX.r3.5HG0433470.1 | 0.358114 | 8.072662 | 3.159287 | 0.002612 | 0.026441 | -2.35919 | Auxin-responsive protein | elongation zone | 12 h |
| HORVU.MOREX.r3.6HG0585520.2 | -0.541 | 2.330826 | -2.83341 | 0.0065 | 0.04774 | -2.36016 | Calmodulin binding protein-like protein | elongation zone | 12 h |
| HORVU.MOREX.r3.1HG0007670.1 | 0.583834 | 3.98467 | 2.921975 | 0.005102 | 0.041056 | -2.36017 | OCS-element binding factor 5 | elongation zone | 12 h |
| HORVU.MOREX.r3.7HG0648520.1 | -1.86787 | 0.765033 | -2.96732 | 0.0045 | 0.037778 | -2.36035 | Retrovirus-related Pol polyprotein from transposon TNT 1-94 | elongation zone | 12 h |
| HORVU.MOREX.r3.3HG0304230.1 | 2.170384 | -2.34493 | 2.955542 | 0.004649 | 0.038656 | -2.3609 | Cysteine proteinase | elongation zone | 12 h |
| HORVU.MOREX.r3.7HG0692070.1 | -0.7256 | 3.172308 | -2.83931 | 0.006397 | 0.047329 | -2.36194 | Pentatricopeptide repeat-containing protein | elongation zone | 12 h |
| HORVU.MOREX.r3.1HG0012330.1 | -0.77273 | 6.207328 | -2.94343 | 0.004808 | 0.039386 | -2.36389 | Peroxisomal membrane 22 kDa (Mpv17/PMP22) family protein | elongation zone | 12 h |
| HORVU.MOREX.r3.5HG0519240.1 | -0.58035 | 3.688408 | -2.85786 | 0.006082 | 0.045953 | -2.36421 | PHD finger-containing protein | elongation zone | 12 h |
| HORVU.MOREX.r3.5HG0487640.1 | -1.04074 | 6.108222 | -2.94972 | 0.004725 | 0.03901 | -2.36676 | Histone H2A | elongation zone | 12 h |
| HORVU.MOREX.r3.5HG0422490.1 | -0.41388 | 4.367008 | -2.90404 | 0.00536 | 0.042352 | -2.36829 | #N/A | elongation zone | 12 h |
| HORVU.MOREX.r3.1HG0059000.1 | -0.95461 | 2.194539 | -2.82618 | 0.006629 | 0.048286 | -2.37195 | Receptor-like kinase | elongation zone | 12 h |
| HORVU.MOREX.r3.6HG0599080.1 | -0.36576 | 8.233534 | -3.08314 | 0.003248 | 0.030632 | -2.37203 | Hsp70-Hsp90 organizing protein 1 | elongation zone | 12 h |
| HORVU.MOREX.r3.4HG0397530.1 | 1.080408 | 3.745067 | 2.950686 | 0.004712 | 0.03896 | -2.3731 | Plant calmodulin-binding-like protein | elongation zone | 12 h |
| HORVU.MOREX.r3.2HG0209910.1 | -0.41547 | 8.205773 | -3.09559 | 0.003135 | 0.029868 | -2.37503 | Pyruvate kinase | elongation zone | 12 h |
| HORVU.MOREX.r3.1HG0063000.1 | -0.43826 | 4.145332 | -2.89506 | 0.005494 | 0.042997 | -2.37533 | Methyl-CpG-binding domain-containing family protein | elongation zone | 12 h |
| HORVU.MOREX.r3.7HG0725060.1 | -1.52027 | 3.350962 | -2.82993 | 0.006562 | 0.048105 | -2.37705 | Microtubule associated family protein | elongation zone | 12 h |
| HORVU.MOREX.r3.7HG0670020.1 | -1.49884 | 7.08952 | -2.92998 | 0.00499 | 0.04038 | -2.37845 | Ribonucleoside-diphosphate reductase small chain | elongation zone | 12 h |
| HORVU.MOREX.r3.7HG0696190.1 | -0.43566 | 6.218033 | -2.96505 | 0.004528 | 0.037933 | -2.37943 | Glucose-1-phosphate adenylyltransferase | elongation zone | 12 h |

**Table S1** Continued.

| ID | logFC | AveExpr | t | P.Value | adj.P.Val | B | gene description | root zone | time point |
|---|---|---|---|---|---|---|---|---|---|
| HORVU.MOREX.r3.2HG0138280.1 | -0.62209 | 5.728929 | -2.94537 | 0.004782 | 0.039253 | -2.38053 | Histone H2A | elongation zone | 12 h |
| HORVU.MOREX.r3.4HG0343540.1 | -1.49432 | -0.26772 | -2.92525 | 0.005056 | 0.040734 | -2.38087 | BSD domain containing protein, expressed | elongation zone | 12 h |
| HORVU.MOREX.r3.4HG0393120.1 | -1.04833 | 1.488884 | -2.81082 | 0.00691 | 0.049645 | -2.38268 | Zinc finger protein | elongation zone | 12 h |
| HORVU.MOREX.r3.2HG0217550.1 | 0.388921 | 5.544285 | 3.056857 | 0.0035 | 0.032032 | -2.38533 | rRNA N-glycosidase | elongation zone | 12 h |
| HORVU.MOREX.r3.6HG0608960.1 | 0.529995 | 3.939137 | 2.91158 | 0.00525 | 0.041806 | -2.38728 | Galactan beta-1,4-galactosyltransferase GALS1 | elongation zone | 12 h |
| HORVU.MOREX.r3.3HG0327710.1 | -0.3268 | 7.225707 | -3.06312 | 0.003438 | 0.03178 | -2.38757 | Cytochrome b5 | elongation zone | 12 h |
| HORVU.MOREX.r3.6HG0567370.2 | 0.79654 | 0.756346 | 2.81843 | 0.006769 | 0.048991 | -2.38789 | #N/A | elongation zone | 12 h |
| HORVU.MOREX.r3.2HG0122140.1 | -0.47674 | 5.407639 | -2.98111 | 0.00433 | 0.036952 | -2.3883 | #N/A | elongation zone | 12 h |
| HORVU.MOREX.r3.1HG0079130.1 | 0.362439 | 7.030038 | 3.075665 | 0.003318 | 0.031043 | -2.38888 | ENTH/ANTH/VHS superfamily protein | elongation zone | 12 h |
| HORVU.MOREX.r3.5HG0475150.1 | 1.030572 | 1.439697 | 2.836485 | 0.006446 | 0.047445 | -2.39003 | ATP-binding cassette transporter subfamily A member | elongation zone | 12 h |
| HORVU.MOREX.r3.2HG0190300.1 | 0.520915 | 3.611092 | 2.950152 | 0.004719 | 0.039002 | -2.39068 | SHR5-receptor-like kinase | elongation zone | 12 h |
| HORVU.MOREX.r3.2HG0114840.1 | -0.66714 | 2.755393 | -2.81998 | 0.006741 | 0.048843 | -2.39232 | #N/A | elongation zone | 12 h |
| HORVU.MOREX.r3.3HG0294310.1 | 0.881653 | 1.665951 | 2.811627 | 0.006895 | 0.049555 | -2.39469 | Breast carcinoma-amplified sequence 3-like protein | elongation zone | 12 h |
| HORVU.MOREX.r3.3HG0320350.1 | 1.859923 | -0.52659 | 2.964422 | 0.004536 | 0.037976 | -2.39756 | Proline synthase co-transcribed bacterial | elongation zone | 12 h |
| HORVU.MOREX.r3.6HG0623830.1 | -1.88605 | 4.072429 | -2.81313 | 0.006867 | 0.049493 | -2.39788 | Disease resistance protein (NBS-LRR class) family | elongation zone | 12 h |
| HORVU.MOREX.r3.6HG0558250.1 | -0.59319 | 4.279829 | -2.89838 | 0.005444 | 0.042719 | -2.40005 | Disease resistance protein (NBS-LRR class) family | elongation zone | 12 h |
| HORVU.MOREX.r3.4HG0384700.1 | -0.92274 | 3.809991 | -2.85194 | 0.006181 | 0.04639 | -2.403 | Protein phosphatase 2C family protein | elongation zone | 12 h |
| HORVU.MOREX.r3.2HG0156590.1 | -0.43263 | 6.191891 | -2.96979 | 0.004469 | 0.037596 | -2.40346 | Protein embryo defective 1674 | elongation zone | 12 h |
| HORVU.MOREX.r3.1HG0064390.1 | 0.382057 | 5.924306 | 3.062444 | 0.003445 | 0.031796 | -2.40454 | Mitochondrial carrier family | elongation zone | 12 h |
| HORVU.MOREX.r3.2HG0142490.1 | -0.27853 | 6.46738 | -2.98918 | 0.004233 | 0.036363 | -2.40801 | Protein FLX-like 1 | elongation zone | 12 h |
| HORVU.MOREX.r3.5HG0460560.1 | -2.33798 | -0.88929 | -3.13181 | 0.002827 | 0.027901 | -2.40809 | Set1/Ash2 histone methyltransferase complex subunit ASH2 | elongation zone | 12 h |
| HORVU.MOREX.r3.7HG0674680.1 | 1.236367 | -0.96251 | 2.827736 | 0.006601 | 0.048163 | -2.40977 | Peroxidase | elongation zone | 12 h |
| HORVU.MOREX.r3.2HG0180060.1 | -1.21111 | 1.155124 | -2.86602 | 0.005949 | 0.045377 | -2.41056 | Receptor-kinase, putative | elongation zone | 12 h |
| HORVU.MOREX.r3.5HG0499620.1 | -0.53511 | 3.333821 | -2.84989 | 0.006216 | 0.046499 | -2.41168 | Methyltransferase, FkbM family protein, expressed | elongation zone | 12 h |
| HORVU.MOREX.r3.5HG0495580.1 | -0.3148 | 5.893655 | -2.971 | 0.004454 | 0.037542 | -2.41224 | Ankyrin repeat domain containing protein | elongation zone | 12 h |
| HORVU.MOREX.r3.1HG0056740.1 | -0.55149 | 8.162243 | -3.06246 | 0.003445 | 0.031796 | -2.41227 | Pyruvate kinase | elongation zone | 12 h |
| HORVU.MOREX.r3.1HG0059850.1 | 0.524134 | 4.1487 | 2.947521 | 0.004754 | 0.039139 | -2.41249 | Lipase | elongation zone | 12 h |
| HORVU.MOREX.r3.3HG0293420.1 | 1.061129 | -1.27948 | 2.871752 | 0.005856 | 0.04486 | -2.41536 | Expansin | elongation zone | 12 h |
| HORVU.MOREX.r3.6HG0554560.1 | -1.00637 | 4.575431 | -2.85861 | 0.00607 | 0.045938 | -2.41668 | BZIP transcription factor | elongation zone | 12 h |
| HORVU.MOREX.r3.1HG0052790.1 | -0.46969 | 6.26243 | -2.98713 | 0.004258 | 0.036533 | -2.41687 | Nucleoid-associated protein Ccur92_18190 | elongation zone | 12 h |
| HORVU.MOREX.r3.4HG0398980.1 | 0.353057 | 5.041369 | 3.010092 | 0.003993 | 0.034968 | -2.41886 | ADP-ribosylation factor, putative | elongation zone | 12 h |
| HORVU.MOREX.r3.4HG0401980.1 | 2.833202 | -2.74383 | 2.882004 | 0.005694 | 0.043944 | -2.42017 | Sulfate transporter | elongation zone | 12 h |
| HORVU.MOREX.r3.6HG0603210.1 | 2.042637 | 0.043239 | 3.152146 | 0.002666 | 0.026789 | -2.42093 | DNA-directed RNA polymerase subunit | elongation zone | 12 h |
| HORVU.MOREX.r3.2HG0096760.1 | -0.80443 | 4.20342 | -2.82015 | 0.006738 | 0.048837 | -2.42139 | TOM1-like protein 2 | elongation zone | 12 h |
| HORVU.MOREX.r3.3HG0320520.1 | 0.680544 | 2.494839 | 2.828518 | 0.006587 | 0.048163 | -2.42162 | DUF241 domain protein, putative (DUF241) | elongation zone | 12 h |
| HORVU.MOREX.r3.3HG0302340.1 | -2.406 | -0.09437 | -3.00838 | 0.004012 | 0.035058 | -2.42405 | Eukaryotic aspartyl protease family protein, expressed | elongation zone | 12 h |
| HORVU.MOREX.r3.7HG0697940.1 | -1.34573 | 0.148475 | -2.90906 | 0.005287 | 0.042 | -2.4243 | UvrABC system protein C | elongation zone | 12 h |
| HORVU.MOREX.r3.5HG0467370.1 | -0.7575 | 7.65806 | -3.00004 | 0.004107 | 0.035609 | -2.42757 | Pleiotropic drug resistance ABC transporter | elongation zone | 12 h |
| HORVU.MOREX.r3.6HG0628930.1 | -0.49945 | 7.959732 | -3.02025 | 0.00388 | 0.034278 | -2.42951 | Alcohol dehydrogenase | elongation zone | 12 h |
| HORVU.MOREX.r3.7HG0668820.1 | 0.500293 | 4.620228 | 3.050435 | 0.003564 | 0.032373 | -2.42981 | Endoglucanase | elongation zone | 12 h |
| HORVU.MOREX.r3.3HG0219750.1 | -1.52347 | -0.71619 | -2.85327 | 0.006159 | 0.046276 | -2.43105 | Disease resistance protein (NBS-LRR class) family | elongation zone | 12 h |

**Table S1** Continued.

| ID | logFC | AveExpr | t | P.Value | adj.P.Val | B | gene description | root zone | time point |
|---|---|---|---|---|---|---|---|---|---|
| HORVU.MOREX.r3.4HG0392590.1 | -0.43266 | 3.653452 | -2.85835 | 0.006074 | 0.045944 | -2.43407 | Calmodulin-lysine N-methyltransferase | elongation zone | 12 h |
| HORVU.MOREX.r3.1HG0032230.1 | -0.84171 | 4.670024 | -2.82824 | 0.006592 | 0.048163 | -2.43605 | Rad3-related DNA helicase family protein | elongation zone | 12 h |
| HORVU.MOREX.r3.3HG0243130.1 | -0.68236 | 2.631713 | -2.83875 | 0.006407 | 0.047359 | -2.43625 | Peroxidase | elongation zone | 12 h |
| HORVU.MOREX.r3.5HG0514790.1 | -0.40292 | 9.909631 | -3.11551 | 0.002962 | 0.028751 | -2.43635 | Elongation factor 1-alpha | elongation zone | 12 h |
| HORVU.MOREX.r3.7HG0637470.1 | -0.47142 | 3.443051 | -2.85158 | 0.006187 | 0.04639 | -2.44073 | NBS-LRR resistance-like protein | elongation zone | 12 h |
| HORVU.MOREX.r3.2HG0208790.1 | -1.66877 | 3.913939 | -2.86043 | 0.00604 | 0.045836 | -2.44432 | Glycosyltransferase | elongation zone | 12 h |
| HORVU.MOREX.r3.4HG0390030.1 | -2.18136 | 0.118063 | -2.82048 | 0.006732 | 0.048811 | -2.44632 | Nodulation-signaling pathway 2 protein | elongation zone | 12 h |
| HORVU.MOREX.r3.5HG0534640.1 | -0.51927 | 8.816317 | -3.04594 | 0.003609 | 0.032683 | -2.44803 | Elongation factor Tu | elongation zone | 12 h |
| HORVU.MOREX.r3.3HG0259660.1 | -0.58929 | 3.912666 | -2.81681 | 0.006799 | 0.049107 | -2.44876 | CRS2-associated factor 1, chloroplastic | elongation zone | 12 h |
| HORVU.MOREX.r3.3HG0276730.1 | 0.273047 | 6.945927 | 3.027033 | 0.003807 | 0.033836 | -2.44887 | Long-Chain Acyl-CoA Synthetase | elongation zone | 12 h |
| HORVU.MOREX.r3.7HG0667660.1 | 0.934211 | 5.314359 | 2.990924 | 0.004213 | 0.036201 | -2.44928 | Peroxidase | elongation zone | 12 h |
| HORVU.MOREX.r3.7HG0744540.1 | -2.06586 | -0.40765 | -2.84133 | 0.006362 | 0.04718 | -2.45362 | Glycosyltransferase | elongation zone | 12 h |
| HORVU.MOREX.r3.1HG0085380.1 | 0.39141 | 4.053078 | 2.890854 | 0.005558 | 0.043299 | -2.45436 | 2-oxoglutarate (2OG) and Fe(II)-dependent oxygenase superfamily protein | elongation zone | 12 h |
| HORVU.MOREX.r3.4HG0397570.1 | 0.75818 | 4.104655 | 2.894398 | 0.005504 | 0.043043 | -2.4557 | Glucan endo-1,3-beta-glucosidase 3 | elongation zone | 12 h |
| HORVU.MOREX.r3.3HG0237330.1 | -1.62893 | 0.337971 | -2.85874 | 0.006068 | 0.045938 | -2.45608 | Protein FAR1-RELATED SEQUENCE 5 | elongation zone | 12 h |
| HORVU.MOREX.r3.7HG0687540.1 | -0.29001 | 5.801521 | -2.97074 | 0.004457 | 0.037543 | -2.45788 | Proteasome subunit alpha type | elongation zone | 12 h |
| HORVU.MOREX.r3.3HG0301930.1 | 0.4544 | 4.019686 | 2.956413 | 0.004638 | 0.03861 | -2.45813 | G patch domain protein | elongation zone | 12 h |
| HORVU.MOREX.r3.5HG0462780.1 | -0.40363 | 7.18947 | -3.01641 | 0.003923 | 0.03453 | -2.45872 | Transporter, putative, expressed | elongation zone | 12 h |
| HORVU.MOREX.r3.4HG0352540.1 | -0.57039 | 2.652903 | -2.847 | 0.006265 | 0.046718 | -2.45879 | Glycosyltransferase | elongation zone | 12 h |
| HORVU.MOREX.r3.7HG0717850.1 | -0.35545 | 6.287604 | -2.97688 | 0.004381 | 0.037198 | -2.45924 | Potassium efflux antiporter | elongation zone | 12 h |
| HORVU.MOREX.r3.1HG0044290.1 | 0.607068 | 3.742696 | 2.811797 | 0.006892 | 0.049549 | -2.45969 | electron protein, putative (Protein of unknown function, DUF547) | elongation zone | 12 h |
| HORVU.MOREX.r3.4HG0394810.1 | -0.98318 | 5.082806 | -2.80831 | 0.006957 | 0.049739 | -2.45993 | U-box domain-containing protein | elongation zone | 12 h |
| HORVU.MOREX.r3.3HG0310920.1 | 0.629205 | 5.137634 | 2.951105 | 0.004707 | 0.03893 | -2.46014 | Histidine kinase | elongation zone | 12 h |
| HORVU.MOREX.r3.3HG0253450.1 | -0.32934 | 5.357439 | -2.94114 | 0.004839 | 0.039574 | -2.46135 | FHA domain containing protein, expressed | elongation zone | 12 h |
| HORVU.MOREX.r3.3HG0293430.1 | 1.207805 | -1.42288 | 2.818337 | 0.006771 | 0.048991 | -2.46182 | Expansin | elongation zone | 12 h |
| HORVU.MOREX.r3.3HG0219800.1 | -0.54266 | 3.630299 | -2.80964 | 0.006932 | 0.049665 | -2.46211 | Dephospho-CoA kinase, putative, expressed | elongation zone | 12 h |
| HORVU.MOREX.r3.4HG0387580.1 | 0.327166 | 4.638536 | 2.959243 | 0.004602 | 0.038401 | -2.46241 | Xylosyltransferase 1 | elongation zone | 12 h |
| HORVU.MOREX.r3.2HG0197710.1 | -1.15813 | 4.270225 | -2.8252 | 0.006646 | 0.048379 | -2.46375 | #N/A | elongation zone | 12 h |
| HORVU.MOREX.r3.7HG0725290.1 | -0.47921 | 3.61352 | -2.85102 | 0.006197 | 0.04639 | -2.46493 | YGL010w-like protein | elongation zone | 12 h |
| HORVU.MOREX.r3.1HG0008640.1 | 0.489631 | 3.733371 | 2.876708 | 0.005777 | 0.044368 | -2.46546 | 7,8-dihydroneopterin aldolase | elongation zone | 12 h |
| HORVU.MOREX.r3.7HG0736870.1 | 0.43898 | 5.259228 | 2.97828 | 0.004364 | 0.037141 | -2.46747 | Molybdenum cofactor sulfurase | elongation zone | 12 h |
| HORVU.MOREX.r3.1HG0073520.1 | -0.78185 | 3.742361 | -2.90372 | 0.005365 | 0.042358 | -2.46779 | Heavy metal transport/detoxification superfamily protein | elongation zone | 12 h |
| HORVU.MOREX.r3.4HG0403510.1 | -0.38327 | 4.248358 | -2.89784 | 0.005452 | 0.042752 | -2.46789 | Protein kinase | elongation zone | 12 h |
| HORVU.MOREX.r3.4HG0364880.1 | -1.25577 | -0.54939 | -2.81539 | 0.006825 | 0.049261 | -2.471 | cysteine-rich RECEPTOR-like kinase | elongation zone | 12 h |
| HORVU.MOREX.r3.3HG0306030.1 | -0.45642 | 3.28785 | -2.8511 | 0.006195 | 0.04639 | -2.47399 | Rhomboid protein | elongation zone | 12 h |
| HORVU.MOREX.r3.1HG0046900.1 | -0.31894 | 3.626727 | -2.88652 | 0.005624 | 0.043667 | -2.47456 | 1-acyl-sn-glycerol-3-phosphate acyltransferase | elongation zone | 12 h |
| HORVU.MOREX.r3.4HG0342090.1 | -0.46843 | 5.870045 | -2.92878 | 0.005007 | 0.040463 | -2.47502 | Remorin | elongation zone | 12 h |
| HORVU.MOREX.r3.7HG0730870.1 | 0.350572 | 7.493886 | 3.075644 | 0.003318 | 0.031043 | -2.47548 | Glycosyltransferases | elongation zone | 12 h |
| HORVU.MOREX.r3.3HG0308980.1 | -0.69354 | 4.452519 | -2.82165 | 0.006711 | 0.048693 | -2.47574 | S-adenosyl-L-methionine-dependent methyltransferase, putative | elongation zone | 12 h |
| HORVU.MOREX.r3.1HG0075340.1 | 0.776156 | 1.815297 | 2.830777 | 0.006547 | 0.04803 | -2.47603 | Alpha/beta-Hydrolases superfamily protein | elongation zone | 12 h |
| HORVU.MOREX.r3.5HG0513120.1 | -0.31344 | 5.944244 | -2.99696 | 0.004142 | 0.035871 | -2.47714 | Cytochrome P450 allene oxide synthase | elongation zone | 12 h |

**Table S1** Continued.

| ID | logFC | AveExpr | t | P.Value | adj.P.Val | B | gene description | root zone | time point |
|---|---|---|---|---|---|---|---|---|---|
| HORVU.MOREX.r3.2HG0167540.1 | -0.22235 | 7.425024 | -3.02086 | 0.003874 | 0.03426 | -2.47785 | Histidine decarboxylase | elongation zone | 12 h |
| HORVU.MOREX.r3.6HG0615630.1 | 0.47469 | 4.54843 | 2.885364 | 0.005642 | 0.043691 | -2.47913 | Rac-like GTP-binding protein | elongation zone | 12 h |
| HORVU.MOREX.r3.7HG0708120.1 | -1.27115 | 6.606021 | -2.88377 | 0.005667 | 0.043799 | -2.48081 | High mobility group family | elongation zone | 12 h |
| HORVU.MOREX.r3.2HG0100360.1 | 0.393214 | 4.261633 | 2.903572 | 0.005367 | 0.04236 | -2.48468 | 3-oxo-5-alpha-steroid 4-dehydrogenase | elongation zone | 12 h |
| HORVU.MOREX.r3.1HG0077170.1 | -0.56227 | 4.830925 | -2.84061 | 0.006374 | 0.047226 | -2.4858 | Protein EMBRYONIC FLOWER 1 | elongation zone | 12 h |
| HORVU.MOREX.r3.2HG0171100.1 | -0.44144 | 4.075179 | -2.84696 | 0.006266 | 0.046718 | -2.48621 | Plectin-like protein | elongation zone | 12 h |
| HORVU.MOREX.r3.3HG0323810.1 | -0.47724 | 6.634621 | -2.93996 | 0.004855 | 0.039601 | -2.48775 | SWAP (Suppressor-of-White-APricot)/surp domain-containing protein / ubiquitin f | elongation zone | 12 h |
| HORVU.MOREX.r3.4HG0387150.1 | -0.87583 | 6.899898 | -2.93963 | 0.004859 | 0.039613 | -2.48913 | Histone H2A | elongation zone | 12 h |
| HORVU.MOREX.r3.7HG0739230.1 | 2.960629 | -3.45098 | 3.051928 | 0.003549 | 0.032342 | -2.48943 | Jacalin-like lectin | elongation zone | 12 h |
| HORVU.MOREX.r3.5HG0468940.1 | -0.32013 | 5.005775 | -2.89247 | 0.005533 | 0.043222 | -2.48965 | NAD(P)-binding Rossmann-fold superfamily protein | elongation zone | 12 h |
| HORVU.MOREX.r3.1HG0007570.1 | -1.36066 | -2.58012 | -2.86021 | 0.006043 | 0.045842 | -2.48985 | Werner Syndrome-like exonuclease | elongation zone | 12 h |
| HORVU.MOREX.r3.5HG0528890.1 | -1.0161 | 5.084337 | -2.80495 | 0.00702 | 0.049999 | -2.49133 | GATA transcription factor, putative | elongation zone | 12 h |
| HORVU.MOREX.r3.2HG0165640.1 | -2.49908 | -0.84713 | -2.99243 | 0.004195 | 0.036159 | -2.49312 | CsAtPR5 | elongation zone | 12 h |
| HORVU.MOREX.r3.7HG0714400.1 | 0.784239 | 2.702391 | 2.809847 | 0.006928 | 0.049654 | -2.49407 | Auxin-responsive protein | elongation zone | 12 h |
| HORVU.MOREX.r3.6HG0610890.1 | 0.467718 | 5.180456 | 2.981307 | 0.004328 | 0.036952 | -2.49642 | KH domain-containing protein | elongation zone | 12 h |
| HORVU.MOREX.r3.6HG0597460.1 | -1.84011 | -0.69687 | -2.95377 | 0.004672 | 0.038786 | -2.49713 | Glucose-6-phosphate isomerase | elongation zone | 12 h |
| HORVU.MOREX.r3.2HG0135860.1 | -0.386 | 5.818016 | -2.90499 | 0.005346 | 0.042285 | -2.49859 | Transcription factor jumonji domain-containing protein | elongation zone | 12 h |
| HORVU.MOREX.r3.7HG0680680.1 | -0.3036 | 5.101819 | -2.93401 | 0.004935 | 0.040075 | -2.49886 | 3-hydroxy-3-methylglutaryl coenzyme A reductase | elongation zone | 12 h |
| HORVU.MOREX.r3.7HG0745700.1 | -0.25833 | 5.461365 | -2.94989 | 0.004723 | 0.03901 | -2.49889 | Phox (PX) domain-containing protein | elongation zone | 12 h |
| HORVU.MOREX.r3.7HG0689490.1 | -0.55096 | 6.098735 | -2.9088 | 0.005291 | 0.042014 | -2.4993 | Peptidyl-prolyl cis-trans isomerase | elongation zone | 12 h |
| HORVU.MOREX.r3.5HG0436900.1 | 0.365132 | 4.896931 | 2.945087 | 0.004786 | 0.039269 | -2.50023 | Syntaxin, putative | elongation zone | 12 h |
| HORVU.MOREX.r3.1HG0086140.1 | -0.79722 | 4.57931 | -2.81957 | 0.006748 | 0.048879 | -2.50048 | Oxidoreductase | elongation zone | 12 h |
| HORVU.MOREX.r3.7HG0688300.1 | -0.42671 | 4.553776 | -2.85602 | 0.006113 | 0.046117 | -2.50187 | O-fucosyltransferase-like protein | elongation zone | 12 h |
| HORVU.MOREX.r3.7HG0684040.1 | -0.52279 | 5.674326 | -2.89605 | 0.005479 | 0.042914 | -2.50195 | Glutamate-1-semialdehyde 2,1-aminomutase | elongation zone | 12 h |
| HORVU.MOREX.r3.3HG0265140.1 | 1.991628 | -1.23714 | 2.827362 | 0.006608 | 0.048183 | -2.5021 | DUF3511 domain protein | elongation zone | 12 h |
| HORVU.MOREX.r3.4HG0401630.1 | 0.679082 | 1.754312 | 2.822335 | 0.006698 | 0.048636 | -2.50386 | Protein kinase | elongation zone | 12 h |
| HORVU.MOREX.r3.7HG0688930.1 | 0.808843 | -0.39064 | 2.817012 | 0.006795 | 0.049097 | -2.50407 | DUF868 family protein | elongation zone | 12 h |
| HORVU.MOREX.r3.5HG0471790.1 | -0.45592 | 8.627392 | -3.04903 | 0.003578 | 0.032459 | -2.50419 | Phosphate translocator | elongation zone | 12 h |
| HORVU.MOREX.r3.3HG0255150.1 | -0.27335 | 5.442871 | -2.93491 | 0.004923 | 0.040007 | -2.50446 | Sec14p-like phosphatidylinositol transfer family protein | elongation zone | 12 h |
| HORVU.MOREX.r3.5HG0444270.1 | 0.445868 | 5.156347 | 3.024224 | 0.003837 | 0.034028 | -2.50446 | Pectin lyase-like superfamily protein | elongation zone | 12 h |
| HORVU.MOREX.r3.5HG0427060.1 | -0.87135 | 5.128579 | -2.83754 | 0.006428 | 0.047408 | -2.50484 | 50S ribosomal protein L14 | elongation zone | 12 h |
| HORVU.MOREX.r3.5HG0526140.1 | -0.45187 | 4.078722 | -2.86153 | 0.006022 | 0.045715 | -2.5051 | NBS-LRR disease resistance protein, putative | elongation zone | 12 h |
| HORVU.MOREX.r3.7HG0663400.1 | -0.28541 | 6.420462 | -2.97039 | 0.004461 | 0.03755 | -2.50604 | LysM domain-containing GPI-anchored protein 1 | elongation zone | 12 h |
| HORVU.MOREX.r3.1HG0058050.1 | -0.5195 | 5.246818 | -2.87915 | 0.005739 | 0.044183 | -2.50885 | 8-amino-7-oxononanoate synthase | elongation zone | 12 h |
| HORVU.MOREX.r3.7HG0636530.1 | -0.39294 | 4.42133 | -2.89149 | 0.005548 | 0.043288 | -2.51003 | MADS-box transcription factor 8 | elongation zone | 12 h |
| HORVU.MOREX.r3.2HG0137620.1 | 0.474327 | 3.671497 | 2.845662 | 0.006288 | 0.046798 | -2.51036 | Phototropic-responsive NPH3 family protein | elongation zone | 12 h |
| HORVU.MOREX.r3.3HG0292680.1 | 0.62085 | 2.459133 | 2.908463 | 0.005295 | 0.042036 | -2.51239 | Replicase polyprotein 1a | elongation zone | 12 h |
| HORVU.MOREX.r3.4HG0383340.1 | -1.59328 | 0.295717 | -2.8988 | 0.005438 | 0.042705 | -2.5133 | Pentatricopeptide repeat-containing protein | elongation zone | 12 h |
| HORVU.MOREX.r3.1HG0078040.1 | 0.450855 | 5.579406 | 2.994948 | 0.004166 | 0.035962 | -2.51592 | Absent in melanoma 1 protein | elongation zone | 12 h |
| HORVU.MOREX.r3.2HG0172050.1 | -0.35655 | 8.585874 | -3.0253 | 0.003826 | 0.033984 | -2.51633 | RNA-binding protein | elongation zone | 12 h |
| HORVU.MOREX.r3.7HG0664130.1 | -0.36054 | 5.138266 | -2.87097 | 0.005869 | 0.044935 | -2.51765 | Dipeptidyl peptidase family member 6 | elongation zone | 12 h |

**Table S1** Continued.

| ID | logFC | AveExpr | t | P.Value | adj.P.Val | B | gene description | root zone | time point |
|---|---|---|---|---|---|---|---|---|---|
| HORVU.MOREX.r3.1HG0071310.1 | -0.48079 | 4.338136 | -2.80651 | 0.006991 | 0.049912 | -2.52033 | WEAK movement UNDER BLUE LIGHT-like protein | elongation zone | 12 h |
| HORVU.MOREX.r3.7HG0644190.1 | 0.564148 | 0.806058 | 2.844558 | 0.006307 | 0.046884 | -2.52037 | Glucan endo-1,3-beta-glucosidase, putative | elongation zone | 12 h |
| HORVU.MOREX.r3.1HG0069110.1 | -0.47836 | 4.126574 | -2.80499 | 0.007019 | 0.049999 | -2.52342 | Zinc finger protein 830 | elongation zone | 12 h |
| HORVU.MOREX.r3.7HG0720720.1 | -0.48704 | 3.515226 | -2.80505 | 0.007018 | 0.049999 | -2.52412 | Vesicle transport protein | elongation zone | 12 h |
| HORVU.MOREX.r3.2HG0204960.1 | -0.33121 | 4.354857 | -2.85083 | 0.0062 | 0.046397 | -2.52437 | RING/U-box superfamily protein | elongation zone | 12 h |
| HORVU.MOREX.r3.5HG0521030.1 | -0.60368 | 4.415413 | -2.81864 | 0.006765 | 0.048986 | -2.52479 | basic helix-loop-helix (bHLH) DNA-binding superfamily protein | elongation zone | 12 h |
| HORVU.MOREX.r3.6HG0583020.1 | -0.37984 | 3.316584 | -2.84616 | 0.006279 | 0.046769 | -2.52556 | Protein phosphatase 2C containing protein | elongation zone | 12 h |
| HORVU.MOREX.r3.6HG0554700.1 | 0.304527 | 7.324166 | 3.027312 | 0.003804 | 0.033836 | -2.5256 | Sec14p-like phosphatidylinositol transfer family protein | elongation zone | 12 h |
| HORVU.MOREX.r3.1HG0080770.1 | -0.94248 | 6.795288 | -2.92534 | 0.005055 | 0.040734 | -2.52777 | Histone H4 | elongation zone | 12 h |
| HORVU.MOREX.r3.3HG0260580.1 | -0.47598 | 4.905647 | -2.84058 | 0.006375 | 0.047226 | -2.52808 | DUF630 family protein, putative (DUF630 and DUF632) | elongation zone | 12 h |
| HORVU.MOREX.r3.2HG0180580.1 | -1.42213 | 7.018787 | -2.85991 | 0.006049 | 0.045842 | -2.52838 | Proliferating cell nuclear antigen | elongation zone | 12 h |
| HORVU.MOREX.r3.3HG0253630.1 | 0.944014 | 3.730748 | 2.830293 | 0.006555 | 0.048076 | -2.5286 | Late embryogenesis abundant protein | elongation zone | 12 h |
| HORVU.MOREX.r3.7HG0703780.1 | -0.54752 | 5.334771 | -2.87017 | 0.005882 | 0.04499 | -2.53103 | Succinate dehydrogenase [ubiquinone] iron-sulfur subunit, mitochondrial | elongation zone | 12 h |
| HORVU.MOREX.r3.7HG0657370.1 | -0.32765 | 5.109469 | -2.90272 | 0.00538 | 0.042386 | -2.53148 | NADH-quinone oxidoreductase subunit C | elongation zone | 12 h |
| HORVU.MOREX.r3.7HG0648600.1 | -0.58197 | 5.915169 | -2.87955 | 0.005733 | 0.044174 | -2.53214 | Elongation factor 4 | elongation zone | 12 h |
| HORVU.MOREX.r3.2HG0116870.1 | -0.82549 | 6.950005 | -2.91392 | 0.005216 | 0.04165 | -2.53652 | Condensin complex subunit 1 | elongation zone | 12 h |
| HORVU.MOREX.r3.3HG0295380.1 | 0.711616 | 1.713149 | 2.855187 | 0.006127 | 0.046171 | -2.53917 | Cytochrome P450 | elongation zone | 12 h |
| HORVU.MOREX.r3.5HG0532200.1 | 0.812732 | 4.356719 | 2.953918 | 0.00467 | 0.038786 | -2.53936 | Patatin | elongation zone | 12 h |
| HORVU.MOREX.r3.5HG0493200.1 | 0.3917 | 8.020324 | 3.056629 | 0.003502 | 0.032038 | -2.54429 | Fasciclin-like arabinogalactan-protein-like | elongation zone | 12 h |
| HORVU.MOREX.r3.1HG0084920.1 | -0.30692 | 5.205134 | -2.90251 | 0.005383 | 0.042386 | -2.54655 | ER membrane protein complex subunit 6 | elongation zone | 12 h |
| HORVU.MOREX.r3.5HG0467880.1 | -0.45989 | 5.367745 | -2.84968 | 0.006219 | 0.046509 | -2.54665 | DNA-directed RNA polymerase family protein | elongation zone | 12 h |
| HORVU.MOREX.r3.3HG0278120.1 | 0.381498 | 4.5822 | 2.863083 | 0.005996 | 0.04559 | -2.54684 | CTD small phosphatase-like protein | elongation zone | 12 h |
| HORVU.MOREX.r3.7HG0706930.1 | -0.49784 | 4.439187 | -2.82686 | 0.006617 | 0.048214 | -2.54753 | F-box family protein | elongation zone | 12 h |
| HORVU.MOREX.r3.1HG0019440.1 | -0.39541 | 5.729684 | -2.88795 | 0.005602 | 0.043562 | -2.55203 | Vegetative cell wall protein gp1 | elongation zone | 12 h |
| HORVU.MOREX.r3.5HG0537780.1 | 0.414884 | 7.666685 | 3.059851 | 0.00347 | 0.031904 | -2.55281 | COP1-interacting-like protein | elongation zone | 12 h |
| HORVU.MOREX.r3.2HG0200730.1 | -0.19684 | 7.663906 | -3.01542 | 0.003933 | 0.034581 | -2.55449 | Protein transport SEC13-like protein | elongation zone | 12 h |
| HORVU.MOREX.r3.6HG0569480.1 | -2.5324 | 2.886546 | -2.85984 | 0.00605 | 0.045842 | -2.55623 | Tubby-like F-box protein | elongation zone | 12 h |
| HORVU.MOREX.r3.2HG0122520.1 | 0.753511 | 0.436988 | 2.812281 | 0.006883 | 0.049549 | -2.55627 | SAUR-like auxin-responsive protein family | elongation zone | 12 h |
| HORVU.MOREX.r3.4HG0384400.1 | 0.507289 | 4.602549 | 2.965347 | 0.004524 | 0.037933 | -2.55742 | Ethylene-overproduction protein 1 | elongation zone | 12 h |
| HORVU.MOREX.r3.5HG0439710.1 | -1.97296 | -0.51069 | -2.97594 | 0.004393 | 0.037263 | -2.55811 | Zinc finger protein | elongation zone | 12 h |
| HORVU.MOREX.r3.3HG0296380.1 | 0.589903 | 4.241542 | 2.942721 | 0.004818 | 0.039433 | -2.55816 | Type I inositol-1,4,5-trisphosphate 5-phosphatase | elongation zone | 12 h |
| HORVU.MOREX.r3.7HG0730350.1 | -0.53824 | 7.382323 | -2.9709 | 0.004455 | 0.037542 | -2.56036 | Histone H2A | elongation zone | 12 h |
| HORVU.MOREX.r3.6HG0584860.1 | -0.49169 | 4.293818 | -2.81189 | 0.00689 | 0.049549 | -2.56408 | 3-oxoacyl-reductase | elongation zone | 12 h |
| HORVU.MOREX.r3.1HG0030200.1 | -0.41757 | 4.198578 | -2.87332 | 0.005831 | 0.044731 | -2.56534 | DUF641 family protein | elongation zone | 12 h |
| HORVU.MOREX.r3.6HG0614110.1 | 0.392329 | 4.183035 | 2.947293 | 0.004757 | 0.039139 | -2.56753 | nucleolar GTP-binding protein | elongation zone | 12 h |
| HORVU.MOREX.r3.4HG0412650.1 | 1.761252 | 1.247202 | 2.845405 | 0.006292 | 0.046811 | -2.56876 | TVP38/TMEM64 family membrane protein | elongation zone | 12 h |
| HORVU.MOREX.r3.5HG0459320.1 | -0.44249 | 4.812268 | -2.86006 | 0.006046 | 0.045842 | -2.56898 | Peptidylprolyl isomerase | elongation zone | 12 h |
| HORVU.MOREX.r3.6HG0555040.1 | 0.520128 | 3.833259 | 2.838256 | 0.006415 | 0.047365 | -2.56959 | Succinate dehydrogenase [ubiquinone] flavoprotein subunit, mitochondrial | elongation zone | 12 h |
| HORVU.MOREX.r3.2HG0205550.1 | -0.80192 | 5.79238 | -2.82065 | 0.006729 | 0.048807 | -2.57089 | Thymidylate synthase | elongation zone | 12 h |
| HORVU.MOREX.r3.3HG0279110.1 | 2.077134 | -0.01382 | 2.81035 | 0.006918 | 0.049654 | -2.5748 | NAC domain-containing protein | elongation zone | 12 h |
| HORVU.MOREX.r3.2HG0104500.1 | -0.42188 | 4.987186 | -2.84908 | 0.006229 | 0.046551 | -2.57494 | INO80 complex subunit C | elongation zone | 12 h |

**Table S1** Continued.

| ID | logFC | AveExpr | t | P.Value | adj.P.Val | B | gene description | root zone | time point |
|---|---|---|---|---|---|---|---|---|---|
| HORVU.MOREX.r3.1HG0003620.1 | 0.54509 | 2.573513 | 2.816352 | 0.006807 | 0.04915 | -2.57579 | Protein ZINC INDUCED FACILITATOR-LIKE 1 | elongation zone | 12 h |
| HORVU.MOREX.r3.7HG0710170.1 | -1.79832 | 0.072486 | -2.87927 | 0.005737 | 0.044183 | -2.57735 | ATP-dependent DNA helicase 2 subunit KU70 | elongation zone | 12 h |
| HORVU.MOREX.r3.4HG0335830.1 | 0.371203 | 6.279353 | 2.946863 | 0.004763 | 0.039158 | -2.57739 | Importin-5 | elongation zone | 12 h |
| HORVU.MOREX.r3.2HG0173210.1 | -0.89218 | 5.578206 | -2.81364 | 0.006857 | 0.049459 | -2.57903 | Kinesin-like protein | elongation zone | 12 h |
| HORVU.MOREX.r3.1HG0060570.1 | -0.54139 | 5.889922 | -2.87684 | 0.005775 | 0.044368 | -2.58097 | Mitochondrial zinc maintenance protein 1, mitochondrial | elongation zone | 12 h |
| HORVU.MOREX.r3.2HG0213850.1 | -0.60794 | 5.02608 | -2.85791 | 0.006082 | 0.045953 | -2.58149 | Eukaryotic aspartyl protease family protein, putative | elongation zone | 12 h |
| HORVU.MOREX.r3.6HG0604810.1 | -0.33432 | 6.104373 | -2.91484 | 0.005203 | 0.041594 | -2.58204 | Calcium-dependent protein kinase | elongation zone | 12 h |
| HORVU.MOREX.r3.5HG0523150.1 | -0.25232 | 6.194464 | -2.90325 | 0.005372 | 0.042365 | -2.58352 | Protein phosphatase 2c, putative | elongation zone | 12 h |
| HORVU.MOREX.r3.7HG0635550.1 | -1.22484 | 6.336423 | -2.84675 | 0.006269 | 0.046728 | -2.58539 | Lipoxygenase domain-containing 1 | elongation zone | 12 h |
| HORVU.MOREX.r3.5HG0533660.1 | -0.59365 | 5.124285 | -2.80612 | 0.006998 | 0.049947 | -2.58581 | CTD small phosphatase-like protein | elongation zone | 12 h |
| HORVU.MOREX.r3.7HG0653740.1 | 0.459249 | 7.130211 | 2.995105 | 0.004164 | 0.035961 | -2.58854 | Long-Chain Acyl-CoA Synthetase | elongation zone | 12 h |
| HORVU.MOREX.r3.5HG0519040.1 | -0.34278 | 4.736037 | -2.88155 | 0.005701 | 0.043982 | -2.59029 | Serine/arginine repetitive matrix protein 2, putative isoform 1 | elongation zone | 12 h |
| HORVU.MOREX.r3.1HG0059870.1 | -0.37757 | 4.771051 | -2.85118 | 0.006194 | 0.04639 | -2.59032 | Succinate dehydrogenase subunit 4 | elongation zone | 12 h |
| HORVU.MOREX.r3.3HG0284780.1 | 0.322096 | 5.323435 | 2.897345 | 0.00546 | 0.042794 | -2.59155 | Protein PLASTID MOVEMENT IMPAIRED 1-RELATED 1 | elongation zone | 12 h |
| HORVU.MOREX.r3.3HG0256300.1 | -0.4583 | 7.085102 | -2.91114 | 0.005257 | 0.041825 | -2.59243 | Glutamate--tRNA ligase | elongation zone | 12 h |
| HORVU.MOREX.r3.4HG0381800.1 | -0.26939 | 7.086406 | -2.97125 | 0.004451 | 0.037537 | -2.59265 | AP-2 complex subunit mu | elongation zone | 12 h |
| HORVU.MOREX.r3.1HG0051020.1 | -0.49511 | 5.895904 | -2.86534 | 0.00596 | 0.045444 | -2.59282 | Membrane protein insertase YidC | elongation zone | 12 h |
| HORVU.MOREX.r3.7HG0651030.1 | -0.27013 | 5.034878 | -2.88256 | 0.005686 | 0.043894 | -2.59384 | BRISC and BRCA1-A complex member 1 | elongation zone | 12 h |
| HORVU.MOREX.r3.5HG0473900.1 | 0.393019 | 4.702091 | 2.883491 | 0.005671 | 0.043815 | -2.59505 | Galactose mutarotase-like superfamily protein | elongation zone | 12 h |
| HORVU.MOREX.r3.4HG0392680.1 | 0.79742 | 4.390666 | 2.853928 | 0.006148 | 0.046261 | -2.59508 | Endo-1,4-beta-xylanase | elongation zone | 12 h |
| HORVU.MOREX.r3.4HG0406960.1 | -0.32571 | 5.198023 | -2.85902 | 0.006063 | 0.045928 | -2.59785 | Mitochondrial carrier protein, expressed | elongation zone | 12 h |
| HORVU.MOREX.r3.5HG0420080.1 | 0.421958 | 3.867078 | 2.871716 | 0.005857 | 0.04486 | -2.59871 | Nodulin-like / Major Facilitator Superfamily protein | elongation zone | 12 h |
| HORVU.MOREX.r3.1HG0061210.1 | 0.324619 | 4.858002 | 2.863051 | 0.005997 | 0.04559 | -2.60005 | Replicase polyprotein 1a | elongation zone | 12 h |
| HORVU.MOREX.r3.4HG0335790.2 | -0.59251 | 4.412211 | -2.95252 | 0.004689 | 0.038809 | -2.60516 | Lipoxygenase | elongation zone | 12 h |
| HORVU.MOREX.r3.1HG0045040.1 | -0.38017 | 4.222961 | -2.85115 | 0.006194 | 0.04639 | -2.60606 | Thioredoxin | elongation zone | 12 h |
| HORVU.MOREX.r3.2HG0210680.1 | -0.27247 | 5.95404 | -2.90403 | 0.00536 | 0.042352 | -2.60679 | Anamorsin homolog | elongation zone | 12 h |
| HORVU.MOREX.r3.6HG0613230.1 | -0.30685 | 6.009357 | -2.93669 | 0.004899 | 0.039842 | -2.60827 | Allergen, putative | elongation zone | 12 h |
| HORVU.MOREX.r3.3HG0249700.1 | -0.55325 | 4.807961 | -2.81461 | 0.006839 | 0.049347 | -2.61371 | Protein DEFECTIVE IN MERISTEM SILENCING 3 | elongation zone | 12 h |
| HORVU.MOREX.r3.4HG0358980.1 | 0.502063 | 6.464101 | 3.06231 | 0.003446 | 0.031796 | -2.61464 | Xyloglucan endotransglucosylase/hydrolase | elongation zone | 12 h |
| HORVU.MOREX.r3.7HG0634470.1 | -0.28863 | 7.084322 | -2.95602 | 0.004643 | 0.038637 | -2.61489 | RING/FYVE/PHD zinc finger protein | elongation zone | 12 h |
| HORVU.MOREX.r3.6HG0539990.1 | -0.34178 | 5.586062 | -2.87012 | 0.005882 | 0.04499 | -2.61657 | #N/A | elongation zone | 12 h |
| HORVU.MOREX.r3.6HG0551650.1 | 0.439979 | 2.987415 | 2.833713 | 0.006495 | 0.047717 | -2.61942 | Cyclin family protein | elongation zone | 12 h |
| HORVU.MOREX.r3.6HG0602290.1 | -0.3294 | 4.970789 | -2.86309 | 0.005996 | 0.04559 | -2.6219 | Transmembrane protein-like protein | elongation zone | 12 h |
| HORVU.MOREX.r3.2HG0149120.1 | -0.3032 | 6.262393 | -2.89692 | 0.005466 | 0.042828 | -2.6246 | WW domain-binding protein 11 | elongation zone | 12 h |
| HORVU.MOREX.r3.2HG0205810.1 | 1.230134 | 3.581905 | 2.853061 | 0.006162 | 0.046285 | -2.62562 | Cytochrome P450 family protein, expressed | elongation zone | 12 h |
| HORVU.MOREX.r3.3HG0310320.1 | -0.68899 | 5.514023 | -2.8669 | 0.005934 | 0.045302 | -2.6275 | Glycosyltransferase | elongation zone | 12 h |
| HORVU.MOREX.r3.5HG0493890.2 | 0.441228 | 3.23102 | 2.82785 | 0.006599 | 0.048163 | -2.63183 | Beta-ureidopropionase | elongation zone | 12 h |
| HORVU.MOREX.r3.2HG0173600.1 | 0.617852 | 2.960526 | 2.919574 | 0.005136 | 0.041216 | -2.63271 | Receptor-like kinase | elongation zone | 12 h |
| HORVU.MOREX.r3.3HG0329830.1 | -0.24261 | 5.545794 | -2.88413 | 0.005661 | 0.043788 | -2.63327 | Pectin acetylesterase | elongation zone | 12 h |
| HORVU.MOREX.r3.6HG0622550.1 | -0.28357 | 5.355981 | -2.8857 | 0.005637 | 0.043682 | -2.63666 | Cyclic nucleotide-gated channel | elongation zone | 12 h |
| HORVU.MOREX.r3.6HG0560710.1 | 0.263064 | 7.131181 | 2.953284 | 0.004679 | 0.038805 | -2.63737 | Protein VERNALIZATION INSENSITIVE 3 | elongation zone | 12 h |

**Table S1** Continued.

| ID | logFC | AveExpr | t | P.Value | adj.P.Val | B | gene description | root zone | time point |
|---|---|---|---|---|---|---|---|---|---|
| HORVU.MOREX.r3.5HG0476330.1 | -0.37461 | 6.914052 | -2.94647 | 0.004768 | 0.039181 | -2.63794 | Major facilitator superfamily transporter | elongation zone | 12 h |
| HORVU.MOREX.r3.5HG0458010.1 | 0.274772 | 5.942045 | 2.901748 | 0.005394 | 0.042442 | -2.63915 | Protein kinase, putative | elongation zone | 12 h |
| HORVU.MOREX.r3.3HG0270500.1 | -0.57376 | 4.937967 | -2.82723 | 0.00661 | 0.048184 | -2.63958 | Amino acid permease | elongation zone | 12 h |
| HORVU.MOREX.r3.3HG0291710.1 | -0.47247 | 6.615068 | -2.89109 | 0.005554 | 0.043297 | -2.63981 | Protein PAT1-1-like protein | elongation zone | 12 h |
| HORVU.MOREX.r3.1HG0013820.1 | 0.522677 | 4.142516 | 2.898267 | 0.005446 | 0.042719 | -2.64045 | Protein IQ-DOMAIN 1 | elongation zone | 12 h |
| HORVU.MOREX.r3.4HG0412430.1 | 0.431772 | 4.229658 | 2.839883 | 0.006387 | 0.047281 | -2.64183 | Sugar transporter, putative | elongation zone | 12 h |
| HORVU.MOREX.r3.1HG0033390.4 | -1.71825 | -0.46879 | -2.82453 | 0.006658 | 0.048416 | -2.64421 | Ferrochelatase | elongation zone | 12 h |
| HORVU.MOREX.r3.4HG0416090.1 | -0.50693 | 3.798358 | -2.80985 | 0.006928 | 0.049654 | -2.64451 | 2-oxoglutarate (2OG) and Fe(II)-dependent oxygenase superfamily protein | elongation zone | 12 h |
| HORVU.MOREX.r3.2HG0179540.1 | -0.41053 | 4.341716 | -2.84247 | 0.006342 | 0.047069 | -2.64509 | Ergosterol biosynthetic protein 28 | elongation zone | 12 h |
| HORVU.MOREX.r3.2HG0121570.1 | -0.4111 | 4.65733 | -2.82314 | 0.006684 | 0.048564 | -2.64678 | Protein LTV1 | elongation zone | 12 h |
| HORVU.MOREX.r3.6HG0624190.1 | -0.33097 | 4.552445 | -2.84299 | 0.006333 | 0.047036 | -2.6504 | Elongation factor 1-alpha | elongation zone | 12 h |
| HORVU.MOREX.r3.3HG0236170.1 | -0.35081 | 4.945465 | -2.8683 | 0.005912 | 0.045179 | -2.65197 | Zinc finger protein | elongation zone | 12 h |
| HORVU.MOREX.r3.5HG0505100.1 | -0.39609 | 5.885938 | -2.90045 | 0.005413 | 0.042561 | -2.65374 | Cytochrome b561 and domon domain-containing protein | elongation zone | 12 h |
| HORVU.MOREX.r3.1HG0069940.1 | -0.74165 | 7.781675 | -2.90188 | 0.005392 | 0.042442 | -2.65422 | 60S ribosomal protein L30 | elongation zone | 12 h |
| HORVU.MOREX.r3.1HG0092880.1 | 0.599481 | 4.897051 | 2.825074 | 0.006649 | 0.048379 | -2.65489 | Calcium-dependent protein kinase | elongation zone | 12 h |
| HORVU.MOREX.r3.6HG0612250.1 | -0.38116 | 5.887822 | -2.86294 | 0.005999 | 0.04559 | -2.65733 | Potassium transporter | elongation zone | 12 h |
| HORVU.MOREX.r3.6HG0616240.1 | 0.741873 | -0.52276 | 2.834332 | 0.006484 | 0.047655 | -2.65988 | Endoglucanase | elongation zone | 12 h |
| HORVU.MOREX.r3.3HG0305600.1 | 0.407397 | 6.61338 | 2.972588 | 0.004434 | 0.037428 | -2.66258 | #N/A | elongation zone | 12 h |
| HORVU.MOREX.r3.3HG0280530.1 | 0.532807 | 9.236452 | 3.048382 | 0.003585 | 0.032498 | -2.66404 | Glutamate synthase, putative | elongation zone | 12 h |
| HORVU.MOREX.r3.2HG0192540.1 | 0.318262 | 6.723009 | 2.981399 | 0.004326 | 0.036952 | -2.66625 | Ubiquitin-specific protease family C19-related protein | elongation zone | 12 h |
| HORVU.MOREX.r3.7HG0737960.2 | -0.67232 | 7.555618 | -2.91125 | 0.005255 | 0.041825 | -2.66847 | Glutathione S-transferase | elongation zone | 12 h |
| HORVU.MOREX.r3.5HG0442460.1 | 0.44212 | 3.367563 | 2.917013 | 0.005172 | 0.041427 | -2.66888 | Protein kinase family protein | elongation zone | 12 h |
| HORVU.MOREX.r3.1HG0064080.1 | -0.59532 | 7.790388 | -2.9392 | 0.004865 | 0.039613 | -2.66943 | Peptide-N4-(N-acetyl-beta-glucosaminyl)asparagine amidase A | elongation zone | 12 h |
| HORVU.MOREX.r3.1HG0016950.1 | -0.38144 | 6.051221 | -2.89071 | 0.00556 | 0.043299 | -2.67412 | O-fucosyltransferase family protein | elongation zone | 12 h |
| HORVU.MOREX.r3.2HG0209030.1 | -1.4887 | 0.629763 | -2.81231 | 0.006882 | 0.049549 | -2.67432 | RING finger protein | elongation zone | 12 h |
| HORVU.MOREX.r3.2HG0186650.1 | -0.73704 | 7.283329 | -2.86668 | 0.005938 | 0.045312 | -2.67643 | Non-specific serine/threonine protein kinase | elongation zone | 12 h |
| HORVU.MOREX.r3.1HG0052940.1 | -0.40742 | 6.11793 | -2.86198 | 0.006014 | 0.045676 | -2.67732 | Nuclear control of ATPase protein 2 | elongation zone | 12 h |
| HORVU.MOREX.r3.6HG0614870.1 | 0.713054 | 2.771326 | 2.938055 | 0.00488 | 0.039708 | -2.67862 | Peroxidase | elongation zone | 12 h |
| HORVU.MOREX.r3.5HG0445020.1 | -0.53028 | 5.533594 | -2.82769 | 0.006602 | 0.048163 | -2.6807 | Riboflavin synthase | elongation zone | 12 h |
| HORVU.MOREX.r3.4HG0380710.1 | -0.36533 | 6.055872 | -2.91574 | 0.005191 | 0.04154 | -2.68199 | Calmodulin | elongation zone | 12 h |
| HORVU.MOREX.r3.7HG0671360.1 | -0.25592 | 9.214028 | -3.04269 | 0.003643 | 0.032929 | -2.69129 | Aldehyde dehydrogenase | elongation zone | 12 h |
| HORVU.MOREX.r3.1HG0087980.1 | -0.56358 | 7.176279 | -2.87811 | 0.005755 | 0.044265 | -2.69401 | Ubiquitin | elongation zone | 12 h |
| HORVU.MOREX.r3.3HG0291100.1 | 0.324167 | 8.24123 | 3.020343 | 0.003879 | 0.034278 | -2.69582 | Glutathione S-transferase | elongation zone | 12 h |
| HORVU.MOREX.r3.5HG0494320.1 | -0.43931 | 7.049375 | -2.89477 | 0.005498 | 0.043015 | -2.69701 | Nuclear transport factor 2 | elongation zone | 12 h |
| HORVU.MOREX.r3.4HG0381680.1 | -0.5434 | 4.604183 | -2.83924 | 0.006398 | 0.047329 | -2.69825 | #N/A | elongation zone | 12 h |
| HORVU.MOREX.r3.3HG0269130.1 | 0.985267 | 4.115188 | 2.975572 | 0.004397 | 0.037281 | -2.69994 | Leucine-rich repeat protein kinase family protein | elongation zone | 12 h |
| HORVU.MOREX.r3.3HG0293040.1 | -0.44278 | 5.431414 | -2.81003 | 0.006924 | 0.049654 | -2.70094 | Purple acid phosphatase | elongation zone | 12 h |
| HORVU.MOREX.r3.1HG0026070.2 | -0.56326 | 8.133614 | -2.91026 | 0.005269 | 0.04191 | -2.7054 | Ribosomal protein | elongation zone | 12 h |
| HORVU.MOREX.r3.1HG0083440.1 | 0.349658 | 5.222069 | 2.892055 | 0.00554 | 0.043254 | -2.70594 | Lactation elevated protein 1 | elongation zone | 12 h |
| HORVU.MOREX.r3.3HG0288080.1 | -0.25833 | 6.241682 | -2.86992 | 0.005886 | 0.044997 | -2.70636 | Kinase family protein | elongation zone | 12 h |
| HORVU.MOREX.r3.4HG0386800.1 | 0.281391 | 5.120324 | 2.870571 | 0.005875 | 0.044967 | -2.70797 | Ankyrin repeat family protein, putative, expressed | elongation zone | 12 h |

**Table S1** Continued.

| ID | logFC | AveExpr | t | P.Value | adj.P.Val | B | gene description | root zone | time point |
|---|---|---|---|---|---|---|---|---|---|
| HORVU.MOREX.r3.7HG0709230.1 | -2.49489 | 0.345142 | -2.8412 | 0.006364 | 0.047181 | -2.71064 | 50S ribosomal protein L10 | elongation zone | 12 h |
| HORVU.MOREX.r3.3HG0304660.1 | 0.419427 | 3.80291 | 2.853427 | 0.006156 | 0.046273 | -2.71194 | Ceramide glucosyltransferase, putative | elongation zone | 12 h |
| HORVU.MOREX.r3.2HG0215550.1 | -0.80091 | 7.917138 | -2.88442 | 0.005657 | 0.043769 | -2.71869 | Peroxidase | elongation zone | 12 h |
| HORVU.MOREX.r3.1HG0021880.1 | 2.304736 | 0.747345 | 2.927569 | 0.005024 | 0.040554 | -2.72121 | SKP1-like protein 4 | elongation zone | 12 h |
| HORVU.MOREX.r3.4HG0378000.1 | -0.27214 | 5.073003 | -2.81804 | 0.006776 | 0.048995 | -2.72127 | Uroporphyrinogen decarboxylase | elongation zone | 12 h |
| HORVU.MOREX.r3.6HG0582950.1 | 0.279462 | 6.154329 | 2.895439 | 0.005488 | 0.042969 | -2.72272 | SNARE-interacting protein KEULE | elongation zone | 12 h |
| HORVU.MOREX.r3.2HG0144230.1 | -0.32593 | 6.071895 | -2.86814 | 0.005914 | 0.045182 | -2.72551 | Nucleobase ascorbate transporter | elongation zone | 12 h |
| HORVU.MOREX.r3.4HG0383070.1 | -0.54647 | 5.448589 | -2.81338 | 0.006862 | 0.049476 | -2.72603 | Calmodulin-binding protein | elongation zone | 12 h |
| HORVU.MOREX.r3.3HG0299540.1 | -0.30403 | 6.662261 | -2.90815 | 0.0053 | 0.04204 | -2.72807 | Protein YIPF | elongation zone | 12 h |
| HORVU.MOREX.r3.2HG0124470.1 | -2.27754 | 0.969558 | -2.80512 | 0.007017 | 0.049999 | -2.72816 | Transposon protein, putative, Mutator sub-class | elongation zone | 12 h |
| HORVU.MOREX.r3.3HG0327200.3 | -0.58346 | 5.932631 | -2.84658 | 0.006272 | 0.046732 | -2.72853 | Sulfite exporter TauE/SafE family protein | elongation zone | 12 h |
| HORVU.MOREX.r3.1HG0014430.1 | -0.20412 | 5.638714 | -2.85853 | 0.006071 | 0.045938 | -2.72868 | Beta 1, 3 galactosyltransferase | elongation zone | 12 h |
| HORVU.MOREX.r3.2HG0197760.1 | 0.537422 | 1.898016 | 2.899987 | 0.00542 | 0.042599 | -2.73353 | IQ domain-containing protein | elongation zone | 12 h |
| HORVU.MOREX.r3.5HG0501380.1 | 0.344965 | 6.789726 | 2.90154 | 0.005397 | 0.04245 | -2.73408 | Phospholipase D | elongation zone | 12 h |
| HORVU.MOREX.r3.4HG0337770.1 | 0.572173 | 5.839766 | 2.926786 | 0.005035 | 0.04061 | -2.7364 | Histidine kinase | elongation zone | 12 h |
| HORVU.MOREX.r3.1HG0070430.1 | 0.600286 | 5.294003 | 2.885805 | 0.005635 | 0.043682 | -2.73722 | LURP-one-like protein | elongation zone | 12 h |
| HORVU.MOREX.r3.3HG0279750.1 | 0.327672 | 6.168153 | 2.899828 | 0.005423 | 0.042601 | -2.73904 | Auxin-responsive protein | elongation zone | 12 h |
| HORVU.MOREX.r3.6HG0617240.1 | 0.560498 | 4.621925 | 2.911764 | 0.005248 | 0.041806 | -2.74077 | RNA-binding protein | elongation zone | 12 h |
| HORVU.MOREX.r3.6HG0545460.1 | 0.505992 | 3.011798 | 2.853439 | 0.006156 | 0.046273 | -2.74515 | Transmembrane protein 97 | elongation zone | 12 h |
| HORVU.MOREX.r3.1HG0048010.1 | -0.3448 | 5.141674 | -2.81068 | 0.006912 | 0.049647 | -2.75344 | Membrane steroid-binding protein | elongation zone | 12 h |
| HORVU.MOREX.r3.4HG0359960.1 | 0.3106 | 4.67912 | 2.838306 | 0.006414 | 0.047365 | -2.75624 | Mannose-6-phosphate isomerase | elongation zone | 12 h |
| HORVU.MOREX.r3.1HG0059120.1 | -0.32844 | 5.049749 | -2.8071 | 0.006979 | 0.04985 | -2.7593 | AP-3 complex subunit mu | elongation zone | 12 h |
| HORVU.MOREX.r3.2HG0176290.1 | -0.38019 | 6.030619 | -2.80791 | 0.006964 | 0.049776 | -2.762 | Mitochondrial inner membrane protease ATP23 | elongation zone | 12 h |
| HORVU.MOREX.r3.5HG0491750.1 | -0.31245 | 6.143123 | -2.82846 | 0.006588 | 0.048163 | -2.76221 | Methyl-CpG-binding domain protein | elongation zone | 12 h |
| HORVU.MOREX.r3.4HG0418460.1 | 0.39911 | 1.759537 | 2.845938 | 0.006283 | 0.04678 | -2.76356 | Cysteine protease | elongation zone | 12 h |
| HORVU.MOREX.r3.6HG0547570.1 | 0.394807 | 5.073232 | 2.857348 | 0.006091 | 0.045984 | -2.76634 | LURP-one-like protein | elongation zone | 12 h |
| HORVU.MOREX.r3.2HG0134430.1 | -2.238 | -1.35112 | -2.84008 | 0.006384 | 0.047273 | -2.7671 | Peptidyl-prolyl cis-trans isomerase, putative | elongation zone | 12 h |
| HORVU.MOREX.r3.3HG0279850.1 | 0.479553 | 3.975384 | 2.842154 | 0.006348 | 0.047092 | -2.77077 | mediator of RNA polymerase II transcription subunit | elongation zone | 12 h |
| HORVU.MOREX.r3.4HG0409180.1 | -0.27181 | 6.8545 | -2.88802 | 0.005601 | 0.043562 | -2.78211 | Glutathione S-transferase | elongation zone | 12 h |
| HORVU.MOREX.r3.1HG0076540.1 | -0.35548 | 7.761396 | -2.89369 | 0.005515 | 0.043111 | -2.78422 | Mitochondrial-processing peptidase subunit alpha | elongation zone | 12 h |
| HORVU.MOREX.r3.4HG0404340.1 | -0.36545 | 5.995886 | -2.83628 | 0.00645 | 0.047454 | -2.79075 | Ubiquitin-fold modifier-conjugating enzyme 1 | elongation zone | 12 h |
| HORVU.MOREX.r3.7HG0669240.1 | -0.32209 | 6.048266 | -2.82949 | 0.00657 | 0.048129 | -2.79194 | Microtubule-associated protein 70-2 | elongation zone | 12 h |
| HORVU.MOREX.r3.1HG0003160.1 | 0.5974 | 3.46226 | 2.8495 | 0.006222 | 0.046515 | -2.79511 | Phosphoinositide phosphatase family protein | elongation zone | 12 h |
| HORVU.MOREX.r3.6HG0624460.1 | 2.059731 | -1.05782 | 2.933604 | 0.004941 | 0.040088 | -2.7989 | Receptor-like protein kinase | elongation zone | 12 h |
| HORVU.MOREX.r3.7HG0703780.2 | -0.24845 | 6.247658 | -2.85443 | 0.006139 | 0.046231 | -2.79899 | Succinate dehydrogenase [ubiquinone] iron-sulfur subunit, mitochondrial | elongation zone | 12 h |
| HORVU.MOREX.r3.7HG0634710.1 | -0.22817 | 6.589249 | -2.86294 | 0.005999 | 0.04559 | -2.80191 | Signal peptide peptidase-like protein | elongation zone | 12 h |
| HORVU.MOREX.r3.1HG0054110.1 | 0.461177 | 4.610299 | 2.825414 | 0.006643 | 0.048369 | -2.80351 | Phosphatidylinositol transfer protein | elongation zone | 12 h |
| HORVU.MOREX.r3.7HG0677770.1 | 0.495465 | 5.664941 | 2.911753 | 0.005248 | 0.041806 | -2.80371 | Alkaline alpha-galactosidase seed imbibition protein | elongation zone | 12 h |
| HORVU.MOREX.r3.5HG0506230.1 | 0.524241 | 5.086085 | 2.909257 | 0.005284 | 0.041993 | -2.80584 | Cysteine protease, putative | elongation zone | 12 h |
| HORVU.MOREX.r3.1HG0055560.1 | -0.42947 | 6.140122 | -2.82833 | 0.00659 | 0.048163 | -2.81073 | Metallo-beta-lactamase-like | elongation zone | 12 h |
| HORVU.MOREX.r3.4HG0345610.1 | -0.24167 | 8.214926 | -2.94029 | 0.00485 | 0.039601 | -2.81225 | Protein transport protein Sec24-like family | elongation zone | 12 h |

**Table S1** Continued.

| ID | logFC | AveExpr | t | P.Value | adj.P.Val | B | gene description | root zone | time point |
|---|---|---|---|---|---|---|---|---|---|
| HORVU.MOREX.r3.1HG0047580.1 | 0.505907 | 5.252581 | 2.830803 | 0.006546 | 0.04803 | -2.81453 | acyl-UDP-N-acetylglucosamine O-acyltransferase | elongation zone | 12 h |
| HORVU.MOREX.r3.7HG0733910.1 | -0.38589 | 7.386459 | -2.88672 | 0.005621 | 0.04366 | -2.81867 | Protein LONGIFOLIA 1 | elongation zone | 12 h |
| HORVU.MOREX.r3.2HG0113660.1 | 0.353248 | 7.044507 | 2.872937 | 0.005837 | 0.044752 | -2.82595 | ATP-dependent zinc metalloprotease FtsH | elongation zone | 12 h |
| HORVU.MOREX.r3.1HG0000040.1 | -0.17735 | 6.675243 | -2.865 | 0.005965 | 0.045453 | -2.83816 | Peroxisomal membrane protein PEX14 | elongation zone | 12 h |
| HORVU.MOREX.r3.5HG0534100.1 | -0.88738 | 3.881038 | -2.85148 | 0.006189 | 0.04639 | -2.84257 | Glycine rich protein 3 | elongation zone | 12 h |
| HORVU.MOREX.r3.2HG0173550.1 | -0.66575 | 7.816522 | -2.82966 | 0.006567 | 0.048124 | -2.84673 | Prohibitin | elongation zone | 12 h |
| HORVU.MOREX.r3.5HG0490520.1 | 0.879449 | 1.851159 | 2.829068 | 0.006577 | 0.048163 | -2.84962 | Tonoplast dicarboxylate transporter | elongation zone | 12 h |
| HORVU.MOREX.r3.4HG0385270.1 | -0.46339 | 9.972412 | -2.94412 | 0.004799 | 0.039359 | -2.85778 | Elongation factor 1-alpha | elongation zone | 12 h |
| HORVU.MOREX.r3.4HG0391130.1 | 0.478829 | 5.666862 | 2.82774 | 0.006601 | 0.048163 | -2.85783 | E3 ubiquitin-protein ligase | elongation zone | 12 h |
| HORVU.MOREX.r3.2HG0157170.1 | 0.285975 | 6.50561 | 2.864414 | 0.005975 | 0.045496 | -2.86604 | Dirigent protein | elongation zone | 12 h |
| HORVU.MOREX.r3.3HG0291160.1 | 0.256716 | 5.548618 | 2.834428 | 0.006482 | 0.047655 | -2.86733 | #N/A | elongation zone | 12 h |
| HORVU.MOREX.r3.3HG0299990.1 | 0.507138 | 6.228943 | 2.853946 | 0.006148 | 0.046261 | -2.8705 | Glutathione-S-transferase | elongation zone | 12 h |
| HORVU.MOREX.r3.5HG0420210.1 | -0.3332 | 6.548288 | -2.80996 | 0.006926 | 0.049654 | -2.8744 | Plasma membrane ATPase | elongation zone | 12 h |
| HORVU.MOREX.r3.7HG0667780.1 | 0.582501 | 5.786042 | 2.853699 | 0.006152 | 0.046273 | -2.87689 | U-box domain-containing protein | elongation zone | 12 h |
| HORVU.MOREX.r3.5HG0522630.1 | 0.583768 | 1.481024 | 2.808587 | 0.006951 | 0.049719 | -2.87926 | Blue copper protein | elongation zone | 12 h |
| HORVU.MOREX.r3.2HG0122040.1 | -0.26255 | 6.676824 | -2.83654 | 0.006445 | 0.047445 | -2.88237 | Transmembrane emp24 domain-containing protein | elongation zone | 12 h |
| HORVU.MOREX.r3.4HG0415030.1 | -0.30111 | 11.24828 | -3.031 | 0.003765 | 0.03365 | -2.88548 | Acid phosphatase 1 | elongation zone | 12 h |
| HORVU.MOREX.r3.5HG0534520.1 | -2.30775 | -1.49675 | -2.84445 | 0.006308 | 0.046884 | -2.88707 | tRNA (guanine-N(7)-)-methyltransferase | elongation zone | 12 h |
| HORVU.MOREX.r3.3HG0300020.1 | -2.1725 | -1.49861 | -2.80753 | 0.006971 | 0.049809 | -2.89027 | Apoptosis-inducing factor-like protein A | elongation zone | 12 h |
| HORVU.MOREX.r3.2HG0206650.1 | 0.558893 | 3.016844 | 2.828498 | 0.006587 | 0.048163 | -2.89955 | basic helix-loop-helix (bHLH) DNA-binding superfamily protein | elongation zone | 12 h |
| HORVU.MOREX.r3.4HG0333650.1 | 0.430095 | 5.11965 | 2.812934 | 0.00687 | 0.049502 | -2.91434 | Transmembrane protein 56 | elongation zone | 12 h |
| HORVU.MOREX.r3.6HG0605160.1 | 0.480223 | 1.355711 | 2.812049 | 0.006887 | 0.049549 | -2.91539 | glucan synthase-like 9 | elongation zone | 12 h |
| HORVU.MOREX.r3.2HG0104520.1 | -0.58744 | 9.641823 | -2.93261 | 0.004954 | 0.040151 | -2.92249 | Acetyl-CoA carboxylase | elongation zone | 12 h |
| HORVU.MOREX.r3.2HG0099260.1 | -0.24184 | 6.601234 | -2.81807 | 0.006776 | 0.048995 | -2.92431 | HR-like lesion-inducing protein-related protein | elongation zone | 12 h |
| HORVU.MOREX.r3.5HG0504720.1 | 0.305288 | 6.757338 | 2.809005 | 0.006944 | 0.049702 | -2.92804 | Octicosapeptide/Phox/Bem1p domain-containing protein / tetratricopeptide repe | elongation zone | 12 h |
| HORVU.MOREX.r3.1HG0073070.1 | -0.25929 | 6.545638 | -2.83773 | 0.006425 | 0.047405 | -2.92843 | Lung seven transmembrane receptor family protein | elongation zone | 12 h |
| HORVU.MOREX.r3.2HG0132590.1 | -0.25893 | 6.039953 | -2.82164 | 0.006711 | 0.048693 | -2.93448 | GDP-fucose protein O-fucosyltransferase 2 | elongation zone | 12 h |
| HORVU.MOREX.r3.3HG0289910.1 | -0.35414 | 7.237133 | -2.83817 | 0.006417 | 0.047365 | -2.94785 | Katanin p60 ATPase-containing subunit, putative | elongation zone | 12 h |
| HORVU.MOREX.r3.4HG0390700.1 | -0.20852 | 6.601984 | -2.80942 | 0.006936 | 0.049677 | -2.95789 | ERAD-associated E3 ubiquitin-protein ligase component HRD3A | elongation zone | 12 h |
| HORVU.MOREX.r3.2HG0119650.1 | 0.697641 | 2.120735 | 2.85531 | 0.006125 | 0.046171 | -2.96967 | Peroxidase | elongation zone | 12 h |
| HORVU.MOREX.r3.2HG0125870.1 | -0.51302 | 9.837843 | -2.88099 | 0.00571 | 0.044033 | -2.9698 | 50S ribosomal protein L15 | elongation zone | 12 h |
| HORVU.MOREX.r3.2HG0215280.1 | -0.57203 | 7.631962 | -2.83737 | 0.006431 | 0.047408 | -2.98307 | Peroxidase | elongation zone | 12 h |
| HORVU.MOREX.r3.7HG0676610.1 | 0.37599 | 6.359219 | 2.828539 | 0.006587 | 0.048163 | -2.98615 | Phospholipid-transporting ATPase | elongation zone | 12 h |
| HORVU.MOREX.r3.7HG0750140.1 | 0.376532 | 6.437734 | 2.837141 | 0.006435 | 0.047408 | -2.99151 | NBS-LRR disease resistance protein | elongation zone | 12 h |
| HORVU.MOREX.r3.3HG0241720.1 | -0.19348 | 9.227312 | -2.90815 | 0.0053 | 0.04204 | -2.99209 | Phospholipase D | elongation zone | 12 h |
| HORVU.MOREX.r3.5HG0517240.1 | 0.39148 | 5.444266 | 2.811797 | 0.006892 | 0.049549 | -2.99985 | myosin-binding protein (Protein of unknown function, DUF593) | elongation zone | 12 h |
| HORVU.MOREX.r3.1HG0067500.1 | 0.427503 | 4.190069 | 2.808973 | 0.006944 | 0.049702 | -3.01454 | GDSL esterase/lipase | elongation zone | 12 h |
| HORVU.MOREX.r3.1HG0078770.1 | -0.14486 | 7.962974 | -2.83536 | 0.006466 | 0.047556 | -3.01493 | #N/A | elongation zone | 12 h |
| HORVU.MOREX.r3.4HG0386190.1 | -0.28042 | 8.332385 | -2.81792 | 0.006779 | 0.048995 | -3.02673 | Glutathione-S-transferase | elongation zone | 12 h |
| HORVU.MOREX.r3.3HG0257310.1 | -0.22557 | 7.693219 | -2.82455 | 0.006658 | 0.048416 | -3.02717 | Glucosidase 2 subunit beta | elongation zone | 12 h |
| HORVU.MOREX.r3.2HG0107900.1 | -0.31468 | 7.940345 | -2.80571 | 0.007005 | 0.049984 | -3.03054 | Elongation factor 1-alpha | elongation zone | 12 h |

**Table S1** Continued.

| ID | logFC | AveExpr | t | P.Value | adj.P.Val | B | gene description | root zone | time point |
|---|---|---|---|---|---|---|---|---|---|
| HORVU.MOREX.r3.1HG0056580.1 | 0.345758 | 7.670039 | 2.837036 | 0.006437 | 0.047408 | -3.03913 | Expansin protein | elongation zone | 12 h |
| HORVU.MOREX.r3.3HG0280900.1 | -0.33472 | 8.096043 | -2.80989 | 0.006927 | 0.049654 | -3.04498 | ATP synthase subunit beta | elongation zone | 12 h |
| HORVU.MOREX.r3.4HG0413800.1 | 0.477723 | 7.955857 | 2.934085 | 0.004934 | 0.040075 | -3.04785 | Rhomboid-like protein | elongation zone | 12 h |
| HORVU.MOREX.r3.3HG0254430.1 | 0.26856 | 8.132394 | 2.851324 | 0.006192 | 0.04639 | -3.07286 | Oxidoreductase | elongation zone | 12 h |
| HORVU.MOREX.r3.1HG0024510.1 | 0.40608 | 5.713563 | 2.8051 | 0.007017 | 0.049999 | -3.07851 | Universal stress protein | elongation zone | 12 h |
| HORVU.MOREX.r3.2HG0105020.1 | 0.691857 | 6.347815 | 2.893409 | 0.005519 | 0.043127 | -3.09739 | Aquaporin | elongation zone | 12 h |
| HORVU.MOREX.r3.1HG0003820.1 | 0.349394 | 7.438968 | 2.847067 | 0.006264 | 0.046718 | -3.11774 | Protein DETOXIFICATION | elongation zone | 12 h |
| HORVU.MOREX.r3.7HG0678940.1 | 0.368765 | 6.707895 | 2.838395 | 0.006413 | 0.047365 | -3.13689 | Fasciclin-like arabinogalactan protein | elongation zone | 12 h |
| HORVU.MOREX.r3.2HG0197260.1 | 0.268839 | 7.526184 | 2.82234 | 0.006698 | 0.048636 | -3.14263 | WEB family protein, chloroplastic | elongation zone | 12 h |
| HORVU.MOREX.r3.3HG0228660.1 | 0.337639 | 7.196618 | 2.82339 | 0.006679 | 0.048549 | -3.17854 | Myosin | elongation zone | 12 h |
| HORVU.MOREX.r3.4HG0412730.1 | -0.42217 | 10.67637 | -2.8332 | 0.006504 | 0.047749 | -3.2732 | Purple acid phosphatase | elongation zone | 12 h |
| HORVU.MOREX.r3.3HG0313690.1 | -0.29347 | 9.880246 | -2.80859 | 0.006951 | 0.049719 | -3.34916 | Glutathione S-transferase | elongation zone | 12 h |

up-regulated
down-regulated

**Table S4b** Enriched biological process terms among differentially expressed genes in the elongation zone of gravistimulated wild type roots.

| No. cluster | GO term | Description | cluster 1 | | | cluster 2 | | | cluster 3 | | | cluster 4 | | | cluster 5 | | | cluster 6 | | | cluster 7 | | | cluster 8 | | | cluster 9 | | |
|---|---|---|---|---|---|---|---|---|---|---|---|---|---|---|---|---|---|---|---|---|---|---|---|---|---|---|---|---|---|
| | | | 3 h | 6 h | 12 h | 3 h | 6 h | 12 h | 3 h | 6 h | 12 h | 3 h | 6 h | 12 h | 3 h | 6 h | 12 h | 3 h | 6 h | 12 h | 3 h | 6 h | 12 h | 3 h | 6 h | 12 h | 3 h | 6 h | 12 h |
| cluster 1 | GO:0009664 | plant-type cell wall organization | | | | | | | | | | | | | | | | | | | | | | | | | | | |
| | GO:0006566 | threonine metabolic process | | | | | | | | | | | | | | | | | | | | | | | | | | | |
| | GO:0006544 | glycine metabolic process | | | | | | | | | | | | | | | | | | | | | | | | | | | |
| | GO:1901606 | alpha-amino acid catabolic process | | | | | | | | | | | | | | | | | | | | | | | | | | | |
| | GO:0009063 | cellular amino acid catabolic process | | | | | | | | | | | | | | | | | | | | | | | | | | | |
| | GO:0016311 | dephosphorylation | | | | | | | | | | | | | | | | | | | | | | | | | | | |
| | GO:0006004 | fucose metabolic process | | | | | | | | | | | | | | | | | | | | | | | | | | | |
| | GO:0019320 | hexose catabolic process | | | | | | | | | | | | | | | | | | | | | | | | | | | |
| | GO:0006012 | galactose metabolic process | | | | | | | | | | | | | | | | | | | | | | | | | | | |
| | GO:0007219 | Notch signaling pathway | | | | | | | | | | | | | | | | | | | | | | | | | | | |
| | GO:0010102 | lateral root morphogenesis | | | | | | | | | | | | | | | | | | | | | | | | | | | |
| | GO:0072595 | maintenance of protein localization in organelle | | | | | | | | | | | | | | | | | | | | | | | | | | | |
| | GO:0007020 | microtubule nucleation | | | | | | | | | | | | | | | | | | | | | | | | | | | |
| | GO:0098930 | axonal transport | | | | | | | | | | | | | | | | | | | | | | | | | | | |
| | GO:0045104 | intermediate filament cytoskeleton organization | | | | | | | | | | | | | | | | | | | | | | | | | | | |
| | GO:0047496 | vesicle transport along microtubule | | | | | | | | | | | | | | | | | | | | | | | | | | | |
| | GO:0006457 | protein folding | | | | | | | | | | | | | | | | | | | | | | | | | | | |
| | GO:0016485 | protein processing | | | | | | | | | | | | | | | | | | | | | | | | | | | |
| | GO:0006511 | ubiquitin-dependent protein catabolic process | | | | | | | | | | | | | | | | | | | | | | | | | | | |
| | GO:0030433 | ubiquitin-dependent ERAD pathway | | | | | | | | | | | | | | | | | | | | | | | | | | | |
| | GO:0008299 | isoprenoid biosynthetic process | | | | | | | | | | | | | | | | | | | | | | | | | | | |
| | GO:0006665 | sphingolipid metabolic process | | | | | | | | | | | | | | | | | | | | | | | | | | | |
| | GO:0005975 | carbohydrate metabolic process | | | | | | | | | | | | | | | | | | | | | | | | | | | |
| | GO:1903036 | positive regulation of response to wounding | | | | | | | | | | | | | | | | | | | | | | | | | | | |
| | GO:0015780 | nucleotide-sugar transmembrane transport | | | | | | | | | | | | | | | | | | | | | | | | | | | |
| | GO:0033157 | regulation of intracellular protein transport | | | | | | | | | | | | | | | | | | | | | | | | | | | |
| | GO:0040008 | regulation of growth | | | | | | | | | | | | | | | | | | | | | | | | | | | |
| | GO:0006486 | protein glycosylation | | | | | | | | | | | | | | | | | | | | | | | | | | | |
| | GO:0036065 | fucosylation | | | | | | | | | | | | | | | | | | | | | | | | | | | |
| | GO:0006493 | protein O-linked glycosylation | | | | | | | | | | | | | | | | | | | | | | | | | | | |
| | GO:0006487 | protein N-linked glycosylation | | | | | | | | | | | | | | | | | | | | | | | | | | | |
| | GO:0031529 | ruffle organization | | | | | | | | | | | | | | | | | | | | | | | | | | | |
| | GO:0031103 | axon regeneration | | | | | | | | | | | | | | | | | | | | | | | | | | | |
| | GO:0006082 | organic acid metabolic process | | | | | | | | | | | | | | | | | | | | | | | | | | | |
| | GO:0045489 | pectin biosynthetic process | | | | | | | | | | | | | | | | | | | | | | | | | | | |
| | GO:0070592 | cell wall polysaccharide biosynthetic process | | | | | | | | | | | | | | | | | | | | | | | | | | | |
| | GO:0009734 | auxin-activated signaling pathway | | | | | | | | | | | | | | | | | | | | | | | | | | | |
| | GO:0008286 | insulin receptor signaling pathway | | | | | | | | | | | | | | | | | | | | | | | | | | | |
| | GO:0046686 | response to cadmium ion | | | | | | | | | | | | | | | | | | | | | | | | | | | |
| | GO:0009414 | response to water deprivation | | | | | | | | | | | | | | | | | | | | | | | | | | | |
| | GO:0009409 | response to cold | | | | | | | | | | | | | | | | | | | | | | | | | | | |
| | GO:0010039 | response to iron ion | | | | | | | | | | | | | | | | | | | | | | | | | | | |
| | GO:0045773 | positive regulation of axon extension | | | | | | | | | | | | | | | | | | | | | | | | | | | |
| | GO:0022029 | telencephalon cell migration | | | | | | | | | | | | | | | | | | | | | | | | | | | |
| | GO:0021954 | central nervous system neuron development | | | | | | | | | | | | | | | | | | | | | | | | | | | |
| | GO:0001764 | neuron migration | | | | | | | | | | | | | | | | | | | | | | | | | | | |
| | GO:0021987 | cerebral cortex development | | | | | | | | | | | | | | | | | | | | | | | | | | | |
| | GO:0009311 | oligosaccharide metabolic process | | | | | | | | | | | | | | | | | | | | | | | | | | | |
| | GO:0032418 | lysosome localization | | | | | | | | | | | | | | | | | | | | | | | | | | | |

**Table S4b** Continued.

| No. cluster | GO term | Description | cluster 1 | | | cluster 2 | | | cluster 3 | | | cluster 4 | | | cluster 5 | | | cluster 6 | | | cluster 7 | | | cluster 8 | | | cluster 9 | | |
|---|---|---|---|---|---|---|---|---|---|---|---|---|---|---|---|---|---|---|---|---|---|---|---|---|---|---|---|---|---|
| | | | 3 h | 6 h | 12 h | 3 h | 6 h | 12 h | 3 h | 6 h | 12 h | 3 h | 6 h | 12 h | 3 h | 6 h | 12 h | 3 h | 6 h | 12 h | 3 h | 6 h | 12 h | 3 h | 6 h | 12 h | 3 h | 6 h | 12 h |
| cluster 4 | GO:0048016 | inositol phosphate-mediated signaling | | | | | | | | | | | blue | | | | | | | | | | | | | | | | |
| | GO:0009268 | response to pH | | | | | | | | | | | blue | | | | | | | | | | | | | | | | |
| | GO:0051602 | response to electrical stimulus | | | | | | | | | | | blue | | | | | | | | | | | | | | | | |
| | GO:0006972 | hyperosmotic response | | | | | | | | | | | blue | blue | | | | | | | | | | | | | | | |
| | GO:0006303 | double-strand break repair via nonhomologous end joining | | | | | | | | | | | blue | blue | | | | | | | | | | | | | | | |
| | GO:0009735 | response to cytokinin | | | | | | | | | | | blue | blue | | | | | | | | | | | | | | | |
| | GO:2000028 | regulation of photoperiodism, flowering | | | | | | | | | | | blue | blue | | | | | | | | | | | | | | | |
| | GO:0009100 | glycoprotein metabolic process | | | | | | | | | | | blue | blue | | | | | | | | | | | | | | | |
| | GO:0006506 | GPI anchor biosynthetic process | | | | | | | | | | | blue | blue | | | | | | | | blue | | | | | | | |
| | GO:0006888 | endoplasmic reticulum to Golgi vesicle-mediated transport | | | | | | | | | | | | blue | | | | | | | | | | | | | | | |
| cluster 5 | GO:0006536 | glutamate metabolic process | | | | | | | | | | | | | | red | red | | | | | | | | | | | | |
| | GO:0009084 | glutamine family amino acid biosynthetic process | | | | | | | | | | | | | | red | red | | | | | | | | | | | | |
| | GO:0043650 | dicarboxylic acid biosynthetic process | | | | | | | | | | | | | | red | red | | | | | | | | | | | | |
| | GO:0009741 | response to brassinosteroid | | | | | | | | | | | | | | | red | | | | | | | | | | | | |
| | GO:0010923 | negative regulation of phosphatase activity | | | | | | | | | | | | | | | red | | | | | | | | | | | | |
| | GO:0043666 | regulation of phosphoprotein phosphatase activity | | | | | | | | | | | | | | | red | | | | | | | | | | | | |
| | GO:0006458 | 'de novo' protein folding | | | | | | | | | | | | | | | red | | | | | | | | | | | | |
| | GO:1990542 | mitochondrial transmembrane transport | | | | | | | | | | | | | | | red | | | | | | | | | | | | |
| | GO:0007015 | actin filament organization | | | | | | | | | | | | | | | red | | | | | | | | | | | | |
| | GO:0007018 | microtubule-based movement | | | | | | | | | | | | | | | red | | | | | | | | | | | | |
| | GO:0009773 | photosynthetic electron transport in photosystem I | | | | | | | | | | | | | | | red | | | | | | | | | | | | |
| | GO:0006807 | nitrogen compound metabolic process | | | | | | | | | | | | | | | red | | | | | | | | | | | | |
| | GO:0046654 | tetrahydrofolate biosynthetic process | | | | | | | | | | | | | | | red | | | | | | | | | | | | |
| | GO:0072529 | pyrimidine-containing compound catabolic process | | | | | | | | | | | | | | | red | | | | | | | | | | | | |
| | GO:0008272 | sulfate transport | | | | | | | | | | | | | | | red | | | | | | | | | | | | |
| | GO:0044743 | protein transmembrane import into intracellular organelle | | | | | | | | | | | | | | | red | | | | | | | | | | | | |
| | GO:0006626 | protein targeting to mitochondrion | | | | | | | | | | | | | | | red | | | | | | | | | | | | |
| | GO:0065002 | intracellular protein transmembrane transport | | | | | | | | | | | | | | | red | | | | | | | | | | | | |
| | GO:0036092 | phosphatidylinositol-3-phosphate biosynthetic process | | | | | | | | | | | | | | | red | | | | | | | | | | | | |
| cluster 6 | GO:0006086 | acetyl-CoA biosynthetic process from pyruvate | | | | | | | | | | | | | | | | | red | | | | | | | | | | |
| | GO:0006431 | methionyl-tRNA aminoacylation | | | | | | | | | | | | | | | | | red | | | | | | | | | | |
| | GO:0016024 | CDP-diacylglycerol biosynthetic process | | | | | | | | | | | | | | | | | | blue | | | | | | | | | |
| | GO:0045892 | negative regulation of transcription, DNA-templated | | | | | | | | | | | | | | | | | | blue | | | | | | | | | |
| | GO:0007031 | peroxisome organization | | | | | | | | | | | | | | | | | | blue | | | | | | | | | |
| | GO:0009056 | catabolic process | | | | | | | | | | | | | | | | | | blue | | | | | | | | | |
| | GO:0045292 | mRNA cis splicing, via spliceosome | | | | | | | | | | | | | | | | | | blue | | | | | | | | | |
| | GO:0009740 | gibberellic acid mediated signaling pathway | | | | | | | | | | | | | | | | | | blue | | | | | | | | | |
| | GO:0051781 | positive regulation of cell division | | | | | | | | | | | | | | | | | | blue | | | | | | | | | |
| | GO:0009411 | response to UV | | | | | | | | | | | | | | | | | | blue | | | | | | | | | |
| | GO:0009157 | deoxyribonucleoside monophosphate biosynthetic process | | | | | | | | | | | | | | | | | | blue | | | | | | | | | |
| | GO:0007131 | reciprocal meiotic recombination | | | | | | | | | | | | | | | | | | blue | | | | | | | | | |
| | GO:0019264 | glycine biosynthetic process from serine | | | | | | | | | | | | | | | | | | blue | | | | | | | | | |
| | GO:0048507 | meristem development | | | | | | | | | | | | | | | | | | blue | | | | | | | | | |
| | GO:0034728 | nucleosome organization | | | | | | | | | | | | | | | | | | blue | | | | | | | | | |
| | GO:0006333 | chromatin assembly or disassembly | | | | | | | | | | | | | | | | | | blue | | | | | | | | | |
| cluster 7 | GO:0010015 | root morphogenesis | | | | | | | | | | | | | | | | | | | | blue | | | | | | | |
| | GO:0060560 | developmental growth involved in morphogenesis | | | | | | | | | | | | | | | | | | | | blue | | | | | | | |
| | GO:0048588 | developmental cell growth | | | | | | | | | | | | | | | | | | | | blue | | | | | | | |
| | GO:0048168 | regulation of neuronal synaptic plasticity | | | | | | | | | | | | | | | | | | | | blue | | | | | | | |

up-regulated
down-regulated

**Table S6b** Enriched biological processes terms among genes differentially expressed between wild type and *egt2* (FDR <5%) meristems after a gravistimulation time-course experiment.

| Go term | description | time points 0h | 3h | 6h | 12h |
|---|---|---|---|---|---|
| GO:0031122 | cytoplasmic microtubule organization | up | up | up | up |
| GO:0030865 | cortical cytoskeleton organization | up | up | up | up |
| GO:0010091 | trichome branching | up | up | up | up |
| GO:0009231 | riboflavin biosynthetic process | up | | | |
| GO:0016049 | cell growth | up | up | up | up |
| GO:0005978 | glycogen biosynthetic process | up | | up | up |
| GO:0009832 | plant-type cell wall biogenesis | up | up | up | up |
| GO:0006730 | one-carbon metabolic process | down | down | down | down |
| GO:0006979 | response to oxidative stress | | up | | |
| GO:0042744 | hydrogen peroxide catabolic process | | up | | |
| GO:0006511 | ubiquitin-dependent protein catabolic process | | up | | |
| GO:0016567 | protein ubiquitination | | up | | |
| GO:0006352 | DNA-templated transcription, initiation | | | up | up |
| GO:0006413 | translational initiation | | | up | up |
| GO:0006813 | potassium ion transport | | | | up |
| GO:0030488 | tRNA methylation | | | | up |
| GO:0030001 | metal ion transport | | | | down |
