## Supporting Information Fig.S2 for "ENHANCED GRAVITROPISM 2 coordinates molecular adaptations to gravistimulation in the elongation zone of barley roots"

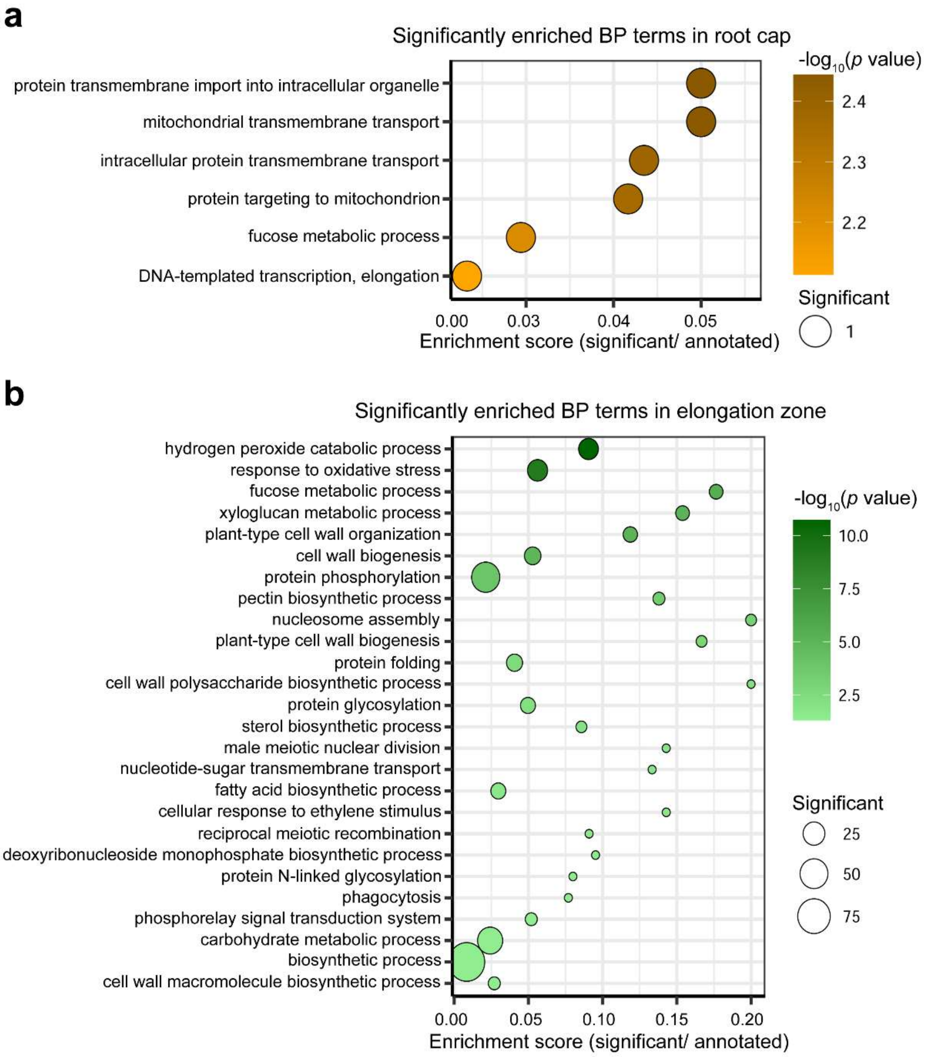


**Fig. S2: Enriched gene ontology (GO) terms for gravity regulated genes that are *EGT2* related.**

**a-b** Significantly enriched GO terms among intersected differentially expressed genes (FDR <5%) in root cap (a) and elongation zone (b).
