## Supporting Information Fig.S3 for "ENHANCED GRAVITROPISM 2 coordinates molecular adaptations to gravistimulation in the elongation zone of barley roots"

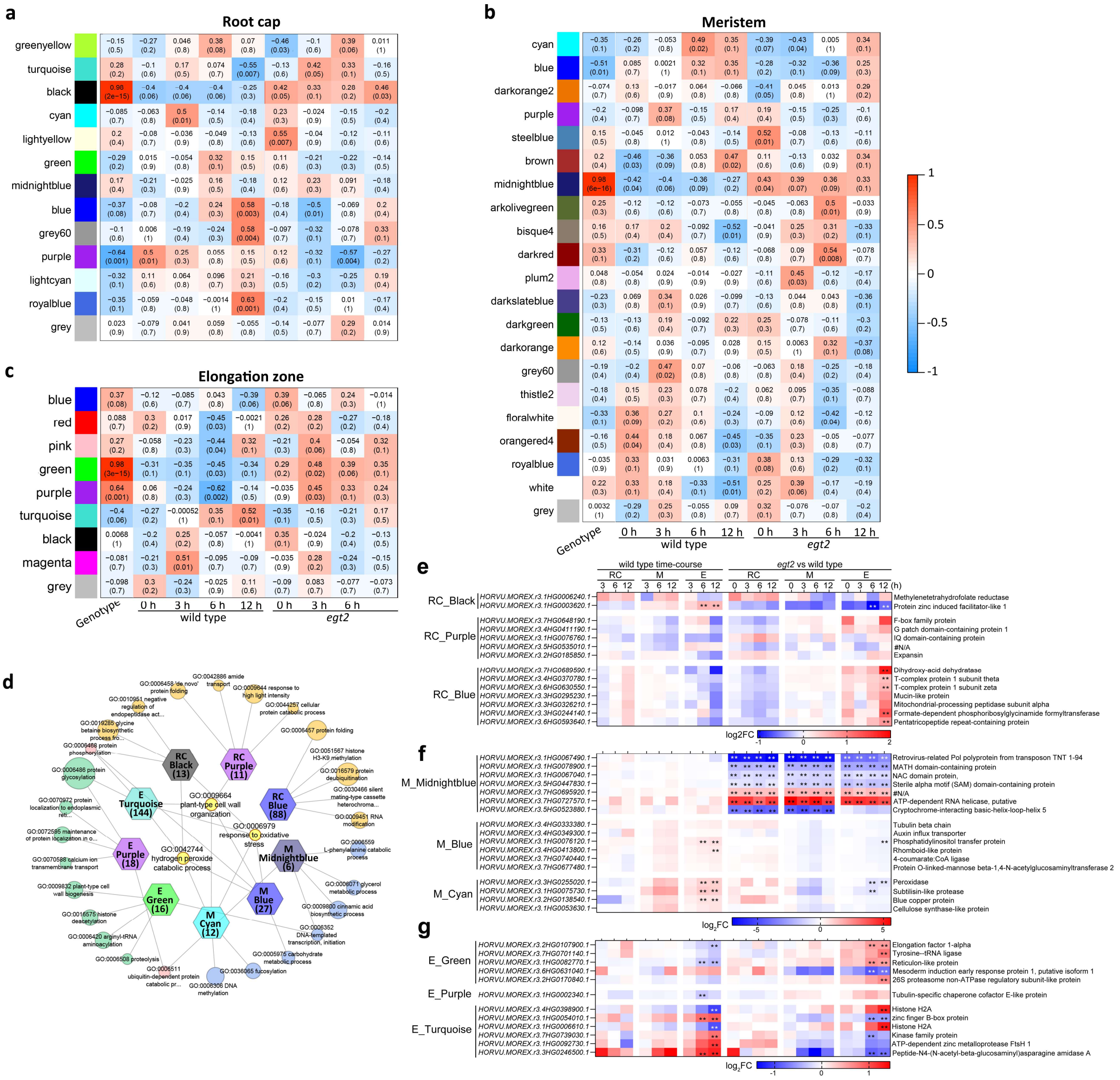

**Fig. S3: Co-expression analyses.**

**a-c** Module-trait relationships in the root cap (a), the meristem (b) and the elongation zone (c). The genotypes (wily type and *egt2*) and time points (0, 3, 6 and 12 h) after gravistimulation are used as traits, each column corresponds to a different trait. Each row corresponds to the characteristic genes of the module. The relationship between the modules and traits is indicated in cell by Pearson correlation coefficients. Numbers in brackets indicate significant levels. Cell color ranges from red (highly positive correlation) to blue (highly negative correction).

**d** Gene ontology analyses of genes in selected significantly correlated modules in the root cap, the meristem and the elongation zone. The five most significantly enriched biological process terms of each module are shown. Hexagonal nodes represent modules, circle nodes represent individual GO terms. Numbers in hexagonal indicate the total number of significantly enriched biological process terms in corresponding module. The size of circle nodes indicates the ratio of the number of genes associated with each GO term to the total number of genes annotated for corresponding term.

**e-g** The expression patterns of hub genes of the black, the purple and the blue modules in the root cap (RC; e), the midnightblue, the blue and cyan modules in the meristem (M; f), and the green, purple and turquoise modules in the elongation zone (E; g) in the WT gravistimulation time-course experiments (wild type time-course) and in the comparisons of *egt2* with wild type (*egt2* vs wild type). **, FDR<5%.
