## Supporting Information Fig.S4 for "ENHANCED GRAVITROPISM 2 coordinates molecular adaptations to gravistimulation in the elongation zone of barley roots"

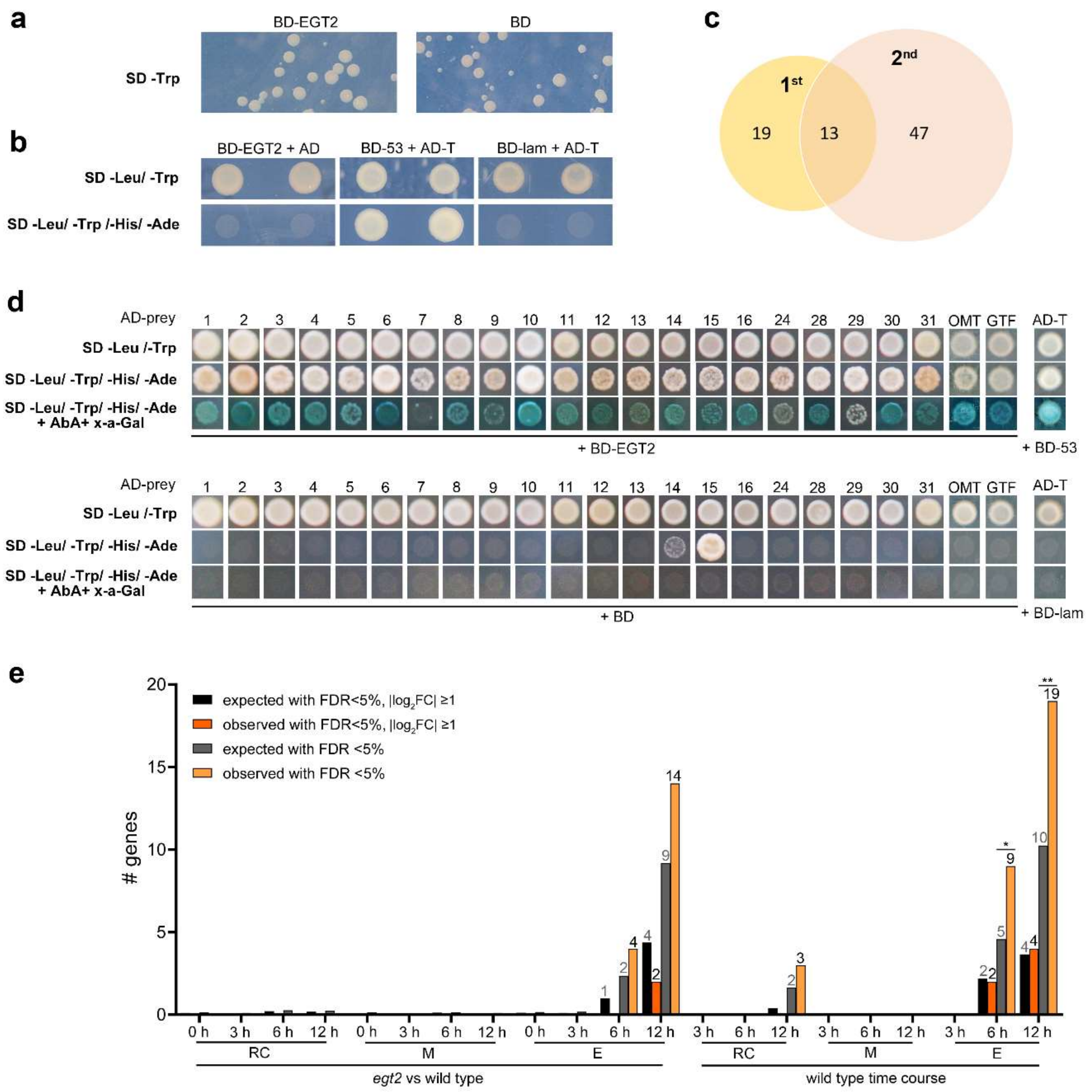


**Fig. S4: Overview of yeast-two-hybrid screening.**

**a**. Toxicity testing of EGT2.

**b**. Autoactivation testing of EGT2.

**c**. Numbers of interaction partners identified by two screens.

**d**. Confirmation of the interaction between EGT2 (BD-EGT2) and the 23 interaction partners (AD-prey) by one-on-one yeast-two-hybrid assays. Co-transformation with each of the AD-preys and pGBKT7 (BD) was used as a negative control. OMT: HORVU.MOREX.r3.3HG0330120.1; GTF: HORVU.MOREX.r3.7HG0736300.1.

**c-d**. Co-transformation of BD-53 and AD-T was shown as positive control, co-transformation of BD-lam and AD-T was shown as negative control,

**e**. Numbers of expected interaction partners and observed interaction candidates in differentially expressed genes. Fisher's exact test, *, *p* <0.05; **, *p* <0.01.
