## Supporting Information Fig.S5 for "ENHANCED GRAVITROPISM 2 coordinates molecular adaptations to gravistimulation in the elongation zone of barley roots"

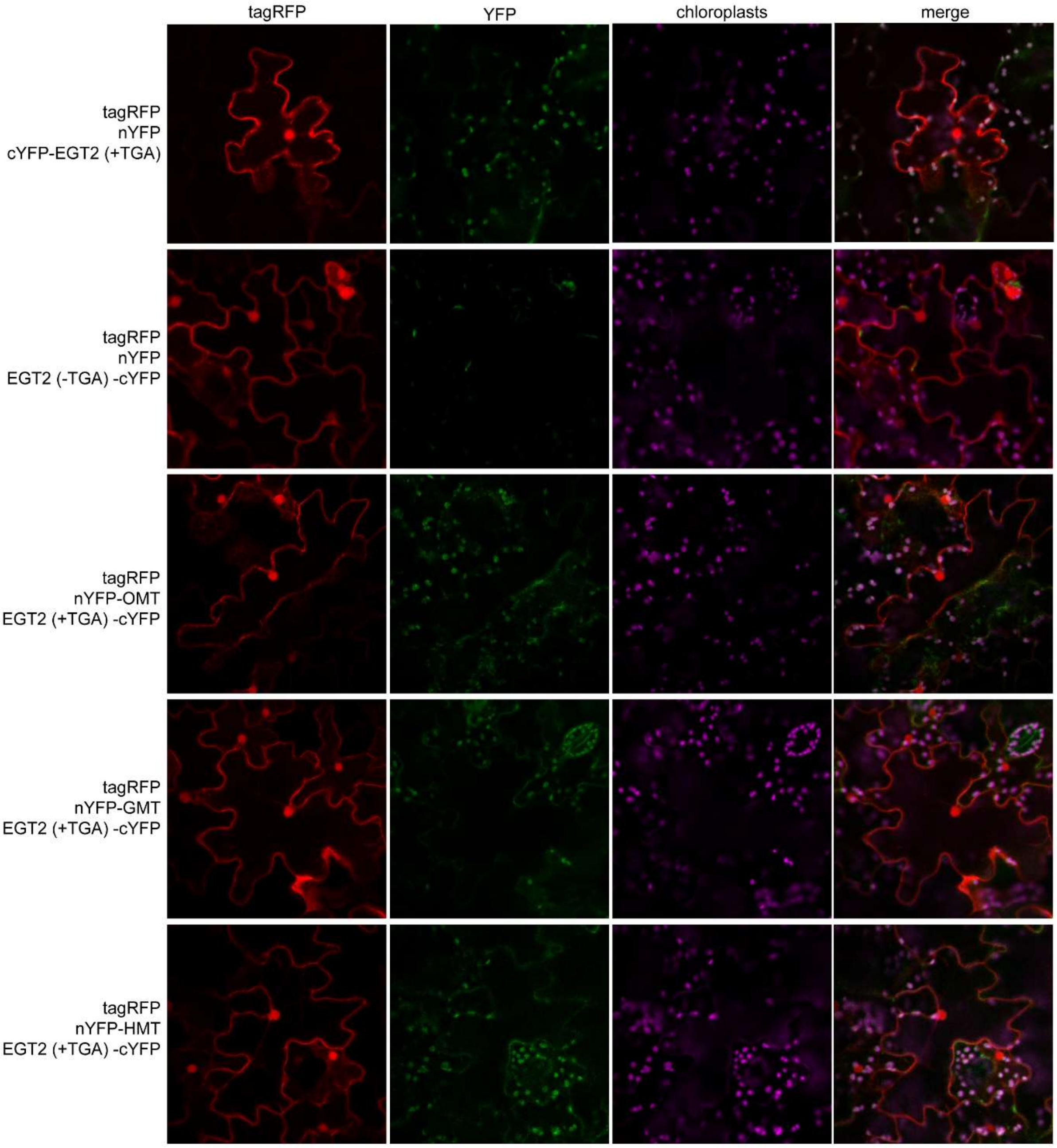


**Fig. S5: Control experiments for bimolecular fluorescence complementation analyses of EGT2 and interaction candidates.**

tagRFP: red fluorescence; YFP: yellow fluorescence; chloroplasts: auto-fluorescence of chloroplasts. nYFP: N-terminal part of YFP; cYFP: C-terminal part of YFP. EGT2 (+TGA): full-length coding sequence of EGT2; EGT2 (-TGA): coding sequence of EGT2 removed stop codon.
