## Supporting Information Fig.S6 for "ENHANCED GRAVITROPISM 2 coordinates molecular adaptations to gravistimulation in the elongation zone of barley roots"

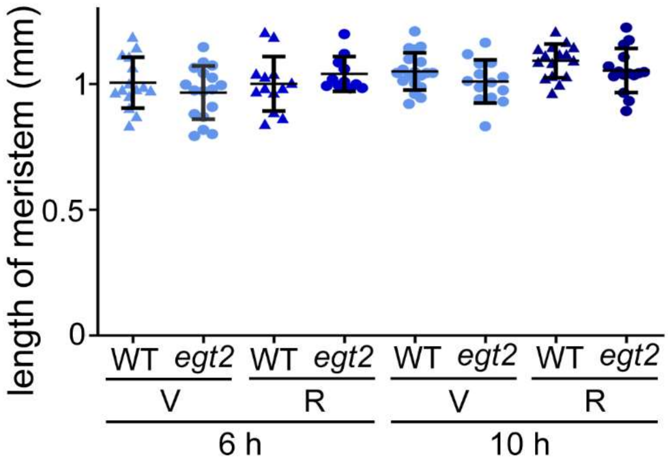


**Fig. S6: Quantification of meristem length of wild type (WT) and *egt2* before and after 6 h or 10 h of gravistimulation.**

(V): vertically grown samples; (R): rotated samples. n = 11-19 per genotype by time point combination.
