## Supporting Information Table 2 for "ENHANCED GRAVITROPISM 2 coordinates molecular adaptations to gravistimulation in the elongation zone of barley roots"

**Table S2** Hierarchical clustering analysis of differentially expressed genes (FDR <5%) in root cap of gravistimulated wild type roots.

| ID | logFC_3 h | logFC_6 h | logFC_12 h | cluster No. |
|---|---|---|---|---|
| HORVU.MOREX.r3.1HG0002330.1 | 0.1707356 | -0.069731 | -0.6646211 | cluster 1 |
| HORVU.MOREX.r3.1HG0008890.1 | 0.1645802 | -0.0130841 | -0.5037172 | cluster 1 |
| HORVU.MOREX.r3.1HG0014220.1 | 0.3493891 | 0.0068183 | -0.8539787 | cluster 1 |
| HORVU.MOREX.r3.1HG0014560.1 | 0.098206 | -0.0579504 | -0.8326012 | cluster 1 |
| HORVU.MOREX.r3.1HG0015430.1 | 0.0140082 | -0.1213651 | -1.5951741 | cluster 1 |
| HORVU.MOREX.r3.1HG0017350.1 | 0.0925823 | -0.0962892 | -0.7454588 | cluster 1 |
| HORVU.MOREX.r3.1HG0019240.1 | -0.465142 | -0.6634696 | -0.9857007 | cluster 1 |
| HORVU.MOREX.r3.1HG0019440.1 | -0.059283 | -0.0980176 | -0.4338449 | cluster 1 |
| HORVU.MOREX.r3.1HG0025520.1 | -0.113161 | -0.2527211 | -1.1252092 | cluster 1 |
| HORVU.MOREX.r3.1HG0027030.1 | -0.201481 | -0.2584751 | -0.7571077 | cluster 1 |
| HORVU.MOREX.r3.1HG0029270.1 | -0.068689 | -0.1856717 | -0.5638445 | cluster 1 |
| HORVU.MOREX.r3.1HG0031820.1 | 0.0602048 | -0.289448 | -0.9447196 | cluster 1 |
| HORVU.MOREX.r3.1HG0036390.1 | 0.0204946 | -0.1146592 | -0.4906721 | cluster 1 |
| HORVU.MOREX.r3.1HG0051880.1 | -0.122715 | -0.280313 | -0.8224489 | cluster 1 |
| HORVU.MOREX.r3.1HG0051890.1 | -0.135185 | -0.2405748 | -0.8744259 | cluster 1 |
| HORVU.MOREX.r3.1HG0053120.1 | 0.1660103 | 0.0451837 | -0.7353026 | cluster 1 |
| HORVU.MOREX.r3.1HG0054960.1 | -0.058411 | -0.1761302 | -0.7343525 | cluster 1 |
| HORVU.MOREX.r3.1HG0055320.1 | 0.1454075 | -0.0141316 | -0.3210414 | cluster 1 |
| HORVU.MOREX.r3.1HG0056360.1 | 0.1753546 | 0.0339342 | -0.4873287 | cluster 1 |
| HORVU.MOREX.r3.1HG0056540.1 | 0.0352941 | 0.0874273 | -0.9744306 | cluster 1 |
| HORVU.MOREX.r3.1HG0057290.1 | 0.0997116 | -0.1008676 | -0.7499372 | cluster 1 |
| HORVU.MOREX.r3.1HG0057610.1 | -0.17338 | -0.2743248 | -0.5220569 | cluster 1 |
| HORVU.MOREX.r3.1HG0057800.1 | 0.2064075 | -0.1769001 | -1.342247 | cluster 1 |
| HORVU.MOREX.r3.1HG0058170.1 | 0.0526902 | -0.0644834 | -0.4953391 | cluster 1 |
| HORVU.MOREX.r3.1HG0059150.1 | 0.1383591 | 0.0409054 | -0.7585698 | cluster 1 |
| HORVU.MOREX.r3.1HG0062030.1 | 0.3329477 | -0.3390041 | -1.455675 | cluster 1 |
| HORVU.MOREX.r3.1HG0064310.1 | 0.0167471 | -0.0834389 | -0.6617663 | cluster 1 |
| HORVU.MOREX.r3.1HG0065120.1 | -0.042207 | -0.1003168 | -0.5499375 | cluster 1 |
| HORVU.MOREX.r3.1HG0069010.1 | -0.107712 | -0.3083214 | -0.8324017 | cluster 1 |
| HORVU.MOREX.r3.1HG0070670.1 | 0.021674 | -0.1568637 | -0.6412713 | cluster 1 |
| HORVU.MOREX.r3.1HG0071320.1 | 0.0857173 | 0.0503672 | -1.037841 | cluster 1 |
| HORVU.MOREX.r3.1HG0072100.1 | 0.2686935 | 0.0831744 | -0.7749167 | cluster 1 |
| HORVU.MOREX.r3.1HG0073140.1 | -0.046214 | -0.2538513 | -0.8635086 | cluster 1 |
| HORVU.MOREX.r3.1HG0075220.1 | 0.2967808 | 0.1601418 | -1.0192773 | cluster 1 |
| HORVU.MOREX.r3.1HG0077230.1 | 0.1899281 | -0.0093486 | -0.789304 | cluster 1 |
| HORVU.MOREX.r3.1HG0077710.1 | 0.1542422 | 0.1803493 | -0.7503547 | cluster 1 |
| HORVU.MOREX.r3.1HG0080260.1 | -0.245798 | -0.2291487 | -0.5082043 | cluster 1 |
| HORVU.MOREX.r3.1HG0081120.1 | -0.272373 | -0.1142109 | -1.4459268 | cluster 1 |
| HORVU.MOREX.r3.1HG0081860.1 | 0.1326211 | -0.11948 | -0.8788293 | cluster 1 |
| HORVU.MOREX.r3.1HG0082670.1 | -0.115995 | -0.3154451 | -0.7413344 | cluster 1 |
| HORVU.MOREX.r3.1HG0083840.1 | -0.108545 | -0.0497991 | -0.4975227 | cluster 1 |
| HORVU.MOREX.r3.1HG0085780.1 | 0.227707 | 0.0763213 | -1.0106562 | cluster 1 |
| HORVU.MOREX.r3.1HG0089180.1 | 0.1740548 | 0.0111709 | -0.6094135 | cluster 1 |
| HORVU.MOREX.r3.1HG0093750.1 | -0.135597 | -0.0185006 | -1.0255918 | cluster 1 |
| HORVU.MOREX.r3.1HG0093880.1 | 0.3636581 | 0.1300544 | -0.77277 | cluster 1 |
| HORVU.MOREX.r3.2HG0096100.1 | 0.0229233 | -0.0662696 | -0.8618271 | cluster 1 |
| HORVU.MOREX.r3.2HG0097600.1 | -0.054917 | -0.1509645 | -0.5279722 | cluster 1 |

**Table S2** Continued.

| ID | logFC_3 h | logFC_6 h | logFC_12 h | cluster No. |
|---|---|---|---|---|
| HORVU.MOREX.r3.2HG0099750.1 | 0.182292 | -0.0954893 | -0.5572858 | cluster 1 |
| HORVU.MOREX.r3.2HG0099980.1 | -0.236216 | -0.2842068 | -0.7613671 | cluster 1 |
| HORVU.MOREX.r3.2HG0100060.1 | 0.0682534 | -0.1090605 | -1.0421373 | cluster 1 |
| HORVU.MOREX.r3.2HG0100270.2 | 0.1435812 | 0.0823961 | -0.5912385 | cluster 1 |
| HORVU.MOREX.r3.2HG0104110.1 | 0.0629003 | 0.0021257 | -0.9242198 | cluster 1 |
| HORVU.MOREX.r3.2HG0109400.1 | 0.1065021 | -0.1021695 | -1.2689128 | cluster 1 |
| HORVU.MOREX.r3.2HG0113410.1 | 0.1233282 | -0.0049988 | -0.5250859 | cluster 1 |
| HORVU.MOREX.r3.2HG0113590.1 | -0.056695 | -0.0157784 | -0.5608697 | cluster 1 |
| HORVU.MOREX.r3.2HG0113860.1 | 0.1942076 | -0.0043177 | -1.2067332 | cluster 1 |
| HORVU.MOREX.r3.2HG0119150.1 | 0.0459465 | 0.0763396 | -0.5504699 | cluster 1 |
| HORVU.MOREX.r3.2HG0119920.1 | 0.1578493 | 0.0488629 | -0.671108 | cluster 1 |
| HORVU.MOREX.r3.2HG0121380.1 | -0.07747 | -0.2040315 | -0.6364525 | cluster 1 |
| HORVU.MOREX.r3.2HG0122840.1 | 0.1800344 | 0.1011832 | -0.5711731 | cluster 1 |
| HORVU.MOREX.r3.2HG0124110.1 | -0.124875 | -0.0871535 | -0.7312259 | cluster 1 |
| HORVU.MOREX.r3.2HG0125600.1 | -0.123034 | -0.3901724 | -1.282441 | cluster 1 |
| HORVU.MOREX.r3.2HG0127370.1 | 0.1575099 | 0.2037384 | -0.6091735 | cluster 1 |
| HORVU.MOREX.r3.2HG0128270.1 | 0.1627538 | 0.1000413 | -0.776112 | cluster 1 |
| HORVU.MOREX.r3.2HG0128490.1 | -0.075366 | -0.1253905 | -0.4289783 | cluster 1 |
| HORVU.MOREX.r3.2HG0130760.1 | -0.043515 | -0.1034422 | -0.6913018 | cluster 1 |
| HORVU.MOREX.r3.2HG0131330.1 | 0.1699821 | 0.0290188 | -0.6502478 | cluster 1 |
| HORVU.MOREX.r3.2HG0133060.1 | 0.355961 | 0.0803982 | -0.870611 | cluster 1 |
| HORVU.MOREX.r3.2HG0133930.1 | -0.936341 | -0.0392541 | -2.1038791 | cluster 1 |
| HORVU.MOREX.r3.2HG0135590.1 | -0.244832 | -0.0869697 | -0.6340419 | cluster 1 |
| HORVU.MOREX.r3.2HG0138270.1 | -0.169465 | -0.230731 | -0.6614251 | cluster 1 |
| HORVU.MOREX.r3.2HG0147190.1 | 0.1119502 | -0.0668936 | -0.7262734 | cluster 1 |
| HORVU.MOREX.r3.2HG0154110.1 | 0.2194787 | 0.0261628 | -1.033996 | cluster 1 |
| HORVU.MOREX.r3.2HG0163200.1 | 0.2192212 | 0.1676308 | -0.5199979 | cluster 1 |
| HORVU.MOREX.r3.2HG0164440.1 | 0.2034692 | 0.3096843 | -0.6251427 | cluster 1 |
| HORVU.MOREX.r3.2HG0165290.2 | -0.095882 | 0.0993993 | -0.7885599 | cluster 1 |
| HORVU.MOREX.r3.2HG0165780.1 | 0.0892943 | 0.1883986 | -0.4339442 | cluster 1 |
| HORVU.MOREX.r3.2HG0168300.1 | 0.054616 | -0.1043369 | -1.1604944 | cluster 1 |
| HORVU.MOREX.r3.2HG0168640.1 | -0.114701 | -0.0881309 | -0.4952482 | cluster 1 |
| HORVU.MOREX.r3.2HG0169000.1 | -0.392723 | -0.769097 | -1.4321648 | cluster 1 |
| HORVU.MOREX.r3.2HG0169420.1 | -0.027819 | -0.0138016 | -0.8419302 | cluster 1 |
| HORVU.MOREX.r3.2HG0172180.1 | -0.121844 | -0.3012642 | -0.7902189 | cluster 1 |
| HORVU.MOREX.r3.2HG0173440.1 | -0.014501 | -0.1274453 | -0.851014 | cluster 1 |
| HORVU.MOREX.r3.2HG0173750.1 | 0.0613164 | -0.0597776 | -0.5074225 | cluster 1 |
| HORVU.MOREX.r3.2HG0174560.1 | -0.114083 | -0.1786777 | -0.5089115 | cluster 1 |
| HORVU.MOREX.r3.2HG0177780.1 | 0.0285171 | -0.0895278 | -0.4935162 | cluster 1 |
| HORVU.MOREX.r3.2HG0180040.2 | 0.0533812 | -0.0608599 | -1.1036712 | cluster 1 |
| HORVU.MOREX.r3.2HG0181020.1 | 0.2204522 | 0.0198243 | -0.6008209 | cluster 1 |
| HORVU.MOREX.r3.2HG0184350.1 | -0.038935 | -0.0857675 | -0.5818051 | cluster 1 |
| HORVU.MOREX.r3.2HG0185660.1 | -0.08651 | -0.1538465 | -0.4095365 | cluster 1 |
| HORVU.MOREX.r3.2HG0186670.1 | 0.0442886 | 0.027498 | -0.6764735 | cluster 1 |
| HORVU.MOREX.r3.2HG0187170.1 | -0.018128 | -0.0817307 | -0.4488681 | cluster 1 |
| HORVU.MOREX.r3.2HG0187980.1 | 0.3392904 | 0.1406879 | -0.5813126 | cluster 1 |
| HORVU.MOREX.r3.2HG0188070.1 | 0.2777649 | 0.1336551 | -0.8322951 | cluster 1 |

**Table S2** Continued.

| ID | logFC_3 h | logFC_6 h | logFC_12 h | cluster No. |
|---|---|---|---|---|
| HORVU.MOREX.r3.2HG0189190.1 | 0.2117247 | 0.0348889 | -0.564115 | cluster 1 |
| HORVU.MOREX.r3.2HG0189340.1 | -0.157698 | -0.2219058 | -0.9130796 | cluster 1 |
| HORVU.MOREX.r3.2HG0191970.1 | 0.0791868 | -0.1581829 | -0.8752616 | cluster 1 |
| HORVU.MOREX.r3.2HG0192860.1 | 0.1482787 | -0.0736532 | -0.5624417 | cluster 1 |
| HORVU.MOREX.r3.2HG0192920.1 | 0.2505769 | -0.0475889 | -0.8141112 | cluster 1 |
| HORVU.MOREX.r3.2HG0192960.1 | 0.1026076 | 0.1015388 | -0.6391422 | cluster 1 |
| HORVU.MOREX.r3.2HG0194600.1 | -0.566806 | -0.5078339 | -0.7637254 | cluster 1 |
| HORVU.MOREX.r3.2HG0194790.1 | 0.1559992 | 0.0290132 | -0.5149481 | cluster 1 |
| HORVU.MOREX.r3.2HG0195580.1 | 0.2191461 | 0.1882456 | -0.6067809 | cluster 1 |
| HORVU.MOREX.r3.2HG0197550.1 | 0.3083522 | 0.1903185 | -0.8592855 | cluster 1 |
| HORVU.MOREX.r3.2HG0201420.1 | -0.138068 | -0.3993381 | -1.0927887 | cluster 1 |
| HORVU.MOREX.r3.2HG0202110.1 | 0.1008244 | 0.0033848 | -0.4397894 | cluster 1 |
| HORVU.MOREX.r3.2HG0203390.1 | -0.111121 | -0.4001883 | -1.0824523 | cluster 1 |
| HORVU.MOREX.r3.2HG0205250.1 | 0.1502304 | 0.1656253 | -0.7819281 | cluster 1 |
| HORVU.MOREX.r3.2HG0207160.1 | 0.2676942 | 0.2612654 | -0.7243435 | cluster 1 |
| HORVU.MOREX.r3.2HG0211680.1 | -0.070159 | -0.347567 | -1.0188098 | cluster 1 |
| HORVU.MOREX.r3.2HG0213480.1 | 0.1684082 | 0.1369867 | -0.4820726 | cluster 1 |
| HORVU.MOREX.r3.2HG0214820.1 | 0.0216719 | -0.3691005 | -0.9907546 | cluster 1 |
| HORVU.MOREX.r3.3HG0220830.1 | 0.2452903 | 0.0003142 | -1.4049309 | cluster 1 |
| HORVU.MOREX.r3.3HG0227860.1 | 0.0905777 | -0.0332528 | -0.3865948 | cluster 1 |
| HORVU.MOREX.r3.3HG0227890.1 | 0.0198392 | -0.0569362 | -1.181551 | cluster 1 |
| HORVU.MOREX.r3.3HG0228600.1 | 0.0683067 | -0.1947412 | -0.8040683 | cluster 1 |
| HORVU.MOREX.r3.3HG0231010.1 | -0.030001 | -0.0410105 | -0.4832806 | cluster 1 |
| HORVU.MOREX.r3.3HG0231330.1 | -0.065927 | -0.3270764 | -0.8240308 | cluster 1 |
| HORVU.MOREX.r3.3HG0231460.1 | 0.168657 | 0.1539204 | -0.7719347 | cluster 1 |
| HORVU.MOREX.r3.3HG0232840.1 | 0.1496139 | 0.1093986 | -0.4678902 | cluster 1 |
| HORVU.MOREX.r3.3HG0234880.1 | 0.1085436 | 0.0271935 | -0.7876179 | cluster 1 |
| HORVU.MOREX.r3.3HG0234890.1 | 0.0597158 | 0.1404151 | -0.7690663 | cluster 1 |
| HORVU.MOREX.r3.3HG0235800.1 | 0.3764253 | -0.5083754 | -2.3271368 | cluster 1 |
| HORVU.MOREX.r3.3HG0236350.1 | -0.514395 | -0.2139384 | -0.8716992 | cluster 1 |
| HORVU.MOREX.r3.3HG0242790.1 | 0.2172794 | -0.0040146 | -0.4991882 | cluster 1 |
| HORVU.MOREX.r3.3HG0244080.1 | 0.0143144 | -0.1989374 | -0.9640661 | cluster 1 |
| HORVU.MOREX.r3.3HG0245010.1 | -0.009652 | -0.119163 | -0.6249435 | cluster 1 |
| HORVU.MOREX.r3.3HG0249710.1 | 0.0964757 | -0.1540475 | -0.680477 | cluster 1 |
| HORVU.MOREX.r3.3HG0250330.1 | 0.0685641 | -0.0833517 | -0.5594949 | cluster 1 |
| HORVU.MOREX.r3.3HG0251700.1 | -0.007638 | -0.2978602 | -0.9078617 | cluster 1 |
| HORVU.MOREX.r3.3HG0251900.1 | -0.021701 | -0.1306439 | -0.5818368 | cluster 1 |
| HORVU.MOREX.r3.3HG0252100.1 | 0.0895496 | -0.0647937 | -0.7727761 | cluster 1 |
| HORVU.MOREX.r3.3HG0252270.1 | 0.1472279 | -0.0650709 | -0.9676407 | cluster 1 |
| HORVU.MOREX.r3.3HG0253860.1 | -0.176805 | -0.6068891 | -1.4092203 | cluster 1 |
| HORVU.MOREX.r3.3HG0256160.1 | 0.1818661 | 0.0931427 | -0.5030953 | cluster 1 |
| HORVU.MOREX.r3.3HG0256430.1 | 0.1617285 | -0.0372638 | -0.6559322 | cluster 1 |
| HORVU.MOREX.r3.3HG0256910.1 | 0.0327792 | -0.0574169 | -0.8204516 | cluster 1 |
| HORVU.MOREX.r3.3HG0266950.1 | -0.129288 | 0.130242 | -0.6905455 | cluster 1 |
| HORVU.MOREX.r3.3HG0271380.1 | 0.1804017 | 0.129765 | -0.545123 | cluster 1 |
| HORVU.MOREX.r3.3HG0271550.1 | 0.1986398 | -0.070624 | -0.5219619 | cluster 1 |
| HORVU.MOREX.r3.3HG0273560.1 | -0.357839 | -0.5254332 | -1.300763 | cluster 1 |

**Table S2** Continued.

| ID | logFC_3 h | logFC_6 h | logFC_12 h | cluster No. |
|---|---|---|---|---|
| HORVU.MOREX.r3.3HG0273730.1 | -0.164858 | -0.1678612 | -0.6509035 | cluster 1 |
| HORVU.MOREX.r3.3HG0275260.1 | 0.0111266 | -0.1864652 | -0.7524535 | cluster 1 |
| HORVU.MOREX.r3.3HG0275450.1 | 0.0212448 | 0.2655736 | -0.9271112 | cluster 1 |
| HORVU.MOREX.r3.3HG0275500.1 | -0.200904 | -0.623191 | -1.6564576 | cluster 1 |
| HORVU.MOREX.r3.3HG0276000.2 | 1.1018298 | 1.8100643 | -0.6585563 | cluster 1 |
| HORVU.MOREX.r3.3HG0277400.1 | 0.3540908 | 0.1408818 | -0.6071427 | cluster 1 |
| HORVU.MOREX.r3.3HG0281000.1 | -0.005218 | 0.0368879 | -1.1362508 | cluster 1 |
| HORVU.MOREX.r3.3HG0281780.1 | -0.207269 | -0.2383308 | -0.6641343 | cluster 1 |
| HORVU.MOREX.r3.3HG0284890.1 | 0.1629977 | 0.2150009 | -0.6938056 | cluster 1 |
| HORVU.MOREX.r3.3HG0286640.1 | -0.106466 | -0.2502943 | -0.921725 | cluster 1 |
| HORVU.MOREX.r3.3HG0288830.1 | 0.0091573 | -0.0170967 | -0.3301356 | cluster 1 |
| HORVU.MOREX.r3.3HG0290130.1 | 0.0707521 | -0.0632943 | -0.76123 | cluster 1 |
| HORVU.MOREX.r3.3HG0290180.1 | 0.0603195 | -0.1700186 | -0.6116577 | cluster 1 |
| HORVU.MOREX.r3.3HG0291590.1 | -0.088436 | -0.2753589 | -0.6288763 | cluster 1 |
| HORVU.MOREX.r3.3HG0291680.1 | 0.1814673 | 0.0711571 | -0.9785149 | cluster 1 |
| HORVU.MOREX.r3.3HG0293070.1 | 0.1025929 | -0.1167899 | -0.6356689 | cluster 1 |
| HORVU.MOREX.r3.3HG0293570.1 | -0.354902 | -0.2515102 | -2.9257568 | cluster 1 |
| HORVU.MOREX.r3.3HG0295550.1 | 0.1507494 | 0.0980024 | -0.5186528 | cluster 1 |
| HORVU.MOREX.r3.3HG0295840.1 | 0.0508498 | 0.0271712 | -0.5843366 | cluster 1 |
| HORVU.MOREX.r3.3HG0296420.1 | 0.0569306 | -0.0195863 | -0.463752 | cluster 1 |
| HORVU.MOREX.r3.3HG0299100.1 | 0.0169666 | -0.1209807 | -0.5434559 | cluster 1 |
| HORVU.MOREX.r3.3HG0300080.1 | 0.2541824 | 0.0574149 | -0.7490972 | cluster 1 |
| HORVU.MOREX.r3.3HG0300580.1 | 0.00154 | -0.2233061 | -1.0561484 | cluster 1 |
| HORVU.MOREX.r3.3HG0303190.1 | 0.1717346 | 0.141804 | -0.4057797 | cluster 1 |
| HORVU.MOREX.r3.3HG0303670.1 | 0.1681824 | -0.2322876 | -0.8676161 | cluster 1 |
| HORVU.MOREX.r3.3HG0305540.1 | -0.098984 | -0.6168426 | -1.3167503 | cluster 1 |
| HORVU.MOREX.r3.3HG0306820.1 | 0.2426683 | 0.0673865 | -0.5940755 | cluster 1 |
| HORVU.MOREX.r3.3HG0308250.1 | 0.0358422 | -0.022312 | -0.5375334 | cluster 1 |
| HORVU.MOREX.r3.3HG0311480.1 | 0.0922847 | 0.0494435 | -0.8640716 | cluster 1 |
| HORVU.MOREX.r3.3HG0315300.1 | 0.0648248 | -0.0581596 | -0.8686528 | cluster 1 |
| HORVU.MOREX.r3.3HG0320710.1 | 0.0459291 | -0.106247 | -0.5763273 | cluster 1 |
| HORVU.MOREX.r3.3HG0323410.1 | 0.3461359 | -0.0157362 | -1.0490819 | cluster 1 |
| HORVU.MOREX.r3.3HG0328340.1 | -0.080608 | -0.3457436 | -0.8072279 | cluster 1 |
| HORVU.MOREX.r3.3HG0329000.1 | 0.1757403 | 0.123947 | -0.5454783 | cluster 1 |
| HORVU.MOREX.r3.4HG0332530.1 | -0.040955 | -0.1235787 | -0.5665792 | cluster 1 |
| HORVU.MOREX.r3.4HG0336330.1 | 0.049546 | 0.0393657 | -0.8291267 | cluster 1 |
| HORVU.MOREX.r3.4HG0338810.1 | 0.2379603 | 0.1118242 | -0.5885998 | cluster 1 |
| HORVU.MOREX.r3.4HG0340420.1 | 0.2455155 | -0.0348712 | -0.4823853 | cluster 1 |
| HORVU.MOREX.r3.4HG0342860.1 | -0.045176 | -0.0185048 | -0.835231 | cluster 1 |
| HORVU.MOREX.r3.4HG0343380.1 | -0.03962 | -0.1308218 | -0.4306572 | cluster 1 |
| HORVU.MOREX.r3.4HG0347330.1 | 0.1528549 | 0.0584248 | -0.5585683 | cluster 1 |
| HORVU.MOREX.r3.4HG0348850.1 | 0.2404947 | 0.1267945 | -0.6904592 | cluster 1 |
| HORVU.MOREX.r3.4HG0352420.1 | -0.031264 | -0.2470611 | -0.641372 | cluster 1 |
| HORVU.MOREX.r3.4HG0353950.1 | -0.15119 | -0.1458492 | -1.6328211 | cluster 1 |
| HORVU.MOREX.r3.4HG0356780.1 | 0.0014687 | 0.0269827 | -0.5457976 | cluster 1 |
| HORVU.MOREX.r3.4HG0365970.1 | -0.1093 | -0.0976246 | -0.4987654 | cluster 1 |
| HORVU.MOREX.r3.4HG0378750.1 | 0.0617481 | -6.277E-05 | -0.5717213 | cluster 1 |

**Table S2** Continued.

| ID | logFC_3 h | logFC_6 h | logFC_12 h | cluster No. |
|---|---|---|---|---|
| HORVU.MOREX.r3.4HG0378900.1 | 0.196064 | 0.0852259 | -0.5236892 | cluster 1 |
| HORVU.MOREX.r3.4HG0381750.1 | -0.117547 | -0.1359577 | -0.6095192 | cluster 1 |
| HORVU.MOREX.r3.4HG0381820.1 | 0.1308354 | 0.0013819 | -0.6168217 | cluster 1 |
| HORVU.MOREX.r3.4HG0382640.1 | 0.2568095 | 0.2242829 | -1.3880395 | cluster 1 |
| HORVU.MOREX.r3.4HG0384300.1 | -0.001681 | -0.0583377 | -1.2011882 | cluster 1 |
| HORVU.MOREX.r3.4HG0384590.1 | -0.332835 | -0.3626826 | -1.4357014 | cluster 1 |
| HORVU.MOREX.r3.4HG0385780.1 | 0.0064918 | -0.1207269 | -0.5374264 | cluster 1 |
| HORVU.MOREX.r3.4HG0389580.1 | 0.0242274 | -0.0821395 | -0.434092 | cluster 1 |
| HORVU.MOREX.r3.4HG0389860.1 | -0.022641 | -0.1038957 | -0.3391206 | cluster 1 |
| HORVU.MOREX.r3.4HG0390580.1 | 0.0983186 | 0.1044467 | -0.7348324 | cluster 1 |
| HORVU.MOREX.r3.4HG0391050.1 | 0.2521625 | -0.0739531 | -0.6706895 | cluster 1 |
| HORVU.MOREX.r3.4HG0391970.1 | 0.0045147 | -0.0289776 | -1.2157536 | cluster 1 |
| HORVU.MOREX.r3.4HG0392440.1 | 0.0293051 | -0.2593695 | -0.6496022 | cluster 1 |
| HORVU.MOREX.r3.4HG0393500.1 | 0.1140934 | 0.1020535 | -0.8301427 | cluster 1 |
| HORVU.MOREX.r3.4HG0394470.1 | -0.028764 | -0.4560075 | -1.1302958 | cluster 1 |
| HORVU.MOREX.r3.4HG0395120.1 | -0.296843 | -0.1032085 | -0.650262 | cluster 1 |
| HORVU.MOREX.r3.4HG0397280.1 | 0.0421174 | -0.0032905 | -0.5443237 | cluster 1 |
| HORVU.MOREX.r3.4HG0397570.1 | 0.062818 | -0.1718389 | -0.9564125 | cluster 1 |
| HORVU.MOREX.r3.4HG0399460.1 | 0.1386627 | -0.3977537 | -2.1109442 | cluster 1 |
| HORVU.MOREX.r3.4HG0400830.1 | -0.283792 | -0.2380419 | -2.2740873 | cluster 1 |
| HORVU.MOREX.r3.4HG0403910.1 | -0.462744 | -0.2304379 | -1.1725856 | cluster 1 |
| HORVU.MOREX.r3.4HG0403930.1 | 0.2191421 | -0.0297276 | -1.0240076 | cluster 1 |
| HORVU.MOREX.r3.4HG0405830.1 | -0.027851 | -0.0722627 | -0.7361311 | cluster 1 |
| HORVU.MOREX.r3.4HG0406720.1 | -0.468853 | -0.6609613 | -1.7474568 | cluster 1 |
| HORVU.MOREX.r3.4HG0406900.1 | -0.052493 | -0.1494114 | -0.5710678 | cluster 1 |
| HORVU.MOREX.r3.4HG0406980.1 | 0.2603271 | 0.1433092 | -0.9154549 | cluster 1 |
| HORVU.MOREX.r3.4HG0407120.1 | 0.0538909 | -0.0180769 | -0.5743709 | cluster 1 |
| HORVU.MOREX.r3.4HG0411320.1 | 0.1112038 | 0.0159783 | -0.4985803 | cluster 1 |
| HORVU.MOREX.r3.4HG0413890.1 | 0.0006388 | -0.1494476 | -0.424155 | cluster 1 |
| HORVU.MOREX.r3.4HG0417410.1 | -0.493011 | -0.3531985 | -1.3495621 | cluster 1 |
| HORVU.MOREX.r3.5HG0426180.1 | 0.002104 | -0.1640506 | -0.5993603 | cluster 1 |
| HORVU.MOREX.r3.5HG0427690.1 | 0.1895993 | 0.0017769 | -0.8550742 | cluster 1 |
| HORVU.MOREX.r3.5HG0436990.1 | 0.1520933 | 0.1580867 | -0.637329 | cluster 1 |
| HORVU.MOREX.r3.5HG0439620.1 | 0.0102061 | -0.1402296 | -0.4894307 | cluster 1 |
| HORVU.MOREX.r3.5HG0440700.1 | 0.0283733 | 0.0290499 | -0.444244 | cluster 1 |
| HORVU.MOREX.r3.5HG0445510.1 | -0.048345 | -0.1288695 | -0.4798599 | cluster 1 |
| HORVU.MOREX.r3.5HG0446490.1 | 0.1124989 | 0.0530041 | -0.8088415 | cluster 1 |
| HORVU.MOREX.r3.5HG0453430.1 | -0.011561 | -0.1353413 | -0.849851 | cluster 1 |
| HORVU.MOREX.r3.5HG0455240.1 | 0.1990786 | 0.1165271 | -0.7857424 | cluster 1 |
| HORVU.MOREX.r3.5HG0455770.1 | 0.2350848 | 0.0527657 | -0.8139052 | cluster 1 |
| HORVU.MOREX.r3.5HG0455820.1 | 0.2251138 | -0.1074046 | -0.9689068 | cluster 1 |
| HORVU.MOREX.r3.5HG0456950.1 | 0.0185592 | 0.1010292 | -0.4289335 | cluster 1 |
| HORVU.MOREX.r3.5HG0457960.1 | 0.0311898 | 0.0335217 | -0.3381911 | cluster 1 |
| HORVU.MOREX.r3.5HG0458100.1 | 0.2157855 | 0.073006 | -0.8015739 | cluster 1 |
| HORVU.MOREX.r3.5HG0460300.1 | -0.170396 | -0.4116975 | -0.859383 | cluster 1 |
| HORVU.MOREX.r3.5HG0461790.1 | 0.3984617 | 0.1073383 | -1.0271729 | cluster 1 |
| HORVU.MOREX.r3.5HG0462780.1 | 0.0685421 | 0.052287 | -0.5322816 | cluster 1 |

**Table S2** Continued.

| ID | logFC_3 h | logFC_6 h | logFC_12 h | cluster No. |
|---|---|---|---|---|
| HORVU.MOREX.r3.5HG0463370.1 | 0.2956809 | -0.1183146 | -1.1099129 | cluster 1 |
| HORVU.MOREX.r3.5HG0464600.1 | 0.124074 | 0.0020526 | -0.6616022 | cluster 1 |
| HORVU.MOREX.r3.5HG0467230.2 | 0.0352216 | -0.1302257 | -0.448615 | cluster 1 |
| HORVU.MOREX.r3.5HG0468320.1 | 0.2729129 | 0.0914121 | -0.8455433 | cluster 1 |
| HORVU.MOREX.r3.5HG0471410.1 | 0.0297267 | -0.0133108 | -0.4627252 | cluster 1 |
| HORVU.MOREX.r3.5HG0471560.1 | 0.0783714 | 0.0665973 | -0.5959327 | cluster 1 |
| HORVU.MOREX.r3.5HG0471940.1 | -0.057601 | -0.1501616 | -0.3613076 | cluster 1 |
| HORVU.MOREX.r3.5HG0476220.1 | 0.1782595 | -0.0710944 | -0.7292878 | cluster 1 |
| HORVU.MOREX.r3.5HG0480430.1 | 0.1411169 | 0.0038158 | -0.7527898 | cluster 1 |
| HORVU.MOREX.r3.5HG0481030.1 | 0.2389392 | 0.1744726 | -0.8003171 | cluster 1 |
| HORVU.MOREX.r3.5HG0481120.1 | 0.1169389 | -0.1089151 | -0.9823108 | cluster 1 |
| HORVU.MOREX.r3.5HG0481570.1 | 0.2363627 | 0.1421559 | -0.8273991 | cluster 1 |
| HORVU.MOREX.r3.5HG0483810.1 | 0.0204578 | -0.0567118 | -0.4489361 | cluster 1 |
| HORVU.MOREX.r3.5HG0487890.1 | 0.0781162 | -0.0644205 | -0.8183504 | cluster 1 |
| HORVU.MOREX.r3.5HG0488620.1 | 0.2344706 | 0.0734827 | -0.5842426 | cluster 1 |
| HORVU.MOREX.r3.5HG0494470.1 | -0.016452 | -0.0193565 | -0.464456 | cluster 1 |
| HORVU.MOREX.r3.5HG0494890.1 | 0.0758095 | -0.1661683 | -0.7030441 | cluster 1 |
| HORVU.MOREX.r3.5HG0495750.1 | -0.05841 | -0.2452742 | -1.9144681 | cluster 1 |
| HORVU.MOREX.r3.5HG0495770.1 | 0.1529297 | 0.0353129 | -0.4807744 | cluster 1 |
| HORVU.MOREX.r3.5HG0496780.1 | 0.1548495 | 0.0326807 | -0.5162585 | cluster 1 |
| HORVU.MOREX.r3.5HG0497170.1 | 0.070234 | -0.0373455 | -0.5609432 | cluster 1 |
| HORVU.MOREX.r3.5HG0501370.1 | 0.1640964 | 0.0254852 | -0.9053354 | cluster 1 |
| HORVU.MOREX.r3.5HG0501950.1 | 0.0456301 | -0.2756289 | -1.1865335 | cluster 1 |
| HORVU.MOREX.r3.5HG0504340.1 | -0.119178 | -0.0650048 | -0.6721698 | cluster 1 |
| HORVU.MOREX.r3.5HG0506790.1 | 0.1216365 | -0.0420748 | -0.6837788 | cluster 1 |
| HORVU.MOREX.r3.5HG0507370.1 | -0.036919 | 0.0483272 | -0.5774916 | cluster 1 |
| HORVU.MOREX.r3.5HG0509060.1 | 0.0658858 | -0.1279841 | -0.9596401 | cluster 1 |
| HORVU.MOREX.r3.5HG0510700.1 | 0.1073467 | -0.0698439 | -0.470715 | cluster 1 |
| HORVU.MOREX.r3.5HG0511040.1 | 0.2626462 | -0.1841417 | -0.9497797 | cluster 1 |
| HORVU.MOREX.r3.5HG0512430.1 | 0.2742379 | -0.1540368 | -0.8102867 | cluster 1 |
| HORVU.MOREX.r3.5HG0513000.1 | -0.037123 | -0.1405624 | -0.4575904 | cluster 1 |
| HORVU.MOREX.r3.5HG0517300.1 | 0.2159498 | 0.0391928 | -1.098873 | cluster 1 |
| HORVU.MOREX.r3.5HG0517330.1 | 0.101533 | -0.1227283 | -0.5071322 | cluster 1 |
| HORVU.MOREX.r3.5HG0517370.1 | -0.162276 | -0.2578441 | -2.3322055 | cluster 1 |
| HORVU.MOREX.r3.5HG0519030.1 | -0.084406 | -0.1435039 | -0.6605251 | cluster 1 |
| HORVU.MOREX.r3.5HG0519200.1 | 0.0015367 | -0.062071 | -0.8082744 | cluster 1 |
| HORVU.MOREX.r3.5HG0527770.1 | 0.2700764 | 0.1916167 | -0.7964304 | cluster 1 |
| HORVU.MOREX.r3.6HG0548800.1 | -0.100672 | -0.3426621 | -1.1749279 | cluster 1 |
| HORVU.MOREX.r3.6HG0552230.1 | 0.1295268 | -0.2060539 | -0.8487039 | cluster 1 |
| HORVU.MOREX.r3.6HG0555250.1 | 0.0951596 | -0.1402506 | -0.6986989 | cluster 1 |
| HORVU.MOREX.r3.6HG0559390.1 | 0.1236001 | -0.1795724 | -1.120197 | cluster 1 |
| HORVU.MOREX.r3.6HG0560030.1 | 0.1700847 | -0.0490231 | -0.5275027 | cluster 1 |
| HORVU.MOREX.r3.6HG0561320.1 | -0.134292 | -0.2913722 | -1.2817497 | cluster 1 |
| HORVU.MOREX.r3.6HG0565880.1 | 0.067755 | -0.1601905 | -0.515877 | cluster 1 |
| HORVU.MOREX.r3.6HG0568690.1 | 0.0482127 | -0.1862989 | -1.0389096 | cluster 1 |
| HORVU.MOREX.r3.6HG0568720.1 | 0.1485002 | -0.0694785 | -0.8142139 | cluster 1 |
| HORVU.MOREX.r3.6HG0570660.1 | -0.251501 | -0.5498413 | -1.3005543 | cluster 1 |

**Table S2** Continued.

| ID | logFC_3 h | logFC_6 h | logFC_12 h | cluster No. |
|---|---|---|---|---|
| HORVU.MOREX.r3.6HG0570850.1 | 0.127417 | 0.0169309 | -0.6008693 | cluster 1 |
| HORVU.MOREX.r3.6HG0577180.1 | 0.2508565 | -0.1256526 | -1.0030752 | cluster 1 |
| HORVU.MOREX.r3.6HG0591720.1 | 0.0498022 | 0.0731967 | -0.4857176 | cluster 1 |
| HORVU.MOREX.r3.6HG0597880.1 | 0.2578119 | 0.1874793 | -0.5704934 | cluster 1 |
| HORVU.MOREX.r3.6HG0598380.1 | -0.120661 | -0.4153604 | -1.0412936 | cluster 1 |
| HORVU.MOREX.r3.6HG0600190.1 | 0.2602727 | 0.0129475 | -0.6702258 | cluster 1 |
| HORVU.MOREX.r3.6HG0601740.1 | 0.1677533 | 0.0638366 | -1.1147798 | cluster 1 |
| HORVU.MOREX.r3.6HG0603150.1 | 0.2919512 | 0.0933604 | -0.7566002 | cluster 1 |
| HORVU.MOREX.r3.6HG0604990.1 | -0.447725 | -0.4972408 | -1.3495248 | cluster 1 |
| HORVU.MOREX.r3.6HG0607470.1 | 0.1319364 | 0.1609648 | -0.4571649 | cluster 1 |
| HORVU.MOREX.r3.6HG0608380.1 | -0.010442 | -0.059872 | -0.473855 | cluster 1 |
| HORVU.MOREX.r3.6HG0609220.1 | 0.1201262 | -0.0377395 | -0.6675735 | cluster 1 |
| HORVU.MOREX.r3.6HG0611930.1 | -0.577187 | -0.6670841 | -1.5392809 | cluster 1 |
| HORVU.MOREX.r3.6HG0614420.1 | -0.004193 | -0.2411698 | -0.7060912 | cluster 1 |
| HORVU.MOREX.r3.6HG0615840.1 | -0.491774 | -0.3090927 | -1.8641589 | cluster 1 |
| HORVU.MOREX.r3.6HG0617830.1 | 0.0998872 | -0.1553865 | -0.9023153 | cluster 1 |
| HORVU.MOREX.r3.6HG0622540.1 | 0.2053204 | -0.1165191 | -0.5823518 | cluster 1 |
| HORVU.MOREX.r3.6HG0623730.1 | 0.1644459 | 0.0709713 | -0.9257004 | cluster 1 |
| HORVU.MOREX.r3.6HG0625490.1 | -0.079074 | -0.2381439 | -0.5004157 | cluster 1 |
| HORVU.MOREX.r3.6HG0626000.1 | -0.044269 | -0.0781342 | -1.0887527 | cluster 1 |
| HORVU.MOREX.r3.6HG0626330.1 | 0.2099912 | 0.1832228 | -0.6671732 | cluster 1 |
| HORVU.MOREX.r3.6HG0630690.1 | 0.0081018 | -0.1090883 | -0.6131988 | cluster 1 |
| HORVU.MOREX.r3.6HG0631080.1 | 0.1026792 | -0.1050413 | -0.4997856 | cluster 1 |
| HORVU.MOREX.r3.6HG0634140.1 | -0.000579 | -0.1146015 | -0.8232646 | cluster 1 |
| HORVU.MOREX.r3.7HG0635050.1 | 0.0880867 | -0.0077246 | -0.7648137 | cluster 1 |
| HORVU.MOREX.r3.7HG0635330.1 | 0.1399067 | -0.632396 | -1.7380487 | cluster 1 |
| HORVU.MOREX.r3.7HG0639840.1 | -0.241984 | -0.8490971 | -1.9875216 | cluster 1 |
| HORVU.MOREX.r3.7HG0644160.1 | 0.346686 | -0.0760785 | -0.928141 | cluster 1 |
| HORVU.MOREX.r3.7HG0646880.1 | 0.1051319 | 0.0621529 | -0.4806318 | cluster 1 |
| HORVU.MOREX.r3.7HG0648140.1 | 0.0304123 | -0.2111924 | -0.5829098 | cluster 1 |
| HORVU.MOREX.r3.7HG0648610.1 | 0.1054259 | 0.0656255 | -0.2977385 | cluster 1 |
| HORVU.MOREX.r3.7HG0656220.1 | 0.0757624 | 0.0514909 | -0.583046 | cluster 1 |
| HORVU.MOREX.r3.7HG0658130.1 | 0.156762 | -0.0498072 | -0.7854154 | cluster 1 |
| HORVU.MOREX.r3.7HG0659980.1 | 0.1938869 | 0.1546256 | -0.7528913 | cluster 1 |
| HORVU.MOREX.r3.7HG0660830.1 | 0.0202442 | -0.0344004 | -0.3930426 | cluster 1 |
| HORVU.MOREX.r3.7HG0663550.1 | -0.524353 | -1.1082988 | -2.1275594 | cluster 1 |
| HORVU.MOREX.r3.7HG0666350.1 | 0.1070443 | -0.0164354 | -0.6601946 | cluster 1 |
| HORVU.MOREX.r3.7HG0668190.1 | 0.005287 | -0.0551883 | -0.5840778 | cluster 1 |
| HORVU.MOREX.r3.7HG0668260.1 | -0.305339 | -0.5685239 | -1.4719424 | cluster 1 |
| HORVU.MOREX.r3.7HG0669600.1 | 0.1709775 | -0.1215489 | -0.6486921 | cluster 1 |
| HORVU.MOREX.r3.7HG0670910.1 | 0.0779755 | -0.0383785 | -0.4293611 | cluster 1 |
| HORVU.MOREX.r3.7HG0671350.1 | -0.027484 | -0.0982013 | -0.4159508 | cluster 1 |
| HORVU.MOREX.r3.7HG0671720.1 | -0.176628 | -0.1787856 | -0.8791208 | cluster 1 |
| HORVU.MOREX.r3.7HG0675390.1 | -0.051485 | -0.209642 | -0.8313616 | cluster 1 |
| HORVU.MOREX.r3.7HG0676620.1 | -0.223148 | -0.4208722 | -1.4446499 | cluster 1 |
| HORVU.MOREX.r3.7HG0677380.1 | 0.2690831 | 0.2664463 | -0.5500272 | cluster 1 |
| HORVU.MOREX.r3.7HG0677710.1 | 0.3548376 | -0.2685487 | -1.3310348 | cluster 1 |

**Table S2** Continued.

| ID | logFC_3 h | logFC_6 h | logFC_12 h | cluster No. |
|---|---|---|---|---|
| HORVU.MOREX.r3.7HG0678310.1 | 0.1227949 | 0.1579376 | -0.6653024 | cluster 1 |
| HORVU.MOREX.r3.7HG0678530.1 | -0.011797 | -0.1279937 | -0.5352814 | cluster 1 |
| HORVU.MOREX.r3.7HG0682660.1 | 0.3288058 | 0.2327431 | -1.9326566 | cluster 1 |
| HORVU.MOREX.r3.7HG0682960.1 | 0.0610082 | -0.0545504 | -0.7044404 | cluster 1 |
| HORVU.MOREX.r3.7HG0685620.1 | 0.0505644 | -0.1332645 | -0.3892556 | cluster 1 |
| HORVU.MOREX.r3.7HG0700200.1 | 0.3314231 | 0.1487501 | -0.4546355 | cluster 1 |
| HORVU.MOREX.r3.7HG0700790.1 | 0.033588 | -0.184451 | -0.6111468 | cluster 1 |
| HORVU.MOREX.r3.7HG0702100.1 | 0.3679663 | 0.1743058 | -0.8553461 | cluster 1 |
| HORVU.MOREX.r3.7HG0702990.1 | 0.0227395 | -0.1430075 | -0.5485105 | cluster 1 |
| HORVU.MOREX.r3.7HG0703000.1 | 0.2292592 | 0.0543605 | -0.7113631 | cluster 1 |
| HORVU.MOREX.r3.7HG0704050.1 | -0.375344 | -0.514671 | -0.7466578 | cluster 1 |
| HORVU.MOREX.r3.7HG0705240.1 | 0.2418222 | -0.0298241 | -0.8731412 | cluster 1 |
| HORVU.MOREX.r3.7HG0706090.1 | 0.0118082 | -0.0816041 | -0.5179133 | cluster 1 |
| HORVU.MOREX.r3.7HG0707460.1 | 0.154681 | -0.0484387 | -0.9166165 | cluster 1 |
| HORVU.MOREX.r3.7HG0708320.1 | -0.242928 | -0.2894651 | -0.6994034 | cluster 1 |
| HORVU.MOREX.r3.7HG0708910.1 | -0.005875 | -0.0218331 | -0.5841389 | cluster 1 |
| HORVU.MOREX.r3.7HG0709580.1 | -0.623111 | -0.392332 | -1.1588578 | cluster 1 |
| HORVU.MOREX.r3.7HG0709700.1 | 0.1132441 | -0.2572299 | -1.0859089 | cluster 1 |
| HORVU.MOREX.r3.7HG0713100.1 | 0.0915034 | 0.0036383 | -0.7228589 | cluster 1 |
| HORVU.MOREX.r3.7HG0713160.1 | 0.1546164 | 0.0117517 | -0.6117935 | cluster 1 |
| HORVU.MOREX.r3.7HG0713610.1 | 0.0515866 | 0.1333606 | -0.4893423 | cluster 1 |
| HORVU.MOREX.r3.7HG0714350.1 | -0.127096 | -0.2561708 | -0.9241313 | cluster 1 |
| HORVU.MOREX.r3.7HG0719310.1 | -1.1903 | -0.3407218 | -2.314998 | cluster 1 |
| HORVU.MOREX.r3.7HG0719840.1 | -0.052685 | 0.0812511 | -0.6876225 | cluster 1 |
| HORVU.MOREX.r3.7HG0720730.1 | 0.3140162 | -0.109002 | -1.036229 | cluster 1 |
| HORVU.MOREX.r3.7HG0720980.1 | 0.2862122 | -0.1200729 | -0.7428184 | cluster 1 |
| HORVU.MOREX.r3.7HG0721080.1 | -0.187276 | -0.3456837 | -0.7337321 | cluster 1 |
| HORVU.MOREX.r3.7HG0721660.1 | 0.1128853 | -0.090692 | -0.7058249 | cluster 1 |
| HORVU.MOREX.r3.7HG0723280.1 | 0.2980273 | -0.0995604 | -1.8839692 | cluster 1 |
| HORVU.MOREX.r3.7HG0723770.1 | -0.195223 | -0.2712737 | -0.5650397 | cluster 1 |
| HORVU.MOREX.r3.7HG0725210.3 | 0.2136778 | -0.0725653 | -0.6957848 | cluster 1 |
| HORVU.MOREX.r3.7HG0728500.1 | 0.3348645 | -0.1300075 | -0.9267793 | cluster 1 |
| HORVU.MOREX.r3.7HG0729010.1 | 0.2698452 | 0.0186518 | -0.8651826 | cluster 1 |
| HORVU.MOREX.r3.7HG0729510.1 | -0.10835 | -0.3466034 | -0.7606226 | cluster 1 |
| HORVU.MOREX.r3.7HG0732860.1 | 0.053409 | -0.1874811 | -0.8497754 | cluster 1 |
| HORVU.MOREX.r3.7HG0733020.1 | 0.1931233 | -0.1032472 | -0.5307724 | cluster 1 |
| HORVU.MOREX.r3.7HG0733490.1 | -0.007564 | -0.1972393 | -0.681121 | cluster 1 |
| HORVU.MOREX.r3.7HG0735280.1 | 0.0500998 | -0.0531069 | -0.4343864 | cluster 1 |
| HORVU.MOREX.r3.7HG0736000.1 | 0.0433785 | -0.0826895 | -0.7408431 | cluster 1 |
| HORVU.MOREX.r3.7HG0743430.1 | -0.030952 | -0.1973464 | -1.2716657 | cluster 1 |
| HORVU.MOREX.r3.7HG0748320.1 | 0.0360236 | -0.1921464 | -0.7514418 | cluster 1 |
| HORVU.MOREX.r3.7HG0749800.1 | 0.1722487 | 0.0275875 | -0.7703866 | cluster 1 |
| HORVU.MOREX.r3.7HG0752830.1 | 0.0725296 | -0.0269739 | -0.4228159 | cluster 1 |
| HORVU.MOREX.r3.UnG0753160.1 | 0.1726371 | 0.1024129 | -0.6114766 | cluster 1 |
| HORVU.MOREX.r3.1HG0003270.1 | 2.8233992 | 1.862729 | 1.634235 | cluster 2 |
| HORVU.MOREX.r3.1HG0051740.1 | 0.0370234 | -0.6811892 | -0.9873424 | cluster 2 |
| HORVU.MOREX.r3.1HG0068060.1 | -0.042873 | -0.2001404 | -0.3550378 | cluster 2 |

**Table S2** Continued.

| ID | logFC_3 h | logFC_6 h | logFC_12 h | cluster No. |
|---|---|---|---|---|
| HORVU.MOREX.r3.1HG0080420.1 | 0.2379561 | -0.2697485 | -0.6518095 | cluster 2 |
| HORVU.MOREX.r3.1HG0080790.1 | -0.167621 | -0.7173333 | -1.0990947 | cluster 2 |
| HORVU.MOREX.r3.1HG0092900.1 | 0.5080365 | -0.4951608 | -1.5485724 | cluster 2 |
| HORVU.MOREX.r3.1HG0095220.1 | 0.136006 | -0.3412548 | -0.7813061 | cluster 2 |
| HORVU.MOREX.r3.2HG0110640.1 | -0.086711 | -0.3776334 | -0.3137127 | cluster 2 |
| HORVU.MOREX.r3.2HG0117540.1 | -0.124083 | -0.4106064 | -0.5624112 | cluster 2 |
| HORVU.MOREX.r3.2HG0119220.1 | -0.052723 | -0.3327517 | -0.6913123 | cluster 2 |
| HORVU.MOREX.r3.2HG0120640.1 | -0.274605 | -1.0532317 | -1.5376216 | cluster 2 |
| HORVU.MOREX.r3.2HG0121950.1 | -0.236053 | -0.5226593 | -0.7069055 | cluster 2 |
| HORVU.MOREX.r3.2HG0125510.1 | -0.286447 | -1.3277697 | -1.4071916 | cluster 2 |
| HORVU.MOREX.r3.2HG0140970.1 | -0.086658 | -0.9432994 | -1.704572 | cluster 2 |
| HORVU.MOREX.r3.2HG0151920.1 | -0.063584 | -0.5804899 | -0.8562623 | cluster 2 |
| HORVU.MOREX.r3.2HG0162620.1 | 0.1360319 | -0.6676713 | -1.5894203 | cluster 2 |
| HORVU.MOREX.r3.2HG0175220.1 | 0.0842263 | -0.1561619 | -0.4366907 | cluster 2 |
| HORVU.MOREX.r3.2HG0183730.1 | 0.1082802 | -0.5671608 | -1.2022611 | cluster 2 |
| HORVU.MOREX.r3.2HG0184620.1 | -0.32692 | -0.7191037 | -0.7873918 | cluster 2 |
| HORVU.MOREX.r3.3HG0221140.1 | 0.1221567 | -0.2749698 | -0.7589752 | cluster 2 |
| HORVU.MOREX.r3.3HG0231430.1 | 0.0445826 | -0.206786 | -0.378026 | cluster 2 |
| HORVU.MOREX.r3.3HG0235720.1 | -0.526825 | -1.2098574 | -1.4772443 | cluster 2 |
| HORVU.MOREX.r3.3HG0249970.1 | 0.0863874 | -0.326294 | -0.5529555 | cluster 2 |
| HORVU.MOREX.r3.3HG0257120.1 | -0.052205 | -0.3750885 | -0.7598041 | cluster 2 |
| HORVU.MOREX.r3.3HG0295500.2 | 0.227079 | -0.365709 | -1.1006335 | cluster 2 |
| HORVU.MOREX.r3.3HG0305840.1 | -0.086915 | -0.6456474 | -0.7956632 | cluster 2 |
| HORVU.MOREX.r3.4HG0340200.1 | 0.0298715 | -0.335933 | -0.6705089 | cluster 2 |
| HORVU.MOREX.r3.4HG0408300.1 | 0.0707924 | -0.0811674 | -0.2722344 | cluster 2 |
| HORVU.MOREX.r3.5HG0459810.1 | -0.112209 | -0.3342213 | -0.5635667 | cluster 2 |
| HORVU.MOREX.r3.5HG0479000.1 | -0.23248 | -1.5089971 | -1.5447502 | cluster 2 |
| HORVU.MOREX.r3.5HG0496740.1 | 0.1128452 | -0.7820818 | -1.662913 | cluster 2 |
| HORVU.MOREX.r3.5HG0500130.1 | -0.068644 | -1.1977382 | -2.2554112 | cluster 2 |
| HORVU.MOREX.r3.5HG0516730.1 | 0.0303497 | -0.346808 | -0.8068943 | cluster 2 |
| HORVU.MOREX.r3.5HG0520850.1 | -0.015761 | -0.2766114 | -0.5416283 | cluster 2 |
| HORVU.MOREX.r3.6HG0551620.1 | 0.0126099 | -0.264654 | -0.5492368 | cluster 2 |
| HORVU.MOREX.r3.6HG0553150.1 | 0.065038 | -0.3376159 | -0.816789 | cluster 2 |
| HORVU.MOREX.r3.6HG0557640.2 | -0.110285 | -0.4558812 | -0.852527 | cluster 2 |
| HORVU.MOREX.r3.6HG0565830.1 | 0.2628817 | -0.2434719 | -0.705834 | cluster 2 |
| HORVU.MOREX.r3.6HG0565890.1 | 0.0716683 | -0.3935353 | -0.8171697 | cluster 2 |
| HORVU.MOREX.r3.6HG0592790.1 | -0.00306 | -0.1774979 | -0.3776622 | cluster 2 |
| HORVU.MOREX.r3.6HG0596240.1 | -0.011888 | -0.5674737 | -1.0823091 | cluster 2 |
| HORVU.MOREX.r3.6HG0607560.1 | 0.1633152 | -0.2593501 | -0.8087621 | cluster 2 |
| HORVU.MOREX.r3.6HG0621190.1 | -0.199352 | -0.9152175 | -1.0412622 | cluster 2 |
| HORVU.MOREX.r3.6HG0630410.1 | 0.1693997 | -0.6981072 | -1.2062575 | cluster 2 |
| HORVU.MOREX.r3.7HG0635410.1 | -0.061666 | -0.6913512 | -1.1357064 | cluster 2 |
| HORVU.MOREX.r3.7HG0664810.1 | -0.59145 | -1.1235193 | -1.2315395 | cluster 2 |
| HORVU.MOREX.r3.7HG0680170.1 | -0.105886 | -0.3539509 | -0.5322635 | cluster 2 |
| HORVU.MOREX.r3.7HG0724830.1 | -0.12338 | -0.4212108 | -0.6871245 | cluster 2 |
| HORVU.MOREX.r3.7HG0733070.1 | 0.2774792 | -0.1652324 | -0.5134848 | cluster 2 |
| HORVU.MOREX.r3.7HG0733110.1 | 0.0655865 | -0.4471865 | -0.701074 | cluster 2 |

**Table S2** Continued.

| ID | logFC_3 h | logFC_6 h | logFC_12 h | cluster No. |
|---|---|---|---|---|
| HORVU.MOREX.r3.7HG0740120.1 | -0.045066 | -0.4939793 | -0.8609598 | cluster 2 |
| HORVU.MOREX.r3.1HG0024440.1 | 0.6240205 | 0.3338753 | 1.7616367 | cluster 3 |
| HORVU.MOREX.r3.1HG0029060.1 | -0.002115 | 0.115194 | 0.3316882 | cluster 3 |
| HORVU.MOREX.r3.1HG0056810.1 | -0.050522 | -0.0760236 | 0.5789208 | cluster 3 |
| HORVU.MOREX.r3.1HG0062480.1 | 0.1769776 | 0.1754143 | 0.8481508 | cluster 3 |
| HORVU.MOREX.r3.1HG0072560.1 | 1.420637 | 2.1085454 | 1.8846069 | cluster 3 |
| HORVU.MOREX.r3.2HG0104400.1 | 0.1392872 | -1.5799965 | 2.8499093 | cluster 3 |
| HORVU.MOREX.r3.2HG0116730.1 | 0.6039269 | 0.8607134 | 3.2531155 | cluster 3 |
| HORVU.MOREX.r3.2HG0141900.1 | 0.1107861 | 0.4220753 | 0.6719383 | cluster 3 |
| HORVU.MOREX.r3.2HG0141980.1 | -0.086386 | -0.0404557 | 0.6743435 | cluster 3 |
| HORVU.MOREX.r3.2HG0181380.1 | 0.1104037 | 0.2767668 | 0.7848569 | cluster 3 |
| HORVU.MOREX.r3.2HG0191150.1 | -0.119802 | 0.0994863 | 0.987218 | cluster 3 |
| HORVU.MOREX.r3.2HG0192180.1 | 0.0106233 | 0.0702629 | 0.2270186 | cluster 3 |
| HORVU.MOREX.r3.2HG0198000.1 | -0.05969 | 0.0875793 | 0.300141 | cluster 3 |
| HORVU.MOREX.r3.3HG0219850.1 | 0.4620524 | 1.6355633 | 3.5424753 | cluster 3 |
| HORVU.MOREX.r3.3HG0300110.1 | 0.811676 | 1.5112064 | 0.6639938 | cluster 3 |
| HORVU.MOREX.r3.3HG0309210.1 | -0.08868 | -0.0479044 | 0.4562355 | cluster 3 |
| HORVU.MOREX.r3.4HG0402190.1 | 1.1812933 | 2.5774222 | 1.9268757 | cluster 3 |
| HORVU.MOREX.r3.4HG0402390.1 | 2.9041334 | 4.4100052 | 5.2262576 | cluster 3 |
| HORVU.MOREX.r3.4HG0405330.1 | 1.0455541 | 1.7983357 | 1.1670497 | cluster 3 |
| HORVU.MOREX.r3.5HG0429390.1 | 2.0002395 | 3.6322175 | 3.1784806 | cluster 3 |
| HORVU.MOREX.r3.5HG0458770.1 | 0.5221025 | 0.6619199 | 0.7160158 | cluster 3 |
| HORVU.MOREX.r3.5HG0482710.1 | 0.226056 | 0.2311383 | 0.6233293 | cluster 3 |
| HORVU.MOREX.r3.5HG0498320.1 | 0.1160828 | 0.1697017 | 0.4293063 | cluster 3 |
| HORVU.MOREX.r3.5HG0504720.1 | -0.10073 | 0.0335607 | 0.464637 | cluster 3 |
| HORVU.MOREX.r3.5HG0510240.1 | -0.048271 | 1.0640468 | 4.8663744 | cluster 3 |
| HORVU.MOREX.r3.5HG0529140.1 | -0.059179 | 0.0726222 | 0.4325476 | cluster 3 |
| HORVU.MOREX.r3.6HG0541580.1 | 0.0976735 | 0.1899155 | 0.2698673 | cluster 3 |
| HORVU.MOREX.r3.6HG0550260.1 | 0.0195363 | 0.1405006 | 0.4032082 | cluster 3 |
| HORVU.MOREX.r3.6HG0604200.1 | 0.7056787 | -0.0247538 | 2.9483156 | cluster 3 |
| HORVU.MOREX.r3.6HG0627900.1 | 0.3557403 | 1.5427197 | 2.4470309 | cluster 3 |
| HORVU.MOREX.r3.7HG0656980.1 | 0.0472123 | 1.4907654 | 1.220906 | cluster 3 |
| HORVU.MOREX.r3.7HG0689920.1 | 0.6722508 | 0.54806 | 1.3527666 | cluster 3 |
| HORVU.MOREX.r3.7HG0705130.1 | 0.7312682 | 0.6193222 | 1.0084752 | cluster 3 |
| HORVU.MOREX.r3.7HG0725140.1 | 0.0342515 | -0.2241711 | 0.6286018 | cluster 3 |
| HORVU.MOREX.r3.7HG0725940.1 | 0.1803747 | 0.8325886 | 1.0925768 | cluster 3 |
| HORVU.MOREX.r3.7HG0733550.1 | 0.5837047 | 1.680915 | 1.5883796 | cluster 3 |
| HORVU.MOREX.r3.7HG0739570.1 | 1.1470843 | 2.5493466 | 2.1625177 | cluster 3 |
| HORVU.MOREX.r3.7HG0750850.1 | 1.4902377 | 3.4536548 | 3.8051856 | cluster 3 |
