## Supporting Information Table 3 for "ENHANCED GRAVITROPISM 2 coordinates molecular adaptations to gravistimulation in the elongation zone of barley roots"

**Table S3** Hierarchical clustering analysis of differentially expressed genes (FDR <5%) in elongation zone of gravistimulated wild type roots.

| ID | logFC_3 h | logFC_6 h | logFC_12 h | cluster No. |
|---|---|---|---|---|
| HORVU.MOREX.r3.1HG0000040.1 | -0.071645 | -0.124524 | -0.1773485 | cluster 1 |
| HORVU.MOREX.r3.1HG0000670.1 | -0.438469 | -0.684321 | -0.8443564 | cluster 1 |
| HORVU.MOREX.r3.1HG0001010.1 | -0.377642 | -0.794982 | -0.8842452 | cluster 1 |
| HORVU.MOREX.r3.1HG0001460.1 | -0.133788 | -0.543799 | -0.836291 | cluster 1 |
| HORVU.MOREX.r3.1HG0001510.1 | -0.612987 | -0.793475 | -1.0657923 | cluster 1 |
| HORVU.MOREX.r3.1HG0002800.1 | -0.331522 | -0.93359 | -1.3595718 | cluster 1 |
| HORVU.MOREX.r3.1HG0003090.4 | -0.183698 | -1.041325 | -1.6979849 | cluster 1 |
| HORVU.MOREX.r3.1HG0003090.7 | -0.577884 | -1.191555 | -1.8866381 | cluster 1 |
| HORVU.MOREX.r3.1HG0003130.1 | -0.142771 | -0.448507 | -0.493319 | cluster 1 |
| HORVU.MOREX.r3.1HG0004290.1 | -0.107222 | -0.376673 | -1.0557282 | cluster 1 |
| HORVU.MOREX.r3.1HG0004300.1 | -0.726547 | -0.889262 | -1.9261646 | cluster 1 |
| HORVU.MOREX.r3.1HG0005750.1 | -0.161951 | -0.328139 | -0.4439202 | cluster 1 |
| HORVU.MOREX.r3.1HG0006610.1 | -0.127119 | -0.498945 | -0.9589396 | cluster 1 |
| HORVU.MOREX.r3.1HG0006650.1 | -0.152638 | -0.430358 | -0.8783079 | cluster 1 |
| HORVU.MOREX.r3.1HG0007360.1 | -0.358003 | -0.434503 | -0.5486322 | cluster 1 |
| HORVU.MOREX.r3.1HG0007370.1 | -0.17099 | -0.276913 | -0.4870731 | cluster 1 |
| HORVU.MOREX.r3.1HG0008140.1 | 0.029832 | -0.045235 | -1.0526349 | cluster 1 |
| HORVU.MOREX.r3.1HG0008190.1 | -0.079562 | -0.49924 | -1.1637413 | cluster 1 |
| HORVU.MOREX.r3.1HG0008200.1 | -0.016943 | -0.950898 | -2.4521858 | cluster 1 |
| HORVU.MOREX.r3.1HG0008400.1 | -0.002467 | -0.483007 | -0.8133046 | cluster 1 |
| HORVU.MOREX.r3.1HG0008440.1 | 0.000127 | -0.155753 | -0.70841 | cluster 1 |
| HORVU.MOREX.r3.1HG0010470.1 | -0.129151 | -0.372956 | -0.5835351 | cluster 1 |
| HORVU.MOREX.r3.1HG0012170.1 | -0.249119 | -0.48712 | -0.841958 | cluster 1 |
| HORVU.MOREX.r3.1HG0012330.1 | -0.181089 | -0.298638 | -0.772728 | cluster 1 |
| HORVU.MOREX.r3.1HG0014410.1 | -0.019009 | -0.303508 | -0.635752 | cluster 1 |
| HORVU.MOREX.r3.1HG0016810.1 | -0.219936 | -0.374805 | -0.8116261 | cluster 1 |
| HORVU.MOREX.r3.1HG0016950.1 | -0.149667 | -0.340014 | -0.3814369 | cluster 1 |
| HORVU.MOREX.r3.1HG0017310.1 | -0.16519 | -0.283884 | -1.1001499 | cluster 1 |
| HORVU.MOREX.r3.1HG0017720.1 | -0.104546 | -0.614856 | -0.7433992 | cluster 1 |
| HORVU.MOREX.r3.1HG0018190.2 | -0.022676 | -0.659745 | -1.1814946 | cluster 1 |
| HORVU.MOREX.r3.1HG0018390.1 | -0.125982 | -0.457016 | -0.9856067 | cluster 1 |
| HORVU.MOREX.r3.1HG0018730.1 | -0.052508 | -0.16995 | -0.320822 | cluster 1 |
| HORVU.MOREX.r3.1HG0018810.1 | -0.378457 | -0.611311 | -0.9691882 | cluster 1 |
| HORVU.MOREX.r3.1HG0019410.1 | -0.225678 | -0.299536 | -0.9646582 | cluster 1 |
| HORVU.MOREX.r3.1HG0019440.1 | -0.105029 | -0.315985 | -0.3954099 | cluster 1 |
| HORVU.MOREX.r3.1HG0021830.1 | -0.079326 | -0.203601 | -0.3393333 | cluster 1 |
| HORVU.MOREX.r3.1HG0022860.1 | -0.088711 | -0.301696 | -0.3503478 | cluster 1 |
| HORVU.MOREX.r3.1HG0023610.1 | -0.273233 | -0.669494 | -1.3268268 | cluster 1 |
| HORVU.MOREX.r3.1HG0023620.1 | -0.167881 | -0.548371 | -0.9138041 | cluster 1 |
| HORVU.MOREX.r3.1HG0024040.1 | -0.11524 | -0.359729 | -0.4670541 | cluster 1 |
| HORVU.MOREX.r3.1HG0024140.1 | -0.141816 | -0.463273 | -1.0286073 | cluster 1 |
| HORVU.MOREX.r3.1HG0024230.1 | -0.062139 | -0.081072 | -0.5736968 | cluster 1 |
| HORVU.MOREX.r3.1HG0024280.1 | -0.149099 | -0.410992 | -0.7193662 | cluster 1 |
| HORVU.MOREX.r3.1HG0026070.2 | -0.164501 | -0.190542 | -0.5632605 | cluster 1 |
| HORVU.MOREX.r3.1HG0026430.1 | -0.21961 | -0.262259 | -0.4150473 | cluster 1 |
| HORVU.MOREX.r3.1HG0028320.1 | -0.169179 | -0.517356 | -0.6028779 | cluster 1 |
| HORVU.MOREX.r3.1HG0029080.1 | -0.1787 | -0.587693 | -0.6492816 | cluster 1 |

**Table S3** Continued.

| ID | logFC_3 h | logFC_6 h | logFC_12 h | cluster No. |
|---|---|---|---|---|
| HORVU.MOREX.r3.1HG0030090.1 | -0.216979 | -0.457384 | -1.0998744 | cluster 1 |
| HORVU.MOREX.r3.1HG0030200.1 | 0.0963493 | -0.098407 | -0.4175741 | cluster 1 |
| HORVU.MOREX.r3.1HG0031180.1 | -0.188618 | -0.26601 | -0.4864687 | cluster 1 |
| HORVU.MOREX.r3.1HG0032230.1 | -0.058604 | -0.33076 | -0.8417067 | cluster 1 |
| HORVU.MOREX.r3.1HG0032910.1 | -0.126575 | -0.418482 | -0.5760219 | cluster 1 |
| HORVU.MOREX.r3.1HG0033360.1 | -0.147892 | -0.192431 | -0.2847466 | cluster 1 |
| HORVU.MOREX.r3.1HG0033390.4 | -0.481493 | -1.038087 | -1.7182452 | cluster 1 |
| HORVU.MOREX.r3.1HG0036370.1 | -0.107118 | -0.242352 | -0.390755 | cluster 1 |
| HORVU.MOREX.r3.1HG0036930.1 | -0.225925 | -0.407071 | -1.0819263 | cluster 1 |
| HORVU.MOREX.r3.1HG0037020.1 | -0.178481 | -0.239307 | -0.4336448 | cluster 1 |
| HORVU.MOREX.r3.1HG0038960.1 | -0.273156 | -0.341301 | -1.0375946 | cluster 1 |
| HORVU.MOREX.r3.1HG0039230.1 | -0.272132 | -0.382149 | -0.661259 | cluster 1 |
| HORVU.MOREX.r3.1HG0039670.1 | -0.031701 | -0.238142 | -0.3364584 | cluster 1 |
| HORVU.MOREX.r3.1HG0042740.1 | -0.098437 | -0.296755 | -0.4789963 | cluster 1 |
| HORVU.MOREX.r3.1HG0044530.1 | -0.143886 | -0.230621 | -0.4752674 | cluster 1 |
| HORVU.MOREX.r3.1HG0045040.1 | -0.030519 | -0.202812 | -0.3801681 | cluster 1 |
| HORVU.MOREX.r3.1HG0046650.1 | -0.119401 | -0.420107 | -0.6038916 | cluster 1 |
| HORVU.MOREX.r3.1HG0046770.1 | -0.217608 | -0.3459 | -0.7078966 | cluster 1 |
| HORVU.MOREX.r3.1HG0046900.1 | -0.055034 | -0.284637 | -0.318943 | cluster 1 |
| HORVU.MOREX.r3.1HG0047130.1 | -0.102935 | -0.290613 | -0.5012279 | cluster 1 |
| HORVU.MOREX.r3.1HG0048010.1 | -0.11181 | -0.215178 | -0.3447995 | cluster 1 |
| HORVU.MOREX.r3.1HG0048030.1 | -0.043702 | -0.179739 | -0.4603345 | cluster 1 |
| HORVU.MOREX.r3.1HG0049840.1 | -0.035429 | -0.460083 | -0.7164691 | cluster 1 |
| HORVU.MOREX.r3.1HG0050990.1 | -0.044326 | -0.331078 | -2.2870311 | cluster 1 |
| HORVU.MOREX.r3.1HG0051020.1 | -0.186997 | -0.295926 | -0.4951078 | cluster 1 |
| HORVU.MOREX.r3.1HG0051740.1 | -0.149533 | -1.907755 | -2.7104377 | cluster 1 |
| HORVU.MOREX.r3.1HG0051860.1 | -0.521655 | -0.64618 | -0.920105 | cluster 1 |
| HORVU.MOREX.r3.1HG0052080.1 | -0.097286 | -0.30255 | -0.4502105 | cluster 1 |
| HORVU.MOREX.r3.1HG0052120.1 | -0.424983 | -0.620213 | -0.7628829 | cluster 1 |
| HORVU.MOREX.r3.1HG0052790.1 | -0.002574 | -0.24127 | -0.4696911 | cluster 1 |
| HORVU.MOREX.r3.1HG0052810.1 | -0.162186 | -0.364431 | -0.6609272 | cluster 1 |
| HORVU.MOREX.r3.1HG0053770.1 | -0.214542 | -0.450222 | -1.3037244 | cluster 1 |
| HORVU.MOREX.r3.1HG0054200.1 | 0.0229659 | -0.112167 | -0.3330798 | cluster 1 |
| HORVU.MOREX.r3.1HG0054500.1 | -0.141381 | -0.502872 | -0.9798408 | cluster 1 |
| HORVU.MOREX.r3.1HG0054770.1 | -0.204379 | -0.460411 | -0.5724173 | cluster 1 |
| HORVU.MOREX.r3.1HG0055230.1 | -0.112802 | -0.512176 | -0.8892409 | cluster 1 |
| HORVU.MOREX.r3.1HG0055380.1 | -0.136801 | -0.476637 | -0.6613041 | cluster 1 |
| HORVU.MOREX.r3.1HG0055400.1 | -0.091803 | -0.233082 | -0.5051363 | cluster 1 |
| HORVU.MOREX.r3.1HG0055580.1 | -0.217535 | -0.316879 | -0.4107946 | cluster 1 |
| HORVU.MOREX.r3.1HG0055730.1 | -0.181182 | -0.18899 | -0.4323416 | cluster 1 |
| HORVU.MOREX.r3.1HG0056280.1 | -0.111364 | -0.524947 | -1.053041 | cluster 1 |
| HORVU.MOREX.r3.1HG0056560.1 | -0.11454 | -0.286266 | -0.3917377 | cluster 1 |
| HORVU.MOREX.r3.1HG0056740.1 | -0.200779 | -0.410893 | -0.551492 | cluster 1 |
| HORVU.MOREX.r3.1HG0057620.1 | -0.149427 | -0.357208 | -0.7477856 | cluster 1 |
| HORVU.MOREX.r3.1HG0057780.1 | -0.08526 | -0.406475 | -0.5546754 | cluster 1 |
| HORVU.MOREX.r3.1HG0057900.1 | -0.305534 | -0.397298 | -0.6624623 | cluster 1 |
| HORVU.MOREX.r3.1HG0057990.1 | -0.332 | -0.607808 | -0.8161959 | cluster 1 |

**Table S3** Continued.

| ID | logFC_3 h | logFC_6 h | logFC_12 h | cluster No. |
|---|---|---|---|---|
| HORVU.MOREX.r3.1HG0058050.1 | -0.188489 | -0.272955 | -0.5194952 | cluster 1 |
| HORVU.MOREX.r3.1HG0058390.1 | -0.046827 | -0.501733 | -1.2016923 | cluster 1 |
| HORVU.MOREX.r3.1HG0058640.1 | -0.146274 | -0.534883 | -1.4259288 | cluster 1 |
| HORVU.MOREX.r3.1HG0058800.1 | -0.136702 | -0.32663 | -0.5382268 | cluster 1 |
| HORVU.MOREX.r3.1HG0059000.1 | -0.410661 | -0.673067 | -0.954609 | cluster 1 |
| HORVU.MOREX.r3.1HG0059120.1 | 0.0082228 | -0.265023 | -0.3284409 | cluster 1 |
| HORVU.MOREX.r3.1HG0059350.1 | -0.310436 | -0.589076 | -0.8252261 | cluster 1 |
| HORVU.MOREX.r3.1HG0059790.1 | -0.214234 | -0.530524 | -0.986547 | cluster 1 |
| HORVU.MOREX.r3.1HG0060240.1 | -0.39667 | -0.734476 | -0.8303838 | cluster 1 |
| HORVU.MOREX.r3.1HG0060260.1 | -0.19775 | -0.498184 | -0.9775598 | cluster 1 |
| HORVU.MOREX.r3.1HG0060270.1 | -0.209438 | -0.627924 | -1.0589566 | cluster 1 |
| HORVU.MOREX.r3.1HG0060320.1 | -0.17793 | -0.540094 | -1.0558507 | cluster 1 |
| HORVU.MOREX.r3.1HG0060340.1 | -0.04652 | -0.503584 | -1.4127406 | cluster 1 |
| HORVU.MOREX.r3.1HG0060570.1 | -0.050198 | -0.153708 | -0.5413945 | cluster 1 |
| HORVU.MOREX.r3.1HG0061220.1 | -0.06833 | -0.411928 | -1.1357942 | cluster 1 |
| HORVU.MOREX.r3.1HG0063870.1 | -0.173316 | -0.397724 | -0.6224312 | cluster 1 |
| HORVU.MOREX.r3.1HG0064080.1 | -0.161441 | -0.257858 | -0.5953225 | cluster 1 |
| HORVU.MOREX.r3.1HG0064220.1 | -0.199911 | -0.335029 | -0.7590624 | cluster 1 |
| HORVU.MOREX.r3.1HG0064320.1 | -0.07699 | -0.180027 | -0.5523171 | cluster 1 |
| HORVU.MOREX.r3.1HG0065020.1 | 0.0348523 | -0.269963 | -1.170304 | cluster 1 |
| HORVU.MOREX.r3.1HG0065370.1 | -0.106742 | -0.29119 | -0.7791662 | cluster 1 |
| HORVU.MOREX.r3.1HG0066530.1 | -0.157526 | -0.392825 | -1.1926228 | cluster 1 |
| HORVU.MOREX.r3.1HG0066730.1 | -0.212473 | -0.512146 | -0.6817678 | cluster 1 |
| HORVU.MOREX.r3.1HG0067410.1 | -0.095465 | -0.223165 | -0.4149666 | cluster 1 |
| HORVU.MOREX.r3.1HG0067550.1 | -0.076405 | -0.416209 | -0.7286452 | cluster 1 |
| HORVU.MOREX.r3.1HG0068370.1 | -0.2881 | -0.85527 | -1.2770371 | cluster 1 |
| HORVU.MOREX.r3.1HG0068710.1 | -0.539862 | -0.644922 | -1.0737803 | cluster 1 |
| HORVU.MOREX.r3.1HG0068890.1 | -0.214045 | -0.365888 | -0.5535155 | cluster 1 |
| HORVU.MOREX.r3.1HG0069290.1 | -0.091143 | -0.204475 | -0.3333073 | cluster 1 |
| HORVU.MOREX.r3.1HG0069940.1 | -0.08985 | -0.200192 | -0.7416549 | cluster 1 |
| HORVU.MOREX.r3.1HG0071310.1 | 0.2367884 | -0.156589 | -0.4807924 | cluster 1 |
| HORVU.MOREX.r3.1HG0072110.1 | -0.130093 | -0.303295 | -0.8139992 | cluster 1 |
| HORVU.MOREX.r3.1HG0072320.1 | -0.562477 | -0.568571 | -0.8266783 | cluster 1 |
| HORVU.MOREX.r3.1HG0072410.1 | -0.076207 | -0.23978 | -0.2694548 | cluster 1 |
| HORVU.MOREX.r3.1HG0072520.1 | -0.290377 | -0.694298 | -1.0645262 | cluster 1 |
| HORVU.MOREX.r3.1HG0072530.1 | -0.341481 | -0.655417 | -1.1449186 | cluster 1 |
| HORVU.MOREX.r3.1HG0072540.1 | -0.3341 | -0.569029 | -1.2075381 | cluster 1 |
| HORVU.MOREX.r3.1HG0073030.1 | -0.084773 | -0.477778 | -0.810206 | cluster 1 |
| HORVU.MOREX.r3.1HG0073050.1 | -0.488801 | -0.547014 | -1.4627183 | cluster 1 |
| HORVU.MOREX.r3.1HG0073070.1 | -0.055412 | -0.182142 | -0.2592944 | cluster 1 |
| HORVU.MOREX.r3.1HG0073170.1 | -0.075394 | -0.612165 | -0.8972417 | cluster 1 |
| HORVU.MOREX.r3.1HG0073320.1 | -0.406974 | -0.606669 | -0.9159297 | cluster 1 |
| HORVU.MOREX.r3.1HG0073330.1 | -0.116256 | -0.410177 | -0.4614661 | cluster 1 |
| HORVU.MOREX.r3.1HG0074260.1 | -0.150953 | -0.29947 | -0.4001584 | cluster 1 |
| HORVU.MOREX.r3.1HG0074490.1 | -0.103373 | -0.514311 | -1.2212446 | cluster 1 |
| HORVU.MOREX.r3.1HG0074500.1 | -0.215708 | -0.554579 | -1.0786443 | cluster 1 |
| HORVU.MOREX.r3.1HG0074600.1 | -0.014879 | -0.376475 | -1.2244556 | cluster 1 |

**Table S3** Continued.

| ID | logFC_3 h | logFC_6 h | logFC_12 h | cluster No. |
|---|---|---|---|---|
| HORVU.MOREX.r3.1HG0075230.1 | -0.181992 | -0.560358 | -1.2093664 | cluster 1 |
| HORVU.MOREX.r3.1HG0075590.1 | -0.097289 | -0.219511 | -0.2765667 | cluster 1 |
| HORVU.MOREX.r3.1HG0076060.1 | -0.175558 | -0.363665 | -0.4204095 | cluster 1 |
| HORVU.MOREX.r3.1HG0076540.1 | -0.072142 | -0.220057 | -0.3554766 | cluster 1 |
| HORVU.MOREX.r3.1HG0076980.1 | -0.078762 | -0.270896 | -0.3444065 | cluster 1 |
| HORVU.MOREX.r3.1HG0077170.1 | 0.1600867 | -0.137227 | -0.5622657 | cluster 1 |
| HORVU.MOREX.r3.1HG0077840.1 | -0.016089 | -0.286722 | -0.4611398 | cluster 1 |
| HORVU.MOREX.r3.1HG0077880.1 | -0.177941 | -0.31771 | -1.0754309 | cluster 1 |
| HORVU.MOREX.r3.1HG0077950.1 | -0.074884 | -0.310284 | -0.3894016 | cluster 1 |
| HORVU.MOREX.r3.1HG0078370.1 | -0.214938 | -0.412978 | -1.7854262 | cluster 1 |
| HORVU.MOREX.r3.1HG0078750.1 | -0.22172 | -0.501184 | -0.5644803 | cluster 1 |
| HORVU.MOREX.r3.1HG0078770.1 | -0.030589 | -0.121128 | -0.1448566 | cluster 1 |
| HORVU.MOREX.r3.1HG0079250.1 | -0.118705 | -0.33521 | -0.8268634 | cluster 1 |
| HORVU.MOREX.r3.1HG0079280.1 | -0.162388 | -0.439595 | -0.9395741 | cluster 1 |
| HORVU.MOREX.r3.1HG0079800.1 | -0.202182 | -0.501695 | -0.7085236 | cluster 1 |
| HORVU.MOREX.r3.1HG0079870.1 | -0.369286 | -0.820073 | -1.1327439 | cluster 1 |
| HORVU.MOREX.r3.1HG0080230.1 | -0.124723 | -0.440592 | -1.0959477 | cluster 1 |
| HORVU.MOREX.r3.1HG0080370.1 | -0.149332 | -0.316705 | -0.4841854 | cluster 1 |
| HORVU.MOREX.r3.1HG0080690.1 | -0.834104 | -0.890462 | -2.1771715 | cluster 1 |
| HORVU.MOREX.r3.1HG0080760.1 | -0.250615 | -0.580502 | -1.0702099 | cluster 1 |
| HORVU.MOREX.r3.1HG0080770.1 | -0.12037 | -0.439135 | -0.9424823 | cluster 1 |
| HORVU.MOREX.r3.1HG0080780.1 | -0.137393 | -0.53814 | -0.9969792 | cluster 1 |
| HORVU.MOREX.r3.1HG0081220.1 | -0.15209 | -0.417209 | -0.6739663 | cluster 1 |
| HORVU.MOREX.r3.1HG0082630.1 | -0.10244 | -0.36116 | -0.470862 | cluster 1 |
| HORVU.MOREX.r3.1HG0082770.1 | -0.088956 | -0.286728 | -0.3556985 | cluster 1 |
| HORVU.MOREX.r3.1HG0083110.1 | 0.015112 | -0.235565 | -0.5496173 | cluster 1 |
| HORVU.MOREX.r3.1HG0084540.1 | -0.142345 | -0.38105 | -0.5155577 | cluster 1 |
| HORVU.MOREX.r3.1HG0084850.1 | -0.259554 | -0.711168 | -1.195516 | cluster 1 |
| HORVU.MOREX.r3.1HG0084920.1 | 0.023018 | -0.198592 | -0.3069225 | cluster 1 |
| HORVU.MOREX.r3.1HG0085140.1 | -0.147025 | -0.343182 | -0.5442868 | cluster 1 |
| HORVU.MOREX.r3.1HG0085420.1 | -0.110057 | -0.423937 | -0.7578857 | cluster 1 |
| HORVU.MOREX.r3.1HG0085700.1 | -0.345336 | -0.468882 | -0.9633685 | cluster 1 |
| HORVU.MOREX.r3.1HG0086010.1 | -0.100527 | -0.179081 | -0.6194021 | cluster 1 |
| HORVU.MOREX.r3.1HG0086140.1 | -0.30656 | -0.455471 | -0.7972206 | cluster 1 |
| HORVU.MOREX.r3.1HG0087150.1 | -0.212304 | -0.595539 | -0.8760306 | cluster 1 |
| HORVU.MOREX.r3.1HG0087190.1 | -0.257724 | -0.395481 | -0.5619376 | cluster 1 |
| HORVU.MOREX.r3.1HG0087980.1 | -0.06494 | -0.212942 | -0.5635841 | cluster 1 |
| HORVU.MOREX.r3.1HG0088870.1 | -0.084584 | -0.170154 | -0.2695324 | cluster 1 |
| HORVU.MOREX.r3.1HG0088900.1 | -0.286546 | -0.525796 | -0.9227434 | cluster 1 |
| HORVU.MOREX.r3.1HG0088990.1 | -0.123571 | -0.366833 | -0.5198763 | cluster 1 |
| HORVU.MOREX.r3.1HG0089130.1 | -0.157362 | -0.353039 | -0.4242991 | cluster 1 |
| HORVU.MOREX.r3.1HG0089220.1 | -0.266468 | -0.849343 | -1.0779379 | cluster 1 |
| HORVU.MOREX.r3.1HG0089750.1 | -0.130187 | -0.411166 | -1.1368502 | cluster 1 |
| HORVU.MOREX.r3.1HG0090760.1 | -0.404317 | -0.416619 | -0.6036512 | cluster 1 |
| HORVU.MOREX.r3.1HG0091330.1 | 0.0901205 | -0.342249 | -0.9788283 | cluster 1 |
| HORVU.MOREX.r3.1HG0091360.1 | -0.122726 | -0.33111 | -0.4780708 | cluster 1 |
| HORVU.MOREX.r3.1HG0091670.1 | -0.217443 | -0.476509 | -0.5635125 | cluster 1 |

**Table S3** Continued.

| ID | logFC_3 h | logFC_6 h | logFC_12 h | cluster No. |
|---|---|---|---|---|
| HORVU.MOREX.r3.1HG0091770.1 | -0.049803 | -0.549965 | -1.2130895 | cluster 1 |
| HORVU.MOREX.r3.1HG0091820.1 | -0.049758 | -0.425615 | -0.6096677 | cluster 1 |
| HORVU.MOREX.r3.1HG0091860.1 | -0.106875 | -0.255762 | -0.3237083 | cluster 1 |
| HORVU.MOREX.r3.1HG0091870.1 | -0.07343 | -0.346194 | -0.6111268 | cluster 1 |
| HORVU.MOREX.r3.1HG0092810.1 | -0.048961 | -0.519802 | -1.1548745 | cluster 1 |
| HORVU.MOREX.r3.1HG0093750.1 | -0.335496 | -0.593977 | -0.9393611 | cluster 1 |
| HORVU.MOREX.r3.1HG0094940.1 | -0.103662 | -0.266 | -0.4251903 | cluster 1 |
| HORVU.MOREX.r3.1HG0095210.1 | -0.069131 | -0.226301 | -0.348763 | cluster 1 |
| HORVU.MOREX.r3.1HG0095440.1 | -0.154967 | -0.256627 | -0.3432853 | cluster 1 |
| HORVU.MOREX.r3.2HG0095520.1 | -0.089357 | -0.359405 | -0.5946982 | cluster 1 |
| HORVU.MOREX.r3.2HG0095650.1 | 0.2344835 | -0.038861 | -1.4585334 | cluster 1 |
| HORVU.MOREX.r3.2HG0095710.1 | -0.165009 | -0.367995 | -0.8961363 | cluster 1 |
| HORVU.MOREX.r3.2HG0095970.1 | 0.0305742 | -0.75694 | -1.272342 | cluster 1 |
| HORVU.MOREX.r3.2HG0096600.1 | -0.159346 | -0.412181 | -0.5964173 | cluster 1 |
| HORVU.MOREX.r3.2HG0096760.1 | -0.027579 | -0.221576 | -0.8044271 | cluster 1 |
| HORVU.MOREX.r3.2HG0097360.1 | -0.196544 | -0.338193 | -0.8257672 | cluster 1 |
| HORVU.MOREX.r3.2HG0097360.3 | -0.270235 | -0.612642 | -1.1548022 | cluster 1 |
| HORVU.MOREX.r3.2HG0097390.1 | -0.241772 | -0.678806 | -0.9076011 | cluster 1 |
| HORVU.MOREX.r3.2HG0097950.1 | -0.210855 | -0.560446 | -0.870493 | cluster 1 |
| HORVU.MOREX.r3.2HG0098420.1 | -0.13539 | -0.330129 | -1.1414945 | cluster 1 |
| HORVU.MOREX.r3.2HG0099260.1 | -0.052397 | -0.160273 | -0.2418427 | cluster 1 |
| HORVU.MOREX.r3.2HG0099740.1 | -0.136727 | -0.304046 | -0.3932536 | cluster 1 |
| HORVU.MOREX.r3.2HG0100040.1 | -0.167265 | -0.45725 | -2.0923434 | cluster 1 |
| HORVU.MOREX.r3.2HG0104260.1 | -0.190136 | -0.461552 | -0.6638723 | cluster 1 |
| HORVU.MOREX.r3.2HG0104480.1 | -0.176111 | -0.466432 | -0.7913353 | cluster 1 |
| HORVU.MOREX.r3.2HG0104500.1 | -0.079758 | -0.258261 | -0.4218833 | cluster 1 |
| HORVU.MOREX.r3.2HG0104510.1 | -0.135732 | -0.187992 | -0.3580829 | cluster 1 |
| HORVU.MOREX.r3.2HG0105680.3 | -0.326239 | -0.486491 | -1.0268083 | cluster 1 |
| HORVU.MOREX.r3.2HG0105980.1 | -0.396491 | -0.435585 | -0.6732587 | cluster 1 |
| HORVU.MOREX.r3.2HG0106430.1 | -0.001113 | -0.20332 | -0.4842964 | cluster 1 |
| HORVU.MOREX.r3.2HG0106750.1 | -0.188562 | -0.806451 | -1.0911822 | cluster 1 |
| HORVU.MOREX.r3.2HG0107450.1 | 0.0259002 | -0.417632 | -0.9297142 | cluster 1 |
| HORVU.MOREX.r3.2HG0107470.1 | -0.18259 | -0.354834 | -0.6522059 | cluster 1 |
| HORVU.MOREX.r3.2HG0107740.1 | -0.142411 | -0.353075 | -0.5643446 | cluster 1 |
| HORVU.MOREX.r3.2HG0107820.1 | -0.227116 | -0.524651 | -0.6777043 | cluster 1 |
| HORVU.MOREX.r3.2HG0107900.1 | -0.107317 | -0.242271 | -0.3146779 | cluster 1 |
| HORVU.MOREX.r3.2HG0110060.1 | -0.177789 | -0.262715 | -0.5774648 | cluster 1 |
| HORVU.MOREX.r3.2HG0110130.1 | -0.130025 | -0.569712 | -1.0201166 | cluster 1 |
| HORVU.MOREX.r3.2HG0110580.1 | -0.251766 | -0.363459 | -0.5595737 | cluster 1 |
| HORVU.MOREX.r3.2HG0110730.1 | -0.108897 | -0.367931 | -0.4073033 | cluster 1 |
| HORVU.MOREX.r3.2HG0110840.1 | -0.137 | -0.392845 | -0.6225663 | cluster 1 |
| HORVU.MOREX.r3.2HG0110920.1 | 0.0093348 | -0.169307 | -1.8863476 | cluster 1 |
| HORVU.MOREX.r3.2HG0111130.1 | 0.248764 | -0.257083 | -0.7758288 | cluster 1 |
| HORVU.MOREX.r3.2HG0113700.1 | 0.0212599 | -0.227534 | -1.1215831 | cluster 1 |
| HORVU.MOREX.r3.2HG0114840.1 | -0.228084 | -0.269203 | -0.6671363 | cluster 1 |
| HORVU.MOREX.r3.2HG0115300.2 | -0.083328 | -0.152473 | -0.2807162 | cluster 1 |
| HORVU.MOREX.r3.2HG0115480.1 | -0.286427 | -0.519993 | -0.6926657 | cluster 1 |

**Table S3** Continued.

| ID | logFC_3 h | logFC_6 h | logFC_12 h | cluster No. |
|---|---|---|---|---|
| HORVU.MOREX.r3.2HG0115780.1 | 0.1134533 | -0.51977 | -1.1816623 | cluster 1 |
| HORVU.MOREX.r3.2HG0115930.1 | -0.214079 | -0.50204 | -0.6523408 | cluster 1 |
| HORVU.MOREX.r3.2HG0115970.1 | 0.0611034 | -0.476529 | -1.0576632 | cluster 1 |
| HORVU.MOREX.r3.2HG0116540.1 | -0.070387 | -0.39954 | -0.5996536 | cluster 1 |
| HORVU.MOREX.r3.2HG0116870.1 | -0.189529 | -0.50958 | -0.8254902 | cluster 1 |
| HORVU.MOREX.r3.2HG0118580.1 | -0.14647 | -0.713252 | -1.0614147 | cluster 1 |
| HORVU.MOREX.r3.2HG0119060.1 | -0.219285 | -0.371699 | -0.4720529 | cluster 1 |
| HORVU.MOREX.r3.2HG0119220.1 | -0.211415 | -0.463912 | -0.5456335 | cluster 1 |
| HORVU.MOREX.r3.2HG0120030.1 | -0.124486 | -0.636145 | -0.7238389 | cluster 1 |
| HORVU.MOREX.r3.2HG0121830.1 | -0.153995 | -0.246546 | -0.3361858 | cluster 1 |
| HORVU.MOREX.r3.2HG0122040.1 | -0.09104 | -0.22291 | -0.2625496 | cluster 1 |
| HORVU.MOREX.r3.2HG0122140.1 | -0.180866 | -0.40154 | -0.4767424 | cluster 1 |
| HORVU.MOREX.r3.2HG0123650.1 | -0.04694 | -0.182486 | -0.3902826 | cluster 1 |
| HORVU.MOREX.r3.2HG0124130.1 | -0.165196 | -0.426497 | -0.7236887 | cluster 1 |
| HORVU.MOREX.r3.2HG0124820.1 | -0.185655 | -0.418642 | -0.8536779 | cluster 1 |
| HORVU.MOREX.r3.2HG0124990.1 | -0.206806 | -0.368187 | -0.6340075 | cluster 1 |
| HORVU.MOREX.r3.2HG0125050.1 | -0.185359 | -0.329336 | -0.4519597 | cluster 1 |
| HORVU.MOREX.r3.2HG0125780.1 | -0.110717 | -0.671355 | -1.2601952 | cluster 1 |
| HORVU.MOREX.r3.2HG0125870.1 | -0.064663 | -0.194271 | -0.5130197 | cluster 1 |
| HORVU.MOREX.r3.2HG0126010.1 | -0.075528 | -0.211478 | -0.3559582 | cluster 1 |
| HORVU.MOREX.r3.2HG0127530.1 | -0.352958 | -0.897155 | -1.488231 | cluster 1 |
| HORVU.MOREX.r3.2HG0128230.1 | 0.025941 | -0.342984 | -0.9415368 | cluster 1 |
| HORVU.MOREX.r3.2HG0128840.1 | -0.052212 | -0.280712 | -0.5947437 | cluster 1 |
| HORVU.MOREX.r3.2HG0128890.1 | -0.203016 | -0.367148 | -0.683621 | cluster 1 |
| HORVU.MOREX.r3.2HG0129200.1 | -0.083783 | -0.139018 | -0.5883573 | cluster 1 |
| HORVU.MOREX.r3.2HG0130000.1 | -0.037357 | -0.228341 | -0.3509321 | cluster 1 |
| HORVU.MOREX.r3.2HG0130180.1 | -0.104604 | -0.237342 | -0.3030084 | cluster 1 |
| HORVU.MOREX.r3.2HG0130200.1 | -0.172676 | -0.389335 | -0.4279482 | cluster 1 |
| HORVU.MOREX.r3.2HG0130770.1 | 0.0153705 | -0.243458 | -0.5732018 | cluster 1 |
| HORVU.MOREX.r3.2HG0132440.2 | -0.127483 | -0.138168 | -0.4546447 | cluster 1 |
| HORVU.MOREX.r3.2HG0132700.1 | -0.152424 | -0.579105 | -1.795339 | cluster 1 |
| HORVU.MOREX.r3.2HG0134220.1 | -0.054179 | -0.284603 | -0.3339927 | cluster 1 |
| HORVU.MOREX.r3.2HG0134430.1 | -0.529686 | -1.333204 | -2.2379963 | cluster 1 |
| HORVU.MOREX.r3.2HG0134700.1 | -0.092173 | -0.502229 | -0.7719558 | cluster 1 |
| HORVU.MOREX.r3.2HG0136570.1 | -0.306474 | -0.455575 | -2.1497726 | cluster 1 |
| HORVU.MOREX.r3.2HG0136670.1 | -0.088922 | -0.986027 | -1.9058988 | cluster 1 |
| HORVU.MOREX.r3.2HG0137670.1 | -0.401046 | -0.980536 | -1.0807166 | cluster 1 |
| HORVU.MOREX.r3.2HG0138280.1 | -0.037415 | -0.155592 | -0.6220855 | cluster 1 |
| HORVU.MOREX.r3.2HG0138300.1 | -0.055584 | -0.352456 | -0.7087585 | cluster 1 |
| HORVU.MOREX.r3.2HG0138620.1 | -0.404916 | -0.546314 | -1.116241 | cluster 1 |
| HORVU.MOREX.r3.2HG0139050.1 | 0.0930897 | -0.406323 | -1.0093808 | cluster 1 |
| HORVU.MOREX.r3.2HG0139370.1 | 0.0264594 | -0.36251 | -0.7941697 | cluster 1 |
| HORVU.MOREX.r3.2HG0139910.1 | -0.047241 | -0.497112 | -1.0090869 | cluster 1 |
| HORVU.MOREX.r3.2HG0141490.1 | -0.263341 | -0.484516 | -0.6681454 | cluster 1 |
| HORVU.MOREX.r3.2HG0146580.1 | -0.156103 | -0.36425 | -0.5505038 | cluster 1 |
| HORVU.MOREX.r3.2HG0147770.1 | -0.163058 | -0.281204 | -0.4436069 | cluster 1 |
| HORVU.MOREX.r3.2HG0149120.1 | -0.153745 | -0.188882 | -0.3032045 | cluster 1 |

**Table S3** Continued.

| ID | logFC_3 h | logFC_6 h | logFC_12 h | cluster No. |
|---|---|---|---|---|
| HORVU.MOREX.r3.2HG0149940.1 | -0.196568 | -0.409185 | -0.4708896 | cluster 1 |
| HORVU.MOREX.r3.2HG0150450.1 | -0.188254 | -0.476474 | -0.6592005 | cluster 1 |
| HORVU.MOREX.r3.2HG0151910.1 | -0.196639 | -0.697255 | -1.0244033 | cluster 1 |
| HORVU.MOREX.r3.2HG0151920.1 | -0.14807 | -0.560621 | -0.937555 | cluster 1 |
| HORVU.MOREX.r3.2HG0152140.1 | -0.09649 | -0.268293 | -0.6634219 | cluster 1 |
| HORVU.MOREX.r3.2HG0152370.1 | -0.071849 | -0.420731 | -0.6082781 | cluster 1 |
| HORVU.MOREX.r3.2HG0154740.1 | -0.512702 | -0.679296 | -0.7835556 | cluster 1 |
| HORVU.MOREX.r3.2HG0154760.1 | -0.133005 | -0.598735 | -0.9130175 | cluster 1 |
| HORVU.MOREX.r3.2HG0155450.1 | -0.15577 | -0.565461 | -0.8543335 | cluster 1 |
| HORVU.MOREX.r3.2HG0155460.1 | -0.172534 | -0.451907 | -0.6529114 | cluster 1 |
| HORVU.MOREX.r3.2HG0155810.1 | -0.270665 | -0.661728 | -0.9430821 | cluster 1 |
| HORVU.MOREX.r3.2HG0156590.1 | -0.053559 | -0.241695 | -0.4326334 | cluster 1 |
| HORVU.MOREX.r3.2HG0157160.1 | -0.307624 | -0.314861 | -0.835145 | cluster 1 |
| HORVU.MOREX.r3.2HG0157290.1 | -0.260212 | -0.678873 | -1.0739248 | cluster 1 |
| HORVU.MOREX.r3.2HG0158020.1 | -0.219433 | -0.330355 | -0.411972 | cluster 1 |
| HORVU.MOREX.r3.2HG0158620.1 | -0.01903 | -0.274628 | -0.4212415 | cluster 1 |
| HORVU.MOREX.r3.2HG0159500.1 | -0.174523 | -0.439394 | -0.7588238 | cluster 1 |
| HORVU.MOREX.r3.2HG0159540.1 | -0.613017 | -0.82915 | -1.0129392 | cluster 1 |
| HORVU.MOREX.r3.2HG0159920.1 | -0.213478 | -0.385033 | -0.5645933 | cluster 1 |
| HORVU.MOREX.r3.2HG0160550.1 | -0.274036 | -0.794337 | -2.716791 | cluster 1 |
| HORVU.MOREX.r3.2HG0160560.1 | -0.530672 | -1.321614 | -1.4680875 | cluster 1 |
| HORVU.MOREX.r3.2HG0160700.1 | 2.0996491 | 2.049175 | 1.32696119 | cluster 1 |
| HORVU.MOREX.r3.2HG0160790.1 | -0.02349 | -0.09619 | -0.5278546 | cluster 1 |
| HORVU.MOREX.r3.2HG0161020.1 | -0.171999 | -0.34568 | -0.8661492 | cluster 1 |
| HORVU.MOREX.r3.2HG0161670.1 | -0.133304 | -0.333342 | -0.5223391 | cluster 1 |
| HORVU.MOREX.r3.2HG0162940.1 | -0.168547 | -0.21352 | -0.3300561 | cluster 1 |
| HORVU.MOREX.r3.2HG0163090.1 | 0.1544503 | -0.235319 | -0.8334186 | cluster 1 |
| HORVU.MOREX.r3.2HG0164650.1 | -0.052745 | -0.199233 | -0.4735161 | cluster 1 |
| HORVU.MOREX.r3.2HG0164700.2 | -0.130199 | -0.361979 | -0.6727413 | cluster 1 |
| HORVU.MOREX.r3.2HG0165250.1 | -0.180908 | -0.342351 | -0.4033817 | cluster 1 |
| HORVU.MOREX.r3.2HG0165840.1 | -0.127416 | -0.559245 | -0.7276125 | cluster 1 |
| HORVU.MOREX.r3.2HG0167500.2 | 0.0473956 | -0.184061 | -1.671877 | cluster 1 |
| HORVU.MOREX.r3.2HG0167570.1 | -0.091062 | -0.569095 | -1.3170208 | cluster 1 |
| HORVU.MOREX.r3.2HG0167620.1 | -0.186775 | -0.242001 | -1.2516465 | cluster 1 |
| HORVU.MOREX.r3.2HG0168010.1 | -0.190579 | -0.340969 | -0.5559603 | cluster 1 |
| HORVU.MOREX.r3.2HG0168090.1 | -0.148237 | -0.339162 | -0.479939 | cluster 1 |
| HORVU.MOREX.r3.2HG0168270.1 | -0.168038 | -0.535135 | -0.6121322 | cluster 1 |
| HORVU.MOREX.r3.2HG0168730.1 | -0.156901 | -0.280808 | -0.5281516 | cluster 1 |
| HORVU.MOREX.r3.2HG0168790.1 | -0.218259 | -0.455337 | -0.7242725 | cluster 1 |
| HORVU.MOREX.r3.2HG0169700.1 | -0.171016 | -0.425236 | -0.7408696 | cluster 1 |
| HORVU.MOREX.r3.2HG0169720.1 | -0.069796 | -0.136412 | -0.4961491 | cluster 1 |
| HORVU.MOREX.r3.2HG0170300.1 | -0.229741 | -0.425032 | -0.5693042 | cluster 1 |
| HORVU.MOREX.r3.2HG0170390.1 | 0.0210103 | -0.194068 | -0.3191257 | cluster 1 |
| HORVU.MOREX.r3.2HG0170560.1 | -0.169778 | -0.244654 | -1.2182252 | cluster 1 |
| HORVU.MOREX.r3.2HG0170880.1 | -0.140018 | -0.456729 | -1.3017766 | cluster 1 |
| HORVU.MOREX.r3.2HG0171550.1 | -0.104165 | -0.64018 | -0.8125073 | cluster 1 |
| HORVU.MOREX.r3.2HG0171860.1 | -0.172697 | -0.260442 | -0.618192 | cluster 1 |

**Table S3** Continued.

| ID | logFC_3 h | logFC_6 h | logFC_12 h | cluster No. |
|---|---|---|---|---|
| HORVU.MOREX.r3.2HG0172050.1 | -0.073516 | -0.095321 | -0.3565514 | cluster 1 |
| HORVU.MOREX.r3.2HG0172400.1 | 0.0232819 | -0.885943 | -1.7502537 | cluster 1 |
| HORVU.MOREX.r3.2HG0172730.1 | -0.16599 | -0.433281 | -0.532916 | cluster 1 |
| HORVU.MOREX.r3.2HG0172790.1 | -0.111495 | -0.465494 | -0.8142454 | cluster 1 |
| HORVU.MOREX.r3.2HG0173190.1 | -0.248269 | -0.456951 | -1.4397215 | cluster 1 |
| HORVU.MOREX.r3.2HG0173210.1 | -0.211964 | -0.43618 | -0.8921819 | cluster 1 |
| HORVU.MOREX.r3.2HG0173340.1 | -0.672694 | -1.024944 | -3.4011705 | cluster 1 |
| HORVU.MOREX.r3.2HG0173550.1 | -0.12745 | -0.142497 | -0.6657476 | cluster 1 |
| HORVU.MOREX.r3.2HG0173680.1 | -0.065129 | -0.212515 | -0.4291996 | cluster 1 |
| HORVU.MOREX.r3.2HG0174820.1 | -0.026387 | -0.315706 | -0.4958504 | cluster 1 |
| HORVU.MOREX.r3.2HG0175320.1 | -0.278855 | -0.341635 | -0.8507718 | cluster 1 |
| HORVU.MOREX.r3.2HG0176290.1 | -0.132214 | -0.213024 | -0.3801937 | cluster 1 |
| HORVU.MOREX.r3.2HG0176680.1 | -0.046792 | -0.118489 | -0.2374491 | cluster 1 |
| HORVU.MOREX.r3.2HG0176920.1 | -0.262881 | -0.359843 | -1.2359358 | cluster 1 |
| HORVU.MOREX.r3.2HG0177200.1 | 0.027253 | -0.431642 | -0.5373577 | cluster 1 |
| HORVU.MOREX.r3.2HG0177440.1 | -0.278161 | -0.547659 | -0.8454242 | cluster 1 |
| HORVU.MOREX.r3.2HG0178160.1 | -0.184445 | -0.569243 | -0.6504268 | cluster 1 |
| HORVU.MOREX.r3.2HG0179410.1 | -0.115718 | -0.364064 | -0.569688 | cluster 1 |
| HORVU.MOREX.r3.2HG0179540.1 | -0.124311 | -0.150309 | -0.4105338 | cluster 1 |
| HORVU.MOREX.r3.2HG0180060.1 | -0.137065 | -0.664672 | -1.211109 | cluster 1 |
| HORVU.MOREX.r3.2HG0180090.1 | -0.041684 | -0.385505 | -0.596626 | cluster 1 |
| HORVU.MOREX.r3.2HG0180210.1 | -0.109261 | -0.643828 | -1.2018793 | cluster 1 |
| HORVU.MOREX.r3.2HG0180260.1 | -0.405119 | -0.522241 | -0.698624 | cluster 1 |
| HORVU.MOREX.r3.2HG0180580.1 | -0.110076 | -0.40217 | -1.4221313 | cluster 1 |
| HORVU.MOREX.r3.2HG0180880.1 | -0.193847 | -0.624619 | -0.9009626 | cluster 1 |
| HORVU.MOREX.r3.2HG0181390.1 | -0.216082 | -0.524876 | -1.4964878 | cluster 1 |
| HORVU.MOREX.r3.2HG0181660.1 | -0.160828 | -0.406047 | -0.8346062 | cluster 1 |
| HORVU.MOREX.r3.2HG0181800.1 | -0.349773 | -0.599705 | -0.7995019 | cluster 1 |
| HORVU.MOREX.r3.2HG0182020.2 | -0.540107 | -0.692229 | -1.3458377 | cluster 1 |
| HORVU.MOREX.r3.2HG0182280.1 | -0.236545 | -0.506593 | -0.7276101 | cluster 1 |
| HORVU.MOREX.r3.2HG0182800.1 | -0.314825 | -0.69223 | -1.1727827 | cluster 1 |
| HORVU.MOREX.r3.2HG0183740.1 | -0.160999 | -0.625029 | -1.2185225 | cluster 1 |
| HORVU.MOREX.r3.2HG0183780.1 | -0.141197 | -0.437104 | -0.5578557 | cluster 1 |
| HORVU.MOREX.r3.2HG0184620.1 | -0.542433 | -1.007238 | -1.0762834 | cluster 1 |
| HORVU.MOREX.r3.2HG0184780.1 | -0.190778 | -0.443959 | -0.7601437 | cluster 1 |
| HORVU.MOREX.r3.2HG0185840.1 | -0.150161 | -0.435802 | -0.9607608 | cluster 1 |
| HORVU.MOREX.r3.2HG0185850.1 | -0.047482 | -0.281538 | -0.4626977 | cluster 1 |
| HORVU.MOREX.r3.2HG0186650.1 | -0.173337 | -0.225558 | -0.7370369 | cluster 1 |
| HORVU.MOREX.r3.2HG0186680.1 | -0.220169 | -0.500706 | -0.6940226 | cluster 1 |
| HORVU.MOREX.r3.2HG0187440.1 | -0.113755 | -0.357394 | -0.5374455 | cluster 1 |
| HORVU.MOREX.r3.2HG0187640.1 | -0.344579 | -0.492267 | -0.7954147 | cluster 1 |
| HORVU.MOREX.r3.2HG0188300.1 | -0.143614 | -0.425722 | -0.6026267 | cluster 1 |
| HORVU.MOREX.r3.2HG0189800.1 | -0.27634 | -0.509165 | -0.6867791 | cluster 1 |
| HORVU.MOREX.r3.2HG0189940.1 | -0.13915 | -0.532157 | -1.3204146 | cluster 1 |
| HORVU.MOREX.r3.2HG0189950.1 | -0.047778 | -0.166634 | -0.4376899 | cluster 1 |
| HORVU.MOREX.r3.2HG0190220.1 | -0.184515 | -0.591466 | -0.998013 | cluster 1 |
| HORVU.MOREX.r3.2HG0190230.1 | -0.251482 | -0.587172 | -1.4737691 | cluster 1 |

**Table S3** Continued.

| ID | logFC_3 h | logFC_6 h | logFC_12 h | cluster No. |
|---|---|---|---|---|
| HORVU.MOREX.r3.2HG0190470.1 | -0.186053 | -0.466043 | -0.5873399 | cluster 1 |
| HORVU.MOREX.r3.2HG0190680.1 | -0.165902 | -0.254017 | -0.4810851 | cluster 1 |
| HORVU.MOREX.r3.2HG0190710.1 | -0.103241 | -0.251953 | -0.343677 | cluster 1 |
| HORVU.MOREX.r3.2HG0192900.1 | -0.14576 | -0.608535 | -1.3209815 | cluster 1 |
| HORVU.MOREX.r3.2HG0193080.1 | -0.115497 | -0.213986 | -0.4413659 | cluster 1 |
| HORVU.MOREX.r3.2HG0193220.1 | -0.21748 | -0.279497 | -0.6454699 | cluster 1 |
| HORVU.MOREX.r3.2HG0193930.1 | -0.118934 | -0.462256 | -0.6992255 | cluster 1 |
| HORVU.MOREX.r3.2HG0194450.1 | -0.065113 | -0.18727 | -0.381859 | cluster 1 |
| HORVU.MOREX.r3.2HG0194840.1 | -0.156688 | -1.263726 | -1.8597546 | cluster 1 |
| HORVU.MOREX.r3.2HG0195110.2 | -0.379744 | -0.677136 | -1.8754575 | cluster 1 |
| HORVU.MOREX.r3.2HG0195190.1 | -0.128876 | -0.328486 | -0.8708187 | cluster 1 |
| HORVU.MOREX.r3.2HG0197560.1 | -0.265836 | -0.500201 | -0.7677648 | cluster 1 |
| HORVU.MOREX.r3.2HG0197680.1 | -0.143467 | -0.163153 | -0.3968303 | cluster 1 |
| HORVU.MOREX.r3.2HG0197710.1 | -0.242123 | -0.711377 | -1.1581298 | cluster 1 |
| HORVU.MOREX.r3.2HG0199520.1 | -0.231423 | -1.19727 | -1.6691928 | cluster 1 |
| HORVU.MOREX.r3.2HG0199560.1 | -0.10429 | -0.706589 | -1.8280722 | cluster 1 |
| HORVU.MOREX.r3.2HG0199590.1 | -0.570944 | -1.056623 | -1.2647195 | cluster 1 |
| HORVU.MOREX.r3.2HG0199600.1 | -0.493 | -1.489763 | -2.217265 | cluster 1 |
| HORVU.MOREX.r3.2HG0200630.1 | -0.109808 | -0.355475 | -0.7775816 | cluster 1 |
| HORVU.MOREX.r3.2HG0200640.1 | -0.16703 | -0.256722 | -0.5216467 | cluster 1 |
| HORVU.MOREX.r3.2HG0200980.1 | -0.045959 | -0.171844 | -0.4061348 | cluster 1 |
| HORVU.MOREX.r3.2HG0203500.1 | -0.123255 | -0.239296 | -0.8256111 | cluster 1 |
| HORVU.MOREX.r3.2HG0204920.1 | -0.162813 | -0.572041 | -1.3845879 | cluster 1 |
| HORVU.MOREX.r3.2HG0204940.1 | -0.160487 | -0.496076 | -0.6690274 | cluster 1 |
| HORVU.MOREX.r3.2HG0204960.1 | -0.182984 | -0.28201 | -0.3312112 | cluster 1 |
| HORVU.MOREX.r3.2HG0205200.1 | -0.211409 | -0.457111 | -0.7688419 | cluster 1 |
| HORVU.MOREX.r3.2HG0205550.1 | -0.201367 | -0.426763 | -0.8019225 | cluster 1 |
| HORVU.MOREX.r3.2HG0205750.1 | -0.195928 | -0.456015 | -0.6426124 | cluster 1 |
| HORVU.MOREX.r3.2HG0206910.1 | -0.196755 | -0.517186 | -1.0029526 | cluster 1 |
| HORVU.MOREX.r3.2HG0208790.1 | -0.349134 | -1.197783 | -1.6687659 | cluster 1 |
| HORVU.MOREX.r3.2HG0209300.1 | -0.092672 | -0.631859 | -1.1824651 | cluster 1 |
| HORVU.MOREX.r3.2HG0209440.1 | -0.115503 | -0.221954 | -0.4802608 | cluster 1 |
| HORVU.MOREX.r3.2HG0209910.1 | 0.0546079 | -0.296826 | -0.4154749 | cluster 1 |
| HORVU.MOREX.r3.2HG0210420.1 | -0.154001 | -0.360597 | -0.5717427 | cluster 1 |
| HORVU.MOREX.r3.2HG0210880.1 | 0.1371645 | 0.0302801 | -1.1557906 | cluster 1 |
| HORVU.MOREX.r3.2HG0212570.1 | -0.193354 | -0.405159 | -0.5208923 | cluster 1 |
| HORVU.MOREX.r3.2HG0212900.1 | -0.060892 | -0.240509 | -0.6129391 | cluster 1 |
| HORVU.MOREX.r3.2HG0213250.1 | -0.15275 | -0.538482 | -0.7092201 | cluster 1 |
| HORVU.MOREX.r3.2HG0213850.1 | -0.011557 | -0.135704 | -0.6079445 | cluster 1 |
| HORVU.MOREX.r3.2HG0214090.1 | -0.194082 | -0.541106 | -1.0790097 | cluster 1 |
| HORVU.MOREX.r3.2HG0215220.1 | 0.3156464 | -0.337822 | -0.7494649 | cluster 1 |
| HORVU.MOREX.r3.2HG0215250.1 | 0.1623403 | -0.523518 | -1.1321166 | cluster 1 |
| HORVU.MOREX.r3.2HG0215280.1 | -0.031021 | -0.358731 | -0.5720262 | cluster 1 |
| HORVU.MOREX.r3.2HG0215310.1 | 0.1124949 | -0.300838 | -1.4518362 | cluster 1 |
| HORVU.MOREX.r3.2HG0215550.1 | 0.1403848 | -0.018232 | -0.8009071 | cluster 1 |
| HORVU.MOREX.r3.2HG0216610.1 | -0.122363 | -0.279792 | -0.3320576 | cluster 1 |
| HORVU.MOREX.r3.2HG0217610.1 | -0.040075 | -0.21691 | -0.3070362 | cluster 1 |

**Table S3** Continued.

| ID | logFC_3 h | logFC_6 h | logFC_12 h | cluster No. |
|---|---|---|---|---|
| HORVU.MOREX.r3.2HG0218010.1 | -0.14466 | -0.383605 | -0.4903498 | cluster 1 |
| HORVU.MOREX.r3.3HG0218100.1 | -0.125372 | -0.330788 | -0.3622098 | cluster 1 |
| HORVU.MOREX.r3.3HG0218330.1 | 0.0072857 | -0.56199 | -1.5749569 | cluster 1 |
| HORVU.MOREX.r3.3HG0219290.1 | -0.267742 | -0.279616 | -0.4249593 | cluster 1 |
| HORVU.MOREX.r3.3HG0219650.1 | -0.070838 | -0.15453 | -0.4423778 | cluster 1 |
| HORVU.MOREX.r3.3HG0219740.1 | -0.172104 | -0.317892 | -0.6152745 | cluster 1 |
| HORVU.MOREX.r3.3HG0219800.1 | -0.029697 | -0.262911 | -0.542665 | cluster 1 |
| HORVU.MOREX.r3.3HG0219810.1 | -0.37275 | -0.442903 | -0.9179098 | cluster 1 |
| HORVU.MOREX.r3.3HG0220250.1 | -0.221401 | -0.34977 | -0.4856685 | cluster 1 |
| HORVU.MOREX.r3.3HG0220360.1 | -0.126673 | -0.496967 | -0.8243858 | cluster 1 |
| HORVU.MOREX.r3.3HG0221460.1 | -0.090241 | -0.098305 | -0.5980666 | cluster 1 |
| HORVU.MOREX.r3.3HG0223950.1 | 0.0829855 | -0.268924 | -0.6635622 | cluster 1 |
| HORVU.MOREX.r3.3HG0223980.1 | -0.22551 | -0.349647 | -0.4959089 | cluster 1 |
| HORVU.MOREX.r3.3HG0224680.1 | -0.072198 | -0.281699 | -0.4108601 | cluster 1 |
| HORVU.MOREX.r3.3HG0225610.1 | -0.288524 | -0.328496 | -0.8636404 | cluster 1 |
| HORVU.MOREX.r3.3HG0226060.1 | -0.229348 | -0.5027 | -1.2634158 | cluster 1 |
| HORVU.MOREX.r3.3HG0229480.1 | -0.093827 | -0.159974 | -0.3893624 | cluster 1 |
| HORVU.MOREX.r3.3HG0230190.1 | -0.236364 | -0.356318 | -1.1796632 | cluster 1 |
| HORVU.MOREX.r3.3HG0230970.1 | -0.109852 | -0.438205 | -0.8163375 | cluster 1 |
| HORVU.MOREX.r3.3HG0231050.1 | -0.131468 | -0.479898 | -0.8809321 | cluster 1 |
| HORVU.MOREX.r3.3HG0231630.1 | -0.13687 | -0.730602 | -1.466273 | cluster 1 |
| HORVU.MOREX.r3.3HG0231780.1 | -0.119593 | -0.437619 | -1.0611853 | cluster 1 |
| HORVU.MOREX.r3.3HG0231900.1 | -0.055668 | -0.187305 | -0.2424899 | cluster 1 |
| HORVU.MOREX.r3.3HG0232780.1 | 0.0663294 | -0.434466 | -0.5689191 | cluster 1 |
| HORVU.MOREX.r3.3HG0233150.1 | -0.096706 | -0.277548 | -0.8277078 | cluster 1 |
| HORVU.MOREX.r3.3HG0234560.1 | -0.134236 | -1.078676 | -1.3408735 | cluster 1 |
| HORVU.MOREX.r3.3HG0234800.1 | -0.124679 | -0.221639 | -0.2789724 | cluster 1 |
| HORVU.MOREX.r3.3HG0236170.1 | -0.055837 | -0.20051 | -0.3508077 | cluster 1 |
| HORVU.MOREX.r3.3HG0237870.1 | -0.103412 | -0.576425 | -1.6593224 | cluster 1 |
| HORVU.MOREX.r3.3HG0237910.1 | -0.354448 | -0.708414 | -0.9922342 | cluster 1 |
| HORVU.MOREX.r3.3HG0238180.1 | -0.245973 | -0.494704 | -0.5746101 | cluster 1 |
| HORVU.MOREX.r3.3HG0239110.1 | -0.354672 | -0.652166 | -0.9254847 | cluster 1 |
| HORVU.MOREX.r3.3HG0240390.2 | -0.662318 | -0.76606 | -1.2581255 | cluster 1 |
| HORVU.MOREX.r3.3HG0240450.1 | -0.158411 | -0.40163 | -0.7534721 | cluster 1 |
| HORVU.MOREX.r3.3HG0240640.1 | -0.139937 | -0.693355 | -1.4793637 | cluster 1 |
| HORVU.MOREX.r3.3HG0240670.1 | -0.177471 | -0.451187 | -1.1615878 | cluster 1 |
| HORVU.MOREX.r3.3HG0240710.1 | -0.253345 | -0.801872 | -1.4033411 | cluster 1 |
| HORVU.MOREX.r3.3HG0241000.1 | -0.23877 | -1.145729 | -1.7322621 | cluster 1 |
| HORVU.MOREX.r3.3HG0241630.1 | -0.220412 | -0.504613 | -0.6515102 | cluster 1 |
| HORVU.MOREX.r3.3HG0241720.1 | -0.03839 | -0.157978 | -0.1934811 | cluster 1 |
| HORVU.MOREX.r3.3HG0242030.1 | -0.059841 | -0.154442 | -1.6653751 | cluster 1 |
| HORVU.MOREX.r3.3HG0242520.1 | -0.098191 | -0.399811 | -0.5407455 | cluster 1 |
| HORVU.MOREX.r3.3HG0242870.1 | -0.116097 | -0.292479 | -0.3235167 | cluster 1 |
| HORVU.MOREX.r3.3HG0244690.1 | -0.080686 | -0.489595 | -0.8390583 | cluster 1 |
| HORVU.MOREX.r3.3HG0245310.1 | 0.0273886 | -0.405969 | -0.7475374 | cluster 1 |
| HORVU.MOREX.r3.3HG0246370.1 | -0.25615 | -0.432403 | -0.6589591 | cluster 1 |
| HORVU.MOREX.r3.3HG0246560.1 | -0.189221 | -0.513876 | -0.7699912 | cluster 1 |

**Table S3** Continued.

| ID | logFC_3 h | logFC_6 h | logFC_12 h | cluster No. |
|---|---|---|---|---|
| HORVU.MOREX.r3.3HG0247250.1 | -0.141916 | -0.283166 | -1.3240752 | cluster 1 |
| HORVU.MOREX.r3.3HG0247260.1 | -0.36892 | -0.888752 | -1.1693937 | cluster 1 |
| HORVU.MOREX.r3.3HG0248280.1 | -0.139838 | -0.216864 | -0.3936771 | cluster 1 |
| HORVU.MOREX.r3.3HG0249590.1 | -0.375729 | -0.430333 | -0.6995825 | cluster 1 |
| HORVU.MOREX.r3.3HG0249700.1 | -0.133678 | -0.457757 | -0.5532478 | cluster 1 |
| HORVU.MOREX.r3.3HG0249810.1 | -0.129046 | -0.145227 | -0.3999752 | cluster 1 |
| HORVU.MOREX.r3.3HG0250170.1 | 0.0153043 | -0.131782 | -0.5871409 | cluster 1 |
| HORVU.MOREX.r3.3HG0250630.1 | -0.254622 | -0.537133 | -0.6985942 | cluster 1 |
| HORVU.MOREX.r3.3HG0251830.1 | -0.356283 | -0.391232 | -1.1215397 | cluster 1 |
| HORVU.MOREX.r3.3HG0251950.1 | -0.242722 | -0.458711 | -0.6915756 | cluster 1 |
| HORVU.MOREX.r3.3HG0252240.1 | -0.131064 | -0.396222 | -0.9017322 | cluster 1 |
| HORVU.MOREX.r3.3HG0252610.1 | -0.160318 | -0.337298 | -0.3771383 | cluster 1 |
| HORVU.MOREX.r3.3HG0253360.1 | -0.475094 | -0.541572 | -1.0775016 | cluster 1 |
| HORVU.MOREX.r3.3HG0253440.1 | -0.032487 | -0.363593 | -0.8320019 | cluster 1 |
| HORVU.MOREX.r3.3HG0254850.1 | -0.015903 | -0.166361 | -0.3370218 | cluster 1 |
| HORVU.MOREX.r3.3HG0255040.1 | -0.225901 | -0.373302 | -0.4099441 | cluster 1 |
| HORVU.MOREX.r3.3HG0255150.1 | -0.146222 | -0.226422 | -0.2733467 | cluster 1 |
| HORVU.MOREX.r3.3HG0255220.1 | -0.241422 | -0.543248 | -0.998544 | cluster 1 |
| HORVU.MOREX.r3.3HG0255900.1 | -0.134357 | -0.201523 | -0.2914825 | cluster 1 |
| HORVU.MOREX.r3.3HG0256300.1 | -0.108122 | -0.265612 | -0.4583029 | cluster 1 |
| HORVU.MOREX.r3.3HG0256360.1 | -0.087076 | -0.280291 | -0.374385 | cluster 1 |
| HORVU.MOREX.r3.3HG0256380.1 | -0.140095 | -0.328872 | -0.8566649 | cluster 1 |
| HORVU.MOREX.r3.3HG0256580.1 | -0.136381 | -0.254027 | -0.4907328 | cluster 1 |
| HORVU.MOREX.r3.3HG0256620.1 | -0.118856 | -0.728438 | -1.0254593 | cluster 1 |
| HORVU.MOREX.r3.3HG0257130.1 | -0.283517 | -0.506689 | -0.7760597 | cluster 1 |
| HORVU.MOREX.r3.3HG0259660.1 | -0.191272 | -0.441023 | -0.5892941 | cluster 1 |
| HORVU.MOREX.r3.3HG0259960.1 | 0.0744353 | -1.297706 | -1.9686553 | cluster 1 |
| HORVU.MOREX.r3.3HG0260580.1 | -0.051405 | -0.162657 | -0.4759776 | cluster 1 |
| HORVU.MOREX.r3.3HG0261980.1 | -0.034974 | -0.271694 | -0.5902238 | cluster 1 |
| HORVU.MOREX.r3.3HG0262210.1 | -0.168114 | -0.280944 | -0.328426 | cluster 1 |
| HORVU.MOREX.r3.3HG0266960.1 | -0.131143 | -0.289426 | -0.6094999 | cluster 1 |
| HORVU.MOREX.r3.3HG0267150.1 | -0.082957 | -0.288586 | -0.352175 | cluster 1 |
| HORVU.MOREX.r3.3HG0270070.1 | -0.194179 | -0.415184 | -0.8724205 | cluster 1 |
| HORVU.MOREX.r3.3HG0270400.1 | -0.301809 | -0.518499 | -0.8898794 | cluster 1 |
| HORVU.MOREX.r3.3HG0270500.1 | -0.220232 | -0.482652 | -0.5737567 | cluster 1 |
| HORVU.MOREX.r3.3HG0270910.1 | -0.072651 | -0.253474 | -0.2953239 | cluster 1 |
| HORVU.MOREX.r3.3HG0272160.2 | -0.274508 | -0.465267 | -0.7060044 | cluster 1 |
| HORVU.MOREX.r3.3HG0272430.1 | -0.056853 | -0.165352 | -0.4036842 | cluster 1 |
| HORVU.MOREX.r3.3HG0275530.1 | -0.113086 | -0.307145 | -0.7536933 | cluster 1 |
| HORVU.MOREX.r3.3HG0277100.1 | -0.127665 | -0.419388 | -0.4620881 | cluster 1 |
| HORVU.MOREX.r3.3HG0277200.1 | -0.272808 | -0.680377 | -1.1215708 | cluster 1 |
| HORVU.MOREX.r3.3HG0279760.1 | -0.503867 | -0.552255 | -0.7878619 | cluster 1 |
| HORVU.MOREX.r3.3HG0280310.1 | -0.212239 | -0.495671 | -0.6812373 | cluster 1 |
| HORVU.MOREX.r3.3HG0280900.1 | -0.036837 | -0.190958 | -0.3347234 | cluster 1 |
| HORVU.MOREX.r3.3HG0281000.1 | -0.094973 | -0.223989 | -0.8512444 | cluster 1 |
| HORVU.MOREX.r3.3HG0282170.1 | -0.577209 | -0.948347 | -1.825963 | cluster 1 |
| HORVU.MOREX.r3.3HG0283990.1 | -0.117159 | -0.316105 | -0.525227 | cluster 1 |

**Table S3** Continued.

| ID | logFC_3 h | logFC_6 h | logFC_12 h | cluster No. |
|---|---|---|---|---|
| HORVU.MOREX.r3.3HG0284170.1 | -0.144863 | -0.489183 | -0.5758377 | cluster 1 |
| HORVU.MOREX.r3.3HG0285010.1 | -0.765206 | -0.943498 | -2.0178816 | cluster 1 |
| HORVU.MOREX.r3.3HG0285270.1 | -0.335414 | -0.824703 | -2.0053754 | cluster 1 |
| HORVU.MOREX.r3.3HG0285570.1 | -0.162313 | -0.601252 | -0.7285259 | cluster 1 |
| HORVU.MOREX.r3.3HG0286160.1 | -0.292278 | -0.811066 | -1.7481436 | cluster 1 |
| HORVU.MOREX.r3.3HG0287070.1 | -0.133152 | -0.838915 | -1.2917272 | cluster 1 |
| HORVU.MOREX.r3.3HG0287800.1 | 0.1339816 | -0.221696 | -0.6885903 | cluster 1 |
| HORVU.MOREX.r3.3HG0287840.1 | -0.199367 | -0.611982 | -0.7099253 | cluster 1 |
| HORVU.MOREX.r3.3HG0288240.1 | -0.205668 | -0.310078 | -0.6990169 | cluster 1 |
| HORVU.MOREX.r3.3HG0288420.1 | 0.0100033 | -0.169455 | -0.9249703 | cluster 1 |
| HORVU.MOREX.r3.3HG0288700.1 | -0.274665 | -0.590487 | -0.6701073 | cluster 1 |
| HORVU.MOREX.r3.3HG0288940.1 | -0.263878 | -0.481211 | -0.7157122 | cluster 1 |
| HORVU.MOREX.r3.3HG0289910.1 | -0.055999 | -0.221969 | -0.3541359 | cluster 1 |
| HORVU.MOREX.r3.3HG0289970.1 | -0.162318 | -0.28301 | -0.6045168 | cluster 1 |
| HORVU.MOREX.r3.3HG0290180.1 | -0.256618 | -0.498685 | -1.1300931 | cluster 1 |
| HORVU.MOREX.r3.3HG0290510.1 | -0.049319 | -0.510759 | -0.8913259 | cluster 1 |
| HORVU.MOREX.r3.3HG0291490.1 | -0.197087 | -0.44142 | -0.9611394 | cluster 1 |
| HORVU.MOREX.r3.3HG0291710.1 | -0.237267 | -0.420506 | -0.4724722 | cluster 1 |
| HORVU.MOREX.r3.3HG0293040.1 | -0.189637 | -0.239068 | -0.4427811 | cluster 1 |
| HORVU.MOREX.r3.3HG0293700.1 | -0.213003 | -0.433337 | -0.6604125 | cluster 1 |
| HORVU.MOREX.r3.3HG0294100.1 | -0.166916 | -0.340918 | -0.3761033 | cluster 1 |
| HORVU.MOREX.r3.3HG0294840.1 | -0.045392 | -0.18564 | -0.5245908 | cluster 1 |
| HORVU.MOREX.r3.3HG0295020.1 | -0.084589 | -0.31783 | -0.56766 | cluster 1 |
| HORVU.MOREX.r3.3HG0295330.1 | -0.140062 | -0.405564 | -0.6056142 | cluster 1 |
| HORVU.MOREX.r3.3HG0295440.1 | -0.175854 | -0.374836 | -0.8104747 | cluster 1 |
| HORVU.MOREX.r3.3HG0295450.1 | -0.087159 | -0.537072 | -0.8221331 | cluster 1 |
| HORVU.MOREX.r3.3HG0296490.1 | -0.012781 | -0.103117 | -0.6310038 | cluster 1 |
| HORVU.MOREX.r3.3HG0296810.1 | -0.330369 | -0.442928 | -0.9073306 | cluster 1 |
| HORVU.MOREX.r3.3HG0296940.1 | -0.404681 | -0.438639 | -0.7780682 | cluster 1 |
| HORVU.MOREX.r3.3HG0297140.1 | -0.128292 | -0.188447 | -0.26881 | cluster 1 |
| HORVU.MOREX.r3.3HG0297180.1 | -0.591426 | -0.843079 | -1.7608999 | cluster 1 |
| HORVU.MOREX.r3.3HG0297940.1 | -0.06286 | -0.960919 | -1.1779962 | cluster 1 |
| HORVU.MOREX.r3.3HG0298340.1 | -0.286744 | -0.396123 | -0.738991 | cluster 1 |
| HORVU.MOREX.r3.3HG0298390.1 | -0.293623 | -0.523294 | -1.155846 | cluster 1 |
| HORVU.MOREX.r3.3HG0298750.1 | -0.033868 | -0.320112 | -0.4786808 | cluster 1 |
| HORVU.MOREX.r3.3HG0299010.1 | -0.382724 | -0.539569 | -1.071483 | cluster 1 |
| HORVU.MOREX.r3.3HG0299200.1 | -0.150773 | -0.358807 | -0.6619428 | cluster 1 |
| HORVU.MOREX.r3.3HG0299280.1 | -0.129571 | -0.361193 | -0.7325662 | cluster 1 |
| HORVU.MOREX.r3.3HG0299300.1 | 0.1040784 | -0.793205 | -1.1079439 | cluster 1 |
| HORVU.MOREX.r3.3HG0299530.1 | 0.1216835 | -0.313153 | -0.5168956 | cluster 1 |
| HORVU.MOREX.r3.3HG0299540.1 | -0.09918 | -0.221253 | -0.3040279 | cluster 1 |
| HORVU.MOREX.r3.3HG0299820.1 | -0.243889 | -0.577134 | -1.061206 | cluster 1 |
| HORVU.MOREX.r3.3HG0300020.1 | 0.2560973 | -0.681768 | -2.172499 | cluster 1 |
| HORVU.MOREX.r3.3HG0301250.1 | -0.268101 | -0.864757 | -1.1706077 | cluster 1 |
| HORVU.MOREX.r3.3HG0301750.1 | -0.10808 | -0.456495 | -0.6238556 | cluster 1 |
| HORVU.MOREX.r3.3HG0302090.1 | -0.06493 | -0.137075 | -0.586395 | cluster 1 |
| HORVU.MOREX.r3.3HG0302140.1 | -0.152563 | -0.312717 | -0.4769035 | cluster 1 |

**Table S3** Continued.

| ID | logFC_3 h | logFC_6 h | logFC_12 h | cluster No. |
|---|---|---|---|---|
| HORVU.MOREX.r3.3HG0302860.1 | 0.8047109 | -0.100145 | -1.9563874 | cluster 1 |
| HORVU.MOREX.r3.3HG0302910.1 | -0.114349 | -0.383689 | -0.9603979 | cluster 1 |
| HORVU.MOREX.r3.3HG0303330.1 | -0.121177 | -0.291992 | -0.8113502 | cluster 1 |
| HORVU.MOREX.r3.3HG0303430.1 | -0.133288 | -0.372844 | -0.4640951 | cluster 1 |
| HORVU.MOREX.r3.3HG0303970.1 | -0.074889 | -0.395173 | -0.6779429 | cluster 1 |
| HORVU.MOREX.r3.3HG0304080.1 | -0.019526 | -0.474277 | -0.8626337 | cluster 1 |
| HORVU.MOREX.r3.3HG0304420.1 | -0.136011 | -0.379394 | -1.0740291 | cluster 1 |
| HORVU.MOREX.r3.3HG0304640.1 | -0.039855 | -0.349929 | -0.5820753 | cluster 1 |
| HORVU.MOREX.r3.3HG0304710.1 | -0.117879 | -0.287969 | -0.346661 | cluster 1 |
| HORVU.MOREX.r3.3HG0305150.1 | -0.230037 | -0.69222 | -0.9446847 | cluster 1 |
| HORVU.MOREX.r3.3HG0305270.1 | -0.217125 | -0.295205 | -0.627447 | cluster 1 |
| HORVU.MOREX.r3.3HG0305500.1 | 0.1561274 | -0.222634 | -1.3145458 | cluster 1 |
| HORVU.MOREX.r3.3HG0305580.1 | 0.1242108 | -0.387167 | -1.3815488 | cluster 1 |
| HORVU.MOREX.r3.3HG0306030.1 | -0.105565 | -0.261625 | -0.4564221 | cluster 1 |
| HORVU.MOREX.r3.3HG0306170.1 | -0.055892 | -0.145347 | -0.32995 | cluster 1 |
| HORVU.MOREX.r3.3HG0306210.1 | -0.131027 | -0.215962 | -0.6006012 | cluster 1 |
| HORVU.MOREX.r3.3HG0306220.1 | -0.088388 | -0.408085 | -0.471634 | cluster 1 |
| HORVU.MOREX.r3.3HG0306420.1 | -0.327989 | -0.398514 | -1.0430224 | cluster 1 |
| HORVU.MOREX.r3.3HG0306440.1 | -0.16585 | -0.812018 | -1.166288 | cluster 1 |
| HORVU.MOREX.r3.3HG0306820.2 | -0.349779 | -0.866324 | -1.3258413 | cluster 1 |
| HORVU.MOREX.r3.3HG0307120.1 | -0.153053 | -0.361715 | -0.6358684 | cluster 1 |
| HORVU.MOREX.r3.3HG0308420.1 | 0.0914717 | -0.604517 | -0.8279024 | cluster 1 |
| HORVU.MOREX.r3.3HG0308820.1 | -0.115504 | -0.276578 | -0.3613674 | cluster 1 |
| HORVU.MOREX.r3.3HG0309410.1 | -0.517402 | -0.972681 | -1.3512535 | cluster 1 |
| HORVU.MOREX.r3.3HG0309460.1 | -0.074743 | -0.284965 | -0.3338557 | cluster 1 |
| HORVU.MOREX.r3.3HG0309820.1 | -0.019137 | -0.227125 | -0.7379945 | cluster 1 |
| HORVU.MOREX.r3.3HG0310210.1 | -0.148528 | -0.278551 | -0.5385161 | cluster 1 |
| HORVU.MOREX.r3.3HG0310320.1 | -0.055874 | -0.306726 | -0.6889858 | cluster 1 |
| HORVU.MOREX.r3.3HG0310540.1 | -0.127021 | -0.498619 | -0.5852403 | cluster 1 |
| HORVU.MOREX.r3.3HG0313040.1 | -0.295317 | -0.636241 | -1.4215096 | cluster 1 |
| HORVU.MOREX.r3.3HG0313490.1 | -0.129221 | -0.440114 | -0.9070372 | cluster 1 |
| HORVU.MOREX.r3.3HG0313690.1 | -0.071126 | -0.12746 | -0.2934731 | cluster 1 |
| HORVU.MOREX.r3.3HG0314270.1 | -0.124534 | -0.534095 | -0.7887598 | cluster 1 |
| HORVU.MOREX.r3.3HG0315000.1 | -0.137706 | -0.258206 | -0.4724098 | cluster 1 |
| HORVU.MOREX.r3.3HG0316000.1 | -0.127668 | -0.60661 | -0.7385573 | cluster 1 |
| HORVU.MOREX.r3.3HG0316110.1 | -0.245983 | -0.534869 | -1.0504752 | cluster 1 |
| HORVU.MOREX.r3.3HG0316280.1 | -0.184719 | -0.374697 | -0.8538376 | cluster 1 |
| HORVU.MOREX.r3.3HG0316640.1 | -0.372364 | -0.640982 | -0.7054226 | cluster 1 |
| HORVU.MOREX.r3.3HG0318400.1 | -0.013993 | -0.202029 | -0.9627932 | cluster 1 |
| HORVU.MOREX.r3.3HG0319760.1 | -0.169943 | -0.291739 | -0.4562695 | cluster 1 |
| HORVU.MOREX.r3.3HG0319960.1 | -0.125226 | -0.453155 | -0.9454177 | cluster 1 |
| HORVU.MOREX.r3.3HG0320040.1 | -0.492825 | -0.740241 | -1.4702151 | cluster 1 |
| HORVU.MOREX.r3.3HG0320840.1 | -0.114514 | -0.246791 | -0.4363357 | cluster 1 |
| HORVU.MOREX.r3.3HG0322660.1 | -0.009946 | -0.227472 | -1.3126946 | cluster 1 |
| HORVU.MOREX.r3.3HG0323530.1 | -0.060302 | -0.21161 | -0.7659833 | cluster 1 |
| HORVU.MOREX.r3.3HG0325760.1 | -0.162663 | -0.304926 | -0.495179 | cluster 1 |
| HORVU.MOREX.r3.3HG0327170.1 | -0.141397 | -0.423876 | -0.5524671 | cluster 1 |

**Table S3** Continued.

| ID | logFC_3 h | logFC_6 h | logFC_12 h | cluster No. |
|---|---|---|---|---|
| HORVU.MOREX.r3.3HG0327200.3 | -0.314218 | -0.474313 | -0.5834595 | cluster 1 |
| HORVU.MOREX.r3.3HG0327630.1 | -0.28005 | -0.360357 | -1.1598417 | cluster 1 |
| HORVU.MOREX.r3.3HG0327710.1 | -0.08448 | -0.238175 | -0.3268001 | cluster 1 |
| HORVU.MOREX.r3.3HG0327810.1 | -0.160024 | -0.485271 | -0.6033858 | cluster 1 |
| HORVU.MOREX.r3.3HG0328340.1 | -0.353535 | -0.527215 | -1.6131196 | cluster 1 |
| HORVU.MOREX.r3.3HG0328480.1 | -0.536475 | -1.031843 | -1.7306506 | cluster 1 |
| HORVU.MOREX.r3.3HG0328990.1 | -0.018362 | -0.297842 | -0.4181607 | cluster 1 |
| HORVU.MOREX.r3.3HG0329830.1 | -0.078209 | -0.157845 | -0.2426092 | cluster 1 |
| HORVU.MOREX.r3.3HG0330980.1 | -0.123833 | -0.381879 | -0.4385183 | cluster 1 |
| HORVU.MOREX.r3.3HG0331030.1 | -0.228502 | -0.424614 | -0.4774815 | cluster 1 |
| HORVU.MOREX.r3.4HG0331850.1 | -0.13993 | -0.402414 | -0.6082197 | cluster 1 |
| HORVU.MOREX.r3.4HG0333130.1 | -0.565435 | -0.879041 | -1.1070301 | cluster 1 |
| HORVU.MOREX.r3.4HG0333840.1 | -0.150543 | -0.423726 | -0.8004422 | cluster 1 |
| HORVU.MOREX.r3.4HG0335110.1 | -0.327801 | -0.607782 | -0.9457045 | cluster 1 |
| HORVU.MOREX.r3.4HG0335150.1 | -0.089601 | -0.227157 | -0.3093424 | cluster 1 |
| HORVU.MOREX.r3.4HG0335650.1 | -0.29059 | -0.70595 | -1.3316759 | cluster 1 |
| HORVU.MOREX.r3.4HG0335690.1 | -0.055254 | -0.419542 | -0.6433416 | cluster 1 |
| HORVU.MOREX.r3.4HG0335790.2 | -0.218619 | -0.237669 | -0.5925053 | cluster 1 |
| HORVU.MOREX.r3.4HG0336310.1 | -0.164011 | -0.371162 | -0.6741489 | cluster 1 |
| HORVU.MOREX.r3.4HG0337120.1 | -0.035982 | -0.212683 | -0.6505132 | cluster 1 |
| HORVU.MOREX.r3.4HG0337500.1 | -0.036754 | -0.133957 | -0.1810437 | cluster 1 |
| HORVU.MOREX.r3.4HG0338180.1 | -0.076914 | -0.245034 | -0.4032402 | cluster 1 |
| HORVU.MOREX.r3.4HG0338210.1 | -0.067749 | -0.360667 | -0.7759765 | cluster 1 |
| HORVU.MOREX.r3.4HG0338400.1 | -0.141062 | -0.366588 | -0.8607311 | cluster 1 |
| HORVU.MOREX.r3.4HG0339180.1 | -0.252025 | -1.173939 | -1.9513218 | cluster 1 |
| HORVU.MOREX.r3.4HG0339430.2 | -0.451211 | -0.836488 | -1.8444257 | cluster 1 |
| HORVU.MOREX.r3.4HG0339750.1 | -0.13566 | -0.186062 | -0.6116615 | cluster 1 |
| HORVU.MOREX.r3.4HG0340200.1 | -0.181384 | -0.626989 | -1.0051655 | cluster 1 |
| HORVU.MOREX.r3.4HG0340920.1 | -0.082401 | -0.169444 | -0.4379429 | cluster 1 |
| HORVU.MOREX.r3.4HG0341080.1 | -0.273763 | -0.67648 | -1.8100013 | cluster 1 |
| HORVU.MOREX.r3.4HG0341810.1 | -0.136861 | -0.257877 | -0.3150077 | cluster 1 |
| HORVU.MOREX.r3.4HG0342090.1 | -0.113438 | -0.150665 | -0.4684307 | cluster 1 |
| HORVU.MOREX.r3.4HG0342160.1 | -0.253861 | -0.363561 | -0.4802373 | cluster 1 |
| HORVU.MOREX.r3.4HG0342640.1 | -0.194333 | -0.528217 | -1.808421 | cluster 1 |
| HORVU.MOREX.r3.4HG0342850.1 | -0.085012 | -0.304409 | -0.4053954 | cluster 1 |
| HORVU.MOREX.r3.4HG0342950.1 | -0.293596 | -0.336706 | -0.957211 | cluster 1 |
| HORVU.MOREX.r3.4HG0344990.1 | -0.215404 | -0.766952 | -1.2309492 | cluster 1 |
| HORVU.MOREX.r3.4HG0346720.1 | -0.172152 | -0.411621 | -0.6839953 | cluster 1 |
| HORVU.MOREX.r3.4HG0349170.1 | -0.336471 | -0.500202 | -1.0234216 | cluster 1 |
| HORVU.MOREX.r3.4HG0349530.1 | -0.238848 | -0.396966 | -0.4404921 | cluster 1 |
| HORVU.MOREX.r3.4HG0350800.1 | 0.4696285 | -0.584577 | -2.5380005 | cluster 1 |
| HORVU.MOREX.r3.4HG0353330.1 | -0.284836 | -0.503407 | -0.9357699 | cluster 1 |
| HORVU.MOREX.r3.4HG0353450.1 | -0.439724 | -0.73863 | -2.115183 | cluster 1 |
| HORVU.MOREX.r3.4HG0354480.1 | -0.046077 | -0.113958 | -0.3004255 | cluster 1 |
| HORVU.MOREX.r3.4HG0354540.1 | -0.11519 | -0.349752 | -2.4083044 | cluster 1 |
| HORVU.MOREX.r3.4HG0354970.1 | -0.087296 | -0.277413 | -0.6802499 | cluster 1 |
| HORVU.MOREX.r3.4HG0355150.1 | -0.14746 | -0.313481 | -0.6148953 | cluster 1 |

**Table S3** Continued.

| ID | logFC_3 h | logFC_6 h | logFC_12 h | cluster No. |
|---|---|---|---|---|
| HORVU.MOREX.r3.4HG0357230.1 | 0.0257401 | -0.445226 | -1.2268151 | cluster 1 |
| HORVU.MOREX.r3.4HG0358310.1 | -0.231043 | -0.352879 | -0.7157995 | cluster 1 |
| HORVU.MOREX.r3.4HG0358580.1 | -0.066706 | -0.374257 | -1.0493184 | cluster 1 |
| HORVU.MOREX.r3.4HG0358880.2 | -0.146453 | -0.300945 | -0.522805 | cluster 1 |
| HORVU.MOREX.r3.4HG0358890.1 | -0.173941 | -0.256255 | -0.5796136 | cluster 1 |
| HORVU.MOREX.r3.4HG0361420.2 | -0.069486 | -0.369699 | -0.4902013 | cluster 1 |
| HORVU.MOREX.r3.4HG0361680.1 | -0.42961 | -0.87521 | -1.2830612 | cluster 1 |
| HORVU.MOREX.r3.4HG0362450.1 | -0.460453 | -0.48639 | -0.9434047 | cluster 1 |
| HORVU.MOREX.r3.4HG0364310.1 | -0.108025 | -0.30134 | -0.5137443 | cluster 1 |
| HORVU.MOREX.r3.4HG0364520.1 | -0.104446 | -0.194497 | -0.380177 | cluster 1 |
| HORVU.MOREX.r3.4HG0367380.1 | -0.67793 | -1.321525 | -2.6938687 | cluster 1 |
| HORVU.MOREX.r3.4HG0369880.1 | -0.195898 | -1.070415 | -1.5680736 | cluster 1 |
| HORVU.MOREX.r3.4HG0372280.1 | -0.32752 | -0.724046 | -1.6445618 | cluster 1 |
| HORVU.MOREX.r3.4HG0374640.1 | -0.152825 | -0.420288 | -0.7195119 | cluster 1 |
| HORVU.MOREX.r3.4HG0375550.1 | -0.675131 | -1.346595 | -2.2677136 | cluster 1 |
| HORVU.MOREX.r3.4HG0375690.1 | -0.065895 | -0.37703 | -1.1521943 | cluster 1 |
| HORVU.MOREX.r3.4HG0375760.1 | -0.084617 | -0.229821 | -0.7723314 | cluster 1 |
| HORVU.MOREX.r3.4HG0376950.1 | -0.521269 | -1.510157 | -2.2875001 | cluster 1 |
| HORVU.MOREX.r3.4HG0377020.1 | -0.13919 | -0.294229 | -0.3552233 | cluster 1 |
| HORVU.MOREX.r3.4HG0377070.1 | -0.184358 | -0.464871 | -0.5346444 | cluster 1 |
| HORVU.MOREX.r3.4HG0378580.1 | -0.240317 | -0.539403 | -0.6870235 | cluster 1 |
| HORVU.MOREX.r3.4HG0378710.1 | -0.169922 | -0.482442 | -0.7722851 | cluster 1 |
| HORVU.MOREX.r3.4HG0378960.1 | -0.266631 | -0.545358 | -0.9943749 | cluster 1 |
| HORVU.MOREX.r3.4HG0379960.1 | -0.018742 | -0.151219 | -0.4266713 | cluster 1 |
| HORVU.MOREX.r3.4HG0380710.1 | -0.071625 | -0.25288 | -0.3653337 | cluster 1 |
| HORVU.MOREX.r3.4HG0381310.1 | -0.078991 | -0.712993 | -1.0288265 | cluster 1 |
| HORVU.MOREX.r3.4HG0381680.1 | -0.133526 | -0.28718 | -0.5433962 | cluster 1 |
| HORVU.MOREX.r3.4HG0381880.1 | -0.116201 | -0.415551 | -0.4782793 | cluster 1 |
| HORVU.MOREX.r3.4HG0382500.1 | -0.141508 | -0.324907 | -0.3950199 | cluster 1 |
| HORVU.MOREX.r3.4HG0382760.1 | -0.291802 | -0.472674 | -0.9330541 | cluster 1 |
| HORVU.MOREX.r3.4HG0383070.1 | -0.206758 | -0.486628 | -0.5464747 | cluster 1 |
| HORVU.MOREX.r3.4HG0383870.1 | -0.199645 | -0.46203 | -0.7961873 | cluster 1 |
| HORVU.MOREX.r3.4HG0384110.1 | -0.078175 | -0.309068 | -0.3775722 | cluster 1 |
| HORVU.MOREX.r3.4HG0384700.1 | -0.478101 | -0.549769 | -0.9227407 | cluster 1 |
| HORVU.MOREX.r3.4HG0385080.1 | -0.106665 | -0.162032 | -0.5753367 | cluster 1 |
| HORVU.MOREX.r3.4HG0385100.1 | -0.098855 | -0.25588 | -0.3381732 | cluster 1 |
| HORVU.MOREX.r3.4HG0385270.1 | -0.100593 | -0.26726 | -0.4633914 | cluster 1 |
| HORVU.MOREX.r3.4HG0385300.1 | -0.122212 | -0.290636 | -0.4992224 | cluster 1 |
| HORVU.MOREX.r3.4HG0385780.1 | -0.035605 | -0.278428 | -0.5608728 | cluster 1 |
| HORVU.MOREX.r3.4HG0385880.1 | -0.249969 | -0.595731 | -0.932234 | cluster 1 |
| HORVU.MOREX.r3.4HG0386000.2 | -0.316867 | -0.323233 | -0.5177861 | cluster 1 |
| HORVU.MOREX.r3.4HG0386190.1 | -0.04106 | -0.109 | -0.2804241 | cluster 1 |
| HORVU.MOREX.r3.4HG0386480.1 | -0.162104 | -0.324584 | -0.3988844 | cluster 1 |
| HORVU.MOREX.r3.4HG0386830.1 | 0.0939924 | -0.46654 | -1.0408594 | cluster 1 |
| HORVU.MOREX.r3.4HG0387120.1 | -0.180567 | -0.49297 | -1.0190293 | cluster 1 |
| HORVU.MOREX.r3.4HG0387150.1 | -0.045952 | -0.392939 | -0.875827 | cluster 1 |
| HORVU.MOREX.r3.4HG0388270.1 | -0.205618 | -0.373403 | -0.4161157 | cluster 1 |

**Table S3** Continued.

| ID | logFC_3 h | logFC_6 h | logFC_12 h | cluster No. |
|---|---|---|---|---|
| HORVU.MOREX.r3.4HG0390030.1 | -0.658142 | -0.810853 | -2.1813579 | cluster 1 |
| HORVU.MOREX.r3.4HG0391330.1 | -0.108701 | -0.4174 | -0.534566 | cluster 1 |
| HORVU.MOREX.r3.4HG0392150.1 | 0.0690786 | -0.421996 | -1.5061598 | cluster 1 |
| HORVU.MOREX.r3.4HG0392160.1 | -0.139579 | -0.611611 | -1.2616238 | cluster 1 |
| HORVU.MOREX.r3.4HG0393120.1 | -0.348134 | -0.371416 | -1.0483317 | cluster 1 |
| HORVU.MOREX.r3.4HG0393260.1 | -0.063712 | -0.095408 | -0.3220089 | cluster 1 |
| HORVU.MOREX.r3.4HG0393630.1 | -0.094206 | -0.251055 | -0.2971404 | cluster 1 |
| HORVU.MOREX.r3.4HG0393890.1 | 0.0591768 | -0.468575 | -0.6528003 | cluster 1 |
| HORVU.MOREX.r3.4HG0394690.1 | -0.143459 | -0.167882 | -0.3297057 | cluster 1 |
| HORVU.MOREX.r3.4HG0394810.1 | -0.279647 | -0.453144 | -0.9831815 | cluster 1 |
| HORVU.MOREX.r3.4HG0394980.1 | -0.307281 | -0.403415 | -0.8316173 | cluster 1 |
| HORVU.MOREX.r3.4HG0395120.1 | -0.069213 | -0.123892 | -0.631625 | cluster 1 |
| HORVU.MOREX.r3.4HG0395220.1 | -0.112752 | -0.464357 | -0.6522316 | cluster 1 |
| HORVU.MOREX.r3.4HG0396220.1 | -0.100954 | -0.308114 | -0.3849942 | cluster 1 |
| HORVU.MOREX.r3.4HG0396660.1 | -0.057927 | -0.451731 | -1.6125181 | cluster 1 |
| HORVU.MOREX.r3.4HG0396710.1 | -0.17085 | -0.584794 | -1.1570608 | cluster 1 |
| HORVU.MOREX.r3.4HG0398420.1 | -0.246197 | -0.423128 | -0.5509643 | cluster 1 |
| HORVU.MOREX.r3.4HG0398900.1 | -0.206199 | -0.617502 | -1.0400882 | cluster 1 |
| HORVU.MOREX.r3.4HG0398910.1 | -0.204757 | -0.52852 | -1.0188622 | cluster 1 |
| HORVU.MOREX.r3.4HG0400800.1 | 0.0106904 | -0.229913 | -0.426318 | cluster 1 |
| HORVU.MOREX.r3.4HG0400990.1 | -0.237998 | -0.526986 | -0.7805544 | cluster 1 |
| HORVU.MOREX.r3.4HG0401450.1 | -0.278717 | -0.394576 | -0.4432732 | cluster 1 |
| HORVU.MOREX.r3.4HG0401710.1 | -0.168331 | -0.544497 | -1.2788517 | cluster 1 |
| HORVU.MOREX.r3.4HG0401720.1 | -0.188296 | -0.565064 | -1.1285892 | cluster 1 |
| HORVU.MOREX.r3.4HG0402030.1 | -0.233811 | -0.488545 | -0.5885886 | cluster 1 |
| HORVU.MOREX.r3.4HG0402600.2 | -0.282409 | -0.31583 | -1.2258447 | cluster 1 |
| HORVU.MOREX.r3.4HG0403070.1 | -0.144869 | -0.438474 | -0.6719997 | cluster 1 |
| HORVU.MOREX.r3.4HG0403250.1 | -0.297497 | -0.661895 | -1.080686 | cluster 1 |
| HORVU.MOREX.r3.4HG0404340.1 | -0.06588 | -0.269008 | -0.3654471 | cluster 1 |
| HORVU.MOREX.r3.4HG0404450.1 | -0.035418 | -0.047512 | -0.2822729 | cluster 1 |
| HORVU.MOREX.r3.4HG0404750.1 | -0.207453 | -0.803663 | -2.0069474 | cluster 1 |
| HORVU.MOREX.r3.4HG0406380.1 | -0.024941 | -0.201893 | -0.3852673 | cluster 1 |
| HORVU.MOREX.r3.4HG0406960.1 | -0.161863 | -0.225985 | -0.3257137 | cluster 1 |
| HORVU.MOREX.r3.4HG0407290.1 | 0.0115131 | -0.428804 | -1.4103224 | cluster 1 |
| HORVU.MOREX.r3.4HG0407410.1 | -0.182641 | -0.297975 | -0.9451391 | cluster 1 |
| HORVU.MOREX.r3.4HG0407980.1 | -0.016514 | -0.148909 | -0.7172955 | cluster 1 |
| HORVU.MOREX.r3.4HG0410720.1 | -0.278606 | -0.437323 | -0.8602379 | cluster 1 |
| HORVU.MOREX.r3.4HG0411270.1 | -0.068719 | -0.302788 | -0.7868665 | cluster 1 |
| HORVU.MOREX.r3.4HG0411960.1 | -0.132768 | -0.383825 | -0.9173044 | cluster 1 |
| HORVU.MOREX.r3.4HG0412030.1 | -0.281051 | -0.529297 | -0.6222541 | cluster 1 |
| HORVU.MOREX.r3.4HG0413410.1 | -0.250306 | -0.462422 | -0.9673097 | cluster 1 |
| HORVU.MOREX.r3.4HG0414200.1 | -0.271174 | -0.380106 | -0.4905726 | cluster 1 |
| HORVU.MOREX.r3.4HG0415030.1 | -0.064407 | -0.158585 | -0.3011072 | cluster 1 |
| HORVU.MOREX.r3.4HG0415230.1 | -0.28607 | -0.668266 | -0.905844 | cluster 1 |
| HORVU.MOREX.r3.4HG0415420.1 | -0.160822 | -0.713833 | -0.9823515 | cluster 1 |
| HORVU.MOREX.r3.4HG0415590.1 | -0.783516 | -1.435547 | -1.6041733 | cluster 1 |
| HORVU.MOREX.r3.4HG0415600.1 | -0.017555 | -0.740146 | -1.6847485 | cluster 1 |

**Table S3** Continued.

| ID | logFC_3 h | logFC_6 h | logFC_12 h | cluster No. |
|---|---|---|---|---|
| HORVU.MOREX.r3.4HG0415720.1 | -0.053527 | -0.432013 | -0.7108292 | cluster 1 |
| HORVU.MOREX.r3.4HG0416070.1 | 0.2637421 | -0.505991 | -2.1461068 | cluster 1 |
| HORVU.MOREX.r3.4HG0416220.1 | -0.103926 | -0.224476 | -0.3157687 | cluster 1 |
| HORVU.MOREX.r3.4HG0416730.1 | -0.125933 | -0.512197 | -0.8084348 | cluster 1 |
| HORVU.MOREX.r3.4HG0417410.1 | -0.574144 | -0.849607 | -1.6895623 | cluster 1 |
| HORVU.MOREX.r3.4HG0418530.1 | -0.501548 | -0.635461 | -1.4742094 | cluster 1 |
| HORVU.MOREX.r3.4HG0418640.1 | -0.145755 | -0.152818 | -0.6914223 | cluster 1 |
| HORVU.MOREX.r3.4HG0418690.1 | -0.154026 | -0.403809 | -1.1884371 | cluster 1 |
| HORVU.MOREX.r3.5HG0420150.1 | -0.08752 | -0.262195 | -0.4410016 | cluster 1 |
| HORVU.MOREX.r3.5HG0420210.1 | -0.024909 | -0.213441 | -0.3332005 | cluster 1 |
| HORVU.MOREX.r3.5HG0421460.1 | -0.082273 | -0.165365 | -0.6818786 | cluster 1 |
| HORVU.MOREX.r3.5HG0422490.1 | -0.04789 | -0.07745 | -0.4138775 | cluster 1 |
| HORVU.MOREX.r3.5HG0425680.1 | -0.214913 | -0.338977 | -0.4555909 | cluster 1 |
| HORVU.MOREX.r3.5HG0426060.1 | -0.227911 | -0.457938 | -0.6257845 | cluster 1 |
| HORVU.MOREX.r3.5HG0426480.1 | -0.233434 | -0.287119 | -1.1734032 | cluster 1 |
| HORVU.MOREX.r3.5HG0427060.1 | -0.258452 | -0.332426 | -0.8713509 | cluster 1 |
| HORVU.MOREX.r3.5HG0427370.1 | -0.144446 | -0.428781 | -0.8348413 | cluster 1 |
| HORVU.MOREX.r3.5HG0428840.1 | -0.133444 | -0.36841 | -0.7318554 | cluster 1 |
| HORVU.MOREX.r3.5HG0429230.1 | -0.165596 | -0.349302 | -0.49938 | cluster 1 |
| HORVU.MOREX.r3.5HG0429930.1 | -0.32797 | -0.512412 | -1.0588528 | cluster 1 |
| HORVU.MOREX.r3.5HG0432040.1 | -0.155799 | -0.559888 | -0.6315268 | cluster 1 |
| HORVU.MOREX.r3.5HG0432640.1 | -0.108819 | -0.295572 | -0.3894179 | cluster 1 |
| HORVU.MOREX.r3.5HG0433490.1 | -0.208955 | -0.266373 | -0.5034093 | cluster 1 |
| HORVU.MOREX.r3.5HG0433570.1 | -0.169745 | -0.222535 | -0.580701 | cluster 1 |
| HORVU.MOREX.r3.5HG0435800.1 | -0.454926 | -0.95736 | -2.0966367 | cluster 1 |
| HORVU.MOREX.r3.5HG0437610.1 | -0.002842 | -0.178124 | -0.8276831 | cluster 1 |
| HORVU.MOREX.r3.5HG0438160.1 | -0.17195 | -0.256853 | -0.5415301 | cluster 1 |
| HORVU.MOREX.r3.5HG0438750.1 | 0.1583185 | -0.493062 | -0.7283115 | cluster 1 |
| HORVU.MOREX.r3.5HG0444060.1 | -0.302357 | -0.885659 | -1.0841803 | cluster 1 |
| HORVU.MOREX.r3.5HG0444860.1 | -0.163494 | -0.706663 | -1.3650608 | cluster 1 |
| HORVU.MOREX.r3.5HG0445020.1 | -0.123422 | -0.358476 | -0.5302803 | cluster 1 |
| HORVU.MOREX.r3.5HG0446210.1 | -0.199827 | -0.464107 | -1.069582 | cluster 1 |
| HORVU.MOREX.r3.5HG0446330.1 | -0.138583 | -0.513316 | -0.7147087 | cluster 1 |
| HORVU.MOREX.r3.5HG0446350.1 | -0.080374 | -0.25548 | -0.3615626 | cluster 1 |
| HORVU.MOREX.r3.5HG0447720.1 | -0.162886 | -0.362749 | -0.6370921 | cluster 1 |
| HORVU.MOREX.r3.5HG0450100.1 | -0.104976 | -0.296985 | -0.5261148 | cluster 1 |
| HORVU.MOREX.r3.5HG0450470.1 | -0.054589 | -0.220525 | -0.5590352 | cluster 1 |
| HORVU.MOREX.r3.5HG0454860.1 | -0.187892 | -0.338099 | -0.4102621 | cluster 1 |
| HORVU.MOREX.r3.5HG0456510.1 | -0.069159 | -0.343121 | -0.4460837 | cluster 1 |
| HORVU.MOREX.r3.5HG0459270.1 | -0.173339 | -0.479544 | -0.8485506 | cluster 1 |
| HORVU.MOREX.r3.5HG0459660.1 | -0.07731 | -0.354216 | -0.7815851 | cluster 1 |
| HORVU.MOREX.r3.5HG0459680.1 | -0.096768 | -0.183233 | -0.3843765 | cluster 1 |
| HORVU.MOREX.r3.5HG0460720.1 | -0.076616 | -0.281629 | -0.5017231 | cluster 1 |
| HORVU.MOREX.r3.5HG0461170.1 | -0.112985 | -0.327771 | -0.6830765 | cluster 1 |
| HORVU.MOREX.r3.5HG0461400.1 | -0.038045 | -0.537371 | -1.3010497 | cluster 1 |
| HORVU.MOREX.r3.5HG0461830.1 | -0.0853 | -0.238625 | -1.4149577 | cluster 1 |
| HORVU.MOREX.r3.5HG0462380.1 | -0.288473 | -0.377294 | -0.6342317 | cluster 1 |

**Table S3** Continued.

| ID | logFC_3 h | logFC_6 h | logFC_12 h | cluster No. |
|---|---|---|---|---|
| HORVU.MOREX.r3.5HG0462780.1 | -0.17759 | -0.349005 | -0.4036322 | cluster 1 |
| HORVU.MOREX.r3.5HG0463350.1 | -0.145433 | -0.734358 | -1.6339683 | cluster 1 |
| HORVU.MOREX.r3.5HG0463760.1 | -0.105654 | -0.49392 | -0.747709 | cluster 1 |
| HORVU.MOREX.r3.5HG0465260.1 | -0.376562 | -0.632548 | -0.7770126 | cluster 1 |
| HORVU.MOREX.r3.5HG0465520.1 | -0.370998 | -0.475167 | -1.8639912 | cluster 1 |
| HORVU.MOREX.r3.5HG0466010.1 | -0.122164 | -0.225025 | -0.6696366 | cluster 1 |
| HORVU.MOREX.r3.5HG0466190.1 | -0.191196 | -0.255447 | -0.7094792 | cluster 1 |
| HORVU.MOREX.r3.5HG0466890.1 | -0.021693 | -0.466145 | -0.5776881 | cluster 1 |
| HORVU.MOREX.r3.5HG0467370.1 | -0.203793 | -0.481914 | -0.7575021 | cluster 1 |
| HORVU.MOREX.r3.5HG0467880.1 | -0.003148 | -0.172388 | -0.4598904 | cluster 1 |
| HORVU.MOREX.r3.5HG0467950.1 | -0.31389 | -0.419203 | -0.4777401 | cluster 1 |
| HORVU.MOREX.r3.5HG0468460.1 | -0.127443 | -0.361307 | -0.6000874 | cluster 1 |
| HORVU.MOREX.r3.5HG0468940.1 | -0.129469 | -0.144926 | -0.3201336 | cluster 1 |
| HORVU.MOREX.r3.5HG0470030.1 | -0.158427 | -0.404541 | -0.4953998 | cluster 1 |
| HORVU.MOREX.r3.5HG0470170.1 | -0.19311 | -0.526318 | -0.592036 | cluster 1 |
| HORVU.MOREX.r3.5HG0470800.1 | 0.7275915 | -0.374609 | -2.1821604 | cluster 1 |
| HORVU.MOREX.r3.5HG0471790.1 | -0.110304 | -0.331141 | -0.4559165 | cluster 1 |
| HORVU.MOREX.r3.5HG0471960.1 | -0.004208 | -0.429914 | -1.2011233 | cluster 1 |
| HORVU.MOREX.r3.5HG0472880.1 | -0.172909 | -0.281933 | -0.5088918 | cluster 1 |
| HORVU.MOREX.r3.5HG0473920.1 | -0.375461 | -0.461909 | -0.4836124 | cluster 1 |
| HORVU.MOREX.r3.5HG0475250.1 | -0.085014 | -0.261249 | -0.6964655 | cluster 1 |
| HORVU.MOREX.r3.5HG0476000.1 | -0.110004 | -0.266129 | -0.3452014 | cluster 1 |
| HORVU.MOREX.r3.5HG0476330.1 | -0.048948 | -0.331058 | -0.3746148 | cluster 1 |
| HORVU.MOREX.r3.5HG0476380.1 | -0.137002 | -0.347556 | -0.6614743 | cluster 1 |
| HORVU.MOREX.r3.5HG0476460.1 | 0.1319611 | -0.182961 | -0.7597641 | cluster 1 |
| HORVU.MOREX.r3.5HG0477040.1 | -0.158882 | -0.408917 | -0.6574259 | cluster 1 |
| HORVU.MOREX.r3.5HG0477180.1 | -0.175929 | -0.389786 | -0.5789351 | cluster 1 |
| HORVU.MOREX.r3.5HG0477360.1 | -0.039631 | -0.499111 | -0.6815793 | cluster 1 |
| HORVU.MOREX.r3.5HG0478390.1 | -0.118401 | -0.269869 | -0.4779025 | cluster 1 |
| HORVU.MOREX.r3.5HG0478450.1 | -0.192386 | -0.388657 | -0.417434 | cluster 1 |
| HORVU.MOREX.r3.5HG0479210.1 | -0.188358 | -0.52051 | -1.0162223 | cluster 1 |
| HORVU.MOREX.r3.5HG0479970.1 | -0.120583 | -0.254801 | -0.3904151 | cluster 1 |
| HORVU.MOREX.r3.5HG0480540.1 | -0.041509 | -0.232904 | -0.3684535 | cluster 1 |
| HORVU.MOREX.r3.5HG0481440.1 | -0.181333 | -0.21774 | -0.4766231 | cluster 1 |
| HORVU.MOREX.r3.5HG0482360.1 | 0.0745564 | -0.46641 | -0.7843659 | cluster 1 |
| HORVU.MOREX.r3.5HG0482590.1 | -0.132298 | -0.559607 | -0.9263454 | cluster 1 |
| HORVU.MOREX.r3.5HG0485220.1 | -0.199026 | -0.507254 | -0.6075654 | cluster 1 |
| HORVU.MOREX.r3.5HG0485800.1 | -0.18989 | -0.417902 | -0.5032049 | cluster 1 |
| HORVU.MOREX.r3.5HG0485860.1 | -0.257651 | -0.48557 | -0.6617041 | cluster 1 |
| HORVU.MOREX.r3.5HG0486070.1 | -0.144918 | -0.271329 | -0.6475072 | cluster 1 |
| HORVU.MOREX.r3.5HG0486330.1 | -0.048573 | -0.135892 | -0.3159013 | cluster 1 |
| HORVU.MOREX.r3.5HG0487040.1 | 0.0566432 | -0.267227 | -0.4943747 | cluster 1 |
| HORVU.MOREX.r3.5HG0487640.1 | -0.093607 | -0.641279 | -1.0407351 | cluster 1 |
| HORVU.MOREX.r3.5HG0487660.1 | -0.217624 | -0.633378 | -1.0582883 | cluster 1 |
| HORVU.MOREX.r3.5HG0489130.1 | -0.110733 | -0.543887 | -1.382232 | cluster 1 |
| HORVU.MOREX.r3.5HG0490720.1 | -1.79272 | -2.350369 | -3.9447452 | cluster 1 |
| HORVU.MOREX.r3.5HG0490810.1 | -0.05943 | -0.197519 | -0.2634756 | cluster 1 |

**Table S3** Continued.

| ID | logFC_3 h | logFC_6 h | logFC_12 h | cluster No. |
|---|---|---|---|---|
| HORVU.MOREX.r3.5HG0490830.1 | -0.085344 | -0.146106 | -0.4166418 | cluster 1 |
| HORVU.MOREX.r3.5HG0491380.1 | -0.110673 | -0.478207 | -0.5388866 | cluster 1 |
| HORVU.MOREX.r3.5HG0491750.1 | -0.092746 | -0.230883 | -0.3124519 | cluster 1 |
| HORVU.MOREX.r3.5HG0492650.1 | -0.022488 | -0.27686 | -1.0364724 | cluster 1 |
| HORVU.MOREX.r3.5HG0493070.1 | -0.122015 | -0.35629 | -0.4342602 | cluster 1 |
| HORVU.MOREX.r3.5HG0493850.1 | -0.088564 | -0.256549 | -0.5279151 | cluster 1 |
| HORVU.MOREX.r3.5HG0494320.1 | -0.056222 | -0.100717 | -0.4393089 | cluster 1 |
| HORVU.MOREX.r3.5HG0494470.1 | 0.0229095 | -0.116089 | -0.521104 | cluster 1 |
| HORVU.MOREX.r3.5HG0495580.1 | -0.079477 | -0.164793 | -0.3147977 | cluster 1 |
| HORVU.MOREX.r3.5HG0495790.1 | -0.160887 | -0.419527 | -0.6347201 | cluster 1 |
| HORVU.MOREX.r3.5HG0495840.1 | 0.0035152 | -0.287856 | -0.3910271 | cluster 1 |
| HORVU.MOREX.r3.5HG0496220.1 | -0.156064 | -0.454739 | -1.1960839 | cluster 1 |
| HORVU.MOREX.r3.5HG0497360.1 | 0.0601987 | -0.140292 | -0.5607761 | cluster 1 |
| HORVU.MOREX.r3.5HG0497940.1 | -0.485998 | -0.791284 | -1.2613961 | cluster 1 |
| HORVU.MOREX.r3.5HG0498150.1 | -0.159561 | -0.646318 | -1.2856173 | cluster 1 |
| HORVU.MOREX.r3.5HG0498770.1 | -0.109113 | -0.340288 | -0.4100756 | cluster 1 |
| HORVU.MOREX.r3.5HG0499000.1 | 2.3647132 | 2.1218982 | 1.50728133 | cluster 1 |
| HORVU.MOREX.r3.5HG0499490.1 | -0.108544 | -0.144045 | -0.2847346 | cluster 1 |
| HORVU.MOREX.r3.5HG0499620.1 | -0.101615 | -0.383001 | -0.5351115 | cluster 1 |
| HORVU.MOREX.r3.5HG0500220.1 | -0.158601 | -0.282739 | -0.3162118 | cluster 1 |
| HORVU.MOREX.r3.5HG0500560.2 | -0.22635 | -0.882928 | -1.0044721 | cluster 1 |
| HORVU.MOREX.r3.5HG0500680.1 | -0.150547 | -0.390053 | -0.6810883 | cluster 1 |
| HORVU.MOREX.r3.5HG0500730.1 | -0.085445 | -0.376971 | -0.6680202 | cluster 1 |
| HORVU.MOREX.r3.5HG0501120.1 | -0.129259 | -0.26871 | -0.3221837 | cluster 1 |
| HORVU.MOREX.r3.5HG0501200.1 | -0.225379 | -0.490892 | -0.5402406 | cluster 1 |
| HORVU.MOREX.r3.5HG0501270.1 | -0.137922 | -0.52833 | -1.4076696 | cluster 1 |
| HORVU.MOREX.r3.5HG0501980.1 | -0.183212 | -0.353262 | -0.4485465 | cluster 1 |
| HORVU.MOREX.r3.5HG0502130.1 | -0.147955 | -0.391233 | -0.7761111 | cluster 1 |
| HORVU.MOREX.r3.5HG0502310.1 | -0.196656 | -0.635931 | -0.710535 | cluster 1 |
| HORVU.MOREX.r3.5HG0502750.1 | -0.096127 | -0.372251 | -0.9094411 | cluster 1 |
| HORVU.MOREX.r3.5HG0502760.1 | -0.073732 | -0.303695 | -0.7464793 | cluster 1 |
| HORVU.MOREX.r3.5HG0503950.1 | -0.133301 | -0.365726 | -0.4823158 | cluster 1 |
| HORVU.MOREX.r3.5HG0504160.1 | -0.105746 | -0.314844 | -1.0721874 | cluster 1 |
| HORVU.MOREX.r3.5HG0504250.1 | -0.160806 | -0.363591 | -0.7365398 | cluster 1 |
| HORVU.MOREX.r3.5HG0504670.1 | -0.456707 | -0.632888 | -1.3021566 | cluster 1 |
| HORVU.MOREX.r3.5HG0504710.1 | -0.114832 | -0.211946 | -0.3962978 | cluster 1 |
| HORVU.MOREX.r3.5HG0504800.1 | -0.035177 | -0.484558 | -0.8134552 | cluster 1 |
| HORVU.MOREX.r3.5HG0505100.1 | -0.138924 | -0.330778 | -0.3960939 | cluster 1 |
| HORVU.MOREX.r3.5HG0508190.1 | -0.186162 | -0.390828 | -0.7119832 | cluster 1 |
| HORVU.MOREX.r3.5HG0510060.1 | -0.075469 | -0.256016 | -0.6564189 | cluster 1 |
| HORVU.MOREX.r3.5HG0510590.4 | -0.243129 | -0.756586 | -1.47342 | cluster 1 |
| HORVU.MOREX.r3.5HG0510920.1 | -0.127388 | -0.308239 | -0.389345 | cluster 1 |
| HORVU.MOREX.r3.5HG0510940.1 | -0.213918 | -0.568567 | -1.4615132 | cluster 1 |
| HORVU.MOREX.r3.5HG0511860.1 | -0.148252 | -0.435656 | -0.9293757 | cluster 1 |
| HORVU.MOREX.r3.5HG0511930.1 | 0.2269265 | -0.67185 | -0.8945959 | cluster 1 |
| HORVU.MOREX.r3.5HG0512320.1 | -0.305306 | -0.546114 | -0.7498745 | cluster 1 |
| HORVU.MOREX.r3.5HG0512510.1 | -0.065184 | -0.157201 | -0.6098173 | cluster 1 |

**Table S3** Continued.

| ID | logFC_3 h | logFC_6 h | logFC_12 h | cluster No. |
|---|---|---|---|---|
| HORVU.MOREX.r3.5HG0513020.1 | -0.093005 | -0.267105 | -0.3339665 | cluster 1 |
| HORVU.MOREX.r3.5HG0513120.1 | -0.046063 | -0.087371 | -0.3134437 | cluster 1 |
| HORVU.MOREX.r3.5HG0513440.1 | -0.058705 | -0.314961 | -0.4091539 | cluster 1 |
| HORVU.MOREX.r3.5HG0514100.1 | -0.060005 | -0.287394 | -0.6292722 | cluster 1 |
| HORVU.MOREX.r3.5HG0514110.1 | -0.103543 | -0.3816 | -0.6477007 | cluster 1 |
| HORVU.MOREX.r3.5HG0514490.1 | -0.135669 | -0.310424 | -0.505075 | cluster 1 |
| HORVU.MOREX.r3.5HG0514790.1 | -0.093666 | -0.242443 | -0.4029237 | cluster 1 |
| HORVU.MOREX.r3.5HG0514790.3 | -0.117309 | -0.296945 | -0.4752577 | cluster 1 |
| HORVU.MOREX.r3.5HG0514950.1 | -0.004511 | -0.196927 | -0.5784034 | cluster 1 |
| HORVU.MOREX.r3.5HG0515370.1 | -0.147268 | -0.364991 | -0.502435 | cluster 1 |
| HORVU.MOREX.r3.5HG0516310.1 | -0.144465 | -0.345284 | -0.978312 | cluster 1 |
| HORVU.MOREX.r3.5HG0516470.1 | -0.389694 | -0.863386 | -1.4075846 | cluster 1 |
| HORVU.MOREX.r3.5HG0516490.2 | -0.067545 | -0.173028 | -0.5224994 | cluster 1 |
| HORVU.MOREX.r3.5HG0516720.1 | -0.099159 | -0.577595 | -1.8266352 | cluster 1 |
| HORVU.MOREX.r3.5HG0517640.1 | -0.334622 | -0.684101 | -0.9409454 | cluster 1 |
| HORVU.MOREX.r3.5HG0518490.1 | -0.270225 | -0.558026 | -0.8727459 | cluster 1 |
| HORVU.MOREX.r3.5HG0519240.1 | -0.250013 | -0.536071 | -0.580351 | cluster 1 |
| HORVU.MOREX.r3.5HG0519810.1 | -0.082753 | -0.391947 | -0.5518064 | cluster 1 |
| HORVU.MOREX.r3.5HG0520260.1 | -0.783208 | -0.855478 | -2.4668028 | cluster 1 |
| HORVU.MOREX.r3.5HG0520670.1 | -0.796169 | -1.034934 | -1.3849627 | cluster 1 |
| HORVU.MOREX.r3.5HG0521030.1 | -0.043173 | -0.386634 | -0.6036831 | cluster 1 |
| HORVU.MOREX.r3.5HG0522060.1 | -0.164633 | -0.433521 | -0.803681 | cluster 1 |
| HORVU.MOREX.r3.5HG0523150.1 | 0.0351798 | -0.138438 | -0.2523233 | cluster 1 |
| HORVU.MOREX.r3.5HG0523560.1 | -0.170678 | -0.490673 | -0.8705797 | cluster 1 |
| HORVU.MOREX.r3.5HG0524010.1 | -0.550485 | -0.70662 | -1.3107967 | cluster 1 |
| HORVU.MOREX.r3.5HG0524800.1 | -0.262463 | -1.372057 | -1.6252392 | cluster 1 |
| HORVU.MOREX.r3.5HG0525020.1 | -0.122096 | -0.244444 | -0.8031354 | cluster 1 |
| HORVU.MOREX.r3.5HG0525440.1 | -0.07228 | -0.247331 | -0.3727253 | cluster 1 |
| HORVU.MOREX.r3.5HG0525560.1 | 0.0536787 | -0.219266 | -0.5084243 | cluster 1 |
| HORVU.MOREX.r3.5HG0525980.1 | -0.083957 | -0.287406 | -0.4414487 | cluster 1 |
| HORVU.MOREX.r3.5HG0526250.1 | -0.090948 | -0.205844 | -0.3253673 | cluster 1 |
| HORVU.MOREX.r3.5HG0526940.1 | -0.133589 | -0.278363 | -0.3609518 | cluster 1 |
| HORVU.MOREX.r3.5HG0527650.1 | -0.20443 | -0.486414 | -0.6945739 | cluster 1 |
| HORVU.MOREX.r3.5HG0527670.1 | -0.046059 | -0.61131 | -1.9486555 | cluster 1 |
| HORVU.MOREX.r3.5HG0528340.1 | -0.308398 | -0.659851 | -0.9797436 | cluster 1 |
| HORVU.MOREX.r3.5HG0528520.1 | -0.182602 | -0.378013 | -0.5191138 | cluster 1 |
| HORVU.MOREX.r3.5HG0528890.1 | -0.119078 | -0.275562 | -1.0161001 | cluster 1 |
| HORVU.MOREX.r3.5HG0529120.1 | -0.06845 | -0.364191 | -0.7132555 | cluster 1 |
| HORVU.MOREX.r3.5HG0529130.1 | -0.176822 | -0.293857 | -0.5865178 | cluster 1 |
| HORVU.MOREX.r3.5HG0529500.1 | -0.112009 | -0.318854 | -0.3493907 | cluster 1 |
| HORVU.MOREX.r3.5HG0530750.1 | -0.121243 | -0.341036 | -0.4972007 | cluster 1 |
| HORVU.MOREX.r3.5HG0531850.1 | -0.100767 | -0.366563 | -0.6401909 | cluster 1 |
| HORVU.MOREX.r3.5HG0532150.1 | 2.5233464 | 2.4596495 | 2.12318072 | cluster 1 |
| HORVU.MOREX.r3.5HG0532630.1 | -0.217936 | -0.480784 | -1.2750969 | cluster 1 |
| HORVU.MOREX.r3.5HG0532960.1 | -0.212279 | -0.407444 | -0.5507634 | cluster 1 |
| HORVU.MOREX.r3.5HG0533630.1 | -0.067051 | -0.725516 | -1.444453 | cluster 1 |
| HORVU.MOREX.r3.5HG0533660.1 | -0.082845 | -0.246383 | -0.593649 | cluster 1 |

**Table S3** Continued.

| ID | logFC_3 h | logFC_6 h | logFC_12 h | cluster No. |
|---|---|---|---|---|
| HORVU.MOREX.r3.5HG0534340.1 | -0.104357 | -0.563631 | -1.0525089 | cluster 1 |
| HORVU.MOREX.r3.5HG0534520.1 | -1.050897 | -1.098633 | -2.3077461 | cluster 1 |
| HORVU.MOREX.r3.5HG0534640.1 | -0.089248 | -0.196648 | -0.5192665 | cluster 1 |
| HORVU.MOREX.r3.5HG0536490.1 | -0.341643 | -0.644494 | -1.5120693 | cluster 1 |
| HORVU.MOREX.r3.5HG0536710.1 | -0.108267 | -0.251241 | -0.4415964 | cluster 1 |
| HORVU.MOREX.r3.5HG0536900.1 | -0.388373 | -0.497828 | -1.4737137 | cluster 1 |
| HORVU.MOREX.r3.6HG0539200.1 | -0.159588 | -0.178725 | -0.3625018 | cluster 1 |
| HORVU.MOREX.r3.6HG0539460.1 | -0.067962 | -0.229186 | -0.3163651 | cluster 1 |
| HORVU.MOREX.r3.6HG0539990.1 | 0.0096768 | -0.163307 | -0.3417815 | cluster 1 |
| HORVU.MOREX.r3.6HG0540280.1 | -0.139952 | -0.387102 | -0.8325777 | cluster 1 |
| HORVU.MOREX.r3.6HG0540620.1 | -0.209532 | -0.711663 | -1.5389765 | cluster 1 |
| HORVU.MOREX.r3.6HG0541250.1 | -0.040144 | -0.209566 | -0.2505157 | cluster 1 |
| HORVU.MOREX.r3.6HG0541940.1 | -0.124719 | -0.45334 | -0.5346758 | cluster 1 |
| HORVU.MOREX.r3.6HG0542970.1 | -0.081606 | -0.481892 | -1.0953234 | cluster 1 |
| HORVU.MOREX.r3.6HG0543250.1 | -0.160758 | -0.516459 | -0.7937058 | cluster 1 |
| HORVU.MOREX.r3.6HG0543320.1 | -0.075517 | -0.601413 | -0.6855234 | cluster 1 |
| HORVU.MOREX.r3.6HG0543400.1 | 1.7121169 | 1.3278864 | 1.26388651 | cluster 1 |
| HORVU.MOREX.r3.6HG0543720.1 | 0.012743 | -0.459471 | -0.9085489 | cluster 1 |
| HORVU.MOREX.r3.6HG0543730.1 | -0.183019 | -0.57209 | -0.8529852 | cluster 1 |
| HORVU.MOREX.r3.6HG0543740.1 | -0.206847 | -0.547469 | -0.9817973 | cluster 1 |
| HORVU.MOREX.r3.6HG0543750.1 | -0.178295 | -0.475736 | -0.8004844 | cluster 1 |
| HORVU.MOREX.r3.6HG0543770.1 | -0.191993 | -0.537121 | -0.9086951 | cluster 1 |
| HORVU.MOREX.r3.6HG0543780.1 | -0.223047 | -0.625823 | -0.9423179 | cluster 1 |
| HORVU.MOREX.r3.6HG0543790.1 | -0.136678 | -0.433918 | -1.0669179 | cluster 1 |
| HORVU.MOREX.r3.6HG0543800.1 | -0.253389 | -0.577924 | -0.9281217 | cluster 1 |
| HORVU.MOREX.r3.6HG0545620.1 | -0.18558 | -0.379591 | -0.5578175 | cluster 1 |
| HORVU.MOREX.r3.6HG0545630.1 | -0.042163 | -0.383086 | -1.4940692 | cluster 1 |
| HORVU.MOREX.r3.6HG0545970.1 | -0.120034 | -0.360244 | -0.4801082 | cluster 1 |
| HORVU.MOREX.r3.6HG0546180.1 | -0.141295 | -0.541487 | -0.7092508 | cluster 1 |
| HORVU.MOREX.r3.6HG0546200.1 | -1.75E-05 | -0.291992 | -0.558433 | cluster 1 |
| HORVU.MOREX.r3.6HG0547550.1 | 0.0503365 | -0.264659 | -0.3602582 | cluster 1 |
| HORVU.MOREX.r3.6HG0547970.1 | -0.241155 | -0.71596 | -0.9269406 | cluster 1 |
| HORVU.MOREX.r3.6HG0548460.1 | -0.07358 | -0.491766 | -1.1085409 | cluster 1 |
| HORVU.MOREX.r3.6HG0548670.1 | -0.193966 | -0.658034 | -1.1623691 | cluster 1 |
| HORVU.MOREX.r3.6HG0548820.1 | -0.658338 | -1.080766 | -1.2353357 | cluster 1 |
| HORVU.MOREX.r3.6HG0549510.1 | -0.106393 | -0.212633 | -0.6713129 | cluster 1 |
| HORVU.MOREX.r3.6HG0549830.1 | 0.0703942 | -0.295495 | -0.7422728 | cluster 1 |
| HORVU.MOREX.r3.6HG0550670.1 | -0.060651 | -0.756972 | -1.8102906 | cluster 1 |
| HORVU.MOREX.r3.6HG0550690.1 | -0.050241 | -0.533345 | -1.0214195 | cluster 1 |
| HORVU.MOREX.r3.6HG0550700.1 | -0.496152 | -0.648986 | -2.2968634 | cluster 1 |
| HORVU.MOREX.r3.6HG0550740.1 | 0.037321 | -0.417627 | -0.5001768 | cluster 1 |
| HORVU.MOREX.r3.6HG0550950.1 | -0.095569 | -0.232834 | -0.486758 | cluster 1 |
| HORVU.MOREX.r3.6HG0551740.1 | -0.053592 | -0.394004 | -0.7025778 | cluster 1 |
| HORVU.MOREX.r3.6HG0552030.1 | -0.191493 | -0.32119 | -0.7303511 | cluster 1 |
| HORVU.MOREX.r3.6HG0552720.1 | -0.167026 | -0.358805 | -0.4105796 | cluster 1 |
| HORVU.MOREX.r3.6HG0552800.1 | -0.215235 | -0.622583 | -1.1339797 | cluster 1 |
| HORVU.MOREX.r3.6HG0552860.1 | -0.305328 | -0.591965 | -1.3249857 | cluster 1 |

**Table S3** Continued.

| ID | logFC_3 h | logFC_6 h | logFC_12 h | cluster No. |
|---|---|---|---|---|
| HORVU.MOREX.r3.6HG0553290.1 | -0.083877 | -0.305043 | -0.6079216 | cluster 1 |
| HORVU.MOREX.r3.6HG0554520.1 | -0.199605 | -0.331028 | -0.9874885 | cluster 1 |
| HORVU.MOREX.r3.6HG0554560.1 | 0.1963069 | -0.038654 | -1.0063706 | cluster 1 |
| HORVU.MOREX.r3.6HG0554790.1 | -0.096292 | -0.233471 | -0.3005031 | cluster 1 |
| HORVU.MOREX.r3.6HG0555500.1 | -0.175097 | -0.461786 | -0.618865 | cluster 1 |
| HORVU.MOREX.r3.6HG0555700.1 | -0.196569 | -0.369213 | -0.5635567 | cluster 1 |
| HORVU.MOREX.r3.6HG0557140.1 | -0.045645 | -0.343177 | -0.6447453 | cluster 1 |
| HORVU.MOREX.r3.6HG0557340.1 | -0.153551 | -0.382255 | -0.4724565 | cluster 1 |
| HORVU.MOREX.r3.6HG0558330.1 | 0.028089 | -0.409693 | -1.1625206 | cluster 1 |
| HORVU.MOREX.r3.6HG0558800.1 | -0.152338 | -0.350004 | -0.6439828 | cluster 1 |
| HORVU.MOREX.r3.6HG0558810.1 | -0.229006 | -0.541518 | -0.9661162 | cluster 1 |
| HORVU.MOREX.r3.6HG0558840.1 | -0.085703 | -0.262509 | -0.291443 | cluster 1 |
| HORVU.MOREX.r3.6HG0558880.1 | -0.114661 | -0.373337 | -1.2919911 | cluster 1 |
| HORVU.MOREX.r3.6HG0559000.1 | -0.268953 | -0.393251 | -0.4307431 | cluster 1 |
| HORVU.MOREX.r3.6HG0560290.1 | 0.4472467 | -0.113597 | -2.4548625 | cluster 1 |
| HORVU.MOREX.r3.6HG0564510.1 | 0.0114972 | -0.190698 | -0.9187131 | cluster 1 |
| HORVU.MOREX.r3.6HG0564590.1 | -0.056744 | -0.331409 | -0.4869545 | cluster 1 |
| HORVU.MOREX.r3.6HG0567330.1 | -0.12387 | -0.448646 | -1.2531021 | cluster 1 |
| HORVU.MOREX.r3.6HG0568870.1 | -0.045602 | -0.465213 | -0.7951865 | cluster 1 |
| HORVU.MOREX.r3.6HG0568880.1 | -0.138397 | -0.584181 | -1.2644704 | cluster 1 |
| HORVU.MOREX.r3.6HG0568900.1 | -0.195344 | -0.578485 | -1.1186505 | cluster 1 |
| HORVU.MOREX.r3.6HG0569490.1 | -0.152879 | -0.280017 | -0.4448282 | cluster 1 |
| HORVU.MOREX.r3.6HG0570960.1 | -0.198814 | -0.605972 | -1.1964963 | cluster 1 |
| HORVU.MOREX.r3.6HG0572960.1 | -0.448195 | -1.037814 | -1.5756738 | cluster 1 |
| HORVU.MOREX.r3.6HG0573340.1 | -0.177053 | -0.32049 | -0.3835374 | cluster 1 |
| HORVU.MOREX.r3.6HG0573850.1 | -0.135246 | -0.202562 | -0.3285277 | cluster 1 |
| HORVU.MOREX.r3.6HG0573870.1 | -0.015555 | -0.181699 | -0.6436575 | cluster 1 |
| HORVU.MOREX.r3.6HG0574510.1 | 0.3521514 | -0.237465 | -1.5347521 | cluster 1 |
| HORVU.MOREX.r3.6HG0575410.1 | -0.432236 | -0.868028 | -2.0557757 | cluster 1 |
| HORVU.MOREX.r3.6HG0575690.1 | -0.119582 | -0.625271 | -0.8532731 | cluster 1 |
| HORVU.MOREX.r3.6HG0576550.1 | -0.195785 | -0.545986 | -0.7001287 | cluster 1 |
| HORVU.MOREX.r3.6HG0577170.1 | -0.174211 | -0.932948 | -1.7329456 | cluster 1 |
| HORVU.MOREX.r3.6HG0581010.1 | -0.270151 | -0.447324 | -0.7952286 | cluster 1 |
| HORVU.MOREX.r3.6HG0581020.1 | -0.084033 | -0.329945 | -0.3831694 | cluster 1 |
| HORVU.MOREX.r3.6HG0582230.1 | 0.0804078 | -0.168859 | -0.5132912 | cluster 1 |
| HORVU.MOREX.r3.6HG0583020.1 | -0.110065 | -0.240644 | -0.3798351 | cluster 1 |
| HORVU.MOREX.r3.6HG0584860.1 | 0.0155137 | -0.305834 | -0.4916906 | cluster 1 |
| HORVU.MOREX.r3.6HG0585520.2 | -0.187737 | -0.376382 | -0.5409994 | cluster 1 |
| HORVU.MOREX.r3.6HG0586290.1 | -0.445832 | -0.580732 | -0.8778517 | cluster 1 |
| HORVU.MOREX.r3.6HG0586850.1 | -0.274042 | -0.844347 | -1.267564 | cluster 1 |
| HORVU.MOREX.r3.6HG0587140.1 | -0.201252 | -0.413123 | -0.7067387 | cluster 1 |
| HORVU.MOREX.r3.6HG0587790.1 | -0.652999 | -0.998668 | -1.5951561 | cluster 1 |
| HORVU.MOREX.r3.6HG0588350.1 | -0.273513 | -0.499329 | -0.7133528 | cluster 1 |
| HORVU.MOREX.r3.6HG0590430.1 | -0.172371 | -0.302229 | -0.4119242 | cluster 1 |
| HORVU.MOREX.r3.6HG0593200.1 | -0.216678 | -0.871244 | -1.0180012 | cluster 1 |
| HORVU.MOREX.r3.6HG0594420.1 | 0.0266503 | -0.0987 | -0.3955817 | cluster 1 |
| HORVU.MOREX.r3.6HG0595810.1 | -0.27463 | -0.33438 | -0.9261727 | cluster 1 |

**Table S3** Continued.

| ID | logFC_3 h | logFC_6 h | logFC_12 h | cluster No. |
|---|---|---|---|---|
| HORVU.MOREX.r3.6HG0595860.1 | -0.048797 | -0.30248 | -0.4568244 | cluster 1 |
| HORVU.MOREX.r3.6HG0596460.1 | -0.127101 | -0.208021 | -0.4154092 | cluster 1 |
| HORVU.MOREX.r3.6HG0597320.1 | -0.148617 | -0.287288 | -0.3111842 | cluster 1 |
| HORVU.MOREX.r3.6HG0597360.1 | -0.103421 | -0.228069 | -0.5113958 | cluster 1 |
| HORVU.MOREX.r3.6HG0597590.1 | -0.162113 | -0.407571 | -0.5068844 | cluster 1 |
| HORVU.MOREX.r3.6HG0597620.1 | -0.092541 | -0.381399 | -0.4709729 | cluster 1 |
| HORVU.MOREX.r3.6HG0597770.1 | -0.169845 | -0.460768 | -0.5070072 | cluster 1 |
| HORVU.MOREX.r3.6HG0600500.1 | -0.196866 | -0.518997 | -0.8637638 | cluster 1 |
| HORVU.MOREX.r3.6HG0601560.1 | -0.863825 | -1.01753 | -1.1885003 | cluster 1 |
| HORVU.MOREX.r3.6HG0601690.1 | -0.185185 | -0.326747 | -0.4785737 | cluster 1 |
| HORVU.MOREX.r3.6HG0602290.1 | -0.12907 | -0.217141 | -0.3294031 | cluster 1 |
| HORVU.MOREX.r3.6HG0603410.1 | -0.109436 | -0.462683 | -0.7603923 | cluster 1 |
| HORVU.MOREX.r3.6HG0603530.1 | 0.0002071 | -0.595904 | -1.0911653 | cluster 1 |
| HORVU.MOREX.r3.6HG0603820.1 | -0.074595 | -0.162005 | -0.247231 | cluster 1 |
| HORVU.MOREX.r3.6HG0604200.1 | -0.003365 | -1.00506 | -1.3983131 | cluster 1 |
| HORVU.MOREX.r3.6HG0604620.1 | -0.15296 | -0.557243 | -1.3441761 | cluster 1 |
| HORVU.MOREX.r3.6HG0604810.1 | -0.064523 | -0.298405 | -0.3343238 | cluster 1 |
| HORVU.MOREX.r3.6HG0604860.1 | -0.627473 | -0.819276 | -1.0953738 | cluster 1 |
| HORVU.MOREX.r3.6HG0605290.1 | -0.387361 | -0.473305 | -0.6542963 | cluster 1 |
| HORVU.MOREX.r3.6HG0607000.1 | -0.505787 | -0.9256 | -1.7327798 | cluster 1 |
| HORVU.MOREX.r3.6HG0607360.1 | -0.135846 | -0.555846 | -0.7976135 | cluster 1 |
| HORVU.MOREX.r3.6HG0608780.1 | -0.217584 | -0.416974 | -0.6072037 | cluster 1 |
| HORVU.MOREX.r3.6HG0608800.1 | -0.127509 | -0.309221 | -0.5685331 | cluster 1 |
| HORVU.MOREX.r3.6HG0609100.1 | -0.46752 | -0.593136 | -1.1145585 | cluster 1 |
| HORVU.MOREX.r3.6HG0609520.1 | -0.131174 | -0.468917 | -0.8377816 | cluster 1 |
| HORVU.MOREX.r3.6HG0609560.1 | -0.368463 | -0.639049 | -1.1261645 | cluster 1 |
| HORVU.MOREX.r3.6HG0609720.1 | -0.180461 | -0.704473 | -0.8075552 | cluster 1 |
| HORVU.MOREX.r3.6HG0609750.1 | -0.037484 | -0.161733 | -0.5140055 | cluster 1 |
| HORVU.MOREX.r3.6HG0610700.1 | -0.012566 | -0.607274 | -0.906836 | cluster 1 |
| HORVU.MOREX.r3.6HG0611060.1 | -0.113073 | -0.315331 | -0.4070372 | cluster 1 |
| HORVU.MOREX.r3.6HG0611290.1 | -0.157683 | -0.226004 | -0.8512012 | cluster 1 |
| HORVU.MOREX.r3.6HG0611300.1 | 0.0457688 | -0.263475 | -0.9823054 | cluster 1 |
| HORVU.MOREX.r3.6HG0612070.1 | -0.152228 | -0.434324 | -0.525986 | cluster 1 |
| HORVU.MOREX.r3.6HG0612180.1 | 0.0134935 | -0.479933 | -0.8275732 | cluster 1 |
| HORVU.MOREX.r3.6HG0612250.1 | -0.045218 | -0.131449 | -0.3811627 | cluster 1 |
| HORVU.MOREX.r3.6HG0613270.1 | -0.039291 | -0.209214 | -0.4492613 | cluster 1 |
| HORVU.MOREX.r3.6HG0614350.1 | -0.062384 | -0.239571 | -0.5243657 | cluster 1 |
| HORVU.MOREX.r3.6HG0614460.1 | -0.21881 | -0.333585 | -0.7104182 | cluster 1 |
| HORVU.MOREX.r3.6HG0614500.1 | -0.095265 | -0.200484 | -0.3856272 | cluster 1 |
| HORVU.MOREX.r3.6HG0615710.1 | -0.047347 | -0.167197 | -0.5360668 | cluster 1 |
| HORVU.MOREX.r3.6HG0616480.1 | -0.141398 | -0.255995 | -0.6056354 | cluster 1 |
| HORVU.MOREX.r3.6HG0616680.1 | -0.181669 | -0.311575 | -1.340675 | cluster 1 |
| HORVU.MOREX.r3.6HG0616710.1 | -0.195746 | -0.650215 | -0.9854517 | cluster 1 |
| HORVU.MOREX.r3.6HG0616790.1 | -0.203734 | -0.365348 | -0.5905028 | cluster 1 |
| HORVU.MOREX.r3.6HG0616960.1 | -0.134653 | -0.199891 | -0.6141932 | cluster 1 |
| HORVU.MOREX.r3.6HG0617860.1 | -0.144798 | -0.186554 | -0.2982219 | cluster 1 |
| HORVU.MOREX.r3.6HG0618100.1 | -0.050866 | -0.45235 | -1.8370623 | cluster 1 |

**Table S3** Continued.

| ID | logFC_3 h | logFC_6 h | logFC_12 h | cluster No. |
|---|---|---|---|---|
| HORVU.MOREX.r3.6HG0618110.1 | -0.053755 | -0.344908 | -0.7043801 | cluster 1 |
| HORVU.MOREX.r3.6HG0618170.1 | 0.0039983 | -0.17359 | -0.2558558 | cluster 1 |
| HORVU.MOREX.r3.6HG0618630.1 | -0.045632 | -0.366512 | -0.6031653 | cluster 1 |
| HORVU.MOREX.r3.6HG0619260.1 | -0.062207 | -0.297993 | -0.4589042 | cluster 1 |
| HORVU.MOREX.r3.6HG0619540.1 | 0.0365632 | -0.146762 | -0.351223 | cluster 1 |
| HORVU.MOREX.r3.6HG0619760.1 | -0.191497 | -0.43564 | -0.4785616 | cluster 1 |
| HORVU.MOREX.r3.6HG0620040.1 | -0.569807 | -1.12627 | -1.2902095 | cluster 1 |
| HORVU.MOREX.r3.6HG0620240.1 | -0.25637 | -0.402508 | -0.5072228 | cluster 1 |
| HORVU.MOREX.r3.6HG0620650.1 | -0.215603 | -0.26121 | -0.5769564 | cluster 1 |
| HORVU.MOREX.r3.6HG0621690.1 | -0.084347 | -0.435478 | -0.7427506 | cluster 1 |
| HORVU.MOREX.r3.6HG0622030.1 | -0.185696 | -0.487685 | -0.5847006 | cluster 1 |
| HORVU.MOREX.r3.6HG0622300.1 | -0.104044 | -0.450725 | -0.7942567 | cluster 1 |
| HORVU.MOREX.r3.6HG0623140.1 | 2.8931926 | 2.3679238 | 1.92254547 | cluster 1 |
| HORVU.MOREX.r3.6HG0623830.1 | 0.1543554 | -0.035479 | -1.8860509 | cluster 1 |
| HORVU.MOREX.r3.6HG0623920.1 | -0.030793 | -0.091294 | -0.2024879 | cluster 1 |
| HORVU.MOREX.r3.6HG0624190.1 | -0.128871 | -0.295866 | -0.3309713 | cluster 1 |
| HORVU.MOREX.r3.6HG0624580.1 | -0.017266 | -0.297154 | -1.1248018 | cluster 1 |
| HORVU.MOREX.r3.6HG0624630.1 | -0.092227 | -0.307584 | -0.4434398 | cluster 1 |
| HORVU.MOREX.r3.6HG0624650.1 | -0.082536 | -0.205596 | -0.3545164 | cluster 1 |
| HORVU.MOREX.r3.6HG0627570.1 | -0.136437 | -0.539164 | -0.7442745 | cluster 1 |
| HORVU.MOREX.r3.6HG0627920.1 | -0.118676 | -0.285747 | -0.3309031 | cluster 1 |
| HORVU.MOREX.r3.6HG0628930.1 | -0.110921 | -0.209581 | -0.4994471 | cluster 1 |
| HORVU.MOREX.r3.6HG0629220.1 | -0.2681 | -0.542515 | -0.7523083 | cluster 1 |
| HORVU.MOREX.r3.6HG0631050.1 | -0.020991 | -0.183303 | -1.1214454 | cluster 1 |
| HORVU.MOREX.r3.6HG0631580.1 | -0.16739 | -0.361011 | -0.5723441 | cluster 1 |
| HORVU.MOREX.r3.6HG0631760.1 | -0.170698 | -0.324464 | -0.5334179 | cluster 1 |
| HORVU.MOREX.r3.6HG0632210.1 | 0.0810498 | -0.101504 | -0.5457316 | cluster 1 |
| HORVU.MOREX.r3.6HG0632780.1 | -0.302065 | -0.38065 | -0.6571739 | cluster 1 |
| HORVU.MOREX.r3.6HG0632980.1 | -0.173918 | -0.578965 | -1.562704 | cluster 1 |
| HORVU.MOREX.r3.6HG0633160.1 | -0.422387 | -0.456114 | -0.7772551 | cluster 1 |
| HORVU.MOREX.r3.6HG0633420.1 | -0.202335 | -0.302528 | -0.7963007 | cluster 1 |
| HORVU.MOREX.r3.6HG0633640.1 | -0.102613 | -0.174541 | -0.2723529 | cluster 1 |
| HORVU.MOREX.r3.6HG0634070.1 | -0.110411 | -0.348775 | -0.4293681 | cluster 1 |
| HORVU.MOREX.r3.6HG0634260.1 | -0.006188 | -0.39369 | -0.7518639 | cluster 1 |
| HORVU.MOREX.r3.7HG0634660.1 | -0.22261 | -0.431385 | -0.5765137 | cluster 1 |
| HORVU.MOREX.r3.7HG0634710.1 | -0.119864 | -0.203271 | -0.2281727 | cluster 1 |
| HORVU.MOREX.r3.7HG0635320.1 | -0.160727 | -0.393076 | -0.5982559 | cluster 1 |
| HORVU.MOREX.r3.7HG0635550.1 | -0.229231 | -0.302084 | -1.2248388 | cluster 1 |
| HORVU.MOREX.r3.7HG0635700.1 | -0.087885 | -0.219946 | -0.3338246 | cluster 1 |
| HORVU.MOREX.r3.7HG0636510.1 | -0.260127 | -0.406094 | -0.5417717 | cluster 1 |
| HORVU.MOREX.r3.7HG0636530.1 | -0.094616 | -0.222455 | -0.3929379 | cluster 1 |
| HORVU.MOREX.r3.7HG0636660.1 | -0.047626 | -0.185569 | -0.4099518 | cluster 1 |
| HORVU.MOREX.r3.7HG0636750.1 | -0.0737 | -0.400214 | -0.8620779 | cluster 1 |
| HORVU.MOREX.r3.7HG0637470.1 | -0.113137 | -0.363452 | -0.4714156 | cluster 1 |
| HORVU.MOREX.r3.7HG0637760.1 | -0.221907 | -0.42383 | -0.632365 | cluster 1 |
| HORVU.MOREX.r3.7HG0638340.1 | 1.3564459 | 1.291694 | 1.23369918 | cluster 1 |
| HORVU.MOREX.r3.7HG0638850.1 | -0.140797 | -0.424577 | -0.5714483 | cluster 1 |

**Table S3** Continued.

| ID | logFC_3 h | logFC_6 h | logFC_12 h | cluster No. |
|---|---|---|---|---|
| HORVU.MOREX.r3.7HG0640570.1 | -0.039458 | -0.201169 | -0.4215153 | cluster 1 |
| HORVU.MOREX.r3.7HG0641160.1 | -0.168173 | -0.213976 | -0.3799753 | cluster 1 |
| HORVU.MOREX.r3.7HG0642550.1 | -0.086905 | -0.24661 | -0.3276198 | cluster 1 |
| HORVU.MOREX.r3.7HG0642770.1 | -0.045991 | -0.270793 | -0.3538325 | cluster 1 |
| HORVU.MOREX.r3.7HG0643230.1 | -0.166227 | -0.517072 | -0.7595003 | cluster 1 |
| HORVU.MOREX.r3.7HG0645350.1 | -0.000586 | -0.142832 | -0.3024828 | cluster 1 |
| HORVU.MOREX.r3.7HG0645480.1 | -0.233966 | -0.319076 | -0.5033557 | cluster 1 |
| HORVU.MOREX.r3.7HG0647520.1 | -0.113709 | -0.401838 | -0.7704195 | cluster 1 |
| HORVU.MOREX.r3.7HG0647550.1 | -0.143077 | -0.462468 | -0.5087458 | cluster 1 |
| HORVU.MOREX.r3.7HG0647930.1 | -0.307559 | -1.095103 | -1.3982835 | cluster 1 |
| HORVU.MOREX.r3.7HG0648520.1 | -0.509666 | -0.534049 | -1.8678721 | cluster 1 |
| HORVU.MOREX.r3.7HG0648600.1 | -0.14247 | -0.313815 | -0.5819736 | cluster 1 |
| HORVU.MOREX.r3.7HG0650210.1 | -0.066146 | -0.34155 | -0.4127646 | cluster 1 |
| HORVU.MOREX.r3.7HG0650250.1 | -0.194615 | -0.513916 | -0.6299226 | cluster 1 |
| HORVU.MOREX.r3.7HG0650970.1 | -0.180302 | -0.588829 | -1.176132 | cluster 1 |
| HORVU.MOREX.r3.7HG0650980.1 | -0.231316 | -0.58227 | -0.9585001 | cluster 1 |
| HORVU.MOREX.r3.7HG0651010.1 | -0.31858 | -0.855394 | -0.9329356 | cluster 1 |
| HORVU.MOREX.r3.7HG0651030.1 | -0.095301 | -0.188984 | -0.2701294 | cluster 1 |
| HORVU.MOREX.r3.7HG0651110.1 | -0.230318 | -0.35594 | -0.4334242 | cluster 1 |
| HORVU.MOREX.r3.7HG0652830.1 | -0.500769 | -1.088361 | -1.9147495 | cluster 1 |
| HORVU.MOREX.r3.7HG0653850.1 | -0.206184 | -0.42687 | -0.6948013 | cluster 1 |
| HORVU.MOREX.r3.7HG0654120.1 | -0.561266 | -0.88787 | -1.1985793 | cluster 1 |
| HORVU.MOREX.r3.7HG0654410.1 | -0.167674 | -0.462908 | -0.8840223 | cluster 1 |
| HORVU.MOREX.r3.7HG0654460.1 | -0.135353 | -0.501507 | -1.0717078 | cluster 1 |
| HORVU.MOREX.r3.7HG0654480.1 | -0.230905 | -0.820741 | -1.0693943 | cluster 1 |
| HORVU.MOREX.r3.7HG0654510.1 | -0.327359 | -0.697149 | -1.3495929 | cluster 1 |
| HORVU.MOREX.r3.7HG0654580.1 | -0.197862 | -0.512079 | -1.5678007 | cluster 1 |
| HORVU.MOREX.r3.7HG0654680.1 | -0.053544 | -0.240875 | -0.9333271 | cluster 1 |
| HORVU.MOREX.r3.7HG0655180.1 | -0.133992 | -0.287163 | -0.5348613 | cluster 1 |
| HORVU.MOREX.r3.7HG0655210.1 | 1.8347523 | 1.7303599 | 1.26129321 | cluster 1 |
| HORVU.MOREX.r3.7HG0656250.1 | -0.073877 | -0.220912 | -0.3968207 | cluster 1 |
| HORVU.MOREX.r3.7HG0656340.1 | -0.274754 | -0.984231 | -1.9722095 | cluster 1 |
| HORVU.MOREX.r3.7HG0656430.1 | -0.076029 | -0.215066 | -0.371474 | cluster 1 |
| HORVU.MOREX.r3.7HG0656750.1 | -0.117691 | -0.419695 | -0.7872498 | cluster 1 |
| HORVU.MOREX.r3.7HG0657370.1 | -0.126133 | -0.227293 | -0.3276496 | cluster 1 |
| HORVU.MOREX.r3.7HG0657400.1 | -0.174464 | -0.286906 | -0.6719245 | cluster 1 |
| HORVU.MOREX.r3.7HG0659420.1 | -0.110464 | -0.608047 | -2.5333763 | cluster 1 |
| HORVU.MOREX.r3.7HG0660310.1 | -0.193217 | -0.239191 | -0.6410892 | cluster 1 |
| HORVU.MOREX.r3.7HG0660660.1 | -0.135902 | -0.223746 | -0.480509 | cluster 1 |
| HORVU.MOREX.r3.7HG0660760.1 | -0.199956 | -0.632249 | -1.1535636 | cluster 1 |
| HORVU.MOREX.r3.7HG0661060.1 | -0.065558 | -0.29189 | -0.4474257 | cluster 1 |
| HORVU.MOREX.r3.7HG0661480.1 | -0.125365 | -0.295651 | -0.3558575 | cluster 1 |
| HORVU.MOREX.r3.7HG0661890.1 | -0.138678 | -0.411495 | -0.5739824 | cluster 1 |
| HORVU.MOREX.r3.7HG0662160.1 | -0.236196 | -0.244414 | -0.586524 | cluster 1 |
| HORVU.MOREX.r3.7HG0662210.1 | -0.091926 | -0.254713 | -0.5790927 | cluster 1 |
| HORVU.MOREX.r3.7HG0662220.1 | -0.108194 | -0.687347 | -1.0076766 | cluster 1 |
| HORVU.MOREX.r3.7HG0662650.1 | -0.182877 | -0.712817 | -1.0096494 | cluster 1 |

**Table S3** Continued.

| ID | logFC_3 h | logFC_6 h | logFC_12 h | cluster No. |
|---|---|---|---|---|
| HORVU.MOREX.r3.7HG0662830.1 | -0.106215 | -0.123595 | -0.9237938 | cluster 1 |
| HORVU.MOREX.r3.7HG0663150.1 | 0.0678366 | -0.156122 | -0.3919222 | cluster 1 |
| HORVU.MOREX.r3.7HG0663180.1 | -0.181334 | -0.695322 | -1.0066708 | cluster 1 |
| HORVU.MOREX.r3.7HG0663400.1 | -0.024533 | -0.114788 | -0.2854097 | cluster 1 |
| HORVU.MOREX.r3.7HG0663730.1 | 0.0127606 | -0.482258 | -1.9576747 | cluster 1 |
| HORVU.MOREX.r3.7HG0664130.1 | -0.128656 | -0.209484 | -0.3605416 | cluster 1 |
| HORVU.MOREX.r3.7HG0665400.1 | -0.202368 | -0.303001 | -0.7806399 | cluster 1 |
| HORVU.MOREX.r3.7HG0665520.1 | -0.053235 | -0.331659 | -0.5399756 | cluster 1 |
| HORVU.MOREX.r3.7HG0666340.1 | -0.083134 | -0.440476 | -0.6611363 | cluster 1 |
| HORVU.MOREX.r3.7HG0667280.1 | -0.174941 | -0.431613 | -0.5394402 | cluster 1 |
| HORVU.MOREX.r3.7HG0668850.1 | -0.123102 | -0.377072 | -0.6130462 | cluster 1 |
| HORVU.MOREX.r3.7HG0669240.1 | -0.171567 | -0.296455 | -0.3220881 | cluster 1 |
| HORVU.MOREX.r3.7HG0670020.1 | -0.03904 | -0.525027 | -1.4988357 | cluster 1 |
| HORVU.MOREX.r3.7HG0671880.1 | -0.11702 | -0.415995 | -0.7475091 | cluster 1 |
| HORVU.MOREX.r3.7HG0673610.1 | -0.131863 | -0.251661 | -0.5587134 | cluster 1 |
| HORVU.MOREX.r3.7HG0674440.1 | -0.044694 | -0.453809 | -0.5554254 | cluster 1 |
| HORVU.MOREX.r3.7HG0674480.1 | -0.086969 | -0.217718 | -0.2995958 | cluster 1 |
| HORVU.MOREX.r3.7HG0674750.1 | -0.158349 | -0.26186 | -0.4465006 | cluster 1 |
| HORVU.MOREX.r3.7HG0674790.1 | -0.024804 | -0.179051 | -0.7409316 | cluster 1 |
| HORVU.MOREX.r3.7HG0676000.1 | -0.135898 | -0.74651 | -1.2417459 | cluster 1 |
| HORVU.MOREX.r3.7HG0676230.1 | -0.081152 | -0.494662 | -0.8694304 | cluster 1 |
| HORVU.MOREX.r3.7HG0676350.1 | -0.090925 | -0.324776 | -0.6021678 | cluster 1 |
| HORVU.MOREX.r3.7HG0676570.1 | -0.105052 | -0.670906 | -0.754483 | cluster 1 |
| HORVU.MOREX.r3.7HG0676890.1 | -0.298882 | -0.57958 | -1.4801034 | cluster 1 |
| HORVU.MOREX.r3.7HG0677230.1 | -0.19451 | -0.846785 | -1.4829404 | cluster 1 |
| HORVU.MOREX.r3.7HG0677510.1 | -0.137647 | -0.432649 | -0.59865 | cluster 1 |
| HORVU.MOREX.r3.7HG0677580.1 | -0.076853 | -0.581738 | -1.499187 | cluster 1 |
| HORVU.MOREX.r3.7HG0677860.1 | -0.655353 | -0.931231 | -2.4003369 | cluster 1 |
| HORVU.MOREX.r3.7HG0677940.1 | -0.156659 | -0.289668 | -0.3575406 | cluster 1 |
| HORVU.MOREX.r3.7HG0679350.1 | -0.124085 | -0.508575 | -0.8090444 | cluster 1 |
| HORVU.MOREX.r3.7HG0679730.1 | -0.655764 | -0.845431 | -1.4007413 | cluster 1 |
| HORVU.MOREX.r3.7HG0679760.1 | -0.112621 | -0.472933 | -0.5682749 | cluster 1 |
| HORVU.MOREX.r3.7HG0680170.1 | -0.19124 | -0.35253 | -0.4792956 | cluster 1 |
| HORVU.MOREX.r3.7HG0680680.1 | -0.184219 | -0.276535 | -0.3036024 | cluster 1 |
| HORVU.MOREX.r3.7HG0682120.1 | -0.13942 | -0.243464 | -0.3825648 | cluster 1 |
| HORVU.MOREX.r3.7HG0684040.1 | -0.106939 | -0.252789 | -0.5227941 | cluster 1 |
| HORVU.MOREX.r3.7HG0685550.1 | -0.100185 | -0.279594 | -0.7309508 | cluster 1 |
| HORVU.MOREX.r3.7HG0686000.1 | -0.202797 | -0.478567 | -0.6570539 | cluster 1 |
| HORVU.MOREX.r3.7HG0686830.1 | -0.121564 | -0.492091 | -0.8245815 | cluster 1 |
| HORVU.MOREX.r3.7HG0687050.1 | -0.119937 | -0.335545 | -0.4636151 | cluster 1 |
| HORVU.MOREX.r3.7HG0687080.1 | -0.229433 | -0.401607 | -0.5547604 | cluster 1 |
| HORVU.MOREX.r3.7HG0687190.1 | -0.170013 | -0.18717 | -0.5190393 | cluster 1 |
| HORVU.MOREX.r3.7HG0687630.1 | -0.169298 | -0.537407 | -0.7622627 | cluster 1 |
| HORVU.MOREX.r3.7HG0688300.1 | -0.014668 | -0.244412 | -0.4267074 | cluster 1 |
| HORVU.MOREX.r3.7HG0689490.1 | -0.146435 | -0.253472 | -0.5509563 | cluster 1 |
| HORVU.MOREX.r3.7HG0695500.1 | -0.108798 | -0.358754 | -0.446606 | cluster 1 |
| HORVU.MOREX.r3.7HG0696190.1 | -0.065495 | -0.141331 | -0.4356585 | cluster 1 |

**Table S3** Continued.

| ID | logFC_3 h | logFC_6 h | logFC_12 h | cluster No. |
|---|---|---|---|---|
| HORVU.MOREX.r3.7HG0696220.1 | -0.121042 | -0.153208 | -0.9496701 | cluster 1 |
| HORVU.MOREX.r3.7HG0698910.1 | -0.069497 | -0.30609 | -0.3927772 | cluster 1 |
| HORVU.MOREX.r3.7HG0700930.1 | -0.054687 | -0.342451 | -1.4382572 | cluster 1 |
| HORVU.MOREX.r3.7HG0703580.1 | -0.154695 | -0.340258 | -0.6269324 | cluster 1 |
| HORVU.MOREX.r3.7HG0703780.1 | -0.163881 | -0.275805 | -0.5475225 | cluster 1 |
| HORVU.MOREX.r3.7HG0703780.2 | -0.093231 | -0.187604 | -0.2484471 | cluster 1 |
| HORVU.MOREX.r3.7HG0705490.1 | -0.338225 | -0.532682 | -1.528505 | cluster 1 |
| HORVU.MOREX.r3.7HG0705700.1 | -0.178573 | -0.368019 | -0.6988959 | cluster 1 |
| HORVU.MOREX.r3.7HG0706140.1 | -0.160804 | -0.357811 | -0.5611145 | cluster 1 |
| HORVU.MOREX.r3.7HG0706330.1 | -0.169117 | -0.280805 | -0.5281658 | cluster 1 |
| HORVU.MOREX.r3.7HG0706860.1 | -0.288051 | -0.359972 | -0.6439385 | cluster 1 |
| HORVU.MOREX.r3.7HG0706930.1 | 0.009728 | -0.355013 | -0.4978365 | cluster 1 |
| HORVU.MOREX.r3.7HG0708120.1 | -0.008908 | -0.315109 | -1.271149 | cluster 1 |
| HORVU.MOREX.r3.7HG0708320.1 | -0.285227 | -0.526197 | -0.6768737 | cluster 1 |
| HORVU.MOREX.r3.7HG0708500.1 | -0.049101 | -0.349748 | -0.9004663 | cluster 1 |
| HORVU.MOREX.r3.7HG0708820.1 | -0.056712 | -0.207484 | -0.6855952 | cluster 1 |
| HORVU.MOREX.r3.7HG0709110.1 | -0.202043 | -0.618362 | -1.5696279 | cluster 1 |
| HORVU.MOREX.r3.7HG0709230.1 | -0.616893 | -0.85014 | -2.4948944 | cluster 1 |
| HORVU.MOREX.r3.7HG0709860.1 | -0.100549 | -0.279522 | -0.6928871 | cluster 1 |
| HORVU.MOREX.r3.7HG0710980.1 | 0.0007696 | -0.329803 | -1.1495816 | cluster 1 |
| HORVU.MOREX.r3.7HG0711850.1 | -0.181235 | -0.337918 | -0.388833 | cluster 1 |
| HORVU.MOREX.r3.7HG0712650.1 | -0.270573 | -0.518248 | -0.8557637 | cluster 1 |
| HORVU.MOREX.r3.7HG0712850.1 | -0.078997 | -0.333915 | -0.512629 | cluster 1 |
| HORVU.MOREX.r3.7HG0714120.1 | -0.102133 | -0.268439 | -0.3313837 | cluster 1 |
| HORVU.MOREX.r3.7HG0714510.1 | -0.317924 | -0.442413 | -1.1559138 | cluster 1 |
| HORVU.MOREX.r3.7HG0714870.1 | -0.082615 | -0.506534 | -0.6194152 | cluster 1 |
| HORVU.MOREX.r3.7HG0714910.1 | -0.115071 | -0.375343 | -0.7035154 | cluster 1 |
| HORVU.MOREX.r3.7HG0714930.1 | -0.151498 | -0.284934 | -0.3979646 | cluster 1 |
| HORVU.MOREX.r3.7HG0714950.1 | -0.1894 | -0.42833 | -0.5935839 | cluster 1 |
| HORVU.MOREX.r3.7HG0715570.1 | -0.197607 | -0.487681 | -0.6433136 | cluster 1 |
| HORVU.MOREX.r3.7HG0716170.1 | -0.116099 | -0.78014 | -1.0136248 | cluster 1 |
| HORVU.MOREX.r3.7HG0716900.1 | -0.096514 | -0.271943 | -0.3549208 | cluster 1 |
| HORVU.MOREX.r3.7HG0718710.1 | -0.090745 | -0.235178 | -0.2781975 | cluster 1 |
| HORVU.MOREX.r3.7HG0719070.1 | -0.130119 | -0.137319 | -0.6855384 | cluster 1 |
| HORVU.MOREX.r3.7HG0719410.1 | -0.173565 | -0.244727 | -0.8934651 | cluster 1 |
| HORVU.MOREX.r3.7HG0719720.1 | -0.123505 | -0.398525 | -0.7186812 | cluster 1 |
| HORVU.MOREX.r3.7HG0720130.1 | -0.161254 | -0.407649 | -0.591589 | cluster 1 |
| HORVU.MOREX.r3.7HG0720670.1 | -0.057808 | -0.359034 | -0.8829704 | cluster 1 |
| HORVU.MOREX.r3.7HG0720790.1 | -0.586399 | -0.876625 | -1.1746271 | cluster 1 |
| HORVU.MOREX.r3.7HG0721760.1 | -0.121892 | -0.283438 | -0.3845036 | cluster 1 |
| HORVU.MOREX.r3.7HG0721790.1 | -0.303067 | -0.688737 | -0.7865252 | cluster 1 |
| HORVU.MOREX.r3.7HG0721800.1 | -0.077775 | -0.287488 | -0.3480656 | cluster 1 |
| HORVU.MOREX.r3.7HG0721930.1 | -0.247642 | -0.313167 | -0.5556501 | cluster 1 |
| HORVU.MOREX.r3.7HG0723420.1 | -0.124151 | -0.196288 | -0.3773619 | cluster 1 |
| HORVU.MOREX.r3.7HG0725060.1 | -0.48511 | -0.629375 | -1.5202651 | cluster 1 |
| HORVU.MOREX.r3.7HG0726210.1 | -0.277083 | -0.480339 | -0.7209696 | cluster 1 |
| HORVU.MOREX.r3.7HG0726350.1 | 0.0035137 | -0.34395 | -0.5869455 | cluster 1 |

**Table S3** Continued.

| ID | logFC_3 h | logFC_6 h | logFC_12 h | cluster No. |
|---|---|---|---|---|
| HORVU.MOREX.r3.7HG0726700.1 | -0.097543 | -0.370089 | -0.8612115 | cluster 1 |
| HORVU.MOREX.r3.7HG0726790.1 | -0.149551 | -0.301441 | -0.4855129 | cluster 1 |
| HORVU.MOREX.r3.7HG0727070.1 | -0.136339 | -0.411342 | -0.6475236 | cluster 1 |
| HORVU.MOREX.r3.7HG0727350.1 | -0.140516 | -0.369975 | -0.4026862 | cluster 1 |
| HORVU.MOREX.r3.7HG0727460.1 | -0.149635 | -0.279721 | -0.8341254 | cluster 1 |
| HORVU.MOREX.r3.7HG0728190.1 | -0.137765 | -0.334828 | -0.4237947 | cluster 1 |
| HORVU.MOREX.r3.7HG0728220.1 | -0.209479 | -0.323363 | -0.5219552 | cluster 1 |
| HORVU.MOREX.r3.7HG0728630.1 | 0.0848225 | -0.653313 | -1.0050638 | cluster 1 |
| HORVU.MOREX.r3.7HG0728910.1 | -0.105664 | -0.152004 | -0.8345446 | cluster 1 |
| HORVU.MOREX.r3.7HG0730070.1 | -0.113278 | -0.625955 | -1.3355518 | cluster 1 |
| HORVU.MOREX.r3.7HG0730310.1 | -0.142713 | -0.508358 | -0.8960971 | cluster 1 |
| HORVU.MOREX.r3.7HG0730350.1 | -0.14344 | -0.346178 | -0.5382438 | cluster 1 |
| HORVU.MOREX.r3.7HG0730800.1 | -0.26359 | -0.63261 | -1.1261643 | cluster 1 |
| HORVU.MOREX.r3.7HG0730820.1 | -0.189394 | -0.617362 | -0.9696442 | cluster 1 |
| HORVU.MOREX.r3.7HG0730840.1 | -0.343589 | -0.636674 | -1.0896918 | cluster 1 |
| HORVU.MOREX.r3.7HG0731490.1 | -0.3258 | -0.452923 | -0.5993975 | cluster 1 |
| HORVU.MOREX.r3.7HG0732270.1 | 0.0796105 | -0.151539 | -0.5649961 | cluster 1 |
| HORVU.MOREX.r3.7HG0732290.1 | -0.03798 | -0.862297 | -1.7091001 | cluster 1 |
| HORVU.MOREX.r3.7HG0732330.1 | -0.870155 | -1.042089 | -1.5236155 | cluster 1 |
| HORVU.MOREX.r3.7HG0732610.1 | -0.19077 | -0.627832 | -1.2427384 | cluster 1 |
| HORVU.MOREX.r3.7HG0733910.1 | -0.112638 | -0.28049 | -0.3858919 | cluster 1 |
| HORVU.MOREX.r3.7HG0734670.1 | -0.115313 | -0.440215 | -0.560796 | cluster 1 |
| HORVU.MOREX.r3.7HG0735840.1 | -0.564004 | -1.04726 | -1.2687473 | cluster 1 |
| HORVU.MOREX.r3.7HG0736960.1 | -0.209552 | -0.806234 | -1.1528079 | cluster 1 |
| HORVU.MOREX.r3.7HG0737440.1 | -0.186253 | -0.813779 | -1.1761342 | cluster 1 |
| HORVU.MOREX.r3.7HG0737870.1 | -0.013682 | -0.605608 | -1.71847 | cluster 1 |
| HORVU.MOREX.r3.7HG0739380.1 | -0.140426 | -0.47347 | -0.9423897 | cluster 1 |
| HORVU.MOREX.r3.7HG0739400.1 | -0.324056 | -0.716058 | -1.2105552 | cluster 1 |
| HORVU.MOREX.r3.7HG0739460.1 | -0.1767 | -0.512547 | -1.0485936 | cluster 1 |
| HORVU.MOREX.r3.7HG0739480.1 | -0.201196 | -0.48366 | -0.8839477 | cluster 1 |
| HORVU.MOREX.r3.7HG0739490.1 | -0.208689 | -0.461091 | -0.9538976 | cluster 1 |
| HORVU.MOREX.r3.7HG0739640.1 | -0.223411 | -0.576685 | -0.9622662 | cluster 1 |
| HORVU.MOREX.r3.7HG0739910.1 | -0.054828 | -0.479194 | -0.678719 | cluster 1 |
| HORVU.MOREX.r3.7HG0741080.1 | -0.672432 | -0.796848 | -0.8366293 | cluster 1 |
| HORVU.MOREX.r3.7HG0741640.1 | -0.325964 | -0.756658 | -0.9646278 | cluster 1 |
| HORVU.MOREX.r3.7HG0741650.1 | -0.125074 | -0.569975 | -0.8381484 | cluster 1 |
| HORVU.MOREX.r3.7HG0742070.1 | -0.178064 | -0.394719 | -0.6542131 | cluster 1 |
| HORVU.MOREX.r3.7HG0742450.1 | -0.104021 | -0.329626 | -0.641118 | cluster 1 |
| HORVU.MOREX.r3.7HG0742470.1 | -0.36422 | -0.439455 | -0.8769606 | cluster 1 |
| HORVU.MOREX.r3.7HG0744250.1 | -0.243621 | -0.430026 | -0.7346591 | cluster 1 |
| HORVU.MOREX.r3.7HG0744260.1 | -0.607973 | -1.32412 | -2.9809703 | cluster 1 |
| HORVU.MOREX.r3.7HG0744540.1 | 0.1405138 | -0.192473 | -2.0658572 | cluster 1 |
| HORVU.MOREX.r3.7HG0744790.1 | 0.0128261 | -0.265496 | -0.581483 | cluster 1 |
| HORVU.MOREX.r3.7HG0744970.1 | -0.070899 | -0.256537 | -0.2950067 | cluster 1 |
| HORVU.MOREX.r3.7HG0745980.1 | -0.081069 | -0.342585 | -0.4031379 | cluster 1 |
| HORVU.MOREX.r3.7HG0746230.1 | -0.070312 | -0.374381 | -0.6037365 | cluster 1 |
| HORVU.MOREX.r3.7HG0746770.1 | -0.227242 | -0.498766 | -0.6970324 | cluster 1 |

**Table S3** Continued.

| ID | logFC_3 h | logFC_6 h | logFC_12 h | cluster No. |
|---|---|---|---|---|
| HORVU.MOREX.r3.7HG0749150.1 | 0.1075723 | -0.097489 | -2.0555127 | cluster 1 |
| HORVU.MOREX.r3.7HG0749170.1 | -0.343989 | -0.626683 | -1.3516979 | cluster 1 |
| HORVU.MOREX.r3.7HG0749660.1 | -0.033838 | -0.180697 | -0.6566934 | cluster 1 |
| HORVU.MOREX.r3.7HG0749940.3 | -0.173912 | -0.586304 | -0.7117558 | cluster 1 |
| HORVU.MOREX.r3.7HG0751000.1 | -0.037894 | -0.469851 | -1.2526513 | cluster 1 |
| HORVU.MOREX.r3.7HG0751110.1 | -0.190387 | -0.510756 | -0.6138129 | cluster 1 |
| HORVU.MOREX.r3.7HG0751290.1 | -0.660763 | -0.908229 | -1.0176798 | cluster 1 |
| HORVU.MOREX.r3.1HG0002570.1 | 0.816034 | 1.3298146 | 1.61969582 | cluster 2 |
| HORVU.MOREX.r3.1HG0002960.1 | 0.8480958 | 1.1891287 | 1.30211439 | cluster 2 |
| HORVU.MOREX.r3.1HG0003160.1 | 0.2815356 | 0.461246 | 0.59740027 | cluster 2 |
| HORVU.MOREX.r3.1HG0003430.1 | 0.1910299 | 0.4586132 | 0.50114337 | cluster 2 |
| HORVU.MOREX.r3.1HG0004560.1 | -0.022642 | 0.8777935 | 0.84834751 | cluster 2 |
| HORVU.MOREX.r3.1HG0006400.1 | 0.1146551 | 0.7031986 | 0.82396251 | cluster 2 |
| HORVU.MOREX.r3.1HG0006660.1 | 0.4186282 | 0.9539727 | 0.94286013 | cluster 2 |
| HORVU.MOREX.r3.1HG0007670.1 | -0.169503 | 0.3231918 | 0.58383366 | cluster 2 |
| HORVU.MOREX.r3.1HG0010920.1 | 0.401162 | 0.5468505 | 0.56432129 | cluster 2 |
| HORVU.MOREX.r3.1HG0011820.1 | 0.4297878 | 0.9730911 | 0.92402335 | cluster 2 |
| HORVU.MOREX.r3.1HG0013820.1 | 0.1688292 | 0.3662798 | 0.52267715 | cluster 2 |
| HORVU.MOREX.r3.1HG0014680.1 | 0.8092587 | 1.3972174 | 1.37838415 | cluster 2 |
| HORVU.MOREX.r3.1HG0015750.1 | 0.6461444 | 1.7678989 | 1.65720455 | cluster 2 |
| HORVU.MOREX.r3.1HG0015960.1 | 0.2752366 | 1.2472685 | 1.20640107 | cluster 2 |
| HORVU.MOREX.r3.1HG0018110.1 | 0.1850191 | 0.2824626 | 0.37471105 | cluster 2 |
| HORVU.MOREX.r3.1HG0018590.1 | 0.377141 | 0.6107542 | 0.84173356 | cluster 2 |
| HORVU.MOREX.r3.1HG0021880.1 | 0.7647028 | 2.4449034 | 2.30473582 | cluster 2 |
| HORVU.MOREX.r3.1HG0023200.1 | 0.3561223 | 1.268951 | 1.27112406 | cluster 2 |
| HORVU.MOREX.r3.1HG0024900.1 | 0.0310827 | 0.1763802 | 0.34858337 | cluster 2 |
| HORVU.MOREX.r3.1HG0025430.1 | 0.3553935 | 1.2727393 | 1.1845806 | cluster 2 |
| HORVU.MOREX.r3.1HG0026010.1 | 0.3545835 | 0.4935198 | 0.64887578 | cluster 2 |
| HORVU.MOREX.r3.1HG0026080.1 | 0.3773057 | 0.6697716 | 0.64487517 | cluster 2 |
| HORVU.MOREX.r3.1HG0026830.1 | 0.3568673 | 0.5057221 | 0.62992631 | cluster 2 |
| HORVU.MOREX.r3.1HG0029730.1 | 0.2412764 | 0.3823849 | 0.47092527 | cluster 2 |
| HORVU.MOREX.r3.1HG0032930.1 | 0.122655 | 0.670299 | 0.67473973 | cluster 2 |
| HORVU.MOREX.r3.1HG0035400.1 | 0.1905985 | 0.5406722 | 0.66107006 | cluster 2 |
| HORVU.MOREX.r3.1HG0038800.1 | 0.1954959 | 0.6345157 | 0.70049348 | cluster 2 |
| HORVU.MOREX.r3.1HG0040930.1 | 0.8895017 | 1.7882863 | 1.80373232 | cluster 2 |
| HORVU.MOREX.r3.1HG0041280.1 | 0.4552104 | 0.8644407 | 0.98091236 | cluster 2 |
| HORVU.MOREX.r3.1HG0044470.1 | 0.2535506 | 0.7005231 | 0.77087652 | cluster 2 |
| HORVU.MOREX.r3.1HG0046180.1 | 0.2889429 | 0.4092173 | 0.49234211 | cluster 2 |
| HORVU.MOREX.r3.1HG0047100.1 | 0.4002883 | 1.1687198 | 1.15087066 | cluster 2 |
| HORVU.MOREX.r3.1HG0047580.1 | 0.161516 | 0.3993397 | 0.50590673 | cluster 2 |
| HORVU.MOREX.r3.1HG0050440.1 | 1.1427038 | 1.6308828 | 1.80523882 | cluster 2 |
| HORVU.MOREX.r3.1HG0050700.1 | 0.68656 | 1.4703829 | 1.41346892 | cluster 2 |
| HORVU.MOREX.r3.1HG0050920.1 | 0.4405744 | 0.8341891 | 1.04312538 | cluster 2 |
| HORVU.MOREX.r3.1HG0051920.1 | 0.3662728 | 0.9571774 | 0.93966354 | cluster 2 |
| HORVU.MOREX.r3.1HG0053450.1 | 0.4461442 | 0.8577287 | 0.92566908 | cluster 2 |
| HORVU.MOREX.r3.1HG0054010.1 | 0.3164991 | 0.5716175 | 0.63374806 | cluster 2 |
| HORVU.MOREX.r3.1HG0054070.1 | 0.6250529 | 0.9386297 | 1.20247035 | cluster 2 |

**Table S3** Continued.

| ID | logFC_3 h | logFC_6 h | logFC_12 h | cluster No. |
|---|---|---|---|---|
| HORVU.MOREX.r3.1HG0054110.1 | 0.2333108 | 0.3679888 | 0.46117724 | cluster 2 |
| HORVU.MOREX.r3.1HG0054510.1 | 0.7755924 | 1.0213169 | 1.11389941 | cluster 2 |
| HORVU.MOREX.r3.1HG0055240.1 | 2.1640636 | 2.2385391 | 2.28817889 | cluster 2 |
| HORVU.MOREX.r3.1HG0056240.1 | 0.3077347 | 0.6718929 | 0.74188459 | cluster 2 |
| HORVU.MOREX.r3.1HG0056570.1 | 0.2650334 | 0.4121611 | 0.51512523 | cluster 2 |
| HORVU.MOREX.r3.1HG0056580.1 | 0.1856858 | 0.2706326 | 0.34575771 | cluster 2 |
| HORVU.MOREX.r3.1HG0056810.1 | 0.1974665 | 0.3538185 | 0.41777063 | cluster 2 |
| HORVU.MOREX.r3.1HG0056990.1 | -0.126232 | 0.1188655 | 0.45777116 | cluster 2 |
| HORVU.MOREX.r3.1HG0057860.1 | 0.2926234 | 0.6115088 | 0.65790933 | cluster 2 |
| HORVU.MOREX.r3.1HG0061210.1 | 0.1557527 | 0.2821924 | 0.32461919 | cluster 2 |
| HORVU.MOREX.r3.1HG0061390.1 | 0.1600274 | 0.5309879 | 0.7218625 | cluster 2 |
| HORVU.MOREX.r3.1HG0061660.1 | 0.5192854 | 0.8859364 | 0.98617271 | cluster 2 |
| HORVU.MOREX.r3.1HG0061710.1 | 0.1552148 | 0.4241786 | 0.50222289 | cluster 2 |
| HORVU.MOREX.r3.1HG0062110.1 | 0.8732405 | 1.3550929 | 1.35788268 | cluster 2 |
| HORVU.MOREX.r3.1HG0062980.1 | 0.4314587 | 0.7410038 | 1.03359978 | cluster 2 |
| HORVU.MOREX.r3.1HG0063430.1 | 0.1334172 | 0.3139 | 0.39567343 | cluster 2 |
| HORVU.MOREX.r3.1HG0064740.1 | 0.1060772 | 0.5370709 | 0.75737343 | cluster 2 |
| HORVU.MOREX.r3.1HG0065520.1 | 0.0545143 | 0.3096202 | 0.52584262 | cluster 2 |
| HORVU.MOREX.r3.1HG0065860.1 | 0.4890558 | 0.8488967 | 1.05962715 | cluster 2 |
| HORVU.MOREX.r3.1HG0066610.1 | 0.7729362 | 1.1409509 | 1.12183008 | cluster 2 |
| HORVU.MOREX.r3.1HG0067500.1 | 0.0757265 | 0.2535178 | 0.42750343 | cluster 2 |
| HORVU.MOREX.r3.1HG0067630.1 | 0.1278233 | 0.3343533 | 0.39612494 | cluster 2 |
| HORVU.MOREX.r3.1HG0067930.1 | 0.1286308 | 0.3808856 | 0.55963106 | cluster 2 |
| HORVU.MOREX.r3.1HG0067980.1 | 1.1432184 | 1.4805668 | 1.56074787 | cluster 2 |
| HORVU.MOREX.r3.1HG0068010.1 | 1.1317354 | 1.6387976 | 1.61049372 | cluster 2 |
| HORVU.MOREX.r3.1HG0069380.1 | 0.2306783 | 0.824246 | 0.81738825 | cluster 2 |
| HORVU.MOREX.r3.1HG0070110.1 | 0.144219 | 0.2750925 | 0.34724864 | cluster 2 |
| HORVU.MOREX.r3.1HG0070340.1 | 1.0667337 | 1.5971119 | 1.5663168 | cluster 2 |
| HORVU.MOREX.r3.1HG0070430.1 | -0.038802 | 0.2344276 | 0.60028624 | cluster 2 |
| HORVU.MOREX.r3.1HG0070850.1 | 0.4133316 | 0.8047114 | 0.89347048 | cluster 2 |
| HORVU.MOREX.r3.1HG0072160.1 | 0.2770799 | 0.6884414 | 0.70504596 | cluster 2 |
| HORVU.MOREX.r3.1HG0072510.1 | 0.5431199 | 1.0432019 | 1.25919789 | cluster 2 |
| HORVU.MOREX.r3.1HG0072680.1 | 0.328617 | 0.746658 | 0.9050929 | cluster 2 |
| HORVU.MOREX.r3.1HG0073010.1 | 0.4497443 | 0.8382302 | 0.95892818 | cluster 2 |
| HORVU.MOREX.r3.1HG0073160.1 | 0.1885816 | 0.4061384 | 0.42731956 | cluster 2 |
| HORVU.MOREX.r3.1HG0073390.1 | 0.9925304 | 1.6807759 | 1.86620372 | cluster 2 |
| HORVU.MOREX.r3.1HG0073500.1 | 0.489418 | 1.6829513 | 1.88635403 | cluster 2 |
| HORVU.MOREX.r3.1HG0073550.1 | 1.4658151 | 2.0831008 | 2.09899008 | cluster 2 |
| HORVU.MOREX.r3.1HG0074010.1 | 0.4620102 | 0.9071867 | 1.0026054 | cluster 2 |
| HORVU.MOREX.r3.1HG0075320.1 | 0.7816868 | 1.1791158 | 1.34370577 | cluster 2 |
| HORVU.MOREX.r3.1HG0075440.1 | 0.4944808 | 0.716354 | 0.8320652 | cluster 2 |
| HORVU.MOREX.r3.1HG0076120.1 | 0.3149319 | 0.5684553 | 0.68528036 | cluster 2 |
| HORVU.MOREX.r3.1HG0077790.1 | 0.3056013 | 0.893749 | 1.00880469 | cluster 2 |
| HORVU.MOREX.r3.1HG0078040.1 | 0.1764706 | 0.3862847 | 0.45085531 | cluster 2 |
| HORVU.MOREX.r3.1HG0078910.1 | 0.1789599 | 0.3371299 | 0.52394093 | cluster 2 |
| HORVU.MOREX.r3.1HG0078970.1 | 1.0924665 | 1.6063089 | 1.99171783 | cluster 2 |
| HORVU.MOREX.r3.1HG0079020.1 | 1.0870269 | 2.1099641 | 2.03949696 | cluster 2 |

**Table S3** Continued.

| ID | logFC_3 h | logFC_6 h | logFC_12 h | cluster No. |
|---|---|---|---|---|
| HORVU.MOREX.r3.1HG0079750.1 | 0.4190078 | 1.0094094 | 0.9744382 | cluster 2 |
| HORVU.MOREX.r3.1HG0079760.1 | 0.4314093 | 0.6363233 | 0.79672809 | cluster 2 |
| HORVU.MOREX.r3.1HG0079920.1 | 0.3361798 | 0.6173308 | 0.6513715 | cluster 2 |
| HORVU.MOREX.r3.1HG0081110.1 | 0.0059002 | 0.3542383 | 0.65001886 | cluster 2 |
| HORVU.MOREX.r3.1HG0082530.1 | 0.1196339 | 0.2679148 | 0.46830668 | cluster 2 |
| HORVU.MOREX.r3.1HG0083440.1 | -0.030056 | 0.1969053 | 0.34965751 | cluster 2 |
| HORVU.MOREX.r3.1HG0084970.1 | 0.8014998 | 1.2288608 | 1.27796679 | cluster 2 |
| HORVU.MOREX.r3.1HG0085380.1 | 0.1572528 | 0.3175546 | 0.3914105 | cluster 2 |
| HORVU.MOREX.r3.1HG0086090.1 | 0.3035259 | 0.6194064 | 1.04015992 | cluster 2 |
| HORVU.MOREX.r3.1HG0086390.1 | 0.2549845 | 0.7967687 | 0.89146002 | cluster 2 |
| HORVU.MOREX.r3.1HG0087590.1 | 0.8094329 | 1.456679 | 1.42436505 | cluster 2 |
| HORVU.MOREX.r3.1HG0087620.1 | 0.6565543 | 1.0056498 | 1.05313113 | cluster 2 |
| HORVU.MOREX.r3.1HG0090100.1 | 0.7165692 | 1.2540656 | 1.31925538 | cluster 2 |
| HORVU.MOREX.r3.1HG0091240.1 | 0.192182 | 0.4585977 | 0.56887539 | cluster 2 |
| HORVU.MOREX.r3.1HG0092010.1 | 0.4914559 | 0.7236539 | 0.81422792 | cluster 2 |
| HORVU.MOREX.r3.1HG0092730.1 | 0.236626 | 0.6166629 | 0.77346805 | cluster 2 |
| HORVU.MOREX.r3.1HG0095170.1 | 0.8601542 | 1.5836265 | 1.57227006 | cluster 2 |
| HORVU.MOREX.r3.2HG0096290.1 | 0.1697319 | 0.4266028 | 0.60815498 | cluster 2 |
| HORVU.MOREX.r3.2HG0099780.1 | 0.4150032 | 0.724752 | 1.10345775 | cluster 2 |
| HORVU.MOREX.r3.2HG0099900.1 | 0.2870693 | 0.5805348 | 0.57411585 | cluster 2 |
| HORVU.MOREX.r3.2HG0100900.2 | 0.7694444 | 1.466048 | 1.59847627 | cluster 2 |
| HORVU.MOREX.r3.2HG0102670.1 | 1.8547983 | 2.2314588 | 2.3913667 | cluster 2 |
| HORVU.MOREX.r3.2HG0104080.1 | 0.5863758 | 0.8885191 | 1.00608694 | cluster 2 |
| HORVU.MOREX.r3.2HG0105020.1 | 0.3320269 | 0.5026094 | 0.69185748 | cluster 2 |
| HORVU.MOREX.r3.2HG0106260.1 | 1.5296484 | 2.1383183 | 2.5738544 | cluster 2 |
| HORVU.MOREX.r3.2HG0107300.1 | 0.8474797 | 1.4586036 | 1.64984172 | cluster 2 |
| HORVU.MOREX.r3.2HG0109340.1 | 0.4941929 | 1.1236998 | 1.2043941 | cluster 2 |
| HORVU.MOREX.r3.2HG0111620.1 | 0.3389687 | 0.6832929 | 0.75725143 | cluster 2 |
| HORVU.MOREX.r3.2HG0112030.1 | 0.1177364 | 0.3804763 | 0.51672242 | cluster 2 |
| HORVU.MOREX.r3.2HG0112670.1 | 1.124314 | 1.793326 | 1.8373569 | cluster 2 |
| HORVU.MOREX.r3.2HG0113880.1 | 0.2605632 | 0.4587301 | 0.62138027 | cluster 2 |
| HORVU.MOREX.r3.2HG0113990.1 | 1.652593 | 2.2789359 | 2.25069031 | cluster 2 |
| HORVU.MOREX.r3.2HG0116990.1 | 0.2554949 | 0.5555379 | 0.56638687 | cluster 2 |
| HORVU.MOREX.r3.2HG0117880.1 | 1.9599887 | 2.6174308 | 2.59952396 | cluster 2 |
| HORVU.MOREX.r3.2HG0119650.1 | 0.2576946 | 0.457482 | 0.69764091 | cluster 2 |
| HORVU.MOREX.r3.2HG0119800.1 | 1.0987614 | 1.4900502 | 1.53377838 | cluster 2 |
| HORVU.MOREX.r3.2HG0120320.1 | 0.377562 | 0.5789971 | 0.82476645 | cluster 2 |
| HORVU.MOREX.r3.2HG0120650.1 | 0.7099929 | 1.0125719 | 1.15429085 | cluster 2 |
| HORVU.MOREX.r3.2HG0121200.1 | 0.5073511 | 0.743407 | 0.90263914 | cluster 2 |
| HORVU.MOREX.r3.2HG0121480.1 | 0.3044656 | 0.687162 | 0.76526272 | cluster 2 |
| HORVU.MOREX.r3.2HG0122510.1 | 0.3761735 | 0.5250866 | 0.70689603 | cluster 2 |
| HORVU.MOREX.r3.2HG0122520.1 | 0.1152367 | 0.5835001 | 0.75351087 | cluster 2 |
| HORVU.MOREX.r3.2HG0123170.1 | 0.3764052 | 0.656127 | 0.9938262 | cluster 2 |
| HORVU.MOREX.r3.2HG0124660.1 | -0.014887 | 0.1596953 | 0.29612374 | cluster 2 |
| HORVU.MOREX.r3.2HG0127020.1 | 0.1809534 | 0.5127101 | 0.68511203 | cluster 2 |
| HORVU.MOREX.r3.2HG0129450.1 | 0.4115037 | 0.9624815 | 0.95965053 | cluster 2 |
| HORVU.MOREX.r3.2HG0132160.1 | 0.5832001 | 2.3261431 | 2.18912656 | cluster 2 |

**Table S3** Continued.

| ID | logFC_3 h | logFC_6 h | logFC_12 h | cluster No. |
|---|---|---|---|---|
| HORVU.MOREX.r3.2HG0134990.1 | 0.4536518 | 0.6062987 | 0.68588902 | cluster 2 |
| HORVU.MOREX.r3.2HG0135210.1 | 0.3780272 | 0.6727723 | 0.77181105 | cluster 2 |
| HORVU.MOREX.r3.2HG0138540.1 | 0.3085631 | 0.5798993 | 0.6000674 | cluster 2 |
| HORVU.MOREX.r3.2HG0140170.1 | 0.2706125 | 0.7297222 | 0.74070197 | cluster 2 |
| HORVU.MOREX.r3.2HG0140380.1 | 1.5615412 | 1.675108 | 1.76626899 | cluster 2 |
| HORVU.MOREX.r3.2HG0141140.1 | 0.9139418 | 1.060458 | 1.11878653 | cluster 2 |
| HORVU.MOREX.r3.2HG0145360.1 | 0.5039365 | 0.9907402 | 1.04013448 | cluster 2 |
| HORVU.MOREX.r3.2HG0150530.1 | 0.1210945 | 0.4404153 | 0.56604141 | cluster 2 |
| HORVU.MOREX.r3.2HG0155750.1 | 0.9979915 | 1.8172943 | 2.01468789 | cluster 2 |
| HORVU.MOREX.r3.2HG0156610.1 | 1.1955286 | 1.9668569 | 1.99590465 | cluster 2 |
| HORVU.MOREX.r3.2HG0157170.1 | 0.0892596 | 0.2925576 | 0.28597544 | cluster 2 |
| HORVU.MOREX.r3.2HG0157340.1 | 0.2365324 | 0.415833 | 0.49672226 | cluster 2 |
| HORVU.MOREX.r3.2HG0158180.1 | 0.5049772 | 0.9209716 | 0.93866018 | cluster 2 |
| HORVU.MOREX.r3.2HG0164160.2 | 0.8756918 | 1.4548757 | 1.55391465 | cluster 2 |
| HORVU.MOREX.r3.2HG0165180.1 | 0.162007 | 0.3244045 | 0.48891405 | cluster 2 |
| HORVU.MOREX.r3.2HG0165920.1 | 0.7383041 | 1.0390414 | 1.32369249 | cluster 2 |
| HORVU.MOREX.r3.2HG0168070.1 | 0.8585737 | 1.3022418 | 1.54383282 | cluster 2 |
| HORVU.MOREX.r3.2HG0168330.1 | 0.1734171 | 0.4352607 | 0.5097686 | cluster 2 |
| HORVU.MOREX.r3.2HG0173600.1 | 0.2766916 | 0.5268451 | 0.61785162 | cluster 2 |
| HORVU.MOREX.r3.2HG0173960.1 | 0.4870302 | 0.6833282 | 0.66725743 | cluster 2 |
| HORVU.MOREX.r3.2HG0174650.1 | 0.8497415 | 0.9243346 | 0.96707212 | cluster 2 |
| HORVU.MOREX.r3.2HG0174750.1 | 0.299016 | 1.205453 | 1.19501899 | cluster 2 |
| HORVU.MOREX.r3.2HG0175380.1 | 0.9053362 | 1.6285147 | 1.65633196 | cluster 2 |
| HORVU.MOREX.r3.2HG0178050.1 | 0.3553613 | 0.6719366 | 0.76918271 | cluster 2 |
| HORVU.MOREX.r3.2HG0179560.1 | 0.5620998 | 1.2650234 | 1.19788231 | cluster 2 |
| HORVU.MOREX.r3.2HG0180500.1 | 0.1028389 | 0.2320499 | 0.36337228 | cluster 2 |
| HORVU.MOREX.r3.2HG0181510.2 | 0.0948066 | 0.3408424 | 0.45152873 | cluster 2 |
| HORVU.MOREX.r3.2HG0181680.1 | 0.9902678 | 1.8076092 | 2.00934686 | cluster 2 |
| HORVU.MOREX.r3.2HG0182210.2 | 0.3718404 | 0.6267738 | 0.86392819 | cluster 2 |
| HORVU.MOREX.r3.2HG0182400.1 | 0.2782616 | 0.7873183 | 0.9242027 | cluster 2 |
| HORVU.MOREX.r3.2HG0184180.1 | 0.449513 | 0.8846691 | 1.13761939 | cluster 2 |
| HORVU.MOREX.r3.2HG0184230.1 | 0.3538529 | 2.484088 | 2.39357162 | cluster 2 |
| HORVU.MOREX.r3.2HG0185370.1 | 2.0892432 | 2.6125066 | 2.71949862 | cluster 2 |
| HORVU.MOREX.r3.2HG0185570.1 | 0.1081284 | 0.5133423 | 0.73647328 | cluster 2 |
| HORVU.MOREX.r3.2HG0188200.1 | 0.5634929 | 0.6913168 | 0.78333772 | cluster 2 |
| HORVU.MOREX.r3.2HG0189980.1 | 0.0695638 | 0.3528309 | 0.50174275 | cluster 2 |
| HORVU.MOREX.r3.2HG0190290.1 | 0.4486419 | 0.6013912 | 0.62647603 | cluster 2 |
| HORVU.MOREX.r3.2HG0192000.1 | 0.4654997 | 0.7865423 | 0.82910508 | cluster 2 |
| HORVU.MOREX.r3.2HG0192270.1 | 1.6090773 | 2.3650346 | 2.55024845 | cluster 2 |
| HORVU.MOREX.r3.2HG0192400.1 | 0.3662454 | 0.6147984 | 0.7856642 | cluster 2 |
| HORVU.MOREX.r3.2HG0192540.1 | 0.1144642 | 0.2114541 | 0.31826157 | cluster 2 |
| HORVU.MOREX.r3.2HG0195910.1 | 0.3344327 | 0.56002 | 0.70352231 | cluster 2 |
| HORVU.MOREX.r3.2HG0197760.1 | 0.2338253 | 0.386019 | 0.53742174 | cluster 2 |
| HORVU.MOREX.r3.2HG0198870.1 | 0.9372408 | 1.1802771 | 1.17767397 | cluster 2 |
| HORVU.MOREX.r3.2HG0200710.1 | 0.7596757 | 1.5594746 | 1.8335563 | cluster 2 |
| HORVU.MOREX.r3.2HG0202220.1 | 0.1173856 | 0.2586358 | 0.34870383 | cluster 2 |
| HORVU.MOREX.r3.2HG0203330.1 | 0.2694214 | 0.3680154 | 0.43296135 | cluster 2 |

**Table S3** Continued.

| ID | logFC_3 h | logFC_6 h | logFC_12 h | cluster No. |
|---|---|---|---|---|
| HORVU.MOREX.r3.2HG0205050.1 | 0.7536856 | 1.4715989 | 2.02591118 | cluster 2 |
| HORVU.MOREX.r3.2HG0205810.1 | 0.3280324 | 1.2016192 | 1.23013379 | cluster 2 |
| HORVU.MOREX.r3.2HG0206470.1 | 0.2694722 | 0.4030391 | 0.55066039 | cluster 2 |
| HORVU.MOREX.r3.2HG0206650.1 | 0.2620155 | 0.4397694 | 0.55889322 | cluster 2 |
| HORVU.MOREX.r3.2HG0207130.1 | 0.4068899 | 1.4490313 | 1.53422862 | cluster 2 |
| HORVU.MOREX.r3.2HG0207720.1 | 0.5067106 | 0.9111347 | 1.14572552 | cluster 2 |
| HORVU.MOREX.r3.2HG0207860.1 | 1.2944609 | 1.4486193 | 1.58711801 | cluster 2 |
| HORVU.MOREX.r3.2HG0208170.1 | 0.2352241 | 1.3862978 | 1.56262846 | cluster 2 |
| HORVU.MOREX.r3.2HG0209810.1 | 0.0662381 | 0.2299367 | 0.43206324 | cluster 2 |
| HORVU.MOREX.r3.2HG0210610.1 | 0.3736037 | 0.9086098 | 0.92546459 | cluster 2 |
| HORVU.MOREX.r3.2HG0217550.1 | 0.2170308 | 0.2891089 | 0.38892147 | cluster 2 |
| HORVU.MOREX.r3.3HG0218800.1 | 1.3408989 | 2.0365772 | 1.97108408 | cluster 2 |
| HORVU.MOREX.r3.3HG0219410.1 | 0.3314618 | 0.7538848 | 0.81639832 | cluster 2 |
| HORVU.MOREX.r3.3HG0221180.1 | 0.6790219 | 1.1981478 | 1.20578344 | cluster 2 |
| HORVU.MOREX.r3.3HG0221240.1 | 0.4260681 | 0.626593 | 0.78135254 | cluster 2 |
| HORVU.MOREX.r3.3HG0221440.1 | 0.3967657 | 0.8068423 | 0.79981404 | cluster 2 |
| HORVU.MOREX.r3.3HG0223160.1 | 0.260881 | 0.5494062 | 0.61297236 | cluster 2 |
| HORVU.MOREX.r3.3HG0224440.1 | 0.2521341 | 0.8101636 | 0.86777827 | cluster 2 |
| HORVU.MOREX.r3.3HG0225760.1 | 0.3849776 | 1.178426 | 1.18855926 | cluster 2 |
| HORVU.MOREX.r3.3HG0230090.1 | 0.3232532 | 0.5851915 | 0.94060617 | cluster 2 |
| HORVU.MOREX.r3.3HG0230500.1 | 0.8450312 | 1.0647175 | 1.22687651 | cluster 2 |
| HORVU.MOREX.r3.3HG0230870.1 | 0.961177 | 1.6638412 | 1.70204986 | cluster 2 |
| HORVU.MOREX.r3.3HG0231000.1 | 0.3280071 | 0.9797396 | 0.92413278 | cluster 2 |
| HORVU.MOREX.r3.3HG0231240.1 | 0.5431899 | 0.810961 | 1.05285024 | cluster 2 |
| HORVU.MOREX.r3.3HG0233990.1 | 0.8122961 | 1.2915571 | 1.24043971 | cluster 2 |
| HORVU.MOREX.r3.3HG0234240.1 | 0.3978413 | 0.9288043 | 0.97395022 | cluster 2 |
| HORVU.MOREX.r3.3HG0236500.1 | 0.2220856 | 0.6874581 | 0.67005911 | cluster 2 |
| HORVU.MOREX.r3.3HG0236690.1 | 0.4484496 | 0.6348102 | 0.66802296 | cluster 2 |
| HORVU.MOREX.r3.3HG0237990.1 | 0.522957 | 0.8426499 | 0.88595667 | cluster 2 |
| HORVU.MOREX.r3.3HG0242150.1 | 1.5930177 | 2.8160289 | 3.27182942 | cluster 2 |
| HORVU.MOREX.r3.3HG0245010.1 | 0.1656037 | 0.3584781 | 0.47089257 | cluster 2 |
| HORVU.MOREX.r3.3HG0246030.1 | 0.2451723 | 0.6744941 | 1.08833755 | cluster 2 |
| HORVU.MOREX.r3.3HG0246500.1 | 0.3732535 | 0.7684238 | 0.86297927 | cluster 2 |
| HORVU.MOREX.r3.3HG0246670.1 | 1.2505966 | 1.6748638 | 1.75670347 | cluster 2 |
| HORVU.MOREX.r3.3HG0248440.1 | 0.1967806 | 0.4395193 | 0.69610435 | cluster 2 |
| HORVU.MOREX.r3.3HG0249400.1 | 0.5126295 | 0.8759675 | 1.21276614 | cluster 2 |
| HORVU.MOREX.r3.3HG0249640.1 | 0.2657831 | 0.5988288 | 0.62351463 | cluster 2 |
| HORVU.MOREX.r3.3HG0249890.1 | 0.1908097 | 0.6359167 | 0.76939864 | cluster 2 |
| HORVU.MOREX.r3.3HG0252170.1 | 0.8956447 | 1.6729829 | 1.65645577 | cluster 2 |
| HORVU.MOREX.r3.3HG0252210.1 | 0.6918987 | 0.9490763 | 1.17295354 | cluster 2 |
| HORVU.MOREX.r3.3HG0252250.1 | 0.6987571 | 1.3986258 | 1.60335922 | cluster 2 |
| HORVU.MOREX.r3.3HG0252380.1 | 0.1710449 | 0.899178 | 1.02187193 | cluster 2 |
| HORVU.MOREX.r3.3HG0254430.1 | -0.025285 | 0.2238274 | 0.2685602 | cluster 2 |
| HORVU.MOREX.r3.3HG0254930.1 | 1.8775805 | 2.620343 | 2.68949759 | cluster 2 |
| HORVU.MOREX.r3.3HG0254940.1 | 0.4143863 | 0.860821 | 0.86871366 | cluster 2 |
| HORVU.MOREX.r3.3HG0254950.1 | 0.7777211 | 1.4584539 | 1.40223712 | cluster 2 |
| HORVU.MOREX.r3.3HG0255020.1 | 0.4974885 | 0.9494325 | 0.98417562 | cluster 2 |

**Table S3** Continued.

| ID | logFC_3 h | logFC_6 h | logFC_12 h | cluster No. |
|---|---|---|---|---|
| HORVU.MOREX.r3.3HG0255580.1 | 0.1565359 | 0.3025266 | 0.48972851 | cluster 2 |
| HORVU.MOREX.r3.3HG0255700.1 | 0.6050989 | 0.9367771 | 0.94903354 | cluster 2 |
| HORVU.MOREX.r3.3HG0257840.1 | 0.7573933 | 1.3827047 | 1.38972203 | cluster 2 |
| HORVU.MOREX.r3.3HG0264650.1 | 0.737846 | 1.8000893 | 1.69217634 | cluster 2 |
| HORVU.MOREX.r3.3HG0266530.1 | 0.276387 | 0.6074802 | 0.85406364 | cluster 2 |
| HORVU.MOREX.r3.3HG0267000.1 | 0.2663401 | 0.504754 | 0.81989737 | cluster 2 |
| HORVU.MOREX.r3.3HG0271380.1 | 0.2859479 | 0.4741105 | 0.60762866 | cluster 2 |
| HORVU.MOREX.r3.3HG0273790.2 | 0.1458978 | 0.6251988 | 0.668958 | cluster 2 |
| HORVU.MOREX.r3.3HG0275060.1 | 0.6942171 | 1.3793674 | 1.55736662 | cluster 2 |
| HORVU.MOREX.r3.3HG0275100.1 | 0.2138621 | 1.2206472 | 1.28314239 | cluster 2 |
| HORVU.MOREX.r3.3HG0275770.1 | 0.2103829 | 0.5128552 | 0.72826346 | cluster 2 |
| HORVU.MOREX.r3.3HG0276130.1 | 0.5630525 | 1.0544702 | 1.09624594 | cluster 2 |
| HORVU.MOREX.r3.3HG0276160.2 | 0.8450076 | 1.2238195 | 1.55791778 | cluster 2 |
| HORVU.MOREX.r3.3HG0276730.1 | 0.1223194 | 0.2069207 | 0.27304692 | cluster 2 |
| HORVU.MOREX.r3.3HG0278120.1 | 0.2083986 | 0.3600599 | 0.38149806 | cluster 2 |
| HORVU.MOREX.r3.3HG0278170.1 | 0.9548513 | 1.7076948 | 1.70670555 | cluster 2 |
| HORVU.MOREX.r3.3HG0278710.1 | 0.296406 | 0.4826987 | 0.66008226 | cluster 2 |
| HORVU.MOREX.r3.3HG0279750.1 | 0.207975 | 0.3143969 | 0.32767197 | cluster 2 |
| HORVU.MOREX.r3.3HG0279850.1 | -0.059459 | 0.3299392 | 0.47955293 | cluster 2 |
| HORVU.MOREX.r3.3HG0280060.1 | 0.6899142 | 1.8979377 | 2.09462376 | cluster 2 |
| HORVU.MOREX.r3.3HG0280160.1 | 0.0395933 | 0.2655924 | 0.42237899 | cluster 2 |
| HORVU.MOREX.r3.3HG0280960.1 | 0.2730958 | 0.7139438 | 0.68640652 | cluster 2 |
| HORVU.MOREX.r3.3HG0283250.1 | 0.1756096 | 0.439437 | 0.6176891 | cluster 2 |
| HORVU.MOREX.r3.3HG0283740.1 | 0.1571979 | 0.3460681 | 0.54575535 | cluster 2 |
| HORVU.MOREX.r3.3HG0284100.1 | 0.1955155 | 0.3840195 | 0.43775086 | cluster 2 |
| HORVU.MOREX.r3.3HG0284780.1 | 0.0401797 | 0.3559578 | 0.3220962 | cluster 2 |
| HORVU.MOREX.r3.3HG0285780.1 | 0.3742395 | 0.8296677 | 1.03195173 | cluster 2 |
| HORVU.MOREX.r3.3HG0285840.1 | 0.4650272 | 0.7566451 | 0.98992789 | cluster 2 |
| HORVU.MOREX.r3.3HG0287000.1 | 0.4184393 | 0.7786466 | 1.00541978 | cluster 2 |
| HORVU.MOREX.r3.3HG0287690.1 | 0.3638968 | 0.9060343 | 0.93261865 | cluster 2 |
| HORVU.MOREX.r3.3HG0288070.1 | 0.1652256 | 0.5652533 | 0.52258319 | cluster 2 |
| HORVU.MOREX.r3.3HG0288710.1 | 0.9306129 | 1.4825416 | 1.44900912 | cluster 2 |
| HORVU.MOREX.r3.3HG0288710.2 | 1.1291964 | 1.6664555 | 1.7527596 | cluster 2 |
| HORVU.MOREX.r3.3HG0288960.1 | 0.5413086 | 1.0478369 | 1.24567004 | cluster 2 |
| HORVU.MOREX.r3.3HG0291030.1 | 0.3112427 | 0.6142111 | 0.84448301 | cluster 2 |
| HORVU.MOREX.r3.3HG0291040.1 | 0.0753592 | 0.7613512 | 0.78598658 | cluster 2 |
| HORVU.MOREX.r3.3HG0291100.1 | 0.1292693 | 0.3142801 | 0.3241665 | cluster 2 |
| HORVU.MOREX.r3.3HG0291160.1 | 0.1120419 | 0.2082623 | 0.25671613 | cluster 2 |
| HORVU.MOREX.r3.3HG0293130.1 | 0.7673112 | 1.0615266 | 1.08817398 | cluster 2 |
| HORVU.MOREX.r3.3HG0293310.1 | 0.8866974 | 1.9219134 | 1.8702952 | cluster 2 |
| HORVU.MOREX.r3.3HG0293320.1 | 1.0305108 | 1.3440974 | 1.50808586 | cluster 2 |
| HORVU.MOREX.r3.3HG0293490.1 | 0.8345457 | 1.0987903 | 1.08783324 | cluster 2 |
| HORVU.MOREX.r3.3HG0293760.1 | 0.6830217 | 1.3727751 | 1.45386285 | cluster 2 |
| HORVU.MOREX.r3.3HG0294040.1 | 0.404668 | 0.6441345 | 0.77944156 | cluster 2 |
| HORVU.MOREX.r3.3HG0294110.1 | 0.4317541 | 0.6554953 | 0.9271765 | cluster 2 |
| HORVU.MOREX.r3.3HG0294310.1 | 0.6319281 | 0.7702235 | 0.88165333 | cluster 2 |
| HORVU.MOREX.r3.3HG0294950.1 | 0.5721569 | 1.1659046 | 1.23153616 | cluster 2 |

**Table S3** Continued.

| ID | logFC_3 h | logFC_6 h | logFC_12 h | cluster No. |
|---|---|---|---|---|
| HORVU.MOREX.r3.3HG0294970.2 | 0.2580354 | 0.531544 | 0.85279494 | cluster 2 |
| HORVU.MOREX.r3.3HG0295380.1 | 0.244323 | 0.7373603 | 0.71161573 | cluster 2 |
| HORVU.MOREX.r3.3HG0296380.1 | 0.2139823 | 0.4967027 | 0.58990296 | cluster 2 |
| HORVU.MOREX.r3.3HG0296990.1 | -0.028178 | 0.9248453 | 0.84704542 | cluster 2 |
| HORVU.MOREX.r3.3HG0298090.1 | -0.054641 | 0.6582396 | 0.80351652 | cluster 2 |
| HORVU.MOREX.r3.3HG0298540.1 | 0.3203182 | 0.5833246 | 0.6584545 | cluster 2 |
| HORVU.MOREX.r3.3HG0301930.1 | 0.2016251 | 0.3889631 | 0.45439954 | cluster 2 |
| HORVU.MOREX.r3.3HG0303210.1 | 0.5020722 | 1.0455784 | 1.17174944 | cluster 2 |
| HORVU.MOREX.r3.3HG0303380.1 | 0.1781205 | 0.6848632 | 0.67693005 | cluster 2 |
| HORVU.MOREX.r3.3HG0303410.1 | 0.3390692 | 0.7666368 | 0.78439139 | cluster 2 |
| HORVU.MOREX.r3.3HG0303560.2 | 0.5327535 | 1.2702987 | 1.61822217 | cluster 2 |
| HORVU.MOREX.r3.3HG0303660.1 | 0.1834872 | 0.3823286 | 0.54769047 | cluster 2 |
| HORVU.MOREX.r3.3HG0304370.1 | 0.1834196 | 0.3751105 | 0.60819535 | cluster 2 |
| HORVU.MOREX.r3.3HG0304490.1 | 0.8293216 | 1.0838474 | 1.31250048 | cluster 2 |
| HORVU.MOREX.r3.3HG0305520.1 | 0.104132 | 0.5599306 | 0.69533456 | cluster 2 |
| HORVU.MOREX.r3.3HG0305800.1 | 0.4568399 | 0.724486 | 0.76647775 | cluster 2 |
| HORVU.MOREX.r3.3HG0306840.1 | 0.0239969 | 0.4021271 | 0.55810977 | cluster 2 |
| HORVU.MOREX.r3.3HG0307280.1 | 0.7045634 | 0.9952811 | 1.1344328 | cluster 2 |
| HORVU.MOREX.r3.3HG0307400.1 | 0.7801407 | 1.2549735 | 1.28158699 | cluster 2 |
| HORVU.MOREX.r3.3HG0308350.1 | 0.5269863 | 0.9100809 | 1.03734012 | cluster 2 |
| HORVU.MOREX.r3.3HG0308840.1 | 0.1481768 | 0.3742646 | 0.6264595 | cluster 2 |
| HORVU.MOREX.r3.3HG0309930.1 | 0.4885017 | 0.975444 | 1.05341105 | cluster 2 |
| HORVU.MOREX.r3.3HG0310920.1 | 0.2619564 | 0.5265518 | 0.62920463 | cluster 2 |
| HORVU.MOREX.r3.3HG0312430.1 | 0.8334222 | 2.0270515 | 1.90003817 | cluster 2 |
| HORVU.MOREX.r3.3HG0313900.1 | 1.5263673 | 1.7945304 | 1.87306177 | cluster 2 |
| HORVU.MOREX.r3.3HG0314070.1 | 0.4156637 | 1.3869242 | 1.44404811 | cluster 2 |
| HORVU.MOREX.r3.3HG0314820.1 | 0.3095286 | 0.5594509 | 0.70393134 | cluster 2 |
| HORVU.MOREX.r3.3HG0315410.1 | 0.1108427 | 0.2769116 | 0.35979591 | cluster 2 |
| HORVU.MOREX.r3.3HG0318700.1 | 0.1403915 | 0.3221063 | 0.50887756 | cluster 2 |
| HORVU.MOREX.r3.3HG0319900.1 | 1.0318526 | 1.5263922 | 1.5319189 | cluster 2 |
| HORVU.MOREX.r3.3HG0323600.1 | 0.1933303 | 0.4262943 | 0.53259787 | cluster 2 |
| HORVU.MOREX.r3.3HG0326330.1 | 0.3923725 | 0.9899287 | 1.03980027 | cluster 2 |
| HORVU.MOREX.r3.3HG0327100.1 | 0.1043931 | 0.9616848 | 1.16764068 | cluster 2 |
| HORVU.MOREX.r3.3HG0329040.1 | 1.5667103 | 2.4205926 | 2.62444447 | cluster 2 |
| HORVU.MOREX.r3.3HG0329870.1 | 1.3405947 | 1.8628077 | 2.00959851 | cluster 2 |
| HORVU.MOREX.r3.3HG0330120.1 | 0.9415784 | 1.5871535 | 2.2897583 | cluster 2 |
| HORVU.MOREX.r3.4HG0333450.1 | 0.4112081 | 0.6776574 | 0.76167709 | cluster 2 |
| HORVU.MOREX.r3.4HG0333650.1 | 0.0390368 | 0.3753369 | 0.43009465 | cluster 2 |
| HORVU.MOREX.r3.4HG0334360.1 | 0.4920948 | 1.0129823 | 1.10859449 | cluster 2 |
| HORVU.MOREX.r3.4HG0334600.1 | 0.2362238 | 0.4697044 | 0.60556625 | cluster 2 |
| HORVU.MOREX.r3.4HG0335160.1 | 0.575435 | 0.8010621 | 0.77857999 | cluster 2 |
| HORVU.MOREX.r3.4HG0335830.1 | 0.1900917 | 0.2945307 | 0.3712031 | cluster 2 |
| HORVU.MOREX.r3.4HG0337040.1 | 0.4629059 | 0.7655924 | 0.99127762 | cluster 2 |
| HORVU.MOREX.r3.4HG0337770.1 | 0.0355463 | 0.463949 | 0.57217308 | cluster 2 |
| HORVU.MOREX.r3.4HG0337870.1 | 0.1402813 | 0.8222738 | 0.75116839 | cluster 2 |
| HORVU.MOREX.r3.4HG0338070.1 | 0.5763057 | 0.9751026 | 1.07352575 | cluster 2 |
| HORVU.MOREX.r3.4HG0339890.1 | 0.9900853 | 1.6938507 | 1.68582548 | cluster 2 |

**Table S3** Continued.

| ID | logFC_3 h | logFC_6 h | logFC_12 h | cluster No. |
|---|---|---|---|---|
| HORVU.MOREX.r3.4HG0340910.2 | 1.5464803 | 2.3316632 | 2.52756548 | cluster 2 |
| HORVU.MOREX.r3.4HG0341050.1 | 0.6468136 | 1.3654148 | 1.50733577 | cluster 2 |
| HORVU.MOREX.r3.4HG0341800.1 | 0.3931678 | 0.6513793 | 0.83585319 | cluster 2 |
| HORVU.MOREX.r3.4HG0342080.1 | 0.2705359 | 0.6939733 | 0.7656478 | cluster 2 |
| HORVU.MOREX.r3.4HG0342660.1 | 0.6725532 | 1.2454115 | 1.266999 | cluster 2 |
| HORVU.MOREX.r3.4HG0343040.1 | 0.1746996 | 0.5204223 | 0.7760771 | cluster 2 |
| HORVU.MOREX.r3.4HG0343050.1 | 0.2833578 | 0.4298264 | 0.49476416 | cluster 2 |
| HORVU.MOREX.r3.4HG0343260.1 | 0.5825461 | 1.1123837 | 1.41752398 | cluster 2 |
| HORVU.MOREX.r3.4HG0344370.1 | 0.8558216 | 1.6664461 | 1.8305369 | cluster 2 |
| HORVU.MOREX.r3.4HG0344790.1 | 0.2268993 | 0.4391813 | 0.45317672 | cluster 2 |
| HORVU.MOREX.r3.4HG0345260.1 | 1.0256867 | 1.1938476 | 1.20978148 | cluster 2 |
| HORVU.MOREX.r3.4HG0345990.1 | -0.064489 | 0.3767458 | 0.51143182 | cluster 2 |
| HORVU.MOREX.r3.4HG0346830.1 | 1.1709522 | 1.9494157 | 2.07744307 | cluster 2 |
| HORVU.MOREX.r3.4HG0347760.1 | 0.1447569 | 0.4451303 | 0.66666929 | cluster 2 |
| HORVU.MOREX.r3.4HG0348380.1 | 0.5362334 | 0.8289037 | 0.91855597 | cluster 2 |
| HORVU.MOREX.r3.4HG0349080.1 | 0.2161735 | 0.4842153 | 0.72832219 | cluster 2 |
| HORVU.MOREX.r3.4HG0351070.1 | 0.0673531 | 0.3588632 | 0.74699814 | cluster 2 |
| HORVU.MOREX.r3.4HG0352200.1 | 0.45411 | 0.6246446 | 0.68749416 | cluster 2 |
| HORVU.MOREX.r3.4HG0352780.1 | 0.2452678 | 0.5738966 | 0.75116078 | cluster 2 |
| HORVU.MOREX.r3.4HG0353390.1 | 0.2763529 | 0.8060027 | 0.84907564 | cluster 2 |
| HORVU.MOREX.r3.4HG0354010.1 | 1.6504168 | 2.1037581 | 2.2176036 | cluster 2 |
| HORVU.MOREX.r3.4HG0354980.1 | 0.477069 | 0.6872052 | 0.97622784 | cluster 2 |
| HORVU.MOREX.r3.4HG0356280.1 | 0.4156574 | 0.955948 | 1.0671736 | cluster 2 |
| HORVU.MOREX.r3.4HG0356690.1 | 0.496617 | 0.8638757 | 1.10307239 | cluster 2 |
| HORVU.MOREX.r3.4HG0356860.1 | 0.4296005 | 0.9932794 | 0.93839516 | cluster 2 |
| HORVU.MOREX.r3.4HG0359960.1 | 0.0481177 | 0.2615211 | 0.3106003 | cluster 2 |
| HORVU.MOREX.r3.4HG0364050.1 | 0.3615742 | 0.7555068 | 0.76165059 | cluster 2 |
| HORVU.MOREX.r3.4HG0378080.1 | 1.3501296 | 2.8965001 | 2.88167628 | cluster 2 |
| HORVU.MOREX.r3.4HG0379260.1 | 0.3001805 | 0.5027635 | 0.63639233 | cluster 2 |
| HORVU.MOREX.r3.4HG0379400.1 | 0.4957518 | 0.8120354 | 1.06799756 | cluster 2 |
| HORVU.MOREX.r3.4HG0379660.1 | 0.6626674 | 1.4889937 | 1.43081562 | cluster 2 |
| HORVU.MOREX.r3.4HG0380540.1 | 0.4686172 | 0.7116699 | 0.8090097 | cluster 2 |
| HORVU.MOREX.r3.4HG0381790.1 | 0.4115763 | 1.1527833 | 1.08853007 | cluster 2 |
| HORVU.MOREX.r3.4HG0381990.1 | 0.3432611 | 0.924176 | 1.46516005 | cluster 2 |
| HORVU.MOREX.r3.4HG0382550.1 | 0.2568968 | 0.6590222 | 0.63661169 | cluster 2 |
| HORVU.MOREX.r3.4HG0382610.1 | 0.6480635 | 1.3044924 | 1.24352216 | cluster 2 |
| HORVU.MOREX.r3.4HG0383280.1 | 0.1986514 | 0.4795672 | 0.70614599 | cluster 2 |
| HORVU.MOREX.r3.4HG0383530.3 | 0.3284265 | 0.7387163 | 0.94778101 | cluster 2 |
| HORVU.MOREX.r3.4HG0383670.1 | 0.5060547 | 1.153631 | 1.29406186 | cluster 2 |
| HORVU.MOREX.r3.4HG0383780.1 | 0.799171 | 1.8433864 | 1.80427655 | cluster 2 |
| HORVU.MOREX.r3.4HG0384390.1 | 0.8730756 | 1.1942553 | 1.39596124 | cluster 2 |
| HORVU.MOREX.r3.4HG0384400.1 | 0.2391783 | 0.4010898 | 0.50728861 | cluster 2 |
| HORVU.MOREX.r3.4HG0384780.1 | 1.8829206 | 3.1804654 | 3.25352083 | cluster 2 |
| HORVU.MOREX.r3.4HG0386800.1 | 0.0156825 | 0.2490754 | 0.28139142 | cluster 2 |
| HORVU.MOREX.r3.4HG0387580.1 | 0.0049719 | 0.2238315 | 0.32716647 | cluster 2 |
| HORVU.MOREX.r3.4HG0388310.1 | 0.3021673 | 0.4118418 | 0.42517248 | cluster 2 |
| HORVU.MOREX.r3.4HG0388440.1 | 0.3905342 | 0.8014992 | 0.87568721 | cluster 2 |

**Table S3** Continued.

| ID | logFC_3 h | logFC_6 h | logFC_12 h | cluster No. |
|---|---|---|---|---|
| HORVU.MOREX.r3.4HG0389200.1 | 0.4199592 | 0.7002946 | 0.81906453 | cluster 2 |
| HORVU.MOREX.r3.4HG0391130.1 | 0.2318906 | 0.4926743 | 0.47882861 | cluster 2 |
| HORVU.MOREX.r3.4HG0391260.1 | 0.2829957 | 0.4690374 | 0.61470029 | cluster 2 |
| HORVU.MOREX.r3.4HG0392230.1 | 0.6502808 | 1.0714124 | 1.38221333 | cluster 2 |
| HORVU.MOREX.r3.4HG0392680.1 | 0.3636547 | 0.5668818 | 0.79741976 | cluster 2 |
| HORVU.MOREX.r3.4HG0392940.1 | 0.2260992 | 0.4908348 | 0.52022384 | cluster 2 |
| HORVU.MOREX.r3.4HG0393520.1 | 0.272282 | 0.7815714 | 0.74873951 | cluster 2 |
| HORVU.MOREX.r3.4HG0393710.1 | 0.2070321 | 0.8328329 | 1.00129397 | cluster 2 |
| HORVU.MOREX.r3.4HG0393870.1 | 0.8422932 | 1.5099747 | 1.68495442 | cluster 2 |
| HORVU.MOREX.r3.4HG0394870.1 | 0.2343377 | 0.8269788 | 1.10620424 | cluster 2 |
| HORVU.MOREX.r3.4HG0395940.1 | 0.9225598 | 1.4318554 | 1.45687047 | cluster 2 |
| HORVU.MOREX.r3.4HG0396030.1 | 0.9705374 | 1.9385054 | 1.83872192 | cluster 2 |
| HORVU.MOREX.r3.4HG0397010.1 | 0.9666427 | 1.5883306 | 1.92837147 | cluster 2 |
| HORVU.MOREX.r3.4HG0397530.1 | 0.3634674 | 0.8982456 | 1.0804083 | cluster 2 |
| HORVU.MOREX.r3.4HG0398050.1 | 0.7947834 | 1.416871 | 1.59561309 | cluster 2 |
| HORVU.MOREX.r3.4HG0398090.1 | 0.2618222 | 0.6719989 | 0.81460377 | cluster 2 |
| HORVU.MOREX.r3.4HG0398550.1 | 0.4094856 | 0.9569696 | 1.04927534 | cluster 2 |
| HORVU.MOREX.r3.4HG0399470.1 | 0.8254288 | 2.2371615 | 2.52968369 | cluster 2 |
| HORVU.MOREX.r3.4HG0399530.1 | 1.2854404 | 1.7500597 | 1.70869347 | cluster 2 |
| HORVU.MOREX.r3.4HG0400040.1 | 0.3088012 | 0.4532839 | 0.60372683 | cluster 2 |
| HORVU.MOREX.r3.4HG0400840.1 | 0.4045477 | 0.7406587 | 0.90631406 | cluster 2 |
| HORVU.MOREX.r3.4HG0400920.1 | 0.7049011 | 1.2452702 | 1.31393698 | cluster 2 |
| HORVU.MOREX.r3.4HG0401630.1 | 0.4531026 | 0.6001547 | 0.67908155 | cluster 2 |
| HORVU.MOREX.r3.4HG0401980.1 | 1.4130215 | 2.7376907 | 2.83320209 | cluster 2 |
| HORVU.MOREX.r3.4HG0402650.1 | 0.3316932 | 0.762461 | 0.82507227 | cluster 2 |
| HORVU.MOREX.r3.4HG0402730.1 | 1.0848712 | 1.9165727 | 1.83720141 | cluster 2 |
| HORVU.MOREX.r3.4HG0404740.1 | 0.181578 | 0.4822729 | 0.80757972 | cluster 2 |
| HORVU.MOREX.r3.4HG0405400.1 | 0.9809624 | 1.8846201 | 1.84235306 | cluster 2 |
| HORVU.MOREX.r3.4HG0405780.1 | 0.2309227 | 0.4082526 | 0.63931514 | cluster 2 |
| HORVU.MOREX.r3.4HG0405920.1 | 0.6550032 | 1.5808902 | 1.53146869 | cluster 2 |
| HORVU.MOREX.r3.4HG0406060.1 | 0.8964474 | 2.2873495 | 2.60646335 | cluster 2 |
| HORVU.MOREX.r3.4HG0406140.1 | 0.0768009 | 0.4622742 | 0.48942009 | cluster 2 |
| HORVU.MOREX.r3.4HG0406600.1 | 0.5288912 | 0.7448671 | 0.88836529 | cluster 2 |
| HORVU.MOREX.r3.4HG0406630.1 | 0.2646697 | 0.5942882 | 0.71951802 | cluster 2 |
| HORVU.MOREX.r3.4HG0407470.1 | 0.4171851 | 0.8652511 | 0.89349686 | cluster 2 |
| HORVU.MOREX.r3.4HG0407990.1 | 0.2373114 | 0.6788043 | 0.87073362 | cluster 2 |
| HORVU.MOREX.r3.4HG0408000.1 | 1.1643976 | 1.8015354 | 2.63989086 | cluster 2 |
| HORVU.MOREX.r3.4HG0408780.1 | -0.943287 | 2.0851534 | 1.97751546 | cluster 2 |
| HORVU.MOREX.r3.4HG0409670.1 | 0.8149989 | 1.0992152 | 1.17181536 | cluster 2 |
| HORVU.MOREX.r3.4HG0410090.1 | 0.3287014 | 0.4503451 | 0.5981346 | cluster 2 |
| HORVU.MOREX.r3.4HG0412090.1 | 0.7596638 | 1.8386347 | 1.81770888 | cluster 2 |
| HORVU.MOREX.r3.4HG0412430.1 | -0.022167 | 0.4239312 | 0.43177193 | cluster 2 |
| HORVU.MOREX.r3.4HG0412790.1 | 0.532284 | 0.6579071 | 0.73362573 | cluster 2 |
| HORVU.MOREX.r3.4HG0413800.1 | 0.1734431 | 0.3904929 | 0.47772268 | cluster 2 |
| HORVU.MOREX.r3.4HG0415170.1 | 0.2963256 | 1.1164611 | 1.54423623 | cluster 2 |
| HORVU.MOREX.r3.4HG0416390.1 | 0.9077139 | 1.2040081 | 1.34727728 | cluster 2 |
| HORVU.MOREX.r3.4HG0417010.1 | 1.1506263 | 1.9261093 | 1.94717723 | cluster 2 |

**Table S3** Continued.

| ID | logFC_3 h | logFC_6 h | logFC_12 h | cluster No. |
|---|---|---|---|---|
| HORVU.MOREX.r3.4HG0417160.1 | 0.2855021 | 0.5117306 | 0.5733361 | cluster 2 |
| HORVU.MOREX.r3.4HG0417320.1 | 0.8115794 | 1.409745 | 1.5882952 | cluster 2 |
| HORVU.MOREX.r3.4HG0417670.1 | 0.8401505 | 1.218021 | 1.19228366 | cluster 2 |
| HORVU.MOREX.r3.4HG0418460.1 | 0.2791984 | 0.3610209 | 0.39910965 | cluster 2 |
| HORVU.MOREX.r3.5HG0419590.1 | 0.1288701 | 0.4173495 | 0.51122227 | cluster 2 |
| HORVU.MOREX.r3.5HG0420080.1 | 0.1730085 | 0.3711592 | 0.42195773 | cluster 2 |
| HORVU.MOREX.r3.5HG0420830.1 | 0.3019412 | 1.1288759 | 1.28831804 | cluster 2 |
| HORVU.MOREX.r3.5HG0420970.1 | 0.3232078 | 0.6296639 | 0.89150456 | cluster 2 |
| HORVU.MOREX.r3.5HG0420980.1 | 0.3949095 | 0.5844236 | 0.79194101 | cluster 2 |
| HORVU.MOREX.r3.5HG0422080.1 | 0.2802278 | 0.4229508 | 0.4267247 | cluster 2 |
| HORVU.MOREX.r3.5HG0425250.1 | 0.7698766 | 1.2337542 | 1.55042174 | cluster 2 |
| HORVU.MOREX.r3.5HG0428270.2 | 0.0671169 | 0.4139552 | 0.69077879 | cluster 2 |
| HORVU.MOREX.r3.5HG0430940.1 | 0.4923077 | 0.9284412 | 1.09685903 | cluster 2 |
| HORVU.MOREX.r3.5HG0432950.1 | 0.1672651 | 0.4212326 | 0.45154427 | cluster 2 |
| HORVU.MOREX.r3.5HG0433470.1 | 0.2047951 | 0.2934084 | 0.35811409 | cluster 2 |
| HORVU.MOREX.r3.5HG0437110.1 | 0.1078287 | 0.2902869 | 0.50892863 | cluster 2 |
| HORVU.MOREX.r3.5HG0442460.1 | 0.1475444 | 0.3101127 | 0.44211993 | cluster 2 |
| HORVU.MOREX.r3.5HG0444350.1 | 0.2384017 | 0.7529188 | 0.8246533 | cluster 2 |
| HORVU.MOREX.r3.5HG0446420.1 | 0.399862 | 0.6221633 | 0.76528343 | cluster 2 |
| HORVU.MOREX.r3.5HG0449620.1 | 0.3227242 | 0.6800839 | 0.72840761 | cluster 2 |
| HORVU.MOREX.r3.5HG0454230.1 | 0.3450621 | 0.7756468 | 0.75101708 | cluster 2 |
| HORVU.MOREX.r3.5HG0457310.1 | 0.3793025 | 1.1601469 | 1.1265514 | cluster 2 |
| HORVU.MOREX.r3.5HG0458010.1 | 0.1073535 | 0.182246 | 0.27477234 | cluster 2 |
| HORVU.MOREX.r3.5HG0461280.1 | 0.3619618 | 0.5180967 | 0.5913279 | cluster 2 |
| HORVU.MOREX.r3.5HG0461950.1 | -0.007288 | 0.8501847 | 0.80316713 | cluster 2 |
| HORVU.MOREX.r3.5HG0462220.1 | 0.368836 | 0.7981703 | 0.94321337 | cluster 2 |
| HORVU.MOREX.r3.5HG0463550.1 | 0.4243225 | 0.8470079 | 1.04395853 | cluster 2 |
| HORVU.MOREX.r3.5HG0464440.1 | 0.8676465 | 1.0462781 | 1.03930402 | cluster 2 |
| HORVU.MOREX.r3.5HG0465270.1 | 0.3607954 | 0.6702523 | 0.74170566 | cluster 2 |
| HORVU.MOREX.r3.5HG0465400.1 | 0.5373646 | 1.1918769 | 1.21332619 | cluster 2 |
| HORVU.MOREX.r3.5HG0465690.1 | 0.2593417 | 0.5542937 | 0.82829633 | cluster 2 |
| HORVU.MOREX.r3.5HG0469120.1 | 1.2168502 | 1.5890204 | 1.73777658 | cluster 2 |
| HORVU.MOREX.r3.5HG0471520.1 | 0.2693765 | 0.3952702 | 0.54907622 | cluster 2 |
| HORVU.MOREX.r3.5HG0472810.1 | 0.4125309 | 0.8835838 | 0.85386721 | cluster 2 |
| HORVU.MOREX.r3.5HG0472840.1 | 0.0019756 | 0.2382994 | 0.5109538 | cluster 2 |
| HORVU.MOREX.r3.5HG0472870.1 | 0.193304 | 0.527352 | 0.63912229 | cluster 2 |
| HORVU.MOREX.r3.5HG0473520.1 | 0.1275543 | 0.2431925 | 0.39199122 | cluster 2 |
| HORVU.MOREX.r3.5HG0477220.1 | 0.7841996 | 1.0146353 | 1.05301851 | cluster 2 |
| HORVU.MOREX.r3.5HG0477800.1 | 0.4977347 | 0.6934595 | 0.74789724 | cluster 2 |
| HORVU.MOREX.r3.5HG0478050.1 | 0.2641038 | 0.45175 | 0.60793165 | cluster 2 |
| HORVU.MOREX.r3.5HG0479500.1 | 1.2590247 | 1.4855937 | 1.60256641 | cluster 2 |
| HORVU.MOREX.r3.5HG0479810.1 | 0.0426092 | 0.1361107 | 0.23748287 | cluster 2 |
| HORVU.MOREX.r3.5HG0480600.1 | -0.152634 | 1.3959017 | 1.84601562 | cluster 2 |
| HORVU.MOREX.r3.5HG0482670.1 | 0.8283812 | 1.3690747 | 1.54939215 | cluster 2 |
| HORVU.MOREX.r3.5HG0484180.1 | 0.301845 | 0.4084648 | 0.44929194 | cluster 2 |
| HORVU.MOREX.r3.5HG0484660.1 | 0.6494517 | 1.4891231 | 1.49141657 | cluster 2 |
| HORVU.MOREX.r3.5HG0485610.1 | 0.8402966 | 1.1851522 | 1.35797644 | cluster 2 |

**Table S3** Continued.

| ID | logFC_3 h | logFC_6 h | logFC_12 h | cluster No. |
|---|---|---|---|---|
| HORVU.MOREX.r3.5HG0486610.1 | 0.3195001 | 0.728249 | 0.83150623 | cluster 2 |
| HORVU.MOREX.r3.5HG0486750.1 | 0.2180509 | 0.7341424 | 0.87387666 | cluster 2 |
| HORVU.MOREX.r3.5HG0490520.1 | 0.1978655 | 0.6820398 | 0.87944905 | cluster 2 |
| HORVU.MOREX.r3.5HG0490610.1 | 0.8011162 | 1.8063093 | 1.71784466 | cluster 2 |
| HORVU.MOREX.r3.5HG0490800.1 | 0.9275562 | 1.2475257 | 1.64621621 | cluster 2 |
| HORVU.MOREX.r3.5HG0493040.1 | 0.2080434 | 0.5060233 | 0.55263335 | cluster 2 |
| HORVU.MOREX.r3.5HG0494040.1 | 0.2203348 | 1.6371662 | 1.85311292 | cluster 2 |
| HORVU.MOREX.r3.5HG0495240.1 | 0.4517982 | 0.773699 | 1.05799994 | cluster 2 |
| HORVU.MOREX.r3.5HG0495730.1 | 0.5876246 | 0.9852875 | 1.15328876 | cluster 2 |
| HORVU.MOREX.r3.5HG0495860.1 | 0.5515498 | 0.7776646 | 0.76335005 | cluster 2 |
| HORVU.MOREX.r3.5HG0496780.1 | 0.173293 | 0.2889691 | 0.43670843 | cluster 2 |
| HORVU.MOREX.r3.5HG0496930.1 | 0.2323162 | 0.6896972 | 0.83641896 | cluster 2 |
| HORVU.MOREX.r3.5HG0497490.1 | 0.8369841 | 1.3476139 | 1.76946037 | cluster 2 |
| HORVU.MOREX.r3.5HG0497900.1 | 0.5995338 | 0.9889431 | 0.99618653 | cluster 2 |
| HORVU.MOREX.r3.5HG0501170.1 | 0.3852307 | 0.5130394 | 0.64276813 | cluster 2 |
| HORVU.MOREX.r3.5HG0501380.1 | 0.0844627 | 0.2130409 | 0.34496489 | cluster 2 |
| HORVU.MOREX.r3.5HG0504720.1 | 0.1014129 | 0.3259423 | 0.30528838 | cluster 2 |
| HORVU.MOREX.r3.5HG0508870.1 | 0.2668091 | 0.5499848 | 0.75040954 | cluster 2 |
| HORVU.MOREX.r3.5HG0509020.1 | 0.2692762 | 0.7428903 | 1.01310401 | cluster 2 |
| HORVU.MOREX.r3.5HG0511090.1 | 0.3390588 | 0.5528819 | 0.69682833 | cluster 2 |
| HORVU.MOREX.r3.5HG0511450.1 | 0.3371689 | 0.4258705 | 0.43400053 | cluster 2 |
| HORVU.MOREX.r3.5HG0513740.1 | 0.6270661 | 1.1857086 | 1.59899486 | cluster 2 |
| HORVU.MOREX.r3.5HG0513780.1 | 0.1784336 | 0.6028146 | 0.64852256 | cluster 2 |
| HORVU.MOREX.r3.5HG0513900.1 | 0.5238724 | 0.6676236 | 0.78546448 | cluster 2 |
| HORVU.MOREX.r3.5HG0513980.1 | 0.7987007 | 1.2576441 | 1.34151688 | cluster 2 |
| HORVU.MOREX.r3.5HG0517190.1 | 0.3748177 | 0.612034 | 0.70295419 | cluster 2 |
| HORVU.MOREX.r3.5HG0517240.1 | 0.1112988 | 0.2599837 | 0.39148009 | cluster 2 |
| HORVU.MOREX.r3.5HG0517260.1 | 0.2473357 | 0.4833552 | 0.54054673 | cluster 2 |
| HORVU.MOREX.r3.5HG0517490.1 | 0.2353696 | 0.4646789 | 0.764187 | cluster 2 |
| HORVU.MOREX.r3.5HG0519360.1 | 0.4833657 | 0.6959955 | 0.96840789 | cluster 2 |
| HORVU.MOREX.r3.5HG0521630.1 | 0.0963929 | 0.513868 | 0.66457976 | cluster 2 |
| HORVU.MOREX.r3.5HG0523860.1 | 0.1661731 | 0.3948824 | 0.5181811 | cluster 2 |
| HORVU.MOREX.r3.5HG0524060.1 | 0.7913655 | 1.2930894 | 1.32015096 | cluster 2 |
| HORVU.MOREX.r3.5HG0525430.1 | 0.3646986 | 0.7086655 | 0.74596087 | cluster 2 |
| HORVU.MOREX.r3.5HG0525850.1 | 0.2909068 | 0.5240224 | 0.68520648 | cluster 2 |
| HORVU.MOREX.r3.5HG0527460.1 | 0.9758297 | 1.4137042 | 1.40692609 | cluster 2 |
| HORVU.MOREX.r3.5HG0527720.1 | 0.7336271 | 0.9795008 | 1.0821862 | cluster 2 |
| HORVU.MOREX.r3.5HG0528280.1 | 0.2352558 | 0.4249636 | 0.51270501 | cluster 2 |
| HORVU.MOREX.r3.5HG0528310.1 | 0.5307114 | 0.5710331 | 0.60443093 | cluster 2 |
| HORVU.MOREX.r3.5HG0531070.1 | 2.5625885 | 3.4057006 | 3.68460448 | cluster 2 |
| HORVU.MOREX.r3.5HG0532200.1 | 0.541597 | 0.6596337 | 0.81273203 | cluster 2 |
| HORVU.MOREX.r3.5HG0532560.1 | 1.4614443 | 1.8418639 | 1.89784638 | cluster 2 |
| HORVU.MOREX.r3.5HG0533760.1 | 0.8064972 | 1.2863139 | 1.36594148 | cluster 2 |
| HORVU.MOREX.r3.5HG0534420.1 | 0.343871 | 0.784436 | 0.84107089 | cluster 2 |
| HORVU.MOREX.r3.5HG0535180.1 | 1.1571559 | 1.6750447 | 1.72444736 | cluster 2 |
| HORVU.MOREX.r3.5HG0536200.1 | 0.8263819 | 1.101929 | 1.076437 | cluster 2 |
| HORVU.MOREX.r3.5HG0536610.1 | 0.1008134 | 0.6016052 | 0.65610442 | cluster 2 |

**Table S3** Continued.

| ID | logFC_3 h | logFC_6 h | logFC_12 h | cluster No. |
|---|---|---|---|---|
| HORVU.MOREX.r3.5HG0537150.1 | 0.7193756 | 1.4313003 | 1.70040162 | cluster 2 |
| HORVU.MOREX.r3.5HG0537360.1 | 1.4914172 | 2.099833 | 2.28994247 | cluster 2 |
| HORVU.MOREX.r3.6HG0539590.1 | 0.5752701 | 0.7797706 | 0.98364708 | cluster 2 |
| HORVU.MOREX.r3.6HG0542030.1 | 0.6563637 | 1.0902273 | 1.30863958 | cluster 2 |
| HORVU.MOREX.r3.6HG0542040.1 | 0.520591 | 0.9251602 | 1.32881351 | cluster 2 |
| HORVU.MOREX.r3.6HG0542050.1 | 0.8135932 | 1.2932291 | 1.55494779 | cluster 2 |
| HORVU.MOREX.r3.6HG0542320.1 | 0.3619709 | 1.2739338 | 1.21269683 | cluster 2 |
| HORVU.MOREX.r3.6HG0544350.1 | 0.1825648 | 0.3996631 | 0.42312297 | cluster 2 |
| HORVU.MOREX.r3.6HG0546160.1 | 0.4494889 | 0.7885519 | 1.07728293 | cluster 2 |
| HORVU.MOREX.r3.6HG0547010.1 | 0.4113203 | 0.6171305 | 0.88044854 | cluster 2 |
| HORVU.MOREX.r3.6HG0547020.1 | 0.3323692 | 0.905998 | 1.15425002 | cluster 2 |
| HORVU.MOREX.r3.6HG0547030.1 | 0.4545781 | 0.7414847 | 1.11423101 | cluster 2 |
| HORVU.MOREX.r3.6HG0547570.1 | 0.0745734 | 0.2591938 | 0.3948071 | cluster 2 |
| HORVU.MOREX.r3.6HG0547660.1 | 1.4830317 | 2.2438891 | 2.39181162 | cluster 2 |
| HORVU.MOREX.r3.6HG0549810.1 | 0.1829943 | 0.3580765 | 0.52235269 | cluster 2 |
| HORVU.MOREX.r3.6HG0550600.1 | 0.8517285 | 1.3576493 | 1.56083185 | cluster 2 |
| HORVU.MOREX.r3.6HG0551650.1 | 0.2633161 | 0.4528928 | 0.43997862 | cluster 2 |
| HORVU.MOREX.r3.6HG0554050.1 | 0.5180914 | 0.8651744 | 0.89134845 | cluster 2 |
| HORVU.MOREX.r3.6HG0554700.1 | 0.0898452 | 0.2560453 | 0.30452703 | cluster 2 |
| HORVU.MOREX.r3.6HG0554840.1 | 0.2615541 | 0.6312156 | 0.87777144 | cluster 2 |
| HORVU.MOREX.r3.6HG0555040.1 | 0.287924 | 0.5201071 | 0.52012828 | cluster 2 |
| HORVU.MOREX.r3.6HG0555250.1 | 0.3416737 | 1.7096039 | 2.16806086 | cluster 2 |
| HORVU.MOREX.r3.6HG0556240.1 | 0.2417086 | 0.3349085 | 0.45552598 | cluster 2 |
| HORVU.MOREX.r3.6HG0557920.1 | 0.4135744 | 0.9231393 | 1.06960137 | cluster 2 |
| HORVU.MOREX.r3.6HG0559990.1 | 0.464411 | 0.8158961 | 0.94593506 | cluster 2 |
| HORVU.MOREX.r3.6HG0560710.1 | 0.1098479 | 0.2120663 | 0.26306374 | cluster 2 |
| HORVU.MOREX.r3.6HG0560930.1 | 0.3078785 | 0.4763428 | 0.64037642 | cluster 2 |
| HORVU.MOREX.r3.6HG0565950.1 | 0.8249902 | 1.1625846 | 1.3935857 | cluster 2 |
| HORVU.MOREX.r3.6HG0566940.1 | 0.3821272 | 0.5756211 | 0.58031974 | cluster 2 |
| HORVU.MOREX.r3.6HG0567150.1 | 0.5391259 | 1.0741839 | 1.15932802 | cluster 2 |
| HORVU.MOREX.r3.6HG0567370.2 | 0.3978744 | 0.7186024 | 0.79653984 | cluster 2 |
| HORVU.MOREX.r3.6HG0568910.1 | 0.3789607 | 0.5723245 | 0.82335041 | cluster 2 |
| HORVU.MOREX.r3.6HG0570430.1 | -0.06227 | 0.224238 | 0.37468298 | cluster 2 |
| HORVU.MOREX.r3.6HG0573150.1 | 0.1594812 | 0.2897869 | 0.3891766 | cluster 2 |
| HORVU.MOREX.r3.6HG0573290.1 | 0.1302786 | 0.3153781 | 0.31968137 | cluster 2 |
| HORVU.MOREX.r3.6HG0573920.1 | 0.766951 | 1.4144927 | 1.47543076 | cluster 2 |
| HORVU.MOREX.r3.6HG0574950.1 | 0.2496861 | 0.4242427 | 0.65298771 | cluster 2 |
| HORVU.MOREX.r3.6HG0582120.1 | 0.20045 | 0.4747158 | 0.47731547 | cluster 2 |
| HORVU.MOREX.r3.6HG0582480.1 | 0.3956409 | 0.7421302 | 0.92358983 | cluster 2 |
| HORVU.MOREX.r3.6HG0585240.1 | 0.5852624 | 1.0308635 | 1.03198216 | cluster 2 |
| HORVU.MOREX.r3.6HG0588520.1 | 0.4279784 | 0.9427609 | 0.97156598 | cluster 2 |
| HORVU.MOREX.r3.6HG0589860.1 | 0.424529 | 0.7956522 | 0.79276116 | cluster 2 |
| HORVU.MOREX.r3.6HG0593050.1 | 0.3094143 | 0.588504 | 0.91078424 | cluster 2 |
| HORVU.MOREX.r3.6HG0593390.1 | 1.0727411 | 1.2641774 | 1.36503238 | cluster 2 |
| HORVU.MOREX.r3.6HG0596770.1 | 0.3412317 | 0.6308943 | 0.73754549 | cluster 2 |
| HORVU.MOREX.r3.6HG0597400.2 | 0.4998391 | 0.90667 | 1.31877775 | cluster 2 |
| HORVU.MOREX.r3.6HG0599900.1 | 0.2909016 | 0.5333152 | 0.62981831 | cluster 2 |

**Table S3** Continued.

| ID | logFC_3 h | logFC_6 h | logFC_12 h | cluster No. |
|---|---|---|---|---|
| HORVU.MOREX.r3.6HG0601160.1 | 0.3158278 | 0.5490306 | 0.68381175 | cluster 2 |
| HORVU.MOREX.r3.6HG0601180.1 | 0.488909 | 0.7011847 | 0.91844231 | cluster 2 |
| HORVU.MOREX.r3.6HG0601260.1 | 0.511754 | 1.0425773 | 1.10705238 | cluster 2 |
| HORVU.MOREX.r3.6HG0601680.1 | 0.415202 | 0.6723902 | 0.8621493 | cluster 2 |
| HORVU.MOREX.r3.6HG0601830.1 | 1.0243086 | 1.6885951 | 1.74978443 | cluster 2 |
| HORVU.MOREX.r3.6HG0602210.1 | 0.5105762 | 0.6931835 | 0.84292741 | cluster 2 |
| HORVU.MOREX.r3.6HG0603210.1 | 0.7727444 | 1.4604009 | 2.04263675 | cluster 2 |
| HORVU.MOREX.r3.6HG0605150.1 | 0.3535078 | 0.6842286 | 0.90491722 | cluster 2 |
| HORVU.MOREX.r3.6HG0605160.1 | 0.1637581 | 0.3544376 | 0.48022311 | cluster 2 |
| HORVU.MOREX.r3.6HG0605940.1 | 0.3250967 | 0.6384968 | 0.71767755 | cluster 2 |
| HORVU.MOREX.r3.6HG0605960.1 | 0.277122 | 0.5627489 | 0.68714776 | cluster 2 |
| HORVU.MOREX.r3.6HG0606140.1 | -0.141648 | 1.0420557 | 2.00505825 | cluster 2 |
| HORVU.MOREX.r3.6HG0606360.1 | 1.031606 | 2.0810804 | 2.19827158 | cluster 2 |
| HORVU.MOREX.r3.6HG0607960.1 | 0.1238032 | 0.3404552 | 0.53025735 | cluster 2 |
| HORVU.MOREX.r3.6HG0608910.1 | 0.762652 | 1.0527539 | 1.21866412 | cluster 2 |
| HORVU.MOREX.r3.6HG0610500.1 | 0.5111657 | 0.9589503 | 1.09267208 | cluster 2 |
| HORVU.MOREX.r3.6HG0610890.1 | 0.1242264 | 0.3310396 | 0.4677181 | cluster 2 |
| HORVU.MOREX.r3.6HG0614110.1 | 0.1992911 | 0.3322784 | 0.39232921 | cluster 2 |
| HORVU.MOREX.r3.6HG0614840.1 | 0.0167346 | 0.3206517 | 0.56924838 | cluster 2 |
| HORVU.MOREX.r3.6HG0614870.1 | 0.3853101 | 0.670507 | 0.71305407 | cluster 2 |
| HORVU.MOREX.r3.6HG0615130.1 | 0.2970304 | 0.663016 | 0.68977244 | cluster 2 |
| HORVU.MOREX.r3.6HG0615630.1 | 0.1912273 | 0.4608209 | 0.47468987 | cluster 2 |
| HORVU.MOREX.r3.6HG0616130.1 | 0.19623 | 0.4546644 | 0.58864528 | cluster 2 |
| HORVU.MOREX.r3.6HG0616300.1 | 0.1392149 | 0.3896496 | 0.5502242 | cluster 2 |
| HORVU.MOREX.r3.6HG0617080.1 | 0.1546483 | 0.3857756 | 0.5070965 | cluster 2 |
| HORVU.MOREX.r3.6HG0619500.1 | 0.6282363 | 1.4464878 | 1.49013116 | cluster 2 |
| HORVU.MOREX.r3.6HG0620520.1 | 0.3558299 | 0.7667271 | 0.9000793 | cluster 2 |
| HORVU.MOREX.r3.6HG0620630.1 | 0.4602424 | 0.9172385 | 1.01213706 | cluster 2 |
| HORVU.MOREX.r3.6HG0620720.1 | 0.6013842 | 0.9252779 | 0.95546276 | cluster 2 |
| HORVU.MOREX.r3.6HG0622080.1 | 0.3824593 | 0.6294396 | 0.69296603 | cluster 2 |
| HORVU.MOREX.r3.6HG0622120.1 | 0.2692973 | 0.4714377 | 0.57379924 | cluster 2 |
| HORVU.MOREX.r3.6HG0622250.1 | 0.9393621 | 1.4391752 | 1.38710931 | cluster 2 |
| HORVU.MOREX.r3.6HG0624240.1 | 0.4357281 | 0.7386809 | 0.86573578 | cluster 2 |
| HORVU.MOREX.r3.6HG0624460.1 | 0.0187827 | 1.2833566 | 2.05973085 | cluster 2 |
| HORVU.MOREX.r3.6HG0625570.1 | 0.3311309 | 0.5716644 | 0.65982525 | cluster 2 |
| HORVU.MOREX.r3.6HG0626070.1 | 0.2709765 | 0.4237318 | 0.56957533 | cluster 2 |
| HORVU.MOREX.r3.6HG0628790.1 | 0.2688086 | 0.4943078 | 0.72855732 | cluster 2 |
| HORVU.MOREX.r3.6HG0629240.1 | 1.3152298 | 1.4994885 | 1.74505029 | cluster 2 |
| HORVU.MOREX.r3.6HG0631600.1 | 0.5506977 | 1.1166081 | 1.12470387 | cluster 2 |
| HORVU.MOREX.r3.6HG0631700.1 | 0.1635609 | 0.3671579 | 0.44161692 | cluster 2 |
| HORVU.MOREX.r3.6HG0631710.1 | 0.6147842 | 1.0170948 | 1.05782311 | cluster 2 |
| HORVU.MOREX.r3.6HG0632020.2 | 0.8245057 | 1.1398974 | 1.44812562 | cluster 2 |
| HORVU.MOREX.r3.6HG0633510.1 | 0.2162093 | 0.3727915 | 0.58087212 | cluster 2 |
| HORVU.MOREX.r3.7HG0636370.1 | -0.154745 | 0.6267003 | 1.41091824 | cluster 2 |
| HORVU.MOREX.r3.7HG0636630.1 | 0.5256918 | 1.0699841 | 1.23352911 | cluster 2 |
| HORVU.MOREX.r3.7HG0638370.1 | 0.323149 | 0.605877 | 0.72929621 | cluster 2 |
| HORVU.MOREX.r3.7HG0639360.1 | 0.3404557 | 0.6165901 | 0.74913007 | cluster 2 |

**Table S3** Continued.

| ID | logFC_3 h | logFC_6 h | logFC_12 h | cluster No. |
|---|---|---|---|---|
| HORVU.MOREX.r3.7HG0640800.1 | 1.1123416 | 1.6968407 | 2.1802692 | cluster 2 |
| HORVU.MOREX.r3.7HG0641030.1 | 0.6312965 | 1.4336324 | 1.34905455 | cluster 2 |
| HORVU.MOREX.r3.7HG0641750.1 | 2.1002884 | 2.8860427 | 2.97912163 | cluster 2 |
| HORVU.MOREX.r3.7HG0642940.1 | 0.2559375 | 1.9017177 | 1.86045285 | cluster 2 |
| HORVU.MOREX.r3.7HG0644190.1 | 0.2202766 | 0.4281783 | 0.56414754 | cluster 2 |
| HORVU.MOREX.r3.7HG0648620.1 | 0.9283434 | 1.3347288 | 1.42138496 | cluster 2 |
| HORVU.MOREX.r3.7HG0649950.1 | 0.2948762 | 0.6041181 | 0.64772697 | cluster 2 |
| HORVU.MOREX.r3.7HG0650300.1 | 0.2487571 | 0.4879088 | 0.67397048 | cluster 2 |
| HORVU.MOREX.r3.7HG0650530.2 | 1.2490283 | 1.6020453 | 1.66694647 | cluster 2 |
| HORVU.MOREX.r3.7HG0650950.1 | 0.0013464 | 0.2622808 | 0.44287508 | cluster 2 |
| HORVU.MOREX.r3.7HG0653380.1 | 0.9012942 | 1.3602381 | 1.43901056 | cluster 2 |
| HORVU.MOREX.r3.7HG0655100.1 | 0.2582812 | 0.4796561 | 0.52609551 | cluster 2 |
| HORVU.MOREX.r3.7HG0656030.1 | 0.1608324 | 0.4685297 | 0.61465027 | cluster 2 |
| HORVU.MOREX.r3.7HG0657010.1 | 0.2492701 | 0.4034795 | 0.52140551 | cluster 2 |
| HORVU.MOREX.r3.7HG0657330.1 | 0.1585255 | 0.2883706 | 0.42685057 | cluster 2 |
| HORVU.MOREX.r3.7HG0659040.1 | 0.2778608 | 0.8372696 | 0.96093387 | cluster 2 |
| HORVU.MOREX.r3.7HG0661470.1 | 0.3378853 | 0.649975 | 0.71511032 | cluster 2 |
| HORVU.MOREX.r3.7HG0661830.1 | 1.1463543 | 2.311953 | 2.19335103 | cluster 2 |
| HORVU.MOREX.r3.7HG0664720.1 | 0.6858335 | 1.1720211 | 1.14764425 | cluster 2 |
| HORVU.MOREX.r3.7HG0665990.1 | 0.1996344 | 0.5791237 | 0.62468426 | cluster 2 |
| HORVU.MOREX.r3.7HG0667540.1 | 0.2050488 | 0.4309773 | 0.62799912 | cluster 2 |
| HORVU.MOREX.r3.7HG0667610.1 | 1.1030737 | 1.5331211 | 1.52933442 | cluster 2 |
| HORVU.MOREX.r3.7HG0667620.1 | 0.893003 | 1.390352 | 1.51892579 | cluster 2 |
| HORVU.MOREX.r3.7HG0667630.1 | 0.7991252 | 1.0749149 | 1.18015378 | cluster 2 |
| HORVU.MOREX.r3.7HG0667660.1 | 0.4127201 | 0.6768874 | 0.93421058 | cluster 2 |
| HORVU.MOREX.r3.7HG0667680.1 | 0.4618684 | 0.7090718 | 1.00940936 | cluster 2 |
| HORVU.MOREX.r3.7HG0667780.1 | 0.1856161 | 0.5576689 | 0.58250129 | cluster 2 |
| HORVU.MOREX.r3.7HG0668440.1 | 0.5007339 | 0.5590214 | 0.58100318 | cluster 2 |
| HORVU.MOREX.r3.7HG0669590.1 | 0.4147112 | 1.4633242 | 1.38950404 | cluster 2 |
| HORVU.MOREX.r3.7HG0670690.1 | 0.4063011 | 0.6171257 | 0.62126112 | cluster 2 |
| HORVU.MOREX.r3.7HG0672960.1 | 0.7646285 | 1.1752983 | 1.53415731 | cluster 2 |
| HORVU.MOREX.r3.7HG0673710.1 | 0.5399536 | 1.2426141 | 1.43695975 | cluster 2 |
| HORVU.MOREX.r3.7HG0674600.1 | 0.3618624 | 0.6940338 | 1.11592901 | cluster 2 |
| HORVU.MOREX.r3.7HG0674720.1 | 1.2608873 | 2.2130867 | 2.11722617 | cluster 2 |
| HORVU.MOREX.r3.7HG0676690.1 | 0.4287134 | 0.8020907 | 0.81600374 | cluster 2 |
| HORVU.MOREX.r3.7HG0677770.1 | 0.1595816 | 0.51013 | 0.49546538 | cluster 2 |
| HORVU.MOREX.r3.7HG0679050.1 | 0.2594055 | 0.5656885 | 0.7228548 | cluster 2 |
| HORVU.MOREX.r3.7HG0679060.1 | -0.009673 | 0.3316562 | 0.79732488 | cluster 2 |
| HORVU.MOREX.r3.7HG0681050.1 | 0.1644495 | 0.2917135 | 0.4296793 | cluster 2 |
| HORVU.MOREX.r3.7HG0683940.1 | 0.5078305 | 0.7645397 | 0.89184404 | cluster 2 |
| HORVU.MOREX.r3.7HG0684180.1 | 0.288567 | 0.6461458 | 0.61388343 | cluster 2 |
| HORVU.MOREX.r3.7HG0684880.1 | 0.3850775 | 0.7208246 | 0.79923951 | cluster 2 |
| HORVU.MOREX.r3.7HG0685360.1 | 0.2818553 | 0.4289727 | 0.60499427 | cluster 2 |
| HORVU.MOREX.r3.7HG0685920.1 | 0.1563281 | 0.4299148 | 0.5382098 | cluster 2 |
| HORVU.MOREX.r3.7HG0685970.1 | 0.2689053 | 0.4909085 | 0.76916821 | cluster 2 |
| HORVU.MOREX.r3.7HG0688930.1 | 0.3464928 | 0.8284437 | 0.80884317 | cluster 2 |
| HORVU.MOREX.r3.7HG0690090.1 | 0.293307 | 0.7387097 | 0.8379731 | cluster 2 |

**Table S3** Continued.

| ID | logFC_3 h | logFC_6 h | logFC_12 h | cluster No. |
|---|---|---|---|---|
| HORVU.MOREX.r3.7HG0699010.1 | 0.5478962 | 1.3781398 | 2.21993619 | cluster 2 |
| HORVU.MOREX.r3.7HG0699020.1 | 0.9413463 | 1.4926843 | 1.85111967 | cluster 2 |
| HORVU.MOREX.r3.7HG0702070.1 | 0.1648653 | 0.8832185 | 1.29303519 | cluster 2 |
| HORVU.MOREX.r3.7HG0704450.1 | 0.2471358 | 0.5251608 | 0.6036671 | cluster 2 |
| HORVU.MOREX.r3.7HG0705500.1 | -0.353763 | 1.0926535 | 1.09689662 | cluster 2 |
| HORVU.MOREX.r3.7HG0705860.1 | 0.0512008 | 0.3069656 | 0.47617723 | cluster 2 |
| HORVU.MOREX.r3.7HG0706310.1 | 0.3041754 | 0.4370852 | 0.48102721 | cluster 2 |
| HORVU.MOREX.r3.7HG0707160.1 | 1.0650226 | 1.6882276 | 1.66223207 | cluster 2 |
| HORVU.MOREX.r3.7HG0707220.1 | 0.1437703 | 0.6143529 | 0.62104562 | cluster 2 |
| HORVU.MOREX.r3.7HG0707700.1 | 0.4027753 | 1.2813403 | 1.299693 | cluster 2 |
| HORVU.MOREX.r3.7HG0709270.1 | 0.6162508 | 0.8876136 | 1.01067133 | cluster 2 |
| HORVU.MOREX.r3.7HG0711270.1 | 0.3761801 | 0.7568498 | 0.80735751 | cluster 2 |
| HORVU.MOREX.r3.7HG0711890.1 | 0.4657853 | 1.0349723 | 1.31182187 | cluster 2 |
| HORVU.MOREX.r3.7HG0715170.1 | 1.3493185 | 1.5050292 | 1.53037937 | cluster 2 |
| HORVU.MOREX.r3.7HG0717610.1 | 0.2193268 | 0.5897254 | 0.7718575 | cluster 2 |
| HORVU.MOREX.r3.7HG0718170.1 | 0.4822731 | 0.8101556 | 1.02239264 | cluster 2 |
| HORVU.MOREX.r3.7HG0718190.1 | 0.3629962 | 0.6908468 | 0.87321273 | cluster 2 |
| HORVU.MOREX.r3.7HG0718200.1 | 0.2861182 | 0.6426695 | 0.85426654 | cluster 2 |
| HORVU.MOREX.r3.7HG0718540.1 | 0.1767034 | 0.6245295 | 0.78858041 | cluster 2 |
| HORVU.MOREX.r3.7HG0718720.1 | 0.3023671 | 0.9275159 | 1.04637524 | cluster 2 |
| HORVU.MOREX.r3.7HG0718830.1 | 0.5322107 | 0.7212304 | 0.84611557 | cluster 2 |
| HORVU.MOREX.r3.7HG0718930.2 | 0.876005 | 1.3337527 | 1.41420998 | cluster 2 |
| HORVU.MOREX.r3.7HG0719170.1 | 0.1369497 | 0.5489585 | 0.6993676 | cluster 2 |
| HORVU.MOREX.r3.7HG0720100.1 | 0.26822 | 0.7850186 | 0.76892662 | cluster 2 |
| HORVU.MOREX.r3.7HG0721690.1 | 0.5945101 | 1.2770139 | 1.25755181 | cluster 2 |
| HORVU.MOREX.r3.7HG0722360.1 | 0.8556942 | 1.6579914 | 1.63620645 | cluster 2 |
| HORVU.MOREX.r3.7HG0722690.1 | 0.1149204 | 0.3309091 | 0.50938211 | cluster 2 |
| HORVU.MOREX.r3.7HG0723730.1 | 0.3540922 | 0.6391943 | 0.90352887 | cluster 2 |
| HORVU.MOREX.r3.7HG0724060.1 | 0.6458837 | 1.4720508 | 1.59710113 | cluster 2 |
| HORVU.MOREX.r3.7HG0724140.1 | 0.5940254 | 1.0527349 | 1.0710578 | cluster 2 |
| HORVU.MOREX.r3.7HG0725450.1 | 0.2488084 | 1.1230295 | 1.58046086 | cluster 2 |
| HORVU.MOREX.r3.7HG0726770.1 | 0.6632144 | 0.9362614 | 1.19564074 | cluster 2 |
| HORVU.MOREX.r3.7HG0726800.1 | 0.5385227 | 0.9064497 | 1.13339153 | cluster 2 |
| HORVU.MOREX.r3.7HG0727290.1 | 0.2858006 | 0.5017598 | 0.69328248 | cluster 2 |
| HORVU.MOREX.r3.7HG0727750.1 | 0.4565523 | 1.2713684 | 1.32364467 | cluster 2 |
| HORVU.MOREX.r3.7HG0727980.1 | 0.1937819 | 0.3807121 | 0.59720063 | cluster 2 |
| HORVU.MOREX.r3.7HG0728060.1 | 0.1216479 | 0.4787506 | 0.48325967 | cluster 2 |
| HORVU.MOREX.r3.7HG0730510.1 | 0.8509991 | 1.3663026 | 1.49259686 | cluster 2 |
| HORVU.MOREX.r3.7HG0731950.1 | 0.2889116 | 0.6582949 | 0.78541346 | cluster 2 |
| HORVU.MOREX.r3.7HG0736150.1 | 0.6526935 | 1.2176069 | 1.32390434 | cluster 2 |
| HORVU.MOREX.r3.7HG0736870.1 | 0.0994965 | 0.251 | 0.43898021 | cluster 2 |
| HORVU.MOREX.r3.7HG0737470.1 | 0.657427 | 1.0278927 | 1.04021175 | cluster 2 |
| HORVU.MOREX.r3.7HG0737890.1 | 0.4605395 | 1.0125301 | 1.13516466 | cluster 2 |
| HORVU.MOREX.r3.7HG0738680.1 | 0.4751268 | 0.700665 | 0.72210838 | cluster 2 |
| HORVU.MOREX.r3.7HG0738770.1 | 0.6228843 | 0.9265767 | 1.21144757 | cluster 2 |
| HORVU.MOREX.r3.7HG0739030.1 | 0.1261805 | 0.4185308 | 0.54952571 | cluster 2 |
| HORVU.MOREX.r3.7HG0739120.1 | 0.6367749 | 1.0566231 | 1.10893669 | cluster 2 |

**Table S3** Continued.

| ID | logFC_3 h | logFC_6 h | logFC_12 h | cluster No. |
|---|---|---|---|---|
| HORVU.MOREX.r3.7HG0742080.1 | 0.8008977 | 1.3684256 | 1.57602568 | cluster 2 |
| HORVU.MOREX.r3.7HG0743900.1 | 0.0221244 | 0.2491488 | 0.51036886 | cluster 2 |
| HORVU.MOREX.r3.7HG0744890.1 | 0.7665333 | 1.0109883 | 1.09590618 | cluster 2 |
| HORVU.MOREX.r3.7HG0747690.1 | 3.0485897 | 3.5558559 | 3.5684823 | cluster 2 |
| HORVU.MOREX.r3.7HG0747940.1 | 0.2078592 | 0.444076 | 0.45911291 | cluster 2 |
| HORVU.MOREX.r3.7HG0748070.1 | 0.8146708 | 1.352132 | 1.31915482 | cluster 2 |
| HORVU.MOREX.r3.7HG0749070.1 | 0.3711935 | 0.9861429 | 1.22317384 | cluster 2 |
| HORVU.MOREX.r3.7HG0749080.1 | 0.0282132 | 1.0131946 | 1.12856448 | cluster 2 |
| HORVU.MOREX.r3.7HG0749090.1 | 0.2502276 | 1.0832272 | 1.18146177 | cluster 2 |
| HORVU.MOREX.r3.7HG0749100.1 | 0.2043081 | 0.8330815 | 0.92186048 | cluster 2 |
| HORVU.MOREX.r3.7HG0749640.1 | 0.0602674 | 0.4354517 | 0.61300476 | cluster 2 |
| HORVU.MOREX.r3.7HG0749870.1 | 0.2883228 | 0.4127493 | 0.54530599 | cluster 2 |
| HORVU.MOREX.r3.7HG0750140.1 | 0.1655989 | 0.3765363 | 0.37653161 | cluster 2 |
| HORVU.MOREX.r3.7HG0750170.1 | 0.8117068 | 1.2800809 | 1.56679351 | cluster 2 |
| HORVU.MOREX.r3.7HG0751300.1 | 0.6516701 | 1.4318477 | 1.47771629 | cluster 2 |
| HORVU.MOREX.r3.7HG0751340.1 | 0.5787381 | 1.0448161 | 1.0172609 | cluster 2 |
| HORVU.MOREX.r3.7HG0752370.1 | 0.2443724 | 0.6179565 | 0.76548163 | cluster 2 |
| HORVU.MOREX.r3.1HG0002630.1 | 0.1711594 | 1.9896814 | 0.46660601 | cluster 3 |
| HORVU.MOREX.r3.1HG0003620.1 | 0.3138326 | 0.5928033 | 0.5450898 | cluster 3 |
| HORVU.MOREX.r3.1HG0005700.1 | 1.0631149 | 2.2824019 | 1.94990724 | cluster 3 |
| HORVU.MOREX.r3.1HG0005730.1 | 0.7849748 | 2.1219919 | 1.67003551 | cluster 3 |
| HORVU.MOREX.r3.1HG0010340.1 | 1.5592583 | 2.4536826 | 2.20032447 | cluster 3 |
| HORVU.MOREX.r3.1HG0012340.1 | 0.6722211 | 1.3412727 | 1.25333711 | cluster 3 |
| HORVU.MOREX.r3.1HG0017280.1 | 0.8374164 | 1.7056726 | 1.59651037 | cluster 3 |
| HORVU.MOREX.r3.1HG0017770.1 | 0.4544104 | 0.7517881 | 0.63944908 | cluster 3 |
| HORVU.MOREX.r3.1HG0024510.1 | 0.2267448 | 0.4286452 | 0.40608049 | cluster 3 |
| HORVU.MOREX.r3.1HG0025320.1 | 0.613492 | 1.2169377 | 1.07232972 | cluster 3 |
| HORVU.MOREX.r3.1HG0027200.1 | 0.7464108 | 1.1617697 | 0.81180175 | cluster 3 |
| HORVU.MOREX.r3.1HG0042290.1 | 0.0574144 | 0.4066945 | 0.34009793 | cluster 3 |
| HORVU.MOREX.r3.1HG0044290.1 | 0.2085412 | 0.8069825 | 0.60706767 | cluster 3 |
| HORVU.MOREX.r3.1HG0044800.1 | 1.2830162 | 2.1568566 | 1.80537799 | cluster 3 |
| HORVU.MOREX.r3.1HG0046210.1 | 1.7563681 | 2.432764 | 2.2640672 | cluster 3 |
| HORVU.MOREX.r3.1HG0047570.1 | 0.1905652 | 0.7051528 | 0.5919737 | cluster 3 |
| HORVU.MOREX.r3.1HG0050220.1 | 0.9252035 | 1.467589 | 1.40353289 | cluster 3 |
| HORVU.MOREX.r3.1HG0053060.1 | 0.6190743 | 1.0949658 | 1.04145333 | cluster 3 |
| HORVU.MOREX.r3.1HG0054170.1 | 0.9433977 | 1.926734 | 1.56146538 | cluster 3 |
| HORVU.MOREX.r3.1HG0054180.1 | 0.8191721 | 1.7437234 | 1.53861711 | cluster 3 |
| HORVU.MOREX.r3.1HG0054190.1 | 1.21786 | 2.2431323 | 1.88452959 | cluster 3 |
| HORVU.MOREX.r3.1HG0055500.1 | 0.6284683 | 1.2057958 | 1.01315561 | cluster 3 |
| HORVU.MOREX.r3.1HG0056620.1 | 1.0613668 | 1.936995 | 1.82998389 | cluster 3 |
| HORVU.MOREX.r3.1HG0062630.1 | 0.2563032 | 0.5211878 | 0.46617722 | cluster 3 |
| HORVU.MOREX.r3.1HG0066800.1 | 0.7309063 | 1.4529727 | 1.17637157 | cluster 3 |
| HORVU.MOREX.r3.1HG0069030.1 | 1.2248662 | 1.6209528 | 1.51342566 | cluster 3 |
| HORVU.MOREX.r3.1HG0069410.1 | 0.6270373 | 1.1136511 | 0.92525564 | cluster 3 |
| HORVU.MOREX.r3.1HG0071420.1 | 0.659344 | 0.9175095 | 0.71187787 | cluster 3 |
| HORVU.MOREX.r3.1HG0073430.1 | 0.1941013 | 0.4226951 | 0.33224878 | cluster 3 |
| HORVU.MOREX.r3.1HG0075260.1 | 2.5500555 | 3.5917174 | 3.10578727 | cluster 3 |

**Table S3** Continued.

| ID | logFC_3 h | logFC_6 h | logFC_12 h | cluster No. |
|---|---|---|---|---|
| HORVU.MOREX.r3.1HG0075340.1 | 0.3320698 | 0.9745518 | 0.77615585 | cluster 3 |
| HORVU.MOREX.r3.1HG0075730.1 | 0.635621 | 1.1428067 | 1.0477762 | cluster 3 |
| HORVU.MOREX.r3.1HG0077960.1 | 0.2938817 | 0.7137327 | 0.5300652 | cluster 3 |
| HORVU.MOREX.r3.1HG0080180.1 | 1.9535087 | 2.3634378 | 2.19937724 | cluster 3 |
| HORVU.MOREX.r3.1HG0081950.1 | 2.2709035 | 3.02783 | 2.57640243 | cluster 3 |
| HORVU.MOREX.r3.1HG0086370.1 | 1.1999165 | 2.4399783 | 2.10522523 | cluster 3 |
| HORVU.MOREX.r3.1HG0088530.1 | 0.3068814 | 0.9592564 | 0.78364279 | cluster 3 |
| HORVU.MOREX.r3.1HG0092200.1 | 0.3910832 | 1.3644708 | 0.56997417 | cluster 3 |
| HORVU.MOREX.r3.1HG0092860.1 | 0.3222543 | 1.3788585 | 1.2111266 | cluster 3 |
| HORVU.MOREX.r3.1HG0093050.1 | 0.5405552 | 1.0930189 | 0.81022969 | cluster 3 |
| HORVU.MOREX.r3.1HG0094110.1 | 1.911838 | 2.6213036 | 2.00917259 | cluster 3 |
| HORVU.MOREX.r3.2HG0096230.1 | 0.5403085 | 1.3345003 | 0.81438353 | cluster 3 |
| HORVU.MOREX.r3.2HG0096520.1 | 0.3392425 | 1.5790662 | 1.4271017 | cluster 3 |
| HORVU.MOREX.r3.2HG0096800.1 | 0.380927 | 0.7518304 | 0.69055058 | cluster 3 |
| HORVU.MOREX.r3.2HG0105830.1 | 0.3618823 | 0.7921987 | 0.68576609 | cluster 3 |
| HORVU.MOREX.r3.2HG0112190.1 | 1.0595431 | 1.7810511 | 1.55388523 | cluster 3 |
| HORVU.MOREX.r3.2HG0112690.1 | 1.1115037 | 1.3132885 | 1.26073711 | cluster 3 |
| HORVU.MOREX.r3.2HG0113720.1 | 0.7358426 | 1.3766037 | 1.16483125 | cluster 3 |
| HORVU.MOREX.r3.2HG0117830.1 | 1.0322649 | 1.8281935 | 1.39659553 | cluster 3 |
| HORVU.MOREX.r3.2HG0118220.1 | 0.6849396 | 1.2005412 | 1.08306497 | cluster 3 |
| HORVU.MOREX.r3.2HG0119080.1 | 0.2754023 | 0.8202671 | 0.60203402 | cluster 3 |
| HORVU.MOREX.r3.2HG0120470.1 | 0.6878973 | 1.3167354 | 1.17548026 | cluster 3 |
| HORVU.MOREX.r3.2HG0120490.1 | 0.6908535 | 1.3151249 | 1.17123373 | cluster 3 |
| HORVU.MOREX.r3.2HG0120560.1 | 0.7204918 | 1.3190281 | 1.161665 | cluster 3 |
| HORVU.MOREX.r3.2HG0121410.1 | 0.5299154 | 0.9256832 | 0.701258 | cluster 3 |
| HORVU.MOREX.r3.2HG0121630.1 | 0.3627151 | 0.7387854 | 0.55379348 | cluster 3 |
| HORVU.MOREX.r3.2HG0121670.1 | 0.4341492 | 0.7372941 | 0.50348341 | cluster 3 |
| HORVU.MOREX.r3.2HG0122880.1 | 0.5541906 | 1.4521582 | 1.32916844 | cluster 3 |
| HORVU.MOREX.r3.2HG0124550.1 | 1.0530838 | 1.3884352 | 1.29305111 | cluster 3 |
| HORVU.MOREX.r3.2HG0124850.1 | 1.1344358 | 1.7515246 | 1.65931916 | cluster 3 |
| HORVU.MOREX.r3.2HG0127260.1 | 0.8566943 | 1.4226129 | 1.31323357 | cluster 3 |
| HORVU.MOREX.r3.2HG0139810.1 | 0.6391969 | 1.0241819 | 0.92747759 | cluster 3 |
| HORVU.MOREX.r3.2HG0144760.1 | 0.4923039 | 1.2240785 | 1.10775071 | cluster 3 |
| HORVU.MOREX.r3.2HG0157410.1 | 0.8008322 | 1.2827141 | 1.21201271 | cluster 3 |
| HORVU.MOREX.r3.2HG0157450.1 | 0.6323934 | 1.0853997 | 1.02574131 | cluster 3 |
| HORVU.MOREX.r3.2HG0160690.1 | 0.4966044 | 1.0381931 | 0.82364778 | cluster 3 |
| HORVU.MOREX.r3.2HG0164020.1 | 0.6279206 | 1.1690404 | 0.75672277 | cluster 3 |
| HORVU.MOREX.r3.2HG0164470.1 | 0.5899381 | 0.8859545 | 0.73787661 | cluster 3 |
| HORVU.MOREX.r3.2HG0170230.1 | 0.4756654 | 0.6205134 | 0.57442443 | cluster 3 |
| HORVU.MOREX.r3.2HG0171780.1 | 1.0559533 | 1.5694528 | 1.16564535 | cluster 3 |
| HORVU.MOREX.r3.2HG0177790.1 | 1.1244072 | 2.1076834 | 1.60997698 | cluster 3 |
| HORVU.MOREX.r3.2HG0178480.1 | 0.5904511 | 1.1582042 | 0.94615385 | cluster 3 |
| HORVU.MOREX.r3.2HG0179320.1 | 0.2457168 | 0.7990075 | 0.58470685 | cluster 3 |
| HORVU.MOREX.r3.2HG0179790.1 | 0.5119616 | 1.0905045 | 0.72312472 | cluster 3 |
| HORVU.MOREX.r3.2HG0180010.1 | 0.8964688 | 1.6113559 | 1.37511721 | cluster 3 |
| HORVU.MOREX.r3.2HG0182650.1 | 0.4272274 | 2.0442696 | 1.12709483 | cluster 3 |
| HORVU.MOREX.r3.2HG0185600.1 | 0.714457 | 1.2198044 | 1.05281393 | cluster 3 |

**Table S3** Continued.

| ID | logFC_3 h | logFC_6 h | logFC_12 h | cluster No. |
|---|---|---|---|---|
| HORVU.MOREX.r3.2HG0186480.1 | 1.1047789 | 2.389649 | 2.16071754 | cluster 3 |
| HORVU.MOREX.r3.2HG0186750.1 | 0.5331418 | 2.9190879 | 0.8443993 | cluster 3 |
| HORVU.MOREX.r3.2HG0187450.1 | 0.7612022 | 1.5025257 | 1.35973473 | cluster 3 |
| HORVU.MOREX.r3.2HG0189010.1 | 0.5339469 | 1.0113907 | 0.94550697 | cluster 3 |
| HORVU.MOREX.r3.2HG0189530.1 | 0.8482455 | 1.5025059 | 1.26936228 | cluster 3 |
| HORVU.MOREX.r3.2HG0189670.1 | 0.9204562 | 1.5866451 | 1.51279104 | cluster 3 |
| HORVU.MOREX.r3.2HG0190300.1 | 0.4676555 | 0.5979554 | 0.52091469 | cluster 3 |
| HORVU.MOREX.r3.2HG0192100.1 | 1.070724 | 1.8982312 | 1.68325945 | cluster 3 |
| HORVU.MOREX.r3.2HG0192410.1 | 0.890758 | 1.9465455 | 1.68137018 | cluster 3 |
| HORVU.MOREX.r3.2HG0193170.1 | 3.6640461 | 5.2023163 | 4.47835727 | cluster 3 |
| HORVU.MOREX.r3.2HG0193490.1 | 0.7376333 | 1.4119295 | 0.96590353 | cluster 3 |
| HORVU.MOREX.r3.2HG0194620.1 | 0.980254 | 2.0662313 | 1.87952744 | cluster 3 |
| HORVU.MOREX.r3.2HG0195690.1 | 0.7582451 | 1.6692054 | 1.07497895 | cluster 3 |
| HORVU.MOREX.r3.2HG0196520.1 | 0.2885222 | 0.736155 | 0.63318261 | cluster 3 |
| HORVU.MOREX.r3.2HG0196800.1 | 0.8724415 | 2.308663 | 2.02439845 | cluster 3 |
| HORVU.MOREX.r3.2HG0197110.1 | 0.2337897 | 0.3555761 | 0.26597198 | cluster 3 |
| HORVU.MOREX.r3.2HG0197230.1 | 0.721065 | 1.2399644 | 1.11077221 | cluster 3 |
| HORVU.MOREX.r3.2HG0197540.1 | 0.2789629 | 1.2485438 | 0.84213138 | cluster 3 |
| HORVU.MOREX.r3.2HG0198320.1 | 0.9876411 | 1.7139737 | 1.55860221 | cluster 3 |
| HORVU.MOREX.r3.2HG0199160.1 | 0.2774651 | 0.5976138 | 0.53367882 | cluster 3 |
| HORVU.MOREX.r3.2HG0202250.1 | 1.0529513 | 1.3503838 | 1.2943979 | cluster 3 |
| HORVU.MOREX.r3.2HG0203070.1 | 0.4929699 | 1.0286154 | 0.87215367 | cluster 3 |
| HORVU.MOREX.r3.2HG0205420.1 | 0.4736651 | 0.8810024 | 0.8222821 | cluster 3 |
| HORVU.MOREX.r3.2HG0205990.1 | 1.6154246 | 2.2776938 | 1.89734079 | cluster 3 |
| HORVU.MOREX.r3.2HG0208520.1 | 1.2428445 | 2.3492031 | 1.90672709 | cluster 3 |
| HORVU.MOREX.r3.2HG0209680.1 | 0.3782002 | 1.5149124 | 1.30868509 | cluster 3 |
| HORVU.MOREX.r3.2HG0210500.1 | 0.7432536 | 1.4401568 | 1.21918749 | cluster 3 |
| HORVU.MOREX.r3.2HG0210510.1 | 0.6554861 | 1.0386631 | 0.96490302 | cluster 3 |
| HORVU.MOREX.r3.2HG0212860.1 | 1.12497 | 1.8819863 | 1.76056432 | cluster 3 |
| HORVU.MOREX.r3.2HG0214070.1 | 1.0739833 | 2.2200397 | 1.61859641 | cluster 3 |
| HORVU.MOREX.r3.2HG0214130.1 | 0.9015364 | 1.4784312 | 1.25669869 | cluster 3 |
| HORVU.MOREX.r3.2HG0217090.1 | 1.3578365 | 2.2543033 | 1.86592829 | cluster 3 |
| HORVU.MOREX.r3.3HG0219380.1 | 0.6293154 | 1.0460411 | 0.94657638 | cluster 3 |
| HORVU.MOREX.r3.3HG0225880.1 | 0.3562515 | 0.9080649 | 0.61147233 | cluster 3 |
| HORVU.MOREX.r3.3HG0228940.1 | 0.8885458 | 1.1608421 | 1.11924368 | cluster 3 |
| HORVU.MOREX.r3.3HG0230640.1 | 1.2626452 | 2.1498645 | 1.84424381 | cluster 3 |
| HORVU.MOREX.r3.3HG0234000.1 | 1.1717256 | 1.9594596 | 1.54213879 | cluster 3 |
| HORVU.MOREX.r3.3HG0234010.1 | 1.0424388 | 1.626236 | 1.46465529 | cluster 3 |
| HORVU.MOREX.r3.3HG0234030.1 | 0.3967579 | 0.9326129 | 0.83863909 | cluster 3 |
| HORVU.MOREX.r3.3HG0234970.1 | 0.2472746 | 0.6808934 | 0.51916076 | cluster 3 |
| HORVU.MOREX.r3.3HG0235320.1 | 0.8330944 | 2.090663 | 1.5069093 | cluster 3 |
| HORVU.MOREX.r3.3HG0235350.1 | 1.2096463 | 2.5296998 | 2.00644621 | cluster 3 |
| HORVU.MOREX.r3.3HG0239900.1 | 0.5999186 | 1.3665604 | 0.7924196 | cluster 3 |
| HORVU.MOREX.r3.3HG0239980.1 | 0.2951308 | 1.0442109 | 0.86909149 | cluster 3 |
| HORVU.MOREX.r3.3HG0240350.1 | 0.3704946 | 0.6667146 | 0.56487049 | cluster 3 |
| HORVU.MOREX.r3.3HG0242770.1 | 0.22128 | 0.6537208 | 0.45196031 | cluster 3 |
| HORVU.MOREX.r3.3HG0244000.1 | 1.062452 | 2.0120722 | 1.68368127 | cluster 3 |

**Table S3** Continued.

| ID | logFC_3 h | logFC_6 h | logFC_12 h | cluster No. |
|---|---|---|---|---|
| HORVU.MOREX.r3.3HG0244040.1 | 0.4836287 | 1.2285902 | 0.893857 | cluster 3 |
| HORVU.MOREX.r3.3HG0245120.1 | 2.3474942 | 3.008632 | 2.75105463 | cluster 3 |
| HORVU.MOREX.r3.3HG0245250.1 | 2.6149236 | 4.0011948 | 3.23905423 | cluster 3 |
| HORVU.MOREX.r3.3HG0246300.1 | 0.1292844 | 0.4852659 | 0.34127403 | cluster 3 |
| HORVU.MOREX.r3.3HG0250060.1 | 1.0479656 | 2.5488608 | 1.99188759 | cluster 3 |
| HORVU.MOREX.r3.3HG0252960.1 | 1.1087823 | 1.5277099 | 1.29909636 | cluster 3 |
| HORVU.MOREX.r3.3HG0253860.1 | 1.2048684 | 2.2359774 | 1.40856828 | cluster 3 |
| HORVU.MOREX.r3.3HG0258240.1 | 0.4599494 | 1.3517616 | 0.93703102 | cluster 3 |
| HORVU.MOREX.r3.3HG0264640.1 | -0.09276 | 1.4500437 | 0.83294059 | cluster 3 |
| HORVU.MOREX.r3.3HG0265140.1 | 1.5471385 | 2.4299566 | 1.99162846 | cluster 3 |
| HORVU.MOREX.r3.3HG0269130.1 | 0.6192941 | 1.2298994 | 0.98526732 | cluster 3 |
| HORVU.MOREX.r3.3HG0275090.1 | 1.7547882 | 2.2925491 | 1.9218451 | cluster 3 |
| HORVU.MOREX.r3.3HG0275490.1 | 1.0334564 | 1.9055257 | 1.70784416 | cluster 3 |
| HORVU.MOREX.r3.3HG0276120.1 | 0.8867794 | 1.93491 | 1.77700969 | cluster 3 |
| HORVU.MOREX.r3.3HG0278630.1 | 0.2406564 | 0.5193078 | 0.45988332 | cluster 3 |
| HORVU.MOREX.r3.3HG0279100.1 | 0.9252075 | 1.1891752 | 1.01137387 | cluster 3 |
| HORVU.MOREX.r3.3HG0285150.1 | 0.5458466 | 0.8589187 | 0.81773433 | cluster 3 |
| HORVU.MOREX.r3.3HG0285440.1 | 0.9720005 | 1.7141319 | 1.49152921 | cluster 3 |
| HORVU.MOREX.r3.3HG0286100.1 | 1.3374496 | 2.0602449 | 1.54984812 | cluster 3 |
| HORVU.MOREX.r3.3HG0290300.1 | 1.1074348 | 2.9956127 | 1.73496857 | cluster 3 |
| HORVU.MOREX.r3.3HG0292490.1 | 0.4252445 | 0.897386 | 0.83439692 | cluster 3 |
| HORVU.MOREX.r3.3HG0292680.1 | 0.3480327 | 0.6873488 | 0.62085004 | cluster 3 |
| HORVU.MOREX.r3.3HG0294390.1 | 0.2485188 | 0.7169626 | 0.30449036 | cluster 3 |
| HORVU.MOREX.r3.3HG0294960.1 | 0.8456551 | 1.7854641 | 1.55170444 | cluster 3 |
| HORVU.MOREX.r3.3HG0295640.1 | 0.5578424 | 1.319709 | 1.21356137 | cluster 3 |
| HORVU.MOREX.r3.3HG0296120.1 | 0.6779504 | 1.531733 | 1.11603476 | cluster 3 |
| HORVU.MOREX.r3.3HG0298080.1 | 0.4345492 | 1.2062768 | 1.08617308 | cluster 3 |
| HORVU.MOREX.r3.3HG0298180.1 | 0.8187417 | 1.7149992 | 1.50350633 | cluster 3 |
| HORVU.MOREX.r3.3HG0298510.1 | 0.329936 | 0.7106195 | 0.5726424 | cluster 3 |
| HORVU.MOREX.r3.3HG0298790.1 | 0.5148781 | 0.8288828 | 0.63477045 | cluster 3 |
| HORVU.MOREX.r3.3HG0302230.1 | 0.3024303 | 0.6112375 | 0.43190034 | cluster 3 |
| HORVU.MOREX.r3.3HG0303240.1 | 0.9116472 | 1.2660741 | 1.11420955 | cluster 3 |
| HORVU.MOREX.r3.3HG0305440.1 | 0.8643178 | 1.5666716 | 0.97526344 | cluster 3 |
| HORVU.MOREX.r3.3HG0307240.1 | 2.6195977 | 3.0473965 | 2.70112743 | cluster 3 |
| HORVU.MOREX.r3.3HG0307250.1 | 1.5947862 | 2.116851 | 1.7424705 | cluster 3 |
| HORVU.MOREX.r3.3HG0309760.1 | 0.540732 | 1.1335568 | 0.81508027 | cluster 3 |
| HORVU.MOREX.r3.3HG0309780.1 | 2.2034485 | 3.0540032 | 2.84011543 | cluster 3 |
| HORVU.MOREX.r3.3HG0310110.1 | 1.6053668 | 1.9320626 | 1.75034226 | cluster 3 |
| HORVU.MOREX.r3.3HG0310390.1 | -0.224291 | 1.633224 | 0.66658981 | cluster 3 |
| HORVU.MOREX.r3.3HG0313090.1 | 0.495117 | 0.8080619 | 0.73737565 | cluster 3 |
| HORVU.MOREX.r3.3HG0313320.1 | 0.383243 | 1.5017975 | 1.33907217 | cluster 3 |
| HORVU.MOREX.r3.3HG0317780.1 | 0.836648 | 1.3484772 | 1.21671848 | cluster 3 |
| HORVU.MOREX.r3.3HG0318590.1 | 1.9662537 | 2.8236098 | 2.63609669 | cluster 3 |
| HORVU.MOREX.r3.3HG0319430.1 | 0.5926847 | 0.9180671 | 0.82386622 | cluster 3 |
| HORVU.MOREX.r3.3HG0321700.1 | 0.4348523 | 0.8269235 | 0.68354288 | cluster 3 |
| HORVU.MOREX.r3.3HG0322650.1 | 0.494499 | 0.8209263 | 0.70024765 | cluster 3 |
| HORVU.MOREX.r3.3HG0323750.1 | 0.7604213 | 0.9168295 | 0.86647867 | cluster 3 |

**Table S3** Continued.

| ID | logFC_3 h | logFC_6 h | logFC_12 h | cluster No. |
|---|---|---|---|---|
| HORVU.MOREX.r3.3HG0325250.1 | 2.8303653 | 3.8272581 | 3.37321448 | cluster 3 |
| HORVU.MOREX.r3.3HG0328430.1 | 0.5000046 | 1.7154025 | 0.87623675 | cluster 3 |
| HORVU.MOREX.r3.3HG0329950.1 | 1.0426997 | 2.0173073 | 1.63146116 | cluster 3 |
| HORVU.MOREX.r3.3HG0330200.1 | 0.7872128 | 1.317891 | 1.23348626 | cluster 3 |
| HORVU.MOREX.r3.3HG0330220.1 | 0.3105887 | 1.170575 | 1.03358316 | cluster 3 |
| HORVU.MOREX.r3.4HG0331440.1 | 1.1050685 | 1.5029967 | 1.22483615 | cluster 3 |
| HORVU.MOREX.r3.4HG0335170.1 | 0.6011743 | 1.3636845 | 1.25029495 | cluster 3 |
| HORVU.MOREX.r3.4HG0337950.1 | 1.6835407 | 2.5528601 | 2.27156529 | cluster 3 |
| HORVU.MOREX.r3.4HG0340030.1 | 0.6666234 | 1.9054957 | 1.39495362 | cluster 3 |
| HORVU.MOREX.r3.4HG0342480.1 | 0.7205496 | 0.9991801 | 0.89056024 | cluster 3 |
| HORVU.MOREX.r3.4HG0342680.1 | 0.6153629 | 1.2654457 | 0.94795787 | cluster 3 |
| HORVU.MOREX.r3.4HG0344550.1 | 0.8312747 | 1.6923254 | 1.06466948 | cluster 3 |
| HORVU.MOREX.r3.4HG0345580.1 | 1.4882406 | 2.4305487 | 1.91477372 | cluster 3 |
| HORVU.MOREX.r3.4HG0348800.1 | 0.3939965 | 1.2995825 | 1.13020082 | cluster 3 |
| HORVU.MOREX.r3.4HG0351290.1 | 0.3928932 | 1.0509139 | 0.47622544 | cluster 3 |
| HORVU.MOREX.r3.4HG0351750.1 | 1.470455 | 2.3362083 | 2.02113489 | cluster 3 |
| HORVU.MOREX.r3.4HG0353120.1 | 0.1511889 | 1.4223427 | 1.1709541 | cluster 3 |
| HORVU.MOREX.r3.4HG0355210.1 | 0.916253 | 2.0940685 | 1.60108941 | cluster 3 |
| HORVU.MOREX.r3.4HG0381150.1 | 0.7934923 | 2.0748825 | 1.85121144 | cluster 3 |
| HORVU.MOREX.r3.4HG0382620.1 | 0.2655961 | 0.6455969 | 0.48102118 | cluster 3 |
| HORVU.MOREX.r3.4HG0383530.1 | 0.6392348 | 1.160899 | 1.09093805 | cluster 3 |
| HORVU.MOREX.r3.4HG0384620.1 | 0.3592195 | 1.0461923 | 0.92359023 | cluster 3 |
| HORVU.MOREX.r3.4HG0386490.1 | 1.1026408 | 2.0003374 | 1.7867265 | cluster 3 |
| HORVU.MOREX.r3.4HG0389250.1 | 0.5581347 | 1.3098591 | 0.87133647 | cluster 3 |
| HORVU.MOREX.r3.4HG0390770.1 | 0.7110921 | 1.1278313 | 1.0822598 | cluster 3 |
| HORVU.MOREX.r3.4HG0394330.1 | 0.7667639 | 1.2983732 | 1.22561783 | cluster 3 |
| HORVU.MOREX.r3.4HG0394460.1 | 0.2243026 | 1.5822182 | 1.08193766 | cluster 3 |
| HORVU.MOREX.r3.4HG0395540.1 | 1.7548703 | 2.0826682 | 1.8518538 | cluster 3 |
| HORVU.MOREX.r3.4HG0399970.1 | 0.8449133 | 1.1084163 | 1.05951791 | cluster 3 |
| HORVU.MOREX.r3.4HG0400520.1 | 0.9972842 | 1.3617709 | 1.26348373 | cluster 3 |
| HORVU.MOREX.r3.4HG0400530.1 | 0.9162412 | 1.3692997 | 1.26424739 | cluster 3 |
| HORVU.MOREX.r3.4HG0403620.1 | 0.8105203 | 1.6808756 | 1.53698126 | cluster 3 |
| HORVU.MOREX.r3.4HG0405860.1 | 0.7839534 | 2.4328059 | 2.05054853 | cluster 3 |
| HORVU.MOREX.r3.4HG0406670.1 | 0.4756937 | 0.8967279 | 0.81404402 | cluster 3 |
| HORVU.MOREX.r3.4HG0407230.1 | 0.6594467 | 1.4212745 | 1.02386225 | cluster 3 |
| HORVU.MOREX.r3.4HG0407310.1 | 0.6173735 | 0.9311402 | 0.72037621 | cluster 3 |
| HORVU.MOREX.r3.4HG0408270.1 | 1.5757383 | 2.186194 | 1.82641241 | cluster 3 |
| HORVU.MOREX.r3.4HG0409230.1 | 1.1281871 | 2.2149577 | 1.85850757 | cluster 3 |
| HORVU.MOREX.r3.4HG0409880.1 | 1.1842185 | 2.0855537 | 1.419768 | cluster 3 |
| HORVU.MOREX.r3.4HG0410830.1 | 0.4722377 | 1.2267372 | 1.08934367 | cluster 3 |
| HORVU.MOREX.r3.4HG0410910.1 | 0.6882354 | 2.2746247 | 0.9723833 | cluster 3 |
| HORVU.MOREX.r3.4HG0412140.1 | 0.7597879 | 1.2239557 | 1.10254828 | cluster 3 |
| HORVU.MOREX.r3.4HG0412370.1 | 1.289062 | 2.2483469 | 2.02792442 | cluster 3 |
| HORVU.MOREX.r3.4HG0413320.1 | 1.6092173 | 2.408495 | 1.85975722 | cluster 3 |
| HORVU.MOREX.r3.4HG0413740.1 | 1.3826066 | 2.3358491 | 2.06676432 | cluster 3 |
| HORVU.MOREX.r3.4HG0415330.1 | 1.7208855 | 2.2981548 | 2.20676924 | cluster 3 |
| HORVU.MOREX.r3.4HG0415490.1 | 1.3712582 | 2.2799411 | 1.91661429 | cluster 3 |

**Table S3** Continued.

| ID | logFC_3 h | logFC_6 h | logFC_12 h | cluster No. |
|---|---|---|---|---|
| HORVU.MOREX.r3.4HG0417710.1 | 0.4850072 | 0.9575886 | 0.8932471 | cluster 3 |
| HORVU.MOREX.r3.5HG0420480.1 | 1.5142313 | 2.1727482 | 1.97952195 | cluster 3 |
| HORVU.MOREX.r3.5HG0420820.1 | 0.2361459 | 1.512927 | 1.18910008 | cluster 3 |
| HORVU.MOREX.r3.5HG0423820.1 | 0.6251739 | 0.9525042 | 0.87832373 | cluster 3 |
| HORVU.MOREX.r3.5HG0429180.1 | 1.1578838 | 1.5883447 | 1.51847088 | cluster 3 |
| HORVU.MOREX.r3.5HG0429900.1 | 0.8309484 | 1.5690068 | 1.29612079 | cluster 3 |
| HORVU.MOREX.r3.5HG0430050.1 | 0.3381948 | 0.6380405 | 0.55480978 | cluster 3 |
| HORVU.MOREX.r3.5HG0438460.1 | 1.0108142 | 1.6370994 | 1.2713 | cluster 3 |
| HORVU.MOREX.r3.5HG0458890.1 | 0.2821128 | 0.6130923 | 0.55484359 | cluster 3 |
| HORVU.MOREX.r3.5HG0462200.1 | 0.8685806 | 1.6681302 | 1.55919222 | cluster 3 |
| HORVU.MOREX.r3.5HG0467870.1 | 0.4080381 | 1.2350319 | 0.99358787 | cluster 3 |
| HORVU.MOREX.r3.5HG0468580.1 | 0.6477837 | 1.4170507 | 1.22714733 | cluster 3 |
| HORVU.MOREX.r3.5HG0469300.1 | 0.4322471 | 1.8371289 | 1.64450062 | cluster 3 |
| HORVU.MOREX.r3.5HG0469340.1 | 0.6755812 | 1.3640911 | 1.28897872 | cluster 3 |
| HORVU.MOREX.r3.5HG0470680.1 | 1.07089 | 1.8095719 | 1.34311909 | cluster 3 |
| HORVU.MOREX.r3.5HG0470700.1 | 1.1686656 | 1.7112834 | 1.62797996 | cluster 3 |
| HORVU.MOREX.r3.5HG0472770.1 | 0.3644496 | 0.9356928 | 0.66081951 | cluster 3 |
| HORVU.MOREX.r3.5HG0480130.1 | 2.2228809 | 2.7550735 | 2.36662323 | cluster 3 |
| HORVU.MOREX.r3.5HG0480980.1 | 2.1262608 | 2.8208384 | 2.46573355 | cluster 3 |
| HORVU.MOREX.r3.5HG0481580.1 | 0.1811226 | 0.7633795 | 0.58267162 | cluster 3 |
| HORVU.MOREX.r3.5HG0483980.1 | 0.4557283 | 1.4910701 | 1.16530945 | cluster 3 |
| HORVU.MOREX.r3.5HG0485780.1 | 1.6887318 | 3.0873953 | 2.53798604 | cluster 3 |
| HORVU.MOREX.r3.5HG0485790.1 | 0.5181595 | 0.7671405 | 0.56319019 | cluster 3 |
| HORVU.MOREX.r3.5HG0485940.1 | 1.324522 | 2.2839296 | 1.85733786 | cluster 3 |
| HORVU.MOREX.r3.5HG0486660.1 | 1.0465157 | 1.3277827 | 1.24955444 | cluster 3 |
| HORVU.MOREX.r3.5HG0488040.1 | 1.0923523 | 1.7732844 | 1.62325311 | cluster 3 |
| HORVU.MOREX.r3.5HG0488050.1 | 1.0785668 | 1.8958393 | 1.44670761 | cluster 3 |
| HORVU.MOREX.r3.5HG0488300.1 | 0.6213391 | 1.4837628 | 1.24871352 | cluster 3 |
| HORVU.MOREX.r3.5HG0488650.1 | 0.4018848 | 0.7007772 | 0.50270261 | cluster 3 |
| HORVU.MOREX.r3.5HG0493080.1 | 0.2535699 | 0.6497137 | 0.6015228 | cluster 3 |
| HORVU.MOREX.r3.5HG0493200.1 | 0.1955649 | 0.4277088 | 0.39170022 | cluster 3 |
| HORVU.MOREX.r3.5HG0493330.1 | 0.7777255 | 1.0852783 | 0.99429843 | cluster 3 |
| HORVU.MOREX.r3.5HG0498240.1 | 1.122667 | 2.6776854 | 2.35906406 | cluster 3 |
| HORVU.MOREX.r3.5HG0507680.1 | 0.7568425 | 1.5093324 | 1.28741254 | cluster 3 |
| HORVU.MOREX.r3.5HG0508880.1 | 1.0132383 | 1.7158097 | 1.50461804 | cluster 3 |
| HORVU.MOREX.r3.5HG0509120.1 | 0.9671099 | 1.3294133 | 1.03666518 | cluster 3 |
| HORVU.MOREX.r3.5HG0511820.1 | 1.3528892 | 2.8631742 | 2.27948748 | cluster 3 |
| HORVU.MOREX.r3.5HG0512210.1 | 2.4911558 | 3.3845249 | 2.9983563 | cluster 3 |
| HORVU.MOREX.r3.5HG0512230.1 | 0.6450639 | 1.4964435 | 1.18310829 | cluster 3 |
| HORVU.MOREX.r3.5HG0512250.1 | 0.5548324 | 1.4973299 | 1.30995135 | cluster 3 |
| HORVU.MOREX.r3.5HG0512490.1 | 0.8751636 | 1.7964817 | 1.66333167 | cluster 3 |
| HORVU.MOREX.r3.5HG0517760.1 | 1.3069569 | 2.0781906 | 1.69639787 | cluster 3 |
| HORVU.MOREX.r3.5HG0518560.1 | 1.8337137 | 2.7412758 | 2.22933652 | cluster 3 |
| HORVU.MOREX.r3.5HG0519120.1 | 1.3017572 | 1.7814097 | 1.49195287 | cluster 3 |
| HORVU.MOREX.r3.5HG0519660.1 | 0.2163523 | 0.466698 | 0.3453212 | cluster 3 |
| HORVU.MOREX.r3.5HG0521610.1 | 0.6700677 | 1.3505614 | 1.23466197 | cluster 3 |
| HORVU.MOREX.r3.5HG0522630.1 | 0.3678684 | 0.7218506 | 0.58376818 | cluster 3 |

**Table S3** Continued.

| ID | logFC_3 h | logFC_6 h | logFC_12 h | cluster No. |
|---|---|---|---|---|
| HORVU.MOREX.r3.5HG0524350.1 | 1.2023506 | 2.2913521 | 2.05571947 | cluster 3 |
| HORVU.MOREX.r3.5HG0527570.1 | 1.8854684 | 2.1718098 | 2.04833397 | cluster 3 |
| HORVU.MOREX.r3.5HG0528570.1 | 0.9358814 | 1.0829495 | 0.96066106 | cluster 3 |
| HORVU.MOREX.r3.5HG0528910.1 | 0.3617325 | 0.9156462 | 0.85130311 | cluster 3 |
| HORVU.MOREX.r3.5HG0532140.1 | 1.5492928 | 2.1014358 | 1.95829789 | cluster 3 |
| HORVU.MOREX.r3.5HG0533430.1 | 0.9031022 | 1.446154 | 1.33072424 | cluster 3 |
| HORVU.MOREX.r3.6HG0540020.1 | 0.8588432 | 1.5402701 | 1.25319459 | cluster 3 |
| HORVU.MOREX.r3.6HG0542640.1 | 0.5218459 | 1.2641997 | 1.1800139 | cluster 3 |
| HORVU.MOREX.r3.6HG0548710.1 | 1.0982311 | 2.3994057 | 2.18060188 | cluster 3 |
| HORVU.MOREX.r3.6HG0558350.1 | 1.9940078 | 3.1572343 | 2.50428293 | cluster 3 |
| HORVU.MOREX.r3.6HG0558430.1 | 0.148306 | 2.2535914 | 1.13767428 | cluster 3 |
| HORVU.MOREX.r3.6HG0560140.1 | 1.4626424 | 2.1736853 | 1.98031687 | cluster 3 |
| HORVU.MOREX.r3.6HG0568170.1 | 0.1992469 | 0.8528938 | 0.46445861 | cluster 3 |
| HORVU.MOREX.r3.6HG0570230.1 | 0.6332285 | 1.1625092 | 0.84009985 | cluster 3 |
| HORVU.MOREX.r3.6HG0571750.1 | 0.4053088 | 1.1426233 | 0.94440251 | cluster 3 |
| HORVU.MOREX.r3.6HG0575530.1 | 0.2038005 | 0.4900524 | 0.37931383 | cluster 3 |
| HORVU.MOREX.r3.6HG0577040.1 | 0.7590438 | 2.1745454 | 2.01778438 | cluster 3 |
| HORVU.MOREX.r3.6HG0577220.1 | 0.737418 | 1.8475852 | 1.43370062 | cluster 3 |
| HORVU.MOREX.r3.6HG0579050.1 | 0.7332115 | 1.2562127 | 0.8746038 | cluster 3 |
| HORVU.MOREX.r3.6HG0593260.1 | 0.1786921 | 0.7811675 | 0.48834429 | cluster 3 |
| HORVU.MOREX.r3.6HG0595190.1 | 0.8667652 | 1.6978509 | 1.51950652 | cluster 3 |
| HORVU.MOREX.r3.6HG0600000.1 | 0.8610614 | 1.6195013 | 1.48165102 | cluster 3 |
| HORVU.MOREX.r3.6HG0600910.1 | 1.1219058 | 1.5254939 | 1.32431676 | cluster 3 |
| HORVU.MOREX.r3.6HG0607590.1 | 0.7460334 | 1.0357551 | 0.93750755 | cluster 3 |
| HORVU.MOREX.r3.6HG0608960.1 | 0.4282294 | 0.6753256 | 0.52999535 | cluster 3 |
| HORVU.MOREX.r3.6HG0609830.1 | 0.4477905 | 0.7712683 | 0.66177301 | cluster 3 |
| HORVU.MOREX.r3.6HG0610980.1 | 0.6627385 | 1.4841871 | 1.29351447 | cluster 3 |
| HORVU.MOREX.r3.6HG0611050.1 | 0.5382335 | 2.4658504 | 1.62772118 | cluster 3 |
| HORVU.MOREX.r3.6HG0612780.1 | 1.1300789 | 1.6260166 | 1.53359698 | cluster 3 |
| HORVU.MOREX.r3.6HG0613360.1 | 1.0940168 | 1.971445 | 1.36987849 | cluster 3 |
| HORVU.MOREX.r3.6HG0613430.1 | 1.2341802 | 1.9262015 | 1.68468874 | cluster 3 |
| HORVU.MOREX.r3.6HG0613460.1 | 0.9048869 | 1.1602723 | 1.0746214 | cluster 3 |
| HORVU.MOREX.r3.6HG0614100.1 | 0.335877 | 0.8316447 | 0.6733021 | cluster 3 |
| HORVU.MOREX.r3.6HG0615840.1 | 2.543944 | 3.0297172 | 2.78659421 | cluster 3 |
| HORVU.MOREX.r3.6HG0616240.1 | 0.5017971 | 0.8374594 | 0.74187336 | cluster 3 |
| HORVU.MOREX.r3.6HG0616450.1 | 0.4330728 | 1.3871709 | 1.12929946 | cluster 3 |
| HORVU.MOREX.r3.6HG0617970.1 | 1.0835867 | 1.7436559 | 1.57859963 | cluster 3 |
| HORVU.MOREX.r3.6HG0618180.1 | 0.854205 | 1.4432678 | 1.30328419 | cluster 3 |
| HORVU.MOREX.r3.6HG0620200.1 | 0.5097715 | 1.4024548 | 1.18997482 | cluster 3 |
| HORVU.MOREX.r3.6HG0621170.1 | 1.0628228 | 2.4977178 | 1.7546666 | cluster 3 |
| HORVU.MOREX.r3.6HG0622110.1 | 0.9349369 | 1.6219918 | 1.42377366 | cluster 3 |
| HORVU.MOREX.r3.6HG0622740.1 | 0.0651294 | 0.4522345 | 0.37215229 | cluster 3 |
| HORVU.MOREX.r3.6HG0624390.1 | 0.2569616 | 1.2637085 | 1.05390227 | cluster 3 |
| HORVU.MOREX.r3.6HG0626020.1 | 0.9358838 | 1.5998957 | 1.42864688 | cluster 3 |
| HORVU.MOREX.r3.6HG0626030.1 | 1.3209903 | 2.1629848 | 1.72243897 | cluster 3 |
| HORVU.MOREX.r3.6HG0629520.1 | 0.9316716 | 1.8149024 | 1.52849703 | cluster 3 |
| HORVU.MOREX.r3.6HG0629810.2 | 0.7057222 | 1.0147678 | 0.92128087 | cluster 3 |

**Table S3** Continued.

| ID | logFC_3 h | logFC_6 h | logFC_12 h | cluster No. |
|---|---|---|---|---|
| HORVU.MOREX.r3.6HG0631660.1 | 0.3410089 | 0.6921533 | 0.559483 | cluster 3 |
| HORVU.MOREX.r3.6HG0633900.1 | 1.2350146 | 2.2505326 | 1.82618881 | cluster 3 |
| HORVU.MOREX.r3.6HG0633960.1 | 0.3834116 | 0.9775832 | 0.88169322 | cluster 3 |
| HORVU.MOREX.r3.7HG0636890.1 | 0.9542233 | 1.9684083 | 1.6790455 | cluster 3 |
| HORVU.MOREX.r3.7HG0637030.1 | 0.37647 | 1.1998429 | 0.97163466 | cluster 3 |
| HORVU.MOREX.r3.7HG0641180.1 | 0.2307557 | 0.4779287 | 0.36897876 | cluster 3 |
| HORVU.MOREX.r3.7HG0641860.1 | 1.520959 | 2.1056133 | 1.96930234 | cluster 3 |
| HORVU.MOREX.r3.7HG0644220.1 | 0.0606799 | 0.5812324 | 0.23246826 | cluster 3 |
| HORVU.MOREX.r3.7HG0647300.1 | 1.1971558 | 2.3443238 | 1.78520597 | cluster 3 |
| HORVU.MOREX.r3.7HG0647950.1 | -0.159238 | 2.3282228 | 1.24488122 | cluster 3 |
| HORVU.MOREX.r3.7HG0648390.1 | 0.5570917 | 0.9313779 | 0.72074745 | cluster 3 |
| HORVU.MOREX.r3.7HG0649240.1 | 1.1203123 | 1.6543708 | 1.54596008 | cluster 3 |
| HORVU.MOREX.r3.7HG0655170.1 | 0.7339463 | 1.3564245 | 1.13663409 | cluster 3 |
| HORVU.MOREX.r3.7HG0656570.1 | 0.4039746 | 0.9104744 | 0.78599654 | cluster 3 |
| HORVU.MOREX.r3.7HG0658560.1 | 0.1447342 | 0.7394279 | 0.51293477 | cluster 3 |
| HORVU.MOREX.r3.7HG0660720.1 | 0.9043602 | 1.6722795 | 1.04166286 | cluster 3 |
| HORVU.MOREX.r3.7HG0663920.1 | 1.8027123 | 2.6206634 | 2.35411352 | cluster 3 |
| HORVU.MOREX.r3.7HG0663940.1 | 0.4773975 | 0.8353165 | 0.78775812 | cluster 3 |
| HORVU.MOREX.r3.7HG0667060.1 | 0.4364638 | 1.6455151 | 0.87384296 | cluster 3 |
| HORVU.MOREX.r3.7HG0668820.1 | 0.1926463 | 0.5634167 | 0.50029293 | cluster 3 |
| HORVU.MOREX.r3.7HG0669410.1 | 0.2278248 | 0.5083021 | 0.40417588 | cluster 3 |
| HORVU.MOREX.r3.7HG0670900.1 | 0.9578828 | 1.6657957 | 1.57099504 | cluster 3 |
| HORVU.MOREX.r3.7HG0671410.1 | 0.6661116 | 1.2920591 | 0.89859128 | cluster 3 |
| HORVU.MOREX.r3.7HG0675310.1 | 0.8329982 | 1.4561707 | 1.23997831 | cluster 3 |
| HORVU.MOREX.r3.7HG0676040.1 | 0.0551884 | 0.6571765 | 0.56005952 | cluster 3 |
| HORVU.MOREX.r3.7HG0680960.1 | 1.0264564 | 2.0012917 | 1.61550618 | cluster 3 |
| HORVU.MOREX.r3.7HG0683600.1 | 1.2594735 | 1.8515597 | 1.62245145 | cluster 3 |
| HORVU.MOREX.r3.7HG0688730.1 | 0.1733679 | 0.5156428 | 0.42665333 | cluster 3 |
| HORVU.MOREX.r3.7HG0688870.1 | 0.5712165 | 0.7818879 | 0.74855596 | cluster 3 |
| HORVU.MOREX.r3.7HG0701080.1 | 0.7074732 | 0.9843881 | 0.8755626 | cluster 3 |
| HORVU.MOREX.r3.7HG0706130.1 | 0.4785005 | 0.6950621 | 0.54304118 | cluster 3 |
| HORVU.MOREX.r3.7HG0711230.1 | 0.3609468 | 0.5896796 | 0.45582649 | cluster 3 |
| HORVU.MOREX.r3.7HG0712000.1 | 1.0152618 | 1.9992251 | 1.73022492 | cluster 3 |
| HORVU.MOREX.r3.7HG0714400.1 | 0.437687 | 0.8739298 | 0.78423884 | cluster 3 |
| HORVU.MOREX.r3.7HG0714660.1 | 0.1668459 | 1.5116325 | 1.15283434 | cluster 3 |
| HORVU.MOREX.r3.7HG0716640.1 | 0.5646628 | 0.8312807 | 0.60267475 | cluster 3 |
| HORVU.MOREX.r3.7HG0721320.1 | 0.3195789 | 1.3864357 | 0.75136531 | cluster 3 |
| HORVU.MOREX.r3.7HG0722260.1 | 1.1134465 | 1.8066255 | 1.63999031 | cluster 3 |
| HORVU.MOREX.r3.7HG0727650.1 | 1.4778651 | 2.7853943 | 2.62748075 | cluster 3 |
| HORVU.MOREX.r3.7HG0728000.1 | 0.3480905 | 0.6297342 | 0.58300689 | cluster 3 |
| HORVU.MOREX.r3.7HG0728080.1 | 0.3138312 | 0.8212387 | 0.4492996 | cluster 3 |
| HORVU.MOREX.r3.7HG0728480.1 | 1.0632378 | 1.5774043 | 1.40350588 | cluster 3 |
| HORVU.MOREX.r3.7HG0728820.1 | 0.6658363 | 1.7114571 | 1.42708304 | cluster 3 |
| HORVU.MOREX.r3.7HG0729000.1 | 0.5445362 | 1.0377436 | 0.97930677 | cluster 3 |
| HORVU.MOREX.r3.7HG0730530.1 | 0.4549969 | 1.3054203 | 1.00940735 | cluster 3 |
| HORVU.MOREX.r3.7HG0730660.1 | 1.1890454 | 1.6958903 | 1.50246046 | cluster 3 |
| HORVU.MOREX.r3.7HG0730870.1 | 0.2037409 | 0.3743778 | 0.35057178 | cluster 3 |

**Table S3** Continued.

| ID | logFC_3 h | logFC_6 h | logFC_12 h | cluster No. |
|---|---|---|---|---|
| HORVU.MOREX.r3.7HG0737340.1 | 0.4232764 | 1.6567824 | 1.39201746 | cluster 3 |
| HORVU.MOREX.r3.7HG0739080.1 | 1.6079733 | 2.592859 | 2.34402343 | cluster 3 |
| HORVU.MOREX.r3.7HG0742370.1 | 0.5386573 | 1.3387466 | 1.24105096 | cluster 3 |
| HORVU.MOREX.r3.7HG0747230.1 | 0.6875564 | 0.9838461 | 0.74613293 | cluster 3 |
| HORVU.MOREX.r3.7HG0747720.1 | 0.3506165 | 1.0227787 | 0.69360431 | cluster 3 |
| HORVU.MOREX.r3.7HG0748270.1 | 1.2442207 | 1.9760816 | 1.72587828 | cluster 3 |
| HORVU.MOREX.r3.7HG0748670.1 | 0.9419249 | 2.3304367 | 1.69523928 | cluster 3 |
| HORVU.MOREX.r3.7HG0750000.1 | 1.7484813 | 2.3774016 | 2.11315091 | cluster 3 |
| HORVU.MOREX.r3.7HG0751450.1 | 1.0234693 | 2.0879019 | 1.42930168 | cluster 3 |
| HORVU.MOREX.r3.1HG0000050.1 | -0.210564 | -0.486688 | -0.4453408 | cluster 4 |
| HORVU.MOREX.r3.1HG0000140.1 | -0.057747 | -0.364258 | -0.353427 | cluster 4 |
| HORVU.MOREX.r3.1HG0002340.1 | -0.090387 | -0.30519 | -0.2024824 | cluster 4 |
| HORVU.MOREX.r3.1HG0006230.1 | -0.265836 | -0.627234 | -0.6510553 | cluster 4 |
| HORVU.MOREX.r3.1HG0014430.1 | -0.117054 | -0.198848 | -0.2041247 | cluster 4 |
| HORVU.MOREX.r3.1HG0020270.1 | -0.148247 | -0.379992 | -0.3739876 | cluster 4 |
| HORVU.MOREX.r3.1HG0037910.1 | -0.171377 | -0.704039 | -0.6912778 | cluster 4 |
| HORVU.MOREX.r3.1HG0039370.1 | -0.069398 | -0.402941 | -0.407958 | cluster 4 |
| HORVU.MOREX.r3.1HG0043520.1 | -0.238147 | -0.519477 | -0.50014 | cluster 4 |
| HORVU.MOREX.r3.1HG0049450.1 | -0.131521 | -0.451091 | -0.4273442 | cluster 4 |
| HORVU.MOREX.r3.1HG0049520.1 | -0.13782 | -0.732898 | -0.7527107 | cluster 4 |
| HORVU.MOREX.r3.1HG0051960.1 | -0.220278 | -1.091 | -1.1754849 | cluster 4 |
| HORVU.MOREX.r3.1HG0052940.1 | -0.090563 | -0.378274 | -0.4074182 | cluster 4 |
| HORVU.MOREX.r3.1HG0053680.1 | -0.078399 | -0.302188 | -0.2049298 | cluster 4 |
| HORVU.MOREX.r3.1HG0055560.1 | -0.002046 | -0.40991 | -0.4294738 | cluster 4 |
| HORVU.MOREX.r3.1HG0057000.1 | -0.154442 | -1.170442 | -1.2252833 | cluster 4 |
| HORVU.MOREX.r3.1HG0057030.1 | -0.195214 | -0.854579 | -0.6987391 | cluster 4 |
| HORVU.MOREX.r3.1HG0058470.1 | -0.134275 | -0.379924 | -0.2626498 | cluster 4 |
| HORVU.MOREX.r3.1HG0059340.1 | -0.171788 | -0.34212 | -0.2890759 | cluster 4 |
| HORVU.MOREX.r3.1HG0063810.1 | -0.087931 | -0.413414 | -0.2988521 | cluster 4 |
| HORVU.MOREX.r3.1HG0066260.1 | -0.11812 | -0.444257 | -0.4658723 | cluster 4 |
| HORVU.MOREX.r3.1HG0069210.1 | -0.028778 | -0.703608 | -0.4911704 | cluster 4 |
| HORVU.MOREX.r3.1HG0069570.1 | -0.134462 | -0.387142 | -0.3771681 | cluster 4 |
| HORVU.MOREX.r3.1HG0072760.1 | -0.139447 | -0.819272 | -0.9126195 | cluster 4 |
| HORVU.MOREX.r3.1HG0072840.1 | -0.10212 | -0.342829 | -0.3691419 | cluster 4 |
| HORVU.MOREX.r3.1HG0072910.2 | -0.222798 | -0.514361 | -0.5098969 | cluster 4 |
| HORVU.MOREX.r3.1HG0073520.1 | -0.227209 | -0.749321 | -0.7818548 | cluster 4 |
| HORVU.MOREX.r3.1HG0081620.1 | -0.44562 | -0.670831 | -0.6509182 | cluster 4 |
| HORVU.MOREX.r3.1HG0083040.1 | -0.377981 | -0.74817 | -0.5874048 | cluster 4 |
| HORVU.MOREX.r3.1HG0087170.1 | -0.246922 | -0.540091 | -0.543782 | cluster 4 |
| HORVU.MOREX.r3.1HG0090170.1 | -0.274432 | -0.587936 | -0.5627546 | cluster 4 |
| HORVU.MOREX.r3.1HG0090200.1 | -0.137066 | -0.510105 | -0.4726677 | cluster 4 |
| HORVU.MOREX.r3.1HG0090710.1 | -0.225109 | -0.39678 | -0.3901458 | cluster 4 |
| HORVU.MOREX.r3.1HG0094770.1 | -0.184493 | -0.551202 | -0.4861682 | cluster 4 |
| HORVU.MOREX.r3.1HG0094980.1 | -0.227694 | -0.595078 | -0.4299233 | cluster 4 |
| HORVU.MOREX.r3.2HG0098090.1 | -0.358358 | -1.227477 | -1.2327465 | cluster 4 |
| HORVU.MOREX.r3.2HG0102600.1 | -0.164277 | -0.410456 | -0.3688374 | cluster 4 |
| HORVU.MOREX.r3.2HG0104520.1 | -0.177553 | -0.553732 | -0.5874395 | cluster 4 |

**Table S3** Continued.

| ID | logFC_3 h | logFC_6 h | logFC_12 h | cluster No. |
|---|---|---|---|---|
| HORVU.MOREX.r3.2HG0108780.1 | -0.065142 | -0.205477 | -0.1530493 | cluster 4 |
| HORVU.MOREX.r3.2HG0112060.1 | -0.280124 | -0.780091 | -0.7503091 | cluster 4 |
| HORVU.MOREX.r3.2HG0117980.1 | -0.154248 | -0.359594 | -0.2791979 | cluster 4 |
| HORVU.MOREX.r3.2HG0121570.1 | -0.21981 | -0.507466 | -0.4110998 | cluster 4 |
| HORVU.MOREX.r3.2HG0122500.2 | -0.384472 | -1.657488 | -1.8314263 | cluster 4 |
| HORVU.MOREX.r3.2HG0124450.1 | -0.119997 | -0.549691 | -0.5896555 | cluster 4 |
| HORVU.MOREX.r3.2HG0128240.1 | -0.203944 | -0.407351 | -0.3836201 | cluster 4 |
| HORVU.MOREX.r3.2HG0128490.1 | -0.094818 | -0.347458 | -0.3261412 | cluster 4 |
| HORVU.MOREX.r3.2HG0132590.1 | -0.131936 | -0.331505 | -0.258935 | cluster 4 |
| HORVU.MOREX.r3.2HG0135510.1 | -0.092381 | -0.318173 | -0.3435911 | cluster 4 |
| HORVU.MOREX.r3.2HG0140700.1 | -0.135934 | -0.314454 | -0.2293707 | cluster 4 |
| HORVU.MOREX.r3.2HG0140970.1 | -0.280162 | -0.804431 | -0.859195 | cluster 4 |
| HORVU.MOREX.r3.2HG0141980.1 | -0.152674 | -0.403651 | -0.4324579 | cluster 4 |
| HORVU.MOREX.r3.2HG0144230.1 | -0.093503 | -0.303229 | -0.3259337 | cluster 4 |
| HORVU.MOREX.r3.2HG0152430.1 | -0.157479 | -0.477822 | -0.3368035 | cluster 4 |
| HORVU.MOREX.r3.2HG0152890.1 | -0.480596 | -0.656631 | -0.6549707 | cluster 4 |
| HORVU.MOREX.r3.2HG0156810.1 | 0.1657929 | -1.605093 | -0.7460263 | cluster 4 |
| HORVU.MOREX.r3.2HG0161410.1 | -0.124115 | -0.33139 | -0.3275692 | cluster 4 |
| HORVU.MOREX.r3.2HG0161580.1 | -0.283235 | -0.552608 | -0.5039145 | cluster 4 |
| HORVU.MOREX.r3.2HG0164850.1 | -0.306654 | -0.837785 | -0.6724567 | cluster 4 |
| HORVU.MOREX.r3.2HG0171810.1 | -0.221987 | -0.489077 | -0.5187989 | cluster 4 |
| HORVU.MOREX.r3.2HG0182990.1 | -0.179004 | -0.792773 | -0.6544065 | cluster 4 |
| HORVU.MOREX.r3.2HG0188490.1 | -0.066109 | -0.808051 | -0.6820208 | cluster 4 |
| HORVU.MOREX.r3.2HG0192910.1 | -0.109653 | -0.787404 | -0.7465992 | cluster 4 |
| HORVU.MOREX.r3.2HG0193000.1 | -0.196324 | -0.71756 | -0.6826582 | cluster 4 |
| HORVU.MOREX.r3.2HG0196940.1 | -0.053432 | -0.341399 | -0.2791273 | cluster 4 |
| HORVU.MOREX.r3.2HG0199550.1 | -0.066644 | -0.521988 | -0.4085273 | cluster 4 |
| HORVU.MOREX.r3.2HG0200730.1 | -0.067512 | -0.196754 | -0.1968357 | cluster 4 |
| HORVU.MOREX.r3.2HG0204340.1 | -0.613565 | -2.387041 | -2.2924992 | cluster 4 |
| HORVU.MOREX.r3.2HG0208960.1 | -0.332485 | -0.850496 | -0.8815784 | cluster 4 |
| HORVU.MOREX.r3.2HG0210680.1 | -0.013726 | -0.28563 | -0.2724734 | cluster 4 |
| HORVU.MOREX.r3.2HG0211730.1 | -0.139536 | -0.501831 | -0.3484092 | cluster 4 |
| HORVU.MOREX.r3.2HG0211880.1 | -0.168738 | -0.456205 | -0.4618274 | cluster 4 |
| HORVU.MOREX.r3.2HG0214240.1 | -0.330191 | -0.720019 | -0.63166 | cluster 4 |
| HORVU.MOREX.r3.3HG0219750.1 | -0.694726 | -1.735553 | -1.523473 | cluster 4 |
| HORVU.MOREX.r3.3HG0221490.1 | -0.11546 | -0.349416 | -0.2641155 | cluster 4 |
| HORVU.MOREX.r3.3HG0222130.1 | -0.11536 | -0.341379 | -0.3573973 | cluster 4 |
| HORVU.MOREX.r3.3HG0222830.1 | -0.229299 | -0.48403 | -0.41894 | cluster 4 |
| HORVU.MOREX.r3.3HG0228010.1 | -0.205685 | -0.397499 | -0.3962321 | cluster 4 |
| HORVU.MOREX.r3.3HG0230880.1 | -0.097587 | -0.213501 | -0.2081264 | cluster 4 |
| HORVU.MOREX.r3.3HG0235030.1 | -0.184656 | -0.516033 | -0.5021107 | cluster 4 |
| HORVU.MOREX.r3.3HG0235720.1 | -0.451454 | -2.293505 | -2.0720386 | cluster 4 |
| HORVU.MOREX.r3.3HG0237330.1 | -0.88099 | -1.616044 | -1.6289269 | cluster 4 |
| HORVU.MOREX.r3.3HG0238250.1 | -0.611268 | -0.985374 | -0.8984795 | cluster 4 |
| HORVU.MOREX.r3.3HG0239940.1 | -0.106152 | -0.564578 | -0.605368 | cluster 4 |
| HORVU.MOREX.r3.3HG0240310.1 | -0.369458 | -1.122443 | -1.1950132 | cluster 4 |
| HORVU.MOREX.r3.3HG0242950.1 | -0.190897 | -0.433324 | -0.4367021 | cluster 4 |

**Table S3** Continued.

| ID | logFC_3 h | logFC_6 h | logFC_12 h | cluster No. |
|---|---|---|---|---|
| HORVU.MOREX.r3.3HG0243130.1 | -0.486099 | -0.756778 | -0.6823611 | cluster 4 |
| HORVU.MOREX.r3.3HG0246540.1 | -0.304757 | -0.834515 | -0.8782421 | cluster 4 |
| HORVU.MOREX.r3.3HG0247440.1 | -0.412838 | -1.038788 | -0.9144083 | cluster 4 |
| HORVU.MOREX.r3.3HG0250870.1 | -0.162252 | -0.503649 | -0.5000841 | cluster 4 |
| HORVU.MOREX.r3.3HG0250940.1 | -0.189537 | -0.60542 | -0.5773945 | cluster 4 |
| HORVU.MOREX.r3.3HG0253450.1 | -0.011765 | -0.368997 | -0.3293449 | cluster 4 |
| HORVU.MOREX.r3.3HG0256400.1 | 0.0735805 | -0.499938 | -0.3785823 | cluster 4 |
| HORVU.MOREX.r3.3HG0257310.1 | -0.098947 | -0.273317 | -0.2255739 | cluster 4 |
| HORVU.MOREX.r3.3HG0264700.1 | -0.285675 | -0.333653 | -0.3400111 | cluster 4 |
| HORVU.MOREX.r3.3HG0274100.1 | -1.067729 | -1.297125 | -1.2686463 | cluster 4 |
| HORVU.MOREX.r3.3HG0280630.1 | -0.172824 | -0.41087 | -0.4177276 | cluster 4 |
| HORVU.MOREX.r3.3HG0280920.1 | -0.416991 | -0.733995 | -0.6264072 | cluster 4 |
| HORVU.MOREX.r3.3HG0280970.1 | -0.077547 | -0.406923 | -0.414523 | cluster 4 |
| HORVU.MOREX.r3.3HG0281730.1 | -0.132377 | -1.52473 | -1.3651815 | cluster 4 |
| HORVU.MOREX.r3.3HG0288080.1 | -0.066388 | -0.254736 | -0.2583273 | cluster 4 |
| HORVU.MOREX.r3.3HG0288360.1 | -0.212077 | -0.287403 | -0.2583098 | cluster 4 |
| HORVU.MOREX.r3.3HG0288370.1 | -0.486641 | -0.972153 | -1.0220741 | cluster 4 |
| HORVU.MOREX.r3.3HG0292080.1 | -0.144733 | -0.49333 | -0.4810081 | cluster 4 |
| HORVU.MOREX.r3.3HG0293430.1 | 1.3180018 | 1.1773024 | 1.20780458 | cluster 4 |
| HORVU.MOREX.r3.3HG0293570.1 | -0.215612 | -2.422838 | -1.605738 | cluster 4 |
| HORVU.MOREX.r3.3HG0294670.1 | -0.848006 | -1.583304 | -1.383073 | cluster 4 |
| HORVU.MOREX.r3.3HG0297330.1 | -0.257665 | -0.634878 | -0.5272362 | cluster 4 |
| HORVU.MOREX.r3.3HG0300950.1 | -0.331933 | -0.855851 | -0.8120216 | cluster 4 |
| HORVU.MOREX.r3.3HG0302340.1 | -0.961716 | -2.742539 | -2.4059971 | cluster 4 |
| HORVU.MOREX.r3.3HG0304780.1 | -0.217809 | -0.513246 | -0.5397324 | cluster 4 |
| HORVU.MOREX.r3.3HG0306060.1 | -0.108451 | -0.40301 | -0.373696 | cluster 4 |
| HORVU.MOREX.r3.3HG0307850.1 | -0.204906 | -0.693866 | -0.7553537 | cluster 4 |
| HORVU.MOREX.r3.3HG0307950.1 | -0.100394 | -0.303004 | -0.271704 | cluster 4 |
| HORVU.MOREX.r3.3HG0310460.1 | -0.18705 | -0.805496 | -0.5478687 | cluster 4 |
| HORVU.MOREX.r3.3HG0313020.1 | -0.155766 | -0.357107 | -0.332207 | cluster 4 |
| HORVU.MOREX.r3.3HG0314240.1 | -0.180125 | -0.417324 | -0.4126967 | cluster 4 |
| HORVU.MOREX.r3.3HG0316130.1 | -0.227113 | -0.695321 | -0.7369012 | cluster 4 |
| HORVU.MOREX.r3.3HG0322740.1 | -0.227097 | -0.902679 | -0.6225242 | cluster 4 |
| HORVU.MOREX.r3.3HG0324180.1 | -0.412241 | -0.868378 | -0.8212219 | cluster 4 |
| HORVU.MOREX.r3.3HG0325670.1 | -0.125844 | -0.621853 | -0.6298433 | cluster 4 |
| HORVU.MOREX.r3.3HG0326430.1 | -0.072333 | -0.365205 | -0.2748693 | cluster 4 |
| HORVU.MOREX.r3.4HG0331630.1 | -0.032278 | -0.369393 | -0.3918205 | cluster 4 |
| HORVU.MOREX.r3.4HG0331730.1 | -0.167803 | -0.308637 | -0.3063395 | cluster 4 |
| HORVU.MOREX.r3.4HG0335180.1 | -0.316712 | -0.622832 | -0.6396912 | cluster 4 |
| HORVU.MOREX.r3.4HG0335700.1 | -0.098305 | -0.239357 | -0.2007837 | cluster 4 |
| HORVU.MOREX.r3.4HG0337110.1 | -0.119306 | -0.40188 | -0.3460329 | cluster 4 |
| HORVU.MOREX.r3.4HG0343540.1 | -0.886864 | -1.868205 | -1.4943153 | cluster 4 |
| HORVU.MOREX.r3.4HG0344830.1 | -0.094413 | -0.449223 | -0.328731 | cluster 4 |
| HORVU.MOREX.r3.4HG0345610.1 | -0.137979 | -0.285953 | -0.2416721 | cluster 4 |
| HORVU.MOREX.r3.4HG0347400.1 | -0.146166 | -0.452852 | -0.3028916 | cluster 4 |
| HORVU.MOREX.r3.4HG0349310.1 | -0.142031 | -0.410999 | -0.2838386 | cluster 4 |
| HORVU.MOREX.r3.4HG0350900.1 | -0.161247 | -0.44244 | -0.4339045 | cluster 4 |

**Table S3** Continued.

| ID | logFC_3 h | logFC_6 h | logFC_12 h | cluster No. |
|---|---|---|---|---|
| HORVU.MOREX.r3.4HG0352540.1 | -0.218739 | -0.583265 | -0.5703874 | cluster 4 |
| HORVU.MOREX.r3.4HG0355390.1 | -0.152652 | -0.745518 | -0.7235153 | cluster 4 |
| HORVU.MOREX.r3.4HG0355460.1 | -0.219534 | -0.463683 | -0.3953742 | cluster 4 |
| HORVU.MOREX.r3.4HG0378000.1 | -0.114183 | -0.285587 | -0.2721397 | cluster 4 |
| HORVU.MOREX.r3.4HG0379290.1 | -0.138201 | -0.316911 | -0.3198258 | cluster 4 |
| HORVU.MOREX.r3.4HG0379530.1 | -0.234909 | -0.402578 | -0.4169981 | cluster 4 |
| HORVU.MOREX.r3.4HG0380330.1 | -0.152072 | -0.499123 | -0.5291229 | cluster 4 |
| HORVU.MOREX.r3.4HG0381800.1 | -0.120611 | -0.279721 | -0.2693879 | cluster 4 |
| HORVU.MOREX.r3.4HG0384230.1 | -0.154127 | -0.579808 | -0.637713 | cluster 4 |
| HORVU.MOREX.r3.4HG0390700.1 | -0.060572 | -0.194416 | -0.2085167 | cluster 4 |
| HORVU.MOREX.r3.4HG0391730.1 | -0.124635 | -0.317151 | -0.2467658 | cluster 4 |
| HORVU.MOREX.r3.4HG0392590.1 | -0.029707 | -0.420046 | -0.432657 | cluster 4 |
| HORVU.MOREX.r3.4HG0400740.1 | -0.099135 | -0.304881 | -0.2874618 | cluster 4 |
| HORVU.MOREX.r3.4HG0401730.1 | -0.246305 | -0.713924 | -0.7434039 | cluster 4 |
| HORVU.MOREX.r3.4HG0402000.1 | -0.241346 | -0.71102 | -0.6381152 | cluster 4 |
| HORVU.MOREX.r3.4HG0403370.1 | -0.130151 | -0.402659 | -0.3882804 | cluster 4 |
| HORVU.MOREX.r3.4HG0403510.1 | -0.111716 | -0.413298 | -0.3832699 | cluster 4 |
| HORVU.MOREX.r3.4HG0403680.1 | -0.225767 | -1.264573 | -1.3828257 | cluster 4 |
| HORVU.MOREX.r3.4HG0409180.1 | -0.074596 | -0.328478 | -0.2718058 | cluster 4 |
| HORVU.MOREX.r3.4HG0410640.1 | -0.124296 | -0.378262 | -0.3970308 | cluster 4 |
| HORVU.MOREX.r3.4HG0416090.1 | -0.111536 | -0.523653 | -0.5069306 | cluster 4 |
| HORVU.MOREX.r3.4HG0416760.1 | -0.239652 | -0.586074 | -0.5290765 | cluster 4 |
| HORVU.MOREX.r3.5HG0420460.1 | 3.0932412 | 2.6734724 | 2.79652069 | cluster 4 |
| HORVU.MOREX.r3.5HG0420630.1 | -0.18813 | -0.480883 | -0.4727335 | cluster 4 |
| HORVU.MOREX.r3.5HG0434380.1 | -0.191479 | -0.482553 | -0.3843015 | cluster 4 |
| HORVU.MOREX.r3.5HG0436660.1 | -0.224065 | -0.476557 | -0.4550899 | cluster 4 |
| HORVU.MOREX.r3.5HG0446900.1 | -0.036148 | -0.218937 | -0.2236596 | cluster 4 |
| HORVU.MOREX.r3.5HG0454660.1 | -0.208314 | -0.40023 | -0.4169716 | cluster 4 |
| HORVU.MOREX.r3.5HG0458440.1 | -0.042315 | -1.067416 | -0.9131093 | cluster 4 |
| HORVU.MOREX.r3.5HG0459320.1 | -0.201293 | -0.450674 | -0.4424902 | cluster 4 |
| HORVU.MOREX.r3.5HG0460630.1 | 0.1977752 | -0.980742 | -0.6336549 | cluster 4 |
| HORVU.MOREX.r3.5HG0463000.1 | -0.214065 | -0.448239 | -0.3731562 | cluster 4 |
| HORVU.MOREX.r3.5HG0466440.1 | -0.253495 | -0.496231 | -0.3984102 | cluster 4 |
| HORVU.MOREX.r3.5HG0466510.1 | -0.281459 | -0.969587 | -0.761974 | cluster 4 |
| HORVU.MOREX.r3.5HG0476690.1 | 0.0280055 | -0.321602 | -0.3075335 | cluster 4 |
| HORVU.MOREX.r3.5HG0480890.1 | 1.8701414 | 1.2649812 | 1.45109118 | cluster 4 |
| HORVU.MOREX.r3.5HG0481760.1 | -0.061848 | -0.290194 | -0.2071981 | cluster 4 |
| HORVU.MOREX.r3.5HG0486960.1 | -0.654483 | -1.501356 | -1.1381945 | cluster 4 |
| HORVU.MOREX.r3.5HG0489720.1 | -0.177447 | -0.431894 | -0.4331791 | cluster 4 |
| HORVU.MOREX.r3.5HG0493380.1 | -0.068113 | -0.320081 | -0.2420716 | cluster 4 |
| HORVU.MOREX.r3.5HG0495490.1 | -0.201213 | -0.423029 | -0.4221545 | cluster 4 |
| HORVU.MOREX.r3.5HG0500120.1 | -0.522798 | -1.061761 | -1.1050016 | cluster 4 |
| HORVU.MOREX.r3.5HG0500180.1 | -0.129879 | -0.41309 | -0.3181799 | cluster 4 |
| HORVU.MOREX.r3.5HG0501500.1 | 1.1804435 | 1.1252077 | 1.12682132 | cluster 4 |
| HORVU.MOREX.r3.5HG0503200.1 | -0.057395 | -0.566785 | -0.609787 | cluster 4 |
| HORVU.MOREX.r3.5HG0510160.1 | -0.055237 | -0.459951 | -0.5016767 | cluster 4 |
| HORVU.MOREX.r3.5HG0510210.1 | -0.150813 | -0.356296 | -0.3731947 | cluster 4 |

**Table S3** Continued.

| ID | logFC_3 h | logFC_6 h | logFC_12 h | cluster No. |
|---|---|---|---|---|
| HORVU.MOREX.r3.5HG0519040.1 | -0.148928 | -0.368191 | -0.3427753 | cluster 4 |
| HORVU.MOREX.r3.5HG0520650.1 | -0.209542 | -0.68725 | -0.6756638 | cluster 4 |
| HORVU.MOREX.r3.5HG0520990.1 | -0.35326 | -0.741574 | -0.6553458 | cluster 4 |
| HORVU.MOREX.r3.5HG0522130.1 | -0.161695 | -0.483941 | -0.5132723 | cluster 4 |
| HORVU.MOREX.r3.5HG0524120.1 | -0.152452 | -0.366911 | -0.3915019 | cluster 4 |
| HORVU.MOREX.r3.5HG0526140.1 | -0.051704 | -0.403299 | -0.4518669 | cluster 4 |
| HORVU.MOREX.r3.5HG0526160.1 | -0.230133 | -0.497186 | -0.3693641 | cluster 4 |
| HORVU.MOREX.r3.5HG0526200.1 | -0.100757 | -0.272216 | -0.2658474 | cluster 4 |
| HORVU.MOREX.r3.5HG0527500.1 | -0.195335 | -0.51146 | -0.5397168 | cluster 4 |
| HORVU.MOREX.r3.5HG0529660.1 | -0.035728 | -0.271686 | -0.1610445 | cluster 4 |
| HORVU.MOREX.r3.5HG0533230.1 | -0.265764 | -0.528208 | -0.5553388 | cluster 4 |
| HORVU.MOREX.r3.5HG0534100.1 | -0.214006 | -0.820121 | -0.8873792 | cluster 4 |
| HORVU.MOREX.r3.5HG0537090.1 | -0.119397 | -0.327596 | -0.3455348 | cluster 4 |
| HORVU.MOREX.r3.6HG0543390.1 | 1.5419602 | 0.9739213 | 1.22556774 | cluster 4 |
| HORVU.MOREX.r3.6HG0546140.1 | -0.070897 | -0.758392 | -0.7023619 | cluster 4 |
| HORVU.MOREX.r3.6HG0548360.1 | -0.021977 | -0.595089 | -0.401464 | cluster 4 |
| HORVU.MOREX.r3.6HG0550940.1 | -0.189132 | -0.390098 | -0.3346286 | cluster 4 |
| HORVU.MOREX.r3.6HG0553150.1 | -0.386615 | -0.593359 | -0.5321773 | cluster 4 |
| HORVU.MOREX.r3.6HG0555660.2 | -0.228528 | -0.430502 | -0.4350808 | cluster 4 |
| HORVU.MOREX.r3.6HG0557070.1 | -0.17213 | -0.56379 | -0.4946179 | cluster 4 |
| HORVU.MOREX.r3.6HG0558250.1 | -0.094468 | -0.551478 | -0.5931937 | cluster 4 |
| HORVU.MOREX.r3.6HG0564910.1 | -0.158965 | -0.985196 | -1.0100623 | cluster 4 |
| HORVU.MOREX.r3.6HG0572400.1 | -0.235159 | -0.413275 | -0.3607975 | cluster 4 |
| HORVU.MOREX.r3.6HG0577360.1 | -0.171359 | -0.661974 | -0.6563386 | cluster 4 |
| HORVU.MOREX.r3.6HG0599080.1 | -0.121656 | -0.386829 | -0.3657578 | cluster 4 |
| HORVU.MOREX.r3.6HG0601670.1 | -0.310073 | -1.038854 | -0.9143986 | cluster 4 |
| HORVU.MOREX.r3.6HG0602230.1 | -0.231684 | -0.514986 | -0.5091945 | cluster 4 |
| HORVU.MOREX.r3.6HG0605090.1 | -0.04292 | -0.285138 | -0.3172045 | cluster 4 |
| HORVU.MOREX.r3.6HG0606850.1 | -0.155525 | -0.459291 | -0.3134643 | cluster 4 |
| HORVU.MOREX.r3.6HG0606930.1 | -0.071743 | -0.374522 | -0.4077279 | cluster 4 |
| HORVU.MOREX.r3.6HG0613230.1 | -0.113996 | -0.367812 | -0.3068486 | cluster 4 |
| HORVU.MOREX.r3.6HG0616890.1 | -0.536575 | -0.917484 | -0.9478679 | cluster 4 |
| HORVU.MOREX.r3.6HG0617730.1 | -0.078532 | -0.339833 | -0.3596403 | cluster 4 |
| HORVU.MOREX.r3.6HG0618760.1 | -0.292818 | -0.747648 | -0.796672 | cluster 4 |
| HORVU.MOREX.r3.6HG0621190.1 | -0.154558 | -0.673173 | -0.6018863 | cluster 4 |
| HORVU.MOREX.r3.6HG0622550.1 | -0.15326 | -0.349934 | -0.2835664 | cluster 4 |
| HORVU.MOREX.r3.6HG0626650.1 | -0.123334 | -0.267557 | -0.2501574 | cluster 4 |
| HORVU.MOREX.r3.6HG0632270.1 | -0.201751 | -0.511546 | -0.4392241 | cluster 4 |
| HORVU.MOREX.r3.6HG0632320.1 | -0.140583 | -0.360637 | -0.3880408 | cluster 4 |
| HORVU.MOREX.r3.6HG0633650.1 | -0.187888 | -1.172822 | -1.176745 | cluster 4 |
| HORVU.MOREX.r3.6HG0634130.1 | -0.228673 | -0.344764 | -0.3096888 | cluster 4 |
| HORVU.MOREX.r3.7HG0634430.1 | -0.05789 | -0.820241 | -0.530186 | cluster 4 |
| HORVU.MOREX.r3.7HG0634470.1 | -0.076647 | -0.276924 | -0.2886309 | cluster 4 |
| HORVU.MOREX.r3.7HG0639930.1 | -0.145969 | -0.513816 | -0.3434781 | cluster 4 |
| HORVU.MOREX.r3.7HG0640560.1 | -1.061191 | -3.052375 | -2.5346574 | cluster 4 |
| HORVU.MOREX.r3.7HG0643260.1 | -0.082668 | -0.639752 | -0.4085841 | cluster 4 |
| HORVU.MOREX.r3.7HG0658270.1 | -0.071725 | -0.604362 | -0.5618305 | cluster 4 |

**Table S3** Continued.

| ID | logFC_3 h | logFC_6 h | logFC_12 h | cluster No. |
|---|---|---|---|---|
| HORVU.MOREX.r3.7HG0661490.1 | -0.147876 | -0.307367 | -0.3085276 | cluster 4 |
| HORVU.MOREX.r3.7HG0666650.1 | -0.609304 | -1.748073 | -1.6004303 | cluster 4 |
| HORVU.MOREX.r3.7HG0671360.1 | -0.142628 | -0.337151 | -0.2559202 | cluster 4 |
| HORVU.MOREX.r3.7HG0674320.1 | -0.262906 | -0.368676 | -0.3746705 | cluster 4 |
| HORVU.MOREX.r3.7HG0674990.1 | -0.171202 | -0.465086 | -0.500011 | cluster 4 |
| HORVU.MOREX.r3.7HG0678160.1 | -0.270922 | -0.702624 | -0.6451568 | cluster 4 |
| HORVU.MOREX.r3.7HG0680970.1 | -0.361888 | -0.798843 | -0.6609786 | cluster 4 |
| HORVU.MOREX.r3.7HG0680990.1 | -0.129828 | -0.348204 | -0.2598531 | cluster 4 |
| HORVU.MOREX.r3.7HG0682460.1 | -0.164134 | -0.35679 | -0.3473942 | cluster 4 |
| HORVU.MOREX.r3.7HG0687540.1 | -0.113748 | -0.311866 | -0.2900132 | cluster 4 |
| HORVU.MOREX.r3.7HG0693590.1 | -0.197214 | -0.409448 | -0.3367645 | cluster 4 |
| HORVU.MOREX.r3.7HG0696770.1 | -0.133347 | -0.329911 | -0.3385414 | cluster 4 |
| HORVU.MOREX.r3.7HG0700270.1 | -0.143956 | -0.39241 | -0.4245597 | cluster 4 |
| HORVU.MOREX.r3.7HG0710170.1 | -0.34057 | -2.238231 | -1.7983181 | cluster 4 |
| HORVU.MOREX.r3.7HG0717850.1 | -0.119137 | -0.348577 | -0.3554462 | cluster 4 |
| HORVU.MOREX.r3.7HG0719830.1 | -0.75286 | -1.733023 | -1.4829856 | cluster 4 |
| HORVU.MOREX.r3.7HG0722380.1 | -0.149099 | -0.503357 | -0.3453409 | cluster 4 |
| HORVU.MOREX.r3.7HG0725290.1 | -0.005522 | -0.60065 | -0.47921 | cluster 4 |
| HORVU.MOREX.r3.7HG0728870.1 | -0.200143 | -0.675879 | -0.5988388 | cluster 4 |
| HORVU.MOREX.r3.7HG0731320.1 | -0.118784 | -0.341523 | -0.2786513 | cluster 4 |
| HORVU.MOREX.r3.7HG0734380.1 | -0.178956 | -0.455402 | -0.4756674 | cluster 4 |
| HORVU.MOREX.r3.7HG0739180.1 | -0.138186 | -0.30909 | -0.2397068 | cluster 4 |
| HORVU.MOREX.r3.7HG0752940.1 | -0.061403 | -0.324838 | -0.196665 | cluster 4 |
| HORVU.MOREX.r3.1HG0003820.1 | 0.1506945 | 0.1903632 | 0.34939377 | cluster 5 |
| HORVU.MOREX.r3.1HG0008510.2 | 0.1315153 | 0.2016694 | 0.54376722 | cluster 5 |
| HORVU.MOREX.r3.1HG0008640.1 | 0.3204557 | 0.345242 | 0.48963061 | cluster 5 |
| HORVU.MOREX.r3.1HG0016710.2 | 1.1890242 | 1.3376758 | 1.75589204 | cluster 5 |
| HORVU.MOREX.r3.1HG0020980.1 | 0.2916687 | 0.3662362 | 0.54445194 | cluster 5 |
| HORVU.MOREX.r3.1HG0025870.1 | 0.5108263 | 0.5636698 | 0.75828619 | cluster 5 |
| HORVU.MOREX.r3.1HG0042440.1 | 1.26164 | 1.2501095 | 1.3921203 | cluster 5 |
| HORVU.MOREX.r3.1HG0043640.1 | 0.1201616 | 0.2422031 | 0.42338664 | cluster 5 |
| HORVU.MOREX.r3.1HG0045770.1 | 1.5447 | 1.7458726 | 2.12094315 | cluster 5 |
| HORVU.MOREX.r3.1HG0049550.1 | 0.1576196 | 0.1479541 | 0.39915593 | cluster 5 |
| HORVU.MOREX.r3.1HG0057610.1 | 0.5288138 | 0.5805584 | 0.69813619 | cluster 5 |
| HORVU.MOREX.r3.1HG0059850.1 | 0.1454644 | 0.2951392 | 0.52413389 | cluster 5 |
| HORVU.MOREX.r3.1HG0064390.1 | 0.1071974 | 0.1965282 | 0.38205732 | cluster 5 |
| HORVU.MOREX.r3.1HG0067720.1 | 1.2160888 | 1.3105495 | 1.65738455 | cluster 5 |
| HORVU.MOREX.r3.1HG0069870.1 | 0.1767651 | 0.3193547 | 0.68052598 | cluster 5 |
| HORVU.MOREX.r3.1HG0070480.1 | 0.0244839 | 0.1380118 | 0.32689078 | cluster 5 |
| HORVU.MOREX.r3.1HG0075650.1 | 0.3233917 | 0.6804061 | 1.18634955 | cluster 5 |
| HORVU.MOREX.r3.1HG0079130.1 | 0.123639 | 0.1546145 | 0.36243892 | cluster 5 |
| HORVU.MOREX.r3.1HG0079140.1 | 0.2935846 | 0.4445291 | 0.69272931 | cluster 5 |
| HORVU.MOREX.r3.1HG0079950.1 | 0.0858294 | 0.1651136 | 0.31117409 | cluster 5 |
| HORVU.MOREX.r3.1HG0082550.1 | 0.8194643 | 1.0342348 | 1.50018093 | cluster 5 |
| HORVU.MOREX.r3.1HG0084890.1 | 0.3478786 | 0.34503 | 0.61071091 | cluster 5 |
| HORVU.MOREX.r3.1HG0092880.1 | 0.4749795 | 0.5257856 | 0.59948098 | cluster 5 |
| HORVU.MOREX.r3.2HG0097010.1 | 0.0877437 | 0.1815887 | 0.39799786 | cluster 5 |

**Table S3** Continued.

| ID | logFC_3 h | logFC_6 h | logFC_12 h | cluster No. |
|---|---|---|---|---|
| HORVU.MOREX.r3.2HG0099010.1 | 0.5486596 | 0.6936151 | 1.17562956 | cluster 5 |
| HORVU.MOREX.r3.2HG0099160.1 | 0.6424212 | 0.686862 | 0.98464269 | cluster 5 |
| HORVU.MOREX.r3.2HG0100360.1 | 0.125064 | 0.1270412 | 0.39321359 | cluster 5 |
| HORVU.MOREX.r3.2HG0100750.1 | 1.2075072 | 1.4933226 | 2.00037087 | cluster 5 |
| HORVU.MOREX.r3.2HG0105050.1 | 0.8090145 | 1.1605794 | 1.85682601 | cluster 5 |
| HORVU.MOREX.r3.2HG0111570.1 | -0.033407 | 0.1424131 | 0.73615542 | cluster 5 |
| HORVU.MOREX.r3.2HG0113660.1 | 0.1554899 | 0.2139374 | 0.35324822 | cluster 5 |
| HORVU.MOREX.r3.2HG0117030.1 | 0.5822317 | 0.6240572 | 0.72265079 | cluster 5 |
| HORVU.MOREX.r3.2HG0122080.1 | 0.2666389 | 0.3454356 | 0.59234208 | cluster 5 |
| HORVU.MOREX.r3.2HG0126380.1 | 0.2457761 | 0.3007804 | 0.47791978 | cluster 5 |
| HORVU.MOREX.r3.2HG0126440.1 | 0.3812549 | 0.6296109 | 1.16229679 | cluster 5 |
| HORVU.MOREX.r3.2HG0131280.1 | -0.04178 | 0.3444847 | 0.89225606 | cluster 5 |
| HORVU.MOREX.r3.2HG0135650.1 | 1.2476173 | 1.5945768 | 2.770416 | cluster 5 |
| HORVU.MOREX.r3.2HG0137620.1 | 0.1669473 | 0.2071834 | 0.47432716 | cluster 5 |
| HORVU.MOREX.r3.2HG0142740.1 | 0.1788709 | 0.2082363 | 0.36584606 | cluster 5 |
| HORVU.MOREX.r3.2HG0153420.1 | 0.5183998 | 0.7413509 | 1.11738569 | cluster 5 |
| HORVU.MOREX.r3.2HG0155290.1 | 0.4689755 | 0.8772635 | 1.482062 | cluster 5 |
| HORVU.MOREX.r3.2HG0157260.1 | 0.8422643 | 1.0459764 | 1.44511468 | cluster 5 |
| HORVU.MOREX.r3.2HG0159580.1 | 0.2014004 | 0.3455338 | 0.65480766 | cluster 5 |
| HORVU.MOREX.r3.2HG0160130.1 | 0.5326993 | 0.5959193 | 0.79332848 | cluster 5 |
| HORVU.MOREX.r3.2HG0160760.1 | 0.6251605 | 0.6294764 | 0.66585156 | cluster 5 |
| HORVU.MOREX.r3.2HG0165200.1 | 0.334928 | 0.4551799 | 0.62889149 | cluster 5 |
| HORVU.MOREX.r3.2HG0172580.1 | 1.1213565 | 1.2401841 | 1.65259497 | cluster 5 |
| HORVU.MOREX.r3.2HG0173380.1 | 0.0191126 | 0.133931 | 0.41562246 | cluster 5 |
| HORVU.MOREX.r3.2HG0177830.1 | 0.2974003 | 0.4238256 | 0.62031522 | cluster 5 |
| HORVU.MOREX.r3.2HG0186980.1 | 0.4348704 | 0.7049726 | 1.09965889 | cluster 5 |
| HORVU.MOREX.r3.2HG0189110.1 | 0.8017171 | 0.8633209 | 1.06107648 | cluster 5 |
| HORVU.MOREX.r3.2HG0190620.1 | 0.7019473 | 0.6991351 | 0.95931523 | cluster 5 |
| HORVU.MOREX.r3.2HG0191510.1 | 0.5490179 | 0.5822354 | 0.84188673 | cluster 5 |
| HORVU.MOREX.r3.2HG0196630.1 | 0.1987878 | 0.2309701 | 0.96600749 | cluster 5 |
| HORVU.MOREX.r3.2HG0197260.1 | 0.0062804 | 0.1096589 | 0.2688385 | cluster 5 |
| HORVU.MOREX.r3.2HG0199330.1 | 0.2860455 | 0.3700994 | 0.62871499 | cluster 5 |
| HORVU.MOREX.r3.2HG0204020.1 | 0.2760942 | 0.4700435 | 0.7417162 | cluster 5 |
| HORVU.MOREX.r3.2HG0207790.1 | 1.0148165 | 1.1822833 | 1.61922448 | cluster 5 |
| HORVU.MOREX.r3.2HG0213430.1 | 0.5039028 | 0.544796 | 0.76088997 | cluster 5 |
| HORVU.MOREX.r3.3HG0218560.1 | 0.872263 | 1.0259185 | 1.41284311 | cluster 5 |
| HORVU.MOREX.r3.3HG0226110.1 | 0.5157394 | 0.7011145 | 1.86209217 | cluster 5 |
| HORVU.MOREX.r3.3HG0228660.1 | 0.0955271 | 0.1285591 | 0.33763937 | cluster 5 |
| HORVU.MOREX.r3.3HG0234210.1 | 0.5296158 | 0.5100511 | 0.98978648 | cluster 5 |
| HORVU.MOREX.r3.3HG0244240.1 | 0.8114471 | 0.8835885 | 1.22221518 | cluster 5 |
| HORVU.MOREX.r3.3HG0245400.1 | 0.2898759 | 0.4244746 | 0.61866946 | cluster 5 |
| HORVU.MOREX.r3.3HG0269960.1 | 0.2215019 | 0.3635156 | 0.56275935 | cluster 5 |
| HORVU.MOREX.r3.3HG0279110.1 | 0.9218836 | 1.296394 | 2.07713439 | cluster 5 |
| HORVU.MOREX.r3.3HG0280530.1 | 0.2267596 | 0.2403021 | 0.53280743 | cluster 5 |
| HORVU.MOREX.r3.3HG0283080.2 | 0.2793467 | 0.5406405 | 1.02444722 | cluster 5 |
| HORVU.MOREX.r3.3HG0284760.1 | 0.0894929 | 0.1566816 | 0.56866413 | cluster 5 |
| HORVU.MOREX.r3.3HG0289020.1 | 1.1492856 | 1.1769054 | 1.30557443 | cluster 5 |

**Table S3** Continued.

| ID | logFC_3 h | logFC_6 h | logFC_12 h | cluster No. |
|---|---|---|---|---|
| HORVU.MOREX.r3.3HG0291570.1 | 0.0639497 | 0.1001521 | 0.40453683 | cluster 5 |
| HORVU.MOREX.r3.3HG0291670.1 | 0.6984078 | 0.7129584 | 1.29457936 | cluster 5 |
| HORVU.MOREX.r3.3HG0295680.1 | 0.3017542 | 0.3989332 | 0.6162365 | cluster 5 |
| HORVU.MOREX.r3.3HG0296010.1 | 0.9557079 | 1.3367467 | 1.96877264 | cluster 5 |
| HORVU.MOREX.r3.3HG0297000.1 | 0.7808635 | 0.9095607 | 1.16065524 | cluster 5 |
| HORVU.MOREX.r3.3HG0299990.1 | 0.1441051 | 0.2615899 | 0.50713829 | cluster 5 |
| HORVU.MOREX.r3.3HG0301000.1 | 0.5097699 | 0.590378 | 0.91653442 | cluster 5 |
| HORVU.MOREX.r3.3HG0304660.1 | 0.0208108 | -0.032096 | 0.41942742 | cluster 5 |
| HORVU.MOREX.r3.3HG0305600.1 | 0.0859495 | 0.1762314 | 0.40739692 | cluster 5 |
| HORVU.MOREX.r3.3HG0309900.1 | 1.0254695 | 1.1486088 | 1.46368758 | cluster 5 |
| HORVU.MOREX.r3.3HG0316870.1 | 0.0681683 | 0.1493181 | 0.36697207 | cluster 5 |
| HORVU.MOREX.r3.3HG0320350.1 | 1.4726565 | 1.5749652 | 1.85992323 | cluster 5 |
| HORVU.MOREX.r3.3HG0320520.1 | 0.4376358 | 0.4291602 | 0.68054367 | cluster 5 |
| HORVU.MOREX.r3.3HG0324480.1 | 0.4735448 | 0.6441702 | 1.35677217 | cluster 5 |
| HORVU.MOREX.r3.4HG0333540.1 | 0.6643687 | 0.7770503 | 0.94212038 | cluster 5 |
| HORVU.MOREX.r3.4HG0333550.2 | 0.195438 | 0.3645787 | 0.70912011 | cluster 5 |
| HORVU.MOREX.r3.4HG0340650.1 | 0.2275561 | 0.2445699 | 0.39966237 | cluster 5 |
| HORVU.MOREX.r3.4HG0349980.1 | 0.1567483 | 0.2679095 | 0.61503478 | cluster 5 |
| HORVU.MOREX.r3.4HG0351440.1 | 0.2172212 | 0.2217873 | 0.46079846 | cluster 5 |
| HORVU.MOREX.r3.4HG0353200.1 | 0.1292698 | 0.223277 | 0.52468811 | cluster 5 |
| HORVU.MOREX.r3.4HG0353950.1 | 0.5221309 | 0.5084894 | 2.94704517 | cluster 5 |
| HORVU.MOREX.r3.4HG0358980.1 | 0.1653312 | 0.1892386 | 0.50206292 | cluster 5 |
| HORVU.MOREX.r3.4HG0378330.1 | 0.6500979 | 0.8954483 | 1.26116524 | cluster 5 |
| HORVU.MOREX.r3.4HG0384350.1 | -0.004781 | 0.0987288 | 0.37871172 | cluster 5 |
| HORVU.MOREX.r3.4HG0385240.1 | 0.2914315 | 0.411096 | 0.62102358 | cluster 5 |
| HORVU.MOREX.r3.4HG0392690.1 | 0.3101105 | 0.3534756 | 0.44460083 | cluster 5 |
| HORVU.MOREX.r3.4HG0394830.1 | 0.6965264 | 0.824201 | 1.25434005 | cluster 5 |
| HORVU.MOREX.r3.4HG0397570.1 | 0.3785475 | 0.5006545 | 0.75817975 | cluster 5 |
| HORVU.MOREX.r3.4HG0398980.1 | 0.0802368 | 0.1704924 | 0.35305742 | cluster 5 |
| HORVU.MOREX.r3.4HG0401130.1 | 0.1270489 | 0.2114731 | 0.36245202 | cluster 5 |
| HORVU.MOREX.r3.4HG0403260.1 | 0.1030162 | 0.0944493 | 0.3463164 | cluster 5 |
| HORVU.MOREX.r3.4HG0403630.1 | 0.8573751 | 1.0227619 | 1.56282623 | cluster 5 |
| HORVU.MOREX.r3.4HG0407560.1 | 0.1921403 | 0.2773518 | 0.41498208 | cluster 5 |
| HORVU.MOREX.r3.4HG0409600.1 | 0.1887469 | 0.2941607 | 0.48745659 | cluster 5 |
| HORVU.MOREX.r3.4HG0414000.1 | 0.1665393 | 0.3454709 | 0.76980864 | cluster 5 |
| HORVU.MOREX.r3.5HG0420410.1 | 0.2534881 | 0.3710897 | 0.5507145 | cluster 5 |
| HORVU.MOREX.r3.5HG0421310.1 | 0.6414658 | 0.7966063 | 1.07325559 | cluster 5 |
| HORVU.MOREX.r3.5HG0426210.3 | 0.4977726 | 0.6222315 | 0.86104193 | cluster 5 |
| HORVU.MOREX.r3.5HG0430460.1 | 0.1856268 | 0.236529 | 0.48946727 | cluster 5 |
| HORVU.MOREX.r3.5HG0432360.1 | 0.28872 | 0.3754987 | 0.63714401 | cluster 5 |
| HORVU.MOREX.r3.5HG0436900.1 | 0.1760451 | 0.1861287 | 0.36513154 | cluster 5 |
| HORVU.MOREX.r3.5HG0444270.1 | 0.1056411 | 0.2175851 | 0.44586786 | cluster 5 |
| HORVU.MOREX.r3.5HG0454720.1 | 0.2017966 | 0.3520529 | 0.5693287 | cluster 5 |
| HORVU.MOREX.r3.5HG0460160.1 | 0.1634637 | 0.2742783 | 0.43094171 | cluster 5 |
| HORVU.MOREX.r3.5HG0462450.1 | 0.7964124 | 0.9036124 | 1.30469581 | cluster 5 |
| HORVU.MOREX.r3.5HG0464630.1 | 0.3210834 | 0.4123414 | 0.56749817 | cluster 5 |
| HORVU.MOREX.r3.5HG0470940.1 | 0.1503364 | 0.2760358 | 0.50070992 | cluster 5 |

**Table S3** Continued.

| ID | logFC_3 h | logFC_6 h | logFC_12 h | cluster No. |
|---|---|---|---|---|
| HORVU.MOREX.r3.5HG0472140.1 | 0.4155581 | 0.4014265 | 0.69239096 | cluster 5 |
| HORVU.MOREX.r3.5HG0478610.1 | 0.0949356 | 0.179812 | 0.39951234 | cluster 5 |
| HORVU.MOREX.r3.5HG0485760.1 | 1.2127406 | 1.3978826 | 1.66332866 | cluster 5 |
| HORVU.MOREX.r3.5HG0487910.1 | 1.743065 | 1.7830562 | 2.00123739 | cluster 5 |
| HORVU.MOREX.r3.5HG0493890.2 | 0.2756003 | 0.2539389 | 0.44122832 | cluster 5 |
| HORVU.MOREX.r3.5HG0495640.1 | 1.541128 | 1.5710812 | 1.90941616 | cluster 5 |
| HORVU.MOREX.r3.5HG0498320.1 | 0.0654798 | 0.0814044 | 0.42141937 | cluster 5 |
| HORVU.MOREX.r3.5HG0506230.1 | 0.1985123 | 0.2556131 | 0.52424111 | cluster 5 |
| HORVU.MOREX.r3.5HG0509390.1 | 0.7920428 | 0.8589774 | 1.09567221 | cluster 5 |
| HORVU.MOREX.r3.5HG0510320.1 | 0.2486342 | 0.3375388 | 0.72221946 | cluster 5 |
| HORVU.MOREX.r3.5HG0510500.1 | 0.0186906 | 0.1532601 | 0.46359741 | cluster 5 |
| HORVU.MOREX.r3.5HG0513810.1 | 0.2686357 | 0.3272462 | 0.62971808 | cluster 5 |
| HORVU.MOREX.r3.5HG0514020.1 | 0.2824591 | 0.3358146 | 0.44111193 | cluster 5 |
| HORVU.MOREX.r3.5HG0515610.1 | 0.354598 | 0.6985583 | 1.23595498 | cluster 5 |
| HORVU.MOREX.r3.5HG0522090.1 | 0.1003735 | 0.1869424 | 0.60113722 | cluster 5 |
| HORVU.MOREX.r3.5HG0531450.1 | 0.5936271 | 0.5556235 | 0.91417864 | cluster 5 |
| HORVU.MOREX.r3.5HG0537780.1 | 0.121621 | 0.2363552 | 0.41488425 | cluster 5 |
| HORVU.MOREX.r3.6HG0545460.1 | 0.1884398 | 0.2478194 | 0.50599213 | cluster 5 |
| HORVU.MOREX.r3.6HG0549520.1 | 0.1886624 | 0.3124328 | 0.634171 | cluster 5 |
| HORVU.MOREX.r3.6HG0564690.1 | 0.1434775 | 0.2110247 | 0.41003985 | cluster 5 |
| HORVU.MOREX.r3.6HG0565800.1 | 0.5641555 | 0.5657139 | 0.77030214 | cluster 5 |
| HORVU.MOREX.r3.6HG0568390.1 | 0.1456539 | 0.2871736 | 0.653533 | cluster 5 |
| HORVU.MOREX.r3.6HG0569990.1 | 0.0073052 | 0.1058387 | 0.27427454 | cluster 5 |
| HORVU.MOREX.r3.6HG0582950.1 | 0.1647378 | 0.1551424 | 0.27946176 | cluster 5 |
| HORVU.MOREX.r3.6HG0587460.1 | 0.3697823 | 0.4815205 | 0.95382752 | cluster 5 |
| HORVU.MOREX.r3.6HG0597640.1 | 0.4515542 | 0.5103614 | 0.76651863 | cluster 5 |
| HORVU.MOREX.r3.6HG0597790.1 | 0.4176998 | 0.5353106 | 0.73827734 | cluster 5 |
| HORVU.MOREX.r3.6HG0598750.1 | 0.4075013 | 0.6800884 | 1.17151566 | cluster 5 |
| HORVU.MOREX.r3.6HG0599450.1 | 1.1916244 | 1.2732576 | 1.72370577 | cluster 5 |
| HORVU.MOREX.r3.6HG0605130.1 | 0.1834353 | 0.3140859 | 1.024442 | cluster 5 |
| HORVU.MOREX.r3.6HG0608180.1 | 0.6316125 | 0.7255867 | 1.10985309 | cluster 5 |
| HORVU.MOREX.r3.6HG0608390.1 | 0.016203 | 0.1408953 | 0.37418566 | cluster 5 |
| HORVU.MOREX.r3.6HG0614470.2 | 0.3174888 | 0.3683412 | 0.50449019 | cluster 5 |
| HORVU.MOREX.r3.6HG0617240.1 | 0.2407525 | 0.3667786 | 0.56049813 | cluster 5 |
| HORVU.MOREX.r3.6HG0627090.1 | 0.4150708 | 0.377575 | 0.66252627 | cluster 5 |
| HORVU.MOREX.r3.6HG0631400.1 | 0.4495172 | 0.7629271 | 1.21136897 | cluster 5 |
| HORVU.MOREX.r3.6HG0634200.1 | 0.739161 | 0.9506689 | 1.28357549 | cluster 5 |
| HORVU.MOREX.r3.7HG0636030.1 | 0.7435151 | 0.8391918 | 1.40370231 | cluster 5 |
| HORVU.MOREX.r3.7HG0644240.1 | 0.6288995 | 0.8654318 | 1.20747996 | cluster 5 |
| HORVU.MOREX.r3.7HG0648460.1 | 0.3065913 | 0.4263577 | 0.62082676 | cluster 5 |
| HORVU.MOREX.r3.7HG0653740.1 | 0.1315017 | 0.2266617 | 0.45924945 | cluster 5 |
| HORVU.MOREX.r3.7HG0661450.1 | 0.1414274 | 0.3598007 | 0.66913788 | cluster 5 |
| HORVU.MOREX.r3.7HG0667150.1 | 0.4012188 | 0.585758 | 0.91635962 | cluster 5 |
| HORVU.MOREX.r3.7HG0667730.1 | 0.0317561 | 0.0615784 | 0.45914324 | cluster 5 |
| HORVU.MOREX.r3.7HG0669580.1 | 0.2534228 | 0.4093086 | 0.71139983 | cluster 5 |
| HORVU.MOREX.r3.7HG0669750.1 | 0.2169147 | 0.197497 | 0.4081658 | cluster 5 |
| HORVU.MOREX.r3.7HG0672400.1 | 0.6814586 | 0.758207 | 1.07071728 | cluster 5 |

**Table S3** Continued.

| ID | logFC_3 h | logFC_6 h | logFC_12 h | cluster No. |
|---|---|---|---|---|
| HORVU.MOREX.r3.7HG0676610.1 | 0.0885464 | 0.1749045 | 0.37598996 | cluster 5 |
| HORVU.MOREX.r3.7HG0678940.1 | 0.1357469 | 0.2300726 | 0.36876502 | cluster 5 |
| HORVU.MOREX.r3.7HG0685350.1 | 0.3166707 | 0.4168246 | 0.60223185 | cluster 5 |
| HORVU.MOREX.r3.7HG0701640.1 | 0.3650906 | 0.5382216 | 0.80746439 | cluster 5 |
| HORVU.MOREX.r3.7HG0702050.1 | 0.104106 | 0.4031577 | 1.01586603 | cluster 5 |
| HORVU.MOREX.r3.7HG0702660.1 | 0.3179493 | 0.2673309 | 1.16124912 | cluster 5 |
| HORVU.MOREX.r3.7HG0704760.1 | 0.4325514 | 0.3761209 | 1.21235137 | cluster 5 |
| HORVU.MOREX.r3.7HG0716320.1 | 0.3082597 | 0.3601296 | 0.53056742 | cluster 5 |
| HORVU.MOREX.r3.7HG0729150.1 | 0.6630894 | 0.8021488 | 1.26014855 | cluster 5 |
| HORVU.MOREX.r3.7HG0745120.1 | 0.2089014 | 0.2337941 | 0.35355569 | cluster 5 |
| HORVU.MOREX.r3.7HG0751030.1 | 0.4365522 | 0.7805611 | 1.31887325 | cluster 5 |
| HORVU.MOREX.r3.1HG0003150.1 | -0.210376 | -0.205283 | -0.6263923 | cluster 6 |
| HORVU.MOREX.r3.1HG0003280.1 | -1.465937 | -1.082383 | -2.2260952 | cluster 6 |
| HORVU.MOREX.r3.1HG0005840.1 | -0.519881 | -0.266103 | -1.4218761 | cluster 6 |
| HORVU.MOREX.r3.1HG0007570.1 | -0.639076 | -0.266017 | -1.3606559 | cluster 6 |
| HORVU.MOREX.r3.1HG0007930.1 | -0.173029 | -0.174301 | -0.766703 | cluster 6 |
| HORVU.MOREX.r3.1HG0018430.1 | -0.46222 | -0.441472 | -1.0676984 | cluster 6 |
| HORVU.MOREX.r3.1HG0037920.1 | -1.074663 | -0.740293 | -1.9231614 | cluster 6 |
| HORVU.MOREX.r3.1HG0041250.1 | -0.092455 | 0.0016478 | -0.3701922 | cluster 6 |
| HORVU.MOREX.r3.1HG0043350.1 | -0.085053 | -0.035176 | -0.4948174 | cluster 6 |
| HORVU.MOREX.r3.1HG0045490.1 | -0.211641 | -0.189946 | -0.4149042 | cluster 6 |
| HORVU.MOREX.r3.1HG0053530.1 | -0.199408 | -0.195196 | -1.4522622 | cluster 6 |
| HORVU.MOREX.r3.1HG0053720.1 | -0.425146 | -0.297207 | -0.9030535 | cluster 6 |
| HORVU.MOREX.r3.1HG0059870.1 | -0.132292 | -0.126445 | -0.3775719 | cluster 6 |
| HORVU.MOREX.r3.1HG0063000.1 | -0.086115 | -0.078976 | -0.4382613 | cluster 6 |
| HORVU.MOREX.r3.1HG0064610.1 | -0.190379 | -0.070031 | -0.9729466 | cluster 6 |
| HORVU.MOREX.r3.1HG0069110.1 | -0.157536 | -0.140838 | -0.4783552 | cluster 6 |
| HORVU.MOREX.r3.1HG0074650.2 | -0.510431 | -0.168715 | -1.0294116 | cluster 6 |
| HORVU.MOREX.r3.1HG0079810.1 | -0.278065 | -0.161205 | -0.5402787 | cluster 6 |
| HORVU.MOREX.r3.1HG0082750.1 | -0.738823 | -0.507772 | -1.8998339 | cluster 6 |
| HORVU.MOREX.r3.1HG0083980.1 | -0.275479 | -0.275687 | -0.9930048 | cluster 6 |
| HORVU.MOREX.r3.1HG0095000.1 | -0.565694 | -0.539614 | -1.5807766 | cluster 6 |
| HORVU.MOREX.r3.2HG0097350.1 | -1.005101 | -0.924292 | -2.0949814 | cluster 6 |
| HORVU.MOREX.r3.2HG0108490.1 | -0.219808 | -0.132454 | -1.2703107 | cluster 6 |
| HORVU.MOREX.r3.2HG0111270.1 | 0.8853755 | 1.1046113 | 0.53938805 | cluster 6 |
| HORVU.MOREX.r3.2HG0117100.1 | 0.6704028 | 0.9147007 | 0.08654056 | cluster 6 |
| HORVU.MOREX.r3.2HG0119360.1 | -0.662395 | -0.383498 | -2.1030955 | cluster 6 |
| HORVU.MOREX.r3.2HG0120830.1 | 2.006445 | 2.4275847 | 1.35478833 | cluster 6 |
| HORVU.MOREX.r3.2HG0121530.1 | -0.586808 | -0.276879 | -1.0529693 | cluster 6 |
| HORVU.MOREX.r3.2HG0122720.1 | -0.417898 | -0.257733 | -0.9808534 | cluster 6 |
| HORVU.MOREX.r3.2HG0124470.1 | -0.872348 | -0.664108 | -2.277542 | cluster 6 |
| HORVU.MOREX.r3.2HG0134300.1 | 1.7005685 | 2.0693025 | 0.24289318 | cluster 6 |
| HORVU.MOREX.r3.2HG0135860.1 | -0.237153 | -0.166929 | -0.3859966 | cluster 6 |
| HORVU.MOREX.r3.2HG0136010.1 | -0.295066 | -0.173897 | -0.7129157 | cluster 6 |
| HORVU.MOREX.r3.2HG0142490.1 | -0.085467 | -0.058264 | -0.2785315 | cluster 6 |
| HORVU.MOREX.r3.2HG0148170.1 | -0.812555 | -0.597135 | -1.3169603 | cluster 6 |
| HORVU.MOREX.r3.2HG0154500.1 | -0.610336 | -0.59653 | -1.0398242 | cluster 6 |

**Table S3** Continued.

| ID | logFC_3 h | logFC_6 h | logFC_12 h | cluster No. |
|---|---|---|---|---|
| HORVU.MOREX.r3.2HG0167540.1 | -0.172394 | -0.165011 | -0.2223481 | cluster 6 |
| HORVU.MOREX.r3.2HG0171100.1 | -0.332299 | -0.317458 | -0.4414421 | cluster 6 |
| HORVU.MOREX.r3.2HG0171850.1 | -0.255421 | -0.146486 | -0.4887234 | cluster 6 |
| HORVU.MOREX.r3.2HG0181550.1 | 0.0751298 | 0.1297928 | -1.0106527 | cluster 6 |
| HORVU.MOREX.r3.2HG0182010.1 | -0.476485 | -0.414783 | -0.8520016 | cluster 6 |
| HORVU.MOREX.r3.2HG0189970.1 | -0.301055 | -0.147908 | -0.7152154 | cluster 6 |
| HORVU.MOREX.r3.2HG0190260.1 | 1.0366098 | 1.0486261 | 0.84157807 | cluster 6 |
| HORVU.MOREX.r3.2HG0196960.1 | -0.967413 | -0.511161 | -2.6121341 | cluster 6 |
| HORVU.MOREX.r3.2HG0201250.1 | -0.272512 | -0.199863 | -0.5651718 | cluster 6 |
| HORVU.MOREX.r3.2HG0205680.1 | -0.035521 | 0.1972397 | -1.9432136 | cluster 6 |
| HORVU.MOREX.r3.2HG0205760.1 | -0.361271 | -0.366731 | -0.9134359 | cluster 6 |
| HORVU.MOREX.r3.2HG0208280.1 | -0.25249 | -0.234284 | -0.4920429 | cluster 6 |
| HORVU.MOREX.r3.2HG0209030.1 | -0.117056 | -0.088912 | -1.4886989 | cluster 6 |
| HORVU.MOREX.r3.2HG0209090.1 | -0.222263 | -0.15666 | -0.6005596 | cluster 6 |
| HORVU.MOREX.r3.2HG0215060.1 | 2.4857767 | 2.5684304 | 2.22913008 | cluster 6 |
| HORVU.MOREX.r3.3HG0225640.1 | -0.421046 | -0.117874 | -1.2981488 | cluster 6 |
| HORVU.MOREX.r3.3HG0231880.1 | -0.614058 | -0.576063 | -1.0812871 | cluster 6 |
| HORVU.MOREX.r3.3HG0236000.1 | -0.195667 | -0.189929 | -0.4465279 | cluster 6 |
| HORVU.MOREX.r3.3HG0243560.1 | -0.134212 | -0.115022 | -0.4068952 | cluster 6 |
| HORVU.MOREX.r3.3HG0243920.1 | -0.125523 | -0.023681 | -1.1859745 | cluster 6 |
| HORVU.MOREX.r3.3HG0244010.1 | -0.067479 | -0.051859 | -0.6610377 | cluster 6 |
| HORVU.MOREX.r3.3HG0245320.1 | -0.15354 | -0.137279 | -0.5011831 | cluster 6 |
| HORVU.MOREX.r3.3HG0253630.1 | 1.1990125 | 1.4139342 | 0.94401369 | cluster 6 |
| HORVU.MOREX.r3.3HG0277330.1 | -0.168801 | -0.172149 | -0.6224506 | cluster 6 |
| HORVU.MOREX.r3.3HG0281860.1 | 0.0321565 | 0.0784197 | -0.5930055 | cluster 6 |
| HORVU.MOREX.r3.3HG0289070.1 | 0.0018635 | -0.00883 | -1.2054247 | cluster 6 |
| HORVU.MOREX.r3.3HG0291340.1 | -0.606158 | -0.546952 | -0.9681079 | cluster 6 |
| HORVU.MOREX.r3.3HG0300810.1 | -0.275664 | -0.221569 | -0.4895947 | cluster 6 |
| HORVU.MOREX.r3.3HG0308980.1 | -0.262001 | -0.253071 | -0.6935355 | cluster 6 |
| HORVU.MOREX.r3.3HG0319570.1 | -0.360375 | -0.351079 | -1.1568333 | cluster 6 |
| HORVU.MOREX.r3.3HG0323810.1 | -0.168794 | -0.15391 | -0.4772449 | cluster 6 |
| HORVU.MOREX.r3.3HG0329510.1 | -0.404317 | -0.279715 | -0.9626113 | cluster 6 |
| HORVU.MOREX.r3.4HG0335450.1 | 2.7351032 | 2.8494273 | 1.80351533 | cluster 6 |
| HORVU.MOREX.r3.4HG0345810.1 | -0.516735 | -0.363656 | -1.2604258 | cluster 6 |
| HORVU.MOREX.r3.4HG0362350.1 | -0.389043 | -0.258159 | -1.2144725 | cluster 6 |
| HORVU.MOREX.r3.4HG0364210.1 | 2.183526 | 2.5067424 | 0.99266889 | cluster 6 |
| HORVU.MOREX.r3.4HG0364880.1 | -1.012113 | -0.854058 | -1.255772 | cluster 6 |
| HORVU.MOREX.r3.4HG0370520.1 | -0.271313 | -0.260419 | -0.7428434 | cluster 6 |
| HORVU.MOREX.r3.4HG0375440.1 | -0.496671 | -0.138871 | -1.167625 | cluster 6 |
| HORVU.MOREX.r3.4HG0388470.1 | -0.113298 | 0.0635349 | -2.2735105 | cluster 6 |
| HORVU.MOREX.r3.4HG0395130.1 | -0.570158 | 0.1452184 | -1.6829943 | cluster 6 |
| HORVU.MOREX.r3.4HG0405830.1 | -0.26024 | -0.129966 | -0.7898168 | cluster 6 |
| HORVU.MOREX.r3.4HG0405990.1 | 1.2352983 | 1.4902374 | 0.90794655 | cluster 6 |
| HORVU.MOREX.r3.4HG0412730.1 | -0.087227 | 0.1044175 | -0.422168 | cluster 6 |
| HORVU.MOREX.r3.4HG0417660.1 | 2.5850014 | 2.7906547 | 1.85509947 | cluster 6 |
| HORVU.MOREX.r3.5HG0420670.1 | -0.30801 | -0.144831 | -0.7737035 | cluster 6 |
| HORVU.MOREX.r3.5HG0439090.1 | -0.370212 | -0.360967 | -0.727011 | cluster 6 |

**Table S3** Continued.

| ID | logFC_3 h | logFC_6 h | logFC_12 h | cluster No. |
|---|---|---|---|---|
| HORVU.MOREX.r3.5HG0455080.1 | -0.482031 | -0.088085 | -0.957045 | cluster 6 |
| HORVU.MOREX.r3.5HG0459600.1 | 0.6656884 | 0.6719967 | 0.63974505 | cluster 6 |
| HORVU.MOREX.r3.5HG0460560.1 | -0.601994 | -0.21141 | -2.3379839 | cluster 6 |
| HORVU.MOREX.r3.5HG0465810.1 | 2.4748372 | 2.5847967 | 2.27103236 | cluster 6 |
| HORVU.MOREX.r3.5HG0466020.1 | -0.005802 | 0.0216804 | -0.7626933 | cluster 6 |
| HORVU.MOREX.r3.5HG0476390.1 | -0.439241 | -0.367227 | -0.85495 | cluster 6 |
| HORVU.MOREX.r3.5HG0480060.1 | -0.439933 | -0.412945 | -1.1842657 | cluster 6 |
| HORVU.MOREX.r3.5HG0483880.1 | -0.573398 | -0.364214 | -1.6012317 | cluster 6 |
| HORVU.MOREX.r3.5HG0489840.1 | -0.47416 | -0.381838 | -1.1150172 | cluster 6 |
| HORVU.MOREX.r3.5HG0490960.1 | -0.437415 | -0.353481 | -0.643194 | cluster 6 |
| HORVU.MOREX.r3.5HG0491230.1 | -0.457861 | -0.347093 | -0.9233452 | cluster 6 |
| HORVU.MOREX.r3.5HG0504700.1 | -0.590943 | -0.362548 | -0.9160539 | cluster 6 |
| HORVU.MOREX.r3.5HG0506290.1 | -0.653223 | -0.484077 | -0.8812599 | cluster 6 |
| HORVU.MOREX.r3.5HG0510370.1 | -0.13359 | -0.125914 | -0.3018527 | cluster 6 |
| HORVU.MOREX.r3.5HG0512350.1 | -0.509796 | -0.512041 | -1.1280956 | cluster 6 |
| HORVU.MOREX.r3.5HG0530240.1 | -0.499727 | -0.448928 | -1.1496825 | cluster 6 |
| HORVU.MOREX.r3.5HG0535780.1 | -0.242318 | -0.180728 | -0.7112602 | cluster 6 |
| HORVU.MOREX.r3.6HG0539410.1 | -0.802439 | -0.71711 | -1.7411792 | cluster 6 |
| HORVU.MOREX.r3.6HG0553310.1 | -0.36968 | -0.189265 | -0.7856658 | cluster 6 |
| HORVU.MOREX.r3.6HG0553580.1 | 1.1866115 | 1.274208 | 0.98142484 | cluster 6 |
| HORVU.MOREX.r3.6HG0555810.1 | -0.682413 | -0.607668 | -1.2217529 | cluster 6 |
| HORVU.MOREX.r3.6HG0566930.1 | -0.22637 | 0.0490535 | -0.7599335 | cluster 6 |
| HORVU.MOREX.r3.6HG0569480.1 | -0.379795 | -0.23017 | -2.5324008 | cluster 6 |
| HORVU.MOREX.r3.6HG0583730.1 | -0.169149 | -0.166424 | -0.3843065 | cluster 6 |
| HORVU.MOREX.r3.6HG0586570.1 | -0.066396 | 0.1873834 | -0.8643434 | cluster 6 |
| HORVU.MOREX.r3.6HG0594290.1 | -0.37741 | -0.143018 | -1.0226001 | cluster 6 |
| HORVU.MOREX.r3.6HG0596240.1 | -0.508059 | -0.427754 | -1.1653371 | cluster 6 |
| HORVU.MOREX.r3.6HG0597460.1 | -0.531191 | 0.0172299 | -1.840106 | cluster 6 |
| HORVU.MOREX.r3.6HG0604310.1 | -0.588063 | -0.520915 | -1.1290382 | cluster 6 |
| HORVU.MOREX.r3.6HG0606940.1 | -0.438108 | -0.176777 | -3.0814176 | cluster 6 |
| HORVU.MOREX.r3.6HG0622980.1 | -0.072754 | -0.050755 | -0.3923796 | cluster 6 |
| HORVU.MOREX.r3.6HG0630650.1 | -0.097713 | 0.2318769 | -1.690516 | cluster 6 |
| HORVU.MOREX.r3.7HG0635160.1 | -0.068051 | -0.028414 | -0.3964848 | cluster 6 |
| HORVU.MOREX.r3.7HG0644540.1 | 0.6256665 | 0.7559992 | 0.30281531 | cluster 6 |
| HORVU.MOREX.r3.7HG0655260.1 | -0.312909 | -0.293031 | -0.9635639 | cluster 6 |
| HORVU.MOREX.r3.7HG0656330.1 | -1.163553 | -0.904951 | -1.4923467 | cluster 6 |
| HORVU.MOREX.r3.7HG0656420.1 | -0.123269 | 0.128882 | -1.063885 | cluster 6 |
| HORVU.MOREX.r3.7HG0656440.1 | -0.027168 | 0.0020899 | -0.4591596 | cluster 6 |
| HORVU.MOREX.r3.7HG0663820.1 | -0.363469 | -0.315317 | -0.8339684 | cluster 6 |
| HORVU.MOREX.r3.7HG0668450.1 | -0.279607 | -0.259426 | -0.4965059 | cluster 6 |
| HORVU.MOREX.r3.7HG0668560.1 | 0.8654427 | 0.9760688 | 0.58199648 | cluster 6 |
| HORVU.MOREX.r3.7HG0676880.1 | -0.200974 | -0.159506 | -0.3829243 | cluster 6 |
| HORVU.MOREX.r3.7HG0688770.1 | 0.002493 | 0.0578383 | -0.688143 | cluster 6 |
| HORVU.MOREX.r3.7HG0689580.1 | -0.242788 | -0.237787 | -0.6068275 | cluster 6 |
| HORVU.MOREX.r3.7HG0692070.1 | -0.30171 | -0.181181 | -0.7256002 | cluster 6 |
| HORVU.MOREX.r3.7HG0697940.1 | 0.4916888 | 0.5754474 | -1.3457295 | cluster 6 |
| HORVU.MOREX.r3.7HG0702190.1 | -0.113268 | -0.119335 | -1.6389848 | cluster 6 |

**Table S3** Continued.

| ID | logFC_3 h | logFC_6 h | logFC_12 h | cluster No. |
|---|---|---|---|---|
| HORVU.MOREX.r3.7HG0704060.1 | 0.3659122 | 0.5559675 | 0.0679357 | cluster 6 |
| HORVU.MOREX.r3.7HG0708050.1 | -1.159545 | -0.485533 | -2.6356189 | cluster 6 |
| HORVU.MOREX.r3.7HG0709720.1 | -0.045605 | -0.048208 | -0.4654158 | cluster 6 |
| HORVU.MOREX.r3.7HG0714900.1 | -0.461434 | -0.307175 | -0.7230933 | cluster 6 |
| HORVU.MOREX.r3.7HG0720720.1 | -0.327706 | -0.213257 | -0.4870422 | cluster 6 |
| HORVU.MOREX.r3.7HG0723240.1 | 3.3718803 | 3.5954892 | 3.03861644 | cluster 6 |
| HORVU.MOREX.r3.7HG0736860.1 | -0.454611 | -0.321158 | -2.1718564 | cluster 6 |
| HORVU.MOREX.r3.7HG0737960.2 | -0.191814 | -0.136333 | -0.6723185 | cluster 6 |
| HORVU.MOREX.r3.1HG0030290.1 | 3.2160988 | 2.9900065 | 3.26082502 | cluster 7 |
| HORVU.MOREX.r3.1HG0042120.1 | -0.15605 | -0.470082 | -0.2554328 | cluster 7 |
| HORVU.MOREX.r3.1HG0042870.1 | -0.170446 | -0.246205 | -0.1303301 | cluster 7 |
| HORVU.MOREX.r3.1HG0051110.1 | -0.212994 | -0.429426 | -0.206373 | cluster 7 |
| HORVU.MOREX.r3.1HG0055320.1 | -0.133147 | -0.277538 | -0.1776111 | cluster 7 |
| HORVU.MOREX.r3.1HG0056510.1 | 0.0287858 | -1.460636 | -0.3552398 | cluster 7 |
| HORVU.MOREX.r3.1HG0058140.1 | -0.214192 | -0.407669 | -0.2639165 | cluster 7 |
| HORVU.MOREX.r3.1HG0062450.1 | -0.427538 | -0.820102 | -0.6131836 | cluster 7 |
| HORVU.MOREX.r3.1HG0063020.1 | 1.2227825 | 0.9115231 | 1.29517293 | cluster 7 |
| HORVU.MOREX.r3.1HG0083250.1 | -0.090481 | -0.792259 | -0.4232434 | cluster 7 |
| HORVU.MOREX.r3.2HG0105990.1 | -0.198432 | -0.412564 | -0.3015049 | cluster 7 |
| HORVU.MOREX.r3.2HG0109580.1 | -0.452354 | -1.391812 | -0.4661135 | cluster 7 |
| HORVU.MOREX.r3.2HG0115650.1 | -0.269629 | -1.041937 | -0.4347862 | cluster 7 |
| HORVU.MOREX.r3.2HG0126960.1 | -0.132265 | -0.56668 | -0.2388556 | cluster 7 |
| HORVU.MOREX.r3.2HG0137140.1 | -0.120656 | -1.13362 | -0.2842848 | cluster 7 |
| HORVU.MOREX.r3.2HG0157710.1 | -0.253123 | -0.85214 | -0.3936226 | cluster 7 |
| HORVU.MOREX.r3.2HG0158260.1 | -0.624144 | -0.99133 | -0.7315255 | cluster 7 |
| HORVU.MOREX.r3.2HG0162300.1 | -0.207332 | -0.431563 | -0.1364607 | cluster 7 |
| HORVU.MOREX.r3.2HG0168700.1 | -0.169481 | -1.93174 | -0.7636944 | cluster 7 |
| HORVU.MOREX.r3.2HG0175350.1 | 2.6333297 | 1.7344105 | 2.26098474 | cluster 7 |
| HORVU.MOREX.r3.2HG0181880.1 | -0.141264 | -0.348116 | -0.2230559 | cluster 7 |
| HORVU.MOREX.r3.2HG0190800.1 | 1.5459298 | 1.0077875 | 1.70810426 | cluster 7 |
| HORVU.MOREX.r3.2HG0193510.2 | -0.201709 | -0.360121 | -0.1759317 | cluster 7 |
| HORVU.MOREX.r3.2HG0195600.1 | -0.116421 | -0.358898 | -0.21699 | cluster 7 |
| HORVU.MOREX.r3.2HG0216470.1 | -0.261736 | -0.58313 | -0.3678178 | cluster 7 |
| HORVU.MOREX.r3.3HG0219860.1 | -0.160323 | -0.444479 | -0.248577 | cluster 7 |
| HORVU.MOREX.r3.3HG0228600.1 | -0.217675 | -0.838092 | -0.253968 | cluster 7 |
| HORVU.MOREX.r3.3HG0234470.1 | -0.244497 | -0.562767 | -0.2254953 | cluster 7 |
| HORVU.MOREX.r3.3HG0241320.1 | -0.235043 | -0.574987 | -0.3063886 | cluster 7 |
| HORVU.MOREX.r3.3HG0246550.1 | -0.378418 | -0.792749 | -0.3262336 | cluster 7 |
| HORVU.MOREX.r3.3HG0249430.1 | 0.4532392 | -2.481498 | -0.9514063 | cluster 7 |
| HORVU.MOREX.r3.3HG0257680.1 | -0.20108 | -0.506971 | -0.2343225 | cluster 7 |
| HORVU.MOREX.r3.3HG0258280.1 | -0.119148 | -0.287422 | -0.1490613 | cluster 7 |
| HORVU.MOREX.r3.3HG0264440.1 | -0.171005 | -0.653189 | -0.4036477 | cluster 7 |
| HORVU.MOREX.r3.3HG0267430.2 | -0.340256 | -0.706942 | -0.449371 | cluster 7 |
| HORVU.MOREX.r3.3HG0273730.1 | -0.147686 | -0.551093 | -0.2891259 | cluster 7 |
| HORVU.MOREX.r3.3HG0293010.1 | -0.212651 | -0.411838 | -0.3017727 | cluster 7 |
| HORVU.MOREX.r3.3HG0293420.1 | 1.1549496 | 0.9357247 | 1.06112938 | cluster 7 |
| HORVU.MOREX.r3.3HG0304230.1 | 2.0220451 | 1.5808034 | 2.17038351 | cluster 7 |

**Table S3** Continued.

| ID | logFC_3 h | logFC_6 h | logFC_12 h | cluster No. |
|---|---|---|---|---|
| HORVU.MOREX.r3.3HG0307310.1 | -0.067888 | -0.307881 | -0.1429791 | cluster 7 |
| HORVU.MOREX.r3.3HG0315130.1 | -0.43211 | -1.200566 | -0.5473714 | cluster 7 |
| HORVU.MOREX.r3.3HG0324070.1 | 1.6702737 | 1.4062936 | 1.76382212 | cluster 7 |
| HORVU.MOREX.r3.3HG0330040.1 | -0.120768 | -0.658763 | -0.3435413 | cluster 7 |
| HORVU.MOREX.r3.4HG0331520.1 | -0.187311 | -0.586644 | -0.2447804 | cluster 7 |
| HORVU.MOREX.r3.4HG0332930.1 | -0.822034 | -1.615903 | -1.0259526 | cluster 7 |
| HORVU.MOREX.r3.4HG0343710.1 | -0.220913 | -0.349295 | -0.1657513 | cluster 7 |
| HORVU.MOREX.r3.4HG0348450.1 | -0.717871 | -1.243245 | -0.8769814 | cluster 7 |
| HORVU.MOREX.r3.4HG0352760.1 | -0.168049 | -0.344307 | -0.2394104 | cluster 7 |
| HORVU.MOREX.r3.4HG0356090.1 | -0.295739 | -0.622584 | -0.4116669 | cluster 7 |
| HORVU.MOREX.r3.4HG0373800.1 | -0.458263 | -1.0656 | -0.7512242 | cluster 7 |
| HORVU.MOREX.r3.4HG0383400.1 | -0.18225 | -0.355746 | -0.231642 | cluster 7 |
| HORVU.MOREX.r3.4HG0389410.1 | 0.615519 | 0.2822746 | 0.75198528 | cluster 7 |
| HORVU.MOREX.r3.4HG0390140.1 | -0.132516 | -0.262139 | -0.1947101 | cluster 7 |
| HORVU.MOREX.r3.4HG0396760.1 | -0.106994 | -0.400321 | -0.2508712 | cluster 7 |
| HORVU.MOREX.r3.5HG0419520.1 | -0.060643 | -0.262126 | -0.1109363 | cluster 7 |
| HORVU.MOREX.r3.5HG0423790.1 | -0.169621 | -0.545146 | -0.1361613 | cluster 7 |
| HORVU.MOREX.r3.5HG0425880.1 | -0.186934 | -0.548493 | -0.2320833 | cluster 7 |
| HORVU.MOREX.r3.5HG0426280.1 | 0.895911 | 0.8026891 | 0.91336719 | cluster 7 |
| HORVU.MOREX.r3.5HG0430480.1 | -0.146778 | -0.272973 | -0.1370712 | cluster 7 |
| HORVU.MOREX.r3.5HG0438280.1 | -0.47725 | -1.805543 | -0.9756747 | cluster 7 |
| HORVU.MOREX.r3.5HG0439190.1 | -0.256617 | -0.507693 | -0.3533273 | cluster 7 |
| HORVU.MOREX.r3.5HG0441840.1 | 1.3633768 | 1.2209914 | 1.3181252 | cluster 7 |
| HORVU.MOREX.r3.5HG0448080.1 | -0.208832 | -0.556157 | -0.3113608 | cluster 7 |
| HORVU.MOREX.r3.5HG0473900.1 | 0.3363969 | 0.2043496 | 0.39301885 | cluster 7 |
| HORVU.MOREX.r3.5HG0474540.1 | -0.176849 | -0.363756 | -0.245905 | cluster 7 |
| HORVU.MOREX.r3.5HG0483860.1 | -0.114833 | -0.54771 | -0.3128479 | cluster 7 |
| HORVU.MOREX.r3.5HG0486810.1 | -0.377942 | -1.001511 | -0.4599156 | cluster 7 |
| HORVU.MOREX.r3.5HG0488950.1 | -0.530402 | -1.081169 | -0.7442717 | cluster 7 |
| HORVU.MOREX.r3.5HG0491250.1 | -0.056524 | -0.439289 | -0.2369434 | cluster 7 |
| HORVU.MOREX.r3.5HG0496980.1 | -0.132492 | -0.627109 | -0.2096685 | cluster 7 |
| HORVU.MOREX.r3.5HG0497990.1 | 1.0351763 | 0.7722732 | 1.11838961 | cluster 7 |
| HORVU.MOREX.r3.5HG0527280.1 | -0.064662 | -0.251611 | -0.1256452 | cluster 7 |
| HORVU.MOREX.r3.5HG0531390.1 | -0.134208 | -0.297476 | -0.1909864 | cluster 7 |
| HORVU.MOREX.r3.5HG0537000.1 | 1.0746695 | 0.9108017 | 1.00821353 | cluster 7 |
| HORVU.MOREX.r3.6HG0549990.1 | -0.134297 | -0.91081 | -0.41267 | cluster 7 |
| HORVU.MOREX.r3.6HG0552230.1 | -0.34495 | -0.528849 | -0.4279295 | cluster 7 |
| HORVU.MOREX.r3.6HG0567780.1 | -0.434157 | -0.731683 | -0.5097421 | cluster 7 |
| HORVU.MOREX.r3.6HG0569610.1 | -0.29027 | -0.723435 | -0.3262721 | cluster 7 |
| HORVU.MOREX.r3.6HG0605120.1 | -0.503633 | -0.941651 | -0.6142515 | cluster 7 |
| HORVU.MOREX.r3.6HG0616880.1 | -0.235822 | -1.496044 | -0.5642134 | cluster 7 |
| HORVU.MOREX.r3.6HG0617750.1 | -0.108153 | -0.585285 | -0.3302267 | cluster 7 |
| HORVU.MOREX.r3.6HG0628410.1 | -0.107487 | -0.322295 | -0.1185619 | cluster 7 |
| HORVU.MOREX.r3.7HG0652820.1 | -0.588486 | -2.251526 | -1.3184711 | cluster 7 |
| HORVU.MOREX.r3.7HG0667180.1 | -0.556496 | -0.798449 | -0.4581096 | cluster 7 |
| HORVU.MOREX.r3.7HG0671470.1 | -0.208542 | -0.924604 | -0.4782527 | cluster 7 |
| HORVU.MOREX.r3.7HG0674680.1 | 1.1070658 | 0.7065894 | 1.2363665 | cluster 7 |

**Table S3** Continued.

| ID | logFC_3 h | logFC_6 h | logFC_12 h | cluster No. |
|---|---|---|---|---|
| HORVU.MOREX.r3.7HG0675770.1 | -0.353457 | -0.599998 | -0.4555224 | cluster 7 |
| HORVU.MOREX.r3.7HG0680410.1 | -0.206222 | -0.647094 | -0.3041446 | cluster 7 |
| HORVU.MOREX.r3.7HG0711200.1 | -0.071671 | -0.380001 | -0.1717068 | cluster 7 |
| HORVU.MOREX.r3.7HG0714530.1 | -0.385011 | -1.361976 | -0.3521423 | cluster 7 |
| HORVU.MOREX.r3.7HG0721080.1 | -0.362552 | -0.888793 | -0.455568 | cluster 7 |
| HORVU.MOREX.r3.7HG0726170.1 | -0.107939 | -0.368825 | -0.2214472 | cluster 7 |
| HORVU.MOREX.r3.7HG0729510.1 | -0.38105 | -0.660762 | -0.4855394 | cluster 7 |
| HORVU.MOREX.r3.7HG0742720.1 | -0.204679 | -0.355489 | -0.2419012 | cluster 7 |
| HORVU.MOREX.r3.7HG0745360.1 | -0.120774 | -0.317561 | -0.1896039 | cluster 7 |
| HORVU.MOREX.r3.7HG0745700.1 | -0.252755 | -0.305338 | -0.2583258 | cluster 7 |
| HORVU.MOREX.r3.7HG0751060.1 | -0.049984 | -0.386301 | -0.1963032 | cluster 7 |
| HORVU.MOREX.r3.1HG0016560.1 | -0.68839 | 0.0457791 | -1.4256237 | cluster 8 |
| HORVU.MOREX.r3.1HG0033450.1 | -1.370905 | -0.973254 | -1.6061167 | cluster 8 |
| HORVU.MOREX.r3.1HG0047250.1 | -0.842719 | -0.335649 | -0.8617388 | cluster 8 |
| HORVU.MOREX.r3.1HG0052250.1 | 0.7624389 | 1.4813074 | 0.77029671 | cluster 8 |
| HORVU.MOREX.r3.1HG0055590.1 | 1.279564 | 1.4617378 | 1.22363957 | cluster 8 |
| HORVU.MOREX.r3.1HG0073460.1 | 0.2357812 | 0.4582906 | 0.25351407 | cluster 8 |
| HORVU.MOREX.r3.1HG0083550.1 | 1.005615 | 2.8081284 | 0.07977259 | cluster 8 |
| HORVU.MOREX.r3.1HG0083730.1 | 0.6875268 | 1.06344 | 0.69693972 | cluster 8 |
| HORVU.MOREX.r3.1HG0086480.1 | 0.9142496 | 1.3676222 | 0.79241127 | cluster 8 |
| HORVU.MOREX.r3.2HG0101220.2 | -0.421702 | -0.342134 | -0.4980625 | cluster 8 |
| HORVU.MOREX.r3.2HG0151790.1 | 0.0187827 | 2.5871455 | 0.07610652 | cluster 8 |
| HORVU.MOREX.r3.2HG0156140.1 | -1.001972 | -0.789672 | -1.146646 | cluster 8 |
| HORVU.MOREX.r3.2HG0165640.1 | -1.879812 | -1.097162 | -2.4990806 | cluster 8 |
| HORVU.MOREX.r3.2HG0170210.1 | 1.0146391 | 1.7031544 | 0.85067223 | cluster 8 |
| HORVU.MOREX.r3.2HG0189780.1 | 0.9533907 | 1.6835911 | 0.92413746 | cluster 8 |
| HORVU.MOREX.r3.2HG0203080.1 | 1.6417226 | 2.0966103 | 1.67264528 | cluster 8 |
| HORVU.MOREX.r3.2HG0206920.1 | 0.7491963 | 0.9379036 | 0.73132938 | cluster 8 |
| HORVU.MOREX.r3.2HG0210650.1 | 1.4265372 | 1.7254115 | 1.43270808 | cluster 8 |
| HORVU.MOREX.r3.2HG0212990.1 | -0.435296 | -0.136573 | -0.6607481 | cluster 8 |
| HORVU.MOREX.r3.2HG0213020.1 | 0.8360542 | 1.6698752 | 0.89323859 | cluster 8 |
| HORVU.MOREX.r3.2HG0213840.1 | 0.7932587 | 1.0372114 | 0.79750973 | cluster 8 |
| HORVU.MOREX.r3.3HG0256860.1 | 1.0448509 | 2.9093608 | 0.61511877 | cluster 8 |
| HORVU.MOREX.r3.3HG0275300.1 | -0.756268 | -0.330074 | -0.9695207 | cluster 8 |
| HORVU.MOREX.r3.3HG0299020.1 | 0.8142026 | 1.221284 | 0.80736762 | cluster 8 |
| HORVU.MOREX.r3.3HG0307340.1 | 0.9516359 | 1.299012 | 0.82274733 | cluster 8 |
| HORVU.MOREX.r3.4HG0331420.1 | 0.6161197 | 1.3603997 | 0.29829575 | cluster 8 |
| HORVU.MOREX.r3.4HG0332710.1 | -2.24356 | -1.189108 | -2.9250651 | cluster 8 |
| HORVU.MOREX.r3.4HG0337720.1 | 1.653155 | 2.7358164 | 1.55437631 | cluster 8 |
| HORVU.MOREX.r3.4HG0338290.1 | 0.6005137 | 0.8366727 | 0.56932138 | cluster 8 |
| HORVU.MOREX.r3.4HG0345840.1 | 1.0853304 | 1.4740281 | 0.77264726 | cluster 8 |
| HORVU.MOREX.r3.4HG0347840.1 | 0.3951563 | 0.4822898 | 0.31282058 | cluster 8 |
| HORVU.MOREX.r3.4HG0381620.1 | 1.0317313 | 1.2708755 | 0.96484377 | cluster 8 |
| HORVU.MOREX.r3.4HG0382280.1 | 0.3856754 | 1.2089321 | 0.2711101 | cluster 8 |
| HORVU.MOREX.r3.4HG0383340.1 | -0.870385 | -0.089032 | -1.5932755 | cluster 8 |
| HORVU.MOREX.r3.4HG0391160.1 | 0.5692179 | 0.9617819 | 0.54210761 | cluster 8 |
| HORVU.MOREX.r3.4HG0391440.1 | 1.7641957 | 2.444945 | 1.76772557 | cluster 8 |

**Table S3** Continued.

| ID | logFC_3 h | logFC_6 h | logFC_12 h | cluster No. |
|---|---|---|---|---|
| HORVU.MOREX.r3.4HG0395770.1 | -0.631838 | -0.472398 | -0.7406453 | cluster 8 |
| HORVU.MOREX.r3.4HG0410820.1 | 0.3757779 | 1.2713538 | 0.25521382 | cluster 8 |
| HORVU.MOREX.r3.4HG0412650.1 | 1.8399537 | 2.405969 | 1.76125167 | cluster 8 |
| HORVU.MOREX.r3.4HG0417920.1 | 2.6693365 | 3.4513226 | 2.62403404 | cluster 8 |
| HORVU.MOREX.r3.5HG0426110.1 | 1.8031372 | 2.4005857 | 1.66149118 | cluster 8 |
| HORVU.MOREX.r3.5HG0439710.1 | -1.381654 | -0.486641 | -1.972956 | cluster 8 |
| HORVU.MOREX.r3.5HG0441420.1 | 1.2605773 | 2.3016515 | 1.05034251 | cluster 8 |
| HORVU.MOREX.r3.5HG0447610.1 | 0.5047409 | 0.9429919 | 0.52926345 | cluster 8 |
| HORVU.MOREX.r3.5HG0470690.1 | 1.2429358 | 1.6905877 | 1.23651628 | cluster 8 |
| HORVU.MOREX.r3.5HG0471320.1 | -1.346566 | -0.79475 | -1.7343734 | cluster 8 |
| HORVU.MOREX.r3.5HG0473970.1 | 1.4828238 | 2.2987644 | 1.5130937 | cluster 8 |
| HORVU.MOREX.r3.5HG0477750.1 | 2.1417084 | 2.4619085 | 1.98143426 | cluster 8 |
| HORVU.MOREX.r3.5HG0485960.1 | 1.3586954 | 1.7677775 | 1.06750445 | cluster 8 |
| HORVU.MOREX.r3.5HG0488200.1 | 0.4976283 | 1.1247694 | 0.34608666 | cluster 8 |
| HORVU.MOREX.r3.5HG0495910.1 | 1.4237489 | 1.9154817 | 1.0961816 | cluster 8 |
| HORVU.MOREX.r3.5HG0500360.1 | 3.0432371 | 3.3969682 | 2.91684266 | cluster 8 |
| HORVU.MOREX.r3.5HG0508610.1 | 0.7048323 | 1.2109206 | 0.58303996 | cluster 8 |
| HORVU.MOREX.r3.5HG0509350.1 | 0.4899925 | 2.0910094 | 0.56851341 | cluster 8 |
| HORVU.MOREX.r3.5HG0509460.1 | -0.685116 | -0.427906 | -0.9448471 | cluster 8 |
| HORVU.MOREX.r3.5HG0512710.1 | 1.3515157 | 1.9535053 | 1.15738349 | cluster 8 |
| HORVU.MOREX.r3.5HG0517740.1 | 2.4264659 | 2.6794845 | 2.36584881 | cluster 8 |
| HORVU.MOREX.r3.5HG0527310.1 | 1.5258793 | 2.5036434 | 1.46706368 | cluster 8 |
| HORVU.MOREX.r3.5HG0532120.1 | 1.8178781 | 1.9583289 | 1.72602711 | cluster 8 |
| HORVU.MOREX.r3.6HG0543540.1 | 1.3863265 | 1.5053508 | 1.3537395 | cluster 8 |
| HORVU.MOREX.r3.6HG0554070.1 | 1.6818835 | 3.6687108 | 1.22141612 | cluster 8 |
| HORVU.MOREX.r3.6HG0576010.1 | 0.3679369 | 0.6885457 | 0.31848861 | cluster 8 |
| HORVU.MOREX.r3.6HG0595580.1 | 1.4276508 | 2.3878808 | 1.0853941 | cluster 8 |
| HORVU.MOREX.r3.6HG0606530.1 | 0.4428105 | 0.6577069 | 0.30875491 | cluster 8 |
| HORVU.MOREX.r3.6HG0607160.1 | 0.5753563 | 0.7585488 | 0.57609991 | cluster 8 |
| HORVU.MOREX.r3.6HG0609390.1 | 0.2668566 | 0.7232304 | 0.25618375 | cluster 8 |
| HORVU.MOREX.r3.6HG0609680.1 | 1.1862527 | 1.5598004 | 1.01676472 | cluster 8 |
| HORVU.MOREX.r3.6HG0613020.1 | 0.5524338 | 0.7327898 | 0.46253984 | cluster 8 |
| HORVU.MOREX.r3.7HG0634520.1 | 1.1504993 | 1.5528927 | 1.16398918 | cluster 8 |
| HORVU.MOREX.r3.7HG0636540.1 | 1.5695804 | 1.9686879 | 1.23916107 | cluster 8 |
| HORVU.MOREX.r3.7HG0638940.1 | 1.3386458 | 2.4034981 | 0.80305954 | cluster 8 |
| HORVU.MOREX.r3.7HG0642520.1 | 1.9370895 | 2.2840032 | 1.83025225 | cluster 8 |
| HORVU.MOREX.r3.7HG0650310.1 | 0.6615734 | 1.3406596 | 0.51987973 | cluster 8 |
| HORVU.MOREX.r3.7HG0654810.1 | -1.942124 | -0.260052 | -2.4042506 | cluster 8 |
| HORVU.MOREX.r3.7HG0662370.1 | 1.3241722 | 1.7259933 | 1.21876266 | cluster 8 |
| HORVU.MOREX.r3.7HG0662420.1 | 0.7603474 | 1.0866451 | 0.67240873 | cluster 8 |
| HORVU.MOREX.r3.7HG0663750.1 | 1.7470975 | 2.3972216 | 1.42992263 | cluster 8 |
| HORVU.MOREX.r3.7HG0665040.1 | 2.4892506 | 3.1369681 | 1.92838846 | cluster 8 |
| HORVU.MOREX.r3.7HG0667260.1 | 0.5586346 | 0.8013911 | 0.52307275 | cluster 8 |
| HORVU.MOREX.r3.7HG0673330.1 | 1.2884278 | 1.6333472 | 1.31253661 | cluster 8 |
| HORVU.MOREX.r3.7HG0697980.1 | -0.910687 | -0.560554 | -0.9376121 | cluster 8 |
| HORVU.MOREX.r3.7HG0708080.1 | 1.3750928 | 1.6864139 | 1.15384371 | cluster 8 |
| HORVU.MOREX.r3.7HG0710180.1 | -0.960132 | -0.461752 | -1.3850899 | cluster 8 |

**Table S3** Continued.

| ID | logFC_3 h | logFC_6 h | logFC_12 h | cluster No. |
|---|---|---|---|---|
| HORVU.MOREX.r3.7HG0731560.1 | 1.2332042 | 1.8283902 | 0.96734845 | cluster 8 |
| HORVU.MOREX.r3.7HG0735700.1 | 1.1785731 | 1.7513368 | 1.01796317 | cluster 8 |
| HORVU.MOREX.r3.7HG0739230.1 | 3.8556548 | 4.7252315 | 2.96062925 | cluster 8 |
| HORVU.MOREX.r3.1HG0072580.1 | 0.5504146 | 0.5074494 | 0.6826394 | cluster 9 |
| HORVU.MOREX.r3.1HG0076800.1 | 0.3709137 | 0.3121963 | 0.64710848 | cluster 9 |
| HORVU.MOREX.r3.1HG0084760.1 | 0.9694768 | 0.7509061 | 1.31581169 | cluster 9 |
| HORVU.MOREX.r3.2HG0101250.1 | 0.5952558 | 0.4208567 | 0.80339888 | cluster 9 |
| HORVU.MOREX.r3.2HG0194850.1 | 0.5105544 | 0.4560437 | 0.68449166 | cluster 9 |
| HORVU.MOREX.r3.2HG0204480.1 | -0.341332 | -0.630844 | -0.0348636 | cluster 9 |
| HORVU.MOREX.r3.3HG0226150.1 | 1.3843313 | 1.0545768 | 1.69469367 | cluster 9 |
| HORVU.MOREX.r3.4HG0376720.1 | 2.8197207 | 2.748408 | 2.94486373 | cluster 9 |
| HORVU.MOREX.r3.4HG0378460.1 | 0.5557031 | 0.4984911 | 0.77886874 | cluster 9 |
| HORVU.MOREX.r3.4HG0388780.1 | 0.8159196 | -0.05139 | 1.83722955 | cluster 9 |
| HORVU.MOREX.r3.5HG0442250.1 | 0.8581869 | 0.8140699 | 0.92212005 | cluster 9 |
| HORVU.MOREX.r3.5HG0475150.1 | 0.6894442 | 0.5314124 | 1.03057189 | cluster 9 |
| HORVU.MOREX.r3.5HG0485410.1 | 2.3398523 | 2.2335101 | 2.42992147 | cluster 9 |
| HORVU.MOREX.r3.5HG0509970.1 | 1.3142047 | 0.6687246 | 1.77801444 | cluster 9 |
| HORVU.MOREX.r3.5HG0527890.1 | 0.1545306 | 0.0248258 | 0.31949894 | cluster 9 |
| HORVU.MOREX.r3.6HG0570520.1 | -0.193203 | -0.251218 | -0.1025166 | cluster 9 |
| HORVU.MOREX.r3.6HG0577770.1 | -1.393416 | -1.916621 | -0.6306346 | cluster 9 |
| HORVU.MOREX.r3.6HG0606990.1 | 1.2102586 | 1.1073845 | 1.42794469 | cluster 9 |
| HORVU.MOREX.r3.7HG0674590.1 | 0.591647 | 0.4832687 | 1.1299294 | cluster 9 |
| HORVU.MOREX.r3.7HG0725440.1 | 0.4876101 | 0.2580707 | 0.8889395 | cluster 9 |
| HORVU.MOREX.r3.7HG0735210.1 | 1.5661608 | 1.5068382 | 1.6566023 | cluster 9 |
