## Supporting Information Table 4 for "ENHANCED GRAVITROPISM 2 coordinates molecular adaptations to gravistimulation in the elongation zone of barley roots"

**Table S4a** Enriched biological process terms among differentially expressed genes in the root cap of gravistimulated wild type roots.

| No. cluster | GO term | Description | cluster 1 | | | cluster 2 | | | cluster 3 | | |
|---|---|---|---|---|---|---|---|---|---|---|---|
| | | | 3 h | 6 h | 12 h | 3 h | 6 h | 12 h | 3 h | 6 h | 12 h |
| cluster 1 | GO:0006635 | fatty acid beta-oxidation | | | down-regulated | | | | | | |
| | GO:0009408 | response to heat | | | down-regulated | | | | | | |
| | GO:0009409 | response to cold | | | down-regulated | | | | | | |
| | GO:0015976 | carbon utilization | | | down-regulated | | | | | | |
| | GO:0050881 | musculoskeletal movement | | | down-regulated | | | | | | |
| | GO:1903426 | regulation of reactive oxygen species biosynthetic process | | | down-regulated | | | | | | |
| | GO:0031122 | cytoplasmic microtubule organization | | | down-regulated | | | | | | |
| | GO:0044743 | protein transmembrane import into intracellular organelle | | | down-regulated | | | | | | |
| | GO:0065002 | intracellular protein transmembrane transport | | | down-regulated | | | | | | |
| | GO:0006470 | protein dephosphorylation | | | down-regulated | | | | | | |
| | GO:0006468 | protein phosphorylation | | | down-regulated | | | | | | |
| | GO:0006809 | nitric oxide biosynthetic process | | | down-regulated | | | | | | |
| | GO:0006629 | lipid metabolic process | | | down-regulated | | | | | | |
| | GO:0005975 | carbohydrate metabolic process | | | down-regulated | | | | | | |
| | GO:0055023 | positive regulation of cardiac muscle tissue growth | | | down-regulated | | | | | | |
| | GO:0050795 | regulation of behavior | | | down-regulated | | | | | | |
| | GO:0051726 | regulation of cell cycle | | | down-regulated | | | | | | |
| | GO:0061041 | regulation of wound healing | | | down-regulated | | | | | | |
| | GO:0006012 | galactose metabolic process | | | down-regulated | | | | | | |
| | GO:0010951 | negative regulation of endopeptidase activity | | | down-regulated | | | | | | |
| | GO:0010507 | negative regulation of autophagy | | | down-regulated | | | | | | |
| | GO:0006359 | regulation of transcription by RNA polymerase III | | | down-regulated | | | | | | |
| | GO:0006570 | tyrosine metabolic process | | | down-regulated | | | | | | |
| | GO:0046496 | nicotinamide nucleotide metabolic process | | | down-regulated | | | | | | |
| | GO:0014009 | glial cell proliferation | | | down-regulated | | | | | | |
| | GO:0010411 | xyloglucan metabolic process | | | down-regulated | | | | | | |
| | GO:0005978 | glycogen biosynthetic process | | | down-regulated | | | | | | |
| | GO:0005977 | glycogen metabolic process | | | down-regulated | | | | | | |
| | GO:0015866 | ADP transport | | | down-regulated | | | | | | |
| | GO:0034198 | cellular response to amino acid starvation | | | down-regulated | | | | | | |
| | GO:0002011 | morphogenesis of an epithelial sheet | | | down-regulated | | | | | | |
| | GO:0019673 | GDP-mannose metabolic process | | | down-regulated | | | | | | |
| | GO:0010717 | regulation of epithelial to mesenchymal transition | | | down-regulated | | | | | | |
| | GO:0016117 | carotenoid biosynthetic process | | | down-regulated | | | | | | |
| | GO:0006914 | autophagy | | | down-regulated | | | | | | |
| | GO:0035307 | positive regulation of protein dephosphorylation | | | down-regulated | | | | | | |
| | GO:0050848 | regulation of calcium-mediated signaling | | | down-regulated | | | | | | |
| cluster 2 | GO:0009308 | amine metabolic process | | | | | | down-regulated | | | |
| cluster 3 | GO:0051491 | positive regulation of filopodium assembly | | | | | | | | | up-regulated |
| | GO:0030833 | regulation of actin filament polymerization | | | | | | | | | up-regulated |
| | GO:0098742 | cell-cell adhesion via plasma-membrane adhesion molecules | | | | | | | | | up-regulated |
| | GO:0019722 | calcium-mediated signaling | | | | | | | | | up-regulated |
| | GO:0042176 | regulation of protein catabolic process | | | | | | | | | up-regulated |
| | GO:0007411 | axon guidance | | | | | | | | | up-regulated |

up-regulated
down-regulated

**Table S4b** Enriched biological process terms among differentially expressed genes in the elongation zone of gravistimulated wild type roots.

| No. cluster | GO term | Description | cluster 1 | | | cluster 2 | | | cluster 3 | | | cluster 4 | | | cluster 5 | | | cluster 6 | | | cluster 7 | | | cluster 8 | | | cluster 9 | | |
|---|---|---|---|---|---|---|---|---|---|---|---|---|---|---|---|---|---|---|---|---|---|---|---|---|---|---|---|---|---|
| | | | 3 h | 6 h | 12 h | 3 h | 6 h | 12 h | 3 h | 6 h | 12 h | 3 h | 6 h | 12 h | 3 h | 6 h | 12 h | 3 h | 6 h | 12 h | 3 h | 6 h | 12 h | 3 h | 6 h | 12 h | 3 h | 6 h | 12 h |
| cluster 1 | GO:0009664 | plant-type cell wall organization | up | up and down | up and down | | up | | | up | up | | | up | | | | | | | | down | up and down | | | up | | | |
| | GO:0006566 | threonine metabolic process | | up | down | | | | | | | | | | | | up | | | | | | | | | | | | |
| | GO:0006544 | glycine metabolic process | | up | | | | | | | | | | | | | | | | | | | | | | | | | |
| | GO:1901606 | alpha-amino acid catabolic process | | up | | | | | | | | | | | | | | | | | | | | | | | | | |
| | GO:0009063 | cellular amino acid catabolic process | | up | | | | | | | | | | | | | | | | | | | | | | | | | |
| | GO:0016311 | dephosphorylation | | up | | | | | | | | | | | | | | | | | | | | | | | | | |
| | GO:0006004 | fucose metabolic process | | down | down | | | | | | | | down | down | | | | | | | | down | | | | | | | |
| | GO:0019320 | hexose catabolic process | | down | | | | | | | | | | | | | | | | | | | | | | | | | |
| | GO:0006012 | galactose metabolic process | | down | down | | | up | | | | | | | | | | | | | | | | | | | | | |
| | GO:0007219 | Notch signaling pathway | | down | down | | | | | | | | | | | | | | | | | | | | | | | | |
| | GO:0010102 | lateral root morphogenesis | | down | down | | | | | | | | | | | | | | | | | | | | | | | | |
| | GO:0072595 | maintenance of protein localization in organelle | | down | | | | | | | | | | | | | | | | | | | | | | | | | |
| | GO:0007020 | microtubule nucleation | | down | down | | | | | | | | | | | | | | | | | | | | | | | | |
| | GO:0098930 | axonal transport | | down | | | | | | | | | | | | | | | | | | | | | | | | | |
| | GO:0045104 | intermediate filament cytoskeleton organization | | down | | | | | | | | | | | | | | | | | | | | | | | | | |
| | GO:0047496 | vesicle transport along microtubule | | down | | | | | | | | | | | | | | | | | | | | | | | | | |
| | GO:0006457 | protein folding | | down | down | | | | | | | | | | | | | | | | | | | | | | | | |
| | GO:0016485 | protein processing | | down | | | | | | | | | | | | | | | | | | | | | | | | | |
| | GO:0006511 | ubiquitin-dependent protein catabolic process | | down | down | | | | | | | | | | | | | | | | | | | | | | | | |
| | GO:0030433 | ubiquitin-dependent ERAD pathway | | down | | | | | | | | | | | | | | | | | | | | | | | | | |
| | GO:0008299 | isoprenoid biosynthetic process | | down | | | | | | | | | | | | | | | | | | | | | | | | | |
| | GO:0006665 | sphingolipid metabolic process | | down | | | | | | | | | | | | | | | | | | | | | | | | | |
| | GO:0005975 | carbohydrate metabolic process | | down | down | | | | | | | | | | | | up | | | | | | | | | | | | |
| | GO:1903036 | positive regulation of response to wounding | | down | | | | | | | | | | | | | | | | | | | | | | | | | |
| | GO:0015780 | nucleotide-sugar transmembrane transport | | down | down | | | up | | | | | | | | | | | | | | | | | | | | | |
| | GO:0033157 | regulation of intracellular protein transport | | down | | | | | | | | | | | | | | | | | | | | | | | | | |
| | GO:0040008 | regulation of growth | | down | | | | | | | | | | | | | | | | | | | | | | | | | |
| | GO:0006486 | protein glycosylation | | down | down | | | | | | | | down | down | | | | | | | | | | | | | | | |
| | GO:0036065 | fucosylation | | down | down | | | | | | | | | | | | | | | | | | | | up | up | | | |
| | GO:0006493 | protein O-linked glycosylation | | down | down | | | | | | | | | | | | | | | | | | | | | | | | |
| | GO:0006487 | protein N-linked glycosylation | | down | down | | | | | | | | | | | | | | | | | | | | | | | | |
| | GO:0031529 | ruffle organization | | down | | | | | | | | | | | | | | | | | | | | | | | | | |
| | GO:0031103 | axon regeneration | | down | | | | | | | | | | | | | | | | | | | | | | | | | |
| | GO:0006082 | organic acid metabolic process | | down | | | | | | | | | | | | | | | | | | | | | | | | | |
| | GO:0045489 | pectin biosynthetic process | | down | down | | | | | | | | | | | | | | | | | | | | | | | | |
| | GO:0070592 | cell wall polysaccharide biosynthetic process | | down | down | | | | | | | | down | | | | | | | | | | | | | | | | |
| | GO:0009734 | auxin-activated signaling pathway | | down | | | | | | | | | | | | | | | | | | | | | | | | | |
| | GO:0008286 | insulin receptor signaling pathway | | down | | | | | | | | | | | | | | | | | | | | | | | | | |
| | GO:0046686 | response to cadmium ion | | down | down | | | | | | | | | | | | | | | | | | | | | | | | |
| | GO:0009414 | response to water deprivation | | down | down | | | | | | | | | | | | | | | | | | | | | | | | |
| | GO:0009409 | response to cold | | | down | | | | | | | | | | | | | | | | | | | | | | | | |
| | GO:0010039 | response to iron ion | | | down | | | | | | | | | | | | up | | | | | | | | | | | | |
| | GO:0045773 | positive regulation of axon extension | | down | | | | | | | | | | | | | | | | | | | | | | | | | |
| | GO:0022029 | telencephalon cell migration | | down | | | | | | | | | | | | | | | | | | | | | | | | | |
| | GO:0021954 | central nervous system neuron development | | down | | | | | | | | | | | | | | | | | | | | | | | | | |
| | GO:0001764 | neuron migration | | down | | | | | | | | | | | | | | | | | | | | | | | | | |
| | GO:0021987 | cerebral cortex development | | down | | | | | | | | | | | | | | | | | | | | | | | | | |
| | GO:0009311 | oligosaccharide metabolic process | | down | down | | | | | | | | | | | | | | | | | | | | | | | | |
| | GO:0032418 | lysosome localization | | down | | | | | | | | | | | | | | | | | | | | | | | | | |

- up-regulated
- down-regulated
- up and down-regulation

**Table S4b** Continued.

| No. cluster | GO term | Description | cluster 1 | | | cluster 2 | | | cluster 3 | | | cluster 4 | | | cluster 5 | | | cluster 6 | | | cluster 7 | | | cluster 8 | | | cluster 9 | | |
|---|---|---|---|---|---|---|---|---|---|---|---|---|---|---|---|---|---|---|---|---|---|---|---|---|---|---|---|---|---|
| | | | 3 h | 6 h | 12 h | 3 h | 6 h | 12 h | 3 h | 6 h | 12 h | 3 h | 6 h | 12 h | 3 h | 6 h | 12 h | 3 h | 6 h | 12 h | 3 h | 6 h | 12 h | 3 h | 6 h | 12 h | 3 h | 6 h | 12 h |
| cluster 1 | GO:0051303 | establishment of chromosome localization | | | | | | | | | | | | | | | | | | | | | | | | | | | |
| | GO:0005984 | disaccharide metabolic process | | | | | | | | | | | | | | | | | | | | | | | | | | | |
| | GO:0005992 | trehalose biosynthetic process | | | | | | | | | | | | | | | | | | | | | | | | | | | |
| | GO:0001824 | blastocyst development | | | | | | | | | | | | | | | | | | | | | | | | | | | |
| | GO:0016579 | protein deubiquitination | | | | | | | | | | | | | | | | | | | | | | | | | | | |
| | GO:1905393 | plant organ formation | | | | | | | | | | | | | | | | | | | | | | | | | | | |
| | GO:0006085 | acetyl-CoA biosynthetic process | | | | | | | | | | | | | | | | | | | | | | | | | | | |
| | GO:0048316 | seed development | | | | | | | | | | | | | | | | | | | | | | | | | | | |
| | GO:0007140 | male meiotic nuclear division | | | | | | | | | | | | | | | | | | | | | | | | | | | |
| | GO:0009832 | plant-type cell wall biogenesis | | | | | | | | | | | | | | | | | | | | | | | | | | | |
| | GO:1990748 | cellular detoxification | | | | | | | | | | | | | | | | | | | | | | | | | | | |
| | GO:0036503 | ERAD pathway | | | | | | | | | | | | | | | | | | | | | | | | | | | |
| | GO:0009873 | ethylene-activated signaling pathway | | | | | | | | | | | | | | | | | | | | | | | | | | | |
| | GO:0009737 | response to abscisic acid | | | | | | | | | | | | | | | | | | | | | | | | | | | |
| | GO:0071369 | cellular response to ethylene stimulus | | | | | | | | | | | | | | | | | | | | | | | | | | | |
| | GO:0006099 | tricarboxylic acid cycle | | | | | | | | | | | | | | | | | | | | | | | | | | | |
| | GO:0006067 | ethanol metabolic process | | | | | | | | | | | | | | | | | | | | | | | | | | | |
| | GO:0009206 | purine ribonucleoside triphosphate biosynthetic process | | | | | | | | | | | | | | | | | | | | | | | | | | | |
| | GO:0015936 | coenzyme A metabolic process | | | | | | | | | | | | | | | | | | | | | | | | | | | |
| | GO:0006165 | nucleoside diphosphate phosphorylation | | | | | | | | | | | | | | | | | | | | | | | | | | | |
| | GO:0046039 | GTP metabolic process | | | | | | | | | | | | | | | | | | | | | | | | | | | |
| | GO:0006633 | fatty acid biosynthetic process | | | | | | | | | | | | | | | | | | | | | | | | | | | |
| | GO:0046470 | phosphatidylcholine metabolic process | | | | | | | | | | | | | | | | | | | | | | | | | | | |
| | GO:0009800 | cinnamic acid biosynthetic process | | | | | | | | | | | | | | | | | | | | | | | | | | | |
| | GO:0009059 | macromolecule biosynthetic process | | | | | | | | | | | | | | | | | | | | | | | | | | | |
| | GO:0006508 | proteolysis | | | | | | | | | | | | | | | | | | | | | | | | | | | |
| | GO:0006334 | nucleosome assembly | | | | | | | | | | | | | | | | | | | | | | | | | | | |
| | GO:0007010 | cytoskeleton organization | | | | | | | | | | | | | | | | | | | | | | | | | | | |
| | GO:0006073 | cellular glucan metabolic process | | | | | | | | | | | | | | | | | | | | | | | | | | | |
| | GO:0030244 | cellulose biosynthetic process | | | | | | | | | | | | | | | | | | | | | | | | | | | |
| | GO:0006559 | L-phenylalanine catabolic process | | | | | | | | | | | | | | | | | | | | | | | | | | | |
| | GO:1901575 | organic substance catabolic process | | | | | | | | | | | | | | | | | | | | | | | | | | | |
| | GO:0006997 | nucleus organization | | | | | | | | | | | | | | | | | | | | | | | | | | | |
| | GO:0000096 | sulfur amino acid metabolic process | | | | | | | | | | | | | | | | | | | | | | | | | | | |
| | GO:0009826 | unidimensional cell growth | | | | | | | | | | | | | | | | | | | | | | | | | | | |
| cluster 2 | GO:0006979 | response to oxidative stress | | | | | | | | | | | | | | | | | | | | | | | | | | | |
| | GO:0035377 | transepithelial water transport | | | | | | | | | | | | | | | | | | | | | | | | | | | |
| | GO:0042744 | hydrogen peroxide catabolic process | | | | | | | | | | | | | | | | | | | | | | | | | | | |
| | GO:0048146 | positive regulation of fibroblast proliferation | | | | | | | | | | | | | | | | | | | | | | | | | | | |
| | GO:0030950 | establishment or maintenance of actin cytoskeleton polarity | | | | | | | | | | | | | | | | | | | | | | | | | | | |
| | GO:0045445 | myoblast differentiation | | | | | | | | | | | | | | | | | | | | | | | | | | | |
| | GO:0035914 | skeletal muscle cell differentiation | | | | | | | | | | | | | | | | | | | | | | | | | | | |
| | GO:0006468 | protein phosphorylation | | | | | | | | | | | | | | | | | | | | | | | | | | | |
| | GO:0032941 | secretion by tissue | | | | | | | | | | | | | | | | | | | | | | | | | | | |
| | GO:0006687 | glycosphingolipid metabolic process | | | | | | | | | | | | | | | | | | | | | | | | | | | |
| | GO:0006672 | ceramide metabolic process | | | | | | | | | | | | | | | | | | | | | | | | | | | |
| | GO:0072488 | ammonium transmembrane transport | | | | | | | | | | | | | | | | | | | | | | | | | | | |
| | GO:0015791 | polyol transport | | | | | | | | | | | | | | | | | | | | | | | | | | | |
| | GO:0010411 | xyloglucan metabolic process | | | | | | | | | | | | | | | | | | | | | | | | | | | |

**Table S4b** Continued.

| No. cluster | GO term | Description |
|---|---|---|
| cluster 2 | GO:0005991 | trehalose metabolic process |
| | GO:0042546 | cell wall biogenesis |
| | GO:0071870 | cellular response to catecholamine stimulus |
| | GO:0021670 | lateral ventricle development |
| | GO:0022600 | digestive system process |
| | GO:0046689 | response to mercury ion |
| | GO:0046274 | lignin catabolic process |
| | GO:0071474 | cellular hyperosmotic response |
| | GO:0043171 | peptide catabolic process |
| | GO:0042219 | cellular modified amino acid catabolic process |
| | GO:0007178 | transmembrane receptor protein serine/threonine kinase signaling pathway |
| | GO:0046364 | monosaccharide biosynthetic process |
| | GO:0008643 | carbohydrate transport |
| | GO:0000160 | phosphorelay signal transduction system |
| cluster3 | GO:0006542 | glutamine biosynthetic process |
| | GO:0015695 | organic cation transport |
| | GO:0034220 | ion transmembrane transport |
| | GO:0046856 | phosphatidylinositol dephosphorylation |
| | GO:0032606 | type I interferon production |
| | GO:0009690 | cytokinin metabolic process |
| | GO:0001959 | regulation of cytokine-mediated signaling pathway |
| | GO:2001233 | regulation of apoptotic signaling pathway |
| | GO:1905477 | positive regulation of protein localization to membrane |
| | GO:0032414 | positive regulation of ion transmembrane transporter activity |
| | GO:0022408 | negative regulation of cell-cell adhesion |
| | GO:0045582 | positive regulation of T cell differentiation |
| | GO:0007041 | lysosomal transport |
| | GO:0006741 | NADP biosynthetic process |
| | GO:0016119 | carotene metabolic process |
| | GO:0050866 | negative regulation of cell activation |
| | GO:0048532 | anatomical structure arrangement |
| | GO:0050905 | neuromuscular process |
| | GO:0010043 | response to zinc ion |
| | GO:0030837 | negative regulation of actin filament polymerization |
| | GO:0030834 | regulation of actin filament depolymerization |
| | GO:1901880 | negative regulation of protein depolymerization |
| | GO:0035315 | hair cell differentiation |
| | GO:0006950 | response to stress |
| cluster 4 | GO:0006813 | potassium ion transport |
| | GO:0070839 | metal ion export |
| | GO:0035725 | sodium ion transmembrane transport |
| | GO:0007006 | mitochondrial membrane organization |
| | GO:0050848 | regulation of calcium-mediated signaling |
| | GO:0071871 | response to epinephrine |
| | GO:0006629 | lipid metabolic process |
| | GO:0051930 | regulation of sensory perception of pain |
| | GO:0010611 | regulation of cardiac muscle hypertrophy |
| | GO:0051453 | regulation of intracellular pH |

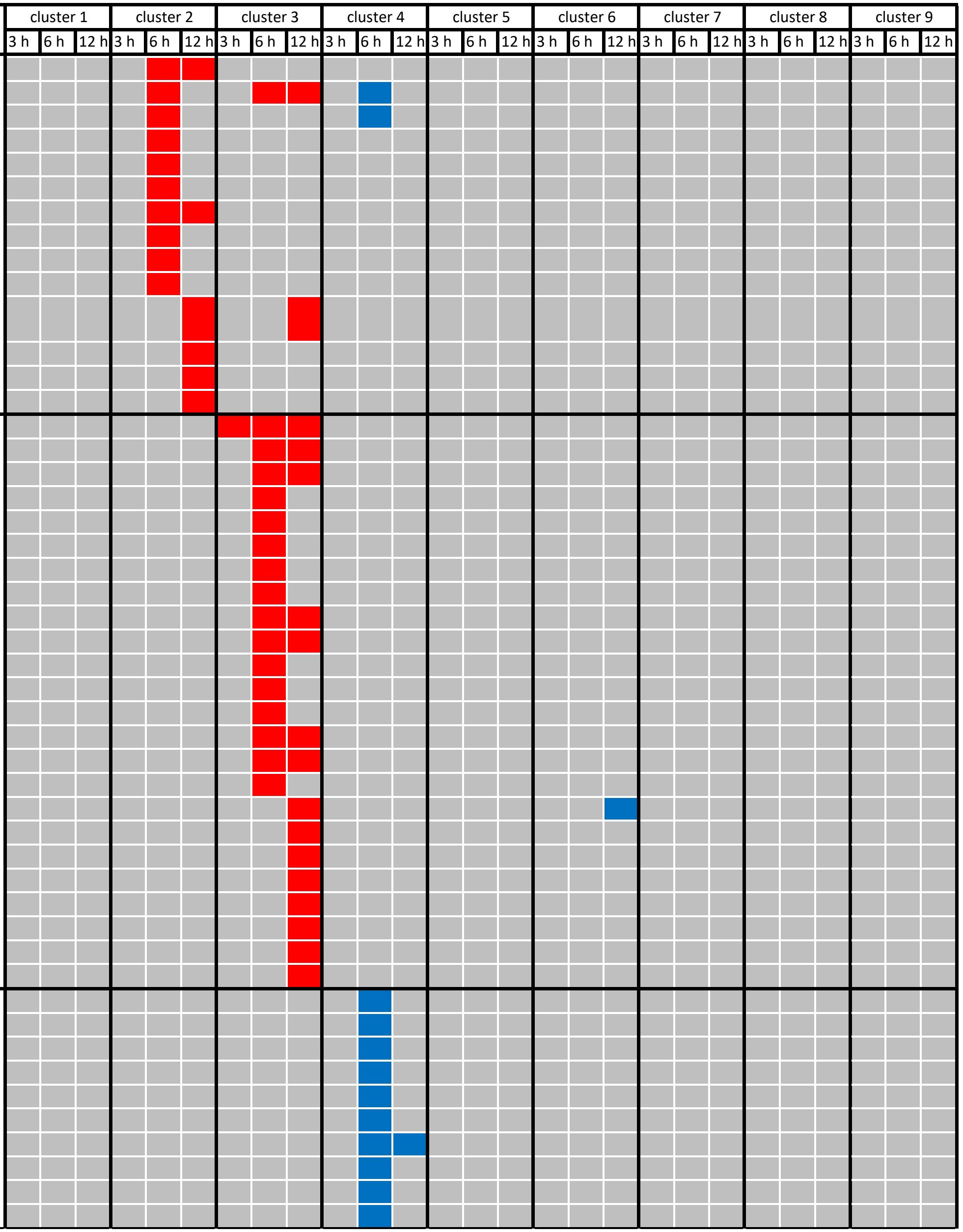

**Table S4b** Continued.

| No. cluster | GO term | Description |
|---|---|---|
| cluster 4 | GO:0048016 | inositol phosphate-mediated signaling |
| | GO:0009268 | response to pH |
| | GO:0051602 | response to electrical stimulus |
| | GO:0006972 | hyperosmotic response |
| | GO:0006303 | double-strand break repair via nonhomologous end joining |
| | GO:0009735 | response to cytokinin |
| | GO:2000028 | regulation of photoperiodism, flowering |
| | GO:0009100 | glycoprotein metabolic process |
| | GO:0006506 | GPI anchor biosynthetic process |
| | GO:0006888 | endoplasmic reticulum to Golgi vesicle-mediated transport |
| cluster 5 | GO:0006536 | glutamate metabolic process |
| | GO:0009084 | glutamine family amino acid biosynthetic process |
| | GO:0043650 | dicarboxylic acid biosynthetic process |
| | GO:0009741 | response to brassinosteroid |
| | GO:0010923 | negative regulation of phosphatase activity |
| | GO:0043666 | regulation of phosphoprotein phosphatase activity |
| | GO:0006458 | 'de novo' protein folding |
| | GO:1990542 | mitochondrial transmembrane transport |
| | GO:0007015 | actin filament organization |
| | GO:0007018 | microtubule-based movement |
| | GO:0009773 | photosynthetic electron transport in photosystem I |
| | GO:0006807 | nitrogen compound metabolic process |
| | GO:0046654 | tetrahydrofolate biosynthetic process |
| | GO:0072529 | pyrimidine-containing compound catabolic process |
| | GO:0008272 | sulfate transport |
| | GO:0044743 | protein transmembrane import into intracellular organelle |
| | GO:0006626 | protein targeting to mitochondrion |
| | GO:0065002 | intracellular protein transmembrane transport |
| | GO:0036092 | phosphatidylinositol-3-phosphate biosynthetic process |
| cluster 6 | GO:0006086 | acetyl-CoA biosynthetic process from pyruvate |
| | GO:0006431 | methionyl-tRNA aminoacylation |
| | GO:0016024 | CDP-diacylglycerol biosynthetic process |
| | GO:0045892 | negative regulation of transcription, DNA-templated |
| | GO:0007031 | peroxisome organization |
| | GO:0009056 | catabolic process |
| | GO:0045292 | mRNA cis splicing, via spliceosome |
| | GO:0009740 | gibberellic acid mediated signaling pathway |
| | GO:0051781 | positive regulation of cell division |
| | GO:0009411 | response to UV |
| | GO:0009157 | deoxyribonucleoside monophosphate biosynthetic process |
| | GO:0007131 | reciprocal meiotic recombination |
| | GO:0019264 | glycine biosynthetic process from serine |
| | GO:0048507 | meristem development |
| | GO:0034728 | nucleosome organization |
| | GO:0006333 | chromatin assembly or disassembly |
| cluster 7 | GO:0010015 | root morphogenesis |
| | GO:0060560 | developmental growth involved in morphogenesis |
| | GO:0048588 | developmental cell growth |
| | GO:0048168 | regulation of neuronal synaptic plasticity |

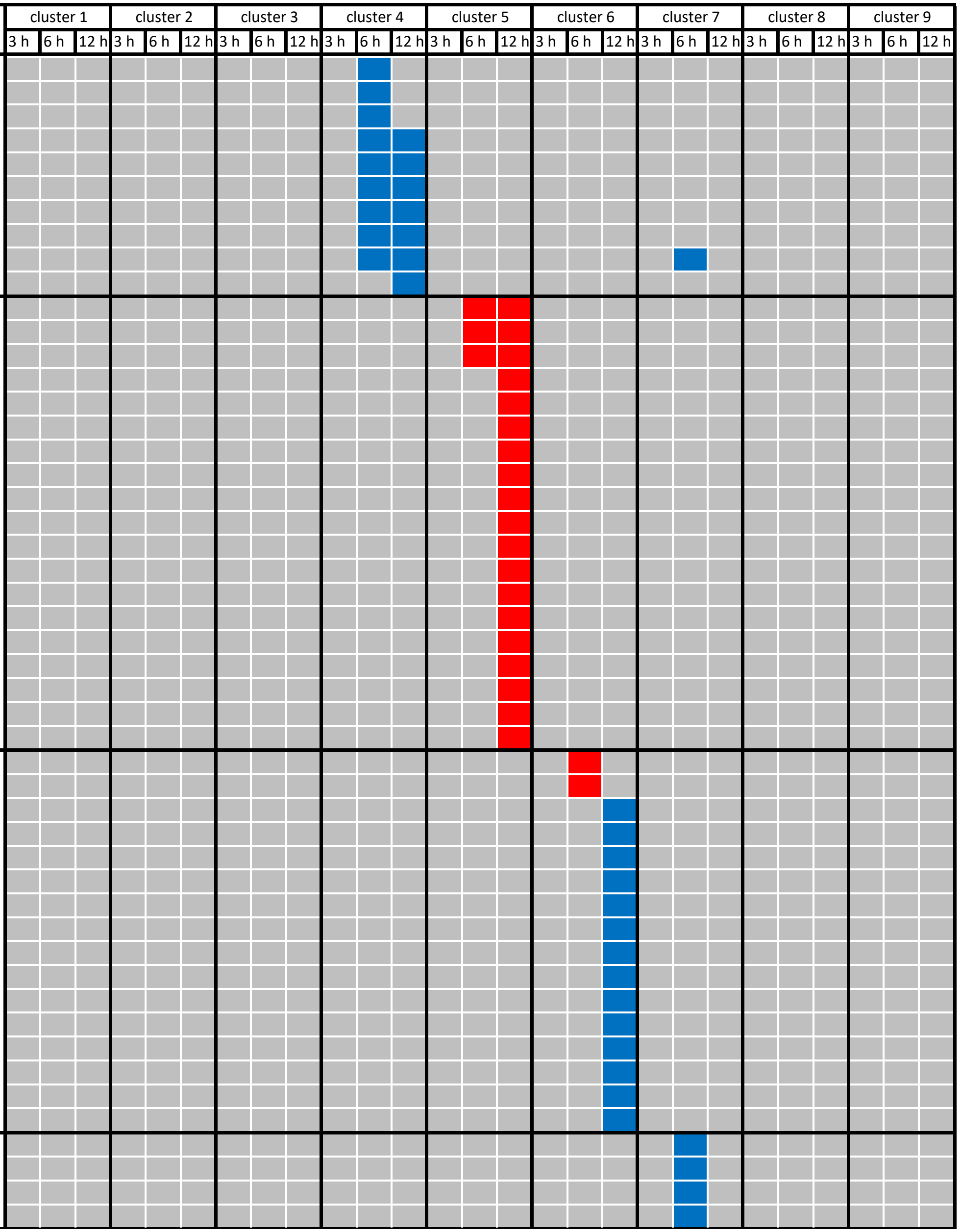

**Table S4b** Continued.

| No. cluster | GO term | Description | cluster 1 | | | cluster 2 | | | cluster 3 | | | cluster 4 | | | cluster 5 | | | cluster 6 | | | cluster 7 | | | cluster 8 | | | cluster 9 | | |
|---|---|---|---|---|---|---|---|---|---|---|---|---|---|---|---|---|---|---|---|---|---|---|---|---|---|---|---|---|---|
| | | | 3 h | 6 h | 12 h | 3 h | 6 h | 12 h | 3 h | 6 h | 12 h | 3 h | 6 h | 12 h | 3 h | 6 h | 12 h | 3 h | 6 h | 12 h | 3 h | 6 h | 12 h | 3 h | 6 h | 12 h | 3 h | 6 h | 12 h |
| cluster 7 | GO:0042545 | cell wall modification | | | | | | | | | | | | | | | | | | | | | | | | | | | |
| | GO:0046149 | pigment catabolic process | | | | | | | | | | | | | | | | | | | | | | | | | | | |
| | GO:0006787 | porphyrin-containing compound catabolic process | | | | | | | | | | | | | | | | | | | | | | | | | | | |
| | GO:0019511 | peptidyl-proline hydroxylation | | | | | | | | | | | | | | | | | | | | | | | | | | | |
| | GO:0007269 | neurotransmitter secretion | | | | | | | | | | | | | | | | | | | | | | | | | | | |
| | GO:0090558 | plant epidermis development | | | | | | | | | | | | | | | | | | | | | | | | | | | |
| | GO:0009103 | lipopolysaccharide biosynthetic process | | | | | | | | | | | | | | | | | | | | | | | | | | | |
| | GO:0030833 | regulation of actin filament polymerization | | | | | | | | | | | | | | | | | | | | | | | | | | | |
| | GO:0001558 | regulation of cell growth | | | | | | | | | | | | | | | | | | | | | | | | | | | |
| | GO:0048469 | cell maturation | | | | | | | | | | | | | | | | | | | | | | | | | | | |
| cluster 8 | GO:0010228 | vegetative to reproductive phase transition of meristem | | | | | | | | | | | | | | | | | | | | | | | | | | | |
| | GO:0009809 | lignin biosynthetic process | | | | | | | | | | | | | | | | | | | | | | | | | | | |
| | GO:0035335 | peptidyl-tyrosine dephosphorylation | | | | | | | | | | | | | | | | | | | | | | | | | | | |
| | GO:0006367 | transcription initiation from RNA polymerase II promoter | | | | | | | | | | | | | | | | | | | | | | | | | | | |
| | GO:0015689 | molybdate ion transport | | | | | | | | | | | | | | | | | | | | | | | | | | | |
| | GO:0016042 | lipid catabolic process | | | | | | | | | | | | | | | | | | | | | | | | | | | |
| | GO:0009252 | peptidoglycan biosynthetic process | | | | | | | | | | | | | | | | | | | | | | | | | | | |
| | GO:0006298 | mismatch repair | | | | | | | | | | | | | | | | | | | | | | | | | | | |
| | GO:0008360 | regulation of cell shape | | | | | | | | | | | | | | | | | | | | | | | | | | | |
| cluster 9 | GO:0006520 | cellular amino acid metabolic process | | | | | | | | | | | | | | | | | | | | | | | | | | | |
| | GO:0006352 | DNA-templated transcription, initiation | | | | | | | | | | | | | | | | | | | | | | | | | | | |
| | GO:0009058 | biosynthetic process | | | | | | | | | | | | | | | | | | | | | | | | | | | |
