## Supporting Information Table 5 for "ENHANCED GRAVITROPISM 2 coordinates molecular adaptations to gravistimulation in the elongation zone of barley roots"

**Table S5** Overview of genes differentially expressed between wild type and *egt2* (FDR <5%) in a gravistimulation time-course experiment.

| ID | logFC | AveExpr | t | P.Value | adj.P.Val | B | gene description | root zone | time point |
|---|---|---|---|---|---|---|---|---|---|
| HORVU.MOREX.r3.1HG0078900.1 | -1.929327809 | 5.8370464 | -23.68173821 | 7.29E-30 | 1.50E-25 | 56.03841628 | MATH domain-containing protein | root cap | 0 h |
| HORVU.MOREX.r3.1HG0067040.1 | -1.308404397 | 7.101906906 | -11.15904817 | 1.58E-15 | 1.08E-11 | 25.11728876 | NAC domain protein, | root cap | 0 h |
| HORVU.MOREX.r3.1HG0067490.1 | -6.554595547 | -1.72649178 | -12.74731843 | 8.95E-18 | 9.19E-14 | 18.16094389 | Retrovirus-related Pol polyprotein from transposon TNT 1-94 | root cap | 0 h |
| HORVU.MOREX.r3.7HG0748980.1 | 3.493254047 | -2.90728761 | 9.609202952 | 3.32E-13 | 1.70E-09 | 15.920196 | 11S seed storage globulin A | root cap | 0 h |
| HORVU.MOREX.r3.3HG0280700.1 | 0.608457798 | 6.27769849 | 7.701208229 | 3.36E-10 | 7.66E-07 | 13.11387248 | Katanin p60 ATPase-containing subunit A1 | root cap | 0 h |
| HORVU.MOREX.r3.5HG0475760.1 | -4.962837145 | -2.872794743 | -8.551939863 | 1.48E-11 | 6.08E-08 | 11.97354268 | Skp1 | root cap | 0 h |
| HORVU.MOREX.r3.7HG0727570.1 | 4.315861861 | -0.569333422 | 8.16872267 | 6.01E-11 | 1.76E-07 | 11.34057257 | ATP-dependent RNA helicase, putative | root cap | 0 h |
| HORVU.MOREX.r3.1HG0007570.1 | -4.739766406 | -2.580123327 | -8.313168127 | 3.54E-11 | 1.21E-07 | 9.983565255 | Werner Syndrome-like exonuclease | root cap | 0 h |
| HORVU.MOREX.r3.2HG0111680.1 | -4.781564999 | -2.082427636 | -7.935560457 | 1.42E-10 | 3.63E-07 | 8.840158354 | Formate--tetrahydrofolate ligase | root cap | 0 h |
| HORVU.MOREX.r3.7HG0679070.1 | -1.43967604 | 9.365871523 | -6.345405901 | 5.08E-08 | 9.47E-05 | 8.104093549 | Rab3 GTPase-activating protein catalytic subunit | root cap | 0 h |
| HORVU.MOREX.r3.1HG0067030.1 | -0.905505797 | 5.504670742 | -6.131950615 | 1.11E-07 | 0.000190617 | 7.527487432 | Protein trichome birefringence | root cap | 0 h |
| HORVU.MOREX.r3.5HG0447830.1 | -0.688518141 | 4.419279148 | -5.9785554 | 1.96E-07 | 0.000308917 | 7.00166162 | Sterile alpha motif (SAM) domain-containing protein | root cap | 0 h |
| HORVU.MOREX.r3.6HG0629160.1 | 1.201077078 | 2.999097227 | 5.800628848 | 3.75E-07 | 0.000480882 | 6.390525245 | Disease resistance protein RPM1 | root cap | 0 h |
| HORVU.MOREX.r3.6HG0559680.1 | -2.865919207 | 0.573580493 | -5.848088929 | 3.15E-07 | 0.000431402 | 6.058192839 | NADPH:quinone oxidoreductase | root cap | 0 h |
| HORVU.MOREX.r3.5HG0523890.1 | -4.305158039 | -2.93597984 | -6.621856604 | 1.83E-08 | 3.75E-05 | 6.053215176 | Protein FAR1-RELATED SEQUENCE 5 | root cap | 0 h |
| HORVU.MOREX.r3.2HG0170640.1 | 3.361663461 | -0.509068665 | 5.677495016 | 5.87E-07 | 0.000692137 | 5.125097007 | Protein FAR1-RELATED SEQUENCE 5 | root cap | 0 h |
| HORVU.MOREX.r3.5HG0523880.1 | -3.723667717 | -2.522148569 | -5.887396499 | 2.73E-07 | 0.000400404 | 4.751039016 | cryptochrome-interacting basic-helix-loop-helix 5 | root cap | 0 h |
| HORVU.MOREX.r3.7HG0739910.1 | 0.892656383 | 4.210713917 | 5.286844618 | 2.40E-06 | 0.002516764 | 4.661580015 | Disease resistance protein (NBS-LRR class) family | root cap | 0 h |
| HORVU.MOREX.r3.7HG0695920.1 | 1.276263915 | 2.660691655 | 5.266940332 | 2.58E-06 | 0.002516764 | 4.603056232 | #N/A | root cap | 0 h |
| HORVU.MOREX.r3.2HG0215060.1 | 4.98419157 | -1.599980119 | 5.66814002 | 6.07E-07 | 0.000692137 | 4.059746903 | WAT1-related protein | root cap | 0 h |
| HORVU.MOREX.r3.4HG0343330.1 | 5.573085555 | -0.333480115 | 5.277388614 | 2.48E-06 | 0.002516764 | 3.838588837 | Endonuclease III homolog | root cap | 0 h |
| HORVU.MOREX.r3.5HG0489790.1 | 1.891910844 | -0.067483895 | 5.063576336 | 5.30E-06 | 0.00473253 | 3.557181376 | SKP1-like protein 4 | root cap | 0 h |
| HORVU.MOREX.r3.7HG0720330.1 | -2.199244835 | -3.241254914 | -5.220048657 | 3.04E-06 | 0.002839724 | 3.546752538 | Phosphatidylinositol 4-phosphate 5-kinase 1 | root cap | 0 h |
| HORVU.MOREX.r3.7HG0663550.1 | -2.348596099 | 0.15599737 | -4.949164396 | 7.94E-06 | 0.006097532 | 3.488292383 | Glycosyltransferase | root cap | 0 h |
| HORVU.MOREX.r3.1HG0049550.1 | 0.544247115 | 3.543555344 | 4.900725739 | 9.40E-06 | 0.006892471 | 3.366511337 | Protein BPS1, chloroplastic | root cap | 0 h |
| HORVU.MOREX.r3.6HG0628890.1 | -1.568007098 | -2.722150267 | -4.949769109 | 7.92E-06 | 0.006097532 | 3.356220853 | Nicotianamine synthase | root cap | 0 h |
| HORVU.MOREX.r3.1HG0068380.1 | 0.559201929 | 5.314485157 | 4.946074853 | 8.02E-06 | 0.006097532 | 3.332950369 | #N/A | root cap | 0 h |
| HORVU.MOREX.r3.6HG0559690.1 | 1.292974786 | 2.590055839 | 4.841196551 | 1.16E-05 | 0.00791968 | 3.210011071 | Cell division cycle 5-like protein | root cap | 0 h |
| HORVU.MOREX.r3.7HG0679060.1 | -2.731993942 | 1.156648697 | -5.024302698 | 6.09E-06 | 0.005209649 | 3.208114798 | Nuclear transport factor 2 family protein, putative | root cap | 0 h |
| HORVU.MOREX.r3.1HG0012570.1 | -1.549175047 | -1.556736336 | -4.821807681 | 1.24E-05 | 0.008199768 | 3.144475753 | Chymotrypsin inhibitor | root cap | 0 h |
| HORVU.MOREX.r3.2HG0196720.1 | 1.344875712 | -0.225277908 | 4.807586579 | 1.30E-05 | 0.008346514 | 3.102298545 | XH/XS domain-containing protein | root cap | 0 h |
| HORVU.MOREX.r3.3HG0229300.1 | -1.500383909 | 1.779653302 | -4.731243511 | 1.70E-05 | 0.010547776 | 2.717609275 | Germin-like protein | root cap | 0 h |
| HORVU.MOREX.r3.1HG0024020.1 | 0.64615229 | 5.072656773 | 4.673785933 | 2.07E-05 | 0.012484071 | 2.481014965 | Heparan-alpha-glucosaminide N-acetyltransferase | root cap | 0 h |
| HORVU.MOREX.r3.5HG0425000.1 | -3.476618373 | -0.918831697 | -4.860106437 | 1.08E-05 | 0.007669821 | 2.36137305 | Transposon protein, putative, Pong sub-class | root cap | 0 h |
| HORVU.MOREX.r3.2HG0215830.1 | 0.909482307 | 2.436514446 | 4.433572634 | 4.70E-05 | 0.027159657 | 1.906518672 | CsAtPR5 | root cap | 0 h |
| HORVU.MOREX.r3.4HG0343000.1 | -0.734055922 | 2.745864292 | -4.421209371 | 4.90E-05 | 0.027159657 | 1.776200143 | Cyclin family protein | root cap | 0 h |
| HORVU.MOREX.r3.1HG0060370.1 | 2.167139952 | -0.902409718 | 4.268831162 | 8.16E-05 | 0.042958019 | 1.275393188 | 1,4-alpha-glucan branching enzyme GlgB | root cap | 0 h |
| HORVU.MOREX.r3.UnG0765280.1 | 0.91177688 | 2.589811572 | 4.223060034 | 9.51E-05 | 0.047576893 | 1.191925829 | F-box family protein | root cap | 0 h |
| HORVU.MOREX.r3.6HG0630920.1 | 2.981017522 | -2.009589986 | 4.410268098 | 5.08E-05 | 0.027439865 | 1.166346435 | NBS-LRR disease resistance protein-like | root cap | 0 h |
| HORVU.MOREX.r3.1HG0069090.1 | 2.855974492 | -3.256645115 | 4.423482203 | 4.86E-05 | 0.027159657 | 1.052861777 | Ribosomal protein S5/Elongation factor G/III/V family protein | root cap | 0 h |
| HORVU.MOREX.r3.1HG0069420.1 | 0.509256845 | 7.626353217 | 4.257942902 | 8.47E-05 | 0.04343025 | 1.002512481 | Cytochrome P450 | root cap | 0 h |
| HORVU.MOREX.r3.1HG0078900.1 | -2.049158051 | 5.8370464 | -25.5913251 | 1.65E-31 | 3.39E-27 | 59.40939911 | MATH domain-containing protein | root cap | 3 h |

**Table S5** Continued.

| ID | logFC | AveExpr | t | P.Value | adj.P.Val | B | gene description | root zone | time point |
|---|---|---|---|---|---|---|---|---|---|
| HORVU.MOREX.r3.1HG0067490.1 | -6.383840185 | -1.72649178 | -12.37876227 | 2.89E-17 | 2.96E-13 | 17.39549212 | Retrovirus-related Pol polyprotein from transposon TNT 1-94 | root cap | 3 h |
| HORVU.MOREX.r3.1HG0067040.1 | -0.948748924 | 7.101906906 | -8.133925411 | 6.83E-11 | 2.00E-07 | 14.64438446 | NAC domain protein, | root cap | 3 h |
| HORVU.MOREX.r3.5HG0475760.1 | -4.7787136 | -2.872794743 | -9.124845659 | 1.87E-12 | 1.28E-08 | 14.10459468 | Skp1 | root cap | 3 h |
| HORVU.MOREX.r3.7HG0748980.1 | 2.574044815 | -2.90728761 | 8.349935208 | 3.10E-11 | 1.06E-07 | 13.49933969 | 11S seed storage globulin A | root cap | 3 h |
| HORVU.MOREX.r3.7HG0679070.1 | -1.771150056 | 9.365871523 | -7.760381157 | 2.70E-10 | 6.93E-07 | 13.26377351 | Rab3 GTPase-activating protein catalytic subunit | root cap | 3 h |
| HORVU.MOREX.r3.7HG0727570.1 | 4.301563852 | -0.569333422 | 8.461585041 | 2.06E-11 | 8.45E-08 | 12.27770433 | ATP-dependent RNA helicase, putative | root cap | 3 h |
| HORVU.MOREX.r3.1HG0007570.1 | -4.984701776 | -2.580123327 | -8.492081443 | 1.84E-11 | 8.45E-08 | 9.988247475 | Werner Syndrome-like exonuclease | root cap | 3 h |
| HORVU.MOREX.r3.5HG0447830.1 | -0.765202966 | 4.419279148 | -6.446617635 | 3.50E-08 | 7.17E-05 | 8.664763017 | Sterile alpha motif (SAM) domain-containing protein | root cap | 3 h |
| HORVU.MOREX.r3.2HG0111680.1 | -4.451956379 | -2.082427636 | -7.310294255 | 1.43E-09 | 3.25E-06 | 7.313765456 | Formate--tetrahydrofolate ligase | root cap | 3 h |
| HORVU.MOREX.r3.3HG0280700.1 | 0.453504859 | 6.27769849 | 5.7487585 | 4.53E-07 | 0.000714932 | 6.090140023 | Katanin p60 ATPase-containing subunit A1 | root cap | 3 h |
| HORVU.MOREX.r3.7HG0679060.1 | -3.838166529 | 1.156648697 | -6.030511322 | 1.62E-07 | 0.000301782 | 5.455380293 | Nuclear transport factor 2 family protein, putative | root cap | 3 h |
| HORVU.MOREX.r3.7HG0695920.1 | 1.355366476 | 2.660691655 | 5.460487952 | 1.29E-06 | 0.001649904 | 5.241127218 | #N/A | root cap | 3 h |
| HORVU.MOREX.r3.2HG0215060.1 | 4.959945214 | -1.599980119 | 5.882627922 | 2.78E-07 | 0.000475349 | 4.727483489 | WAT1-related protein | root cap | 3 h |
| HORVU.MOREX.r3.3HG0306460.1 | 2.437108867 | -3.648540528 | 5.472775832 | 1.23E-06 | 0.001649904 | 4.390093937 | #N/A | root cap | 3 h |
| HORVU.MOREX.r3.2HG0186210.1 | -1.805650424 | 1.781727841 | -5.314095923 | 2.18E-06 | 0.002626649 | 4.245888499 | DUF1191 superfamily protein | root cap | 3 h |
| HORVU.MOREX.r3.6HG0559680.1 | -2.537697339 | 0.573580493 | -5.188963938 | 3.40E-06 | 0.0038769 | 4.02962174 | NADPH:quinone oxidoreductase | root cap | 3 h |
| HORVU.MOREX.r3.5HG0523880.1 | -3.666992223 | -2.522148569 | -5.578441515 | 8.40E-07 | 0.001231731 | 3.772507218 | cryptochrome-interacting basic-helix-loop-helix 5 | root cap | 3 h |
| HORVU.MOREX.r3.7HG0718080.1 | 1.111041113 | -3.103532351 | 5.125626538 | 4.26E-06 | 0.004369302 | 3.697404394 | Amino acid permease | root cap | 3 h |
| HORVU.MOREX.r3.2HG0170640.1 | 3.087621572 | -0.509068665 | 5.168782711 | 3.65E-06 | 0.003946062 | 3.623831661 | Protein FAR1-RELATED SEQUENCE 5 | root cap | 3 h |
| HORVU.MOREX.r3.6HG0628890.1 | -1.552503511 | -2.722150267 | -4.945152987 | 8.05E-06 | 0.007865085 | 3.340133295 | Nicotianamine synthase | root cap | 3 h |
| HORVU.MOREX.r3.4HG0363330.1 | 1.15043771 | 2.648390841 | 4.853547214 | 1.11E-05 | 0.010344293 | 3.24056058 | F-box/RNI-like superfamily protein | root cap | 3 h |
| HORVU.MOREX.r3.6HG0629160.1 | 0.964971655 | 2.999097227 | 4.647292824 | 2.27E-05 | 0.019374489 | 2.483946442 | Disease resistance protein RPM1 | root cap | 3 h |
| HORVU.MOREX.r3.6HG0559690.1 | 1.177159453 | 2.590055839 | 4.598857558 | 2.68E-05 | 0.020336863 | 2.436512865 | Cell division cycle 5-like protein | root cap | 3 h |
| HORVU.MOREX.r3.4HG0343330.1 | 3.794018323 | -0.333480115 | 4.605535487 | 2.62E-05 | 0.020336863 | 2.299370177 | Endonuclease III homolog | root cap | 3 h |
| HORVU.MOREX.r3.3HG0260150.1 | 1.812591182 | -0.115050093 | 4.722613126 | 1.75E-05 | 0.015592341 | 2.255830577 | Acylamino-acid-releasing enzyme | root cap | 3 h |
| HORVU.MOREX.r3.5HG0448120.1 | 1.262846698 | 1.313003386 | 4.501170221 | 3.73E-05 | 0.026427435 | 2.136736162 | Polyol transporter | root cap | 3 h |
| HORVU.MOREX.r3.7HG0649080.1 | 0.767022726 | 5.069910261 | 4.605261218 | 2.62E-05 | 0.020336863 | 2.091891208 | Peroxidase | root cap | 3 h |
| HORVU.MOREX.r3.2HG0135600.1 | -2.229351087 | -2.598497762 | -4.517288548 | 3.54E-05 | 0.025910754 | 1.975692355 | Chlorophyll a-b binding protein, chloroplastic | root cap | 3 h |
| HORVU.MOREX.r3.1HG0024020.1 | 0.596143047 | 5.072656773 | 4.337732035 | 6.49E-05 | 0.044360004 | 1.393976869 | Heparan-alpha-glucosaminide N-acetyltransferase | root cap | 3 h |
| HORVU.MOREX.r3.1HG0078900.1 | -1.905105788 | 5.8370464 | -23.73087579 | 6.59E-30 | 1.35E-25 | 57.10526724 | MATH domain-containing protein | root cap | 6 h |
| HORVU.MOREX.r3.1HG0067490.1 | -6.473047274 | -1.72649178 | -12.57987883 | 1.52E-17 | 1.56E-13 | 21.32052513 | Retrovirus-related Pol polyprotein from transposon TNT 1-94 | root cap | 6 h |
| HORVU.MOREX.r3.7HG0748980.1 | 3.078420767 | -2.90728761 | 9.576066553 | 3.73E-13 | 2.55E-09 | 18.03062108 | 11S seed storage globulin A | root cap | 6 h |
| HORVU.MOREX.r3.1HG0067040.1 | -1.060515911 | 7.101906906 | -9.063050876 | 2.33E-12 | 9.57E-09 | 17.88324005 | NAC domain protein, | root cap | 6 h |
| HORVU.MOREX.r3.5HG0447830.1 | -1.002200885 | 4.419279148 | -8.565800294 | 1.41E-11 | 4.82E-08 | 16.21493576 | Sterile alpha motif (SAM) domain-containing protein | root cap | 6 h |
| HORVU.MOREX.r3.5HG0475760.1 | -5.342607838 | -2.872794743 | -9.146451526 | 1.73E-12 | 8.86E-09 | 15.12050511 | Skp1 | root cap | 6 h |
| HORVU.MOREX.r3.3HG0225800.1 | 2.77197395 | -2.677835456 | 7.354417545 | 1.21E-09 | 3.55E-06 | 11.40915567 | Core-2/I-branching beta-1,6-N-acetylglucosaminyltransferase far | root cap | 6 h |
| HORVU.MOREX.r3.7HG0727570.1 | 3.259990996 | -0.569333422 | 7.306371149 | 1.45E-09 | 3.71E-06 | 10.79206579 | ATP-dependent RNA helicase, putative | root cap | 6 h |
| HORVU.MOREX.r3.1HG0007570.1 | -3.855234082 | -2.580123327 | -7.091254218 | 3.21E-09 | 7.32E-06 | 8.857110138 | Werner Syndrome-like exonuclease | root cap | 6 h |
| HORVU.MOREX.r3.6HG0576610.1 | 1.42396027 | 3.781355355 | 6.435078271 | 3.65E-08 | 5.76E-05 | 8.626959924 | LURP-one-like protein (DUF567) | root cap | 6 h |
| HORVU.MOREX.r3.5HG0523890.1 | -4.530447334 | -2.93597984 | -7.052336049 | 3.71E-09 | 7.61E-06 | 8.489291638 | Protein FAR1-RELATED SEQUENCE 5 | root cap | 6 h |
| HORVU.MOREX.r3.5HG0523880.1 | -4.368236389 | -2.522148569 | -6.849619152 | 7.87E-09 | 1.47E-05 | 8.286613515 | cryptochrome-interacting basic-helix-loop-helix 5 | root cap | 6 h |

**Table S5** Continued.

| ID | logFC | AveExpr | t | P.Value | adj.P.Val | B | gene description | root zone | time point |
|---|---|---|---|---|---|---|---|---|---|
| HORVU.MOREX.r3.2HG0111680.1 | -4.144731809 | -2.082427636 | -6.726225566 | 1.24E-08 | 2.12E-05 | 7.333570058 | Formate--tetrahydrofolate ligase | root cap | 6 h |
| HORVU.MOREX.r3.6HG0559680.1 | -3.143773538 | 0.573580493 | -6.077871376 | 1.36E-07 | 0.00017438 | 7.07251255 | NADPH:quinone oxidoreductase | root cap | 6 h |
| HORVU.MOREX.r3.7HG0679070.1 | -1.378115686 | 9.365871523 | -6.080124984 | 1.35E-07 | 0.00017438 | 6.977712719 | Rab3 GTPase-activating protein catalytic subunit | root cap | 6 h |
| HORVU.MOREX.r3.6HG0552590.1 | 1.643000496 | -2.145556214 | 5.909263246 | 2.52E-07 | 0.000304414 | 6.776063715 | Phototropic-responsive NPH3 family protein | root cap | 6 h |
| HORVU.MOREX.r3.2HG0215060.1 | 4.81737596 | -1.599980119 | 6.122529353 | 1.15E-07 | 0.000169142 | 6.51244959 | WAT1-related protein | root cap | 6 h |
| HORVU.MOREX.r3.7HG0644170.1 | 1.029409495 | -1.791009835 | 5.67257598 | 5.97E-07 | 0.000645235 | 5.902947131 | Transposon protein, putative, CACTA, En/Spm sub-class | root cap | 6 h |
| HORVU.MOREX.r3.7HG0649080.1 | 0.965171841 | 5.069910261 | 5.733061223 | 4.80E-07 | 0.000546688 | 5.80844451 | Peroxidase | root cap | 6 h |
| HORVU.MOREX.r3.1HG0060370.1 | 2.815729541 | -0.902409718 | 5.432737203 | 1.42E-06 | 0.001389128 | 4.949288258 | 1,4-alpha-glucan branching enzyme GlgB | root cap | 6 h |
| HORVU.MOREX.r3.7HG0723140.1 | 1.293486809 | -2.014032702 | 5.341215776 | 1.97E-06 | 0.001761756 | 4.790818327 | #N/A | root cap | 6 h |
| HORVU.MOREX.r3.2HG0170640.1 | 3.9698512 | -0.509068665 | 5.481843511 | 1.19E-06 | 0.001222164 | 4.621636959 | Protein FAR1-RELATED SEQUENCE 5 | root cap | 6 h |
| HORVU.MOREX.r3.7HG0695920.1 | 1.291739565 | 2.660691655 | 5.255225341 | 2.69E-06 | 0.002041083 | 4.541485572 | #N/A | root cap | 6 h |
| HORVU.MOREX.r3.4HG0343330.1 | 6.268033718 | -0.333480115 | 5.416526571 | 1.51E-06 | 0.001405587 | 4.539731684 | Endonuclease III homolog | root cap | 6 h |
| HORVU.MOREX.r3.3HG0280700.1 | 0.422045286 | 6.27769849 | 5.322885841 | 2.11E-06 | 0.001802915 | 4.434245491 | Katanin p60 ATPase-containing subunit A1 | root cap | 6 h |
| HORVU.MOREX.r3.6HG0550820.2 | 5.104844661 | -0.93362822 | 5.265526537 | 2.59E-06 | 0.002041083 | 4.373755268 | GDSL esterase/lipase | root cap | 6 h |
| HORVU.MOREX.r3.3HG0225830.1 | 1.202776609 | -1.658623447 | 5.183301698 | 3.47E-06 | 0.002542992 | 4.2321717 | Core-2/I-branching beta-1,6-N-acetylglucosaminyltransferase far | root cap | 6 h |
| HORVU.MOREX.r3.1HG0076850.1 | 1.853257222 | -2.352399703 | 5.144859825 | 3.98E-06 | 0.002814589 | 4.155886879 | Avr9/Cf-9 rapidly elicited protein | root cap | 6 h |
| HORVU.MOREX.r3.3HG0220770.1 | -0.909853286 | 6.392414316 | -5.284763285 | 2.42E-06 | 0.001983673 | 4.146893812 | Dipeptidyl peptidase family member 6 | root cap | 6 h |
| HORVU.MOREX.r3.6HG0559690.1 | 1.235844928 | 2.590055839 | 4.975707487 | 7.23E-06 | 0.004495764 | 3.592677419 | Cell division cycle 5-like protein | root cap | 6 h |
| HORVU.MOREX.r3.1HG0003270.1 | 1.481880615 | -2.891112437 | 4.955464158 | 7.76E-06 | 0.004685173 | 3.555318877 | Core-2/I-branching beta-1,6-N-acetylglucosaminyltransferase far | root cap | 6 h |
| HORVU.MOREX.r3.5HG0423130.1 | 2.044171189 | -2.684878217 | 4.942810317 | 8.12E-06 | 0.004758 | 3.498680716 | Thaumatin-like protein | root cap | 6 h |
| HORVU.MOREX.r3.1HG0078890.1 | 2.618879091 | -0.071217622 | 5.021417054 | 6.16E-06 | 0.004137923 | 3.493487467 | Ubiquitin carboxyl-terminal hydrolase 2 | root cap | 6 h |
| HORVU.MOREX.r3.3HG0306460.1 | 2.408791676 | -3.648540528 | 5.017035472 | 6.25E-06 | 0.004137923 | 3.415391408 | #N/A | root cap | 6 h |
| HORVU.MOREX.r3.5HG0479830.1 | -1.885238911 | -1.947529218 | -4.900460761 | 9.41E-06 | 0.005365784 | 3.398538424 | Cytochrome P450 | root cap | 6 h |
| HORVU.MOREX.r3.4HG0409190.1 | 2.740609543 | 1.165329523 | 4.993357397 | 6.79E-06 | 0.004357194 | 3.117221626 | Glutathione S-transferase | root cap | 6 h |
| HORVU.MOREX.r3.5HG0425000.1 | -2.745319596 | -0.918831697 | -4.819566889 | 1.25E-05 | 0.006568806 | 3.000563542 | Transposon protein, putative, Pong sub-class | root cap | 6 h |
| HORVU.MOREX.r3.5HG0489790.1 | 1.729913955 | -0.067483895 | 4.792584234 | 1.37E-05 | 0.007034697 | 2.997183911 | SKP1-like protein 4 | root cap | 6 h |
| HORVU.MOREX.r3.4HG0403870.1 | 1.115887457 | -2.784620446 | 4.76329599 | 1.52E-05 | 0.007597555 | 2.960366468 | Cysteine proteinase inhibitor | root cap | 6 h |
| HORVU.MOREX.r3.6HG0628890.1 | -1.353334416 | -2.722150267 | -4.700587535 | 1.89E-05 | 0.00899955 | 2.759250428 | Nicotianamine synthase | root cap | 6 h |
| HORVU.MOREX.r3.7HG0676800.1 | -0.440173088 | 6.777542145 | -4.867072001 | 1.06E-05 | 0.005867065 | 2.756918908 | formate dehydrogenase | root cap | 6 h |
| HORVU.MOREX.r3.1HG0069420.1 | 0.588761032 | 7.626353217 | 4.822387998 | 1.24E-05 | 0.006568806 | 2.692197104 | Cytochrome P450 | root cap | 6 h |
| HORVU.MOREX.r3.6HG0609250.1 | 1.551098974 | -2.575857699 | 4.678834511 | 2.03E-05 | 0.009269715 | 2.685685375 | Stigma-specific Stig1 family protein | root cap | 6 h |
| HORVU.MOREX.r3.7HG0679060.1 | -2.624088415 | 1.156648697 | -4.693586671 | 1.93E-05 | 0.009010121 | 2.553271747 | Nuclear transport factor 2 family protein, putative | root cap | 6 h |
| HORVU.MOREX.r3.1HG0076760.1 | 0.739718445 | 3.88919764 | 4.708824294 | 1.83E-05 | 0.008955432 | 2.53375467 | IQ domain-containing protein | root cap | 6 h |
| HORVU.MOREX.r3.7HG0724080.1 | 1.505414211 | 2.23197157 | 4.623830511 | 2.46E-05 | 0.010500511 | 2.483291017 | 6,7-dimethyl-8-ribityllumazine synthase | root cap | 6 h |
| HORVU.MOREX.r3.3HG0244320.1 | -2.674012215 | -2.865168374 | -4.660187334 | 2.17E-05 | 0.009598564 | 2.451635624 | Lachrymatory-factor synthase | root cap | 6 h |
| HORVU.MOREX.r3.7HG0749060.1 | 1.729470321 | 0.00150025 | 4.516383674 | 3.55E-05 | 0.014554787 | 2.179762807 | Transducin/WD40 repeat-like superfamily protein | root cap | 6 h |
| HORVU.MOREX.r3.1HG0069100.1 | 1.846276758 | -2.641840701 | 4.492769873 | 3.84E-05 | 0.015165223 | 2.056725151 | BZIP transcription factor | root cap | 6 h |
| HORVU.MOREX.r3.2HG0215830.1 | 0.926109433 | 2.436514446 | 4.506786099 | 3.66E-05 | 0.014743103 | 2.05260992 | CsAtPR5 | root cap | 6 h |
| HORVU.MOREX.r3.2HG0191320.1 | 2.9640787 | 1.356813235 | 4.656086323 | 2.20E-05 | 0.009598564 | 1.911901745 | Glucan endo-1,3-beta-glucosidase-like protein 3 | root cap | 6 h |
| HORVU.MOREX.r3.5HG0446460.1 | 1.613594534 | 0.590554613 | 4.424241984 | 4.85E-05 | 0.017854203 | 1.864622834 | Mitochondrial carrier protein, expressed | root cap | 6 h |
| HORVU.MOREX.r3.7HG0725700.1 | -3.356564102 | -1.181670666 | -4.54949582 | 3.17E-05 | 0.013266691 | 1.772500128 | Receptor-like protein kinase | root cap | 6 h |

**Table S5** Continued.

| ID | logFC | AveExpr | t | P.Value | adj.P.Val | B | gene description | root zone | time point |
|---|---|---|---|---|---|---|---|---|---|
| HORVU.MOREX.r3.2HG0175580.1 | 2.216988136 | 1.299950378 | 4.422707585 | 4.87E-05 | 0.017854203 | 1.676043836 | Aldo/keto reductase family oxidoreductase | root cap | 6 h |
| HORVU.MOREX.r3.2HG0132440.2 | -0.498359392 | 4.455296999 | -4.446445915 | 4.50E-05 | 0.017409918 | 1.612179926 | #N/A | root cap | 6 h |
| HORVU.MOREX.r3.5HG0502990.1 | 3.538883067 | 3.237955153 | 4.311887062 | 7.07E-05 | 0.024410973 | 1.459453342 | NAC domain protein, | root cap | 6 h |
| HORVU.MOREX.r3.1HG0034450.1 | -1.311503219 | 0.621391691 | -4.250713897 | 8.67E-05 | 0.028245893 | 1.369687759 | Retrovirus-related Pol polyprotein from transposon TNT 1-94 | root cap | 6 h |
| HORVU.MOREX.r3.1HG0067030.1 | -0.642872967 | 5.504670742 | -4.357508726 | 6.07E-05 | 0.021471697 | 1.320329842 | Protein trichome birefringence | root cap | 6 h |
| HORVU.MOREX.r3.6HG0601210.1 | 2.959072381 | -0.829190823 | 4.42327686 | 4.86E-05 | 0.017854203 | 1.248866905 | En/Spm transposon protein-like | root cap | 6 h |
| HORVU.MOREX.r3.6HG0544580.1 | 3.253885945 | 0.367989846 | 4.402636782 | 5.21E-05 | 0.018770204 | 1.225924613 | Werner Syndrome-like exonuclease | root cap | 6 h |
| HORVU.MOREX.r3.1HG0081920.1 | 0.922012939 | -2.160509685 | 4.233961362 | 9.17E-05 | 0.029396124 | 1.223479246 | Pollen Ole e 1 allergen/extensin | root cap | 6 h |
| HORVU.MOREX.r3.2HG0113590.1 | 0.571972113 | 3.8003454 | 4.284378752 | 7.75E-05 | 0.026077776 | 1.171842723 | Zinc finger CCCH domain protein, putative | root cap | 6 h |
| HORVU.MOREX.r3.3HG0245590.1 | 1.127912611 | -1.410291341 | 4.216621335 | 9.71E-05 | 0.03065756 | 1.132263772 | Ring finger protein, putative | root cap | 6 h |
| HORVU.MOREX.r3.6HG0572410.1 | 2.39959512 | -1.952913174 | 4.176637771 | 0.00011 | 0.034462619 | 1.060234282 | Peptidylprolyl isomerase | root cap | 6 h |
| HORVU.MOREX.r3.7HG0745510.1 | -0.401626073 | 5.787158675 | -4.309105143 | 7.14E-05 | 0.024410973 | 0.952275663 | Transcriptional corepressor SEUSS | root cap | 6 h |
| HORVU.MOREX.r3.1HG0062030.1 | 1.228525962 | -2.693593678 | 4.095262378 | 0.00014 | 0.040170981 | 0.912607799 | Dehydration responsive element binding transcription factor | root cap | 6 h |
| HORVU.MOREX.r3.7HG0739910.1 | 0.725625834 | 4.210713917 | 4.163210122 | 0.00012 | 0.034964133 | 0.911102394 | Disease resistance protein (NBS-LRR class) family | root cap | 6 h |
| HORVU.MOREX.r3.7HG0644060.1 | 4.132576443 | 3.296548784 | 4.266268571 | 8.23E-05 | 0.027253602 | 0.87667231 | Starch synthase, chloroplastic/amyloplastic | root cap | 6 h |
| HORVU.MOREX.r3.5HG0518370.1 | 0.848757546 | -1.963228378 | 4.170237538 | 0.00011 | 0.03467297 | 0.866667165 | BTB/POZ domain containing protein | root cap | 6 h |
| HORVU.MOREX.r3.1HG0069580.1 | 2.380284628 | -2.817290135 | 4.100231981 | 0.00014 | 0.040170981 | 0.835294348 | Gag-Pol polyprotein | root cap | 6 h |
| HORVU.MOREX.r3.7HG0678660.1 | 2.06524295 | 0.844122535 | 4.103859273 | 0.00014 | 0.040170981 | 0.830998435 | Protein kinase family protein | root cap | 6 h |
| HORVU.MOREX.r3.6HG0561420.1 | 1.5545289 | -3.217292483 | 4.055437995 | 0.00016 | 0.045137355 | 0.736120328 | Transposon protein, putative, Mutator sub-class | root cap | 6 h |
| HORVU.MOREX.r3.3HG0298070.1 | 3.218684852 | -2.093355386 | 4.132359661 | 0.00013 | 0.038140595 | 0.717396318 | Amino acid permease | root cap | 6 h |
| HORVU.MOREX.r3.5HG0509930.1 | 0.76678881 | 3.497515567 | 4.107053411 | 0.00014 | 0.040170981 | 0.66276904 | Zinc-transporting ATPase | root cap | 6 h |
| HORVU.MOREX.r3.5HG0520080.1 | 2.127642877 | -2.977064176 | 4.025094675 | 0.00018 | 0.049167601 | 0.576853223 | Kinase-like | root cap | 6 h |
| HORVU.MOREX.r3.3HG0220340.1 | 0.878729448 | 2.953247644 | 4.095222952 | 0.00014 | 0.040170981 | 0.511634139 | Disease resistance protein RPM1 | root cap | 6 h |
| HORVU.MOREX.r3.1HG0078900.1 | -1.860688339 | 5.8370464 | -22.36883507 | 1.14E-28 | 2.35E-24 | 54.25575787 | MATH domain-containing protein | root cap | 12 h |
| HORVU.MOREX.r3.1HG0067490.1 | -7.017751104 | -1.72649178 | -13.807562 | 3.40E-19 | 3.48E-15 | 23.23124234 | Retrovirus-related Pol polyprotein from transposon TNT 1-94 | root cap | 12 h |
| HORVU.MOREX.r3.7HG0748980.1 | 4.992720797 | -2.90728761 | 11.79582486 | 1.91E-16 | 1.31E-12 | 21.50283895 | 11S seed storage globulin A | root cap | 12 h |
| HORVU.MOREX.r3.1HG0067040.1 | -1.107535402 | 7.101906906 | -9.153177808 | 1.69E-12 | 8.65E-09 | 18.22984053 | NAC domain protein, | root cap | 12 h |
| HORVU.MOREX.r3.7HG0727570.1 | 4.231410401 | -0.569333422 | 8.279332645 | 4.01E-11 | 1.65E-07 | 13.0655516 | ATP-dependent RNA helicase, putative | root cap | 12 h |
| HORVU.MOREX.r3.5HG0475760.1 | -4.953424396 | -2.872794743 | -8.18757804 | 5.61E-11 | 1.92E-07 | 12.155674 | Skp1 | root cap | 12 h |
| HORVU.MOREX.r3.2HG0175580.1 | 3.354116578 | 1.299950378 | 7.121002037 | 2.88E-09 | 6.56E-06 | 11.0508038 | Aldo/keto reductase family oxidoreductase | root cap | 12 h |
| HORVU.MOREX.r3.3HG0280700.1 | 0.550692265 | 6.27769849 | 6.918609376 | 6.09E-09 | 1.14E-05 | 10.18636156 | Katanin p60 ATPase-containing subunit A1 | root cap | 12 h |
| HORVU.MOREX.r3.3HG0225800.1 | 2.932023177 | -2.677835456 | 7.032958629 | 3.99E-09 | 8.18E-06 | 10.09410244 | Core-2/I-branching beta-1,6-N-acetylglucosaminyltransferase far | root cap | 12 h |
| HORVU.MOREX.r3.1HG0007570.1 | -4.400085394 | -2.580123327 | -7.348284787 | 1.24E-09 | 3.63E-06 | 8.859679054 | Werner Syndrome-like exonuclease | root cap | 12 h |
| HORVU.MOREX.r3.2HG0111680.1 | -4.370135615 | -2.082427636 | -7.166342333 | 2.43E-09 | 6.24E-06 | 8.388492305 | Formate--tetrahydrofolate ligase | root cap | 12 h |
| HORVU.MOREX.r3.3HG0225830.1 | 1.640390941 | -1.658623447 | 6.009233187 | 1.75E-07 | 0.000251122 | 7.136309612 | Core-2/I-branching beta-1,6-N-acetylglucosaminyltransferase far | root cap | 12 h |
| HORVU.MOREX.r3.5HG0523880.1 | -4.163324659 | -2.522148569 | -6.476266222 | 3.13E-08 | 5.36E-05 | 7.099554497 | cryptochrome-interacting basic-helix-loop-helix 5 | root cap | 12 h |
| HORVU.MOREX.r3.4HG0382640.1 | 1.790785999 | -2.507767436 | 5.996034154 | 1.84E-07 | 0.000251122 | 6.874460067 | Mitochondrial import inner membrane translocase subunit Tim1 | root cap | 12 h |
| HORVU.MOREX.r3.5HG0447830.1 | -0.686440852 | 4.419279148 | -5.911403457 | 2.50E-07 | 0.00029337 | 6.68456388 | Sterile alpha motif (SAM) domain-containing protein | root cap | 12 h |
| HORVU.MOREX.r3.5HG0425000.1 | -5.175252186 | -0.918831697 | -6.356427969 | 4.88E-08 | 7.70E-05 | 6.593019236 | Transposon protein, putative, Pong sub-class | root cap | 12 h |
| HORVU.MOREX.r3.7HG0679070.1 | -1.389938158 | 9.365871523 | -5.903737311 | 2.57E-07 | 0.00029337 | 6.392269163 | Rab3 GTPase-activating protein catalytic subunit | root cap | 12 h |
| HORVU.MOREX.r3.7HG0695920.1 | 1.313948132 | 2.660691655 | 5.76748266 | 4.23E-07 | 0.000456931 | 6.287331113 | #N/A | root cap | 12 h |

**Table S5** Continued.

| ID | logFC | AveExpr | t | P.Value | adj.P.Val | B | gene description | root zone | time point |
|---|---|---|---|---|---|---|---|---|---|
| HORVU.MOREX.r3.6HG0559680.1 | -2.805018916 | 0.573580493 | -5.707287109 | 5.27E-07 | 0.000540359 | 5.903888502 | NADPH:quinone oxidoreductase | root cap | 12 h |
| HORVU.MOREX.r3.5HG0489790.1 | 1.908347936 | -0.067483895 | 5.651049133 | 6.46E-07 | 0.000631218 | 5.748701508 | SKP1-like protein 4 | root cap | 12 h |
| HORVU.MOREX.r3.5HG0523890.1 | -3.89806797 | -2.93597984 | -5.924538516 | 2.38E-07 | 0.00029337 | 5.238937919 | Protein FAR1-RELATED SEQUENCE 5 | root cap | 12 h |
| HORVU.MOREX.r3.2HG0113590.1 | 0.786254625 | 3.8003454 | 5.390998159 | 1.65E-06 | 0.001540615 | 4.93093729 | Zinc finger CCCH domain protein, putative | root cap | 12 h |
| HORVU.MOREX.r3.1HG0060370.1 | 3.217276087 | -0.902409718 | 5.377109252 | 1.74E-06 | 0.001548973 | 4.541704472 | 1,4-alpha-glucan branching enzyme GlgB | root cap | 12 h |
| HORVU.MOREX.r3.7HG0725940.1 | -1.639235011 | -1.945827342 | -5.128839728 | 4.21E-06 | 0.003455844 | 4.140817413 | protein kinase family protein | root cap | 12 h |
| HORVU.MOREX.r3.4HG0343330.1 | 5.767619043 | -0.333480115 | 5.205803365 | 3.20E-06 | 0.00273855 | 3.951141896 | Endonuclease III homolog | root cap | 12 h |
| HORVU.MOREX.r3.6HG0559690.1 | 1.278339865 | 2.590055839 | 4.989977941 | 6.88E-06 | 0.00542685 | 3.659731614 | Cell division cycle 5-like protein | root cap | 12 h |
| HORVU.MOREX.r3.6HG0540710.1 | 0.951452883 | 1.044028848 | 4.862110717 | 1.08E-05 | 0.00745542 | 3.263468096 | Disease resistance protein (TIR-NBS-LRR class) family | root cap | 12 h |
| HORVU.MOREX.r3.2HG0215060.1 | 3.287258171 | -1.599980119 | 4.927538272 | 8.56E-06 | 0.006507102 | 3.240359655 | WAT1-related protein | root cap | 12 h |
| HORVU.MOREX.r3.7HG0724080.1 | 1.461771759 | 2.23197157 | 4.845016446 | 1.14E-05 | 0.007562815 | 3.185745841 | 6,7-dimethyl-8-ribityllumazine synthase | root cap | 12 h |
| HORVU.MOREX.r3.UnG0783730.1 | 1.374590715 | 6.739233425 | 4.889572135 | 9.78E-06 | 0.007166702 | 2.981034384 | Histone H2B | root cap | 12 h |
| HORVU.MOREX.r3.5HG0511210.1 | -1.672657715 | 1.506867328 | -4.735462662 | 1.67E-05 | 0.010089164 | 2.861889321 | MADS box transcription factor | root cap | 12 h |
| HORVU.MOREX.r3.5HG0518370.1 | 1.241226206 | -1.963228378 | 4.746033134 | 1.61E-05 | 0.010021345 | 2.861746518 | BTB/POZ domain containing protein | root cap | 12 h |
| HORVU.MOREX.r3.6HG0630920.1 | 3.184249492 | -2.009589986 | 4.858516336 | 1.09E-05 | 0.00745542 | 2.797978419 | NBS-LRR disease resistance protein-like | root cap | 12 h |
| HORVU.MOREX.r3.5HG0498610.1 | 1.772925053 | -3.32898876 | 4.788734549 | 1.39E-05 | 0.008911768 | 2.593032399 | Protein Mei2 | root cap | 12 h |
| HORVU.MOREX.r3.1HG0070000.1 | 0.947071992 | 1.808570831 | 4.582695162 | 2.83E-05 | 0.015314464 | 2.368356651 | Histone H3 | root cap | 12 h |
| HORVU.MOREX.r3.2HG0197550.1 | 1.009479028 | -0.290355644 | 4.67852934 | 2.03E-05 | 0.011930748 | 2.321526759 | Transmembrane protein, putative | root cap | 12 h |
| HORVU.MOREX.r3.6HG0633910.1 | 1.705820626 | -2.820374223 | 4.581885972 | 2.84E-05 | 0.015314464 | 2.318850106 | Protein SAWADEE HOMEODOMAIN-like protein 2 | root cap | 12 h |
| HORVU.MOREX.r3.2HG0132820.1 | -3.434720032 | 1.249014446 | -4.627649298 | 2.42E-05 | 0.013818314 | 2.285329858 | Phenylalanine ammonia-lyase | root cap | 12 h |
| HORVU.MOREX.r3.7HG0749060.1 | 1.918352159 | 0.00150025 | 4.506366673 | 3.67E-05 | 0.017927897 | 2.114753098 | Transducin/WD40 repeat-like superfamily protein | root cap | 12 h |
| HORVU.MOREX.r3.2HG0178180.1 | 1.883069531 | -3.517770368 | 4.539256759 | 3.28E-05 | 0.017260972 | 1.967631887 | Proteasome subunit beta type-9 | root cap | 12 h |
| HORVU.MOREX.r3.7HG0635050.1 | 0.859260477 | 0.346305785 | 4.509043283 | 3.64E-05 | 0.017927897 | 1.846757133 | NBS-LRR-like resistance protein | root cap | 12 h |
| HORVU.MOREX.r3.1HG0095330.1 | 0.908300605 | 2.512196306 | 4.491379774 | 3.86E-05 | 0.018426154 | 1.774637506 | Zinc finger CCCH domain protein | root cap | 12 h |
| HORVU.MOREX.r3.6HG0629160.1 | 0.954897274 | 2.999097227 | 4.404505237 | 5.18E-05 | 0.023112653 | 1.595978201 | Disease resistance protein RPM1 | root cap | 12 h |
| HORVU.MOREX.r3.6HG0614490.1 | 0.800966667 | 2.880979216 | 4.443934219 | 4.54E-05 | 0.020680043 | 1.51683545 | Type I inositol-1,4,5-trisphosphate 5-phosphatase | root cap | 12 h |
| HORVU.MOREX.r3.1HG0070730.1 | -3.52859849 | -0.4130893 | -4.454534191 | 4.38E-05 | 0.020404574 | 1.41874584 | Glycosyltransferase | root cap | 12 h |
| HORVU.MOREX.r3.2HG0191320.1 | -3.06833176 | 1.356813235 | -4.52472845 | 3.45E-05 | 0.017683779 | 1.378952424 | Glucan endo-1,3-beta-glucosidase-like protein 3 | root cap | 12 h |
| HORVU.MOREX.r3.2HG0215830.1 | 0.825847403 | 2.436514446 | 4.290078812 | 7.61E-05 | 0.031852538 | 1.342456969 | CsAtPR5 | root cap | 12 h |
| HORVU.MOREX.r3.6HG0565970.1 | 0.78319336 | 4.099013656 | 4.308926272 | 7.14E-05 | 0.030531971 | 1.31008281 | Early nodulin-like protein | root cap | 12 h |
| HORVU.MOREX.r3.3HG0291580.1 | 0.686909508 | 2.805578274 | 4.221238262 | 9.56E-05 | 0.037739864 | 1.135751902 | Lysosomal Pro-X carboxypeptidase | root cap | 12 h |
| HORVU.MOREX.r3.1HG0092900.1 | 1.626394045 | -2.962290826 | 4.200740505 | 0.0001 | 0.039629343 | 1.066277888 | C3HC4-type RING finger-containing protein | root cap | 12 h |
| HORVU.MOREX.r3.3HG0240580.1 | 0.969073804 | 2.986398292 | 4.272358538 | 8.07E-05 | 0.033115789 | 1.059836153 | Glycosyltransferase | root cap | 12 h |
| HORVU.MOREX.r3.6HG0548800.1 | 1.27034093 | -1.579734173 | 4.180293118 | 0.00011 | 0.040252358 | 1.039195703 | transcription factor IIIA | root cap | 12 h |
| HORVU.MOREX.r3.3HG0220340.1 | 1.000045067 | 2.953247644 | 4.233693225 | 9.18E-05 | 0.036922107 | 1.038302764 | Disease resistance protein RPM1 | root cap | 12 h |
| HORVU.MOREX.r3.6HG0552590.1 | 1.388142618 | -2.145556214 | 4.12711859 | 0.00013 | 0.044608301 | 0.945928611 | Phototropic-responsive NPH3 family protein | root cap | 12 h |
| HORVU.MOREX.r3.2HG0099100.1 | 1.945926633 | -3.276482409 | 4.180054467 | 0.00011 | 0.040252358 | 0.941271562 | Serine/threonine-protein phosphatase 7 long form-like protein | root cap | 12 h |
| HORVU.MOREX.r3.5HG0495750.1 | 2.159968158 | -2.868788994 | 4.151707368 | 0.00012 | 0.043322914 | 0.914374044 | B3 domain-containing protein | root cap | 12 h |
| HORVU.MOREX.r3.4HG0363330.1 | 1.075073102 | 2.648390841 | 4.127755767 | 0.00013 | 0.044608301 | 0.898258513 | F-box/RNI-like superfamily protein | root cap | 12 h |
| HORVU.MOREX.r3.3HG0286140.1 | 1.000293083 | 3.423023169 | 4.335507385 | 6.53E-05 | 0.028526829 | 0.840601977 | Pollen Ole e 1 allergen/extensin | root cap | 12 h |
| HORVU.MOREX.r3.3HG0258640.1 | 1.069871576 | -1.96360197 | 4.072865425 | 0.00016 | 0.04951996 | 0.83868399 | Kelch repeat protein, putative | root cap | 12 h |

**Table S5** Continued.

| ID | logFC | AveExpr | t | P.Value | adj.P.Val | B | gene description | root zone | time point |
|---|---|---|---|---|---|---|---|---|---|
| HORVU.MOREX.r3.1HG0068380.1 | 0.486285361 | 5.314485157 | 4.179366439 | 0.00011 | 0.040252358 | 0.689411461 | #N/A | root cap | 12 h |
| HORVU.MOREX.r3.3HG0311610.1 | 1.596124021 | -1.025604193 | 4.061610029 | 0.00016 | 0.04951996 | 0.661533667 | Glutathione S-transferase | root cap | 12 h |
| HORVU.MOREX.r3.4HG0353070.1 | 1.038153513 | 0.247089748 | 4.107219961 | 0.00014 | 0.04536637 | 0.607847918 | Phototropic-responsive NPH3 family protein | root cap | 12 h |
| HORVU.MOREX.r3.6HG0575750.1 | 3.449962273 | -0.706156741 | 4.122189733 | 0.00013 | 0.044608301 | 0.554110927 | Cytochrome P450 family protein | root cap | 12 h |
| HORVU.MOREX.r3.7HG0728160.1 | 0.720644627 | 5.080435682 | 4.110797805 | 0.00014 | 0.04536637 | 0.457283644 | Mitogen-activated protein kinase | root cap | 12 h |
| HORVU.MOREX.r3.2HG0125600.1 | 0.841028064 | -0.842945351 | 4.064614401 | 0.00016 | 0.04951996 | 0.401703192 | DNA-directed RNA polymerase subunit alpha | root cap | 12 h |
| HORVU.MOREX.r3.7HG0651330.1 | 0.933042619 | 5.54057482 | 4.145306698 | 0.00012 | 0.043482632 | 0.382147406 | Apyrase | root cap | 12 h |
| HORVU.MOREX.r3.5HG0455240.1 | 0.852403177 | 2.696541261 | 4.063246318 | 0.00016 | 0.04951996 | 0.323663583 | O-fucosyltransferase family protein | root cap | 12 h |
| HORVU.MOREX.r3.1HG0078900.1 | -2.18941927 | 5.8370464 | -24.93737725 | 5.88E-31 | 1.21E-26 | 57.7066814 | MATH domain-containing protein | meristem | 0 h |
| HORVU.MOREX.r3.3HG0280700.1 | 0.753498871 | 6.27769849 | 10.0313638 | 7.52E-14 | 5.14E-10 | 21.32651553 | Katanin p60 ATPase-containing subunit A1 | meristem | 0 h |
| HORVU.MOREX.r3.5HG0447830.1 | -1.109883294 | 4.419279148 | -9.636797126 | 3.01E-13 | 1.54E-09 | 19.85482732 | Sterile alpha motif (SAM) domain-containing protein | meristem | 0 h |
| HORVU.MOREX.r3.1HG0067490.1 | -6.929389502 | -1.72649178 | -13.54705862 | 7.49E-19 | 7.68E-15 | 19.12319322 | Retrovirus-related Pol polyprotein from transposon TNT 1-94 | meristem | 0 h |
| HORVU.MOREX.r3.1HG0067040.1 | -1.016390302 | 7.101906906 | -8.9505017 | 3.50E-12 | 1.43E-08 | 17.55857713 | NAC domain protein, | meristem | 0 h |
| HORVU.MOREX.r3.7HG0727570.1 | 5.579475085 | -0.569333422 | 8.809521826 | 5.82E-12 | 1.99E-08 | 11.2003243 | ATP-dependent RNA helicase, putative | meristem | 0 h |
| HORVU.MOREX.r3.7HG0695920.1 | 1.723301085 | 2.660691655 | 7.137694518 | 2.70E-09 | 6.94E-06 | 10.96505618 | #N/A | meristem | 0 h |
| HORVU.MOREX.r3.2HG0111680.1 | -4.776616021 | -2.082427636 | -7.878325928 | 1.75E-10 | 5.13E-07 | 8.550309124 | Formate--tetrahydrofolate ligase | meristem | 0 h |
| HORVU.MOREX.r3.1HG0011440.1 | -3.170933054 | -1.581106574 | -6.619075132 | 1.85E-08 | 4.21E-05 | 7.636262692 | Glucan endo-1,3-beta-glucosidase, putative | meristem | 0 h |
| HORVU.MOREX.r3.7HG0739910.1 | 0.835184976 | 4.210713917 | 6.140077736 | 1.08E-07 | 0.000181892 | 7.572146015 | Disease resistance protein (NBS-LRR class) family | meristem | 0 h |
| HORVU.MOREX.r3.6HG0559690.1 | 1.572989065 | 2.590055839 | 6.102748126 | 1.24E-07 | 0.000181892 | 7.405751997 | Cell division cycle 5-like protein | meristem | 0 h |
| HORVU.MOREX.r3.7HG0640470.1 | 0.489930266 | 6.260306545 | 5.833312239 | 3.33E-07 | 0.000426848 | 6.354846771 | WPP domain-interacting tail-anchored protein 1 | meristem | 0 h |
| HORVU.MOREX.r3.2HG0170640.1 | 3.723898226 | -0.509068665 | 5.994626223 | 1.85E-07 | 0.000252421 | 5.847058386 | Protein FAR1-RELATED SEQUENCE 5 | meristem | 0 h |
| HORVU.MOREX.r3.6HG0546230.1 | 5.085801563 | -1.679648171 | 6.366968932 | 4.69E-08 | 8.75E-05 | 5.558806213 | Disease resistance protein RPM1 | meristem | 0 h |
| HORVU.MOREX.r3.1HG0007570.1 | -4.032039242 | -2.580123327 | -6.559325583 | 2.30E-08 | 4.73E-05 | 5.460801058 | Werner Syndrome-like exonuclease | meristem | 0 h |
| HORVU.MOREX.r3.1HG0024020.1 | 0.741911468 | 5.072656773 | 5.425302687 | 1.46E-06 | 0.001762489 | 5.028488511 | Heparan-alpha-glucosaminide N-acetyltransferase | meristem | 0 h |
| HORVU.MOREX.r3.2HG0215830.1 | 1.087989707 | 2.436514446 | 5.344623867 | 1.95E-06 | 0.002223807 | 4.863223805 | CsAtPR5 | meristem | 0 h |
| HORVU.MOREX.r3.5HG0523880.1 | -4.277013001 | -2.522148569 | -6.115176206 | 1.19E-07 | 0.000181892 | 4.754025334 | cryptochrome-interacting basic-helix-loop-helix 5 | meristem | 0 h |
| HORVU.MOREX.r3.1HG0026900.1 | -1.247644411 | 1.690413985 | -5.315417037 | 2.17E-06 | 0.002311762 | 4.702054786 | Protein apaG | meristem | 0 h |
| HORVU.MOREX.r3.2HG0216110.1 | 0.47731075 | 5.350102781 | 5.304364239 | 2.25E-06 | 0.002311762 | 4.58568552 | #N/A | meristem | 0 h |
| HORVU.MOREX.r3.3HG0220340.1 | 1.557242415 | 2.953247644 | 5.081125008 | 4.99E-06 | 0.004871474 | 3.989898487 | Disease resistance protein RPM1 | meristem | 0 h |
| HORVU.MOREX.r3.1HG0068380.1 | 0.583137921 | 5.314485157 | 4.863947993 | 1.07E-05 | 0.009975551 | 3.102506262 | #N/A | meristem | 0 h |
| HORVU.MOREX.r3.1HG0049550.1 | 0.527322497 | 3.543555344 | 4.737558429 | 1.66E-05 | 0.01362209 | 2.836982872 | Protein BPS1, chloroplastic | meristem | 0 h |
| HORVU.MOREX.r3.6HG0559680.1 | -2.33163652 | 0.573580493 | -4.747859363 | 1.60E-05 | 0.01362209 | 2.707498062 | NADPH:quinone oxidoreductase | meristem | 0 h |
| HORVU.MOREX.r3.7HG0724080.1 | 1.482851842 | 2.23197157 | 4.511674398 | 3.60E-05 | 0.023854751 | 2.169251024 | 6,7-dimethyl-8-ribityllumazine synthase | meristem | 0 h |
| HORVU.MOREX.r3.3HG0288140.1 | 0.412434522 | 6.173911395 | 4.616621976 | 2.52E-05 | 0.019871122 | 2.167480328 | Golgin candidate 1 | meristem | 0 h |
| HORVU.MOREX.r3.1HG0060370.1 | 2.553201532 | -0.902409718 | 4.601640437 | 2.65E-05 | 0.020143851 | 2.077762197 | 1,4-alpha-glucan branching enzyme GlgB | meristem | 0 h |
| HORVU.MOREX.r3.1HG0067030.1 | -0.62418381 | 5.504670742 | -4.532564211 | 3.36E-05 | 0.022957163 | 2.010991743 | Protein trichome birefringence | meristem | 0 h |
| HORVU.MOREX.r3.5HG0502990.1 | 3.908091076 | 3.237955153 | 4.420153102 | 4.91E-05 | 0.029661712 | 1.886697685 | NAC domain protein, | meristem | 0 h |
| HORVU.MOREX.r3.6HG0568640.1 | 0.463735712 | 4.543924675 | 4.477023678 | 4.05E-05 | 0.025996858 | 1.863384785 | Methyltransferase domain protein | meristem | 0 h |
| HORVU.MOREX.r3.5HG0425000.1 | -2.988900341 | -0.918831697 | -4.572067762 | 2.93E-05 | 0.021493761 | 1.7831053 | Transposon protein, putative, Pong sub-class | meristem | 0 h |
| HORVU.MOREX.r3.6HG0629160.1 | 1.106297761 | 2.999097227 | 4.39709699 | 5.31E-05 | 0.031144818 | 1.768107482 | Disease resistance protein RPM1 | meristem | 0 h |
| HORVU.MOREX.r3.5HG0471070.1 | -0.341124774 | 6.510820563 | -4.467849354 | 4.18E-05 | 0.026005847 | 1.657009205 | Oxidoreductase family, NAD-binding rossmann fold protein | meristem | 0 h |

**Table S5** Continued.

| ID | logFC | AveExpr | t | P.Value | adj.P.Val | B | gene description | root zone | time point |
|---|---|---|---|---|---|---|---|---|---|
| HORVU.MOREX.r3.3HG0285150.1 | 1.81137796 | 1.826068882 | 4.332369067 | 6.60E-05 | 0.037637117 | 1.568274079 | Tryptophan aminotransferase | meristem | 0 h |
| HORVU.MOREX.r3.5HG0523890.1 | -3.412504681 | -2.93597984 | -4.810357757 | 1.29E-05 | 0.01150114 | 1.432936023 | Protein FAR1-RELATED SEQUENCE 5 | meristem | 0 h |
| HORVU.MOREX.r3.5HG0516660.1 | 0.469172516 | 4.126867737 | 4.31814388 | 6.93E-05 | 0.03840644 | 1.405890637 | LEM3 (Ligand-effect modulator 3) family protein | meristem | 0 h |
| HORVU.MOREX.r3.1HG0086120.1 | 1.215770892 | 4.055301306 | 4.292834704 | 7.54E-05 | 0.04049761 | 1.302483156 | Histone H2B | meristem | 0 h |
| HORVU.MOREX.r3.1HG0044150.1 | 0.452833383 | 5.220215331 | 4.286522696 | 7.70E-05 | 0.04049761 | 1.21974384 | Ran-binding protein in the microtubule-organising centre protein | meristem | 0 h |
| HORVU.MOREX.r3.6HG0572410.1 | 3.500578192 | -1.952913174 | 4.559597791 | 3.06E-05 | 0.021656115 | 1.168643326 | Peptidylprolyl isomerase | meristem | 0 h |
| HORVU.MOREX.r3.1HG0078900.1 | -2.187273592 | 5.8370464 | -24.75184593 | 8.47E-31 | 1.74E-26 | 56.49450737 | MATH domain-containing protein | meristem | 3 h |
| HORVU.MOREX.r3.5HG0447830.1 | -1.334440822 | 4.419279148 | -11.27170688 | 1.08E-15 | 7.40E-12 | 24.95374435 | Sterile alpha motif (SAM) domain-containing protein | meristem | 3 h |
| HORVU.MOREX.r3.1HG0067040.1 | -1.069173544 | 7.101906906 | -9.224199954 | 1.31E-12 | 6.71E-09 | 18.53434344 | NAC domain protein, | meristem | 3 h |
| HORVU.MOREX.r3.1HG0067490.1 | -6.573128045 | -1.72649178 | -12.83803928 | 6.73E-18 | 6.90E-14 | 16.21875274 | Retrovirus-related Pol polyprotein from transposon TNT 1-94 | meristem | 3 h |
| HORVU.MOREX.r3.3HG0280700.1 | 0.560771909 | 6.27769849 | 7.448451638 | 8.55E-10 | 2.51E-06 | 12.22075425 | Katanin p60 ATPase-containing subunit A1 | meristem | 3 h |
| HORVU.MOREX.r3.7HG0727570.1 | 4.484418745 | -0.569333422 | 8.255716277 | 4.37E-11 | 1.79E-07 | 10.36056584 | ATP-dependent RNA helicase, putative | meristem | 3 h |
| HORVU.MOREX.r3.1HG0026900.1 | -1.674423898 | 1.690413985 | -6.954094546 | 5.34E-09 | 1.37E-05 | 9.851473109 | Protein apaG | meristem | 3 h |
| HORVU.MOREX.r3.7HG0695920.1 | 1.514843807 | 2.660691655 | 6.691892865 | 1.41E-08 | 2.90E-05 | 9.395962711 | #N/A | meristem | 3 h |
| HORVU.MOREX.r3.2HG0111680.1 | -4.821699509 | -2.082427636 | -8.067416399 | 8.72E-11 | 2.98E-07 | 8.043258442 | Formate--tetrahydrofolate ligase | meristem | 3 h |
| HORVU.MOREX.r3.6HG0559680.1 | -3.65460718 | 0.573580493 | -6.282987397 | 6.39E-08 | 0.000119268 | 6.464081395 | NADPH:quinone oxidoreductase | meristem | 3 h |
| HORVU.MOREX.r3.5HG0523880.1 | -4.815067675 | -2.522148569 | -6.832129409 | 8.39E-09 | 1.91E-05 | 5.582442447 | cryptochrome-interacting basic-helix-loop-helix 5 | meristem | 3 h |
| HORVU.MOREX.r3.1HG0067030.1 | -0.763654599 | 5.504670742 | -5.422627172 | 1.47E-06 | 0.0020168 | 5.073605184 | Protein trichome birefringence | meristem | 3 h |
| HORVU.MOREX.r3.7HG0739910.1 | 0.701562223 | 4.210713917 | 5.199965078 | 3.27E-06 | 0.003728078 | 4.342699681 | Disease resistance protein (NBS-LRR class) family | meristem | 3 h |
| HORVU.MOREX.r3.6HG0546230.1 | 5.188005922 | -1.679648171 | 6.199783949 | 8.69E-08 | 0.00014853 | 4.258821364 | Disease resistance protein RPM1 | meristem | 3 h |
| HORVU.MOREX.r3.2HG0170640.1 | 3.199505282 | -0.509068665 | 5.440581317 | 1.38E-06 | 0.0020168 | 4.126352675 | Protein FAR1-RELATED SEQUENCE 5 | meristem | 3 h |
| HORVU.MOREX.r3.5HG0478990.1 | -1.133688844 | 3.637153099 | -5.000159182 | 6.63E-06 | 0.006806588 | 3.710624046 | Kinase family protein | meristem | 3 h |
| HORVU.MOREX.r3.5HG0471070.1 | -0.380360853 | 6.510820563 | -4.985255031 | 6.99E-06 | 0.006831523 | 3.4600243 | Oxidoreductase family, NAD-binding rossmann fold protein | meristem | 3 h |
| HORVU.MOREX.r3.1HG0011440.1 | -2.584902877 | -1.581106574 | -5.172151125 | 3.61E-06 | 0.003899108 | 3.1456579 | Glucan endo-1,3-beta-glucosidase, putative | meristem | 3 h |
| HORVU.MOREX.r3.3HG0223920.1 | 3.476345648 | -2.112939632 | 5.271772231 | 2.53E-06 | 0.003055756 | 3.102856736 | E3 ubiquitin-protein ligase | meristem | 3 h |
| HORVU.MOREX.r3.1HG0049550.1 | 0.508661675 | 3.543555344 | 4.632014235 | 2.39E-05 | 0.022275341 | 2.525616419 | Protein BPS1, chloroplastic | meristem | 3 h |
| HORVU.MOREX.r3.1HG0007570.1 | -3.409555231 | -2.580123327 | -5.463055643 | 1.27E-06 | 0.002011957 | 2.396128103 | Werner Syndrome-like exonuclease | meristem | 3 h |
| HORVU.MOREX.r3.5HG0523890.1 | -3.678873668 | -2.93597984 | -5.318514045 | 2.14E-06 | 0.00274703 | 2.157912709 | Protein FAR1-RELATED SEQUENCE 5 | meristem | 3 h |
| HORVU.MOREX.r3.1HG0017480.1 | 2.554456701 | 0.957708265 | 4.515443616 | 3.56E-05 | 0.031742252 | 2.129593347 | Peroxidase | meristem | 3 h |
| HORVU.MOREX.r3.6HG0559690.1 | 1.063940877 | 2.590055839 | 4.435813166 | 4.66E-05 | 0.038260777 | 1.932199372 | Cell division cycle 5-like protein | meristem | 3 h |
| HORVU.MOREX.r3.5HG0425000.1 | -2.937718782 | -0.918831697 | -4.493478279 | 3.83E-05 | 0.032778993 | 1.348701347 | Transposon protein, putative, Pong sub-class | meristem | 3 h |
| HORVU.MOREX.r3.1HG0078900.1 | -2.227245044 | 5.8370464 | -25.10525122 | 4.23E-31 | 8.69E-27 | 58.88620202 | MATH domain-containing protein | meristem | 6 h |
| HORVU.MOREX.r3.1HG0067040.1 | -1.236115214 | 7.101906906 | -10.63411971 | 9.35E-15 | 6.40E-11 | 23.36988965 | NAC domain protein, | meristem | 6 h |
| HORVU.MOREX.r3.5HG0447830.1 | -1.175827184 | 4.419279148 | -10.00606528 | 8.22E-14 | 4.21E-10 | 21.19439009 | Sterile alpha motif (SAM) domain-containing protein | meristem | 6 h |
| HORVU.MOREX.r3.1HG0067490.1 | -6.610847604 | -1.72649178 | -12.85971473 | 6.29E-18 | 6.45E-14 | 20.65873986 | Retrovirus-related Pol polyprotein from transposon TNT 1-94 | meristem | 6 h |
| HORVU.MOREX.r3.3HG0280700.1 | 0.613176273 | 6.27769849 | 8.081329267 | 8.29E-11 | 2.83E-07 | 14.42197547 | Katanin p60 ATPase-containing subunit A1 | meristem | 6 h |
| HORVU.MOREX.r3.1HG0067030.1 | -1.085004777 | 5.504670742 | -7.643174947 | 4.16E-10 | 1.07E-06 | 12.90352646 | Protein trichome birefringence | meristem | 6 h |
| HORVU.MOREX.r3.7HG0727570.1 | 5.25170781 | -0.569333422 | 8.180207516 | 5.76E-11 | 2.37E-07 | 11.02020211 | ATP-dependent RNA helicase, putative | meristem | 6 h |
| HORVU.MOREX.r3.2HG0111680.1 | -4.83556055 | -2.082427636 | -8.020814481 | 1.03E-10 | 3.03E-07 | 10.21321374 | Formate--tetrahydrofolate ligase | meristem | 6 h |
| HORVU.MOREX.r3.7HG0695920.1 | 1.595960863 | 2.660691655 | 6.642378699 | 1.69E-08 | 3.86E-05 | 9.344165472 | #N/A | meristem | 6 h |
| HORVU.MOREX.r3.5HG0471070.1 | -0.472783869 | 6.510820563 | -6.131912256 | 1.11E-07 | 0.000228773 | 7.311705316 | Oxidoreductase family, NAD-binding rossmann fold protein | meristem | 6 h |

**Table S5** Continued.

| ID | logFC | AveExpr | t | P.Value | adj.P.Val | B | gene description | root zone | time point |
|---|---|---|---|---|---|---|---|---|---|
| HORVU.MOREX.r3.6HG0559680.1 | -2.917450981 | 0.573580493 | -5.446020584 | 1.36E-06 | 0.002139234 | 4.907623967 | NADPH:quinone oxidoreductase | meristem | 6 h |
| HORVU.MOREX.r3.6HG0559690.1 | 1.387729973 | 2.590055839 | 5.2743685 | 2.51E-06 | 0.003107881 | 4.626466926 | Cell division cycle 5-like protein | meristem | 6 h |
| HORVU.MOREX.r3.6HG0546230.1 | 4.655478069 | -1.679648171 | 5.719925291 | 5.03E-07 | 0.000860158 | 4.586915689 | Disease resistance protein RPM1 | meristem | 6 h |
| HORVU.MOREX.r3.5HG0523880.1 | -4.184534518 | -2.522148569 | -5.740752464 | 4.66E-07 | 0.000860158 | 4.399657341 | cryptochrome-interacting basic-helix-loop-helix 5 | meristem | 6 h |
| HORVU.MOREX.r3.7HG0739910.1 | 0.730435435 | 4.210713917 | 5.220261438 | 3.04E-06 | 0.003468143 | 4.302367512 | Disease resistance protein (NBS-LRR class) family | meristem | 6 h |
| HORVU.MOREX.r3.1HG0026900.1 | -1.267020115 | 1.690413985 | -5.148713981 | 3.92E-06 | 0.004025705 | 4.20242939 | Protein apaG | meristem | 6 h |
| HORVU.MOREX.r3.1HG0011440.1 | -2.707612035 | -1.581106574 | -5.267036101 | 2.57E-06 | 0.003107881 | 4.020869579 | Glucan endo-1,3-beta-glucosidase, putative | meristem | 6 h |
| HORVU.MOREX.r3.5HG0425000.1 | -3.88143041 | -0.918831697 | -5.313227343 | 2.18E-06 | 0.003107881 | 3.896764527 | Transposon protein, putative, Pong sub-class | meristem | 6 h |
| HORVU.MOREX.r3.5HG0489790.1 | 1.870434419 | -0.067483895 | 5.09663292 | 4.72E-06 | 0.004307903 | 3.862876945 | SKP1-like protein 4 | meristem | 6 h |
| HORVU.MOREX.r3.2HG0170640.1 | 3.205760675 | -0.509068665 | 5.158560514 | 3.79E-06 | 0.004025705 | 3.819457129 | Protein FAR1-RELATED SEQUENCE 5 | meristem | 6 h |
| HORVU.MOREX.r3.7HG0640470.1 | 0.434325274 | 6.260306545 | 5.13002745 | 4.19E-06 | 0.004096809 | 3.781484078 | WPP domain-interacting tail-anchored protein 1 | meristem | 6 h |
| HORVU.MOREX.r3.1HG0024020.1 | 0.696729666 | 5.072656773 | 5.090163043 | 4.83E-06 | 0.004307903 | 3.777962551 | Heparan-alpha-glucosaminide N-acetyltransferase | meristem | 6 h |
| HORVU.MOREX.r3.2HG0215830.1 | 0.967978927 | 2.436514446 | 4.9904093 | 6.87E-06 | 0.005870172 | 3.646293647 | CsAtPR5 | meristem | 6 h |
| HORVU.MOREX.r3.3HG0220340.1 | 1.433547088 | 2.953247644 | 4.922847986 | 8.70E-06 | 0.007144143 | 3.424124614 | Disease resistance protein RPM1 | meristem | 6 h |
| HORVU.MOREX.r3.1HG0007570.1 | -3.340096404 | -2.580123327 | -5.29140569 | 2.36E-06 | 0.003107881 | 3.132270009 | Werner Syndrome-like exonuclease | meristem | 6 h |
| HORVU.MOREX.r3.1HG0060370.1 | 2.945280259 | -0.902409718 | 4.862312934 | 1.08E-05 | 0.008489167 | 2.919950252 | 1,4-alpha-glucan branching enzyme GlgB | meristem | 6 h |
| HORVU.MOREX.r3.6HG0629160.1 | 1.240259326 | 2.999097227 | 4.683304963 | 2.00E-05 | 0.01456431 | 2.641039924 | Disease resistance protein RPM1 | meristem | 6 h |
| HORVU.MOREX.r3.2HG0178600.1 | -1.55256008 | 0.128064074 | -4.672365139 | 2.08E-05 | 0.01456431 | 2.624451379 | auxin canalization protein (DUF828) | meristem | 6 h |
| HORVU.MOREX.r3.2HG0165670.1 | -2.846409734 | 0.059715459 | -4.795387704 | 1.36E-05 | 0.010320737 | 2.267923637 | Cytochrome P450 | meristem | 6 h |
| HORVU.MOREX.r3.4HG0343330.1 | 5.692113458 | -0.333480115 | 4.665379318 | 2.13E-05 | 0.01456431 | 2.156336471 | Endonuclease III homolog | meristem | 6 h |
| HORVU.MOREX.r3.5HG0502990.1 | 4.234823953 | 3.237955153 | 4.428004351 | 4.79E-05 | 0.029760706 | 1.902041287 | NAC domain protein, | meristem | 6 h |
| HORVU.MOREX.r3.6HG0614470.2 | -1.516668008 | 0.971508223 | -4.410237313 | 5.08E-05 | 0.03067127 | 1.849982946 | Beta-xylosidase, putative | meristem | 6 h |
| HORVU.MOREX.r3.2HG0120320.1 | -2.228314567 | 1.685244219 | -4.386755463 | 5.50E-05 | 0.032248892 | 1.768206866 | Peroxidase | meristem | 6 h |
| HORVU.MOREX.r3.2HG0181120.1 | -2.430420067 | -0.308323987 | -4.475133889 | 4.08E-05 | 0.02700808 | 1.732342873 | CheY-like two-component responsive regulator family protein | meristem | 6 h |
| HORVU.MOREX.r3.2HG0190780.1 | -2.941339907 | 0.027076117 | -4.457271067 | 4.33E-05 | 0.027797407 | 1.515180368 | Lectin protein kinase family protein | meristem | 6 h |
| HORVU.MOREX.r3.1HG0046230.1 | -1.800046565 | -0.059533313 | -4.325177321 | 6.76E-05 | 0.037512763 | 1.41805808 | Chloroplast inner envelope protein, putative, expressed | meristem | 6 h |
| HORVU.MOREX.r3.7HG0738460.1 | -1.714850255 | 2.451203072 | -4.281371669 | 7.83E-05 | 0.042283677 | 1.335424861 | Cortical cell-delineating protein | meristem | 6 h |
| HORVU.MOREX.r3.4HG0407340.1 | -1.451619444 | 4.417710977 | -4.358560748 | 6.05E-05 | 0.034471281 | 1.231992455 | Receptor-like protein kinase | meristem | 6 h |
| HORVU.MOREX.r3.3HG0297940.1 | 0.942706379 | 3.507795484 | 4.207508857 | 0.0001 | 0.049839617 | 1.060080699 | Transcription initiation factor TFIID subunit 9 | meristem | 6 h |
| HORVU.MOREX.r3.7HG0723600.1 | -1.780246331 | 3.100437987 | -4.201762404 | 0.0001 | 0.049839617 | 0.986151857 | Peroxidase | meristem | 6 h |
| HORVU.MOREX.r3.6HG0615550.1 | -0.520631395 | 6.486073771 | -4.225619412 | 9.43E-05 | 0.049593927 | 0.764136131 | Auxin efflux carrier component | meristem | 6 h |
| HORVU.MOREX.r3.2HG0136150.1 | 3.558062051 | -1.262543021 | 4.21427047 | 9.79E-05 | 0.049839617 | 0.751063599 | Cyclin family protein | meristem | 6 h |
| HORVU.MOREX.r3.1HG0078900.1 | -2.356943162 | 5.8370464 | -26.67159165 | 2.16E-32 | 4.42E-28 | 61.18748575 | MATH domain-containing protein | meristem | 12 h |
| HORVU.MOREX.r3.5HG0447830.1 | -1.323993218 | 4.419279148 | -11.57068945 | 4.00E-16 | 2.74E-12 | 26.28868029 | Sterile alpha motif (SAM) domain-containing protein | meristem | 12 h |
| HORVU.MOREX.r3.1HG0067040.1 | -1.166006839 | 7.101906906 | -10.21502612 | 3.97E-14 | 2.03E-10 | 21.9504842 | NAC domain protein, | meristem | 12 h |
| HORVU.MOREX.r3.1HG0067490.1 | -6.64671345 | -1.72649178 | -12.99379168 | 4.13E-18 | 4.24E-14 | 20.05467657 | Retrovirus-related Pol polyprotein from transposon TNT 1-94 | meristem | 12 h |
| HORVU.MOREX.r3.3HG0280700.1 | 0.604546298 | 6.27769849 | 8.150181955 | 6.44E-11 | 2.20E-07 | 14.68533822 | Katanin p60 ATPase-containing subunit A1 | meristem | 12 h |
| HORVU.MOREX.r3.2HG0111680.1 | -5.098858127 | -2.082427636 | -8.602040897 | 1.24E-11 | 5.07E-08 | 11.21508203 | Formate--tetrahydrofolate ligase | meristem | 12 h |
| HORVU.MOREX.r3.7HG0727570.1 | 4.174950339 | -0.569333422 | 7.805185343 | 2.29E-10 | 6.71E-07 | 10.88033053 | ATP-dependent RNA helicase, putative | meristem | 12 h |
| HORVU.MOREX.r3.7HG0695920.1 | 1.622784768 | 2.660691655 | 6.878807223 | 7.06E-09 | 1.81E-05 | 10.15654296 | #N/A | meristem | 12 h |
| HORVU.MOREX.r3.1HG0067030.1 | -0.862966737 | 5.504670742 | -6.333594859 | 5.31E-08 | 9.90E-05 | 8.17836392 | Protein trichome birefringence | meristem | 12 h |

**Table S5** Continued.

| ID | logFC | AveExpr | t | P.Value | adj.P.Val | B | gene description | root zone | time point |
|---|---|---|---|---|---|---|---|---|---|
| HORVU.MOREX.r3.6HG0546230.1 | 5.459334402 | -1.679648171 | 6.842051948 | 8.09E-09 | 1.84E-05 | 7.30519672 | Disease resistance protein RPM1 | meristem | 12 h |
| HORVU.MOREX.r3.3HG0260130.1 | -1.599399816 | 3.376690694 | -5.928102502 | 2.35E-07 | 0.000371577 | 6.843899026 | Metal transporter, putative | meristem | 12 h |
| HORVU.MOREX.r3.5HG0523880.1 | -4.722789843 | -2.522148569 | -6.676633239 | 1.49E-08 | 3.06E-05 | 6.601525934 | cryptochrome-interacting basic-helix-loop-helix 5 | meristem | 12 h |
| HORVU.MOREX.r3.6HG0559680.1 | -3.215558858 | 0.573580493 | -5.997855654 | 1.82E-07 | 0.000311814 | 6.518765098 | NADPH:quinone oxidoreductase | meristem | 12 h |
| HORVU.MOREX.r3.5HG0471070.1 | -0.401247133 | 6.510820563 | -5.297305196 | 2.31E-06 | 0.002943382 | 4.376261495 | Oxidoreductase family, NAD-binding rossmann fold protein | meristem | 12 h |
| HORVU.MOREX.r3.7HG0739910.1 | 0.71406699 | 4.210713917 | 5.233952919 | 2.90E-06 | 0.003302976 | 4.367441517 | Disease resistance protein (NBS-LRR class) family | meristem | 12 h |
| HORVU.MOREX.r3.5HG0425000.1 | -4.319248879 | -0.918831697 | -5.574245672 | 8.53E-07 | 0.001250572 | 4.252879251 | Transposon protein, putative, Pong sub-class | meristem | 12 h |
| HORVU.MOREX.r3.3HG0297940.1 | 1.129542517 | 3.507795484 | 5.055647179 | 5.46E-06 | 0.005597007 | 3.846979247 | Transcription initiation factor TFIID subunit 9 | meristem | 12 h |
| HORVU.MOREX.r3.1HG0067420.1 | 2.325264852 | 2.43369759 | 5.022866059 | 6.12E-06 | 0.005984115 | 3.7959806 | Cyclic nucleotide-gated channel | meristem | 12 h |
| HORVU.MOREX.r3.1HG0060370.1 | 3.279898541 | -0.902409718 | 5.21807718 | 3.07E-06 | 0.003311276 | 3.762431345 | 1,4-alpha-glucan branching enzyme GlgB | meristem | 12 h |
| HORVU.MOREX.r3.1HG0026900.1 | -1.154634422 | 1.690413985 | -4.945755063 | 8.03E-06 | 0.007166009 | 3.543833674 | Protein apaG | meristem | 12 h |
| HORVU.MOREX.r3.3HG0223800.1 | 2.63548691 | -2.080959444 | 5.419661895 | 1.49E-06 | 0.002038419 | 3.527682514 | E3 ubiquitin-protein ligase | meristem | 12 h |
| HORVU.MOREX.r3.2HG0109620.1 | -2.445135025 | 0.349033917 | -4.967114766 | 7.45E-06 | 0.006950404 | 3.274846815 | Retrotransposon protein, putative, Ty3-gypsy subclass | meristem | 12 h |
| HORVU.MOREX.r3.5HG0523890.1 | -3.65555498 | -2.93597984 | -5.282259871 | 2.44E-06 | 0.002943382 | 2.982928747 | Protein FAR1-RELATED SEQUENCE 5 | meristem | 12 h |
| HORVU.MOREX.r3.5HG0502990.1 | 4.493857223 | 3.237955153 | 4.643211834 | 2.30E-05 | 0.018862338 | 2.580120455 | NAC domain protein, | meristem | 12 h |
| HORVU.MOREX.r3.6HG0559690.1 | 1.138810387 | 2.590055839 | 4.5844711 | 2.81E-05 | 0.021363852 | 2.362103233 | Cell division cycle 5-like protein | meristem | 12 h |
| HORVU.MOREX.r3.1HG0086120.1 | 1.367385604 | 4.055301306 | 4.631457791 | 2.39E-05 | 0.018884426 | 2.355806089 | Histone H2B | meristem | 12 h |
| HORVU.MOREX.r3.3HG0220340.1 | 1.363613788 | 2.953247644 | 4.472439375 | 4.12E-05 | 0.030176366 | 2.000332721 | Disease resistance protein RPM1 | meristem | 12 h |
| HORVU.MOREX.r3.5HG0455140.1 | 3.789572114 | -2.159476557 | 4.677506067 | 2.04E-05 | 0.017460482 | 1.503725036 | tRNA (guanine(37)-N1)-methyltransferase | meristem | 12 h |
| HORVU.MOREX.r3.4HG0343330.1 | 5.520532154 | -0.333480115 | 4.432941053 | 4.71E-05 | 0.03330527 | 1.401491537 | Endonuclease III homolog | meristem | 12 h |
| HORVU.MOREX.r3.1HG0078900.1 | -2.422614082 | 5.8370464 | -28.51895383 | 7.77E-34 | 1.59E-29 | 64.98084387 | MATH domain-containing protein | elongation zone | 0 h |
| HORVU.MOREX.r3.5HG0447830.1 | -1.739209628 | 4.419279148 | -15.07217186 | 8.24E-21 | 8.45E-17 | 36.70962545 | Sterile alpha motif (SAM) domain-containing protein | elongation zone | 0 h |
| HORVU.MOREX.r3.1HG0067040.1 | -1.148278282 | 7.101906906 | -10.76607327 | 5.96E-15 | 3.62E-11 | 23.78021089 | NAC domain protein, | elongation zone | 0 h |
| HORVU.MOREX.r3.3HG0280700.1 | 0.77344826 | 6.27769849 | 10.71690425 | 7.05E-15 | 3.62E-11 | 23.64510567 | Katanin p60 ATPase-containing subunit A1 | elongation zone | 0 h |
| HORVU.MOREX.r3.6HG0546230.1 | 5.587987042 | -1.679648171 | 10.1529121 | 4.92E-14 | 2.02E-10 | 19.114336 | Disease resistance protein RPM1 | elongation zone | 0 h |
| HORVU.MOREX.r3.1HG0070480.1 | -0.999693512 | 6.291080843 | -9.110723368 | 1.96E-12 | 5.76E-09 | 18.09361735 | 60 kDa chaperonin | elongation zone | 0 h |
| HORVU.MOREX.r3.2HG0111680.1 | -5.827259149 | -2.082427636 | -10.01921086 | 7.85E-14 | 2.68E-10 | 15.56820658 | Formate--tetrahydrofolate ligase | elongation zone | 0 h |
| HORVU.MOREX.r3.7HG0739910.1 | 0.970181061 | 4.210713917 | 7.839195746 | 2.02E-10 | 4.60E-07 | 13.59220078 | Disease resistance protein (NBS-LRR class) family | elongation zone | 0 h |
| HORVU.MOREX.r3.7HG0727570.1 | 4.344454469 | -0.569333422 | 8.018617137 | 1.04E-10 | 2.68E-07 | 11.99875711 | ATP-dependent RNA helicase, putative | elongation zone | 0 h |
| HORVU.MOREX.r3.1HG0067420.1 | 2.113663726 | 2.43369759 | 7.255677299 | 1.75E-09 | 3.26E-06 | 11.49215906 | Cyclic nucleotide-gated channel | elongation zone | 0 h |
| HORVU.MOREX.r3.1HG0024020.1 | 0.881576746 | 5.072656773 | 6.611655844 | 1.90E-08 | 3.00E-05 | 9.126149482 | Heparan-alpha-glucosaminide N-acetyltransferase | elongation zone | 0 h |
| HORVU.MOREX.r3.1HG0067490.1 | -4.109176405 | -1.72649178 | -7.351158696 | 1.23E-09 | 2.52E-06 | 8.734196801 | Retrovirus-related Pol polyprotein from transposon TNT 1-94 | elongation zone | 0 h |
| HORVU.MOREX.r3.7HG0729150.1 | -4.08685209 | -2.071842766 | -6.486030244 | 3.02E-08 | 4.43E-05 | 8.273992271 | GDSL esterase/lipase | elongation zone | 0 h |
| HORVU.MOREX.r3.1HG0067030.1 | -0.70170823 | 5.504670742 | -6.244083069 | 7.38E-08 | 0.000100942 | 7.671149471 | Protein trichome birefringence | elongation zone | 0 h |
| HORVU.MOREX.r3.1HG0007570.1 | -4.176564388 | -2.580123327 | -6.871743302 | 7.25E-09 | 1.24E-05 | 7.532448969 | Werner Syndrome-like exonuclease | elongation zone | 0 h |
| HORVU.MOREX.r3.3HG0298020.1 | 1.559034982 | 3.677613517 | 6.089064833 | 1.30E-07 | 0.000167358 | 7.414162165 | Transcription initiation factor TFIID subunit 9 | elongation zone | 0 h |
| HORVU.MOREX.r3.7HG0695920.1 | 1.655379288 | 2.660691655 | 5.881841045 | 2.79E-07 | 0.000336506 | 6.682821932 | #N/A | elongation zone | 0 h |
| HORVU.MOREX.r3.3HG0297940.1 | 1.681302659 | 3.507795484 | 5.84300731 | 3.21E-07 | 0.000366232 | 6.560477311 | Transcription initiation factor TFIID subunit 9 | elongation zone | 0 h |
| HORVU.MOREX.r3.6HG0559680.1 | -2.825403355 | 0.573580493 | -5.537876655 | 9.73E-07 | 0.000998411 | 5.316461587 | NADPH:quinone oxidoreductase | elongation zone | 0 h |
| HORVU.MOREX.r3.1HG0011440.1 | -2.432512003 | -1.581106574 | -5.503821648 | 1.10E-06 | 0.001075269 | 5.078537785 | Glucan endo-1,3-beta-glucosidase, putative | elongation zone | 0 h |
| HORVU.MOREX.r3.1HG0069040.1 | 2.736631601 | 0.814092704 | 5.552625688 | 9.23E-07 | 0.00099641 | 4.913083956 | #N/A | elongation zone | 0 h |

**Table S5** Continued.

| ID | logFC | AveExpr | t | P.Value | adj.P.Val | B | gene description | root zone | time point |
|---|---|---|---|---|---|---|---|---|---|
| HORVU.MOREX.r3.1HG0026900.1 | -1.537272869 | 1.690413985 | -5.372606052 | 1.76E-06 | 0.001645764 | 4.894176493 | Protein apaG | elongation zone | 0 h |
| HORVU.MOREX.r3.7HG0713840.1 | -1.729173965 | -2.185081282 | -5.323087466 | 2.11E-06 | 0.001879945 | 4.67913266 | Cytochrome P450 | elongation zone | 0 h |
| HORVU.MOREX.r3.5HG0471070.1 | -0.38511 | 6.510820563 | -5.270083685 | 2.55E-06 | 0.002177587 | 4.16886943 | Oxidoreductase family, NAD-binding rossmann fold protein | elongation zone | 0 h |
| HORVU.MOREX.r3.1HG0008190.1 | -2.015856807 | -0.63381752 | -5.145343793 | 3.97E-06 | 0.00325932 | 4.153817343 | Cytochrome P450 | elongation zone | 0 h |
| HORVU.MOREX.r3.1HG0070000.1 | 1.18351875 | 1.808570831 | 4.968347796 | 7.42E-06 | 0.005728582 | 3.617173342 | Histone H3 | elongation zone | 0 h |
| HORVU.MOREX.r3.5HG0456970.1 | 0.756861983 | 5.8820687 | 4.963852047 | 7.54E-06 | 0.005728582 | 3.297343114 | Methylthioribose-1-phosphate isomerase | elongation zone | 0 h |
| HORVU.MOREX.r3.5HG0502990.1 | 3.613022113 | 3.237955153 | 4.833050057 | 1.19E-05 | 0.008428702 | 3.110484948 | NAC domain protein, | elongation zone | 0 h |
| HORVU.MOREX.r3.5HG0425000.1 | -3.369231289 | -0.918831697 | -4.879950122 | 1.01E-05 | 0.007411861 | 2.902544643 | Transposon protein, putative, Pong sub-class | elongation zone | 0 h |
| HORVU.MOREX.r3.7HG0642740.1 | 0.566304994 | 4.413737834 | 4.684904102 | 1.99E-05 | 0.013616628 | 2.478305116 | SET domain-containing protein | elongation zone | 0 h |
| HORVU.MOREX.r3.1HG0049550.1 | 0.507585192 | 3.543555344 | 4.647304544 | 2.27E-05 | 0.014999 | 2.433088554 | Protein BPS1, chloroplastic | elongation zone | 0 h |
| HORVU.MOREX.r3.1HG0070730.1 | -1.966552983 | -0.4130893 | -4.569639661 | 2.96E-05 | 0.018963815 | 2.338886368 | Glycosyltransferase | elongation zone | 0 h |
| HORVU.MOREX.r3.6HG0559690.1 | 1.26091266 | 2.590055839 | 4.547933073 | 3.18E-05 | 0.019804371 | 2.2461328 | Cell division cycle 5-like protein | elongation zone | 0 h |
| HORVU.MOREX.r3.2HG0170640.1 | 2.884435418 | -0.509068665 | 4.513814427 | 3.58E-05 | 0.021592098 | 1.979093494 | Protein FAR1-RELATED SEQUENCE 5 | elongation zone | 0 h |
| HORVU.MOREX.r3.2HG0215830.1 | 1.269991868 | 2.436514446 | 4.389339633 | 5.45E-05 | 0.028690561 | 1.792378738 | CsAtPR5 | elongation zone | 0 h |
| HORVU.MOREX.r3.1HG0070100.1 | -0.529189266 | 4.109158355 | -4.442367971 | 4.56E-05 | 0.025670408 | 1.660912197 | Prolyl 4-hydroxylase alpha subunit, putative | elongation zone | 0 h |
| HORVU.MOREX.r3.1HG0086120.1 | 1.87007834 | 4.055301306 | 4.352790109 | 6.17E-05 | 0.031631148 | 1.604095686 | Histone H2B | elongation zone | 0 h |
| HORVU.MOREX.r3.7HG0672900.1 | 2.368462875 | 0.425915719 | 4.410644072 | 5.08E-05 | 0.027405079 | 1.569048605 | Pentatricopeptide repeat-containing protein | elongation zone | 0 h |
| HORVU.MOREX.r3.5HG0462380.1 | -0.804499194 | 3.998698155 | -4.43789627 | 4.63E-05 | 0.025670408 | 1.558859351 | LYR family of Fe/S cluster biogenesis protein | elongation zone | 0 h |
| HORVU.MOREX.r3.3HG0222160.1 | 1.645322602 | -1.846318567 | 4.264610651 | 8.28E-05 | 0.040454234 | 1.415475584 | Nuclease S1 | elongation zone | 0 h |
| HORVU.MOREX.r3.UnG0765280.1 | 1.461373095 | 2.589811572 | 4.211707084 | 9.87E-05 | 0.045010155 | 1.249952442 | F-box family protein | elongation zone | 0 h |
| HORVU.MOREX.r3.5HG0485410.1 | 2.807732376 | -0.444670009 | 4.448551505 | 4.46E-05 | 0.025670408 | 1.23452293 | #N/A | elongation zone | 0 h |
| HORVU.MOREX.r3.1HG0069380.1 | -1.252691628 | 2.129642878 | -4.222105907 | 9.54E-05 | 0.044473516 | 1.102014108 | Cysteine proteinase inhibitor | elongation zone | 0 h |
| HORVU.MOREX.r3.1HG0068410.1 | -0.396632513 | 4.707522572 | -4.227371005 | 9.37E-05 | 0.044473516 | 0.901219617 | E3 ubiquitin-protein ligase | elongation zone | 0 h |
| HORVU.MOREX.r3.7HG0640470.1 | 0.360908747 | 6.260306545 | 4.268354115 | 8.18E-05 | 0.040454234 | 0.863727652 | WPP domain-interacting tail-anchored protein 1 | elongation zone | 0 h |
| HORVU.MOREX.r3.1HG0078900.1 | -2.367787064 | 5.8370464 | -27.16858464 | 8.65E-33 | 1.78E-28 | 61.4975963 | MATH domain-containing protein | elongation zone | 3 h |
| HORVU.MOREX.r3.3HG0280700.1 | 0.955350063 | 6.27769849 | 13.06592538 | 3.30E-18 | 3.38E-14 | 31.11992727 | Katanin p60 ATPase-containing subunit A1 | elongation zone | 3 h |
| HORVU.MOREX.r3.5HG0447830.1 | -1.385731994 | 4.419279148 | -12.00606175 | 9.61E-17 | 6.57E-13 | 27.52191979 | Sterile alpha motif (SAM) domain-containing protein | elongation zone | 3 h |
| HORVU.MOREX.r3.1HG0070480.1 | -1.055905972 | 6.291080843 | -9.525129096 | 4.47E-13 | 1.84E-09 | 19.58432076 | 60 kDa chaperonin | elongation zone | 3 h |
| HORVU.MOREX.r3.6HG0546230.1 | 8.223902991 | -1.679648171 | 10.89599429 | 3.83E-15 | 1.97E-11 | 16.04555447 | Disease resistance protein RPM1 | elongation zone | 3 h |
| HORVU.MOREX.r3.7HG0739910.1 | 1.058388899 | 4.210713917 | 8.463391419 | 2.05E-11 | 6.00E-08 | 15.84797849 | Disease resistance protein (NBS-LRR class) family | elongation zone | 3 h |
| HORVU.MOREX.r3.1HG0067040.1 | -0.860334376 | 7.101906906 | -8.072437153 | 8.56E-11 | 2.20E-07 | 14.37828129 | NAC domain protein, | elongation zone | 3 h |
| HORVU.MOREX.r3.1HG0024020.1 | 1.007859267 | 5.072656773 | 7.548999833 | 5.90E-10 | 1.10E-06 | 12.58535786 | Heparan-alpha-glucosaminide N-acetyltransferase | elongation zone | 3 h |
| HORVU.MOREX.r3.7HG0727570.1 | 4.507550537 | -0.569333422 | 8.57418003 | 1.37E-11 | 4.67E-08 | 12.4573334 | ATP-dependent RNA helicase, putative | elongation zone | 3 h |
| HORVU.MOREX.r3.7HG0729150.1 | -4.941180132 | -2.071842766 | -7.64021831 | 4.21E-10 | 8.64E-07 | 11.13257623 | GDSL esterase/lipase | elongation zone | 3 h |
| HORVU.MOREX.r3.2HG0111680.1 | -4.589353007 | -2.082427636 | -7.886653376 | 1.70E-10 | 3.87E-07 | 9.094847114 | Formate--tetrahydrofolate ligase | elongation zone | 3 h |
| HORVU.MOREX.r3.7HG0640470.1 | 0.528792269 | 6.260306545 | 6.249446952 | 7.23E-08 | 0.000114194 | 7.835006347 | WPP domain-interacting tail-anchored protein 1 | elongation zone | 3 h |
| HORVU.MOREX.r3.1HG0070000.1 | 1.471763849 | 1.808570831 | 6.201305183 | 8.64E-08 | 0.000126601 | 7.523035759 | Histone H3 | elongation zone | 3 h |
| HORVU.MOREX.r3.1HG0067030.1 | -0.681642754 | 5.504670742 | -6.029899435 | 1.62E-07 | 0.000221804 | 7.029741265 | Protein trichome birefringence | elongation zone | 3 h |
| HORVU.MOREX.r3.1HG0067420.1 | 1.693409973 | 2.43369759 | 5.992921729 | 1.86E-07 | 0.000229369 | 7.013538098 | Cyclic nucleotide-gated channel | elongation zone | 3 h |
| HORVU.MOREX.r3.6HG0559690.1 | 1.701878582 | 2.590055839 | 5.986602932 | 1.90E-07 | 0.000229369 | 6.968088164 | Cell division cycle 5-like protein | elongation zone | 3 h |
| HORVU.MOREX.r3.1HG0067490.1 | -3.871019017 | -1.72649178 | -6.885558721 | 6.89E-09 | 1.18E-05 | 6.35207369 | Retrovirus-related Pol polyprotein from transposon TNT 1-94 | elongation zone | 3 h |

**Table S5** Continued.

| ID | logFC | AveExpr | t | P.Value | adj.P.Val | B | gene description | root zone | time point |
|---|---|---|---|---|---|---|---|---|---|
| HORVU.MOREX.r3.1HG0069380.1 | -1.723970494 | 2.129642878 | -5.596127384 | 7.88E-07 | 0.000851314 | 5.714386819 | Cysteine proteinase inhibitor | elongation zone | 3 h |
| HORVU.MOREX.r3.6HG0559680.1 | -3.241273008 | 0.573580493 | -5.695060318 | 5.51E-07 | 0.000627676 | 5.288905706 | NADPH:quinone oxidoreductase | elongation zone | 3 h |
| HORVU.MOREX.r3.1HG0049550.1 | 0.553787039 | 3.543555344 | 5.202462527 | 3.24E-06 | 0.002994663 | 4.358235879 | Protein BPS1, chloroplastic | elongation zone | 3 h |
| HORVU.MOREX.r3.5HG0502990.1 | 3.912615301 | 3.237955153 | 5.192641224 | 3.36E-06 | 0.002994663 | 4.356523379 | NAC domain protein, | elongation zone | 3 h |
| HORVU.MOREX.r3.2HG0216110.1 | 0.462249853 | 5.350102781 | 5.139171845 | 4.06E-06 | 0.003470302 | 4.004733356 | #N/A | elongation zone | 3 h |
| HORVU.MOREX.r3.1HG0011440.1 | -2.859919008 | -1.581106574 | -5.341069817 | 1.98E-06 | 0.001930552 | 3.787414724 | Glucan endo-1,3-beta-glucosidase, putative | elongation zone | 3 h |
| HORVU.MOREX.r3.5HG0471070.1 | -0.36625879 | 6.510820563 | -4.929120818 | 8.51E-06 | 0.006471105 | 3.155351816 | Oxidoreductase family, NAD-binding rossmann fold protein | elongation zone | 3 h |
| HORVU.MOREX.r3.1HG0007570.1 | -3.437776806 | -2.580123327 | -5.489837761 | 1.16E-06 | 0.001187441 | 3.075820548 | Werner Syndrome-like exonuclease | elongation zone | 3 h |
| HORVU.MOREX.r3.1HG0008190.1 | -1.885579918 | -0.63381752 | -4.751862413 | 1.58E-05 | 0.011175626 | 2.805536576 | Cytochrome P450 | elongation zone | 3 h |
| HORVU.MOREX.r3.3HG0297940.1 | 1.409703042 | 3.507795484 | 4.699491836 | 1.89E-05 | 0.012530555 | 2.75751099 | Transcription initiation factor TFIID subunit 9 | elongation zone | 3 h |
| HORVU.MOREX.r3.4HG0344830.1 | 0.618537725 | 4.535567699 | 4.732773499 | 1.69E-05 | 0.011541293 | 2.685874488 | NBS-LRR disease resistance protein, putative, expressed | elongation zone | 3 h |
| HORVU.MOREX.r3.2HG0170640.1 | 3.7419642 | -0.509068665 | 4.987331845 | 6.94E-06 | 0.005477606 | 2.614080447 | Protein FAR1-RELATED SEQUENCE 5 | elongation zone | 3 h |
| HORVU.MOREX.r3.1HG0069030.1 | -2.774203235 | 1.480380355 | -4.650823875 | 2.24E-05 | 0.014355463 | 2.553387438 | Calcium-transporting ATPase | elongation zone | 3 h |
| HORVU.MOREX.r3.2HG0131630.1 | -3.211907792 | -3.028166822 | -4.765991826 | 1.50E-05 | 0.011021467 | 2.482398226 | vacuolar sorting-associated protein (DUF946) | elongation zone | 3 h |
| HORVU.MOREX.r3.6HG0581570.1 | 0.473463599 | 4.929735327 | 4.619585589 | 2.49E-05 | 0.015497632 | 2.320262592 | Leucine-rich repeat-containing protein 1 | elongation zone | 3 h |
| HORVU.MOREX.r3.5HG0516660.1 | 0.484063951 | 4.126867737 | 4.578655188 | 2.87E-05 | 0.017306414 | 2.213175948 | LEM3 (Ligand-effect modulator 3) family protein | elongation zone | 3 h |
| HORVU.MOREX.r3.3HG0283300.1 | 0.864415145 | 2.473566507 | 4.516716492 | 3.54E-05 | 0.02076901 | 2.177606725 | Farnesyl diphosphate synthase | elongation zone | 3 h |
| HORVU.MOREX.r3.4HG0408780.1 | 3.311828691 | 0.649042183 | 5.099689871 | 4.67E-06 | 0.003831858 | 2.138941056 | Pentatricopeptide repeat-containing protein | elongation zone | 3 h |
| HORVU.MOREX.r3.1HG0026900.1 | -1.314590096 | 1.690413985 | -4.426931743 | 4.80E-05 | 0.025938662 | 1.834003757 | Protein apaG | elongation zone | 3 h |
| HORVU.MOREX.r3.2HG0101220.2 | 0.715197686 | 2.581198545 | 4.352859232 | 6.16E-05 | 0.030465058 | 1.641381682 | Formin-like protein | elongation zone | 3 h |
| HORVU.MOREX.r3.3HG0298020.1 | 1.093419967 | 3.677613517 | 4.31722943 | 6.95E-05 | 0.0323953 | 1.543507182 | Transcription initiation factor TFIID subunit 9 | elongation zone | 3 h |
| HORVU.MOREX.r3.1HG0068410.1 | -0.417149359 | 4.707522572 | -4.376301115 | 5.70E-05 | 0.029977701 | 1.525312309 | E3 ubiquitin-protein ligase | elongation zone | 3 h |
| HORVU.MOREX.r3.5HG0490030.1 | 0.303902563 | 7.481265074 | 4.481675984 | 3.99E-05 | 0.022746333 | 1.521349586 | Vacuolar protein sorting-associated protein 52 | elongation zone | 3 h |
| HORVU.MOREX.r3.4HG0413690.1 | 0.622272005 | 4.40336351 | 4.337224121 | 6.50E-05 | 0.03100158 | 1.470809108 | 26S proteasome non-ATPase regulatory subunit-like protein | elongation zone | 3 h |
| HORVU.MOREX.r3.7HG0695920.1 | 1.148034454 | 2.660691655 | 4.277746903 | 7.93E-05 | 0.036140397 | 1.449527232 | #N/A | elongation zone | 3 h |
| HORVU.MOREX.r3.1HG0044150.1 | 0.429249479 | 5.220215331 | 4.349444765 | 6.24E-05 | 0.030465058 | 1.356020546 | Ran-binding protein in the microtubule-organising centre protein | elongation zone | 3 h |
| HORVU.MOREX.r3.7HG0635110.1 | 1.280557244 | 1.994165679 | 4.222670585 | 9.52E-05 | 0.042056332 | 1.289237533 | NBS-LRR-like resistance protein | elongation zone | 3 h |
| HORVU.MOREX.r3.4HG0374910.2 | 0.345737887 | 6.337717806 | 4.368329306 | 5.85E-05 | 0.030022376 | 1.278032186 | Beta-adaptin-like protein | elongation zone | 3 h |
| HORVU.MOREX.r3.5HG0515610.1 | -2.401569095 | -3.366672775 | -4.44048086 | 4.59E-05 | 0.025446978 | 1.107059229 | Collagen alpha chain | elongation zone | 3 h |
| HORVU.MOREX.r3.5HG0443920.1 | 0.579301829 | 3.840341821 | 4.161039685 | 0.00012 | 0.048870078 | 0.940454887 | Calmodulin-binding protein, putative, expressed | elongation zone | 3 h |
| HORVU.MOREX.r3.7HG0685870.1 | 0.282506125 | 7.40990073 | 4.219067623 | 9.63E-05 | 0.042056332 | 0.714242914 | Ankyrin repeat domain containing protein | elongation zone | 3 h |
| HORVU.MOREX.r3.3HG0309270.1 | 0.357546443 | 8.53260482 | 4.179899551 | 0.00011 | 0.04687848 | 0.561145832 | Proline--tRNA ligase | elongation zone | 3 h |
| HORVU.MOREX.r3.1HG0078900.1 | -2.455370206 | 5.8370464 | -28.11453467 | 1.58E-33 | 3.25E-29 | 63.65412911 | MATH domain-containing protein | elongation zone | 6 h |
| HORVU.MOREX.r3.5HG0447830.1 | -1.690771799 | 4.419279148 | -14.01906786 | 1.80E-19 | 1.84E-15 | 33.53649068 | Sterile alpha motif (SAM) domain-containing protein | elongation zone | 6 h |
| HORVU.MOREX.r3.3HG0280700.1 | 0.95254549 | 6.27769849 | 12.9611432 | 4.57E-18 | 3.13E-14 | 30.85679699 | Katanin p60 ATPase-containing subunit A1 | elongation zone | 6 h |
| HORVU.MOREX.r3.1HG0070480.1 | -1.174994142 | 6.291080843 | -10.68035051 | 7.98E-15 | 3.28E-11 | 23.53342025 | 60 kDa chaperonin | elongation zone | 6 h |
| HORVU.MOREX.r3.6HG0546230.1 | 6.651659642 | -1.679648171 | 10.71297343 | 7.14E-15 | 3.28E-11 | 18.94302896 | Disease resistance protein RPM1 | elongation zone | 6 h |
| HORVU.MOREX.r3.1HG0067040.1 | -0.981175584 | 7.101906906 | -9.180873495 | 1.53E-12 | 5.22E-09 | 18.31447218 | NAC domain protein, | elongation zone | 6 h |
| HORVU.MOREX.r3.7HG0739910.1 | 1.19638186 | 4.210713917 | 9.0268823 | 2.66E-12 | 7.78E-09 | 17.83509786 | Disease resistance protein (NBS-LRR class) family | elongation zone | 6 h |
| HORVU.MOREX.r3.1HG0024020.1 | 1.079933311 | 5.072656773 | 7.891375987 | 1.67E-10 | 3.42E-07 | 13.80313715 | Heparan-alpha-glucosaminide N-acetyltransferase | elongation zone | 6 h |
| HORVU.MOREX.r3.7HG0729150.1 | -5.192955515 | -2.071842766 | -7.905081967 | 1.58E-10 | 3.42E-07 | 12.28495143 | GDSL esterase/lipase | elongation zone | 6 h |

**Table S5** Continued.

| ID | logFC | AveExpr | t | P.Value | adj.P.Val | B | gene description | root zone | time point |
|---|---|---|---|---|---|---|---|---|---|
| HORVU.MOREX.r3.7HG0727570.1 | 4.338282709 | -0.569333422 | 7.775303779 | 2.56E-10 | 4.77E-07 | 10.67323007 | ATP-dependent RNA helicase, putative | elongation zone | 6 h |
| HORVU.MOREX.r3.3HG0297940.1 | 2.412350516 | 3.507795484 | 7.017407454 | 4.22E-09 | 6.67E-06 | 10.41416616 | Transcription initiation factor TFIID subunit 9 | elongation zone | 6 h |
| HORVU.MOREX.r3.2HG0111680.1 | -4.61053162 | -2.082427636 | -7.946881235 | 1.36E-10 | 3.42E-07 | 10.04730863 | Formate--tetrahydrofolate ligase | elongation zone | 6 h |
| HORVU.MOREX.r3.7HG0640470.1 | 0.58174285 | 6.260306545 | 6.794966612 | 9.63E-09 | 1.41E-05 | 9.759209171 | WPP domain-interacting tail-anchored protein 1 | elongation zone | 6 h |
| HORVU.MOREX.r3.3HG0307310.1 | 0.614982179 | 5.321616586 | 6.774594255 | 1.04E-08 | 1.42E-05 | 9.723133442 | magnesium transporter, putative (DUF803) | elongation zone | 6 h |
| HORVU.MOREX.r3.1HG0069380.1 | -1.763094232 | 2.129642878 | -6.491279587 | 2.96E-08 | 3.58E-05 | 8.820532232 | Cysteine proteinase inhibitor | elongation zone | 6 h |
| HORVU.MOREX.r3.1HG0067030.1 | -0.726019821 | 5.504670742 | -6.539553565 | 2.48E-08 | 3.18E-05 | 8.794661825 | Protein trichome birefringence | elongation zone | 6 h |
| HORVU.MOREX.r3.2HG0121410.1 | -2.139380697 | 3.844655715 | -6.341546329 | 5.15E-08 | 5.87E-05 | 8.21376623 | Cytochrome P450 | elongation zone | 6 h |
| HORVU.MOREX.r3.1HG0067490.1 | -4.025300541 | -1.72649178 | -7.211767514 | 2.06E-09 | 3.51E-06 | 7.834831591 | Retrovirus-related Pol polyprotein from transposon TNT 1-94 | elongation zone | 6 h |
| HORVU.MOREX.r3.1HG0067420.1 | 1.639056173 | 2.43369759 | 6.197460773 | 8.76E-08 | 8.99E-05 | 7.661526291 | Cyclic nucleotide-gated channel | elongation zone | 6 h |
| HORVU.MOREX.r3.7HG0713840.1 | -1.835945375 | -2.185081282 | -6.139491032 | 1.08E-07 | 0.000105947 | 7.315556406 | Cytochrome P450 | elongation zone | 6 h |
| HORVU.MOREX.r3.3HG0298020.1 | 1.532913962 | 3.677613517 | 5.884167085 | 2.76E-07 | 0.000246617 | 6.700175248 | Transcription initiation factor TFIID subunit 9 | elongation zone | 6 h |
| HORVU.MOREX.r3.7HG0717610.1 | -1.021543256 | 4.008622047 | -5.780761256 | 4.03E-07 | 0.000330872 | 6.22234164 | Pollen-specific protein SF21 | elongation zone | 6 h |
| HORVU.MOREX.r3.3HG0246300.1 | -0.761004763 | 4.385187463 | -5.749778683 | 4.51E-07 | 0.000356141 | 6.14000964 | RNA-binding protein, putative | elongation zone | 6 h |
| HORVU.MOREX.r3.7HG0721320.1 | -2.888168968 | -3.129421625 | -6.054431513 | 1.48E-07 | 0.000138215 | 6.131854686 | Auxin-responsive family protein | elongation zone | 6 h |
| HORVU.MOREX.r3.5HG0467870.1 | -1.53632014 | 2.049337326 | -5.669167377 | 6.05E-07 | 0.000459707 | 5.948759622 | Kynurenine formamidase | elongation zone | 6 h |
| HORVU.MOREX.r3.7HG0750000.1 | -1.961665845 | -2.120414367 | -5.638396638 | 6.76E-07 | 0.000495644 | 5.84507816 | Disease resistance protein | elongation zone | 6 h |
| HORVU.MOREX.r3.3HG0265690.1 | 2.604079702 | 0.540508683 | 5.854804527 | 3.08E-07 | 0.000263097 | 5.434526612 | Oxidoreductase/transition metal ion-binding protein | elongation zone | 6 h |
| HORVU.MOREX.r3.1HG0045510.1 | 0.88273122 | 5.275918097 | 5.552347984 | 9.24E-07 | 0.000653476 | 5.43350822 | Prefoldin subunit, putative | elongation zone | 6 h |
| HORVU.MOREX.r3.1HG0007570.1 | -3.810028916 | -2.580123327 | -6.205618112 | 8.50E-08 | 8.99E-05 | 5.345121288 | Werner Syndrome-like exonuclease | elongation zone | 6 h |
| HORVU.MOREX.r3.1HG0070000.1 | 1.31894142 | 1.808570831 | 5.450424803 | 1.33E-06 | 0.00091242 | 5.145583959 | Histone H3 | elongation zone | 6 h |
| HORVU.MOREX.r3.6HG0576010.1 | -0.862268712 | 4.663155786 | -5.384744763 | 1.69E-06 | 0.001083225 | 4.915571494 | Protein kinase family protein | elongation zone | 6 h |
| HORVU.MOREX.r3.6HG0559690.1 | 1.539799486 | 2.590055839 | 5.324074131 | 2.10E-06 | 0.001240485 | 4.789298387 | Cell division cycle 5-like protein | elongation zone | 6 h |
| HORVU.MOREX.r3.7HG0751110.1 | 0.903579811 | 4.358148537 | 5.321938433 | 2.12E-06 | 0.001240485 | 4.686130966 | Kinesin-like | elongation zone | 6 h |
| HORVU.MOREX.r3.1HG0069030.1 | -3.029787188 | 1.480380355 | -5.297280366 | 2.31E-06 | 0.001281665 | 4.6572396 | Calcium-transporting ATPase | elongation zone | 6 h |
| HORVU.MOREX.r3.1HG0066800.1 | -1.713700121 | -2.502872481 | -5.304873224 | 2.25E-06 | 0.001281665 | 4.648832848 | Potassium channel | elongation zone | 6 h |
| HORVU.MOREX.r3.3HG0257680.1 | 0.606888573 | 4.018531146 | 5.287662346 | 2.39E-06 | 0.001291596 | 4.58263926 | GDSL esterase/lipase | elongation zone | 6 h |
| HORVU.MOREX.r3.2HG0191490.1 | -1.218138174 | 4.031252139 | -5.271671375 | 2.53E-06 | 0.001324316 | 4.578022461 | Protein kinase family protein | elongation zone | 6 h |
| HORVU.MOREX.r3.3HG0250060.1 | -2.507206312 | -2.506423371 | -5.395287951 | 1.63E-06 | 0.001076624 | 4.536257979 | CASP-like protein | elongation zone | 6 h |
| HORVU.MOREX.r3.3HG0234470.1 | 0.554525675 | 6.523252025 | 5.337159015 | 2.00E-06 | 0.001240485 | 4.492703754 | Kinase interacting (KIP1-like) family protein | elongation zone | 6 h |
| HORVU.MOREX.r3.5HG0456970.1 | 0.821729809 | 5.8820687 | 5.266302155 | 2.58E-06 | 0.001324316 | 4.410059397 | Methylthioribose-1-phosphate isomerase | elongation zone | 6 h |
| HORVU.MOREX.r3.5HG0502990.1 | 4.9082101 | 3.237955153 | 5.187439488 | 3.42E-06 | 0.001670563 | 4.333541478 | NAC domain protein, | elongation zone | 6 h |
| HORVU.MOREX.r3.7HG0695920.1 | 1.447766581 | 2.660691655 | 5.173306978 | 3.60E-06 | 0.001710089 | 4.290523171 | #N/A | elongation zone | 6 h |
| HORVU.MOREX.r3.2HG0121570.1 | 0.750639648 | 4.65733005 | 5.204131254 | 3.22E-06 | 0.001612625 | 4.246480902 | Protein LTV1 | elongation zone | 6 h |
| HORVU.MOREX.r3.1HG0003620.1 | -1.057878604 | 2.573512603 | -5.123715687 | 4.29E-06 | 0.001955112 | 4.073214787 | Protein ZINC INDUCED FACILITATOR-LIKE 1 | elongation zone | 6 h |
| HORVU.MOREX.r3.2HG0096230.1 | -1.969780166 | -2.935502699 | -5.16777579 | 3.67E-06 | 0.001710089 | 3.94862768 | Phosphatidylinositol 3- and 4-kinase family protein | elongation zone | 6 h |
| HORVU.MOREX.r3.7HG0749460.1 | 0.74573892 | 5.222321052 | 5.089005984 | 4.85E-06 | 0.002125793 | 3.773305094 | Thioredoxin | elongation zone | 6 h |
| HORVU.MOREX.r3.6HG0549990.1 | 1.245683575 | 2.338995774 | 4.96765541 | 7.44E-06 | 0.002962634 | 3.611201186 | Pentatricopeptide repeat-containing protein | elongation zone | 6 h |
| HORVU.MOREX.r3.5HG0472700.1 | -0.583699361 | 4.865209235 | -5.004481291 | 6.53E-06 | 0.002793246 | 3.532137528 | Lipid phosphate phosphatase-like protein | elongation zone | 6 h |
| HORVU.MOREX.r3.1HG0082770.1 | 0.418014229 | 7.702552295 | 5.087803984 | 4.87E-06 | 0.002125793 | 3.527496667 | Reticulon-like protein | elongation zone | 6 h |
| HORVU.MOREX.r3.5HG0513990.1 | -1.907823345 | -1.75785159 | -4.99251821 | 6.81E-06 | 0.00285393 | 3.516572003 | Nitrate transporter 1.1 | elongation zone | 6 h |

**Table S5** Continued.

| ID | logFC | AveExpr | t | P.Value | adj.P.Val | B | gene description | root zone | time point |
|---|---|---|---|---|---|---|---|---|---|
| HORVU.MOREX.r3.3HG0280960.1 | -1.085878314 | 1.80535629 | -4.959397535 | 7.66E-06 | 0.002962634 | 3.483252383 | Glycosyltransferase | elongation zone | 6 h |
| HORVU.MOREX.r3.7HG0636630.1 | -1.615620741 | -0.128553353 | -4.934860399 | 8.34E-06 | 0.002991092 | 3.435965215 | WAT1-related protein | elongation zone | 6 h |
| HORVU.MOREX.r3.6HG0546140.1 | 1.16624523 | 0.639454141 | 4.892027425 | 9.70E-06 | 0.003309051 | 3.373914025 | F-box protein | elongation zone | 6 h |
| HORVU.MOREX.r3.5HG0518560.1 | -2.120765461 | 2.148281347 | -4.95423955 | 7.80E-06 | 0.002962634 | 3.356430992 | Leucine-rich repeat protein kinase family protein | elongation zone | 6 h |
| HORVU.MOREX.r3.7HG0706100.1 | 0.454452673 | 5.89851201 | 4.964950579 | 7.51E-06 | 0.002962634 | 3.317552742 | Survival of motor neuron-related-splicing factor 30 | elongation zone | 6 h |
| HORVU.MOREX.r3.7HG0649950.1 | -0.814788896 | 3.836694774 | -4.926479766 | 8.59E-06 | 0.002991092 | 3.297380017 | Nudix hydrolase | elongation zone | 6 h |
| HORVU.MOREX.r3.6HG0629160.1 | 1.937870345 | 2.999097227 | 4.874145492 | 1.03E-05 | 0.00335152 | 3.279240853 | Disease resistance protein RPM1 | elongation zone | 6 h |
| HORVU.MOREX.r3.6HG0626030.1 | -1.96809199 | -0.562096779 | -4.926270129 | 8.60E-06 | 0.002991092 | 3.26239866 | Peroxidase | elongation zone | 6 h |
| HORVU.MOREX.r3.3HG0278540.1 | -0.579158692 | 6.253662754 | -4.977925143 | 7.17E-06 | 0.00294416 | 3.261890915 | Hydroxyacylglutathione hydrolase | elongation zone | 6 h |
| HORVU.MOREX.r3.4HG0417920.1 | -2.121932948 | -1.699022369 | -4.86825044 | 1.05E-05 | 0.00335152 | 3.249244029 | Peroxidase | elongation zone | 6 h |
| HORVU.MOREX.r3.1HG0068380.1 | 0.602322 | 5.314485157 | 4.926879574 | 8.58E-06 | 0.002991092 | 3.247150527 | #N/A | elongation zone | 6 h |
| HORVU.MOREX.r3.1HG0070730.1 | -2.143795304 | -0.4130893 | -4.867084457 | 1.06E-05 | 0.00335152 | 3.209379901 | Glycosyltransferase | elongation zone | 6 h |
| HORVU.MOREX.r3.2HG0217090.1 | -1.775775231 | -0.857487868 | -4.887889031 | 9.84E-06 | 0.003309051 | 3.209130287 | C2 calcium/lipid-binding and GRAM domain protein | elongation zone | 6 h |
| HORVU.MOREX.r3.1HG0010340.1 | -1.651723824 | -2.094132866 | -4.825228968 | 1.22E-05 | 0.003693864 | 3.150066422 | Non-specific lipid-transfer protein-like protein | elongation zone | 6 h |
| HORVU.MOREX.r3.3HG0302100.1 | 0.386270197 | 5.746504247 | 4.928421174 | 8.54E-06 | 0.002991092 | 3.142443404 | Mannose-1-phosphate guanyltransferase 1 | elongation zone | 6 h |
| HORVU.MOREX.r3.5HG0504840.1 | 0.990941715 | 2.172085864 | 4.805590901 | 1.31E-05 | 0.003897807 | 3.092352702 | transmembrane protein, putative (DUF594) | elongation zone | 6 h |
| HORVU.MOREX.r3.7HG0748070.1 | -1.973762015 | -0.496958177 | -4.796514172 | 1.35E-05 | 0.003955741 | 2.972546252 | Heavy metal transport/detoxification superfamily protein, putati | elongation zone | 6 h |
| HORVU.MOREX.r3.6HG0600000.1 | -1.648680404 | -0.219047099 | -4.793127069 | 1.37E-05 | 0.003955741 | 2.891052276 | Blue copper protein | elongation zone | 6 h |
| HORVU.MOREX.r3.7HG0680960.1 | -2.642732462 | 0.61149754 | -4.749076613 | 1.59E-05 | 0.004422113 | 2.884297231 | electron transporter, putative (Protein of unknown function, DUI | elongation zone | 6 h |
| HORVU.MOREX.r3.5HG0468700.1 | -1.511673438 | 1.52369506 | -4.754597548 | 1.56E-05 | 0.004422113 | 2.874546671 | #N/A | elongation zone | 6 h |
| HORVU.MOREX.r3.3HG0270190.1 | 0.468703429 | 5.543138586 | 4.829850583 | 1.20E-05 | 0.003689135 | 2.865858569 | dihydroflavonol 4-reductase/flavanone protein | elongation zone | 6 h |
| HORVU.MOREX.r3.2HG0207860.1 | -2.002251609 | -2.164021772 | -4.741937003 | 1.63E-05 | 0.004446848 | 2.846789904 | Serine/threonine-protein kinase | elongation zone | 6 h |
| HORVU.MOREX.r3.5HG0519810.1 | 0.573419215 | 7.61792539 | 4.866061342 | 1.06E-05 | 0.00335152 | 2.776597205 | Diphosphomevalonate decarboxylase | elongation zone | 6 h |
| HORVU.MOREX.r3.5HG0532140.1 | -1.781995241 | -1.001133554 | -4.750013417 | 1.59E-05 | 0.004422113 | 2.745681458 | Expansin | elongation zone | 6 h |
| HORVU.MOREX.r3.4HG0413320.1 | -2.076701937 | 2.316272111 | -4.739765317 | 1.65E-05 | 0.004446848 | 2.711744461 | Subtilisin-like protease | elongation zone | 6 h |
| HORVU.MOREX.r3.1HG0079800.1 | 0.612737305 | 7.794068 | 4.843388053 | 1.15E-05 | 0.003572457 | 2.670162339 | Farnesyl diphosphate synthase | elongation zone | 6 h |
| HORVU.MOREX.r3.3HG0329870.1 | -1.626871202 | -1.903215009 | -4.679555651 | 2.03E-05 | 0.005267108 | 2.65195878 | Pectin acetylesterase | elongation zone | 6 h |
| HORVU.MOREX.r3.7HG0685920.1 | -0.691070243 | 4.571683277 | -4.726928497 | 1.72E-05 | 0.00458846 | 2.492304 | Solute carrier family 35 protein | elongation zone | 6 h |
| HORVU.MOREX.r3.4HG0380950.1 | -0.687805181 | 4.409828858 | -4.701192657 | 1.88E-05 | 0.004950936 | 2.452080927 | ABC(ATP-binding) family transporter | elongation zone | 6 h |
| HORVU.MOREX.r3.7HG0634520.1 | -1.65769725 | -2.239871453 | -4.603537684 | 2.63E-05 | 0.006432844 | 2.450305755 | NBS-LRR-like resistance protein | elongation zone | 6 h |
| HORVU.MOREX.r3.1HG0034450.1 | -1.866428293 | 0.621391691 | -4.645227969 | 2.28E-05 | 0.005793016 | 2.407088615 | Retrovirus-related Pol polyprotein from transposon TNT 1-94 | elongation zone | 6 h |
| HORVU.MOREX.r3.7HG0724080.1 | 1.764464637 | 2.23197157 | 4.581041349 | 2.84E-05 | 0.006612707 | 2.381617739 | 6,7-dimethyl-8-ribityllumazine synthase | elongation zone | 6 h |
| HORVU.MOREX.r3.7HG0661290.1 | -0.837154875 | 0.704847423 | -4.633055835 | 2.38E-05 | 0.005954966 | 2.348395921 | Inositol-tetrakisphosphate 1-kinase | elongation zone | 6 h |
| HORVU.MOREX.r3.3HG0254260.1 | -0.825521242 | 4.868531812 | -4.644648949 | 2.29E-05 | 0.005793016 | 2.279995286 | Kinesin, putative | elongation zone | 6 h |
| HORVU.MOREX.r3.7HG0647950.1 | -2.682472799 | -1.041996542 | -4.559469655 | 3.06E-05 | 0.006755938 | 2.255867347 | Peroxidase | elongation zone | 6 h |
| HORVU.MOREX.r3.1HG0026900.1 | -1.412420154 | 1.690413985 | -4.54195166 | 3.25E-05 | 0.006941783 | 2.204195351 | Protein apaG | elongation zone | 6 h |
| HORVU.MOREX.r3.1HG0038800.1 | -0.796912567 | 2.879103431 | -4.572879316 | 2.92E-05 | 0.006612707 | 2.187208787 | Protein kinase | elongation zone | 6 h |
| HORVU.MOREX.r3.2HG0196800.1 | -2.258666615 | -0.209944285 | -4.540646951 | 3.27E-05 | 0.006941783 | 2.182634132 | Multidrug resistance protein ABC transporter family | elongation zone | 6 h |
| HORVU.MOREX.r3.4HG0408270.1 | -1.422349983 | -2.271186585 | -4.51595474 | 3.55E-05 | 0.007361644 | 2.166218441 | LOB domain-containing protein, putative | elongation zone | 6 h |
| HORVU.MOREX.r3.2HG0210510.1 | -1.067113774 | -0.619193937 | -4.573037577 | 2.92E-05 | 0.006612707 | 2.165181322 | Rapid alkalinization factor 1 | elongation zone | 6 h |
| HORVU.MOREX.r3.1HG0068410.1 | -0.43540793 | 4.707522572 | -4.583131491 | 2.82E-05 | 0.006612707 | 2.124589145 | E3 ubiquitin-protein ligase | elongation zone | 6 h |

**Table S5** Continued.

| ID | logFC | AveExpr | t | P.Value | adj.P.Val | B | gene description | root zone | time point |
|---|---|---|---|---|---|---|---|---|---|
| HORVU.MOREX.r3.1HG0069410.1 | -1.737504924 | 3.463734972 | -4.561018935 | 3.05E-05 | 0.006755938 | 2.056849284 | Aldehyde dehydrogenase | elongation zone | 6 h |
| HORVU.MOREX.r3.7HG0676040.1 | -0.946662139 | -0.533828415 | -4.494385217 | 3.82E-05 | 0.007613546 | 2.052938204 | Decaprenyl-phosphate N-acetylglucosaminephosphotransferase | elongation zone | 6 h |
| HORVU.MOREX.r3.6HG0559680.1 | -2.581853641 | 0.573580493 | -4.527845665 | 3.41E-05 | 0.007141623 | 2.04209267 | NADPH:quinone oxidoreductase | elongation zone | 6 h |
| HORVU.MOREX.r3.4HG0393520.1 | -1.081828956 | 3.798189813 | -4.488754278 | 3.90E-05 | 0.007613546 | 1.957383638 | Microtubule associated protein family protein, putative, express | elongation zone | 6 h |
| HORVU.MOREX.r3.1HG0088530.1 | -1.252124111 | 0.05264283 | -4.440442789 | 4.59E-05 | 0.008407674 | 1.948881935 | Cytochrome b561 domain-containing protein | elongation zone | 6 h |
| HORVU.MOREX.r3.5HG0513020.1 | 0.430410759 | 6.516881897 | 4.58660244 | 2.79E-05 | 0.006612707 | 1.936110168 | Transmembrane protein 147 | elongation zone | 6 h |
| HORVU.MOREX.r3.1HG0011440.1 | -2.281968739 | -1.581106574 | -4.539180506 | 3.28E-05 | 0.006941783 | 1.931966525 | Glucan endo-1,3-beta-glucosidase, putative | elongation zone | 6 h |
| HORVU.MOREX.r3.3HG0297330.1 | 0.539603786 | 5.598321606 | 4.551018823 | 3.15E-05 | 0.006879758 | 1.911558133 | Receptor-like kinase | elongation zone | 6 h |
| HORVU.MOREX.r3.6HG0615840.1 | -2.744951602 | -2.27410401 | -4.627031686 | 2.43E-05 | 0.00600621 | 1.910875403 | Metalloendoproteinase 1 | elongation zone | 6 h |
| HORVU.MOREX.r3.2HG0191180.1 | 0.580323494 | 4.405018466 | 4.48919171 | 3.89E-05 | 0.007613546 | 1.902204661 | Xanthine phosphoribosyltransferase | elongation zone | 6 h |
| HORVU.MOREX.r3.6HG0613230.1 | 0.476650804 | 6.00935735 | 4.572101272 | 2.93E-05 | 0.006612707 | 1.895853911 | Allergen, putative | elongation zone | 6 h |
| HORVU.MOREX.r3.3HG0329040.1 | -2.146745974 | -0.454710292 | -4.440881524 | 4.58E-05 | 0.008407674 | 1.877809408 | Vacuolar-processing enzyme | elongation zone | 6 h |
| HORVU.MOREX.r3.6HG0599090.1 | -1.310914408 | -2.428665549 | -4.409162925 | 5.10E-05 | 0.008964956 | 1.842611423 | Vacuolar iron transporter | elongation zone | 6 h |
| HORVU.MOREX.r3.3HG0291580.1 | 0.782303409 | 2.805578274 | 4.416594877 | 4.97E-05 | 0.008875539 | 1.831090662 | Lysosomal Pro-X carboxypeptidase | elongation zone | 6 h |
| HORVU.MOREX.r3.6HG0627570.1 | 0.718674028 | 4.766956665 | 4.483648763 | 3.96E-05 | 0.007673608 | 1.814434455 | Immunoglobulin A1 protease autotransporter | elongation zone | 6 h |
| HORVU.MOREX.r3.1HG0086370.1 | -2.172626164 | -3.209110522 | -4.461466808 | 4.27E-05 | 0.008042578 | 1.810709604 | Glyoxal oxidase | elongation zone | 6 h |
| HORVU.MOREX.r3.1HG0091820.1 | 0.807918085 | 5.321522929 | 4.470272771 | 4.15E-05 | 0.00794144 | 1.778044025 | Glutamyl-tRNA (Gln) amidotransferase subunit A | elongation zone | 6 h |
| HORVU.MOREX.r3.4HG0342160.1 | 0.522337716 | 5.308472026 | 4.489007835 | 3.89E-05 | 0.007613546 | 1.768949566 | Presenilin | elongation zone | 6 h |
| HORVU.MOREX.r3.3HG0234970.1 | -0.755117158 | 0.091121871 | -4.458879424 | 4.31E-05 | 0.008042578 | 1.766942527 | 6,7-dimethyl-8-ribityllumazine synthase | elongation zone | 6 h |
| HORVU.MOREX.r3.5HG0494040.1 | -1.831724794 | -3.000857515 | -4.402406455 | 5.22E-05 | 0.009074007 | 1.738029167 | Auxin efflux carrier component | elongation zone | 6 h |
| HORVU.MOREX.r3.3HG0242880.1 | -1.131554244 | -0.890008185 | -4.380148607 | 5.62E-05 | 0.009382875 | 1.732053631 | Phloem protein 2-like protein | elongation zone | 6 h |
| HORVU.MOREX.r3.5HG0425000.1 | -3.529383361 | -0.918831697 | -4.573038409 | 2.92E-05 | 0.006612707 | 1.700066573 | Transposon protein, putative, Pong sub-class | elongation zone | 6 h |
| HORVU.MOREX.r3.6HG0601830.1 | -1.481295163 | -2.059555704 | -4.354773318 | 6.12E-05 | 0.00981914 | 1.657143854 | C2 calcium/lipid-binding and GRAM domain protein | elongation zone | 6 h |
| HORVU.MOREX.r3.4HG0351750.1 | -1.818178955 | -1.823804439 | -4.368974247 | 5.84E-05 | 0.009607858 | 1.649633027 | BTB/POZ domain-containing protein | elongation zone | 6 h |
| HORVU.MOREX.r3.6HG0624390.1 | -1.524798697 | 1.652699025 | -4.340103303 | 6.43E-05 | 0.010155862 | 1.630072027 | Two-component response regulator | elongation zone | 6 h |
| HORVU.MOREX.r3.7HG0655210.1 | -1.841854981 | -1.689922301 | -4.365743885 | 5.90E-05 | 0.009607858 | 1.626060229 | Peroxidase | elongation zone | 6 h |
| HORVU.MOREX.r3.3HG0330200.1 | -1.243251393 | 1.169812259 | -4.468027223 | 4.18E-05 | 0.00794144 | 1.592256168 | Aquaporin | elongation zone | 6 h |
| HORVU.MOREX.r3.1HG0017280.1 | -1.765314558 | -0.330109732 | -4.383406651 | 5.56E-05 | 0.009382875 | 1.571695713 | Peroxidase | elongation zone | 6 h |
| HORVU.MOREX.r3.3HG0275090.1 | -2.036079506 | -3.212508152 | -4.389313057 | 5.45E-05 | 0.009325267 | 1.567163157 | SWI/SNF-related matrix-associated actin-dependent regulator of | elongation zone | 6 h |
| HORVU.MOREX.r3.1HG0072580.1 | -0.653030101 | 5.490114394 | -4.421404573 | 4.89E-05 | 0.008809167 | 1.555926018 | Aminotransferase-related family protein | elongation zone | 6 h |
| HORVU.MOREX.r3.2HG0170230.1 | -0.688713481 | 3.492538265 | -4.343120367 | 6.37E-05 | 0.010131561 | 1.55421533 | Smr domain containing protein | elongation zone | 6 h |
| HORVU.MOREX.r3.2HG0096800.1 | -0.940855915 | 4.691800049 | -4.408515067 | 5.11E-05 | 0.008964956 | 1.55195701 | Flavin-containing monooxygenase | elongation zone | 6 h |
| HORVU.MOREX.r3.5HG0483860.1 | 0.643096146 | 3.316156884 | 4.316997505 | 6.95E-05 | 0.010725562 | 1.49802978 | RING/U-box superfamily protein | elongation zone | 6 h |
| HORVU.MOREX.r3.1HG0075440.1 | -0.953320761 | 1.650312221 | -4.366973554 | 5.88E-05 | 0.009607858 | 1.491539976 | Protein NRT1/ PTR FAMILY 5.5 | elongation zone | 6 h |
| HORVU.MOREX.r3.4HG0350900.1 | 0.398899633 | 7.97599371 | 4.507261227 | 3.66E-05 | 0.007506841 | 1.488266334 | UDP-glucose 6-dehydrogenase | elongation zone | 6 h |
| HORVU.MOREX.r3.7HG0674720.1 | -1.950135509 | -1.463275625 | -4.290977406 | 7.58E-05 | 0.011115 | 1.487489686 | Epoxide hydrolase 2 | elongation zone | 6 h |
| HORVU.MOREX.r3.6HG0620820.1 | -0.936220749 | 3.246758451 | -4.298752377 | 7.39E-05 | 0.011107173 | 1.470768152 | Pectin lyase-like superfamily protein | elongation zone | 6 h |
| HORVU.MOREX.r3.6HG0626650.1 | 0.326313709 | 8.259288679 | 4.493677457 | 3.83E-05 | 0.007613546 | 1.461863311 | 26S protease regulatory subunit-like protein | elongation zone | 6 h |
| HORVU.MOREX.r3.7HG0684940.1 | -0.720838457 | 4.72025707 | -4.392633116 | 5.39E-05 | 0.009299057 | 1.458140361 | Zinc finger constans-like protein | elongation zone | 6 h |
| HORVU.MOREX.r3.5HG0510830.1 | 0.980424002 | 5.120306759 | 4.319319864 | 6.90E-05 | 0.010723158 | 1.406130191 | #N/A | elongation zone | 6 h |
| HORVU.MOREX.r3.2HG0136150.1 | 3.428126635 | -1.262543021 | 4.432126313 | 4.72E-05 | 0.008570943 | 1.391408015 | Cyclin family protein | elongation zone | 6 h |

**Table S5** Continued.

| ID | logFC | AveExpr | t | P.Value | adj.P.Val | B | gene description | root zone | time point |
|---|---|---|---|---|---|---|---|---|---|
| HORVU.MOREX.r3.7HG0648610.1 | 0.373099131 | 6.01586743 | 4.363584643 | 5.95E-05 | 0.009607858 | 1.29140352 | Carboxypeptidase | elongation zone | 6 h |
| HORVU.MOREX.r3.4HG0355210.1 | -2.465957966 | -0.137693228 | -4.229300421 | 9.31E-05 | 0.012247977 | 1.277185858 | transcription factor-like protein | elongation zone | 6 h |
| HORVU.MOREX.r3.7HG0655680.1 | -2.026967996 | -1.898894256 | -4.238220662 | 9.04E-05 | 0.012075989 | 1.2730143 | Serine/threonine-protein kinase | elongation zone | 6 h |
| HORVU.MOREX.r3.4HG0354980.1 | -0.873012559 | 2.412831318 | -4.257532155 | 8.48E-05 | 0.011833919 | 1.266977237 | #N/A | elongation zone | 6 h |
| HORVU.MOREX.r3.1HG0025320.1 | -1.57590765 | 0.572544846 | -4.203541999 | 0.0001 | 0.012845495 | 1.233028551 | Receptor-like kinase, putative | elongation zone | 6 h |
| HORVU.MOREX.r3.3HG0330980.1 | 0.382819961 | 6.458067382 | 4.380404172 | 5.62E-05 | 0.009382875 | 1.224501131 | Protein IQ-DOMAIN 1 | elongation zone | 6 h |
| HORVU.MOREX.r3.5HG0537150.1 | -1.141844089 | -2.246878812 | -4.174842061 | 0.00011 | 0.013700934 | 1.147008989 | Protein DETOXIFICATION | elongation zone | 6 h |
| HORVU.MOREX.r3.4HG0337950.1 | -2.281421129 | -0.296307238 | -4.286700326 | 7.69E-05 | 0.01119483 | 1.142024029 | 65-kDa microtubule-associated-like protein | elongation zone | 6 h |
| HORVU.MOREX.r3.4HG0390450.1 | -2.322171728 | -2.315588818 | -4.181763093 | 0.00011 | 0.013472048 | 1.141178733 | #N/A | elongation zone | 6 h |
| HORVU.MOREX.r3.4HG0344830.1 | 0.590822337 | 4.535567699 | 4.263337935 | 8.32E-05 | 0.011767538 | 1.135671865 | NBS-LRR disease resistance protein, putative, expressed | elongation zone | 6 h |
| HORVU.MOREX.r3.4HG0372230.1 | -2.125195208 | -1.629808544 | -4.246237208 | 8.80E-05 | 0.01204116 | 1.131489209 | Transposon protein, putative, CACTA, En/Spm sub-class, expresse | elongation zone | 6 h |
| HORVU.MOREX.r3.5HG0519120.1 | -1.326655212 | -0.875101352 | -4.240277143 | 8.98E-05 | 0.012075989 | 1.122887799 | CASP-like protein | elongation zone | 6 h |
| HORVU.MOREX.r3.6HG0550600.1 | -1.046749539 | 1.720715964 | -4.331227687 | 6.63E-05 | 0.010382654 | 1.113060496 | Protein ROOT HAIR DEFECTIVE 3 homolog | elongation zone | 6 h |
| HORVU.MOREX.r3.7HG0724090.1 | 3.11047028 | 0.704867316 | 4.237021147 | 9.08E-05 | 0.012075989 | 1.105027383 | RING/U-box superfamily protein | elongation zone | 6 h |
| HORVU.MOREX.r3.6HG0619410.1 | -1.017910577 | 1.820333919 | -4.235497898 | 9.12E-05 | 0.012075989 | 1.104114927 | Flavonol synthase | elongation zone | 6 h |
| HORVU.MOREX.r3.4HG0407230.1 | -1.748100229 | -0.944900136 | -4.164125519 | 0.00012 | 0.013781372 | 1.082116014 | Flowering-promoting factor 1-like protein 1 | elongation zone | 6 h |
| HORVU.MOREX.r3.1HG0050700.1 | -1.382940743 | 2.994863261 | -4.264135513 | 8.29E-05 | 0.011767538 | 1.076018524 | ATP-dependent zinc metalloprotease FtsH | elongation zone | 6 h |
| HORVU.MOREX.r3.1HG0055240.1 | -1.931442357 | -3.14720018 | -4.186154225 | 0.00011 | 0.013427958 | 1.052779012 | Acetyl-coenzyme A carboxylase carboxyl transferase subunit beta | elongation zone | 6 h |
| HORVU.MOREX.r3.5HG0511820.1 | -2.949386381 | -3.38085013 | -4.222424557 | 9.53E-05 | 0.012294138 | 1.049066927 | Cyclic nucleotide-gated channel | elongation zone | 6 h |
| HORVU.MOREX.r3.3HG0329950.1 | -1.399744779 | 1.75889036 | -4.148155103 | 0.00012 | 0.014195784 | 1.032670336 | Pectin acetylesterase | elongation zone | 6 h |
| HORVU.MOREX.r3.3HG0293570.1 | 2.45269119 | -0.387941199 | 4.242544469 | 8.91E-05 | 0.012075989 | 1.005691981 | BTB/POZ domain-containing protein | elongation zone | 6 h |
| HORVU.MOREX.r3.6HG0540020.1 | -2.305406832 | -2.701095093 | -4.119142437 | 0.00013 | 0.015184882 | 0.976942379 | Ubiquitin carboxyl-terminal hydrolase 2 | elongation zone | 6 h |
| HORVU.MOREX.r3.2HG0161630.1 | -1.020148712 | 1.307163675 | -4.131270819 | 0.00013 | 0.01483784 | 0.97498662 | basic helix-loop-helix (bHLH) DNA-binding superfamily protein | elongation zone | 6 h |
| HORVU.MOREX.r3.3HG0272430.1 | 0.478577692 | 5.817079707 | 4.252535379 | 8.62E-05 | 0.011950976 | 0.919624673 | Pectin lyase-like superfamily protein | elongation zone | 6 h |
| HORVU.MOREX.r3.2HG0131630.1 | -2.665484743 | -3.028166822 | -4.132888102 | 0.00013 | 0.01483784 | 0.910190016 | vacuolar sorting-associated protein (DUF946) | elongation zone | 6 h |
| HORVU.MOREX.r3.7HG0661480.1 | 0.335871265 | 7.365151763 | 4.29333575 | 7.52E-05 | 0.011107173 | 0.887681238 | Alpha-amylase 2 | elongation zone | 6 h |
| HORVU.MOREX.r3.4HG0378580.1 | 0.539006381 | 5.353083837 | 4.224416317 | 9.46E-05 | 0.01229049 | 0.876971332 | Transcription factor | elongation zone | 6 h |
| HORVU.MOREX.r3.7HG0737450.1 | 1.000026479 | 2.956707885 | 4.090076162 | 0.00015 | 0.015995535 | 0.876334311 | Phosphoglycerate kinase | elongation zone | 6 h |
| HORVU.MOREX.r3.3HG0246670.1 | -1.447321914 | -1.917828013 | -4.08210546 | 0.00015 | 0.016166055 | 0.873698317 | DUF4228 domain protein | elongation zone | 6 h |
| HORVU.MOREX.r3.7HG0707700.1 | -1.269395343 | -2.42616811 | -4.081352052 | 0.00015 | 0.016166055 | 0.872980507 | Late embryogenesis abundant (LEA) hydroxyproline-rich glycopro | elongation zone | 6 h |
| HORVU.MOREX.r3.2HG0142850.1 | 0.310553124 | 6.438473516 | 4.249259576 | 8.71E-05 | 0.012000787 | 0.872282947 | Organic solute transporter-like | elongation zone | 6 h |
| HORVU.MOREX.r3.6HG0575710.1 | 0.521056028 | 4.807664746 | 4.184584419 | 0.00011 | 0.013427958 | 0.863389314 | Kynurenine formamidase | elongation zone | 6 h |
| HORVU.MOREX.r3.2HG0120320.1 | -0.877368159 | 1.685244219 | -4.271644222 | 8.09E-05 | 0.011606539 | 0.858856116 | Peroxidase | elongation zone | 6 h |
| HORVU.MOREX.r3.7HG0744040.1 | -1.057940304 | 1.177534674 | -4.080591753 | 0.00015 | 0.016166055 | 0.858414615 | Transcription factor | elongation zone | 6 h |
| HORVU.MOREX.r3.7HG0684880.1 | -0.853998862 | 3.761266823 | -4.158815352 | 0.00012 | 0.013943663 | 0.842152766 | meiosis chromosome segregation family protein | elongation zone | 6 h |
| HORVU.MOREX.r3.1HG0081950.1 | -2.141802047 | -2.05546379 | -4.068492722 | 0.00016 | 0.016551728 | 0.830369243 | Peroxidase | elongation zone | 6 h |
| HORVU.MOREX.r3.3HG0253860.1 | -2.836916007 | -0.092758097 | -4.097648553 | 0.00014 | 0.015896607 | 0.820722141 | Rapid alkalinization factor (RALF) family protein | elongation zone | 6 h |
| HORVU.MOREX.r3.1HG0065130.1 | -1.554171213 | 1.689963316 | -4.090138529 | 0.00015 | 0.015995535 | 0.810438353 | NAC domain-containing protein | elongation zone | 6 h |
| HORVU.MOREX.r3.4HG0344370.1 | -1.430817839 | -2.722566782 | -4.061756204 | 0.00016 | 0.0166646 | 0.8087306 | Glucuronoxylan 4-O-methyltransferase | elongation zone | 6 h |
| HORVU.MOREX.r3.5HG0426110.1 | -2.085521811 | -3.224786885 | -4.104777158 | 0.00014 | 0.015743485 | 0.797852564 | NBS-LRR class disease resistance protein | elongation zone | 6 h |
| HORVU.MOREX.r3.4HG0403620.1 | -1.524790505 | 0.126434613 | -4.226188264 | 9.41E-05 | 0.01229049 | 0.78400056 | #N/A | elongation zone | 6 h |

**Table S5** Continued.

| ID | logFC | AveExpr | t | P.Value | adj.P.Val | B | gene description | root zone | time point |
|---|---|---|---|---|---|---|---|---|---|
| HORVU.MOREX.r3.1HG0075590.1 | 0.32236971 | 8.23086424 | 4.293977264 | 7.51E-05 | 0.011107173 | 0.772006037 | Coiled-coil domain-containing protein 47 | elongation zone | 6 h |
| HORVU.MOREX.r3.4HG0402730.1 | -1.733601611 | -1.081620137 | -4.112811115 | 0.00014 | 0.01541845 | 0.738749119 | Homeobox protein BEL1 like | elongation zone | 6 h |
| HORVU.MOREX.r3.6HG0552040.1 | 0.237654174 | 8.106568618 | 4.276364677 | 7.96E-05 | 0.011505838 | 0.738357035 | #N/A | elongation zone | 6 h |
| HORVU.MOREX.r3.1HG0006230.1 | 0.526358707 | 5.574000008 | 4.16887669 | 0.00011 | 0.013703395 | 0.727987234 | Disease resistance protein RPM1 | elongation zone | 6 h |
| HORVU.MOREX.r3.3HG0222130.1 | 0.363013413 | 8.795276054 | 4.29700533 | 7.43E-05 | 0.011107173 | 0.721759669 | Transmembrane protein 214 | elongation zone | 6 h |
| HORVU.MOREX.r3.7HG0721650.1 | 0.374062496 | 5.578373097 | 4.190751872 | 0.00011 | 0.013318454 | 0.716487556 | Novel plant snare, putative | elongation zone | 6 h |
| HORVU.MOREX.r3.3HG0313090.1 | -0.973454075 | 2.944420659 | -4.080430752 | 0.00015 | 0.016166055 | 0.707330289 | Thiamin pyrophosphokinase 1 | elongation zone | 6 h |
| HORVU.MOREX.r3.2HG0181680.1 | -1.917355076 | -1.908817715 | -4.054967922 | 0.00017 | 0.016720589 | 0.707040615 | CASP-like protein | elongation zone | 6 h |
| HORVU.MOREX.r3.5HG0466320.1 | -0.391907084 | 6.324949057 | -4.204694022 | 0.0001 | 0.012845495 | 0.697018543 | Phosphoinositide phospholipase C | elongation zone | 6 h |
| HORVU.MOREX.r3.6HG0612180.1 | 0.675229267 | 6.264662645 | 4.169267023 | 0.00011 | 0.013703395 | 0.677123183 | NHL repeat-containing protein-like | elongation zone | 6 h |
| HORVU.MOREX.r3.1HG0094770.1 | 0.715074383 | 3.670278309 | 4.053118196 | 0.00017 | 0.016720589 | 0.667316171 | Transmembrane protein, putative | elongation zone | 6 h |
| HORVU.MOREX.r3.4HG0337110.1 | 0.349044245 | 7.514078085 | 4.219504648 | 9.62E-05 | 0.012336161 | 0.642767094 | Coatomer subunit alpha | elongation zone | 6 h |
| HORVU.MOREX.r3.6HG0589860.1 | -0.936026482 | 4.121931249 | -4.09685916 | 0.00014 | 0.015896607 | 0.632999008 | Kinase family protein | elongation zone | 6 h |
| HORVU.MOREX.r3.7HG0667060.1 | -1.308924041 | 2.126304133 | -4.007883844 | 0.00019 | 0.018284709 | 0.619126833 | target of early activation tagged (EAT) 2 | elongation zone | 6 h |
| HORVU.MOREX.r3.7HG0641750.1 | -2.672995114 | -2.739012037 | -3.996522601 | 0.0002 | 0.018702211 | 0.614380054 | Aspartic proteinase nepenthesin-2 | elongation zone | 6 h |
| HORVU.MOREX.r3.4HG0415490.1 | -1.836030917 | -1.784637332 | -4.054821235 | 0.00017 | 0.016720589 | 0.611857737 | 14 kDa proline-rich protein DC2.15 | elongation zone | 6 h |
| HORVU.MOREX.r3.4HG0412370.1 | -2.127962639 | -1.425115928 | -4.010430538 | 0.00019 | 0.018284709 | 0.605327166 | Homeobox protein knotted-1, putative | elongation zone | 6 h |
| HORVU.MOREX.r3.3HG0313320.1 | -2.109205885 | -2.456522524 | -3.983412742 | 0.00021 | 0.018906489 | 0.58880125 | Potassium transporter | elongation zone | 6 h |
| HORVU.MOREX.r3.6HG0610980.1 | -1.396593689 | -1.883261679 | -4.008762756 | 0.00019 | 0.018284709 | 0.586320574 | Nucleotide/sugar transporter family protein | elongation zone | 6 h |
| HORVU.MOREX.r3.7HG0722360.1 | -1.607779447 | 0.66493112 | -4.038013658 | 0.00017 | 0.017309173 | 0.584736752 | Receptor-like protein kinase | elongation zone | 6 h |
| HORVU.MOREX.r3.2HG0187430.1 | 0.526471942 | 5.203928279 | 4.12778367 | 0.00013 | 0.01492498 | 0.576617883 | Transmembrane emp24 domain-containing protein | elongation zone | 6 h |
| HORVU.MOREX.r3.3HG0252610.1 | 0.43712499 | 7.233438842 | 4.167794809 | 0.00011 | 0.013703395 | 0.573042954 | Protein kinase | elongation zone | 6 h |
| HORVU.MOREX.r3.7HG0679050.1 | -0.692966191 | 4.002867978 | -4.047329903 | 0.00017 | 0.016955841 | 0.550752884 | Upstream activation factor subunit spp27 | elongation zone | 6 h |
| HORVU.MOREX.r3.2HG0168110.1 | -3.142973101 | -1.271051778 | -4.260614149 | 8.39E-05 | 0.011793388 | 0.548390176 | Protein SET DOMAIN GROUP 41 | elongation zone | 6 h |
| HORVU.MOREX.r3.4HG0369570.1 | 0.242070995 | 10.41352761 | 4.312996419 | 7.05E-05 | 0.010788971 | 0.546739372 | Alpha-1,4-glucan-protein synthase [UDP-forming] | elongation zone | 6 h |
| HORVU.MOREX.r3.2HG0194620.1 | -2.514314519 | -2.147983455 | -4.009796329 | 0.00019 | 0.018284709 | 0.524522797 | F-box protein | elongation zone | 6 h |
| HORVU.MOREX.r3.6HG0567780.1 | 0.716660177 | 3.016752833 | 3.992282346 | 0.0002 | 0.018702211 | 0.514919098 | disease resistance protein (TIR-NBS-LRR class) | elongation zone | 6 h |
| HORVU.MOREX.r3.3HG0245250.1 | -2.379427167 | -1.643307554 | -3.996452879 | 0.0002 | 0.018702211 | 0.489587225 | Cysteine protease, putative | elongation zone | 6 h |
| HORVU.MOREX.r3.7HG0694090.1 | 0.353560018 | 5.778504944 | 4.123141007 | 0.00013 | 0.015070113 | 0.483992615 | Rer1 protein, putative | elongation zone | 6 h |
| HORVU.MOREX.r3.6HG0541250.1 | 0.273508196 | 7.143194567 | 4.149322056 | 0.00012 | 0.014195784 | 0.469710791 | Acetate kinase | elongation zone | 6 h |
| HORVU.MOREX.r3.2HG0205990.1 | -2.146107509 | -2.557658252 | -3.951496218 | 0.00023 | 0.020527378 | 0.46736824 | Glutaredoxin family protein, putative | elongation zone | 6 h |
| HORVU.MOREX.r3.5HG0526250.1 | 0.345237407 | 12.0381843 | 4.304781961 | 7.24E-05 | 0.011007254 | 0.461181975 | 70 kDa heat shock protein | elongation zone | 6 h |
| HORVU.MOREX.r3.3HG0309270.1 | 0.356544664 | 8.53260482 | 4.167614186 | 0.00011 | 0.013703395 | 0.433361136 | Proline--tRNA ligase | elongation zone | 6 h |
| HORVU.MOREX.r3.4HG0342080.1 | -0.842240504 | 4.067325035 | -4.07310794 | 0.00016 | 0.016387742 | 0.427787963 | Adenine/guanine permease | elongation zone | 6 h |
| HORVU.MOREX.r3.6HG0581570.1 | 0.419759835 | 4.929735327 | 4.043151288 | 0.00017 | 0.017104733 | 0.421590684 | Leucine-rich repeat-containing protein 1 | elongation zone | 6 h |
| HORVU.MOREX.r3.2HG0218010.1 | 0.401955758 | 8.228324079 | 4.154422105 | 0.00012 | 0.01406566 | 0.362793053 | Dolichyl-diphosphooligosaccharide--protein glycosyltransferase s | elongation zone | 6 h |
| HORVU.MOREX.r3.3HG0305440.1 | -1.669627587 | -2.501750371 | -3.902353607 | 0.00027 | 0.021949523 | 0.357981485 | Late embryogenesis abundant protein | elongation zone | 6 h |
| HORVU.MOREX.r3.6HG0632270.1 | 0.570025075 | 3.993704775 | 3.987539398 | 0.00021 | 0.018842535 | 0.356438417 | Leucine-rich repeat family protein | elongation zone | 6 h |
| HORVU.MOREX.r3.5HG0485610.1 | -1.500600981 | 0.794719635 | -3.933336494 | 0.00024 | 0.021333038 | 0.355238729 | Cytochrome P450, putative | elongation zone | 6 h |
| HORVU.MOREX.r3.3HG0311410.1 | -0.49145949 | 5.699117557 | -4.031450942 | 0.00018 | 0.017513475 | 0.347655058 | F box protein | elongation zone | 6 h |
| HORVU.MOREX.r3.5HG0458010.1 | -0.393277383 | 5.942045141 | -4.062742441 | 0.00016 | 0.0166646 | 0.336334701 | Protein kinase, putative | elongation zone | 6 h |

**Table S5** Continued.

| ID | logFC | AveExpr | t | P.Value | adj.P.Val | B | gene description | root zone | time point |
|---|---|---|---|---|---|---|---|---|---|
| HORVU.MOREX.r3.6HG0605940.1 | -0.868259182 | 3.17964847 | -3.991464562 | 0.0002 | 0.018702211 | 0.328231967 | Xyloglucan endotransglucosylase/hydrolase | elongation zone | 6 h |
| HORVU.MOREX.r3.6HG0541940.1 | 0.39006774 | 6.76425912 | 4.094472053 | 0.00015 | 0.015935666 | 0.324753005 | S-adenosyl-L-methionine-dependent methyltransferases superfa | elongation zone | 6 h |
| HORVU.MOREX.r3.7HG0658560.1 | -0.963861332 | 3.664515124 | -3.905393734 | 0.00027 | 0.021822902 | 0.284550176 | Kinase family protein | elongation zone | 6 h |
| HORVU.MOREX.r3.6HG0631040.1 | -0.92405237 | 2.526682694 | -3.93830977 | 0.00024 | 0.021271955 | 0.278154919 | Mesoderm induction early response protein 1, putative isoform 1 | elongation zone | 6 h |
| HORVU.MOREX.r3.2HG0178160.1 | 0.666237379 | 7.879945441 | 4.099295252 | 0.00014 | 0.015896607 | 0.274563851 | HTH-type transcriptional regulator YidZ | elongation zone | 6 h |
| HORVU.MOREX.r3.2HG0213020.1 | -1.229744423 | 3.221450174 | -3.893688323 | 0.00028 | 0.022216127 | 0.269418936 | Protein F12F1.11-, putative | elongation zone | 6 h |
| HORVU.MOREX.r3.6HG0561180.1 | -1.669539359 | -0.688407242 | -3.886100072 | 0.00029 | 0.02238184 | 0.267984347 | Receptor kinase | elongation zone | 6 h |
| HORVU.MOREX.r3.6HG0615600.1 | -2.016650283 | -2.882863566 | -3.912004746 | 0.00026 | 0.021709616 | 0.263883037 | Cytochrome P450, putative | elongation zone | 6 h |
| HORVU.MOREX.r3.6HG0606930.1 | 0.423800413 | 5.680326244 | 4.029103626 | 0.00018 | 0.017563621 | 0.259367434 | 3-ketodihydrosphingosine reductase | elongation zone | 6 h |
| HORVU.MOREX.r3.6HG0573150.1 | -0.522561598 | 5.184389048 | -3.982859121 | 0.00021 | 0.018906489 | 0.248353199 | Copper-transporting ATPase | elongation zone | 6 h |
| HORVU.MOREX.r3.7HG0649240.1 | -1.639045242 | -2.995715777 | -3.906493893 | 0.00027 | 0.021822902 | 0.240879612 | protein kinase family protein | elongation zone | 6 h |
| HORVU.MOREX.r3.2HG0180010.1 | -1.330955496 | 1.125047742 | -4.011410437 | 0.00019 | 0.018284709 | 0.221892077 | #N/A | elongation zone | 6 h |
| HORVU.MOREX.r3.7HG0737340.1 | -1.545629428 | -0.106662143 | -4.032281836 | 0.00018 | 0.017513475 | 0.212505384 | GDA1/CD39 nucleoside phosphatase family protein | elongation zone | 6 h |
| HORVU.MOREX.r3.2HG0162900.2 | 0.80856924 | 2.320158811 | 3.869265595 | 0.0003 | 0.023276244 | 0.20541253 | Aldo-keto reductase | elongation zone | 6 h |
| HORVU.MOREX.r3.2HG0214070.1 | -1.744830889 | -2.623970249 | -3.850904434 | 0.00032 | 0.02415331 | 0.20528132 | AT hook motif DNA-binding family protein, putative | elongation zone | 6 h |
| HORVU.MOREX.r3.2HG0171550.1 | 0.790489854 | 4.111529363 | 3.919551005 | 0.00026 | 0.021417658 | 0.202629984 | Major facilitator superfamily transporter | elongation zone | 6 h |
| HORVU.MOREX.r3.4HG0374910.2 | 0.324275456 | 6.337717806 | 4.053576916 | 0.00017 | 0.016720589 | 0.201981915 | Beta-adaptin-like protein | elongation zone | 6 h |
| HORVU.MOREX.r3.3HG0241320.1 | 0.568118576 | 3.585417651 | 3.931494927 | 0.00025 | 0.021333038 | 0.192702002 | Prolyl 4-hydroxylase alpha subunit, putative | elongation zone | 6 h |
| HORVU.MOREX.r3.3HG0239980.1 | -1.334910759 | 1.292153679 | -3.928287338 | 0.00025 | 0.021333038 | 0.192548623 | Alpha/beta-Hydrolases superfamily protein, putative | elongation zone | 6 h |
| HORVU.MOREX.r3.1HG0039670.1 | 0.289079396 | 7.408424513 | 4.075040613 | 0.00015 | 0.016368541 | 0.191788058 | Calcium ion-binding protein | elongation zone | 6 h |
| HORVU.MOREX.r3.7HG0718170.1 | -1.209497807 | 1.273374933 | -3.845474054 | 0.00032 | 0.024305588 | 0.187861058 | Pleiotropic drug resistance ABC transporter | elongation zone | 6 h |
| HORVU.MOREX.r3.4HG0370380.1 | 0.47958889 | 5.905711064 | 4.009024064 | 0.00019 | 0.018284709 | 0.185123738 | Erlin-2 | elongation zone | 6 h |
| HORVU.MOREX.r3.4HG0383780.1 | -2.419761971 | -0.062687302 | -3.881835714 | 0.00029 | 0.022463016 | 0.183864602 | Laccase | elongation zone | 6 h |
| HORVU.MOREX.r3.4HG0380430.1 | -0.42600792 | 3.638981565 | -3.924639025 | 0.00025 | 0.021375796 | 0.177321242 | Membrane steroid-binding protein | elongation zone | 6 h |
| HORVU.MOREX.r3.5HG0523860.1 | -0.679560428 | 4.167580678 | -3.965166912 | 0.00022 | 0.019727211 | 0.159267997 | Calcium-dependent protein kinase | elongation zone | 6 h |
| HORVU.MOREX.r3.3HG0257310.1 | 0.324842068 | 7.69321866 | 4.065720413 | 0.00016 | 0.016617419 | 0.126582353 | Glucosidase 2 subunit beta | elongation zone | 6 h |
| HORVU.MOREX.r3.6HG0581020.1 | 0.391806348 | 5.648770754 | 3.991230852 | 0.0002 | 0.018702211 | 0.119704078 | Xyloglucan 6-xylosyltransferase 1 | elongation zone | 6 h |
| HORVU.MOREX.r3.6HG0557550.1 | -0.94365046 | 1.605488567 | -3.815945699 | 0.00036 | 0.025487626 | 0.113231556 | CRS2-associated factor 1 | elongation zone | 6 h |
| HORVU.MOREX.r3.5HG0462220.1 | -1.097383812 | 1.070356343 | -3.932592536 | 0.00025 | 0.021333038 | 0.112529195 | Calmodulin protein | elongation zone | 6 h |
| HORVU.MOREX.r3.7HG0664780.1 | -0.859211583 | 4.573395079 | -4.054149436 | 0.00017 | 0.016720589 | 0.110526867 | Patatin | elongation zone | 6 h |
| HORVU.MOREX.r3.6HG0595190.1 | -1.711180164 | -2.072106303 | -3.809083482 | 0.00036 | 0.025778846 | 0.095865769 | GDSL esterase/lipase | elongation zone | 6 h |
| HORVU.MOREX.r3.3HG0293130.1 | -1.124496679 | 3.772343167 | -3.923174218 | 0.00025 | 0.021375796 | 0.075048991 | Eukaryotic aspartyl protease family protein | elongation zone | 6 h |
| HORVU.MOREX.r3.4HG0334360.1 | -0.924191945 | 4.121321037 | -3.982380184 | 0.00021 | 0.018906489 | 0.073898099 | Homeobox protein BEL1 like | elongation zone | 6 h |
| HORVU.MOREX.r3.3HG0276120.1 | -2.051419046 | -1.330753495 | -3.788007048 | 0.00039 | 0.026638848 | 0.035650793 | Phospholipase A1 | elongation zone | 6 h |
| HORVU.MOREX.r3.5HG0483980.1 | -1.424145581 | -1.844541926 | -3.78294743 | 0.0004 | 0.026712078 | 0.019506026 | Cinnamoyl CoA reductase | elongation zone | 6 h |
| HORVU.MOREX.r3.4HG0416170.1 | -0.703163406 | 3.384356338 | -3.935850914 | 0.00024 | 0.021311375 | 0.019177217 | Methionyl-tRNA formyltransferase | elongation zone | 6 h |
| HORVU.MOREX.r3.2HG0141140.1 | -1.286792871 | -2.857994565 | -3.783448513 | 0.0004 | 0.026712078 | 0.016069937 | DUF4228 domain protein | elongation zone | 6 h |
| HORVU.MOREX.r3.2HG0190560.1 | -1.205550001 | -0.224848231 | -3.868342361 | 0.0003 | 0.023276244 | 0.014443599 | #N/A | elongation zone | 6 h |
| HORVU.MOREX.r3.2HG0197920.1 | 2.857684379 | -1.098587388 | 3.847341169 | 0.00032 | 0.024250025 | -0.00147563 | BTB/POZ domain containing protein, expressed | elongation zone | 6 h |
| HORVU.MOREX.r3.4HG0346830.1 | -1.425774146 | -2.756213705 | -3.770713393 | 0.00041 | 0.027115872 | -0.01269217 | WEB family protein | elongation zone | 6 h |
| HORVU.MOREX.r3.5HG0419590.1 | -0.566743523 | 6.133126066 | -4.006614848 | 0.00019 | 0.018284709 | -0.03148458 | #N/A | elongation zone | 6 h |

**Table S5** Continued.

| ID | logFC | AveExpr | t | P.Value | adj.P.Val | B | gene description | root zone | time point |
|---|---|---|---|---|---|---|---|---|---|
| HORVU.MOREX.r3.5HG0527310.1 | -1.840336786 | -3.15597805 | -3.778567267 | 0.0004 | 0.026862598 | -0.03817673 | Protein NRT1/ PTR FAMILY 5.5 | elongation zone | 6 h |
| HORVU.MOREX.r3.2HG0144000.1 | -0.410227815 | 5.798118458 | -3.937757137 | 0.00024 | 0.021271955 | -0.04164743 | Zeta-carotene desaturase | elongation zone | 6 h |
| HORVU.MOREX.r3.1HG0081120.1 | 1.63134628 | 1.005476641 | 3.757974735 | 0.00043 | 0.027892198 | -0.04594231 | DNA mismatch repair protein MutL | elongation zone | 6 h |
| HORVU.MOREX.r3.5HG0472810.1 | -1.006116412 | 2.371977796 | -3.977873609 | 0.00021 | 0.019099888 | -0.04710299 | Aquaporin | elongation zone | 6 h |
| HORVU.MOREX.r3.3HG0312780.1 | 0.69776507 | 3.639594263 | 3.828028142 | 0.00034 | 0.024875194 | -0.04726394 | NBS-LRR disease resistance protein, putative, expressed | elongation zone | 6 h |
| HORVU.MOREX.r3.2HG0130960.1 | -0.838721146 | 1.762022677 | -3.770976152 | 0.00041 | 0.027115872 | -0.06646704 | Benzene 1,2-dioxygenase subunit alpha | elongation zone | 6 h |
| HORVU.MOREX.r3.1HG0015750.1 | -1.893034039 | -2.912149854 | -3.750892135 | 0.00044 | 0.028211116 | -0.06888109 | transcription factor, putative (Protein of unknown function, DUF | elongation zone | 6 h |
| HORVU.MOREX.r3.3HG0299740.1 | -0.455980635 | 5.806671811 | -3.927662719 | 0.00025 | 0.021333038 | -0.06929315 | Tubby-like F-box protein | elongation zone | 6 h |
| HORVU.MOREX.r3.1HG0050920.1 | -1.023400497 | 2.478719451 | -3.924456478 | 0.00025 | 0.021375796 | -0.08119197 | Trehalase | elongation zone | 6 h |
| HORVU.MOREX.r3.5HG0469300.1 | -1.57608715 | 0.250215668 | -3.796291663 | 0.00038 | 0.026284234 | -0.08800144 | CCCH-type zinc finger protein with ARM repeat domain-containir | elongation zone | 6 h |
| HORVU.MOREX.r3.4HG0388310.1 | -0.459123738 | 3.870759069 | -3.901117138 | 0.00027 | 0.021949918 | -0.09417258 | Protein kinase | elongation zone | 6 h |
| HORVU.MOREX.r3.6HG0622740.1 | -0.566048291 | 4.313476147 | -3.918745153 | 0.00026 | 0.021417658 | -0.09633858 | Protein DETOXIFICATION | elongation zone | 6 h |
| HORVU.MOREX.r3.2HG0152530.1 | -0.458855554 | 4.851529635 | -3.912345687 | 0.00026 | 0.021709616 | -0.09671932 | Fasciclin-like arabinogalactan protein | elongation zone | 6 h |
| HORVU.MOREX.r3.1HG0027200.1 | -0.971650424 | 4.502879298 | -3.858019392 | 0.00031 | 0.023872158 | -0.1073209 | Auxin-responsive protein | elongation zone | 6 h |
| HORVU.MOREX.r3.3HG0248480.1 | -1.700583687 | -1.004527296 | -3.743242814 | 0.00045 | 0.028496623 | -0.12054271 | Cytochrome P450 | elongation zone | 6 h |
| HORVU.MOREX.r3.2HG0170640.1 | 2.802997303 | -0.509068665 | 3.810867568 | 0.00036 | 0.025722162 | -0.12998571 | Protein FAR1-RELATED SEQUENCE 5 | elongation zone | 6 h |
| HORVU.MOREX.r3.7HG0640410.1 | -2.963594707 | -2.747397168 | -3.77009813 | 0.00041 | 0.027115872 | -0.13863515 | Cytochrome P450 | elongation zone | 6 h |
| HORVU.MOREX.r3.5HG0488050.1 | -1.401771625 | -2.926199569 | -3.7255621 | 0.00047 | 0.029390657 | -0.14268426 | Blue copper protein | elongation zone | 6 h |
| HORVU.MOREX.r3.7HG0641230.1 | -1.297823001 | -1.229188244 | -3.823029539 | 0.00035 | 0.025184263 | -0.15373962 | Retrovirus-related Pol polyprotein from transposon TNT 1-94 | elongation zone | 6 h |
| HORVU.MOREX.r3.1HG0053060.1 | -1.032894535 | 4.011377898 | -3.829500534 | 0.00034 | 0.024875194 | -0.1577276 | PLATZ transcription factor | elongation zone | 6 h |
| HORVU.MOREX.r3.2HG0165250.1 | 0.452839604 | 6.077437333 | 3.908426596 | 0.00027 | 0.021784575 | -0.17968031 | OJ000315_02.19 protein | elongation zone | 6 h |
| HORVU.MOREX.r3.3HG0254950.1 | -1.160163954 | 1.019759768 | -3.890488091 | 0.00028 | 0.022218304 | -0.17975102 | Peroxidase | elongation zone | 6 h |
| HORVU.MOREX.r3.5HG0459320.1 | 0.587882775 | 4.812268395 | 3.839517452 | 0.00033 | 0.02468067 | -0.19106983 | Peptidylprolyl isomerase | elongation zone | 6 h |
| HORVU.MOREX.r3.3HG0248540.2 | -0.547608481 | 3.626501132 | -3.789221256 | 0.00039 | 0.026638848 | -0.19375322 | Peroxisomal membrane 22 kDa (Mpv17/PMP22) family protein | elongation zone | 6 h |
| HORVU.MOREX.r3.4HG0333450.1 | -0.638331526 | 3.618972788 | -3.797819768 | 0.00038 | 0.026284234 | -0.20059046 | Type I inositol-1,4,5-trisphosphate 5-phosphatase CVP2 | elongation zone | 6 h |
| HORVU.MOREX.r3.7HG0639300.1 | 0.626346912 | 3.626615674 | 3.7787741 | 0.0004 | 0.026862598 | -0.20692234 | F-box protein | elongation zone | 6 h |
| HORVU.MOREX.r3.5HG0511090.1 | -0.638230452 | 5.583411147 | -3.85994922 | 0.00031 | 0.023818477 | -0.20702057 | Phytochrome | elongation zone | 6 h |
| HORVU.MOREX.r3.3HG0278170.1 | -1.651193094 | 0.346838504 | -3.705523556 | 0.0005 | 0.030363091 | -0.21208383 | CCCH-type zinc fingerfamily protein with RNA-binding domain-co | elongation zone | 6 h |
| HORVU.MOREX.r3.2HG0140700.1 | 0.347921757 | 5.904167686 | 3.890032383 | 0.00028 | 0.022218304 | -0.21274615 | #N/A | elongation zone | 6 h |
| HORVU.MOREX.r3.4HG0354010.1 | -2.332775953 | -0.997515384 | -3.696415728 | 0.00052 | 0.030363091 | -0.2142125 | LURP-one-like protein | elongation zone | 6 h |
| HORVU.MOREX.r3.4HG0379290.1 | 0.349308761 | 5.287980079 | 3.851134773 | 0.00032 | 0.02415331 | -0.23622023 | COP1-interacting-like protein, putative | elongation zone | 6 h |
| HORVU.MOREX.r3.5HG0470030.1 | 0.514512176 | 7.999201135 | 3.967759405 | 0.00022 | 0.019648199 | -0.23766636 | 3-ketoacyl-CoA thiolase | elongation zone | 6 h |
| HORVU.MOREX.r3.5HG0430050.1 | -0.664882426 | 4.495639375 | -3.828167793 | 0.00034 | 0.024875194 | -0.23920045 | Stomatal closure-related actin-binding protein 1 | elongation zone | 6 h |
| HORVU.MOREX.r3.3HG0269810.1 | 0.693680639 | 4.116724617 | 3.798381617 | 0.00038 | 0.026284234 | -0.24160016 | Aspartyl protease family protein | elongation zone | 6 h |
| HORVU.MOREX.r3.6HG0558430.1 | -2.918678757 | -1.969157859 | -3.698319925 | 0.00052 | 0.030363091 | -0.26003925 | SAUR-like auxin-responsive protein family, putative | elongation zone | 6 h |
| HORVU.MOREX.r3.1HG0056620.1 | -1.589470634 | 0.189161761 | -3.892462406 | 0.00028 | 0.022217049 | -0.26192557 | Expansin protein | elongation zone | 6 h |
| HORVU.MOREX.r3.4HG0405920.1 | -1.252631866 | 0.085137287 | -3.828240129 | 0.00034 | 0.024875194 | -0.2652035 | Hexosyltransferase | elongation zone | 6 h |
| HORVU.MOREX.r3.4HG0387180.1 | -0.933286469 | 1.821336783 | -3.682540464 | 0.00054 | 0.030744687 | -0.26576925 | Proton pump-interactor 1 | elongation zone | 6 h |
| HORVU.MOREX.r3.4HG0391330.1 | 0.43530433 | 6.457455724 | 3.885345551 | 0.00029 | 0.02238184 | -0.27199816 | Phosphomevalonate kinase | elongation zone | 6 h |
| HORVU.MOREX.r3.2HG0126550.1 | -1.618153587 | 1.396372437 | -3.678957948 | 0.00055 | 0.031004434 | -0.27281395 | Alpha/beta hydrolase, putative | elongation zone | 6 h |
| HORVU.MOREX.r3.3HG0264440.1 | 0.748483772 | 2.161098312 | 3.691094948 | 0.00053 | 0.030429659 | -0.27487653 | Condensin complex subunit 2 | elongation zone | 6 h |

**Table S5** Continued.

| ID | logFC | AveExpr | t | P.Value | adj.P.Val | B | gene description | root zone | time point |
|---|---|---|---|---|---|---|---|---|---|
| HORVU.MOREX.r3.4HG0343050.1 | -0.56982995 | 6.230075431 | -3.894244508 | 0.00028 | 0.022216127 | -0.27831619 | N-acetyl-gamma-glutamyl-phosphate reductase | elongation zone | 6 h |
| HORVU.MOREX.r3.4HG0335450.1 | -2.457946646 | -2.959169593 | -3.692737263 | 0.00053 | 0.03036885 | -0.27885736 | Nucleobase ascorbate transporter | elongation zone | 6 h |
| HORVU.MOREX.r3.5HG0420480.1 | -2.100792208 | 1.423524837 | -3.693701107 | 0.00052 | 0.030363091 | -0.29138024 | BEL1-like homeodomain protein 6 | elongation zone | 6 h |
| HORVU.MOREX.r3.2HG0195690.1 | -1.08462304 | -2.810433048 | -3.666669525 | 0.00057 | 0.031690477 | -0.29505302 | filamentous hemagglutinin transporter | elongation zone | 6 h |
| HORVU.MOREX.r3.1HG0080180.1 | -1.339686867 | -2.352460779 | -3.687901429 | 0.00053 | 0.030483629 | -0.30078029 | Jacalin lectin family protein | elongation zone | 6 h |
| HORVU.MOREX.r3.7HG0666740.1 | -1.042417761 | 1.938696651 | -3.773454641 | 0.00041 | 0.027115872 | -0.30648019 | Glycosyltransferase | elongation zone | 6 h |
| HORVU.MOREX.r3.6HG0628790.1 | -0.927189607 | 2.686977072 | -3.694667111 | 0.00052 | 0.030363091 | -0.32305914 | Myb-like transcription factor family protein | elongation zone | 6 h |
| HORVU.MOREX.r3.2HG0112690.1 | -0.952480116 | 1.247172627 | -3.894301923 | 0.00028 | 0.022216127 | -0.32372335 | Peroxidase | elongation zone | 6 h |
| HORVU.MOREX.r3.2HG0195020.1 | 1.090477608 | 2.396301121 | 3.65352303 | 0.00059 | 0.032312416 | -0.33052753 | tRNA (Guanine-N(7)-)-methyltransferase | elongation zone | 6 h |
| HORVU.MOREX.r3.3HG0307630.1 | 2.051403981 | 2.923305984 | 3.665506728 | 0.00057 | 0.031719413 | -0.33099415 | SET-domain protein lysine methyltransferase family protein | elongation zone | 6 h |
| HORVU.MOREX.r3.3HG0284780.1 | -0.427976595 | 5.323435331 | -3.828892723 | 0.00034 | 0.024875194 | -0.33302062 | Protein PLASTID MOVEMENT IMPAIRED 1-RELATED 1 | elongation zone | 6 h |
| HORVU.MOREX.r3.3HG0303380.1 | -0.900310102 | -0.867377041 | -3.723517822 | 0.00048 | 0.029402036 | -0.33883151 | Zinc finger protein CONSTANS | elongation zone | 6 h |
| HORVU.MOREX.r3.1HG0094110.1 | -1.723953107 | 0.561026983 | -3.734174008 | 0.00046 | 0.029138607 | -0.34109787 | Sigma factor sigB regulation protein rsbQ | elongation zone | 6 h |
| HORVU.MOREX.r3.4HG0413690.1 | 0.538253757 | 4.40336351 | 3.769966532 | 0.00041 | 0.027115872 | -0.34376599 | 26S proteasome non-ATPase regulatory subunit-like protein | elongation zone | 6 h |
| HORVU.MOREX.r3.7HG0716900.1 | 0.384875417 | 8.192568258 | 3.918867731 | 0.00026 | 0.021417658 | -0.35974345 | Transmembrane 9 superfamily member | elongation zone | 6 h |
| HORVU.MOREX.r3.3HG0234000.1 | -1.578710628 | -0.274430722 | -3.837001274 | 0.00033 | 0.024767988 | -0.35991519 | Xyloglucan endotransglucosylase/hydrolase | elongation zone | 6 h |
| HORVU.MOREX.r3.7HG0712850.1 | 0.414444229 | 4.698390518 | 3.783720401 | 0.00039 | 0.026712078 | -0.36713344 | Fiber protein Fb34 | elongation zone | 6 h |
| HORVU.MOREX.r3.6HG0614970.1 | -0.422492877 | 5.579922669 | -3.836128908 | 0.00033 | 0.024767988 | -0.36936213 | Kelch repeat-containing protein | elongation zone | 6 h |
| HORVU.MOREX.r3.3HG0248280.1 | 0.403275481 | 8.506150471 | 3.928980425 | 0.00025 | 0.021333038 | -0.37002643 | Reticulon-like protein | elongation zone | 6 h |
| HORVU.MOREX.r3.3HG0287000.1 | -0.944549102 | 3.76861142 | -3.778059237 | 0.0004 | 0.026862598 | -0.37571845 | Respiratory burst oxidase | elongation zone | 6 h |
| HORVU.MOREX.r3.2HG0165110.1 | -0.983853664 | 3.128473967 | -3.745545292 | 0.00045 | 0.028496623 | -0.37936024 | Acetyl-coenzyme A synthetase | elongation zone | 6 h |
| HORVU.MOREX.r3.2HG0179560.1 | -0.949413077 | 1.523229319 | -3.764365667 | 0.00042 | 0.027510832 | -0.38486509 | Nodulin-like / Major Facilitator Superfamily protein | elongation zone | 6 h |
| HORVU.MOREX.r3.5HG0518640.1 | 1.156564549 | 2.499571845 | 3.658649333 | 0.00058 | 0.031886221 | -0.39063303 | Methionine S-methyltransferase | elongation zone | 6 h |
| HORVU.MOREX.r3.2HG0216110.1 | 0.347778795 | 5.350102781 | 3.798219655 | 0.00038 | 0.026284234 | -0.39295764 | #N/A | elongation zone | 6 h |
| HORVU.MOREX.r3.4HG0354140.1 | -1.358408329 | -2.197267006 | -3.627412003 | 0.00064 | 0.033489087 | -0.40152952 | CheY-like two-component responsive regulator family protein | elongation zone | 6 h |
| HORVU.MOREX.r3.2HG0189800.1 | 0.706385586 | 3.770834829 | 3.732655821 | 0.00046 | 0.029138607 | -0.40488431 | ethylene-responsive transcription factor | elongation zone | 6 h |
| HORVU.MOREX.r3.5HG0471070.1 | -0.288264241 | 6.510820563 | -3.847725377 | 0.00032 | 0.024250025 | -0.41521631 | Oxidoreductase family, NAD-binding rossmann fold protein | elongation zone | 6 h |
| HORVU.MOREX.r3.2HG0133220.1 | -0.852435556 | 3.407204844 | -3.831540632 | 0.00034 | 0.024875194 | -0.41732151 | Blue copper protein | elongation zone | 6 h |
| HORVU.MOREX.r3.7HG0728060.1 | -0.550522633 | 5.512336647 | -3.79689628 | 0.00038 | 0.026284234 | -0.4262837 | Aminodeoxychorismate synthase | elongation zone | 6 h |
| HORVU.MOREX.r3.2HG0182150.1 | 0.819024346 | 4.062913174 | 3.694669058 | 0.00052 | 0.030363091 | -0.4300998 | Charged multivesicular body protein 7 | elongation zone | 6 h |
| HORVU.MOREX.r3.7HG0679350.1 | 0.598814537 | 3.668407943 | 3.797777138 | 0.00038 | 0.026284234 | -0.43805717 | Leucine-rich repeat receptor-like protein kinase family | elongation zone | 6 h |
| HORVU.MOREX.r3.2HG0113590.1 | 0.496439718 | 3.8003454 | 3.728199413 | 0.00047 | 0.029236824 | -0.44983752 | Zinc finger CCCH domain protein, putative | elongation zone | 6 h |
| HORVU.MOREX.r3.3HG0245120.1 | -3.163524487 | -2.694279726 | -3.643811494 | 0.00061 | 0.032525759 | -0.45584644 | Cysteine protease, putative | elongation zone | 6 h |
| HORVU.MOREX.r3.3HG0294740.1 | 0.963770586 | 3.043041266 | 3.643440809 | 0.00061 | 0.032525759 | -0.45918477 | HTH-type transcriptional activator RhaR | elongation zone | 6 h |
| HORVU.MOREX.r3.5HG0500680.1 | 0.717233715 | 6.785876601 | 3.819409787 | 0.00035 | 0.025297158 | -0.4591963 | Acyl carrier protein | elongation zone | 6 h |
| HORVU.MOREX.r3.2HG0114290.1 | 0.342421226 | 6.856362561 | 3.820931096 | 0.00035 | 0.025263639 | -0.45995938 | #N/A | elongation zone | 6 h |
| HORVU.MOREX.r3.3HG0316280.1 | 0.954495209 | 4.501603233 | 3.696277107 | 0.00052 | 0.030363091 | -0.46557911 | Transcription initiation factor TFIID subunit 9 | elongation zone | 6 h |
| HORVU.MOREX.r3.5HG0504520.1 | -1.196472882 | 0.68442194 | -3.664415445 | 0.00057 | 0.031741526 | -0.4688817 | Phytoene synthase | elongation zone | 6 h |
| HORVU.MOREX.r3.7HG0723060.1 | -2.459985507 | -1.228512714 | -3.659185966 | 0.00058 | 0.031886221 | -0.47135088 | 4-hydroxy-3-methylbut-2-enyl diphosphate synthase | elongation zone | 6 h |
| HORVU.MOREX.r3.6HG0545620.1 | 0.388113795 | 11.02319041 | 3.992994465 | 0.0002 | 0.018702211 | -0.48035237 | Heat shock 70 kDa protein | elongation zone | 6 h |
| HORVU.MOREX.r3.4HG0415130.1 | -1.642907647 | -1.494103874 | -3.630040661 | 0.00064 | 0.033339257 | -0.49324883 | rRNA N-glycosidase | elongation zone | 6 h |

**Table S5** Continued.

| ID | logFC | AveExpr | t | P.Value | adj.P.Val | B | gene description | root zone | time point |
|---|---|---|---|---|---|---|---|---|---|
| HORVU.MOREX.r3.5HG0491230.1 | 1.006784728 | 2.284227707 | 3.61856007 | 0.00066 | 0.033752681 | -0.49364031 | Dihydroflavonol-4-reductase | elongation zone | 6 h |
| HORVU.MOREX.r3.5HG0474540.1 | 0.39529092 | 7.609555508 | 3.856902765 | 0.00031 | 0.023872158 | -0.49561125 | Protein transport protein Sec61 subunit alpha | elongation zone | 6 h |
| HORVU.MOREX.r3.7HG0736960.1 | 0.718182229 | 3.878659947 | 3.669247569 | 0.00057 | 0.031591904 | -0.4973797 | BTB/POZ domain containing protein, expressed | elongation zone | 6 h |
| HORVU.MOREX.r3.7HG0748670.1 | -2.591030527 | -1.854958195 | -3.58887907 | 0.00073 | 0.0350911 | -0.50467662 | GDSL esterase/lipase | elongation zone | 6 h |
| HORVU.MOREX.r3.1HG0076980.1 | 0.373230846 | 8.500485602 | 3.883703439 | 0.00029 | 0.022414154 | -0.50539665 | Dolichyl-diphosphooligosaccharide--protein glycosyltransferase s | elongation zone | 6 h |
| HORVU.MOREX.r3.3HG0307240.1 | -2.09381985 | -3.376882233 | -3.603992908 | 0.00069 | 0.033986772 | -0.50835229 | Amino acid permease | elongation zone | 6 h |
| HORVU.MOREX.r3.5HG0490760.1 | 0.632810261 | 4.107029811 | 3.690297667 | 0.00053 | 0.030429659 | -0.50896684 | Glucosamine 6-phosphate N-acetyltransferase | elongation zone | 6 h |
| HORVU.MOREX.r3.3HG0283300.1 | 0.722989541 | 2.473566507 | 3.626440182 | 0.00065 | 0.033489087 | -0.50896878 | Farnesyl diphosphate synthase | elongation zone | 6 h |
| HORVU.MOREX.r3.3HG0235030.1 | 0.532422271 | 3.06592017 | 3.643892372 | 0.00061 | 0.032525759 | -0.51204366 | Dehydration-responsive element binding protein | elongation zone | 6 h |
| HORVU.MOREX.r3.5HG0428840.1 | 0.728581537 | 5.358223693 | 3.719947148 | 0.00048 | 0.029555507 | -0.51595812 | BTB/POZ domain-containing protein | elongation zone | 6 h |
| HORVU.MOREX.r3.7HG0664720.1 | -1.177477733 | 0.421259005 | -3.788989372 | 0.00039 | 0.026638848 | -0.51793575 | Blue copper protein | elongation zone | 6 h |
| HORVU.MOREX.r3.1HG0000050.1 | 0.441729568 | 5.048249047 | 3.744159449 | 0.00045 | 0.028496623 | -0.53205209 | RING-finger ubiquitin ligase | elongation zone | 6 h |
| HORVU.MOREX.r3.3HG0321700.1 | -0.63411323 | 3.776225864 | -3.68628843 | 0.00054 | 0.030483629 | -0.53690083 | FMN-dependent NADPH-azoreductase | elongation zone | 6 h |
| HORVU.MOREX.r3.4HG0348800.1 | -0.971944539 | 2.044228231 | -3.644323952 | 0.00061 | 0.032525759 | -0.53751171 | TCP transcription factor | elongation zone | 6 h |
| HORVU.MOREX.r3.4HG0364210.1 | -2.68297036 | -2.342478207 | -3.731323783 | 0.00047 | 0.029138607 | -0.54538209 | #N/A | elongation zone | 6 h |
| HORVU.MOREX.r3.2HG0156390.1 | 0.327521307 | 6.734888733 | 3.795421762 | 0.00038 | 0.026284234 | -0.54741284 | Protein kinase | elongation zone | 6 h |
| HORVU.MOREX.r3.6HG0560140.1 | -2.494514576 | -1.916702632 | -3.573884082 | 0.00076 | 0.035856081 | -0.54896686 | Endoglucanase | elongation zone | 6 h |
| HORVU.MOREX.r3.1HG0000030.1 | 1.360598273 | 0.806284125 | 3.56865402 | 0.00077 | 0.035934872 | -0.56838418 | Serine/threonine-protein kinase ATM | elongation zone | 6 h |
| HORVU.MOREX.r3.1HG0024040.1 | 0.369160348 | 9.734967723 | 3.910725207 | 0.00026 | 0.02171138 | -0.57237238 | Disulfide-isomerase | elongation zone | 6 h |
| HORVU.MOREX.r3.2HG0204820.1 | -0.76047504 | 3.904121549 | -3.706778099 | 0.0005 | 0.030363091 | -0.5804904 | Ethylene-responsive transcription factor | elongation zone | 6 h |
| HORVU.MOREX.r3.5HG0465260.1 | 0.74295925 | 2.270828878 | 3.572646327 | 0.00076 | 0.035872795 | -0.58904827 | Zinc finger HIT domain-containing protein 1 | elongation zone | 6 h |
| HORVU.MOREX.r3.6HG0570230.1 | -1.000381402 | 0.733369112 | -3.558218256 | 0.0008 | 0.035982407 | -0.61064421 | Aldose 1-epimerase family protein | elongation zone | 6 h |
| HORVU.MOREX.r3.2HG0140170.1 | -0.68427086 | 2.981208713 | -3.638962134 | 0.00062 | 0.032755332 | -0.61886727 | Ras-related protein Rab-21 | elongation zone | 6 h |
| HORVU.MOREX.r3.6HG0607590.1 | -1.193495013 | 1.682477401 | -3.567888833 | 0.00077 | 0.035934872 | -0.62609016 | U-box domain-containing family protein | elongation zone | 6 h |
| HORVU.MOREX.r3.7HG0731370.1 | 2.689394922 | 0.369074602 | 3.703249293 | 0.00051 | 0.030363091 | -0.63027933 | #N/A | elongation zone | 6 h |
| HORVU.MOREX.r3.6HG0625340.1 | 1.048708262 | 1.518599241 | 3.542127047 | 0.00084 | 0.036935434 | -0.64307007 | GPI transamidase component PIG-T | elongation zone | 6 h |
| HORVU.MOREX.r3.4HG0397940.1 | 0.793378071 | 3.598811101 | 3.61197521 | 0.00068 | 0.033870225 | -0.64427415 | Alpha-L-fucosidase 2 | elongation zone | 6 h |
| HORVU.MOREX.r3.2HG0170120.1 | -0.840175754 | 2.445884193 | -3.596547714 | 0.00071 | 0.034460123 | -0.65222647 | Protein NRT1/ PTR FAMILY 1.2 | elongation zone | 6 h |
| HORVU.MOREX.r3.7HG0674250.1 | 0.889694467 | 0.727477133 | 3.606543226 | 0.00069 | 0.033986772 | -0.6548979 | hydroxysteroid dehydrogenase 3 | elongation zone | 6 h |
| HORVU.MOREX.r3.5HG0536200.1 | -1.056301773 | -0.55460225 | -3.69671955 | 0.00052 | 0.030363091 | -0.65989239 | Xyloglucan endotransglucosylase/hydrolase | elongation zone | 6 h |
| HORVU.MOREX.r3.5HG0472770.1 | -0.692807154 | 4.33132919 | -3.67525642 | 0.00055 | 0.031192351 | -0.66400056 | SIN3-like 2 | elongation zone | 6 h |
| HORVU.MOREX.r3.4HG0402000.1 | 0.684436036 | 8.180685451 | 3.813120715 | 0.00036 | 0.02562773 | -0.6643136 | Kinase interacting (KIP1-like) family protein, putative | elongation zone | 6 h |
| HORVU.MOREX.r3.4HG0351790.1 | -1.686942525 | 0.588108997 | -3.532894505 | 0.00086 | 0.037429607 | -0.66442234 | Calcium-activated potassium channel subunit alpha-1 | elongation zone | 6 h |
| HORVU.MOREX.r3.2HG0203080.1 | -2.08098082 | 0.039802538 | -3.527511402 | 0.00088 | 0.037809868 | -0.66675358 | NA | elongation zone | 6 h |
| HORVU.MOREX.r3.3HG0296140.1 | -1.842794583 | 2.111797459 | -3.540115325 | 0.00084 | 0.036941016 | -0.67336962 | F-box protein | elongation zone | 6 h |
| HORVU.MOREX.r3.4HG0383070.1 | 0.703084535 | 5.44858875 | 3.696120929 | 0.00052 | 0.030363091 | -0.6740717 | Calmodulin-binding protein | elongation zone | 6 h |
| HORVU.MOREX.r3.2HG0192000.1 | -0.928972045 | 0.357063756 | -3.699565237 | 0.00051 | 0.030363091 | -0.67463474 | HXXXD-type acyl-transferase family protein, putative | elongation zone | 6 h |
| HORVU.MOREX.r3.3HG0222660.1 | 1.45576785 | 1.364356123 | 3.525669448 | 0.00088 | 0.037863675 | -0.67573833 | F-box family protein | elongation zone | 6 h |
| HORVU.MOREX.r3.2HG0178990.1 | 0.583066437 | 4.094270236 | 3.611790956 | 0.00068 | 0.033870225 | -0.67730136 | DNA-3-methyladenine glycosylase, putative | elongation zone | 6 h |
| HORVU.MOREX.r3.7HG0677270.1 | 0.958468865 | 2.721723616 | 3.560056717 | 0.00079 | 0.035972648 | -0.6781955 | Glycosyltransferase | elongation zone | 6 h |
| HORVU.MOREX.r3.5HG0513980.1 | -1.294486606 | -0.364729073 | -3.568674049 | 0.00077 | 0.035934872 | -0.68041604 | Auxin-responsive protein | elongation zone | 6 h |

**Table S5** Continued.

| ID | logFC | AveExpr | t | P.Value | adj.P.Val | B | gene description | root zone | time point |
|---|---|---|---|---|---|---|---|---|---|
| HORVU.MOREX.r3.2HG0165840.1 | 0.634056901 | 6.204491334 | 3.696444482 | 0.00052 | 0.030363091 | -0.69413382 | Protein kinase | elongation zone | 6 h |
| HORVU.MOREX.r3.5HG0480410.1 | -1.051496101 | 3.257253448 | -3.624739494 | 0.00065 | 0.033489087 | -0.69619772 | Copper transporter | elongation zone | 6 h |
| HORVU.MOREX.r3.1HG0082240.1 | 0.366360194 | 5.659889983 | 3.71771623 | 0.00049 | 0.029586232 | -0.6977039 | ER lumen protein-retaining receptor | elongation zone | 6 h |
| HORVU.MOREX.r3.7HG0731320.1 | 0.374477594 | 5.022738423 | 3.70762296 | 0.0005 | 0.030363091 | -0.69843493 | O-fucosyltransferase family protein | elongation zone | 6 h |
| HORVU.MOREX.r3.3HG0323600.1 | -0.583734233 | 4.35559343 | -3.700742657 | 0.00051 | 0.030363091 | -0.70277397 | Phosphatidylserine decarboxylase proenzyme 2 | elongation zone | 6 h |
| HORVU.MOREX.r3.6HG0575530.1 | -0.566987084 | 6.131400028 | -3.743191102 | 0.00045 | 0.028496623 | -0.70369773 | Alcohol dehydrogenase, putative | elongation zone | 6 h |
| HORVU.MOREX.r3.6HG0601260.1 | -1.051180842 | 3.05036359 | -3.721457477 | 0.00048 | 0.029503984 | -0.71122968 | Sulfotransferase | elongation zone | 6 h |
| HORVU.MOREX.r3.5HG0513810.1 | -0.615029302 | 4.245645455 | -3.686160405 | 0.00054 | 0.030483629 | -0.71724418 | Malate dehydrogenase | elongation zone | 6 h |
| HORVU.MOREX.r3.3HG0282320.1 | 2.080196058 | -0.605168808 | 3.61112935 | 0.00068 | 0.033870225 | -0.71994872 | Fatty acid desaturase 1 | elongation zone | 6 h |
| HORVU.MOREX.r3.7HG0735840.1 | 0.946956833 | 4.848168125 | 3.630224652 | 0.00064 | 0.033339257 | -0.72108942 | serine/arginine repetitive matrix-like protein | elongation zone | 6 h |
| HORVU.MOREX.r3.2HG0175420.1 | 0.358964588 | 6.198454241 | 3.754375615 | 0.00043 | 0.028032118 | -0.73036186 | Leucine-rich repeat (LRR) family protein | elongation zone | 6 h |
| HORVU.MOREX.r3.2HG0108450.1 | -0.549824363 | 5.259081061 | -3.731404575 | 0.00047 | 0.029138607 | -0.73210624 | Protein kinase family protein, putative, expressed | elongation zone | 6 h |
| HORVU.MOREX.r3.5HG0443920.1 | 0.522928884 | 3.840341821 | 3.61670286 | 0.00067 | 0.033798115 | -0.73308867 | Calmodulin-binding protein, putative, expressed | elongation zone | 6 h |
| HORVU.MOREX.r3.2HG0176960.1 | 0.363367851 | 4.988465286 | 3.674100112 | 0.00056 | 0.031219405 | -0.73550279 | Kelch repeat-containing protein | elongation zone | 6 h |
| HORVU.MOREX.r3.2HG0171780.1 | -1.701302517 | -1.262454147 | -3.500850504 | 0.00095 | 0.039256264 | -0.74702175 | transcription repressor | elongation zone | 6 h |
| HORVU.MOREX.r3.4HG0332930.1 | 1.522278035 | 0.588837805 | 3.545192731 | 0.00083 | 0.036748557 | -0.74950134 | Short-chain dehydrogenase/reductase family protein | elongation zone | 6 h |
| HORVU.MOREX.r3.7HG0731560.1 | -1.405468965 | -0.874013134 | -3.49580604 | 0.00096 | 0.039433228 | -0.75032113 | Regulator of Vps4 activity in the MVB pathway protein, putative | elongation zone | 6 h |
| HORVU.MOREX.r3.2HG0119030.1 | -0.698232704 | 3.539432921 | -3.571709606 | 0.00076 | 0.035893875 | -0.75266859 | Protein phosphatase 2c, putative | elongation zone | 6 h |
| HORVU.MOREX.r3.6HG0609720.1 | 0.682038548 | 3.853973881 | 3.579837988 | 0.00075 | 0.035624994 | -0.76587547 | Glucosamine 6-phosphate N-acetyltransferase | elongation zone | 6 h |
| HORVU.MOREX.r3.6HG0614830.1 | 0.30840065 | 6.98068667 | 3.761610612 | 0.00042 | 0.027662361 | -0.77935148 | Exocyst complex component SEC6 | elongation zone | 6 h |
| HORVU.MOREX.r3.5HG0441420.1 | -2.585222576 | -0.428611158 | -3.603872772 | 0.00069 | 0.033986772 | -0.78230944 | Ankyrin repeat-containing protein, putative | elongation zone | 6 h |
| HORVU.MOREX.r3.6HG0610500.1 | -0.957271298 | 1.298618032 | -3.730238866 | 0.00047 | 0.029138607 | -0.79337631 | Eukaryotic aspartyl protease family protein | elongation zone | 6 h |
| HORVU.MOREX.r3.7HG0663940.1 | -0.900157677 | 0.764913385 | -3.647889889 | 0.0006 | 0.032525759 | -0.79541432 | Alpha/beta-Hydrolases superfamily protein, putative | elongation zone | 6 h |
| HORVU.MOREX.r3.5HG0487070.1 | -0.516555601 | 4.654929378 | -3.659515179 | 0.00058 | 0.031886221 | -0.80031021 | Alpha/beta-Hydrolases superfamily protein, putative | elongation zone | 6 h |
| HORVU.MOREX.r3.6HG0631710.1 | -0.959376174 | 2.878977233 | -3.783375877 | 0.0004 | 0.026712078 | -0.80057324 | Aquaporin | elongation zone | 6 h |
| HORVU.MOREX.r3.1HG0062110.1 | -1.259960551 | 0.941666543 | -3.474226753 | 0.00103 | 0.040615136 | -0.80969219 | Pentatricopeptide repeat-containing protein | elongation zone | 6 h |
| HORVU.MOREX.r3.5HG0477750.1 | -1.444196348 | 0.65835179 | -3.515556988 | 0.00091 | 0.038564824 | -0.81277538 | Homeobox protein, putative | elongation zone | 6 h |
| HORVU.MOREX.r3.4HG0363330.1 | 0.910417121 | 2.648390841 | 3.528949615 | 0.00087 | 0.037723585 | -0.81637404 | F-box/RNI-like superfamily protein | elongation zone | 6 h |
| HORVU.MOREX.r3.7HG0742080.1 | -1.06199179 | -1.326097459 | -3.561850966 | 0.00079 | 0.035934872 | -0.8261079 | Receptor-like kinase | elongation zone | 6 h |
| HORVU.MOREX.r3.1HG0086480.1 | -1.447292219 | 0.131690052 | -3.480074337 | 0.00101 | 0.040367299 | -0.82923389 | UV-B-induced protein, chloroplastic | elongation zone | 6 h |
| HORVU.MOREX.r3.4HG0378450.1 | -1.291613682 | 0.244687178 | -3.465269268 | 0.00106 | 0.040858317 | -0.82966076 | #N/A | elongation zone | 6 h |
| HORVU.MOREX.r3.7HG0674320.1 | 0.402260818 | 4.709886156 | 3.658876055 | 0.00058 | 0.031886221 | -0.83327215 | Vesicle transport protein | elongation zone | 6 h |
| HORVU.MOREX.r3.7HG0666340.1 | 0.537842407 | 4.728875506 | 3.613710367 | 0.00067 | 0.033870225 | -0.83648536 | Hexosyltransferase | elongation zone | 6 h |
| HORVU.MOREX.r3.5HG0462450.1 | -1.394727884 | -0.846663356 | -3.461772173 | 0.00107 | 0.040858317 | -0.83838501 | S-acyltransferase | elongation zone | 6 h |
| HORVU.MOREX.r3.7HG0641890.1 | -3.630607398 | -3.164564754 | -3.499499842 | 0.00095 | 0.039256264 | -0.83933149 | Aspartic proteinase nepenthesin-2 | elongation zone | 6 h |
| HORVU.MOREX.r3.7HG0665990.1 | -0.652882016 | 6.15668184 | -3.756723745 | 0.00043 | 0.027913688 | -0.84302483 | Shikimate kinase | elongation zone | 6 h |
| HORVU.MOREX.r3.2HG0120830.1 | -1.628373815 | -3.157076886 | -3.465788933 | 0.00106 | 0.040858317 | -0.846427 | Peroxidase | elongation zone | 6 h |
| HORVU.MOREX.r3.2HG0096710.1 | -1.393826839 | -1.753677392 | -3.49030699 | 0.00098 | 0.039681075 | -0.84744167 | Cytochrome P450 | elongation zone | 6 h |
| HORVU.MOREX.r3.2HG0158180.1 | -1.185681008 | 3.620949743 | -3.565858465 | 0.00078 | 0.035934872 | -0.84890612 | Proline transporter | elongation zone | 6 h |
| HORVU.MOREX.r3.7HG0714660.1 | -1.422186979 | -2.488173436 | -3.456992539 | 0.00108 | 0.040886579 | -0.85015479 | Retrovirus-related Pol polyprotein from transposon TNT 1-94 | elongation zone | 6 h |
| HORVU.MOREX.r3.5HG0473030.1 | -1.059884994 | 0.923732427 | -3.457801879 | 0.00108 | 0.040886579 | -0.85267083 | 2-aminoethanethiol dioxygenase | elongation zone | 6 h |

**Table S5** Continued.

| ID | logFC | AveExpr | t | P.Value | adj.P.Val | B | gene description | root zone | time point |
|---|---|---|---|---|---|---|---|---|---|
| HORVU.MOREX.r3.2HG0187450.1 | -1.304715428 | 0.03595626 | -3.648391175 | 0.0006 | 0.032525759 | -0.85559221 | Rhomboid-like protein | elongation zone | 6 h |
| HORVU.MOREX.r3.6HG0544290.1 | -1.268733394 | -0.109959598 | -3.538990002 | 0.00085 | 0.036941016 | -0.85561716 | Glycerol-3-phosphate acyltransferase | elongation zone | 6 h |
| HORVU.MOREX.r3.4HG0395540.1 | -1.166388737 | 4.838553225 | -3.717960741 | 0.00049 | 0.029586232 | -0.85807641 | Glutamine synthetase | elongation zone | 6 h |
| HORVU.MOREX.r3.5HG0493850.1 | 0.498347115 | 7.419096029 | 3.72418554 | 0.00048 | 0.029402036 | -0.86904307 | Acyl-[acyl-carrier-protein] hydrolase | elongation zone | 6 h |
| HORVU.MOREX.r3.2HG0165300.1 | 0.904904533 | 2.849287283 | 3.495565925 | 0.00096 | 0.039433228 | -0.88626077 | UPF0496 protein | elongation zone | 6 h |
| HORVU.MOREX.r3.6HG0632320.1 | 0.418590435 | 5.523777089 | 3.642972055 | 0.00061 | 0.032525759 | -0.88952204 | Leucine-rich repeat protein | elongation zone | 6 h |
| HORVU.MOREX.r3.6HG0616850.1 | -3.754573666 | -2.90585403 | -3.511515923 | 0.00092 | 0.038757609 | -0.89250604 | Blue copper protein | elongation zone | 6 h |
| HORVU.MOREX.r3.3HG0285440.1 | -1.107104217 | -1.285046886 | -3.50492844 | 0.00094 | 0.039105108 | -0.89527843 | #N/A | elongation zone | 6 h |
| HORVU.MOREX.r3.3HG0241360.1 | -1.993906581 | -0.960116855 | -3.547700149 | 0.00082 | 0.036546446 | -0.90061759 | ATPase E1-E2 type family protein / haloacid dehalogenase-like hy | elongation zone | 6 h |
| HORVU.MOREX.r3.7HG0665040.1 | -2.217504527 | -3.452636637 | -3.479239655 | 0.00101 | 0.040391023 | -0.90215185 | Protein EXORDIUM-like 1 | elongation zone | 6 h |
| HORVU.MOREX.r3.1HG0073430.1 | -0.471739597 | 3.344558802 | -3.538581471 | 0.00085 | 0.036941016 | -0.91328672 | O-fucosyltransferase family protein | elongation zone | 6 h |
| HORVU.MOREX.r3.5HG0500590.1 | -0.994355921 | 1.729937267 | -3.459786133 | 0.00108 | 0.040858317 | -0.91421064 | Receptor-like kinase | elongation zone | 6 h |
| HORVU.MOREX.r3.7HG0706310.1 | -0.550087908 | 5.228140664 | -3.611910418 | 0.00068 | 0.033870225 | -0.91854111 | Alcohol dehydrogenase | elongation zone | 6 h |
| HORVU.MOREX.r3.2HG0168700.1 | 1.549193918 | 0.03517856 | 3.46325728 | 0.00106 | 0.040858317 | -0.92097701 | ADP-ribosylation factor GTPase-activating protein AGD4 | elongation zone | 6 h |
| HORVU.MOREX.r3.3HG0273790.2 | -0.81426197 | 2.302444592 | -3.465564406 | 0.00106 | 0.040858317 | -0.92147104 | CTD small phosphatase-like protein | elongation zone | 6 h |
| HORVU.MOREX.r3.4HG0344640.1 | 0.462989296 | 3.700508827 | 3.556327339 | 0.0008 | 0.035982407 | -0.92260367 | Glycerophosphodiester phosphodiesterase | elongation zone | 6 h |
| HORVU.MOREX.r3.7HG0726360.1 | -0.324310801 | 5.450154375 | -3.649467232 | 0.0006 | 0.032525759 | -0.92782625 | 12-oxophytodienoate reductase | elongation zone | 6 h |
| HORVU.MOREX.r3.5HG0469340.1 | -1.193268551 | 0.552227437 | -3.52567578 | 0.00088 | 0.037863675 | -0.93868516 | Fasciclin-like arabinogalactan-protein-like | elongation zone | 6 h |
| HORVU.MOREX.r3.2HG0190710.1 | 0.371257775 | 4.642344889 | 3.577699835 | 0.00075 | 0.035692712 | -0.94334853 | #N/A | elongation zone | 6 h |
| HORVU.MOREX.r3.7HG0720790.1 | 0.946175021 | 3.444490231 | 3.460103171 | 0.00107 | 0.040858317 | -0.94644246 | Myb family transcription factor family protein | elongation zone | 6 h |
| HORVU.MOREX.r3.2HG0189670.1 | -1.073772461 | 0.283223271 | -3.604750059 | 0.00069 | 0.033986772 | -0.95031813 | Aquaporin-like protein | elongation zone | 6 h |
| HORVU.MOREX.r3.7HG0714400.1 | -0.991274792 | 2.70239141 | -3.492481004 | 0.00097 | 0.039646685 | -0.95441574 | Auxin-responsive protein | elongation zone | 6 h |
| HORVU.MOREX.r3.7HG0721580.1 | -1.515038816 | -1.146006065 | -3.416245266 | 0.00123 | 0.043449664 | -0.95634402 | Ras-related protein Rab-25 | elongation zone | 6 h |
| HORVU.MOREX.r3.6HG0626020.1 | -1.231138459 | 1.680187154 | -3.687830054 | 0.00053 | 0.030483629 | -0.95645817 | Peroxidase | elongation zone | 6 h |
| HORVU.MOREX.r3.1HG0079870.1 | 0.934129884 | 5.830169089 | 3.643051603 | 0.00061 | 0.032525759 | -0.96063528 | Pectin lyase-like superfamily protein | elongation zone | 6 h |
| HORVU.MOREX.r3.1HG0087170.1 | 0.425257402 | 5.424188381 | 3.607381734 | 0.00068 | 0.033986772 | -0.96750056 | 3-deoxy-manno-octulosonate cytidylyltransferase | elongation zone | 6 h |
| HORVU.MOREX.r3.4HG0418460.1 | -0.51671108 | 1.75953737 | -3.625547656 | 0.00065 | 0.033489087 | -0.97573261 | Cysteine protease | elongation zone | 6 h |
| HORVU.MOREX.r3.2HG0173040.1 | -0.485780804 | 4.911334977 | -3.624509234 | 0.00065 | 0.033489087 | -0.97932127 | Loricrin-like protein | elongation zone | 6 h |
| HORVU.MOREX.r3.6HG0609210.1 | -1.26804156 | 1.030285608 | -3.407881803 | 0.00126 | 0.043818088 | -0.97942582 | RNA binding protein, putative | elongation zone | 6 h |
| HORVU.MOREX.r3.5HG0532120.1 | -1.623815914 | -1.755607472 | -3.484974076 | 0.001 | 0.039927046 | -0.98211399 | Expansin | elongation zone | 6 h |
| HORVU.MOREX.r3.5HG0524060.1 | -1.343682815 | 3.909293568 | -3.567210384 | 0.00078 | 0.035934872 | -0.98473941 | Remorin | elongation zone | 6 h |
| HORVU.MOREX.r3.4HG0404530.1 | -1.393158917 | -1.741402323 | -3.405694845 | 0.00127 | 0.043818088 | -0.98727692 | Transmembrane protein, putative | elongation zone | 6 h |
| HORVU.MOREX.r3.2HG0168090.1 | 0.385930594 | 5.385495042 | 3.608058076 | 0.00068 | 0.033986772 | -0.98973551 | Sphingomyelin synthase-like domain-containing protein | elongation zone | 6 h |
| HORVU.MOREX.r3.2HG0125330.1 | -0.38679831 | 4.679437814 | -3.583008571 | 0.00074 | 0.035480233 | -0.98996649 | RNA-binding protein | elongation zone | 6 h |
| HORVU.MOREX.r3.3HG0244040.1 | -1.278156136 | -2.293841422 | -3.43082716 | 0.00117 | 0.042465578 | -0.99026365 | ethylene-responsive transcription factor | elongation zone | 6 h |
| HORVU.MOREX.r3.4HG0368770.2 | 1.469603179 | 2.047876559 | 3.406129094 | 0.00126 | 0.043818088 | -0.99334241 | ATP-dependent Clp protease proteolytic subunit | elongation zone | 6 h |
| HORVU.MOREX.r3.3HG0311870.1 | 1.933191096 | 0.203024233 | 3.443659131 | 0.00113 | 0.04173815 | -1.00169377 | EamA-like transporter family protein | elongation zone | 6 h |
| HORVU.MOREX.r3.6HG0545970.1 | 0.426895373 | 5.654969195 | 3.613211486 | 0.00067 | 0.033870225 | -1.00206576 | Short-chain dehydrogenase/reductase family protein | elongation zone | 6 h |
| HORVU.MOREX.r3.5HG0512230.1 | -1.108969121 | 2.048290165 | -3.46497461 | 0.00106 | 0.040858317 | -1.00515546 | Protein ENHANCED DISEASE RESISTANCE 2-like | elongation zone | 6 h |
| HORVU.MOREX.r3.1HG0060370.1 | 3.203141388 | -0.902409718 | 3.566632158 | 0.00078 | 0.035934872 | -1.00526719 | 1,4-alpha-glucan branching enzyme GlgB | elongation zone | 6 h |
| HORVU.MOREX.r3.2HG0160690.1 | -1.204767415 | -0.418955769 | -3.490105445 | 0.00098 | 0.039681075 | -1.01013463 | RALF | elongation zone | 6 h |

**Table S5** Continued.

| ID | logFC | AveExpr | t | P.Value | adj.P.Val | B | gene description | root zone | time point |
|---|---|---|---|---|---|---|---|---|---|
| HORVU.MOREX.r3.5HG0526200.1 | 0.288792942 | 5.48572856 | 3.59290839 | 0.00072 | 0.034740116 | -1.0152164 | Sec14 cytosolic factor | elongation zone | 6 h |
| HORVU.MOREX.r3.3HG0307120.1 | 0.512115268 | 8.36463409 | 3.706192838 | 0.0005 | 0.030363091 | -1.01550934 | S-adenosyl-L-methionine-dependent methyltransferases superfa | elongation zone | 6 h |
| HORVU.MOREX.r3.4HG0394460.1 | -1.206231266 | 0.734389124 | -3.41876347 | 0.00122 | 0.043346752 | -1.02179996 | Protein DETOXIFICATION | elongation zone | 6 h |
| HORVU.MOREX.r3.3HG0246500.1 | -0.864018577 | 3.040521827 | -3.644179625 | 0.00061 | 0.032525759 | -1.02829179 | Peptide-N4-(N-acetyl-beta-glucosaminyl)asparagine amidase A | elongation zone | 6 h |
| HORVU.MOREX.r3.5HG0477980.1 | -0.607312768 | 3.756005664 | -3.556936486 | 0.0008 | 0.035982407 | -1.02981776 | Acetyl-coenzyme A synthetase | elongation zone | 6 h |
| HORVU.MOREX.r3.4HG0364050.1 | -0.739879198 | 2.580176455 | -3.468885514 | 0.00105 | 0.040858317 | -1.03252295 | Sodium/hydrogen exchanger | elongation zone | 6 h |
| HORVU.MOREX.r3.7HG0678020.1 | 0.99519063 | 1.442697113 | 3.430070132 | 0.00118 | 0.042487533 | -1.03271959 | WPP domain-interacting protein 1 | elongation zone | 6 h |
| HORVU.MOREX.r3.3HG0280630.1 | 0.34881626 | 5.914468206 | 3.610454703 | 0.00068 | 0.033870225 | -1.03381986 | Glycosylphosphatidylinositol anchor attachment 1 protein | elongation zone | 6 h |
| HORVU.MOREX.r3.5HG0480980.1 | -1.803200929 | -1.63821174 | -3.435979877 | 0.00116 | 0.042177981 | -1.03969479 | Spermidine/putrescine import ATP-binding protein PotA | elongation zone | 6 h |
| HORVU.MOREX.r3.5HG0516660.1 | 0.381593632 | 4.126867737 | 3.53898501 | 0.00085 | 0.036941016 | -1.04134984 | LEM3 (Ligand-effect modulator 3) family protein | elongation zone | 6 h |
| HORVU.MOREX.r3.5HG0461950.1 | -0.783808851 | 2.971610109 | -3.518334502 | 0.0009 | 0.038318953 | -1.04145443 | 2-isopropylmalate synthase 2 | elongation zone | 6 h |
| HORVU.MOREX.r3.5HG0458300.1 | 1.70854909 | 0.796200207 | 3.406250817 | 0.00126 | 0.043818088 | -1.04180535 | transcription repressor | elongation zone | 6 h |
| HORVU.MOREX.r3.2HG0117880.1 | -2.863040425 | -0.66122785 | -3.389228939 | 0.00133 | 0.045015937 | -1.0542159 | GDSL esterase/lipase | elongation zone | 6 h |
| HORVU.MOREX.r3.5HG0462880.1 | -2.02694221 | -1.070584552 | -3.496708272 | 0.00096 | 0.039433228 | -1.05478873 | Serine/threonine-protein phosphatase 7 long form-like protein | elongation zone | 6 h |
| HORVU.MOREX.r3.6HG0582480.1 | -0.728474771 | 2.535480373 | -3.463267635 | 0.00106 | 0.040858317 | -1.05481458 | Ethylene-responsive transcription factor-like protein | elongation zone | 6 h |
| HORVU.MOREX.r3.3HG0330150.1 | -2.128516404 | -0.302066567 | -3.376170856 | 0.00138 | 0.046197603 | -1.05692469 | MYB transcription factor | elongation zone | 6 h |
| HORVU.MOREX.r3.3HG0297870.1 | -2.494816956 | -0.912086094 | -3.395964009 | 0.0013 | 0.044484558 | -1.05777805 | BAG family molecular chaperone regulator 2 | elongation zone | 6 h |
| HORVU.MOREX.r3.3HG0292680.1 | -0.764760996 | 2.459132616 | -3.559836052 | 0.00079 | 0.035972648 | -1.0693981 | Replicase polyprotein 1a | elongation zone | 6 h |
| HORVU.MOREX.r3.5HG0515030.1 | 0.620798381 | 4.511137793 | 3.492575917 | 0.00097 | 0.039646685 | -1.07204352 | Obg-like ATPase 1 | elongation zone | 6 h |
| HORVU.MOREX.r3.3HG0280060.1 | -1.561307679 | -2.863950573 | -3.368182939 | 0.00142 | 0.046697345 | -1.07712742 | Heavy metal transport/detoxification superfamily protein | elongation zone | 6 h |
| HORVU.MOREX.r3.7HG0736150.1 | -1.017119524 | 8.171407337 | -3.730428017 | 0.00047 | 0.029138607 | -1.07832428 | Phosphatidylinositol transfer protein | elongation zone | 6 h |
| HORVU.MOREX.r3.5HG0486660.1 | -1.069270516 | -2.069765734 | -3.413251048 | 0.00124 | 0.043691234 | -1.0797734 | Calcium-binding protein | elongation zone | 6 h |
| HORVU.MOREX.r3.2HG0096600.1 | 0.526825678 | 8.083914516 | 3.677340916 | 0.00055 | 0.031075507 | -1.08052466 | Pyruvate dehydrogenase E1 component subunit alpha | elongation zone | 6 h |
| HORVU.MOREX.r3.4HG0398070.1 | 0.688446993 | 3.523000734 | 3.429443814 | 0.00118 | 0.042490959 | -1.08128542 | Protein kinase | elongation zone | 6 h |
| HORVU.MOREX.r3.5HG0493070.1 | 0.430186531 | 6.006757924 | 3.605737926 | 0.00069 | 0.033986772 | -1.08318631 | Peptidylprolyl isomerase | elongation zone | 6 h |
| HORVU.MOREX.r3.7HG0704450.1 | -0.487948587 | 5.126566504 | -3.579407991 | 0.00075 | 0.035624994 | -1.08334799 | Hexosyltransferase | elongation zone | 6 h |
| HORVU.MOREX.r3.6HG0601670.1 | 0.911809891 | 2.45037985 | 3.392503188 | 0.00132 | 0.044798804 | -1.08476692 | Phosphatidic acid phosphatase | elongation zone | 6 h |
| HORVU.MOREX.r3.3HG0219380.1 | -1.008140873 | 4.367645105 | -3.640953483 | 0.00062 | 0.032645728 | -1.08733599 | Glutamate receptor ionotropic, NMDA 3A | elongation zone | 6 h |
| HORVU.MOREX.r3.3HG0302570.1 | -2.508745825 | -3.210356093 | -3.367652034 | 0.00142 | 0.046697345 | -1.09062584 | Laccase | elongation zone | 6 h |
| HORVU.MOREX.r3.6HG0577220.1 | -1.96185772 | -2.233904068 | -3.362637819 | 0.00144 | 0.046971179 | -1.09188501 | Glycosyltransferase | elongation zone | 6 h |
| HORVU.MOREX.r3.1HG0046180.1 | -0.463043475 | 5.918664759 | -3.630207163 | 0.00064 | 0.033339257 | -1.09701758 | Ser/Thr protein phosphatase family protein, expressed | elongation zone | 6 h |
| HORVU.MOREX.r3.4HG0380540.1 | -0.617719899 | 3.365321911 | -3.506978351 | 0.00093 | 0.039007436 | -1.09899924 | Aquaporin-1 | elongation zone | 6 h |
| HORVU.MOREX.r3.1HG0077950.1 | 0.363648708 | 4.840037428 | 3.542765721 | 0.00084 | 0.036935434 | -1.10307781 | BSD domain containing protein | elongation zone | 6 h |
| HORVU.MOREX.r3.2HG0204940.1 | 0.531883725 | 6.058216754 | 3.565556499 | 0.00078 | 0.035934872 | -1.10327728 | Crossover junction endonuclease EME1 | elongation zone | 6 h |
| HORVU.MOREX.r3.3HG0222830.1 | 0.327945155 | 7.515891005 | 3.668542031 | 0.00057 | 0.031591904 | -1.10427142 | Heavy metal transport/detoxification superfamily protein | elongation zone | 6 h |
| HORVU.MOREX.r3.5HG0473970.1 | -2.387634429 | -3.048444071 | -3.377725252 | 0.00138 | 0.04605869 | -1.10640124 | Glycosyltransferase | elongation zone | 6 h |
| HORVU.MOREX.r3.3HG0321140.1 | -2.032977367 | -2.331879343 | -3.374358192 | 0.00139 | 0.046297461 | -1.1071247 | Glutamate-1-semialdehyde 2,1-aminomutase 2 | elongation zone | 6 h |
| HORVU.MOREX.r3.5HG0518390.1 | -2.60642644 | -0.172193667 | -3.499203216 | 0.00095 | 0.039256264 | -1.10790825 | BTB/POZ domain containing protein | elongation zone | 6 h |
| HORVU.MOREX.r3.4HG0415420.1 | 0.6046561 | 6.947213857 | 3.618530517 | 0.00066 | 0.033752681 | -1.11775158 | Tubulin beta chain | elongation zone | 6 h |
| HORVU.MOREX.r3.7HG0727350.1 | 0.461459796 | 10.95622204 | 3.750354884 | 0.00044 | 0.028211116 | -1.1187487 | #N/A | elongation zone | 6 h |
| HORVU.MOREX.r3.6HG0564590.1 | 0.482683245 | 6.449269546 | 3.576529748 | 0.00075 | 0.035692712 | -1.1214732 | O-fucosyltransferase family protein | elongation zone | 6 h |

**Table S5** Continued.

| ID | logFC | AveExpr | t | P.Value | adj.P.Val | B | gene description | root zone | time point |
|---|---|---|---|---|---|---|---|---|---|
| HORVU.MOREX.r3.6HG0573930.1 | -0.423991807 | 3.609831823 | -3.477204752 | 0.00102 | 0.04048377 | -1.12530013 | Mediator of RNA polymerase II transcription subunit 18 | elongation zone | 6 h |
| HORVU.MOREX.r3.6HG0627380.3 | -1.669154308 | 1.232115316 | -3.402710744 | 0.00128 | 0.043833962 | -1.12727549 | 2-oxoglutarate (2OG) and Fe(II)-dependent oxygenase superfami | elongation zone | 6 h |
| HORVU.MOREX.r3.4HG0410640.1 | 0.387804262 | 8.151713261 | 3.659867614 | 0.00058 | 0.031886221 | -1.12878455 | Dolichyl-diphosphooligosaccharide--protein glycosyltransferase s | elongation zone | 6 h |
| HORVU.MOREX.r3.5HG0532150.1 | -1.551110063 | -2.26204476 | -3.387437424 | 0.00134 | 0.04518326 | -1.13462054 | Expansin | elongation zone | 6 h |
| HORVU.MOREX.r3.5HG0536710.1 | 0.483148693 | 6.624728725 | 3.619464838 | 0.00066 | 0.033752681 | -1.134657 | NAD/NADP-dependent betaine aldehyde dehydrogenase | elongation zone | 6 h |
| HORVU.MOREX.r3.2HG0215830.1 | 0.948562118 | 2.436514446 | 3.362510313 | 0.00144 | 0.046971179 | -1.1379331 | CsAtPR5 | elongation zone | 6 h |
| HORVU.MOREX.r3.2HG0107960.1 | -0.550273195 | 3.123366617 | -3.460354707 | 0.00107 | 0.040858317 | -1.13869949 | Phosphoinositide phospholipase C | elongation zone | 6 h |
| HORVU.MOREX.r3.3HG0220360.1 | 0.851891693 | 4.616464522 | 3.458617057 | 0.00108 | 0.040886579 | -1.13958331 | Disease resistance protein | elongation zone | 6 h |
| HORVU.MOREX.r3.6HG0615130.1 | -0.745681479 | 3.141376461 | -3.565193759 | 0.00078 | 0.035934872 | -1.14154508 | TCP transcription factor | elongation zone | 6 h |
| HORVU.MOREX.r3.4HG0417010.1 | -1.06888801 | -1.713964929 | -3.40506063 | 0.00127 | 0.043818088 | -1.14272089 | Receptor protein kinase, putative | elongation zone | 6 h |
| HORVU.MOREX.r3.7HG0747230.1 | -0.591534859 | 2.606944875 | -3.438354855 | 0.00115 | 0.041957464 | -1.14391566 | Non-specific serine/threonine protein kinase | elongation zone | 6 h |
| HORVU.MOREX.r3.7HG0652950.1 | 1.188317314 | 1.220754935 | 3.353216324 | 0.00148 | 0.047599823 | -1.14555958 | NBS-LRR-like resistance protein | elongation zone | 6 h |
| HORVU.MOREX.r3.1HG0026080.1 | -0.627323442 | 4.010773585 | -3.488945486 | 0.00098 | 0.039681075 | -1.15197786 | RING finger protein | elongation zone | 6 h |
| HORVU.MOREX.r3.2HG0214240.1 | 0.591760427 | 3.183258219 | 3.420927059 | 0.00121 | 0.043205753 | -1.15214231 | Disease resistance protein (NBS-LRR class) family | elongation zone | 6 h |
| HORVU.MOREX.r3.2HG0168770.1 | -0.60205077 | 3.537711265 | -3.490154737 | 0.00098 | 0.039681075 | -1.16276627 | Scarecrow transcription factor family protein | elongation zone | 6 h |
| HORVU.MOREX.r3.3HG0298680.1 | -0.778583864 | 1.911503731 | -3.445868675 | 0.00112 | 0.041610797 | -1.16431576 | Myb transcription factor | elongation zone | 6 h |
| HORVU.MOREX.r3.1HG0086120.1 | 1.659779471 | 4.055301306 | 3.370417593 | 0.00141 | 0.046697345 | -1.16826027 | Histone H2B | elongation zone | 6 h |
| HORVU.MOREX.r3.1HG0086390.1 | -0.906213328 | -0.260888993 | -3.444991364 | 0.00112 | 0.041645779 | -1.17445583 | #N/A | elongation zone | 6 h |
| HORVU.MOREX.r3.5HG0513340.1 | -1.599288247 | -3.018568987 | -3.338593156 | 0.00155 | 0.048879452 | -1.17573303 | Basic-leucine zipper (bZIP) transcription factor family protein | elongation zone | 6 h |
| HORVU.MOREX.r3.1HG0072910.2 | 0.405567947 | 4.342104799 | 3.460615287 | 0.00107 | 0.040858317 | -1.18214329 | Serine/threonine-protein kinase ATM | elongation zone | 6 h |
| HORVU.MOREX.r3.5HG0470170.1 | 0.552753579 | 3.126834047 | 3.40833289 | 0.00126 | 0.043818088 | -1.19137647 | F-box family protein | elongation zone | 6 h |
| HORVU.MOREX.r3.1HG0091360.1 | 0.490690398 | 6.214276876 | 3.557310498 | 0.0008 | 0.035982407 | -1.19657145 | Delta(14)-sterol reductase | elongation zone | 6 h |
| HORVU.MOREX.r3.4HG0342000.1 | -0.717688692 | 0.190754791 | -3.416497126 | 0.00123 | 0.043449664 | -1.20642895 | 4-coumarate:CoA ligase-like protein | elongation zone | 6 h |
| HORVU.MOREX.r3.5HG0476380.1 | 0.590576758 | 4.634572865 | 3.466944592 | 0.00105 | 0.040858317 | -1.21230453 | Major facilitator superfamily transporter | elongation zone | 6 h |
| HORVU.MOREX.r3.5HG0501980.1 | 0.406622867 | 5.561069419 | 3.522936604 | 0.00089 | 0.038100943 | -1.2137628 | Rop guanine nucleotide exchange factor, putative | elongation zone | 6 h |
| HORVU.MOREX.r3.7HG0739230.1 | -2.284631643 | -3.450977617 | -3.326417649 | 0.0016 | 0.049756476 | -1.21437207 | Jacalin-like lectin | elongation zone | 6 h |
| HORVU.MOREX.r3.5HG0461280.1 | -0.488690851 | 6.014759518 | -3.552695926 | 0.00081 | 0.036184546 | -1.21476499 | Calcium-binding EF-hand protein | elongation zone | 6 h |
| HORVU.MOREX.r3.6HG0554050.1 | -0.910612159 | -1.569574336 | -3.381902507 | 0.00136 | 0.045711209 | -1.216031 | cyclin-dependent kinase inhibitor | elongation zone | 6 h |
| HORVU.MOREX.r3.2HG0156810.1 | 1.708782825 | 1.309433717 | 3.324454835 | 0.00161 | 0.049971 | -1.22382897 | Sodium/potassium/calcium exchanger 3 | elongation zone | 6 h |
| HORVU.MOREX.r3.5HG0517740.1 | -1.190237811 | -0.307208015 | -3.533567508 | 0.00086 | 0.037429607 | -1.22712874 | Peroxidase | elongation zone | 6 h |
| HORVU.MOREX.r3.3HG0245440.1 | -0.99505647 | 0.700858326 | -3.39082436 | 0.00132 | 0.044875615 | -1.22935823 | Cellulose synthase | elongation zone | 6 h |
| HORVU.MOREX.r3.1HG0079750.1 | -1.123857566 | -1.155378593 | -3.364004037 | 0.00143 | 0.046971179 | -1.23041424 | MYB transcription factor | elongation zone | 6 h |
| HORVU.MOREX.r3.3HG0254940.1 | -0.88368612 | 2.451212431 | -3.601933452 | 0.0007 | 0.034109179 | -1.23159154 | Peroxidase | elongation zone | 6 h |
| HORVU.MOREX.r3.3HG0288140.1 | 0.311331045 | 6.173911395 | 3.563158785 | 0.00078 | 0.035934872 | -1.23653531 | Golgin candidate 1 | elongation zone | 6 h |
| HORVU.MOREX.r3.2HG0172110.1 | -0.577920929 | 5.282401422 | -3.561790827 | 0.00079 | 0.035934872 | -1.24485477 | Dihydrofolate reductase | elongation zone | 6 h |
| HORVU.MOREX.r3.1HG0067720.1 | -1.073963716 | -2.106023761 | -3.350940847 | 0.00149 | 0.047781359 | -1.24906573 | #N/A | elongation zone | 6 h |
| HORVU.MOREX.r3.4HG0348670.1 | 0.32495001 | 7.764170114 | 3.638214948 | 0.00062 | 0.032755332 | -1.24995178 | #N/A | elongation zone | 6 h |
| HORVU.MOREX.r3.1HG0061710.1 | -0.490520508 | 5.632385509 | -3.577179121 | 0.00075 | 0.035692712 | -1.25831528 | Trichome birefringence-like protein | elongation zone | 6 h |
| HORVU.MOREX.r3.1HG0002960.1 | -0.907623908 | 0.278872787 | -3.502006835 | 0.00095 | 0.039256264 | -1.26945082 | NBS-LRR disease resistance protein, putative, expressed | elongation zone | 6 h |
| HORVU.MOREX.r3.3HG0275910.1 | 3.05451757 | -0.86003395 | 3.442746676 | 0.00113 | 0.041777902 | -1.2728986 | Transcription factor GTE4-like protein | elongation zone | 6 h |
| HORVU.MOREX.r3.5HG0420630.1 | 0.513343995 | 3.720926699 | 3.447817025 | 0.00111 | 0.041517003 | -1.27366929 | Ankyrin repeat family protein | elongation zone | 6 h |

**Table S5** Continued.

| ID | logFC | AveExpr | t | P.Value | adj.P.Val | B | gene description | root zone | time point |
|---|---|---|---|---|---|---|---|---|---|
| HORVU.MOREX.r3.3HG0224680.1 | 0.378954852 | 5.580543005 | 3.514757053 | 0.00091 | 0.038579099 | -1.27368031 | Ferredoxin reductase-like protein | elongation zone | 6 h |
| HORVU.MOREX.r3.7HG0734670.1 | 0.481562973 | 6.626504106 | 3.57354513 | 0.00076 | 0.035856081 | -1.27401052 | L-ascorbate oxidase | elongation zone | 6 h |
| HORVU.MOREX.r3.7HG0751060.1 | 0.406171284 | 4.948457618 | 3.503663188 | 0.00094 | 0.039176352 | -1.27432509 | Disease resistance protein (NBS-LRR class) family | elongation zone | 6 h |
| HORVU.MOREX.r3.4HG0349310.1 | 0.368968984 | 5.375878972 | 3.499016835 | 0.00095 | 0.039256264 | -1.27626465 | Sec1 family domain-containing protein 2 | elongation zone | 6 h |
| HORVU.MOREX.r3.4HG0354970.1 | 0.580167076 | 6.732308281 | 3.55928249 | 0.00079 | 0.035972648 | -1.27643003 | Omega-3 fatty acid desaturase | elongation zone | 6 h |
| HORVU.MOREX.r3.7HG0718190.1 | -0.737788924 | 6.009060521 | -3.617939424 | 0.00066 | 0.033752681 | -1.28265013 | ABC transporter family protein | elongation zone | 6 h |
| HORVU.MOREX.r3.2HG0192610.1 | -0.56292893 | 5.913188646 | -3.46943483 | 0.00104 | 0.040858317 | -1.28514394 | DNA ligase | elongation zone | 6 h |
| HORVU.MOREX.r3.1HG0073460.1 | -0.492927168 | 2.975475398 | -3.398807763 | 0.00129 | 0.044254508 | -1.2878058 | Microneme/rhoptry antigen | elongation zone | 6 h |
| HORVU.MOREX.r3.4HG0389250.1 | -1.39026176 | 3.018685792 | -3.352746352 | 0.00148 | 0.047599823 | -1.29554757 | lipase, putative (DUF620) | elongation zone | 6 h |
| HORVU.MOREX.r3.6HG0616240.1 | -0.913902885 | -0.522759558 | -3.465007106 | 0.00106 | 0.040858317 | -1.30130591 | Endoglucanase | elongation zone | 6 h |
| HORVU.MOREX.r3.2HG0187940.1 | -0.732937167 | 5.04269045 | -3.499592904 | 0.00095 | 0.039256264 | -1.30288943 | Niemann-Pick C1 protein | elongation zone | 6 h |
| HORVU.MOREX.r3.7HG0739030.1 | -0.562701519 | 5.253411819 | -3.51993301 | 0.0009 | 0.038291477 | -1.30478675 | Kinase family protein | elongation zone | 6 h |
| HORVU.MOREX.r3.7HG0642520.1 | -1.37787495 | -0.871699978 | -3.474361026 | 0.00103 | 0.040615136 | -1.30806933 | Peroxidase | elongation zone | 6 h |
| HORVU.MOREX.r3.3HG0312490.1 | 0.660330948 | 2.855498414 | 3.334247866 | 0.00157 | 0.049210449 | -1.30856954 | Short-chain dehydrogenase/reductase family protein | elongation zone | 6 h |
| HORVU.MOREX.r3.4HG0346720.1 | 0.475678784 | 10.60860825 | 3.694266969 | 0.00052 | 0.030363091 | -1.31381815 | Protein disulfide-isomerase | elongation zone | 6 h |
| HORVU.MOREX.r3.4HG0384390.1 | -0.898818765 | 1.57490258 | -3.391797497 | 0.00132 | 0.044819198 | -1.32193356 | Protein kinase family protein, putative, expressed | elongation zone | 6 h |
| HORVU.MOREX.r3.1HG0023970.1 | -1.093184483 | 0.759180505 | -3.368641305 | 0.00141 | 0.046697345 | -1.33028324 | GATA transcription factor, putative | elongation zone | 6 h |
| HORVU.MOREX.r3.1HG0057030.1 | 0.67712633 | 6.546045506 | 3.511214245 | 0.00092 | 0.038757609 | -1.33666601 | Peroxidase | elongation zone | 6 h |
| HORVU.MOREX.r3.1HG0063810.1 | 0.400247305 | 4.973905479 | 3.451084979 | 0.0011 | 0.041410027 | -1.34007455 | Cyclin family protein | elongation zone | 6 h |
| HORVU.MOREX.r3.1HG0008200.1 | 1.528341759 | 1.354368368 | 3.386105239 | 0.00134 | 0.045289016 | -1.34512433 | Cytochrome P450 | elongation zone | 6 h |
| HORVU.MOREX.r3.7HG0711890.1 | -1.024141786 | 2.553034049 | -3.448104206 | 0.00111 | 0.041517003 | -1.34543805 | O-methyltransferase | elongation zone | 6 h |
| HORVU.MOREX.r3.7HG0715570.1 | 0.394750591 | 6.466382964 | 3.520264115 | 0.00089 | 0.038291477 | -1.35382994 | Transmembrane protein | elongation zone | 6 h |
| HORVU.MOREX.r3.5HG0525700.1 | 0.366859152 | 7.295514432 | 3.530427997 | 0.00087 | 0.037633014 | -1.35605801 | Leukotriene a-4 hydrolase aminopeptidase | elongation zone | 6 h |
| HORVU.MOREX.r3.4HG0332920.1 | 0.30273669 | 7.313310024 | 3.55236508 | 0.00081 | 0.036184546 | -1.3578162 | 26S proteasome non-atpase regulatory subunit, putative | elongation zone | 6 h |
| HORVU.MOREX.r3.6HG0568640.1 | 0.365740231 | 4.543924675 | 3.440499803 | 0.00114 | 0.041925776 | -1.35959638 | Methyltransferase domain protein | elongation zone | 6 h |
| HORVU.MOREX.r3.7HG0688870.1 | -0.678369579 | 6.011072385 | -3.540228487 | 0.00084 | 0.036941016 | -1.36162601 | Monodehydroascorbate reductase | elongation zone | 6 h |
| HORVU.MOREX.r3.6HG0574990.1 | -0.585422916 | 7.228108567 | -3.584936172 | 0.00073 | 0.035436154 | -1.36498367 | ABC transporter B family protein | elongation zone | 6 h |
| HORVU.MOREX.r3.7HG0668820.1 | -0.584137731 | 4.620228035 | -3.562845071 | 0.00079 | 0.035934872 | -1.3683326 | Endoglucanase | elongation zone | 6 h |
| HORVU.MOREX.r3.3HG0298750.1 | 0.54170438 | 5.760333104 | 3.489058941 | 0.00098 | 0.039681075 | -1.36976485 | magnesium transporter NIPA (DUF803) | elongation zone | 6 h |
| HORVU.MOREX.r3.4HG0400740.1 | 0.23990593 | 8.244974267 | 3.582253051 | 0.00074 | 0.035480233 | -1.37074328 | Cell division cycle protein 48-like protein | elongation zone | 6 h |
| HORVU.MOREX.r3.3HG0238180.1 | 0.422413375 | 4.144586965 | 3.407385077 | 0.00126 | 0.043818088 | -1.37129481 | #N/A | elongation zone | 6 h |
| HORVU.MOREX.r3.6HG0555700.1 | 0.376158393 | 8.875975145 | 3.618782469 | 0.00066 | 0.033752681 | -1.37820655 | Calreticulin/calnexin | elongation zone | 6 h |
| HORVU.MOREX.r3.7HG0654330.1 | -0.520051316 | 7.269216004 | -3.562970179 | 0.00079 | 0.035934872 | -1.38182486 | ABC transporter, putative | elongation zone | 6 h |
| HORVU.MOREX.r3.3HG0326430.1 | 0.317758576 | 5.095188036 | 3.456508471 | 0.00109 | 0.040886579 | -1.38687321 | Diphosphomevalonate decarboxylase | elongation zone | 6 h |
| HORVU.MOREX.r3.7HG0688040.1 | 0.721205581 | 2.581060956 | 3.327645305 | 0.0016 | 0.049650942 | -1.39067514 | DUF241 domain protein, putative (DUF241) | elongation zone | 6 h |
| HORVU.MOREX.r3.5HG0527650.1 | 0.637056037 | 4.818573339 | 3.420425644 | 0.00121 | 0.043205753 | -1.39463394 | Succinate--CoA ligase [ADP-forming] subunit alpha-1, mitochond | elongation zone | 6 h |
| HORVU.MOREX.r3.1HG0073170.1 | 0.624287957 | 5.178735655 | 3.407966884 | 0.00126 | 0.043818088 | -1.39670533 | F-box protein | elongation zone | 6 h |
| HORVU.MOREX.r3.5HG0482570.1 | -0.321675158 | 4.680679207 | -3.402551342 | 0.00128 | 0.043833962 | -1.40711225 | UDP-N-acetylglucosamine--N-acetylmuramyl-(pentapeptide) pyr | elongation zone | 6 h |
| HORVU.MOREX.r3.5HG0533230.1 | 0.487085107 | 5.643176833 | 3.47511291 | 0.00103 | 0.040615136 | -1.40969915 | Pirin-like protein | elongation zone | 6 h |
| HORVU.MOREX.r3.1HG0079140.1 | -0.554467319 | 4.43581546 | -3.456581276 | 0.00109 | 0.040886579 | -1.41177469 | Kinase family protein | elongation zone | 6 h |
| HORVU.MOREX.r3.3HG0307850.1 | 0.652468311 | 6.891758697 | 3.555567126 | 0.0008 | 0.035987623 | -1.41423869 | Basic 7S globulin | elongation zone | 6 h |

**Table S5** Continued.

| ID | logFC | AveExpr | t | P.Value | adj.P.Val | B | gene description | root zone | time point |
|---|---|---|---|---|---|---|---|---|---|
| HORVU.MOREX.r3.7HG0730510.1 | -0.92235097 | -1.831161525 | -3.329009471 | 0.00159 | 0.049525669 | -1.42151805 | Germin-like protein 1 | elongation zone | 6 h |
| HORVU.MOREX.r3.2HG0117980.1 | 0.380715929 | 6.65430886 | 3.507293776 | 0.00093 | 0.039007436 | -1.42154302 | Transmembrane protein 115 | elongation zone | 6 h |
| HORVU.MOREX.r3.5HG0488040.1 | -1.312871965 | -1.230138021 | -3.378883603 | 0.00137 | 0.045974621 | -1.43006309 | Blue copper protein | elongation zone | 6 h |
| HORVU.MOREX.r3.1HG0094980.1 | 0.486850051 | 4.287158731 | 3.374791352 | 0.00139 | 0.046297461 | -1.43033665 | Mitochondrial outer membrane porin | elongation zone | 6 h |
| HORVU.MOREX.r3.2HG0213250.1 | 0.542454716 | 6.347478748 | 3.506418383 | 0.00093 | 0.039007436 | -1.43203225 | Choline transporter-related family protein | elongation zone | 6 h |
| HORVU.MOREX.r3.4HG0363180.1 | 0.258002357 | 9.157327695 | 3.596301548 | 0.00071 | 0.034460123 | -1.43222046 | Translationally-controlled tumor protein | elongation zone | 6 h |
| HORVU.MOREX.r3.1HG0070340.1 | -0.999266551 | 4.307026595 | -3.486595093 | 0.00099 | 0.039808834 | -1.43395633 | Protein kinase-like protein | elongation zone | 6 h |
| HORVU.MOREX.r3.7HG0674990.1 | 0.4790589 | 3.286294703 | 3.34356581 | 0.00152 | 0.048382678 | -1.43598953 | Mannan endo-1,4-beta-mannosidase-like protein | elongation zone | 6 h |
| HORVU.MOREX.r3.5HG0490030.1 | 0.243342609 | 7.481265074 | 3.547777977 | 0.00082 | 0.036546446 | -1.45729909 | Vacuolar protein sorting-associated protein 52 | elongation zone | 6 h |
| HORVU.MOREX.r3.2HG0134400.1 | -0.370541306 | 6.098222844 | -3.478229521 | 0.00102 | 0.040436406 | -1.45938479 | Protein DETOXIFICATION | elongation zone | 6 h |
| HORVU.MOREX.r3.4HG0382500.1 | 0.419481917 | 5.221396135 | 3.422721373 | 0.0012 | 0.043133621 | -1.46115554 | Seed maturation protein PM36, putative, expressed | elongation zone | 6 h |
| HORVU.MOREX.r3.6HG0555660.2 | 0.386082859 | 7.208637701 | 3.513993449 | 0.00091 | 0.038589259 | -1.46352661 | Calreticulin/calnexin | elongation zone | 6 h |
| HORVU.MOREX.r3.7HG0726770.1 | -0.912777323 | 2.292249938 | -3.370036361 | 0.00141 | 0.046697345 | -1.47054762 | Glycine-rich protein | elongation zone | 6 h |
| HORVU.MOREX.r3.1HG0050780.1 | 0.319692238 | 6.71627124 | 3.518659058 | 0.0009 | 0.038318953 | -1.47234215 | Major facilitator superfamily domain-containing protein | elongation zone | 6 h |
| HORVU.MOREX.r3.4HG0411870.1 | 0.737951282 | 4.541599175 | 3.367149856 | 0.00142 | 0.046697345 | -1.47706039 | Cysteine-rich repeat secretory protein | elongation zone | 6 h |
| HORVU.MOREX.r3.2HG0111880.1 | 2.361155522 | -1.175412118 | 3.364418186 | 0.00143 | 0.046971179 | -1.48260999 | DNA/RNA helicase | elongation zone | 6 h |
| HORVU.MOREX.r3.6HG0604210.1 | 0.365611674 | 6.058555923 | 3.451678315 | 0.0011 | 0.041410027 | -1.48362219 | Protease HtpX | elongation zone | 6 h |
| HORVU.MOREX.r3.4HG0405400.1 | -1.240438052 | 2.976905729 | -3.393652076 | 0.00131 | 0.044719161 | -1.49830958 | DUF936 family protein | elongation zone | 6 h |
| HORVU.MOREX.r3.7HG0661490.1 | 0.306509178 | 6.094894031 | 3.463143467 | 0.00106 | 0.040858317 | -1.50090614 | Stromal cell-derived factor 2 | elongation zone | 6 h |
| HORVU.MOREX.r3.7HG0749870.1 | -0.585558346 | 4.569999785 | -3.464286228 | 0.00106 | 0.040858317 | -1.50914651 | Glucuronoxylan 4-O-methyltransferase | elongation zone | 6 h |
| HORVU.MOREX.r3.7HG0677510.1 | 0.443940306 | 5.971823823 | 3.446115077 | 0.00112 | 0.041610797 | -1.52115952 | Glycosyltransferase | elongation zone | 6 h |
| HORVU.MOREX.r3.2HG0185340.1 | -0.848064 | 1.427350358 | -3.354002437 | 0.00148 | 0.047599823 | -1.53081582 | Cationic amino acid transporter, putative | elongation zone | 6 h |
| HORVU.MOREX.r3.7HG0728190.1 | 0.438885105 | 4.089813032 | 3.358257464 | 0.00146 | 0.047382603 | -1.53391466 | Purple acid phosphatase | elongation zone | 6 h |
| HORVU.MOREX.r3.1HG0085380.1 | -0.465559834 | 4.053078396 | -3.348340733 | 0.0015 | 0.048034173 | -1.53404288 | 2-oxoglutarate (2OG) and Fe(II)-dependent oxygenase superfami | elongation zone | 6 h |
| HORVU.MOREX.r3.2HG0177290.1 | 2.196059937 | -1.308802463 | 3.342778792 | 0.00153 | 0.048382678 | -1.53801593 | C3H4 type zinc finger protein (DUF23) | elongation zone | 6 h |
| HORVU.MOREX.r3.2HG0127260.1 | -1.06885664 | -1.061970634 | -3.355454706 | 0.00147 | 0.047574989 | -1.54274819 | COBRA-like protein | elongation zone | 6 h |
| HORVU.MOREX.r3.5HG0513440.1 | 0.357121751 | 5.701505149 | 3.431055316 | 0.00117 | 0.042465578 | -1.54893642 | #N/A | elongation zone | 6 h |
| HORVU.MOREX.r3.3HG0280970.1 | 0.456301625 | 5.971205795 | 3.420887364 | 0.00121 | 0.043205753 | -1.55811501 | Protein methyltransferase | elongation zone | 6 h |
| HORVU.MOREX.r3.1HG0093710.1 | 0.233333322 | 6.865906621 | 3.466695224 | 0.00105 | 0.040858317 | -1.55946627 | dihydroflavonol 4-reductase/flavanone protein | elongation zone | 6 h |
| HORVU.MOREX.r3.4HG0404740.1 | -0.702204995 | 2.889075204 | -3.412341999 | 0.00124 | 0.04373537 | -1.57021574 | Hydrolase, alpha/beta fold family protein, expressed | elongation zone | 6 h |
| HORVU.MOREX.r3.2HG0212570.1 | 0.383682747 | 10.53927453 | 3.599788197 | 0.0007 | 0.03425392 | -1.57479522 | Calreticulin | elongation zone | 6 h |
| HORVU.MOREX.r3.7HG0736930.1 | 0.253405497 | 7.557950747 | 3.50834383 | 0.00093 | 0.039007436 | -1.57740298 | Embryo yellow protein | elongation zone | 6 h |
| HORVU.MOREX.r3.2HG0107900.1 | 0.38593915 | 7.940344995 | 3.48736894 | 0.00099 | 0.039793297 | -1.57988236 | Elongation factor 1-alpha | elongation zone | 6 h |
| HORVU.MOREX.r3.4HG0352200.1 | -0.676587684 | 3.627838098 | -3.342552995 | 0.00153 | 0.048382678 | -1.59052589 | Methyltransferase-related protein, putative | elongation zone | 6 h |
| HORVU.MOREX.r3.4HG0406140.1 | -0.478817603 | 5.424724713 | -3.402773487 | 0.00128 | 0.043833962 | -1.59164637 | Transmembrane protein 131 | elongation zone | 6 h |
| HORVU.MOREX.r3.2HG0182400.1 | -0.757431318 | 5.649464396 | -3.449697557 | 0.00111 | 0.041507984 | -1.59662405 | Aquaporin | elongation zone | 6 h |
| HORVU.MOREX.r3.4HG0416190.1 | 0.267107679 | 9.750154012 | 3.582246804 | 0.00074 | 0.035480233 | -1.61030611 | Coatomer subunit alpha | elongation zone | 6 h |
| HORVU.MOREX.r3.2HG0124660.1 | -0.343600778 | 4.762294428 | -3.368817039 | 0.00141 | 0.046697345 | -1.61109905 | Dihydropteroate synthase, putative | elongation zone | 6 h |
| HORVU.MOREX.r3.1HG0067410.1 | 0.362918449 | 5.64638285 | 3.385052511 | 0.00135 | 0.045357194 | -1.61633554 | 1-acyl-sn-glycerol-3-phosphate acyltransferase | elongation zone | 6 h |
| HORVU.MOREX.r3.6HG0616300.1 | -0.521691506 | 5.182691489 | -3.431754011 | 0.00117 | 0.042465578 | -1.61857136 | Histidine kinase | elongation zone | 6 h |
| HORVU.MOREX.r3.3HG0329760.1 | -0.583435797 | 4.295442546 | -3.346474417 | 0.00151 | 0.048194138 | -1.63224842 | Ankyrin repeat family protein | elongation zone | 6 h |

**Table S5** Continued.

| ID | logFC | AveExpr | t | P.Value | adj.P.Val | B | gene description | root zone | time point |
|---|---|---|---|---|---|---|---|---|---|
| HORVU.MOREX.r3.3HG0326240.1 | 0.502747604 | 5.727778629 | 3.369659365 | 0.00141 | 0.046697345 | -1.63722507 | Beta-fructofuranosidase, insoluble protein | elongation zone | 6 h |
| HORVU.MOREX.r3.6HG0591570.1 | 0.313338275 | 7.825511756 | 3.468402619 | 0.00105 | 0.040858317 | -1.63766134 | 26S proteasome non-atpase regulatory subunit, putative | elongation zone | 6 h |
| HORVU.MOREX.r3.2HG0200920.1 | -0.689918431 | 3.047317061 | -3.435434301 | 0.00116 | 0.042177981 | -1.65165954 | Agmatine coumaroyltransferase-2 | elongation zone | 6 h |
| HORVU.MOREX.r3.5HG0511450.1 | -0.341933839 | 5.650127206 | -3.398241577 | 0.00129 | 0.044255756 | -1.65308296 | ABC transporter B family protein | elongation zone | 6 h |
| HORVU.MOREX.r3.7HG0685870.1 | 0.234160522 | 7.40990073 | 3.46249263 | 0.00107 | 0.040858317 | -1.65881293 | Ankyrin repeat domain containing protein | elongation zone | 6 h |
| HORVU.MOREX.r3.1HG0054010.1 | -0.561367347 | 5.84165697 | -3.448003074 | 0.00111 | 0.041517003 | -1.65896447 | zinc finger B-box protein | elongation zone | 6 h |
| HORVU.MOREX.r3.7HG0728000.1 | -0.571629244 | 5.007313233 | -3.418074448 | 0.00122 | 0.043361315 | -1.66557838 | Glycosyltransferase | elongation zone | 6 h |
| HORVU.MOREX.r3.5HG0529780.1 | 0.400934389 | 6.056073639 | 3.42843689 | 0.00118 | 0.042490959 | -1.6680417 | Pectin lyase-like superfamily protein | elongation zone | 6 h |
| HORVU.MOREX.r3.6HG0566940.1 | -0.559709745 | 4.63219243 | -3.33370285 | 0.00157 | 0.049214696 | -1.68170997 | Sulfurtransferase | elongation zone | 6 h |
| HORVU.MOREX.r3.1HG0093050.1 | -0.806018447 | 4.062497425 | -3.329621542 | 0.00159 | 0.049511037 | -1.6867662 | Indole-3-acetic acid-amido synthetase GH3.3 | elongation zone | 6 h |
| HORVU.MOREX.r3.6HG0600910.1 | -1.106033705 | 1.060688972 | -3.405701632 | 0.00127 | 0.043818088 | -1.68952789 | Aquaporin | elongation zone | 6 h |
| HORVU.MOREX.r3.6HG0557920.1 | -0.830719314 | 4.693871457 | -3.335785062 | 0.00156 | 0.04913705 | -1.69248614 | Auxin response factor | elongation zone | 6 h |
| HORVU.MOREX.r3.6HG0611250.1 | 0.2039617 | 6.631497812 | 3.428293185 | 0.00118 | 0.042490959 | -1.70104563 | BSD domain containing protein | elongation zone | 6 h |
| HORVU.MOREX.r3.1HG0075730.1 | -0.837858635 | 1.539739381 | -3.438983634 | 0.00114 | 0.041952802 | -1.70506589 | Subtilisin-like protease | elongation zone | 6 h |
| HORVU.MOREX.r3.2HG0119160.1 | 0.464707348 | 6.243872664 | 3.33508557 | 0.00156 | 0.049163604 | -1.74308926 | #N/A | elongation zone | 6 h |
| HORVU.MOREX.r3.1HG0028150.1 | -0.821982076 | 4.589078583 | -3.345426561 | 0.00151 | 0.048269373 | -1.76276913 | Leucine-rich repeat-containing protein | elongation zone | 6 h |
| HORVU.MOREX.r3.7HG0687630.1 | 0.540226522 | 7.969316852 | 3.434823879 | 0.00116 | 0.042180694 | -1.7672578 | Acyl carrier protein | elongation zone | 6 h |
| HORVU.MOREX.r3.3HG0318700.1 | -0.475844535 | 6.088388292 | -3.404910549 | 0.00127 | 0.043818088 | -1.76963284 | Phosphotransferase | elongation zone | 6 h |
| HORVU.MOREX.r3.2HG0172730.1 | 0.416197187 | 7.54861275 | 3.403443169 | 0.00127 | 0.043833962 | -1.79656365 | Alpha-ketoglutarate-dependent dioxygenase alkB-like protein 2 | elongation zone | 6 h |
| HORVU.MOREX.r3.7HG0721760.1 | 0.323215645 | 8.421100262 | 3.440384213 | 0.00114 | 0.041925776 | -1.82924354 | Translocon-associated protein subunit alpha | elongation zone | 6 h |
| HORVU.MOREX.r3.6HG0597590.1 | 0.357031684 | 6.568013907 | 3.357453481 | 0.00146 | 0.047382603 | -1.84109689 | Protein disulfide-isomerase | elongation zone | 6 h |
| HORVU.MOREX.r3.3HG0233990.1 | -0.988127748 | 1.6033308 | -3.411160726 | 0.00124 | 0.043815453 | -1.85735123 | Xyloglucan endotransglucosylase/hydrolase | elongation zone | 6 h |
| HORVU.MOREX.r3.2HG0160130.1 | -0.553722704 | 3.662810319 | -3.331517994 | 0.00158 | 0.049383572 | -1.86644381 | Basic helix-loop-helix transcription factor | elongation zone | 6 h |
| HORVU.MOREX.r3.3HG0287690.1 | -0.784211902 | 2.64681478 | -3.331711403 | 0.00158 | 0.049383572 | -1.86824124 | Profilin | elongation zone | 6 h |
| HORVU.MOREX.r3.3HG0255020.1 | -0.800215572 | 3.485610317 | -3.415129866 | 0.00123 | 0.043520317 | -1.87307188 | Peroxidase | elongation zone | 6 h |
| HORVU.MOREX.r3.2HG0150450.1 | 0.575217754 | 7.245663801 | 3.355032254 | 0.00147 | 0.047574989 | -1.87619477 | Pyruvate kinase | elongation zone | 6 h |
| HORVU.MOREX.r3.2HG0170160.1 | 0.305417895 | 8.212366788 | 3.408198121 | 0.00126 | 0.043818088 | -1.88485399 | Cleft lip and palate transmembrane 1 | elongation zone | 6 h |
| HORVU.MOREX.r3.6HG0624650.1 | 0.330061968 | 8.612307726 | 3.439772286 | 0.00114 | 0.041928054 | -1.89889949 | Transmembrane 9 superfamily member | elongation zone | 6 h |
| HORVU.MOREX.r3.5HG0425110.1 | -0.616496562 | 6.461930984 | -3.348116638 | 0.0015 | 0.048034173 | -1.906502 | Cationic amino acid transporter, putative | elongation zone | 6 h |
| HORVU.MOREX.r3.7HG0661890.1 | 0.490000707 | 7.9160487 | 3.381256402 | 0.00136 | 0.045724589 | -1.92201615 | 3-oxoacyl-[acyl-carrier-protein] synthase | elongation zone | 6 h |
| HORVU.MOREX.r3.1HG0067630.1 | -0.387958686 | 5.282271317 | -3.330596087 | 0.00158 | 0.049443359 | -1.95208936 | Serine/threonine-protein kinase | elongation zone | 6 h |
| HORVU.MOREX.r3.3HG0299220.1 | -0.46464755 | 5.799007839 | -3.353096602 | 0.00148 | 0.047599823 | -1.96161521 | Kinase family protein | elongation zone | 6 h |
| HORVU.MOREX.r3.6HG0608780.1 | 0.432310842 | 7.822442469 | 3.363336781 | 0.00144 | 0.046971179 | -1.96503675 | Heat shock 70 kDa protein, putative | elongation zone | 6 h |
| HORVU.MOREX.r3.5HG0472960.1 | 0.327077857 | 10.61648489 | 3.48385005 | 0.001 | 0.039985373 | -1.98938514 | Phosphoenolpyruvate carboxylase | elongation zone | 6 h |
| HORVU.MOREX.r3.2HG0206470.1 | -0.461365836 | 6.277699834 | -3.336479466 | 0.00156 | 0.049111355 | -1.99955379 | Loricrin-like protein | elongation zone | 6 h |
| HORVU.MOREX.r3.7HG0700270.1 | 0.336032051 | 7.724734938 | 3.34441238 | 0.00152 | 0.04833998 | -2.01023999 | Transaldolase | elongation zone | 6 h |
| HORVU.MOREX.r3.7HG0746770.1 | 0.433597928 | 10.27031467 | 3.35784766 | 0.00146 | 0.047382603 | -2.24500156 | Chaperone protein htpG family protein | elongation zone | 6 h |
| HORVU.MOREX.r3.2HG0134530.1 | 0.499993384 | 9.627861055 | 3.357656866 | 0.00146 | 0.047382603 | -2.26664854 | Brefeldin A-inhibited guanine nucleotide-exchange protein | elongation zone | 6 h |
| HORVU.MOREX.r3.1HG0078900.1 | -2.540241299 | 5.8370464 | -28.86545308 | 4.26E-34 | 8.73E-30 | 65.7525437 | MATH domain-containing protein | elongation zone | 12 h |
| HORVU.MOREX.r3.5HG0447830.1 | -1.596342521 | 4.419279148 | -13.40207822 | 1.17E-18 | 1.20E-14 | 32.05124676 | Sterile alpha motif (SAM) domain-containing protein | elongation zone | 12 h |
| HORVU.MOREX.r3.1HG0070480.1 | -1.337952657 | 6.291080843 | -12.26785472 | 4.12E-17 | 2.82E-13 | 28.72182961 | 60 kDa chaperonin | elongation zone | 12 h |

**Table S5** Continued.

| ID | logFC | AveExpr | t | P.Value | adj.P.Val | B | gene description | root zone | time point |
|---|---|---|---|---|---|---|---|---|---|
| HORVU.MOREX.r3.3HG0280700.1 | 0.85887016 | 6.27769849 | 11.60700024 | 3.55E-16 | 1.82E-12 | 26.5911742 | Katanin p60 ATPase-containing subunit A1 | elongation zone | 12 h |
| HORVU.MOREX.r3.1HG0067040.1 | -1.091220419 | 7.101906906 | -10.16812717 | 4.67E-14 | 1.60E-10 | 21.70255432 | NAC domain protein, | elongation zone | 12 h |
| HORVU.MOREX.r3.6HG0546230.1 | 5.950588965 | -1.679648171 | 10.63420577 | 9.35E-15 | 3.84E-11 | 20.82196479 | Disease resistance protein RPM1 | elongation zone | 12 h |
| HORVU.MOREX.r3.7HG0739910.1 | 1.271815913 | 4.210713917 | 9.222405424 | 1.32E-12 | 3.86E-09 | 18.52866828 | Disease resistance protein (NBS-LRR class) family | elongation zone | 12 h |
| HORVU.MOREX.r3.7HG0729150.1 | -5.401087623 | -2.071842766 | -8.638439118 | 1.08E-11 | 2.47E-08 | 15.34778252 | GDSL esterase/lipase | elongation zone | 12 h |
| HORVU.MOREX.r3.2HG0111680.1 | -5.398840898 | -2.082427636 | -9.151045792 | 1.70E-12 | 4.36E-09 | 14.14453901 | Formate--tetrahydrofolate ligase | elongation zone | 12 h |
| HORVU.MOREX.r3.1HG0067030.1 | -0.856183796 | 5.504670742 | -7.670438386 | 3.76E-10 | 7.02E-07 | 12.82369273 | Protein trichome birefringence | elongation zone | 12 h |
| HORVU.MOREX.r3.7HG0727570.1 | 3.835277904 | -0.569333422 | 7.693927943 | 3.45E-10 | 7.02E-07 | 11.7838467 | ATP-dependent RNA helicase, putative | elongation zone | 12 h |
| HORVU.MOREX.r3.1HG0024020.1 | 1.005817158 | 5.072656773 | 7.319316428 | 1.38E-09 | 2.36E-06 | 11.67339862 | Heparan-alpha-glucosaminide N-acetyltransferase | elongation zone | 12 h |
| HORVU.MOREX.r3.3HG0297940.1 | 2.581895273 | 3.507795484 | 7.160235006 | 2.49E-09 | 3.93E-06 | 11.0374204 | Transcription initiation factor TFIID subunit 9 | elongation zone | 12 h |
| HORVU.MOREX.r3.3HG0298020.1 | 2.037662695 | 3.677613517 | 6.710096779 | 1.32E-08 | 1.93E-05 | 9.568584072 | Transcription initiation factor TFIID subunit 9 | elongation zone | 12 h |
| HORVU.MOREX.r3.1HG0067420.1 | 1.84151765 | 2.43369759 | 6.374136726 | 4.57E-08 | 6.25E-05 | 8.253160805 | Cyclic nucleotide-gated channel | elongation zone | 12 h |
| HORVU.MOREX.r3.7HG0713840.1 | -1.832686304 | -2.185081282 | -6.306367795 | 5.87E-08 | 7.52E-05 | 8.070272429 | Cytochrome P450 | elongation zone | 12 h |
| HORVU.MOREX.r3.7HG0751110.1 | 1.059820126 | 4.358148537 | 6.129681947 | 1.12E-07 | 0.000128142 | 7.451255415 | Kinesin-like | elongation zone | 12 h |
| HORVU.MOREX.r3.5HG0456970.1 | 0.990795929 | 5.8820687 | 6.147814613 | 1.05E-07 | 0.000126931 | 7.440748241 | Methylthioribose-1-phosphate isomerase | elongation zone | 12 h |
| HORVU.MOREX.r3.3HG0307630.1 | 3.377981627 | 2.923305984 | 5.930290449 | 2.34E-07 | 0.000239599 | 6.484933655 | SET-domain protein lysine methyltransferase family protein | elongation zone | 12 h |
| HORVU.MOREX.r3.5HG0526250.1 | 0.489819837 | 12.0381843 | 6.038535948 | 1.57E-07 | 0.000169649 | 6.336778005 | 70 kDa heat shock protein | elongation zone | 12 h |
| HORVU.MOREX.r3.5HG0510830.1 | 1.353050301 | 5.120306759 | 5.795042125 | 3.83E-07 | 0.000341408 | 6.308574636 | #N/A | elongation zone | 12 h |
| HORVU.MOREX.r3.2HG0216110.1 | 0.534201675 | 5.350102781 | 5.823024013 | 3.46E-07 | 0.000327709 | 6.236049967 | #N/A | elongation zone | 12 h |
| HORVU.MOREX.r3.3HG0254260.1 | -1.024300514 | 4.868531812 | -5.818467545 | 3.51E-07 | 0.000327709 | 6.20842364 | Kinesin, putative | elongation zone | 12 h |
| HORVU.MOREX.r3.2HG0109590.1 | -2.395378782 | -0.155486123 | -5.503521966 | 1.10E-06 | 0.000934091 | 5.178525478 | Retrotransposon protein, putative, unclassified | elongation zone | 12 h |
| HORVU.MOREX.r3.7HG0640470.1 | 0.46879748 | 6.260306545 | 5.480749813 | 1.20E-06 | 0.000943673 | 4.912110149 | WPP domain-interacting tail-anchored protein 1 | elongation zone | 12 h |
| HORVU.MOREX.r3.6HG0559680.1 | -3.320513025 | 0.573580493 | -5.437616958 | 1.40E-06 | 0.001023714 | 4.879594033 | NADPH:quinone oxidoreductase | elongation zone | 12 h |
| HORVU.MOREX.r3.4HG0404760.1 | 1.896274552 | 6.564561937 | 5.339261959 | 1.99E-06 | 0.001316292 | 4.67332883 | Tubulin-tyrosine ligase family protein | elongation zone | 12 h |
| HORVU.MOREX.r3.6HG0606940.1 | 3.732215166 | 2.180676744 | 5.470326841 | 1.24E-06 | 0.000943673 | 4.62123451 | Pentatricopeptide repeat-containing protein | elongation zone | 12 h |
| HORVU.MOREX.r3.1HG0069380.1 | -1.371913292 | 2.129642878 | -5.322865985 | 2.11E-06 | 0.001352282 | 4.614832683 | Cysteine proteinase inhibitor | elongation zone | 12 h |
| HORVU.MOREX.r3.2HG0170640.1 | 4.636531871 | -0.509068665 | 5.494507009 | 1.14E-06 | 0.000934091 | 4.393312442 | Protein FAR1-RELATED SEQUENCE 5 | elongation zone | 12 h |
| HORVU.MOREX.r3.1HG0045510.1 | 0.85869774 | 5.275918097 | 5.27932849 | 2.46E-06 | 0.001532253 | 4.392164637 | Prefoldin subunit, putative | elongation zone | 12 h |
| HORVU.MOREX.r3.7HG0724080.1 | 2.194948753 | 2.23197157 | 5.18961423 | 3.39E-06 | 0.001919114 | 4.333751953 | 6,7-dimethyl-8-ribityllumazine synthase | elongation zone | 12 h |
| HORVU.MOREX.r3.3HG0309270.1 | 0.459903889 | 8.53260482 | 5.34519028 | 1.95E-06 | 0.001316292 | 4.252144859 | Proline--tRNA ligase | elongation zone | 12 h |
| HORVU.MOREX.r3.2HG0215830.1 | 1.683128359 | 2.436514446 | 5.133523283 | 4.14E-06 | 0.002178793 | 4.129151422 | CsAtPR5 | elongation zone | 12 h |
| HORVU.MOREX.r3.1HG0075440.1 | -1.139139937 | 1.650312221 | -5.189663251 | 3.39E-06 | 0.001919114 | 4.124086934 | Protein NRT1/ PTR FAMILY 5.5 | elongation zone | 12 h |
| HORVU.MOREX.r3.1HG0068410.1 | -0.489548583 | 4.707522572 | -5.189375642 | 3.40E-06 | 0.001919114 | 4.045607964 | E3 ubiquitin-protein ligase | elongation zone | 12 h |
| HORVU.MOREX.r3.5HG0502990.1 | 5.339440458 | 3.237955153 | 5.083423799 | 4.95E-06 | 0.00247494 | 3.997473504 | NAC domain protein, | elongation zone | 12 h |
| HORVU.MOREX.r3.1HG0067930.1 | -0.991011169 | 2.533288297 | -5.088688699 | 4.85E-06 | 0.00247494 | 3.893418191 | Peroxidase | elongation zone | 12 h |
| HORVU.MOREX.r3.5HG0489790.1 | 2.647632598 | -0.067483895 | 5.184079096 | 3.46E-06 | 0.001919114 | 3.849624559 | SKP1-like protein 4 | elongation zone | 12 h |
| HORVU.MOREX.r3.1HG0050990.1 | 2.93628127 | 2.072399061 | 5.159396263 | 3.78E-06 | 0.002039933 | 3.782246362 | Membrane protein insertase YidC | elongation zone | 12 h |
| HORVU.MOREX.r3.3HG0316280.1 | 1.411227691 | 4.501603233 | 5.044295035 | 5.68E-06 | 0.002774234 | 3.757894752 | Transcription initiation factor TFIID subunit 9 | elongation zone | 12 h |
| HORVU.MOREX.r3.1HG0067490.1 | -3.145796854 | -1.72649178 | -5.376799382 | 1.74E-06 | 0.001229863 | 3.755370642 | Retrovirus-related Pol polyprotein from transposon TNT 1-94 | elongation zone | 12 h |
| HORVU.MOREX.r3.7HG0696190.1 | 0.726630316 | 6.218032617 | 5.011348897 | 6.38E-06 | 0.003043527 | 3.391944882 | Glucose-1-phosphate adenylyltransferase | elongation zone | 12 h |
| HORVU.MOREX.r3.3HG0315690.1 | 0.659686542 | 5.614263673 | 4.971371976 | 7.34E-06 | 0.0034236 | 3.334600351 | E3 ubiquitin-protein ligase BRE1-like 2 | elongation zone | 12 h |

**Table S5** Continued.

| ID | logFC | AveExpr | t | P.Value | adj.P.Val | B | gene description | root zone | time point |
|---|---|---|---|---|---|---|---|---|---|
| HORVU.MOREX.r3.5HG0513490.1 | -0.631143265 | 3.90283549 | -4.931276729 | 8.45E-06 | 0.003650963 | 3.141090749 | Myb transcription factor | elongation zone | 12 h |
| HORVU.MOREX.r3.2HG0205760.1 | 1.24598584 | 5.328219399 | 4.859959287 | 1.08E-05 | 0.004279598 | 3.08080027 | Glutamyl-tRNA(Gln) amidotransferase subunit A | elongation zone | 12 h |
| HORVU.MOREX.r3.1HG0019170.1 | 2.293781213 | 4.041071012 | 4.790280922 | 1.38E-05 | 0.004612029 | 3.041330526 | Replication protein A 70 kDa DNA-binding subunit | elongation zone | 12 h |
| HORVU.MOREX.r3.7HG0717610.1 | -0.830116841 | 4.008622047 | -4.902508367 | 9.35E-06 | 0.003835781 | 3.030989115 | Pollen-specific protein SF21 | elongation zone | 12 h |
| HORVU.MOREX.r3.4HG0337320.1 | 2.78145313 | -0.56622922 | 4.955635496 | 7.76E-06 | 0.003460871 | 3.027816713 | Ribonuclease H | elongation zone | 12 h |
| HORVU.MOREX.r3.3HG0291580.1 | 0.896393635 | 2.805578274 | 4.79582302 | 1.36E-05 | 0.004612029 | 3.009036343 | Lysosomal Pro-X carboxypeptidase | elongation zone | 12 h |
| HORVU.MOREX.r3.2HG0120320.1 | -0.999894142 | 1.685244219 | -4.959201663 | 7.66E-06 | 0.003460871 | 2.983590905 | Peroxidase | elongation zone | 12 h |
| HORVU.MOREX.r3.7HG0655680.1 | -2.39862148 | -1.898894256 | -4.786547179 | 1.40E-05 | 0.004612029 | 2.964654663 | Serine/threonine-protein kinase | elongation zone | 12 h |
| HORVU.MOREX.r3.6HG0550390.1 | 1.883693505 | 2.694299352 | 4.774444594 | 1.46E-05 | 0.004612029 | 2.954586941 | Nitroreductase family protein, putative | elongation zone | 12 h |
| HORVU.MOREX.r3.5HG0471070.1 | -0.368137022 | 6.510820563 | -4.928262656 | 8.54E-06 | 0.003650963 | 2.940632911 | Oxidoreductase family, NAD-binding rossmann fold protein | elongation zone | 12 h |
| HORVU.MOREX.r3.5HG0513740.1 | -1.89889643 | -3.016736504 | -4.782019719 | 1.42E-05 | 0.004612029 | 2.925278368 | Alpha/beta-Hydrolases superfamily protein, putative | elongation zone | 12 h |
| HORVU.MOREX.r3.6HG0628240.1 | 0.939462944 | 6.047595769 | 4.844780568 | 1.14E-05 | 0.004345188 | 2.893522627 | Translation initiation factor eIF-2B subunit epsilon | elongation zone | 12 h |
| HORVU.MOREX.r3.5HG0538060.1 | 3.456849261 | 1.093162331 | 4.902584859 | 9.34E-06 | 0.003835781 | 2.873350663 | Pentatricopeptide repeat-containing protein | elongation zone | 12 h |
| HORVU.MOREX.r3.2HG0193110.1 | 2.023112426 | 1.962681787 | 4.76134155 | 1.53E-05 | 0.004612029 | 2.869355834 | Pentatricopeptide repeat-containing protein | elongation zone | 12 h |
| HORVU.MOREX.r3.1HG0069030.1 | -2.568152033 | 1.480380355 | -4.729127378 | 1.71E-05 | 0.004790957 | 2.840166912 | Calcium-transporting ATPase | elongation zone | 12 h |
| HORVU.MOREX.r3.6HG0551980.1 | 2.213727095 | 0.689536511 | 4.848495075 | 1.13E-05 | 0.004345188 | 2.839319577 | Pentatricopeptide repeat-containing protein | elongation zone | 12 h |
| HORVU.MOREX.r3.6HG0559690.1 | 1.410678913 | 2.590055839 | 4.723247988 | 1.74E-05 | 0.004790957 | 2.807654895 | Cell division cycle 5-like protein | elongation zone | 12 h |
| HORVU.MOREX.r3.5HG0525700.1 | 0.515361168 | 7.295514432 | 4.879946823 | 1.01E-05 | 0.004069304 | 2.79062218 | Leukotriene a-4 hydrolase aminopeptidase | elongation zone | 12 h |
| HORVU.MOREX.r3.5HG0489840.1 | 1.327480706 | 2.789323341 | 4.679336777 | 2.03E-05 | 0.005205195 | 2.69344849 | #N/A | elongation zone | 12 h |
| HORVU.MOREX.r3.7HG0702190.1 | 2.268550826 | 3.843343484 | 4.671747013 | 2.08E-05 | 0.005212813 | 2.661083535 | Fertilization independent endosperm 1 protein | elongation zone | 12 h |
| HORVU.MOREX.r3.2HG0166370.1 | 3.427632951 | 2.267354347 | 4.763054475 | 1.52E-05 | 0.004612029 | 2.630674368 | ATP-binding ABC transporter | elongation zone | 12 h |
| HORVU.MOREX.r3.6HG0612180.1 | 0.812170486 | 6.264662645 | 4.794361092 | 1.36E-05 | 0.004612029 | 2.621485693 | NHL repeat-containing protein-like | elongation zone | 12 h |
| HORVU.MOREX.r3.2HG0178640.1 | 4.143304352 | 2.291364412 | 4.811721353 | 1.28E-05 | 0.004612029 | 2.591941617 | Peroxidase | elongation zone | 12 h |
| HORVU.MOREX.r3.5HG0531850.1 | 0.979608794 | 4.723484706 | 4.695217341 | 1.92E-05 | 0.005054112 | 2.569814967 | #N/A | elongation zone | 12 h |
| HORVU.MOREX.r3.2HG0155210.1 | 0.875539715 | 6.20005446 | 4.747570772 | 1.60E-05 | 0.004767359 | 2.548862812 | #N/A | elongation zone | 12 h |
| HORVU.MOREX.r3.5HG0513810.1 | -0.774180088 | 4.245645455 | -4.76510475 | 1.51E-05 | 0.004612029 | 2.545108289 | Malate dehydrogenase | elongation zone | 12 h |
| HORVU.MOREX.r3.2HG0194450.1 | 0.495017707 | 5.50815323 | 4.74092719 | 1.64E-05 | 0.004790957 | 2.535972093 | Secretory carrier-associated membrane protein | elongation zone | 12 h |
| HORVU.MOREX.r3.2HG0170560.1 | 1.300683704 | 3.704875465 | 4.626651656 | 2.43E-05 | 0.005838526 | 2.516684176 | Ras-like protein | elongation zone | 12 h |
| HORVU.MOREX.r3.5HG0510120.1 | 1.524143941 | 5.694545128 | 4.675006627 | 2.06E-05 | 0.005212813 | 2.490020213 | tRNA (guanine-N(7)-)-methyltransferase non-catalytic subunit | elongation zone | 12 h |
| HORVU.MOREX.r3.5HG0425000.1 | -3.795281094 | -0.918831697 | -4.766804116 | 1.50E-05 | 0.004612029 | 2.47751583 | Transposon protein, putative, Pong sub-class | elongation zone | 12 h |
| HORVU.MOREX.r3.5HG0453710.1 | 0.78968192 | 6.149187887 | 4.732048282 | 1.69E-05 | 0.004790957 | 2.473559635 | 3'(2'),5'-bisphosphate nucleotidase 1 | elongation zone | 12 h |
| HORVU.MOREX.r3.3HG0236260.1 | 2.482622963 | 3.359778756 | 4.624823496 | 2.45E-05 | 0.005838526 | 2.456733322 | Protein IQ-DOMAIN 1 | elongation zone | 12 h |
| HORVU.MOREX.r3.6HG0539520.1 | 0.850672384 | 7.571274396 | 4.769053686 | 1.49E-05 | 0.004612029 | 2.439113139 | tetratricopeptide repeat (TPR)-containing protein | elongation zone | 12 h |
| HORVU.MOREX.r3.7HG0695920.1 | 1.324775331 | 2.660691655 | 4.607088889 | 2.60E-05 | 0.005931341 | 2.43445929 | #N/A | elongation zone | 12 h |
| HORVU.MOREX.r3.6HG0574510.1 | 1.995355805 | 7.321758484 | 4.722050834 | 1.75E-05 | 0.004790957 | 2.43351653 | #N/A | elongation zone | 12 h |
| HORVU.MOREX.r3.5HG0488640.1 | 2.870267469 | 1.861985275 | 4.702638806 | 1.87E-05 | 0.004990255 | 2.418960471 | Protease 2 | elongation zone | 12 h |
| HORVU.MOREX.r3.7HG0750390.1 | 2.012056838 | 1.508596769 | 4.618827285 | 2.50E-05 | 0.005838526 | 2.357979301 | F-box family protein | elongation zone | 12 h |
| HORVU.MOREX.r3.1HG0079800.1 | 0.623931197 | 7.794068 | 4.782954511 | 1.42E-05 | 0.004612029 | 2.347123537 | Farnesyl diphosphate synthase | elongation zone | 12 h |
| HORVU.MOREX.r3.7HG0719070.1 | 0.964232265 | 5.233223615 | 4.632098104 | 2.39E-05 | 0.005838526 | 2.325052807 | SUMO-activating enzyme subunit 1A | elongation zone | 12 h |
| HORVU.MOREX.r3.4HG0372280.1 | 1.357728128 | 3.216376663 | 4.562042287 | 3.04E-05 | 0.006487548 | 2.307807428 | xylulose kinase-2 | elongation zone | 12 h |
| HORVU.MOREX.r3.1HG0079040.1 | 1.622011088 | 2.231395169 | 4.562771861 | 3.03E-05 | 0.006487548 | 2.295564715 | (R)-specific enoyl-CoA hydratase | elongation zone | 12 h |

**Table S5** Continued.

| ID | logFC | AveExpr | t | P.Value | adj.P.Val | B | gene description | root zone | time point |
|---|---|---|---|---|---|---|---|---|---|
| HORVU.MOREX.r3.6HG0550700.1 | 1.949711604 | 1.442253166 | 4.586581605 | 2.79E-05 | 0.006131428 | 2.295526225 | F-box family protein | elongation zone | 12 h |
| HORVU.MOREX.r3.1HG0072580.1 | -0.678852299 | 5.490114394 | -4.685482267 | 1.99E-05 | 0.005160572 | 2.279448692 | Aminotransferase-related family protein | elongation zone | 12 h |
| HORVU.MOREX.r3.1HG0053530.1 | 1.511032401 | 2.523226342 | 4.551577879 | 3.15E-05 | 0.006654271 | 2.26761454 | Glycerol-3-phosphate acyltransferase, chloroplastic | elongation zone | 12 h |
| HORVU.MOREX.r3.7HG0736860.1 | 2.706731769 | 3.347913234 | 4.541494976 | 3.26E-05 | 0.006816924 | 2.236455856 | Aspartyl/glutamyl-tRNA(Asn/Gln) amidotransferase subunit B, pu | elongation zone | 12 h |
| HORVU.MOREX.r3.5HG0441860.1 | 0.657645919 | 9.817417187 | 4.779827325 | 1.43E-05 | 0.004612029 | 2.198779 | 70 kDa heat shock protein | elongation zone | 12 h |
| HORVU.MOREX.r3.5HG0490770.1 | 3.350338224 | 1.191244216 | 4.730529692 | 1.70E-05 | 0.004790957 | 2.185818224 | Inactive rhomboid 1 | elongation zone | 12 h |
| HORVU.MOREX.r3.3HG0220340.1 | 2.161152826 | 2.953247644 | 4.517080222 | 3.54E-05 | 0.007193905 | 2.185490509 | Disease resistance protein RPM1 | elongation zone | 12 h |
| HORVU.MOREX.r3.7HG0701140.1 | 0.536748951 | 7.955915849 | 4.715403956 | 1.79E-05 | 0.004837811 | 2.157177058 | Tyrosine--tRNA ligase | elongation zone | 12 h |
| HORVU.MOREX.r3.3HG0289070.1 | 1.802384306 | 2.310763629 | 4.521803499 | 3.48E-05 | 0.007193905 | 2.152366504 | Protein TIC 22-like, chloroplastic | elongation zone | 12 h |
| HORVU.MOREX.r3.2HG0117010.1 | 1.684308619 | 3.380763648 | 4.494678551 | 3.82E-05 | 0.007270797 | 2.11518682 | #N/A | elongation zone | 12 h |
| HORVU.MOREX.r3.7HG0749460.1 | 0.69582768 | 5.222321052 | 4.618239152 | 2.50E-05 | 0.005838526 | 2.083063357 | Thioredoxin | elongation zone | 12 h |
| HORVU.MOREX.r3.7HG0705220.1 | 0.99333491 | 5.990205142 | 4.587108075 | 2.79E-05 | 0.006131428 | 2.08114599 | Squamous cell carcinoma antigen recognized by T-cells 3 | elongation zone | 12 h |
| HORVU.MOREX.r3.3HG0312510.1 | -1.426682866 | -1.358373367 | -4.50063275 | 3.74E-05 | 0.007270797 | 2.07306892 | cDNA clone:J013058P10, full insert sequence | elongation zone | 12 h |
| HORVU.MOREX.r3.4HG0369090.1 | 1.464715253 | 2.892329497 | 4.474069638 | 4.09E-05 | 0.007570079 | 2.05293318 | 30S ribosomal protein S5 | elongation zone | 12 h |
| HORVU.MOREX.r3.5HG0500650.1 | 1.753345297 | 3.978679874 | 4.488390967 | 3.90E-05 | 0.007343205 | 2.042697766 | Protein MCM10-like protein | elongation zone | 12 h |
| HORVU.MOREX.r3.4HG0418640.1 | 0.831752212 | 4.260686402 | 4.508248921 | 3.65E-05 | 0.007193905 | 1.986747413 | C2 domain-containing protein | elongation zone | 12 h |
| HORVU.MOREX.r3.6HG0564590.1 | 0.632636227 | 6.449269546 | 4.60817632 | 2.59E-05 | 0.005931341 | 1.97543396 | O-fucosyltransferase family protein | elongation zone | 12 h |
| HORVU.MOREX.r3.1HG0080590.1 | 0.956117052 | 6.502287926 | 4.584709135 | 2.81E-05 | 0.006131428 | 1.959890266 | 60 kDa chaperonin | elongation zone | 12 h |
| HORVU.MOREX.r3.1HG0062050.1 | 0.918185403 | 4.541885731 | 4.479074138 | 4.03E-05 | 0.007510284 | 1.940935392 | Tocopherol cyclase | elongation zone | 12 h |
| HORVU.MOREX.r3.3HG0306210.1 | 0.800598064 | 7.280706243 | 4.593898556 | 2.72E-05 | 0.006131428 | 1.898462157 | RAN GTPase activating protein 2 | elongation zone | 12 h |
| HORVU.MOREX.r3.2HG0121270.1 | 2.391501674 | 1.148782547 | 4.50926895 | 3.63E-05 | 0.007193905 | 1.89441405 | DNA helicase | elongation zone | 12 h |
| HORVU.MOREX.r3.5HG0519810.1 | 0.559589673 | 7.61792539 | 4.625851799 | 2.44E-05 | 0.005838526 | 1.846523776 | Diphosphomevalonate decarboxylase | elongation zone | 12 h |
| HORVU.MOREX.r3.7HG0666340.1 | 0.694955925 | 4.728875506 | 4.50867853 | 3.64E-05 | 0.007193905 | 1.832190513 | Hexosyltransferase | elongation zone | 12 h |
| HORVU.MOREX.r3.7HG0634460.1 | 1.755818124 | 3.57799637 | 4.397512789 | 5.30E-05 | 0.008749778 | 1.81717179 | WD repeat-containing protein | elongation zone | 12 h |
| HORVU.MOREX.r3.6HG0607000.1 | 2.375111223 | 0.642043492 | 4.495925972 | 3.80E-05 | 0.007270797 | 1.723583653 | Pectin acetylesterase | elongation zone | 12 h |
| HORVU.MOREX.r3.2HG0196130.1 | 3.364349198 | 1.315960104 | 4.494021162 | 3.83E-05 | 0.007270797 | 1.72351579 | Pentatricopeptide repeat-containing protein | elongation zone | 12 h |
| HORVU.MOREX.r3.2HG0204940.1 | 0.68750571 | 6.058216754 | 4.514502714 | 3.57E-05 | 0.007193905 | 1.717558121 | Crossover junction endonuclease EME1 | elongation zone | 12 h |
| HORVU.MOREX.r3.1HG0084760.1 | -1.726027674 | -3.229685836 | -4.391089299 | 5.42E-05 | 0.008749778 | 1.695446641 | Myb transcription factor | elongation zone | 12 h |
| HORVU.MOREX.r3.6HG0542140.1 | 3.496397909 | 1.647553312 | 4.455551439 | 4.36E-05 | 0.007988512 | 1.695149944 | Pentatricopeptide repeat-containing family protein | elongation zone | 12 h |
| HORVU.MOREX.r3.4HG0414710.1 | 2.292008753 | 1.532728058 | 4.380499213 | 5.62E-05 | 0.008749778 | 1.676296539 | CENP-C | elongation zone | 12 h |
| HORVU.MOREX.r3.5HG0515930.1 | 2.308579427 | 2.608899599 | 4.357265128 | 6.07E-05 | 0.009232427 | 1.641744917 | Pentatricopeptide repeat-containing protein | elongation zone | 12 h |
| HORVU.MOREX.r3.7HG0636630.1 | -1.352768357 | -0.128553353 | -4.426834311 | 4.81E-05 | 0.008500545 | 1.626304839 | WAT1-related protein | elongation zone | 12 h |
| HORVU.MOREX.r3.2HG0098420.1 | 1.673601771 | 5.161437075 | 4.394942065 | 5.35E-05 | 0.008749778 | 1.624062378 | CDT1-like protein a, chloroplastic | elongation zone | 12 h |
| HORVU.MOREX.r3.5HG0455830.1 | 1.089503503 | 3.143860089 | 4.329529237 | 6.67E-05 | 0.00998459 | 1.587883468 | Dynamin family protein, putative, expressed | elongation zone | 12 h |
| HORVU.MOREX.r3.7HG0679060.1 | -0.977683057 | 1.156648697 | -4.405242508 | 5.17E-05 | 0.008691469 | 1.586169175 | Nuclear transport factor 2 family protein, putative | elongation zone | 12 h |
| HORVU.MOREX.r3.4HG0350800.1 | 2.735540074 | 1.296960322 | 4.382179125 | 5.59E-05 | 0.008749778 | 1.584573783 | FBD-associated F-box protein | elongation zone | 12 h |
| HORVU.MOREX.r3.6HG0569480.1 | 3.671851508 | 2.886545793 | 4.381981337 | 5.59E-05 | 0.008749778 | 1.57923168 | Tubby-like F-box protein | elongation zone | 12 h |
| HORVU.MOREX.r3.3HG0231780.1 | 1.215955735 | 5.499283637 | 4.408670257 | 5.11E-05 | 0.008691469 | 1.57665909 | Charged multivesicular body protein 5 | elongation zone | 12 h |
| HORVU.MOREX.r3.2HG0212990.1 | 0.976137725 | 5.148472217 | 4.38699698 | 5.50E-05 | 0.008749778 | 1.57445818 | Acyl-protein thioesterase 1 | elongation zone | 12 h |
| HORVU.MOREX.r3.1HG0091820.1 | 0.830782436 | 5.321522929 | 4.426813448 | 4.81E-05 | 0.008500545 | 1.542499669 | Glutamyl-tRNA (Gln) amidotransferase subunit A | elongation zone | 12 h |
| HORVU.MOREX.r3.4HG0354540.1 | 3.135429099 | 4.811312014 | 4.307915711 | 7.17E-05 | 0.010284717 | 1.538068974 | Receptor-like kinase | elongation zone | 12 h |

**Table S5** Continued.

| ID | logFC | AveExpr | t | P.Value | adj.P.Val | B | gene description | root zone | time point |
|---|---|---|---|---|---|---|---|---|---|
| HORVU.MOREX.r3.5HG0485200.1 | 3.250539611 | 1.510380714 | 4.402872747 | 5.21E-05 | 0.008691469 | 1.5023303 | ATP-dependent zinc metalloprotease FtsH | elongation zone | 12 h |
| HORVU.MOREX.r3.2HG0116440.1 | 1.01636209 | 6.979177665 | 4.43937838 | 4.61E-05 | 0.008363327 | 1.501496415 | Ubiquitin carboxyl-terminal hydrolase | elongation zone | 12 h |
| HORVU.MOREX.r3.6HG0629160.1 | 1.729513557 | 2.999097227 | 4.295569853 | 7.47E-05 | 0.010284717 | 1.501211421 | Disease resistance protein RPM1 | elongation zone | 12 h |
| HORVU.MOREX.r3.3HG0301030.1 | 1.267160968 | 6.005129777 | 4.379922862 | 5.63E-05 | 0.008749778 | 1.461744205 | Nuclear pore complex protein | elongation zone | 12 h |
| HORVU.MOREX.r3.6HG0615710.1 | 0.740825625 | 6.427381525 | 4.435098138 | 4.67E-05 | 0.008410831 | 1.461400198 | Nucleobase ascorbate transporter | elongation zone | 12 h |
| HORVU.MOREX.r3.3HG0231940.1 | 1.898204353 | 2.008606845 | 4.270823453 | 8.11E-05 | 0.011021723 | 1.415280701 | Shikimate kinase family protein | elongation zone | 12 h |
| HORVU.MOREX.r3.6HG0559140.1 | 1.986336 | 0.485291142 | 4.301642401 | 7.32E-05 | 0.010284717 | 1.38375808 | Sn1-specific diacylglycerol lipase alpha | elongation zone | 12 h |
| HORVU.MOREX.r3.1HG0065020.1 | 1.342195953 | 5.184531424 | 4.322233671 | 6.83E-05 | 0.010084309 | 1.380913953 | BZIP transcription factor | elongation zone | 12 h |
| HORVU.MOREX.r3.1HG0079140.1 | -0.693066624 | 4.43581546 | -4.405447208 | 5.16E-05 | 0.008691469 | 1.33134306 | Kinase family protein | elongation zone | 12 h |
| HORVU.MOREX.r3.1HG0079810.1 | 0.637003405 | 3.696599347 | 4.298948608 | 7.38E-05 | 0.010284717 | 1.326134536 | WD repeat protein-like | elongation zone | 12 h |
| HORVU.MOREX.r3.5HG0432550.1 | 1.326517537 | 3.011061408 | 4.24116913 | 8.95E-05 | 0.011688702 | 1.312418621 | Protein ROOT HAIR DEFECTIVE 3 homolog | elongation zone | 12 h |
| HORVU.MOREX.r3.1HG0051190.1 | 0.921638363 | 5.372233168 | 4.334241392 | 6.56E-05 | 0.009900454 | 1.311833874 | COP9 signalosome complex subunit 1 | elongation zone | 12 h |
| HORVU.MOREX.r3.5HG0447870.1 | 1.190871632 | 7.933348257 | 4.416867172 | 4.97E-05 | 0.008641942 | 1.311333453 | Chaperone protein htpG family protein | elongation zone | 12 h |
| HORVU.MOREX.r3.3HG0232890.1 | -2.60709898 | -2.364614764 | -4.297104937 | 7.43E-05 | 0.010284717 | 1.298530058 | NADP-dependent alkenal double bond reductase | elongation zone | 12 h |
| HORVU.MOREX.r3.2HG0114290.1 | 0.398510779 | 6.856362561 | 4.419393446 | 4.93E-05 | 0.008641782 | 1.283958309 | #N/A | elongation zone | 12 h |
| HORVU.MOREX.r3.5HG0443920.1 | 0.642104899 | 3.840341821 | 4.309441591 | 7.13E-05 | 0.010284717 | 1.276854516 | Calmodulin-binding protein, putative, expressed | elongation zone | 12 h |
| HORVU.MOREX.r3.6HG0552860.1 | 1.28060679 | 6.655909595 | 4.359674838 | 6.03E-05 | 0.009226352 | 1.262682419 | Histone H2B | elongation zone | 12 h |
| HORVU.MOREX.r3.6HG0552800.1 | 1.203396933 | 7.010448703 | 4.387620576 | 5.48E-05 | 0.008749778 | 1.258259644 | Histone H2B | elongation zone | 12 h |
| HORVU.MOREX.r3.7HG0749150.1 | 2.336801621 | 3.418594548 | 4.208041678 | 9.99E-05 | 0.012270488 | 1.24590653 | DUF241 domain protein (DUF241) | elongation zone | 12 h |
| HORVU.MOREX.r3.2HG0178430.1 | 1.140746232 | 4.94211731 | 4.271828496 | 8.08E-05 | 0.011021723 | 1.237854657 | RNA binding protein, putative | elongation zone | 12 h |
| HORVU.MOREX.r3.1HG0052120.1 | 0.726326777 | 3.548831498 | 4.245585438 | 8.82E-05 | 0.011603516 | 1.227982636 | GDP-mannose transporter | elongation zone | 12 h |
| HORVU.MOREX.r3.4HG0342950.1 | 1.279335892 | 6.384536094 | 4.32381816 | 6.79E-05 | 0.010084309 | 1.204190614 | 50S ribosomal protein L25 | elongation zone | 12 h |
| HORVU.MOREX.r3.3HG0247250.1 | 1.52225923 | 3.952989106 | 4.220459804 | 9.59E-05 | 0.01214534 | 1.203620212 | V-type proton ATPase subunit E | elongation zone | 12 h |
| HORVU.MOREX.r3.3HG0238570.1 | 1.445357224 | 3.98707821 | 4.197357631 | 0.0001 | 0.012421033 | 1.166756111 | Remodeling and spacing factor 1 | elongation zone | 12 h |
| HORVU.MOREX.r3.7HG0655210.1 | -2.064080552 | -1.689922301 | -4.214144288 | 9.79E-05 | 0.012270488 | 1.142463393 | Peroxidase | elongation zone | 12 h |
| HORVU.MOREX.r3.7HG0664130.1 | 0.53598335 | 5.138266164 | 4.298533007 | 7.39E-05 | 0.010284717 | 1.135134778 | Dipeptidyl peptidase family member 6 | elongation zone | 12 h |
| HORVU.MOREX.r3.1HG0082770.1 | 0.368524786 | 7.702552295 | 4.411369363 | 5.06E-05 | 0.008691469 | 1.12100928 | Reticulon-like protein | elongation zone | 12 h |
| HORVU.MOREX.r3.3HG0329040.1 | -1.88709672 | -0.454710292 | -4.245575901 | 8.82E-05 | 0.011603516 | 1.117758658 | Vacuolar-processing enzyme | elongation zone | 12 h |
| HORVU.MOREX.r3.4HG0343760.1 | 2.624383724 | 1.366971799 | 4.261952425 | 8.35E-05 | 0.011233153 | 1.115721846 | Pentatricopeptide repeat-containing protein | elongation zone | 12 h |
| HORVU.MOREX.r3.6HG0629830.1 | 1.582798338 | 3.458001717 | 4.165922894 | 0.00011 | 0.012462233 | 1.110068276 | Riboflavin biosynthesis protein RibD | elongation zone | 12 h |
| HORVU.MOREX.r3.2HG0122440.1 | 2.902775269 | 2.054200931 | 4.209939475 | 9.93E-05 | 0.012270488 | 1.102772372 | Pentatricopeptide repeat-containing protein | elongation zone | 12 h |
| HORVU.MOREX.r3.4HG0406270.1 | 3.729681347 | 2.424364145 | 4.191873371 | 0.00011 | 0.012430307 | 1.093536158 | RING/FYVE/PHD-type zinc finger family protein | elongation zone | 12 h |
| HORVU.MOREX.r3.2HG0207790.1 | -1.513396937 | -2.594215752 | -4.155398424 | 0.00012 | 0.012462233 | 1.090188244 | Serine/threonine-protein kinase | elongation zone | 12 h |
| HORVU.MOREX.r3.2HG0211900.1 | 1.687710081 | 2.028349871 | 4.156779012 | 0.00012 | 0.012462233 | 1.071436815 | Phytol kinase 1 | elongation zone | 12 h |
| HORVU.MOREX.r3.4HG0372230.1 | -1.986087807 | -1.629808544 | -4.164200917 | 0.00012 | 0.012462233 | 1.067852713 | Transposon protein, putative, CACTA, En/Spm sub-class, expresse | elongation zone | 12 h |
| HORVU.MOREX.r3.2HG0126850.1 | 3.148296787 | 2.517894712 | 4.195570061 | 0.0001 | 0.012422054 | 1.067806823 | RNA-binding protein | elongation zone | 12 h |
| HORVU.MOREX.r3.4HG0334130.1 | 2.371042704 | 2.234967175 | 4.16185419 | 0.00012 | 0.012462233 | 1.065912588 | Thymocyte nuclear protein 1 | elongation zone | 12 h |
| HORVU.MOREX.r3.7HG0634680.1 | 2.623417965 | 5.267359917 | 4.156682163 | 0.00012 | 0.012462233 | 1.043120819 | disease resistance protein (TIR-NBS-LRR class) | elongation zone | 12 h |
| HORVU.MOREX.r3.5HG0518640.1 | 1.37911009 | 2.499571845 | 4.160617535 | 0.00012 | 0.012462233 | 1.037923113 | Methionine S-methyltransferase | elongation zone | 12 h |
| HORVU.MOREX.r3.3HG0244140.1 | 1.316798238 | 5.689186186 | 4.230904961 | 9.26E-05 | 0.011878355 | 1.027798767 | Formate-dependent phosphoribosylglycinamide formyltransfera | elongation zone | 12 h |
| HORVU.MOREX.r3.3HG0320840.1 | 0.372905525 | 7.26652398 | 4.363400925 | 5.95E-05 | 0.009180086 | 1.027503909 | Signal peptide peptidase family protein | elongation zone | 12 h |

**Table S5** Continued.

| ID | logFC | AveExpr | t | P.Value | adj.P.Val | B | gene description | root zone | time point |
|---|---|---|---|---|---|---|---|---|---|
| HORVU.MOREX.r3.3HG0274100.1 | 1.703767433 | 1.731697294 | 4.141845225 | 0.00012 | 0.012462233 | 1.020507371 | 50S ribosomal protein L31 | elongation zone | 12 h |
| HORVU.MOREX.r3.1HG0081540.1 | 2.095482321 | 3.160387272 | 4.131168898 | 0.00013 | 0.012462233 | 1.016515007 | Acid phosphatase/vanadium-dependent haloperoxidase-related | elongation zone | 12 h |
| HORVU.MOREX.r3.6HG0623830.1 | 2.642047754 | 4.072429437 | 4.128436509 | 0.00013 | 0.012462233 | 1.010427748 | Disease resistance protein (NBS-LRR class) family | elongation zone | 12 h |
| HORVU.MOREX.r3.7HG0709860.1 | 0.544776359 | 6.368460733 | 4.304578139 | 7.25E-05 | 0.010284717 | 1.010038994 | Polyadenylate-binding protein | elongation zone | 12 h |
| HORVU.MOREX.r3.3HG0315050.1 | 1.031870078 | 5.388583561 | 4.221540467 | 9.55E-05 | 0.01214534 | 1.007124038 | SET domain containing protein | elongation zone | 12 h |
| HORVU.MOREX.r3.5HG0463310.1 | 2.142007927 | 4.179176168 | 4.134572711 | 0.00013 | 0.012462233 | 0.999401757 | Mitotic spindle checkpoint protein BUBR1 | elongation zone | 12 h |
| HORVU.MOREX.r3.5HG0515020.1 | 1.550591317 | 4.49745777 | 4.15507704 | 0.00012 | 0.012462233 | 0.994012727 | Alpha N-terminal protein methyltransferase 1 | elongation zone | 12 h |
| HORVU.MOREX.r3.5HG0492650.1 | 1.327756263 | 7.121607235 | 4.296643677 | 7.44E-05 | 0.010284717 | 0.989306445 | O-fucosyltransferase family protein | elongation zone | 12 h |
| HORVU.MOREX.r3.6HG0546590.1 | 2.982566107 | 1.264052484 | 4.184121856 | 0.00011 | 0.012462233 | 0.981802876 | Glutamate receptor | elongation zone | 12 h |
| HORVU.MOREX.r3.3HG0318400.1 | 1.092536172 | 6.359365817 | 4.258134665 | 8.46E-05 | 0.011273388 | 0.970371471 | S-formylglutathione hydrolase | elongation zone | 12 h |
| HORVU.MOREX.r3.7HG0722660.1 | -1.052031306 | 1.474969139 | -4.145997163 | 0.00012 | 0.012462233 | 0.958541961 | Diacylglycerol kinase | elongation zone | 12 h |
| HORVU.MOREX.r3.3HG0272430.1 | 0.501110097 | 5.817079707 | 4.297405009 | 7.42E-05 | 0.010284717 | 0.9537935 | Pectin lyase-like superfamily protein | elongation zone | 12 h |
| HORVU.MOREX.r3.3HG0261960.1 | 1.031364808 | 4.490303333 | 4.168088279 | 0.00011 | 0.012462233 | 0.951792287 | ATP-dependent Clp protease proteolytic subunit | elongation zone | 12 h |
| HORVU.MOREX.r3.2HG0111130.1 | 0.99516057 | 3.835075352 | 4.153889908 | 0.00012 | 0.012462233 | 0.948883313 | Short chain dehydrogenase/reductase | elongation zone | 12 h |
| HORVU.MOREX.r3.5HG0431780.1 | 0.982307281 | 5.673780891 | 4.211620936 | 9.87E-05 | 0.012270488 | 0.947400092 | Homocysteine s-methyltransferase | elongation zone | 12 h |
| HORVU.MOREX.r3.5HG0462220.1 | -1.167193216 | 1.070356343 | -4.235962576 | 9.11E-05 | 0.01175404 | 0.918316081 | Calmodulin protein | elongation zone | 12 h |
| HORVU.MOREX.r3.1HG0056980.1 | 2.276631998 | 2.210568558 | 4.108052197 | 0.00014 | 0.013074743 | 0.915391732 | #N/A | elongation zone | 12 h |
| HORVU.MOREX.r3.7HG0719120.1 | 1.123157408 | 6.876683069 | 4.261166421 | 8.38E-05 | 0.011233153 | 0.900272017 | 50S ribosomal protein L1 | elongation zone | 12 h |
| HORVU.MOREX.r3.2HG0106750.1 | 1.452546652 | 0.325037267 | 4.088724303 | 0.00015 | 0.013386102 | 0.882330973 | Alpha-1,4 glucan phosphorylase | elongation zone | 12 h |
| HORVU.MOREX.r3.3HG0265690.1 | 1.849101387 | 0.540508683 | 4.129136259 | 0.00013 | 0.012462233 | 0.881294824 | Oxidoreductase/transition metal ion-binding protein | elongation zone | 12 h |
| HORVU.MOREX.r3.4HG0394830.1 | -1.038348436 | 0.997429304 | -4.166999038 | 0.00011 | 0.012462233 | 0.868168821 | Lecithin-cholesterol acyltransferase-like 1 | elongation zone | 12 h |
| HORVU.MOREX.r3.3HG0268350.1 | 1.076230438 | 2.559660591 | 4.088081906 | 0.00015 | 0.013386102 | 0.867824838 | Exocyst complex component SEC3A | elongation zone | 12 h |
| HORVU.MOREX.r3.4HG0354140.1 | -1.578908935 | -2.197267006 | -4.076833751 | 0.00015 | 0.013607526 | 0.858620091 | CheY-like two-component responsive regulator family protein | elongation zone | 12 h |
| HORVU.MOREX.r3.2HG0173190.1 | 1.780538455 | 7.090068817 | 4.203897447 | 0.0001 | 0.012270488 | 0.844842343 | ABC transporter ATP-binding protein | elongation zone | 12 h |
| HORVU.MOREX.r3.5HG0526320.1 | 1.595123908 | 5.285699775 | 4.125247608 | 0.00013 | 0.012462233 | 0.828624845 | Condensin-2 complex subunit D3 | elongation zone | 12 h |
| HORVU.MOREX.r3.6HG0540200.1 | 0.90587335 | 7.189746969 | 4.239540632 | 9.00E-05 | 0.011688702 | 0.805401952 | 60 kDa chaperonin | elongation zone | 12 h |
| HORVU.MOREX.r3.1HG0024230.1 | 0.705352839 | 5.422255112 | 4.179025214 | 0.00011 | 0.012462233 | 0.770717216 | Tubulin beta chain | elongation zone | 12 h |
| HORVU.MOREX.r3.5HG0437710.1 | 1.153255721 | 2.135740553 | 4.044943812 | 0.00017 | 0.014133454 | 0.761631519 | Protein FAR1-RELATED SEQUENCE 5 | elongation zone | 12 h |
| HORVU.MOREX.r3.3HG0302860.1 | 2.415955213 | 4.061597972 | 4.053418044 | 0.00017 | 0.014078728 | 0.76129776 | Pathogenesis-related thaumatin-like protein | elongation zone | 12 h |
| HORVU.MOREX.r3.1HG0093950.1 | 3.30445539 | -0.165782629 | 4.206236198 | 0.0001 | 0.012270488 | 0.752630371 | Ubiquitin carboxyl-terminal hydrolase 2 | elongation zone | 12 h |
| HORVU.MOREX.r3.3HG0221000.1 | 3.505444984 | 0.305296124 | 4.193394521 | 0.0001 | 0.012430307 | 0.736729495 | Xyloglucan galactosyltransferase KATAMARI1-like protein | elongation zone | 12 h |
| HORVU.MOREX.r3.7HG0647290.1 | 1.606308967 | 4.111332047 | 4.050948207 | 0.00017 | 0.014078728 | 0.729759632 | ATP-dependent Clp protease proteolytic subunit | elongation zone | 12 h |
| HORVU.MOREX.r3.3HG0290660.1 | 0.896981389 | 5.959969367 | 4.174054036 | 0.00011 | 0.012462233 | 0.712759967 | Ribosomal protein S24/S35, mitochondrial | elongation zone | 12 h |
| HORVU.MOREX.r3.6HG0546330.1 | 1.946105904 | 7.259910966 | 4.158473198 | 0.00012 | 0.012462233 | 0.708735779 | GMP synthase [glutamine-hydrolyzing] | elongation zone | 12 h |
| HORVU.MOREX.r3.6HG0610960.1 | 1.066482718 | 3.725371239 | 4.055208355 | 0.00017 | 0.014078728 | 0.702630692 | DNA-directed RNA polymerase II subunit rpb4 | elongation zone | 12 h |
| HORVU.MOREX.r3.3HG0280960.1 | -0.892882973 | 1.80535629 | -4.139160324 | 0.00013 | 0.012462233 | 0.697293869 | Glycosyltransferase | elongation zone | 12 h |
| HORVU.MOREX.r3.1HG0007570.1 | -2.737520062 | -2.580123327 | -4.202815134 | 0.0001 | 0.012270488 | 0.694215041 | Werner Syndrome-like exonuclease | elongation zone | 12 h |
| HORVU.MOREX.r3.5HG0487790.1 | 3.180964661 | 2.300592707 | 4.073039122 | 0.00016 | 0.013659522 | 0.692832291 | Cyclin family protein | elongation zone | 12 h |
| HORVU.MOREX.r3.2HG0105650.1 | 1.309918635 | 6.31756936 | 4.1597246 | 0.00012 | 0.012462233 | 0.691950364 | Cleavage and polyadenylation specificity factor subunit 1 | elongation zone | 12 h |
| HORVU.MOREX.r3.3HG0264580.1 | 1.162985702 | 5.433253416 | 4.12809563 | 0.00013 | 0.012462233 | 0.691495684 | Protein HASTY 1 | elongation zone | 12 h |
| HORVU.MOREX.r3.1HG0090300.1 | 1.263832133 | 5.866767988 | 4.150485178 | 0.00012 | 0.012462233 | 0.689963652 | Topoisomerase 1-associated factor 1 | elongation zone | 12 h |

**Table S5** Continued.

| ID | logFC | AveExpr | t | P.Value | adj.P.Val | B | gene description | root zone | time point |
|---|---|---|---|---|---|---|---|---|---|
| HORVU.MOREX.r3.5HG0440410.1 | 2.757532816 | 1.14377569 | 4.095472538 | 0.00014 | 0.013319392 | 0.685255675 | Pentatricopeptide repeat-containing protein | elongation zone | 12 h |
| HORVU.MOREX.r3.1HG0057040.1 | 1.75610823 | 4.042158307 | 4.035403024 | 0.00018 | 0.014282864 | 0.683309861 | 10 kDa chaperonin | elongation zone | 12 h |
| HORVU.MOREX.r3.2HG0106870.1 | 3.573121745 | 1.977348744 | 4.124076005 | 0.00013 | 0.012462233 | 0.682440269 | Amine oxidase | elongation zone | 12 h |
| HORVU.MOREX.r3.5HG0514360.1 | 2.245961786 | 2.884542848 | 4.015446853 | 0.00019 | 0.014546646 | 0.681118001 | Aspartyl/glutamyl-tRNA(Asn/Gln) amidotransferase subunit B, pu | elongation zone | 12 h |
| HORVU.MOREX.r3.7HG0665250.1 | 2.990950353 | 2.254845156 | 4.030864102 | 0.00018 | 0.014382161 | 0.663372564 | B3 domain-containing protein | elongation zone | 12 h |
| HORVU.MOREX.r3.1HG0049620.1 | 2.851045367 | 0.745244638 | 4.10512282 | 0.00014 | 0.01314062 | 0.661390691 | CRS1 / YhbY (CRM) domain-containing protein | elongation zone | 12 h |
| HORVU.MOREX.r3.1HG0064740.1 | -0.757267995 | 3.101051154 | -4.143530675 | 0.00012 | 0.012462233 | 0.661258042 | Phospholipase A1 | elongation zone | 12 h |
| HORVU.MOREX.r3.3HG0301400.1 | 2.297555535 | 2.640514127 | 4.012030201 | 0.00019 | 0.014582974 | 0.65166826 | Pentatricopeptide repeat-containing protein | elongation zone | 12 h |
| HORVU.MOREX.r3.2HG0173350.1 | 1.179167308 | 5.060871752 | 4.08213258 | 0.00015 | 0.013490162 | 0.642868065 | ATPase ASNA1 | elongation zone | 12 h |
| HORVU.MOREX.r3.6HG0586570.1 | 1.175433995 | 0.923323863 | 4.000259029 | 0.0002 | 0.014672221 | 0.636955991 | Transposon Ty3-G Gag-Pol polyprotein | elongation zone | 12 h |
| HORVU.MOREX.r3.7HG0664400.1 | 1.619263438 | 1.704982407 | 4.002341161 | 0.0002 | 0.014672221 | 0.633690907 | Protein curvature thylakoid chloroplastic-like | elongation zone | 12 h |
| HORVU.MOREX.r3.6HG0566530.1 | 1.566984487 | 5.563778173 | 4.083161556 | 0.00015 | 0.013490162 | 0.626026746 | DNA-directed RNA polymerase subunit | elongation zone | 12 h |
| HORVU.MOREX.r3.3HG0256040.1 | 3.021985858 | 4.80598677 | 3.999939009 | 0.0002 | 0.014672221 | 0.622883431 | DNA gyrase subunit B | elongation zone | 12 h |
| HORVU.MOREX.r3.2HG0204340.1 | 2.629191461 | -0.821479425 | 4.173860679 | 0.00011 | 0.012462233 | 0.611673865 | ERD (early-responsive to dehydration stress) family protein | elongation zone | 12 h |
| HORVU.MOREX.r3.7HG0681830.1 | 1.801603165 | 2.778247346 | 3.991976434 | 0.0002 | 0.014738733 | 0.610161163 | DUF1666 family protein | elongation zone | 12 h |
| HORVU.MOREX.r3.4HG0397510.1 | 2.838349777 | -0.24383738 | 4.163917185 | 0.00012 | 0.012462233 | 0.607051609 | Pentatricopeptide repeat-containing protein | elongation zone | 12 h |
| HORVU.MOREX.r3.3HG0280910.1 | 1.886481541 | 6.11959171 | 4.089693269 | 0.00015 | 0.013386102 | 0.604251116 | Microtubule-associated protein-like | elongation zone | 12 h |
| HORVU.MOREX.r3.5HG0453030.1 | -0.568462346 | 4.568693004 | -4.140433561 | 0.00012 | 0.012462233 | 0.604202656 | P-loop containing nucleoside triphosphate hydrolases superfamil | elongation zone | 12 h |
| HORVU.MOREX.r3.2HG0196960.1 | 2.899581685 | 1.756153365 | 4.035947122 | 0.00018 | 0.014282864 | 0.600427549 | Y' element ATP-dependent helicase YEL077C | elongation zone | 12 h |
| HORVU.MOREX.r3.2HG0121570.1 | 0.59748518 | 4.65733005 | 4.134215372 | 0.00013 | 0.012462233 | 0.598716397 | Protein LTV1 | elongation zone | 12 h |
| HORVU.MOREX.r3.1HG0070730.1 | -1.689617293 | -0.4130893 | -3.990848613 | 0.0002 | 0.014738733 | 0.596657584 | Glycosyltransferase | elongation zone | 12 h |
| HORVU.MOREX.r3.7HG0725060.1 | 2.066410813 | 3.350962126 | 3.985113179 | 0.00021 | 0.014738733 | 0.594174154 | Microtubule associated family protein | elongation zone | 12 h |
| HORVU.MOREX.r3.4HG0396370.1 | 1.025997318 | 7.480547656 | 4.186674514 | 0.00011 | 0.012462233 | 0.59388767 | Methionine--tRNA ligase | elongation zone | 12 h |
| HORVU.MOREX.r3.2HG0190240.1 | 1.57266757 | 2.851074202 | 3.982592424 | 0.00021 | 0.014789061 | 0.586773693 | Smr domain-containing protein, putative | elongation zone | 12 h |
| HORVU.MOREX.r3.1HG0056230.1 | 0.893528829 | 6.035858906 | 4.128785131 | 0.00013 | 0.012462233 | 0.577113195 | 3-hydroxyisobutyryl-CoA hydrolase-like protein | elongation zone | 12 h |
| HORVU.MOREX.r3.7HG0738300.1 | 1.652097342 | 4.034657467 | 4.017353991 | 0.00019 | 0.014546646 | 0.561787932 | Allantoate amidohydrolase | elongation zone | 12 h |
| HORVU.MOREX.r3.1HG0074730.1 | 0.880902142 | 4.444359724 | 4.030852125 | 0.00018 | 0.014382161 | 0.561211183 | Cell division protein ftsZ | elongation zone | 12 h |
| HORVU.MOREX.r3.7HG0685520.1 | 2.273718823 | 0.843971763 | 4.000851375 | 0.0002 | 0.014672221 | 0.556365546 | Pentatricopeptide repeat-containing protein | elongation zone | 12 h |
| HORVU.MOREX.r3.7HG0730510.1 | -1.130110195 | -1.831161525 | -4.046416528 | 0.00017 | 0.014133454 | 0.552971552 | Germin-like protein 1 | elongation zone | 12 h |
| HORVU.MOREX.r3.7HG0720650.1 | 1.214033194 | 6.481386112 | 4.125907375 | 0.00013 | 0.012462233 | 0.551876098 | Elongation factor like protein | elongation zone | 12 h |
| HORVU.MOREX.r3.3HG0287790.1 | 3.401051345 | 1.995670245 | 4.043381228 | 0.00017 | 0.014140313 | 0.532623689 | rRNA methyltransferase | elongation zone | 12 h |
| HORVU.MOREX.r3.5HG0528930.1 | 1.930300707 | 4.365037424 | 3.975204645 | 0.00021 | 0.014889852 | 0.520006393 | GPI-anchored adhesin-like protein | elongation zone | 12 h |
| HORVU.MOREX.r3.7HG0663730.1 | 2.607266074 | 2.179590294 | 3.959292465 | 0.00023 | 0.014924834 | 0.51970011 | Galactoside 2-alpha-L-fucosyltransferase | elongation zone | 12 h |
| HORVU.MOREX.r3.6HG0555600.1 | 2.485035762 | -0.223969439 | 4.010128121 | 0.00019 | 0.014582974 | 0.516379838 | MADS-box transcription factor | elongation zone | 12 h |
| HORVU.MOREX.r3.4HG0345000.1 | 1.053739417 | 3.705245563 | 4.019985102 | 0.00019 | 0.014546646 | 0.515746414 | Kinase | elongation zone | 12 h |
| HORVU.MOREX.r3.3HG0281860.1 | 0.788939478 | 3.606297573 | 4.011062184 | 0.00019 | 0.014582974 | 0.514231466 | Peptidyl-tRNA hydrolase | elongation zone | 12 h |
| HORVU.MOREX.r3.7HG0702450.1 | 3.27209467 | 1.274342504 | 4.053398052 | 0.00017 | 0.014078728 | 0.513769047 | Pentatricopeptide repeat-containing protein | elongation zone | 12 h |
| HORVU.MOREX.r3.4HG0342640.1 | 2.124148368 | 4.773274899 | 3.989876945 | 0.0002 | 0.014738733 | 0.497099001 | Cyclin | elongation zone | 12 h |
| HORVU.MOREX.r3.6HG0585670.1 | 1.094434762 | 4.325989876 | 4.014352627 | 0.00019 | 0.014546646 | 0.494373871 | E2F transcription factor | elongation zone | 12 h |
| HORVU.MOREX.r3.2HG0214090.1 | 1.1209979 | 7.152074623 | 4.134832121 | 0.00013 | 0.012462233 | 0.493385686 | C2 calcium/lipid-binding and GRAM domain protein | elongation zone | 12 h |
| HORVU.MOREX.r3.2HG0141000.1 | 1.36045243 | 5.662430793 | 4.048562308 | 0.00017 | 0.014130693 | 0.49106694 | ATP-dependent RNA helicase DeaD | elongation zone | 12 h |

**Table S5** Continued.

| ID | logFC | AveExpr | t | P.Value | adj.P.Val | B | gene description | root zone | time point |
|---|---|---|---|---|---|---|---|---|---|
| HORVU.MOREX.r3.7HG0662210.1 | 0.638567643 | 8.387753705 | 4.21168385 | 9.87E-05 | 0.012270488 | 0.489415571 | T-complex protein 1 subunit delta | elongation zone | 12 h |
| HORVU.MOREX.r3.2HG0211010.1 | 2.854219095 | 1.796615013 | 3.984713952 | 0.00021 | 0.014738733 | 0.4838759 | Pentatricopeptide repeat-containing protein | elongation zone | 12 h |
| HORVU.MOREX.r3.7HG0724610.1 | 1.215371359 | 3.83877381 | 3.965832278 | 0.00022 | 0.014924834 | 0.477252974 | Ribose-phosphate pyrophosphokinase | elongation zone | 12 h |
| HORVU.MOREX.r3.2HG0197180.2 | 1.331193706 | 1.736102374 | 3.944402677 | 0.00024 | 0.015309918 | 0.477057857 | Exocyst complex component SEC15A | elongation zone | 12 h |
| HORVU.MOREX.r3.2HG0141200.1 | 1.36562103 | 5.885442827 | 4.056797663 | 0.00016 | 0.014078728 | 0.458410769 | Endoribonuclease L-PSP family protein | elongation zone | 12 h |
| HORVU.MOREX.r3.3HG0283730.1 | 1.002996977 | 8.227085107 | 4.17471352 | 0.00011 | 0.012462233 | 0.453158063 | 60S ribosomal protein l28 | elongation zone | 12 h |
| HORVU.MOREX.r3.5HG0527050.1 | 1.761539039 | 2.801057467 | 3.933837846 | 0.00024 | 0.015587775 | 0.44747613 | Mitochondrial carrier protein, putative | elongation zone | 12 h |
| HORVU.MOREX.r3.1HG0038800.1 | -0.694589742 | 2.879103431 | -4.066640726 | 0.00016 | 0.013829713 | 0.443357093 | Protein kinase | elongation zone | 12 h |
| HORVU.MOREX.r3.5HG0509060.1 | -0.621564372 | 4.702327944 | -4.141901922 | 0.00012 | 0.012462233 | 0.433591767 | Non-specific serine/threonine protein kinase | elongation zone | 12 h |
| HORVU.MOREX.r3.1HG0036930.1 | 1.216360965 | 6.161597548 | 4.079416764 | 0.00015 | 0.013551547 | 0.427634346 | Histone H2A | elongation zone | 12 h |
| HORVU.MOREX.r3.4HG0354980.1 | -0.769667545 | 2.412831318 | -4.035410208 | 0.00018 | 0.014282864 | 0.422315108 | #N/A | elongation zone | 12 h |
| HORVU.MOREX.r3.2HG0113740.1 | 2.339588086 | 1.01695296 | 3.994167326 | 0.0002 | 0.014738733 | 0.418320764 | Metallothiol transferase FosB | elongation zone | 12 h |
| HORVU.MOREX.r3.7HG0728280.1 | 2.235417335 | 1.355492103 | 3.955944486 | 0.00023 | 0.014924834 | 0.416262186 | Pentatricopeptide repeat-containing protein | elongation zone | 12 h |
| HORVU.MOREX.r3.5HG0485840.1 | 2.110059694 | 1.087632654 | 3.964021133 | 0.00022 | 0.014924834 | 0.415429099 | DUF1644 family protein | elongation zone | 12 h |
| HORVU.MOREX.r3.1HG0069870.1 | -0.876101801 | 2.55224648 | -4.066663361 | 0.00016 | 0.013829713 | 0.415316576 | #N/A | elongation zone | 12 h |
| HORVU.MOREX.r3.4HG0402070.1 | -0.601716247 | 3.388938852 | -4.127388282 | 0.00013 | 0.012462233 | 0.404636846 | IQ domain-containing protein | elongation zone | 12 h |
| HORVU.MOREX.r3.7HG0660840.1 | 1.851344092 | 2.942614733 | 3.917741598 | 0.00026 | 0.015692177 | 0.39592039 | Pectinesterase | elongation zone | 12 h |
| HORVU.MOREX.r3.3HG0224260.1 | 1.287856406 | 3.431079477 | 3.923531615 | 0.00025 | 0.015692177 | 0.378660523 | ABC transporter G family member | elongation zone | 12 h |
| HORVU.MOREX.r3.6HG0569490.1 | 0.603495331 | 6.255599873 | 4.095496917 | 0.00014 | 0.013319392 | 0.373953699 | Rad23 UV excision repair protein family | elongation zone | 12 h |
| HORVU.MOREX.r3.5HG0529630.1 | 5.015427678 | 8.94083134 | 4.003432086 | 0.0002 | 0.014672221 | 0.37067545 | Lipoxygenase | elongation zone | 12 h |
| HORVU.MOREX.r3.7HG0728120.1 | 1.723432037 | 1.131059546 | 3.913186113 | 0.00026 | 0.015692177 | 0.368207032 | Macrophage migration inhibitory factor | elongation zone | 12 h |
| HORVU.MOREX.r3.1HG0085960.1 | 0.654406567 | 5.291265231 | 4.028253021 | 0.00018 | 0.014447555 | 0.355361994 | ARM repeat superfamily protein | elongation zone | 12 h |
| HORVU.MOREX.r3.5HG0526940.1 | 0.467894156 | 7.556680019 | 4.157312039 | 0.00012 | 0.012462233 | 0.3552297 | Alpha-mannosidase | elongation zone | 12 h |
| HORVU.MOREX.r3.3HG0231730.1 | 2.263734712 | 2.401658147 | 3.908868536 | 0.00027 | 0.015692177 | 0.345585742 | Receptor-like protein kinase | elongation zone | 12 h |
| HORVU.MOREX.r3.2HG0183320.1 | 2.064327138 | 2.199260923 | 3.900691152 | 0.00027 | 0.015692177 | 0.34290641 | mRNA cap guanine-N7 methyltransferase | elongation zone | 12 h |
| HORVU.MOREX.r3.1HG0086360.1 | 1.732565978 | 4.548392115 | 3.919366556 | 0.00026 | 0.015692177 | 0.299628965 | Ribonuclease PH | elongation zone | 12 h |
| HORVU.MOREX.r3.2HG0111790.1 | 2.186526824 | 4.937039867 | 3.887145418 | 0.00028 | 0.015800471 | 0.292911719 | Receptor-like kinase | elongation zone | 12 h |
| HORVU.MOREX.r3.2HG0191230.1 | 1.845140313 | 3.939439112 | 3.890390937 | 0.00028 | 0.015756066 | 0.292486212 | Pentatricopeptide repeat-containing protein | elongation zone | 12 h |
| HORVU.MOREX.r3.1HG0061220.1 | 1.485404067 | 4.835260837 | 3.944287818 | 0.00024 | 0.015309918 | 0.292015443 | DNA primase | elongation zone | 12 h |
| HORVU.MOREX.r3.1HG0085840.1 | 2.885479614 | 1.255424894 | 3.960029088 | 0.00022 | 0.014924834 | 0.291688295 | Maturase-like protein | elongation zone | 12 h |
| HORVU.MOREX.r3.7HG0665640.1 | 1.65993869 | 4.705582024 | 3.908548103 | 0.00027 | 0.015692177 | 0.286678838 | electron protein, putative (Protein of unknown function, DUF547 | elongation zone | 12 h |
| HORVU.MOREX.r3.2HG0167500.2 | 1.993793502 | 4.031521806 | 3.898746046 | 0.00027 | 0.015692177 | 0.281536426 | Repetitive proline-rich cell wall protein | elongation zone | 12 h |
| HORVU.MOREX.r3.1HG0033310.1 | 1.031709046 | 5.652380892 | 4.003540798 | 0.0002 | 0.014672221 | 0.27431727 | DNA mismatch repair protein mutS | elongation zone | 12 h |
| HORVU.MOREX.r3.3HG0322660.1 | 1.497013775 | 5.027667147 | 3.961619121 | 0.00022 | 0.014924834 | 0.271695603 | Receptor protein kinase, putative | elongation zone | 12 h |
| HORVU.MOREX.r3.2HG0184630.1 | 2.226495219 | 2.780348233 | 3.874868521 | 0.0003 | 0.016177895 | 0.268543102 | Elongation factor G | elongation zone | 12 h |
| HORVU.MOREX.r3.6HG0560290.1 | 2.278392923 | -0.187239249 | 3.927949231 | 0.00025 | 0.015638414 | 0.259546777 | Cyclin | elongation zone | 12 h |
| HORVU.MOREX.r3.5HG0460720.1 | 0.588358277 | 6.55235057 | 4.073672635 | 0.00016 | 0.013659522 | 0.250596267 | L-galactono-1,4-lactone dehydrogenase | elongation zone | 12 h |
| HORVU.MOREX.r3.2HG0165840.1 | 0.712182054 | 6.204491334 | 4.044765985 | 0.00017 | 0.014133454 | 0.2454484 | Protein kinase | elongation zone | 12 h |
| HORVU.MOREX.r3.4HG0354780.1 | 1.406092835 | 5.895136905 | 3.976681278 | 0.00021 | 0.014881142 | 0.244867087 | Proline-, glutamic acid-and leucine-rich protein 1 | elongation zone | 12 h |
| HORVU.MOREX.r3.4HG0405600.1 | 0.63958473 | 7.101161055 | 4.087170778 | 0.00015 | 0.013386102 | 0.238037687 | Eukaryotic translation initiation factor 3 subunit K | elongation zone | 12 h |
| HORVU.MOREX.r3.1HG0020820.1 | 2.113618289 | 3.048519021 | 3.858988051 | 0.00031 | 0.016585421 | 0.235307214 | Serine protease | elongation zone | 12 h |

**Table S5** Continued.

| ID | logFC | AveExpr | t | P.Value | adj.P.Val | B | gene description | root zone | time point |
|---|---|---|---|---|---|---|---|---|---|
| HORVU.MOREX.r3.3HG0304280.1 | 2.564599159 | 2.903876701 | 3.866085627 | 0.0003 | 0.016454463 | 0.230102946 | Exocyst complex component, putative | elongation zone | 12 h |
| HORVU.MOREX.r3.7HG0716100.1 | 2.550954856 | 0.460823771 | 3.907599331 | 0.00027 | 0.015692177 | 0.218408403 | magnesium-protoporphyrin IX methyltransferase | elongation zone | 12 h |
| HORVU.MOREX.r3.7HG0648600.1 | 0.804735658 | 5.91516897 | 4.019189025 | 0.00019 | 0.014546646 | 0.213149344 | Elongation factor 4 | elongation zone | 12 h |
| HORVU.MOREX.r3.5HG0486060.1 | 3.712550108 | 1.638161843 | 3.936173228 | 0.00024 | 0.015535305 | 0.21181594 | Protease 2 | elongation zone | 12 h |
| HORVU.MOREX.r3.4HG0416290.1 | 1.760615777 | 3.543238154 | 3.852268775 | 0.00032 | 0.016710658 | 0.204801419 | Sphingomyelin phosphodiesterase 4 | elongation zone | 12 h |
| HORVU.MOREX.r3.2HG0173340.1 | 3.30603414 | 1.078294475 | 3.980028468 | 0.00021 | 0.014824212 | 0.196086443 | Pentatricopeptide repeat-containing protein, putative | elongation zone | 12 h |
| HORVU.MOREX.r3.1HG0073670.1 | 2.981109236 | -0.381423856 | 3.879796169 | 0.00029 | 0.016089033 | 0.193654181 | Aldo-keto reductase | elongation zone | 12 h |
| HORVU.MOREX.r3.1HG0060080.1 | 2.28538379 | 1.496119533 | 3.853469367 | 0.00032 | 0.016689644 | 0.189903092 | Pentatricopeptide repeat-containing protein | elongation zone | 12 h |
| HORVU.MOREX.r3.7HG0704310.1 | 2.545096699 | 3.178531848 | 3.842902668 | 0.00033 | 0.016953892 | 0.18934172 | Receptor-like protein kinase | elongation zone | 12 h |
| HORVU.MOREX.r3.4HG0381360.1 | 1.349447717 | 6.018407731 | 3.97075027 | 0.00022 | 0.014924834 | 0.188206109 | Adenylosuccinate lyase | elongation zone | 12 h |
| HORVU.MOREX.r3.2HG0185190.1 | 1.031435224 | 3.839589672 | 3.891062602 | 0.00028 | 0.015756066 | 0.187209911 | Yellow stripe-like 9 transporter | elongation zone | 12 h |
| HORVU.MOREX.r3.6HG0583670.1 | 0.644399157 | 4.442077678 | 3.95938608 | 0.00023 | 0.014924834 | 0.185310934 | WD repeat protein | elongation zone | 12 h |
| HORVU.MOREX.r3.3HG0263410.1 | 1.21577542 | 4.597647088 | 3.899610073 | 0.00027 | 0.015692177 | 0.162172134 | Bromo-adjacent-like (BAH) domain protein | elongation zone | 12 h |
| HORVU.MOREX.r3.7HG0678230.1 | 0.646995834 | 6.137580535 | 4.022528808 | 0.00018 | 0.014546646 | 0.158155308 | Eukaryotic translation initiation factor 3 subunit F | elongation zone | 12 h |
| HORVU.MOREX.r3.6HG0616960.1 | 0.751845658 | 4.492256721 | 3.937999013 | 0.00024 | 0.015525106 | 0.157449453 | Polyadenylate-binding protein family protein | elongation zone | 12 h |
| HORVU.MOREX.r3.5HG0514790.1 | 0.538341919 | 9.909631182 | 4.175373618 | 0.00011 | 0.012462233 | 0.15732349 | Elongation factor 1-alpha | elongation zone | 12 h |
| HORVU.MOREX.r3.3HG0329300.1 | 1.620498345 | 4.353107466 | 3.861336588 | 0.00031 | 0.016574489 | 0.156678179 | Dual-specificity RNA methyltransferase RlmN | elongation zone | 12 h |
| HORVU.MOREX.r3.3HG0303970.1 | 0.739326708 | 5.516060256 | 3.999208334 | 0.0002 | 0.014672221 | 0.150266923 | Internal alternative NAD(P)H-ubiquinone oxidoreductase A1, mit | elongation zone | 12 h |
| HORVU.MOREX.r3.5HG0448120.1 | 1.587689475 | 1.313003386 | 3.8280143 | 0.00034 | 0.017220216 | 0.14618403 | Polyol transporter | elongation zone | 12 h |
| HORVU.MOREX.r3.2HG0195640.1 | 0.73376492 | 4.790343179 | 3.9407358 | 0.00024 | 0.015437333 | 0.14442023 | Imidazoleglycerol-phosphate dehydratase | elongation zone | 12 h |
| HORVU.MOREX.r3.3HG0228790.1 | 1.323301162 | 7.11644995 | 4.014715227 | 0.00019 | 0.014546646 | 0.142473385 | Mitochondrial glycoprotein | elongation zone | 12 h |
| HORVU.MOREX.r3.5HG0420980.1 | -0.846719469 | 2.046048282 | -3.92111939 | 0.00025 | 0.015692177 | 0.141177259 | ERD (Early-responsive to dehydration stress) family protein | elongation zone | 12 h |
| HORVU.MOREX.r3.7HG0644560.1 | 1.303103331 | 3.358673269 | 3.841832776 | 0.00033 | 0.016953892 | 0.140921191 | Cell cycle regulated microtubule associated protein | elongation zone | 12 h |
| HORVU.MOREX.r3.1HG0039730.1 | 0.832354754 | 5.716966668 | 3.953311465 | 0.00023 | 0.014965579 | 0.128273391 | Glutamine synthetase | elongation zone | 12 h |
| HORVU.MOREX.r3.4HG0404740.1 | -0.805475298 | 2.889075204 | -4.041634671 | 0.00017 | 0.014164068 | 0.127797275 | Hydrolase, alpha/beta fold family protein, expressed | elongation zone | 12 h |
| HORVU.MOREX.r3.1HG0012330.1 | 1.034950725 | 6.207327614 | 3.98762397 | 0.00021 | 0.014738733 | 0.126686707 | Peroxisomal membrane 22 kDa (Mpv17/PMP22) family protein | elongation zone | 12 h |
| HORVU.MOREX.r3.3HG0218560.1 | -1.177863642 | 0.309747036 | -3.922919815 | 0.00025 | 0.015692177 | 0.126065287 | Receptor-like protein kinase | elongation zone | 12 h |
| HORVU.MOREX.r3.6HG0548350.1 | -1.271959689 | 1.250846794 | -3.827107967 | 0.00034 | 0.017220216 | 0.122929005 | Jacalin-related lectin | elongation zone | 12 h |
| HORVU.MOREX.r3.5HG0439710.1 | 2.52221901 | -0.510687428 | 3.907245313 | 0.00027 | 0.015692177 | 0.122368224 | Zinc finger protein | elongation zone | 12 h |
| HORVU.MOREX.r3.4HG0366210.2 | 2.950360036 | 1.77805932 | 3.858676987 | 0.00031 | 0.016585421 | 0.118805445 | Agenet domain containing protein | elongation zone | 12 h |
| HORVU.MOREX.r3.3HG0305500.1 | 1.221664499 | 2.378582061 | 3.825237191 | 0.00035 | 0.01726164 | 0.116050851 | 50S ribosomal protein-related, putative | elongation zone | 12 h |
| HORVU.MOREX.r3.5HG0508890.1 | 3.176031881 | 0.546502989 | 3.926140707 | 0.00025 | 0.015638414 | 0.115290497 | Pentatricopeptide repeat-containing protein | elongation zone | 12 h |
| HORVU.MOREX.r3.7HG0727460.1 | 0.953900379 | 8.013727164 | 4.056161393 | 0.00016 | 0.014078728 | 0.099655865 | 60S acidic ribosomal protein P3 | elongation zone | 12 h |
| HORVU.MOREX.r3.5HG0487850.1 | 1.657061676 | 4.243040388 | 3.835553602 | 0.00033 | 0.017055324 | 0.096485334 | Phosphoribosylaminoimidazole-succinocarboxamide synthase | elongation zone | 12 h |
| HORVU.MOREX.r3.6HG0628050.1 | 2.321577804 | 7.370628441 | 3.955284516 | 0.00023 | 0.014924834 | 0.096189441 | Cytosine-specific methyltransferase | elongation zone | 12 h |
| HORVU.MOREX.r3.1HG0008380.1 | 2.127712065 | 2.770842782 | 3.810393631 | 0.00036 | 0.017580172 | 0.094616763 | methyl-coenzyme M reductase II subunit gamma, putative (DUF3 | elongation zone | 12 h |
| HORVU.MOREX.r3.2HG0122060.1 | 0.691827666 | 5.958624172 | 3.987741712 | 0.00021 | 0.014738733 | 0.09001685 | #N/A | elongation zone | 12 h |
| HORVU.MOREX.r3.5HG0477040.1 | 0.464050238 | 5.558139817 | 3.990802462 | 0.0002 | 0.014738733 | 0.08118903 | MND1-interacting protein 1 | elongation zone | 12 h |
| HORVU.MOREX.r3.6HG0590190.1 | 0.732725588 | 4.321131534 | 3.88217705 | 0.00029 | 0.016010182 | 0.074687373 | #N/A | elongation zone | 12 h |
| HORVU.MOREX.r3.3HG0307120.1 | 0.586916242 | 8.36463409 | 4.10050642 | 0.00014 | 0.013220171 | 0.073661056 | S-adenosyl-L-methionine-dependent methyltransferases superfa | elongation zone | 12 h |
| HORVU.MOREX.r3.3HG0256300.1 | 0.629949286 | 7.085102404 | 4.023383297 | 0.00018 | 0.014546646 | 0.073448068 | Glutamate--tRNA ligase | elongation zone | 12 h |

**Table S5** Continued.

| ID | logFC | AveExpr | t | P.Value | adj.P.Val | B | gene description | root zone | time point |
|---|---|---|---|---|---|---|---|---|---|
| HORVU.MOREX.r3.5HG0534640.1 | 0.693334145 | 8.816317071 | 4.089169467 | 0.00015 | 0.013386102 | 0.063185292 | Elongation factor Tu | elongation zone | 12 h |
| HORVU.MOREX.r3.6HG0540620.1 | 1.909573227 | 4.022997546 | 3.815005313 | 0.00036 | 0.017468576 | 0.057654537 | Malignant brain tumor repeat protein 1 | elongation zone | 12 h |
| HORVU.MOREX.r3.5HG0527750.1 | 0.643499104 | 5.066219563 | 3.926153742 | 0.00025 | 0.015638414 | 0.05756769 | Kinetochore protein NDC80 | elongation zone | 12 h |
| HORVU.MOREX.r3.4HG0407860.1 | -3.4950011 | -1.046602776 | -3.794006546 | 0.00038 | 0.017696476 | 0.054127743 | Cytochrome P450, putative | elongation zone | 12 h |
| HORVU.MOREX.r3.6HG0541240.1 | 1.935447584 | 3.029974348 | 3.794454298 | 0.00038 | 0.017696476 | 0.053139255 | Pentatricopeptide repeat-containing protein | elongation zone | 12 h |
| HORVU.MOREX.r3.2HG0199560.1 | 1.752865944 | 5.029113972 | 3.910702542 | 0.00026 | 0.015692177 | 0.048198084 | Organic cation transporter-like protein | elongation zone | 12 h |
| HORVU.MOREX.r3.7HG0687220.1 | -1.437481623 | -1.361787113 | -3.834512678 | 0.00034 | 0.017059301 | 0.04626063 | Cytochrome P450 | elongation zone | 12 h |
| HORVU.MOREX.r3.6HG0578170.1 | 0.908877578 | 6.59140041 | 3.979733215 | 0.00021 | 0.014824212 | 0.044541824 | Histone deacetylase | elongation zone | 12 h |
| HORVU.MOREX.r3.5HG0528100.1 | 2.000493752 | 1.953644846 | 3.795614441 | 0.00038 | 0.017696476 | 0.044112405 | 50S ribosomal protein L18 | elongation zone | 12 h |
| HORVU.MOREX.r3.3HG0328480.1 | 1.861367417 | 1.933391674 | 3.79067411 | 0.00039 | 0.017696476 | 0.040876071 | Rotundifolia-like protein | elongation zone | 12 h |
| HORVU.MOREX.r3.2HG0121290.1 | 1.522528007 | 2.766171938 | 3.792687356 | 0.00038 | 0.017696476 | 0.037697827 | DNA/RNA helicase protein | elongation zone | 12 h |
| HORVU.MOREX.r3.5HG0514980.1 | 1.767692517 | 7.445624912 | 3.961651264 | 0.00022 | 0.014924834 | 0.036857073 | Inosine-5'-monophosphate dehydrogenase | elongation zone | 12 h |
| HORVU.MOREX.r3.4HG0345400.1 | 1.449205441 | 6.016920639 | 3.900559522 | 0.00027 | 0.015692177 | 0.026885518 | DAG protein, chloroplastic | elongation zone | 12 h |
| HORVU.MOREX.r3.5HG0528920.1 | 1.497277525 | 6.19180472 | 3.908618796 | 0.00027 | 0.015692177 | 0.018279211 | Phosphoribosylformylglycinamidine cyclo-ligase | elongation zone | 12 h |
| HORVU.MOREX.r3.2HG0198550.1 | 3.043469204 | 0.396712397 | 3.888873026 | 0.00028 | 0.015756066 | 0.015855512 | Pentatricopeptide repeat-containing protein | elongation zone | 12 h |
| HORVU.MOREX.r3.7HG0674720.1 | -1.721705558 | -1.463275625 | -3.818964056 | 0.00035 | 0.017458382 | 0.014867266 | Epoxide hydrolase 2 | elongation zone | 12 h |
| HORVU.MOREX.r3.1HG0064610.1 | 1.057723138 | 5.770306439 | 3.918724558 | 0.00026 | 0.015692177 | 0.008106542 | membrane-associated kinase regulator | elongation zone | 12 h |
| HORVU.MOREX.r3.4HG0369200.1 | 0.790980446 | 4.224154659 | 3.858934594 | 0.00031 | 0.016585421 | 0.000952895 | RNA-binding protein Nova | elongation zone | 12 h |
| HORVU.MOREX.r3.1HG0015760.1 | 1.936266237 | 4.576039572 | 3.797523707 | 0.00038 | 0.017696476 | -0.00814129 | Regulator of chromosome condensation (RCC1) family protein | elongation zone | 12 h |
| HORVU.MOREX.r3.4HG0386280.1 | 0.683923146 | 6.657437418 | 3.986663577 | 0.00021 | 0.014738733 | -0.01154922 | #N/A | elongation zone | 12 h |
| HORVU.MOREX.r3.5HG0490180.1 | 0.699261414 | 11.07313729 | 4.143585582 | 0.00012 | 0.012462233 | -0.0121694 | Heat shock protein 90 | elongation zone | 12 h |
| HORVU.MOREX.r3.1HG0058260.1 | 1.846807755 | 2.383540747 | 3.770802674 | 0.00041 | 0.017914792 | -0.01487359 | 50S ribosomal protein L18 | elongation zone | 12 h |
| HORVU.MOREX.r3.6HG0555830.1 | 1.090011624 | 5.364563421 | 3.893068357 | 0.00028 | 0.015751512 | -0.0148904 | Aspartyl/glutamyl-tRNA(Asn/Gln) amidotransferase subunit B | elongation zone | 12 h |
| HORVU.MOREX.r3.4HG0357750.1 | 0.412432361 | 9.218441158 | 4.100637669 | 0.00014 | 0.013220171 | -0.01660595 | Alanine--tRNA ligase | elongation zone | 12 h |
| HORVU.MOREX.r3.6HG0540280.1 | 0.979932544 | 5.482684853 | 3.898767998 | 0.00027 | 0.015692177 | -0.01661585 | F-box family protein | elongation zone | 12 h |
| HORVU.MOREX.r3.1HG0011880.1 | -0.939593396 | 0.780281969 | -3.959492059 | 0.00023 | 0.014924834 | -0.01693149 | Voltage-dependent T-type calcium channel subunit alpha-1G | elongation zone | 12 h |
| HORVU.MOREX.r3.5HG0522760.1 | 3.213034747 | 0.069285095 | 3.870176349 | 0.0003 | 0.016327144 | -0.01886347 | imidazolonepropionase (Protein of unknown function, DUF642) | elongation zone | 12 h |
| HORVU.MOREX.r3.7HG0669580.1 | -0.707492592 | 1.55116404 | -3.993062076 | 0.0002 | 0.014738733 | -0.022011 | Caleosin | elongation zone | 12 h |
| HORVU.MOREX.r3.3HG0310680.1 | 0.757497523 | 7.418942266 | 4.005623884 | 0.00019 | 0.014672221 | -0.02580974 | Mitochondrial import receptor subunit TOM20 | elongation zone | 12 h |
| HORVU.MOREX.r3.4HG0342160.1 | 0.472826079 | 5.308472026 | 3.957640712 | 0.00023 | 0.014924834 | -0.02924309 | Presenilin | elongation zone | 12 h |
| HORVU.MOREX.r3.4HG0346720.1 | 0.553394526 | 10.60860825 | 4.151088987 | 0.00012 | 0.012462233 | -0.03028004 | Protein disulfide-isomerase | elongation zone | 12 h |
| HORVU.MOREX.r3.4HG0413690.1 | 0.579686778 | 4.40336351 | 3.905640063 | 0.00027 | 0.015692177 | -0.03428482 | 26S proteasome non-ATPase regulatory subunit-like protein | elongation zone | 12 h |
| HORVU.MOREX.r3.5HG0494740.1 | 1.43924588 | 7.598359813 | 3.956615456 | 0.00023 | 0.014924834 | -0.03488462 | DAG protein, chloroplastic | elongation zone | 12 h |
| HORVU.MOREX.r3.1HG0067410.1 | 0.436932936 | 5.64638285 | 3.962454563 | 0.00022 | 0.014924834 | -0.03500155 | 1-acyl-sn-glycerol-3-phosphate acyltransferase | elongation zone | 12 h |
| HORVU.MOREX.r3.3HG0298120.1 | 1.338941445 | 3.348184436 | 3.771819366 | 0.00041 | 0.017914792 | -0.04138534 | BTB/POZ domain containing protein | elongation zone | 12 h |
| HORVU.MOREX.r3.1HG0077880.1 | 1.055812612 | 3.552074142 | 3.805976685 | 0.00037 | 0.017598677 | -0.04263445 | BAG family molecular chaperone regulator 7 | elongation zone | 12 h |
| HORVU.MOREX.r3.4HG0338340.1 | 1.827924217 | 3.481593478 | 3.768939688 | 0.00041 | 0.017948348 | -0.04736242 | GRAS transcription factor | elongation zone | 12 h |
| HORVU.MOREX.r3.5HG0439640.1 | 1.002119824 | 3.420116187 | 3.800622124 | 0.00037 | 0.017696476 | -0.05034008 | Expansin protein | elongation zone | 12 h |
| HORVU.MOREX.r3.6HG0543100.1 | 3.124336468 | 1.312256801 | 3.824819411 | 0.00035 | 0.01726164 | -0.05258822 | Pentatricopeptide repeat-containing protein | elongation zone | 12 h |
| HORVU.MOREX.r3.5HG0517490.1 | -0.858383105 | 1.614894636 | -3.897186614 | 0.00028 | 0.015726837 | -0.05618385 | Protein kinase | elongation zone | 12 h |
| HORVU.MOREX.r3.6HG0632320.1 | 0.458402212 | 5.523777089 | 3.962546686 | 0.00022 | 0.014924834 | -0.06492276 | Leucine-rich repeat protein | elongation zone | 12 h |

**Table S5** Continued.

| ID | logFC | AveExpr | t | P.Value | adj.P.Val | B | gene description | root zone | time point |
|---|---|---|---|---|---|---|---|---|---|
| HORVU.MOREX.r3.1HG0050880.1 | 1.48303653 | 5.389192002 | 3.838027078 | 0.00033 | 0.017006082 | -0.06657299 | Methyltransferase-like protein 17, mitochondrial | elongation zone | 12 h |
| HORVU.MOREX.r3.5HG0493850.1 | 0.558372719 | 7.419096029 | 4.025347836 | 0.00018 | 0.014527878 | -0.06780218 | Acyl-[acyl-carrier-protein] hydrolase | elongation zone | 12 h |
| HORVU.MOREX.r3.5HG0427080.1 | 1.524451711 | 5.258314535 | 3.849603062 | 0.00032 | 0.016767189 | -0.06795918 | Kinesin-like protein | elongation zone | 12 h |
| HORVU.MOREX.r3.2HG0124910.1 | 0.886909253 | 9.424538243 | 4.054701509 | 0.00017 | 0.014078728 | -0.06884262 | 40S ribosomal protein SA | elongation zone | 12 h |
| HORVU.MOREX.r3.2HG0141900.1 | 0.668246315 | 5.9402647 | 3.976436261 | 0.00021 | 0.014881142 | -0.07121645 | L-ascorbate oxidase-like protein | elongation zone | 12 h |
| HORVU.MOREX.r3.2HG0142760.1 | 1.530216188 | 5.505958362 | 3.839235726 | 0.00033 | 0.016983175 | -0.07470097 | #N/A | elongation zone | 12 h |
| HORVU.MOREX.r3.4HG0386830.1 | 1.11565737 | 3.806923626 | 3.803685478 | 0.00037 | 0.017603779 | -0.07637784 | UDP-glucuronate decarboxylase protein 1 | elongation zone | 12 h |
| HORVU.MOREX.r3.3HG0288420.1 | 0.969784624 | 7.285552254 | 3.96225572 | 0.00022 | 0.014924834 | -0.07729412 | Mannan endo-1,4-beta-mannosidase-like protein | elongation zone | 12 h |
| HORVU.MOREX.r3.5HG0462450.1 | -1.399345648 | -0.846663356 | -3.752410566 | 0.00044 | 0.018288218 | -0.08802032 | S-acyltransferase | elongation zone | 12 h |
| HORVU.MOREX.r3.4HG0375550.1 | 2.704694656 | 0.494963378 | 3.807832092 | 0.00037 | 0.017580172 | -0.08927401 | Rapid ALkalinization Factor family protein | elongation zone | 12 h |
| HORVU.MOREX.r3.7HG0656360.1 | 1.76887709 | 7.497142351 | 3.935852718 | 0.00024 | 0.015535305 | -0.09078711 | Ribonucleoside-diphosphate reductase | elongation zone | 12 h |
| HORVU.MOREX.r3.5HG0510940.1 | 1.392537918 | 5.282693497 | 3.848005824 | 0.00032 | 0.01680982 | -0.09545177 | HMG-Y-related protein A | elongation zone | 12 h |
| HORVU.MOREX.r3.3HG0230090.1 | -0.896099977 | 1.525121802 | -4.014886429 | 0.00019 | 0.014546646 | -0.09557855 | Cysteine protease, putative | elongation zone | 12 h |
| HORVU.MOREX.r3.3HG0240450.1 | 0.82868465 | 4.298635098 | 3.841353568 | 0.00033 | 0.016953892 | -0.09620189 | Protein gamma response 1 | elongation zone | 12 h |
| HORVU.MOREX.r3.2HG0182750.1 | 0.914582292 | 5.83880247 | 3.914227872 | 0.00026 | 0.015692177 | -0.09730277 | Protein RETICULATA, chloroplastic | elongation zone | 12 h |
| HORVU.MOREX.r3.5HG0424440.1 | 1.696287723 | 2.104628459 | 3.739677556 | 0.00045 | 0.018562843 | -0.09809411 | #N/A | elongation zone | 12 h |
| HORVU.MOREX.r3.7HG0639380.1 | 1.178003064 | 7.629623995 | 3.958914122 | 0.00023 | 0.014924834 | -0.09918018 | 60 kDa chaperonin | elongation zone | 12 h |
| HORVU.MOREX.r3.6HG0541490.1 | 2.005880897 | 2.862577415 | 3.737877706 | 0.00046 | 0.018607144 | -0.10144701 | Mitochondrial transcription termination factor-like | elongation zone | 12 h |
| HORVU.MOREX.r3.7HG0689040.1 | 2.001654105 | 2.151352586 | 3.745522922 | 0.00045 | 0.018421721 | -0.10278845 | RNA-binding protein | elongation zone | 12 h |
| HORVU.MOREX.r3.1HG0090820.1 | 0.58786292 | 4.907388768 | 3.89421745 | 0.00028 | 0.015751512 | -0.10397451 | Protein CHUP1, chloroplastic | elongation zone | 12 h |
| HORVU.MOREX.r3.7HG0659030.1 | 0.789550842 | 7.54316044 | 3.964215859 | 0.00022 | 0.014924834 | -0.10948842 | Ubiquitin carboxyl-terminal hydrolase, putative | elongation zone | 12 h |
| HORVU.MOREX.r3.7HG0634320.1 | 0.475201555 | 4.501085155 | 3.898891273 | 0.00027 | 0.015692177 | -0.10980924 | NAC domain-containing protein | elongation zone | 12 h |
| HORVU.MOREX.r3.5HG0450100.1 | 0.604748966 | 5.024744254 | 3.889522479 | 0.00028 | 0.015756066 | -0.11062129 | Quinolinate synthase A | elongation zone | 12 h |
| HORVU.MOREX.r3.2HG0096760.1 | 1.066723121 | 4.203419621 | 3.80389811 | 0.00037 | 0.017603779 | -0.11541609 | TOM1-like protein 2 | elongation zone | 12 h |
| HORVU.MOREX.r3.7HG0663790.1 | 1.898648736 | 4.045385183 | 3.753995234 | 0.00043 | 0.018288218 | -0.12581478 | Galactoside 2-alpha-L-fucosyltransferase | elongation zone | 12 h |
| HORVU.MOREX.r3.2HG0119160.1 | 0.562141433 | 6.243872664 | 3.929229704 | 0.00025 | 0.015626386 | -0.12629021 | #N/A | elongation zone | 12 h |
| HORVU.MOREX.r3.3HG0257850.1 | 1.86381901 | 2.089146786 | 3.733073283 | 0.00046 | 0.018667357 | -0.12677276 | Magnesium-protoporphyrin IX monomethyl ester [oxidative] cycl | elongation zone | 12 h |
| HORVU.MOREX.r3.5HG0491230.1 | 1.19002053 | 2.284227707 | 3.752975009 | 0.00043 | 0.018288218 | -0.12737294 | Dihydroflavonol-4-reductase | elongation zone | 12 h |
| HORVU.MOREX.r3.1HG0054000.1 | 1.172877099 | 4.221868706 | 3.778182727 | 0.0004 | 0.017833387 | -0.12813916 | ATP-dependent Clp protease proteolytic subunit | elongation zone | 12 h |
| HORVU.MOREX.r3.4HG0385300.1 | 0.593895166 | 9.307024474 | 4.051588368 | 0.00017 | 0.014078728 | -0.13494948 | Elongation factor 1-alpha | elongation zone | 12 h |
| HORVU.MOREX.r3.4HG0418530.1 | 1.226654501 | 4.704048819 | 3.794315962 | 0.00038 | 0.017696476 | -0.13775081 | ATP-dependent chaperone ClpB | elongation zone | 12 h |
| HORVU.MOREX.r3.1HG0086010.1 | 0.560533979 | 5.131586677 | 3.90033892 | 0.00027 | 0.015692177 | -0.14386209 | TSA: Wollemia nobilis Ref_Wollemi_Transcript_14910_3291 tran | elongation zone | 12 h |
| HORVU.MOREX.r3.2HG0205680.1 | 2.359673438 | 1.160765307 | 3.77305137 | 0.00041 | 0.017914792 | -0.14665082 | Pentatricopeptide repeat-containing protein | elongation zone | 12 h |
| HORVU.MOREX.r3.6HG0573900.1 | 1.042973827 | 4.176459117 | 3.772368707 | 0.00041 | 0.017914792 | -0.14744881 | SWI/SNF complex subunit SWI3B | elongation zone | 12 h |
| HORVU.MOREX.r3.2HG0095970.1 | 1.56109742 | 3.317154222 | 3.733378805 | 0.00046 | 0.018667357 | -0.16905724 | NBS-LRR disease resistance protein-like protein | elongation zone | 12 h |
| HORVU.MOREX.r3.3HG0242880.1 | -0.928528605 | -0.890008185 | -3.789830288 | 0.00039 | 0.017696476 | -0.17344614 | Phloem protein 2-like protein | elongation zone | 12 h |
| HORVU.MOREX.r3.6HG0606040.1 | 1.730174942 | 3.594810934 | 3.718386509 | 0.00048 | 0.019097198 | -0.17638392 | Survival motor neuron | elongation zone | 12 h |
| HORVU.MOREX.r3.6HG0549930.1 | 1.893574699 | 3.393056211 | 3.714759561 | 0.00049 | 0.019131832 | -0.17678405 | #N/A | elongation zone | 12 h |
| HORVU.MOREX.r3.7HG0640560.1 | 2.450937624 | 0.442208837 | 3.751660882 | 0.00044 | 0.018290689 | -0.17977253 | NBS-LRR disease resistance protein-like protein | elongation zone | 12 h |
| HORVU.MOREX.r3.5HG0508680.1 | 1.740392098 | 2.582186545 | 3.749329245 | 0.00044 | 0.018313253 | -0.18012033 | Eukaryotic aspartyl protease family protein | elongation zone | 12 h |
| HORVU.MOREX.r3.7HG0725700.1 | -2.273909445 | -1.181670666 | -3.721765866 | 0.00048 | 0.019002693 | -0.1816591 | Receptor-like protein kinase | elongation zone | 12 h |

**Table S5** Continued.

| ID | logFC | AveExpr | t | P.Value | adj.P.Val | B | gene description | root zone | time point |
|---|---|---|---|---|---|---|---|---|---|
| HORVU.MOREX.r3.3HG0304490.1 | -1.029896928 | -1.400694178 | -3.786732496 | 0.00039 | 0.017712844 | -0.19029298 | Myosin IC heavy chain | elongation zone | 12 h |
| HORVU.MOREX.r3.1HG0092430.1 | 1.177273546 | 4.997512913 | 3.790293047 | 0.00039 | 0.017696476 | -0.19457707 | Asparagine synthetase | elongation zone | 12 h |
| HORVU.MOREX.r3.4HG0416070.1 | 2.257225879 | 7.030233778 | 3.893243038 | 0.00028 | 0.015751512 | -0.19466809 | GATA transcription factor | elongation zone | 12 h |
| HORVU.MOREX.r3.5HG0460170.1 | 1.961910199 | 2.2284743 | 3.70138203 | 0.00051 | 0.019449812 | -0.21040802 | Pentatricopeptide repeat-containing protein | elongation zone | 12 h |
| HORVU.MOREX.r3.1HG0095390.1 | 1.399258203 | 8.175356882 | 3.929632038 | 0.00025 | 0.015626386 | -0.21122047 | Histone deacetylase, putative | elongation zone | 12 h |
| HORVU.MOREX.r3.4HG0347310.1 | 1.665646513 | 5.304851749 | 3.775917177 | 0.0004 | 0.017855434 | -0.21578795 | DNA polymerase | elongation zone | 12 h |
| HORVU.MOREX.r3.1HG0094480.1 | 1.348983487 | 4.237351174 | 3.746610293 | 0.00044 | 0.018395923 | -0.21816165 | ATP-dependent Clp protease proteolytic subunit | elongation zone | 12 h |
| HORVU.MOREX.r3.7HG0677580.1 | 1.377613874 | 5.782252606 | 3.853974711 | 0.00032 | 0.016689644 | -0.2182241 | Protein O-linked-mannose beta-1,4-N-acetylglucosaminyltransfe | elongation zone | 12 h |
| HORVU.MOREX.r3.5HG0468930.1 | 2.64618113 | 3.393289477 | 3.695109452 | 0.00052 | 0.019449812 | -0.21900988 | Argonaute | elongation zone | 12 h |
| HORVU.MOREX.r3.1HG0005840.1 | 1.465162812 | 1.736344477 | 3.691891215 | 0.00053 | 0.019449812 | -0.22802844 | Embryo defective 2735 | elongation zone | 12 h |
| HORVU.MOREX.r3.4HG0387040.1 | 1.285401259 | 5.073389367 | 3.786857757 | 0.00039 | 0.017712844 | -0.22873255 | RNA-binding protein | elongation zone | 12 h |
| HORVU.MOREX.r3.5HG0497180.1 | 2.540756619 | 0.857668817 | 3.740014119 | 0.00045 | 0.018562843 | -0.23505186 | Pentatricopeptide repeat-containing protein, putative | elongation zone | 12 h |
| HORVU.MOREX.r3.2HG0114930.1 | 2.325672271 | 8.18750701 | 3.877999598 | 0.00029 | 0.016094925 | -0.24873057 | Nucleolar protein | elongation zone | 12 h |
| HORVU.MOREX.r3.3HG0303110.1 | 0.981411394 | 7.531826753 | 3.91444678 | 0.00026 | 0.015692177 | -0.25280806 | 30S ribosomal protein S8, putative | elongation zone | 12 h |
| HORVU.MOREX.r3.4HG0336830.1 | 0.529865777 | 6.56218357 | 3.901668516 | 0.00027 | 0.015692177 | -0.25822231 | ATP-dependent RNA helicase | elongation zone | 12 h |
| HORVU.MOREX.r3.4HG0342440.1 | 2.425563027 | 3.291704004 | 3.680189758 | 0.00055 | 0.019711469 | -0.25911867 | myosin-2 heavy chain-like protein | elongation zone | 12 h |
| HORVU.MOREX.r3.1HG0057990.1 | 0.639221577 | 4.4289102 | 3.809259891 | 0.00036 | 0.017580172 | -0.26720386 | O-fucosyltransferase family protein | elongation zone | 12 h |
| HORVU.MOREX.r3.5HG0516720.1 | 1.967991277 | 6.439287689 | 3.776146284 | 0.0004 | 0.017855434 | -0.26861529 | Glutathione-S-transferase | elongation zone | 12 h |
| HORVU.MOREX.r3.2HG0163290.1 | 1.627642012 | 1.166577297 | 3.679501944 | 0.00055 | 0.019711469 | -0.27748172 | RecQ-mediated genome instability protein 1 | elongation zone | 12 h |
| HORVU.MOREX.r3.7HG0749600.1 | 1.238952692 | 3.788333413 | 3.715171331 | 0.00049 | 0.019131832 | -0.27812263 | Gamma-tubulin complex component | elongation zone | 12 h |
| HORVU.MOREX.r3.1HG0048070.1 | 2.094624759 | 2.001188019 | 3.67270249 | 0.00056 | 0.019889362 | -0.29265493 | Folylpolyglutamate synthase | elongation zone | 12 h |
| HORVU.MOREX.r3.4HG0331260.1 | 1.231570587 | 3.068963796 | 3.673199341 | 0.00056 | 0.019889362 | -0.29553709 | ABC transporter ATP-binding protein ARB1 | elongation zone | 12 h |
| HORVU.MOREX.r3.6HG0554800.1 | 1.080418235 | 4.238669575 | 3.742204212 | 0.00045 | 0.018540177 | -0.29684055 | Transmembrane protein 53 | elongation zone | 12 h |
| HORVU.MOREX.r3.3HG0300870.1 | 1.787806976 | 5.149023989 | 3.736258396 | 0.00046 | 0.018625563 | -0.30234637 | #N/A | elongation zone | 12 h |
| HORVU.MOREX.r3.6HG0611290.1 | 1.079786181 | 7.158294058 | 3.871002518 | 0.0003 | 0.016327144 | -0.30334899 | Acetolactate synthase | elongation zone | 12 h |
| HORVU.MOREX.r3.2HG0215550.1 | 1.076161527 | 7.917137824 | 3.919469534 | 0.00026 | 0.015692177 | -0.31860406 | Peroxidase | elongation zone | 12 h |
| HORVU.MOREX.r3.2HG0200640.1 | 0.440489098 | 7.509809216 | 3.929752623 | 0.00025 | 0.015626386 | -0.32453776 | Carboxypeptidase | elongation zone | 12 h |
| HORVU.MOREX.r3.1HG0027380.1 | 2.623071012 | 1.136430417 | 3.7098177 | 0.0005 | 0.019247224 | -0.32668582 | Pentatricopeptide repeat-containing protein | elongation zone | 12 h |
| HORVU.MOREX.r3.3HG0295680.1 | -0.585662433 | 4.416665159 | -3.892494031 | 0.00028 | 0.015751512 | -0.32957069 | Dihydropyrimidinase | elongation zone | 12 h |
| HORVU.MOREX.r3.7HG0652830.1 | 1.930400883 | 0.863490792 | 3.674539948 | 0.00056 | 0.019889362 | -0.33204205 | Disease resistance protein RPP13 | elongation zone | 12 h |
| HORVU.MOREX.r3.6HG0558800.1 | 0.608218993 | 5.008690522 | 3.808543161 | 0.00036 | 0.017580172 | -0.33352505 | DNA-binding protein BIN4 | elongation zone | 12 h |
| HORVU.MOREX.r3.3HG0329870.1 | -1.188891159 | -1.903215009 | -3.767460619 | 0.00042 | 0.017977071 | -0.33877421 | Pectin acetylesterase | elongation zone | 12 h |
| HORVU.MOREX.r3.2HG0213240.1 | 1.297868338 | 2.83023571 | 3.664953315 | 0.00057 | 0.020165373 | -0.33927185 | Actin-related protein | elongation zone | 12 h |
| HORVU.MOREX.r3.2HG0200450.1 | 1.043475115 | 0.466638302 | 3.723058016 | 0.00048 | 0.019002693 | -0.3394893 | Beta-fructofuranosidase, insoluble isoenzyme 1 | elongation zone | 12 h |
| HORVU.MOREX.r3.1HG0070640.1 | 1.080389264 | 5.376818556 | 3.762943092 | 0.00042 | 0.01805759 | -0.33949757 | KH domain-containing protein | elongation zone | 12 h |
| HORVU.MOREX.r3.3HG0299990.1 | -0.712534083 | 6.228942935 | -3.93016709 | 0.00025 | 0.015626386 | -0.33959842 | Glutathione-S-transferase | elongation zone | 12 h |
| HORVU.MOREX.r3.6HG0633420.1 | 0.974342945 | 7.127371427 | 3.874716956 | 0.0003 | 0.016177895 | -0.34587705 | Pentatricopeptide repeat-containing protein | elongation zone | 12 h |
| HORVU.MOREX.r3.7HG0662670.1 | 0.695345569 | 4.911988083 | 3.815819148 | 0.00036 | 0.017465122 | -0.34994719 | aminoalcoholphosphotransferase 1 | elongation zone | 12 h |
| HORVU.MOREX.r3.7HG0676040.1 | -0.792488955 | -0.533828415 | -3.739711873 | 0.00045 | 0.018562843 | -0.3551845 | Decaprenyl-phosphate N-acetylglucosaminephosphotransferase | elongation zone | 12 h |
| HORVU.MOREX.r3.1HG0051020.1 | 0.660151682 | 5.895903782 | 3.84206365 | 0.00033 | 0.016953892 | -0.35596042 | Membrane protein insertase YidC | elongation zone | 12 h |
| HORVU.MOREX.r3.3HG0221460.1 | 0.622848705 | 7.255590658 | 3.909916342 | 0.00026 | 0.015692177 | -0.35694119 | Glycosyltransferase | elongation zone | 12 h |

**Table S5** Continued.

| ID | logFC | AveExpr | t | P.Value | adj.P.Val | B | gene description | root zone | time point |
|---|---|---|---|---|---|---|---|---|---|
| HORVU.MOREX.r3.7HG0732610.1 | 1.500245523 | 4.827784988 | 3.711734336 | 0.0005 | 0.019227921 | -0.35943804 | DNA polymerase alpha subunit B | elongation zone | 12 h |
| HORVU.MOREX.r3.2HG0175590.1 | 2.94068047 | 3.241311809 | 3.644971384 | 0.00061 | 0.020507565 | -0.36176498 | ATP-dependent zinc metalloprotease FtsH 2 | elongation zone | 12 h |
| HORVU.MOREX.r3.1HG0093710.1 | 0.266430777 | 6.865906621 | 3.916464361 | 0.00026 | 0.015692177 | -0.36629462 | dihydroflavonol 4-reductase/flavanone protein | elongation zone | 12 h |
| HORVU.MOREX.r3.2HG0174030.1 | 2.561601845 | 1.901242042 | 3.672923982 | 0.00056 | 0.019889362 | -0.36991449 | RNA-dependent RNA polymerase | elongation zone | 12 h |
| HORVU.MOREX.r3.7HG0643280.1 | 0.98822067 | 7.144031345 | 3.862066649 | 0.00031 | 0.016574489 | -0.37095603 | Pentatricopeptide repeat-containing protein | elongation zone | 12 h |
| HORVU.MOREX.r3.7HG0671880.1 | 0.886956638 | 3.67043834 | 3.702082327 | 0.00051 | 0.019449812 | -0.37753819 | RING/U-box protein | elongation zone | 12 h |
| HORVU.MOREX.r3.5HG0442360.1 | 1.081916712 | 6.472159574 | 3.827397511 | 0.00034 | 0.017220216 | -0.38222113 | Diaminopimelate epimerase | elongation zone | 12 h |
| HORVU.MOREX.r3.5HG0498530.1 | 3.055988022 | 3.096808835 | 3.633380806 | 0.00063 | 0.020646627 | -0.38549414 | Subtilisin-like protease | elongation zone | 12 h |
| HORVU.MOREX.r3.7HG0708820.1 | 0.838151521 | 5.143802841 | 3.761765209 | 0.00042 | 0.018086982 | -0.38566705 | PHD finger family protein | elongation zone | 12 h |
| HORVU.MOREX.r3.3HG0256580.1 | 0.483826913 | 6.795309482 | 3.878374605 | 0.00029 | 0.016094925 | -0.3896554 | NEDD8-activating enzyme E1 catalytic subunit | elongation zone | 12 h |
| HORVU.MOREX.r3.5HG0489810.1 | 1.088823816 | 3.755916949 | 3.697314584 | 0.00052 | 0.019449812 | -0.39051409 | Caffeoyl-CoA O-methyltransferase | elongation zone | 12 h |
| HORVU.MOREX.r3.6HG0577760.2 | 3.221193491 | 1.21790513 | 3.700850403 | 0.00051 | 0.019449812 | -0.39273698 | Ran-binding zinc finger protein | elongation zone | 12 h |
| HORVU.MOREX.r3.3HG0312290.1 | 1.828693254 | 2.160544907 | 3.63480839 | 0.00063 | 0.020623325 | -0.39454269 | Receptor-like kinase | elongation zone | 12 h |
| HORVU.MOREX.r3.4HG0387850.1 | 1.434073074 | 4.670908916 | 3.692599869 | 0.00053 | 0.019449812 | -0.3983981 | #N/A | elongation zone | 12 h |
| HORVU.MOREX.r3.6HG0581570.1 | 0.397031917 | 4.929735327 | 3.81736707 | 0.00035 | 0.017465122 | -0.39878496 | Leucine-rich repeat-containing protein 1 | elongation zone | 12 h |
| HORVU.MOREX.r3.4HG0345330.1 | 1.597331619 | 3.738426759 | 3.658871306 | 0.00058 | 0.020375761 | -0.40246251 | DNA-directed RNA polymerase | elongation zone | 12 h |
| HORVU.MOREX.r3.7HG0640430.1 | 0.890259147 | 3.689469005 | 3.68166363 | 0.00054 | 0.019711469 | -0.40381677 | Disease resistance protein (TIR-NBS-LRR class) | elongation zone | 12 h |
| HORVU.MOREX.r3.1HG0084850.1 | 1.402388705 | 4.250961882 | 3.689079015 | 0.00053 | 0.019449812 | -0.40397038 | Histone chaperone | elongation zone | 12 h |
| HORVU.MOREX.r3.4HG0370380.1 | 0.472861239 | 5.905711064 | 3.855983905 | 0.00031 | 0.016685099 | -0.40456454 | Erlin-2 | elongation zone | 12 h |
| HORVU.MOREX.r3.1HG0071050.1 | -1.15528135 | -0.746820209 | -3.715672568 | 0.00049 | 0.019131832 | -0.40479133 | Expansin | elongation zone | 12 h |
| HORVU.MOREX.r3.6HG0551300.1 | 1.885463903 | 0.944462851 | 3.639449283 | 0.00062 | 0.020516782 | -0.40483502 | F-box family protein | elongation zone | 12 h |
| HORVU.MOREX.r3.5HG0487010.1 | 1.878015489 | 2.469699433 | 3.625507145 | 0.00065 | 0.020713759 | -0.40595576 | Pentatricopeptide repeat-containing protein | elongation zone | 12 h |
| HORVU.MOREX.r3.7HG0703580.1 | 0.716774355 | 4.485598753 | 3.794504145 | 0.00038 | 0.017696476 | -0.40706487 | Glyceraldehyde-3-phosphate dehydrogenase | elongation zone | 12 h |
| HORVU.MOREX.r3.1HG0046180.1 | -0.496732635 | 5.918664759 | -3.910461387 | 0.00026 | 0.015692177 | -0.40909638 | Ser/Thr protein phosphatase family protein, expressed | elongation zone | 12 h |
| HORVU.MOREX.r3.5HG0478010.1 | 0.790400761 | 3.664153717 | 3.696398391 | 0.00052 | 0.019449812 | -0.41128551 | Chaperone protein DnaJ | elongation zone | 12 h |
| HORVU.MOREX.r3.2HG0194120.1 | 1.035455542 | 4.519929846 | 3.706266591 | 0.0005 | 0.019342524 | -0.41198606 | Phospholipid scramblase | elongation zone | 12 h |
| HORVU.MOREX.r3.6HG0605940.1 | -0.811550697 | 3.17964847 | -3.792933316 | 0.00038 | 0.017696476 | -0.41325016 | Xyloglucan endotransglucosylase/hydrolase | elongation zone | 12 h |
| HORVU.MOREX.r3.2HG0170840.1 | 0.441757548 | 6.055499187 | 3.842260554 | 0.00033 | 0.016953892 | -0.41690148 | 26S proteasome non-ATPase regulatory subunit-like protein | elongation zone | 12 h |
| HORVU.MOREX.r3.3HG0248280.1 | 0.416878457 | 8.506150471 | 3.95514316 | 0.00023 | 0.014924834 | -0.41869663 | Reticulon-like protein | elongation zone | 12 h |
| HORVU.MOREX.r3.4HG0335990.1 | 2.896453733 | 2.777740737 | 3.628032557 | 0.00064 | 0.020713759 | -0.41999475 | Pentatricopeptide repeat-containing protein | elongation zone | 12 h |
| HORVU.MOREX.r3.1HG0079750.1 | -1.279614574 | -1.155378593 | -3.680163317 | 0.00055 | 0.019711469 | -0.42028166 | MYB transcription factor | elongation zone | 12 h |
| HORVU.MOREX.r3.2HG0197560.1 | 0.747212548 | 5.10495621 | 3.763249268 | 0.00042 | 0.01805759 | -0.42354107 | RNA-binding protein 39 | elongation zone | 12 h |
| HORVU.MOREX.r3.5HG0478390.1 | 0.588109754 | 5.551366827 | 3.807547332 | 0.00037 | 0.017580172 | -0.42416109 | PCI domain-containing protein 2 | elongation zone | 12 h |
| HORVU.MOREX.r3.5HG0501270.1 | 1.777626755 | 6.40008053 | 3.767020967 | 0.00042 | 0.017977071 | -0.42464445 | Chromatin assembly factor-1 | elongation zone | 12 h |
| HORVU.MOREX.r3.4HG0396660.1 | 1.968604424 | 6.770443008 | 3.778517059 | 0.0004 | 0.017833387 | -0.42705547 | Replication protein A 70 kDa DNA-binding subunit | elongation zone | 12 h |
| HORVU.MOREX.r3.1HG0086120.1 | 1.964556698 | 4.055301306 | 3.651439682 | 0.0006 | 0.020507565 | -0.43030629 | Histone H2B | elongation zone | 12 h |
| HORVU.MOREX.r3.1HG0050860.1 | 1.656957356 | 2.488742376 | 3.615029906 | 0.00067 | 0.020839563 | -0.43401747 | HIT zinc finger family protein, expressed | elongation zone | 12 h |
| HORVU.MOREX.r3.5HG0461170.1 | 0.678107149 | 4.201237205 | 3.725532012 | 0.00047 | 0.018928454 | -0.44293964 | Protein FAR1-RELATED SEQUENCE 5 | elongation zone | 12 h |
| HORVU.MOREX.r3.1HG0003930.1 | 0.962297706 | 6.637383562 | 3.814120626 | 0.00036 | 0.017476052 | -0.45274573 | 50S ribosomal protein L22 | elongation zone | 12 h |
| HORVU.MOREX.r3.2HG0100770.1 | 0.543533917 | 4.593259975 | 3.781860548 | 0.0004 | 0.017833387 | -0.45479275 | Glutamyl-tRNA(Gln) amidotransferase subunit A | elongation zone | 12 h |
| HORVU.MOREX.r3.1HG0072540.1 | 1.086136261 | 5.751312553 | 3.788414291 | 0.00039 | 0.017712844 | -0.45659796 | Histone H4 | elongation zone | 12 h |

**Table S5** Continued.

| ID | logFC | AveExpr | t | P.Value | adj.P.Val | B | gene description | root zone | time point |
|---|---|---|---|---|---|---|---|---|---|
| HORVU.MOREX.r3.1HG0017310.1 | 1.300190628 | 6.737539169 | 3.790038435 | 0.00039 | 0.017696476 | -0.45710296 | Auxin influx transporter | elongation zone | 12 h |
| HORVU.MOREX.r3.5HG0526170.1 | -1.105496647 | -1.761923852 | -3.659614159 | 0.00058 | 0.02036338 | -0.45786442 | Actin depolymerizing factor | elongation zone | 12 h |
| HORVU.MOREX.r3.3HG0288320.1 | 1.061262251 | 6.036276973 | 3.773713739 | 0.00041 | 0.017914792 | -0.46570255 | Nuclear pore complex protein NUP85 | elongation zone | 12 h |
| HORVU.MOREX.r3.6HG0564510.1 | 1.00389942 | 4.576465099 | 3.698714608 | 0.00052 | 0.019449812 | -0.46609928 | Arabinose 5-phosphate isomerase, putative | elongation zone | 12 h |
| HORVU.MOREX.r3.6HG0542170.1 | 2.675167317 | -0.57519434 | 3.719074923 | 0.00048 | 0.019092689 | -0.46848266 | #N/A | elongation zone | 12 h |
| HORVU.MOREX.r3.5HG0466010.1 | 0.600223118 | 3.637083856 | 3.688886776 | 0.00053 | 0.019449812 | -0.47359046 | Glutaredoxin | elongation zone | 12 h |
| HORVU.MOREX.r3.7HG0656370.1 | 3.700888403 | 3.059766906 | 3.605225397 | 0.00069 | 0.021013719 | -0.4789416 | Bifunctional inhibitor/lipid-transfer protein/seed storage 2S albu | elongation zone | 12 h |
| HORVU.MOREX.r3.7HG0667510.1 | 0.999018287 | 1.198678787 | 3.608937762 | 0.00068 | 0.020937187 | -0.48220901 | ADP-ribosylation factor family protein | elongation zone | 12 h |
| HORVU.MOREX.r3.3HG0307240.1 | -2.293989484 | -3.376882233 | -3.603978199 | 0.00069 | 0.021039776 | -0.48249504 | Amino acid permease | elongation zone | 12 h |
| HORVU.MOREX.r3.4HG0405960.1 | 1.28664455 | 4.60369269 | 3.668070952 | 0.00057 | 0.020073937 | -0.48495636 | Origin recognition complex subunit 2 | elongation zone | 12 h |
| HORVU.MOREX.r3.6HG0631400.1 | -0.920202146 | 2.336066158 | -3.787352023 | 0.00039 | 0.017712844 | -0.48766214 | Disease resistance protein RPM1 | elongation zone | 12 h |
| HORVU.MOREX.r3.2HG0181010.1 | 1.410523911 | 4.533256071 | 3.651280628 | 0.0006 | 0.020507565 | -0.49513039 | RNA binding protein | elongation zone | 12 h |
| HORVU.MOREX.r3.3HG0297180.1 | 1.537818976 | 2.307651608 | 3.594693845 | 0.00071 | 0.021178576 | -0.49674734 | Pentatricopeptide repeat-containing protein | elongation zone | 12 h |
| HORVU.MOREX.r3.2HG0195110.2 | 1.838439388 | 0.674913375 | 3.596566636 | 0.00071 | 0.021178576 | -0.50014782 | Remorin family protein | elongation zone | 12 h |
| HORVU.MOREX.r3.4HG0403200.1 | 2.533786693 | 0.946991273 | 3.627980753 | 0.00064 | 0.020713759 | -0.50037778 | Pentatricopeptide repeat-containing protein | elongation zone | 12 h |
| HORVU.MOREX.r3.3HG0277200.1 | 1.200202633 | 6.184721678 | 3.796644156 | 0.00038 | 0.017696476 | -0.50205984 | Pyruvate kinase | elongation zone | 12 h |
| HORVU.MOREX.r3.2HG0113870.1 | 1.548708327 | 2.572538505 | 3.593993109 | 0.00071 | 0.021178576 | -0.50346811 | Pentatricopeptide repeat-containing protein | elongation zone | 12 h |
| HORVU.MOREX.r3.4HG0416570.1 | 1.975996615 | 1.341219886 | 3.591318713 | 0.00072 | 0.021247809 | -0.50419438 | Major facilitator superfamily domain-containing protein | elongation zone | 12 h |
| HORVU.MOREX.r3.7HG0685290.1 | 0.875490369 | 7.739023195 | 3.850005913 | 0.00032 | 0.016767189 | -0.5045831 | Glycine--tRNA ligase | elongation zone | 12 h |
| HORVU.MOREX.r3.2HG0173270.1 | 1.70675105 | 4.153097441 | 3.614840613 | 0.00067 | 0.020839563 | -0.50646911 | ATP-dependent RNA helicase | elongation zone | 12 h |
| HORVU.MOREX.r3.7HG0736160.2 | 2.379610321 | 1.391443228 | 3.616830888 | 0.00067 | 0.020839563 | -0.50946837 | DNA primase/helicase | elongation zone | 12 h |
| HORVU.MOREX.r3.5HG0514790.3 | 0.568863209 | 10.92395731 | 3.995506629 | 0.0002 | 0.014738733 | -0.51168087 | Elongation factor 1-alpha | elongation zone | 12 h |
| HORVU.MOREX.r3.2HG0100900.1 | -0.379504376 | 4.579241165 | -3.799164817 | 0.00038 | 0.017696476 | -0.5122824 | Phosphoglycerate mutase-like protein | elongation zone | 12 h |
| HORVU.MOREX.r3.2HG0183640.1 | 1.585572849 | 4.446959637 | 3.641817687 | 0.00062 | 0.020516782 | -0.51919242 | P-loop containing nucleoside triphosphate hydrolases superfami | elongation zone | 12 h |
| HORVU.MOREX.r3.2HG0188760.1 | 1.860179119 | 0.265515101 | 3.625507807 | 0.00065 | 0.020713759 | -0.51960045 | Glycosyltransferases | elongation zone | 12 h |
| HORVU.MOREX.r3.3HG0329590.1 | 1.265359979 | 1.948546577 | 3.587410104 | 0.00073 | 0.021412735 | -0.52013584 | 30S ribosomal protein S17 | elongation zone | 12 h |
| HORVU.MOREX.r3.3HG0219410.1 | -0.869457323 | 3.869903081 | -3.780708403 | 0.0004 | 0.017833387 | -0.52674691 | Hydroxyproline-rich glycoprotein family protein | elongation zone | 12 h |
| HORVU.MOREX.r3.5HG0467960.1 | 1.455384419 | 2.390386799 | 3.576156653 | 0.00075 | 0.021691049 | -0.53886412 | DNA-directed RNA polymerase subunit | elongation zone | 12 h |
| HORVU.MOREX.r3.4HG0334890.1 | 1.096206981 | 7.019132846 | 3.791184535 | 0.00039 | 0.017696476 | -0.53998225 | Glycine-rich family protein | elongation zone | 12 h |
| HORVU.MOREX.r3.6HG0611730.1 | 1.935721726 | 6.381854124 | 3.699304715 | 0.00051 | 0.019449812 | -0.54512436 | H/ACA ribonucleoprotein complex subunit 2-like protein | elongation zone | 12 h |
| HORVU.MOREX.r3.3HG0243420.1 | 1.232380764 | 4.335089839 | 3.640045562 | 0.00062 | 0.020516782 | -0.55572377 | DNA polymerase delta subunit 3 | elongation zone | 12 h |
| HORVU.MOREX.r3.3HG0283990.1 | 0.522128773 | 7.009142279 | 3.840389447 | 0.00033 | 0.01696336 | -0.55578352 | Glycylpeptide N-tetradecanoyltransferase | elongation zone | 12 h |
| HORVU.MOREX.r3.3HG0294270.1 | 2.959702511 | -0.244504286 | 3.65244831 | 0.0006 | 0.020507565 | -0.55646035 | Protein BIG GRAIN 1 | elongation zone | 12 h |
| HORVU.MOREX.r3.2HG0118800.1 | 0.856877637 | 4.917895689 | 3.71074039 | 0.0005 | 0.019227921 | -0.56245996 | Mitochondrial inner membrane translocase subunit Tim44 | elongation zone | 12 h |
| HORVU.MOREX.r3.3HG0268790.1 | 2.653313743 | 1.412428125 | 3.611058505 | 0.00068 | 0.020915275 | -0.56880837 | Pentatricopeptide repeat-containing protein | elongation zone | 12 h |
| HORVU.MOREX.r3.2HG0199600.1 | 2.118480391 | 2.139351569 | 3.566133052 | 0.00078 | 0.021941854 | -0.57051048 | Organic cation transporter protein | elongation zone | 12 h |
| HORVU.MOREX.r3.2HG0186650.1 | 0.969578529 | 7.283329417 | 3.804676665 | 0.00037 | 0.017603779 | -0.57060133 | Non-specific serine/threonine protein kinase | elongation zone | 12 h |
| HORVU.MOREX.r3.1HG0039000.1 | 1.168896713 | 5.928790739 | 3.725835651 | 0.00047 | 0.018928454 | -0.5720679 | Translin family protein, expressed | elongation zone | 12 h |
| HORVU.MOREX.r3.3HG0259970.1 | 0.590626069 | 6.103470349 | 3.781597633 | 0.0004 | 0.017833387 | -0.57430802 | Bromo-adjacent-like (BAH) domain protein | elongation zone | 12 h |
| HORVU.MOREX.r3.4HG0388320.1 | 0.718737277 | 3.811059557 | 3.647024897 | 0.00061 | 0.020507565 | -0.57487862 | ubiquitin carboxyl-terminal hydrolase | elongation zone | 12 h |
| HORVU.MOREX.r3.5HG0486780.1 | 0.770020347 | 2.89699251 | 3.61226515 | 0.00067 | 0.020885636 | -0.5877369 | Zinc-binding dehydrogenase family protein | elongation zone | 12 h |

**Table S5** Continued.

| ID | logFC | AveExpr | t | P.Value | adj.P.Val | B | gene description | root zone | time point |
|---|---|---|---|---|---|---|---|---|---|
| HORVU.MOREX.r3.2HG0152530.1 | -0.452677668 | 4.851529635 | -3.792132056 | 0.00038 | 0.017696476 | -0.58778282 | Fasciclin-like arabinogalactan protein | elongation zone | 12 h |
| HORVU.MOREX.r3.2HG0117880.1 | -3.132515163 | -0.66122785 | -3.561528127 | 0.00079 | 0.022039057 | -0.58784169 | GDSL esterase/lipase | elongation zone | 12 h |
| HORVU.MOREX.r3.3HG0306870.1 | 2.819877444 | 0.992463659 | 3.597778949 | 0.00071 | 0.021178576 | -0.5885416 | nuclear pore anchor | elongation zone | 12 h |
| HORVU.MOREX.r3.3HG0309820.1 | 0.80884575 | 5.758874854 | 3.757695938 | 0.00043 | 0.018168927 | -0.58958121 | Small ubiquitin-related modifier | elongation zone | 12 h |
| HORVU.MOREX.r3.1HG0024910.1 | 0.893573348 | 9.026974973 | 3.869037902 | 0.0003 | 0.016343249 | -0.5916618 | Ubiquitin | elongation zone | 12 h |
| HORVU.MOREX.r3.7HG0731780.1 | 1.083508591 | 5.146448477 | 3.662368559 | 0.00058 | 0.020275771 | -0.59251743 | Gag polyprotein | elongation zone | 12 h |
| HORVU.MOREX.r3.6HG0624580.1 | 1.362308894 | 4.00677816 | 3.612167884 | 0.00067 | 0.020885636 | -0.59385685 | Translation initiation factor IF-2, putative isoform 2 | elongation zone | 12 h |
| HORVU.MOREX.r3.4HG0416450.1 | 0.693097767 | 6.139964885 | 3.763879698 | 0.00042 | 0.01805759 | -0.59623987 | Protein ARABIDILLO 1 | elongation zone | 12 h |
| HORVU.MOREX.r3.5HG0520260.1 | 2.544275733 | 1.284181653 | 3.593278217 | 0.00072 | 0.021178576 | -0.59800101 | GTPase obg | elongation zone | 12 h |
| HORVU.MOREX.r3.5HG0471960.1 | 1.307390422 | 3.222285397 | 3.594784106 | 0.00071 | 0.021178576 | -0.59897355 | Chromosome transmission fidelity protein 8 | elongation zone | 12 h |
| HORVU.MOREX.r3.6HG0548670.1 | 0.980441257 | 5.252080241 | 3.727247317 | 0.00047 | 0.018928454 | -0.59923073 | Histone H4 | elongation zone | 12 h |
| HORVU.MOREX.r3.3HG0284120.1 | 0.835427551 | 7.524666369 | 3.808600202 | 0.00036 | 0.017580172 | -0.60135103 | WD-40 repeat protein | elongation zone | 12 h |
| HORVU.MOREX.r3.1HG0015270.1 | 2.158625882 | 5.354810827 | 3.610247774 | 0.00068 | 0.020915275 | -0.60288145 | RRP15-like protein | elongation zone | 12 h |
| HORVU.MOREX.r3.4HG0377830.1 | 0.479953561 | 7.17360811 | 3.827974997 | 0.00034 | 0.017220216 | -0.60381843 | Nuclear cap-binding protein subunit 1 | elongation zone | 12 h |
| HORVU.MOREX.r3.4HG0380360.1 | -0.783196742 | -1.11903663 | -3.641297931 | 0.00062 | 0.020516782 | -0.60777248 | Peroxidase | elongation zone | 12 h |
| HORVU.MOREX.r3.7HG0650300.1 | -0.643332762 | 3.514135404 | -3.739269814 | 0.00045 | 0.018562843 | -0.60842153 | Protein kinase | elongation zone | 12 h |
| HORVU.MOREX.r3.4HG0335750.1 | 1.871155301 | 7.585427625 | 3.735669592 | 0.00046 | 0.018625563 | -0.61113628 | Pescadillo homolog | elongation zone | 12 h |
| HORVU.MOREX.r3.1HG0059770.1 | 0.693039334 | 8.691483921 | 3.864931839 | 0.00031 | 0.016471701 | -0.61133086 | Nascent polypeptide-associated complex subunit alpha-like prot | elongation zone | 12 h |
| HORVU.MOREX.r3.2HG0109970.1 | 0.664064513 | 7.705771322 | 3.830469881 | 0.00034 | 0.017205109 | -0.61423782 | Lon protease homolog | elongation zone | 12 h |
| HORVU.MOREX.r3.5HG0507820.1 | 0.623243424 | 8.522605219 | 3.854439286 | 0.00032 | 0.016689644 | -0.61716243 | Methyl-CpG binding domain containing protein, expressed | elongation zone | 12 h |
| HORVU.MOREX.r3.7HG0638400.1 | 0.999912254 | 3.0032628 | 3.586061436 | 0.00073 | 0.021412967 | -0.62005256 | 30S ribosomal protein S13 | elongation zone | 12 h |
| HORVU.MOREX.r3.1HG0071300.1 | 1.71242917 | 7.640000377 | 3.760547537 | 0.00042 | 0.018118761 | -0.62304888 | DNA helicase | elongation zone | 12 h |
| HORVU.MOREX.r3.5HG0465060.1 | 2.386785555 | 0.671256507 | 3.593242466 | 0.00072 | 0.021178576 | -0.62318215 | Pentatricopeptide repeat-containing family protein | elongation zone | 12 h |
| HORVU.MOREX.r3.UnG0753320.1 | 1.228288324 | 4.78278288 | 3.631976929 | 0.00063 | 0.020646627 | -0.62429652 | Nucleoporin protein Ndc1-Nup | elongation zone | 12 h |
| HORVU.MOREX.r3.7HG0642890.1 | 2.046639053 | 5.571712758 | 3.655557198 | 0.00059 | 0.020437054 | -0.63054682 | Acid phosphatase 1 | elongation zone | 12 h |
| HORVU.MOREX.r3.5HG0489530.1 | 2.793480429 | 0.386435685 | 3.623741246 | 0.00065 | 0.020713759 | -0.63238918 | Pentatricopeptide repeat-containing protein | elongation zone | 12 h |
| HORVU.MOREX.r3.6HG0585150.1 | 0.314629888 | 5.680949913 | 3.778460211 | 0.0004 | 0.017833387 | -0.63449807 | NHL repeat-containing protein | elongation zone | 12 h |
| HORVU.MOREX.r3.2HG0116790.1 | 1.280270779 | 3.25502819 | 3.5612267 | 0.00079 | 0.022039057 | -0.63704295 | E3 ubiquitin-protein ligase | elongation zone | 12 h |
| HORVU.MOREX.r3.2HG0215310.1 | 1.689559112 | 7.090699835 | 3.749375438 | 0.00044 | 0.018313253 | -0.63805518 | Peroxidase | elongation zone | 12 h |
| HORVU.MOREX.r3.4HG0395060.1 | 2.586894382 | 0.480504378 | 3.62599001 | 0.00065 | 0.020713759 | -0.63888005 | #N/A | elongation zone | 12 h |
| HORVU.MOREX.r3.4HG0332620.1 | 2.538542806 | -0.394695086 | 3.595267176 | 0.00071 | 0.021178576 | -0.64038205 | DUF868 family protein (DUF868) | elongation zone | 12 h |
| HORVU.MOREX.r3.3HG0231880.1 | 1.03549124 | 2.985187022 | 3.574196831 | 0.00076 | 0.021740481 | -0.64143974 | Leucine-rich repeat (LRR) family protein | elongation zone | 12 h |
| HORVU.MOREX.r3.4HG0396240.1 | 1.898892028 | 0.669237595 | 3.551247416 | 0.00081 | 0.022303809 | -0.64234881 | Ferredoxin--NAD(P)(+) reductase (naphthalene dioxygenase/salic | elongation zone | 12 h |
| HORVU.MOREX.r3.6HG0589360.1 | 1.246358777 | 7.208700986 | 3.75904465 | 0.00043 | 0.018129366 | -0.64365421 | Nucleosome assembly protein 1-like 1 | elongation zone | 12 h |
| HORVU.MOREX.r3.7HG0703750.1 | 0.886785189 | 6.844981761 | 3.759800942 | 0.00043 | 0.018123749 | -0.65017147 | Programmed cell death protein 4 | elongation zone | 12 h |
| HORVU.MOREX.r3.3HG0288020.1 | 1.124944208 | 3.529030436 | 3.584633173 | 0.00073 | 0.021473531 | -0.65068389 | Tryptophan--tRNA ligase | elongation zone | 12 h |
| HORVU.MOREX.r3.3HG0323600.1 | -0.58841575 | 4.35559343 | -3.76886603 | 0.00041 | 0.017948348 | -0.6510716 | Phosphatidylserine decarboxylase proenzyme 2 | elongation zone | 12 h |
| HORVU.MOREX.r3.1HG0032230.1 | 1.067916356 | 4.67002403 | 3.636560013 | 0.00063 | 0.020577395 | -0.65476163 | Rad3-related DNA helicase family protein | elongation zone | 12 h |
| HORVU.MOREX.r3.6HG0573870.1 | 0.705197149 | 4.150119935 | 3.649157157 | 0.0006 | 0.020507565 | -0.65535517 | Histone H3 | elongation zone | 12 h |
| HORVU.MOREX.r3.3HG0237870.1 | 1.990211986 | 5.343811141 | 3.627558375 | 0.00064 | 0.020713759 | -0.65617286 | Histone-lysine N-methyltransferase | elongation zone | 12 h |
| HORVU.MOREX.r3.3HG0260150.1 | 1.175350512 | -0.115050093 | 3.530873509 | 0.00087 | 0.022907481 | -0.66031948 | Acylamino-acid-releasing enzyme | elongation zone | 12 h |

**Table S5** Continued.

| ID | logFC | AveExpr | t | P.Value | adj.P.Val | B | gene description | root zone | time point |
|---|---|---|---|---|---|---|---|---|---|
| HORVU.MOREX.r3.3HG0286530.1 | -1.381196215 | -2.561378469 | -3.534879354 | 0.00086 | 0.022899718 | -0.66307293 | Transposon protein, putative, mutator sub-class | elongation zone | 12 h |
| HORVU.MOREX.r3.3HG0278540.1 | -0.434545725 | 6.253662754 | -3.815891163 | 0.00036 | 0.017465122 | -0.66566399 | Hydroxyacylglutathione hydrolase | elongation zone | 12 h |
| HORVU.MOREX.r3.4HG0401040.1 | 2.549656401 | 6.811727505 | 3.640146058 | 0.00062 | 0.020516782 | -0.66900757 | Expansin | elongation zone | 12 h |
| HORVU.MOREX.r3.3HG0308530.1 | 0.896321964 | 4.112501755 | 3.596985871 | 0.00071 | 0.021178576 | -0.66944322 | Histone deacetylase, putative | elongation zone | 12 h |
| HORVU.MOREX.r3.3HG0316000.1 | 0.836221039 | 6.5484869 | 3.752352936 | 0.00044 | 0.018288218 | -0.67065001 | Salicylate O-methyltransferase | elongation zone | 12 h |
| HORVU.MOREX.r3.6HG0552190.1 | 1.343656741 | 4.860782531 | 3.605572036 | 0.00069 | 0.021013719 | -0.67108719 | Ribose-phosphate pyrophosphokinase family protein | elongation zone | 12 h |
| HORVU.MOREX.r3.6HG0551860.1 | 0.58560237 | 7.532070736 | 3.807045214 | 0.00037 | 0.017580172 | -0.67120079 | Eukaryotic translation initiation factor 3 subunit F | elongation zone | 12 h |
| HORVU.MOREX.r3.2HG0214200.1 | -0.735289243 | 1.301757812 | -3.777673881 | 0.0004 | 0.017833387 | -0.67155456 | Dihydroflavonol 4-reductase | elongation zone | 12 h |
| HORVU.MOREX.r3.6HG0634260.1 | 0.846884139 | 4.589475239 | 3.639723874 | 0.00062 | 0.020516782 | -0.67745659 | TSA: Wollemia nobilis Ref_Wollemi_Transcript_29242_1275 tran | elongation zone | 12 h |
| HORVU.MOREX.r3.5HG0500680.1 | 0.740789255 | 6.785876601 | 3.782640615 | 0.0004 | 0.017833387 | -0.6792495 | Acyl carrier protein | elongation zone | 12 h |
| HORVU.MOREX.r3.7HG0684300.1 | 1.344741376 | 5.071242343 | 3.630431686 | 0.00064 | 0.020706437 | -0.68072537 | Bifunctional uridylyltransferase/uridylyl-removing enzyme | elongation zone | 12 h |
| HORVU.MOREX.r3.2HG0207380.1 | 1.765994394 | 4.363598916 | 3.562883522 | 0.00079 | 0.022017161 | -0.68744243 | Double-strand break repair protein MRE11 | elongation zone | 12 h |
| HORVU.MOREX.r3.1HG0065100.1 | 1.664313313 | 7.852856437 | 3.75046428 | 0.00044 | 0.018313253 | -0.68778342 | DNA helicase | elongation zone | 12 h |
| HORVU.MOREX.r3.6HG0552180.1 | 1.143322381 | 5.108310854 | 3.63923938 | 0.00062 | 0.020516782 | -0.69065943 | Pentatricopeptide repeat-containing protein | elongation zone | 12 h |
| HORVU.MOREX.r3.6HG0564520.1 | 1.297245436 | 5.49826859 | 3.635741221 | 0.00063 | 0.020596683 | -0.6966649 | Single-stranded DNA-binding protein WHY3 | elongation zone | 12 h |
| HORVU.MOREX.r3.7HG0676160.1 | 1.306034251 | 6.283730528 | 3.694505564 | 0.00052 | 0.019449812 | -0.70549152 | Mitochondrial glycoprotein-like | elongation zone | 12 h |
| HORVU.MOREX.r3.3HG0295530.1 | 1.92527103 | 3.813404323 | 3.516147947 | 0.00091 | 0.023235104 | -0.70607561 | UDP-glycosyltransferase | elongation zone | 12 h |
| HORVU.MOREX.r3.3HG0296610.1 | 1.325569062 | 2.287341464 | 3.51379045 | 0.00091 | 0.023235104 | -0.70704702 | Pentatricopeptide repeat-containing protein | elongation zone | 12 h |
| HORVU.MOREX.r3.5HG0496170.1 | 0.968711547 | 5.175044795 | 3.654214434 | 0.00059 | 0.020437054 | -0.71618303 | RNA polymerase II transcriptional coactivator | elongation zone | 12 h |
| HORVU.MOREX.r3.1HG0013020.1 | 2.655846757 | 0.328708637 | 3.576280844 | 0.00075 | 0.021691049 | -0.71685901 | Pentatricopeptide repeat-containing protein | elongation zone | 12 h |
| HORVU.MOREX.r3.7HG0650970.1 | 1.096035255 | 7.468679104 | 3.766050822 | 0.00042 | 0.017994261 | -0.71826672 | Histone H3 | elongation zone | 12 h |
| HORVU.MOREX.r3.5HG0467500.1 | 1.873668171 | 3.330218247 | 3.516319512 | 0.00091 | 0.023235104 | -0.71908279 | Pseudouridine synthase family protein | elongation zone | 12 h |
| HORVU.MOREX.r3.3HG0304420.1 | 1.219362598 | 5.851634764 | 3.693286143 | 0.00052 | 0.019449812 | -0.72045741 | #N/A | elongation zone | 12 h |
| HORVU.MOREX.r3.1HG0053940.1 | 1.447428463 | 5.672823401 | 3.629890158 | 0.00064 | 0.020708432 | -0.72065532 | stress response protein | elongation zone | 12 h |
| HORVU.MOREX.r3.1HG0077010.1 | 0.917491844 | 4.063866452 | 3.594361948 | 0.00071 | 0.021178576 | -0.72167211 | Transcriptional regulatory plant protein, putative | elongation zone | 12 h |
| HORVU.MOREX.r3.1HG0023610.1 | 1.209616487 | 5.36944383 | 3.675654945 | 0.00055 | 0.019889362 | -0.72272311 | #N/A | elongation zone | 12 h |
| HORVU.MOREX.r3.2HG0121130.1 | 2.737765063 | 2.407479747 | 3.51595798 | 0.00091 | 0.023235104 | -0.72320185 | Pentatricopeptide repeat-containing protein | elongation zone | 12 h |
| HORVU.MOREX.r3.7HG0737460.1 | 1.244885953 | 4.569245525 | 3.576174427 | 0.00075 | 0.021691049 | -0.72332934 | Peroxidase | elongation zone | 12 h |
| HORVU.MOREX.r3.1HG0075430.1 | 2.095082013 | 4.723898757 | 3.542456467 | 0.00084 | 0.022605629 | -0.72495976 | Amidophosphoribosyltransferase | elongation zone | 12 h |
| HORVU.MOREX.r3.7HG0703170.1 | 1.389539465 | 4.025119613 | 3.543648012 | 0.00083 | 0.022553062 | -0.72533215 | BTB/POZ and MATH domain-containing protein 2 | elongation zone | 12 h |
| HORVU.MOREX.r3.7HG0704180.1 | 0.67134139 | 6.317889073 | 3.725966277 | 0.00047 | 0.018928454 | -0.72596677 | COP9 signalosome complex subunit 3 | elongation zone | 12 h |
| HORVU.MOREX.r3.3HG0316210.1 | 1.848147722 | 7.250167183 | 3.691100706 | 0.00053 | 0.019449812 | -0.7266567 | Basic 7S globulin 2 | elongation zone | 12 h |
| HORVU.MOREX.r3.5HG0513920.1 | 1.353469816 | 4.916144619 | 3.594546702 | 0.00071 | 0.021178576 | -0.72706557 | NADPH--cytochrome P450 reductase | elongation zone | 12 h |
| HORVU.MOREX.r3.2HG0211230.1 | 1.884151653 | 1.198269179 | 3.515718181 | 0.00091 | 0.023235104 | -0.72724164 | Pentatricopeptide repeat-containing protein | elongation zone | 12 h |
| HORVU.MOREX.r3.1HG0005590.1 | 0.725566297 | 3.576430878 | 3.596957576 | 0.00071 | 0.021178576 | -0.72842318 | Mitochondrial import inner membrane translocase subunit tim2: | elongation zone | 12 h |
| HORVU.MOREX.r3.2HG0170060.1 | 1.237013122 | 5.035774752 | 3.617125895 | 0.00066 | 0.020839563 | -0.72851056 | Protein TIC 40, chloroplastic | elongation zone | 12 h |
| HORVU.MOREX.r3.4HG0417410.1 | 1.397608728 | 0.91597262 | 3.508273205 | 0.00093 | 0.023255319 | -0.73079491 | Cold regulated protein 27 | elongation zone | 12 h |
| HORVU.MOREX.r3.2HG0204590.1 | 1.106220574 | 4.606565659 | 3.610247555 | 0.00068 | 0.020915275 | -0.73091601 | Receptor protein kinase, putative | elongation zone | 12 h |
| HORVU.MOREX.r3.6HG0616490.1 | 1.378969231 | 4.602188135 | 3.580669907 | 0.00074 | 0.021613876 | -0.73314051 | Peptidyl-prolyl cis-trans isomerase | elongation zone | 12 h |
| HORVU.MOREX.r3.4HG0343330.1 | 4.629059127 | -0.333480115 | 3.529919452 | 0.00087 | 0.022944774 | -0.73355675 | Endonuclease III homolog | elongation zone | 12 h |
| HORVU.MOREX.r3.5HG0526830.1 | 0.416496166 | 5.623951381 | 3.72128394 | 0.00048 | 0.019002693 | -0.73431601 | Hsp70 nucleotide exchange factor fes1 | elongation zone | 12 h |

**Table S5** Continued.

| ID | logFC | AveExpr | t | P.Value | adj.P.Val | B | gene description | root zone | time point |
|---|---|---|---|---|---|---|---|---|---|
| HORVU.MOREX.r3.7HG0749870.1 | -0.633155909 | 4.569999785 | -3.782156275 | 0.0004 | 0.017833387 | -0.73521981 | Glucuronoxylan 4-O-methyltransferase | elongation zone | 12 h |
| HORVU.MOREX.r3.3HG0267680.1 | 2.728600722 | 4.551509802 | 3.51458863 | 0.00091 | 0.023235104 | -0.74340314 | UBP1-associated proteins 1C | elongation zone | 12 h |
| HORVU.MOREX.r3.1HG0063560.1 | 2.214197615 | -0.256820188 | 3.532154967 | 0.00086 | 0.022899718 | -0.74636583 | Sulfotransferase | elongation zone | 12 h |
| HORVU.MOREX.r3.4HG0338270.1 | 2.127970295 | 7.140700386 | 3.654563815 | 0.00059 | 0.020437054 | -0.74660103 | ATP-dependent DNA helicase DDM1 | elongation zone | 12 h |
| HORVU.MOREX.r3.2HG0173240.1 | 0.905322125 | 4.646167723 | 3.605135315 | 0.00069 | 0.021013719 | -0.75021337 | Nucleosome assembly protein 1-like 1 | elongation zone | 12 h |
| HORVU.MOREX.r3.1HG0039710.1 | 0.980529664 | 3.112736137 | 3.53192345 | 0.00086 | 0.022899718 | -0.75102951 | Protein fluG | elongation zone | 12 h |
| HORVU.MOREX.r3.2HG0161020.1 | 0.899356595 | 7.262436364 | 3.770969004 | 0.00041 | 0.017914792 | -0.75125719 | #N/A | elongation zone | 12 h |
| HORVU.MOREX.r3.6HG0627970.1 | 0.70359547 | 6.26071971 | 3.734035243 | 0.00046 | 0.018667357 | -0.75414021 | #N/A | elongation zone | 12 h |
| HORVU.MOREX.r3.2HG0180580.1 | 1.801057555 | 7.018787121 | 3.680355735 | 0.00055 | 0.019711469 | -0.75475998 | Proliferating cell nuclear antigen | elongation zone | 12 h |
| HORVU.MOREX.r3.6HG0624620.1 | 1.571442515 | 7.088282878 | 3.693291944 | 0.00052 | 0.019449812 | -0.75677566 | DNA helicase | elongation zone | 12 h |
| HORVU.MOREX.r3.2HG0108490.1 | 1.344716739 | 3.170050036 | 3.548906472 | 0.00082 | 0.022369939 | -0.75877838 | Xyloglucan endotransglucosylase/hydrolase | elongation zone | 12 h |
| HORVU.MOREX.r3.5HG0529140.1 | 0.413909922 | 10.35930424 | 3.906179673 | 0.00027 | 0.015692177 | -0.76045175 | Elongation factor | elongation zone | 12 h |
| HORVU.MOREX.r3.7HG0665990.1 | -0.666005138 | 6.15668184 | -3.833923275 | 0.00034 | 0.017059301 | -0.76399587 | Shikimate kinase | elongation zone | 12 h |
| HORVU.MOREX.r3.5HG0460900.1 | 0.967734088 | 6.775177122 | 3.721196446 | 0.00048 | 0.019002693 | -0.76430828 | 2-isopropylmalate synthase | elongation zone | 12 h |
| HORVU.MOREX.r3.5HG0534510.1 | 0.962585172 | 7.454533079 | 3.744172722 | 0.00045 | 0.018462873 | -0.76544312 | Nascent polypeptide-associated complex subunit beta | elongation zone | 12 h |
| HORVU.MOREX.r3.4HG0401740.1 | 2.406224252 | 1.996606862 | 3.49947913 | 0.00095 | 0.023511243 | -0.76827912 | Eukaryotic translation initiation factor 3 subunit H | elongation zone | 12 h |
| HORVU.MOREX.r3.2HG0172050.1 | 0.450273143 | 8.585873782 | 3.822945782 | 0.00035 | 0.017322598 | -0.77184881 | RNA-binding protein | elongation zone | 12 h |
| HORVU.MOREX.r3.3HG0306420.1 | 0.922603897 | 3.782406003 | 3.575847243 | 0.00075 | 0.021691049 | -0.77640611 | TPR repeat-containing protein ZIP4 | elongation zone | 12 h |
| HORVU.MOREX.r3.6HG0556980.1 | 1.789584424 | 5.184914753 | 3.557528695 | 0.0008 | 0.022169819 | -0.78264814 | RNA binding protein | elongation zone | 12 h |
| HORVU.MOREX.r3.1HG0086110.1 | 1.163300713 | 5.469120462 | 3.62493285 | 0.00065 | 0.020713759 | -0.78545373 | 50S ribosomal protein L21 | elongation zone | 12 h |
| HORVU.MOREX.r3.2HG0130960.1 | -0.751241114 | 1.762022677 | -3.558003947 | 0.0008 | 0.022167528 | -0.78716425 | Benzene 1,2-dioxygenase subunit alpha | elongation zone | 12 h |
| HORVU.MOREX.r3.4HG0334580.1 | 1.904393791 | 3.64777309 | 3.50394945 | 0.00094 | 0.023362135 | -0.78733196 | #N/A | elongation zone | 12 h |
| HORVU.MOREX.r3.7HG0690370.1 | 0.844051772 | 2.109174244 | 3.501530999 | 0.00095 | 0.023426471 | -0.78937633 | Malate dehydrogenase | elongation zone | 12 h |
| HORVU.MOREX.r3.2HG0106340.1 | 2.716736448 | 1.552213778 | 3.512392812 | 0.00092 | 0.023235104 | -0.78978037 | Pentatricopeptide repeat-containing protein | elongation zone | 12 h |
| HORVU.MOREX.r3.5HG0505140.1 | -0.547640677 | 6.833596665 | -3.81654762 | 0.00036 | 0.017465122 | -0.78989101 | Phosphatase 2C family protein | elongation zone | 12 h |
| HORVU.MOREX.r3.7HG0735200.1 | 0.449670967 | 10.46904646 | 3.892377852 | 0.00028 | 0.015751512 | -0.7901656 | Elongation factor | elongation zone | 12 h |
| HORVU.MOREX.r3.2HG0152140.1 | 0.688200135 | 5.200813775 | 3.645652475 | 0.00061 | 0.020507565 | -0.79127253 | D-amino acid dehydrogenase | elongation zone | 12 h |
| HORVU.MOREX.r3.1HG0067720.1 | -1.057154991 | -2.106023761 | -3.569687492 | 0.00077 | 0.021876787 | -0.79211567 | #N/A | elongation zone | 12 h |
| HORVU.MOREX.r3.1HG0020280.1 | 0.839107842 | 7.185819282 | 3.736351811 | 0.00046 | 0.018625563 | -0.79417599 | Pentatricopeptide repeat superfamily protein | elongation zone | 12 h |
| HORVU.MOREX.r3.3HG0281850.1 | 0.971727967 | 5.846586489 | 3.645411502 | 0.00061 | 0.020507565 | -0.79487605 | Threonine synthase | elongation zone | 12 h |
| HORVU.MOREX.r3.7HG0684630.1 | 1.250228158 | 6.041630602 | 3.645869628 | 0.00061 | 0.020507565 | -0.79719195 | DAG protein, chloroplastic | elongation zone | 12 h |
| HORVU.MOREX.r3.1HG0069800.1 | 2.350056869 | 0.744516388 | 3.532762613 | 0.00086 | 0.022899718 | -0.7982619 | Histone-lysine N-methyltransferase, H3 lysine-9 specific | elongation zone | 12 h |
| HORVU.MOREX.r3.7HG0680060.1 | 2.095002975 | 2.004315332 | 3.482070341 | 0.00101 | 0.024008795 | -0.7983037 | Mitochondrial transcription termination factor family protein, pu | elongation zone | 12 h |
| HORVU.MOREX.r3.5HG0532630.1 | 1.471979904 | 4.235353088 | 3.551177825 | 0.00081 | 0.022303809 | -0.80051171 | Sister chromatid cohesion protein DCC1 | elongation zone | 12 h |
| HORVU.MOREX.r3.2HG0155140.1 | 1.432451139 | 3.347249234 | 3.490547967 | 0.00098 | 0.023786521 | -0.80056725 | tRNA dimethylallyltransferase | elongation zone | 12 h |
| HORVU.MOREX.r3.2HG0127160.1 | 1.895807616 | 2.05404778 | 3.47598243 | 0.00102 | 0.024166531 | -0.80077995 | WD40 repeat-like protein | elongation zone | 12 h |
| HORVU.MOREX.r3.2HG0163090.1 | 1.019459254 | 7.015883321 | 3.71114994 | 0.0005 | 0.019227921 | -0.80488218 | Heavy metal-associated protein | elongation zone | 12 h |
| HORVU.MOREX.r3.6HG0541690.1 | 2.219268074 | -0.276490972 | 3.518910329 | 0.0009 | 0.023235104 | -0.80778142 | Mitochondrial transcription termination factor-like | elongation zone | 12 h |
| HORVU.MOREX.r3.4HG0406380.1 | 0.475265826 | 5.163918803 | 3.672727036 | 0.00056 | 0.019889362 | -0.81255923 | GEM-like protein 1 | elongation zone | 12 h |
| HORVU.MOREX.r3.5HG0468290.1 | 0.973879791 | 9.889642173 | 3.819167586 | 0.00035 | 0.017458382 | -0.81476109 | Polyadenylate-binding protein | elongation zone | 12 h |
| HORVU.MOREX.r3.2HG0119360.1 | 1.93374086 | -1.10964552 | 3.472776936 | 0.00103 | 0.024215578 | -0.81795375 | Gibberellin receptor GID1A | elongation zone | 12 h |

**Table S5** Continued.

| ID | logFC | AveExpr | t | P.Value | adj.P.Val | B | gene description | root zone | time point |
|---|---|---|---|---|---|---|---|---|---|
| HORVU.MOREX.r3.4HG0342990.1 | 1.626563212 | 4.752270826 | 3.555318129 | 0.0008 | 0.022260348 | -0.82011429 | Condensin-2 complex subunit G2 | elongation zone | 12 h |
| HORVU.MOREX.r3.1HG0022460.1 | 0.484459438 | 7.308294084 | 3.74672678 | 0.00044 | 0.018395923 | -0.82114811 | Transport inhibitor response 1 | elongation zone | 12 h |
| HORVU.MOREX.r3.4HG0402600.2 | 1.120008014 | 2.451455995 | 3.482830803 | 0.001 | 0.023981387 | -0.82215209 | 5-dehydro-2-deoxygluconokinase | elongation zone | 12 h |
| HORVU.MOREX.r3.3HG0323490.1 | 1.214523625 | 4.264274151 | 3.533212071 | 0.00086 | 0.022899718 | -0.8245389 | DNA mismatch repair protein MutL | elongation zone | 12 h |
| HORVU.MOREX.r3.6HG0588300.1 | 1.627355075 | 2.046655197 | 3.466147534 | 0.00105 | 0.024538512 | -0.8262988 | #N/A | elongation zone | 12 h |
| HORVU.MOREX.r3.3HG0291380.1 | 2.328969179 | 4.361758282 | 3.488805179 | 0.00098 | 0.023815524 | -0.82670011 | Peptidoglycan-recognition protein SB2 | elongation zone | 12 h |
| HORVU.MOREX.r3.2HG0125870.1 | 0.679690857 | 9.837842611 | 3.836127739 | 0.00033 | 0.017055324 | -0.82718556 | 50S ribosomal protein L15 | elongation zone | 12 h |
| HORVU.MOREX.r3.2HG0173960.1 | -0.72336846 | 2.814914619 | -3.571208587 | 0.00077 | 0.021819151 | -0.82732264 | Peroxidase | elongation zone | 12 h |
| HORVU.MOREX.r3.2HG0177450.1 | 1.270710518 | 4.715585642 | 3.565784884 | 0.00078 | 0.021941854 | -0.82895407 | Ribosome-recycling factor | elongation zone | 12 h |
| HORVU.MOREX.r3.7HG0750000.1 | -1.153031864 | -2.120414367 | -3.5522524 | 0.00081 | 0.022303809 | -0.83386982 | Disease resistance protein | elongation zone | 12 h |
| HORVU.MOREX.r3.5HG0502580.1 | 1.764688339 | 4.788366911 | 3.520741339 | 0.00089 | 0.023235104 | -0.83724914 | Regulator of Vps4 activity in the MVB pathway protein | elongation zone | 12 h |
| HORVU.MOREX.r3.2HG0119970.1 | 3.117197447 | 3.337857733 | 3.458811692 | 0.00108 | 0.024669554 | -0.84539319 | Heat shock transcription factor | elongation zone | 12 h |
| HORVU.MOREX.r3.3HG0313070.1 | 1.975475577 | 6.095175499 | 3.604997811 | 0.00069 | 0.021013719 | -0.8476503 | #N/A | elongation zone | 12 h |
| HORVU.MOREX.r3.5HG0467900.1 | 3.109191459 | 1.256467408 | 3.511574702 | 0.00092 | 0.023235104 | -0.84933997 | Cyclin | elongation zone | 12 h |
| HORVU.MOREX.r3.1HG0032910.1 | 0.658230142 | 5.350158872 | 3.666206083 | 0.00057 | 0.020121419 | -0.8569504 | Lysine-specific demethylase | elongation zone | 12 h |
| HORVU.MOREX.r3.2HG0204440.1 | 0.886612016 | 4.696803122 | 3.572602382 | 0.00076 | 0.021766161 | -0.85755461 | 4-hydroxy-tetrahydrodipicolinate synthase | elongation zone | 12 h |
| HORVU.MOREX.r3.3HG0236000.1 | 0.434116877 | 6.001989313 | 3.692931383 | 0.00053 | 0.019449812 | -0.85866516 | Ectonucleotide pyrophosphatase/phosphodiesterase, putative | elongation zone | 12 h |
| HORVU.MOREX.r3.5HG0478280.1 | 1.775642454 | 3.919076773 | 3.48416512 | 0.001 | 0.023938404 | -0.85916159 | ATP-dependent RNA helicase | elongation zone | 12 h |
| HORVU.MOREX.r3.2HG0113180.1 | 0.789913887 | 4.229499943 | 3.566324392 | 0.00078 | 0.021941854 | -0.86158599 | DNA-directed RNA polymerases I, II, and III subunit rpabc3 | elongation zone | 12 h |
| HORVU.MOREX.r3.2HG0186750.1 | 2.547989855 | 1.34331984 | 3.490151936 | 0.00098 | 0.023786999 | -0.86344391 | Growth-regulating factor | elongation zone | 12 h |
| HORVU.MOREX.r3.UnG0765280.1 | 1.289981496 | 2.589811572 | 3.478150622 | 0.00102 | 0.024127373 | -0.86546819 | F-box family protein | elongation zone | 12 h |
| HORVU.MOREX.r3.7HG0752350.1 | 2.502256874 | 3.186534982 | 3.450523484 | 0.00111 | 0.025099251 | -0.86689727 | AT hook motif DNA-binding family protein | elongation zone | 12 h |
| HORVU.MOREX.r3.7HG0741780.1 | 0.952452873 | 6.389609859 | 3.674522214 | 0.00056 | 0.019889362 | -0.86792621 | 40S ribosomal protein S27 | elongation zone | 12 h |
| HORVU.MOREX.r3.5HG0522140.1 | 0.863971331 | 5.949304073 | 3.645755854 | 0.00061 | 0.020507565 | -0.86893197 | Cleavage and polyadenylation specificity factor subunit 2 | elongation zone | 12 h |
| HORVU.MOREX.r3.6HG0572260.1 | 2.26777119 | -0.289254551 | 3.489084156 | 0.00098 | 0.023815524 | -0.87631201 | Exocyst complex component SEC3A | elongation zone | 12 h |
| HORVU.MOREX.r3.7HG0672900.1 | 1.995551471 | 0.425915719 | 3.466538142 | 0.00105 | 0.02453734 | -0.8770273 | Pentatricopeptide repeat-containing protein | elongation zone | 12 h |
| HORVU.MOREX.r3.2HG0113530.1 | 1.440138181 | 3.744005308 | 3.475714253 | 0.00102 | 0.024166531 | -0.8772812 | THUMP domain-containing protein 1 like | elongation zone | 12 h |
| HORVU.MOREX.r3.6HG0567330.1 | 1.262892859 | 5.748868839 | 3.615181719 | 0.00067 | 0.020839563 | -0.88212548 | aspartyl/glutamyl-tRNA(Asn/Gln) amidotransferase subunit | elongation zone | 12 h |
| HORVU.MOREX.r3.2HG0194130.1 | 0.993908903 | 5.878463741 | 3.613339417 | 0.00067 | 0.020873129 | -0.88452115 | Glycosyltransferase | elongation zone | 12 h |
| HORVU.MOREX.r3.7HG0724060.1 | -1.225376035 | 0.588187119 | -3.624484261 | 0.00065 | 0.020713759 | -0.88507511 | U-box domain-containing protein | elongation zone | 12 h |
| HORVU.MOREX.r3.5HG0438300.1 | 3.028000787 | -0.03895989 | 3.483806422 | 0.001 | 0.023938404 | -0.88717202 | Protein FAR1-RELATED SEQUENCE 5 | elongation zone | 12 h |
| HORVU.MOREX.r3.2HG0172190.1 | 2.096060395 | 1.141861715 | 3.454723216 | 0.00109 | 0.024865394 | -0.88989135 | Meiosis-specific with OB domain-containing protein | elongation zone | 12 h |
| HORVU.MOREX.r3.5HG0493060.1 | 0.811596358 | 7.550677769 | 3.716429376 | 0.00049 | 0.019131832 | -0.89159709 | 60S ribosomal protein L32 | elongation zone | 12 h |
| HORVU.MOREX.r3.1HG0031730.1 | 1.232462337 | 7.573338482 | 3.696258422 | 0.00052 | 0.019449812 | -0.89174928 | DNA helicase | elongation zone | 12 h |
| HORVU.MOREX.r3.1HG0073600.1 | 1.287806815 | 3.523923593 | 3.474247022 | 0.00103 | 0.024201764 | -0.89187327 | #N/A | elongation zone | 12 h |
| HORVU.MOREX.r3.2HG0204920.1 | 1.186495269 | 3.638603207 | 3.485849927 | 0.00099 | 0.023901967 | -0.89241444 | Crossover junction endonuclease EME1B | elongation zone | 12 h |
| HORVU.MOREX.r3.3HG0232070.1 | 2.076589318 | 5.516136915 | 3.533270699 | 0.00086 | 0.022899718 | -0.89945128 | Protein O-linked-mannose beta-1,4-N-acetylglucosaminyltransfe | elongation zone | 12 h |
| HORVU.MOREX.r3.3HG0325830.1 | 2.963445661 | 1.73334501 | 3.457962177 | 0.00108 | 0.024705447 | -0.90468215 | Uridylate kinase | elongation zone | 12 h |
| HORVU.MOREX.r3.7HG0740180.1 | 1.357187766 | 5.365923975 | 3.562414063 | 0.00079 | 0.022018823 | -0.90631132 | MMS19 nucleotide excision repair protein-like protein | elongation zone | 12 h |
| HORVU.MOREX.r3.2HG0118530.1 | 1.81166156 | 5.0118675 | 3.504236396 | 0.00094 | 0.023362135 | -0.907091 | ATP-dependent RNA helicase | elongation zone | 12 h |
| HORVU.MOREX.r3.3HG0303330.1 | 0.887815573 | 6.555668972 | 3.68280774 | 0.00054 | 0.019711469 | -0.90821332 | #N/A | elongation zone | 12 h |

**Table S5** Continued.

| ID | logFC | AveExpr | t | P.Value | adj.P.Val | B | gene description | root zone | time point |
|---|---|---|---|---|---|---|---|---|---|
| HORVU.MOREX.r3.2HG0125550.1 | 0.647263091 | 8.919649801 | 3.779174832 | 0.0004 | 0.017833387 | -0.90863472 | Elongation factor 1-beta | elongation zone | 12 h |
| HORVU.MOREX.r3.2HG0179940.1 | 1.006834788 | 5.507339313 | 3.600532801 | 0.0007 | 0.021178576 | -0.90958903 | #N/A | elongation zone | 12 h |
| HORVU.MOREX.r3.1HG0066530.1 | 1.129152114 | 4.906097437 | 3.541231878 | 0.00084 | 0.022660635 | -0.91056253 | DUF630 family protein, putative (DUF630 and DUF632) | elongation zone | 12 h |
| HORVU.MOREX.r3.5HG0514490.1 | 0.599303615 | 5.559766109 | 3.64757304 | 0.0006 | 0.020507565 | -0.91689691 | Hexosyltransferase | elongation zone | 12 h |
| HORVU.MOREX.r3.5HG0514100.1 | 0.651308964 | 4.963873023 | 3.604860016 | 0.00069 | 0.021013719 | -0.91986501 | #N/A | elongation zone | 12 h |
| HORVU.MOREX.r3.7HG0659670.1 | 1.28852847 | 5.414107691 | 3.554459596 | 0.00081 | 0.022275873 | -0.92097586 | Nucleoporin NUP188 | elongation zone | 12 h |
| HORVU.MOREX.r3.7HG0674840.1 | 2.741855128 | 0.633419814 | 3.478848895 | 0.00101 | 0.024104211 | -0.92220056 | #N/A | elongation zone | 12 h |
| HORVU.MOREX.r3.6HG0622440.1 | 0.572347423 | 6.187280102 | 3.659992255 | 0.00058 | 0.02036338 | -0.92327465 | Protein FLX-like 1 | elongation zone | 12 h |
| HORVU.MOREX.r3.6HG0591270.1 | -0.345735511 | 5.135492747 | -3.689968114 | 0.00053 | 0.019449812 | -0.92521127 | Vacuolar protein sorting-associated protein 20 | elongation zone | 12 h |
| HORVU.MOREX.r3.2HG0183740.1 | 1.046224556 | 5.211569422 | 3.56448663 | 0.00078 | 0.021955355 | -0.92911056 | Sugar transporter, putative | elongation zone | 12 h |
| HORVU.MOREX.r3.1HG0088350.1 | 2.567967827 | -0.24071049 | 3.497833905 | 0.00096 | 0.02354153 | -0.93566289 | F-box protein (DUF295) | elongation zone | 12 h |
| HORVU.MOREX.r3.5HG0428840.1 | 0.759195213 | 5.358223693 | 3.60139784 | 0.0007 | 0.021176603 | -0.93606715 | BTB/POZ domain-containing protein | elongation zone | 12 h |
| HORVU.MOREX.r3.6HG0616300.1 | -0.557978062 | 5.182691489 | -3.721608299 | 0.00048 | 0.019002693 | -0.93796601 | Histidine kinase | elongation zone | 12 h |
| HORVU.MOREX.r3.1HG0080320.1 | 1.738536443 | 1.65312479 | 3.426508229 | 0.00119 | 0.025731015 | -0.9383112 | IQ domain-containing protein | elongation zone | 12 h |
| HORVU.MOREX.r3.6HG0626050.1 | 1.084176504 | 4.114338477 | 3.533583557 | 0.00086 | 0.022899718 | -0.94528891 | RNA binding | elongation zone | 12 h |
| HORVU.MOREX.r3.5HG0471400.1 | 2.303806854 | 3.16323989 | 3.42165568 | 0.00121 | 0.025782931 | -0.94659532 | QWRF motif protein (DUF566) | elongation zone | 12 h |
| HORVU.MOREX.r3.6HG0540980.1 | 1.756296565 | 4.62525369 | 3.46570845 | 0.00106 | 0.024543314 | -0.94956655 | Trihelix transcription factor GT-2 | elongation zone | 12 h |
| HORVU.MOREX.r3.5HG0529120.1 | 0.785666683 | 7.801286727 | 3.705841279 | 0.0005 | 0.019342524 | -0.95118095 | Elongation factor | elongation zone | 12 h |
| HORVU.MOREX.r3.2HG0158900.1 | 1.629803963 | 1.613344832 | 3.41725039 | 0.00122 | 0.02596429 | -0.9526706 | Ribonuclease P/MRP protein subunit POP5 | elongation zone | 12 h |
| HORVU.MOREX.r3.1HG0077170.1 | 0.702719956 | 4.830924904 | 3.569120098 | 0.00077 | 0.021876787 | -0.95579365 | Protein EMBRYONIC FLOWER 1 | elongation zone | 12 h |
| HORVU.MOREX.r3.6HG0616480.1 | 0.590987208 | 5.832985524 | 3.655962644 | 0.00059 | 0.020437054 | -0.95655354 | #N/A | elongation zone | 12 h |
| HORVU.MOREX.r3.4HG0401320.1 | -0.741179368 | 0.708055432 | -3.520151491 | 0.0009 | 0.023235104 | -0.96039096 | Tetraspanin | elongation zone | 12 h |
| HORVU.MOREX.r3.1HG0044670.1 | 1.500055821 | 5.661428894 | 3.552384268 | 0.00081 | 0.022303809 | -0.96638823 | Pentatricopeptide repeat-containing protein | elongation zone | 12 h |
| HORVU.MOREX.r3.7HG0714570.1 | 1.188959822 | 4.671992391 | 3.513028138 | 0.00091 | 0.023235104 | -0.96716648 | 50S ribosomal protein L19, putative | elongation zone | 12 h |
| HORVU.MOREX.r3.4HG0404110.1 | 0.956882915 | 7.328767239 | 3.666978988 | 0.00057 | 0.020107621 | -0.96720693 | Adenylosuccinate synthetase | elongation zone | 12 h |
| HORVU.MOREX.r3.6HG0552550.1 | 1.714292627 | 4.44139878 | 3.459234121 | 0.00108 | 0.024665565 | -0.96894145 | Upstream activation factor subunit spp27 | elongation zone | 12 h |
| HORVU.MOREX.r3.3HG0298200.1 | 2.006165546 | 3.96464743 | 3.428566098 | 0.00118 | 0.025704276 | -0.97682762 | Cobalt ion binding | elongation zone | 12 h |
| HORVU.MOREX.r3.3HG0236810.1 | 2.740145429 | 3.500823851 | 3.414553898 | 0.00123 | 0.026021602 | -0.9776118 | #N/A | elongation zone | 12 h |
| HORVU.MOREX.r3.5HG0523860.1 | -0.610291748 | 4.167580678 | -3.637505378 | 0.00062 | 0.020550106 | -0.97803048 | Calcium-dependent protein kinase | elongation zone | 12 h |
| HORVU.MOREX.r3.3HG0322190.1 | 1.081504368 | 4.857803386 | 3.531083939 | 0.00087 | 0.022907481 | -0.97807547 | 30S ribosomal protein S11 | elongation zone | 12 h |
| HORVU.MOREX.r3.6HG0543780.1 | 0.808781272 | 7.397821422 | 3.707600353 | 0.0005 | 0.019342524 | -0.97852186 | Histone H2B | elongation zone | 12 h |
| HORVU.MOREX.r3.6HG0606460.1 | 1.645821798 | 7.91527308 | 3.65409772 | 0.00059 | 0.020437054 | -0.98121405 | DNA topoisomerase 2 | elongation zone | 12 h |
| HORVU.MOREX.r3.2HG0206380.1 | 0.911940865 | 5.584402928 | 3.564244859 | 0.00078 | 0.021955355 | -0.98466227 | DNA-directed RNA polymerase subunit beta | elongation zone | 12 h |
| HORVU.MOREX.r3.5HG0466780.1 | 0.915659588 | 7.10069964 | 3.648197127 | 0.0006 | 0.020507565 | -0.99139738 | Carbamoyl-phosphate synthase small chain | elongation zone | 12 h |
| HORVU.MOREX.r3.7HG0719410.1 | 1.02288595 | 6.134815643 | 3.583676864 | 0.00074 | 0.021475647 | -0.99235335 | Trichome birefringence-like protein | elongation zone | 12 h |
| HORVU.MOREX.r3.5HG0470640.1 | 0.68030928 | 4.647151278 | 3.57342482 | 0.00076 | 0.021761735 | -0.99515527 | WD40 repeat-containing protein | elongation zone | 12 h |
| HORVU.MOREX.r3.3HG0330120.1 | -2.266911887 | -3.009670865 | -3.41122508 | 0.00124 | 0.026060689 | -0.99757556 | O-methyltransferase | elongation zone | 12 h |
| HORVU.MOREX.r3.4HG0390380.1 | 0.917972135 | 6.629358353 | 3.62123954 | 0.00066 | 0.020810194 | -1.0007143 | Importin-4 | elongation zone | 12 h |
| HORVU.MOREX.r3.7HG0643190.1 | 1.123931854 | 4.36385392 | 3.499893979 | 0.00095 | 0.023509855 | -1.00436266 | Ribosomal protein L29 family protein | elongation zone | 12 h |
| HORVU.MOREX.r3.6HG0624450.1 | 2.008515761 | 3.649637416 | 3.407890101 | 0.00126 | 0.026083278 | -1.0055563 | Zinc finger protein VAR3, chloroplastic | elongation zone | 12 h |
| HORVU.MOREX.r3.4HG0417850.1 | 0.644846445 | 4.694211226 | 3.564307902 | 0.00078 | 0.021955355 | -1.00606073 | Cyclin delta-3, putative | elongation zone | 12 h |

**Table S5** Continued.

| ID | logFC | AveExpr | t | P.Value | adj.P.Val | B | gene description | root zone | time point |
|---|---|---|---|---|---|---|---|---|---|
| HORVU.MOREX.r3.7HG0700320.1 | 2.966417693 | 0.851795789 | 3.433958153 | 0.00116 | 0.025647345 | -1.00742906 | Pentatricopeptide repeat-containing protein | elongation zone | 12 h |
| HORVU.MOREX.r3.2HG0196800.1 | -1.670335251 | -0.209944285 | -3.408917045 | 0.00125 | 0.026083278 | -1.00789161 | Multidrug resistance protein ABC transporter family | elongation zone | 12 h |
| HORVU.MOREX.r3.6HG0578070.1 | 1.072244483 | 7.144692186 | 3.638216799 | 0.00062 | 0.020537791 | -1.01040076 | Nuclear pore complex protein NUP205 | elongation zone | 12 h |
| HORVU.MOREX.r3.5HG0504520.1 | -1.109472727 | 0.68442194 | -3.522433208 | 0.00089 | 0.023206422 | -1.01083909 | Phytoene synthase | elongation zone | 12 h |
| HORVU.MOREX.r3.4HG0343850.1 | 0.44044438 | 6.579849823 | 3.662097786 | 0.00058 | 0.020275771 | -1.01416639 | La | elongation zone | 12 h |
| HORVU.MOREX.r3.2HG0125940.1 | 1.40781657 | 2.626858817 | 3.402881118 | 0.00128 | 0.026178272 | -1.01446012 | Methyltransferase-like protein | elongation zone | 12 h |
| HORVU.MOREX.r3.5HG0507770.1 | 2.050249052 | 2.818084292 | 3.393559257 | 0.00131 | 0.026348744 | -1.01720196 | Cytosolic 5-nucleotidase | elongation zone | 12 h |
| HORVU.MOREX.r3.5HG0519930.1 | 1.386614023 | 8.04952968 | 3.656951234 | 0.00059 | 0.020428272 | -1.0218576 | 30S ribosomal protein S19 | elongation zone | 12 h |
| HORVU.MOREX.r3.6HG0617940.1 | 1.634656725 | 4.532901735 | 3.447627087 | 0.00112 | 0.025172156 | -1.02476215 | Replication factor C subunit | elongation zone | 12 h |
| HORVU.MOREX.r3.3HG0277330.1 | 0.625401879 | 4.294396239 | 3.547829376 | 0.00082 | 0.022414047 | -1.02654741 | CASP-like protein | elongation zone | 12 h |
| HORVU.MOREX.r3.6HG0553310.1 | 0.905923064 | 4.173624101 | 3.476918578 | 0.00102 | 0.024161792 | -1.02685913 | Pentatricopeptide repeat-containing protein | elongation zone | 12 h |
| HORVU.MOREX.r3.5HG0493430.1 | 1.975598498 | 1.099113261 | 3.394806784 | 0.00131 | 0.026348744 | -1.02917053 | tRNA-splicing endonuclease subunit SEN54 | elongation zone | 12 h |
| HORVU.MOREX.r3.4HG0370520.1 | 0.678991664 | 3.910657525 | 3.514589464 | 0.00091 | 0.023235104 | -1.03259812 | inactive purple acid phosphatase-like protein | elongation zone | 12 h |
| HORVU.MOREX.r3.4HG0344470.1 | 1.51268705 | 6.296849234 | 3.538707266 | 0.00085 | 0.022755386 | -1.0338962 | #N/A | elongation zone | 12 h |
| HORVU.MOREX.r3.6HG0543480.1 | 1.200963145 | 3.781413567 | 3.448021002 | 0.00111 | 0.025172156 | -1.03486582 | Disease resistance protein (NBS-LRR class) family | elongation zone | 12 h |
| HORVU.MOREX.r3.5HG0486070.1 | 0.665887224 | 6.53847179 | 3.639862118 | 0.00062 | 0.020516782 | -1.03734953 | Aminotransferase | elongation zone | 12 h |
| HORVU.MOREX.r3.3HG0302510.1 | 1.599628644 | 2.670886155 | 3.386757167 | 0.00134 | 0.026498466 | -1.03767675 | ATP-dependent zinc metalloprotease FtsH | elongation zone | 12 h |
| HORVU.MOREX.r3.1HG0081090.1 | 1.897166645 | 1.704255649 | 3.384991504 | 0.00135 | 0.026498466 | -1.03811704 | 30S ribosomal protein S1 | elongation zone | 12 h |
| HORVU.MOREX.r3.2HG0157610.1 | 1.29444426 | 5.355379445 | 3.514436111 | 0.00091 | 0.023235104 | -1.04019947 | Aspartate carbamoyltransferase | elongation zone | 12 h |
| HORVU.MOREX.r3.6HG0607150.1 | -0.361471359 | 5.329149771 | -3.671839727 | 0.00056 | 0.019908381 | -1.04104397 | Syntaxin family protein | elongation zone | 12 h |
| HORVU.MOREX.r3.1HG0061390.1 | -0.695262848 | 1.721091073 | -3.706815462 | 0.0005 | 0.019342524 | -1.04408875 | Trichome birefringence-like protein | elongation zone | 12 h |
| HORVU.MOREX.r3.6HG0573930.1 | -0.431494247 | 3.609831823 | -3.551744913 | 0.00081 | 0.022303809 | -1.0449799 | Mediator of RNA polymerase II transcription subunit 18 | elongation zone | 12 h |
| HORVU.MOREX.r3.2HG0124470.1 | 2.708125132 | 0.969557895 | 3.404593523 | 0.00127 | 0.026178272 | -1.04875238 | Transposon protein, putative, Mutator sub-class | elongation zone | 12 h |
| HORVU.MOREX.r3.4HG0399210.1 | 0.841593427 | 6.281577577 | 3.609143046 | 0.00068 | 0.020937187 | -1.05152409 | Serine--tRNA ligase | elongation zone | 12 h |
| HORVU.MOREX.r3.3HG0320580.1 | 1.214034251 | 2.137390256 | 3.391730891 | 0.00132 | 0.026441594 | -1.05260963 | Pentatricopeptide repeat-containing protein | elongation zone | 12 h |
| HORVU.MOREX.r3.3HG0278440.1 | 0.807832441 | 8.62320918 | 3.696898407 | 0.00052 | 0.019449812 | -1.05422291 | 60S ribosomal protein L18a | elongation zone | 12 h |
| HORVU.MOREX.r3.7HG0636580.1 | 0.818019731 | 9.366307276 | 3.732414477 | 0.00046 | 0.018669328 | -1.05576776 | 50S ribosomal protein L6 | elongation zone | 12 h |
| HORVU.MOREX.r3.3HG0269960.1 | -0.45547752 | 4.098981318 | -3.625547518 | 0.00065 | 0.020713759 | -1.05894867 | Protein kinase family protein | elongation zone | 12 h |
| HORVU.MOREX.r3.5HG0469740.1 | 0.83750342 | 6.005540796 | 3.593382958 | 0.00072 | 0.021178576 | -1.05910802 | WD repeat-containing protein | elongation zone | 12 h |
| HORVU.MOREX.r3.7HG0659760.1 | 1.18125835 | 6.289693825 | 3.586013691 | 0.00073 | 0.021412967 | -1.06012315 | Origin recognition complex subunit 1 | elongation zone | 12 h |
| HORVU.MOREX.r3.7HG0680880.1 | 1.910552467 | 2.289472989 | 3.374208039 | 0.00139 | 0.026563254 | -1.06250401 | Pentatricopeptide repeat-containing protein | elongation zone | 12 h |
| HORVU.MOREX.r3.5HG0459530.1 | 2.247849206 | 2.402607807 | 3.379710803 | 0.00137 | 0.026563254 | -1.06299172 | AT hook motif DNA-binding family protein | elongation zone | 12 h |
| HORVU.MOREX.r3.4HG0341080.1 | 2.112717987 | 4.721757022 | 3.436681862 | 0.00115 | 0.025575256 | -1.06333057 | interferon-activable protein | elongation zone | 12 h |
| HORVU.MOREX.r3.5HG0494890.1 | -0.664869431 | 5.12004239 | -3.618801508 | 0.00066 | 0.020839563 | -1.06457378 | Esterase/lipase/thioesterase family protein | elongation zone | 12 h |
| HORVU.MOREX.r3.2HG0096440.1 | 1.6064863 | 6.317526302 | 3.526937323 | 0.00088 | 0.023065637 | -1.0656414 | tRNA-dihydrouridine(47) synthase [NAD(P)(+)] | elongation zone | 12 h |
| HORVU.MOREX.r3.5HG0519510.1 | 1.664279744 | 2.934455383 | 3.37763276 | 0.00138 | 0.026563254 | -1.06973596 | Pseudouridine synthase | elongation zone | 12 h |
| HORVU.MOREX.r3.5HG0447720.1 | 0.539129137 | 7.798320353 | 3.700000381 | 0.00051 | 0.019449812 | -1.07020796 | Dicarboxylate transporter 2.1, chloroplastic | elongation zone | 12 h |
| HORVU.MOREX.r3.5HG0499910.1 | 1.230508401 | 2.676451276 | 3.390220856 | 0.00133 | 0.026498466 | -1.07408954 | BTB/POZ domain containing protein | elongation zone | 12 h |
| HORVU.MOREX.r3.1HG0021330.1 | 0.650527053 | 7.729068078 | 3.669535672 | 0.00056 | 0.020017114 | -1.07476353 | Asparagine--tRNA ligase | elongation zone | 12 h |
| HORVU.MOREX.r3.6HG0627090.1 | -0.81166539 | 1.382859959 | -3.449523932 | 0.00111 | 0.025121413 | -1.07483575 | Alcohol dehydrogenase 3 | elongation zone | 12 h |
| HORVU.MOREX.r3.3HG0277760.1 | 2.36500543 | 1.654351235 | 3.38954525 | 0.00133 | 0.026498466 | -1.07556134 | Mannan endo-1,4-beta-mannosidase 1 | elongation zone | 12 h |

**Table S5** Continued.

| ID | logFC | AveExpr | t | P.Value | adj.P.Val | B | gene description | root zone | time point |
|---|---|---|---|---|---|---|---|---|---|
| HORVU.MOREX.r3.7HG0704790.1 | 1.911314694 | 5.362516783 | 3.428223621 | 0.00118 | 0.025704276 | -1.07764814 | IQ domain-containing protein | elongation zone | 12 h |
| HORVU.MOREX.r3.7HG0664110.1 | 2.218394835 | -1.073195206 | 3.462913738 | 0.00107 | 0.024644437 | -1.07849664 | Polynucleotidyl transferase, ribonuclease H-like superfamily prot | elongation zone | 12 h |
| HORVU.MOREX.r3.4HG0416730.1 | 0.85423567 | 6.869698753 | 3.62411018 | 0.00065 | 0.020713759 | -1.07973951 | Kinesin-like protein | elongation zone | 12 h |
| HORVU.MOREX.r3.2HG0173210.1 | 1.111091884 | 5.578205888 | 3.539961957 | 0.00084 | 0.02271895 | -1.0819638 | Kinesin-like protein | elongation zone | 12 h |
| HORVU.MOREX.r3.6HG0588750.2 | 1.26222121 | 2.298912834 | 3.376592968 | 0.00138 | 0.026563254 | -1.08249707 | Basic-leucine zipper (bZIP) transcription factor family protein | elongation zone | 12 h |
| HORVU.MOREX.r3.2HG0109620.1 | -1.446558504 | 0.349033917 | -3.37695475 | 0.00138 | 0.026563254 | -1.08546241 | Retrotransposon protein, putative, Ty3-gypsy subclass | elongation zone | 12 h |
| HORVU.MOREX.r3.6HG0634100.1 | 2.066987701 | 5.582248586 | 3.45722133 | 0.00108 | 0.02473329 | -1.08627858 | O-linked-mannose beta-1,4-N-acetylglucosaminyltransferase-like | elongation zone | 12 h |
| HORVU.MOREX.r3.6HG0627010.1 | 1.58409629 | 5.519545424 | 3.498313891 | 0.00096 | 0.023538121 | -1.08843018 | Proliferating cell nuclear antigen | elongation zone | 12 h |
| HORVU.MOREX.r3.2HG0217550.1 | -0.475797058 | 5.544284998 | -3.695275224 | 0.00052 | 0.019449812 | -1.09064051 | rRNA N-glycosidase | elongation zone | 12 h |
| HORVU.MOREX.r3.5HG0482220.1 | 0.689982232 | 8.482975852 | 3.689193429 | 0.00053 | 0.019449812 | -1.09139437 | Eukaryotic translation initiation factor 3 subunit C | elongation zone | 12 h |
| HORVU.MOREX.r3.2HG0116210.1 | 0.718196638 | 8.951948174 | 3.714038705 | 0.00049 | 0.019138691 | -1.09208365 | Elongation factor 1-beta | elongation zone | 12 h |
| HORVU.MOREX.r3.2HG0116870.1 | 1.018721167 | 6.95000519 | 3.619639122 | 0.00066 | 0.020839563 | -1.09220361 | Condensin complex subunit 1 | elongation zone | 12 h |
| HORVU.MOREX.r3.3HG0249410.1 | 2.578480722 | 4.478527107 | 3.384632382 | 0.00135 | 0.026498466 | -1.09540557 | Cyclin family protein | elongation zone | 12 h |
| HORVU.MOREX.r3.7HG0731860.1 | 0.712410723 | 4.962633306 | 3.505565961 | 0.00094 | 0.023303902 | -1.0960883 | Ran-binding protein 1 domain-containing | elongation zone | 12 h |
| HORVU.MOREX.r3.2HG0182150.1 | 0.781089015 | 4.062913174 | 3.501457914 | 0.00095 | 0.023426471 | -1.0975967 | Charged multivesicular body protein 7 | elongation zone | 12 h |
| HORVU.MOREX.r3.6HG0572280.1 | -0.572635585 | 3.513127043 | -3.628906255 | 0.00064 | 0.020713759 | -1.09809626 | Kinase family protein | elongation zone | 12 h |
| HORVU.MOREX.r3.2HG0122720.1 | 1.091868594 | 3.012374132 | 3.411816032 | 0.00124 | 0.026060689 | -1.09864406 | Serine hydroxymethyltransferase | elongation zone | 12 h |
| HORVU.MOREX.r3.4HG0334360.1 | -0.833176487 | 4.121321037 | -3.645756693 | 0.00061 | 0.020507565 | -1.10034344 | Homeobox protein BEL1 like | elongation zone | 12 h |
| HORVU.MOREX.r3.3HG0289860.1 | 0.611581705 | 6.345885753 | 3.598668328 | 0.0007 | 0.021178576 | -1.1011817 | choice-of-anchor C domain protein, putative (Protein of unknowi | elongation zone | 12 h |
| HORVU.MOREX.r3.5HG0524250.1 | 0.788521005 | 3.283171645 | 3.41189489 | 0.00124 | 0.026060689 | -1.10300581 | Mg-protoporphyrin IX chelatase | elongation zone | 12 h |
| HORVU.MOREX.r3.1HG0029840.1 | -0.713231069 | 1.808873342 | -3.420571613 | 0.00121 | 0.025813316 | -1.10975042 | WEB family protein, chloroplastic | elongation zone | 12 h |
| HORVU.MOREX.r3.2HG0193460.1 | 0.800175958 | 3.83689201 | 3.449133809 | 0.00111 | 0.025121413 | -1.10975584 | #N/A | elongation zone | 12 h |
| HORVU.MOREX.r3.6HG0549050.1 | 2.379044016 | 0.900308474 | 3.369921822 | 0.00141 | 0.026719424 | -1.11058562 | Pentatricopeptide repeat-containing protein, putative | elongation zone | 12 h |
| HORVU.MOREX.r3.5HG0530810.1 | 0.993019373 | 5.275506896 | 3.519787048 | 0.0009 | 0.023235104 | -1.11178591 | Ammonium transporter | elongation zone | 12 h |
| HORVU.MOREX.r3.4HG0395410.1 | 1.44187097 | 6.47047948 | 3.547071901 | 0.00082 | 0.022436302 | -1.11401768 | Nuclear pore complex protein Nup155 | elongation zone | 12 h |
| HORVU.MOREX.r3.1HG0059190.1 | 2.709941602 | -1.006758538 | 3.432341862 | 0.00117 | 0.025668159 | -1.11889279 | Cytochrome P450 | elongation zone | 12 h |
| HORVU.MOREX.r3.7HG0727510.1 | 0.460193597 | 9.85060615 | 3.753609002 | 0.00043 | 0.018288218 | -1.12174761 | ATP-dependent RNA helicase | elongation zone | 12 h |
| HORVU.MOREX.r3.4HG0408030.1 | 1.033347969 | 5.109700092 | 3.488201388 | 0.00099 | 0.023815524 | -1.12184322 | Methyltransferase-like protein 1 | elongation zone | 12 h |
| HORVU.MOREX.r3.5HG0505990.1 | 1.40584563 | 3.985167002 | 3.430342907 | 0.00118 | 0.025679564 | -1.12378004 | Molybdenum cofactor sulfurase protein-like | elongation zone | 12 h |
| HORVU.MOREX.r3.5HG0478260.1 | 1.331750045 | 5.335995353 | 3.507386557 | 0.00093 | 0.023256457 | -1.12410464 | Protein kinase | elongation zone | 12 h |
| HORVU.MOREX.r3.2HG0160800.1 | 0.621035745 | 8.570133613 | 3.692194023 | 0.00053 | 0.019449812 | -1.12555701 | Nuclear transport factor 2 family protein with RNA binding doma | elongation zone | 12 h |
| HORVU.MOREX.r3.3HG0221630.1 | 1.475227092 | 6.479981921 | 3.521625634 | 0.00089 | 0.023234115 | -1.12870372 | Ethylene-responsive transcription factor | elongation zone | 12 h |
| HORVU.MOREX.r3.3HG0277810.1 | 0.698695993 | 5.684125322 | 3.568380858 | 0.00077 | 0.021876787 | -1.13249872 | Kinase-related protein DUF1296 | elongation zone | 12 h |
| HORVU.MOREX.r3.1HG0078160.1 | 1.172223493 | 5.033832594 | 3.469209021 | 0.00104 | 0.024422909 | -1.13433519 | S-adenosyl-L-methionine-dependent methyltransferases superfa | elongation zone | 12 h |
| HORVU.MOREX.r3.1HG0081960.1 | 2.114628418 | 1.77002833 | 3.352377834 | 0.00148 | 0.027239638 | -1.13606143 | Mitochondrial carrier protein, expressed | elongation zone | 12 h |
| HORVU.MOREX.r3.2HG0166740.1 | 2.418362485 | 1.177029533 | 3.366856917 | 0.00142 | 0.026855068 | -1.13726922 | Pentatricopeptide repeat-containing protein | elongation zone | 12 h |
| HORVU.MOREX.r3.6HG0567530.1 | 1.568641845 | 4.356508323 | 3.41042699 | 0.00125 | 0.026062006 | -1.13783391 | Ribonuclease H2 subunit C | elongation zone | 12 h |
| HORVU.MOREX.r3.7HG0662160.1 | 0.502710996 | 5.721315271 | 3.577400145 | 0.00075 | 0.021691049 | -1.13886547 | Phosphopantothenoylcysteine decarboxylase | elongation zone | 12 h |
| HORVU.MOREX.r3.5HG0462370.1 | 0.546230719 | 4.87439825 | 3.524888297 | 0.00088 | 0.023172964 | -1.13907195 | F-box protein | elongation zone | 12 h |
| HORVU.MOREX.r3.4HG0335740.1 | 2.079892843 | 3.576868841 | 3.360765569 | 0.00145 | 0.027076709 | -1.14036767 | WD-repeat protein, putative | elongation zone | 12 h |
| HORVU.MOREX.r3.6HG0579370.1 | 0.565234994 | 8.57000807 | 3.6868512 | 0.00054 | 0.019538954 | -1.14413714 | Aminopeptidase | elongation zone | 12 h |

**Table S5** Continued.

| ID | logFC | AveExpr | t | P.Value | adj.P.Val | B | gene description | root zone | time point |
|---|---|---|---|---|---|---|---|---|---|
| HORVU.MOREX.r3.5HG0501970.1 | 3.18254406 | -0.116227004 | 3.410189894 | 0.00125 | 0.026062006 | -1.14592347 | DUF674 family protein | elongation zone | 12 h |
| HORVU.MOREX.r3.2HG0107850.2 | 0.554666623 | 6.853519311 | 3.632072836 | 0.00063 | 0.020646627 | -1.14929405 | Elongation factor 1-alpha | elongation zone | 12 h |
| HORVU.MOREX.r3.1HG0031890.1 | 0.885965694 | 7.824402431 | 3.63187664 | 0.00063 | 0.020646627 | -1.15078973 | 40S ribosomal protein S7 | elongation zone | 12 h |
| HORVU.MOREX.r3.1HG0083980.1 | 1.084638627 | 4.41183861 | 3.428046199 | 0.00118 | 0.025704276 | -1.15316499 | Thymidine kinase | elongation zone | 12 h |
| HORVU.MOREX.r3.6HG0617390.1 | 1.914060433 | 2.152837321 | 3.33831123 | 0.00155 | 0.027506911 | -1.15408272 | 50S ribosomal protein L29 | elongation zone | 12 h |
| HORVU.MOREX.r3.4HG0343750.1 | 2.0083277 | 3.214895301 | 3.345904674 | 0.00151 | 0.027239638 | -1.15595341 | Protein FAM136A | elongation zone | 12 h |
| HORVU.MOREX.r3.5HG0483980.1 | -1.353018539 | -1.844541926 | -3.361908844 | 0.00144 | 0.02703001 | -1.15679418 | Cinnamoyl CoA reductase | elongation zone | 12 h |
| HORVU.MOREX.r3.5HG0529130.1 | 0.618811142 | 9.571785293 | 3.715966585 | 0.00049 | 0.019131832 | -1.15928889 | Elongation factor | elongation zone | 12 h |
| HORVU.MOREX.r3.6HG0619920.1 | -2.584684009 | -1.999158359 | -3.35051079 | 0.00149 | 0.027239638 | -1.16230954 | Tetratricopeptide repeat protein 38 | elongation zone | 12 h |
| HORVU.MOREX.r3.3HG0254910.1 | 1.495185032 | 1.641129906 | 3.336462679 | 0.00156 | 0.027532387 | -1.163242 | Replication protein A 14 kDa subunit B | elongation zone | 12 h |
| HORVU.MOREX.r3.2HG0128190.1 | 0.690217678 | 6.703219641 | 3.588870382 | 0.00073 | 0.021347237 | -1.16541663 | Pentatricopeptide repeat-containing protein | elongation zone | 12 h |
| HORVU.MOREX.r3.6HG0556580.1 | 1.842655323 | 2.247275253 | 3.333519781 | 0.00157 | 0.02762467 | -1.16666953 | Ribosomal RNA small subunit methyltransferase H | elongation zone | 12 h |
| HORVU.MOREX.r3.5HG0505730.1 | 2.384222949 | 4.302003717 | 3.355512878 | 0.00147 | 0.027209299 | -1.16713608 | #N/A | elongation zone | 12 h |
| HORVU.MOREX.r3.5HG0473860.1 | 1.926104239 | 0.945240803 | 3.339200371 | 0.00154 | 0.027475011 | -1.1683866 | Pentatricopeptide repeat-containing protein | elongation zone | 12 h |
| HORVU.MOREX.r3.7HG0705960.1 | 0.951065763 | 7.124115161 | 3.577985613 | 0.00075 | 0.021691049 | -1.16854197 | Far upstream element-binding protein | elongation zone | 12 h |
| HORVU.MOREX.r3.3HG0294930.1 | 2.318261919 | 4.769733248 | 3.377735652 | 0.00138 | 0.026563254 | -1.16975761 | Serine/Threonine-kinase WNK (WNK)-like protein | elongation zone | 12 h |
| HORVU.MOREX.r3.4HG0390010.1 | 0.442855906 | 6.991406485 | 3.61818404 | 0.00066 | 0.020839563 | -1.17038033 | Splicing factor 3A subunit 2 | elongation zone | 12 h |
| HORVU.MOREX.r3.4HG0341160.1 | 1.703517194 | 5.238782121 | 3.423052395 | 0.0012 | 0.025763058 | -1.17162608 | WD40 repeat-like protein | elongation zone | 12 h |
| HORVU.MOREX.r3.1HG0045360.1 | 0.843088485 | 4.090890615 | 3.426832198 | 0.00119 | 0.025731015 | -1.1741422 | Nijmegen breakage syndrome 1 protein | elongation zone | 12 h |
| HORVU.MOREX.r3.6HG0626100.1 | 0.903852548 | 7.271720005 | 3.592845182 | 0.00072 | 0.021178576 | -1.17549345 | 30S ribosomal protein S15 | elongation zone | 12 h |
| HORVU.MOREX.r3.4HG0407610.1 | 1.265532049 | 3.868431006 | 3.408225257 | 0.00126 | 0.026083278 | -1.17607345 | Cyclin-dependent kinases regulatory subunit | elongation zone | 12 h |
| HORVU.MOREX.r3.3HG0280990.1 | 0.847072406 | 9.328992522 | 3.686068748 | 0.00054 | 0.019551933 | -1.17804698 | Guanine nucleotide-binding protein beta subunit-like protein | elongation zone | 12 h |
| HORVU.MOREX.r3.2HG0097190.1 | 2.048954594 | 0.265000574 | 3.374715825 | 0.00139 | 0.026563254 | -1.17840335 | Fructokinase-2 | elongation zone | 12 h |
| HORVU.MOREX.r3.6HG0587460.1 | -1.080911799 | -2.657116661 | -3.344182717 | 0.00152 | 0.027297085 | -1.17874159 | D-alanine--D-alanine ligase | elongation zone | 12 h |
| HORVU.MOREX.r3.5HG0510150.1 | 2.637673921 | 3.882701354 | 3.339344509 | 0.00154 | 0.027475011 | -1.18222132 | Sigma non-opioid intracellular receptor 1 | elongation zone | 12 h |
| HORVU.MOREX.r3.2HG0142740.1 | -0.330259531 | 6.623244558 | -3.679788536 | 0.00055 | 0.019711469 | -1.18229607 | Protein phosphatase 2c | elongation zone | 12 h |
| HORVU.MOREX.r3.1HG0073350.1 | 1.659865315 | 2.87898944 | 3.349200204 | 0.0015 | 0.027239638 | -1.18397802 | 70 kDa heat shock protein | elongation zone | 12 h |
| HORVU.MOREX.r3.1HG0086370.1 | -1.627311857 | -3.209110522 | -3.326156628 | 0.0016 | 0.027832573 | -1.18455705 | Glyoxal oxidase | elongation zone | 12 h |
| HORVU.MOREX.r3.5HG0534960.1 | 0.799757165 | 5.864296719 | 3.534208371 | 0.00086 | 0.022899718 | -1.18617365 | #N/A | elongation zone | 12 h |
| HORVU.MOREX.r3.3HG0218330.1 | 1.65411808 | 8.028797595 | 3.589521004 | 0.00072 | 0.02133511 | -1.18625284 | #N/A | elongation zone | 12 h |
| HORVU.MOREX.r3.7HG0659390.1 | 3.146265022 | 5.47393541 | 3.380028249 | 0.00137 | 0.026563254 | -1.18667922 | Hydroxycinnamoyl-CoA shikimate/quinate hydroxycinnamoyltrar | elongation zone | 12 h |
| HORVU.MOREX.r3.3HG0231630.1 | 1.504429394 | 3.593294948 | 3.355757536 | 0.00147 | 0.027209299 | -1.18981068 | Zinc-finger domain of monoamine-oxidase A repressor R1 protei | elongation zone | 12 h |
| HORVU.MOREX.r3.7HG0750100.1 | 2.369542258 | 0.414991589 | 3.331696127 | 0.00158 | 0.02769762 | -1.1922457 | Basic-leucine zipper (BZIP) transcription factor family protein | elongation zone | 12 h |
| HORVU.MOREX.r3.4HG0384390.1 | -0.88994331 | 1.57490258 | -3.492814661 | 0.00097 | 0.023707462 | -1.19622145 | Protein kinase family protein, putative, expressed | elongation zone | 12 h |
| HORVU.MOREX.r3.3HG0270450.1 | 2.180534089 | 1.889574407 | 3.325701724 | 0.00161 | 0.027832573 | -1.19721121 | Bifunctional inhibitor/lipid-transfer protein/seed storage 2S albu | elongation zone | 12 h |
| HORVU.MOREX.r3.7HG0643620.1 | 2.222015504 | 3.330782685 | 3.329055955 | 0.00159 | 0.027806246 | -1.19735813 | B3 domain-containing protein | elongation zone | 12 h |
| HORVU.MOREX.r3.1HG0004300.1 | 1.954975263 | 1.490538886 | 3.322919173 | 0.00162 | 0.027925695 | -1.19753941 | Disease resistance protein (TIR-NBS-LRR class) family | elongation zone | 12 h |
| HORVU.MOREX.r3.2HG0133400.1 | 1.727526417 | 2.606200127 | 3.32427213 | 0.00161 | 0.027860864 | -1.20417087 | Methyltransferase family protein, putative | elongation zone | 12 h |
| HORVU.MOREX.r3.4HG0358890.1 | 0.635070137 | 6.541550372 | 3.574210034 | 0.00076 | 0.021740481 | -1.20551821 | #N/A | elongation zone | 12 h |
| HORVU.MOREX.r3.6HG0598360.1 | 2.777783718 | 0.051977317 | 3.362566619 | 0.00144 | 0.02703001 | -1.20670779 | Nucleoporin autopeptidase | elongation zone | 12 h |
| HORVU.MOREX.r3.7HG0679050.1 | -0.576508162 | 4.002867978 | -3.506988586 | 0.00093 | 0.023256457 | -1.20798971 | Upstream activation factor subunit spp27 | elongation zone | 12 h |

**Table S5** Continued.

| ID | logFC | AveExpr | t | P.Value | adj.P.Val | B | gene description | root zone | time point |
|---|---|---|---|---|---|---|---|---|---|
| HORVU.MOREX.r3.1HG0054500.1 | 0.891881558 | 5.920468629 | 3.56018618 | 0.00079 | 0.022079483 | -1.21005105 | #N/A | elongation zone | 12 h |
| HORVU.MOREX.r3.5HG0474550.1 | 1.228456363 | 5.604885166 | 3.477517405 | 0.00102 | 0.024145821 | -1.21014307 | Histone acetyltransferase type B catalytic subunit | elongation zone | 12 h |
| HORVU.MOREX.r3.2HG0122750.1 | -2.712376774 | -2.143475904 | -3.348151152 | 0.0015 | 0.027239638 | -1.21440398 | Regulator of chromosome condensation (RCC1) family protein | elongation zone | 12 h |
| HORVU.MOREX.r3.5HG0485610.1 | -1.205983685 | 0.794719635 | -3.444646667 | 0.00113 | 0.025241276 | -1.21441974 | Cytochrome P450, putative | elongation zone | 12 h |
| HORVU.MOREX.r3.1HG0056420.1 | 0.845240947 | 8.535003487 | 3.691500598 | 0.00053 | 0.019449812 | -1.2150864 | Bifunctional inhibitor/lipid-transfer protein/seed storage 2S albu | elongation zone | 12 h |
| HORVU.MOREX.r3.2HG0105270.1 | 1.53147355 | 3.902906721 | 3.353496853 | 0.00148 | 0.027239638 | -1.21691731 | UPF0307 protein | elongation zone | 12 h |
| HORVU.MOREX.r3.7HG0700220.1 | 1.223983362 | 2.018022978 | 3.324822227 | 0.00161 | 0.027838944 | -1.21708689 | Carboxypeptidase Y homolog A | elongation zone | 12 h |
| HORVU.MOREX.r3.2HG0167620.1 | 1.427622797 | 2.679480768 | 3.327623116 | 0.0016 | 0.027806246 | -1.22282317 | Plastid-lipid associated protein PAP / fibrillin family protein | elongation zone | 12 h |
| HORVU.MOREX.r3.7HG0725470.1 | 1.179637565 | 2.027520928 | 3.32768951 | 0.0016 | 0.027806246 | -1.22303341 | Pentatricopeptide repeat-containing protein | elongation zone | 12 h |
| HORVU.MOREX.r3.1HG0078040.1 | -0.55511889 | 5.57940583 | -3.639065738 | 0.00062 | 0.020516782 | -1.22384796 | Absent in melanoma 1 protein | elongation zone | 12 h |
| HORVU.MOREX.r3.1HG0055550.1 | 2.566910767 | 2.424299381 | 3.309632205 | 0.00168 | 0.028265516 | -1.22473441 | histone acetyltransferase of the CBP family 12 | elongation zone | 12 h |
| HORVU.MOREX.r3.3HG0302080.1 | 0.848266177 | 4.735593914 | 3.455602183 | 0.00109 | 0.024826998 | -1.22502572 | Aldo-keto reductase | elongation zone | 12 h |
| HORVU.MOREX.r3.3HG0323440.1 | 2.587780903 | 0.455380393 | 3.345266198 | 0.00152 | 0.027239638 | -1.22612532 | Pentatricopeptide repeat-containing protein | elongation zone | 12 h |
| HORVU.MOREX.r3.3HG0270270.1 | 1.333470411 | 5.100425589 | 3.461185756 | 0.00107 | 0.024644437 | -1.2284944 | Kinesin-like protein | elongation zone | 12 h |
| HORVU.MOREX.r3.5HG0512510.1 | 0.711116537 | 7.202032188 | 3.632512279 | 0.00063 | 0.020646627 | -1.23015064 | Subtilisin-like protease | elongation zone | 12 h |
| HORVU.MOREX.r3.3HG0234670.1 | 1.924714108 | 5.483714808 | 3.413998681 | 0.00123 | 0.026021602 | -1.23208638 | Mini-chromosome maintenance complex-binding protein | elongation zone | 12 h |
| HORVU.MOREX.r3.6HG0625200.1 | 2.678486856 | 0.649401402 | 3.364810783 | 0.00143 | 0.026979742 | -1.23219403 | defensin-like protein | elongation zone | 12 h |
| HORVU.MOREX.r3.5HG0519170.1 | 0.694592572 | 8.3989122 | 3.639693017 | 0.00062 | 0.020516782 | -1.23401591 | 60S ribosomal protein L4 | elongation zone | 12 h |
| HORVU.MOREX.r3.5HG0450470.1 | 0.602233179 | 3.631343249 | 3.431085236 | 0.00117 | 0.025676928 | -1.23486055 | Basic-leucine zipper (bZIP) transcription factor family protein | elongation zone | 12 h |
| HORVU.MOREX.r3.3HG0296590.1 | 1.71475109 | 4.817113451 | 3.384112008 | 0.00135 | 0.026498466 | -1.2353634 | Zinc finger protein VAR3, chloroplastic | elongation zone | 12 h |
| HORVU.MOREX.r3.4HG0343180.1 | 1.735315704 | 1.398218777 | 3.304191768 | 0.00171 | 0.028348165 | -1.23802048 | Diphthamide biosynthesis protein | elongation zone | 12 h |
| HORVU.MOREX.r3.7HG0702560.1 | 0.793080412 | 10.1678559 | 3.701958447 | 0.00051 | 0.019449812 | -1.24254761 | 60S acidic ribosomal protein P0 | elongation zone | 12 h |
| HORVU.MOREX.r3.2HG0164770.1 | 2.0566667 | 2.242803081 | 3.301592542 | 0.00172 | 0.028397697 | -1.24526155 | DNA cross-link repair protein | elongation zone | 12 h |
| HORVU.MOREX.r3.3HG0306150.1 | -1.239948703 | 0.587040762 | -3.34682628 | 0.00151 | 0.027239638 | -1.24658326 | RING/FYVE/PHD zinc finger superfamily protein | elongation zone | 12 h |
| HORVU.MOREX.r3.6HG0601210.1 | 1.287582464 | -0.829190823 | 3.310097621 | 0.00168 | 0.028265516 | -1.24674841 | En/Spm transposon protein-like | elongation zone | 12 h |
| HORVU.MOREX.r3.1HG0053640.1 | 1.83911017 | 1.981161553 | 3.300334093 | 0.00173 | 0.028397697 | -1.24730053 | Pentatricopeptide repeat-containing protein, putative | elongation zone | 12 h |
| HORVU.MOREX.r3.3HG0317800.1 | 0.778179081 | 3.512446305 | 3.383571044 | 0.00135 | 0.026499651 | -1.24818296 | Protein kinase superfamily protein | elongation zone | 12 h |
| HORVU.MOREX.r3.6HG0540020.1 | -1.878886981 | -2.701095093 | -3.325536257 | 0.00161 | 0.027832573 | -1.24970787 | Ubiquitin carboxyl-terminal hydrolase 2 | elongation zone | 12 h |
| HORVU.MOREX.r3.1HG0069480.1 | 1.509475144 | 3.796517346 | 3.358145485 | 0.00146 | 0.027115369 | -1.25040432 | Cyclin | elongation zone | 12 h |
| HORVU.MOREX.r3.1HG0095030.1 | 1.250789137 | 2.631178482 | 3.315322627 | 0.00166 | 0.028097383 | -1.25146634 | Tudor/PWWP/MBT superfamily protein | elongation zone | 12 h |
| HORVU.MOREX.r3.5HG0512380.1 | 1.28692172 | 6.515264522 | 3.511783212 | 0.00092 | 0.023235104 | -1.25169521 | Aspartate-semialdehyde dehydrogenase, N-acetyl-gamma-glutar | elongation zone | 12 h |
| HORVU.MOREX.r3.5HG0504000.1 | 2.080494257 | -0.714764534 | 3.330959331 | 0.00158 | 0.027734466 | -1.2540319 | DEP domain-containing protein 5 | elongation zone | 12 h |
| HORVU.MOREX.r3.1HG0073480.1 | 1.701909389 | 3.492165186 | 3.325461683 | 0.00161 | 0.027832573 | -1.2571554 | Myb-like protein | elongation zone | 12 h |
| HORVU.MOREX.r3.5HG0502760.1 | 0.818558062 | 7.161461629 | 3.595078462 | 0.00071 | 0.021178576 | -1.26053522 | Histone H4 | elongation zone | 12 h |
| HORVU.MOREX.r3.2HG0205550.1 | 0.981826331 | 5.792379962 | 3.47982564 | 0.00101 | 0.024088685 | -1.26233207 | Thymidylate synthase | elongation zone | 12 h |
| HORVU.MOREX.r3.6HG0560180.1 | 1.233905824 | 7.215113646 | 3.544832135 | 0.00083 | 0.022530917 | -1.2629593 | Diaminopimelate decarboxylase | elongation zone | 12 h |
| HORVU.MOREX.r3.4HG0403380.1 | 2.295288904 | 0.78477187 | 3.311114027 | 0.00168 | 0.028253315 | -1.26323494 | Pentatricopeptide repeat-containing protein | elongation zone | 12 h |
| HORVU.MOREX.r3.5HG0420330.1 | 2.574031917 | 4.123826071 | 3.303108096 | 0.00172 | 0.028392091 | -1.26602652 | Receptor-like protein kinase | elongation zone | 12 h |
| HORVU.MOREX.r3.1HG0078760.1 | 2.135829127 | 4.578212279 | 3.340404335 | 0.00154 | 0.02745794 | -1.26643268 | tRNA pseudouridine synthase d, putative | elongation zone | 12 h |
| HORVU.MOREX.r3.2HG0122110.1 | 2.012891559 | 3.790294672 | 3.313340541 | 0.00167 | 0.028155399 | -1.26816631 | Exosome complex component | elongation zone | 12 h |
| HORVU.MOREX.r3.4HG0399930.1 | 0.954229737 | 4.063960499 | 3.393628686 | 0.00131 | 0.026348744 | -1.2687485 | Ribosome biogenesis protein TSR3-like protein | elongation zone | 12 h |

**Table S5** Continued.

| ID | logFC | AveExpr | t | P.Value | adj.P.Val | B | gene description | root zone | time point |
|---|---|---|---|---|---|---|---|---|---|
| HORVU.MOREX.r3.6HG0594410.1 | 0.900513879 | 4.144620416 | 3.402998274 | 0.00128 | 0.026178272 | -1.26934675 | Protein TIC 20, chloroplastic | elongation zone | 12 h |
| HORVU.MOREX.r3.2HG0140980.1 | 1.736325207 | 4.118667294 | 3.33906157 | 0.00154 | 0.027475011 | -1.2699497 | RNA-binding family protein | elongation zone | 12 h |
| HORVU.MOREX.r3.7HG0702770.1 | 2.656959315 | 0.424380843 | 3.347785083 | 0.0015 | 0.027239638 | -1.27124054 | Tetratricopeptide repeat (TPR)-like superfamily protein | elongation zone | 12 h |
| HORVU.MOREX.r3.1HG0039230.1 | 0.740005367 | 5.567693484 | 3.49189848 | 0.00098 | 0.023723522 | -1.27188511 | AT-rich interactive domain-containing protein 2 | elongation zone | 12 h |
| HORVU.MOREX.r3.3HG0310510.1 | 1.165415981 | 5.599216728 | 3.460526911 | 0.00107 | 0.024651876 | -1.27234322 | RNA-binding protein | elongation zone | 12 h |
| HORVU.MOREX.r3.3HG0255530.1 | 0.92403902 | 3.072891497 | 3.32747384 | 0.0016 | 0.027806246 | -1.27617131 | Phenazine biosynthesis PhzC/PhzF family protein | elongation zone | 12 h |
| HORVU.MOREX.r3.7HG0648100.1 | 1.499895174 | 0.956664255 | 3.29126948 | 0.00178 | 0.028617771 | -1.27633699 | CsAtPR5 | elongation zone | 12 h |
| HORVU.MOREX.r3.4HG0347760.1 | -0.787430845 | 2.347769018 | -3.394918139 | 0.00131 | 0.026348744 | -1.27668966 | MYB transcription factor | elongation zone | 12 h |
| HORVU.MOREX.r3.2HG0151710.1 | 1.379481714 | 7.682528676 | 3.550259249 | 0.00082 | 0.022336842 | -1.27794238 | Cytosine-specific methyltransferase | elongation zone | 12 h |
| HORVU.MOREX.r3.2HG0123800.1 | 0.883290569 | 4.716102679 | 3.430482835 | 0.00117 | 0.025679564 | -1.27938011 | Multifunctional methyltransferase subunit TRM112-like protein | elongation zone | 12 h |
| HORVU.MOREX.r3.3HG0310280.1 | -0.349579612 | 5.67062812 | -3.61945494 | 0.00066 | 0.020839563 | -1.28167284 | CBS domain containing protein | elongation zone | 12 h |
| HORVU.MOREX.r3.7HG0700930.1 | 1.584575673 | 4.508976778 | 3.348902645 | 0.0015 | 0.027239638 | -1.28436417 | RING/U-box superfamily protein | elongation zone | 12 h |
| HORVU.MOREX.r3.3HG0264800.1 | 1.648470552 | 4.015971181 | 3.349796386 | 0.0015 | 0.027239638 | -1.28521671 | DNA helicase | elongation zone | 12 h |
| HORVU.MOREX.r3.3HG0230190.1 | 1.016060523 | 2.957930782 | 3.320998563 | 0.00163 | 0.02796111 | -1.28562121 | #N/A | elongation zone | 12 h |
| HORVU.MOREX.r3.2HG0149820.1 | 1.155499115 | 6.513488761 | 3.510950278 | 0.00092 | 0.023235104 | -1.2858033 | Small nuclear ribonucleoprotein-associated protein | elongation zone | 12 h |
| HORVU.MOREX.r3.4HG0380430.1 | -0.384317714 | 3.638981565 | -3.461152769 | 0.00107 | 0.024644437 | -1.28627434 | Membrane steroid-binding protein | elongation zone | 12 h |
| HORVU.MOREX.r3.7HG0717720.1 | 0.684651634 | 9.161719424 | 3.657250653 | 0.00059 | 0.020428272 | -1.2895325 | T-complex protein 1 subunit epsilon | elongation zone | 12 h |
| HORVU.MOREX.r3.4HG0415180.1 | 3.14881883 | 3.10939606 | 3.283566644 | 0.00182 | 0.028793645 | -1.29052386 | Jasmonate-zim-domain protein | elongation zone | 12 h |
| HORVU.MOREX.r3.4HG0367550.1 | 0.66593969 | 6.581101357 | 3.531829126 | 0.00086 | 0.022899718 | -1.29066392 | Apoptotic chromatin condensation inducer in the nucleus | elongation zone | 12 h |
| HORVU.MOREX.r3.3HG0258440.1 | 1.912620465 | 2.205284764 | 3.283377309 | 0.00182 | 0.028793645 | -1.29107945 | CRS2-associated factor 1, chloroplastic | elongation zone | 12 h |
| HORVU.MOREX.r3.1HG0079060.1 | 1.846404669 | 2.517249753 | 3.283399914 | 0.00182 | 0.028793645 | -1.2941246 | Pentatricopeptide repeat-containing protein | elongation zone | 12 h |
| HORVU.MOREX.r3.1HG0069940.1 | 0.911205604 | 7.781675396 | 3.583853866 | 0.00074 | 0.021475647 | -1.29521547 | 60S ribosomal protein L30 | elongation zone | 12 h |
| HORVU.MOREX.r3.1HG0051960.1 | 1.175623672 | 1.90386557 | 3.285958682 | 0.00181 | 0.028744755 | -1.29538381 | DNA topoisomerase 2 | elongation zone | 12 h |
| HORVU.MOREX.r3.7HG0650090.1 | 1.287463902 | 3.089010373 | 3.309581521 | 0.00168 | 0.028265516 | -1.29683625 | Single-stranded DNA-binding protein WHY1, chloroplastic | elongation zone | 12 h |
| HORVU.MOREX.r3.3HG0314030.1 | 2.546285916 | -0.769398738 | 3.355552209 | 0.00147 | 0.027209299 | -1.29693158 | F-box domain containing protein | elongation zone | 12 h |
| HORVU.MOREX.r3.4HG0410210.1 | 1.412892596 | 6.140943336 | 3.451140337 | 0.0011 | 0.025080286 | -1.30046353 | Protein O-linked-mannose beta-1,4-N-acetylglucosaminyltransfe | elongation zone | 12 h |
| HORVU.MOREX.r3.3HG0243130.1 | 0.817868586 | 2.631713432 | 3.418546487 | 0.00122 | 0.025897833 | -1.30504598 | Peroxidase | elongation zone | 12 h |
| HORVU.MOREX.r3.7HG0692650.1 | 3.325650074 | 3.60398256 | 3.276754088 | 0.00186 | 0.028861865 | -1.30592496 | Ribonuclease E/G family protein | elongation zone | 12 h |
| HORVU.MOREX.r3.4HG0349950.1 | 0.808773249 | 5.968153859 | 3.487816709 | 0.00099 | 0.023815524 | -1.30595037 | #N/A | elongation zone | 12 h |
| HORVU.MOREX.r3.6HG0633020.1 | 1.965152676 | 1.254863943 | 3.277272349 | 0.00185 | 0.028861865 | -1.30903558 | Disease resistance protein RPM1 | elongation zone | 12 h |
| HORVU.MOREX.r3.1HG0029280.1 | 0.466737879 | 7.393407536 | 3.581823905 | 0.00074 | 0.02156782 | -1.30934303 | FACT complex subunit SSRP1 | elongation zone | 12 h |
| HORVU.MOREX.r3.7HG0647980.1 | 0.97944166 | 3.018864795 | 3.316338859 | 0.00165 | 0.028050531 | -1.30935248 | Coiled-coil domain-containing protein 97 | elongation zone | 12 h |
| HORVU.MOREX.r3.7HG0720810.1 | -0.377117687 | 4.98512073 | -3.535131178 | 0.00086 | 0.022899718 | -1.31070591 | Alpha/beta-Hydrolases superfamily protein | elongation zone | 12 h |
| HORVU.MOREX.r3.2HG0099780.1 | -1.093983314 | -0.017792574 | -3.352075201 | 0.00149 | 0.027239638 | -1.31100805 | Maternal effect embryo arrest protein | elongation zone | 12 h |
| HORVU.MOREX.r3.7HG0685920.1 | -0.509432712 | 4.571683277 | -3.568115053 | 0.00077 | 0.021876787 | -1.31143453 | Solute carrier family 35 protein | elongation zone | 12 h |
| HORVU.MOREX.r3.3HG0323810.1 | 0.575497566 | 6.634620504 | 3.549008731 | 0.00082 | 0.022369939 | -1.31160054 | SWAP (Suppressor-of-White-APricot)/surp domain-containing pr | elongation zone | 12 h |
| HORVU.MOREX.r3.7HG0675050.1 | 1.351939474 | 5.035182354 | 3.398595662 | 0.00129 | 0.026336616 | -1.31194822 | Histone deacetylase, putative | elongation zone | 12 h |
| HORVU.MOREX.r3.6HG0555240.1 | 1.182276985 | 4.912990377 | 3.398773056 | 0.00129 | 0.026336616 | -1.31261179 | Kinase-like protein | elongation zone | 12 h |
| HORVU.MOREX.r3.7HG0718150.1 | 0.744538449 | 8.476808649 | 3.615519222 | 0.00067 | 0.020839563 | -1.31346587 | 40S ribosomal protein S24 | elongation zone | 12 h |
| HORVU.MOREX.r3.3HG0232420.1 | 0.758925227 | 9.487579567 | 3.65020675 | 0.0006 | 0.020507565 | -1.3164565 | 30S ribosomal protein S7 | elongation zone | 12 h |
| HORVU.MOREX.r3.5HG0421470.1 | 0.957935006 | 6.457449928 | 3.50827182 | 0.00093 | 0.023255319 | -1.31766769 | Pyridine nucleotide-disulfide oxidoreductase domain-containing | elongation zone | 12 h |

**Table S5** Continued.

| ID | logFC | AveExpr | t | P.Value | adj.P.Val | B | gene description | root zone | time point |
|---|---|---|---|---|---|---|---|---|---|
| HORVU.MOREX.r3.1HG0041030.1 | 1.826212576 | 0.101262697 | 3.300589142 | 0.00173 | 0.028397697 | -1.31781036 | #N/A | elongation zone | 12 h |
| HORVU.MOREX.r3.6HG0599760.1 | 0.901687226 | 4.051408872 | 3.372552572 | 0.0014 | 0.026598639 | -1.3210378 | U6 snRNA phosphodiesterase | elongation zone | 12 h |
| HORVU.MOREX.r3.3HG0228020.1 | -1.268718884 | -1.433568275 | -3.299017652 | 0.00174 | 0.02840648 | -1.32150655 | Glycogen synthase | elongation zone | 12 h |
| HORVU.MOREX.r3.6HG0556190.1 | 1.958129273 | 1.897109288 | 3.270023435 | 0.00189 | 0.029006839 | -1.32313964 | 50S ribosomal protein L3 | elongation zone | 12 h |
| HORVU.MOREX.r3.2HG0168730.1 | 0.497700447 | 7.047952151 | 3.558353828 | 0.0008 | 0.022167528 | -1.32322967 | Kinase family protein | elongation zone | 12 h |
| HORVU.MOREX.r3.1HG0078370.1 | 1.926874695 | 4.28064148 | 3.310546193 | 0.00168 | 0.028265516 | -1.32365481 | breast cancer associated RING 1 | elongation zone | 12 h |
| HORVU.MOREX.r3.6HG0632980.1 | 1.810827275 | 5.615727708 | 3.414439127 | 0.00123 | 0.026021602 | -1.32385076 | Replication protein A 32 kDa subunit | elongation zone | 12 h |
| HORVU.MOREX.r3.6HG0545630.1 | 1.397239824 | 3.709729912 | 3.352400299 | 0.00148 | 0.027239638 | -1.32408339 | Histone H2A | elongation zone | 12 h |
| HORVU.MOREX.r3.3HG0255500.2 | 1.442309649 | 2.726992 | 3.282698345 | 0.00182 | 0.028808148 | -1.32524777 | La related protein-like | elongation zone | 12 h |
| HORVU.MOREX.r3.6HG0595190.1 | -1.508815449 | -2.072106303 | -3.298361759 | 0.00174 | 0.02840648 | -1.32614332 | GDSL esterase/lipase | elongation zone | 12 h |
| HORVU.MOREX.r3.7HG0656260.1 | 0.550169552 | 5.217076705 | 3.484134534 | 0.001 | 0.023938404 | -1.3289912 | Metaxin-related | elongation zone | 12 h |
| HORVU.MOREX.r3.2HG0109850.1 | 0.970283881 | 5.983148067 | 3.495518511 | 0.00096 | 0.023625697 | -1.3302296 | Protein kinase | elongation zone | 12 h |
| HORVU.MOREX.r3.4HG0364440.1 | 0.901708156 | 6.096424332 | 3.496374452 | 0.00096 | 0.02359246 | -1.33111211 | translation initiation factor | elongation zone | 12 h |
| HORVU.MOREX.r3.7HG0652110.1 | 0.49536997 | 5.273679743 | 3.508824066 | 0.00093 | 0.023255319 | -1.33122155 | Ubiquinone biosynthesis O-methyltransferase | elongation zone | 12 h |
| HORVU.MOREX.r3.2HG0181680.1 | -1.439485141 | -1.908817715 | -3.418448011 | 0.00122 | 0.025897833 | -1.33131411 | CASP-like protein | elongation zone | 12 h |
| HORVU.MOREX.r3.2HG0194580.1 | 2.852123438 | 5.284996064 | 3.320894229 | 0.00163 | 0.02796111 | -1.33221704 | High mobility group family protein | elongation zone | 12 h |
| HORVU.MOREX.r3.4HG0392970.1 | 2.003431615 | 6.133609232 | 3.405040054 | 0.00127 | 0.026178272 | -1.33282091 | Ribosomal protein l7ae | elongation zone | 12 h |
| HORVU.MOREX.r3.3HG0318700.1 | -0.49615998 | 6.088388292 | -3.614987456 | 0.00067 | 0.020839563 | -1.3343867 | Phosphotransferase | elongation zone | 12 h |
| HORVU.MOREX.r3.6HG0630920.1 | 2.885365412 | -2.009589986 | 3.330289659 | 0.00158 | 0.027765848 | -1.33624466 | NBS-LRR disease resistance protein-like | elongation zone | 12 h |
| HORVU.MOREX.r3.5HG0432030.1 | 0.684027373 | 7.313410269 | 3.557060736 | 0.0008 | 0.022171653 | -1.33723249 | Tesmin/TSO1-like CXC domain containing protein, expressed | elongation zone | 12 h |
| HORVU.MOREX.r3.6HG0591340.1 | 1.471461505 | 5.612016666 | 3.40333054 | 0.00127 | 0.026178272 | -1.33816969 | Ribosomal RNA small subunit methyltransferase NEP1 | elongation zone | 12 h |
| HORVU.MOREX.r3.6HG0629990.1 | 1.642455445 | 2.855652822 | 3.273130062 | 0.00188 | 0.028923495 | -1.33865952 | Pentatricopeptide repeat-containing protein | elongation zone | 12 h |
| HORVU.MOREX.r3.7HG0720670.1 | 0.658099568 | 2.879624117 | 3.352980574 | 0.00148 | 0.027239638 | -1.34095122 | Nucleoside diphosphate kinase | elongation zone | 12 h |
| HORVU.MOREX.r3.3HG0311510.1 | 1.259732943 | 6.871599224 | 3.498525186 | 0.00096 | 0.023538121 | -1.34211503 | Dek protein | elongation zone | 12 h |
| HORVU.MOREX.r3.3HG0309580.1 | 1.209304949 | 8.340368599 | 3.568096873 | 0.00077 | 0.021876787 | -1.34265266 | RNA binding protein-like protein | elongation zone | 12 h |
| HORVU.MOREX.r3.5HG0426480.1 | 1.319211091 | 2.819296038 | 3.297898876 | 0.00174 | 0.02840648 | -1.34318211 | 1-deoxy-D-xylulose 5-phosphate synthase 1 | elongation zone | 12 h |
| HORVU.MOREX.r3.3HG0327630.1 | 1.325185715 | 4.246104341 | 3.345186396 | 0.00152 | 0.027239638 | -1.34621219 | Cyclin family protein | elongation zone | 12 h |
| HORVU.MOREX.r3.3HG0270750.1 | -0.399886973 | 6.689944506 | -3.645656779 | 0.00061 | 0.020507565 | -1.34719258 | Receptor-like kinase | elongation zone | 12 h |
| HORVU.MOREX.r3.3HG0266920.1 | 0.926810316 | 7.650919912 | 3.538579927 | 0.00085 | 0.022755386 | -1.34763488 | Carbamoyl-phosphate synthase large chain | elongation zone | 12 h |
| HORVU.MOREX.r3.4HG0416120.1 | 1.791294944 | 5.199636546 | 3.367049947 | 0.00142 | 0.026855068 | -1.34802173 | #N/A | elongation zone | 12 h |
| HORVU.MOREX.r3.2HG0176040.1 | -0.814884789 | 2.922347917 | -3.514181162 | 0.00091 | 0.023235104 | -1.34816536 | Transducin/WD40 repeat protein | elongation zone | 12 h |
| HORVU.MOREX.r3.2HG0187470.1 | 2.030335101 | 4.280507622 | 3.298369252 | 0.00174 | 0.02840648 | -1.3484807 | Replication factor C subunit | elongation zone | 12 h |
| HORVU.MOREX.r3.1HG0058500.1 | 1.992898173 | 1.953197262 | 3.260432619 | 0.00195 | 0.029482217 | -1.34868233 | Carbon catabolite repressor protein 4 | elongation zone | 12 h |
| HORVU.MOREX.r3.6HG0566930.1 | 0.865248955 | 4.26253915 | 3.387153869 | 0.00134 | 0.026498466 | -1.349377 | Biotin holocarboxylase synthetase 1-like protein | elongation zone | 12 h |
| HORVU.MOREX.r3.2HG0114200.1 | 2.465974992 | 0.611042475 | 3.297693387 | 0.00174 | 0.02840648 | -1.34955267 | Pentatricopeptide repeat-containing protein | elongation zone | 12 h |
| HORVU.MOREX.r3.2HG0143770.1 | 1.469078755 | 2.829580346 | 3.267579241 | 0.00191 | 0.029085354 | -1.3500481 | Phosphate-responsive 1 family protein | elongation zone | 12 h |
| HORVU.MOREX.r3.2HG0136010.1 | 0.690939171 | 2.50549698 | 3.318835793 | 0.00164 | 0.028033028 | -1.35028584 | DNA repair family protein | elongation zone | 12 h |
| HORVU.MOREX.r3.4HG0386590.1 | 0.721570799 | 6.15515178 | 3.508020028 | 0.00093 | 0.023255319 | -1.35211512 | Ribosomal protein S10 | elongation zone | 12 h |
| HORVU.MOREX.r3.7HG0686470.1 | 1.369501361 | 6.33041116 | 3.441476994 | 0.00114 | 0.025400464 | -1.354557 | 60S ribosomal protein L7 | elongation zone | 12 h |
| HORVU.MOREX.r3.7HG0650590.1 | 2.10462261 | -0.768965688 | 3.272532822 | 0.00188 | 0.028923495 | -1.35469993 | RNA-directed DNA polymerase (reverse transcriptase)-related fa | elongation zone | 12 h |
| HORVU.MOREX.r3.6HG0612340.1 | -1.304867094 | -2.225003498 | -3.277046998 | 0.00185 | 0.028861865 | -1.35593485 | AWPM-19-like membrane family protein | elongation zone | 12 h |

**Table S5** Continued.

| ID | logFC | AveExpr | t | P.Value | adj.P.Val | B | gene description | root zone | time point |
|---|---|---|---|---|---|---|---|---|---|
| HORVU.MOREX.r3.1HG0080170.1 | 0.797193624 | 7.403972972 | 3.544949043 | 0.00083 | 0.022530917 | -1.35683686 | 60S ribosomal protein l28 | elongation zone | 12 h |
| HORVU.MOREX.r3.7HG0708120.1 | 1.516365587 | 6.60602094 | 3.473584155 | 0.00103 | 0.024207459 | -1.36059132 | High mobility group family | elongation zone | 12 h |
| HORVU.MOREX.r3.5HG0504790.1 | 1.156457223 | 7.009436866 | 3.51320682 | 0.00091 | 0.023235104 | -1.36183717 | Zinc finger protein hangover | elongation zone | 12 h |
| HORVU.MOREX.r3.3HG0275910.1 | 2.820172993 | -0.86003395 | 3.303210787 | 0.00172 | 0.028392091 | -1.36444646 | Transcription factor GTE4-like protein | elongation zone | 12 h |
| HORVU.MOREX.r3.3HG0291490.1 | 0.692292499 | 7.190585134 | 3.59368278 | 0.00071 | 0.021178576 | -1.37034066 | #N/A | elongation zone | 12 h |
| HORVU.MOREX.r3.2HG0111570.1 | -0.733658094 | 1.289253696 | -3.369429238 | 0.00141 | 0.026723686 | -1.37151979 | Polynucleotidyl transferase, ribonuclease H-like superfamily prot | elongation zone | 12 h |
| HORVU.MOREX.r3.4HG0392160.1 | 1.068172747 | 6.711088276 | 3.508505981 | 0.00093 | 0.023255319 | -1.37210916 | Glucan endo-1,3-beta-glucosidase, putative | elongation zone | 12 h |
| HORVU.MOREX.r3.5HG0502740.1 | 2.508017898 | 5.228316694 | 3.292333949 | 0.00177 | 0.028575945 | -1.37386288 | LysM domain GPI-anchored protein | elongation zone | 12 h |
| HORVU.MOREX.r3.2HG0177000.1 | 1.349283287 | 4.493234445 | 3.326957691 | 0.0016 | 0.027806246 | -1.37564305 | DNA-directed RNA polymerase II | elongation zone | 12 h |
| HORVU.MOREX.r3.5HG0441980.1 | 3.418686043 | 5.375222678 | 3.275294034 | 0.00186 | 0.028906691 | -1.37612299 | Glutamate-1-semialdehyde 2,1-aminomutase | elongation zone | 12 h |
| HORVU.MOREX.r3.1HG0025870.1 | -0.564845069 | 4.903163827 | -3.513316951 | 0.00091 | 0.023235104 | -1.37748925 | 6,7-dimethyl-8-ribityllumazine synthase | elongation zone | 12 h |
| HORVU.MOREX.r3.6HG0629700.1 | -0.691106023 | 3.612143683 | -3.524577786 | 0.00088 | 0.023172964 | -1.37749925 | Protein PLASTID MOVEMENT IMPAIRED 1 | elongation zone | 12 h |
| HORVU.MOREX.r3.6HG0605130.1 | -0.961971866 | -2.052059751 | -3.299676412 | 0.00173 | 0.02840648 | -1.38052313 | Invertase inhibitor | elongation zone | 12 h |
| HORVU.MOREX.r3.2HG0216050.1 | 0.922092235 | 3.582358028 | 3.345978798 | 0.00151 | 0.027239638 | -1.38054997 | 2-oxoglutarate (2OG) and Fe(II)-dependent oxygenase-like protei | elongation zone | 12 h |
| HORVU.MOREX.r3.5HG0477490.1 | 1.979070139 | 2.081852442 | 3.24455422 | 0.00204 | 0.030344376 | -1.38497826 | CST complex subunit CTC1 | elongation zone | 12 h |
| HORVU.MOREX.r3.3HG0263400.1 | 0.489316639 | 6.016519655 | 3.49472968 | 0.00097 | 0.023626021 | -1.38587833 | evolutionarily conserved C-terminal region 2 | elongation zone | 12 h |
| HORVU.MOREX.r3.5HG0531560.1 | 2.441650516 | -0.689139632 | 3.299986509 | 0.00173 | 0.028404126 | -1.38643798 | Pentatricopeptide repeat-containing protein | elongation zone | 12 h |
| HORVU.MOREX.r3.1HG0066050.1 | 1.332033621 | 5.419215562 | 3.391904154 | 0.00132 | 0.026441594 | -1.3880472 | Serine protease | elongation zone | 12 h |
| HORVU.MOREX.r3.4HG0371500.1 | 1.411647236 | 7.969743783 | 3.52346949 | 0.00089 | 0.02319596 | -1.38807042 | H/ACA ribonucleoprotein complex subunit 1-like protein 1 | elongation zone | 12 h |
| HORVU.MOREX.r3.7HG0670020.1 | 1.757151711 | 7.089520342 | 3.468633163 | 0.00105 | 0.024437696 | -1.3884825 | Ribonucleoside-diphosphate reductase small chain | elongation zone | 12 h |
| HORVU.MOREX.r3.2HG0105120.1 | 0.478786289 | 6.067458615 | 3.51651975 | 0.00091 | 0.023235104 | -1.3893308 | Prostaglandin E synthase 2 | elongation zone | 12 h |
| HORVU.MOREX.r3.4HG0353450.1 | 2.059580251 | 2.283432519 | 3.242811082 | 0.00205 | 0.030414208 | -1.39030356 | Cell division control protein 45-like protein | elongation zone | 12 h |
| HORVU.MOREX.r3.6HG0631210.1 | 0.419192357 | 7.433011137 | 3.572452248 | 0.00076 | 0.021766161 | -1.39051373 | RNA-binding protein Nova | elongation zone | 12 h |
| HORVU.MOREX.r3.6HG0593330.2 | 1.629471518 | 3.765007283 | 3.292937012 | 0.00177 | 0.028575945 | -1.39185577 | Acetolactate synthase small subunit | elongation zone | 12 h |
| HORVU.MOREX.r3.5HG0486180.1 | 1.703277632 | 5.053805616 | 3.325184058 | 0.00161 | 0.027832573 | -1.39381087 | F-box protein-like | elongation zone | 12 h |
| HORVU.MOREX.r3.5HG0500740.1 | 1.302758204 | 5.554889337 | 3.402656453 | 0.00128 | 0.026178272 | -1.39600238 | WD-repeat protein, putative | elongation zone | 12 h |
| HORVU.MOREX.r3.5HG0462850.1 | 2.626048181 | 2.419844835 | 3.240400439 | 0.00206 | 0.03054087 | -1.40223032 | Peroxisomal membrane MPV17/PMP22-like protein | elongation zone | 12 h |
| HORVU.MOREX.r3.3HG0251470.1 | 2.90132338 | 2.189387907 | 3.243085527 | 0.00205 | 0.030411811 | -1.40261172 | Kinesin-like protein | elongation zone | 12 h |
| HORVU.MOREX.r3.2HG0210950.1 | 1.870397621 | 0.661852208 | 3.236836159 | 0.00209 | 0.030699736 | -1.40492934 | #N/A | elongation zone | 12 h |
| HORVU.MOREX.r3.7HG0725580.1 | -1.288733552 | -0.151355091 | -3.236401072 | 0.00209 | 0.030700472 | -1.40937935 | Protein TRANSPARENT TESTA 1 | elongation zone | 12 h |
| HORVU.MOREX.r3.1HG0057870.1 | 0.709880123 | 6.695838064 | 3.494731337 | 0.00097 | 0.023626021 | -1.40958534 | Serine hydroxymethyltransferase | elongation zone | 12 h |
| HORVU.MOREX.r3.1HG0069550.1 | 1.954977375 | 5.304263931 | 3.338086743 | 0.00155 | 0.027506911 | -1.41032047 | Cyclin | elongation zone | 12 h |
| HORVU.MOREX.r3.5HG0477290.1 | 1.212023439 | 0.968509278 | 3.245330706 | 0.00203 | 0.030322567 | -1.41034035 | cDNA clone:J013058P10, full insert sequence | elongation zone | 12 h |
| HORVU.MOREX.r3.7HG0667340.1 | 3.325714743 | 1.732788988 | 3.287356566 | 0.0018 | 0.028691351 | -1.41349359 | Trichome birefringence-like protein | elongation zone | 12 h |
| HORVU.MOREX.r3.7HG0647520.1 | 0.584244134 | 6.632237001 | 3.508259546 | 0.00093 | 0.023255319 | -1.41534347 | GDP-mannose 4,6 dehydratase-like protein | elongation zone | 12 h |
| HORVU.MOREX.r3.4HG0342000.1 | -0.736962917 | 0.190754791 | -3.37323121 | 0.00139 | 0.026569474 | -1.41751269 | 4-coumarate:CoA ligase-like protein | elongation zone | 12 h |
| HORVU.MOREX.r3.7HG0689590.1 | 1.806974217 | 6.156729257 | 3.385419958 | 0.00134 | 0.026498466 | -1.41866563 | Dihydroxy-acid dehydratase | elongation zone | 12 h |
| HORVU.MOREX.r3.5HG0524730.1 | 0.866683327 | 8.875281474 | 3.586838383 | 0.00073 | 0.021412967 | -1.42310351 | 40S ribosomal protein S26 | elongation zone | 12 h |
| HORVU.MOREX.r3.4HG0374570.1 | 1.351811777 | 3.614922227 | 3.276594686 | 0.00186 | 0.028861865 | -1.42452837 | glycine-rich RNA-binding-like protein | elongation zone | 12 h |
| HORVU.MOREX.r3.3HG0218930.1 | 3.264383974 | 2.999856259 | 3.229174387 | 0.00213 | 0.030911827 | -1.42584332 | telomeric DNA binding protein 1 | elongation zone | 12 h |
| HORVU.MOREX.r3.5HG0507950.2 | -2.477107944 | -2.471541906 | -3.280765679 | 0.00183 | 0.028819302 | -1.4262269 | Protein IQ-DOMAIN 1 | elongation zone | 12 h |

**Table S5** Continued.

| ID | logFC | AveExpr | t | P.Value | adj.P.Val | B | gene description | root zone | time point |
|---|---|---|---|---|---|---|---|---|---|
| HORVU.MOREX.r3.7HG0653210.1 | 0.81790727 | 3.132795054 | 3.279183686 | 0.00184 | 0.028839684 | -1.42918644 | F-box family protein | elongation zone | 12 h |
| HORVU.MOREX.r3.4HG0399920.1 | 1.788665191 | 6.69753872 | 3.415810959 | 0.00123 | 0.026021602 | -1.43027286 | RNA cytidine acetyltransferase | elongation zone | 12 h |
| HORVU.MOREX.r3.5HG0440860.1 | -0.36216103 | 4.818593256 | -3.49747852 | 0.00096 | 0.02354153 | -1.43181803 | Foldase protein prsA 1 | elongation zone | 12 h |
| HORVU.MOREX.r3.2HG0181310.1 | 1.784088474 | 4.106143284 | 3.281311466 | 0.00183 | 0.028819302 | -1.4330203 | Aspartyl/glutamyl-tRNA(Asn/Gln) amidotransferase subunit B, pu | elongation zone | 12 h |
| HORVU.MOREX.r3.1HG0080230.1 | 1.1587159 | 6.650953832 | 3.486491605 | 0.00099 | 0.023883457 | -1.43605039 | #N/A | elongation zone | 12 h |
| HORVU.MOREX.r3.6HG0592230.1 | 2.23646925 | 0.020065053 | 3.248261139 | 0.00202 | 0.030129448 | -1.43835115 | Mitochondrial transcription termination factor family protein | elongation zone | 12 h |
| HORVU.MOREX.r3.2HG0188300.1 | 0.474727034 | 4.693991942 | 3.428748469 | 0.00118 | 0.025704276 | -1.44011474 | CsAtPR5 | elongation zone | 12 h |
| HORVU.MOREX.r3.1HG0001510.1 | 0.930978676 | 3.140126593 | 3.308722834 | 0.00169 | 0.028279404 | -1.44330839 | NBS-LRR-like resistance protein | elongation zone | 12 h |
| HORVU.MOREX.r3.5HG0430710.1 | 1.74109837 | 1.861597982 | 3.220349988 | 0.00219 | 0.031406458 | -1.44410335 | Pentatricopeptide repeat-containing protein | elongation zone | 12 h |
| HORVU.MOREX.r3.1HG0040570.1 | 2.760688027 | 1.96316218 | 3.229528786 | 0.00213 | 0.030911827 | -1.44678023 | Pentatricopeptide repeat-containing protein | elongation zone | 12 h |
| HORVU.MOREX.r3.3HG0229650.1 | 0.587474431 | 8.40993175 | 3.580201082 | 0.00074 | 0.021614457 | -1.44820399 | Threonine--tRNA ligase | elongation zone | 12 h |
| HORVU.MOREX.r3.3HG0314250.1 | 1.345933616 | 4.306407266 | 3.297800542 | 0.00174 | 0.02840648 | -1.44956938 | Spermatogenesis-associated protein 20 | elongation zone | 12 h |
| HORVU.MOREX.r3.2HG0174250.1 | 1.09335977 | 4.093580531 | 3.306551478 | 0.0017 | 0.028288147 | -1.45143152 | Poly(A)-specific ribonuclease PARN | elongation zone | 12 h |
| HORVU.MOREX.r3.1HG0056950.1 | 1.846283004 | 2.268962637 | 3.217474825 | 0.00221 | 0.031443869 | -1.45163011 | Maturase | elongation zone | 12 h |
| HORVU.MOREX.r3.5HG0463350.1 | 1.6574046 | 5.140919247 | 3.360623372 | 0.00145 | 0.027076709 | -1.45256085 | Cell division control 45-like protein | elongation zone | 12 h |
| HORVU.MOREX.r3.6HG0549010.1 | 0.647661085 | 4.591957387 | 3.391146083 | 0.00132 | 0.02646207 | -1.45642336 | Protein DETOXIFICATION | elongation zone | 12 h |
| HORVU.MOREX.r3.4HG0410520.1 | 1.35289199 | 2.028956772 | 3.223741233 | 0.00217 | 0.031206894 | -1.45893953 | Cytochrome c-type biogenesis CcmH | elongation zone | 12 h |
| HORVU.MOREX.r3.7HG0701760.1 | 2.153092672 | 3.061025791 | 3.213518869 | 0.00223 | 0.031515925 | -1.460761 | NAD(P)H dehydrogenase (Quinone) | elongation zone | 12 h |
| HORVU.MOREX.r3.6HG0578020.1 | 1.693959855 | 6.079593404 | 3.374394766 | 0.00139 | 0.026563254 | -1.46182316 | Alba DNA/RNA-binding protein | elongation zone | 12 h |
| HORVU.MOREX.r3.1HG0021110.1 | 2.122004324 | -1.06471218 | 3.288678224 | 0.00179 | 0.028679559 | -1.46318219 | Disease resistance protein (NBS-LRR class) family | elongation zone | 12 h |
| HORVU.MOREX.r3.3HG0235960.1 | 0.781497577 | 5.136869032 | 3.384272 | 0.00135 | 0.026498466 | -1.4632093 | ATP-dependent RNA helicase p62 | elongation zone | 12 h |
| HORVU.MOREX.r3.2HG0105680.1 | 2.009935424 | 1.363898856 | 3.21261123 | 0.00224 | 0.031520621 | -1.46325292 | Pyridoxamine 5-phosphate oxidase family protein | elongation zone | 12 h |
| HORVU.MOREX.r3.7HG0669710.1 | 1.6226739 | 5.508591997 | 3.348426047 | 0.0015 | 0.027239638 | -1.46556975 | Condensin complex subunit 3 | elongation zone | 12 h |
| HORVU.MOREX.r3.1HG0062650.1 | 0.823249391 | 3.036967678 | 3.279759883 | 0.00184 | 0.028819302 | -1.46745343 | Pentatricopeptide repeat-containing protein | elongation zone | 12 h |
| HORVU.MOREX.r3.5HG0478780.1 | 3.107922423 | 1.412999456 | 3.241183104 | 0.00206 | 0.030515114 | -1.47070845 | Myb family transcription factor-like | elongation zone | 12 h |
| HORVU.MOREX.r3.2HG0210320.1 | 1.485105044 | 2.321433783 | 3.218602089 | 0.0022 | 0.031443869 | -1.4708572 | Pentatricopeptide repeat-containing protein | elongation zone | 12 h |
| HORVU.MOREX.r3.2HG0169540.1 | 1.796493985 | 1.072754222 | 3.217121069 | 0.00221 | 0.031443869 | -1.47330539 | Pentatricopeptide repeat-containing protein | elongation zone | 12 h |
| HORVU.MOREX.r3.2HG0196720.1 | 2.285326565 | -0.225277908 | 3.226782745 | 0.00215 | 0.031018589 | -1.47478417 | XH/XS domain-containing protein | elongation zone | 12 h |
| HORVU.MOREX.r3.5HG0489440.1 | 2.416177932 | 2.658027414 | 3.20728373 | 0.00227 | 0.031692167 | -1.47582247 | Subtilisin-like protease | elongation zone | 12 h |
| HORVU.MOREX.r3.1HG0014350.1 | 1.71320114 | 4.738056098 | 3.313273922 | 0.00167 | 0.028155399 | -1.47701096 | P-loop containing nucleoside triphosphate hydrolases superfamil | elongation zone | 12 h |
| HORVU.MOREX.r3.6HG0593990.1 | 1.078794311 | 4.654306111 | 3.350011581 | 0.00149 | 0.027239638 | -1.47761841 | 30S ribosomal protein S17 | elongation zone | 12 h |
| HORVU.MOREX.r3.5HG0455760.1 | 1.430929068 | 6.696936172 | 3.425349789 | 0.00119 | 0.025739392 | -1.4778073 | RNA helicase | elongation zone | 12 h |
| HORVU.MOREX.r3.5HG0447150.1 | 3.468525469 | 2.535633305 | 3.219615963 | 0.00219 | 0.031443869 | -1.48113701 | DNA binding protein | elongation zone | 12 h |
| HORVU.MOREX.r3.4HG0337010.1 | 2.820212361 | 1.286313481 | 3.249187633 | 0.00201 | 0.030069779 | -1.4835474 | Homologous-pairing protein 2-like protein | elongation zone | 12 h |
| HORVU.MOREX.r3.1HG0081780.1 | 1.302033728 | 4.770668925 | 3.323709868 | 0.00162 | 0.027883821 | -1.48562947 | Cleavage and polyadenylation specificity factor subunit 5 | elongation zone | 12 h |
| HORVU.MOREX.r3.7HG0712100.1 | 1.41484454 | 1.596116844 | 3.205337124 | 0.00229 | 0.031785508 | -1.48663635 | Aminotransferase-related family protein | elongation zone | 12 h |
| HORVU.MOREX.r3.7HG0657930.1 | 0.727475558 | 4.769359272 | 3.379356875 | 0.00137 | 0.026563254 | -1.49104013 | Aberrant root formation protein 4 | elongation zone | 12 h |
| HORVU.MOREX.r3.1HG0070100.1 | -0.403123107 | 4.109158355 | -3.432154654 | 0.00117 | 0.025668159 | -1.49116313 | Prolyl 4-hydroxylase alpha subunit, putative | elongation zone | 12 h |
| HORVU.MOREX.r3.7HG0689320.1 | 0.998810283 | 7.120907428 | 3.488532796 | 0.00099 | 0.023815524 | -1.49208519 | Kinesin-like protein | elongation zone | 12 h |
| HORVU.MOREX.r3.3HG0256890.1 | 0.929127756 | 6.520171866 | 3.42382213 | 0.0012 | 0.025763058 | -1.49453214 | Argonaute | elongation zone | 12 h |
| HORVU.MOREX.r3.6HG0609520.1 | 0.489028146 | 8.96663263 | 3.605140168 | 0.00069 | 0.021013719 | -1.49517216 | Fatty acid desaturase | elongation zone | 12 h |

**Table S5** Continued.

| ID | logFC | AveExpr | t | P.Value | adj.P.Val | B | gene description | root zone | time point |
|---|---|---|---|---|---|---|---|---|---|
| HORVU.MOREX.r3.1HG0059000.1 | 1.081616838 | 2.194539462 | 3.217375102 | 0.00221 | 0.031443869 | -1.49672425 | Receptor-like kinase | elongation zone | 12 h |
| HORVU.MOREX.r3.1HG0040400.1 | 3.004563624 | 4.472815111 | 3.233965552 | 0.0021 | 0.030821023 | -1.49690313 | Xyloglucan galactosyltransferase KATAMARI1-like protein | elongation zone | 12 h |
| HORVU.MOREX.r3.2HG0200330.1 | 1.3747188 | 4.462805147 | 3.274453146 | 0.00187 | 0.028923495 | -1.49704159 | Cell division protein ftsZ | elongation zone | 12 h |
| HORVU.MOREX.r3.5HG0527890.1 | -0.331735724 | 6.194702471 | -3.527789888 | 0.00087 | 0.0230352 | -1.49709478 | Anthranilate synthase | elongation zone | 12 h |
| HORVU.MOREX.r3.2HG0204960.1 | 0.396247647 | 4.354857362 | 3.404489483 | 0.00127 | 0.026178272 | -1.50004266 | RING/U-box superfamily protein | elongation zone | 12 h |
| HORVU.MOREX.r3.5HG0443620.1 | 1.456175913 | 2.064194108 | 3.203372188 | 0.0023 | 0.031887924 | -1.50047568 | Pentatricopeptide repeat-containing protein, putative | elongation zone | 12 h |
| HORVU.MOREX.r3.1HG0050700.1 | -1.105818947 | 2.994863261 | -3.460997221 | 0.00107 | 0.024644437 | -1.501271 | ATP-dependent zinc metalloprotease FtsH | elongation zone | 12 h |
| HORVU.MOREX.r3.6HG0551230.1 | 3.200642751 | 2.012376965 | 3.21695496 | 0.00221 | 0.031443869 | -1.50301339 | Peroxidase | elongation zone | 12 h |
| HORVU.MOREX.r3.3HG0275170.1 | 1.718419022 | 4.459730287 | 3.257908646 | 0.00196 | 0.029613672 | -1.50479547 | Single-stranded DNA-binding protein | elongation zone | 12 h |
| HORVU.MOREX.r3.5HG0516840.1 | 1.748235842 | 2.497565851 | 3.199676787 | 0.00232 | 0.031903923 | -1.50503335 | DNA topoisomerase | elongation zone | 12 h |
| HORVU.MOREX.r3.2HG0181590.1 | 0.810437736 | 7.96866194 | 3.5226954 | 0.00089 | 0.023206422 | -1.50653453 | 60S ribosomal protein L14, putative | elongation zone | 12 h |
| HORVU.MOREX.r3.6HG0555690.1 | 0.832504036 | 4.095938839 | 3.312448829 | 0.00167 | 0.028188358 | -1.50705797 | F-box family protein | elongation zone | 12 h |
| HORVU.MOREX.r3.1HG0008200.1 | 2.040115576 | 1.354368368 | 3.287723384 | 0.0018 | 0.028691351 | -1.50764395 | Cytochrome P450 | elongation zone | 12 h |
| HORVU.MOREX.r3.1HG0062130.1 | 1.488851614 | 6.63509952 | 3.397884294 | 0.0013 | 0.026344787 | -1.51067691 | Importin subunit beta-1 | elongation zone | 12 h |
| HORVU.MOREX.r3.7HG0685970.1 | -0.694211882 | 4.801919798 | -3.577570512 | 0.00075 | 0.021691049 | -1.51079553 | Kinase family protein | elongation zone | 12 h |
| HORVU.MOREX.r3.2HG0102500.1 | 2.412016545 | -0.677166593 | 3.259173537 | 0.00195 | 0.029569389 | -1.51247587 | 60 kDa chaperonin | elongation zone | 12 h |
| HORVU.MOREX.r3.6HG0570960.1 | 1.140601422 | 6.423530915 | 3.439066276 | 0.00114 | 0.025432695 | -1.51247849 | Histone H3 | elongation zone | 12 h |
| HORVU.MOREX.r3.4HG0385270.1 | 0.569214878 | 9.972411854 | 3.621485849 | 0.00066 | 0.020810194 | -1.51700703 | Elongation factor 1-alpha | elongation zone | 12 h |
| HORVU.MOREX.r3.7HG0650920.1 | 1.5399166 | 7.40240725 | 3.434552357 | 0.00116 | 0.02562904 | -1.51755779 | Nucleosome assembly protein 1-1 | elongation zone | 12 h |
| HORVU.MOREX.r3.6HG0629800.1 | 0.903217131 | 6.18269492 | 3.427555677 | 0.00119 | 0.025704276 | -1.51762799 | 50S ribosomal protein L13 | elongation zone | 12 h |
| HORVU.MOREX.r3.2HG0174140.1 | 0.60914574 | 4.885352835 | 3.414258818 | 0.00123 | 0.026021602 | -1.51788141 | Agmatine deiminase | elongation zone | 12 h |
| HORVU.MOREX.r3.2HG0131630.1 | -2.018281512 | -3.028166822 | -3.19470538 | 0.00236 | 0.032009236 | -1.51940428 | vacuolar sorting-associated protein (DUF946) | elongation zone | 12 h |
| HORVU.MOREX.r3.2HG0212900.1 | 0.610083511 | 6.207724717 | 3.449278009 | 0.00111 | 0.025121413 | -1.52060411 | Methylthioribose kinase | elongation zone | 12 h |
| HORVU.MOREX.r3.7HG0637330.1 | 1.05958791 | 3.159355737 | 3.238943823 | 0.00207 | 0.03060905 | -1.52094846 | ATP-dependent RNA helicase | elongation zone | 12 h |
| HORVU.MOREX.r3.4HG0338450.1 | 2.140081472 | 0.677849295 | 3.212570923 | 0.00224 | 0.031520621 | -1.52194805 | Pentatricopeptide repeat-containing protein | elongation zone | 12 h |
| HORVU.MOREX.r3.4HG0396270.1 | -1.929387802 | -0.551483003 | -3.186601695 | 0.00241 | 0.032503127 | -1.52605448 | RNA-directed DNA polymerase (reverse transcriptase)-related fa | elongation zone | 12 h |
| HORVU.MOREX.r3.4HG0362830.1 | 1.191014308 | 6.738047017 | 3.434881048 | 0.00116 | 0.02562904 | -1.52625622 | TRAF-like superfamily protein | elongation zone | 12 h |
| HORVU.MOREX.r3.5HG0499490.1 | 0.309273954 | 6.586701245 | 3.502714709 | 0.00094 | 0.023393436 | -1.52845569 | Transcription initiation factor IIB | elongation zone | 12 h |
| HORVU.MOREX.r3.6HG0631040.1 | -0.766829263 | 2.526682694 | -3.362239601 | 0.00144 | 0.02703001 | -1.52973654 | Mesoderm induction early response protein 1, putative isoform 1 | elongation zone | 12 h |
| HORVU.MOREX.r3.4HG0342880.1 | 1.292016604 | 0.599719418 | 3.184700396 | 0.00243 | 0.032575632 | -1.53058729 | Oxidoreductase, aldo/keto reductase family protein, expressed | elongation zone | 12 h |
| HORVU.MOREX.r3.2HG0176650.1 | 0.993957609 | 4.874812167 | 3.340569543 | 0.00154 | 0.02745794 | -1.53168115 | Anaphase-promoting complex subunit 2 | elongation zone | 12 h |
| HORVU.MOREX.r3.6HG0573240.1 | 1.535164211 | 5.752755014 | 3.348069463 | 0.0015 | 0.027239638 | -1.53178478 | Chaperone protein dnaJ, putative | elongation zone | 12 h |
| HORVU.MOREX.r3.4HG0368770.2 | 1.250433891 | 2.047876559 | 3.197848831 | 0.00234 | 0.03195973 | -1.53195499 | ATP-dependent Clp protease proteolytic subunit | elongation zone | 12 h |
| HORVU.MOREX.r3.6HG0581990.1 | -1.058423062 | 0.470811621 | -3.212438893 | 0.00224 | 0.031520621 | -1.53228992 | Mitochondrial carrier protein-like | elongation zone | 12 h |
| HORVU.MOREX.r3.3HG0294470.1 | 1.546152475 | 2.140692119 | 3.187998302 | 0.0024 | 0.032414492 | -1.53326301 | Pentatricopeptide repeat-containing protein | elongation zone | 12 h |
| HORVU.MOREX.r3.7HG0674340.1 | 0.924617841 | 5.899987322 | 3.413178011 | 0.00124 | 0.026041204 | -1.53798666 | Ribosomal protein L46 | elongation zone | 12 h |
| HORVU.MOREX.r3.2HG0203060.1 | 0.814666089 | 6.067213545 | 3.420679972 | 0.00121 | 0.025813316 | -1.53806742 | COP9 signalosome complex subunit 5 | elongation zone | 12 h |
| HORVU.MOREX.r3.5HG0526750.1 | 2.129235516 | 1.870786541 | 3.180904227 | 0.00245 | 0.032706849 | -1.54067913 | Ribosomal RNA large subunit methyltransferase E | elongation zone | 12 h |
| HORVU.MOREX.r3.4HG0389790.1 | -0.491829151 | 3.234107873 | -3.385183946 | 0.00135 | 0.026498466 | -1.54071197 | Receptor-like kinase | elongation zone | 12 h |
| HORVU.MOREX.r3.1HG0079250.1 | 0.889095276 | 7.964832516 | 3.51249491 | 0.00092 | 0.023235104 | -1.54172393 | Histone H2B | elongation zone | 12 h |
| HORVU.MOREX.r3.7HG0726810.1 | 2.811028686 | -1.043010381 | 3.203435191 | 0.0023 | 0.031887924 | -1.54200714 | HAT transposon superfamily protein, putative | elongation zone | 12 h |

**Table S5** Continued.

| ID | logFC | AveExpr | t | P.Value | adj.P.Val | B | gene description | root zone | time point |
|---|---|---|---|---|---|---|---|---|---|
| HORVU.MOREX.r3.4HG0366460.1 | 0.71957863 | 6.79957525 | 3.462360239 | 0.00107 | 0.024644437 | -1.54358873 | Embryo defective 2765 | elongation zone | 12 h |
| HORVU.MOREX.r3.3HG0254850.1 | 0.350760332 | 4.808158728 | 3.412564775 | 0.00124 | 0.026060689 | -1.54368608 | sucrose synthase 1 | elongation zone | 12 h |
| HORVU.MOREX.r3.3HG0268950.1 | 0.760510843 | 5.053714073 | 3.379031103 | 0.00137 | 0.026563254 | -1.54533237 | Helicase-related family protein | elongation zone | 12 h |
| HORVU.MOREX.r3.3HG0293310.1 | -1.093013618 | -2.571332228 | -3.217296831 | 0.00221 | 0.031443869 | -1.54559388 | Protein upstream of flc | elongation zone | 12 h |
| HORVU.MOREX.r3.3HG0248130.1 | 1.314345608 | 4.234048933 | 3.258585474 | 0.00196 | 0.029576765 | -1.54710066 | 3-methyl-2-oxobutanoate hydroxymethyltransferase | elongation zone | 12 h |
| HORVU.MOREX.r3.4HG0333550.2 | -0.781069804 | 1.181426652 | -3.306139442 | 0.0017 | 0.028288147 | -1.54719699 | senescence-associated family protein, putative (DUF581) | elongation zone | 12 h |
| HORVU.MOREX.r3.2HG0141420.1 | 1.048923897 | 5.442729657 | 3.355421845 | 0.00147 | 0.027209299 | -1.54767197 | tRNA wybutosine-synthesizing protein 2/3/4 | elongation zone | 12 h |
| HORVU.MOREX.r3.7HG0722350.1 | 0.708974997 | 5.076598366 | 3.353518273 | 0.00148 | 0.027239638 | -1.54771781 | UBX domain-containing protein, putative | elongation zone | 12 h |
| HORVU.MOREX.r3.6HG0625100.1 | 0.592122328 | 2.539221075 | 3.278574652 | 0.00185 | 0.028847361 | -1.55089264 | E3 ubiquitin-protein ligase | elongation zone | 12 h |
| HORVU.MOREX.r3.2HG0194250.1 | 0.752651335 | 7.844284678 | 3.506634333 | 0.00093 | 0.023256457 | -1.55119514 | Ribosomal protein L11 | elongation zone | 12 h |
| HORVU.MOREX.r3.4HG0401710.1 | 1.388840025 | 4.660057465 | 3.282682399 | 0.00182 | 0.028808148 | -1.55328038 | Histone H2B | elongation zone | 12 h |
| HORVU.MOREX.r3.1HG0071940.1 | 0.568570628 | 6.762417038 | 3.471695115 | 0.00104 | 0.024267361 | -1.55533705 | Sorting and assembly machinery component 50 like | elongation zone | 12 h |
| HORVU.MOREX.r3.2HG0167500.1 | 1.554086969 | 3.557899401 | 3.2308992 | 0.00212 | 0.030866089 | -1.55659075 | Repetitive proline-rich cell wall protein | elongation zone | 12 h |
| HORVU.MOREX.r3.2HG0104890.1 | 2.455617258 | 3.602032019 | 3.183739084 | 0.00243 | 0.032618994 | -1.55755373 | Rho GTPase-activating protein | elongation zone | 12 h |
| HORVU.MOREX.r3.2HG0183440.1 | 1.24000503 | 2.73877678 | 3.206450111 | 0.00228 | 0.031725905 | -1.56026995 | Mitochondrial intermembrane space import and assembly protei | elongation zone | 12 h |
| HORVU.MOREX.r3.4HG0405590.1 | 0.82005499 | 9.149514262 | 3.552524553 | 0.00081 | 0.022303809 | -1.56053541 | 40S ribosomal protein SA | elongation zone | 12 h |
| HORVU.MOREX.r3.4HG0336000.1 | -0.713337329 | 2.469297952 | -3.382291565 | 0.00136 | 0.026553975 | -1.56160741 | Transcription activator of gluconeogenesis ERT1 | elongation zone | 12 h |
| HORVU.MOREX.r3.4HG0396550.1 | 0.736924067 | 5.452542851 | 3.394055434 | 0.00131 | 0.026348744 | -1.56182599 | Mitotic checkpoint protein bub3.1 | elongation zone | 12 h |
| HORVU.MOREX.r3.7HG0667510.2 | 0.67946036 | 2.422869832 | 3.267732199 | 0.00191 | 0.029085354 | -1.56266974 | ADP-ribosylation factor family protein | elongation zone | 12 h |
| HORVU.MOREX.r3.7HG0746770.1 | 0.482956335 | 10.27031467 | 3.640535763 | 0.00062 | 0.020516782 | -1.56337738 | Chaperone protein htpG family protein | elongation zone | 12 h |
| HORVU.MOREX.r3.7HG0718540.1 | -0.700790426 | 3.33578157 | -3.334336088 | 0.00157 | 0.027599669 | -1.56365483 | Heat shock transcription factor | elongation zone | 12 h |
| HORVU.MOREX.r3.6HG0556470.1 | 0.982873866 | 6.2652176 | 3.406412969 | 0.00126 | 0.026146241 | -1.56491585 | Splicing factor 3B subunit-like protein | elongation zone | 12 h |
| HORVU.MOREX.r3.4HG0391330.1 | 0.399991626 | 6.457455724 | 3.491810853 | 0.00098 | 0.023723522 | -1.5656444 | Phosphomevalonate kinase | elongation zone | 12 h |
| HORVU.MOREX.r3.3HG0291320.1 | 1.916177767 | 0.885868778 | 3.183563413 | 0.00244 | 0.032618994 | -1.56729787 | Pentatricopeptide repeat-containing protein | elongation zone | 12 h |
| HORVU.MOREX.r3.7HG0669620.1 | -0.926534689 | 0.714004188 | -3.201821922 | 0.00231 | 0.031894634 | -1.56733662 | Receptor-like protein kinase | elongation zone | 12 h |
| HORVU.MOREX.r3.1HG0044850.1 | 1.063034834 | 2.603766468 | 3.207390591 | 0.00227 | 0.031692167 | -1.56765083 | Metal-dependent phosphohydrolase, HD subdomain | elongation zone | 12 h |
| HORVU.MOREX.r3.5HG0509880.1 | 0.583624351 | 6.303653309 | 3.443639017 | 0.00113 | 0.025290479 | -1.57160938 | Kinase | elongation zone | 12 h |
| HORVU.MOREX.r3.5HG0532090.1 | 0.64678767 | 8.117226923 | 3.518211665 | 0.0009 | 0.023235104 | -1.57164729 | Prohibitin | elongation zone | 12 h |
| HORVU.MOREX.r3.5HG0443530.1 | 1.646144069 | 6.796564929 | 3.384234475 | 0.00135 | 0.026498466 | -1.5748829 | DNA helicase | elongation zone | 12 h |
| HORVU.MOREX.r3.3HG0328320.1 | 2.281814822 | 1.36086251 | 3.169854728 | 0.00253 | 0.033099534 | -1.57531773 | Heavy metal transport/detoxification superfamily protein | elongation zone | 12 h |
| HORVU.MOREX.r3.5HG0444350.1 | -0.812737967 | 2.597522561 | -3.310333897 | 0.00168 | 0.028265516 | -1.57535509 | Sn1-specific diacylglycerol lipase alpha | elongation zone | 12 h |
| HORVU.MOREX.r3.1HG0081380.1 | -0.653245533 | 2.215537802 | -3.413770537 | 0.00124 | 0.026021602 | -1.57609489 | transcription factor BIG PETAL P (BPE) | elongation zone | 12 h |
| HORVU.MOREX.r3.3HG0295540.1 | 1.733907399 | 6.067104647 | 3.346177784 | 0.00151 | 0.027239638 | -1.5771653 | Cyclin | elongation zone | 12 h |
| HORVU.MOREX.r3.2HG0196830.1 | 2.182084546 | 0.59388019 | 3.186957915 | 0.00241 | 0.032490942 | -1.5779072 | #N/A | elongation zone | 12 h |
| HORVU.MOREX.r3.3HG0308420.1 | 0.689894839 | 3.844691287 | 3.302575195 | 0.00172 | 0.028392091 | -1.58000238 | Integrator complex subunit | elongation zone | 12 h |
| HORVU.MOREX.r3.5HG0490320.1 | 0.665321716 | 7.95248412 | 3.50784982 | 0.00093 | 0.023255319 | -1.58092324 | Eukaryotic translation initiation factor 3 subunit E | elongation zone | 12 h |
| HORVU.MOREX.r3.7HG0729940.1 | 2.104260395 | 1.804584133 | 3.163650052 | 0.00258 | 0.033395092 | -1.58139886 | Leucine-rich repeat receptor-like protein kinase family protein | elongation zone | 12 h |
| HORVU.MOREX.r3.4HG0389210.1 | 2.135309706 | 3.54018037 | 3.176487468 | 0.00249 | 0.032843155 | -1.58281059 | Mitochondrial carrier protein, expressed | elongation zone | 12 h |
| HORVU.MOREX.r3.1HG0041060.1 | 0.821521649 | 7.230752246 | 3.45966094 | 0.00108 | 0.024665565 | -1.58301035 | WD40 repeat-like protein | elongation zone | 12 h |
| HORVU.MOREX.r3.5HG0458010.1 | -0.332189141 | 5.942045141 | -3.473264642 | 0.00103 | 0.024207459 | -1.58341349 | Protein kinase, putative | elongation zone | 12 h |
| HORVU.MOREX.r3.7HG0648460.1 | -0.51023731 | 2.500172303 | -3.386335039 | 0.00134 | 0.026498466 | -1.58439952 | Glycosyltransferase | elongation zone | 12 h |

**Table S5** Continued.

| ID | logFC | AveExpr | t | P.Value | adj.P.Val | B | gene description | root zone | time point |
|---|---|---|---|---|---|---|---|---|---|
| HORVU.MOREX.r3.2HG0124120.1 | 2.626675424 | 1.689838316 | 3.170938164 | 0.00253 | 0.033099534 | -1.5846469 | Pentatricopeptide repeat-containing protein | elongation zone | 12 h |
| HORVU.MOREX.r3.7HG0659440.1 | 3.610178308 | 3.492381947 | 3.166057706 | 0.00256 | 0.033247885 | -1.58565333 | Anthocyanin 5-aromatic acyltransferase-like protein | elongation zone | 12 h |
| HORVU.MOREX.r3.4HG0416620.1 | 0.537579496 | 5.629102912 | 3.409584512 | 0.00125 | 0.026082866 | -1.58780087 | Post-GPI attachment to proteins factor 3 | elongation zone | 12 h |
| HORVU.MOREX.r3.5HG0431350.1 | 1.79605298 | 0.368564414 | 3.165117214 | 0.00257 | 0.033296092 | -1.58825586 | 5'-3' exoribonuclease 4 | elongation zone | 12 h |
| HORVU.MOREX.r3.6HG0606410.1 | 0.921813121 | 8.824885632 | 3.523415893 | 0.00089 | 0.02319596 | -1.58828314 | 50S ribosomal protein L11 | elongation zone | 12 h |
| HORVU.MOREX.r3.5HG0450950.1 | 0.498013774 | 4.96403423 | 3.363693066 | 0.00143 | 0.027010031 | -1.5889725 | GDP-L-galactose phosphorylase 1 | elongation zone | 12 h |
| HORVU.MOREX.r3.1HG0061740.1 | 0.851517377 | 6.215496352 | 3.394486805 | 0.00131 | 0.026348744 | -1.589863 | Ran-binding protein 1 domain-containing | elongation zone | 12 h |
| HORVU.MOREX.r3.7HG0708050.1 | 2.474527973 | -1.430799155 | 3.205598874 | 0.00228 | 0.031782886 | -1.59165324 | Cytochrome b561 | elongation zone | 12 h |
| HORVU.MOREX.r3.3HG0220360.1 | 0.884069747 | 4.616464522 | 3.318652135 | 0.00164 | 0.028033028 | -1.59277186 | Disease resistance protein | elongation zone | 12 h |
| HORVU.MOREX.r3.7HG0659410.1 | 2.834496914 | 7.290126011 | 3.347791098 | 0.0015 | 0.027239638 | -1.59321334 | #N/A | elongation zone | 12 h |
| HORVU.MOREX.r3.5HG0479950.1 | 1.800870716 | 1.83204156 | 3.158843941 | 0.00262 | 0.033669786 | -1.59498454 | #N/A | elongation zone | 12 h |
| HORVU.MOREX.r3.2HG0211800.1 | 1.697089691 | 2.154731344 | 3.173650102 | 0.00251 | 0.03299418 | -1.5965961 | Serine/threonine-protein kinase | elongation zone | 12 h |
| HORVU.MOREX.r3.7HG0666710.1 | 2.828660165 | 0.417146587 | 3.211669609 | 0.00224 | 0.031545502 | -1.59706212 | Pentatricopeptide repeat-containing protein | elongation zone | 12 h |
| HORVU.MOREX.r3.6HG0569700.1 | 1.198830649 | 6.267855445 | 3.381150053 | 0.00136 | 0.026563254 | -1.60114835 | Protein CYPRO4 | elongation zone | 12 h |
| HORVU.MOREX.r3.2HG0113700.1 | 1.156906256 | 4.987099523 | 3.388340131 | 0.00133 | 0.026498466 | -1.6018124 | Rubber elongation factor family protein | elongation zone | 12 h |
| HORVU.MOREX.r3.7HG0661290.1 | -0.612448358 | 0.704847423 | -3.385140433 | 0.00135 | 0.026498466 | -1.60240633 | Inositol-tetrakisphosphate 1-kinase | elongation zone | 12 h |
| HORVU.MOREX.r3.5HG0508190.1 | 0.536178871 | 8.082339826 | 3.54381911 | 0.00083 | 0.022553062 | -1.60258159 | 1-deoxy-D-xylulose 5-phosphate synthase 1 | elongation zone | 12 h |
| HORVU.MOREX.r3.2HG0100670.1 | 0.590353213 | 8.076646951 | 3.513382299 | 0.00091 | 0.023235104 | -1.603626 | Eukaryotic translation initiation factor 3 subunit M | elongation zone | 12 h |
| HORVU.MOREX.r3.1HG0063870.1 | 0.632869755 | 6.880931169 | 3.46278757 | 0.00107 | 0.024644437 | -1.60377565 | Kinesin-like protein | elongation zone | 12 h |
| HORVU.MOREX.r3.6HG0546810.1 | 2.348725546 | -0.232723193 | 3.201478974 | 0.00231 | 0.031894634 | -1.60500495 | Pentatricopeptide repeat-containing protein | elongation zone | 12 h |
| HORVU.MOREX.r3.2HG0108540.1 | 0.938404779 | 6.78478822 | 3.424250521 | 0.0012 | 0.025763058 | -1.60509689 | Eukaryotic translation initiation factor 2 subunit alpha | elongation zone | 12 h |
| HORVU.MOREX.r3.5HG0441010.1 | 1.331248543 | 3.529366075 | 3.19002748 | 0.00239 | 0.032288187 | -1.60709864 | Asparagine synthetase, putative | elongation zone | 12 h |
| HORVU.MOREX.r3.2HG0099120.1 | 1.258845032 | 3.152513512 | 3.201635118 | 0.00231 | 0.031894634 | -1.60757698 | Disease resistance protein (NBS-LRR class) family | elongation zone | 12 h |
| HORVU.MOREX.r3.2HG0181550.1 | 1.06870708 | 3.831201973 | 3.28553055 | 0.00181 | 0.028744755 | -1.61160417 | Imidazole glycerol phosphate synthase subunit HisF | elongation zone | 12 h |
| HORVU.MOREX.r3.1HG0062390.3 | 0.953275202 | 1.005907134 | 3.17677448 | 0.00248 | 0.032837066 | -1.61233657 | Dehydration responsive element-binding factor protein | elongation zone | 12 h |
| HORVU.MOREX.r3.3HG0298810.1 | 2.492829581 | -2.743442253 | 3.210376636 | 0.00225 | 0.031580217 | -1.61314169 | Protein NRT1/ PTR FAMILY 5.5 | elongation zone | 12 h |
| HORVU.MOREX.r3.6HG0544400.1 | 1.066368588 | 6.141473387 | 3.359244913 | 0.00145 | 0.027115369 | -1.61503026 | RNA-binding NOB1 | elongation zone | 12 h |
| HORVU.MOREX.r3.2HG0209030.1 | 1.66268062 | 0.629763314 | 3.157392791 | 0.00263 | 0.033716726 | -1.61661081 | RING finger protein | elongation zone | 12 h |
| HORVU.MOREX.r3.1HG0073440.1 | 1.348767502 | 4.629259021 | 3.271253183 | 0.00189 | 0.028966331 | -1.61779147 | Kinesin-like protein | elongation zone | 12 h |
| HORVU.MOREX.r3.5HG0466020.1 | 0.770205985 | 7.677986698 | 3.502833419 | 0.00094 | 0.023393436 | -1.61798729 | Omega-3 fatty acid desaturase | elongation zone | 12 h |
| HORVU.MOREX.r3.6HG0632020.2 | -1.168745607 | -2.239832204 | -3.207822851 | 0.00227 | 0.031692167 | -1.61802624 | Photosystem II 22 kDa, chloroplastic | elongation zone | 12 h |
| HORVU.MOREX.r3.5HG0524930.1 | 0.601179016 | 4.635265951 | 3.345725462 | 0.00151 | 0.027239638 | -1.61866359 | Myosin heavy chain-like protein, putative | elongation zone | 12 h |
| HORVU.MOREX.r3.3HG0328740.1 | 2.27274107 | 4.710593389 | 3.21529146 | 0.00222 | 0.031454344 | -1.62145896 | WD-repeat protein, putative | elongation zone | 12 h |
| HORVU.MOREX.r3.4HG0344990.1 | 1.115807489 | 2.816049021 | 3.196329741 | 0.00235 | 0.032009236 | -1.62180695 | NBS-LRR disease resistance protein family-1 | elongation zone | 12 h |
| HORVU.MOREX.r3.2HG0172320.3 | 0.498810237 | 5.240874315 | 3.402139569 | 0.00128 | 0.0261927 | -1.62435763 | Histone H3 | elongation zone | 12 h |
| HORVU.MOREX.r3.5HG0455140.1 | 2.713511499 | -2.159476557 | 3.200880619 | 0.00232 | 0.031894634 | -1.62448195 | tRNA (guanine(37)-N1)-methyltransferase | elongation zone | 12 h |
| HORVU.MOREX.r3.5HG0517790.1 | 1.504194555 | 5.254951629 | 3.269169611 | 0.0019 | 0.029041385 | -1.62546594 | Evolutionarily conserved C-terminal region 2 | elongation zone | 12 h |
| HORVU.MOREX.r3.4HG0365450.1 | 0.626849291 | 4.01791179 | 3.293279333 | 0.00177 | 0.028575945 | -1.62636105 | Thiol:disulfide interchange protein txlA | elongation zone | 12 h |
| HORVU.MOREX.r3.6HG0631760.1 | 0.381575359 | 5.489051319 | 3.422258782 | 0.0012 | 0.025763058 | -1.62662171 | Signal peptide peptidase-like protein | elongation zone | 12 h |
| HORVU.MOREX.r3.4HG0397300.1 | 1.403854057 | 3.644537391 | 3.191882368 | 0.00238 | 0.032199922 | -1.62811129 | Tyrosine-specific transport protein, putative | elongation zone | 12 h |
| HORVU.MOREX.r3.4HG0397850.1 | 1.373476022 | 3.722866272 | 3.199720094 | 0.00232 | 0.031903923 | -1.62916563 | Methionine--tRNA ligase | elongation zone | 12 h |

**Table S5** Continued.

| ID | logFC | AveExpr | t | P.Value | adj.P.Val | B | gene description | root zone | time point |
|---|---|---|---|---|---|---|---|---|---|
| HORVU.MOREX.r3.1HG0011440.1 | -1.416850113 | -1.581106574 | -3.149303357 | 0.00269 | 0.033864708 | -1.63042014 | Glucan endo-1,3-beta-glucosidase, putative | elongation zone | 12 h |
| HORVU.MOREX.r3.3HG0303920.1 | 1.5828651 | 2.04413203 | 3.145843236 | 0.00272 | 0.033883993 | -1.63154271 | HIT zinc finger family protein | elongation zone | 12 h |
| HORVU.MOREX.r3.7HG0715150.1 | 1.702654321 | 4.93366821 | 3.230165768 | 0.00213 | 0.030888027 | -1.63199886 | WD-repeat protein, putative | elongation zone | 12 h |
| HORVU.MOREX.r3.1HG0067160.1 | -1.140144512 | 0.452737078 | -3.322422841 | 0.00162 | 0.02794328 | -1.63245935 | Wound-induced protease inhibitor | elongation zone | 12 h |
| HORVU.MOREX.r3.3HG0310210.1 | 0.548042606 | 7.031206457 | 3.466586196 | 0.00105 | 0.02453734 | -1.63275224 | Acyl-[acyl-carrier-protein] desaturase | elongation zone | 12 h |
| HORVU.MOREX.r3.2HG0188620.1 | -0.513568777 | 4.015975413 | -3.369116667 | 0.00141 | 0.026723934 | -1.6332952 | #N/A | elongation zone | 12 h |
| HORVU.MOREX.r3.2HG0156390.1 | 0.301545598 | 6.734888733 | 3.474097558 | 0.00103 | 0.024201764 | -1.6347325 | Protein kinase | elongation zone | 12 h |
| HORVU.MOREX.r3.3HG0250640.1 | 0.782772682 | 5.868281794 | 3.388248548 | 0.00133 | 0.026498466 | -1.63726367 | Leucine--tRNA ligase | elongation zone | 12 h |
| HORVU.MOREX.r3.7HG0677860.1 | 2.460081871 | 2.903040617 | 3.146319209 | 0.00271 | 0.033864708 | -1.63829257 | RNA polymerase II elongation factor | elongation zone | 12 h |
| HORVU.MOREX.r3.3HG0269890.1 | 1.935473054 | 5.144673374 | 3.224484588 | 0.00216 | 0.031161246 | -1.63927829 | Activating transcription factor 7-interacting 1 | elongation zone | 12 h |
| HORVU.MOREX.r3.7HG0659310.1 | 0.806402105 | 3.341339311 | 3.215705797 | 0.00222 | 0.031454344 | -1.6395403 | Pentatricopeptide repeat-containing protein | elongation zone | 12 h |
| HORVU.MOREX.r3.3HG0294550.1 | 1.283227749 | 5.803277229 | 3.317902035 | 0.00164 | 0.028035058 | -1.63974538 | Pentatricopeptide repeat-containing protein | elongation zone | 12 h |
| HORVU.MOREX.r3.3HG0219810.1 | 0.939116149 | 3.950557979 | 3.228613477 | 0.00214 | 0.030940575 | -1.64071029 | Agenet domain containing protein | elongation zone | 12 h |
| HORVU.MOREX.r3.7HG0659400.1 | 2.182969104 | 7.513960163 | 3.377107178 | 0.00138 | 0.026563254 | -1.64142362 | #N/A | elongation zone | 12 h |
| HORVU.MOREX.r3.2HG0096600.1 | 0.520400134 | 8.083914516 | 3.529406006 | 0.00087 | 0.022951249 | -1.64197878 | Pyruvate dehydrogenase E1 component subunit alpha | elongation zone | 12 h |
| HORVU.MOREX.r3.5HG0537150.1 | -0.793424641 | -2.246878812 | -3.226821963 | 0.00215 | 0.031018589 | -1.64221185 | Protein DETOXIFICATION | elongation zone | 12 h |
| HORVU.MOREX.r3.4HG0412630.1 | 2.655384143 | 3.079994324 | 3.138467986 | 0.00277 | 0.03425211 | -1.64291345 | Peroxidase | elongation zone | 12 h |
| HORVU.MOREX.r3.6HG0605290.1 | 0.623397675 | 4.144725964 | 3.278627161 | 0.00185 | 0.028847361 | -1.64427736 | Protein phosphatase 2C | elongation zone | 12 h |
| HORVU.MOREX.r3.1HG0046280.1 | 0.532859514 | 7.208401788 | 3.454335524 | 0.00109 | 0.024866938 | -1.64428801 | Chloroplast inner envelope protein, putative, expressed | elongation zone | 12 h |
| HORVU.MOREX.r3.5HG0421570.1 | 2.401053783 | 5.345365617 | 3.226843255 | 0.00215 | 0.031018589 | -1.64484266 | Deoxyuridine 5'-triphosphate nucleotidohydrolase | elongation zone | 12 h |
| HORVU.MOREX.r3.7HG0636750.1 | 0.875519168 | 7.072570729 | 3.463413884 | 0.00106 | 0.024644437 | -1.64491188 | carbohydrate esterase, putative (DUF303) | elongation zone | 12 h |
| HORVU.MOREX.r3.2HG0200630.1 | 0.81646507 | 9.635164688 | 3.554211376 | 0.00081 | 0.022275873 | -1.64583121 | Adenosylhomocysteinase | elongation zone | 12 h |
| HORVU.MOREX.r3.4HG0337770.1 | -0.69669679 | 5.839765801 | -3.511272801 | 0.00092 | 0.023235104 | -1.6464253 | Histidine kinase | elongation zone | 12 h |
| HORVU.MOREX.r3.7HG0659510.1 | 2.162882906 | 0.823748924 | 3.146760675 | 0.00271 | 0.033864708 | -1.65159108 | Pentatricopeptide repeat-containing protein | elongation zone | 12 h |
| HORVU.MOREX.r3.4HG0363330.1 | 0.867156344 | 2.648390841 | 3.25539201 | 0.00198 | 0.029636522 | -1.65329341 | F-box/RNI-like superfamily protein | elongation zone | 12 h |
| HORVU.MOREX.r3.4HG0410710.1 | 2.484405258 | 1.150606136 | 3.150091823 | 0.00268 | 0.033864708 | -1.65408219 | DNA polymerase delta small subunit | elongation zone | 12 h |
| HORVU.MOREX.r3.3HG0231840.1 | 1.551122172 | 1.492682319 | 3.139362944 | 0.00277 | 0.03425211 | -1.65450751 | Disease resistance protein (NBS-LRR class) family | elongation zone | 12 h |
| HORVU.MOREX.r3.2HG0108460.1 | 0.759543968 | 5.625884288 | 3.336465077 | 0.00156 | 0.027532387 | -1.65474486 | Ubiquitin activating enzyme E1 | elongation zone | 12 h |
| HORVU.MOREX.r3.7HG0662090.1 | 1.116703086 | 1.086674019 | 3.141796864 | 0.00275 | 0.034124913 | -1.65632729 | F-box protein | elongation zone | 12 h |
| HORVU.MOREX.r3.7HG0641550.1 | 0.758881155 | 5.040372267 | 3.307658324 | 0.00169 | 0.02828778 | -1.6581353 | Chaperone protein dnaJ | elongation zone | 12 h |
| HORVU.MOREX.r3.4HG0385160.1 | 1.580699868 | 4.698386898 | 3.230335505 | 0.00213 | 0.030888027 | -1.65818749 | Transmembrane protein 209 | elongation zone | 12 h |
| HORVU.MOREX.r3.5HG0533330.1 | 0.619518397 | 3.431932283 | 3.255568626 | 0.00197 | 0.029636522 | -1.65832024 | Long-chain-fatty-acid CoA ligase, putative | elongation zone | 12 h |
| HORVU.MOREX.r3.4HG0383590.1 | 0.927599408 | 7.128541216 | 3.422794432 | 0.0012 | 0.025763058 | -1.65854878 | Aminotransferase | elongation zone | 12 h |
| HORVU.MOREX.r3.5HG0507540.1 | 0.799592838 | 3.379116795 | 3.232105343 | 0.00211 | 0.030866089 | -1.65892083 | Aspartyl/glutamyl-tRNA(Asn/Gln) amidotransferase,C subunit | elongation zone | 12 h |
| HORVU.MOREX.r3.2HG0212570.1 | 0.392966711 | 10.53927453 | 3.618065703 | 0.00066 | 0.020839563 | -1.66275482 | Calreticulin | elongation zone | 12 h |
| HORVU.MOREX.r3.2HG0117150.1 | 1.6051674 | 1.019556079 | 3.130811118 | 0.00283 | 0.034668194 | -1.66294188 | Pentatricopeptide repeat-containing protein | elongation zone | 12 h |
| HORVU.MOREX.r3.2HG0110060.1 | 0.520632972 | 6.526409036 | 3.442912292 | 0.00113 | 0.025318336 | -1.66620033 | Aldehyde dehydrogenase | elongation zone | 12 h |
| HORVU.MOREX.r3.5HG0503530.1 | -0.539750374 | 0.239465893 | -3.336118252 | 0.00156 | 0.027532387 | -1.6662299 | transmembrane protein, putative (DUF594) | elongation zone | 12 h |
| HORVU.MOREX.r3.5HG0504820.1 | 2.074411625 | 4.749265507 | 3.217303673 | 0.00221 | 0.031443869 | -1.66657383 | Uveal autoantigen with coiled-coil domains and ankyrin repeats i | elongation zone | 12 h |
| HORVU.MOREX.r3.2HG0097390.1 | 0.522564444 | 4.644505302 | 3.379711989 | 0.00137 | 0.026563254 | -1.66933783 | Inosine-5'-monophosphate dehydrogenase | elongation zone | 12 h |
| HORVU.MOREX.r3.1HG0073890.1 | 1.986788952 | 3.385036735 | 3.146447218 | 0.00271 | 0.033864708 | -1.66969166 | Microtubule binding protein | elongation zone | 12 h |

**Table S5** Continued.

| ID | logFC | AveExpr | t | P.Value | adj.P.Val | B | gene description | root zone | time point |
|---|---|---|---|---|---|---|---|---|---|
| HORVU.MOREX.r3.4HG0407540.1 | 0.798812629 | 6.028730279 | 3.375269388 | 0.00139 | 0.026563254 | -1.67239178 | Ankyrin repeat domain protein, putative | elongation zone | 12 h |
| HORVU.MOREX.r3.6HG0543770.1 | 0.787434127 | 7.792354173 | 3.480776787 | 0.00101 | 0.024060672 | -1.68199249 | Histone H2A | elongation zone | 12 h |
| HORVU.MOREX.r3.2HG0171930.1 | 1.328774792 | 3.812478286 | 3.160945312 | 0.0026 | 0.033587945 | -1.68383277 | Phosphate carrier protein, mitochondrial | elongation zone | 12 h |
| HORVU.MOREX.r3.3HG0256600.1 | 1.150024475 | 3.275742589 | 3.178076067 | 0.00247 | 0.032834207 | -1.68457035 | ATP-dependent Clp protease proteolytic subunit | elongation zone | 12 h |
| HORVU.MOREX.r3.2HG0169030.1 | 1.469232821 | 5.125078024 | 3.257004702 | 0.00197 | 0.029636522 | -1.68643862 | ATP-dependent RNA helicase DeaD | elongation zone | 12 h |
| HORVU.MOREX.r3.2HG0201770.1 | 1.104991427 | 4.924549228 | 3.263237962 | 0.00193 | 0.029341049 | -1.68829885 | Ribosomal protein S4 | elongation zone | 12 h |
| HORVU.MOREX.r3.2HG0115290.1 | 2.386481221 | 3.505789531 | 3.142473224 | 0.00274 | 0.034095864 | -1.68860019 | Kinase interacting (KIP1-like) family protein | elongation zone | 12 h |
| HORVU.MOREX.r3.2HG0161340.1 | 0.905476128 | 6.240076424 | 3.358101635 | 0.00146 | 0.027115369 | -1.6905995 | Methyltransferase-like protein 13 | elongation zone | 12 h |
| HORVU.MOREX.r3.7HG0745430.1 | 5.182605326 | 4.479660325 | 3.118761011 | 0.00293 | 0.035349763 | -1.6917984 | Bifunctional inhibitor/lipid-transfer protein/seed storage 2S albu | elongation zone | 12 h |
| HORVU.MOREX.r3.2HG0207860.1 | -1.162842427 | -2.164021772 | -3.19103761 | 0.00238 | 0.032236279 | -1.69202811 | Serine/threonine-protein kinase | elongation zone | 12 h |
| HORVU.MOREX.r3.3HG0321260.1 | 1.120790738 | 7.27024282 | 3.395014349 | 0.00131 | 0.026348744 | -1.69375507 | Orotate phosphoribosyltransferase, Orotidine-5'-phosphate deca | elongation zone | 12 h |
| HORVU.MOREX.r3.6HG0542360.1 | 0.671027945 | 7.686684687 | 3.446643878 | 0.00112 | 0.025172156 | -1.69419755 | U5 small nuclear ribonucleoprotein helicase | elongation zone | 12 h |
| HORVU.MOREX.r3.6HG0624720.1 | 0.435425905 | 7.91443531 | 3.480592051 | 0.00101 | 0.024060672 | -1.6948411 | ABC transporter ATP-binding protein ARB1 | elongation zone | 12 h |
| HORVU.MOREX.r3.6HG0547570.1 | -0.48105541 | 5.07323181 | -3.440238539 | 0.00114 | 0.025432695 | -1.69504986 | LURP-one-like protein | elongation zone | 12 h |
| HORVU.MOREX.r3.3HG0283250.1 | -0.615820559 | 3.003628567 | -3.419515599 | 0.00121 | 0.02586849 | -1.69572504 | MYB transcription factor | elongation zone | 12 h |
| HORVU.MOREX.r3.7HG0648740.1 | 1.179625464 | 2.892311845 | 3.14745735 | 0.0027 | 0.033864708 | -1.69618456 | Pentatricopeptide repeat-containing family protein | elongation zone | 12 h |
| HORVU.MOREX.r3.2HG0122420.1 | 1.606556212 | 3.865411996 | 3.175556613 | 0.00249 | 0.032910434 | -1.697613 | Origin recognition complex subunit 6 | elongation zone | 12 h |
| HORVU.MOREX.r3.6HG0591300.1 | 1.544720939 | 6.018441169 | 3.283803547 | 0.00182 | 0.028793645 | -1.69794499 | CCAAT/enhancer-binding protein zeta | elongation zone | 12 h |
| HORVU.MOREX.r3.3HG0319570.1 | 1.260387179 | 3.275640979 | 3.151247282 | 0.00267 | 0.033864708 | -1.69865468 | Crossover junction endonuclease mus81 | elongation zone | 12 h |
| HORVU.MOREX.r3.4HG0401720.1 | 1.035943904 | 7.267357119 | 3.42546233 | 0.00119 | 0.025739392 | -1.69885216 | Histone H3 | elongation zone | 12 h |
| HORVU.MOREX.r3.5HG0488290.1 | 1.100009367 | 6.25888895 | 3.337239404 | 0.00155 | 0.027528501 | -1.69956432 | Ribosomal RNA small subunit methyltransferase F | elongation zone | 12 h |
| HORVU.MOREX.r3.4HG0354970.1 | 0.593744529 | 6.732308281 | 3.445026429 | 0.00112 | 0.025239957 | -1.70472186 | Omega-3 fatty acid desaturase | elongation zone | 12 h |
| HORVU.MOREX.r3.4HG0381070.1 | -0.485400896 | 3.312494191 | -3.296198222 | 0.00175 | 0.028450551 | -1.70606863 | helicase domain-containing protein / IBR domain-containing prot | elongation zone | 12 h |
| HORVU.MOREX.r3.1HG0068380.1 | 0.409960461 | 5.314485157 | 3.38944548 | 0.00133 | 0.026498466 | -1.70748111 | #N/A | elongation zone | 12 h |
| HORVU.MOREX.r3.5HG0530750.1 | 0.404234396 | 5.706473606 | 3.399064024 | 0.00129 | 0.026336616 | -1.70813704 | Post-GPI attachment to proteins factor 3 | elongation zone | 12 h |
| HORVU.MOREX.r3.2HG0160160.1 | 1.228122433 | 6.43005085 | 3.350577717 | 0.00149 | 0.027239638 | -1.70831627 | Kinesin-like protein | elongation zone | 12 h |
| HORVU.MOREX.r3.5HG0516650.1 | 0.318338905 | 6.060187193 | 3.416241656 | 0.00123 | 0.026016162 | -1.70845172 | Nuclear-interacting partner of ALK | elongation zone | 12 h |
| HORVU.MOREX.r3.5HG0507230.1 | 1.46258788 | 4.207892853 | 3.1722202 | 0.00252 | 0.033037836 | -1.71074148 | RNA-binding protein | elongation zone | 12 h |
| HORVU.MOREX.r3.3HG0252850.1 | 0.864332523 | 7.821717991 | 3.43175649 | 0.00117 | 0.025668159 | -1.71330107 | Importin subunit alpha | elongation zone | 12 h |
| HORVU.MOREX.r3.2HG0176570.1 | 1.117925721 | 5.576785683 | 3.288391179 | 0.00179 | 0.028681437 | -1.71415539 | Kelch domain-containing 4 | elongation zone | 12 h |
| HORVU.MOREX.r3.5HG0497370.1 | 0.956888463 | 9.177672535 | 3.475171192 | 0.00103 | 0.024178546 | -1.71509921 | ATP-dependent RNA helicase | elongation zone | 12 h |
| HORVU.MOREX.r3.6HG0545970.1 | 0.411901732 | 5.654969195 | 3.408640039 | 0.00125 | 0.026083278 | -1.71555455 | Short-chain dehydrogenase/reductase family protein | elongation zone | 12 h |
| HORVU.MOREX.r3.7HG0714720.1 | 2.59689246 | 4.54757859 | 3.169736415 | 0.00253 | 0.033099534 | -1.71597355 | 3'-N-debenzoyl-2'-deoxytaxol N-benzoyltransferase | elongation zone | 12 h |
| HORVU.MOREX.r3.5HG0523140.1 | 1.685796788 | 5.884927616 | 3.289811729 | 0.00179 | 0.028628646 | -1.71969631 | DNA ligase-like protein | elongation zone | 12 h |
| HORVU.MOREX.r3.3HG0311730.1 | -0.296824223 | 5.970093793 | -3.447336669 | 0.00112 | 0.025172156 | -1.72038392 | S-acyltransferase | elongation zone | 12 h |
| HORVU.MOREX.r3.5HG0509260.1 | -0.532830892 | 4.152888421 | -3.434657733 | 0.00116 | 0.02562904 | -1.72283174 | BTB/POZ domain-containing protein | elongation zone | 12 h |
| HORVU.MOREX.r3.2HG0104510.1 | 0.360638277 | 7.315942183 | 3.440708681 | 0.00114 | 0.025431673 | -1.72321344 | FACT complex subunit SPT16 | elongation zone | 12 h |
| HORVU.MOREX.r3.3HG0238730.1 | 2.14467479 | 4.16441507 | 3.139930261 | 0.00276 | 0.034221402 | -1.72373985 | DNA mismatch repair protein mutS | elongation zone | 12 h |
| HORVU.MOREX.r3.1HG0053990.1 | 0.520060747 | 5.229624499 | 3.335250368 | 0.00156 | 0.027548584 | -1.72487506 | Ankyrin repeat family protein | elongation zone | 12 h |
| HORVU.MOREX.r3.7HG0669190.2 | 1.903734334 | 0.693680419 | 3.108321973 | 0.00302 | 0.035840683 | -1.72586921 | Adenylyltransferase and sulfurtransferase MOCS3 | elongation zone | 12 h |
| HORVU.MOREX.r3.5HG0470890.1 | 1.097784667 | 6.799446968 | 3.396426623 | 0.0013 | 0.026348744 | -1.72749001 | #N/A | elongation zone | 12 h |

**Table S5** Continued.

| ID | logFC | AveExpr | t | P.Value | adj.P.Val | B | gene description | root zone | time point |
|---|---|---|---|---|---|---|---|---|---|
| HORVU.MOREX.r3.7HG0655530.1 | 0.844974947 | 5.262456683 | 3.280937823 | 0.00183 | 0.028819302 | -1.73081839 | Serine/threonine-protein phosphatase | elongation zone | 12 h |
| HORVU.MOREX.r3.3HG0243180.1 | 0.808237474 | 8.802526067 | 3.476378339 | 0.00102 | 0.024166531 | -1.73093636 | 50S ribosomal protein L5 | elongation zone | 12 h |
| HORVU.MOREX.r3.5HG0458890.1 | -0.450872992 | 3.514885037 | -3.342887311 | 0.00153 | 0.027378374 | -1.73285406 | senescence regulator (Protein of unknown function, DUF584) | elongation zone | 12 h |
| HORVU.MOREX.r3.7HG0634400.1 | 1.436626187 | 6.04833754 | 3.305096009 | 0.00171 | 0.028318301 | -1.73383656 | Kinesin-like protein | elongation zone | 12 h |
| HORVU.MOREX.r3.5HG0506300.1 | -0.359045766 | 5.425044791 | -3.439072733 | 0.00114 | 0.025432695 | -1.73566219 | Potassium transporter | elongation zone | 12 h |
| HORVU.MOREX.r3.4HG0353200.1 | -0.553168164 | 2.886286462 | -3.382247329 | 0.00136 | 0.026553975 | -1.73626245 | heat-inducible transcription repressor | elongation zone | 12 h |
| HORVU.MOREX.r3.7HG0642830.1 | 2.150902026 | 5.18214762 | 3.200753578 | 0.00232 | 0.031894634 | -1.73644378 | Scarecrow transcription factor family protein | elongation zone | 12 h |
| HORVU.MOREX.r3.5HG0507460.1 | 1.702438321 | 6.376315971 | 3.293072384 | 0.00177 | 0.028575945 | -1.73727844 | #N/A | elongation zone | 12 h |
| HORVU.MOREX.r3.3HG0322720.1 | 1.69978198 | 4.211989362 | 3.1598411 | 0.00261 | 0.033615353 | -1.73830582 | Protein RETICULATA, chloroplastic | elongation zone | 12 h |
| HORVU.MOREX.r3.3HG0315280.1 | 2.501590913 | 6.440086434 | 3.19637771 | 0.00235 | 0.032009236 | -1.74195774 | Eukaryotic aspartyl protease family protein | elongation zone | 12 h |
| HORVU.MOREX.r3.5HG0470230.1 | 0.823376227 | 9.108223054 | 3.48454413 | 0.001 | 0.023938404 | -1.74294624 | 40S ribosomal protein S19 | elongation zone | 12 h |
| HORVU.MOREX.r3.4HG0383340.1 | 1.700056526 | 0.2957173 | 3.100228114 | 0.00309 | 0.036259291 | -1.7434892 | Pentatricopeptide repeat-containing protein | elongation zone | 12 h |
| HORVU.MOREX.r3.4HG0333190.1 | 2.660670593 | 0.389975525 | 3.145698041 | 0.00272 | 0.033883993 | -1.74397172 | N utilization substance protein B-like protein | elongation zone | 12 h |
| HORVU.MOREX.r3.3HG0252460.1 | 0.730766996 | 6.678630411 | 3.383905679 | 0.00135 | 0.026498466 | -1.74503895 | 50S ribosomal protein L3, putative | elongation zone | 12 h |
| HORVU.MOREX.r3.3HG0248810.1 | 0.471633611 | 8.679714325 | 3.493746947 | 0.00097 | 0.023668509 | -1.74572243 | Eukaryotic translation initiation factor 3 subunit L | elongation zone | 12 h |
| HORVU.MOREX.r3.4HG0383870.1 | 0.686038888 | 7.088831513 | 3.425884594 | 0.00119 | 0.025739392 | -1.74589167 | Malonyl CoA-acyl carrier protein transacylase containing protein | elongation zone | 12 h |
| HORVU.MOREX.r3.7HG0739930.1 | 0.837618542 | 6.017002486 | 3.346584208 | 0.00151 | 0.027239638 | -1.74605986 | 60S ribosomal protein L30 | elongation zone | 12 h |
| HORVU.MOREX.r3.3HG0320600.1 | 1.00039752 | 4.369351337 | 3.198904771 | 0.00233 | 0.031930285 | -1.74606258 | RNA-dependent RNA polymerase | elongation zone | 12 h |
| HORVU.MOREX.r3.5HG0439450.1 | 2.027562996 | 3.124129671 | 3.11118636 | 0.003 | 0.035630935 | -1.74628745 | Chromosome alignment-maintaining phosphoprotein 1 | elongation zone | 12 h |
| HORVU.MOREX.r3.1HG0019240.1 | -0.547581682 | 2.336885411 | -3.292478453 | 0.00177 | 0.028575945 | -1.74739787 | N-acetyltransferase, putative | elongation zone | 12 h |
| HORVU.MOREX.r3.2HG0191650.1 | 2.203705514 | -0.087996826 | 3.121183815 | 0.00291 | 0.035234858 | -1.74895762 | Methionine aminopeptidase | elongation zone | 12 h |
| HORVU.MOREX.r3.4HG0379470.1 | 1.106455011 | 2.488131719 | 3.120810505 | 0.00292 | 0.035234858 | -1.75162005 | Endoglucanase | elongation zone | 12 h |
| HORVU.MOREX.r3.7HG0725660.1 | 1.063667199 | 1.229915497 | 3.113199408 | 0.00298 | 0.035529393 | -1.75197097 | FBD-associated F-box protein | elongation zone | 12 h |
| HORVU.MOREX.r3.1HG0067200.1 | 1.54096817 | 4.612988993 | 3.215302564 | 0.00222 | 0.031454344 | -1.75322618 | ATP-dependent DNA helicase Hel308 | elongation zone | 12 h |
| HORVU.MOREX.r3.1HG0088590.1 | 0.679097962 | 5.072719668 | 3.306126653 | 0.0017 | 0.028288147 | -1.7541448 | Urease accessory protein ureG | elongation zone | 12 h |
| HORVU.MOREX.r3.3HG0236710.1 | 2.732706935 | 3.114508496 | 3.090117929 | 0.00318 | 0.036772996 | -1.7574599 | IQ domain-containing protein | elongation zone | 12 h |
| HORVU.MOREX.r3.2HG0197700.1 | 1.084066192 | 3.163704659 | 3.153997796 | 0.00265 | 0.033864708 | -1.75836111 | Rho GTPase-activating protein | elongation zone | 12 h |
| HORVU.MOREX.r3.1HG0069490.1 | 1.92790817 | 2.67154507 | 3.102514514 | 0.00307 | 0.036128346 | -1.75881105 | Cyclin | elongation zone | 12 h |
| HORVU.MOREX.r3.5HG0498890.1 | 1.557417299 | 1.232074563 | 3.088148273 | 0.0032 | 0.036875974 | -1.76031478 | Pentatricopeptide repeat-containing protein | elongation zone | 12 h |
| HORVU.MOREX.r3.2HG0183630.1 | 2.273522283 | 0.168588997 | 3.114862768 | 0.00297 | 0.035453096 | -1.76090859 | Hus1-like protein | elongation zone | 12 h |
| HORVU.MOREX.r3.3HG0221030.1 | 2.501449767 | 3.659641944 | 3.091559505 | 0.00317 | 0.036725711 | -1.76227166 | Xyloglucan galactosyltransferase KATAMARI1-like protein | elongation zone | 12 h |
| HORVU.MOREX.r3.2HG0204270.1 | 2.217588978 | 3.872721928 | 3.112768079 | 0.00299 | 0.035540654 | -1.76289292 | Small RNA degrading nuclease | elongation zone | 12 h |
| HORVU.MOREX.r3.2HG0183080.1 | 0.978634145 | 4.793681433 | 3.271008543 | 0.00189 | 0.028966331 | -1.76289298 | Carboxypeptidase | elongation zone | 12 h |
| HORVU.MOREX.r3.1HG0064320.1 | 0.557267225 | 6.073377779 | 3.384981324 | 0.00135 | 0.026498466 | -1.76361504 | Argininosuccinate synthase | elongation zone | 12 h |
| HORVU.MOREX.r3.6HG0545620.1 | 0.361655666 | 11.02319041 | 3.613496652 | 0.00067 | 0.020873129 | -1.76398976 | Heat shock 70 kDa protein | elongation zone | 12 h |
| HORVU.MOREX.r3.4HG0398400.1 | 0.923046348 | 7.165881715 | 3.380757994 | 0.00136 | 0.026563254 | -1.7661431 | BTB/POZ domain-containing protein | elongation zone | 12 h |
| HORVU.MOREX.r3.2HG0187800.1 | 0.459672722 | 5.669988682 | 3.358478021 | 0.00146 | 0.027115369 | -1.76637582 | DNA/RNA helicase protein | elongation zone | 12 h |
| HORVU.MOREX.r3.5HG0535890.1 | 1.464014798 | 4.676851416 | 3.180977273 | 0.00245 | 0.032706849 | -1.76778893 | Protein BCCIP homolog | elongation zone | 12 h |
| HORVU.MOREX.r3.2HG0123850.1 | 1.430508719 | 2.154095266 | 3.099275686 | 0.0031 | 0.036288602 | -1.76964121 | Pentatricopeptide repeat-containing protein, putative | elongation zone | 12 h |
| HORVU.MOREX.r3.6HG0540060.1 | 1.912387354 | 3.856993307 | 3.120878159 | 0.00292 | 0.035234858 | -1.769743 | DAG, chloroplastic | elongation zone | 12 h |
| HORVU.MOREX.r3.5HG0420970.1 | -0.767277527 | 2.17073124 | -3.291425337 | 0.00178 | 0.028617771 | -1.77124361 | ERD (Early-responsive to dehydration stress) family protein | elongation zone | 12 h |

**Table S5** Continued.

| ID | logFC | AveExpr | t | P.Value | adj.P.Val | B | gene description | root zone | time point |
|---|---|---|---|---|---|---|---|---|---|
| HORVU.MOREX.r3.5HG0524260.1 | 1.645563045 | 3.901736522 | 3.139120938 | 0.00277 | 0.03425211 | -1.77309688 | #N/A | elongation zone | 12 h |
| HORVU.MOREX.r3.6HG0628810.1 | 0.881553059 | 9.237220048 | 3.478914295 | 0.00101 | 0.024104211 | -1.77341385 | 50S ribosomal protein L14 | elongation zone | 12 h |
| HORVU.MOREX.r3.4HG0413030.1 | 0.852856504 | 3.465134839 | 3.192158578 | 0.00238 | 0.03219541 | -1.77394521 | Plant/F12P19-7 protein | elongation zone | 12 h |
| HORVU.MOREX.r3.6HG0601810.1 | 2.113645424 | 0.425545697 | 3.092741384 | 0.00316 | 0.036685118 | -1.77474032 | Pentatricopeptide repeat-containing protein | elongation zone | 12 h |
| HORVU.MOREX.r3.7HG0711300.1 | -1.37202072 | -2.949290087 | -3.082217962 | 0.00326 | 0.036983216 | -1.77619775 | DNA topoisomerase | elongation zone | 12 h |
| HORVU.MOREX.r3.2HG0196740.1 | 1.738788674 | 4.447891765 | 3.171885266 | 0.00252 | 0.033048664 | -1.7772245 | DNA replication helicase dna2, putative | elongation zone | 12 h |
| HORVU.MOREX.r3.2HG0120930.1 | 1.926190855 | 9.035111364 | 3.401144803 | 0.00128 | 0.026244722 | -1.77789505 | Glycine rich protein 3 | elongation zone | 12 h |
| HORVU.MOREX.r3.4HG0333740.1 | 1.304451093 | 6.115426012 | 3.292869328 | 0.00177 | 0.028575945 | -1.77888022 | #N/A | elongation zone | 12 h |
| HORVU.MOREX.r3.3HG0291030.1 | -0.625522459 | 1.945137651 | -3.429530272 | 0.00118 | 0.025704276 | -1.78119528 | Glucan endo-1,3-beta-glucosidase 1 | elongation zone | 12 h |
| HORVU.MOREX.r3.5HG0502130.1 | 0.743880485 | 6.543409785 | 3.35973054 | 0.00145 | 0.027106935 | -1.7818264 | T-complex protein 1 subunit beta | elongation zone | 12 h |
| HORVU.MOREX.r3.1HG0089390.1 | 0.817704058 | 7.416473196 | 3.398318825 | 0.00129 | 0.026336616 | -1.78211477 | Kinase-related protein DUF1296 | elongation zone | 12 h |
| HORVU.MOREX.r3.1HG0031980.1 | 1.337742465 | 3.013898901 | 3.115000484 | 0.00297 | 0.035453096 | -1.78270687 | Trichome birefringence-like protein | elongation zone | 12 h |
| HORVU.MOREX.r3.5HG0496660.1 | 1.793506321 | 2.466297521 | 3.086539571 | 0.00322 | 0.036928455 | -1.78419333 | Sister chromatid cohesion 1 protein 3 | elongation zone | 12 h |
| HORVU.MOREX.r3.2HG0171550.1 | 0.693456337 | 4.111529363 | 3.253109234 | 0.00199 | 0.029748302 | -1.78590454 | Major facilitator superfamily transporter | elongation zone | 12 h |
| HORVU.MOREX.r3.7HG0706330.1 | 0.379726576 | 4.543866555 | 3.314757925 | 0.00166 | 0.028112171 | -1.78715163 | TOM1-like protein 2 | elongation zone | 12 h |
| HORVU.MOREX.r3.6HG0609730.1 | 0.861109311 | 8.141507493 | 3.427768638 | 0.00118 | 0.025704276 | -1.78816942 | 60S ribosomal protein L31 | elongation zone | 12 h |
| HORVU.MOREX.r3.5HG0468500.1 | 0.6853649 | 5.407656046 | 3.316524759 | 0.00165 | 0.028050531 | -1.78835407 | Diphthine synthase, putative, expressed | elongation zone | 12 h |
| HORVU.MOREX.r3.7HG0702400.1 | 0.373907307 | 6.100976213 | 3.396253652 | 0.0013 | 0.026348744 | -1.78837942 | Nucleotide/sugar transporter family protein | elongation zone | 12 h |
| HORVU.MOREX.r3.4HG0378580.1 | 0.444561012 | 5.353083837 | 3.373642335 | 0.00139 | 0.026563254 | -1.78959456 | Transcription factor | elongation zone | 12 h |
| HORVU.MOREX.r3.2HG0134290.1 | 2.21027021 | -1.166968614 | 3.098648576 | 0.00311 | 0.036312806 | -1.78960322 | SKP1-like protein 4 | elongation zone | 12 h |
| HORVU.MOREX.r3.3HG0307400.1 | -1.278950751 | 1.770342762 | -3.195657377 | 0.00235 | 0.032009236 | -1.79020936 | GRAS family transcription factor | elongation zone | 12 h |
| HORVU.MOREX.r3.2HG0111990.1 | 2.053261086 | 1.973815438 | 3.075191061 | 0.00332 | 0.037190743 | -1.79039361 | tRNA (Guanine(26)-N(2))-dimethyltransferase | elongation zone | 12 h |
| HORVU.MOREX.r3.4HG0400520.1 | -1.138277317 | -0.785996541 | -3.275627218 | 0.00186 | 0.028906691 | -1.79169687 | Lipid transfer protein | elongation zone | 12 h |
| HORVU.MOREX.r3.7HG0751620.1 | 2.010350862 | 5.432363147 | 3.206757343 | 0.00228 | 0.031719121 | -1.79289888 | Cyclin | elongation zone | 12 h |
| HORVU.MOREX.r3.7HG0641230.1 | -1.088674119 | -1.229188244 | -3.305744865 | 0.0017 | 0.028288147 | -1.79462854 | Retrovirus-related Pol polyprotein from transposon TNT 1-94 | elongation zone | 12 h |
| HORVU.MOREX.r3.2HG0105350.1 | 2.500687248 | 3.890755994 | 3.098311702 | 0.00311 | 0.036312806 | -1.79557473 | Pentatricopeptide repeat-containing protein | elongation zone | 12 h |
| HORVU.MOREX.r3.4HG0384120.1 | 0.843507092 | 2.064852893 | 3.147997473 | 0.0027 | 0.033864708 | -1.7991017 | serine/arginine repetitive matrix-like protein | elongation zone | 12 h |
| HORVU.MOREX.r3.1HG0026830.1 | -0.547525637 | 3.566935325 | -3.371672239 | 0.0014 | 0.026619212 | -1.80133095 | Leucine-rich repeat receptor-like protein kinase family | elongation zone | 12 h |
| HORVU.MOREX.r3.2HG0205050.1 | -1.768321951 | -2.492233883 | -3.084494752 | 0.00324 | 0.036928455 | -1.80143107 | Receptor-like protein kinase | elongation zone | 12 h |
| HORVU.MOREX.r3.4HG0397020.1 | 2.230720084 | 3.896750304 | 3.10249891 | 0.00307 | 0.036128346 | -1.80160474 | TPX2 (Targeting protein for Xklp2) family protein | elongation zone | 12 h |
| HORVU.MOREX.r3.4HG0383330.1 | 0.732586125 | 3.253673369 | 3.168173687 | 0.00255 | 0.033171389 | -1.8021846 | DNA polymerase III polC-type | elongation zone | 12 h |
| HORVU.MOREX.r3.6HG0621750.1 | 1.189750639 | 3.464859945 | 3.137972556 | 0.00278 | 0.034261617 | -1.80310887 | Mitochondrial transcription termination factor | elongation zone | 12 h |
| HORVU.MOREX.r3.2HG0196310.1 | -2.198046866 | -1.066921445 | -3.090995839 | 0.00318 | 0.036736339 | -1.80562193 | Glycosyltransferase | elongation zone | 12 h |
| HORVU.MOREX.r3.6HG0557770.1 | 2.724783425 | -1.181003261 | 3.115088967 | 0.00297 | 0.035453096 | -1.80589102 | FBD-associated F-box protein | elongation zone | 12 h |
| HORVU.MOREX.r3.2HG0111630.1 | -1.533920536 | -2.706877531 | -3.079563747 | 0.00328 | 0.036983216 | -1.80703395 | carboxyl-terminal peptidase (DUF239) | elongation zone | 12 h |
| HORVU.MOREX.r3.1HG0003150.1 | 0.693728718 | 4.554905765 | 3.231751194 | 0.00212 | 0.030866089 | -1.80782537 | Chaperone protein DnaJ | elongation zone | 12 h |
| HORVU.MOREX.r3.5HG0503930.1 | 1.64538256 | 4.348826856 | 3.154903175 | 0.00265 | 0.033831284 | -1.80821625 | Helicase-related family protein | elongation zone | 12 h |
| HORVU.MOREX.r3.1HG0090940.1 | 0.62038187 | 8.802081872 | 3.463629194 | 0.00106 | 0.024644437 | -1.81142345 | GTP-binding nuclear protein | elongation zone | 12 h |
| HORVU.MOREX.r3.2HG0166530.1 | 0.687476585 | 8.752681538 | 3.459326535 | 0.00108 | 0.024665565 | -1.81219179 | 30S ribosomal protein S11 | elongation zone | 12 h |
| HORVU.MOREX.r3.3HG0243990.1 | 0.758546317 | 9.014711028 | 3.461380317 | 0.00107 | 0.024644437 | -1.81289896 | 50S ribosomal protein L24 | elongation zone | 12 h |
| HORVU.MOREX.r3.4HG0357230.1 | 0.991021518 | 5.25636344 | 3.265834409 | 0.00192 | 0.029184417 | -1.81370058 | Pyridoxal 5'-phosphate synthase subunit PdxS | elongation zone | 12 h |

**Table S5** Continued.

| ID | logFC | AveExpr | t | P.Value | adj.P.Val | B | gene description | root zone | time point |
|---|---|---|---|---|---|---|---|---|---|
| HORVU.MOREX.r3.6HG0567060.1 | 2.809092855 | 1.964019752 | 3.067834992 | 0.00339 | 0.03736784 | -1.8152174 | Tyrosyl-DNA phosphodiesterase 2 | elongation zone | 12 h |
| HORVU.MOREX.r3.5HG0497930.1 | 1.877324252 | 2.406259764 | 3.069695464 | 0.00337 | 0.037331432 | -1.8169582 | Protein ROOT PRIMORDIUM DEFECTIVE 1 | elongation zone | 12 h |
| HORVU.MOREX.r3.6HG0616800.1 | 3.025961981 | 0.815040624 | 3.093507209 | 0.00315 | 0.036644453 | -1.81707179 | Pentatricopeptide repeat-containing protein | elongation zone | 12 h |
| HORVU.MOREX.r3.4HG0404070.1 | 0.884289898 | 10.53082143 | 3.513698537 | 0.00091 | 0.023235104 | -1.81888186 | 60S ribosomal protein L4 | elongation zone | 12 h |
| HORVU.MOREX.r3.5HG0536490.1 | 1.070401832 | 4.431143848 | 3.283687208 | 0.00182 | 0.028793645 | -1.819588 | Acyl carrier protein | elongation zone | 12 h |
| HORVU.MOREX.r3.6HG0581690.1 | 1.02504988 | 5.816798645 | 3.273573221 | 0.00187 | 0.028923495 | -1.82039895 | 60S ribosomal protein L6 | elongation zone | 12 h |
| HORVU.MOREX.r3.2HG0112910.1 | 0.899254479 | 4.17912055 | 3.23089499 | 0.00212 | 0.030866089 | -1.82091539 | Pectinesterase | elongation zone | 12 h |
| HORVU.MOREX.r3.6HG0559400.1 | 0.813045807 | 3.632007207 | 3.191149046 | 0.00238 | 0.032236279 | -1.82154537 | Exocyst complex component EXO70A1 | elongation zone | 12 h |
| HORVU.MOREX.r3.3HG0286160.1 | 1.508599307 | 6.290215457 | 3.29428897 | 0.00176 | 0.028575945 | -1.82232817 | Pollen Ole e 1 allergen/extensin | elongation zone | 12 h |
| HORVU.MOREX.r3.7HG0674860.1 | -2.292838694 | -0.715552338 | -3.061975207 | 0.00345 | 0.037649994 | -1.82349515 | Acyl-coenzyme A oxidase | elongation zone | 12 h |
| HORVU.MOREX.r3.7HG0662830.1 | 1.021418636 | 4.730197892 | 3.198187364 | 0.00233 | 0.031954117 | -1.82462655 | Mechanosensitive ion channel family protein | elongation zone | 12 h |
| HORVU.MOREX.r3.5HG0535750.1 | 0.804652482 | 6.040528497 | 3.296827544 | 0.00175 | 0.028442815 | -1.82552626 | Cleavage and polyadenylation specificity factor subunit 3 | elongation zone | 12 h |
| HORVU.MOREX.r3.4HG0380110.1 | 0.791828939 | 7.38894714 | 3.394110764 | 0.00131 | 0.026348744 | -1.82563306 | Mitochondrial import receptor subunit TOM40-1-like protein | elongation zone | 12 h |
| HORVU.MOREX.r3.7HG0675950.1 | 2.040727342 | 2.306247473 | 3.059931195 | 0.00347 | 0.037761197 | -1.82584263 | Pentatricopeptide repeat-containing protein | elongation zone | 12 h |
| HORVU.MOREX.r3.7HG0723410.1 | 2.073124644 | 2.947363407 | 3.067331719 | 0.0034 | 0.037381068 | -1.82597601 | A/G-specific adenine DNA glycosylase | elongation zone | 12 h |
| HORVU.MOREX.r3.7HG0697940.1 | 1.417248786 | 0.148475214 | 3.060127518 | 0.00347 | 0.037761197 | -1.82647507 | UvrABC system protein C | elongation zone | 12 h |
| HORVU.MOREX.r3.2HG0165190.1 | 0.864639839 | 5.986651179 | 3.308076501 | 0.00169 | 0.028279404 | -1.82650526 | Mitochondrial import inner membrane translocase subunit TIM2 | elongation zone | 12 h |
| HORVU.MOREX.r3.1HG0058800.1 | 0.588892113 | 3.603852967 | 3.211525537 | 0.00225 | 0.031545502 | -1.82767393 | Glycosyltransferase | elongation zone | 12 h |
| HORVU.MOREX.r3.7HG0698550.1 | 0.587840312 | 4.953445471 | 3.272580165 | 0.00188 | 0.028923495 | -1.82816091 | Zinc finger CCCH domain protein | elongation zone | 12 h |
| HORVU.MOREX.r3.4HG0395010.1 | 1.350537666 | 5.440099558 | 3.223048404 | 0.00217 | 0.031226117 | -1.82909353 | Nuclear pore complex protein NUP133 | elongation zone | 12 h |
| HORVU.MOREX.r3.5HG0465080.1 | 1.468895996 | 2.12956565 | 3.069126451 | 0.00338 | 0.037331432 | -1.82910296 | Pentatricopeptide repeat-containing family protein | elongation zone | 12 h |
| HORVU.MOREX.r3.7HG0647930.1 | 1.266580186 | 1.498899177 | 3.070651779 | 0.00337 | 0.037331001 | -1.82971555 | S-adenosyl-L-methionine-dependent methyltransferases superfa | elongation zone | 12 h |
| HORVU.MOREX.r3.7HG0695930.1 | 1.258162734 | 0.297229831 | 3.059281915 | 0.00348 | 0.037764638 | -1.83067289 | Succinate--CoA ligase [ADP-forming] subunit beta | elongation zone | 12 h |
| HORVU.MOREX.r3.2HG0147770.1 | 0.439116515 | 5.593653187 | 3.346892527 | 0.00151 | 0.027239638 | -1.8333241 | Growth-regulating factor | elongation zone | 12 h |
| HORVU.MOREX.r3.5HG0438160.1 | 0.407802487 | 5.336636433 | 3.32095254 | 0.00163 | 0.02796111 | -1.83590477 | #N/A | elongation zone | 12 h |
| HORVU.MOREX.r3.7HG0744860.1 | 0.876076148 | 8.187638293 | 3.408179291 | 0.00126 | 0.026083278 | -1.83705061 | Isoleucine--tRNA ligase | elongation zone | 12 h |
| HORVU.MOREX.r3.5HG0479210.1 | 1.084684885 | 7.833408605 | 3.387710523 | 0.00134 | 0.026498466 | -1.83705065 | HMG-Y-related protein A | elongation zone | 12 h |
| HORVU.MOREX.r3.1HG0023800.1 | 0.837157363 | 8.954713759 | 3.439059142 | 0.00114 | 0.025432695 | -1.8386441 | 60S ribosomal protein L18-like protein | elongation zone | 12 h |
| HORVU.MOREX.r3.7HG0728000.1 | -0.575742542 | 5.007313233 | -3.407020394 | 0.00126 | 0.026125006 | -1.83880177 | Glycosyltransferase | elongation zone | 12 h |
| HORVU.MOREX.r3.4HG0349500.1 | 2.071397304 | 3.603219849 | 3.068669491 | 0.00338 | 0.037339648 | -1.84003626 | Meiosis arrest female protein 1 | elongation zone | 12 h |
| HORVU.MOREX.r3.2HG0104690.1 | 2.190110422 | 4.777741937 | 3.130485093 | 0.00284 | 0.034669947 | -1.84042329 | Coiled-coil domain-containing protein 6 | elongation zone | 12 h |
| HORVU.MOREX.r3.2HG0099010.1 | -0.870444269 | -2.649782075 | -3.099362912 | 0.0031 | 0.036288602 | -1.84053881 | Glutamyl-tRNA (Gln) amidotransferase subunit A | elongation zone | 12 h |
| HORVU.MOREX.r3.5HG0524340.1 | 1.815526167 | 0.089050807 | 3.079912317 | 0.00328 | 0.036983216 | -1.84081671 | Mitochondrial transcription termination factor-like | elongation zone | 12 h |
| HORVU.MOREX.r3.4HG0338400.1 | 0.840377395 | 8.453676741 | 3.434692882 | 0.00116 | 0.02562904 | -1.84105547 | Histone H2B | elongation zone | 12 h |
| HORVU.MOREX.r3.2HG0146690.1 | 2.438126631 | 1.304324057 | 3.072122551 | 0.00335 | 0.037256101 | -1.84159382 | Pentatricopeptide repeat-containing protein | elongation zone | 12 h |
| HORVU.MOREX.r3.2HG0186490.1 | 0.978130843 | 3.316357094 | 3.110670309 | 0.003 | 0.035642237 | -1.84275119 | Transmembrane protein 14C | elongation zone | 12 h |
| HORVU.MOREX.r3.4HG0343350.1 | 1.050046888 | 5.871663592 | 3.290820002 | 0.00178 | 0.02862189 | -1.84403034 | 40S ribosomal protein S27 | elongation zone | 12 h |
| HORVU.MOREX.r3.6HG0590420.1 | 1.708161893 | 3.163300203 | 3.075969356 | 0.00332 | 0.037190743 | -1.84437015 | Pentatricopeptide repeat-containing protein | elongation zone | 12 h |
| HORVU.MOREX.r3.2HG0178110.1 | 2.648682964 | 5.730948227 | 3.132775704 | 0.00282 | 0.034576444 | -1.84443921 | transmembrane protein, putative (Protein of unknown function, | elongation zone | 12 h |
| HORVU.MOREX.r3.4HG0352920.1 | 0.687636454 | 4.352513751 | 3.23889739 | 0.00207 | 0.03060905 | -1.84543875 | Cysteine desulfurase, putative, expressed | elongation zone | 12 h |
| HORVU.MOREX.r3.5HG0511960.1 | 2.364742463 | -0.602478814 | 3.053008861 | 0.00354 | 0.037872249 | -1.84671564 | Disease resistance protein RPM1 | elongation zone | 12 h |

**Table S5** Continued.

| ID | logFC | AveExpr | t | P.Value | adj.P.Val | B | gene description | root zone | time point |
|---|---|---|---|---|---|---|---|---|---|
| HORVU.MOREX.r3.7HG0735580.1 | 2.670958571 | 5.931976842 | 3.186112138 | 0.00242 | 0.032527883 | -1.84691578 | Xyloglucan endotransglucosylase/hydrolase | elongation zone | 12 h |
| HORVU.MOREX.r3.7HG0692820.1 | 0.406934392 | 7.244663118 | 3.414953754 | 0.00123 | 0.026021602 | -1.84729198 | 2-oxoglutarate/malate translocator, chloroplastic | elongation zone | 12 h |
| HORVU.MOREX.r3.1HG0025320.1 | -1.144218237 | 0.572544846 | -3.124106943 | 0.00289 | 0.035131319 | -1.84748228 | Receptor-like kinase, putative | elongation zone | 12 h |
| HORVU.MOREX.r3.2HG0120940.1 | 2.16753448 | 6.482268735 | 3.221331993 | 0.00218 | 0.031338693 | -1.84751152 | P-loop containing nucleoside triphosphate hydrolases superfami | elongation zone | 12 h |
| HORVU.MOREX.r3.1HG0049180.1 | 0.576448547 | 5.507193954 | 3.299001942 | 0.00174 | 0.02840648 | -1.84809582 | Chaperone protein dnaJ, putative | elongation zone | 12 h |
| HORVU.MOREX.r3.3HG0230970.1 | 0.806004081 | 7.763065022 | 3.411280489 | 0.00124 | 0.026060689 | -1.84848072 | Histone H2B | elongation zone | 12 h |
| HORVU.MOREX.r3.2HG0215250.1 | 1.119017439 | 6.918385048 | 3.374409361 | 0.00139 | 0.026563254 | -1.84967117 | Peroxidase | elongation zone | 12 h |
| HORVU.MOREX.r3.6HG0631790.1 | 1.844833356 | 0.262211449 | 3.052534457 | 0.00354 | 0.037882054 | -1.85040865 | 50S ribosomal protein L9 | elongation zone | 12 h |
| HORVU.MOREX.r3.4HG0414860.1 | 2.127947158 | -0.918019416 | 3.075038886 | 0.00332 | 0.037190743 | -1.85083972 | Cysteine proteinase | elongation zone | 12 h |
| HORVU.MOREX.r3.5HG0420010.1 | -0.27669041 | 6.947534958 | -3.446709504 | 0.00112 | 0.025172156 | -1.85172151 | Sucrose-phosphatase | elongation zone | 12 h |
| HORVU.MOREX.r3.7HG0697480.1 | 0.683947457 | 4.08465238 | 3.223338387 | 0.00217 | 0.03122163 | -1.85300726 | LETM1-like protein | elongation zone | 12 h |
| HORVU.MOREX.r3.1HG0054200.1 | 0.373278625 | 5.861579496 | 3.356975909 | 0.00146 | 0.027181772 | -1.85334886 | E2 ubiquitin-conjugating-like enzyme | elongation zone | 12 h |
| HORVU.MOREX.r3.6HG0601260.1 | -0.943026534 | 3.05036359 | -3.381105698 | 0.00136 | 0.026563254 | -1.85448583 | Sulfotransferase | elongation zone | 12 h |
| HORVU.MOREX.r3.5HG0424560.1 | -0.555903246 | 7.021103992 | -3.446023956 | 0.00112 | 0.025191667 | -1.85510447 | Glutathione S-transferase | elongation zone | 12 h |
| HORVU.MOREX.r3.2HG0169670.1 | 2.468836937 | 0.536677325 | 3.068755366 | 0.00338 | 0.037339648 | -1.85570653 | Ribosome maturation factor RimP | elongation zone | 12 h |
| HORVU.MOREX.r3.7HG0635160.1 | 0.414987003 | 6.815127135 | 3.373620924 | 0.00139 | 0.026563254 | -1.85646438 | Monocopper oxidase-like protein SKU5 | elongation zone | 12 h |
| HORVU.MOREX.r3.7HG0725050.1 | -0.456077258 | 4.544757744 | -3.385193648 | 0.00135 | 0.026498466 | -1.8581384 | Protein IQ-DOMAIN 1 | elongation zone | 12 h |
| HORVU.MOREX.r3.1HG0085420.1 | 0.630699934 | 3.718026856 | 3.199585868 | 0.00232 | 0.031903923 | -1.85975936 | DNA replication complex GINS protein PSF2 | elongation zone | 12 h |
| HORVU.MOREX.r3.5HG0441110.1 | 1.585913436 | 5.425609299 | 3.195371638 | 0.00235 | 0.032009236 | -1.86059082 | #N/A | elongation zone | 12 h |
| HORVU.MOREX.r3.3HG0242030.1 | 1.339096872 | 2.331267829 | 3.072598264 | 0.00335 | 0.037256101 | -1.86116627 | Alpha/beta-Hydrolases superfamily protein | elongation zone | 12 h |
| HORVU.MOREX.r3.7HG0738670.1 | 0.826153265 | 6.751476612 | 3.332791001 | 0.00157 | 0.027655159 | -1.86141763 | Hydroxyproline-rich glycoprotein-like | elongation zone | 12 h |
| HORVU.MOREX.r3.7HG0691910.1 | 0.72962165 | 6.395808606 | 3.333452009 | 0.00157 | 0.02762467 | -1.86248752 | Regulator of chromosome condensation (RCC1) family protein | elongation zone | 12 h |
| HORVU.MOREX.r3.7HG0674590.1 | -0.974602162 | 1.003958233 | -3.127676362 | 0.00286 | 0.034856493 | -1.86273271 | Retrovirus-related Pol polyprotein from transposon TNT 1-94 | elongation zone | 12 h |
| HORVU.MOREX.r3.5HG0495520.1 | 0.946042616 | 4.992603158 | 3.229196468 | 0.00213 | 0.030911827 | -1.86283415 | Hsp70 nucleotide exchange factor fes1 | elongation zone | 12 h |
| HORVU.MOREX.r3.3HG0290890.1 | 0.895259763 | 5.267096293 | 3.237037575 | 0.00208 | 0.030699736 | -1.86331283 | TSA: Wollemia nobilis Ref_Wollemi_Transcript_16418_819 trans | elongation zone | 12 h |
| HORVU.MOREX.r3.2HG0096710.1 | -1.20833013 | -1.753677392 | -3.156147874 | 0.00264 | 0.033771061 | -1.86459458 | Cytochrome P450 | elongation zone | 12 h |
| HORVU.MOREX.r3.2HG0157500.1 | 1.647382548 | 3.758991925 | 3.087487085 | 0.00321 | 0.036913075 | -1.86667059 | Peroxidase | elongation zone | 12 h |
| HORVU.MOREX.r3.2HG0113710.1 | 0.940133773 | 6.360748126 | 3.302778984 | 0.00172 | 0.028392091 | -1.86887888 | Protein arginine methyltransferase | elongation zone | 12 h |
| HORVU.MOREX.r3.7HG0664240.1 | -0.560753126 | 2.462412603 | -3.292298787 | 0.00177 | 0.028575945 | -1.86902753 | 12-oxophytodienoate reductase-like protein | elongation zone | 12 h |
| HORVU.MOREX.r3.3HG0299220.1 | -0.478757376 | 5.799007839 | -3.438895511 | 0.00115 | 0.025432695 | -1.87001234 | Kinase family protein | elongation zone | 12 h |
| HORVU.MOREX.r3.6HG0593640.1 | 0.802997426 | 6.680974514 | 3.341030922 | 0.00153 | 0.02745794 | -1.87046502 | Pentatricopeptide repeat-containing protein | elongation zone | 12 h |
| HORVU.MOREX.r3.1HG0021320.1 | 1.46535533 | 5.526590149 | 3.208667092 | 0.00226 | 0.031672617 | -1.87101665 | 4-hydroxy-tetrahydrodipicolinate reductase | elongation zone | 12 h |
| HORVU.MOREX.r3.7HG0749060.1 | 1.552546345 | 0.00150025 | 3.043946753 | 0.00363 | 0.038433444 | -1.87373036 | Transducin/WD40 repeat-like superfamily protein | elongation zone | 12 h |
| HORVU.MOREX.r3.1HG0081810.1 | 1.610575161 | 3.256327076 | 3.083283097 | 0.00325 | 0.036948704 | -1.87508155 | tRNA-2-methylthio-N(6)-dimethylallyladenosine synthase | elongation zone | 12 h |
| HORVU.MOREX.r3.6HG0540370.1 | 1.357108844 | 3.67357023 | 3.081788786 | 0.00326 | 0.036983216 | -1.87518589 | Myb family transcription factor-like protein | elongation zone | 12 h |
| HORVU.MOREX.r3.7HG0749430.1 | 1.173471481 | 3.338386635 | 3.088426746 | 0.0032 | 0.03686745 | -1.87810903 | #N/A | elongation zone | 12 h |
| HORVU.MOREX.r3.6HG0541700.1 | 0.824878651 | 2.333953765 | 3.083827295 | 0.00324 | 0.036928455 | -1.87871996 | Mitochondrial transcription termination factor-like | elongation zone | 12 h |
| HORVU.MOREX.r3.2HG0178050.1 | -0.54577067 | 4.22962604 | -3.326929695 | 0.0016 | 0.027806246 | -1.87903004 | F-box protein | elongation zone | 12 h |
| HORVU.MOREX.r3.3HG0256960.1 | 0.936205818 | 6.374900604 | 3.293715109 | 0.00177 | 0.028575945 | -1.88047984 | 60S ribosomal protein L30 | elongation zone | 12 h |
| HORVU.MOREX.r3.2HG0107900.1 | 0.385160643 | 7.940344995 | 3.431553344 | 0.00117 | 0.025668159 | -1.88055009 | Elongation factor 1-alpha | elongation zone | 12 h |
| HORVU.MOREX.r3.4HG0333810.1 | 0.827244061 | 10.90713435 | 3.532199699 | 0.00086 | 0.022899718 | -1.88065551 | RNA-binding protein | elongation zone | 12 h |

**Table S5** Continued.

| ID | logFC | AveExpr | t | P.Value | adj.P.Val | B | gene description | root zone | time point |
|---|---|---|---|---|---|---|---|---|---|
| HORVU.MOREX.r3.2HG0128940.1 | 2.546350025 | -1.078037265 | 3.069432375 | 0.00338 | 0.037331432 | -1.88099552 | GTP cyclohydrolase 1 | elongation zone | 12 h |
| HORVU.MOREX.r3.5HG0516590.1 | 2.053432233 | 0.441151426 | 3.044809165 | 0.00362 | 0.038379585 | -1.88321796 | Exostosin family protein | elongation zone | 12 h |
| HORVU.MOREX.r3.7HG0723530.1 | -0.765526903 | 3.176738798 | -3.36437759 | 0.00143 | 0.026979742 | -1.8837425 | Cinnamoyl-CoA reductase family protein | elongation zone | 12 h |
| HORVU.MOREX.r3.5HG0427780.1 | 1.959838613 | 6.100942308 | 3.218590008 | 0.0022 | 0.031443869 | -1.88390927 | Kinesin-related protein | elongation zone | 12 h |
| HORVU.MOREX.r3.7HG0710130.1 | 0.869820404 | 7.526331028 | 3.371678477 | 0.0014 | 0.026619212 | -1.88440168 | 60S ribosomal protein L22, putative | elongation zone | 12 h |
| HORVU.MOREX.r3.3HG0254240.1 | 1.962307527 | 0.725794066 | 3.038295124 | 0.00369 | 0.038671495 | -1.88567436 | Pentatricopeptide repeat-containing protein | elongation zone | 12 h |
| HORVU.MOREX.r3.6HG0604930.1 | 1.256208466 | 6.186846553 | 3.261629238 | 0.00194 | 0.029444171 | -1.88575495 | UPF0160 protein MYG1, mitochondrial | elongation zone | 12 h |
| HORVU.MOREX.r3.2HG0162920.1 | 1.057308764 | 3.457721936 | 3.11534192 | 0.00296 | 0.035453096 | -1.8864712 | Mediator of RNA polymerase II transcription subunit 30, putative | elongation zone | 12 h |
| HORVU.MOREX.r3.2HG0153800.1 | 2.614697409 | 0.271349874 | 3.057419577 | 0.00349 | 0.037796967 | -1.88728278 | Aldo/keto reductase family-like protein | elongation zone | 12 h |
| HORVU.MOREX.r3.1HG0014320.1 | 1.732487335 | 2.690834854 | 3.039725556 | 0.00367 | 0.038641262 | -1.88786294 | Importin subunit beta-1 | elongation zone | 12 h |
| HORVU.MOREX.r3.2HG0150450.1 | 0.598216908 | 7.245663801 | 3.396372869 | 0.0013 | 0.026348744 | -1.88799092 | Pyruvate kinase | elongation zone | 12 h |
| HORVU.MOREX.r3.4HG0400700.1 | 1.428305391 | 3.339687411 | 3.07376144 | 0.00334 | 0.037244624 | -1.88888607 | DNA polymerase III PolC-type | elongation zone | 12 h |
| HORVU.MOREX.r3.3HG0307000.1 | 2.377338765 | 1.0320042 | 3.039125529 | 0.00368 | 0.038641262 | -1.88967331 | Pentatricopeptide repeat-containing protein | elongation zone | 12 h |
| HORVU.MOREX.r3.6HG0575770.1 | -0.34335707 | 5.020687417 | -3.335926473 | 0.00156 | 0.027532387 | -1.88971289 | Protein FAR1-RELATED SEQUENCE 5 | elongation zone | 12 h |
| HORVU.MOREX.r3.3HG0273010.1 | 0.914653021 | 4.355122207 | 3.173568518 | 0.00251 | 0.03299418 | -1.89165217 | Protein transport protein SEC13 | elongation zone | 12 h |
| HORVU.MOREX.r3.5HG0427050.1 | 1.197723312 | 5.45206071 | 3.195367689 | 0.00235 | 0.032009236 | -1.89184424 | 60S ribosomal protein L24 | elongation zone | 12 h |
| HORVU.MOREX.r3.7HG0735360.1 | 0.770471401 | 5.209545448 | 3.234534246 | 0.0021 | 0.030802035 | -1.89406901 | Zinc finger CCCH domain-containing protein 45 | elongation zone | 12 h |
| HORVU.MOREX.r3.6HG0543730.1 | 0.765746152 | 7.651801236 | 3.395290729 | 0.00131 | 0.026348744 | -1.89510777 | Histone H2B | elongation zone | 12 h |
| HORVU.MOREX.r3.2HG0192900.1 | 1.113319702 | 6.268762483 | 3.279741684 | 0.00184 | 0.028819302 | -1.89597683 | Xyloglucan endotransglucosylase/hydrolase | elongation zone | 12 h |
| HORVU.MOREX.r3.4HG0391960.1 | 1.059660853 | 4.485614922 | 3.165400748 | 0.00257 | 0.0332899 | -1.89607429 | ATP-dependent Clp protease ATP-binding subunit | elongation zone | 12 h |
| HORVU.MOREX.r3.2HG0119710.1 | 1.758257811 | 3.695173693 | 3.065542727 | 0.00341 | 0.037489533 | -1.89671183 | DNA/RNA helicase protein | elongation zone | 12 h |
| HORVU.MOREX.r3.1HG0014190.1 | -1.199302143 | 0.720317016 | -3.035814158 | 0.00371 | 0.038729899 | -1.89778136 | Transducin/WD-like repeat-protein | elongation zone | 12 h |
| HORVU.MOREX.r3.7HG0731600.1 | 1.192350592 | 1.21065695 | 3.029431482 | 0.00378 | 0.039090877 | -1.89794303 | NBS-LRR disease resistance protein, putative, expressed | elongation zone | 12 h |
| HORVU.MOREX.r3.3HG0223950.1 | 0.633337466 | 4.288990681 | 3.202027304 | 0.00231 | 0.031894634 | -1.89932249 | #N/A | elongation zone | 12 h |
| HORVU.MOREX.r3.2HG0160590.1 | 2.333429814 | 1.119229386 | 3.035777573 | 0.00371 | 0.038729899 | -1.89938373 | Pentatricopeptide repeat-containing protein | elongation zone | 12 h |
| HORVU.MOREX.r3.1HG0040910.1 | 0.812327561 | 8.88503314 | 3.422270767 | 0.0012 | 0.025763058 | -1.89982219 | 60S ribosomal protein L21 | elongation zone | 12 h |
| HORVU.MOREX.r3.1HG0073160.1 | -0.475802396 | 5.054880201 | -3.373968094 | 0.00139 | 0.026563254 | -1.90147676 | Histone H2A | elongation zone | 12 h |
| HORVU.MOREX.r3.3HG0316960.1 | 0.708721785 | 5.757042157 | 3.268583337 | 0.0019 | 0.029064599 | -1.90210309 | Nascent polypeptide-associated complex subunit alpha-like prot | elongation zone | 12 h |
| HORVU.MOREX.r3.3HG0285840.1 | -0.863360921 | -0.694836738 | -3.221436316 | 0.00218 | 0.031338693 | -1.90427459 | Senescence/dehydration-associated protein-like protein | elongation zone | 12 h |
| HORVU.MOREX.r3.5HG0535780.1 | 0.786219099 | 6.219128882 | 3.306284686 | 0.0017 | 0.028288147 | -1.90464601 | Transmembrane protein 70, mitochondrial | elongation zone | 12 h |
| HORVU.MOREX.r3.2HG0135650.1 | -2.205372175 | -3.487652168 | -3.027293151 | 0.0038 | 0.039110161 | -1.90518101 | Laccase | elongation zone | 12 h |
| HORVU.MOREX.r3.6HG0616160.1 | 1.235466699 | 2.633093791 | 3.046900652 | 0.0036 | 0.038272175 | -1.90842012 | ATP-dependent RNA helicase family protein | elongation zone | 12 h |
| HORVU.MOREX.r3.2HG0163860.1 | 2.196353102 | 5.202440706 | 3.105064335 | 0.00305 | 0.0360299 | -1.9091126 | Histone H2A.Z-specific chaperone CHZ1 | elongation zone | 12 h |
| HORVU.MOREX.r3.6HG0579510.1 | -1.599692123 | -0.908498026 | -3.023406093 | 0.00385 | 0.039284359 | -1.91006765 | LINE-1 reverse transcriptase like | elongation zone | 12 h |
| HORVU.MOREX.r3.4HG0371790.1 | 0.951756361 | 1.111486812 | 3.084027791 | 0.00324 | 0.036928455 | -1.91043658 | ROP binding protein kinases 1 | elongation zone | 12 h |
| HORVU.MOREX.r3.2HG0209340.1 | 0.981093056 | 5.833737893 | 3.244084009 | 0.00204 | 0.030345139 | -1.91204768 | Clathrin light chain | elongation zone | 12 h |
| HORVU.MOREX.r3.4HG0406290.1 | 1.755855789 | 4.255608694 | 3.079480867 | 0.00328 | 0.036983216 | -1.912638 | DNA-directed RNA polymerase subunit | elongation zone | 12 h |
| HORVU.MOREX.r3.3HG0222390.1 | 3.573051669 | 3.41727625 | 3.024371245 | 0.00384 | 0.03925602 | -1.91357755 | Trypsin inhibitor | elongation zone | 12 h |
| HORVU.MOREX.r3.3HG0314070.1 | -1.202460791 | -1.612808778 | -3.120020302 | 0.00292 | 0.035293865 | -1.9147704 | Protein kinase family protein | elongation zone | 12 h |
| HORVU.MOREX.r3.3HG0228860.1 | 3.492194609 | 2.735909792 | 3.025125864 | 0.00383 | 0.03925602 | -1.91621741 | Mitochondrial glycoprotein | elongation zone | 12 h |
| HORVU.MOREX.r3.3HG0302410.1 | 0.910267216 | 8.959776352 | 3.405849364 | 0.00126 | 0.026164076 | -1.91647051 | 30S ribosomal protein S5 | elongation zone | 12 h |

**Table S5** Continued.

| ID | logFC | AveExpr | t | P.Value | adj.P.Val | B | gene description | root zone | time point |
|---|---|---|---|---|---|---|---|---|---|
| HORVU.MOREX.r3.5HG0498810.1 | 0.69698823 | 6.020060381 | 3.296377117 | 0.00175 | 0.028450551 | -1.91972868 | 60S ribosomal protein L44 | elongation zone | 12 h |
| HORVU.MOREX.r3.3HG0270170.1 | 1.016309305 | 2.933164716 | 3.085400975 | 0.00323 | 0.036928455 | -1.92012612 | Anthranilate phosphoribosyltransferase | elongation zone | 12 h |
| HORVU.MOREX.r3.3HG0239160.1 | 1.507870861 | 3.929564631 | 3.10133095 | 0.00308 | 0.036191331 | -1.92096572 | DNA polymerase epsilon subunit | elongation zone | 12 h |
| HORVU.MOREX.r3.6HG0582230.1 | 0.548537175 | 3.581325437 | 3.209292196 | 0.00226 | 0.031636696 | -1.92224216 | GDSL esterase/lipase | elongation zone | 12 h |
| HORVU.MOREX.r3.3HG0228120.1 | -0.594905974 | 2.921074778 | -3.211031712 | 0.00225 | 0.031545502 | -1.92277267 | MYB-related transcription factor | elongation zone | 12 h |
| HORVU.MOREX.r3.4HG0340710.1 | 1.471725754 | 3.663251486 | 3.079821337 | 0.00328 | 0.036983216 | -1.92300507 | WD40 repeat-containing protein | elongation zone | 12 h |
| HORVU.MOREX.r3.2HG0206470.1 | -0.466690454 | 6.277699834 | -3.422636789 | 0.0012 | 0.025763058 | -1.92517253 | Loricrin-like protein | elongation zone | 12 h |
| HORVU.MOREX.r3.5HG0446090.1 | 1.599034996 | 3.10746457 | 3.051692243 | 0.00355 | 0.037906495 | -1.92581835 | rRNA-processing protein FYV7 | elongation zone | 12 h |
| HORVU.MOREX.r3.3HG0327680.1 | 1.861832812 | 2.040746476 | 3.019839113 | 0.00388 | 0.039444701 | -1.92601678 | Pentatricopeptide repeat-containing protein | elongation zone | 12 h |
| HORVU.MOREX.r3.2HG0205020.1 | 1.780853014 | 1.613831825 | 3.020852772 | 0.00387 | 0.039407591 | -1.92620632 | Glycosyltransferase | elongation zone | 12 h |
| HORVU.MOREX.r3.6HG0627910.1 | 0.891987141 | 6.146155843 | 3.278004741 | 0.00185 | 0.028860537 | -1.92640855 | Kinesin like protein | elongation zone | 12 h |
| HORVU.MOREX.r3.5HG0465780.1 | 1.282633811 | 6.661609217 | 3.247666372 | 0.00202 | 0.030159961 | -1.92705209 | 60S acidic ribosomal protein P0 | elongation zone | 12 h |
| HORVU.MOREX.r3.1HG0076540.1 | 0.418591389 | 7.761395839 | 3.403238302 | 0.00127 | 0.026178272 | -1.9292884 | Mitochondrial-processing peptidase subunit alpha | elongation zone | 12 h |
| HORVU.MOREX.r3.5HG0527230.1 | 1.022326258 | 3.048538089 | 3.069393582 | 0.00338 | 0.037331432 | -1.92933284 | DCD (Development and cell death) domain protein | elongation zone | 12 h |
| HORVU.MOREX.r3.1HG0093330.1 | 1.743233156 | 1.327218247 | 3.014958332 | 0.00394 | 0.039714168 | -1.93055096 | Protein N-methyltransferase | elongation zone | 12 h |
| HORVU.MOREX.r3.4HG0412190.1 | 1.406649064 | 2.746246921 | 3.045296464 | 0.00362 | 0.038346619 | -1.93101342 | 50S ribosomal protein L5 | elongation zone | 12 h |
| HORVU.MOREX.r3.2HG0099920.1 | 0.840652582 | 2.370731385 | 3.093527985 | 0.00315 | 0.036644453 | -1.93242924 | Glyoxylate reductase/hydroxypyruvate reductase | elongation zone | 12 h |
| HORVU.MOREX.r3.7HG0684880.1 | -0.649635025 | 3.761266823 | -3.269109531 | 0.0019 | 0.029041385 | -1.93358931 | meiosis chromosome segregation family protein | elongation zone | 12 h |
| HORVU.MOREX.r3.7HG0646210.1 | 0.709898698 | 4.882966115 | 3.215857825 | 0.00222 | 0.031454344 | -1.93399297 | tRNA-modifying protein YgfZ | elongation zone | 12 h |
| HORVU.MOREX.r3.3HG0324310.1 | 1.852990761 | -0.544565087 | 3.023076304 | 0.00385 | 0.039301249 | -1.93421854 | Disease resistance protein RPP13 | elongation zone | 12 h |
| HORVU.MOREX.r3.3HG0305200.1 | 1.634880304 | 3.248140536 | 3.044596326 | 0.00362 | 0.038382831 | -1.9356399 | Monooxygenase | elongation zone | 12 h |
| HORVU.MOREX.r3.6HG0577950.1 | -1.425750963 | -1.781795644 | -3.047402306 | 0.00359 | 0.038237792 | -1.93585958 | F-box/kelch-repeat protein | elongation zone | 12 h |
| HORVU.MOREX.r3.4HG0409600.1 | -0.536529812 | 4.470888926 | -3.375088533 | 0.00139 | 0.026563254 | -1.9362235 | Protein kinase family protein, putative, expressed | elongation zone | 12 h |
| HORVU.MOREX.r3.2HG0139810.1 | -0.839989314 | 0.636540026 | -3.340339499 | 0.00154 | 0.02745794 | -1.93622675 | Heavy metal transport/detoxification superfamily protein | elongation zone | 12 h |
| HORVU.MOREX.r3.2HG0124930.1 | 0.623321904 | 6.004755837 | 3.28117205 | 0.00183 | 0.028819302 | -1.93671942 | DNA-directed RNA polymerase II family protein | elongation zone | 12 h |
| HORVU.MOREX.r3.4HG0400870.1 | 2.983484565 | 0.562362935 | 3.048267519 | 0.00359 | 0.03818403 | -1.93802044 | Ribosomal large subunit pseudouridine synthase B | elongation zone | 12 h |
| HORVU.MOREX.r3.1HG0065030.2 | 0.316154394 | 5.236932218 | 3.287205741 | 0.0018 | 0.028691351 | -1.93802664 | Nucleic acid binding protein | elongation zone | 12 h |
| HORVU.MOREX.r3.2HG0117180.1 | 0.838449881 | 7.219922766 | 3.335738548 | 0.00156 | 0.027532387 | -1.93986486 | 30S ribosomal protein S9 | elongation zone | 12 h |
| HORVU.MOREX.r3.7HG0683960.1 | 1.19900695 | 5.237797772 | 3.178988735 | 0.00247 | 0.032817853 | -1.94087579 | #N/A | elongation zone | 12 h |
| HORVU.MOREX.r3.4HG0388470.1 | 1.962704821 | 0.659213336 | 3.015535082 | 0.00393 | 0.039669391 | -1.94144872 | Rotundifolia-like protein | elongation zone | 12 h |
| HORVU.MOREX.r3.4HG0359890.1 | 0.671120889 | 3.726697938 | 3.126025365 | 0.00287 | 0.035000997 | -1.94168221 | tRNA-2-methylthio-N(6)-dimethylallyladenosine synthase | elongation zone | 12 h |
| HORVU.MOREX.r3.5HG0462720.1 | 0.860241772 | 4.47715704 | 3.151101263 | 0.00267 | 0.033864708 | -1.94273687 | WD-repeat protein, putative | elongation zone | 12 h |
| HORVU.MOREX.r3.7HG0658660.1 | -0.809424112 | 2.135277695 | -3.157826583 | 0.00262 | 0.033716726 | -1.94405439 | Ethylene-responsive transcription factor | elongation zone | 12 h |
| HORVU.MOREX.r3.2HG0116060.1 | 0.740923569 | 4.617910192 | 3.181165681 | 0.00245 | 0.032706849 | -1.9441837 | Pre-rRNA-processing protein ESF2 | elongation zone | 12 h |
| HORVU.MOREX.r3.6HG0557980.1 | 1.040467687 | 6.414210139 | 3.268303239 | 0.0019 | 0.029066843 | -1.94520601 | #N/A | elongation zone | 12 h |
| HORVU.MOREX.r3.2HG0114280.1 | 2.39465624 | 0.876588196 | 3.027293626 | 0.0038 | 0.039110161 | -1.94676949 | GTPase obg | elongation zone | 12 h |
| HORVU.MOREX.r3.1HG0050920.1 | -0.840532016 | 2.478719451 | -3.348718584 | 0.0015 | 0.027239638 | -1.94892274 | Trehalase | elongation zone | 12 h |
| HORVU.MOREX.r3.3HG0291600.1 | 0.879756792 | 6.498873317 | 3.279871126 | 0.00184 | 0.028819302 | -1.95075239 | ATP-dependent RNA helicase, putative | elongation zone | 12 h |
| HORVU.MOREX.r3.5HG0442460.1 | -0.516510715 | 3.36756339 | -3.369792428 | 0.00141 | 0.026719424 | -1.95144848 | Protein kinase family protein | elongation zone | 12 h |
| HORVU.MOREX.r3.2HG0173660.1 | 2.260590702 | 1.241411952 | 3.018537658 | 0.0039 | 0.03951103 | -1.95202785 | Pentatricopeptide repeat-containing family protein | elongation zone | 12 h |
| HORVU.MOREX.r3.1HG0095070.2 | 0.772874098 | 5.248934381 | 3.233499258 | 0.00211 | 0.030829102 | -1.95355988 | ATP-dependent RNA helicase | elongation zone | 12 h |

**Table S5** Continued.

| ID | logFC | AveExpr | t | P.Value | adj.P.Val | B | gene description | root zone | time point |
|---|---|---|---|---|---|---|---|---|---|
| HORVU.MOREX.r3.4HG0331540.1 | 1.368698864 | 1.537888525 | 3.014185878 | 0.00395 | 0.039746874 | -1.95381613 | ACT domain-containing protein | elongation zone | 12 h |
| HORVU.MOREX.r3.7HG0715110.1 | 3.035833241 | -1.38874377 | 3.0391258 | 0.00368 | 0.038641262 | -1.95533259 | Double-strand break repair protein MRE11 | elongation zone | 12 h |
| HORVU.MOREX.r3.1HG0054820.1 | 1.821744079 | 1.976629801 | 3.008976096 | 0.00401 | 0.040012973 | -1.95737717 | Pentatricopeptide repeat-containing protein | elongation zone | 12 h |
| HORVU.MOREX.r3.4HG0402030.1 | 0.390885137 | 5.321937852 | 3.290484751 | 0.00178 | 0.02862189 | -1.96074654 | Basic helix-loop-helix (BHLH) family transcription factor | elongation zone | 12 h |
| HORVU.MOREX.r3.4HG0406710.1 | -0.536291124 | 2.527858084 | -3.29036666 | 0.00178 | 0.02862189 | -1.96127138 | Homeobox protein, putative | elongation zone | 12 h |
| HORVU.MOREX.r3.7HG0669190.1 | 1.680398279 | 2.070139492 | 3.009840933 | 0.004 | 0.039974405 | -1.96226298 | Adenylyltransferase and sulfurtransferase MOCS3 | elongation zone | 12 h |
| HORVU.MOREX.r3.5HG0426820.1 | 2.193150953 | -0.173769083 | 3.026645546 | 0.00381 | 0.039161886 | -1.96286972 | Pentatricopeptide repeat-containing protein | elongation zone | 12 h |
| HORVU.MOREX.r3.7HG0722300.1 | 2.361106359 | -0.952880022 | 3.032897019 | 0.00374 | 0.038936762 | -1.9629701 | Lipase | elongation zone | 12 h |
| HORVU.MOREX.r3.5HG0432300.1 | 0.776798545 | 6.175259496 | 3.280545405 | 0.00184 | 0.028819302 | -1.96508784 | Tetratricopeptide repeat-containing family protein | elongation zone | 12 h |
| HORVU.MOREX.r3.6HG0539850.1 | 1.832443221 | 4.423652728 | 3.076687214 | 0.00331 | 0.037174732 | -1.96566272 | Kinesin-like protein | elongation zone | 12 h |
| HORVU.MOREX.r3.7HG0666080.1 | 2.439192345 | 1.746087834 | 2.99961963 | 0.00411 | 0.040307075 | -1.96686959 | Pentatricopeptide repeat-containing protein | elongation zone | 12 h |
| HORVU.MOREX.r3.5HG0478950.1 | 1.610976482 | 3.390125726 | 3.031137565 | 0.00376 | 0.03899655 | -1.96883031 | Centromere protein V | elongation zone | 12 h |
| HORVU.MOREX.r3.4HG0349170.1 | 1.092584494 | 4.64522884 | 3.255451972 | 0.00198 | 0.029636522 | -1.97288724 | Long-Chain Acyl-CoA Synthetase | elongation zone | 12 h |
| HORVU.MOREX.r3.6HG0613230.1 | 0.351954535 | 6.00935735 | 3.360410818 | 0.00145 | 0.027076709 | -1.97352793 | Allergen, putative | elongation zone | 12 h |
| HORVU.MOREX.r3.4HG0384570.1 | 0.787788514 | 4.311382607 | 3.139942695 | 0.00276 | 0.034221402 | -1.97540936 | Transcription factor bHLH140 | elongation zone | 12 h |
| HORVU.MOREX.r3.4HG0402670.1 | 1.588832253 | 4.040868248 | 3.057467253 | 0.00349 | 0.037796967 | -1.9759917 | DNA topoisomerase | elongation zone | 12 h |
| HORVU.MOREX.r3.2HG0113160.1 | 0.749053454 | 8.454057913 | 3.380620149 | 0.00136 | 0.026563254 | -1.9766184 | 60S ribosomal protein L22, putative | elongation zone | 12 h |
| HORVU.MOREX.r3.4HG0405370.1 | 2.181412764 | 0.021144763 | 3.012204696 | 0.00397 | 0.039849721 | -1.97669429 | Pentatricopeptide repeat-containing protein | elongation zone | 12 h |
| HORVU.MOREX.r3.4HG0377890.1 | 0.936541272 | 8.002111855 | 3.32791172 | 0.0016 | 0.027806246 | -1.97676155 | Nucleolar GTP-binding protein 2 | elongation zone | 12 h |
| HORVU.MOREX.r3.3HG0272120.1 | 1.457174544 | 4.043401107 | 3.058353692 | 0.00348 | 0.037779464 | -1.97684592 | DVA-1 polyprotein | elongation zone | 12 h |
| HORVU.MOREX.r3.7HG0661600.1 | 1.314929305 | 7.721715927 | 3.285659619 | 0.00181 | 0.028744755 | -1.97761955 | Nucleolar GTP-binding protein 1 | elongation zone | 12 h |
| HORVU.MOREX.r3.4HG0393980.1 | 2.254339775 | 1.396411503 | 2.997710006 | 0.00413 | 0.040391635 | -1.97876604 | DNA-directed RNA polymerase subunit beta | elongation zone | 12 h |
| HORVU.MOREX.r3.1HG0044230.1 | 0.774774547 | 8.794273122 | 3.394478445 | 0.00131 | 0.026348744 | -1.98058872 | 40S ribosomal protein S17, putative, expressed | elongation zone | 12 h |
| HORVU.MOREX.r3.5HG0528890.1 | 1.137532095 | 5.084336588 | 3.155498856 | 0.00264 | 0.033804921 | -1.98195742 | GATA transcription factor, putative | elongation zone | 12 h |
| HORVU.MOREX.r3.1HG0082060.1 | -0.662969886 | 2.494333616 | -3.114791948 | 0.00297 | 0.035453096 | -1.98245689 | epstein-barr nuclear antigen | elongation zone | 12 h |
| HORVU.MOREX.r3.4HG0374180.1 | 1.239260768 | 0.480598598 | 2.99893698 | 0.00412 | 0.040329959 | -1.98272281 | TSC22 domain family protein 1 | elongation zone | 12 h |
| HORVU.MOREX.r3.7HG0641750.1 | -1.784449415 | -2.739012037 | -3.054734652 | 0.00352 | 0.037808566 | -1.98526921 | Aspartic proteinase nepenthesin-2 | elongation zone | 12 h |
| HORVU.MOREX.r3.5HG0527180.1 | 1.152590365 | 3.570770348 | 3.065184531 | 0.00342 | 0.037489533 | -1.98528999 | HAT transposon superfamily, putative | elongation zone | 12 h |
| HORVU.MOREX.r3.7HG0664470.1 | 1.976558575 | 2.932550253 | 3.00307849 | 0.00407 | 0.040229538 | -1.9863202 | KAT8 regulatory NSL complex subunit 3 | elongation zone | 12 h |
| HORVU.MOREX.r3.5HG0514110.1 | 0.559242217 | 6.244309832 | 3.304545409 | 0.00171 | 0.02834146 | -1.98703621 | #N/A | elongation zone | 12 h |
| HORVU.MOREX.r3.4HG0407520.1 | 1.088490314 | 2.035180635 | 3.001467573 | 0.00409 | 0.040265883 | -1.98744656 | Mitochondrial import inner membrane translocase subunit TIM2 | elongation zone | 12 h |
| HORVU.MOREX.r3.3HG0308640.1 | -0.335025446 | 7.154318973 | -3.432491967 | 0.00117 | 0.025668159 | -1.98876882 | lysine ketoglutarate reductase trans-splicing protein (DUF707) | elongation zone | 12 h |
| HORVU.MOREX.r3.5HG0419590.1 | -0.474850288 | 6.133126066 | -3.404065661 | 0.00127 | 0.026178272 | -1.99018642 | #N/A | elongation zone | 12 h |
| HORVU.MOREX.r3.5HG0452940.1 | 2.13309728 | 1.671260459 | 2.989021152 | 0.00424 | 0.040956128 | -1.99109355 | Pentatricopeptide repeat-containing protein | elongation zone | 12 h |
| HORVU.MOREX.r3.3HG0255400.1 | 1.167708274 | 5.346293272 | 3.173114539 | 0.00251 | 0.032994826 | -1.99180251 | Translin family protein | elongation zone | 12 h |
| HORVU.MOREX.r3.1HG0069950.1 | 1.131950512 | 4.849761786 | 3.137107914 | 0.00278 | 0.034272194 | -1.99193763 | Protein KRI1 | elongation zone | 12 h |
| HORVU.MOREX.r3.1HG0028320.1 | 0.388656968 | 6.850868413 | 3.35390038 | 0.00148 | 0.027239638 | -1.99354767 | 70 kDa heat shock protein | elongation zone | 12 h |
| HORVU.MOREX.r3.1HG0082500.1 | 0.918974402 | 4.721829471 | 3.159597962 | 0.00261 | 0.033617799 | -1.99376074 | DNA replication complex GINS protein PSF3 | elongation zone | 12 h |
| HORVU.MOREX.r3.5HG0511950.1 | 1.749451765 | 4.233239305 | 3.057996806 | 0.00349 | 0.037779464 | -1.99492776 | Kinesin-like protein | elongation zone | 12 h |
| HORVU.MOREX.r3.5HG0512440.1 | 1.397386263 | 5.921251417 | 3.189414974 | 0.00239 | 0.032324245 | -1.99496849 | #N/A | elongation zone | 12 h |
| HORVU.MOREX.r3.3HG0303380.1 | -0.758451374 | -0.867377041 | -3.177082926 | 0.00248 | 0.032834207 | -1.99528868 | Zinc finger protein CONSTANS | elongation zone | 12 h |

**Table S5** Continued.

| ID | logFC | AveExpr | t | P.Value | adj.P.Val | B | gene description | root zone | time point |
|---|---|---|---|---|---|---|---|---|---|
| HORVU.MOREX.r3.4HG0379050.1 | -0.349419688 | 5.079597644 | -3.277785047 | 0.00185 | 0.028860537 | -1.99694732 | Receptor-like kinase | elongation zone | 12 h |
| HORVU.MOREX.r3.4HG0345810.1 | 1.296974257 | 1.299714382 | 2.994563035 | 0.00417 | 0.040632794 | -1.99894516 | #N/A | elongation zone | 12 h |
| HORVU.MOREX.r3.7HG0684940.1 | -0.556588218 | 4.72025707 | -3.27968366 | 0.00184 | 0.028819302 | -2.00001161 | Zinc finger constans-like protein | elongation zone | 12 h |
| HORVU.MOREX.r3.5HG0516470.1 | 1.478568679 | 4.720536401 | 3.077446986 | 0.0033 | 0.037119446 | -2.00103448 | GDSL esterase/lipase | elongation zone | 12 h |
| HORVU.MOREX.r3.3HG0266740.1 | 0.602464238 | 4.583959119 | 3.197667162 | 0.00234 | 0.03195973 | -2.00181721 | Pentatricopeptide repeat-containing protein | elongation zone | 12 h |
| HORVU.MOREX.r3.6HG0617080.1 | -0.509498602 | 4.452280294 | -3.317262259 | 0.00165 | 0.028043664 | -2.00497971 | Lipid transfer protein | elongation zone | 12 h |
| HORVU.MOREX.r3.4HG0337200.1 | 1.258608172 | 4.676780151 | 3.095126944 | 0.00314 | 0.036539994 | -2.00856472 | Receptor-like kinase | elongation zone | 12 h |
| HORVU.MOREX.r3.6HG0631440.1 | 1.290996814 | 2.780766611 | 3.036311371 | 0.00371 | 0.038729899 | -2.00994738 | Janus kinase and microtubule-interacting protein 3 | elongation zone | 12 h |
| HORVU.MOREX.r3.5HG0508640.1 | 2.689467447 | 1.961761663 | 2.982020516 | 0.00432 | 0.04129762 | -2.01023655 | Protein recA | elongation zone | 12 h |
| HORVU.MOREX.r3.1HG0047710.1 | 1.09909545 | 1.697525089 | 2.99927559 | 0.00412 | 0.040310975 | -2.01071857 | Ribosomal RNA small subunit methyltransferase H | elongation zone | 12 h |
| HORVU.MOREX.r3.6HG0614700.1 | 0.935297452 | 2.36393809 | 3.055497352 | 0.00351 | 0.037808566 | -2.01090611 | ATP synthase delta-subunit protein | elongation zone | 12 h |
| HORVU.MOREX.r3.3HG0319710.1 | 3.585050404 | 1.482728189 | 3.005869954 | 0.00404 | 0.040178373 | -2.0111656 | Glutathione S-transferase | elongation zone | 12 h |
| HORVU.MOREX.r3.1HG0080610.1 | 1.640693674 | 4.932591417 | 3.094795761 | 0.00314 | 0.036553757 | -2.01214036 | Flap endonuclease 1 | elongation zone | 12 h |
| HORVU.MOREX.r3.1HG0005370.1 | 0.593479208 | 4.817106872 | 3.194615452 | 0.00236 | 0.032009236 | -2.01328079 | Replication protein A 70 kDa DNA-binding subunit | elongation zone | 12 h |
| HORVU.MOREX.r3.1HG0006610.1 | 0.908887879 | 8.693329567 | 3.378025123 | 0.00137 | 0.026563254 | -2.01344622 | Histone H2A | elongation zone | 12 h |
| HORVU.MOREX.r3.5HG0537290.1 | 0.714790323 | 7.867863715 | 3.340148531 | 0.00154 | 0.02745794 | -2.01395241 | 60 kDa chaperonin | elongation zone | 12 h |
| HORVU.MOREX.r3.3HG0325290.1 | 0.758564558 | 7.787813279 | 3.337378813 | 0.00155 | 0.027528501 | -2.01418639 | Ribosomal protein | elongation zone | 12 h |
| HORVU.MOREX.r3.2HG0149130.1 | 0.373987121 | 5.403092628 | 3.275419855 | 0.00186 | 0.028906691 | -2.0144363 | Ubiquitin-conjugating enzyme (E2) | elongation zone | 12 h |
| HORVU.MOREX.r3.6HG0604200.1 | 0.925619712 | 0.7623065 | 3.081636822 | 0.00326 | 0.036983216 | -2.01472793 | transcription repressor | elongation zone | 12 h |
| HORVU.MOREX.r3.2HG0117160.1 | 2.052973629 | 5.56954998 | 3.118646001 | 0.00294 | 0.035349763 | -2.01605537 | Receptor-like kinase | elongation zone | 12 h |
| HORVU.MOREX.r3.3HG0290180.1 | 1.170073441 | 3.615357964 | 3.0411389 | 0.00366 | 0.038619667 | -2.01645666 | Non-specific serine/threonine protein kinase | elongation zone | 12 h |
| HORVU.MOREX.r3.7HG0649260.1 | 1.377077601 | 6.056376679 | 3.181807724 | 0.00245 | 0.032706849 | -2.01661306 | #N/A | elongation zone | 12 h |
| HORVU.MOREX.r3.2HG0151050.1 | 0.562562452 | 6.080741385 | 3.271558388 | 0.00188 | 0.028962927 | -2.01894622 | AT-rich interactive domain protein | elongation zone | 12 h |
| HORVU.MOREX.r3.5HG0522390.1 | 2.480903918 | -0.232655981 | 3.004364547 | 0.00406 | 0.040229538 | -2.01942697 | Pheophorbide a oxygenase, chloroplastic | elongation zone | 12 h |
| HORVU.MOREX.r3.6HG0552540.1 | 0.80561097 | 3.876707041 | 3.090863795 | 0.00318 | 0.036736339 | -2.01951769 | Pyridoxal 5'-phosphate synthase subunit PdxT | elongation zone | 12 h |
| HORVU.MOREX.r3.3HG0286150.1 | -1.851403181 | -0.866760921 | -2.977273365 | 0.00438 | 0.041519226 | -2.02142077 | PHD finger protein 3 | elongation zone | 12 h |
| HORVU.MOREX.r3.2HG0172350.1 | 1.990114237 | 5.213074774 | 3.072944245 | 0.00334 | 0.037254747 | -2.0214784 | WEB family protein, chloroplastic | elongation zone | 12 h |
| HORVU.MOREX.r3.7HG0637270.2 | 1.271661389 | 2.556416211 | 3.004559437 | 0.00406 | 0.040229538 | -2.02222424 | Ribonuclease H | elongation zone | 12 h |
| HORVU.MOREX.r3.6HG0611880.1 | -0.306103832 | 7.734249847 | -3.410772282 | 0.00125 | 0.026062006 | -2.02236237 | Homeobox-leucine zipper protein | elongation zone | 12 h |
| HORVU.MOREX.r3.3HG0221390.1 | 0.978317211 | 2.076992594 | 2.999908263 | 0.00411 | 0.040307075 | -2.02603324 | LisH and RanBPM domains containing protein | elongation zone | 12 h |
| HORVU.MOREX.r3.1HG0054010.1 | -0.546819176 | 5.84165697 | -3.374103962 | 0.00139 | 0.026563254 | -2.026295 | zinc finger B-box protein | elongation zone | 12 h |
| HORVU.MOREX.r3.7HG0728630.1 | 0.978589621 | 1.895271753 | 3.017722096 | 0.00391 | 0.039547618 | -2.02906501 | Receptor-like protein kinase | elongation zone | 12 h |
| HORVU.MOREX.r3.3HG0330150.1 | -1.578195 | -0.302066567 | -3.033797758 | 0.00374 | 0.03886761 | -2.0298084 | MYB transcription factor | elongation zone | 12 h |
| HORVU.MOREX.r3.5HG0512410.1 | 1.275319316 | 5.302905234 | 3.146374751 | 0.00271 | 0.033864708 | -2.03038299 | Upstream activation factor subunit spp27 | elongation zone | 12 h |
| HORVU.MOREX.r3.6HG0634090.1 | 3.03988742 | 2.903382382 | 2.971967203 | 0.00444 | 0.041943839 | -2.03059916 | Protein O-linked-mannose beta-1,4-N-acetylglucosaminyltransfe | elongation zone | 12 h |
| HORVU.MOREX.r3.6HG0588350.1 | 0.504168184 | 4.125370776 | 3.170403179 | 0.00253 | 0.033099534 | -2.03358646 | RING finger family protein | elongation zone | 12 h |
| HORVU.MOREX.r3.7HG0716900.1 | 0.339952165 | 8.192568258 | 3.399172031 | 0.00129 | 0.026336616 | -2.03370085 | Transmembrane 9 superfamily member | elongation zone | 12 h |
| HORVU.MOREX.r3.4HG0357930.1 | -0.668610941 | 2.087250112 | -3.112239928 | 0.00299 | 0.035565004 | -2.03453356 | Scarecrow transcription factor family protein | elongation zone | 12 h |
| HORVU.MOREX.r3.5HG0501840.1 | 0.900721552 | 5.034273177 | 3.149345809 | 0.00269 | 0.033864708 | -2.03718885 | DNA repair protein-like protein | elongation zone | 12 h |
| HORVU.MOREX.r3.2HG0099880.2 | 1.839223161 | 1.089601057 | 2.974813334 | 0.00441 | 0.041702155 | -2.03726711 | caspase-6 protein | elongation zone | 12 h |
| HORVU.MOREX.r3.5HG0471060.1 | 0.793411069 | 5.409891759 | 3.211406304 | 0.00225 | 0.031545502 | -2.03739805 | topoisomerase 6 subunit B | elongation zone | 12 h |

**Table S5** Continued.

| ID | logFC | AveExpr | t | P.Value | adj.P.Val | B | gene description | root zone | time point |
|---|---|---|---|---|---|---|---|---|---|
| HORVU.MOREX.r3.7HG0652090.1 | 1.562525304 | 2.809206881 | 2.990464023 | 0.00422 | 0.040887644 | -2.03785072 | Pentatricopeptide repeat-containing protein | elongation zone | 12 h |
| HORVU.MOREX.r3.7HG0723540.1 | -0.954100183 | 3.755359913 | -3.317646652 | 0.00164 | 0.028035058 | -2.03846759 | Cinnamoyl-CoA reductase family protein | elongation zone | 12 h |
| HORVU.MOREX.r3.2HG0217620.1 | 0.871340955 | 4.501749172 | 3.10731828 | 0.00303 | 0.03590208 | -2.03973234 | Pentatricopeptide repeat-containing family protein | elongation zone | 12 h |
| HORVU.MOREX.r3.4HG0381080.1 | 0.838831122 | 2.763058409 | 3.059808538 | 0.00347 | 0.037761197 | -2.03977168 | Fe/S biogenesis protein nfuA | elongation zone | 12 h |
| HORVU.MOREX.r3.6HG0620520.1 | -0.627597106 | 3.094519972 | -3.307047388 | 0.0017 | 0.028288147 | -2.04004377 | Aminotransferase-related family protein | elongation zone | 12 h |
| HORVU.MOREX.r3.4HG0397940.1 | 0.673610665 | 3.598811101 | 3.157172736 | 0.00263 | 0.033716726 | -2.0405278 | Alpha-L-fucosidase 2 | elongation zone | 12 h |
| HORVU.MOREX.r3.7HG0724090.1 | 2.030323108 | 0.704867316 | 2.974702572 | 0.00441 | 0.041702155 | -2.04112952 | RING/U-box superfamily protein | elongation zone | 12 h |
| HORVU.MOREX.r3.6HG0607360.1 | 0.564019848 | 4.502873455 | 3.177520544 | 0.00248 | 0.032834207 | -2.0417192 | WAT1-related protein | elongation zone | 12 h |
| HORVU.MOREX.r3.5HG0479520.1 | 3.250399284 | 3.438825405 | 2.965396346 | 0.00452 | 0.042347835 | -2.04400335 | DNA mismatch repair protein MutS | elongation zone | 12 h |
| HORVU.MOREX.r3.2HG0182220.1 | 1.989149063 | 1.158860243 | 2.963958354 | 0.00454 | 0.04241279 | -2.04538683 | Pentatricopeptide repeat-containing protein | elongation zone | 12 h |
| HORVU.MOREX.r3.6HG0626980.1 | 2.115976167 | -1.590959919 | 2.996306248 | 0.00415 | 0.040512048 | -2.04604321 | DNA topoisomerase 2 | elongation zone | 12 h |
| HORVU.MOREX.r3.6HG0555700.1 | 0.367696615 | 8.875975145 | 3.432800022 | 0.00117 | 0.025668159 | -2.04613132 | Calreticulin/calnexin | elongation zone | 12 h |
| HORVU.MOREX.r3.7HG0701910.1 | 0.963597551 | 5.137401888 | 3.167832282 | 0.00255 | 0.03318302 | -2.04634802 | RNA binding protein, putative | elongation zone | 12 h |
| HORVU.MOREX.r3.4HG0407060.1 | 0.39061983 | 7.117015257 | 3.329968533 | 0.00159 | 0.027768577 | -2.04748106 | Cysteine--tRNA ligase | elongation zone | 12 h |
| HORVU.MOREX.r3.1HG0057980.1 | 1.864758059 | 0.759657688 | 2.963543166 | 0.00455 | 0.04241279 | -2.04878264 | Sugar transporter, putative | elongation zone | 12 h |
| HORVU.MOREX.r3.7HG0719480.1 | 1.002874789 | 4.687776647 | 3.140483133 | 0.00276 | 0.034208419 | -2.04951961 | Ribosomal RNA small subunit methyltransferase B | elongation zone | 12 h |
| HORVU.MOREX.r3.7HG0709230.1 | 2.604818496 | 0.345142249 | 2.971716099 | 0.00444 | 0.041953879 | -2.05088937 | 50S ribosomal protein L10 | elongation zone | 12 h |
| HORVU.MOREX.r3.7HG0735450.1 | 0.907225051 | 2.173354891 | 3.003222927 | 0.00407 | 0.040229538 | -2.05130169 | Small nuclear ribonucleoprotein | elongation zone | 12 h |
| HORVU.MOREX.r3.1HG0065530.1 | 1.110268532 | 8.837409506 | 3.342393579 | 0.00153 | 0.027394605 | -2.05141552 | Cold shock protein | elongation zone | 12 h |
| HORVU.MOREX.r3.3HG0242830.1 | 2.027614179 | -0.152434766 | 2.978246181 | 0.00436 | 0.041506896 | -2.05190612 | Pentatricopeptide repeat-containing protein | elongation zone | 12 h |
| HORVU.MOREX.r3.3HG0244010.1 | 0.555740157 | 4.561972282 | 3.158088038 | 0.00262 | 0.033716726 | -2.05253456 | Kinase superfamily protein | elongation zone | 12 h |
| HORVU.MOREX.r3.3HG0239970.1 | 1.897354341 | 4.731301793 | 3.062943224 | 0.00344 | 0.03764455 | -2.05335821 | Zinc-finger domain of monoamine-oxidase A repressor R1, putati | elongation zone | 12 h |
| HORVU.MOREX.r3.1HG0092480.1 | 2.220149559 | 4.164362454 | 2.986655001 | 0.00426 | 0.041089957 | -2.05346677 | Glycosyltransferase | elongation zone | 12 h |
| HORVU.MOREX.r3.2HG0146330.1 | 0.958610886 | 8.982026161 | 3.354889066 | 0.00147 | 0.027227922 | -2.05447387 | Ribosomal protein L15 | elongation zone | 12 h |
| HORVU.MOREX.r3.6HG0542950.1 | 1.173904894 | 6.71326239 | 3.23543799 | 0.00209 | 0.030742864 | -2.05755568 | Pentatricopeptide repeat-containing protein | elongation zone | 12 h |
| HORVU.MOREX.r3.3HG0277390.1 | 1.722381585 | 3.530687581 | 3.001043335 | 0.0041 | 0.040265883 | -2.05774532 | Exosome complex component rrp4 | elongation zone | 12 h |
| HORVU.MOREX.r3.7HG0640730.1 | 2.058626598 | -0.0905309 | 2.968851265 | 0.00448 | 0.042098534 | -2.05883045 | Methionine S-methyltransferase | elongation zone | 12 h |
| HORVU.MOREX.r3.7HG0635000.1 | 1.2210249 | 3.997366286 | 3.038322977 | 0.00369 | 0.038671495 | -2.05984442 | Kelch repeat-containing protein | elongation zone | 12 h |
| HORVU.MOREX.r3.3HG0320550.1 | 0.996701232 | 0.545189534 | 2.999711778 | 0.00411 | 0.040307075 | -2.06005288 | DUF241 domain protein, putative (DUF241) | elongation zone | 12 h |
| HORVU.MOREX.r3.5HG0498150.1 | 1.005250588 | 5.564120273 | 3.198362435 | 0.00233 | 0.031954117 | -2.06005942 | 2-keto-3-deoxy-L-rhamnonate aldolase | elongation zone | 12 h |
| HORVU.MOREX.r3.1HG0022820.1 | 1.104556243 | 6.444771319 | 3.218943817 | 0.0022 | 0.031443869 | -2.06147565 | Translocase of chloroplast 159, chloroplastic | elongation zone | 12 h |
| HORVU.MOREX.r3.5HG0505220.1 | 0.797887831 | 2.286205005 | 3.035335582 | 0.00372 | 0.038738812 | -2.06150297 | #N/A | elongation zone | 12 h |
| HORVU.MOREX.r3.1HG0057620.1 | 0.681794992 | 6.017315999 | 3.255507952 | 0.00198 | 0.029636522 | -2.06252929 | Kinase like protein | elongation zone | 12 h |
| HORVU.MOREX.r3.3HG0224450.1 | 0.516114757 | 8.858079699 | 3.377349028 | 0.00138 | 0.026563254 | -2.06263232 | Eukaryotic translation initiation factor 3 subunit A | elongation zone | 12 h |
| HORVU.MOREX.r3.3HG0238180.1 | 0.406473549 | 4.144586965 | 3.197027161 | 0.00234 | 0.031997794 | -2.0627373 | #N/A | elongation zone | 12 h |
| HORVU.MOREX.r3.2HG0183430.1 | 0.601949455 | 5.136892041 | 3.200193466 | 0.00232 | 0.031903923 | -2.06347843 | Mitochondrial carrier protein | elongation zone | 12 h |
| HORVU.MOREX.r3.3HG0288660.1 | 1.852839739 | 4.962352093 | 3.084104735 | 0.00324 | 0.036928455 | -2.06365135 | Dynamin-like | elongation zone | 12 h |
| HORVU.MOREX.r3.1HG0087580.1 | 1.575732398 | 0.956329379 | 2.95804543 | 0.00462 | 0.042747009 | -2.06436736 | Pentatricopeptide repeat-containing protein | elongation zone | 12 h |
| HORVU.MOREX.r3.2HG0137600.1 | 1.409294069 | 0.781391696 | 2.955155396 | 0.00465 | 0.042962871 | -2.06526581 | Peptide chain release factor, putative | elongation zone | 12 h |
| HORVU.MOREX.r3.3HG0219180.1 | -0.647179777 | 2.142648299 | -3.204412616 | 0.00229 | 0.031849386 | -2.06554984 | Adenine nucleotide alpha hydrolases-like superfamily protein | elongation zone | 12 h |
| HORVU.MOREX.r3.4HG0335300.1 | -0.379727831 | 5.365809149 | -3.300525854 | 0.00173 | 0.028397697 | -2.06631493 | Protein ENHANCED DISEASE RESISTANCE 2 | elongation zone | 12 h |

**Table S5** Continued.

| ID | logFC | AveExpr | t | P.Value | adj.P.Val | B | gene description | root zone | time point |
|---|---|---|---|---|---|---|---|---|---|
| HORVU.MOREX.r3.5HG0530720.1 | 0.743843522 | 3.050857477 | 3.066468749 | 0.00341 | 0.037432489 | -2.0680446 | DNA ligase-like protein | elongation zone | 12 h |
| HORVU.MOREX.r3.6HG0616790.1 | 0.540627176 | 7.277734621 | 3.320349328 | 0.00163 | 0.027974391 | -2.07083917 | Transport inhibitor response 1-like protein | elongation zone | 12 h |
| HORVU.MOREX.r3.2HG0195910.1 | -0.66128342 | 2.102421817 | -3.301259451 | 0.00173 | 0.028397697 | -2.07299822 | Protease inhibitor/seed storage/lipid transfer family protein | elongation zone | 12 h |
| HORVU.MOREX.r3.1HG0000630.1 | 1.2833122 | 7.118166762 | 3.232972296 | 0.00211 | 0.030854536 | -2.07435669 | RING-finger E3 ubiquitin ligase, putative | elongation zone | 12 h |
| HORVU.MOREX.r3.6HG0564630.1 | 0.572634907 | 3.887737518 | 3.154484644 | 0.00265 | 0.033840457 | -2.07437337 | 5'-nucleotidase SurE | elongation zone | 12 h |
| HORVU.MOREX.r3.2HG0209800.1 | 0.571004237 | 3.215079802 | 3.086028031 | 0.00322 | 0.036928455 | -2.07441065 | Elongation factor 1-alpha | elongation zone | 12 h |
| HORVU.MOREX.r3.6HG0621690.1 | 0.518847361 | 4.664752545 | 3.182433388 | 0.00244 | 0.032683152 | -2.07445566 | E3 ubiquitin-protein ligase | elongation zone | 12 h |
| HORVU.MOREX.r3.5HG0427090.1 | 2.4155895 | 1.115796108 | 2.957942211 | 0.00462 | 0.042747009 | -2.07498995 | Pentatricopeptide repeat-containing protein | elongation zone | 12 h |
| HORVU.MOREX.r3.2HG0173120.1 | -0.560595997 | 5.602428159 | -3.384147896 | 0.00135 | 0.026498466 | -2.07512168 | Aldose 1-epimerase | elongation zone | 12 h |
| HORVU.MOREX.r3.5HG0503070.1 | 1.088876191 | 2.533422681 | 2.993929089 | 0.00418 | 0.040647009 | -2.07566176 | RNA binding protein | elongation zone | 12 h |
| HORVU.MOREX.r3.4HG0335110.1 | 0.871642321 | 5.132212442 | 3.173984458 | 0.0025 | 0.03299418 | -2.07575256 | BHLH family protein, putative, expressed | elongation zone | 12 h |
| HORVU.MOREX.r3.5HG0488160.1 | 1.549190382 | 2.826332785 | 3.012978426 | 0.00396 | 0.039818139 | -2.07608849 | Dicarboxylate transporter 2.1, chloroplastic | elongation zone | 12 h |
| HORVU.MOREX.r3.2HG0126050.1 | 0.761745574 | 6.555399703 | 3.236654723 | 0.00209 | 0.030699736 | -2.07614722 | 30S ribosomal protein S3 | elongation zone | 12 h |
| HORVU.MOREX.r3.3HG0242050.1 | 1.956369722 | 3.962990486 | 2.988085154 | 0.00425 | 0.04102313 | -2.07824499 | Alpha/beta-Hydrolases superfamily protein | elongation zone | 12 h |
| HORVU.MOREX.r3.5HG0510770.1 | 1.40538738 | 4.300400413 | 3.039894598 | 0.00367 | 0.038641262 | -2.0785931 | Protein-lysine N-methyltransferase | elongation zone | 12 h |
| HORVU.MOREX.r3.2HG0125780.1 | 1.266764621 | 0.405475347 | 2.953621498 | 0.00467 | 0.043107427 | -2.07953629 | #N/A | elongation zone | 12 h |
| HORVU.MOREX.r3.2HG0173230.1 | 1.680893866 | 1.507460003 | 2.949005019 | 0.00473 | 0.043487212 | -2.08047537 | RAN guanine nucleotide release factor | elongation zone | 12 h |
| HORVU.MOREX.r3.7HG0678440.1 | 2.354237576 | 0.782612295 | 2.96795077 | 0.00449 | 0.042132428 | -2.08296386 | Pentatricopeptide repeat-containing protein | elongation zone | 12 h |
| HORVU.MOREX.r3.1HG0067140.1 | -3.375826458 | 1.103422222 | -2.992828985 | 0.00419 | 0.040685115 | -2.08577278 | Wound-induced protease inhibitor | elongation zone | 12 h |
| HORVU.MOREX.r3.2HG0187970.1 | 1.400917779 | 4.77957446 | 3.079855984 | 0.00328 | 0.036983216 | -2.08589262 | Cyclin | elongation zone | 12 h |
| HORVU.MOREX.r3.4HG0392430.1 | 1.345804522 | 2.568155848 | 2.968191393 | 0.00449 | 0.042132428 | -2.08595037 | 4-hydroxy-tetrahydrodipicolinate reductase | elongation zone | 12 h |
| HORVU.MOREX.r3.5HG0515030.1 | 0.561430394 | 4.511137793 | 3.172446241 | 0.00251 | 0.033037412 | -2.0861361 | Obg-like ATPase 1 | elongation zone | 12 h |
| HORVU.MOREX.r3.3HG0284290.1 | 1.146344329 | 6.057281649 | 3.170711377 | 0.00253 | 0.033099534 | -2.08670206 | Mitochondrial processing peptidase | elongation zone | 12 h |
| HORVU.MOREX.r3.7HG0649950.1 | -0.517679198 | 3.836694774 | -3.233833586 | 0.0021 | 0.030821023 | -2.08692029 | Nudix hydrolase | elongation zone | 12 h |
| HORVU.MOREX.r3.6HG0605640.1 | 1.39160131 | 4.881506983 | 3.095158303 | 0.00314 | 0.036539994 | -2.08710103 | Ubiquitin carboxyl-terminal hydrolase-like protein | elongation zone | 12 h |
| HORVU.MOREX.r3.3HG0302570.1 | -2.098302178 | -3.210356093 | -2.962910586 | 0.00456 | 0.04244798 | -2.08839219 | Laccase | elongation zone | 12 h |
| HORVU.MOREX.r3.5HG0493400.1 | 1.619602529 | 1.705240672 | 2.948369398 | 0.00474 | 0.043524966 | -2.08848882 | tRNA-splicing endonuclease subunit SEN54 | elongation zone | 12 h |
| HORVU.MOREX.r3.2HG0183750.1 | 2.013013363 | 1.148914994 | 2.94545981 | 0.00478 | 0.043696105 | -2.08984219 | Transcription factor tfiiic, tau55-like protein, putative | elongation zone | 12 h |
| HORVU.MOREX.r3.2HG0173550.1 | 0.777051278 | 7.816521867 | 3.308571186 | 0.00169 | 0.028279404 | -2.0911123 | Prohibitin | elongation zone | 12 h |
| HORVU.MOREX.r3.5HG0535060.1 | 2.125065002 | 1.687849838 | 2.943098328 | 0.00481 | 0.043773947 | -2.09214979 | Pentatricopeptide repeat-containing protein | elongation zone | 12 h |
| HORVU.MOREX.r3.7HG0711890.1 | -0.915948832 | 2.553034049 | -3.246096645 | 0.00203 | 0.030276719 | -2.09314464 | O-methyltransferase | elongation zone | 12 h |
| HORVU.MOREX.r3.5HG0507920.1 | 0.728159155 | 10.41613396 | 3.424777187 | 0.00119 | 0.02575668 | -2.09400804 | 60S ribosomal protein L5 | elongation zone | 12 h |
| HORVU.MOREX.r3.3HG0234950.1 | 1.901205322 | 2.174772641 | 2.946066662 | 0.00477 | 0.043696105 | -2.0963198 | Multidrug resistance 3 | elongation zone | 12 h |
| HORVU.MOREX.r3.6HG0614110.1 | -0.444665409 | 4.183035474 | -3.308034169 | 0.00169 | 0.028279404 | -2.09692052 | nucleolar GTP-binding protein | elongation zone | 12 h |
| HORVU.MOREX.r3.5HG0528860.1 | 1.957230528 | 5.888709477 | 3.101021285 | 0.00309 | 0.036198029 | -2.09838193 | Pumilio | elongation zone | 12 h |
| HORVU.MOREX.r3.2HG0193700.1 | 2.87064665 | -2.231369651 | 2.972134807 | 0.00444 | 0.041943564 | -2.09862599 | Glucan endo-1,3-beta-glucosidase | elongation zone | 12 h |
| HORVU.MOREX.r3.3HG0291310.1 | 1.873599411 | -0.134112607 | 2.951674515 | 0.0047 | 0.043263203 | -2.09891881 | Acetamidase/formamidase, putative | elongation zone | 12 h |
| HORVU.MOREX.r3.3HG0280080.1 | 0.783587557 | 5.980620424 | 3.217566551 | 0.00221 | 0.031443869 | -2.099079 | 50S ribosomal protein L15, putative, expressed | elongation zone | 12 h |
| HORVU.MOREX.r3.5HG0476620.1 | 1.019012431 | 3.286126116 | 3.024890891 | 0.00383 | 0.03925602 | -2.10042626 | Protein SCO1, mitochondrial-like protein | elongation zone | 12 h |
| HORVU.MOREX.r3.1HG0061750.1 | 0.825644694 | 5.583468809 | 3.18491801 | 0.00243 | 0.032575632 | -2.10136907 | Chlorophyll synthase | elongation zone | 12 h |
| HORVU.MOREX.r3.7HG0745340.1 | 1.081972822 | 7.247395442 | 3.256816628 | 0.00197 | 0.029636522 | -2.10171384 | 30S ribosomal protein S5 | elongation zone | 12 h |

**Table S5** Continued.

| ID | logFC | AveExpr | t | P.Value | adj.P.Val | B | gene description | root zone | time point |
|---|---|---|---|---|---|---|---|---|---|
| HORVU.MOREX.r3.4HG0370780.1 | 0.50022066 | 8.874988886 | 3.375716301 | 0.00138 | 0.026563254 | -2.10365734 | T-complex protein 1 subunit theta | elongation zone | 12 h |
| HORVU.MOREX.r3.5HG0536240.1 | -0.786361529 | 0.742329041 | -3.092112109 | 0.00317 | 0.036688668 | -2.10376384 | Beta-fructofuranosidase, insoluble protein | elongation zone | 12 h |
| HORVU.MOREX.r3.1HG0066370.1 | 2.367068988 | 5.852246704 | 3.08537556 | 0.00323 | 0.036928455 | -2.10380544 | Glutathione S-transferase | elongation zone | 12 h |
| HORVU.MOREX.r3.1HG0006230.1 | 0.417570967 | 5.574000008 | 3.254262705 | 0.00198 | 0.029691287 | -2.10436104 | Disease resistance protein RPM1 | elongation zone | 12 h |
| HORVU.MOREX.r3.1HG0065370.1 | 0.670128057 | 5.483156066 | 3.21683202 | 0.00221 | 0.031443869 | -2.10553307 | TPX2 (Targeting protein for Xklp2) family protein | elongation zone | 12 h |
| HORVU.MOREX.r3.2HG0109530.1 | -0.285313523 | 6.710852914 | -3.364647523 | 0.00143 | 0.026979742 | -2.10644992 | Protein phosphatase 2C-like | elongation zone | 12 h |
| HORVU.MOREX.r3.7HG0751130.1 | 2.277606029 | -0.911602578 | 2.964584255 | 0.00453 | 0.042404918 | -2.10692377 | Ubiquitin carboxyl-terminal hydrolase 38 | elongation zone | 12 h |
| HORVU.MOREX.r3.2HG0160550.1 | 2.144311073 | 2.252307779 | 2.937849142 | 0.00488 | 0.044199271 | -2.10793266 | At4g33800-like protein | elongation zone | 12 h |
| HORVU.MOREX.r3.2HG0196710.1 | 1.27981585 | 4.927717077 | 3.080615516 | 0.00327 | 0.036983216 | -2.10906448 | Ribosome biogenesis protein BRX1-like protein | elongation zone | 12 h |
| HORVU.MOREX.r3.6HG0557080.1 | 1.573002231 | 6.140814891 | 3.134809894 | 0.0028 | 0.034437001 | -2.10921426 | Protein arginine N-methyltransferase | elongation zone | 12 h |
| HORVU.MOREX.r3.1HG0058530.1 | 1.055116627 | 2.661369894 | 3.024525409 | 0.00383 | 0.03925602 | -2.10939542 | Small nuclear ribonucleoprotein family protein | elongation zone | 12 h |
| HORVU.MOREX.r3.7HG0667140.1 | 1.026603044 | 6.545087438 | 3.208016318 | 0.00227 | 0.031692167 | -2.11027334 | #N/A | elongation zone | 12 h |
| HORVU.MOREX.r3.6HG0573070.1 | 1.397165107 | 3.004821099 | 2.96469277 | 0.00453 | 0.042404918 | -2.11053751 | MLO-like protein | elongation zone | 12 h |
| HORVU.MOREX.r3.6HG0619410.1 | -0.769958523 | 1.820333919 | -3.180839126 | 0.00245 | 0.032706849 | -2.11075791 | Flavonol synthase | elongation zone | 12 h |
| HORVU.MOREX.r3.1HG0080750.1 | 1.792758606 | 5.902530715 | 3.104485558 | 0.00306 | 0.036052722 | -2.1108203 | Phosphoribosylformylglycinamidine synthase family protein | elongation zone | 12 h |
| HORVU.MOREX.r3.7HG0706580.1 | 0.873462395 | 4.99575294 | 3.148099671 | 0.0027 | 0.033864708 | -2.11234568 | WD40 repeat-containing protein | elongation zone | 12 h |
| HORVU.MOREX.r3.7HG0711410.1 | 1.143605932 | 5.613777061 | 3.168368652 | 0.00254 | 0.033171389 | -2.11240771 | 60S ribosomal protein L37a | elongation zone | 12 h |
| HORVU.MOREX.r3.6HG0609720.1 | 0.618286162 | 3.853973881 | 3.123278535 | 0.0029 | 0.035162818 | -2.11280976 | Glucosamine 6-phosphate N-acetyltransferase | elongation zone | 12 h |
| HORVU.MOREX.r3.5HG0466320.1 | -0.310574002 | 6.324949057 | -3.313122906 | 0.00167 | 0.028155399 | -2.11289487 | Phosphoinositide phospholipase C | elongation zone | 12 h |
| HORVU.MOREX.r3.2HG0123890.1 | 1.0627663 | 8.179504177 | 3.287260006 | 0.0018 | 0.028691351 | -2.11293394 | 40S ribosomal protein S6 | elongation zone | 12 h |
| HORVU.MOREX.r3.1HG0086390.1 | -0.807705644 | -0.260888993 | -3.156063515 | 0.00264 | 0.033771061 | -2.11316877 | #N/A | elongation zone | 12 h |
| HORVU.MOREX.r3.6HG0596310.1 | 0.817704169 | 8.298151592 | 3.318695482 | 0.00164 | 0.028033028 | -2.11565295 | 60S ribosomal protein L14, putative | elongation zone | 12 h |
| HORVU.MOREX.r3.4HG0412170.1 | 0.8059807 | 9.2702411 | 3.362113754 | 0.00144 | 0.02703001 | -2.11584083 | 60S ribosomal protein L21 | elongation zone | 12 h |
| HORVU.MOREX.r3.7HG0731150.1 | 1.859218618 | 4.133250031 | 2.981254578 | 0.00433 | 0.041308987 | -2.11632466 | F-box and JmjC domain-containing protein | elongation zone | 12 h |
| HORVU.MOREX.r3.6HG0597130.1 | 0.916440949 | 4.149561941 | 3.069446369 | 0.00338 | 0.037331432 | -2.11634002 | Replication protein A subunit | elongation zone | 12 h |
| HORVU.MOREX.r3.1HG0084680.1 | 1.940615311 | 6.696342312 | 3.150358559 | 0.00268 | 0.033864708 | -2.11638309 | Nucleolar family protein | elongation zone | 12 h |
| HORVU.MOREX.r3.3HG0312220.1 | 1.024509771 | 4.580442378 | 3.073559231 | 0.00334 | 0.037245743 | -2.11818743 | Histone-lysine N-methyltransferase, H3 lysine-9 specific | elongation zone | 12 h |
| HORVU.MOREX.r3.5HG0427060.1 | 0.966933541 | 5.128578703 | 3.153766925 | 0.00265 | 0.033864708 | -2.11880283 | 50S ribosomal protein L14 | elongation zone | 12 h |
| HORVU.MOREX.r3.3HG0298340.1 | 0.777507118 | 4.243388819 | 3.102972293 | 0.00307 | 0.036128346 | -2.11885751 | Beta-galactosidase | elongation zone | 12 h |
| HORVU.MOREX.r3.3HG0237590.1 | 1.447131086 | 4.756362292 | 3.052364293 | 0.00354 | 0.037882054 | -2.11898042 | ATP-dependent RNA helicase | elongation zone | 12 h |
| HORVU.MOREX.r3.2HG0178630.1 | 2.350094012 | 3.452461861 | 2.945548447 | 0.00478 | 0.043696105 | -2.11921848 | DNA mismatch repair protein mutS | elongation zone | 12 h |
| HORVU.MOREX.r3.7HG0704390.1 | 0.594795308 | 7.743233053 | 3.308158105 | 0.00169 | 0.028279404 | -2.1199122 | 40S ribosomal protein S13 | elongation zone | 12 h |
| HORVU.MOREX.r3.7HG0682130.1 | 0.797678299 | 5.527659552 | 3.170004569 | 0.00253 | 0.033099534 | -2.12005611 | #N/A | elongation zone | 12 h |
| HORVU.MOREX.r3.6HG0542970.1 | 0.827944026 | 4.467056131 | 3.147006181 | 0.00271 | 0.033864708 | -2.12159214 | Receptor-like kinase | elongation zone | 12 h |
| HORVU.MOREX.r3.6HG0569630.1 | 1.535856622 | 4.808423924 | 3.062603759 | 0.00344 | 0.03764455 | -2.12233335 | DNA-directed RNA polymerase subunit beta | elongation zone | 12 h |
| HORVU.MOREX.r3.6HG0625860.1 | 0.679208124 | 4.74429219 | 3.123739798 | 0.00289 | 0.035147457 | -2.12243374 | CobW-domain-containing protein | elongation zone | 12 h |
| HORVU.MOREX.r3.3HG0315120.1 | 2.282648031 | 0.551320449 | 2.945046229 | 0.00479 | 0.043697279 | -2.12329952 | Ribonuclease P/MRP protein subunit POP5 | elongation zone | 12 h |
| HORVU.MOREX.r3.1HG0090330.1 | 1.183159684 | 3.818381072 | 3.014154662 | 0.00395 | 0.039746874 | -2.12553915 | Aldose 1-epimerase-like | elongation zone | 12 h |
| HORVU.MOREX.r3.3HG0305790.1 | 0.867390699 | 6.433470179 | 3.224747085 | 0.00216 | 0.031159303 | -2.12622072 | Pentatricopeptide repeat protein | elongation zone | 12 h |
| HORVU.MOREX.r3.5HG0461830.1 | 1.051449648 | 1.937434209 | 2.956342005 | 0.00464 | 0.042860101 | -2.12629019 | F-box/kelch-repeat protein | elongation zone | 12 h |
| HORVU.MOREX.r3.3HG0273130.1 | 0.682430886 | 5.87661788 | 3.200673833 | 0.00232 | 0.031894634 | -2.13005362 | Extra-large G-protein-like | elongation zone | 12 h |

**Table S5** Continued.

| ID | logFC | AveExpr | t | P.Value | adj.P.Val | B | gene description | root zone | time point |
|---|---|---|---|---|---|---|---|---|---|
| HORVU.MOREX.r3.4HG0344830.1 | 0.442692961 | 4.535567699 | 3.203118567 | 0.0023 | 0.031887924 | -2.13064922 | NBS-LRR disease resistance protein, putative, expressed | elongation zone | 12 h |
| HORVU.MOREX.r3.7HG0726360.1 | -0.295771045 | 5.450154375 | -3.273026022 | 0.00188 | 0.028923495 | -2.13239046 | 12-oxophytodienoate reductase | elongation zone | 12 h |
| HORVU.MOREX.r3.6HG0619290.1 | 1.329604511 | 3.250452134 | 3.028668412 | 0.00379 | 0.039108923 | -2.13348769 | Glutaredoxin-like protein | elongation zone | 12 h |
| HORVU.MOREX.r3.6HG0600220.1 | 0.904161051 | 5.927792261 | 3.173343165 | 0.00251 | 0.03299418 | -2.13378324 | 3-isopropylmalate dehydratase small subunit | elongation zone | 12 h |
| HORVU.MOREX.r3.3HG0236160.1 | 0.612581313 | 4.863001087 | 3.147396792 | 0.0027 | 0.033864708 | -2.13476289 | Transcription factor, putative | elongation zone | 12 h |
| HORVU.MOREX.r3.3HG0246560.1 | 0.609718985 | 5.025368236 | 3.143253799 | 0.00274 | 0.034060761 | -2.1348304 | Multiprotein-bridging factor, putative | elongation zone | 12 h |
| HORVU.MOREX.r3.5HG0527340.1 | 2.016785111 | -0.841855617 | 2.946576962 | 0.00477 | 0.04368319 | -2.13538302 | DNA-directed RNA polymerase subunit | elongation zone | 12 h |
| HORVU.MOREX.r3.3HG0297400.1 | 2.575622397 | 0.76599868 | 2.928531968 | 0.00501 | 0.044779816 | -2.13634401 | Pentatricopeptide repeat-containing protein | elongation zone | 12 h |
| HORVU.MOREX.r3.1HG0082750.1 | 1.673796478 | 0.305019362 | 2.923020941 | 0.00509 | 0.045226145 | -2.13693549 | Ring finger protein, putative | elongation zone | 12 h |
| HORVU.MOREX.r3.5HG0517610.1 | 1.469173376 | 4.899893762 | 3.054411711 | 0.00352 | 0.037808566 | -2.13706699 | #N/A | elongation zone | 12 h |
| HORVU.MOREX.r3.7HG0646060.1 | 1.63321774 | 4.424298678 | 2.987723049 | 0.00425 | 0.04102761 | -2.1394188 | Receptor-like protein kinase | elongation zone | 12 h |
| HORVU.MOREX.r3.2HG0119870.1 | 0.912473858 | 2.245991143 | 2.960905522 | 0.00458 | 0.042615343 | -2.13970226 | Methyltransferase family protein, putative | elongation zone | 12 h |
| HORVU.MOREX.r3.7HG0661890.1 | 0.497848527 | 7.9160487 | 3.350392863 | 0.00149 | 0.027239638 | -2.14103121 | 3-oxoacyl-[acyl-carrier-protein] synthase | elongation zone | 12 h |
| HORVU.MOREX.r3.5HG0424490.1 | 2.367041664 | -0.073596929 | 2.957066606 | 0.00463 | 0.042812485 | -2.14380923 | Glycosyltransferase | elongation zone | 12 h |
| HORVU.MOREX.r3.2HG0196140.1 | 0.356427435 | 5.377901779 | 3.235664006 | 0.00209 | 0.030742864 | -2.14556568 | Ras-related protein Rab-8B | elongation zone | 12 h |
| HORVU.MOREX.r3.6HG0608180.1 | -0.891621293 | 0.307405102 | -3.062609238 | 0.00344 | 0.03764455 | -2.14778296 | Protein MIZU-KUSSEI 1 | elongation zone | 12 h |
| HORVU.MOREX.r3.2HG0194310.1 | 1.612816839 | 7.045455879 | 3.188111663 | 0.0024 | 0.032414492 | -2.14837676 | Midasin | elongation zone | 12 h |
| HORVU.MOREX.r3.7HG0714910.1 | 0.584632921 | 7.216849728 | 3.305730749 | 0.0017 | 0.028288147 | -2.15082905 | #N/A | elongation zone | 12 h |
| HORVU.MOREX.r3.3HG0305290.1 | 1.807124267 | 1.243753956 | 2.916972146 | 0.00517 | 0.045490951 | -2.15083308 | Pentatricopeptide repeat-containing protein | elongation zone | 12 h |
| HORVU.MOREX.r3.4HG0380980.1 | 1.021205617 | 4.039743688 | 3.04193472 | 0.00365 | 0.0386017 | -2.1522853 | 30S ribosomal protein S1 | elongation zone | 12 h |
| HORVU.MOREX.r3.7HG0720790.1 | 0.900041605 | 3.444490231 | 3.018714453 | 0.0039 | 0.039510918 | -2.15407406 | Myb family transcription factor family protein | elongation zone | 12 h |
| HORVU.MOREX.r3.3HG0310640.1 | -0.626855199 | 2.038725307 | -3.258902654 | 0.00196 | 0.029571063 | -2.15513528 | Sulfotransferase | elongation zone | 12 h |
| HORVU.MOREX.r3.6HG0615360.1 | 1.519973951 | 2.604486893 | 2.966860685 | 0.00451 | 0.042233448 | -2.15550124 | Geranylgeranyl reductase | elongation zone | 12 h |
| HORVU.MOREX.r3.5HG0434210.1 | 1.674218279 | 2.431638709 | 2.923173144 | 0.00509 | 0.045226145 | -2.15561596 | #N/A | elongation zone | 12 h |
| HORVU.MOREX.r3.5HG0513380.1 | 1.512039694 | 2.861847727 | 2.936288963 | 0.0049 | 0.044321273 | -2.15569425 | RNA-binding protein-like | elongation zone | 12 h |
| HORVU.MOREX.r3.6HG0618690.1 | 1.526832417 | 4.702412058 | 3.011743659 | 0.00397 | 0.03986915 | -2.15718028 | DNA-binding storekeeper protein-related transcriptional regulat | elongation zone | 12 h |
| HORVU.MOREX.r3.3HG0246190.1 | 1.057414556 | 1.250282278 | 3.039299085 | 0.00368 | 0.038641262 | -2.15838347 | Cationic amino acid transporter | elongation zone | 12 h |
| HORVU.MOREX.r3.5HG0536500.1 | 0.745089958 | 5.276862926 | 3.114654074 | 0.00297 | 0.035453096 | -2.15852348 | ATP-dependent DNA helicase 2 subunit KU80 | elongation zone | 12 h |
| HORVU.MOREX.r3.6HG0564690.1 | -0.405320666 | 5.759195311 | -3.315215224 | 0.00166 | 0.028097383 | -2.15886757 | Regulator of Vps4 activity in the MVB pathway protein, putative | elongation zone | 12 h |
| HORVU.MOREX.r3.7HG0638370.3 | -0.451616422 | 6.33021084 | -3.348790009 | 0.0015 | 0.027239638 | -2.16348578 | Membrane protein of er body-like protein | elongation zone | 12 h |
| HORVU.MOREX.r3.4HG0335680.1 | -0.634198095 | 1.198730266 | -3.041486154 | 0.00366 | 0.0386017 | -2.16349677 | Protein yippee-like | elongation zone | 12 h |
| HORVU.MOREX.r3.5HG0507610.1 | 1.205052735 | 4.190859194 | 3.006622855 | 0.00403 | 0.040178373 | -2.16372452 | Elongation factor G | elongation zone | 12 h |
| HORVU.MOREX.r3.6HG0556130.1 | 1.580081363 | 0.544616443 | 2.913382245 | 0.00522 | 0.045731286 | -2.16450088 | cytomegalovirus UL139 protein | elongation zone | 12 h |
| HORVU.MOREX.r3.5HG0430050.1 | -0.560300028 | 4.495639375 | -3.200782633 | 0.00232 | 0.031894634 | -2.16451603 | Stomatal closure-related actin-binding protein 1 | elongation zone | 12 h |
| HORVU.MOREX.r3.7HG0745890.1 | 0.904661915 | 4.093881544 | 3.049316907 | 0.00358 | 0.038090743 | -2.1647783 | Elongator complex protein 4 | elongation zone | 12 h |
| HORVU.MOREX.r3.3HG0315210.1 | 2.199490287 | -0.621766802 | 2.922466226 | 0.0051 | 0.045240439 | -2.16628563 | F-box family protein | elongation zone | 12 h |
| HORVU.MOREX.r3.4HG0388770.1 | 0.83802273 | 6.564233193 | 3.200775947 | 0.00232 | 0.031894634 | -2.16639778 | Protein TOC75-3, chloroplastic | elongation zone | 12 h |
| HORVU.MOREX.r3.4HG0411400.1 | 1.440583033 | 4.951130643 | 3.071592668 | 0.00336 | 0.037271777 | -2.16691807 | eisosome protein | elongation zone | 12 h |
| HORVU.MOREX.r3.2HG0105680.3 | 0.972635593 | 4.721414669 | 3.102221595 | 0.00308 | 0.036136245 | -2.16778046 | Translation initiation factor IF-2 | elongation zone | 12 h |
| HORVU.MOREX.r3.2HG0165110.1 | -0.821019484 | 3.128473967 | -3.159909515 | 0.00261 | 0.033615353 | -2.16827757 | Acetyl-coenzyme A synthetase | elongation zone | 12 h |
| HORVU.MOREX.r3.1HG0064870.1 | 2.02698732 | 0.288676864 | 2.91378545 | 0.00522 | 0.045722475 | -2.16865214 | Pentatricopeptide repeat-containing protein | elongation zone | 12 h |

**Table S5** Continued.

| ID | logFC | AveExpr | t | P.Value | adj.P.Val | B | gene description | root zone | time point |
|---|---|---|---|---|---|---|---|---|---|
| HORVU.MOREX.r3.3HG0249790.1 | 0.683757417 | 6.336137063 | 3.216880531 | 0.00221 | 0.031443869 | -2.16937246 | Exportin-2 | elongation zone | 12 h |
| HORVU.MOREX.r3.3HG0309340.1 | 2.721059844 | 0.362849458 | 2.91610103 | 0.00519 | 0.045568238 | -2.17161152 | Pentatricopeptide repeat-containing protein | elongation zone | 12 h |
| HORVU.MOREX.r3.1HG0018810.1 | 0.740788894 | 5.360038976 | 3.14686191 | 0.00271 | 0.033864708 | -2.17246015 | #N/A | elongation zone | 12 h |
| HORVU.MOREX.r3.3HG0292680.1 | -0.696467591 | 2.459132616 | -3.214251791 | 0.00223 | 0.031484433 | -2.17248382 | Replicase polyprotein 1a | elongation zone | 12 h |
| HORVU.MOREX.r3.4HG0345340.1 | -0.33029701 | 5.038124181 | -3.263321909 | 0.00193 | 0.029341049 | -2.17273643 | Basic-leucine zipper (bZIP) transcription factor family protein | elongation zone | 12 h |
| HORVU.MOREX.r3.5HG0502360.1 | 1.329209779 | 1.108342302 | 2.919788063 | 0.00513 | 0.045371176 | -2.17322014 | DNA-directed RNA polymerase subunit alpha | elongation zone | 12 h |
| HORVU.MOREX.r3.6HG0551750.1 | 0.83374449 | 4.011205702 | 3.054342511 | 0.00352 | 0.037808566 | -2.17359503 | General transcription factor IIH subunit, putative | elongation zone | 12 h |
| HORVU.MOREX.r3.2HG0210420.1 | 0.445677043 | 7.910636809 | 3.32748501 | 0.0016 | 0.027806246 | -2.17488265 | Dolichyl-diphosphooligosaccharide--protein glycosyltransferase s | elongation zone | 12 h |
| HORVU.MOREX.r3.4HG0333450.1 | -0.515700817 | 3.618972788 | -3.150232673 | 0.00268 | 0.033864708 | -2.17807599 | Type I inositol-1,4,5-trisphosphate 5-phosphatase CVP2 | elongation zone | 12 h |
| HORVU.MOREX.r3.3HG0299370.1 | 1.398384676 | 6.44411695 | 3.166804894 | 0.00256 | 0.033218309 | -2.1795015 | DNA polymerase | elongation zone | 12 h |
| HORVU.MOREX.r3.5HG0435800.1 | 1.830471823 | 0.267695951 | 2.904601238 | 0.00535 | 0.046230732 | -2.17979449 | Leucine-rich repeat protein kinase family protein | elongation zone | 12 h |
| HORVU.MOREX.r3.5HG0512540.1 | -0.269491756 | 5.508031516 | -3.274282224 | 0.00187 | 0.028923495 | -2.18196739 | Prolyl 4-hydroxylase alpha subunit, putative | elongation zone | 12 h |
| HORVU.MOREX.r3.2HG0207000.1 | 0.811976897 | 5.903816857 | 3.147614848 | 0.0027 | 0.033864708 | -2.18249579 | RNA binding protein | elongation zone | 12 h |
| HORVU.MOREX.r3.1HG0003620.1 | -0.619013368 | 2.573512603 | -3.133236654 | 0.00282 | 0.034551409 | -2.18289139 | Protein ZINC INDUCED FACILITATOR-LIKE 1 | elongation zone | 12 h |
| HORVU.MOREX.r3.1HG0061850.1 | 0.69606166 | 8.584058175 | 3.311638496 | 0.00167 | 0.028232736 | -2.18299239 | Methionine aminopeptidase 2 | elongation zone | 12 h |
| HORVU.MOREX.r3.6HG0597360.1 | 0.492954474 | 5.075212238 | 3.162518401 | 0.00259 | 0.033483118 | -2.18305298 | SET domain-containing protein 9 | elongation zone | 12 h |
| HORVU.MOREX.r3.1HG0063790.1 | 1.427912575 | 3.096652959 | 2.94182019 | 0.00483 | 0.043890279 | -2.18338372 | Sister chromatid cohesion 1 protein 1 | elongation zone | 12 h |
| HORVU.MOREX.r3.2HG0174320.1 | 2.885712424 | 4.916755342 | 2.96299401 | 0.00455 | 0.04244798 | -2.18548216 | Kinesin-like protein | elongation zone | 12 h |
| HORVU.MOREX.r3.3HG0280610.1 | 0.546198489 | 6.601541701 | 3.231693631 | 0.00212 | 0.030866089 | -2.1855176 | Splicing factor 3A subunit 3 | elongation zone | 12 h |
| HORVU.MOREX.r3.7HG0744540.1 | 2.112785902 | -0.407648584 | 2.905691325 | 0.00534 | 0.04618095 | -2.18782606 | Glycosyltransferase | elongation zone | 12 h |
| HORVU.MOREX.r3.1HG0093060.1 | -0.535815937 | 4.965521065 | -3.297529886 | 0.00175 | 0.02840648 | -2.19034364 | Extra-large G-protein-like | elongation zone | 12 h |
| HORVU.MOREX.r3.5HG0454920.1 | 0.645586991 | 3.842449 | 3.068444669 | 0.00339 | 0.037343384 | -2.19077419 | RNA polymerase II subunit A C-terminal domain phosphatase SSU | elongation zone | 12 h |
| HORVU.MOREX.r3.6HG0618170.1 | 0.27450393 | 6.062292159 | 3.242440988 | 0.00205 | 0.030425122 | -2.19158352 | 1-aminocyclopropane-1-carboxylate deaminase/D-cysteine desu | elongation zone | 12 h |
| HORVU.MOREX.r3.5HG0444920.1 | 0.656449317 | 5.104322172 | 3.132321445 | 0.00282 | 0.034584718 | -2.19227377 | Uveal autoantigen with coiled-coil domains and ankyrin repeats i | elongation zone | 12 h |
| HORVU.MOREX.r3.4HG0341950.1 | 1.449947115 | 5.083003475 | 3.037215193 | 0.0037 | 0.038671495 | -2.19238128 | Exosome complex component | elongation zone | 12 h |
| HORVU.MOREX.r3.2HG0150290.1 | 0.637520356 | 8.717704517 | 3.320792146 | 0.00163 | 0.02796111 | -2.19280283 | 30S ribosomal protein S19 | elongation zone | 12 h |
| HORVU.MOREX.r3.7HG0725460.1 | 1.715981507 | 1.80349634 | 2.905818448 | 0.00533 | 0.04618095 | -2.19300455 | RNA-binding (RRM/RBD/RNP motifs) family protein | elongation zone | 12 h |
| HORVU.MOREX.r3.1HG0025500.2 | 0.750009134 | 5.09562282 | 3.148429067 | 0.0027 | 0.033864708 | -2.19433092 | Arginine--tRNA ligase | elongation zone | 12 h |
| HORVU.MOREX.r3.3HG0223960.1 | 1.198448361 | 2.937552338 | 2.927182672 | 0.00503 | 0.044862884 | -2.19467382 | #N/A | elongation zone | 12 h |
| HORVU.MOREX.r3.6HG0628790.1 | -0.708927991 | 2.686977072 | -3.069454898 | 0.00338 | 0.037331432 | -2.19496869 | Myb-like transcription factor family protein | elongation zone | 12 h |
| HORVU.MOREX.r3.3HG0290220.1 | 2.781249547 | 4.744100239 | 2.960516311 | 0.00459 | 0.042615343 | -2.19561629 | Nucleobase-ascorbate transporter-like protein | elongation zone | 12 h |
| HORVU.MOREX.r3.6HG0572880.1 | 0.677216355 | 6.302994109 | 3.183336355 | 0.00244 | 0.03261914 | -2.1975852 | Alpha/beta-Hydrolases superfamily protein | elongation zone | 12 h |
| HORVU.MOREX.r3.3HG0309000.1 | 0.783842918 | 3.431372321 | 3.000264042 | 0.0041 | 0.040307075 | -2.1983204 | snRNA-activating protein complex subunit | elongation zone | 12 h |
| HORVU.MOREX.r3.3HG0310880.1 | 1.973317232 | 0.788604478 | 2.895173314 | 0.00549 | 0.046765582 | -2.19897793 | #N/A | elongation zone | 12 h |
| HORVU.MOREX.r3.3HG0295850.1 | 1.609314137 | 4.42296407 | 2.993154332 | 0.00419 | 0.040685115 | -2.19986333 | U3 small nucleolar RNA-associated protein 11 | elongation zone | 12 h |
| HORVU.MOREX.r3.3HG0226750.1 | -0.53703631 | 2.949436288 | -3.300442799 | 0.00173 | 0.028397697 | -2.200668 | GDSL esterase/lipase | elongation zone | 12 h |
| HORVU.MOREX.r3.1HG0051810.1 | 2.25608808 | 3.342547612 | 2.911151714 | 0.00526 | 0.045859795 | -2.20185807 | Glutathione S-transferase | elongation zone | 12 h |
| HORVU.MOREX.r3.3HG0265190.1 | 0.540318501 | 6.392012157 | 3.21343482 | 0.00223 | 0.031515925 | -2.20253125 | Exonuclease 3'-5' domain-containing protein 1 | elongation zone | 12 h |
| HORVU.MOREX.r3.7HG0712610.1 | 0.565503236 | 8.251441205 | 3.313554426 | 0.00166 | 0.028155399 | -2.20359163 | Elongation factor G | elongation zone | 12 h |
| HORVU.MOREX.r3.3HG0252240.1 | 0.861572603 | 7.225249207 | 3.244340405 | 0.00204 | 0.030344376 | -2.20516122 | Histone H2B | elongation zone | 12 h |
| HORVU.MOREX.r3.2HG0145780.1 | 2.087711078 | -0.995256061 | 2.904670482 | 0.00535 | 0.046230732 | -2.20531261 | Cinnamoyl-CoA reductase family protein | elongation zone | 12 h |

**Table S5** Continued.

| ID | logFC | AveExpr | t | P.Value | adj.P.Val | B | gene description | root zone | time point |
|---|---|---|---|---|---|---|---|---|---|
| HORVU.MOREX.r3.3HG0246500.1 | -0.768614772 | 3.040521827 | -3.289217346 | 0.00179 | 0.028656416 | -2.20657319 | Peptide-N4-(N-acetyl-beta-glucosaminyl)asparagine amidase A | elongation zone | 12 h |
| HORVU.MOREX.r3.2HG0180060.1 | 1.229438208 | 1.155123868 | 2.896355057 | 0.00547 | 0.046711054 | -2.20764981 | Receptor-kinase, putative | elongation zone | 12 h |
| HORVU.MOREX.r3.6HG0594100.1 | 2.520825687 | 2.846373641 | 2.894101319 | 0.00551 | 0.046806243 | -2.20830044 | H/ACA ribonucleoprotein complex non-core subunit NAF1 | elongation zone | 12 h |
| HORVU.MOREX.r3.7HG0648190.1 | 1.439825943 | 4.830917175 | 3.001725185 | 0.00409 | 0.040265883 | -2.20849887 | F-box family protein | elongation zone | 12 h |
| HORVU.MOREX.r3.1HG0083320.1 | 1.485998598 | 6.710998818 | 3.145380041 | 0.00272 | 0.033887701 | -2.20854348 | Threonine synthase | elongation zone | 12 h |
| HORVU.MOREX.r3.2HG0178560.2 | 1.749762628 | 3.749491986 | 2.957957347 | 0.00462 | 0.042747009 | -2.2090637 | LA RNA-binding protein | elongation zone | 12 h |
| HORVU.MOREX.r3.3HG0312020.1 | -0.394709051 | 6.244935241 | -3.348312616 | 0.0015 | 0.027239638 | -2.21090975 | glycosyltransferase family exostosin protein | elongation zone | 12 h |
| HORVU.MOREX.r3.3HG0286070.1 | 1.844419595 | 0.985153469 | 2.889141253 | 0.00558 | 0.047136585 | -2.2118841 | Pleiotropic drug resistance ABC transporter | elongation zone | 12 h |
| HORVU.MOREX.r3.3HG0316860.1 | 1.660544183 | 0.949417462 | 2.889128983 | 0.00558 | 0.047136585 | -2.21215991 | Pentatricopeptide repeat-containing protein | elongation zone | 12 h |
| HORVU.MOREX.r3.4HG0339290.1 | 1.574426636 | 5.458049088 | 3.055686077 | 0.00351 | 0.037808566 | -2.21248968 | ATP dependent RNA helicase | elongation zone | 12 h |
| HORVU.MOREX.r3.1HG0043060.1 | 1.800835406 | 1.58085499 | 2.888225006 | 0.0056 | 0.047175894 | -2.21426803 | Pentatricopeptide repeat-containing protein | elongation zone | 12 h |
| HORVU.MOREX.r3.1HG0058050.1 | 0.572747785 | 5.246817852 | 3.168404492 | 0.00254 | 0.033171389 | -2.21459844 | 8-amino-7-oxononanoate synthase | elongation zone | 12 h |
| HORVU.MOREX.r3.6HG0590960.1 | 1.94980183 | 0.923831685 | 2.887815887 | 0.0056 | 0.047190057 | -2.21605617 | NAC domain-containing protein | elongation zone | 12 h |
| HORVU.MOREX.r3.3HG0258630.1 | 1.5076939 | 6.575402356 | 3.130372957 | 0.00284 | 0.034669947 | -2.21710878 | Pentatricopeptide repeat-containing protein | elongation zone | 12 h |
| HORVU.MOREX.r3.2HG0134300.1 | 1.900723637 | 1.307766187 | 2.886663147 | 0.00562 | 0.04722308 | -2.21724321 | Skp1, putative | elongation zone | 12 h |
| HORVU.MOREX.r3.7HG0729090.1 | 1.149909236 | 1.917852249 | 2.898000902 | 0.00545 | 0.046636203 | -2.21777534 | Pentatricopeptide repeat-containing protein | elongation zone | 12 h |
| HORVU.MOREX.r3.1HG0015750.1 | -1.44040578 | -2.912149854 | -2.933371328 | 0.00494 | 0.044514543 | -2.21848572 | transcription factor, putative (Protein of unknown function, DUF | elongation zone | 12 h |
| HORVU.MOREX.r3.5HG0522770.1 | 0.824701905 | 9.471285898 | 3.331713308 | 0.00158 | 0.02769762 | -2.21911844 | 30S ribosomal protein S5 | elongation zone | 12 h |
| HORVU.MOREX.r3.2HG0193220.1 | 0.660836492 | 4.020109938 | 3.056042948 | 0.00351 | 0.037808566 | -2.21985127 | Putrescine-binding periplasmic protein | elongation zone | 12 h |
| HORVU.MOREX.r3.4HG0383780.1 | -1.735090175 | -0.062687302 | -3.08900952 | 0.00319 | 0.036827029 | -2.22040415 | Laccase | elongation zone | 12 h |
| HORVU.MOREX.r3.3HG0302050.1 | 2.189760715 | 0.747970046 | 2.887942114 | 0.0056 | 0.047190057 | -2.22124049 | Pentatricopeptide repeat-containing protein | elongation zone | 12 h |
| HORVU.MOREX.r3.1HG0084890.1 | -0.635883785 | 1.342406063 | -3.107115368 | 0.00303 | 0.035902171 | -2.22213536 | Protein DETOXIFICATION | elongation zone | 12 h |
| HORVU.MOREX.r3.3HG0243220.1 | 2.36333043 | 1.638276816 | 2.883411648 | 0.00567 | 0.047427345 | -2.22648321 | Pentatricopeptide repeat-containing protein | elongation zone | 12 h |
| HORVU.MOREX.r3.5HG0525230.1 | 0.872093551 | 2.592012409 | 2.926758719 | 0.00504 | 0.044862884 | -2.22656172 | Glucose-1-phosphate adenylyltransferase | elongation zone | 12 h |
| HORVU.MOREX.r3.5HG0504160.1 | 1.073974377 | 6.884631382 | 3.209923508 | 0.00226 | 0.031600255 | -2.22695957 | #N/A | elongation zone | 12 h |
| HORVU.MOREX.r3.6HG0550430.1 | 0.853436989 | 5.615355597 | 3.132092647 | 0.00282 | 0.034584718 | -2.22714949 | 50S ribosomal protein L4 | elongation zone | 12 h |
| HORVU.MOREX.r3.3HG0224510.1 | -3.116507893 | -2.729166608 | -2.905391556 | 0.00534 | 0.046199549 | -2.22791688 | O-methyltransferase | elongation zone | 12 h |
| HORVU.MOREX.r3.4HG0401530.2 | 0.897375312 | 3.547060924 | 3.021714551 | 0.00386 | 0.039354132 | -2.22815709 | evolutionarily conserved C-terminal region 2 | elongation zone | 12 h |
| HORVU.MOREX.r3.5HG0472880.1 | 0.488383975 | 4.87983831 | 3.151759747 | 0.00267 | 0.033864708 | -2.2292235 | GDP-mannose transporter, putative | elongation zone | 12 h |
| HORVU.MOREX.r3.4HG0333040.1 | -0.583305297 | 3.140672069 | -3.196041873 | 0.00235 | 0.032009236 | -2.23012051 | Ankyrin repeat-containing protein | elongation zone | 12 h |
| HORVU.MOREX.r3.1HG0075590.1 | 0.255489545 | 8.23086424 | 3.348470826 | 0.0015 | 0.027239638 | -2.23112837 | Coiled-coil domain-containing protein 47 | elongation zone | 12 h |
| HORVU.MOREX.r3.7HG0689220.1 | 0.809500216 | 8.338861275 | 3.272135425 | 0.00188 | 0.028935564 | -2.23284249 | Ribosomal protein | elongation zone | 12 h |
| HORVU.MOREX.r3.7HG0640980.1 | 2.178049671 | 5.107500163 | 2.986244585 | 0.00427 | 0.041090601 | -2.23357612 | Histone-lysine N-methyltransferase, H3 lysine-36 specific | elongation zone | 12 h |
| HORVU.MOREX.r3.2HG0135860.1 | 0.424564003 | 5.818016201 | 3.18526106 | 0.00242 | 0.032575632 | -2.23422506 | Transcription factor jumonji domain-containing protein | elongation zone | 12 h |
| HORVU.MOREX.r3.4HG0393080.1 | 1.069969998 | 5.199963551 | 3.084169461 | 0.00324 | 0.036928455 | -2.23656191 | Translocase of chloroplast | elongation zone | 12 h |
| HORVU.MOREX.r3.5HG0513020.1 | 0.312561365 | 6.516881897 | 3.263078166 | 0.00193 | 0.029341049 | -2.23760849 | Transmembrane protein 147 | elongation zone | 12 h |
| HORVU.MOREX.r3.4HG0382670.1 | -0.731048261 | 0.489338732 | -3.160778558 | 0.0026 | 0.033587945 | -2.23820591 | Xyloglucan 6-xylosyltransferase 1 | elongation zone | 12 h |
| HORVU.MOREX.r3.6HG0576550.1 | 0.410575492 | 4.943295042 | 3.190240218 | 0.00239 | 0.032288187 | -2.23954932 | F-box protein | elongation zone | 12 h |
| HORVU.MOREX.r3.7HG0680610.1 | 1.521163888 | 6.192253708 | 3.114619258 | 0.00297 | 0.035453096 | -2.2396383 | Protein kinase | elongation zone | 12 h |
| HORVU.MOREX.r3.2HG0203500.1 | 0.77350863 | 6.139707515 | 3.201893331 | 0.00231 | 0.031894634 | -2.24065692 | UPF0136 membrane protein | elongation zone | 12 h |
| HORVU.MOREX.r3.5HG0483020.1 | 0.861524531 | 5.236287099 | 3.119461137 | 0.00293 | 0.035308848 | -2.24108811 | Kinesin-like protein | elongation zone | 12 h |

**Table S5** Continued.

| ID | logFC | AveExpr | t | P.Value | adj.P.Val | B | gene description | root zone | time point |
|---|---|---|---|---|---|---|---|---|---|
| HORVU.MOREX.r3.1HG0038960.1 | 0.83681179 | 2.949856776 | 2.981650134 | 0.00432 | 0.041301836 | -2.24192919 | #N/A | elongation zone | 12 h |
| HORVU.MOREX.r3.5HG0474720.1 | 2.007273662 | -0.474518298 | 2.901738922 | 0.00539 | 0.046372512 | -2.24200804 | Short-chain dehydrogenase/reductase family protein | elongation zone | 12 h |
| HORVU.MOREX.r3.1HG0045520.1 | 2.041508145 | 1.200266662 | 2.875502219 | 0.0058 | 0.047903389 | -2.24283798 | zinc finger MYM-type-like protein | elongation zone | 12 h |
| HORVU.MOREX.r3.5HG0524520.1 | 1.144436756 | 6.419522117 | 3.149838878 | 0.00268 | 0.033864708 | -2.24301433 | 60S ribosomal protein L24 | elongation zone | 12 h |
| HORVU.MOREX.r3.6HG0614660.1 | 2.251970492 | -0.348204795 | 2.909283439 | 0.00528 | 0.045978938 | -2.24302295 | Pentatricopeptide repeat-containing protein | elongation zone | 12 h |
| HORVU.MOREX.r3.6HG0593480.1 | 0.766220052 | 6.095309536 | 3.166544461 | 0.00256 | 0.033222239 | -2.24390502 | Arginine/serine-rich splicing factor, putative | elongation zone | 12 h |
| HORVU.MOREX.r3.5HG0486260.1 | 3.324378483 | 3.929099308 | 2.893510421 | 0.00552 | 0.046829376 | -2.2445433 | Heat shock transcription factor | elongation zone | 12 h |
| HORVU.MOREX.r3.5HG0493030.1 | 0.802374771 | 7.858922883 | 3.25318045 | 0.00199 | 0.029748302 | -2.24579727 | 60S ribosomal protein L32 | elongation zone | 12 h |
| HORVU.MOREX.r3.6HG0600870.1 | -0.40472359 | 4.403104904 | -3.215339751 | 0.00222 | 0.031454344 | -2.24657073 | Acetylglutamate kinase-like protein | elongation zone | 12 h |
| HORVU.MOREX.r3.7HG0715500.1 | 2.269196132 | 0.698143251 | 2.883000704 | 0.00568 | 0.047427345 | -2.2472865 | nuclear pore anchor | elongation zone | 12 h |
| HORVU.MOREX.r3.2HG0215220.1 | 0.715792721 | 4.050855349 | 3.075708123 | 0.00332 | 0.037190743 | -2.24739646 | Tubby-like F-box protein | elongation zone | 12 h |
| HORVU.MOREX.r3.7HG0713670.1 | 0.710373737 | 5.990306123 | 3.149616976 | 0.00269 | 0.033864708 | -2.24793616 | Alba DNA/RNA-binding protein | elongation zone | 12 h |
| HORVU.MOREX.r3.6HG0553290.1 | 0.618537389 | 5.490398459 | 3.149262752 | 0.00269 | 0.033864708 | -2.24915035 | Glycosyltransferase | elongation zone | 12 h |
| HORVU.MOREX.r3.6HG0544580.1 | 1.062300143 | 0.367989846 | 2.943273257 | 0.00481 | 0.043773947 | -2.24924052 | Werner Syndrome-like exonuclease | elongation zone | 12 h |
| HORVU.MOREX.r3.7HG0726240.1 | -0.82271414 | -0.068180582 | -3.089240941 | 0.00319 | 0.036823467 | -2.24985318 | Retrotransposon protein SINE subclass | elongation zone | 12 h |
| HORVU.MOREX.r3.1HG0022710.1 | 0.825038705 | 10.37598935 | 3.358676936 | 0.00146 | 0.027115369 | -2.25180384 | 40S ribosomal protein S4 | elongation zone | 12 h |
| HORVU.MOREX.r3.7HG0696340.1 | 0.441696898 | 6.08013092 | 3.194723399 | 0.00236 | 0.032009236 | -2.25182091 | RNA-binding family protein | elongation zone | 12 h |
| HORVU.MOREX.r3.1HG0060410.1 | 1.15778268 | 3.806642977 | 2.961979897 | 0.00457 | 0.042538646 | -2.25375157 | F21O3.15 protein | elongation zone | 12 h |
| HORVU.MOREX.r3.5HG0420370.1 | 2.577187288 | 2.703090315 | 2.878905463 | 0.00574 | 0.04776728 | -2.25421221 | receptor kinase 1 | elongation zone | 12 h |
| HORVU.MOREX.r3.2HG0172550.1 | 1.198592076 | 2.80726014 | 2.930155384 | 0.00499 | 0.044637748 | -2.25429582 | Synaptonemal complex protein 1 | elongation zone | 12 h |
| HORVU.MOREX.r3.4HG0415590.1 | 1.417263094 | -0.042390657 | 2.973709916 | 0.00442 | 0.041779146 | -2.25645446 | Bifunctional inhibitor/lipid-transfer protein/seed storage 2S albu | elongation zone | 12 h |
| HORVU.MOREX.r3.4HG0358390.1 | 1.559887161 | 4.871920016 | 3.001041531 | 0.0041 | 0.040265883 | -2.25662626 | Selenoprotein H | elongation zone | 12 h |
| HORVU.MOREX.r3.7HG0733600.1 | 0.838951582 | 8.484163878 | 3.270749002 | 0.00189 | 0.028966748 | -2.25666936 | 30S ribosomal protein S11 | elongation zone | 12 h |
| HORVU.MOREX.r3.5HG0519760.1 | 0.65610332 | 8.912276545 | 3.301331907 | 0.00173 | 0.028397697 | -2.25686681 | Eukaryotic translation initiation factor 3 subunit B | elongation zone | 12 h |
| HORVU.MOREX.r3.6HG0565380.1 | 0.502605971 | 5.049690937 | 3.146972377 | 0.00271 | 0.033864708 | -2.25699495 | Zinc finger CCCH domain-containing protein 13 | elongation zone | 12 h |
| HORVU.MOREX.r3.3HG0250080.1 | 0.692220495 | 5.216335383 | 3.121158307 | 0.00291 | 0.035234858 | -2.2578827 | 30S ribosomal protein S8, putative | elongation zone | 12 h |
| HORVU.MOREX.r3.3HG0249590.1 | 0.703218326 | 3.607050289 | 3.005700607 | 0.00404 | 0.040178373 | -2.25847241 | Vesicle transport protein-like | elongation zone | 12 h |
| HORVU.MOREX.r3.1HG0031520.1 | 0.737716091 | 4.885472537 | 3.092226371 | 0.00317 | 0.036688668 | -2.25928732 | Myb | elongation zone | 12 h |
| HORVU.MOREX.r3.1HG0054150.1 | 0.911144648 | 5.965194148 | 3.128004293 | 0.00286 | 0.034844441 | -2.25945818 | N-alpha-acetyltransferase 25, NatB auxiliary subunit | elongation zone | 12 h |
| HORVU.MOREX.r3.7HG0751140.1 | 0.99393265 | 2.824962476 | 2.924351019 | 0.00507 | 0.045122895 | -2.26120163 | F-box family protein | elongation zone | 12 h |
| HORVU.MOREX.r3.2HG0191100.1 | 0.760830552 | 8.580211615 | 3.280121765 | 0.00184 | 0.028819302 | -2.2614823 | 30S ribosomal protein S17 | elongation zone | 12 h |
| HORVU.MOREX.r3.7HG0712850.1 | 0.359882432 | 4.698390518 | 3.151417845 | 0.00267 | 0.033864708 | -2.26157925 | Fiber protein Fb34 | elongation zone | 12 h |
| HORVU.MOREX.r3.2HG0113380.1 | 0.525816602 | 7.165397632 | 3.227236414 | 0.00215 | 0.031018589 | -2.26333793 | splicing factor-like protein | elongation zone | 12 h |
| HORVU.MOREX.r3.2HG0185640.1 | 2.385259077 | 1.439291744 | 2.86596624 | 0.00595 | 0.048522465 | -2.26334968 | Protein ROOT PRIMORDIUM DEFECTIVE 1 | elongation zone | 12 h |
| HORVU.MOREX.r3.6HG0566480.1 | 2.309414619 | -1.265754862 | 2.887174491 | 0.00561 | 0.047204707 | -2.26421565 | Retrotransposon protein, putative, unclassified | elongation zone | 12 h |
| HORVU.MOREX.r3.6HG0568390.1 | -0.439704749 | 4.337128784 | -3.214604915 | 0.00223 | 0.031473786 | -2.26608131 | Kinase family protein | elongation zone | 12 h |
| HORVU.MOREX.r3.7HG0724270.1 | -0.494398676 | 2.618489122 | -3.034735887 | 0.00373 | 0.038784658 | -2.2674011 | F-box/kelch-repeat protein | elongation zone | 12 h |
| HORVU.MOREX.r3.2HG0136670.1 | 1.414639516 | 4.71390114 | 3.265661057 | 0.00192 | 0.029184417 | -2.26748348 | Cellulose synthase-like protein | elongation zone | 12 h |
| HORVU.MOREX.r3.2HG0158080.1 | 2.186685325 | 0.623463989 | 2.865329728 | 0.00596 | 0.048546205 | -2.26759889 | Pentatricopeptide repeat-containing family protein | elongation zone | 12 h |
| HORVU.MOREX.r3.7HG0707410.1 | 1.84466427 | -0.496688719 | 2.863895549 | 0.00598 | 0.048636131 | -2.26819657 | GRF zinc finger / Zinc knuckle protein | elongation zone | 12 h |
| HORVU.MOREX.r3.5HG0455490.1 | 0.879649417 | 1.390809013 | 2.896703289 | 0.00547 | 0.046711054 | -2.27167605 | Aspartic proteinase Asp1 | elongation zone | 12 h |

**Table S5** Continued.

| ID | logFC | AveExpr | t | P.Value | adj.P.Val | B | gene description | root zone | time point |
|---|---|---|---|---|---|---|---|---|---|
| HORVU.MOREX.r3.4HG0411600.1 | 0.563670925 | 7.727342404 | 3.244552117 | 0.00204 | 0.030344376 | -2.27182674 | 50S ribosomal protein L14 | elongation zone | 12 h |
| HORVU.MOREX.r3.7HG0697550.1 | 0.354853515 | 9.219800437 | 3.32868405 | 0.00159 | 0.027806246 | -2.27282758 | ABC transporter ATP-binding protein ARB1 | elongation zone | 12 h |
| HORVU.MOREX.r3.3HG0270190.1 | 0.313518027 | 5.543138586 | 3.203064696 | 0.0023 | 0.031887924 | -2.27310215 | dihydroflavonol 4-reductase/flavanone protein | elongation zone | 12 h |
| HORVU.MOREX.r3.3HG0325650.1 | 1.276170392 | 2.127342509 | 2.8778587 | 0.00576 | 0.04776728 | -2.27479877 | Cupredoxin superfamily protein, putative | elongation zone | 12 h |
| HORVU.MOREX.r3.3HG0327830.1 | 2.137846479 | 4.138764232 | 2.913633033 | 0.00522 | 0.045722475 | -2.27527267 | Pentatricopeptide repeat-containing protein | elongation zone | 12 h |
| HORVU.MOREX.r3.7HG0685410.1 | 0.962560261 | 5.553236747 | 3.098437234 | 0.00311 | 0.036312806 | -2.27537999 | Retinoblastoma-related protein | elongation zone | 12 h |
| HORVU.MOREX.r3.1HG0085140.1 | 0.383226484 | 6.163123216 | 3.216500631 | 0.00221 | 0.031452409 | -2.27631572 | Serine/threonine protein phosphatase 2A regulatory subunit B | elongation zone | 12 h |
| HORVU.MOREX.r3.6HG0552890.1 | 1.993175621 | 0.455902627 | 2.860005763 | 0.00605 | 0.048928619 | -2.27657729 | transmembrane protein, putative (DUF594) | elongation zone | 12 h |
| HORVU.MOREX.r3.7HG0713340.1 | 1.151722849 | 3.465224865 | 2.984477492 | 0.00429 | 0.041182464 | -2.27710077 | Peroxidase | elongation zone | 12 h |
| HORVU.MOREX.r3.4HG0415160.1 | 0.407774584 | 4.127593268 | 3.089248152 | 0.00319 | 0.036823467 | -2.27938096 | DUF3527 domain protein | elongation zone | 12 h |
| HORVU.MOREX.r3.1HG0079930.1 | -2.638199949 | -2.565871354 | -2.884401993 | 0.00566 | 0.047404516 | -2.28155431 | Pectin lyase-like superfamily protein | elongation zone | 12 h |
| HORVU.MOREX.r3.7HG0721450.1 | -0.699863446 | 1.569350469 | -3.277642928 | 0.00185 | 0.028860537 | -2.28164004 | Peroxidase | elongation zone | 12 h |
| HORVU.MOREX.r3.6HG0623920.1 | 0.208032036 | 9.176211122 | 3.362615964 | 0.00144 | 0.02703001 | -2.28197464 | Leucine aminopeptidase, putative | elongation zone | 12 h |
| HORVU.MOREX.r3.5HG0508060.1 | 1.026380894 | 6.113062349 | 3.122597821 | 0.0029 | 0.035200058 | -2.28307525 | GTP binding protein | elongation zone | 12 h |
| HORVU.MOREX.r3.3HG0278550.1 | 0.982292361 | 2.026307923 | 2.979918031 | 0.00434 | 0.041386317 | -2.28368761 | Peroxidase | elongation zone | 12 h |
| HORVU.MOREX.r3.6HG0549900.1 | 2.443521097 | 1.036021217 | 2.868035948 | 0.00592 | 0.048306767 | -2.28798672 | Hydroxycinnamoyl-CoA shikimate/quinate hydroxycinnamoyltrar | elongation zone | 12 h |
| HORVU.MOREX.r3.1HG0065540.1 | 1.130671554 | 10.14930621 | 3.321352924 | 0.00163 | 0.02796111 | -2.28977454 | NA | elongation zone | 12 h |
| HORVU.MOREX.r3.6HG0600630.1 | 1.496651395 | -0.318534445 | 2.852649739 | 0.00617 | 0.049512065 | -2.29213933 | cotton fiber-like protein (DUF761) | elongation zone | 12 h |
| HORVU.MOREX.r3.2HG0122450.1 | 0.880003501 | 8.533148489 | 3.255005892 | 0.00198 | 0.029648381 | -2.29228777 | 30S ribosomal protein S4 | elongation zone | 12 h |
| HORVU.MOREX.r3.1HG0056560.1 | 0.318483658 | 5.466996869 | 3.19506386 | 0.00236 | 0.032009236 | -2.29469223 | AP-3 complex subunit delta | elongation zone | 12 h |
| HORVU.MOREX.r3.2HG0107960.1 | -0.486438392 | 3.123366617 | -3.084192572 | 0.00324 | 0.036928455 | -2.29497315 | Phosphoinositide phospholipase C | elongation zone | 12 h |
| HORVU.MOREX.r3.7HG0671800.1 | 1.312801338 | 5.765569408 | 3.071059255 | 0.00336 | 0.037308039 | -2.29575993 | DNA topoisomerase family | elongation zone | 12 h |
| HORVU.MOREX.r3.4HG0417170.1 | 1.902631521 | 5.34885237 | 2.993954226 | 0.00418 | 0.040647009 | -2.29648126 | WD repeat-containing protein | elongation zone | 12 h |
| HORVU.MOREX.r3.5HG0513980.1 | -1.03971357 | -0.364729073 | -3.020526241 | 0.00388 | 0.039407591 | -2.29776697 | Auxin-responsive protein | elongation zone | 12 h |
| HORVU.MOREX.r3.2HG0201910.1 | 0.760158569 | 6.429805807 | 3.177066458 | 0.00248 | 0.032834207 | -2.29931708 | Myosin-H heavy chain | elongation zone | 12 h |
| HORVU.MOREX.r3.2HG0132450.1 | -0.462628978 | 4.0992375 | -3.180280669 | 0.00246 | 0.032737911 | -2.30120387 | L-rhamnose isomerase | elongation zone | 12 h |
| HORVU.MOREX.r3.5HG0501430.1 | 1.539580101 | 1.709795684 | 2.984358844 | 0.00429 | 0.041182464 | -2.30169855 | Agmatine coumaroyltransferase-2 | elongation zone | 12 h |
| HORVU.MOREX.r3.3HG0255500.1 | 1.210363698 | 3.245772951 | 2.91687243 | 0.00517 | 0.045490951 | -2.30188893 | La related protein-like | elongation zone | 12 h |
| HORVU.MOREX.r3.7HG0650810.1 | 1.482593165 | 1.931516835 | 2.863548354 | 0.00599 | 0.048636131 | -2.30503837 | Acyl-CoA synthetase family protein | elongation zone | 12 h |
| HORVU.MOREX.r3.1HG0079280.1 | 0.932577532 | 8.411998552 | 3.255880596 | 0.00197 | 0.029636522 | -2.30782291 | Histone H2B | elongation zone | 12 h |
| HORVU.MOREX.r3.3HG0315620.1 | 1.288139403 | 1.678500581 | 2.851381963 | 0.00619 | 0.049563437 | -2.30803657 | #N/A | elongation zone | 12 h |
| HORVU.MOREX.r3.7HG0751330.1 | 2.61572078 | 4.028668411 | 2.876953057 | 0.00577 | 0.047810689 | -2.30919417 | Core-2/I-branching beta-1,6-N-acetylglucosaminyltransferase far | elongation zone | 12 h |
| HORVU.MOREX.r3.5HG0463700.1 | 0.553087549 | 4.020835168 | 3.039490544 | 0.00368 | 0.038641262 | -2.30962491 | Scarecrow transcription factor family protein | elongation zone | 12 h |
| HORVU.MOREX.r3.5HG0487070.1 | -0.452303843 | 4.654929378 | -3.170361001 | 0.00253 | 0.033099534 | -2.31148453 | Alpha/beta-Hydrolases superfamily protein, putative | elongation zone | 12 h |
| HORVU.MOREX.r3.6HG0611090.1 | 1.921649698 | 4.272440344 | 2.913065656 | 0.00523 | 0.04573542 | -2.31160216 | tRNA (Guanine(9)-N1)-methyltransferase | elongation zone | 12 h |
| HORVU.MOREX.r3.5HG0427370.1 | 0.882588634 | 1.30418145 | 3.011424483 | 0.00398 | 0.03986915 | -2.31208128 | #N/A | elongation zone | 12 h |
| HORVU.MOREX.r3.2HG0111240.1 | 1.381362073 | 5.668492978 | 3.058677097 | 0.00348 | 0.037779464 | -2.31228711 | Nuclear pore complex protein, putative | elongation zone | 12 h |
| HORVU.MOREX.r3.4HG0364050.1 | -0.642243266 | 2.580176455 | -3.03970572 | 0.00367 | 0.038641262 | -2.3126369 | Sodium/hydrogen exchanger | elongation zone | 12 h |
| HORVU.MOREX.r3.7HG0663940.1 | -0.779305635 | 0.764913385 | -3.157056281 | 0.00263 | 0.033716726 | -2.31366258 | Alpha/beta-Hydrolases superfamily protein, putative | elongation zone | 12 h |
| HORVU.MOREX.r3.5HG0525290.1 | -0.689189282 | 5.237284387 | -3.231487593 | 0.00212 | 0.030866089 | -2.31525468 | Amino acid transporter, putative | elongation zone | 12 h |
| HORVU.MOREX.r3.3HG0265880.1 | 1.173138703 | 2.376813197 | 2.891980283 | 0.00554 | 0.046904576 | -2.31628249 | Pentatricopeptide repeat-containing protein | elongation zone | 12 h |

**Table S5** Continued.

| ID | logFC | AveExpr | t | P.Value | adj.P.Val | B | gene description | root zone | time point |
|---|---|---|---|---|---|---|---|---|---|
| HORVU.MOREX.r3.2HG0113730.1 | 1.496439827 | 5.221019143 | 3.005586597 | 0.00404 | 0.040178373 | -2.31761402 | 60S ribosome subunit biogenesis protein NIP7 homolog | elongation zone | 12 h |
| HORVU.MOREX.r3.2HG0178240.1 | 0.632328958 | 4.274946849 | 3.038394129 | 0.00369 | 0.038671495 | -2.31967898 | Transcription factor | elongation zone | 12 h |
| HORVU.MOREX.r3.6HG0560680.1 | -1.279253164 | -0.976729155 | -2.853702639 | 0.00615 | 0.049405649 | -2.31983088 | #N/A | elongation zone | 12 h |
| HORVU.MOREX.r3.2HG0154600.1 | 2.306870289 | 1.249898666 | 2.846052117 | 0.00628 | 0.04989211 | -2.3201747 | Plastid division protein PDV2 | elongation zone | 12 h |
| HORVU.MOREX.r3.2HG0107450.1 | 0.863007992 | 3.576514789 | 2.995386716 | 0.00416 | 0.040578362 | -2.32225816 | methyl-coenzyme M reductase II subunit gamma, putative (DUF3 | elongation zone | 12 h |
| HORVU.MOREX.r3.7HG0736300.1 | 2.221076572 | 3.861077284 | 2.879522548 | 0.00573 | 0.047754823 | -2.3228241 | Glycosyltransferase | elongation zone | 12 h |
| HORVU.MOREX.r3.2HG0187640.1 | 0.650240909 | 2.595046236 | 2.949509127 | 0.00473 | 0.043465368 | -2.32371355 | Gamma-tubulin complex component | elongation zone | 12 h |
| HORVU.MOREX.r3.2HG0191180.1 | 0.414106004 | 4.405018466 | 3.097901948 | 0.00311 | 0.036334581 | -2.32388382 | Xanthine phosphoribosyltransferase | elongation zone | 12 h |
| HORVU.MOREX.r3.5HG0438550.1 | 1.326017211 | 3.407298471 | 2.899724969 | 0.00542 | 0.046551797 | -2.32420098 | Cyclin family protein | elongation zone | 12 h |
| HORVU.MOREX.r3.2HG0140950.1 | 1.457654007 | 3.361687211 | 2.883123296 | 0.00568 | 0.047427345 | -2.32715005 | Abnormal spindle-like microcephaly-associated protein-like prot | elongation zone | 12 h |
| HORVU.MOREX.r3.4HG0411620.1 | 1.445576433 | 6.015681025 | 3.055425695 | 0.00351 | 0.037808566 | -2.32756499 | Thyroid adenoma-associated protein-like protein | elongation zone | 12 h |
| HORVU.MOREX.r3.1HG0070000.1 | 0.72621818 | 1.808570831 | 2.91965 | 0.00513 | 0.045371176 | -2.32816545 | Histone H3 | elongation zone | 12 h |
| HORVU.MOREX.r3.5HG0451540.1 | 0.702683489 | 7.656228475 | 3.215007303 | 0.00222 | 0.031458646 | -2.3283289 | Chaperone DnaK | elongation zone | 12 h |
| HORVU.MOREX.r3.7HG0714510.1 | 0.970958513 | 5.004338229 | 3.029660251 | 0.00378 | 0.039090877 | -2.32935329 | Protein phosphatase 2C | elongation zone | 12 h |
| HORVU.MOREX.r3.1HG0000380.1 | 1.547131122 | 4.008859542 | 2.958164085 | 0.00462 | 0.042747009 | -2.3333101 | Pectin lyase-like superfamily protein | elongation zone | 12 h |
| HORVU.MOREX.r3.5HG0425730.1 | -0.500574564 | 4.697689499 | -3.284696598 | 0.00181 | 0.028793086 | -2.33730309 | Receptor-like kinase | elongation zone | 12 h |
| HORVU.MOREX.r3.3HG0324930.1 | 0.611359194 | 9.79870902 | 3.316894398 | 0.00165 | 0.028050531 | -2.33976477 | 40S ribosomal protein S10 | elongation zone | 12 h |
| HORVU.MOREX.r3.1HG0075830.1 | 1.204140936 | 3.021026521 | 2.883317341 | 0.00567 | 0.047427345 | -2.33981085 | Polypyrimidine tract-binding protein-like protein | elongation zone | 12 h |
| HORVU.MOREX.r3.1HG0094010.1 | 2.369618966 | -1.304615689 | 2.857974511 | 0.00608 | 0.049103351 | -2.34134709 | Kinase family protein | elongation zone | 12 h |
| HORVU.MOREX.r3.7HG0687630.1 | 0.532710823 | 7.969316852 | 3.274007835 | 0.00187 | 0.028923495 | -2.3416885 | Acyl carrier protein | elongation zone | 12 h |
| HORVU.MOREX.r3.4HG0336310.1 | 0.665749298 | 6.502870264 | 3.167117048 | 0.00255 | 0.033216773 | -2.3425458 | Zinc finger family protein | elongation zone | 12 h |
| HORVU.MOREX.r3.2HG0124990.1 | 0.621722156 | 6.152886279 | 3.177039163 | 0.00248 | 0.032834207 | -2.34413468 | 3-oxoacyl-[acyl-carrier-protein] synthase | elongation zone | 12 h |
| HORVU.MOREX.r3.6HG0586220.1 | 0.896956745 | 5.61880713 | 3.099144584 | 0.0031 | 0.036288602 | -2.34543033 | Prohibitin | elongation zone | 12 h |
| HORVU.MOREX.r3.1HG0034450.1 | -1.058291733 | 0.621391691 | -2.869552888 | 0.00589 | 0.048306767 | -2.34556041 | Retrovirus-related Pol polyprotein from transposon TNT 1-94 | elongation zone | 12 h |
| HORVU.MOREX.r3.7HG0751790.1 | 1.724851416 | 0.792911227 | 2.875576883 | 0.0058 | 0.047903389 | -2.34558588 | DUF241 domain protein (DUF241) | elongation zone | 12 h |
| HORVU.MOREX.r3.4HG0384230.1 | 0.496041279 | 3.949265758 | 3.080778888 | 0.00327 | 0.036983216 | -2.34713495 | RING/U-box superfamily protein | elongation zone | 12 h |
| HORVU.MOREX.r3.3HG0243920.1 | 1.076796871 | 3.290866018 | 2.9679441 | 0.00449 | 0.042132428 | -2.34768507 | VQ motif-containing protein | elongation zone | 12 h |
| HORVU.MOREX.r3.7HG0641160.1 | 0.366193133 | 5.342895251 | 3.141755174 | 0.00275 | 0.034124913 | -2.34833116 | Guanine nucleotide-binding protein subunit alpha-like protein | elongation zone | 12 h |
| HORVU.MOREX.r3.3HG0289250.1 | 0.638368448 | 7.766476009 | 3.231515012 | 0.00212 | 0.030866089 | -2.34858156 | Elongation factor 4 | elongation zone | 12 h |
| HORVU.MOREX.r3.2HG0167030.1 | 0.726699799 | 3.461475439 | 2.94312599 | 0.00481 | 0.043773947 | -2.35098497 | Mis18-binding protein 1 | elongation zone | 12 h |
| HORVU.MOREX.r3.2HG0128410.1 | 0.885062207 | 4.563754238 | 3.014140507 | 0.00395 | 0.039746874 | -2.3524001 | DNA/RNA helicase protein | elongation zone | 12 h |
| HORVU.MOREX.r3.5HG0534140.1 | 1.269077477 | 8.004889694 | 3.210990048 | 0.00225 | 0.031545502 | -2.35255146 | Glycine rich protein 3 | elongation zone | 12 h |
| HORVU.MOREX.r3.5HG0446850.1 | 1.931822485 | 6.103829376 | 3.010880798 | 0.00398 | 0.039893403 | -2.35281947 | Myosin-1 | elongation zone | 12 h |
| HORVU.MOREX.r3.1HG0020980.1 | -0.505075261 | 7.252012736 | -3.317749012 | 0.00164 | 0.028035058 | -2.35345235 | Endo-1,31,4-beta-D-glucanase | elongation zone | 12 h |
| HORVU.MOREX.r3.4HG0382090.1 | 0.495931996 | 6.628026251 | 3.180833339 | 0.00245 | 0.032706849 | -2.35365503 | #N/A | elongation zone | 12 h |
| HORVU.MOREX.r3.1HG0060370.1 | 2.617985893 | -0.902409718 | 2.848505941 | 0.00624 | 0.049739166 | -2.35368169 | 1,4-alpha-glucan branching enzyme GlgB | elongation zone | 12 h |
| HORVU.MOREX.r3.1HG0060320.1 | 0.832555173 | 7.084889179 | 3.184959932 | 0.00243 | 0.032575632 | -2.3561673 | Histone H3 | elongation zone | 12 h |
| HORVU.MOREX.r3.1HG0051900.1 | -0.536150041 | 4.657334527 | -3.224923282 | 0.00216 | 0.031159303 | -2.35735927 | Glutathione S-transferase | elongation zone | 12 h |
| HORVU.MOREX.r3.7HG0746570.1 | 0.860675161 | 9.010351969 | 3.261296092 | 0.00194 | 0.029451158 | -2.35980052 | 60S ribosomal protein L27 | elongation zone | 12 h |
| HORVU.MOREX.r3.5HG0494480.1 | 0.721096419 | 9.280053353 | 3.273313194 | 0.00187 | 0.028923495 | -2.36237551 | 60S ribosomal protein L7a | elongation zone | 12 h |
| HORVU.MOREX.r3.4HG0396600.1 | -0.637932571 | 1.930667574 | -2.965772275 | 0.00452 | 0.042342202 | -2.36364607 | Digalactosyldiacylglycerol synthase (Chloroplastic) | elongation zone | 12 h |

**Table S5** Continued.

| ID | logFC | AveExpr | t | P.Value | adj.P.Val | B | gene description | root zone | time point |
|---|---|---|---|---|---|---|---|---|---|
| HORVU.MOREX.r3.3HG0246030.1 | -0.906906744 | 0.793865926 | -3.039398299 | 0.00368 | 0.038641262 | -2.36590994 | Blue copper protein | elongation zone | 12 h |
| HORVU.MOREX.r3.7HG0677270.1 | 0.836675182 | 2.721723616 | 2.946374994 | 0.00477 | 0.04368814 | -2.36848583 | Glycosyltransferase | elongation zone | 12 h |
| HORVU.MOREX.r3.3HG0327170.1 | 0.422533198 | 7.966567016 | 3.257597214 | 0.00196 | 0.02961891 | -2.37120169 | Bifunctional polymyxin resistance arnA protein | elongation zone | 12 h |
| HORVU.MOREX.r3.6HG0630680.1 | 0.58352657 | 4.625356458 | 3.039122022 | 0.00368 | 0.038641262 | -2.37302004 | F-box family protein | elongation zone | 12 h |
| HORVU.MOREX.r3.5HG0439040.1 | 0.581906443 | 7.004585033 | 3.17954824 | 0.00246 | 0.032786051 | -2.37417641 | Elongation factor Tu | elongation zone | 12 h |
| HORVU.MOREX.r3.3HG0271710.1 | 0.446505564 | 4.526521133 | 3.048046653 | 0.00359 | 0.038188048 | -2.3760517 | Zinc finger CCCH domain protein | elongation zone | 12 h |
| HORVU.MOREX.r3.2HG0189960.1 | 1.648420983 | 8.037428277 | 3.148288341 | 0.0027 | 0.033864708 | -2.37757216 | U3 small nucleolar RNA-associated protein 10 | elongation zone | 12 h |
| HORVU.MOREX.r3.5HG0518160.1 | 0.729845077 | 8.607332408 | 3.239329723 | 0.00207 | 0.03060905 | -2.37766762 | 30S ribosomal protein S13 | elongation zone | 12 h |
| HORVU.MOREX.r3.6HG0549510.1 | 0.66828419 | 8.771368712 | 3.290157227 | 0.00178 | 0.02862189 | -2.37890894 | Peptidyl-prolyl cis-trans isomerase | elongation zone | 12 h |
| HORVU.MOREX.r3.4HG0418790.1 | 1.385314891 | 5.632641367 | 3.008195557 | 0.00401 | 0.040081151 | -2.37991148 | tRNA/rRNA methyltransferase family protein | elongation zone | 12 h |
| HORVU.MOREX.r3.1HG0037990.1 | 0.949840918 | 2.905819651 | 2.890150059 | 0.00557 | 0.047062906 | -2.38095707 | SKP1-like protein | elongation zone | 12 h |
| HORVU.MOREX.r3.1HG0078010.1 | 0.355055091 | 5.540918622 | 3.138599434 | 0.00277 | 0.03425211 | -2.38265716 | Translation initiation factor IF-2 | elongation zone | 12 h |
| HORVU.MOREX.r3.5HG0494040.1 | -1.048179492 | -3.000857515 | -2.872306514 | 0.00585 | 0.04818291 | -2.3835304 | Auxin efflux carrier component | elongation zone | 12 h |
| HORVU.MOREX.r3.2HG0196780.1 | 1.391479302 | 4.987352017 | 2.98917198 | 0.00423 | 0.040956128 | -2.38384064 | Multidrug resistance protein ABC transporter family | elongation zone | 12 h |
| HORVU.MOREX.r3.5HG0497200.1 | 0.494750829 | 6.465584768 | 3.151842017 | 0.00267 | 0.033864708 | -2.38386363 | Cell division cycle 5-like protein | elongation zone | 12 h |
| HORVU.MOREX.r3.4HG0349380.1 | 0.689664029 | 10.03245682 | 3.302095643 | 0.00172 | 0.028397697 | -2.38700952 | Ribosomal protein L3 | elongation zone | 12 h |
| HORVU.MOREX.r3.7HG0702050.1 | -1.003240082 | -0.450446856 | -2.963932855 | 0.00454 | 0.04241279 | -2.38822628 | Blue copper protein | elongation zone | 12 h |
| HORVU.MOREX.r3.1HG0001130.1 | 0.742941765 | 5.638908604 | 3.067383262 | 0.0034 | 0.037381068 | -2.38825143 | Ankyrin repeat family protein | elongation zone | 12 h |
| HORVU.MOREX.r3.5HG0429360.1 | 0.71137268 | 4.758495845 | 3.062514099 | 0.00344 | 0.03764455 | -2.38900224 | Pectinesterase | elongation zone | 12 h |
| HORVU.MOREX.r3.7HG0680960.1 | -1.48703112 | 0.61149754 | -2.909084134 | 0.00529 | 0.045984662 | -2.389173 | electron transporter, putative (Protein of unknown function, DUI | elongation zone | 12 h |
| HORVU.MOREX.r3.3HG0330210.1 | 1.850783216 | 4.048874756 | 2.872196102 | 0.00585 | 0.04818291 | -2.39096341 | GDSL esterase/lipase | elongation zone | 12 h |
| HORVU.MOREX.r3.2HG0133550.1 | -0.262828495 | 5.768893736 | -3.199419762 | 0.00233 | 0.031903923 | -2.39153707 | Calcium-dependent protein kinase | elongation zone | 12 h |
| HORVU.MOREX.r3.4HG0343870.1 | 0.987754443 | 3.294242033 | 2.906245246 | 0.00533 | 0.04618095 | -2.39197122 | Transducin/WD40 repeat-like superfamily protein | elongation zone | 12 h |
| HORVU.MOREX.r3.3HG0242160.1 | 0.50994665 | 8.436022046 | 3.238384831 | 0.00208 | 0.030632868 | -2.39270616 | Arginine/serine-rich splicing factor, putative | elongation zone | 12 h |
| HORVU.MOREX.r3.7HG0701300.1 | 1.704012836 | 3.314094522 | 2.848754051 | 0.00624 | 0.049724941 | -2.39372518 | Exosome complex component | elongation zone | 12 h |
| HORVU.MOREX.r3.4HG0363440.1 | 1.173351947 | 5.032921778 | 2.984864597 | 0.00428 | 0.041182136 | -2.39417077 | DNA-directed RNA polymerase I subunit rpa49 | elongation zone | 12 h |
| HORVU.MOREX.r3.2HG0157290.1 | 0.843304417 | 3.397681153 | 2.940264507 | 0.00485 | 0.044001805 | -2.39514597 | Calcineurin-like metallo-phosphoesterase superfamily | elongation zone | 12 h |
| HORVU.MOREX.r3.5HG0421560.1 | 1.675078945 | 3.372970967 | 2.846280126 | 0.00628 | 0.049885976 | -2.39731054 | GTPase family protein | elongation zone | 12 h |
| HORVU.MOREX.r3.4HG0418210.1 | -0.762238867 | -0.146163126 | -3.027885888 | 0.0038 | 0.039108923 | -2.39867665 | Methylsterol monooxygenase 1-2 | elongation zone | 12 h |
| HORVU.MOREX.r3.1HG0069760.1 | -0.647297473 | 1.711709527 | -3.145237304 | 0.00272 | 0.033887701 | -2.40166617 | Calmodulin | elongation zone | 12 h |
| HORVU.MOREX.r3.5HG0452250.1 | -0.320648299 | 4.804561684 | -3.150252277 | 0.00268 | 0.033864708 | -2.40223754 | AP-2 complex subunit sigma | elongation zone | 12 h |
| HORVU.MOREX.r3.1HG0072360.1 | 0.827910352 | 2.892056241 | 2.901743492 | 0.00539 | 0.046372512 | -2.40273107 | F-box protein | elongation zone | 12 h |
| HORVU.MOREX.r3.2HG0104480.1 | 0.556317193 | 3.899783698 | 3.000410605 | 0.0041 | 0.040307075 | -2.40602023 | F-box protein | elongation zone | 12 h |
| HORVU.MOREX.r3.5HG0443560.1 | 0.422712504 | 3.753350984 | 3.018002116 | 0.0039 | 0.039547618 | -2.40615642 | antigenic heat-stable protein | elongation zone | 12 h |
| HORVU.MOREX.r3.3HG0308980.1 | 0.741012427 | 4.452518559 | 3.003694701 | 0.00406 | 0.040229538 | -2.40707756 | S-adenosyl-L-methionine-dependent methyltransferase, putative | elongation zone | 12 h |
| HORVU.MOREX.r3.5HG0527400.1 | 0.641563614 | 6.40973231 | 3.116775644 | 0.00295 | 0.035453096 | -2.40780612 | Nucleobase ascorbate transporter | elongation zone | 12 h |
| HORVU.MOREX.r3.7HG0723500.1 | 0.665332952 | 3.424822759 | 2.993088881 | 0.00419 | 0.040685115 | -2.41065629 | Cinnamoyl-CoA reductase family protein | elongation zone | 12 h |
| HORVU.MOREX.r3.6HG0603370.1 | -0.613715281 | 2.324583294 | -3.024695189 | 0.00383 | 0.03925602 | -2.41072082 | P-loop containing nucleoside triphosphate hydrolases superfamil | elongation zone | 12 h |
| HORVU.MOREX.r3.2HG0204040.1 | -1.163851706 | 1.864217479 | -2.939145524 | 0.00487 | 0.044099329 | -2.41136679 | Benzyl alcohol O-benzoyltransferase | elongation zone | 12 h |
| HORVU.MOREX.r3.4HG0339880.1 | 0.684031125 | 5.464296399 | 3.06166099 | 0.00345 | 0.037663482 | -2.41165003 | Zinc finger BED domain-containing protein DAYSLEEPER | elongation zone | 12 h |
| HORVU.MOREX.r3.2HG0217790.1 | 0.672477859 | 5.897336985 | 3.095894034 | 0.00313 | 0.036522359 | -2.41195509 | DNA/RNA helicase protein | elongation zone | 12 h |

**Table S5** Continued.

| ID | logFC | AveExpr | t | P.Value | adj.P.Val | B | gene description | root zone | time point |
|---|---|---|---|---|---|---|---|---|---|
| HORVU.MOREX.r3.2HG0218010.1 | 0.323501157 | 8.228324079 | 3.266856378 | 0.00191 | 0.02912546 | -2.41220445 | Dolichyl-diphosphooligosaccharide--protein glycosyltransferase s | elongation zone | 12 h |
| HORVU.MOREX.r3.5HG0524440.1 | 2.468436538 | 5.517230328 | 2.905802843 | 0.00533 | 0.04618095 | -2.41411861 | DNA gyrase subunit A | elongation zone | 12 h |
| HORVU.MOREX.r3.5HG0532380.1 | 0.526092009 | 3.73217286 | 3.0077102 | 0.00402 | 0.040116223 | -2.41415849 | #N/A | elongation zone | 12 h |
| HORVU.MOREX.r3.2HG0202070.1 | 0.40501178 | 5.440379588 | 3.099849256 | 0.0031 | 0.03627777 | -2.41537176 | Chaperone protein DnaJ | elongation zone | 12 h |
| HORVU.MOREX.r3.2HG0136790.1 | -0.427063674 | 6.404608075 | -3.286952699 | 0.0018 | 0.028691351 | -2.41655915 | Serine/threonine-protein kinase | elongation zone | 12 h |
| HORVU.MOREX.r3.6HG0554610.1 | 0.856449733 | 4.967810293 | 3.019285598 | 0.00389 | 0.039467053 | -2.41881249 | Pentatricopeptide repeat-containing protein | elongation zone | 12 h |
| HORVU.MOREX.r3.5HG0447730.1 | 0.591274089 | 6.269812166 | 3.130170491 | 0.00284 | 0.034669947 | -2.41925018 | Phenylalanine--tRNA ligase beta subunit | elongation zone | 12 h |
| HORVU.MOREX.r3.2HG0099270.1 | 0.671436002 | 4.423054289 | 2.995211978 | 0.00416 | 0.040578362 | -2.42332757 | Actin-related protein | elongation zone | 12 h |
| HORVU.MOREX.r3.3HG0218960.1 | 2.875912683 | 4.276243736 | 2.863299455 | 0.00599 | 0.048636131 | -2.4239297 | Cysteine/Histidine-rich C1 domain family protein | elongation zone | 12 h |
| HORVU.MOREX.r3.4HG0402170.1 | -1.133222673 | -0.055773461 | -2.931075095 | 0.00498 | 0.044603947 | -2.4264232 | Heat shock transcription factor | elongation zone | 12 h |
| HORVU.MOREX.r3.5HG0427110.1 | 0.719314275 | 3.005946314 | 2.887195683 | 0.00561 | 0.047204707 | -2.42675553 | tRNA modification GTPase MnmE | elongation zone | 12 h |
| HORVU.MOREX.r3.4HG0362850.1 | 0.937180675 | 2.700858012 | 2.86867784 | 0.00591 | 0.048306767 | -2.42711657 | Acyl-CoA-binding domain-containing protein 4 | elongation zone | 12 h |
| HORVU.MOREX.r3.2HG0196330.1 | 0.589706793 | 3.925350182 | 2.981749718 | 0.00432 | 0.041301836 | -2.42881483 | zinc finger matrin-type protein | elongation zone | 12 h |
| HORVU.MOREX.r3.2HG0173920.1 | 0.944338783 | 3.666148661 | 2.910781153 | 0.00526 | 0.045865572 | -2.43000042 | Chloroplastic group IIA intron splicing facilitator CRS1, chloroplas | elongation zone | 12 h |
| HORVU.MOREX.r3.3HG0304800.1 | 0.872629746 | 6.736708067 | 3.13194 | 0.00283 | 0.034584718 | -2.43099468 | Structural maintenance of chromosomes protein | elongation zone | 12 h |
| HORVU.MOREX.r3.5HG0438440.1 | 1.606556396 | 6.171046967 | 3.031549595 | 0.00376 | 0.03899655 | -2.43202681 | Kinesin-like protein | elongation zone | 12 h |
| HORVU.MOREX.r3.6HG0551740.1 | 0.538513784 | 5.295694184 | 3.075200597 | 0.00332 | 0.037190743 | -2.43446021 | Early light-inducible protein | elongation zone | 12 h |
| HORVU.MOREX.r3.7HG0635400.1 | 1.439567385 | 5.566493882 | 2.982038493 | 0.00432 | 0.04129762 | -2.43469085 | ATP-dependent RNA helicase family protein | elongation zone | 12 h |
| HORVU.MOREX.r3.7HG0643820.1 | 1.418937762 | 4.511304822 | 2.910642912 | 0.00526 | 0.045865572 | -2.43604906 | 6,7-dimethyl-8-ribityllumazine synthase | elongation zone | 12 h |
| HORVU.MOREX.r3.4HG0399020.1 | 0.7596266 | 10.02639982 | 3.281770895 | 0.00183 | 0.028819302 | -2.43788318 | 40S ribosomal protein S3a | elongation zone | 12 h |
| HORVU.MOREX.r3.6HG0604860.1 | 0.819524889 | 1.903057556 | 2.846850511 | 0.00627 | 0.049847313 | -2.43834375 | Glycosyltransferase | elongation zone | 12 h |
| HORVU.MOREX.r3.4HG0375480.1 | 0.933069324 | 5.989901741 | 3.09332942 | 0.00316 | 0.036644453 | -2.44075118 | 40S ribosomal protein S21 | elongation zone | 12 h |
| HORVU.MOREX.r3.1HG0080760.1 | 0.796271209 | 7.939029897 | 3.193469914 | 0.00237 | 0.032094481 | -2.44081765 | Histone H3 | elongation zone | 12 h |
| HORVU.MOREX.r3.4HG0385140.1 | 1.694291486 | 3.56681819 | 2.84594996 | 0.00628 | 0.04989211 | -2.44140948 | Carbamoyl-phosphate synthase large chain | elongation zone | 12 h |
| HORVU.MOREX.r3.4HG0380540.1 | -0.52880238 | 3.365321911 | -3.072899746 | 0.00334 | 0.037254747 | -2.44291172 | Aquaporin-1 | elongation zone | 12 h |
| HORVU.MOREX.r3.4HG0337170.1 | 0.931609903 | 5.504147714 | 3.039707692 | 0.00367 | 0.038641262 | -2.44382227 | Zinc-finger domain of monoamine-oxidase A repressor R1, putati | elongation zone | 12 h |
| HORVU.MOREX.r3.4HG0348470.1 | 0.587997936 | 6.0852859 | 3.113506963 | 0.00298 | 0.035522336 | -2.44384259 | ATP-dependent RNA helicase family protein | elongation zone | 12 h |
| HORVU.MOREX.r3.2HG0181390.1 | 1.435243101 | 5.181001175 | 2.977607929 | 0.00437 | 0.041519226 | -2.44676239 | Protein BREAST CANCER SUSCEPTIBILITY 1-like protein | elongation zone | 12 h |
| HORVU.MOREX.r3.7HG0726770.1 | -0.786701712 | 2.292249938 | -3.076537295 | 0.00331 | 0.037174732 | -2.4494717 | Glycine-rich protein | elongation zone | 12 h |
| HORVU.MOREX.r3.2HG0164540.1 | -1.082261358 | 0.757754706 | -2.846681217 | 0.00627 | 0.049850932 | -2.44979039 | Ethylene-responsive transcription factor, putative | elongation zone | 12 h |
| HORVU.MOREX.r3.1HG0082020.1 | 1.667012203 | 5.076015324 | 2.922894547 | 0.00509 | 0.045226145 | -2.45317515 | WD-repeat protein, putative | elongation zone | 12 h |
| HORVU.MOREX.r3.5HG0508150.1 | -0.952664491 | 2.451835168 | -2.996726992 | 0.00415 | 0.040483637 | -2.45358822 | 9-cis-epoxycarotenoid dioxygenase | elongation zone | 12 h |
| HORVU.MOREX.r3.1HG0019380.1 | 1.139935512 | 6.322023156 | 3.052596511 | 0.00354 | 0.037882054 | -2.45404855 | Glioma tumor suppressor-like protein | elongation zone | 12 h |
| HORVU.MOREX.r3.6HG0616130.1 | -0.486894962 | 6.042280496 | -3.256079508 | 0.00197 | 0.029636522 | -2.45499387 | Hexosyltransferase | elongation zone | 12 h |
| HORVU.MOREX.r3.6HG0546160.1 | -0.683514128 | 2.97610997 | -3.152216613 | 0.00267 | 0.033864708 | -2.45626533 | Acylamino-acid-releasing enzyme | elongation zone | 12 h |
| HORVU.MOREX.r3.7HG0747940.1 | -0.362410136 | 7.327177402 | -3.260539579 | 0.00195 | 0.029482217 | -2.4576376 | Polyamine oxidase | elongation zone | 12 h |
| HORVU.MOREX.r3.5HG0467870.1 | -0.775210511 | 2.049337326 | -3.020887034 | 0.00387 | 0.039407591 | -2.45813318 | Kynurenine formamidase | elongation zone | 12 h |
| HORVU.MOREX.r3.3HG0287640.1 | 0.522662408 | 5.945191223 | 3.102781652 | 0.00307 | 0.036128346 | -2.46007369 | Endoribonuclease YbeY | elongation zone | 12 h |
| HORVU.MOREX.r3.2HG0116280.1 | 1.955955558 | 7.824467532 | 3.079320387 | 0.00328 | 0.036983216 | -2.46196643 | Nucleolar family protein | elongation zone | 12 h |
| HORVU.MOREX.r3.5HG0472840.1 | -0.521599718 | 2.862571295 | -3.092453211 | 0.00316 | 0.036688668 | -2.46272011 | NBS-LRR disease resistance protein | elongation zone | 12 h |
| HORVU.MOREX.r3.6HG0540430.3 | -0.758792887 | 2.730080782 | -3.003769326 | 0.00406 | 0.040229538 | -2.46330518 | F-box and associated interaction domains-containing protein | elongation zone | 12 h |

**Table S5** Continued.

| ID | logFC | AveExpr | t | P.Value | adj.P.Val | B | gene description | root zone | time point |
|---|---|---|---|---|---|---|---|---|---|
| HORVU.MOREX.r3.1HG0059790.1 | 0.632238552 | 4.636699296 | 3.02896506 | 0.00379 | 0.039108923 | -2.46332967 | Hydroxyproline-rich glycoprotein-like | elongation zone | 12 h |
| HORVU.MOREX.r3.2HG0155810.1 | 0.743493165 | 3.567203574 | 2.970936915 | 0.00445 | 0.041967805 | -2.46455786 | Replication protein A 70 kDa DNA-binding subunit A | elongation zone | 12 h |
| HORVU.MOREX.r3.2HG0160930.1 | 0.642755589 | 9.218628834 | 3.240570734 | 0.00206 | 0.03054087 | -2.46456014 | 50S ribosomal protein L29 | elongation zone | 12 h |
| HORVU.MOREX.r3.3HG0290900.1 | 0.551515443 | 4.951870278 | 3.051483994 | 0.00355 | 0.037906495 | -2.46662441 | Rhomboid family protein | elongation zone | 12 h |
| HORVU.MOREX.r3.7HG0647360.1 | 1.430705396 | 5.375391659 | 2.951003454 | 0.00471 | 0.043324391 | -2.46699921 | Ribosome production factor 1 | elongation zone | 12 h |
| HORVU.MOREX.r3.6HG0610140.1 | 0.800890002 | 4.922162889 | 3.003377306 | 0.00407 | 0.040229538 | -2.46765292 | tRNA uridine 5-carboxymethylaminomethyl modification enzyme | elongation zone | 12 h |
| HORVU.MOREX.r3.4HG0347230.1 | 0.847182336 | 2.503173448 | 2.883525248 | 0.00567 | 0.047427345 | -2.46777527 | Polygalacturonase-1 non-catalytic beta subunit | elongation zone | 12 h |
| HORVU.MOREX.r3.1HG0064190.1 | 0.885868019 | 4.191166014 | 2.944289341 | 0.0048 | 0.043746094 | -2.46899801 | 30S ribosomal protein S18 | elongation zone | 12 h |
| HORVU.MOREX.r3.7HG0728060.1 | -0.451888306 | 5.512336647 | -3.129129075 | 0.00285 | 0.034752936 | -2.47173513 | Aminodeoxychorismate synthase | elongation zone | 12 h |
| HORVU.MOREX.r3.5HG0530420.1 | -0.876440271 | -0.668836111 | -3.01708203 | 0.00392 | 0.03957523 | -2.47176319 | Leucine-rich repeat receptor-like protein kinase | elongation zone | 12 h |
| HORVU.MOREX.r3.4HG0415420.1 | 0.554077964 | 6.947213857 | 3.17817799 | 0.00247 | 0.032834207 | -2.47182963 | Tubulin beta chain | elongation zone | 12 h |
| HORVU.MOREX.r3.3HG0234340.1 | 0.822425084 | 2.809522003 | 2.85530491 | 0.00612 | 0.049306414 | -2.47284495 | Protein FRA10AC1 | elongation zone | 12 h |
| HORVU.MOREX.r3.2HG0142190.1 | 0.318515243 | 5.689012541 | 3.110725767 | 0.003 | 0.035642237 | -2.47409286 | Potassium transporter | elongation zone | 12 h |
| HORVU.MOREX.r3.2HG0119860.2 | 0.75005228 | 6.022614408 | 3.095158274 | 0.00314 | 0.036539994 | -2.47449001 | 10 kDa chaperonin | elongation zone | 12 h |
| HORVU.MOREX.r3.7HG0656030.1 | -0.601122951 | 3.605834777 | -3.124233039 | 0.00289 | 0.035131319 | -2.47627326 | Receptor-like kinase | elongation zone | 12 h |
| HORVU.MOREX.r3.3HG0267880.1 | 1.381119207 | 6.864792855 | 3.061106597 | 0.00346 | 0.037702619 | -2.47673924 | Small subunit processome component 20-like protein | elongation zone | 12 h |
| HORVU.MOREX.r3.4HG0403800.1 | 0.929709129 | 5.863635554 | 3.032447644 | 0.00375 | 0.038936762 | -2.47702004 | Nuclear pore complex protein NUP96 | elongation zone | 12 h |
| HORVU.MOREX.r3.1HG0083020.1 | -0.861629366 | 0.461711884 | -2.884700112 | 0.00565 | 0.047400081 | -2.47752707 | aconitase 2 | elongation zone | 12 h |
| HORVU.MOREX.r3.2HG0166640.1 | 1.129147215 | 8.213121972 | 3.152913947 | 0.00266 | 0.033864708 | -2.47813601 | 60S ribosomal protein L5 | elongation zone | 12 h |
| HORVU.MOREX.r3.7HG0674480.1 | 0.265081919 | 6.37117646 | 3.167040472 | 0.00255 | 0.033216773 | -2.47873693 | RING/U-box superfamily protein | elongation zone | 12 h |
| HORVU.MOREX.r3.3HG0238450.1 | 0.463274518 | 6.670539813 | 3.135611734 | 0.0028 | 0.03437844 | -2.48093723 | Protein FRIGIDA | elongation zone | 12 h |
| HORVU.MOREX.r3.5HG0462460.1 | 0.807227957 | 3.212594193 | 2.913253284 | 0.00523 | 0.045731286 | -2.48180584 | complex 1 protein, LYR family protein | elongation zone | 12 h |
| HORVU.MOREX.r3.6HG0558880.1 | 1.224520591 | 5.027918082 | 2.978774161 | 0.00436 | 0.041480063 | -2.48410224 | Histone H2A | elongation zone | 12 h |
| HORVU.MOREX.r3.3HG0271870.2 | 1.224268008 | 3.842763992 | 2.871090281 | 0.00587 | 0.048231861 | -2.48448462 | CTP synthase | elongation zone | 12 h |
| HORVU.MOREX.r3.6HG0599090.1 | -0.825820153 | -2.428665549 | -2.849934744 | 0.00622 | 0.049681395 | -2.48953778 | Vacuolar iron transporter | elongation zone | 12 h |
| HORVU.MOREX.r3.2HG0163890.1 | 0.693978992 | 3.526119037 | 2.892895735 | 0.00553 | 0.046871369 | -2.49178048 | Multifunctional CCA protein | elongation zone | 12 h |
| HORVU.MOREX.r3.7HG0736960.1 | 0.63693711 | 3.878659947 | 2.952116093 | 0.00469 | 0.043259525 | -2.49185388 | BTB/POZ domain containing protein, expressed | elongation zone | 12 h |
| HORVU.MOREX.r3.7HG0695960.1 | 0.568643106 | 9.98141728 | 3.274791529 | 0.00187 | 0.028923495 | -2.491919 | 30S ribosomal protein S3 | elongation zone | 12 h |
| HORVU.MOREX.r3.6HG0555630.1 | 1.447928109 | 6.693514895 | 3.027838801 | 0.0038 | 0.039108923 | -2.49604393 | DNA polymerase V | elongation zone | 12 h |
| HORVU.MOREX.r3.3HG0231480.1 | 0.63377134 | 3.882014662 | 2.951866877 | 0.0047 | 0.043259525 | -2.49651388 | Receptor-like kinase | elongation zone | 12 h |
| HORVU.MOREX.r3.7HG0715570.1 | 0.363397787 | 6.466382964 | 3.154795117 | 0.00265 | 0.033831284 | -2.49774843 | Transmembrane protein | elongation zone | 12 h |
| HORVU.MOREX.r3.2HG0176920.1 | 1.216255934 | 3.864016179 | 2.877316766 | 0.00577 | 0.047801075 | -2.49825069 | ATP-dependent RNA helicase DeaD | elongation zone | 12 h |
| HORVU.MOREX.r3.4HG0393640.1 | 0.894147341 | 5.209210225 | 3.005600432 | 0.00404 | 0.040178373 | -2.49940389 | Anaphase-promoting complex subunit 6-like protein | elongation zone | 12 h |
| HORVU.MOREX.r3.6HG0615820.1 | -0.399539888 | 5.121094299 | -3.120830135 | 0.00292 | 0.035234858 | -2.49972963 | Chaperone protein dnaJ | elongation zone | 12 h |
| HORVU.MOREX.r3.3HG0292240.1 | 1.214740143 | 4.324488603 | 2.902665477 | 0.00538 | 0.046332224 | -2.50109105 | Mitochondrial glycoprotein | elongation zone | 12 h |
| HORVU.MOREX.r3.2HG0203510.1 | -0.516836427 | 5.236895283 | -3.118355755 | 0.00294 | 0.035358381 | -2.50238258 | Jasmonate ZIM domain-containing protein | elongation zone | 12 h |
| HORVU.MOREX.r3.6HG0554520.1 | 0.912284876 | 5.131198847 | 2.965424499 | 0.00452 | 0.042347835 | -2.50466 | BRCT domain-containing protein | elongation zone | 12 h |
| HORVU.MOREX.r3.6HG0596980.1 | 0.887438626 | 5.451026715 | 3.005427438 | 0.00405 | 0.040178373 | -2.50544384 | B-block binding subunit of TFIIIC | elongation zone | 12 h |
| HORVU.MOREX.r3.1HG0066210.1 | 0.829811916 | 2.989216297 | 2.863269315 | 0.00599 | 0.048636131 | -2.50561589 | NBS-LRR-like resistance protein | elongation zone | 12 h |
| HORVU.MOREX.r3.6HG0632380.1 | 0.422825523 | 7.961705372 | 3.183654531 | 0.00243 | 0.032618994 | -2.50717336 | Splicing factor 3B subunit 2 | elongation zone | 12 h |
| HORVU.MOREX.r3.1HG0082450.1 | 2.145164155 | 4.566459078 | 2.855909474 | 0.00611 | 0.049244636 | -2.50858999 | Early nodulin-like protein | elongation zone | 12 h |

**Table S5** Continued.

| ID | logFC | AveExpr | t | P.Value | adj.P.Val | B | gene description | root zone | time point |
|---|---|---|---|---|---|---|---|---|---|
| HORVU.MOREX.r3.5HG0430720.1 | 0.967473326 | 4.903886981 | 2.933263619 | 0.00495 | 0.044514543 | -2.50922702 | ATP-dependent DNA helicase 2 subunit KU70 | elongation zone | 12 h |
| HORVU.MOREX.r3.1HG0038170.1 | 1.162106729 | 6.453681313 | 3.054039253 | 0.00353 | 0.037821239 | -2.51014482 | DNA polymerase epsilon, catalytic subunit, putative | elongation zone | 12 h |
| HORVU.MOREX.r3.1HG0078630.1 | 1.160326316 | 4.590043583 | 2.931199118 | 0.00497 | 0.044603947 | -2.51067249 | Single-stranded DNA-binding protein | elongation zone | 12 h |
| HORVU.MOREX.r3.5HG0449030.1 | 1.188222306 | 4.657038603 | 2.910096109 | 0.00527 | 0.045915159 | -2.51084008 | GRF1-interacting factor-like protein | elongation zone | 12 h |
| HORVU.MOREX.r3.6HG0577130.1 | 0.593831568 | 6.344518727 | 3.103852365 | 0.00306 | 0.036088436 | -2.51115888 | RNA-binding family protein, putative | elongation zone | 12 h |
| HORVU.MOREX.r3.4HG0403250.1 | 0.926361383 | 4.254712987 | 2.934934737 | 0.00492 | 0.044439465 | -2.51218378 | 50S ribosomal L32 | elongation zone | 12 h |
| HORVU.MOREX.r3.4HG0399600.1 | 0.503686721 | 5.665530959 | 3.072284202 | 0.00335 | 0.037256101 | -2.51330607 | COP9 signalosome complex subunit 4 | elongation zone | 12 h |
| HORVU.MOREX.r3.1HG0090780.1 | 0.870631822 | 9.407223354 | 3.207320208 | 0.00227 | 0.031692167 | -2.51621324 | Ketol-acid reductoisomerase (NADP(+)) | elongation zone | 12 h |
| HORVU.MOREX.r3.3HG0296490.1 | 0.560713394 | 6.526216847 | 3.138358444 | 0.00277 | 0.03425211 | -2.51786541 | NADH-cytochrome b5 reductase | elongation zone | 12 h |
| HORVU.MOREX.r3.1HG0049980.1 | -0.324181039 | 8.42398264 | -3.318514373 | 0.00164 | 0.028033028 | -2.52028845 | Actin | elongation zone | 12 h |
| HORVU.MOREX.r3.4HG0398610.1 | 2.056786602 | 4.947656787 | 2.860889807 | 0.00603 | 0.048830126 | -2.52095053 | Urb2/Npa2 family protein | elongation zone | 12 h |
| HORVU.MOREX.r3.7HG0724580.2 | 0.737017445 | 3.515564828 | 2.894728912 | 0.0055 | 0.046792558 | -2.5213583 | Zinc finger CCCH domain-containing protein | elongation zone | 12 h |
| HORVU.MOREX.r3.6HG0589920.1 | 0.73371016 | 9.535234774 | 3.231230094 | 0.00212 | 0.030866089 | -2.5225892 | 60S ribosomal protein L6 | elongation zone | 12 h |
| HORVU.MOREX.r3.1HG0085240.1 | 1.031075396 | 5.724080625 | 2.98686426 | 0.00426 | 0.041087503 | -2.52268879 | RuvB-like helicase | elongation zone | 12 h |
| HORVU.MOREX.r3.4HG0411240.1 | 1.795323233 | 4.73256603 | 2.850343999 | 0.00621 | 0.049645466 | -2.52281442 | WD-repeat protein, putative | elongation zone | 12 h |
| HORVU.MOREX.r3.5HG0497770.1 | -0.459784356 | 6.416725076 | -3.237779903 | 0.00208 | 0.030664986 | -2.5251608 | BAG family molecular chaperone regulator-like protein | elongation zone | 12 h |
| HORVU.MOREX.r3.2HG0185840.1 | 0.883363535 | 7.644996465 | 3.148435691 | 0.0027 | 0.033864708 | -2.52643229 | #N/A | elongation zone | 12 h |
| HORVU.MOREX.r3.5HG0474900.1 | 1.364429733 | 6.343269298 | 3.003434193 | 0.00407 | 0.040229538 | -2.52705383 | #N/A | elongation zone | 12 h |
| HORVU.MOREX.r3.7HG0682750.1 | 0.898401815 | 7.225422779 | 3.106527387 | 0.00304 | 0.035941769 | -2.5277776 | 60S ribosomal protein L32 | elongation zone | 12 h |
| HORVU.MOREX.r3.4HG0403070.1 | 0.496770997 | 6.742400409 | 3.136493561 | 0.00279 | 0.034312101 | -2.53094894 | Serine/threonine-protein phosphatase | elongation zone | 12 h |
| HORVU.MOREX.r3.7HG0704060.1 | 0.521273442 | 4.974659306 | 2.98733573 | 0.00426 | 0.041052717 | -2.53140652 | AT hook motif DNA-binding family protein | elongation zone | 12 h |
| HORVU.MOREX.r3.3HG0255580.1 | -0.435316006 | 5.497551901 | -3.173481952 | 0.00251 | 0.03299418 | -2.53224662 | Respiratory burst oxidase, putative | elongation zone | 12 h |
| HORVU.MOREX.r3.2HG0128230.1 | 0.768068763 | 5.522586618 | 3.037743044 | 0.00369 | 0.038671495 | -2.53270341 | Thaumatin | elongation zone | 12 h |
| HORVU.MOREX.r3.2HG0114570.1 | 0.827674359 | 6.160354978 | 3.054562535 | 0.00352 | 0.037808566 | -2.53354735 | Adenylyl cyclase | elongation zone | 12 h |
| HORVU.MOREX.r3.7HG0714900.1 | 0.626634738 | 4.447401071 | 2.980872249 | 0.00433 | 0.041314646 | -2.5347502 | S-acyltransferase | elongation zone | 12 h |
| HORVU.MOREX.r3.2HG0156960.1 | 0.88211348 | 8.772916108 | 3.177027596 | 0.00248 | 0.032834207 | -2.53476277 | 50S ribosomal protein L13 | elongation zone | 12 h |
| HORVU.MOREX.r3.3HG0237990.1 | -0.519217488 | 4.191673583 | -3.087069463 | 0.00321 | 0.036927181 | -2.53705966 | #N/A | elongation zone | 12 h |
| HORVU.MOREX.r3.5HG0510860.1 | 2.057230763 | 6.298847295 | 2.957942621 | 0.00462 | 0.042747009 | -2.53761038 | Kinesin-like protein | elongation zone | 12 h |
| HORVU.MOREX.r3.4HG0407670.1 | 0.594925846 | 5.172481666 | 3.017505872 | 0.00391 | 0.039547618 | -2.53793188 | Pentatricopeptide repeat-containing protein | elongation zone | 12 h |
| HORVU.MOREX.r3.6HG0572330.1 | 0.434142676 | 5.22316415 | 3.055918842 | 0.00351 | 0.037808566 | -2.53859974 | Actin-related family protein | elongation zone | 12 h |
| HORVU.MOREX.r3.3HG0251830.1 | 0.804572214 | 3.41268951 | 2.877723091 | 0.00576 | 0.04776728 | -2.53934545 | Plant regulator RWP-RK | elongation zone | 12 h |
| HORVU.MOREX.r3.3HG0321700.1 | -0.527573174 | 3.776225864 | -2.999480669 | 0.00411 | 0.040307075 | -2.54073008 | FMN-dependent NADPH-azoreductase | elongation zone | 12 h |
| HORVU.MOREX.r3.3HG0293040.1 | 0.482822666 | 5.431413902 | 3.055367421 | 0.00351 | 0.037808566 | -2.54374111 | Purple acid phosphatase | elongation zone | 12 h |
| HORVU.MOREX.r3.1HG0037800.1 | 1.372893762 | 6.016136386 | 2.974136407 | 0.00441 | 0.041748755 | -2.5467904 | 3'-5' exonuclease domain-containing family protein | elongation zone | 12 h |
| HORVU.MOREX.r3.2HG0176660.1 | 0.932480964 | 4.897976898 | 2.945905833 | 0.00478 | 0.043696105 | -2.5478462 | Elongator complex protein 3 | elongation zone | 12 h |
| HORVU.MOREX.r3.2HG0174520.1 | -0.583238854 | 4.527587866 | -3.131859666 | 0.00283 | 0.034584718 | -2.54791335 | Cytochrome P450 protein | elongation zone | 12 h |
| HORVU.MOREX.r3.1HG0085250.1 | 1.38975589 | 4.749802186 | 2.876058981 | 0.00579 | 0.047903389 | -2.54921963 | 60S ribosomal protein l35a | elongation zone | 12 h |
| HORVU.MOREX.r3.6HG0630650.1 | 1.715248133 | 3.863030917 | 2.875339084 | 0.0058 | 0.047905463 | -2.54948814 | Protein DETOXIFICATION | elongation zone | 12 h |
| HORVU.MOREX.r3.5HG0467880.1 | 0.489782479 | 5.367744504 | 3.023454512 | 0.00385 | 0.039284359 | -2.55151975 | DNA-directed RNA polymerase family protein | elongation zone | 12 h |
| HORVU.MOREX.r3.6HG0595860.1 | 0.358663363 | 5.620864894 | 3.081007753 | 0.00327 | 0.036983216 | -2.5519206 | E3 ubiquitin-protein ligase | elongation zone | 12 h |
| HORVU.MOREX.r3.7HG0739490.1 | 0.871371761 | 7.817057502 | 3.151161398 | 0.00267 | 0.033864708 | -2.55310706 | Histone H2B | elongation zone | 12 h |

**Table S5** Continued.

| ID | logFC | AveExpr | t | P.Value | adj.P.Val | B | gene description | root zone | time point |
|---|---|---|---|---|---|---|---|---|---|
| HORVU.MOREX.r3.6HG0620220.1 | 0.793928373 | 5.661795132 | 3.015815649 | 0.00393 | 0.039657644 | -2.55384138 | Isoleucine--tRNA ligase | elongation zone | 12 h |
| HORVU.MOREX.r3.2HG0174160.1 | 0.934592382 | 5.210032118 | 2.959121249 | 0.0046 | 0.042722832 | -2.55629235 | Testis-expressed sequence 10 protein-like protein | elongation zone | 12 h |
| HORVU.MOREX.r3.4HG0333940.1 | 1.435094108 | 6.071660271 | 2.992582286 | 0.00419 | 0.040685115 | -2.55731485 | Transmembrane protein, putative | elongation zone | 12 h |
| HORVU.MOREX.r3.1HG0039910.1 | 0.781508648 | 5.062796176 | 2.968882323 | 0.00448 | 0.042098534 | -2.55900084 | Polyadenylate-binding protein | elongation zone | 12 h |
| HORVU.MOREX.r3.2HG0162900.2 | 0.629737189 | 2.320158811 | 2.868289978 | 0.00591 | 0.048306767 | -2.56257374 | Aldo-keto reductase | elongation zone | 12 h |
| HORVU.MOREX.r3.6HG0607740.1 | 1.295555813 | 5.336911121 | 2.931803234 | 0.00497 | 0.044592291 | -2.56605856 | ATPase family AAA domain-containing protein 3 | elongation zone | 12 h |
| HORVU.MOREX.r3.6HG0597640.1 | -0.597444683 | 3.081428676 | -3.005766988 | 0.00404 | 0.040178373 | -2.56831141 | Nitrilase | elongation zone | 12 h |
| HORVU.MOREX.r3.7HG0636740.1 | -0.809303603 | 0.502771414 | -2.84740516 | 0.00626 | 0.049810831 | -2.56870925 | NBS-LRR-like resistance protein | elongation zone | 12 h |
| HORVU.MOREX.r3.6HG0598750.1 | -0.893142952 | 0.858146068 | -3.163780312 | 0.00258 | 0.033395092 | -2.56975004 | Remorin | elongation zone | 12 h |
| HORVU.MOREX.r3.7HG0723670.1 | 0.82816187 | 3.921322603 | 2.879740743 | 0.00573 | 0.047754823 | -2.5700048 | Mitochondrial 39S ribosomal protein L53 | elongation zone | 12 h |
| HORVU.MOREX.r3.2HG0207720.1 | -0.657276104 | 0.894336802 | -3.029493704 | 0.00378 | 0.039090877 | -2.57144163 | Flavin-containing monooxygenase | elongation zone | 12 h |
| HORVU.MOREX.r3.3HG0313340.1 | 0.491306581 | 4.971353448 | 3.012544395 | 0.00397 | 0.039847155 | -2.57345802 | nuclear factor Y, subunit B13 | elongation zone | 12 h |
| HORVU.MOREX.r3.2HG0139050.1 | 1.014867435 | 3.432927884 | 2.862648841 | 0.006 | 0.048692565 | -2.57363638 | Protein kinase | elongation zone | 12 h |
| HORVU.MOREX.r3.2HG0143570.1 | 0.773965363 | 5.102737064 | 2.969783222 | 0.00447 | 0.042044934 | -2.57412737 | Asparagine-tRNA ligase | elongation zone | 12 h |
| HORVU.MOREX.r3.5HG0528940.1 | 0.395096768 | 5.397306806 | 3.054655357 | 0.00352 | 0.037808566 | -2.57456727 | GPI-anchored adhesin-like protein | elongation zone | 12 h |
| HORVU.MOREX.r3.6HG0569990.1 | -0.284743152 | 6.145643496 | -3.1572907 | 0.00263 | 0.033716726 | -2.57686103 | Calmodulin binding protein | elongation zone | 12 h |
| HORVU.MOREX.r3.1HG0091360.1 | 0.440320086 | 6.214276876 | 3.104439543 | 0.00306 | 0.036052722 | -2.57800134 | Delta(14)-sterol reductase | elongation zone | 12 h |
| HORVU.MOREX.r3.4HG0393260.1 | 0.332632262 | 7.041152087 | 3.110331701 | 0.00301 | 0.035656113 | -2.57896832 | Oxidoreductase, 2OG-Fe(II) oxygenase family protein | elongation zone | 12 h |
| HORVU.MOREX.r3.3HG0280160.1 | -0.40439279 | 4.904831015 | -3.115950296 | 0.00296 | 0.035453096 | -2.57981466 | Evolutionarily conserved C-terminal region 2 | elongation zone | 12 h |
| HORVU.MOREX.r3.4HG0415600.1 | 1.370804841 | 5.888146948 | 2.992947168 | 0.00419 | 0.040685115 | -2.58075247 | HXXXD-type acyl-transferase family protein | elongation zone | 12 h |
| HORVU.MOREX.r3.1HG0030670.1 | 0.800379012 | 8.049604649 | 3.133403642 | 0.00281 | 0.034551409 | -2.582793 | 50S ribosomal protein L5 | elongation zone | 12 h |
| HORVU.MOREX.r3.5HG0491040.1 | 0.731377343 | 3.434830866 | 2.875556933 | 0.0058 | 0.047903389 | -2.58593307 | DNA-directed RNA polymerase | elongation zone | 12 h |
| HORVU.MOREX.r3.3HG0309410.1 | 0.919093153 | 3.365237867 | 2.849223818 | 0.00623 | 0.049700096 | -2.58673133 | Subtilisin-like protease | elongation zone | 12 h |
| HORVU.MOREX.r3.4HG0411040.1 | 0.446605954 | 5.23830385 | 3.022041123 | 0.00386 | 0.039354132 | -2.58779872 | Chaperone protein dnaJ | elongation zone | 12 h |
| HORVU.MOREX.r3.7HG0744870.1 | 0.617379516 | 4.001055586 | 2.903288225 | 0.00537 | 0.046274876 | -2.58920119 | Mis12 protein | elongation zone | 12 h |
| HORVU.MOREX.r3.6HG0588960.1 | 0.93167083 | 4.77862855 | 2.930662085 | 0.00498 | 0.044603947 | -2.58943701 | Small nuclear ribonucleoprotein Sm D1 | elongation zone | 12 h |
| HORVU.MOREX.r3.5HG0430740.1 | 0.528396134 | 5.807467475 | 3.030891847 | 0.00377 | 0.03899655 | -2.59176146 | #N/A | elongation zone | 12 h |
| HORVU.MOREX.r3.1HG0010370.1 | 0.844421371 | 6.023951457 | 3.013362326 | 0.00396 | 0.03979475 | -2.59343728 | Nuclear pore complex protein Nup107 | elongation zone | 12 h |
| HORVU.MOREX.r3.7HG0711940.2 | 0.527083598 | 2.784849309 | 2.917503675 | 0.00517 | 0.045490951 | -2.59452348 | Aminotransferase | elongation zone | 12 h |
| HORVU.MOREX.r3.5HG0437670.1 | 0.645420414 | 4.311696419 | 2.895541991 | 0.00549 | 0.046758121 | -2.59763662 | Histone deacetylase | elongation zone | 12 h |
| HORVU.MOREX.r3.4HG0395910.1 | 0.766599059 | 6.108491299 | 3.030826544 | 0.00377 | 0.03899655 | -2.59855587 | Glucan endo-1,3-beta-glucosidase 3 | elongation zone | 12 h |
| HORVU.MOREX.r3.1HG0023650.1 | 0.676192423 | 5.863281229 | 3.027306938 | 0.0038 | 0.039110161 | -2.59980577 | Small nuclear ribonucleoprotein | elongation zone | 12 h |
| HORVU.MOREX.r3.7HG0686530.1 | 0.509591542 | 5.623092015 | 3.02626003 | 0.00382 | 0.039184772 | -2.6008783 | La protein like | elongation zone | 12 h |
| HORVU.MOREX.r3.1HG0013050.1 | 1.158848842 | 6.187182499 | 2.971042141 | 0.00445 | 0.041967805 | -2.60109319 | Ribosome biogenesis regulatory protein-like protein | elongation zone | 12 h |
| HORVU.MOREX.r3.5HG0444030.1 | 1.774767835 | 6.014101665 | 2.9450132 | 0.00479 | 0.043697279 | -2.60301465 | Cellulose synthase-like protein | elongation zone | 12 h |
| HORVU.MOREX.r3.3HG0219160.1 | 0.539703859 | 7.250382019 | 3.119673686 | 0.00293 | 0.035308123 | -2.60542186 | Protein decapping 5 | elongation zone | 12 h |
| HORVU.MOREX.r3.2HG0140960.1 | 0.814068008 | 4.296824079 | 2.910840436 | 0.00526 | 0.045865572 | -2.60609167 | Condensin-2 complex subunit H2 | elongation zone | 12 h |
| HORVU.MOREX.r3.6HG0543800.1 | 0.70438517 | 7.420452333 | 3.138457012 | 0.00277 | 0.03425211 | -2.60638832 | Histone H2A | elongation zone | 12 h |
| HORVU.MOREX.r3.6HG0628850.1 | 0.819008686 | 8.487070812 | 3.14990647 | 0.00268 | 0.033864708 | -2.60700111 | Ribosomal protein L37 | elongation zone | 12 h |
| HORVU.MOREX.r3.1HG0065860.1 | -0.555639358 | 4.236426412 | -3.082397634 | 0.00326 | 0.036983216 | -2.60835361 | MLO-like protein | elongation zone | 12 h |
| HORVU.MOREX.r3.5HG0466770.1 | 0.671678036 | 8.962400466 | 3.178180437 | 0.00247 | 0.032834207 | -2.60994875 | Ribosomal protein L15 | elongation zone | 12 h |

**Table S5** Continued.

| ID | logFC | AveExpr | t | P.Value | adj.P.Val | B | gene description | root zone | time point |
|---|---|---|---|---|---|---|---|---|---|
| HORVU.MOREX.r3.4HG0383530.3 | -0.856017228 | 1.457398746 | -3.037381313 | 0.0037 | 0.038671495 | -2.6135996 | Pathogenesis-related protein 1 | elongation zone | 12 h |
| HORVU.MOREX.r3.2HG0192000.1 | -0.755359295 | 0.357063756 | -3.062402032 | 0.00345 | 0.03764455 | -2.61529299 | HXXXD-type acyl-transferase family protein, putative | elongation zone | 12 h |
| HORVU.MOREX.r3.1HG0042740.1 | 0.29319926 | 6.415284526 | 3.11635128 | 0.00295 | 0.035453096 | -2.61530071 | Adaptin ear-binding coat-associated protein, putative | elongation zone | 12 h |
| HORVU.MOREX.r3.3HG0275530.1 | 0.710212482 | 3.985890373 | 2.89585475 | 0.00548 | 0.046755804 | -2.61602734 | Sodium Bile acid symporter family | elongation zone | 12 h |
| HORVU.MOREX.r3.3HG0304660.1 | -0.456799472 | 3.802909764 | -3.078122101 | 0.00329 | 0.037068643 | -2.61692531 | Ceramide glucosyltransferase, putative | elongation zone | 12 h |
| HORVU.MOREX.r3.6HG0541810.1 | 1.017732188 | 4.305459183 | 2.884341613 | 0.00566 | 0.047404516 | -2.61993918 | 28S ribosomal S34, mitochondrial | elongation zone | 12 h |
| HORVU.MOREX.r3.5HG0428100.1 | 0.644473392 | 7.163221856 | 3.08198573 | 0.00326 | 0.036983216 | -2.62115755 | Auxin response factor | elongation zone | 12 h |
| HORVU.MOREX.r3.4HG0398900.1 | 0.868716167 | 8.042634537 | 3.137842889 | 0.00278 | 0.034261617 | -2.62133383 | Histone H2A | elongation zone | 12 h |
| HORVU.MOREX.r3.1HG0083440.1 | -0.379636324 | 5.222068752 | -3.115255062 | 0.00296 | 0.035453096 | -2.62261466 | Lactation elevated protein 1 | elongation zone | 12 h |
| HORVU.MOREX.r3.7HG0683780.1 | 0.741662068 | 7.910611567 | 3.112682452 | 0.00299 | 0.035540654 | -2.62457481 | 50S ribosomal protein L22 | elongation zone | 12 h |
| HORVU.MOREX.r3.7HG0726790.1 | 0.314734579 | 7.796481048 | 3.161136136 | 0.0026 | 0.033587945 | -2.62464242 | Dolichyl-diphosphooligosaccharide--protein glycosyltransferase | elongation zone | 12 h |
| HORVU.MOREX.r3.1HG0001410.1 | 1.361470383 | 3.89097838 | 2.86433404 | 0.00598 | 0.048603777 | -2.62934798 | Transmembrane protein, putative (DUF247) | elongation zone | 12 h |
| HORVU.MOREX.r3.2HG0164150.1 | 0.63614831 | 4.261815439 | 2.920712838 | 0.00512 | 0.045330047 | -2.63047286 | Protein TSSC4 | elongation zone | 12 h |
| HORVU.MOREX.r3.5HG0437610.1 | 0.840850922 | 5.573525032 | 2.983436635 | 0.0043 | 0.041269326 | -2.63480199 | caspase-6 protein | elongation zone | 12 h |
| HORVU.MOREX.r3.4HG0375240.1 | 0.833639464 | 5.249290851 | 2.975616871 | 0.0044 | 0.041634412 | -2.63482154 | Acyl carrier protein | elongation zone | 12 h |
| HORVU.MOREX.r3.1HG0014410.1 | 0.607903059 | 5.218845032 | 2.998358292 | 0.00413 | 0.040356839 | -2.63499285 | Receptor-like kinase | elongation zone | 12 h |
| HORVU.MOREX.r3.1HG0032200.1 | -0.546035013 | 3.740470542 | -3.056109241 | 0.00351 | 0.037808566 | -2.63514346 | S-type anion channel | elongation zone | 12 h |
| HORVU.MOREX.r3.7HG0710600.1 | 0.537689638 | 5.301457053 | 3.003432806 | 0.00407 | 0.040229538 | -2.63571642 | Protein kinase | elongation zone | 12 h |
| HORVU.MOREX.r3.4HG0353330.1 | 0.891525709 | 5.045545148 | 2.944322701 | 0.0048 | 0.043746094 | -2.6371162 | Protein kinase | elongation zone | 12 h |
| HORVU.MOREX.r3.5HG0430460.1 | -0.505765954 | 1.634629182 | -3.046005791 | 0.00361 | 0.03830948 | -2.63775877 | Proline-rich family protein | elongation zone | 12 h |
| HORVU.MOREX.r3.4HG0413410.1 | 0.886347356 | 7.171547875 | 3.115850778 | 0.00296 | 0.035453096 | -2.63847456 | Hydroxymethylglutaryl-CoA synthase | elongation zone | 12 h |
| HORVU.MOREX.r3.1HG0052080.1 | 0.364882537 | 6.314334235 | 3.073776721 | 0.00334 | 0.037244624 | -2.63932527 | C-terminal binding protein | elongation zone | 12 h |
| HORVU.MOREX.r3.1HG0041180.1 | 0.791541723 | 8.516186529 | 3.137318215 | 0.00278 | 0.034272194 | -2.64225357 | 50S ribosomal protein L14 | elongation zone | 12 h |
| HORVU.MOREX.r3.3HG0259660.1 | 0.606329168 | 3.912666018 | 2.882188609 | 0.00569 | 0.047513562 | -2.64619201 | CRS2-associated factor 1, chloroplastic | elongation zone | 12 h |
| HORVU.MOREX.r3.4HG0411740.1 | 1.238878192 | 6.110179775 | 2.949177637 | 0.00473 | 0.043485864 | -2.64816112 | ATP-dependent RNA helicase | elongation zone | 12 h |
| HORVU.MOREX.r3.3HG0239980.1 | -0.999714079 | 1.292153679 | -2.961761134 | 0.00457 | 0.042545213 | -2.64868342 | Alpha/beta-Hydrolases superfamily protein, putative | elongation zone | 12 h |
| HORVU.MOREX.r3.1HG0094520.1 | 0.528825462 | 7.217789849 | 3.083724899 | 0.00324 | 0.036928455 | -2.64870873 | DNA damage-binding protein 1 | elongation zone | 12 h |
| HORVU.MOREX.r3.5HG0455100.1 | 0.9100108 | 9.647836002 | 3.169585512 | 0.00254 | 0.033099534 | -2.65083467 | Ribosomal protein L3 | elongation zone | 12 h |
| HORVU.MOREX.r3.7HG0635550.1 | 1.291484076 | 6.336423435 | 3.001542256 | 0.00409 | 0.040265883 | -2.65222601 | Lipoxygenase domain-containing 1 | elongation zone | 12 h |
| HORVU.MOREX.r3.5HG0492480.1 | 0.742684496 | 5.7458778 | 3.012172661 | 0.00397 | 0.039849721 | -2.65314529 | 30S ribosomal protein S16 | elongation zone | 12 h |
| HORVU.MOREX.r3.7HG0704330.1 | 0.748307947 | 8.747152339 | 3.142950533 | 0.00274 | 0.034069783 | -2.65327898 | 40S ribosomal protein S13 | elongation zone | 12 h |
| HORVU.MOREX.r3.7HG0730720.1 | 0.917177613 | 6.181638114 | 2.981441822 | 0.00433 | 0.041306621 | -2.65330149 | RNA polymerase I-specific transcription initiation factor RRN3 | elongation zone | 12 h |
| HORVU.MOREX.r3.3HG0218440.1 | 0.592758563 | 5.530527629 | 2.978043685 | 0.00437 | 0.041506896 | -2.65368864 | Serine/threonine-protein kinase ATM | elongation zone | 12 h |
| HORVU.MOREX.r3.5HG0533630.1 | 1.171668092 | 5.193357564 | 3.063566969 | 0.00343 | 0.037620683 | -2.65402163 | Expansin | elongation zone | 12 h |
| HORVU.MOREX.r3.1HG0024280.1 | 0.620755139 | 7.969902425 | 3.147187425 | 0.0027 | 0.033864708 | -2.65476049 | Dihydrolipoyl dehydrogenase | elongation zone | 12 h |
| HORVU.MOREX.r3.6HG0558580.1 | 1.007775594 | 4.169275691 | 2.849021981 | 0.00623 | 0.049708047 | -2.65560573 | Ribosomal protein L37, mitochondrial | elongation zone | 12 h |
| HORVU.MOREX.r3.1HG0074490.1 | 1.172620623 | 4.901731214 | 2.904213313 | 0.00536 | 0.046251967 | -2.65975686 | #N/A | elongation zone | 12 h |
| HORVU.MOREX.r3.2HG0099160.1 | -0.955595031 | -0.22735804 | -2.931106774 | 0.00497 | 0.044603947 | -2.66084311 | 2-oxoglutarate (2OG) and Fe(II)-dependent oxygenase superfami | elongation zone | 12 h |
| HORVU.MOREX.r3.6HG0623070.1 | -0.424527683 | 3.799863701 | -3.016576923 | 0.00392 | 0.039611908 | -2.66186463 | Ankyrin repeat-containing protein, putative | elongation zone | 12 h |
| HORVU.MOREX.r3.3HG0314820.1 | -0.614023021 | 3.291171778 | -3.072193846 | 0.00335 | 0.037256101 | -2.66272343 | 26S proteasome regulatory subunit RPN2 | elongation zone | 12 h |
| HORVU.MOREX.r3.3HG0246810.1 | 0.623290955 | 5.554940352 | 2.999629571 | 0.00411 | 0.040307075 | -2.66286539 | 30S ribosomal protein S6 | elongation zone | 12 h |

**Table S5** Continued.

| ID | logFC | AveExpr | t | P.Value | adj.P.Val | B | gene description | root zone | time point |
|---|---|---|---|---|---|---|---|---|---|
| HORVU.MOREX.r3.7HG0696930.2 | 1.782333554 | 6.181980146 | 2.896413332 | 0.00547 | 0.046711054 | -2.66386514 | RNA-binding protein | elongation zone | 12 h |
| HORVU.MOREX.r3.4HG0400040.1 | -0.48670251 | 1.083905502 | -3.072930384 | 0.00334 | 0.037254747 | -2.66399436 | Cytochrome b561 and DOMON domain-containing protein | elongation zone | 12 h |
| HORVU.MOREX.r3.2HG0173380.1 | -0.415020896 | 4.283255367 | -3.058004798 | 0.00349 | 0.037779464 | -2.66463043 | Amino acid transporter, putative | elongation zone | 12 h |
| HORVU.MOREX.r3.3HG0287070.1 | 0.888676954 | 6.192472525 | 3.074320749 | 0.00333 | 0.037226047 | -2.66572486 | Glycosyltransferase | elongation zone | 12 h |
| HORVU.MOREX.r3.3HG0269430.1 | 0.534839276 | 6.166272467 | 3.011302328 | 0.00398 | 0.03986915 | -2.66850694 | Chaperone protein dnaJ | elongation zone | 12 h |
| HORVU.MOREX.r3.3HG0309800.1 | 0.630885253 | 5.007429359 | 2.977295509 | 0.00438 | 0.041519226 | -2.67027326 | Small ubiquitin-related modifier | elongation zone | 12 h |
| HORVU.MOREX.r3.1HG0068810.1 | -0.347513093 | 4.949596997 | -3.103690073 | 0.00306 | 0.036088436 | -2.67212607 | #N/A | elongation zone | 12 h |
| HORVU.MOREX.r3.4HG0352780.1 | -0.652795797 | 1.138744691 | -2.963535589 | 0.00455 | 0.04241279 | -2.67358106 | WRKY transcription factor | elongation zone | 12 h |
| HORVU.MOREX.r3.3HG0298790.1 | 0.590235249 | 4.931630051 | 2.917076475 | 0.00517 | 0.045490951 | -2.67376613 | Amidophosphoribosyltransferase | elongation zone | 12 h |
| HORVU.MOREX.r3.6HG0543740.1 | 0.770980734 | 7.842284834 | 3.124700493 | 0.00289 | 0.035113249 | -2.67523405 | Histone H2A | elongation zone | 12 h |
| HORVU.MOREX.r3.7HG0692740.1 | 0.347805226 | 6.659164505 | 3.066580903 | 0.0034 | 0.037432489 | -2.67614653 | RNA-binding protein 25 | elongation zone | 12 h |
| HORVU.MOREX.r3.2HG0187940.1 | -0.618914677 | 5.04269045 | -3.064987882 | 0.00342 | 0.037489533 | -2.67769091 | Niemann-Pick C1 protein | elongation zone | 12 h |
| HORVU.MOREX.r3.3HG0298750.1 | 0.48894195 | 5.760333104 | 3.056461457 | 0.0035 | 0.037808566 | -2.67838938 | magnesium transporter NIPA (DUF803) | elongation zone | 12 h |
| HORVU.MOREX.r3.1HG0015470.1 | -0.867341334 | 2.791232652 | -2.983058802 | 0.00431 | 0.041274283 | -2.67941499 | 2-oxoglutarate (2OG) and Fe(II)-dependent oxygenase superfami | elongation zone | 12 h |
| HORVU.MOREX.r3.2HG0183390.1 | -0.673212153 | 2.00100842 | -3.002381104 | 0.00408 | 0.040250079 | -2.68012193 | GRAM domain-containing protein / ABA-responsive | elongation zone | 12 h |
| HORVU.MOREX.r3.2HG0196630.1 | -0.938337318 | -0.811111303 | -2.901830233 | 0.00539 | 0.046372512 | -2.68097371 | Invertase/pectin methylesterase inhibitor family protein | elongation zone | 12 h |
| HORVU.MOREX.r3.3HG0307310.1 | 0.278875991 | 5.321616586 | 3.059287423 | 0.00348 | 0.037764638 | -2.68296916 | magnesium transporter, putative (DUF803) | elongation zone | 12 h |
| HORVU.MOREX.r3.6HG0555870.1 | 1.723763945 | 5.140784525 | 2.851936445 | 0.00618 | 0.049512564 | -2.68350568 | Kinesin-like protein | elongation zone | 12 h |
| HORVU.MOREX.r3.5HG0505300.1 | 1.491501307 | 6.157248139 | 2.921000553 | 0.00512 | 0.045330047 | -2.68459906 | Ribosome biogenesis protein BOP1 homolog | elongation zone | 12 h |
| HORVU.MOREX.r3.4HG0398910.1 | 0.904080276 | 7.143679884 | 3.068027622 | 0.00339 | 0.037367493 | -2.68541389 | Histone H2B | elongation zone | 12 h |
| HORVU.MOREX.r3.1HG0094770.1 | 0.516483447 | 3.670278309 | 2.886791844 | 0.00562 | 0.04722308 | -2.6868407 | Transmembrane protein, putative | elongation zone | 12 h |
| HORVU.MOREX.r3.6HG0610110.1 | 0.886783337 | 4.84135302 | 2.876947955 | 0.00577 | 0.047810689 | -2.68932737 | FKBP-like peptidyl-prolyl cis-trans isomerase family protein | elongation zone | 12 h |
| HORVU.MOREX.r3.5HG0529180.1 | 0.584180011 | 6.543237435 | 3.053010829 | 0.00354 | 0.037872249 | -2.68941016 | Huntingtin interacting protein K | elongation zone | 12 h |
| HORVU.MOREX.r3.2HG0192180.1 | 0.195825764 | 9.138579 | 3.212890053 | 0.00224 | 0.031520621 | -2.69027996 | 26S proteasome non-ATPase regulatory subunit 1 | elongation zone | 12 h |
| HORVU.MOREX.r3.3HG0313090.1 | -0.684053204 | 2.944420659 | -2.92156782 | 0.00511 | 0.045296428 | -2.69108713 | Thiamin pyrophosphokinase 1 | elongation zone | 12 h |
| HORVU.MOREX.r3.2HG0204570.1 | -0.458359572 | 5.322525667 | -3.120902231 | 0.00292 | 0.035234858 | -2.69305496 | glucuronoxylan 4-O-methyltransferase-like protein (DUF579) | elongation zone | 12 h |
| HORVU.MOREX.r3.2HG0124660.1 | -0.303025569 | 4.762294428 | -3.027855229 | 0.0038 | 0.039108923 | -2.69320797 | Dihydropteroate synthase, putative | elongation zone | 12 h |
| HORVU.MOREX.r3.7HG0730070.1 | 1.046749275 | 6.723991887 | 3.030940021 | 0.00377 | 0.03899655 | -2.69380606 | Thaumatin-like protein | elongation zone | 12 h |
| HORVU.MOREX.r3.4HG0340230.1 | 0.651971583 | 6.673601725 | 3.032725178 | 0.00375 | 0.038936762 | -2.69417171 | Alba DNA/RNA-binding protein | elongation zone | 12 h |
| HORVU.MOREX.r3.7HG0676250.1 | 0.778801625 | 3.644955906 | 2.896404078 | 0.00547 | 0.046711054 | -2.69442031 | Protein DETOXIFICATION | elongation zone | 12 h |
| HORVU.MOREX.r3.1HG0029460.1 | -0.476249944 | 2.800322038 | -2.893680663 | 0.00551 | 0.046829376 | -2.69628304 | Fatty acid oxidation complex subunit alpha | elongation zone | 12 h |
| HORVU.MOREX.r3.1HG0091240.1 | -0.463161552 | 4.910103678 | -3.086793841 | 0.00321 | 0.036928455 | -2.69966775 | Mitogen-activated protein kinase | elongation zone | 12 h |
| HORVU.MOREX.r3.6HG0630200.1 | 0.358235467 | 6.20685434 | 3.036734589 | 0.0037 | 0.038704216 | -2.70390934 | Protein FAR1-RELATED SEQUENCE 3 | elongation zone | 12 h |
| HORVU.MOREX.r3.3HG0240180.1 | 0.444765694 | 6.812791234 | 3.054971105 | 0.00352 | 0.037808566 | -2.70448689 | Ubiquitinyl hydrolase 1 | elongation zone | 12 h |
| HORVU.MOREX.r3.2HG0134400.1 | -0.329250881 | 6.098222844 | -3.078627516 | 0.00329 | 0.037035763 | -2.70627825 | Protein DETOXIFICATION | elongation zone | 12 h |
| HORVU.MOREX.r3.6HG0546130.1 | 0.553661511 | 4.48574317 | 2.920817329 | 0.00512 | 0.045330047 | -2.70753992 | BTB/POZ domain-containing protein | elongation zone | 12 h |
| HORVU.MOREX.r3.4HG0387120.1 | 0.964614023 | 6.92928788 | 3.0310846 | 0.00376 | 0.03899655 | -2.70993498 | Histone H2A | elongation zone | 12 h |
| HORVU.MOREX.r3.7HG0654170.1 | 1.123031841 | 4.7582708 | 2.849462314 | 0.00622 | 0.049700096 | -2.71162339 | rRNA-processing protein FCF1-like protein | elongation zone | 12 h |
| HORVU.MOREX.r3.3HG0252610.1 | 0.329636418 | 7.233438842 | 3.09136114 | 0.00317 | 0.036725735 | -2.712318 | Protein kinase | elongation zone | 12 h |
| HORVU.MOREX.r3.4HG0418290.1 | -0.24042844 | 6.480281927 | -3.107563214 | 0.00303 | 0.035897692 | -2.71720742 | Charged multivesicular body protein 3 | elongation zone | 12 h |
| HORVU.MOREX.r3.5HG0447330.1 | 1.010880355 | 4.611100684 | 2.941610925 | 0.00483 | 0.043896282 | -2.71835592 | Aquaporin | elongation zone | 12 h |

**Table S5** Continued.

| ID | logFC | AveExpr | t | P.Value | adj.P.Val | B | gene description | root zone | time point |
|---|---|---|---|---|---|---|---|---|---|
| HORVU.MOREX.r3.7HG0640580.1 | 0.765500507 | 6.013252126 | 2.945344079 | 0.00478 | 0.043696105 | -2.71879948 | Phosphoglucan, water dikinase | elongation zone | 12 h |
| HORVU.MOREX.r3.5HG0471150.1 | -0.514698599 | 3.92040144 | -3.045745789 | 0.00361 | 0.038317796 | -2.71914599 | Cobyric acid synthase | elongation zone | 12 h |
| HORVU.MOREX.r3.2HG0170160.1 | 0.28568758 | 8.212366788 | 3.160150504 | 0.00261 | 0.033615353 | -2.71943818 | Cleft lip and palate transmembrane 1 | elongation zone | 12 h |
| HORVU.MOREX.r3.7HG0721930.1 | 0.507493414 | 4.955957757 | 2.947978032 | 0.00475 | 0.043552747 | -2.71963443 | B-cell receptor-associated protein 31-like containing protein | elongation zone | 12 h |
| HORVU.MOREX.r3.1HG0089160.1 | 0.617887456 | 7.967866402 | 3.083992217 | 0.00324 | 0.036928455 | -2.72760932 | Eukaryotic translation initiation factor 3 subunit D | elongation zone | 12 h |
| HORVU.MOREX.r3.6HG0568910.1 | -0.626591428 | 3.99891951 | -3.085929937 | 0.00322 | 0.036928455 | -2.73070601 | L-gulonolactone oxidase | elongation zone | 12 h |
| HORVU.MOREX.r3.2HG0178420.1 | 1.726238559 | 5.745692204 | 2.880314042 | 0.00572 | 0.04769973 | -2.73398729 | transmembrane protein | elongation zone | 12 h |
| HORVU.MOREX.r3.6HG0548420.1 | 1.193184457 | 4.579374657 | 2.852337184 | 0.00617 | 0.049512065 | -2.73431295 | Histone H4 | elongation zone | 12 h |
| HORVU.MOREX.r3.1HG0032650.1 | -0.40982402 | 2.965330327 | -3.019497226 | 0.00389 | 0.039463094 | -2.73537803 | #N/A | elongation zone | 12 h |
| HORVU.MOREX.r3.7HG0743900.1 | -0.498606898 | 5.143712272 | -3.080568147 | 0.00327 | 0.036983216 | -2.73708154 | Kinase family protein | elongation zone | 12 h |
| HORVU.MOREX.r3.5HG0460120.1 | -0.494044967 | 1.893832399 | -2.907278603 | 0.00531 | 0.04613544 | -2.74261283 | Alpha/beta-Hydrolases superfamily protein | elongation zone | 12 h |
| HORVU.MOREX.r3.1HG0022680.1 | 0.844078 | 6.559679477 | 2.984866514 | 0.00428 | 0.041182136 | -2.74409913 | Histidine--tRNA ligase | elongation zone | 12 h |
| HORVU.MOREX.r3.5HG0487950.1 | -0.406072418 | 5.865567848 | -3.105693336 | 0.00305 | 0.035985971 | -2.74602874 | Purine permease-like protein | elongation zone | 12 h |
| HORVU.MOREX.r3.7HG0746810.1 | 0.32669202 | 5.647823023 | 3.028303405 | 0.00379 | 0.039108923 | -2.74657248 | Transcription factor | elongation zone | 12 h |
| HORVU.MOREX.r3.6HG0545460.1 | -0.544741871 | 3.011798115 | -3.043076424 | 0.00364 | 0.038508115 | -2.74753612 | Transmembrane protein 97 | elongation zone | 12 h |
| HORVU.MOREX.r3.5HG0508600.1 | 0.727568582 | 5.919054895 | 2.941300718 | 0.00484 | 0.043914566 | -2.74904692 | WD-40 repeat protein | elongation zone | 12 h |
| HORVU.MOREX.r3.5HG0420410.1 | -0.384877321 | 5.049102333 | -3.037237122 | 0.0037 | 0.038671495 | -2.75048915 | Ubiquitin-conjugating enzyme E2 | elongation zone | 12 h |
| HORVU.MOREX.r3.4HG0355430.1 | 1.2985144 | 5.48935292 | 2.858595008 | 0.00607 | 0.049059031 | -2.75704231 | ATP-dependent RNA helicase | elongation zone | 12 h |
| HORVU.MOREX.r3.3HG0298330.1 | -0.457575836 | 4.860908662 | -3.022419993 | 0.00386 | 0.039334712 | -2.75716631 | Chaperone protein dnaJ | elongation zone | 12 h |
| HORVU.MOREX.r3.7HG0707860.1 | -0.464131508 | 3.600785372 | -2.970644465 | 0.00446 | 0.041982723 | -2.75828223 | CASP-like protein | elongation zone | 12 h |
| HORVU.MOREX.r3.3HG0254940.1 | -0.759562086 | 2.451212431 | -3.113472109 | 0.00298 | 0.035522336 | -2.7590522 | Peroxidase | elongation zone | 12 h |
| HORVU.MOREX.r3.2HG0176290.1 | 0.407436024 | 6.030618554 | 2.997929852 | 0.00413 | 0.040386019 | -2.76005131 | Mitochondrial inner membrane protease ATP23 | elongation zone | 12 h |
| HORVU.MOREX.r3.4HG0414480.1 | 0.728552235 | 4.891800711 | 2.903822522 | 0.00536 | 0.046274876 | -2.76034111 | Small nuclear ribonucleoprotein-like protein | elongation zone | 12 h |
| HORVU.MOREX.r3.4HG0379960.1 | 0.366579445 | 5.175067919 | 2.982184928 | 0.00432 | 0.04129762 | -2.76061691 | Exostosin family protein | elongation zone | 12 h |
| HORVU.MOREX.r3.7HG0684410.1 | 0.759824335 | 9.154592415 | 3.117807378 | 0.00294 | 0.03539314 | -2.76334386 | 60S ribosomal protein L7a | elongation zone | 12 h |
| HORVU.MOREX.r3.1HG0072520.1 | 0.878179753 | 4.709600772 | 2.916894946 | 0.00517 | 0.045490951 | -2.76760036 | #N/A | elongation zone | 12 h |
| HORVU.MOREX.r3.1HG0007670.1 | -0.592060024 | 3.984670291 | -2.929867974 | 0.00499 | 0.044653737 | -2.77024224 | OCS-element binding factor 5 | elongation zone | 12 h |
| HORVU.MOREX.r3.6HG0630420.1 | 0.342163539 | 4.567048698 | 2.93400715 | 0.00494 | 0.044514442 | -2.77132896 | Myosin type-2 heavy chain 2 | elongation zone | 12 h |
| HORVU.MOREX.r3.7HG0712780.1 | 1.181868359 | 6.591425476 | 2.945384629 | 0.00478 | 0.043696105 | -2.77132952 | Pre-rRNA-processing protein TSR1-like protein | elongation zone | 12 h |
| HORVU.MOREX.r3.2HG0159500.1 | 0.749873777 | 5.303207265 | 2.907324958 | 0.00531 | 0.04613544 | -2.77365647 | tRNA (Adenine-N(1)-)-methyltransferase catalytic subunit trmt61 | elongation zone | 12 h |
| HORVU.MOREX.r3.1HG0044060.2 | 0.744066386 | 4.318494758 | 2.868390034 | 0.00591 | 0.048306767 | -2.7755288 | 28S ribosomal protein S29, mitochondrial | elongation zone | 12 h |
| HORVU.MOREX.r3.6HG0608110.1 | 0.577701005 | 6.16140251 | 2.979385367 | 0.00435 | 0.041428624 | -2.77619538 | Histidinol-phosphate aminotransferase | elongation zone | 12 h |
| HORVU.MOREX.r3.5HG0485990.1 | 0.944825522 | 5.054014433 | 2.865059261 | 0.00596 | 0.048546205 | -2.77665588 | Nuclear pore complex protein NUP58 | elongation zone | 12 h |
| HORVU.MOREX.r3.3HG0299820.1 | 0.872151401 | 7.690345431 | 3.053832854 | 0.00353 | 0.037823562 | -2.77795237 | Histone H3 | elongation zone | 12 h |
| HORVU.MOREX.r3.3HG0310540.1 | 0.490138688 | 5.76487875 | 3.00248009 | 0.00408 | 0.040250079 | -2.77919365 | Mannosyl-oligosaccharide glucosidase | elongation zone | 12 h |
| HORVU.MOREX.r3.2HG0203840.1 | 0.923463679 | 7.328278373 | 3.003886563 | 0.00406 | 0.040229538 | -2.77943118 | Nuclease S1 | elongation zone | 12 h |
| HORVU.MOREX.r3.3HG0303520.1 | 0.742021506 | 8.112661201 | 3.06919691 | 0.00338 | 0.037331432 | -2.77966762 | 30S ribosomal protein S12 | elongation zone | 12 h |
| HORVU.MOREX.r3.3HG0244240.1 | -0.894845174 | 3.630904343 | -2.94211804 | 0.00483 | 0.043873519 | -2.78046385 | Alpha/beta-Hydrolases superfamily protein | elongation zone | 12 h |
| HORVU.MOREX.r3.1HG0006650.1 | 0.739642108 | 8.905369094 | 3.114561294 | 0.00297 | 0.035453096 | -2.78327632 | Histone H2A | elongation zone | 12 h |
| HORVU.MOREX.r3.7HG0674750.1 | 0.341449749 | 5.335659117 | 2.9953152 | 0.00416 | 0.040578362 | -2.78710085 | Dual specificity protein phosphatase, putative | elongation zone | 12 h |
| HORVU.MOREX.r3.5HG0463410.1 | 0.837641248 | 7.370416612 | 3.024667286 | 0.00383 | 0.03925602 | -2.79141806 | 30S ribosomal protein S9 | elongation zone | 12 h |

**Table S5** Continued.

| ID | logFC | AveExpr | t | P.Value | adj.P.Val | B | gene description | root zone | time point |
|---|---|---|---|---|---|---|---|---|---|
| HORVU.MOREX.r3.7HG0667730.1 | -0.376986414 | 5.42923688 | -3.046584842 | 0.0036 | 0.038286492 | -2.79196385 | Chaperone protein dnaJ | elongation zone | 12 h |
| HORVU.MOREX.r3.6HG0619320.1 | 0.524889287 | 6.069339622 | 2.976901156 | 0.00438 | 0.04154313 | -2.79246115 | Prolyl-tRNA synthetase associated domain-containing protein 1 | elongation zone | 12 h |
| HORVU.MOREX.r3.5HG0467810.1 | 0.478186104 | 7.61265314 | 3.056216133 | 0.00351 | 0.037808566 | -2.79372706 | Aspartate--tRNA ligase | elongation zone | 12 h |
| HORVU.MOREX.r3.6HG0550670.1 | 1.788517458 | 7.005257995 | 2.944036869 | 0.0048 | 0.043752599 | -2.79405762 | Glycine-rich cell wall structural protein 2 | elongation zone | 12 h |
| HORVU.MOREX.r3.5HG0521570.1 | 0.641181619 | 5.72311113 | 2.93178869 | 0.00497 | 0.044592291 | -2.79623829 | Cell division cycle 5-like protein | elongation zone | 12 h |
| HORVU.MOREX.r3.6HG0581010.1 | 0.544923574 | 4.497989908 | 2.891438705 | 0.00555 | 0.046935568 | -2.79626834 | Poly(A) polymerase | elongation zone | 12 h |
| HORVU.MOREX.r3.7HG0709680.1 | 0.616348841 | 8.315627784 | 3.084704192 | 0.00323 | 0.036928455 | -2.79657734 | T-complex protein 1 subunit gamma | elongation zone | 12 h |
| HORVU.MOREX.r3.2HG0104980.1 | 0.816566773 | 8.609674328 | 3.074720838 | 0.00333 | 0.037204054 | -2.79803673 | 40S ribosomal protein S6 | elongation zone | 12 h |
| HORVU.MOREX.r3.5HG0420210.1 | 0.361202649 | 6.548287697 | 3.035422698 | 0.00372 | 0.038738812 | -2.79922739 | Plasma membrane ATPase | elongation zone | 12 h |
| HORVU.MOREX.r3.5HG0534180.1 | 0.562587711 | 5.30730743 | 2.915247784 | 0.0052 | 0.045616889 | -2.80082444 | KH domain-containing protein | elongation zone | 12 h |
| HORVU.MOREX.r3.2HG0112970.1 | 0.808747478 | 8.618139404 | 3.079324071 | 0.00328 | 0.036983216 | -2.80254688 | 60S ribosomal protein L18 | elongation zone | 12 h |
| HORVU.MOREX.r3.4HG0340180.1 | 0.921748292 | 7.057162416 | 2.976277005 | 0.00439 | 0.041577055 | -2.80307387 | Transducin/WD-like repeat-protein | elongation zone | 12 h |
| HORVU.MOREX.r3.6HG0610500.1 | -0.769965883 | 1.298618032 | -3.080565787 | 0.00327 | 0.036983216 | -2.80341927 | Eukaryotic aspartyl protease family protein | elongation zone | 12 h |
| HORVU.MOREX.r3.3HG0233150.1 | 0.719573793 | 5.953619348 | 2.971518462 | 0.00445 | 0.04195768 | -2.8085982 | Histone H2B | elongation zone | 12 h |
| HORVU.MOREX.r3.7HG0719170.1 | -0.655002581 | 5.542866148 | -3.083141987 | 0.00325 | 0.036948704 | -2.80895407 | O-fucosyltransferase-like protein | elongation zone | 12 h |
| HORVU.MOREX.r3.1HG0081790.1 | 0.58634546 | 5.419367045 | 2.918117634 | 0.00516 | 0.045451924 | -2.80995113 | Phosphotransferase | elongation zone | 12 h |
| HORVU.MOREX.r3.4HG0389240.1 | 0.460233016 | 4.623069502 | 2.9332577 | 0.00495 | 0.044514543 | -2.81073176 | DUF3527 domain protein | elongation zone | 12 h |
| HORVU.MOREX.r3.3HG0288140.1 | 0.26824346 | 6.173911395 | 3.046146136 | 0.00361 | 0.03830948 | -2.81075513 | Golgin candidate 1 | elongation zone | 12 h |
| HORVU.MOREX.r3.4HG0331980.1 | 0.843644191 | 8.08863791 | 3.051468417 | 0.00355 | 0.037906495 | -2.81200432 | 40S ribosomal protein S25, putative | elongation zone | 12 h |
| HORVU.MOREX.r3.4HG0382500.1 | 0.371141106 | 5.221396135 | 2.96788538 | 0.00449 | 0.042132428 | -2.81358977 | Seed maturation protein PM36, putative, expressed | elongation zone | 12 h |
| HORVU.MOREX.r3.5HG0455740.1 | 0.710395368 | 6.88962885 | 2.986164928 | 0.00427 | 0.041090601 | -2.81382246 | ATP dependent RNA helicase | elongation zone | 12 h |
| HORVU.MOREX.r3.4HG0333130.2 | 0.636394116 | 4.143810939 | 2.84823833 | 0.00624 | 0.049750126 | -2.81483091 | #N/A | elongation zone | 12 h |
| HORVU.MOREX.r3.2HG0214780.1 | 0.642009635 | 7.100366582 | 3.016096865 | 0.00393 | 0.039645835 | -2.81701574 | 3-isopropylmalate dehydrogenase | elongation zone | 12 h |
| HORVU.MOREX.r3.5HG0511480.1 | 0.763476908 | 9.452511158 | 3.111927426 | 0.00299 | 0.035576158 | -2.81738704 | 50S ribosomal protein L13 | elongation zone | 12 h |
| HORVU.MOREX.r3.1HG0085120.1 | 0.832060281 | 6.204576906 | 2.919710843 | 0.00513 | 0.045371176 | -2.81879265 | Asparagine synthetase | elongation zone | 12 h |
| HORVU.MOREX.r3.5HG0470030.1 | 0.41422234 | 7.999201135 | 3.123173952 | 0.0029 | 0.035162818 | -2.81954342 | 3-ketoacyl-CoA thiolase | elongation zone | 12 h |
| HORVU.MOREX.r3.5HG0454500.1 | 1.501025022 | 6.350657597 | 2.87511062 | 0.0058 | 0.0479161 | -2.82015067 | Nucleolar protein 6 | elongation zone | 12 h |
| HORVU.MOREX.r3.1HG0058640.1 | 1.26786368 | 5.976990264 | 2.896805672 | 0.00547 | 0.046711054 | -2.82192373 | Histone H2B | elongation zone | 12 h |
| HORVU.MOREX.r3.3HG0304370.1 | -0.437780749 | 4.367817245 | -2.984626745 | 0.00429 | 0.041182464 | -2.82330507 | Glutamyl-tRNA(Gln) amidotransferase subunit A | elongation zone | 12 h |
| HORVU.MOREX.r3.6HG0626930.1 | 0.961169494 | 4.840472652 | 2.851967879 | 0.00618 | 0.049512564 | -2.82751766 | Ribonucleoside-diphosphate reductase | elongation zone | 12 h |
| HORVU.MOREX.r3.1HG0026070.2 | 0.593610938 | 8.133614113 | 3.057955956 | 0.00349 | 0.037779464 | -2.82951359 | Ribosomal protein | elongation zone | 12 h |
| HORVU.MOREX.r3.3HG0323330.1 | -0.711983369 | 1.397008086 | -2.91170089 | 0.00525 | 0.04584899 | -2.83074051 | RNA binding protein, putative | elongation zone | 12 h |
| HORVU.MOREX.r3.4HG0378160.1 | 0.638140267 | 8.929191593 | 3.084225279 | 0.00324 | 0.036928455 | -2.83579339 | Ribosomal protein L19 | elongation zone | 12 h |
| HORVU.MOREX.r3.6HG0630550.1 | 0.59599659 | 8.38745822 | 3.071862554 | 0.00335 | 0.037263411 | -2.83738558 | T-complex protein 1 subunit zeta | elongation zone | 12 h |
| HORVU.MOREX.r3.3HG0293700.1 | 0.651253647 | 6.606906202 | 2.982670239 | 0.00431 | 0.041282541 | -2.83810272 | Protein downstream neighbor of Son | elongation zone | 12 h |
| HORVU.MOREX.r3.2HG0114370.1 | -0.432365803 | 4.135311302 | -3.003748196 | 0.00406 | 0.040229538 | -2.83966091 | Kinase family protein | elongation zone | 12 h |
| HORVU.MOREX.r3.7HG0656440.1 | 0.459037497 | 5.106986071 | 2.897239242 | 0.00546 | 0.046711054 | -2.83971021 | Adenylyltransferase and sulfurtransferase MOCS3 | elongation zone | 12 h |
| HORVU.MOREX.r3.6HG0600000.1 | -0.95245349 | -0.219047099 | -2.93321235 | 0.00495 | 0.044514543 | -2.84159878 | Blue copper protein | elongation zone | 12 h |
| HORVU.MOREX.r3.3HG0253650.1 | 0.591472893 | 6.25534348 | 2.957531088 | 0.00462 | 0.042776562 | -2.84245131 | WD40 repeat-like protein | elongation zone | 12 h |
| HORVU.MOREX.r3.6HG0606400.1 | 2.034984385 | 6.56878441 | 2.871575686 | 0.00586 | 0.048206582 | -2.84638857 | Replicase polyprotein 1a | elongation zone | 12 h |
| HORVU.MOREX.r3.2HG0208810.1 | 0.276196684 | 6.69596435 | 3.02012744 | 0.00388 | 0.03943226 | -2.84654953 | Ubiquitin-conjugating enzyme E2 | elongation zone | 12 h |

**Table S5** Continued.

| ID | logFC | AveExpr | t | P.Value | adj.P.Val | B | gene description | root zone | time point |
|---|---|---|---|---|---|---|---|---|---|
| HORVU.MOREX.r3.4HG0418460.1 | -0.424970757 | 1.75953737 | -3.005435829 | 0.00405 | 0.040178373 | -2.84670388 | Cysteine protease | elongation zone | 12 h |
| HORVU.MOREX.r3.4HG0416710.1 | 0.554628132 | 7.831417977 | 3.037378067 | 0.0037 | 0.038671495 | -2.84898358 | Alanine--tRNA ligase | elongation zone | 12 h |
| HORVU.MOREX.r3.7HG0695870.1 | 0.712695283 | 8.389980015 | 3.058398033 | 0.00348 | 0.037779464 | -2.85061169 | 60S ribosomal protein L13 | elongation zone | 12 h |
| HORVU.MOREX.r3.5HG0480100.1 | -0.34122956 | 8.360963836 | -3.149010616 | 0.00269 | 0.033864708 | -2.85237155 | Cysteine protease | elongation zone | 12 h |
| HORVU.MOREX.r3.6HG0596440.1 | -0.412734698 | 4.52323359 | -3.017585429 | 0.00391 | 0.039547618 | -2.85270277 | Hexosyltransferase | elongation zone | 12 h |
| HORVU.MOREX.r3.2HG0122080.1 | -0.455747725 | 3.732881583 | -3.009593546 | 0.004 | 0.039982662 | -2.85450455 | DUF630 family protein, putative (DUF630 and DUF632) | elongation zone | 12 h |
| HORVU.MOREX.r3.4HG0378960.1 | 0.910911618 | 5.668628466 | 2.870631134 | 0.00587 | 0.048268753 | -2.85504299 | Peroxidase | elongation zone | 12 h |
| HORVU.MOREX.r3.6HG0608390.1 | -0.319042356 | 8.330662034 | -3.137182221 | 0.00278 | 0.034272194 | -2.85531538 | Nuclear matrix constituent protein 1 | elongation zone | 12 h |
| HORVU.MOREX.r3.3HG0236670.1 | 0.682635025 | 5.680641273 | 2.919451102 | 0.00514 | 0.045371176 | -2.85647188 | Protein kinase | elongation zone | 12 h |
| HORVU.MOREX.r3.7HG0645470.1 | 0.683109905 | 7.918742491 | 3.032554556 | 0.00375 | 0.038936762 | -2.85659313 | 30S ribosomal protein S10 | elongation zone | 12 h |
| HORVU.MOREX.r3.1HG0022910.1 | -0.43691274 | 3.569908814 | -2.898564533 | 0.00544 | 0.046616466 | -2.85756711 | Villin | elongation zone | 12 h |
| HORVU.MOREX.r3.6HG0591570.1 | 0.28005563 | 7.825511756 | 3.079774169 | 0.00328 | 0.036983216 | -2.85790118 | 26S proteasome non-atpase regulatory subunit, putative | elongation zone | 12 h |
| HORVU.MOREX.r3.3HG0230070.1 | 0.67076755 | 7.835026485 | 3.02268638 | 0.00385 | 0.039324804 | -2.85915907 | DNA/RNA helicase protein | elongation zone | 12 h |
| HORVU.MOREX.r3.4HG0411870.1 | 0.656925363 | 4.541599175 | 2.880712998 | 0.00571 | 0.047667054 | -2.86426448 | Cysteine-rich repeat secretory protein | elongation zone | 12 h |
| HORVU.MOREX.r3.2HG0185570.1 | -0.675673331 | 1.316705398 | -2.907282046 | 0.00531 | 0.04613544 | -2.86698039 | transcription repressor | elongation zone | 12 h |
| HORVU.MOREX.r3.7HG0706100.1 | 0.273418969 | 5.89851201 | 2.983093033 | 0.00431 | 0.041274283 | -2.86788506 | Survival of motor neuron-related-splicing factor 30 | elongation zone | 12 h |
| HORVU.MOREX.r3.3HG0234970.1 | -0.496604695 | 0.091121871 | -2.915528379 | 0.00519 | 0.045616889 | -2.86822932 | 6,7-dimethyl-8-ribityllumazine synthase | elongation zone | 12 h |
| HORVU.MOREX.r3.7HG0690090.1 | -0.723829946 | 0.298553998 | -2.900723305 | 0.00541 | 0.046443727 | -2.86982328 | PLATZ transcription factor family protein | elongation zone | 12 h |
| HORVU.MOREX.r3.1HG0087990.1 | 0.704487159 | 7.580271477 | 3.013703653 | 0.00395 | 0.039776135 | -2.87315016 | Ubiquitin | elongation zone | 12 h |
| HORVU.MOREX.r3.5HG0445500.1 | 0.344881635 | 6.829254429 | 3.005940732 | 0.00404 | 0.040178373 | -2.87926483 | Tryptophan--tRNA ligase | elongation zone | 12 h |
| HORVU.MOREX.r3.6HG0584160.1 | 1.037805479 | 6.882873618 | 2.938354705 | 0.00488 | 0.044176448 | -2.8801219 | 40S ribosomal protein S24 | elongation zone | 12 h |
| HORVU.MOREX.r3.6HG0613040.1 | -0.415035408 | 5.647164704 | -3.062198398 | 0.00345 | 0.037646224 | -2.88093822 | 2-oxoglutarate-dependent dioxygenase-related family protein | elongation zone | 12 h |
| HORVU.MOREX.r3.3HG0285730.1 | 0.954238618 | 8.817427966 | 3.041549026 | 0.00365 | 0.0386017 | -2.88271662 | RNA binding protein | elongation zone | 12 h |
| HORVU.MOREX.r3.7HG0679370.1 | 0.630843443 | 7.662871406 | 3.024632055 | 0.00383 | 0.03925602 | -2.88362663 | 60S ribosomal protein L31 | elongation zone | 12 h |
| HORVU.MOREX.r3.7HG0637320.1 | 0.696451143 | 5.613003444 | 2.879241274 | 0.00574 | 0.047754823 | -2.88479224 | ATP-dependent zinc metalloprotease FtsH | elongation zone | 12 h |
| HORVU.MOREX.r3.1HG0085100.1 | -0.311119626 | 6.524251428 | -3.101286551 | 0.00308 | 0.036191331 | -2.88575985 | GDSL esterase/lipase | elongation zone | 12 h |
| HORVU.MOREX.r3.6HG0619730.1 | -0.447968364 | 8.659854696 | -3.175085713 | 0.0025 | 0.032934053 | -2.88710254 | Nitrite reductase | elongation zone | 12 h |
| HORVU.MOREX.r3.5HG0485430.1 | 0.647163297 | 6.220484243 | 2.935545378 | 0.00491 | 0.044383997 | -2.88813927 | #N/A | elongation zone | 12 h |
| HORVU.MOREX.r3.4HG0386800.1 | -0.295259663 | 5.120323725 | -2.991421115 | 0.00421 | 0.040797666 | -2.89010381 | Ankyrin repeat family protein, putative, expressed | elongation zone | 12 h |
| HORVU.MOREX.r3.6HG0543520.1 | 0.473619808 | 7.261198846 | 3.002118419 | 0.00408 | 0.040260354 | -2.89029655 | #N/A | elongation zone | 12 h |
| HORVU.MOREX.r3.5HG0510490.1 | 0.681661916 | 5.205188872 | 2.861357987 | 0.00602 | 0.048806356 | -2.89270771 | ATP-dependent RNA helicase, putative | elongation zone | 12 h |
| HORVU.MOREX.r3.2HG0113580.1 | 0.906906348 | 7.071697111 | 2.940567376 | 0.00485 | 0.043984368 | -2.89344786 | Ribosomal protein S8e/ribosomal biogenesis NSA2 | elongation zone | 12 h |
| HORVU.MOREX.r3.2HG0117790.1 | -0.491307534 | 4.935391398 | -2.963672269 | 0.00455 | 0.04241279 | -2.89536306 | Choline transporter-like protein | elongation zone | 12 h |
| HORVU.MOREX.r3.4HG0337120.1 | 0.503894442 | 5.989451331 | 2.968831873 | 0.00448 | 0.042098534 | -2.89589202 | Histone H1, putative | elongation zone | 12 h |
| HORVU.MOREX.r3.6HG0552010.1 | 1.017414298 | 6.596518414 | 2.907100028 | 0.00532 | 0.046138583 | -2.89602545 | Ribosome biogenesis regulatory protein-like protein | elongation zone | 12 h |
| HORVU.MOREX.r3.3HG0219360.1 | 0.939198836 | 5.80749316 | 2.868921764 | 0.0059 | 0.048306767 | -2.89798652 | 50S ribosomal protein L6 | elongation zone | 12 h |
| HORVU.MOREX.r3.2HG0172110.1 | -0.482866133 | 5.282401422 | -3.021876529 | 0.00386 | 0.039354132 | -2.90053527 | Dihydrofolate reductase | elongation zone | 12 h |
| HORVU.MOREX.r3.6HG0626650.1 | 0.22636237 | 8.259288679 | 3.090303959 | 0.00318 | 0.036772996 | -2.90116889 | 26S protease regulatory subunit-like protein | elongation zone | 12 h |
| HORVU.MOREX.r3.2HG0170030.1 | -0.579536885 | 0.532968135 | -2.926805909 | 0.00503 | 0.044862884 | -2.90361559 | Kinase family protein | elongation zone | 12 h |
| HORVU.MOREX.r3.5HG0498770.1 | 0.348105143 | 6.313632909 | 2.989252708 | 0.00423 | 0.040956128 | -2.90402112 | Hexosyltransferase | elongation zone | 12 h |
| HORVU.MOREX.r3.2HG0199260.1 | -0.331633452 | 7.417796497 | -3.122065114 | 0.00291 | 0.035232963 | -2.90613251 | Glycosyltransferase | elongation zone | 12 h |

**Table S5** Continued.

| ID | logFC | AveExpr | t | P.Value | adj.P.Val | B | gene description | root zone | time point |
|---|---|---|---|---|---|---|---|---|---|
| HORVU.MOREX.r3.7HG0732650.1 | -0.56450035 | 3.272852308 | -3.05921494 | 0.00348 | 0.037764638 | -2.90712768 | Peroxidase | elongation zone | 12 h |
| HORVU.MOREX.r3.3HG0314880.1 | -0.379770918 | 4.386487876 | -2.987930534 | 0.00425 | 0.04102313 | -2.9074713 | Transmembrane protein 45B | elongation zone | 12 h |
| HORVU.MOREX.r3.7HG0666290.1 | -0.477354603 | 4.739589313 | -3.002801122 | 0.00408 | 0.040241438 | -2.90811092 | Sphingoid base hydroxylase 2 | elongation zone | 12 h |
| HORVU.MOREX.r3.5HG0502750.1 | 0.808616124 | 6.985366873 | 2.98036559 | 0.00434 | 0.041353873 | -2.90967097 | #N/A | elongation zone | 12 h |
| HORVU.MOREX.r3.7HG0710760.1 | 0.284793048 | 4.651152884 | 2.894660888 | 0.0055 | 0.046792558 | -2.91131601 | DGCR14 protein | elongation zone | 12 h |
| HORVU.MOREX.r3.3HG0231330.1 | -0.457414868 | 4.950298797 | -2.988120485 | 0.00425 | 0.04102313 | -2.91293253 | Transferase family protein | elongation zone | 12 h |
| HORVU.MOREX.r3.1HG0084340.1 | 0.812336969 | 9.276473428 | 3.065058229 | 0.00342 | 0.037489533 | -2.91353575 | Guanine nucleotide-binding protein subunit beta-1 | elongation zone | 12 h |
| HORVU.MOREX.r3.5HG0429680.1 | -0.330450654 | 4.534926532 | -2.937246053 | 0.00489 | 0.044253573 | -2.91415648 | GDP-L-galactose phosphorylase 1 | elongation zone | 12 h |
| HORVU.MOREX.r3.4HG0333090.1 | 0.550400892 | 5.57582127 | 2.89839826 | 0.00544 | 0.046616466 | -2.91552992 | Protein FAR1-RELATED SEQUENCE 5 | elongation zone | 12 h |
| HORVU.MOREX.r3.1HG0008510.2 | -0.386472281 | 3.338317116 | -2.865080812 | 0.00596 | 0.048546205 | -2.9207363 | Beta-hexosaminidase | elongation zone | 12 h |
| HORVU.MOREX.r3.2HG0097950.1 | 0.655599546 | 10.61692335 | 3.140654391 | 0.00276 | 0.034208419 | -2.92438622 | Apyrase | elongation zone | 12 h |
| HORVU.MOREX.r3.5HG0494320.1 | 0.456871429 | 7.049375017 | 2.998691461 | 0.00412 | 0.040338444 | -2.93024445 | Nuclear transport factor 2 | elongation zone | 12 h |
| HORVU.MOREX.r3.7HG0736870.1 | -0.443466772 | 5.259227513 | -2.989072074 | 0.00423 | 0.040956128 | -2.93035009 | Molybdenum cofactor sulfurase | elongation zone | 12 h |
| HORVU.MOREX.r3.6HG0591080.1 | 0.933793576 | 6.734278286 | 2.911175258 | 0.00526 | 0.045859795 | -2.93042939 | Reticulocyte-binding protein 2 a | elongation zone | 12 h |
| HORVU.MOREX.r3.1HG0039940.1 | 0.486137443 | 5.166253007 | 2.872013185 | 0.00585 | 0.048187648 | -2.9310405 | Inosine triphosphate pyrophosphatase | elongation zone | 12 h |
| HORVU.MOREX.r3.6HG0587140.1 | 0.447733402 | 4.797451362 | 2.869951424 | 0.00589 | 0.048306767 | -2.935158 | Transmembrane 9 superfamily member | elongation zone | 12 h |
| HORVU.MOREX.r3.1HG0001460.1 | 0.666875739 | 5.41419522 | 2.932998531 | 0.00495 | 0.044521339 | -2.93541852 | Transmembrane protein, putative (DUF247) | elongation zone | 12 h |
| HORVU.MOREX.r3.4HG0393520.1 | -0.65822398 | 3.798189813 | -2.852357472 | 0.00617 | 0.049512065 | -2.93626128 | Microtubule associated protein family protein, putative, express | elongation zone | 12 h |
| HORVU.MOREX.r3.4HG0362040.1 | -0.205139225 | 7.194760833 | -3.051400042 | 0.00355 | 0.037906495 | -2.9374866 | Armadillo repeat-containing protein 8 | elongation zone | 12 h |
| HORVU.MOREX.r3.4HG0406600.1 | -0.635228829 | 0.071866549 | -2.934446058 | 0.00493 | 0.044479997 | -2.9392917 | Heavy metal transport/detoxification superfamily protein | elongation zone | 12 h |
| HORVU.MOREX.r3.4HG0337460.1 | 0.87336602 | 6.171367141 | 2.869721411 | 0.00589 | 0.048306767 | -2.94302396 | Threonine dehydratase | elongation zone | 12 h |
| HORVU.MOREX.r3.2HG0185340.1 | -0.712015963 | 1.427350358 | -2.87803426 | 0.00576 | 0.04776728 | -2.94466808 | Cationic amino acid transporter, putative | elongation zone | 12 h |
| HORVU.MOREX.r3.5HG0536710.1 | 0.415450209 | 6.624728725 | 3.010737739 | 0.00399 | 0.039893403 | -2.94512669 | NAD/NADP-dependent betaine aldehyde dehydrogenase | elongation zone | 12 h |
| HORVU.MOREX.r3.4HG0331420.1 | 1.206447566 | 6.43121596 | 2.84790724 | 0.00625 | 0.049762195 | -2.94519079 | Peptidyl-prolyl cis-trans isomerase | elongation zone | 12 h |
| HORVU.MOREX.r3.4HG0388500.1 | 0.660947219 | 6.470028413 | 2.912484895 | 0.00524 | 0.045789106 | -2.94524325 | tRNA (Guanine(26)-N(2))-dimethyltransferase | elongation zone | 12 h |
| HORVU.MOREX.r3.4HG0355680.1 | 0.702851985 | 6.142242249 | 2.912128613 | 0.00524 | 0.04581453 | -2.94593008 | stress response NST1-like protein | elongation zone | 12 h |
| HORVU.MOREX.r3.5HG0429940.1 | 1.312146121 | 6.908001713 | 2.886416706 | 0.00563 | 0.047232185 | -2.94826134 | ATP-dependent RNA helicase | elongation zone | 12 h |
| HORVU.MOREX.r3.1HG0040610.1 | 0.566160565 | 8.863532503 | 3.050881459 | 0.00356 | 0.037942384 | -2.94916757 | 60 kDa chaperonin | elongation zone | 12 h |
| HORVU.MOREX.r3.2HG0144390.1 | 0.408694918 | 5.614147136 | 2.898654652 | 0.00544 | 0.046616466 | -2.95178166 | COP9 signalosome complex subunit-like protein | elongation zone | 12 h |
| HORVU.MOREX.r3.1HG0016810.1 | 0.606331134 | 5.521688944 | 2.864696748 | 0.00597 | 0.048574968 | -2.95352833 | Agenet domain-containing protein | elongation zone | 12 h |
| HORVU.MOREX.r3.7HG0643670.1 | 2.197344058 | 8.84959673 | 2.922164796 | 0.0051 | 0.045258495 | -2.95492219 | Peroxidase | elongation zone | 12 h |
| HORVU.MOREX.r3.7HG0643520.1 | -0.513502199 | 3.174361277 | -2.888356517 | 0.0056 | 0.047175894 | -2.95567336 | 5'-methylthioadenosine/S-adenosylhomocysteine nucleosidase | elongation zone | 12 h |
| HORVU.MOREX.r3.6HG0618740.1 | 0.775927809 | 6.100205577 | 2.900829266 | 0.00541 | 0.046443727 | -2.95579339 | Separase, putative | elongation zone | 12 h |
| HORVU.MOREX.r3.7HG0677230.1 | 0.912450606 | 4.893497944 | 3.054853471 | 0.00352 | 0.037808566 | -2.95885902 | Receptor protein kinase, putative | elongation zone | 12 h |
| HORVU.MOREX.r3.3HG0229480.1 | 0.365025398 | 5.272770002 | 2.879150301 | 0.00574 | 0.047754823 | -2.95895302 | Actin/actin-like family protein | elongation zone | 12 h |
| HORVU.MOREX.r3.5HG0525600.1 | 0.615274527 | 5.809975351 | 2.888878364 | 0.00559 | 0.047149582 | -2.96102131 | 30S ribosomal protein S12 | elongation zone | 12 h |
| HORVU.MOREX.r3.4HG0402650.1 | -0.726134963 | 1.552903388 | -2.85655529 | 0.0061 | 0.049235401 | -2.96248241 | Annexin | elongation zone | 12 h |
| HORVU.MOREX.r3.7HG0677570.1 | -0.698420479 | 2.939774147 | -2.899356602 | 0.00543 | 0.046579264 | -2.96283712 | Glycosyltransferase | elongation zone | 12 h |
| HORVU.MOREX.r3.3HG0219650.1 | 0.420483928 | 6.043803629 | 2.914158735 | 0.00521 | 0.045695322 | -2.96303976 | RNA polymerase II-associated factor 1 | elongation zone | 12 h |
| HORVU.MOREX.r3.7HG0656060.1 | 0.638811873 | 8.486727413 | 3.024000896 | 0.00384 | 0.039277346 | -2.96343398 | Pre-mRNA-processing-splicing factor 8 | elongation zone | 12 h |
| HORVU.MOREX.r3.2HG0173750.1 | -0.334616471 | 8.547686714 | -3.084475138 | 0.00324 | 0.036928455 | -2.97060877 | Aldehyde dehydrogenase | elongation zone | 12 h |

**Table S5** Continued.

| ID | logFC | AveExpr | t | P.Value | adj.P.Val | B | gene description | root zone | time point |
|---|---|---|---|---|---|---|---|---|---|
| HORVU.MOREX.r3.4HG0411270.1 | 0.719156992 | 6.58190466 | 2.921547679 | 0.00511 | 0.045296428 | -2.97417688 | N-acetyltransferase | elongation zone | 12 h |
| HORVU.MOREX.r3.4HG0361420.2 | 0.403242517 | 5.485836237 | 2.888462465 | 0.00559 | 0.047175894 | -2.97453951 | ATP dependent RNA helicase | elongation zone | 12 h |
| HORVU.MOREX.r3.2HG0181660.1 | 0.650683403 | 5.953972389 | 2.905970936 | 0.00533 | 0.04618095 | -2.97872697 | 3-hydroxyacyl-[acyl-carrier-protein] dehydratase FabZ | elongation zone | 12 h |
| HORVU.MOREX.r3.7HG0748070.1 | -1.038129431 | -0.496958177 | -2.856195625 | 0.00611 | 0.049242146 | -2.97880041 | Heavy metal transport/detoxification superfamily protein, putati | elongation zone | 12 h |
| HORVU.MOREX.r3.2HG0130030.1 | -0.537928889 | 1.30207633 | -2.903268795 | 0.00537 | 0.046274876 | -2.98207669 | #N/A | elongation zone | 12 h |
| HORVU.MOREX.r3.7HG0635720.1 | 0.826733187 | 5.955493906 | 2.861521059 | 0.00602 | 0.048803932 | -2.98271477 | 50S ribosomal protein L9 | elongation zone | 12 h |
| HORVU.MOREX.r3.7HG0721650.1 | 0.265891299 | 5.578373097 | 2.960815936 | 0.00458 | 0.042615343 | -2.98385671 | Novel plant snare, putative | elongation zone | 12 h |
| HORVU.MOREX.r3.5HG0484510.1 | 0.562714841 | 7.930415335 | 2.989785373 | 0.00423 | 0.040945934 | -2.984392 | 60S ribosomal protein L34 | elongation zone | 12 h |
| HORVU.MOREX.r3.4HG0353520.1 | 0.656135916 | 9.059200888 | 3.04177951 | 0.00365 | 0.0386017 | -2.98477117 | 30S ribosomal protein S9 | elongation zone | 12 h |
| HORVU.MOREX.r3.5HG0468420.1 | 0.292475491 | 6.568899246 | 2.959683777 | 0.0046 | 0.042694723 | -2.98518932 | #N/A | elongation zone | 12 h |
| HORVU.MOREX.r3.1HG0076900.1 | 1.026661097 | 6.223225304 | 2.857979163 | 0.00608 | 0.049103351 | -2.98562024 | GATA transcription factor | elongation zone | 12 h |
| HORVU.MOREX.r3.2HG0191450.1 | 0.799065903 | 6.226559771 | 2.8632191 | 0.00599 | 0.048636131 | -2.98679148 | Serine/threonine-protein phosphatase 2A 56 kDa regulatory subu | elongation zone | 12 h |
| HORVU.MOREX.r3.3HG0254430.1 | -0.292963101 | 8.132393812 | -3.087400698 | 0.00321 | 0.036913075 | -2.9872216 | Oxidoreductase | elongation zone | 12 h |
| HORVU.MOREX.r3.5HG0441770.1 | 0.803760799 | 6.5387839 | 2.89205986 | 0.00554 | 0.046904576 | -2.98815294 | Nuclear pore complex protein NUP93A | elongation zone | 12 h |
| HORVU.MOREX.r3.1HG0061880.1 | 0.48140786 | 7.374139112 | 2.977462183 | 0.00437 | 0.041519226 | -2.99010359 | Ran-binding protein 1 | elongation zone | 12 h |
| HORVU.MOREX.r3.3HG0315410.1 | -0.327532629 | 5.906295463 | -2.978045673 | 0.00437 | 0.041506896 | -2.99070801 | Vesicle-associated protein 1-2 | elongation zone | 12 h |
| HORVU.MOREX.r3.1HG0073190.1 | 0.685119959 | 7.62223663 | 2.970400499 | 0.00446 | 0.041991983 | -2.99093282 | 30S ribosomal protein S8, putative | elongation zone | 12 h |
| HORVU.MOREX.r3.2HG0115970.1 | 0.892241871 | 3.355367576 | 2.86923845 | 0.0059 | 0.048306767 | -2.9953444 | NAD(P)H dehydrogenase (Quinone) | elongation zone | 12 h |
| HORVU.MOREX.r3.5HG0457550.1 | -0.391770239 | 4.931610725 | -2.964344002 | 0.00454 | 0.04241279 | -2.99801787 | Plant/F1M20-13 protein | elongation zone | 12 h |
| HORVU.MOREX.r3.1HG0084910.1 | 0.558826204 | 5.655310184 | 2.873604319 | 0.00583 | 0.048075075 | -3.00229015 | 50S ribosomal protein L7/L12 | elongation zone | 12 h |
| HORVU.MOREX.r3.2HG0120820.1 | 0.282294884 | 4.679691645 | 2.8696963 | 0.00589 | 0.048306767 | -3.00379175 | #N/A | elongation zone | 12 h |
| HORVU.MOREX.r3.4HG0363150.1 | 0.552003928 | 5.976929467 | 2.885805319 | 0.00564 | 0.047276131 | -3.00562762 | WD40 repeat-like protein | elongation zone | 12 h |
| HORVU.MOREX.r3.2HG0179540.1 | 0.418720548 | 4.341715765 | 2.88421854 | 0.00566 | 0.047404516 | -3.00810298 | Ergosterol biosynthetic protein 28 | elongation zone | 12 h |
| HORVU.MOREX.r3.2HG0129980.1 | 0.580935985 | 5.921548911 | 2.875607441 | 0.00579 | 0.047903389 | -3.01073136 | SUMO-activating enzyme 2 | elongation zone | 12 h |
| HORVU.MOREX.r3.3HG0294340.1 | -0.585854853 | 2.811533245 | -2.863577618 | 0.00599 | 0.048636131 | -3.01455225 | Peroxidase | elongation zone | 12 h |
| HORVU.MOREX.r3.5HG0512350.1 | 0.888676789 | 7.007305835 | 2.906237449 | 0.00533 | 0.04618095 | -3.01640224 | Cytochrome P450 | elongation zone | 12 h |
| HORVU.MOREX.r3.4HG0397450.1 | 0.720702425 | 7.797749659 | 2.960598863 | 0.00458 | 0.042615343 | -3.01701975 | T-complex protein 1 subunit beta | elongation zone | 12 h |
| HORVU.MOREX.r3.3HG0280140.1 | 0.853977415 | 6.992266616 | 2.914392842 | 0.00521 | 0.045685378 | -3.02469987 | 60S ribosomal protein L37a | elongation zone | 12 h |
| HORVU.MOREX.r3.6HG0621950.1 | 0.789782524 | 7.314803764 | 2.933377255 | 0.00494 | 0.044514543 | -3.02563555 | 60S ribosomal protein L35a-like protein | elongation zone | 12 h |
| HORVU.MOREX.r3.1HG0092810.1 | 0.791697614 | 7.116496563 | 2.895472183 | 0.00549 | 0.046758121 | -3.02733781 | Late embryogenesis abundant protein Lea14 | elongation zone | 12 h |
| HORVU.MOREX.r3.4HG0351070.1 | -0.680823961 | 2.367322112 | -2.859267049 | 0.00606 | 0.048988577 | -3.02810769 | STRUBBELIG-receptor family 5 | elongation zone | 12 h |
| HORVU.MOREX.r3.2HG0108680.1 | -0.499951742 | 3.066286477 | -2.922706815 | 0.00509 | 0.045229989 | -3.02887277 | Demethylmenaquinone methyltransferase | elongation zone | 12 h |
| HORVU.MOREX.r3.2HG0112180.1 | -0.53789941 | 4.377395587 | -2.887234133 | 0.00561 | 0.047204707 | -3.02969937 | Receptor-like kinase | elongation zone | 12 h |
| HORVU.MOREX.r3.5HG0477180.1 | 0.309915771 | 5.597570777 | 2.914786526 | 0.0052 | 0.045655368 | -3.02985482 | LAG1 longevity assurance-like protein | elongation zone | 12 h |
| HORVU.MOREX.r3.2HG0215610.1 | 0.929375841 | 6.340912174 | 2.959324862 | 0.0046 | 0.042717988 | -3.02998687 | Peroxidase | elongation zone | 12 h |
| HORVU.MOREX.r3.2HG0168090.1 | 0.320024673 | 5.385495042 | 2.904533395 | 0.00535 | 0.046230732 | -3.03029409 | Sphingomyelin synthase-like domain-containing protein | elongation zone | 12 h |
| HORVU.MOREX.r3.7HG0689490.1 | 0.549083273 | 6.098734945 | 2.883114873 | 0.00568 | 0.047427345 | -3.03126766 | Peptidyl-prolyl cis-trans isomerase | elongation zone | 12 h |
| HORVU.MOREX.r3.7HG0739640.1 | 0.779701561 | 8.298465212 | 2.982652606 | 0.00431 | 0.041282541 | -3.03234696 | Auxin efflux carrier component | elongation zone | 12 h |
| HORVU.MOREX.r3.1HG0063880.1 | -0.566652859 | 4.61444207 | -2.906214244 | 0.00533 | 0.04618095 | -3.03457094 | Glycosyltransferase | elongation zone | 12 h |
| HORVU.MOREX.r3.7HG0656430.1 | 0.300035593 | 6.915191761 | 2.943914995 | 0.0048 | 0.043752599 | -3.03531619 | Zinc finger CCCH domain protein | elongation zone | 12 h |
| HORVU.MOREX.r3.1HG0078910.1 | -0.459548855 | 4.676362005 | -2.93960023 | 0.00486 | 0.044063319 | -3.03579908 | p-loop containing nucleoside triphosphate hydrolases superfami | elongation zone | 12 h |

**Table S5** Continued.

| ID | logFC | AveExpr | t | P.Value | adj.P.Val | B | gene description | root zone | time point |
|---|---|---|---|---|---|---|---|---|---|
| HORVU.MOREX.r3.4HG0388980.2 | 0.342050984 | 5.157208042 | 2.883260022 | 0.00567 | 0.047427345 | -3.03615129 | #N/A | elongation zone | 12 h |
| HORVU.MOREX.r3.4HG0389850.1 | -0.380673133 | 6.082211287 | -3.00105967 | 0.0041 | 0.040265883 | -3.03687727 | #N/A | elongation zone | 12 h |
| HORVU.MOREX.r3.6HG0619540.1 | 0.339176199 | 5.67347569 | 2.869433815 | 0.00589 | 0.048306767 | -3.03868711 | THO complex subunit 5 | elongation zone | 12 h |
| HORVU.MOREX.r3.2HG0160270.2 | 0.607985534 | 9.915803777 | 3.060189158 | 0.00347 | 0.037761197 | -3.04068471 | 40S ribosomal protein S8 | elongation zone | 12 h |
| HORVU.MOREX.r3.4HG0387150.1 | 0.871467136 | 6.899898072 | 2.911459842 | 0.00525 | 0.045859795 | -3.04154365 | Histone H2A | elongation zone | 12 h |
| HORVU.MOREX.r3.3HG0312360.1 | -0.239135493 | 7.147206035 | -3.037752985 | 0.00369 | 0.038671495 | -3.04442983 | Coiled-coil domain-containing protein SCD2 | elongation zone | 12 h |
| HORVU.MOREX.r3.7HG0726700.1 | 0.73170601 | 7.183776553 | 2.918621601 | 0.00515 | 0.045408324 | -3.04638955 | Leucine-rich repeat protein kinase family protein | elongation zone | 12 h |
| HORVU.MOREX.r3.6HG0630670.1 | 2.093795787 | 8.145324526 | 2.856288785 | 0.00611 | 0.049242146 | -3.0481649 | rRNA 2'-O-methyltransferase fibrillarin 2 | elongation zone | 12 h |
| HORVU.MOREX.r3.7HG0655280.1 | -0.455658489 | 1.566144439 | -2.926338054 | 0.00504 | 0.044895531 | -3.04949994 | Galactoside 2-alpha-L-fucosyltransferase | elongation zone | 12 h |
| HORVU.MOREX.r3.6HG0608780.1 | 0.399007469 | 7.822442469 | 3.009416855 | 0.004 | 0.039983 | -3.05031984 | Heat shock 70 kDa protein, putative | elongation zone | 12 h |
| HORVU.MOREX.r3.2HG0160130.1 | -0.478213173 | 3.662810319 | -2.954697699 | 0.00466 | 0.042998159 | -3.05115306 | Basic helix-loop-helix transcription factor | elongation zone | 12 h |
| HORVU.MOREX.r3.5HG0511110.3 | -0.398906574 | 4.233551779 | -2.927048358 | 0.00503 | 0.044862884 | -3.0520262 | Potassium channel | elongation zone | 12 h |
| HORVU.MOREX.r3.5HG0530200.1 | -0.284987088 | 8.300591094 | -3.106295829 | 0.00304 | 0.035944821 | -3.05248227 | Actin | elongation zone | 12 h |
| HORVU.MOREX.r3.7HG0654410.1 | 0.60383893 | 6.620820722 | 2.926893326 | 0.00503 | 0.044862884 | -3.0527049 | Histone H3 | elongation zone | 12 h |
| HORVU.MOREX.r3.1HG0035400.1 | -0.412076659 | 4.194434053 | -2.904954322 | 0.00535 | 0.046230732 | -3.05298171 | Ras-related protein, expressed | elongation zone | 12 h |
| HORVU.MOREX.r3.6HG0614500.1 | 0.321876284 | 5.054555082 | 2.861019963 | 0.00603 | 0.048830126 | -3.05752066 | 2,3-bisphosphoglycerate-dependent phosphoglycerate mutase | elongation zone | 12 h |
| HORVU.MOREX.r3.2HG0199300.1 | 1.062718027 | 7.23323174 | 2.871224724 | 0.00586 | 0.048231861 | -3.05920162 | Protein SDA1-like protein | elongation zone | 12 h |
| HORVU.MOREX.r3.6HG0614840.1 | -0.406983683 | 5.332590306 | -2.956722796 | 0.00463 | 0.042834083 | -3.06198672 | Shikimate kinase | elongation zone | 12 h |
| HORVU.MOREX.r3.7HG0656790.1 | 0.693229773 | 6.625384114 | 2.890726519 | 0.00556 | 0.047007942 | -3.06284911 | 30S ribosomal protein S10 | elongation zone | 12 h |
| HORVU.MOREX.r3.1HG0076980.1 | 0.297279412 | 8.500485602 | 3.035850604 | 0.00371 | 0.038729899 | -3.06393726 | Dolichyl-diphosphooligosaccharide--protein glycosyltransferase s | elongation zone | 12 h |
| HORVU.MOREX.r3.3HG0305860.1 | 0.79863207 | 6.802235483 | 2.870517047 | 0.00588 | 0.048268753 | -3.07892799 | Nuclear transport factor 2 family protein with RNA binding doma | elongation zone | 12 h |
| HORVU.MOREX.r3.2HG0213250.1 | 0.470164991 | 6.347478748 | 2.946782096 | 0.00476 | 0.043677865 | -3.08025968 | Choline transporter-related family protein | elongation zone | 12 h |
| HORVU.MOREX.r3.4HG0405780.1 | -0.413383852 | 7.951420898 | -3.075359572 | 0.00332 | 0.037190743 | -3.08229569 | Receptor-like protein kinase | elongation zone | 12 h |
| HORVU.MOREX.r3.1HG0017750.1 | -0.345287872 | 4.005500278 | -2.854091136 | 0.00615 | 0.04938991 | -3.08271584 | Kinase family protein | elongation zone | 12 h |
| HORVU.MOREX.r3.6HG0625570.1 | -0.457769965 | 5.941941183 | -2.985102018 | 0.00428 | 0.041182136 | -3.09233418 | Transmembrane protein | elongation zone | 12 h |
| HORVU.MOREX.r3.5HG0515330.1 | 0.645838988 | 6.09421809 | 2.855085326 | 0.00613 | 0.049316553 | -3.09511582 | 60S ribosomal protein L44 | elongation zone | 12 h |
| HORVU.MOREX.r3.2HG0126380.1 | -0.453337732 | 5.874038041 | -3.027913762 | 0.0038 | 0.039108923 | -3.09826364 | #N/A | elongation zone | 12 h |
| HORVU.MOREX.r3.6HG0546420.1 | 0.811902893 | 9.042284244 | 2.976670348 | 0.00438 | 0.041550671 | -3.09998281 | 60S ribosomal protein L13 | elongation zone | 12 h |
| HORVU.MOREX.r3.2HG0113880.1 | -0.453881692 | 4.823614042 | -2.872876746 | 0.00584 | 0.048151385 | -3.1025005 | Heavy metal transport/detoxification superfamily protein, putati | elongation zone | 12 h |
| HORVU.MOREX.r3.1HG0056740.1 | 0.541835962 | 8.162242671 | 2.992578221 | 0.00419 | 0.040685115 | -3.10546117 | Pyruvate kinase | elongation zone | 12 h |
| HORVU.MOREX.r3.3HG0263050.1 | 0.355124609 | 6.674660346 | 2.898306954 | 0.00545 | 0.046616466 | -3.10673343 | 2-oxoglutarate (2OG) and Fe(II)-dependent oxygenase superfami | elongation zone | 12 h |
| HORVU.MOREX.r3.4HG0393930.1 | 0.5709058 | 8.844580814 | 2.994184921 | 0.00417 | 0.040647009 | -3.10797217 | Ubiquitin | elongation zone | 12 h |
| HORVU.MOREX.r3.2HG0176960.1 | 0.283907521 | 4.988465286 | 2.854672893 | 0.00614 | 0.049352603 | -3.10993465 | Kelch repeat-containing protein | elongation zone | 12 h |
| HORVU.MOREX.r3.2HG0187990.1 | -0.353201691 | 5.49990496 | -2.936215758 | 0.00491 | 0.044321273 | -3.11068085 | O-fucosyltransferase family protein | elongation zone | 12 h |
| HORVU.MOREX.r3.5HG0518560.1 | -1.199400096 | 2.148281347 | -2.853963524 | 0.00615 | 0.04938991 | -3.11372973 | Leucine-rich repeat protein kinase family protein | elongation zone | 12 h |
| HORVU.MOREX.r3.7HG0721760.1 | 0.289413786 | 8.421100262 | 3.02053987 | 0.00388 | 0.039407591 | -3.11650123 | Translocon-associated protein subunit alpha | elongation zone | 12 h |
| HORVU.MOREX.r3.7HG0752370.1 | -0.606400625 | 2.08824866 | -2.919250656 | 0.00514 | 0.045371176 | -3.11705672 | NBS-LRR-like resistance protein | elongation zone | 12 h |
| HORVU.MOREX.r3.7HG0705860.1 | -0.397078963 | 7.982365784 | -3.065353788 | 0.00342 | 0.037489533 | -3.11753886 | Non-lysosomal glucosylceramidase | elongation zone | 12 h |
| HORVU.MOREX.r3.7HG0681100.1 | -0.280121191 | 5.610348477 | -2.930832179 | 0.00498 | 0.044603947 | -3.11893017 | Glycine-rich domain-containing protein 2 | elongation zone | 12 h |
| HORVU.MOREX.r3.1HG0044310.1 | 0.603045083 | 7.299346006 | 2.90796892 | 0.0053 | 0.046106442 | -3.11979322 | Nascent polypeptide-associated complex subunit beta | elongation zone | 12 h |
| HORVU.MOREX.r3.6HG0631360.1 | -0.33504033 | 4.355721511 | -2.86812664 | 0.00591 | 0.048306767 | -3.12279946 | Disease resistance protein RPM1 | elongation zone | 12 h |

**Table S5** Continued.

| ID | logFC | AveExpr | t | P.Value | adj.P.Val | B | gene description | root zone | time point |
|---|---|---|---|---|---|---|---|---|---|
| HORVU.MOREX.r3.1HG0055230.1 | 0.667815911 | 7.59025634 | 2.955843608 | 0.00465 | 0.042900147 | -3.12451366 | Methyltransferase | elongation zone | 12 h |
| HORVU.MOREX.r3.5HG0457110.1 | 0.56952416 | 9.82719665 | 3.028770609 | 0.00379 | 0.039108923 | -3.1301289 | 30S ribosomal protein S4 | elongation zone | 12 h |
| HORVU.MOREX.r3.4HG0389620.1 | -0.361026935 | 4.010754431 | -2.84958727 | 0.00622 | 0.049700096 | -3.13055524 | Glycogen synthase | elongation zone | 12 h |
| HORVU.MOREX.r3.7HG0739480.1 | 0.656194298 | 7.565794341 | 2.92065256 | 0.00512 | 0.045330047 | -3.13163572 | Histone H2B | elongation zone | 12 h |
| HORVU.MOREX.r3.2HG0189070.1 | 1.219226798 | 1.242983665 | 2.919860015 | 0.00513 | 0.045371176 | -3.13231304 | 14 kDa proline-rich protein dc2.15 | elongation zone | 12 h |
| HORVU.MOREX.r3.3HG0301930.1 | -0.445321284 | 4.019686283 | -2.879328914 | 0.00574 | 0.047754823 | -3.13447671 | G patch domain protein | elongation zone | 12 h |
| HORVU.MOREX.r3.4HG0390230.1 | 0.393026691 | 6.080893711 | 2.862045233 | 0.00601 | 0.048753496 | -3.13671945 | Eukaryotic translation initiation factor 4E | elongation zone | 12 h |
| HORVU.MOREX.r3.7HG0648810.1 | -0.256091208 | 6.330560766 | -2.971082351 | 0.00445 | 0.041967805 | -3.13702423 | Acyl-[acyl-carrier-protein] hydrolase | elongation zone | 12 h |
| HORVU.MOREX.r3.3HG0223160.1 | -0.547742168 | 5.278306557 | -3.01161857 | 0.00398 | 0.03986915 | -3.14083548 | aberrant root formation protein | elongation zone | 12 h |
| HORVU.MOREX.r3.7HG0727290.1 | -0.57864768 | 3.998416026 | -2.894177742 | 0.00551 | 0.046806243 | -3.14307341 | #N/A | elongation zone | 12 h |
| HORVU.MOREX.r3.1HG0074500.1 | 0.799384988 | 7.370525428 | 2.891596627 | 0.00555 | 0.046934592 | -3.14670496 | Histone H3 | elongation zone | 12 h |
| HORVU.MOREX.r3.2HG0200920.1 | -0.59318877 | 3.047317061 | -2.937004883 | 0.00489 | 0.044263597 | -3.14692729 | Agmatine coumaroyltransferase-2 | elongation zone | 12 h |
| HORVU.MOREX.r3.2HG0112690.1 | -0.717408979 | 1.247172627 | -2.951919897 | 0.0047 | 0.043259525 | -3.14743163 | Peroxidase | elongation zone | 12 h |
| HORVU.MOREX.r3.3HG0255020.1 | -0.699286777 | 3.485610317 | -3.005991358 | 0.00404 | 0.040178373 | -3.15157012 | Peroxidase | elongation zone | 12 h |
| HORVU.MOREX.r3.7HG0691650.1 | 0.333773703 | 7.880664191 | 2.948593932 | 0.00474 | 0.043517337 | -3.15534981 | RNA binding protein | elongation zone | 12 h |
| HORVU.MOREX.r3.5HG0424290.1 | -0.341661281 | 9.703425568 | -3.069823574 | 0.00337 | 0.037331432 | -3.15654538 | Cysteine synthase | elongation zone | 12 h |
| HORVU.MOREX.r3.3HG0254950.1 | -0.84685387 | 1.019759768 | -2.878040936 | 0.00576 | 0.04776728 | -3.17129481 | Peroxidase | elongation zone | 12 h |
| HORVU.MOREX.r3.1HG0022060.1 | 0.47671216 | 7.428714781 | 2.901011166 | 0.00541 | 0.046443727 | -3.17705313 | Cullin-1 | elongation zone | 12 h |
| HORVU.MOREX.r3.4HG0396220.1 | 0.304230092 | 8.114651828 | 2.980936502 | 0.00433 | 0.041314646 | -3.17831158 | Alpha-glucosidase like protein | elongation zone | 12 h |
| HORVU.MOREX.r3.6HG0566190.1 | -0.513018484 | 5.134408532 | -3.023600209 | 0.00384 | 0.039284359 | -3.18368411 | Peroxidase | elongation zone | 12 h |
| HORVU.MOREX.r3.6HG0624630.1 | 0.330304524 | 6.985603223 | 2.928929207 | 0.005 | 0.044750195 | -3.18488271 | Aspartate aminotransferase | elongation zone | 12 h |
| HORVU.MOREX.r3.5HG0489130.1 | 0.950825725 | 6.654428009 | 2.931957877 | 0.00496 | 0.044592291 | -3.18490221 | Receptor protein kinase, putative | elongation zone | 12 h |
| HORVU.MOREX.r3.4HG0337740.1 | -0.361726469 | 6.566981129 | -2.943359643 | 0.00481 | 0.043773947 | -3.18694458 | Receptor protein kinase, putative | elongation zone | 12 h |
| HORVU.MOREX.r3.1HG0041260.1 | 0.305520107 | 7.367448887 | 2.923765837 | 0.00508 | 0.045176231 | -3.18999341 | Eukaryotic translation initiation factor 4E | elongation zone | 12 h |
| HORVU.MOREX.r3.6HG0614970.1 | -0.315253944 | 5.579922669 | -2.877932734 | 0.00576 | 0.04776728 | -3.1916892 | Kelch repeat-containing protein | elongation zone | 12 h |
| HORVU.MOREX.r3.1HG0051180.1 | -0.428635896 | 5.081434726 | -2.927453157 | 0.00503 | 0.044862884 | -3.19442782 | Rhomboid-like protein | elongation zone | 12 h |
| HORVU.MOREX.r3.3HG0272030.1 | 0.477768938 | 8.079674682 | 2.930587373 | 0.00498 | 0.044603947 | -3.19450287 | N-alpha-acetyltransferase 16, NatA auxiliary subunit | elongation zone | 12 h |
| HORVU.MOREX.r3.6HG0549380.1 | 0.404296281 | 7.237694492 | 2.877918751 | 0.00576 | 0.04776728 | -3.19525184 | Gamma-glutamylcyclotransferase | elongation zone | 12 h |
| HORVU.MOREX.r3.7HG0738190.1 | -0.335818143 | 5.38770736 | -2.915275459 | 0.0052 | 0.045616889 | -3.20860932 | Copper-transporting ATPase | elongation zone | 12 h |
| HORVU.MOREX.r3.5HG0523150.1 | 0.248718051 | 6.194464004 | 2.845307162 | 0.00629 | 0.049959998 | -3.2111347 | Protein phosphatase 2c, putative | elongation zone | 12 h |
| HORVU.MOREX.r3.1HG0006400.1 | -0.551255078 | 6.387241341 | -2.986506564 | 0.00427 | 0.041089957 | -3.21165373 | Beta-1,3-galactosyltransferase-like protein | elongation zone | 12 h |
| HORVU.MOREX.r3.4HG0379400.1 | -0.694825149 | 1.850080654 | -2.937905133 | 0.00488 | 0.044199271 | -3.22032492 | Protein kinase-like protein | elongation zone | 12 h |
| HORVU.MOREX.r3.5HG0508450.1 | -0.369809707 | 5.166935854 | -2.865356915 | 0.00596 | 0.048546205 | -3.22195598 | Calmodulin-binding family protein, putative, expressed | elongation zone | 12 h |
| HORVU.MOREX.r3.6HG0609420.1 | -0.379903983 | 5.023973234 | -2.903433263 | 0.00537 | 0.046274876 | -3.22947899 | phosphoenolpyruvate carboxykinase 2 | elongation zone | 12 h |
| HORVU.MOREX.r3.5HG0458450.1 | -0.32439936 | 5.682873141 | -2.8671452 | 0.00593 | 0.048385844 | -3.23168701 | TLD domain-containing protein 1 | elongation zone | 12 h |
| HORVU.MOREX.r3.1HG0066850.1 | -0.389248488 | 4.235298003 | -2.854277143 | 0.00614 | 0.04938644 | -3.23343777 | Adenine nucleotide alpha hydrolases-like superfamily | elongation zone | 12 h |
| HORVU.MOREX.r3.5HG0507470.1 | -0.34858237 | 5.264725422 | -2.893468731 | 0.00552 | 0.046829376 | -3.23578036 | Arogenate dehydratase | elongation zone | 12 h |
| HORVU.MOREX.r3.3HG0325580.1 | -0.256987513 | 5.824048637 | -2.871851942 | 0.00585 | 0.048189541 | -3.23913609 | MACPF domain-containing protein | elongation zone | 12 h |
| HORVU.MOREX.r3.7HG0710920.1 | -0.267048239 | 5.393550621 | -2.852359858 | 0.00617 | 0.049512065 | -3.24075752 | TOM1-like protein 2 | elongation zone | 12 h |
| HORVU.MOREX.r3.6HG0602990.1 | 0.692651589 | 7.353646827 | 2.847211543 | 0.00626 | 0.04981773 | -3.24091281 | High mobility group family | elongation zone | 12 h |
| HORVU.MOREX.r3.2HG0173880.1 | 0.201180376 | 6.785205192 | 2.887104102 | 0.00562 | 0.047204707 | -3.24664055 | UBX domain-containing protein, putative | elongation zone | 12 h |

**Table S5** Continued.

| ID | logFC | AveExpr | t | P.Value | adj.P.Val | B | gene description | root zone | time point |
|---|---|---|---|---|---|---|---|---|---|
| HORVU.MOREX.r3.6HG0551180.1 | 0.35153076 | 7.241444096 | 2.859294997 | 0.00606 | 0.048988577 | -3.25988646 | 3-isopropylmalate dehydratase large subunit | elongation zone | 12 h |
| HORVU.MOREX.r3.1HG0022440.1 | 0.524341881 | 7.971656134 | 2.889309318 | 0.00558 | 0.047136585 | -3.26086177 | Chromatin remodeling factor, putative | elongation zone | 12 h |
| HORVU.MOREX.r3.2HG0165200.1 | -0.599333802 | 5.318500517 | -2.958450791 | 0.00461 | 0.042747009 | -3.263942 | #N/A | elongation zone | 12 h |
| HORVU.MOREX.r3.3HG0254150.1 | -0.276211082 | 7.075489408 | -2.949776176 | 0.00472 | 0.043452654 | -3.26725358 | Phosphoadenosine phosphosulfate reductase family protein | elongation zone | 12 h |
| HORVU.MOREX.r3.3HG0292280.1 | 0.528467151 | 7.335521365 | 2.856072199 | 0.00611 | 0.049242146 | -3.27711245 | Ubiquitin carboxyl-terminal hydrolase-like protein | elongation zone | 12 h |
| HORVU.MOREX.r3.4HG0369570.1 | 0.173927817 | 10.41352761 | 3.075501904 | 0.00332 | 0.037190743 | -3.28443463 | Alpha-1,4-glucan-protein synthase [UDP-forming] | elongation zone | 12 h |
| HORVU.MOREX.r3.1HG0072860.1 | 0.733714287 | 8.053944148 | 2.868057399 | 0.00592 | 0.048306767 | -3.28509427 | 30S ribosomal protein S12 | elongation zone | 12 h |
| HORVU.MOREX.r3.3HG0296600.1 | 0.583286833 | 9.192372739 | 2.930761361 | 0.00498 | 0.044603947 | -3.29675842 | 60S ribosomal protein L24 | elongation zone | 12 h |
| HORVU.MOREX.r3.6HG0578930.1 | 0.203231026 | 7.54149509 | 2.892840608 | 0.00553 | 0.046871369 | -3.29793007 | Histone deacetylase | elongation zone | 12 h |
| HORVU.MOREX.r3.3HG0314490.2 | -0.490389483 | 3.653110454 | -2.84923063 | 0.00623 | 0.049700096 | -3.30769096 | Receptor-like kinase | elongation zone | 12 h |
| HORVU.MOREX.r3.7HG0704470.1 | 0.623393968 | 8.481284146 | 2.895382676 | 0.00549 | 0.046758121 | -3.313952 | 60S acidic ribosomal protein P1 | elongation zone | 12 h |
| HORVU.MOREX.r3.6HG0550950.1 | 0.391071811 | 7.177293507 | 2.85042772 | 0.00621 | 0.049645466 | -3.31624312 | NADP dependent sorbitol 6-phosphate dehydrogenase family pro | elongation zone | 12 h |
| HORVU.MOREX.r3.6HG0610930.1 | 0.405844507 | 7.991073157 | 2.899205799 | 0.00543 | 0.046579264 | -3.31673128 | Enhancer of mRNA-decapping protein 4 | elongation zone | 12 h |
| HORVU.MOREX.r3.6HG0628930.1 | 0.477342458 | 7.959731866 | 2.868692565 | 0.00591 | 0.048306767 | -3.31743782 | Alcohol dehydrogenase | elongation zone | 12 h |
| HORVU.MOREX.r3.4HG0402000.1 | 0.526499416 | 8.180685451 | 2.91919627 | 0.00514 | 0.045371176 | -3.32030656 | Kinase interacting (KIP1-like) family protein, putative | elongation zone | 12 h |
| HORVU.MOREX.r3.1HG0081060.1 | 0.429929803 | 7.924567077 | 2.881412582 | 0.0057 | 0.047595228 | -3.3233397 | Mitochondrial outer membrane porin | elongation zone | 12 h |
| HORVU.MOREX.r3.5HG0491130.1 | 0.478289324 | 8.223604151 | 2.87775393 | 0.00576 | 0.04776728 | -3.3376914 | Chaperone DnaK | elongation zone | 12 h |
| HORVU.MOREX.r3.2HG0167300.1 | -0.472868125 | 4.945499856 | -2.872404772 | 0.00585 | 0.04818291 | -3.34747866 | Late embryogenesis abundant (LEA) hydroxyproline-rich glycopro | elongation zone | 12 h |
| HORVU.MOREX.r3.1HG0024510.1 | -0.423164755 | 5.713562753 | -2.90219879 | 0.00539 | 0.046372228 | -3.35198611 | Universal stress protein | elongation zone | 12 h |
| HORVU.MOREX.r3.2HG0112820.1 | 0.592113834 | 9.728760634 | 2.932589139 | 0.00495 | 0.044552225 | -3.35443581 | Ribosomal protein L19 | elongation zone | 12 h |
| HORVU.MOREX.r3.6HG0593920.1 | 0.418633688 | 8.297449459 | 2.874308177 | 0.00582 | 0.048001992 | -3.36223612 | #N/A | elongation zone | 12 h |
| HORVU.MOREX.r3.7HG0714620.1 | -0.403076027 | 6.402884248 | -2.886294075 | 0.00563 | 0.047232185 | -3.36583951 | 3'-N-debenzoyl-2'-deoxytaxol N-benzoyltransferase | elongation zone | 12 h |
| HORVU.MOREX.r3.1HG0039670.1 | 0.208126993 | 7.408424513 | 2.869064101 | 0.0059 | 0.048306767 | -3.37822814 | Calcium ion-binding protein | elongation zone | 12 h |
| HORVU.MOREX.r3.6HG0564480.1 | 0.565485578 | 8.296340254 | 2.856799754 | 0.0061 | 0.04922198 | -3.3869467 | T-complex protein 1 subunit eta | elongation zone | 12 h |
| HORVU.MOREX.r3.5HG0444520.1 | -0.612373717 | 5.327750335 | -2.909647079 | 0.00528 | 0.045952435 | -3.38935312 | NADP-dependent alkenal double bond reductase | elongation zone | 12 h |
| HORVU.MOREX.r3.1HG0073380.1 | 0.685789953 | 8.454580573 | 2.851902871 | 0.00618 | 0.049512564 | -3.39480067 | 60S ribosomal protein L36 | elongation zone | 12 h |
| HORVU.MOREX.r3.5HG0470270.1 | 0.56900555 | 9.530865348 | 2.903417876 | 0.00537 | 0.046274876 | -3.39926061 | 50S ribosomal protein L22 | elongation zone | 12 h |
| HORVU.MOREX.r3.1HG0088510.1 | 0.695413918 | 9.089763599 | 2.867305181 | 0.00593 | 0.048383961 | -3.41933361 | 60S ribosomal protein L18a | elongation zone | 12 h |
| HORVU.MOREX.r3.7HG0641060.1 | -0.398401199 | 7.954456485 | -2.927079513 | 0.00503 | 0.044862884 | -3.42045596 | Kinase-like protein | elongation zone | 12 h |
| HORVU.MOREX.r3.1HG0094710.1 | 0.331758591 | 8.999934807 | 2.892619549 | 0.00553 | 0.046880427 | -3.43915997 | Translation initiation factor if-2, putative | elongation zone | 12 h |
| HORVU.MOREX.r3.7HG0718190.1 | -0.572568363 | 6.009060521 | -2.894367508 | 0.0055 | 0.046806243 | -3.4554614 | ABC transporter family protein | elongation zone | 12 h |
| HORVU.MOREX.r3.3HG0254990.1 | -0.389976397 | 7.164090125 | -2.919074338 | 0.00514 | 0.045371176 | -3.47010269 | Peroxidase | elongation zone | 12 h |
| HORVU.MOREX.r3.3HG0283130.1 | -0.689769038 | 6.080795139 | -2.850785274 | 0.0062 | 0.049624591 | -3.47494432 | Subtilisin-like protease | elongation zone | 12 h |
| HORVU.MOREX.r3.2HG0197260.1 | -0.277286229 | 7.52618409 | -2.892396638 | 0.00553 | 0.046889731 | -3.49218591 | WEB family protein, chloroplastic | elongation zone | 12 h |
| HORVU.MOREX.r3.3HG0236850.1 | -0.234602448 | 8.690188681 | -2.896704704 | 0.00547 | 0.046711054 | -3.55190634 | Malic enzyme | elongation zone | 12 h |
| HORVU.MOREX.r3.7HG0711070.1 | -0.323777753 | 7.633823382 | -2.877825963 | 0.00576 | 0.04776728 | -3.55382313 | Receptor-like kinase, putative | elongation zone | 12 h |
| HORVU.MOREX.r3.1HG0069420.1 | -0.24401767 | 7.626353217 | -2.86506656 | 0.00596 | 0.048546205 | -3.5751131 | Cytochrome P450 | elongation zone | 12 h |
| HORVU.MOREX.r3.2HG0202670.1 | -0.28364566 | 8.97883709 | -2.916950289 | 0.00517 | 0.045490951 | -3.57983942 | Plasma membrane ATPase | elongation zone | 12 h |
| HORVU.MOREX.r3.3HG0222130.1 | 0.244033212 | 8.795276054 | 2.848139225 | 0.00625 | 0.049750126 | -3.62747935 | Transmembrane protein 214 | elongation zone | 12 h |
| HORVU.MOREX.r3.7HG0723600.1 | -0.5677823 | 3.100437987 | -2.857416629 | 0.00609 | 0.049158681 | -3.64420562 | Peroxidase | elongation zone | 12 h |
