## Supporting Information Table 6 for "ENHANCED GRAVITROPISM 2 coordinates molecular adaptations to gravistimulation in the elongation zone of barley roots"

**Table S6a** Enriched biological processes terms among genes differentially expressed between wild type and *egt2* (FDR <5%) root caps after a gravistimulation time-course experiment.

| Go term | description | time points 0h | 3h | 6h | 12h |
|---|---|---|---|---|---|
| GO:0031047 | gene silencing by RNA | up | | | |
| GO:0031122 | cytoplasmic microtubule organization | up | up | up | up |
| GO:0030865 | cortical cytoskeleton organization | up | up | up | up |
| GO:0010091 | trichome branching | up | up | up | up |
| GO:0005978 | glycogen biosynthetic process | up | | up | up |
| GO:0016049 | cell growth | up | up | | |
| GO:0009832 | plant-type cell wall biogenesis | up | up | up | up |
| GO:0006730 | one-carbon metabolic process | down | down | down | |
| GO:0010951 | negative regulation of endopeptidase act... | down | | | |
| GO:0009611 | response to wounding | down | | | |
| GO:0006979 | response to oxidative stress | | up | | |
| GO:0042744 | hydrogen peroxide catabolic process | | up | | |
| GO:0006511 | ubiquitin-dependent protein catabolic pr... | | | up | |
| GO:0006829 | zinc ion transport | | | up | |
| GO:0039651 | induction by virus of host cysteine-type endopeptidase activity involved in apoptotic process | | | up | |
| GO:0075732 | viral penetration into host nucleus | | | up | |
| GO:0009231 | riboflavin biosynthetic process | | | up | up |
| GO:0006839 | mitochondrial transport | | | up | |
| GO:0090502 | RNA phosphodiester bond hydrolysis, endonucleolytic | | | up | |
| GO:0009606 | tropism | | | | up |
| GO:0019882 | antigen processing and presentation | | | | up |
| GO:1990542 | mitochondrial transmembrane transport | | | | up |
| GO:0051603 | proteolysis involved in cellular protein catabolic process | | | | up |
| GO:0009637 | response to blue light | | | | up |
| GO:0006004 | fucose metabolic process | | | | up |
| GO:0044743 | protein transmembrane import into intracellular organelle | | | | up |
| GO:0006626 | protein targeting to mitochondrion | | | | up |
| GO:0065002 | intracellular protein transmembrane transport | | | | up |
| GO:0006354 | DNA-templated transcription, elongation | | | | up |
| GO:0009800 | cinnamic acid biosynthetic process | | | | down |
| GO:0006559 | L-phenylalanine catabolic process | | | | down |

up-regulated
down-regulated

**Table S6b** Enriched biological processes terms among genes differentially expressed between wild type and *egt2* (FDR <5%) meristems after a gravistimulation time-course experiment.

| **Go term** | **description** | time points | | | |
|---|---|---|---|---|---|
| | | 0h | 3h | 6h | 12h |
| GO:0031122 | cytoplasmic microtubule organization | [red] | [red] | [red] | [red] |
| GO:0030865 | cortical cytoskeleton organization | [red] | [red] | [red] | [red] |
| GO:0010091 | trichome branching | [red] | [red] | [red] | [red] |
| GO:0009231 | riboflavin biosynthetic process | [red] | [grey] | [grey] | [grey] |
| GO:0016049 | cell growth | [red] | [red] | [red] | [red] |
| GO:0005978 | glycogen biosynthetic process | [red] | [grey] | [red] | [red] |
| GO:0009832 | plant-type cell wall biogenesis | [red] | [red] | [red] | [red] |
| GO:0006730 | one-carbon metabolic process | [blue] | [blue] | [blue] | [blue] |
| GO:0006979 | response to oxidative stress | [grey] | [red] | [grey] | [grey] |
| GO:0042744 | hydrogen peroxide catabolic process | [grey] | [red] | [grey] | [grey] |
| GO:0006511 | ubiquitin-dependent protein catabolic process | [grey] | [red] | [grey] | [grey] |
| GO:0016567 | protein ubiquitination | [grey] | [red] | [grey] | [grey] |
| GO:0006352 | DNA-templated transcription, initiation | [grey] | [grey] | [red] | [red] |
| GO:0006413 | translational initiation | [grey] | [grey] | [red] | [red] |
| GO:0006813 | potassium ion transport | [grey] | [grey] | [grey] | [red] |
| GO:0030488 | tRNA methylation | [grey] | [grey] | [grey] | [red] |
| GO:0030001 | metal ion transport | [grey] | [grey] | [grey] | [blue] |

[red] up-regulated
[blue] down-regulated

**Table S6c** Enriched biological processes terms among genes differentially expressed between wild type and *egt2* (FDR <5%) elongation zones after a gravistimulation time-course experiment.

| Go term | description | time points | | | |
|---|---|---|---|---|---|
| | | 0h | 3h | 6h | 12h |
| GO:0006352 | DNA-templated transcription, initiation | up | up | up | |
| GO:0006813 | potassium ion transport | up | | | |
| GO:0010091 | trichome branching | up | up | up | |
| GO:0031122 | cytoplasmic microtubule organization | up | up | up | |
| GO:0030865 | cortical cytoskeleton organization | up | up | | |
| GO:0016049 | cell growth | up | up | | |
| GO:0009832 | plant-type cell wall biogenesis | up | up | up | |
| GO:0006308 | DNA catabolic process | up | | | |
| GO:0006413 | translational initiation | up | up | up | |
| GO:0006458 | 'de novo' protein folding | down | down | | up |
| GO:0010951 | negative regulation of endopeptidase activity | down | down | | down |
| GO:0019511 | peptidyl-proline hydroxylation | down | | up | down |
| GO:1990542 | mitochondrial transmembrane transport | down | down | | up |
| GO:0006730 | one-carbon metabolic process | down | down | | up |
| GO:0044743 | protein transmembrane import into intracellular organelle | down | down | | up |
| GO:0006626 | protein targeting to mitochondrion | down | down | | up |
| GO:0065002 | intracellular protein transmembrane transport | down | down | | up |
| GO:0006433 | prolyl-tRNA aminoacylation | | up | up | |
| GO:0070588 | calcium ion transmembrane transport | | down | | |
| GO:0009744 | response to sucrose | | | up | up |
| GO:0009664 | plant-type cell wall organization | | | up and down | |
| GO:0045489 | pectin biosynthetic process | | | up | up |
| GO:0070592 | cell wall polysaccharide biosynthetic process | | | up | |
| GO:0006457 | protein folding | | | up | up |
| GO:0032507 | maintenance of protein location in cell | | | up | |
| GO:0006886 | intracellular protein transport | | | up | |
| GO:0072595 | maintenance of protein localization in organelle | | | up | |
| GO:0046149 | pigment catabolic process | | | up | |
| GO:0006099 | tricarboxylic acid cycle | | | up | |
| GO:0008299 | isoprenoid biosynthetic process | | | up | |
| GO:0006636 | unsaturated fatty acid biosynthetic process | | | up | |
| GO:0016126 | sterol biosynthetic process | | | up | |
| GO:0001676 | long-chain fatty acid metabolic process | | | up | |
| GO:0019285 | glycine betaine biosynthetic process from choline | | | up | |
| GO:0006487 | protein N-linked glycosylation | | | up | |
| GO:0006004 | fucose metabolic process | | | up | down |
| GO:0006044 | N-acetylglucosamine metabolic process | | | up | |
| GO:0007030 | Golgi organization | | | up | |
| GO:0048532 | anatomical structure arrangement | | | up | |
| GO:0010016 | shoot system morphogenesis | | | up | |
| GO:0006637 | acyl-CoA metabolic process | | | up | |
| GO:0043101 | purine-containing compound salvage | | | up | |
| GO:0009100 | glycoprotein metabolic process | | | up | |
| GO:0006979 | response to oxidative stress | | | down | down |
| GO:0042744 | hydrogen peroxide catabolic process | | | down | down |
| GO:0048146 | positive regulation of fibroblast proliferation | | | down | |
| GO:0071805 | potassium ion transmembrane transport | | | down | |
| GO:0034220 | ion transmembrane transport | | | down | |
| GO:0016119 | carotene metabolic process | | | down | |
| GO:0010411 | xyloglucan metabolic process | | | down | |
| GO:0042546 | cell wall biogenesis | | | down | |
| GO:0005975 | carbohydrate metabolic process | | | down | |
| GO:0035377 | transepithelial water transport | | | down | |
| GO:0006468 | protein phosphorylation | | | down | down |
| GO:0032941 | secretion by tissue | | | down | |
| GO:0031960 | response to corticosteroid | | | down | |
| GO:0021670 | lateral ventricle development | | | down | |
| GO:0006696 | ergosterol biosynthetic process | | | down | |
| GO:0071474 | cellular hyperosmotic response | | | down | |
| GO:0046689 | response to mercury ion | | | down | |
| GO:0046274 | lignin catabolic process | | | down | down |

up-regulated
down-regulated
up and down-regulation

**Table S6c** Continued.

| Go term | description | time points | | | |
|---|---|---|---|---|---|
| | | 0h | 3h | 6h | 12h |
| GO:0043525 | positive regulation of neuron apoptotic process | | | | |
| GO:0006359 | regulation of transcription by RNA polymerase III | | | | |
| GO:0009737 | response to abscisic acid | | | | |
| GO:0071369 | cellular response to ethylene stimulus | | | | |
| GO:0002183 | cytoplasmic translational initiation | | | | |
| GO:0006913 | nucleocytoplasmic transport | | | | |
| GO:0007018 | microtubule-based movement | | | | |
| GO:0042254 | ribosome biogenesis | | | | |
| GO:0042274 | ribosomal small subunit biogenesis | | | | |
| GO:0006364 | rRNA processing | | | | |
| GO:0009451 | RNA modification | | | | |
| GO:0006189 | 'de novo' IMP biosynthetic process | | | | |
| GO:0009116 | nucleoside metabolic process | | | | |
| GO:0009113 | purine nucleobase biosynthetic process | | | | |
| GO:0009165 | nucleotide biosynthetic process | | | | |
| GO:0006541 | glutamine metabolic process | | | | |
| GO:0006529 | asparagine biosynthetic process | | | | |
| GO:0006542 | glutamine biosynthetic process | | | | |
| GO:0009089 | lysine biosynthetic process via diaminopimelate | | | | |
| GO:0016575 | histone deacetylation | | | | |
| GO:0006260 | DNA replication | | | | |
| GO:0006270 | DNA replication initiation | | | | |
| GO:0006271 | DNA strand elongation involved in DNA replication | | | | |
| GO:0006269 | DNA replication, synthesis of RNA primer | | | | |
| GO:0009658 | chloroplast organization | | | | |
| GO:0018022 | peptidyl-lysine methylation | | | | |
| GO:0051567 | histone H3-K9 methylation | | | | |
| GO:0030488 | tRNA methylation | | | | |
| GO:0034227 | tRNA thio-modification | | | | |
| GO:0006418 | tRNA aminoacylation for protein translation | | | | |
| GO:0002098 | tRNA wobble uridine modification | | | | |
| GO:0006412 | translation | | | | |
| GO:0010039 | response to iron ion | | | | |
| GO:0046654 | tetrahydrofolate biosynthetic process | | | | |
| GO:0009231 | riboflavin biosynthetic process | | | | |
| GO:0016579 | protein deubiquitination | | | | |
| GO:0043628 | ncRNA 3'-end processing | | | | |
| GO:0006265 | DNA topological change | | | | |
| GO:0048285 | organelle fission | | | | |
| GO:0031509 | subtelomeric heterochromatin assembly | | | | |
| GO:0006334 | nucleosome assembly | | | | |
| GO:0030466 | silent mating-type cassette heterochromatin assembly | | | | |
| GO:0007076 | mitotic chromosome condensation | | | | |
| GO:0032508 | DNA duplex unwinding | | | | |
| GO:0000183 | rDNA heterochromatin assembly | | | | |
| GO:0030261 | chromosome condensation | | | | |
| GO:0006414 | translational elongation | | | | |
| GO:0006302 | double-strand break repair | | | | |
| GO:0000377 | nucleophile | | | | |
| GO:0071549 | cellular response to dexamethasone stimulus | | | | |
| GO:0007178 | transmembrane receptor protein serine/threonine kinase signaling pathway | | | | |
| GO:0009103 | lipopolysaccharide biosynthetic process | | | | |
