## Supporting Information Table 7 for "ENHANCED GRAVITROPISM 2 coordinates molecular adaptations to gravistimulation in the elongation zone of barley roots"

**Table S7** Overlapping genes among differentially expressed genes in the wild type time course experiment and *egt2* vs wild type comparisons.

| ID | WT_3 h | WT_6 h | WT_12 h | *egt2* vsWT_0 h | *egt2* vsWT_3 h | *egt* 2vsWT_6 h | *egt2* vsWT_12 h | root zone |
|---|---|---|---|---|---|---|---|---|
| HORVU.MOREX.r3.1HG0003270.1 | yes | | yes | | | yes | | root cap |
| HORVU.MOREX.r3.1HG0062030.1 | | | yes | | | yes | | root cap |
| HORVU.MOREX.r3.1HG0092900.1 | | | yes | | | | yes | root cap |
| HORVU.MOREX.r3.2HG0113590.1 | | | yes | | | yes | yes | root cap |
| HORVU.MOREX.r3.2HG0125600.1 | | | yes | | | | yes | root cap |
| HORVU.MOREX.r3.2HG0197550.1 | | | yes | | | | yes | root cap |
| HORVU.MOREX.r3.4HG0382640.1 | | | yes | | | | yes | root cap |
| HORVU.MOREX.r3.5HG0455240.1 | | | yes | | | | yes | root cap |
| HORVU.MOREX.r3.5HG0495750.1 | | | yes | | | | yes | root cap |
| HORVU.MOREX.r3.6HG0548800.1 | | | yes | | | | yes | root cap |
| HORVU.MOREX.r3.7HG0635050.1 | | | yes | | | | yes | root cap |
| HORVU.MOREX.r3.7HG0663550.1 | | | yes | yes | | | | root cap |
| HORVU.MOREX.r3.7HG0725940.1 | | | yes | | | | yes | root cap |
| HORVU.MOREX.r3.7HG0752370.1 | | | yes | | | | yes | elongation zone |
| HORVU.MOREX.r3.7HG0751110.1 | | | yes | | | yes | yes | elongation zone |
| HORVU.MOREX.r3.7HG0751060.1 | | yes | | | | yes | | elongation zone |
| HORVU.MOREX.r3.7HG0750000.1 | | yes | yes | | | yes | yes | elongation zone |
| HORVU.MOREX.r3.7HG0749870.1 | | | yes | | | yes | yes | elongation zone |
| HORVU.MOREX.r3.7HG0749150.1 | | | yes | | | | yes | elongation zone |
| HORVU.MOREX.r3.7HG0748670.1 | | yes | | | | yes | | elongation zone |
| HORVU.MOREX.r3.7HG0748070.1 | | yes | yes | | | yes | yes | elongation zone |
| HORVU.MOREX.r3.7HG0747940.1 | | yes | yes | | | | yes | elongation zone |
| HORVU.MOREX.r3.7HG0747230.1 | | yes | yes | | | yes | | elongation zone |
| HORVU.MOREX.r3.7HG0746770.1 | | yes | yes | | | yes | yes | elongation zone |
| HORVU.MOREX.r3.7HG0744540.1 | | | yes | | | | yes | elongation zone |
| HORVU.MOREX.r3.7HG0743900.1 | | | yes | | | | yes | elongation zone |
| HORVU.MOREX.r3.7HG0742080.1 | | yes | yes | | | yes | | elongation zone |
| HORVU.MOREX.r3.7HG0739910.1 | | yes | yes | yes | yes | yes | yes | elongation zone |
| HORVU.MOREX.r3.7HG0739640.1 | | | yes | | | | yes | elongation zone |
| HORVU.MOREX.r3.7HG0739490.1 | | | yes | | | | yes | elongation zone |
| HORVU.MOREX.r3.7HG0739480.1 | | | yes | | | | yes | elongation zone |
| HORVU.MOREX.r3.7HG0739230.1 | | yes | yes | | | yes | | elongation zone |
| HORVU.MOREX.r3.7HG0739030.1 | | | yes | | | yes | | elongation zone |
| HORVU.MOREX.r3.7HG0737340.1 | | yes | yes | | | yes | | elongation zone |
| HORVU.MOREX.r3.7HG0736960.1 | | yes | yes | | | yes | yes | elongation zone |

up-regulated

down-regualted

**Table S7** Continued.

| ID | WT_3 h | WT_6 h | WT_12 h | *egt2* vsWT_0 h | *egt2* vsWT_3 h | *egt* 2vsWT_6 h | *egt2* vsWT_12 h | root zone |
|---|---|---|---|---|---|---|---|---|
| HORVU.MOREX.r3.7HG0736870.1 | | | yes | | | | yes | elongation zone |
| HORVU.MOREX.r3.7HG0736860.1 | | | yes | | | | yes | elongation zone |
| HORVU.MOREX.r3.7HG0736150.1 | | yes | yes | | | yes | | elongation zone |
| HORVU.MOREX.r3.7HG0735840.1 | | yes | yes | | | yes | | elongation zone |
| HORVU.MOREX.r3.7HG0734670.1 | | yes | yes | | | yes | | elongation zone |
| HORVU.MOREX.r3.7HG0732610.1 | | | yes | | | | yes | elongation zone |
| HORVU.MOREX.r3.7HG0731560.1 | | yes | | | | yes | | elongation zone |
| HORVU.MOREX.r3.7HG0731320.1 | | yes | | | | yes | | elongation zone |
| HORVU.MOREX.r3.7HG0730510.1 | | yes | yes | | | yes | yes | elongation zone |
| HORVU.MOREX.r3.7HG0730070.1 | | | yes | | | | yes | elongation zone |
| HORVU.MOREX.r3.7HG0729150.1 | | | yes | yes | yes | yes | yes | elongation zone |
| HORVU.MOREX.r3.7HG0728630.1 | | | yes | | | | yes | elongation zone |
| HORVU.MOREX.r3.7HG0728190.1 | | | yes | | | yes | | elongation zone |
| HORVU.MOREX.r3.7HG0728060.1 | | yes | yes | | | yes | yes | elongation zone |
| HORVU.MOREX.r3.7HG0728000.1 | | yes | yes | | | yes | yes | elongation zone |
| HORVU.MOREX.r3.7HG0727460.1 | | | yes | | | | yes | elongation zone |
| HORVU.MOREX.r3.7HG0727350.1 | | | yes | | | yes | | elongation zone |
| HORVU.MOREX.r3.7HG0727290.1 | | | yes | | | | yes | elongation zone |
| HORVU.MOREX.r3.7HG0726790.1 | | yes | yes | | | | yes | elongation zone |
| HORVU.MOREX.r3.7HG0726770.1 | | yes | yes | | | yes | yes | elongation zone |
| HORVU.MOREX.r3.7HG0726700.1 | | | yes | | | | yes | elongation zone |
| HORVU.MOREX.r3.7HG0725060.1 | | | yes | | | | yes | elongation zone |
| HORVU.MOREX.r3.7HG0724060.1 | | yes | yes | | | | yes | elongation zone |
| HORVU.MOREX.r3.7HG0722360.1 | | yes | yes | | | yes | | elongation zone |
| HORVU.MOREX.r3.7HG0721930.1 | | | yes | | | | yes | elongation zone |
| HORVU.MOREX.r3.7HG0721760.1 | | | yes | | | yes | yes | elongation zone |
| HORVU.MOREX.r3.7HG0721320.1 | | yes | | | | yes | | elongation zone |
| HORVU.MOREX.r3.7HG0720790.1 | | yes | yes | | | yes | yes | elongation zone |
| HORVU.MOREX.r3.7HG0720670.1 | | | yes | | | | yes | elongation zone |
| HORVU.MOREX.r3.7HG0719410.1 | | | yes | | | | yes | elongation zone |
| HORVU.MOREX.r3.7HG0719170.1 | | | yes | | | | yes | elongation zone |
| HORVU.MOREX.r3.7HG0719070.1 | | | yes | | | | yes | elongation zone |
| HORVU.MOREX.r3.7HG0718540.1 | | | yes | | | | yes | elongation zone |
| HORVU.MOREX.r3.7HG0718190.1 | | yes | yes | | | yes | yes | elongation zone |
| HORVU.MOREX.r3.7HG0718170.1 | | | yes | | | yes | | elongation zone |

**Table S7** Continued.

| ID | WT_3 h | WT_6 h | WT_12 h | *egt2* vsWT_0 h | *egt2* vsWT_3 h | *egt* 2vsWT_6 h | *egt2* vsWT_12 h | root zone |
|---|---|---|---|---|---|---|---|---|
| HORVU.MOREX.r3.7HG0717610.1 | | yes | yes | | | yes | yes | elongation zone |
| HORVU.MOREX.r3.7HG0716900.1 | | | yes | | | yes | yes | elongation zone |
| HORVU.MOREX.r3.7HG0715570.1 | | yes | yes | | | yes | yes | elongation zone |
| HORVU.MOREX.r3.7HG0714910.1 | | | yes | | | | yes | elongation zone |
| HORVU.MOREX.r3.7HG0714900.1 | | | yes | | | | yes | elongation zone |
| HORVU.MOREX.r3.7HG0714660.1 | | yes | | | | yes | | elongation zone |
| HORVU.MOREX.r3.7HG0714510.1 | | | yes | | | | yes | elongation zone |
| HORVU.MOREX.r3.7HG0714400.1 | | yes | yes | | | yes | | elongation zone |
| HORVU.MOREX.r3.7HG0712850.1 | | | yes | | | yes | yes | elongation zone |
| HORVU.MOREX.r3.7HG0711890.1 | | yes | yes | | | yes | yes | elongation zone |
| HORVU.MOREX.r3.7HG0709860.1 | | | yes | | | | yes | elongation zone |
| HORVU.MOREX.r3.7HG0709230.1 | | | yes | | | | yes | elongation zone |
| HORVU.MOREX.r3.7HG0708820.1 | | | yes | | | | yes | elongation zone |
| HORVU.MOREX.r3.7HG0708120.1 | | | yes | | | | yes | elongation zone |
| HORVU.MOREX.r3.7HG0708050.1 | | | yes | | | | yes | elongation zone |
| HORVU.MOREX.r3.7HG0707700.1 | | yes | yes | | | yes | | elongation zone |
| HORVU.MOREX.r3.7HG0706330.1 | | | yes | | | | yes | elongation zone |
| HORVU.MOREX.r3.7HG0706310.1 | | | yes | | | yes | | elongation zone |
| HORVU.MOREX.r3.7HG0705860.1 | | | yes | | | | yes | elongation zone |
| HORVU.MOREX.r3.7HG0704450.1 | | yes | yes | | | yes | | elongation zone |
| HORVU.MOREX.r3.7HG0704060.1 | | yes | | | | | yes | elongation zone |
| HORVU.MOREX.r3.7HG0703580.1 | | | yes | | | | yes | elongation zone |
| HORVU.MOREX.r3.7HG0702190.1 | | | yes | | | | yes | elongation zone |
| HORVU.MOREX.r3.7HG0702050.1 | | | yes | | | | yes | elongation zone |
| HORVU.MOREX.r3.7HG0700930.1 | | | yes | | | | yes | elongation zone |
| HORVU.MOREX.r3.7HG0700270.1 | | yes | yes | | | yes | | elongation zone |
| HORVU.MOREX.r3.7HG0697940.1 | | | yes | | | | yes | elongation zone |
| HORVU.MOREX.r3.7HG0696190.1 | | | yes | | | | yes | elongation zone |
| HORVU.MOREX.r3.7HG0690090.1 | | | yes | | | | yes | elongation zone |
| HORVU.MOREX.r3.7HG0689490.1 | | | yes | | | | yes | elongation zone |
| HORVU.MOREX.r3.7HG0688870.1 | | yes | yes | | | yes | | elongation zone |
| HORVU.MOREX.r3.7HG0687630.1 | | yes | yes | | | yes | yes | elongation zone |
| HORVU.MOREX.r3.7HG0685970.1 | | | yes | | | | yes | elongation zone |
| HORVU.MOREX.r3.7HG0685920.1 | | | yes | | | yes | yes | elongation zone |
| HORVU.MOREX.r3.7HG0684880.1 | | yes | yes | | | yes | yes | elongation zone |

**Table S7** Continued.

| ID | WT_3 h | WT_6 h | WT_12 h | *egt2* vsWT_0 h | *egt2* vsWT_3 h | *egt* 2vsWT_6 h | *egt2* vsWT_12 h | root zone |
|---|---|---|---|---|---|---|---|---|
| HORVU.MOREX.r3.7HG0680960.1 | | yes | yes | | | yes | yes | elongation zone |
| HORVU.MOREX.r3.7HG0679350.1 | | yes | yes | | | yes | | elongation zone |
| HORVU.MOREX.r3.7HG0679060.1 | | | yes | | | | yes | elongation zone |
| HORVU.MOREX.r3.7HG0679050.1 | | yes | yes | | | yes | yes | elongation zone |
| HORVU.MOREX.r3.7HG0677860.1 | | | yes | | | | yes | elongation zone |
| HORVU.MOREX.r3.7HG0677580.1 | | | yes | | | | yes | elongation zone |
| HORVU.MOREX.r3.7HG0677510.1 | | yes | yes | | | yes | | elongation zone |
| HORVU.MOREX.r3.7HG0677230.1 | | | yes | | | | yes | elongation zone |
| HORVU.MOREX.r3.7HG0676040.1 | | yes | | | | yes | yes | elongation zone |
| HORVU.MOREX.r3.7HG0674990.1 | | yes | yes | | | yes | | elongation zone |
| HORVU.MOREX.r3.7HG0674750.1 | | | yes | | | | yes | elongation zone |
| HORVU.MOREX.r3.7HG0674720.1 | | yes | yes | | | yes | yes | elongation zone |
| HORVU.MOREX.r3.7HG0674590.1 | | | yes | | | | yes | elongation zone |
| HORVU.MOREX.r3.7HG0674480.1 | | | yes | | | | yes | elongation zone |
| HORVU.MOREX.r3.7HG0674320.1 | | yes | yes | | | yes | | elongation zone |
| HORVU.MOREX.r3.7HG0671880.1 | | | yes | | | | yes | elongation zone |
| HORVU.MOREX.r3.7HG0670020.1 | | | yes | | | | yes | elongation zone |
| HORVU.MOREX.r3.7HG0669580.1 | | | yes | | | | yes | elongation zone |
| HORVU.MOREX.r3.7HG0668820.1 | | yes | yes | | | yes | | elongation zone |
| HORVU.MOREX.r3.7HG0667730.1 | | | yes | | | | yes | elongation zone |
| HORVU.MOREX.r3.7HG0667060.1 | | yes | | | | yes | | elongation zone |
| HORVU.MOREX.r3.7HG0666340.1 | | | yes | | | yes | yes | elongation zone |
| HORVU.MOREX.r3.7HG0665990.1 | | yes | yes | | | yes | yes | elongation zone |
| HORVU.MOREX.r3.7HG0665040.1 | | yes | | | | yes | | elongation zone |
| HORVU.MOREX.r3.7HG0664720.1 | | yes | yes | | | yes | | elongation zone |
| HORVU.MOREX.r3.7HG0664130.1 | | | yes | | | | yes | elongation zone |
| HORVU.MOREX.r3.7HG0663940.1 | | yes | yes | | | yes | yes | elongation zone |
| HORVU.MOREX.r3.7HG0663730.1 | | | yes | | | | yes | elongation zone |
| HORVU.MOREX.r3.7HG0662830.1 | | | yes | | | | yes | elongation zone |
| HORVU.MOREX.r3.7HG0662210.1 | | | yes | | | | yes | elongation zone |
| HORVU.MOREX.r3.7HG0662160.1 | | | yes | | | | yes | elongation zone |
| HORVU.MOREX.r3.7HG0661890.1 | | | yes | | | yes | yes | elongation zone |
| HORVU.MOREX.r3.7HG0661490.1 | | yes | yes | | | yes | | elongation zone |
| HORVU.MOREX.r3.7HG0661480.1 | | yes | yes | | | yes | | elongation zone |
| HORVU.MOREX.r3.7HG0658560.1 | | yes | | | | yes | | elongation zone |

**Table S7** Continued.

| ID | WT_3 h | WT_6 h | WT_12 h | *egt2* vsWT_0 h | *egt2* vsWT_3 h | *egt* 2vsWT_6 h | *egt2* vsWT_12 h | root zone |
|---|---|---|---|---|---|---|---|---|
| HORVU.MOREX.r3.7HG0656440.1 | | | yes | | | | yes | elongation zone |
| HORVU.MOREX.r3.7HG0656430.1 | | | yes | | | | yes | elongation zone |
| HORVU.MOREX.r3.7HG0656030.1 | | | yes | | | | yes | elongation zone |
| HORVU.MOREX.r3.7HG0655210.1 | | yes | yes | | | yes | yes | elongation zone |
| HORVU.MOREX.r3.7HG0654410.1 | | | yes | | | | yes | elongation zone |
| HORVU.MOREX.r3.7HG0652830.1 | | | yes | | | | yes | elongation zone |
| HORVU.MOREX.r3.7HG0650970.1 | | | yes | | | | yes | elongation zone |
| HORVU.MOREX.r3.7HG0650300.1 | | | yes | | | | yes | elongation zone |
| HORVU.MOREX.r3.7HG0649950.1 | | yes | yes | | | yes | yes | elongation zone |
| HORVU.MOREX.r3.7HG0649240.1 | | yes | yes | | | yes | | elongation zone |
| HORVU.MOREX.r3.7HG0648600.1 | | | yes | | | | yes | elongation zone |
| HORVU.MOREX.r3.7HG0648460.1 | | | yes | | | | yes | elongation zone |
| HORVU.MOREX.r3.7HG0647950.1 | | yes | | | | yes | | elongation zone |
| HORVU.MOREX.r3.7HG0647930.1 | | | yes | | | | yes | elongation zone |
| HORVU.MOREX.r3.7HG0647520.1 | | | yes | | | | yes | elongation zone |
| HORVU.MOREX.r3.7HG0642520.1 | | yes | yes | | | yes | | elongation zone |
| HORVU.MOREX.r3.7HG0641750.1 | | yes | yes | | | yes | yes | elongation zone |
| HORVU.MOREX.r3.7HG0641160.1 | | | yes | | | | yes | elongation zone |
| HORVU.MOREX.r3.7HG0640560.1 | | yes | yes | | | | yes | elongation zone |
| HORVU.MOREX.r3.7HG0636750.1 | | | yes | | | | yes | elongation zone |
| HORVU.MOREX.r3.7HG0636630.1 | | yes | yes | | | yes | yes | elongation zone |
| HORVU.MOREX.r3.7HG0635550.1 | | | yes | | | | yes | elongation zone |
| HORVU.MOREX.r3.7HG0635160.1 | | | yes | | | | yes | elongation zone |
| HORVU.MOREX.r3.7HG0634520.1 | | yes | yes | | | yes | | elongation zone |
| HORVU.MOREX.r3.6HG0634260.1 | | | yes | | | | yes | elongation zone |
| HORVU.MOREX.r3.6HG0633420.1 | | | yes | | | | yes | elongation zone |
| HORVU.MOREX.r3.6HG0632980.1 | | | yes | | | | yes | elongation zone |
| HORVU.MOREX.r3.6HG0632320.1 | | yes | yes | | | yes | yes | elongation zone |
| HORVU.MOREX.r3.6HG0632270.1 | | yes | yes | | | yes | | elongation zone |
| HORVU.MOREX.r3.6HG0632020.2 | | | yes | | | | yes | elongation zone |
| HORVU.MOREX.r3.6HG0631760.1 | | | yes | | | | yes | elongation zone |
| HORVU.MOREX.r3.6HG0631710.1 | | yes | yes | | | yes | | elongation zone |
| HORVU.MOREX.r3.6HG0631400.1 | | | yes | | | | yes | elongation zone |
| HORVU.MOREX.r3.6HG0630650.1 | | | yes | | | | yes | elongation zone |
| HORVU.MOREX.r3.6HG0628930.1 | | | yes | | | | yes | elongation zone |

**Table S7** Continued.

| ID | WT_3 h | WT_6 h | WT_12 h | *egt2* vsWT_0 h | *egt2* vsWT_3 h | *egt* 2vsWT_6 h | *egt2* vsWT_12 h | root zone |
|---|---|---|---|---|---|---|---|---|
| HORVU.MOREX.r3.6HG0628790.1 | | | yes | | | yes | yes | elongation zone |
| HORVU.MOREX.r3.6HG0627570.1 | | yes | yes | | | yes | | elongation zone |
| HORVU.MOREX.r3.6HG0627090.1 | | | yes | | | | yes | elongation zone |
| HORVU.MOREX.r3.6HG0626650.1 | | yes | yes | | | yes | yes | elongation zone |
| HORVU.MOREX.r3.6HG0626030.1 | | yes | yes | | | yes | | elongation zone |
| HORVU.MOREX.r3.6HG0626020.1 | | yes | yes | | | yes | | elongation zone |
| HORVU.MOREX.r3.6HG0625570.1 | | yes | yes | | | | yes | elongation zone |
| HORVU.MOREX.r3.6HG0624650.1 | | | yes | | | yes | | elongation zone |
| HORVU.MOREX.r3.6HG0624630.1 | | | yes | | | | yes | elongation zone |
| HORVU.MOREX.r3.6HG0624580.1 | | | yes | | | | yes | elongation zone |
| HORVU.MOREX.r3.6HG0624390.1 | | yes | yes | | | yes | | elongation zone |
| HORVU.MOREX.r3.6HG0623920.1 | | | yes | | | | yes | elongation zone |
| HORVU.MOREX.r3.6HG0623830.1 | | | yes | | | | yes | elongation zone |
| HORVU.MOREX.r3.6HG0622740.1 | | yes | | | | yes | | elongation zone |
| HORVU.MOREX.r3.6HG0621690.1 | | | yes | | | | yes | elongation zone |
| HORVU.MOREX.r3.6HG0620520.1 | | yes | yes | | | | yes | elongation zone |
| HORVU.MOREX.r3.6HG0619540.1 | | | yes | | | | yes | elongation zone |
| HORVU.MOREX.r3.6HG0618170.1 | | | yes | | | | yes | elongation zone |
| HORVU.MOREX.r3.6HG0617080.1 | | | yes | | | | yes | elongation zone |
| HORVU.MOREX.r3.6HG0616960.1 | | | yes | | | | yes | elongation zone |
| HORVU.MOREX.r3.6HG0616790.1 | | | yes | | | | yes | elongation zone |
| HORVU.MOREX.r3.6HG0616480.1 | | | yes | | | | yes | elongation zone |
| HORVU.MOREX.r3.6HG0616300.1 | | | yes | | | yes | yes | elongation zone |
| HORVU.MOREX.r3.6HG0616240.1 | | yes | yes | | | yes | | elongation zone |
| HORVU.MOREX.r3.6HG0616130.1 | | | yes | | | | yes | elongation zone |
| HORVU.MOREX.r3.6HG0615840.1 | | yes | yes | | | yes | | elongation zone |
| HORVU.MOREX.r3.6HG0615710.1 | | | yes | | | | yes | elongation zone |
| HORVU.MOREX.r3.6HG0615130.1 | | yes | yes | | | yes | | elongation zone |
| HORVU.MOREX.r3.6HG0614840.1 | | | yes | | | | yes | elongation zone |
| HORVU.MOREX.r3.6HG0614500.1 | | | yes | | | | yes | elongation zone |
| HORVU.MOREX.r3.6HG0614110.1 | | | yes | | | | yes | elongation zone |
| HORVU.MOREX.r3.6HG0613230.1 | | yes | yes | | | yes | yes | elongation zone |
| HORVU.MOREX.r3.6HG0612180.1 | | | yes | | | yes | yes | elongation zone |
| HORVU.MOREX.r3.6HG0611290.1 | | | yes | | | | yes | elongation zone |
| HORVU.MOREX.r3.6HG0610980.1 | | yes | yes | | | yes | | elongation zone |

**Table S7** Continued.

| ID | WT_3 h | WT_6 h | WT_12 h | *egt2* vsWT_0 h | *egt2* vsWT_3 h | *egt* 2vsWT_6 h | *egt2* vsWT_12 h | root zone |
|---|---|---|---|---|---|---|---|---|
| HORVU.MOREX.r3.6HG0610500.1 | | yes | yes | | | yes | yes | elongation zone |
| HORVU.MOREX.r3.6HG0609720.1 | | yes | yes | | | yes | yes | elongation zone |
| HORVU.MOREX.r3.6HG0609520.1 | | yes | yes | | | | yes | elongation zone |
| HORVU.MOREX.r3.6HG0608780.1 | | yes | yes | | | yes | yes | elongation zone |
| HORVU.MOREX.r3.6HG0608390.1 | | | yes | | | | yes | elongation zone |
| HORVU.MOREX.r3.6HG0608180.1 | | | yes | | | | yes | elongation zone |
| HORVU.MOREX.r3.6HG0607590.1 | | yes | yes | | | yes | | elongation zone |
| HORVU.MOREX.r3.6HG0607360.1 | | yes | yes | | | | yes | elongation zone |
| HORVU.MOREX.r3.6HG0607000.1 | | | yes | | | | yes | elongation zone |
| HORVU.MOREX.r3.6HG0606940.1 | | | yes | | | | yes | elongation zone |
| HORVU.MOREX.r3.6HG0606930.1 | | yes | yes | | | yes | | elongation zone |
| HORVU.MOREX.r3.6HG0605940.1 | | | yes | | | yes | yes | elongation zone |
| HORVU.MOREX.r3.6HG0605290.1 | | | yes | | | | yes | elongation zone |
| HORVU.MOREX.r3.6HG0605130.1 | | | yes | | | | yes | elongation zone |
| HORVU.MOREX.r3.6HG0604860.1 | | | yes | | | | yes | elongation zone |
| HORVU.MOREX.r3.6HG0604200.1 | | yes | yes | | | | yes | elongation zone |
| HORVU.MOREX.r3.6HG0601830.1 | | yes | yes | | | yes | | elongation zone |
| HORVU.MOREX.r3.6HG0601670.1 | | yes | yes | | | yes | | elongation zone |
| HORVU.MOREX.r3.6HG0601260.1 | | yes | yes | | | yes | yes | elongation zone |
| HORVU.MOREX.r3.6HG0600910.1 | | yes | yes | | | yes | | elongation zone |
| HORVU.MOREX.r3.6HG0600000.1 | | yes | yes | | | yes | yes | elongation zone |
| HORVU.MOREX.r3.6HG0598750.1 | | | yes | | | | yes | elongation zone |
| HORVU.MOREX.r3.6HG0597640.1 | | | yes | | | | yes | elongation zone |
| HORVU.MOREX.r3.6HG0597590.1 | | yes | yes | | | yes | | elongation zone |
| HORVU.MOREX.r3.6HG0597360.1 | | | yes | | | | yes | elongation zone |
| HORVU.MOREX.r3.6HG0595860.1 | | | yes | | | | yes | elongation zone |
| HORVU.MOREX.r3.6HG0595190.1 | | yes | yes | | | yes | yes | elongation zone |
| HORVU.MOREX.r3.6HG0589860.1 | | yes | yes | | | yes | | elongation zone |
| HORVU.MOREX.r3.6HG0588350.1 | | yes | yes | | | | yes | elongation zone |
| HORVU.MOREX.r3.6HG0587460.1 | | | yes | | | | yes | elongation zone |
| HORVU.MOREX.r3.6HG0587140.1 | | | yes | | | | yes | elongation zone |
| HORVU.MOREX.r3.6HG0586570.1 | | | yes | | | | yes | elongation zone |
| HORVU.MOREX.r3.6HG0582480.1 | | yes | yes | | | yes | | elongation zone |
| HORVU.MOREX.r3.6HG0582230.1 | | | yes | | | | yes | elongation zone |
| HORVU.MOREX.r3.6HG0581020.1 | | yes | yes | | | yes | | elongation zone |

**Table S7** Continued.

| ID | WT_3 h | WT_6 h | WT_12 h | *egt2* vsWT_0 h | *egt2* vsWT_3 h | *egt* 2vsWT_6 h | *egt2* vsWT_12 h | root zone |
|---|---|---|---|---|---|---|---|---|
| HORVU.MOREX.r3.6HG0581010.1 | | | yes | | | | yes | elongation zone |
| HORVU.MOREX.r3.6HG0577220.1 | | yes | | | | yes | | elongation zone |
| HORVU.MOREX.r3.6HG0576550.1 | | yes | yes | | | | yes | elongation zone |
| HORVU.MOREX.r3.6HG0576010.1 | | yes | | | | yes | | elongation zone |
| HORVU.MOREX.r3.6HG0575530.1 | | yes | | | | yes | | elongation zone |
| HORVU.MOREX.r3.6HG0574510.1 | | | yes | | | | yes | elongation zone |
| HORVU.MOREX.r3.6HG0573870.1 | | | yes | | | | yes | elongation zone |
| HORVU.MOREX.r3.6HG0573150.1 | | | yes | | | yes | | elongation zone |
| HORVU.MOREX.r3.6HG0570960.1 | | | yes | | | | yes | elongation zone |
| HORVU.MOREX.r3.6HG0570230.1 | | yes | yes | | | yes | | elongation zone |
| HORVU.MOREX.r3.6HG0569990.1 | | | yes | | | | yes | elongation zone |
| HORVU.MOREX.r3.6HG0569490.1 | | | yes | | | | yes | elongation zone |
| HORVU.MOREX.r3.6HG0569480.1 | | | yes | | | | yes | elongation zone |
| HORVU.MOREX.r3.6HG0568910.1 | | | yes | | | | yes | elongation zone |
| HORVU.MOREX.r3.6HG0568390.1 | | | yes | | | | yes | elongation zone |
| HORVU.MOREX.r3.6HG0567780.1 | | yes | yes | | | yes | | elongation zone |
| HORVU.MOREX.r3.6HG0567330.1 | | | yes | | | | yes | elongation zone |
| HORVU.MOREX.r3.6HG0566940.1 | | yes | yes | | | yes | | elongation zone |
| HORVU.MOREX.r3.6HG0566930.1 | | | yes | | | | yes | elongation zone |
| HORVU.MOREX.r3.6HG0564690.1 | | | yes | | | | yes | elongation zone |
| HORVU.MOREX.r3.6HG0564590.1 | | | yes | | | yes | yes | elongation zone |
| HORVU.MOREX.r3.6HG0564510.1 | | | yes | | | | yes | elongation zone |
| HORVU.MOREX.r3.6HG0560290.1 | | | yes | | | | yes | elongation zone |
| HORVU.MOREX.r3.6HG0560140.1 | | yes | yes | | | yes | | elongation zone |
| HORVU.MOREX.r3.6HG0558880.1 | | | yes | | | | yes | elongation zone |
| HORVU.MOREX.r3.6HG0558800.1 | | | yes | | | | yes | elongation zone |
| HORVU.MOREX.r3.6HG0558430.1 | | yes | | | | yes | | elongation zone |
| HORVU.MOREX.r3.6HG0557920.1 | | yes | yes | | | yes | | elongation zone |
| HORVU.MOREX.r3.6HG0555700.1 | | yes | yes | | | yes | yes | elongation zone |
| HORVU.MOREX.r3.6HG0555660.2 | | yes | yes | | | yes | | elongation zone |
| HORVU.MOREX.r3.6HG0554520.1 | | | yes | | | | yes | elongation zone |
| HORVU.MOREX.r3.6HG0554050.1 | | yes | yes | | | yes | | elongation zone |
| HORVU.MOREX.r3.6HG0553310.1 | | | yes | | | | yes | elongation zone |
| HORVU.MOREX.r3.6HG0553290.1 | | | yes | | | | yes | elongation zone |
| HORVU.MOREX.r3.6HG0552860.1 | | | yes | | | | yes | elongation zone |

**Table S7** Continued.

| ID | WT_3 h | WT_6 h | WT_12 h | *egt2* vsWT_0 h | *egt2* vsWT_3 h | *egt* 2vsWT_6 h | *egt2* vsWT_12 h | root zone |
|---|---|---|---|---|---|---|---|---|
| HORVU.MOREX.r3.6HG0552800.1 | | | yes | | | | yes | elongation zone |
| HORVU.MOREX.r3.6HG0551740.1 | | | yes | | | | yes | elongation zone |
| HORVU.MOREX.r3.6HG0550950.1 | | | yes | | | | yes | elongation zone |
| HORVU.MOREX.r3.6HG0550700.1 | | | yes | | | | yes | elongation zone |
| HORVU.MOREX.r3.6HG0550670.1 | | | yes | | | | yes | elongation zone |
| HORVU.MOREX.r3.6HG0550600.1 | | yes | yes | | | yes | | elongation zone |
| HORVU.MOREX.r3.6HG0549990.1 | | yes | | | | yes | | elongation zone |
| HORVU.MOREX.r3.6HG0549510.1 | | | yes | | | | yes | elongation zone |
| HORVU.MOREX.r3.6HG0548670.1 | | | yes | | | | yes | elongation zone |
| HORVU.MOREX.r3.6HG0547570.1 | | | yes | | | | yes | elongation zone |
| HORVU.MOREX.r3.6HG0546160.1 | | yes | yes | | | | yes | elongation zone |
| HORVU.MOREX.r3.6HG0546140.1 | | | yes | | | yes | | elongation zone |
| HORVU.MOREX.r3.6HG0545970.1 | | | yes | | | yes | yes | elongation zone |
| HORVU.MOREX.r3.6HG0545630.1 | | | yes | | | | yes | elongation zone |
| HORVU.MOREX.r3.6HG0545620.1 | | yes | yes | | | yes | yes | elongation zone |
| HORVU.MOREX.r3.6HG0545460.1 | | | yes | | | | yes | elongation zone |
| HORVU.MOREX.r3.6HG0543800.1 | | | yes | | | | yes | elongation zone |
| HORVU.MOREX.r3.6HG0543780.1 | | | yes | | | | yes | elongation zone |
| HORVU.MOREX.r3.6HG0543770.1 | | | yes | | | | yes | elongation zone |
| HORVU.MOREX.r3.6HG0543740.1 | | | yes | | | | yes | elongation zone |
| HORVU.MOREX.r3.6HG0543730.1 | | | yes | | | | yes | elongation zone |
| HORVU.MOREX.r3.6HG0542970.1 | | | yes | | | | yes | elongation zone |
| HORVU.MOREX.r3.6HG0541940.1 | | yes | yes | | | yes | | elongation zone |
| HORVU.MOREX.r3.6HG0541250.1 | | yes | yes | | | yes | | elongation zone |
| HORVU.MOREX.r3.6HG0540620.1 | | | yes | | | | yes | elongation zone |
| HORVU.MOREX.r3.6HG0540280.1 | | | yes | | | | yes | elongation zone |
| HORVU.MOREX.r3.6HG0540020.1 | | yes | | | | yes | yes | elongation zone |
| HORVU.MOREX.r3.5HG0537150.1 | | yes | yes | | | yes | yes | elongation zone |
| HORVU.MOREX.r3.5HG0536710.1 | | | yes | | | yes | yes | elongation zone |
| HORVU.MOREX.r3.5HG0536490.1 | | | yes | | | | yes | elongation zone |
| HORVU.MOREX.r3.5HG0536200.1 | | yes | yes | | | yes | | elongation zone |
| HORVU.MOREX.r3.5HG0535780.1 | | | yes | | | | yes | elongation zone |
| HORVU.MOREX.r3.5HG0534640.1 | | | yes | | | | yes | elongation zone |
| HORVU.MOREX.r3.5HG0533630.1 | | | yes | | | | yes | elongation zone |
| HORVU.MOREX.r3.5HG0533230.1 | | yes | yes | | | yes | | elongation zone |

**Table S7** Continued.

| ID | WT_3 h | WT_6 h | WT_12 h | *egt2* vsWT_0 h | *egt2* vsWT_3 h | *egt* 2vsWT_6 h | *egt2* vsWT_12 h | root zone |
|---|---|---|---|---|---|---|---|---|
| HORVU.MOREX.r3.5HG0532630.1 | | | yes | | | | yes | elongation zone |
| HORVU.MOREX.r3.5HG0532150.1 | yes | yes | yes | | | yes | | elongation zone |
| HORVU.MOREX.r3.5HG0532140.1 | | yes | yes | | | yes | | elongation zone |
| HORVU.MOREX.r3.5HG0532120.1 | | yes | yes | | | yes | | elongation zone |
| HORVU.MOREX.r3.5HG0531850.1 | | | yes | | | | yes | elongation zone |
| HORVU.MOREX.r3.5HG0530750.1 | | | yes | | | | yes | elongation zone |
| HORVU.MOREX.r3.5HG0529130.1 | | | yes | | | | yes | elongation zone |
| HORVU.MOREX.r3.5HG0529120.1 | | | yes | | | | yes | elongation zone |
| HORVU.MOREX.r3.5HG0528890.1 | | | yes | | | | yes | elongation zone |
| HORVU.MOREX.r3.5HG0527890.1 | | | yes | | | | yes | elongation zone |
| HORVU.MOREX.r3.5HG0527650.1 | | | yes | | | yes | | elongation zone |
| HORVU.MOREX.r3.5HG0527310.1 | | yes | | | | yes | | elongation zone |
| HORVU.MOREX.r3.5HG0526940.1 | | | yes | | | | yes | elongation zone |
| HORVU.MOREX.r3.5HG0526250.1 | | | yes | | | yes | yes | elongation zone |
| HORVU.MOREX.r3.5HG0526200.1 | | yes | yes | | | yes | | elongation zone |
| HORVU.MOREX.r3.5HG0524060.1 | | yes | yes | | | yes | | elongation zone |
| HORVU.MOREX.r3.5HG0523860.1 | | | yes | | | yes | yes | elongation zone |
| HORVU.MOREX.r3.5HG0523150.1 | | | yes | | | | yes | elongation zone |
| HORVU.MOREX.r3.5HG0520260.1 | | | yes | | | | yes | elongation zone |
| HORVU.MOREX.r3.5HG0519810.1 | | yes | yes | | | yes | yes | elongation zone |
| HORVU.MOREX.r3.5HG0519120.1 | | yes | yes | | | yes | | elongation zone |
| HORVU.MOREX.r3.5HG0518560.1 | | yes | yes | | | yes | yes | elongation zone |
| HORVU.MOREX.r3.5HG0517740.1 | yes | yes | yes | | | yes | | elongation zone |
| HORVU.MOREX.r3.5HG0517490.1 | | | yes | | | | yes | elongation zone |
| HORVU.MOREX.r3.5HG0516720.1 | | | yes | | | | yes | elongation zone |
| HORVU.MOREX.r3.5HG0516470.1 | | | yes | | | | yes | elongation zone |
| HORVU.MOREX.r3.5HG0515610.1 | | | yes | | yes | | | elongation zone |
| HORVU.MOREX.r3.5HG0514790.3 | | | yes | | | | yes | elongation zone |
| HORVU.MOREX.r3.5HG0514790.1 | | | yes | | | | yes | elongation zone |
| HORVU.MOREX.r3.5HG0514490.1 | | | yes | | | | yes | elongation zone |
| HORVU.MOREX.r3.5HG0514110.1 | | | yes | | | | yes | elongation zone |
| HORVU.MOREX.r3.5HG0514100.1 | | | yes | | | | yes | elongation zone |
| HORVU.MOREX.r3.5HG0513980.1 | | yes | yes | | | yes | yes | elongation zone |
| HORVU.MOREX.r3.5HG0513810.1 | | | yes | | | yes | yes | elongation zone |
| HORVU.MOREX.r3.5HG0513740.1 | | | yes | | | | yes | elongation zone |

**Table S7** Continued.

| ID | WT_3 h | WT_6 h | WT_12 h | *egt2* vsWT_0 h | *egt2* vsWT_3 h | *egt* 2vsWT_6 h | *egt2* vsWT_12 h | root zone |
|---|---|---|---|---|---|---|---|---|
| HORVU.MOREX.r3.5HG0513440.1 | | | yes | | | yes | | elongation zone |
| HORVU.MOREX.r3.5HG0513020.1 | | | yes | | | yes | yes | elongation zone |
| HORVU.MOREX.r3.5HG0512510.1 | | | yes | | | | yes | elongation zone |
| HORVU.MOREX.r3.5HG0512350.1 | | | yes | | | | yes | elongation zone |
| HORVU.MOREX.r3.5HG0512230.1 | | yes | yes | | | yes | | elongation zone |
| HORVU.MOREX.r3.5HG0511820.1 | | yes | yes | | | yes | | elongation zone |
| HORVU.MOREX.r3.5HG0511450.1 | | yes | yes | | | yes | | elongation zone |
| HORVU.MOREX.r3.5HG0511090.1 | | yes | yes | | | yes | | elongation zone |
| HORVU.MOREX.r3.5HG0510940.1 | | | yes | | | | yes | elongation zone |
| HORVU.MOREX.r3.5HG0508190.1 | | | yes | | | | yes | elongation zone |
| HORVU.MOREX.r3.5HG0504160.1 | | | yes | | | | yes | elongation zone |
| HORVU.MOREX.r3.5HG0502760.1 | | | yes | | | | yes | elongation zone |
| HORVU.MOREX.r3.5HG0502750.1 | | | yes | | | | yes | elongation zone |
| HORVU.MOREX.r3.5HG0502130.1 | | | yes | | | | yes | elongation zone |
| HORVU.MOREX.r3.5HG0501980.1 | | | yes | | | yes | | elongation zone |
| HORVU.MOREX.r3.5HG0501270.1 | | | yes | | | | yes | elongation zone |
| HORVU.MOREX.r3.5HG0500680.1 | | | yes | | | yes | yes | elongation zone |
| HORVU.MOREX.r3.5HG0499490.1 | | | yes | | | | yes | elongation zone |
| HORVU.MOREX.r3.5HG0498770.1 | | | yes | | | | yes | elongation zone |
| HORVU.MOREX.r3.5HG0498150.1 | | | yes | | | | yes | elongation zone |
| HORVU.MOREX.r3.5HG0494320.1 | | | yes | | | | yes | elongation zone |
| HORVU.MOREX.r3.5HG0494040.1 | | yes | yes | | | yes | yes | elongation zone |
| HORVU.MOREX.r3.5HG0493850.1 | | | yes | | | yes | yes | elongation zone |
| HORVU.MOREX.r3.5HG0493070.1 | | | yes | | | yes | | elongation zone |
| HORVU.MOREX.r3.5HG0492650.1 | | | yes | | | | yes | elongation zone |
| HORVU.MOREX.r3.5HG0491230.1 | | | yes | | | yes | yes | elongation zone |
| HORVU.MOREX.r3.5HG0489840.1 | | | yes | | | | yes | elongation zone |
| HORVU.MOREX.r3.5HG0489130.1 | | | yes | | | | yes | elongation zone |
| HORVU.MOREX.r3.5HG0488050.1 | | yes | yes | | | yes | | elongation zone |
| HORVU.MOREX.r3.5HG0488040.1 | | yes | yes | | | yes | | elongation zone |
| HORVU.MOREX.r3.5HG0486660.1 | | yes | yes | | | yes | | elongation zone |
| HORVU.MOREX.r3.5HG0486070.1 | | | yes | | | | yes | elongation zone |
| HORVU.MOREX.r3.5HG0485610.1 | | yes | yes | | | yes | yes | elongation zone |
| HORVU.MOREX.r3.5HG0485410.1 | | yes | yes | yes | | | | elongation zone |
| HORVU.MOREX.r3.5HG0483980.1 | | yes | yes | | | yes | yes | elongation zone |

**Table S7** Continued.

| ID | WT_3 h | WT_6 h | WT_12 h | *egt2* vsWT_0 h | *egt2* vsWT_3 h | *egt* 2vsWT_6 h | *egt2* vsWT_12 h | root zone |
|---|---|---|---|---|---|---|---|---|
| HORVU.MOREX.r3.5HG0483860.1 | | yes | | | | yes | | elongation zone |
| HORVU.MOREX.r3.5HG0480980.1 | | yes | yes | | | yes | | elongation zone |
| HORVU.MOREX.r3.5HG0479210.1 | | | yes | | | | yes | elongation zone |
| HORVU.MOREX.r3.5HG0478390.1 | | | yes | | | | yes | elongation zone |
| HORVU.MOREX.r3.5HG0477750.1 | | yes | yes | | | yes | | elongation zone |
| HORVU.MOREX.r3.5HG0477180.1 | | yes | yes | | | | yes | elongation zone |
| HORVU.MOREX.r3.5HG0477040.1 | | yes | yes | | | | yes | elongation zone |
| HORVU.MOREX.r3.5HG0476380.1 | | | yes | | | yes | | elongation zone |
| HORVU.MOREX.r3.5HG0474540.1 | | yes | | | | yes | | elongation zone |
| HORVU.MOREX.r3.5HG0473970.1 | | yes | | | | yes | | elongation zone |
| HORVU.MOREX.r3.5HG0472880.1 | | | yes | | | | yes | elongation zone |
| HORVU.MOREX.r3.5HG0472840.1 | | | yes | | | | yes | elongation zone |
| HORVU.MOREX.r3.5HG0472810.1 | | yes | yes | | | yes | | elongation zone |
| HORVU.MOREX.r3.5HG0472770.1 | | yes | yes | | | yes | | elongation zone |
| HORVU.MOREX.r3.5HG0471960.1 | | | yes | | | | yes | elongation zone |
| HORVU.MOREX.r3.5HG0470170.1 | | yes | yes | | | yes | | elongation zone |
| HORVU.MOREX.r3.5HG0470030.1 | | yes | yes | | | yes | yes | elongation zone |
| HORVU.MOREX.r3.5HG0469340.1 | | yes | yes | | | yes | | elongation zone |
| HORVU.MOREX.r3.5HG0469300.1 | | yes | yes | | | yes | | elongation zone |
| HORVU.MOREX.r3.5HG0467880.1 | | | yes | | | | yes | elongation zone |
| HORVU.MOREX.r3.5HG0467870.1 | | yes | yes | | | yes | yes | elongation zone |
| HORVU.MOREX.r3.5HG0466020.1 | | | yes | | | | yes | elongation zone |
| HORVU.MOREX.r3.5HG0466010.1 | | | yes | | | | yes | elongation zone |
| HORVU.MOREX.r3.5HG0465260.1 | | | yes | | | yes | | elongation zone |
| HORVU.MOREX.r3.5HG0463350.1 | | | yes | | | | yes | elongation zone |
| HORVU.MOREX.r3.5HG0462450.1 | | | yes | | | yes | yes | elongation zone |
| HORVU.MOREX.r3.5HG0462380.1 | | | yes | yes | | | | elongation zone |
| HORVU.MOREX.r3.5HG0462220.1 | | | yes | | | yes | yes | elongation zone |
| HORVU.MOREX.r3.5HG0461950.1 | | yes | yes | | | yes | | elongation zone |
| HORVU.MOREX.r3.5HG0461830.1 | | | yes | | | | yes | elongation zone |
| HORVU.MOREX.r3.5HG0461280.1 | | yes | yes | | | yes | | elongation zone |
| HORVU.MOREX.r3.5HG0461170.1 | | | yes | | | | yes | elongation zone |
| HORVU.MOREX.r3.5HG0460720.1 | | | yes | | | | yes | elongation zone |
| HORVU.MOREX.r3.5HG0459320.1 | | | yes | | | yes | | elongation zone |
| HORVU.MOREX.r3.5HG0458890.1 | | yes | yes | | | | yes | elongation zone |

**Table S7** Continued.

| ID | WT_3 h | WT_6 h | WT_12 h | *egt2* vsWT_0 h | *egt2* vsWT_3 h | *egt* 2vsWT_6 h | *egt2* vsWT_12 h | root zone |
|---|---|---|---|---|---|---|---|---|
| HORVU.MOREX.r3.5HG0458010.1 | | | yes | | | yes | yes | elongation zone |
| HORVU.MOREX.r3.5HG0450470.1 | | | yes | | | | yes | elongation zone |
| HORVU.MOREX.r3.5HG0450100.1 | | | yes | | | | yes | elongation zone |
| HORVU.MOREX.r3.5HG0447720.1 | | | yes | | | | yes | elongation zone |
| HORVU.MOREX.r3.5HG0444350.1 | | yes | yes | | | | yes | elongation zone |
| HORVU.MOREX.r3.5HG0442460.1 | | | yes | | | | yes | elongation zone |
| HORVU.MOREX.r3.5HG0441420.1 | | yes | | | | yes | | elongation zone |
| HORVU.MOREX.r3.5HG0439710.1 | | | yes | | | | yes | elongation zone |
| HORVU.MOREX.r3.5HG0438160.1 | | | yes | | | | yes | elongation zone |
| HORVU.MOREX.r3.5HG0437610.1 | | | yes | | | | yes | elongation zone |
| HORVU.MOREX.r3.5HG0435800.1 | | | yes | | | | yes | elongation zone |
| HORVU.MOREX.r3.5HG0430460.1 | | | yes | | | | yes | elongation zone |
| HORVU.MOREX.r3.5HG0430050.1 | | yes | yes | | | yes | yes | elongation zone |
| HORVU.MOREX.r3.5HG0428840.1 | | | yes | | | yes | yes | elongation zone |
| HORVU.MOREX.r3.5HG0427370.1 | | | yes | | | | yes | elongation zone |
| HORVU.MOREX.r3.5HG0427060.1 | | | yes | | | | yes | elongation zone |
| HORVU.MOREX.r3.5HG0426480.1 | | | yes | | | | yes | elongation zone |
| HORVU.MOREX.r3.5HG0426110.1 | | yes | yes | | | yes | | elongation zone |
| HORVU.MOREX.r3.5HG0420980.1 | | | yes | | | | yes | elongation zone |
| HORVU.MOREX.r3.5HG0420970.1 | | | yes | | | | yes | elongation zone |
| HORVU.MOREX.r3.5HG0420630.1 | | yes | yes | | | yes | | elongation zone |
| HORVU.MOREX.r3.5HG0420480.1 | | yes | yes | | | yes | | elongation zone |
| HORVU.MOREX.r3.5HG0420410.1 | | | yes | | | | yes | elongation zone |
| HORVU.MOREX.r3.5HG0420210.1 | | | yes | | | | yes | elongation zone |
| HORVU.MOREX.r3.5HG0419590.1 | | | yes | | | yes | yes | elongation zone |
| HORVU.MOREX.r3.4HG0418640.1 | | | yes | | | | yes | elongation zone |
| HORVU.MOREX.r3.4HG0418530.1 | | | yes | | | | yes | elongation zone |
| HORVU.MOREX.r3.4HG0418460.1 | | | yes | | | yes | yes | elongation zone |
| HORVU.MOREX.r3.4HG0417920.1 | yes | yes | yes | | | yes | | elongation zone |
| HORVU.MOREX.r3.4HG0417410.1 | | | yes | | | | yes | elongation zone |
| HORVU.MOREX.r3.4HG0417010.1 | | yes | yes | | | yes | | elongation zone |
| HORVU.MOREX.r3.4HG0416730.1 | | | yes | | | | yes | elongation zone |
| HORVU.MOREX.r3.4HG0416070.1 | | | yes | | | | yes | elongation zone |
| HORVU.MOREX.r3.4HG0415600.1 | | | yes | | | | yes | elongation zone |
| HORVU.MOREX.r3.4HG0415590.1 | | yes | yes | | | | yes | elongation zone |

**Table S7** Continued.

| ID | WT_3 h | WT_6 h | WT_12 h | *egt2* vsWT_0 h | *egt2* vsWT_3 h | *egt* 2vsWT_6 h | *egt2* vsWT_12 h | root zone |
|---|---|---|---|---|---|---|---|---|
| HORVU.MOREX.r3.4HG0415490.1 | | yes | yes | | | yes | | elongation zone |
| HORVU.MOREX.r3.4HG0415420.1 | | yes | yes | | | yes | yes | elongation zone |
| HORVU.MOREX.r3.4HG0413410.1 | | | yes | | | | yes | elongation zone |
| HORVU.MOREX.r3.4HG0413320.1 | | yes | yes | | | yes | | elongation zone |
| HORVU.MOREX.r3.4HG0412370.1 | | yes | yes | | | yes | | elongation zone |
| HORVU.MOREX.r3.4HG0411270.1 | | | yes | | | | yes | elongation zone |
| HORVU.MOREX.r3.4HG0410640.1 | | yes | yes | | | yes | | elongation zone |
| HORVU.MOREX.r3.4HG0409600.1 | | | yes | | | | yes | elongation zone |
| HORVU.MOREX.r3.4HG0408780.1 | | yes | yes | | yes | | | elongation zone |
| HORVU.MOREX.r3.4HG0408270.1 | | yes | yes | | | yes | | elongation zone |
| HORVU.MOREX.r3.4HG0407230.1 | | yes | | | | yes | | elongation zone |
| HORVU.MOREX.r3.4HG0406600.1 | | yes | yes | | | | yes | elongation zone |
| HORVU.MOREX.r3.4HG0406380.1 | | | yes | | | | yes | elongation zone |
| HORVU.MOREX.r3.4HG0406140.1 | | yes | yes | | | yes | | elongation zone |
| HORVU.MOREX.r3.4HG0405920.1 | | yes | yes | | | yes | | elongation zone |
| HORVU.MOREX.r3.4HG0405780.1 | | | yes | | | | yes | elongation zone |
| HORVU.MOREX.r3.4HG0405400.1 | | yes | yes | | | yes | | elongation zone |
| HORVU.MOREX.r3.4HG0404740.1 | | | yes | | | yes | yes | elongation zone |
| HORVU.MOREX.r3.4HG0403620.1 | | yes | yes | | | yes | | elongation zone |
| HORVU.MOREX.r3.4HG0403250.1 | | | yes | | | | yes | elongation zone |
| HORVU.MOREX.r3.4HG0403070.1 | | | yes | | | | yes | elongation zone |
| HORVU.MOREX.r3.4HG0402730.1 | | yes | yes | | | yes | | elongation zone |
| HORVU.MOREX.r3.4HG0402650.1 | | | yes | | | | yes | elongation zone |
| HORVU.MOREX.r3.4HG0402600.2 | | | yes | | | | yes | elongation zone |
| HORVU.MOREX.r3.4HG0402030.1 | | yes | yes | | | | yes | elongation zone |
| HORVU.MOREX.r3.4HG0402000.1 | | yes | yes | | | yes | yes | elongation zone |
| HORVU.MOREX.r3.4HG0401720.1 | | | yes | | | | yes | elongation zone |
| HORVU.MOREX.r3.4HG0401710.1 | | | yes | | | | yes | elongation zone |
| HORVU.MOREX.r3.4HG0400740.1 | | yes | yes | | | yes | | elongation zone |
| HORVU.MOREX.r3.4HG0400520.1 | | yes | yes | | | | yes | elongation zone |
| HORVU.MOREX.r3.4HG0400040.1 | | | yes | | | | yes | elongation zone |
| HORVU.MOREX.r3.4HG0398910.1 | | | yes | | | | yes | elongation zone |
| HORVU.MOREX.r3.4HG0398900.1 | | | yes | | | | yes | elongation zone |
| HORVU.MOREX.r3.4HG0396660.1 | | | yes | | | | yes | elongation zone |
| HORVU.MOREX.r3.4HG0396220.1 | | | yes | | | | yes | elongation zone |

**Table S7** Continued.

| ID | WT_3 h | WT_6 h | WT_12 h | *egt2* vsWT_0 h | *egt2* vsWT_3 h | *egt* 2vsWT_6 h | *egt2* vsWT_12 h | root zone |
|---|---|---|---|---|---|---|---|---|
| HORVU.MOREX.r3.4HG0395540.1 | yes | yes | yes | | | yes | | elongation zone |
| HORVU.MOREX.r3.4HG0394830.1 | | yes | yes | | | | yes | elongation zone |
| HORVU.MOREX.r3.4HG0394460.1 | | yes | | | | yes | | elongation zone |
| HORVU.MOREX.r3.4HG0393520.1 | | yes | yes | | | yes | yes | elongation zone |
| HORVU.MOREX.r3.4HG0393260.1 | | | yes | | | | yes | elongation zone |
| HORVU.MOREX.r3.4HG0392160.1 | | | yes | | | | yes | elongation zone |
| HORVU.MOREX.r3.4HG0391330.1 | | yes | yes | | | yes | yes | elongation zone |
| HORVU.MOREX.r3.4HG0389250.1 | | yes | | | | yes | | elongation zone |
| HORVU.MOREX.r3.4HG0388470.1 | | | yes | | | | yes | elongation zone |
| HORVU.MOREX.r3.4HG0388310.1 | | yes | yes | | | yes | | elongation zone |
| HORVU.MOREX.r3.4HG0387150.1 | | | yes | | | | yes | elongation zone |
| HORVU.MOREX.r3.4HG0387120.1 | | | yes | | | | yes | elongation zone |
| HORVU.MOREX.r3.4HG0386830.1 | | | yes | | | | yes | elongation zone |
| HORVU.MOREX.r3.4HG0386800.1 | | | yes | | | | yes | elongation zone |
| HORVU.MOREX.r3.4HG0385300.1 | | | yes | | | | yes | elongation zone |
| HORVU.MOREX.r3.4HG0385270.1 | | | yes | | | | yes | elongation zone |
| HORVU.MOREX.r3.4HG0384390.1 | | yes | yes | | | yes | yes | elongation zone |
| HORVU.MOREX.r3.4HG0384230.1 | | yes | yes | | | | yes | elongation zone |
| HORVU.MOREX.r3.4HG0383870.1 | | | yes | | | | yes | elongation zone |
| HORVU.MOREX.r3.4HG0383780.1 | | yes | yes | | | yes | yes | elongation zone |
| HORVU.MOREX.r3.4HG0383530.3 | | | yes | | | | yes | elongation zone |
| HORVU.MOREX.r3.4HG0383340.1 | | | yes | | | | yes | elongation zone |
| HORVU.MOREX.r3.4HG0383070.1 | | | yes | | | yes | | elongation zone |
| HORVU.MOREX.r3.4HG0382500.1 | | | yes | | | yes | yes | elongation zone |
| HORVU.MOREX.r3.4HG0380540.1 | | yes | yes | | | yes | yes | elongation zone |
| HORVU.MOREX.r3.4HG0379960.1 | | | yes | | | | yes | elongation zone |
| HORVU.MOREX.r3.4HG0379400.1 | | yes | yes | | | | yes | elongation zone |
| HORVU.MOREX.r3.4HG0379290.1 | | yes | yes | | | yes | | elongation zone |
| HORVU.MOREX.r3.4HG0378960.1 | | | yes | | | | yes | elongation zone |
| HORVU.MOREX.r3.4HG0378580.1 | | yes | yes | | | yes | yes | elongation zone |
| HORVU.MOREX.r3.4HG0375550.1 | | | yes | | | | yes | elongation zone |
| HORVU.MOREX.r3.4HG0372280.1 | | | yes | | | | yes | elongation zone |
| HORVU.MOREX.r3.4HG0370520.1 | | | yes | | | | yes | elongation zone |
| HORVU.MOREX.r3.4HG0364210.1 | | yes | | | | yes | | elongation zone |
| HORVU.MOREX.r3.4HG0364050.1 | | yes | yes | | | yes | yes | elongation zone |

**Table S7** Continued.

| ID | WT_3 h | WT_6 h | WT_12 h | *egt2* vsWT_0 h | *egt2* vsWT_3 h | *egt* 2vsWT_6 h | *egt2* vsWT_12 h | root zone |
|---|---|---|---|---|---|---|---|---|
| HORVU.MOREX.r3.4HG0361420.2 | | | yes | | | | yes | elongation zone |
| HORVU.MOREX.r3.4HG0358890.1 | | | yes | | | | yes | elongation zone |
| HORVU.MOREX.r3.4HG0357230.1 | | | yes | | | | yes | elongation zone |
| HORVU.MOREX.r3.4HG0355210.1 | | yes | | | | yes | | elongation zone |
| HORVU.MOREX.r3.4HG0354980.1 | | yes | yes | | | yes | yes | elongation zone |
| HORVU.MOREX.r3.4HG0354970.1 | | | yes | | | yes | yes | elongation zone |
| HORVU.MOREX.r3.4HG0354540.1 | | | yes | | | | yes | elongation zone |
| HORVU.MOREX.r3.4HG0354010.1 | | yes | yes | | | yes | | elongation zone |
| HORVU.MOREX.r3.4HG0353450.1 | | | yes | | | | yes | elongation zone |
| HORVU.MOREX.r3.4HG0353330.1 | | | yes | | | | yes | elongation zone |
| HORVU.MOREX.r3.4HG0353200.1 | | | yes | | | | yes | elongation zone |
| HORVU.MOREX.r3.4HG0352780.1 | | | yes | | | | yes | elongation zone |
| HORVU.MOREX.r3.4HG0352200.1 | | yes | yes | | | yes | | elongation zone |
| HORVU.MOREX.r3.4HG0351750.1 | | yes | yes | | | yes | | elongation zone |
| HORVU.MOREX.r3.4HG0351070.1 | | | yes | | | | yes | elongation zone |
| HORVU.MOREX.r3.4HG0350900.1 | | yes | yes | | | yes | | elongation zone |
| HORVU.MOREX.r3.4HG0350800.1 | | | yes | | | | yes | elongation zone |
| HORVU.MOREX.r3.4HG0349310.1 | | yes | | | | yes | | elongation zone |
| HORVU.MOREX.r3.4HG0349170.1 | | | yes | | | | yes | elongation zone |
| HORVU.MOREX.r3.4HG0348800.1 | | yes | yes | | | yes | | elongation zone |
| HORVU.MOREX.r3.4HG0347760.1 | | | yes | | | | yes | elongation zone |
| HORVU.MOREX.r3.4HG0346830.1 | | yes | yes | | | yes | | elongation zone |
| HORVU.MOREX.r3.4HG0346720.1 | | yes | yes | | | yes | yes | elongation zone |
| HORVU.MOREX.r3.4HG0345810.1 | | | yes | | | | yes | elongation zone |
| HORVU.MOREX.r3.4HG0344990.1 | | | yes | | | | yes | elongation zone |
| HORVU.MOREX.r3.4HG0344830.1 | | yes | | | yes | yes | yes | elongation zone |
| HORVU.MOREX.r3.4HG0344370.1 | | yes | yes | | | yes | | elongation zone |
| HORVU.MOREX.r3.4HG0343050.1 | | | yes | | | yes | | elongation zone |
| HORVU.MOREX.r3.4HG0342950.1 | | | yes | | | | yes | elongation zone |
| HORVU.MOREX.r3.4HG0342640.1 | | | yes | | | | yes | elongation zone |
| HORVU.MOREX.r3.4HG0342160.1 | | yes | yes | | | yes | yes | elongation zone |
| HORVU.MOREX.r3.4HG0342080.1 | | yes | yes | | | yes | | elongation zone |
| HORVU.MOREX.r3.4HG0341080.1 | | | yes | | | | yes | elongation zone |
| HORVU.MOREX.r3.4HG0338400.1 | | | yes | | | | yes | elongation zone |
| HORVU.MOREX.r3.4HG0337950.1 | | yes | yes | | | yes | | elongation zone |

**Table S7** Continued.

| ID | WT_3 h | WT_6 h | WT_12 h | *egt2* vsWT_0 h | *egt2* vsWT_3 h | *egt* 2vsWT_6 h | *egt2* vsWT_12 h | root zone |
|---|---|---|---|---|---|---|---|---|
| HORVU.MOREX.r3.4HG0337770.1 | | | yes | | | | yes | elongation zone |
| HORVU.MOREX.r3.4HG0337120.1 | | | yes | | | | yes | elongation zone |
| HORVU.MOREX.r3.4HG0337110.1 | | yes | yes | | | yes | | elongation zone |
| HORVU.MOREX.r3.4HG0336310.1 | | | yes | | | | yes | elongation zone |
| HORVU.MOREX.r3.4HG0335450.1 | | yes | | | | yes | | elongation zone |
| HORVU.MOREX.r3.4HG0335110.1 | | | yes | | | | yes | elongation zone |
| HORVU.MOREX.r3.4HG0334360.1 | | yes | yes | | | yes | yes | elongation zone |
| HORVU.MOREX.r3.4HG0333550.2 | | | yes | | | | yes | elongation zone |
| HORVU.MOREX.r3.4HG0333450.1 | | yes | yes | | | yes | yes | elongation zone |
| HORVU.MOREX.r3.4HG0332930.1 | | yes | | | | yes | | elongation zone |
| HORVU.MOREX.r3.4HG0331420.1 | | yes | | | | | yes | elongation zone |
| HORVU.MOREX.r3.3HG0330980.1 | | yes | yes | | | yes | | elongation zone |
| HORVU.MOREX.r3.3HG0330200.1 | | yes | yes | | | yes | | elongation zone |
| HORVU.MOREX.r3.3HG0330120.1 | | | yes | | | | yes | elongation zone |
| HORVU.MOREX.r3.3HG0329950.1 | | yes | yes | | | yes | | elongation zone |
| HORVU.MOREX.r3.3HG0329870.1 | | yes | yes | | | yes | yes | elongation zone |
| HORVU.MOREX.r3.3HG0329040.1 | | yes | yes | | | yes | yes | elongation zone |
| HORVU.MOREX.r3.3HG0328480.1 | | | yes | | | | yes | elongation zone |
| HORVU.MOREX.r3.3HG0327630.1 | | | yes | | | | yes | elongation zone |
| HORVU.MOREX.r3.3HG0327170.1 | | yes | yes | | | | yes | elongation zone |
| HORVU.MOREX.r3.3HG0326430.1 | | yes | yes | | | yes | | elongation zone |
| HORVU.MOREX.r3.3HG0323810.1 | | | yes | | | | yes | elongation zone |
| HORVU.MOREX.r3.3HG0323600.1 | | | yes | | | yes | yes | elongation zone |
| HORVU.MOREX.r3.3HG0322660.1 | | | yes | | | | yes | elongation zone |
| HORVU.MOREX.r3.3HG0321700.1 | | yes | yes | | | yes | yes | elongation zone |
| HORVU.MOREX.r3.3HG0320840.1 | | | yes | | | | yes | elongation zone |
| HORVU.MOREX.r3.3HG0319570.1 | | | yes | | | | yes | elongation zone |
| HORVU.MOREX.r3.3HG0318700.1 | | | yes | | | yes | yes | elongation zone |
| HORVU.MOREX.r3.3HG0318400.1 | | | yes | | | | yes | elongation zone |
| HORVU.MOREX.r3.3HG0316280.1 | | | yes | | | yes | yes | elongation zone |
| HORVU.MOREX.r3.3HG0316000.1 | | | yes | | | | yes | elongation zone |
| HORVU.MOREX.r3.3HG0315410.1 | | | yes | | | | yes | elongation zone |
| HORVU.MOREX.r3.3HG0314820.1 | | | yes | | | | yes | elongation zone |
| HORVU.MOREX.r3.3HG0314070.1 | | yes | yes | | | | yes | elongation zone |
| HORVU.MOREX.r3.3HG0313320.1 | | yes | | | | yes | | elongation zone |

**Table S7** Continued.

| ID | WT_3 h | WT_6 h | WT_12 h | *egt2* vsWT_0 h | *egt2* vsWT_3 h | *egt* 2vsWT_6 h | *egt2* vsWT_12 h | root zone |
|---|---|---|---|---|---|---|---|---|
| HORVU.MOREX.r3.3HG0313090.1 | | yes | yes | | | yes | yes | elongation zone |
| HORVU.MOREX.r3.3HG0310540.1 | | yes | yes | | | | yes | elongation zone |
| HORVU.MOREX.r3.3HG0310210.1 | | | yes | | | | yes | elongation zone |
| HORVU.MOREX.r3.3HG0309820.1 | | | yes | | | | yes | elongation zone |
| HORVU.MOREX.r3.3HG0309410.1 | | yes | yes | | | | yes | elongation zone |
| HORVU.MOREX.r3.3HG0308980.1 | | | yes | | | | yes | elongation zone |
| HORVU.MOREX.r3.3HG0308420.1 | | | yes | | | | yes | elongation zone |
| HORVU.MOREX.r3.3HG0307850.1 | | yes | yes | | | yes | | elongation zone |
| HORVU.MOREX.r3.3HG0307400.1 | | yes | yes | | | | yes | elongation zone |
| HORVU.MOREX.r3.3HG0307310.1 | | yes | | | | yes | yes | elongation zone |
| HORVU.MOREX.r3.3HG0307240.1 | | yes | yes | | | yes | yes | elongation zone |
| HORVU.MOREX.r3.3HG0307120.1 | | | yes | | | yes | yes | elongation zone |
| HORVU.MOREX.r3.3HG0306420.1 | | | yes | | | | yes | elongation zone |
| HORVU.MOREX.r3.3HG0306210.1 | | | yes | | | | yes | elongation zone |
| HORVU.MOREX.r3.3HG0305500.1 | | | yes | | | | yes | elongation zone |
| HORVU.MOREX.r3.3HG0305440.1 | | yes | | | | yes | | elongation zone |
| HORVU.MOREX.r3.3HG0304660.1 | | | yes | | | | yes | elongation zone |
| HORVU.MOREX.r3.3HG0304490.1 | | yes | yes | | | | yes | elongation zone |
| HORVU.MOREX.r3.3HG0304420.1 | | | yes | | | | yes | elongation zone |
| HORVU.MOREX.r3.3HG0304370.1 | | | yes | | | | yes | elongation zone |
| HORVU.MOREX.r3.3HG0303970.1 | | | yes | | | | yes | elongation zone |
| HORVU.MOREX.r3.3HG0303380.1 | | | yes | | | yes | yes | elongation zone |
| HORVU.MOREX.r3.3HG0303330.1 | | | yes | | | | yes | elongation zone |
| HORVU.MOREX.r3.3HG0302860.1 | | | yes | | | | yes | elongation zone |
| HORVU.MOREX.r3.3HG0301930.1 | | | yes | | | | yes | elongation zone |
| HORVU.MOREX.r3.3HG0299990.1 | | | yes | | | | yes | elongation zone |
| HORVU.MOREX.r3.3HG0299820.1 | | | yes | | | | yes | elongation zone |
| HORVU.MOREX.r3.3HG0298790.1 | | yes | | | | | yes | elongation zone |
| HORVU.MOREX.r3.3HG0298750.1 | | | yes | | | yes | yes | elongation zone |
| HORVU.MOREX.r3.3HG0298340.1 | | | yes | | | | yes | elongation zone |
| HORVU.MOREX.r3.3HG0297940.1 | | | yes | yes | yes | yes | yes | elongation zone |
| HORVU.MOREX.r3.3HG0297330.1 | | yes | yes | | | yes | | elongation zone |
| HORVU.MOREX.r3.3HG0297180.1 | | | yes | | | | yes | elongation zone |
| HORVU.MOREX.r3.3HG0296490.1 | | | yes | | | | yes | elongation zone |
| HORVU.MOREX.r3.3HG0295680.1 | | | yes | | | | yes | elongation zone |

**Table S7** Continued.

| ID | WT_3 h | WT_6 h | WT_12 h | *egt2* vsWT_0 h | *egt2* vsWT_3 h | *egt* 2vsWT_6 h | *egt2* vsWT_12 h | root zone |
|---|---|---|---|---|---|---|---|---|
| HORVU.MOREX.r3.3HG0293700.1 | | | yes | | | | yes | elongation zone |
| HORVU.MOREX.r3.3HG0293570.1 | | yes | yes | | | yes | | elongation zone |
| HORVU.MOREX.r3.3HG0293310.1 | | yes | yes | | | | yes | elongation zone |
| HORVU.MOREX.r3.3HG0293130.1 | | yes | yes | | | yes | | elongation zone |
| HORVU.MOREX.r3.3HG0293040.1 | | | yes | | | | yes | elongation zone |
| HORVU.MOREX.r3.3HG0292680.1 | | yes | yes | | | yes | yes | elongation zone |
| HORVU.MOREX.r3.3HG0291490.1 | | | yes | | | | yes | elongation zone |
| HORVU.MOREX.r3.3HG0291030.1 | | yes | yes | | | | yes | elongation zone |
| HORVU.MOREX.r3.3HG0290180.1 | | | yes | | | | yes | elongation zone |
| HORVU.MOREX.r3.3HG0289070.1 | | | yes | | | | yes | elongation zone |
| HORVU.MOREX.r3.3HG0288420.1 | | | yes | | | | yes | elongation zone |
| HORVU.MOREX.r3.3HG0287690.1 | | yes | yes | | | yes | | elongation zone |
| HORVU.MOREX.r3.3HG0287070.1 | | yes | yes | | | | yes | elongation zone |
| HORVU.MOREX.r3.3HG0287000.1 | | yes | yes | | | yes | | elongation zone |
| HORVU.MOREX.r3.3HG0286160.1 | | | yes | | | | yes | elongation zone |
| HORVU.MOREX.r3.3HG0285840.1 | | | yes | | | | yes | elongation zone |
| HORVU.MOREX.r3.3HG0285440.1 | | yes | yes | | | yes | | elongation zone |
| HORVU.MOREX.r3.3HG0284780.1 | | yes | yes | | | yes | | elongation zone |
| HORVU.MOREX.r3.3HG0283990.1 | | | yes | | | | yes | elongation zone |
| HORVU.MOREX.r3.3HG0283250.1 | | | yes | | | | yes | elongation zone |
| HORVU.MOREX.r3.3HG0281860.1 | | | yes | | | | yes | elongation zone |
| HORVU.MOREX.r3.3HG0280970.1 | | | yes | | | yes | | elongation zone |
| HORVU.MOREX.r3.3HG0280960.1 | | yes | yes | | | yes | yes | elongation zone |
| HORVU.MOREX.r3.3HG0280630.1 | | yes | yes | | | yes | | elongation zone |
| HORVU.MOREX.r3.3HG0280160.1 | | | yes | | | | yes | elongation zone |
| HORVU.MOREX.r3.3HG0280060.1 | | yes | yes | | | yes | | elongation zone |
| HORVU.MOREX.r3.3HG0278170.1 | | yes | yes | | | yes | | elongation zone |
| HORVU.MOREX.r3.3HG0277330.1 | | | yes | | | | yes | elongation zone |
| HORVU.MOREX.r3.3HG0277200.1 | | | yes | | | | yes | elongation zone |
| HORVU.MOREX.r3.3HG0276120.1 | | yes | yes | | | yes | | elongation zone |
| HORVU.MOREX.r3.3HG0275530.1 | | | yes | | | | yes | elongation zone |
| HORVU.MOREX.r3.3HG0275090.1 | | yes | yes | | | yes | | elongation zone |
| HORVU.MOREX.r3.3HG0274100.1 | | | yes | | | | yes | elongation zone |
| HORVU.MOREX.r3.3HG0273790.2 | | | yes | | | yes | | elongation zone |
| HORVU.MOREX.r3.3HG0272430.1 | | | yes | | | yes | yes | elongation zone |

**Table S7** Continued.

| ID | WT_3 h | WT_6 h | WT_12 h | *egt2* vsWT_0 h | *egt2* vsWT_3 h | *egt* 2vsWT_6 h | *egt2* vsWT_12 h | root zone |
|---|---|---|---|---|---|---|---|---|
| HORVU.MOREX.r3.3HG0269960.1 | | | yes | | | | yes | elongation zone |
| HORVU.MOREX.r3.3HG0264440.1 | | yes | | | | yes | | elongation zone |
| HORVU.MOREX.r3.3HG0259660.1 | | | yes | | | | yes | elongation zone |
| HORVU.MOREX.r3.3HG0257680.1 | | yes | | | | yes | | elongation zone |
| HORVU.MOREX.r3.3HG0257310.1 | | yes | yes | | | yes | | elongation zone |
| HORVU.MOREX.r3.3HG0256580.1 | | | yes | | | | yes | elongation zone |
| HORVU.MOREX.r3.3HG0256300.1 | | | yes | | | | yes | elongation zone |
| HORVU.MOREX.r3.3HG0255580.1 | | | yes | | | | yes | elongation zone |
| HORVU.MOREX.r3.3HG0255020.1 | | yes | yes | | | yes | yes | elongation zone |
| HORVU.MOREX.r3.3HG0254950.1 | | yes | yes | | | yes | yes | elongation zone |
| HORVU.MOREX.r3.3HG0254940.1 | | yes | yes | | | yes | yes | elongation zone |
| HORVU.MOREX.r3.3HG0254850.1 | | | yes | | | | yes | elongation zone |
| HORVU.MOREX.r3.3HG0254430.1 | | | yes | | | | yes | elongation zone |
| HORVU.MOREX.r3.3HG0253860.1 | | yes | | | | yes | | elongation zone |
| HORVU.MOREX.r3.3HG0252610.1 | | yes | yes | | | yes | yes | elongation zone |
| HORVU.MOREX.r3.3HG0252240.1 | | | yes | | | | yes | elongation zone |
| HORVU.MOREX.r3.3HG0251830.1 | | | yes | | | | yes | elongation zone |
| HORVU.MOREX.r3.3HG0250060.1 | | yes | yes | | | yes | | elongation zone |
| HORVU.MOREX.r3.3HG0249590.1 | | | yes | | | | yes | elongation zone |
| HORVU.MOREX.r3.3HG0248280.1 | | | yes | | | yes | yes | elongation zone |
| HORVU.MOREX.r3.3HG0247250.1 | | | yes | | | | yes | elongation zone |
| HORVU.MOREX.r3.3HG0246670.1 | | yes | yes | | | yes | | elongation zone |
| HORVU.MOREX.r3.3HG0246560.1 | | | yes | | | | yes | elongation zone |
| HORVU.MOREX.r3.3HG0246500.1 | | yes | yes | | | yes | yes | elongation zone |
| HORVU.MOREX.r3.3HG0246300.1 | | yes | | | | yes | | elongation zone |
| HORVU.MOREX.r3.3HG0246030.1 | | | yes | | | | yes | elongation zone |
| HORVU.MOREX.r3.3HG0245250.1 | | yes | yes | | | yes | | elongation zone |
| HORVU.MOREX.r3.3HG0245120.1 | | yes | yes | | | yes | | elongation zone |
| HORVU.MOREX.r3.3HG0244240.1 | | | yes | | | | yes | elongation zone |
| HORVU.MOREX.r3.3HG0244040.1 | | yes | | | | yes | | elongation zone |
| HORVU.MOREX.r3.3HG0244010.1 | | | yes | | | | yes | elongation zone |
| HORVU.MOREX.r3.3HG0243920.1 | | | yes | | | | yes | elongation zone |
| HORVU.MOREX.r3.3HG0243130.1 | | yes | yes | | | | yes | elongation zone |
| HORVU.MOREX.r3.3HG0242030.1 | | | yes | | | | yes | elongation zone |
| HORVU.MOREX.r3.3HG0241320.1 | | yes | | | | yes | | elongation zone |

**Table S7** Continued.

| ID | WT_3 h | WT_6 h | WT_12 h | *egt2* vsWT_0 h | *egt2* vsWT_3 h | *egt* 2vsWT_6 h | *egt2* vsWT_12 h | root zone |
|---|---|---|---|---|---|---|---|---|
| HORVU.MOREX.r3.3HG0240450.1 | | | yes | | | | yes | elongation zone |
| HORVU.MOREX.r3.3HG0239980.1 | | yes | | | | yes | yes | elongation zone |
| HORVU.MOREX.r3.3HG0238180.1 | | yes | yes | | | yes | yes | elongation zone |
| HORVU.MOREX.r3.3HG0237990.1 | | yes | yes | | | | yes | elongation zone |
| HORVU.MOREX.r3.3HG0237870.1 | | | yes | | | | yes | elongation zone |
| HORVU.MOREX.r3.3HG0236000.1 | | | yes | | | | yes | elongation zone |
| HORVU.MOREX.r3.3HG0235030.1 | | yes | yes | | | yes | | elongation zone |
| HORVU.MOREX.r3.3HG0234970.1 | | yes | yes | | | yes | yes | elongation zone |
| HORVU.MOREX.r3.3HG0234470.1 | | yes | | | | yes | | elongation zone |
| HORVU.MOREX.r3.3HG0234000.1 | | yes | yes | | | yes | | elongation zone |
| HORVU.MOREX.r3.3HG0233990.1 | | yes | yes | | | yes | | elongation zone |
| HORVU.MOREX.r3.3HG0233150.1 | | | yes | | | | yes | elongation zone |
| HORVU.MOREX.r3.3HG0231880.1 | | | yes | | | | yes | elongation zone |
| HORVU.MOREX.r3.3HG0231780.1 | | | yes | | | | yes | elongation zone |
| HORVU.MOREX.r3.3HG0231630.1 | | | yes | | | | yes | elongation zone |
| HORVU.MOREX.r3.3HG0230970.1 | | | yes | | | | yes | elongation zone |
| HORVU.MOREX.r3.3HG0230190.1 | | | yes | | | | yes | elongation zone |
| HORVU.MOREX.r3.3HG0230090.1 | | | yes | | | | yes | elongation zone |
| HORVU.MOREX.r3.3HG0229480.1 | | | yes | | | | yes | elongation zone |
| HORVU.MOREX.r3.3HG0224680.1 | | | yes | | | yes | | elongation zone |
| HORVU.MOREX.r3.3HG0223950.1 | | | yes | | | | yes | elongation zone |
| HORVU.MOREX.r3.3HG0223160.1 | | | yes | | | | yes | elongation zone |
| HORVU.MOREX.r3.3HG0222830.1 | | yes | yes | | | yes | | elongation zone |
| HORVU.MOREX.r3.3HG0222130.1 | | yes | yes | | | yes | yes | elongation zone |
| HORVU.MOREX.r3.3HG0221460.1 | | | yes | | | | yes | elongation zone |
| HORVU.MOREX.r3.3HG0220360.1 | | | yes | | | yes | yes | elongation zone |
| HORVU.MOREX.r3.3HG0219810.1 | | | yes | | | | yes | elongation zone |
| HORVU.MOREX.r3.3HG0219650.1 | | | yes | | | | yes | elongation zone |
| HORVU.MOREX.r3.3HG0219410.1 | | yes | yes | | | | yes | elongation zone |
| HORVU.MOREX.r3.3HG0219380.1 | | yes | yes | | | yes | | elongation zone |
| HORVU.MOREX.r3.3HG0218560.1 | | yes | yes | | | | yes | elongation zone |
| HORVU.MOREX.r3.3HG0218330.1 | | | yes | | | | yes | elongation zone |
| HORVU.MOREX.r3.2HG0218010.1 | | yes | yes | | | yes | yes | elongation zone |
| HORVU.MOREX.r3.2HG0217550.1 | | | yes | | | | yes | elongation zone |
| HORVU.MOREX.r3.2HG0217090.1 | | yes | yes | | | yes | | elongation zone |

**Table S7** Continued.

| ID | WT_3 h | WT_6 h | WT_12 h | *egt2* vsWT_0 h | *egt2* vsWT_3 h | *egt* 2vsWT_6 h | *egt2* vsWT_12 h | root zone |
|---|---|---|---|---|---|---|---|---|
| HORVU.MOREX.r3.2HG0215550.1 | | | yes | | | | yes | elongation zone |
| HORVU.MOREX.r3.2HG0215310.1 | | | yes | | | | yes | elongation zone |
| HORVU.MOREX.r3.2HG0215250.1 | | | yes | | | | yes | elongation zone |
| HORVU.MOREX.r3.2HG0215220.1 | | | yes | | | | yes | elongation zone |
| HORVU.MOREX.r3.2HG0214240.1 | | yes | yes | | | yes | | elongation zone |
| HORVU.MOREX.r3.2HG0214090.1 | | | yes | | | | yes | elongation zone |
| HORVU.MOREX.r3.2HG0214070.1 | | yes | yes | | | yes | | elongation zone |
| HORVU.MOREX.r3.2HG0213250.1 | | yes | yes | | | yes | yes | elongation zone |
| HORVU.MOREX.r3.2HG0213020.1 | | yes | | | | yes | | elongation zone |
| HORVU.MOREX.r3.2HG0212990.1 | | | yes | | | | yes | elongation zone |
| HORVU.MOREX.r3.2HG0212900.1 | | | yes | | | | yes | elongation zone |
| HORVU.MOREX.r3.2HG0212570.1 | | yes | yes | | | yes | yes | elongation zone |
| HORVU.MOREX.r3.2HG0210510.1 | | yes | yes | | | yes | | elongation zone |
| HORVU.MOREX.r3.2HG0210420.1 | | | yes | | | | yes | elongation zone |
| HORVU.MOREX.r3.2HG0209030.1 | | | yes | | | | yes | elongation zone |
| HORVU.MOREX.r3.2HG0207860.1 | | yes | yes | | | yes | yes | elongation zone |
| HORVU.MOREX.r3.2HG0207790.1 | | yes | yes | | | | yes | elongation zone |
| HORVU.MOREX.r3.2HG0207720.1 | | yes | yes | | | | yes | elongation zone |
| HORVU.MOREX.r3.2HG0206470.1 | | | yes | | | yes | yes | elongation zone |
| HORVU.MOREX.r3.2HG0205990.1 | | yes | yes | | | yes | | elongation zone |
| HORVU.MOREX.r3.2HG0205760.1 | | | yes | | | | yes | elongation zone |
| HORVU.MOREX.r3.2HG0205680.1 | | | yes | | | | yes | elongation zone |
| HORVU.MOREX.r3.2HG0205550.1 | | | yes | | | | yes | elongation zone |
| HORVU.MOREX.r3.2HG0205050.1 | | | yes | | | | yes | elongation zone |
| HORVU.MOREX.r3.2HG0204960.1 | | | yes | | | | yes | elongation zone |
| HORVU.MOREX.r3.2HG0204940.1 | | yes | yes | | | yes | yes | elongation zone |
| HORVU.MOREX.r3.2HG0204920.1 | | | yes | | | | yes | elongation zone |
| HORVU.MOREX.r3.2HG0204340.1 | | yes | yes | | | | yes | elongation zone |
| HORVU.MOREX.r3.2HG0203500.1 | | | yes | | | | yes | elongation zone |
| HORVU.MOREX.r3.2HG0203080.1 | | yes | | | | yes | | elongation zone |
| HORVU.MOREX.r3.2HG0200640.1 | | | yes | | | | yes | elongation zone |
| HORVU.MOREX.r3.2HG0200630.1 | | | yes | | | | yes | elongation zone |
| HORVU.MOREX.r3.2HG0199600.1 | | | yes | | | | yes | elongation zone |
| HORVU.MOREX.r3.2HG0199560.1 | | | yes | | | | yes | elongation zone |
| HORVU.MOREX.r3.2HG0197560.1 | | | yes | | | | yes | elongation zone |

**Table S7** Continued.

| ID | WT_3 h | WT_6 h | WT_12 h | *egt2* vsWT_0 h | *egt2* vsWT_3 h | *egt* 2vsWT_6 h | *egt2* vsWT_12 h | root zone |
|---|---|---|---|---|---|---|---|---|
| HORVU.MOREX.r3.2HG0197260.1 | | | yes | | | | yes | elongation zone |
| HORVU.MOREX.r3.2HG0196960.1 | | | yes | | | | yes | elongation zone |
| HORVU.MOREX.r3.2HG0196800.1 | | yes | yes | | | yes | yes | elongation zone |
| HORVU.MOREX.r3.2HG0196630.1 | | | yes | | | | yes | elongation zone |
| HORVU.MOREX.r3.2HG0195910.1 | | | yes | | | | yes | elongation zone |
| HORVU.MOREX.r3.2HG0195690.1 | | yes | yes | | | yes | | elongation zone |
| HORVU.MOREX.r3.2HG0195110.2 | | | yes | | | | yes | elongation zone |
| HORVU.MOREX.r3.2HG0194620.1 | | yes | yes | | | yes | | elongation zone |
| HORVU.MOREX.r3.2HG0194450.1 | | | yes | | | | yes | elongation zone |
| HORVU.MOREX.r3.2HG0193220.1 | | | yes | | | | yes | elongation zone |
| HORVU.MOREX.r3.2HG0192900.1 | | | yes | | | | yes | elongation zone |
| HORVU.MOREX.r3.2HG0192000.1 | | yes | yes | | | yes | yes | elongation zone |
| HORVU.MOREX.r3.2HG0190710.1 | | | yes | | | yes | | elongation zone |
| HORVU.MOREX.r3.2HG0189800.1 | | | yes | | | yes | | elongation zone |
| HORVU.MOREX.r3.2HG0189670.1 | | yes | yes | | | yes | | elongation zone |
| HORVU.MOREX.r3.2HG0188300.1 | | yes | yes | | | | yes | elongation zone |
| HORVU.MOREX.r3.2HG0187640.1 | | | yes | | | | yes | elongation zone |
| HORVU.MOREX.r3.2HG0187450.1 | | yes | yes | | | yes | | elongation zone |
| HORVU.MOREX.r3.2HG0186750.1 | | yes | | | | | yes | elongation zone |
| HORVU.MOREX.r3.2HG0186650.1 | | | yes | | | | yes | elongation zone |
| HORVU.MOREX.r3.2HG0185840.1 | | | yes | | | | yes | elongation zone |
| HORVU.MOREX.r3.2HG0185570.1 | | | yes | | | | yes | elongation zone |
| HORVU.MOREX.r3.2HG0183740.1 | | | yes | | | | yes | elongation zone |
| HORVU.MOREX.r3.2HG0182400.1 | | yes | yes | | | yes | | elongation zone |
| HORVU.MOREX.r3.2HG0181680.1 | | yes | yes | | | yes | yes | elongation zone |
| HORVU.MOREX.r3.2HG0181660.1 | | | yes | | | | yes | elongation zone |
| HORVU.MOREX.r3.2HG0181550.1 | | | yes | | | | yes | elongation zone |
| HORVU.MOREX.r3.2HG0181390.1 | | | yes | | | | yes | elongation zone |
| HORVU.MOREX.r3.2HG0180580.1 | | | yes | | | | yes | elongation zone |
| HORVU.MOREX.r3.2HG0180060.1 | | | yes | | | | yes | elongation zone |
| HORVU.MOREX.r3.2HG0180010.1 | | yes | yes | | | yes | | elongation zone |
| HORVU.MOREX.r3.2HG0179560.1 | | yes | yes | | | yes | | elongation zone |
| HORVU.MOREX.r3.2HG0179540.1 | | | yes | | | | yes | elongation zone |
| HORVU.MOREX.r3.2HG0178160.1 | | yes | yes | | | yes | | elongation zone |
| HORVU.MOREX.r3.2HG0178050.1 | | yes | yes | | | | yes | elongation zone |

**Table S7** Continued.

| ID | WT_3 h | WT_6 h | WT_12 h | *egt2* vsWT_0 h | *egt2* vsWT_3 h | *egt* 2vsWT_6 h | *egt2* vsWT_12 h | root zone |
|---|---|---|---|---|---|---|---|---|
| HORVU.MOREX.r3.2HG0176920.1 | | | yes | | | | yes | elongation zone |
| HORVU.MOREX.r3.2HG0176290.1 | | | yes | | | | yes | elongation zone |
| HORVU.MOREX.r3.2HG0173960.1 | | yes | yes | | | | yes | elongation zone |
| HORVU.MOREX.r3.2HG0173550.1 | | | yes | | | | yes | elongation zone |
| HORVU.MOREX.r3.2HG0173380.1 | | | yes | | | | yes | elongation zone |
| HORVU.MOREX.r3.2HG0173340.1 | | | yes | | | | yes | elongation zone |
| HORVU.MOREX.r3.2HG0173210.1 | | | yes | | | | yes | elongation zone |
| HORVU.MOREX.r3.2HG0173190.1 | | | yes | | | | yes | elongation zone |
| HORVU.MOREX.r3.2HG0172730.1 | | yes | yes | | | yes | | elongation zone |
| HORVU.MOREX.r3.2HG0172050.1 | | | yes | | | | yes | elongation zone |
| HORVU.MOREX.r3.2HG0171780.1 | | yes | | | | yes | | elongation zone |
| HORVU.MOREX.r3.2HG0171550.1 | | yes | yes | | | yes | yes | elongation zone |
| HORVU.MOREX.r3.2HG0170560.1 | | | yes | | | | yes | elongation zone |
| HORVU.MOREX.r3.2HG0170230.1 | | yes | yes | | | yes | | elongation zone |
| HORVU.MOREX.r3.2HG0168730.1 | | | yes | | | | yes | elongation zone |
| HORVU.MOREX.r3.2HG0168700.1 | | yes | | | | yes | | elongation zone |
| HORVU.MOREX.r3.2HG0168090.1 | | yes | yes | | | yes | yes | elongation zone |
| HORVU.MOREX.r3.2HG0167620.1 | | | yes | | | | yes | elongation zone |
| HORVU.MOREX.r3.2HG0167500.2 | | | yes | | | | yes | elongation zone |
| HORVU.MOREX.r3.2HG0165840.1 | | yes | yes | | | yes | yes | elongation zone |
| HORVU.MOREX.r3.2HG0165250.1 | | | yes | | | yes | | elongation zone |
| HORVU.MOREX.r3.2HG0165200.1 | | | yes | | | | yes | elongation zone |
| HORVU.MOREX.r3.2HG0163090.1 | | | yes | | | | yes | elongation zone |
| HORVU.MOREX.r3.2HG0161020.1 | | | yes | | | | yes | elongation zone |
| HORVU.MOREX.r3.2HG0160690.1 | | yes | | | | yes | | elongation zone |
| HORVU.MOREX.r3.2HG0160550.1 | | | yes | | | | yes | elongation zone |
| HORVU.MOREX.r3.2HG0160130.1 | | yes | yes | | | yes | yes | elongation zone |
| HORVU.MOREX.r3.2HG0159500.1 | | | yes | | | | yes | elongation zone |
| HORVU.MOREX.r3.2HG0158180.1 | | | yes | | | yes | | elongation zone |
| HORVU.MOREX.r3.2HG0157290.1 | | | yes | | | | yes | elongation zone |
| HORVU.MOREX.r3.2HG0156810.1 | | yes | | | | yes | | elongation zone |
| HORVU.MOREX.r3.2HG0155810.1 | | | yes | | | | yes | elongation zone |
| HORVU.MOREX.r3.2HG0152140.1 | | | yes | | | | yes | elongation zone |
| HORVU.MOREX.r3.2HG0150450.1 | | | yes | | | yes | yes | elongation zone |
| HORVU.MOREX.r3.2HG0147770.1 | | | yes | | | | yes | elongation zone |

**Table S7** Continued.

| ID | WT_3 h | WT_6 h | WT_12 h | *egt2* vsWT_0 h | *egt2* vsWT_3 h | *egt* 2vsWT_6 h | *egt2* vsWT_12 h | root zone |
|---|---|---|---|---|---|---|---|---|
| HORVU.MOREX.r3.2HG0142740.1 | | | yes | | | | yes | elongation zone |
| HORVU.MOREX.r3.2HG0141140.1 | | yes | yes | | | yes | | elongation zone |
| HORVU.MOREX.r3.2HG0140700.1 | | yes | | | | yes | | elongation zone |
| HORVU.MOREX.r3.2HG0140170.1 | | yes | yes | | | yes | | elongation zone |
| HORVU.MOREX.r3.2HG0139810.1 | | yes | yes | | | | yes | elongation zone |
| HORVU.MOREX.r3.2HG0139050.1 | | | yes | | | | yes | elongation zone |
| HORVU.MOREX.r3.2HG0136670.1 | | | yes | | | | yes | elongation zone |
| HORVU.MOREX.r3.2HG0136010.1 | | | yes | | | | yes | elongation zone |
| HORVU.MOREX.r3.2HG0135860.1 | | | yes | | | | yes | elongation zone |
| HORVU.MOREX.r3.2HG0135650.1 | | | yes | | | | yes | elongation zone |
| HORVU.MOREX.r3.2HG0134300.1 | | yes | | | | | yes | elongation zone |
| HORVU.MOREX.r3.2HG0128230.1 | | | yes | | | | yes | elongation zone |
| HORVU.MOREX.r3.2HG0127260.1 | | yes | yes | | | yes | | elongation zone |
| HORVU.MOREX.r3.2HG0126380.1 | | | yes | | | | yes | elongation zone |
| HORVU.MOREX.r3.2HG0125870.1 | | | yes | | | | yes | elongation zone |
| HORVU.MOREX.r3.2HG0125780.1 | | | yes | | | | yes | elongation zone |
| HORVU.MOREX.r3.2HG0124990.1 | | | yes | | | | yes | elongation zone |
| HORVU.MOREX.r3.2HG0124660.1 | | | yes | | | yes | yes | elongation zone |
| HORVU.MOREX.r3.2HG0124470.1 | | | yes | | | | yes | elongation zone |
| HORVU.MOREX.r3.2HG0122720.1 | | | yes | | | | yes | elongation zone |
| HORVU.MOREX.r3.2HG0122080.1 | | | yes | | | | yes | elongation zone |
| HORVU.MOREX.r3.2HG0121570.1 | | yes | yes | | | yes | yes | elongation zone |
| HORVU.MOREX.r3.2HG0121410.1 | | yes | | | | yes | | elongation zone |
| HORVU.MOREX.r3.2HG0120830.1 | | yes | | | | yes | | elongation zone |
| HORVU.MOREX.r3.2HG0120320.1 | | | yes | | | yes | yes | elongation zone |
| HORVU.MOREX.r3.2HG0119360.1 | | | yes | | | | yes | elongation zone |
| HORVU.MOREX.r3.2HG0117980.1 | | yes | | | | yes | | elongation zone |
| HORVU.MOREX.r3.2HG0117880.1 | | yes | yes | | | yes | yes | elongation zone |
| HORVU.MOREX.r3.2HG0116870.1 | | | yes | | | | yes | elongation zone |
| HORVU.MOREX.r3.2HG0115970.1 | | | yes | | | | yes | elongation zone |
| HORVU.MOREX.r3.2HG0113880.1 | | | yes | | | | yes | elongation zone |
| HORVU.MOREX.r3.2HG0113700.1 | | | yes | | | | yes | elongation zone |
| HORVU.MOREX.r3.2HG0112690.1 | | yes | yes | | | yes | yes | elongation zone |
| HORVU.MOREX.r3.2HG0111570.1 | | | yes | | | | yes | elongation zone |
| HORVU.MOREX.r3.2HG0111130.1 | | | yes | | | | yes | elongation zone |

**Table S7** Continued.

| ID | WT_3 h | WT_6 h | WT_12 h | *egt2* vsWT_0 h | *egt2* vsWT_3 h | *egt* 2vsWT_6 h | *egt2* vsWT_12 h | root zone |
|---|---|---|---|---|---|---|---|---|
| HORVU.MOREX.r3.2HG0110060.1 | | | yes | | | | yes | elongation zone |
| HORVU.MOREX.r3.2HG0108490.1 | | | yes | | | | yes | elongation zone |
| HORVU.MOREX.r3.2HG0107900.1 | | | yes | | | yes | yes | elongation zone |
| HORVU.MOREX.r3.2HG0107450.1 | | | yes | | | | yes | elongation zone |
| HORVU.MOREX.r3.2HG0106750.1 | | | yes | | | | yes | elongation zone |
| HORVU.MOREX.r3.2HG0105680.3 | | | yes | | | | yes | elongation zone |
| HORVU.MOREX.r3.2HG0104510.1 | | | yes | | | | yes | elongation zone |
| HORVU.MOREX.r3.2HG0104480.1 | | | yes | | | | yes | elongation zone |
| HORVU.MOREX.r3.2HG0101220.2 | | | yes | | yes | | | elongation zone |
| HORVU.MOREX.r3.2HG0099780.1 | | | yes | | | | yes | elongation zone |
| HORVU.MOREX.r3.2HG0099160.1 | | | yes | | | | yes | elongation zone |
| HORVU.MOREX.r3.2HG0099010.1 | | | yes | | | | yes | elongation zone |
| HORVU.MOREX.r3.2HG0098420.1 | | | yes | | | | yes | elongation zone |
| HORVU.MOREX.r3.2HG0097950.1 | | | yes | | | | yes | elongation zone |
| HORVU.MOREX.r3.2HG0097390.1 | | yes | yes | | | | yes | elongation zone |
| HORVU.MOREX.r3.2HG0096800.1 | | yes | yes | | | yes | | elongation zone |
| HORVU.MOREX.r3.2HG0096760.1 | | | yes | | | | yes | elongation zone |
| HORVU.MOREX.r3.2HG0096600.1 | | | yes | | | yes | yes | elongation zone |
| HORVU.MOREX.r3.2HG0096230.1 | | yes | | | | yes | | elongation zone |
| HORVU.MOREX.r3.2HG0095970.1 | | | yes | | | | yes | elongation zone |
| HORVU.MOREX.r3.1HG0094980.1 | | yes | yes | | | yes | | elongation zone |
| HORVU.MOREX.r3.1HG0094770.1 | | yes | | | | yes | yes | elongation zone |
| HORVU.MOREX.r3.1HG0094110.1 | | yes | yes | | | yes | | elongation zone |
| HORVU.MOREX.r3.1HG0093050.1 | | yes | yes | | | yes | | elongation zone |
| HORVU.MOREX.r3.1HG0092810.1 | | | yes | | | | yes | elongation zone |
| HORVU.MOREX.r3.1HG0091820.1 | | | yes | | | yes | yes | elongation zone |
| HORVU.MOREX.r3.1HG0091360.1 | | | yes | | | yes | yes | elongation zone |
| HORVU.MOREX.r3.1HG0091240.1 | | | yes | | | | yes | elongation zone |
| HORVU.MOREX.r3.1HG0088530.1 | | yes | yes | | | yes | | elongation zone |
| HORVU.MOREX.r3.1HG0087170.1 | | yes | yes | | | yes | | elongation zone |
| HORVU.MOREX.r3.1HG0086480.1 | | yes | | | | yes | | elongation zone |
| HORVU.MOREX.r3.1HG0086390.1 | | yes | yes | | | yes | yes | elongation zone |
| HORVU.MOREX.r3.1HG0086370.1 | | yes | yes | | | yes | yes | elongation zone |
| HORVU.MOREX.r3.1HG0086010.1 | | | yes | | | | yes | elongation zone |
| HORVU.MOREX.r3.1HG0085420.1 | | | yes | | | | yes | elongation zone |

**Table S7** Continued.

| ID | WT_3 h | WT_6 h | WT_12 h | *egt2* vsWT_0 h | *egt2* vsWT_3 h | *egt* 2vsWT_6 h | *egt2* vsWT_12 h | root zone |
|---|---|---|---|---|---|---|---|---|
| HORVU.MOREX.r3.1HG0085380.1 | | | yes | | | yes | | elongation zone |
| HORVU.MOREX.r3.1HG0085140.1 | | | yes | | | | yes | elongation zone |
| HORVU.MOREX.r3.1HG0084890.1 | | | yes | | | | yes | elongation zone |
| HORVU.MOREX.r3.1HG0084850.1 | | | yes | | | | yes | elongation zone |
| HORVU.MOREX.r3.1HG0084760.1 | | | yes | | | | yes | elongation zone |
| HORVU.MOREX.r3.1HG0083980.1 | | | yes | | | | yes | elongation zone |
| HORVU.MOREX.r3.1HG0083440.1 | | | yes | | | | yes | elongation zone |
| HORVU.MOREX.r3.1HG0082770.1 | | yes | yes | | | yes | yes | elongation zone |
| HORVU.MOREX.r3.1HG0082750.1 | | | yes | | | | yes | elongation zone |
| HORVU.MOREX.r3.1HG0081950.1 | | yes | yes | | | yes | | elongation zone |
| HORVU.MOREX.r3.1HG0080760.1 | | | yes | | | | yes | elongation zone |
| HORVU.MOREX.r3.1HG0080230.1 | | | yes | | | | yes | elongation zone |
| HORVU.MOREX.r3.1HG0080180.1 | | yes | yes | | | yes | | elongation zone |
| HORVU.MOREX.r3.1HG0079870.1 | | yes | yes | | | yes | | elongation zone |
| HORVU.MOREX.r3.1HG0079810.1 | | | yes | | | | yes | elongation zone |
| HORVU.MOREX.r3.1HG0079800.1 | | yes | yes | | | yes | yes | elongation zone |
| HORVU.MOREX.r3.1HG0079750.1 | | yes | yes | | | yes | yes | elongation zone |
| HORVU.MOREX.r3.1HG0079280.1 | | | yes | | | | yes | elongation zone |
| HORVU.MOREX.r3.1HG0079250.1 | | | yes | | | | yes | elongation zone |
| HORVU.MOREX.r3.1HG0079140.1 | | | yes | | | yes | yes | elongation zone |
| HORVU.MOREX.r3.1HG0078910.1 | | | yes | | | | yes | elongation zone |
| HORVU.MOREX.r3.1HG0078370.1 | | | yes | | | | yes | elongation zone |
| HORVU.MOREX.r3.1HG0078040.1 | | | yes | | | | yes | elongation zone |
| HORVU.MOREX.r3.1HG0077950.1 | | | yes | | | yes | | elongation zone |
| HORVU.MOREX.r3.1HG0077880.1 | | | yes | | | | yes | elongation zone |
| HORVU.MOREX.r3.1HG0077170.1 | | | yes | | | | yes | elongation zone |
| HORVU.MOREX.r3.1HG0076980.1 | | | yes | | | yes | yes | elongation zone |
| HORVU.MOREX.r3.1HG0076540.1 | | | yes | | | | yes | elongation zone |
| HORVU.MOREX.r3.1HG0075730.1 | | yes | yes | | | yes | | elongation zone |
| HORVU.MOREX.r3.1HG0075590.1 | | | yes | | | yes | yes | elongation zone |
| HORVU.MOREX.r3.1HG0075440.1 | | yes | yes | | | yes | yes | elongation zone |
| HORVU.MOREX.r3.1HG0074500.1 | | | yes | | | | yes | elongation zone |
| HORVU.MOREX.r3.1HG0074490.1 | | | yes | | | | yes | elongation zone |
| HORVU.MOREX.r3.1HG0073460.1 | | yes | | | | yes | | elongation zone |
| HORVU.MOREX.r3.1HG0073430.1 | | yes | | | | yes | | elongation zone |

**Table S7** Continued.

| ID | WT_3 h | WT_6 h | WT_12 h | *egt2* vsWT_0 h | *egt2* vsWT_3 h | *egt* 2vsWT_6 h | *egt2* vsWT_12 h | root zone |
|---|---|---|---|---|---|---|---|---|
| HORVU.MOREX.r3.1HG0073170.1 | | yes | yes | | | yes | | elongation zone |
| HORVU.MOREX.r3.1HG0073160.1 | | | yes | | | | yes | elongation zone |
| HORVU.MOREX.r3.1HG0072910.2 | | yes | yes | | | yes | | elongation zone |
| HORVU.MOREX.r3.1HG0072580.1 | | yes | yes | | | yes | yes | elongation zone |
| HORVU.MOREX.r3.1HG0072540.1 | | | yes | | | | yes | elongation zone |
| HORVU.MOREX.r3.1HG0072520.1 | | | yes | | | | yes | elongation zone |
| HORVU.MOREX.r3.1HG0070480.1 | | | yes | yes | yes | yes | yes | elongation zone |
| HORVU.MOREX.r3.1HG0070340.1 | | yes | yes | | | yes | | elongation zone |
| HORVU.MOREX.r3.1HG0069940.1 | | | yes | | | | yes | elongation zone |
| HORVU.MOREX.r3.1HG0069870.1 | | | yes | | | | yes | elongation zone |
| HORVU.MOREX.r3.1HG0069410.1 | | yes | | | | yes | | elongation zone |
| HORVU.MOREX.r3.1HG0069380.1 | | yes | yes | yes | yes | yes | yes | elongation zone |
| HORVU.MOREX.r3.1HG0069030.1 | | yes | yes | | yes | yes | yes | elongation zone |
| HORVU.MOREX.r3.1HG0067930.1 | | | yes | | | | yes | elongation zone |
| HORVU.MOREX.r3.1HG0067720.1 | | yes | yes | | | yes | yes | elongation zone |
| HORVU.MOREX.r3.1HG0067630.1 | | | yes | | | yes | | elongation zone |
| HORVU.MOREX.r3.1HG0067410.1 | | | yes | | | yes | yes | elongation zone |
| HORVU.MOREX.r3.1HG0066800.1 | | yes | yes | | | yes | | elongation zone |
| HORVU.MOREX.r3.1HG0066530.1 | | | yes | | | | yes | elongation zone |
| HORVU.MOREX.r3.1HG0065860.1 | | yes | yes | | | | yes | elongation zone |
| HORVU.MOREX.r3.1HG0065370.1 | | | yes | | | | yes | elongation zone |
| HORVU.MOREX.r3.1HG0065020.1 | | | yes | | | | yes | elongation zone |
| HORVU.MOREX.r3.1HG0064740.1 | | | yes | | | | yes | elongation zone |
| HORVU.MOREX.r3.1HG0064610.1 | | | yes | | | | yes | elongation zone |
| HORVU.MOREX.r3.1HG0064320.1 | | | yes | | | | yes | elongation zone |
| HORVU.MOREX.r3.1HG0063870.1 | | | yes | | | | yes | elongation zone |
| HORVU.MOREX.r3.1HG0063810.1 | | yes | | | | yes | | elongation zone |
| HORVU.MOREX.r3.1HG0062110.1 | | yes | yes | | | yes | | elongation zone |
| HORVU.MOREX.r3.1HG0061710.1 | | yes | yes | | | yes | | elongation zone |
| HORVU.MOREX.r3.1HG0061390.1 | | | yes | | | | yes | elongation zone |
| HORVU.MOREX.r3.1HG0061220.1 | | | yes | | | | yes | elongation zone |
| HORVU.MOREX.r3.1HG0060320.1 | | | yes | | | | yes | elongation zone |
| HORVU.MOREX.r3.1HG0059790.1 | | | yes | | | | yes | elongation zone |
| HORVU.MOREX.r3.1HG0059000.1 | | | yes | | | | yes | elongation zone |
| HORVU.MOREX.r3.1HG0058800.1 | | | yes | | | | yes | elongation zone |

**Table S7** Continued.

| ID | WT_3 h | WT_6 h | WT_12 h | *egt2* vsWT_0 h | *egt2* vsWT_3 h | *egt* 2vsWT_6 h | *egt2* vsWT_12 h | root zone |
|---|---|---|---|---|---|---|---|---|
| HORVU.MOREX.r3.1HG0058640.1 | | | yes | | | | yes | elongation zone |
| HORVU.MOREX.r3.1HG0058050.1 | | | yes | | | | yes | elongation zone |
| HORVU.MOREX.r3.1HG0057990.1 | | yes | yes | | | | yes | elongation zone |
| HORVU.MOREX.r3.1HG0057620.1 | | | yes | | | | yes | elongation zone |
| HORVU.MOREX.r3.1HG0057030.1 | | yes | yes | | | yes | | elongation zone |
| HORVU.MOREX.r3.1HG0056740.1 | | | yes | | | | yes | elongation zone |
| HORVU.MOREX.r3.1HG0056620.1 | | yes | yes | | | yes | | elongation zone |
| HORVU.MOREX.r3.1HG0056560.1 | | | yes | | | | yes | elongation zone |
| HORVU.MOREX.r3.1HG0055240.1 | | yes | yes | | | yes | | elongation zone |
| HORVU.MOREX.r3.1HG0055230.1 | | | yes | | | | yes | elongation zone |
| HORVU.MOREX.r3.1HG0054500.1 | | | yes | | | | yes | elongation zone |
| HORVU.MOREX.r3.1HG0054200.1 | | | yes | | | | yes | elongation zone |
| HORVU.MOREX.r3.1HG0054010.1 | | yes | yes | | | yes | yes | elongation zone |
| HORVU.MOREX.r3.1HG0053530.1 | | | yes | | | | yes | elongation zone |
| HORVU.MOREX.r3.1HG0053060.1 | | yes | yes | | | yes | | elongation zone |
| HORVU.MOREX.r3.1HG0052120.1 | | yes | yes | | | | yes | elongation zone |
| HORVU.MOREX.r3.1HG0052080.1 | | | yes | | | | yes | elongation zone |
| HORVU.MOREX.r3.1HG0051960.1 | | yes | yes | | | | yes | elongation zone |
| HORVU.MOREX.r3.1HG0051020.1 | | | yes | | | | yes | elongation zone |
| HORVU.MOREX.r3.1HG0050990.1 | | | yes | | | | yes | elongation zone |
| HORVU.MOREX.r3.1HG0050920.1 | | yes | yes | | | yes | yes | elongation zone |
| HORVU.MOREX.r3.1HG0050700.1 | | yes | yes | | | yes | yes | elongation zone |
| HORVU.MOREX.r3.1HG0049550.1 | | | yes | yes | yes | | | elongation zone |
| HORVU.MOREX.r3.1HG0046180.1 | | yes | yes | | | yes | yes | elongation zone |
| HORVU.MOREX.r3.1HG0042740.1 | | yes | yes | | | | yes | elongation zone |
| HORVU.MOREX.r3.1HG0039670.1 | | yes | yes | | | yes | yes | elongation zone |
| HORVU.MOREX.r3.1HG0039230.1 | | | yes | | | | yes | elongation zone |
| HORVU.MOREX.r3.1HG0038960.1 | | | yes | | | | yes | elongation zone |
| HORVU.MOREX.r3.1HG0038800.1 | | yes | yes | | | yes | yes | elongation zone |
| HORVU.MOREX.r3.1HG0036930.1 | | | yes | | | | yes | elongation zone |
| HORVU.MOREX.r3.1HG0035400.1 | | yes | yes | | | | yes | elongation zone |
| HORVU.MOREX.r3.1HG0032910.1 | | | yes | | | | yes | elongation zone |
| HORVU.MOREX.r3.1HG0032230.1 | | | yes | | | | yes | elongation zone |
| HORVU.MOREX.r3.1HG0028320.1 | | yes | yes | | | | yes | elongation zone |
| HORVU.MOREX.r3.1HG0027200.1 | | yes | yes | | | yes | | elongation zone |

**Table S7** Continued.

| ID | WT_3 h | WT_6 h | WT_12 h | *egt2* vsWT_0 h | *egt2* vsWT_3 h | *egt* 2vsWT_6 h | *egt2* vsWT_12 h | root zone |
|---|---|---|---|---|---|---|---|---|
| HORVU.MOREX.r3.1HG0026830.1 | | yes | yes | | | | yes | elongation zone |
| HORVU.MOREX.r3.1HG0026080.1 | | yes | yes | | | yes | | elongation zone |
| HORVU.MOREX.r3.1HG0026070.2 | | | yes | | | | yes | elongation zone |
| HORVU.MOREX.r3.1HG0025870.1 | | yes | yes | | | | yes | elongation zone |
| HORVU.MOREX.r3.1HG0025320.1 | | yes | yes | | | yes | yes | elongation zone |
| HORVU.MOREX.r3.1HG0024510.1 | | | yes | | | | yes | elongation zone |
| HORVU.MOREX.r3.1HG0024280.1 | | | yes | | | | yes | elongation zone |
| HORVU.MOREX.r3.1HG0024230.1 | | | yes | | | | yes | elongation zone |
| HORVU.MOREX.r3.1HG0024040.1 | | yes | yes | | | yes | | elongation zone |
| HORVU.MOREX.r3.1HG0023610.1 | | | yes | | | | yes | elongation zone |
| HORVU.MOREX.r3.1HG0020980.1 | | | yes | | | | yes | elongation zone |
| HORVU.MOREX.r3.1HG0018810.1 | | | yes | | | | yes | elongation zone |
| HORVU.MOREX.r3.1HG0017310.1 | | | yes | | | | yes | elongation zone |
| HORVU.MOREX.r3.1HG0017280.1 | | yes | yes | | | yes | | elongation zone |
| HORVU.MOREX.r3.1HG0016810.1 | | | yes | | | | yes | elongation zone |
| HORVU.MOREX.r3.1HG0015750.1 | | yes | yes | | | yes | yes | elongation zone |
| HORVU.MOREX.r3.1HG0014410.1 | | | yes | | | | yes | elongation zone |
| HORVU.MOREX.r3.1HG0012330.1 | | | yes | | | | yes | elongation zone |
| HORVU.MOREX.r3.1HG0010340.1 | | yes | yes | | | yes | | elongation zone |
| HORVU.MOREX.r3.1HG0008510.2 | | | yes | | | | yes | elongation zone |
| HORVU.MOREX.r3.1HG0008200.1 | | | yes | | | yes | yes | elongation zone |
| HORVU.MOREX.r3.1HG0008190.1 | | | yes | yes | yes | | | elongation zone |
| HORVU.MOREX.r3.1HG0007670.1 | | | yes | | | | yes | elongation zone |
| HORVU.MOREX.r3.1HG0007570.1 | | | yes | yes | yes | yes | yes | elongation zone |
| HORVU.MOREX.r3.1HG0006650.1 | | | yes | | | | yes | elongation zone |
| HORVU.MOREX.r3.1HG0006610.1 | | | yes | | | | yes | elongation zone |
| HORVU.MOREX.r3.1HG0006400.1 | | yes | yes | | | | yes | elongation zone |
| HORVU.MOREX.r3.1HG0006230.1 | | yes | yes | | | yes | yes | elongation zone |
| HORVU.MOREX.r3.1HG0005840.1 | | | yes | | | | yes | elongation zone |
| HORVU.MOREX.r3.1HG0004300.1 | | | yes | | | | yes | elongation zone |
| HORVU.MOREX.r3.1HG0003620.1 | | yes | yes | | | yes | yes | elongation zone |
| HORVU.MOREX.r3.1HG0003150.1 | | | yes | | | | yes | elongation zone |
| HORVU.MOREX.r3.1HG0002960.1 | | yes | yes | | | yes | | elongation zone |
| HORVU.MOREX.r3.1HG0001510.1 | | | yes | | | | yes | elongation zone |
| HORVU.MOREX.r3.1HG0001460.1 | | | yes | | | | yes | elongation zone |

**Table S7** Continued.

| ID | WT_3 h | WT_6 h | WT_12 h | *egt2* vsWT_0 h | *egt2* vsWT_3 h | *egt* 2vsWT_6 h | *egt2* vsWT_12 h | root zone |
|---|---|---|---|---|---|---|---|---|
| HORVU.MOREX.r3.1HG0000050.1 | | yes | yes | | | yes | | elongation zone |
