## Supporting Information Table 8 for "ENHANCED GRAVITROPISM 2 coordinates molecular adaptations to gravistimulation in the elongation zone of barley roots"

**Table S8** Overview of the candidates interacting with EGT2 as identified by Y2H screening.

| ID_version 3(2021) | Description | cuptered times | | confirmation | | differentially expressed genes in *egt2* | | differentially expressed genes in rotated WT | |
|---|---|---|---|---|---|---|---|---|---|
| | | 1st | 2nd | by 1-no-1 Y2H | by BiFC | FDR < 0.05 | FDR < 5%; Log2FC ≥ 1 | FDR < 0.05 | FDR < 5%; Log2FC ≥ 1 |
| HORVU.MOREX.r3.1HG0078770.1 | NADH-quinone oxidoreductase, subunit E | 7 | 12 | Yes | | | | yes | |
| HORVU.MOREX.r3.3HG0330120.1 | O-methyltransferase | 0 | 2 | yes | yes | yes | yes | yes | yes |
| HORVU.MOREX.r3.6HG0548050.1 | Acetyl-coenzyme A synthetase | 2 | 10 | Yes | | | | | |
| HORVU.MOREX.r3.5HG0480100.1 | Cysteine protease | 2 | 1 | yes | | yes | | | |
| HORVU.MOREX.r3.1HG0053480.1 | Cinnamoyl-CoA reductase-like protein | 1 | 3 | yes | | | | | |
| HORVU.MOREX.r3.7HG0726360.1 | 12-oxophytodienoate reductase | 1 | 1 | yes | | yes | | | |
| HORVU.MOREX.r3.1HG0085590.1 | Non specific phospholipase C | 1 | 3 | yes | | | | | |
| HORVU.MOREX.r3.5HG0475360.1 | Mitochondrial outer membrane porin | 1 | 5 | yes | | | | | |
| HORVU.MOREX.r3.7HG0721830.1/HORVU.MOREX.r3.7HG0721840.1(two genes with high similarity) | Germin-like protein | 1 | 3 | yes | | | | | |
| HORVU.MOREX.r3.6HG0599140.1 | Kinesin-like protein | 1 | 1 | yes | | | | | |
| HORVU.MOREX.r3.7HG0736300.1 | Glycosyltransferase | 0 | 2 | yes | | yes | yes | | |
| HORVU.MOREX.r3.1HG0006660.1 | Inorganic pyrophosphatase family protein | 0 | 3 | yes | | | | yes | |
| HORVU.MOREX.r3.2HG0197110.1 | AT hook motif DNA-binding family protein | 0 | 4 | yes | | | | yes | |
| HORVU.MOREX.r3.3HG0269000.1 | zinc finger FYVE domain protein | 1 | 1 | yes | | | | | |
| HORVU.MOREX.r3.3HG0244960.1 | TPR repeat-containing thioredoxin TTL1 | 0 | 2 | yes | | | | | |
| HORVU.MOREX.r3.5HG0423100.1 | Pathogenesis-related thaumatin family protein | 1 | 2 | yes | | | | | |
| HORVU.MOREX.r3.5HG0523150.1/ HORVU.MOREX.r3.5HG0523140.1 | Protein phosphatase 2c, putative/ DNA ligase-like protein | 1 | 1 | yes | | yes | | yes | |
| HORVU.MOREX.r3.5HG0517090.1 | Chaperone protein dnaJ, putative | 0 | 4 | yes | | | | | |
| HORVU.MOREX.r3.6HG0607930.1 | Ring box protein | 0 | 6 | yes | | | | | |
| HORVU.MOREX.r3.2HG0108780.1 | 26S protease regulatory subunit | 7 | 0 | yes | | | | yes | |
| HORVU.MOREX.r3.6HG0571770.1 | Glycine cleavage system H protein | 1 | 2 | yes | | | | | |
| HORVU.MOREX.r3.3HG0292400.1 | Histone-lysine N-methyltransferase | 3 | 0 | yes | | | | | |
| HORVU.MOREX.r3.6HG0608910.1 | Heavy metal transport/detoxification superfamily protein | 0 | 7 | yes | | | | yes | yes |
| HORVU.MOREX.r3.6HG0572880.1 | Alpha/beta-Hydrolases superfamily protein | 0 | 2 | | | yes | | | |
| HORVU.MOREX.r3.6HG0626440.1 | Aldehyde dehydrogenase | 0 | 1 | | | | | | |
| HORVU.MOREX.r3.5HG0462200.1 | Metacaspase | 0 | 1 | | | | | yes | yes |
| HORVU.MOREX.r3.7HG0663950.1 | Alpha/beta-Hydrolases superfamily protein | 0 | 2 | | | | | | |
| HORVU.MOREX.r3.1HG0016210.1 | Endoglucanase | 0 | 1 | | | | | | |
| HORVU.MOREX.r3.6HG0630640.1 | Aconitate hydratase | 0 | 2 | | | | | | |
| HORVU.MOREX.r3.4HG0336620.1 | GAI-like protein 1 | 0 | 1 | | | | | | |
| HORVU.MOREX.r3.4HG0410090.1 | Hypersensitive-induced response protein 1 | 0 | 2 | | | | | yes | |
| HORVU.MOREX.r3.7HG0709860.1 | Polyadenylate-binding protein | 0 | 1 | | no | yes | | yes | |
| HORVU.MOREX.r3.3HG0282230.1 | 4-hydroxy-3-methylbut-2-enyl diphosphate reductase | 0 | 2 | | | | | | |
| HORVU.MOREX.r3.6HG0630520.1 | AT hook motif DNA-binding family protein | 0 | 1 | | | | | | |
| HORVU.MOREX.r3.6HG0613040.1 | 2-oxoglutarate-dependent dioxygenase-related family protein | 0 | 1 | | | yes | | | |
| HORVU.MOREX.r3.4HG0413890.1 | Alpha/beta-Hydrolases superfamily protein | 0 | 2 | | | | | yes | |
| HORVU.MOREX.r3.6HG0608920.2 | DegP protease-like | 0 | 7 | | | | | | |
| HORVU.MOREX.r3.3HG0223160.1 | aberrant root formation protein | 0 | 1 | | | yes | | yes | |
| HORVU.MOREX.r3.4HG0383750.2 | 26S protease regulatory subunit | 0 | 1 | | | | | | |
| HORVU.MOREX.r3.7HG0721230.1 | UPF0183 protein | 0 | 1 | | | | | | |
| HORVU.MOREX.r3.3HG0292280.1 | Ubiquitin carboxyl-terminal hydrolase-like protein | 0 | 1 | | | yes | | | |
| HORVU.MOREX.r3.4HG0333780.1 | Glucan endo-1,3-beta-glucosidase | 0 | 1 | | | | | | |
| HORVU.MOREX.r3.5HG0423110.1 | Thaumatin-like protein | 0 | 1 | | | | | | |
| HORVU.MOREX.r3.5HG0511040.1 | Heavy metal transport/detoxification superfamily protein | 0 | 1 | | | | | yes | |

**Table S8** Continued.

| ID_version 3(2021) | Description | cuptered times | | confirmation | | differentially expressed genes in *egt2* | | differentially expressed genes in rotated WT | |
|---|---|---|---|---|---|---|---|---|---|
| | | 1st | 2nd | by 1-no-1 Y2H | by BiFC | FDR < 0.05 | FDR < 5%; Log2FC ≥ 1 | FDR < 0.05 | FDR < 5%; Log2FC ≥ 1 |
| HORVU.MOREX.r3.3HG0278140.1 | WRKY transcription factor | 0 | 1 | | | | | | |
| HORVU.MOREX.r3.1HG0000050.1 | RING-finger ubiquitin ligase | 0 | 1 | | no | yes | | yes | |
| HORVU.MOREX.r3.6HG0564530.1 | glucuronoxylan 4-O-methyltransferase-like protein (DUF579) | 0 | 1 | | | | | | |
| HORVU.MOREX.r3.6HG0620720.1 | Ribonuclease | 0 | 1 | | | | | yes | |
| HORVU.MOREX.r3.7HG0749870.1 | Glucuronoxylan 4-O-methyltransferase | 0 | 1 | | yes | yes | | yes | |
| HORVU.MOREX.r3.1HG0001180.1 | Leucine-rich repeat protein kinase family protein | 0 | 1 | | | | | | |
| HORVU.MOREX.r3.2HG0191150.1 | Polygalacturonase QRT3 | 0 | 1 | | | | | yes | |
| HORVU.MOREX.r3.7HG0732480.1 | Zinc finger CCCH domain protein | 0 | 1 | | | | | | |
| HORVU.MOREX.r3.3HG0230590.1 | Triosephosphate isomerase | 0 | 1 | | | | | | |
| HORVU.MOREX.r3.2HG0205420.1 | Heavy metal transport/detoxification superfamily protein | 0 | 1 | | | | | yes | |
| HORVU.MOREX.r3.6HG0572960.1 | Chlorophyll a-b binding protein, chloroplastic | 0 | 1 | | | | | yes | yes |
| HORVU.MOREX.r3.5HG0512400.1 | GDSL esterase/lipase | 0 | 1 | | | | | | |
| HORVU.MOREX.r3.6HG0625570.1 | Transmembrane protein | 0 | 2 | | | yes | | yes | |
| HORVU.MOREX.r3.5HG0487040.1 | Alpha/beta-Hydrolases superfamily protein, putative | 0 | 1 | | | | | yes | |
| HORVU.MOREX.r3.5HG0516310.1 | Patatin | 0 | 1 | | | | | yes | |
| HORVU.MOREX.r3.2HG0163850.1 | Phosphoribosylformylglycinamidine synthase | 0 | 1 | | | | | | |
| HORVU.MOREX.r3.2HG0126380.1 | Heavy metal transport/detoxification superfamily protein | 0 | 1 | | yes | yes | | yes | |
| HORVU.MOREX.r3.5HG0499760.1 | Cysteine proteinase | 0 | 2 | | | | | | |
| HORVU.MOREX.r3.1HG0074750.1 | Ferredoxin | 2 | 0 | | | | | | |
| HORVU.MOREX.r3.7HG0721250.1 | NADH dehydrogenase [ubiquinone] 1 beta subcomplex subunit 9 | 1 | 0 | | | | | | |
| HORVU.MOREX.r3.1HG0057560.1 | RNA-binding KH domain-containing protein | 1 | 0 | | | | | | |
| HORVU.MOREX.r3.2HG0146530.1 | ATP synthase epsilon chain | 1 | 0 | | | | | | |
| HORVU.MOREX.r3.2HG0161130.1 | Eukaryotic translation initiation factor 3 subunit H | 2 | 0 | | | | | | |
| HORVU.MOREX.r3.2HG0176810.2 | F-box family protein | 2 | 0 | | | | | | |
| HORVU.MOREX.r3.4HG0395480.1 | calcium/calcium/calmodulin-dependent Serine/Threonine-kinase | 1 | 0 | | | | | | |
| HORVU.MOREX.r3.3HG0291750.1 | Lipoyl synthase | 1 | 0 | | | | | | |
| HORVU.MOREX.r3.7HG0655800.1 | Caffeoyl-CoA O-methyltransferase | 1 | 0 | | | | | | |
| HORVU.MOREX.r3.1HG0043190.1 | Malate dehydrogenase | 1 | 0 | | | | | | |
| HORVU.MOREX.r3.1HG0018270.1 | Bushy growth protein | 1 | 0 | | | | | | |
| HORVU.MOREX.r3.7HG0636530.1 | MADS-box transcription factor 8 | 1 | 0 | | | | | yes | |
| HORVU.MOREX.r3.2HG0107060.1 | IAA-amino acid hydrolase ILR1 | 1 | 0 | | | | | | |
| HORVU.MOREX.r3.4HG0413910.1 | Cell wall invertase | 1 | 0 | | | | | | |
| HORVU.MOREX.r3.5HG0444520.1 | NADP-dependent alkenal double bond reductase | 1 | 0 | | | yes | | | |
| HORVU.MOREX.r3.4HG0416190.1 | Coatomer subunit alpha | 1 | 0 | | | yes | | | |
| HORVU.MOREX.r3.6HG0571760.1 | Splicing factor, arginine/serine-rich 12 | 1 | 0 | | | | | | |
