## Supporting Information Table 9 for "ENHANCED GRAVITROPISM 2 coordinates molecular adaptations to gravistimulation in the elongation zone of barley roots"

**Table S9** List of oligonucleotide primers.

| Name | Purpose | Barley identifier | Sequence 5'-3' |
|---|---|---|---|
| EGT2-AD-fw (EcoRI) | EGT2 cloning (pGADT7/ pGBKT7 plasmids) | HORVU.MOREX.r3.5HG0447830.1 | CGGAATTCATGTCGGCTAAACGATCACAG |
| EGT2-AD-rv (BamHI) | | | CGGGATCCTCAAGGTTCCAGCTTTAGG |
| EGT2-attB1-fw | EGT2 cloning (2in1 system) | HORVU.MOREX.r3.5HG0447830.1 | GGGGACA AGT TTG TAC AAA AAA GCA GGC TTA ATGTCGGCTAAACGATCAC |
| EGT2-attB4-rv | EGT2 cloning (2in1 system, with stop codon) | | GGGGAC AAC TTT GTA TAG AAA AGT TGG GTG AGGTTCCAGCTTTAGGGAG |
| EGT2-attB4-rv(n-tag) | EGT2 cloning (2in1 system, without stop codon) | | GGGGACCACTTTGTACAAGAAAGCTGGGT TCAAGGCTTGAGAGCAATCG |
| OMT- attb3-fw | OMT cloning (2in1 system) | HORVU.MOREX.r3.3HG0330120.1 | GGGGACA ACT TTG TAT AAT AAA GTT GGA ATGGGGTCCATCGCCGCC |
| OMT- attb2-rv | OMT cloning (2in1 system, with stop codon) | | GGGGACCACTTTGTACAAGAAAGCTGGGT CTACTTGGTGAACTCGATG |
| GXM - attb3-fw | GXM cloning (2in1 system) | HORVU.MOREX.r3.7HG0749870.1 | GGGGACA ACT TTG TAT AAT AAA GTT GGA ATGTCGAGCCCCGTGCAC |
| GXM - attb2-rv | GXM cloning (2in1 system, with stop codon) | | GGGGACCACTTTGTACAAGAAAGCTGGGT CTAATTGAGAGGGCAGAA |
| PBP-attb3-fw | PBP cloning (2in1 system) | HORVU.MOREX.r3.7HG0709860.1 | GGGGACA ACT TTG TAT AAT AAA GTT GGA ATGGTTGCCGTGGCGGCC |
| PBP-attb2-rv | PBP cloning (2in1 system, with stop codon) | | GGGGACCACTTTGTACAAGAAAGCTGGGT TCAGTGCGACACCACACGAG |
| HMT-attb3-fw | HMT cloning (2in1 system) | HORVU.MOREX.r3.2HG0126380.1 | GGGGACA ACT TTG TAT AAT AAA GTT GGA ATGGCGGACAAGATCTCCAC |
| HMT-attb2-rv | HMT cloning (2in1 system, with stop codon) | | GGGGACCACTTTGTACAAGAAAGCTGGGT TCACATGACGCTGCACCCGG |
| HORVU.MOREX.r3.1HG0000050.1_attb3-fw | HORVU.MOREX.r3.1HG0000050.1 cloning (2in1 system) | HORVU.MOREX.r3.1HG0000050.1 | GGGGACA ACT TTG TAT AAT AAA GTT GGA ATGGCGATGCGGGGCGTC |
| HORVU.MOREX.r3.1HG0000050.1 _attb2-rv (+TAG) | HORVU.MOREX.r3.1HG0000050.1 cloning (2in1 system, with stop codon) | | GGGGACCACTTTGTACAAGAAAGCTGGGT TCAGCGATCCCGCAGCTCAT |
